# *Ptf1a* robustly drives the gliogenic switch in the rodent embryonic cortex in a dosage-dependent manner by activating pro-glial gene expression programs

**DOI:** 10.1101/2025.10.14.682493

**Authors:** Haixiang Li, Kaiqin Lu, Xiaoxuan Wang, Can Jiao, Kang Huang, Shibo Xu, Yan Liu, She Chen, Shuijin He

**Affiliations:** School of Life Science and Technology, ShanghaiTech University, Shanghai, 201210, China; University of Chinese Academy of Sciences, Beijing, 100049, China; NHC Key Laboratory of Glycoconjugates Research, Department of Biochemistry and Molecular Biology, School of Basic Medical Sciences, Fudan University, Shanghai 200032, China; Institute for Stem Cell and Neural Regeneration, School of Pharmacy, Nanjing Medical University, Nanjing, 211166, China; State Key Laboratory of Reproductive Medicine, Nanjing Medical University, Nanjing, 211166, China; Shanghai Clinical Research and Trial Center, Shanghai, 201210, China

## Abstract

It is widely believed that the gliogenic switch during rodent embryonic development is governed by the orchestrated crosstalk between a cohort of genes and extracellular cues. Here we report that ectopic expression of the single bHLH factor, pancreas transcription factor 1α (PTF1A), is sufficient to drive radial glial progenitors (RGPs)-regardless of progenitor heterogeneity and developmental stage-to differentiate into glial cells. *Ptf1a*- expressing RGPs exhibit progenitor behaviors indicative of a neurogenic-to-gliogenic fate transition, resembling endogenous progenitors at the late stages of embryonic development, and preferentially produce OLIG2+ oligodendrocytes *in vivo*, some of which are positive for PDGFRα or CC1. This robust gliogenic competency depends on the dosage of stable *Ptf1a* expression in RGPs. RNAseq reveals ‘the glial transcriptome’, including upregulated expression of several notch signaling components and the endothelin receptors in *Ptf1a* RGPs. We further identify *Ednrb* as one of the downstream targets of *Ptf1a* that directs RGPs toward gliogenesis via the ERK1/2 signaling pathway. In summary, our study uncovers a novel and robust role for *Ptf1a* in glial fate specification, offering a potential strategy for generating human oligodendrocytes *in vitro*.

## Introduction

The mammalian cortex is a complex structure comprising diverse populations of neurons and glial cells. Increasing evidence shows that glial cells not merely passive structural components as brain “glue” but play crucial roles in sculpting synaptic connectivity, propagating rapid impulses, and maintaining neural homeostasis (Fields *et al*, 2015; Zuchero & Barres, 2013). Macroglia—including astrocytes and oligodendrocytes—are derived from radial glial progenitors (RGPs) in the embryonic forebrain through spatially and temporally regulated developmental gene programs (Rowitch & Kriegstein, 2010). In the dorsal telencephalon, RGPs acquire gliogenic competence and generate both astrocytes and oligodendrocytes following a ‘gliogenic switch’ that occurs during late embryogenesis (Gotz & Huttner, 2005; Molofsky *et al*, 2012; Qian *et al*, 2000). In contrast, ventral RGPs begin to produce oligodendrocytes at early embryonic stages (Kessaris *et al*, 2006). However, recent single-cell RNA sequencing studies have identified a distinct population of astrocytic progenitors with a ventral origin, challenging this longstanding view (Bandler *et al*, 2022; Liu *et al*, 2022).

Despite advances in identifying glial lineage trajectories, the molecular mechanisms by which RGPs gain gliogenic potential remain incompletely understood. Both extracellular cues and intrinsic genetic programs are believed to orchestrate the gliogenic switch in the dorsal and ventral telencephalon (Kang *et al*, 2012; Lin *et al*, 2021; Rowitch & Kriegstein, 2010; Sauvageot & Stiles, 2002; Tchieu *et al*, 2019; Zuchero & Barres, 2013). For example, neurotrophic cytokines such as cardiotrophin- 1 (CT-1), ciliary neurotrophic factor (CNTF), and fibroblast growth factor-2 (FGF2) act as extracellular cues to activate the JAK-STAT and MEK/ERK signaling pathways, which then promotes glial fate commitment (Barnabe-Heider *et al*, 2005; Bonni *et al*, 1997; Gao *et al*, 2025; Li *et al*, 2012; Morrow *et al*, 2001; Song & Ghosh, 2004).

Transcription factors of the basic helix-loop-helix (bHLH) family orchestrate with the Notch signaling to regulate the balance between neuronal and glial differentiation across both invertebrate and vertebrate nervous systems (Anderson, 1999; Anderson *et al*, 1997; Guillemot, 1995; Jan & Jan, 1994; Kageyama & Nakanishi, 1997; Lee, 1997). These genes are functionally categorized into pro-neuronal and pro-glial genes, and their spatiotemporal expression is regulated by several signaling pathways, including NOTCH, bone morphogenetic protein (BMP), fibroblast growth factor (FGF) signaling, among others (Aguirre *et al*, 2010; Gaiano & Fishell, 2002; Gross *et al*, 1996; Morrison *et al*, 2000; Qian *et al*, 1997; Zhou *et al*, 2000; Zirlinger *et al*, 2002). Loss-of-function studies in the retina have shown that deletion of pro-neuronal genes such as *Ascl1* (formerly *Mash1*) or *NeuroD* redirects progenitors toward glial fates at the expense of neurons (Lo *et al*, 1998; Morrow *et al*, 1999; Perez *et al*, 1999; Tomita *et al*, 1996). Similarly, double knockout of *Ascl1* and *Neurod4* (formerly *Math3*) induces glial commitment in regions like the tectum and cerebellum, where these progenitors normally give rise to neurons (Tomita *et al*, 2000). Conversely, pro-glial bHLH genes, including *Hes1*, *Hes5* and *Hesr2*, promote gliogenesis via interaction with the Notch signaling pathway (Furukawa *et al*, 2000; Gaiano & Fishell, 2002; Hojo *et al*, 2000; Morrison *et al*., 2000). Notch signaling is a highly conserved mechanism that contributes to the maintenance of neural stem cell identity, inhibition of neurogenesis, and promotion of glial differentiation (Gaiano & Fishell, 2002; Taylor *et al*, 2007). SoxE transcription factors are also essential for glial lineage progression, in part through their interactions with bHLH factors. Sox9 promotes the initial differentiation of neural stem cells into glia and later cooperates with Olig2 to induce Sox10 expression. Sox10, another SoxE family member, is required for the terminal differentiation of oligodendrocytes in conjunction with Olig1 (Dehm *et al*, 2024; Glasgow *et al*, 2017; Stolt *et al*, 2003; Stolt *et al*, 2002).

PTF1A is a bHLH transcriptional factor first identified as a key regulator of pancreatic cell fate (Kawaguchi *et al*, 2002). In the nervous system, PTF1A expression is normally confined to the cerebellum, brainstem, and spinal cord, and is not detected in the developing cortex. In these regions, *Ptf1a* directs the specification of GABAergic neurons (Glasgow *et al*, 2005; Hoshino *et al*, 2005; Iskusnykh *et al*, 2016). More recently, *Ptf1a* has been implicated in early sexual differentiation of the brain, due to its robust expression in neuroepithelial cells surrounding the third ventricle (Fujiyama *et al*, 2018). Furthermore, ectopic *Ptf1a* expression has been shown to reprogram both mouse and human fibroblasts into induced neural stem cells *in vitro*, capable of differentiating into neurons, astrocytes, and oligodendrocytes (Xiao *et al*, 2018). These findings suggest that *Ptf1a* may regulate cell fate decisions in a context-dependent manner across various brain regions.

In this study, we unexpectedly discovered that ectopic expression of *Ptf1a* strongly promotes the neurogenic-to-gliogenic switch in the embryonic mouse cortex.

## Results

### Virally ectopic expression of PTF1A efficiently directs gliogenesis in both the embryonic cortex and adult hippocampus

While ventral RGPs can generate glia at early embryonic stages, dorsal RGPs acquire gliogenic potential only at later stages, following the completion of neurogenesis (Kessaris *et al*., 2006; Kriegstein & Alvarez-Buylla, 2009; Lin *et al*., 2021). To examine whether *Ptf1a* can induce a cell fate switch in RGPs, we performed in utero retroviral injections (titer ∼10⁹ IFU) into the lateral ventricles of mouse embryos at embryonic days (E) 11, 13, and 15. The retrovirus ectopically expressed PTF1A under the control of the CAG promoter, which drives robust and stable gene expression. Control viruses expressed EGFP alone. Brain sections from postnatal day (P) 30 mice were immunostained for EGFP and the neuronal marker NEUN (Fig. 1A). In controls, approximately 81.9%, 70.6%, and 56.7% of EGFP+ cells in the cortex were NEUN+ at E11, E13, and E15, respectively—consistent with the progressive decline in neurogenic capacity during development. Strikingly, nearly all *Ptf1a*-expressing cells were NEUN- at P30, regardless of the injection time point (Fig. 1B, C). These cells were distributed broadly across the brain, with high densities in the piriform cortex, auditory cortex, amygdala, thalamus, and hippocampus (Suppl. Fig. 1A, B), aligning with known regions of normal glial development. Similarly, at postnatal stages (P3 and P5), *Ptf1a*- expressing cells were NEUN- and widely distributed across multiple brain regions, including the cortex, hippocampus, and striatum (Suppl. Fig. 2A–H). Importantly, these cells were negative for cleaved caspase-3, indicating that their absence of neuronal identity is not due to apoptosis (Suppl. Fig. 3).

**Fig. 1.**
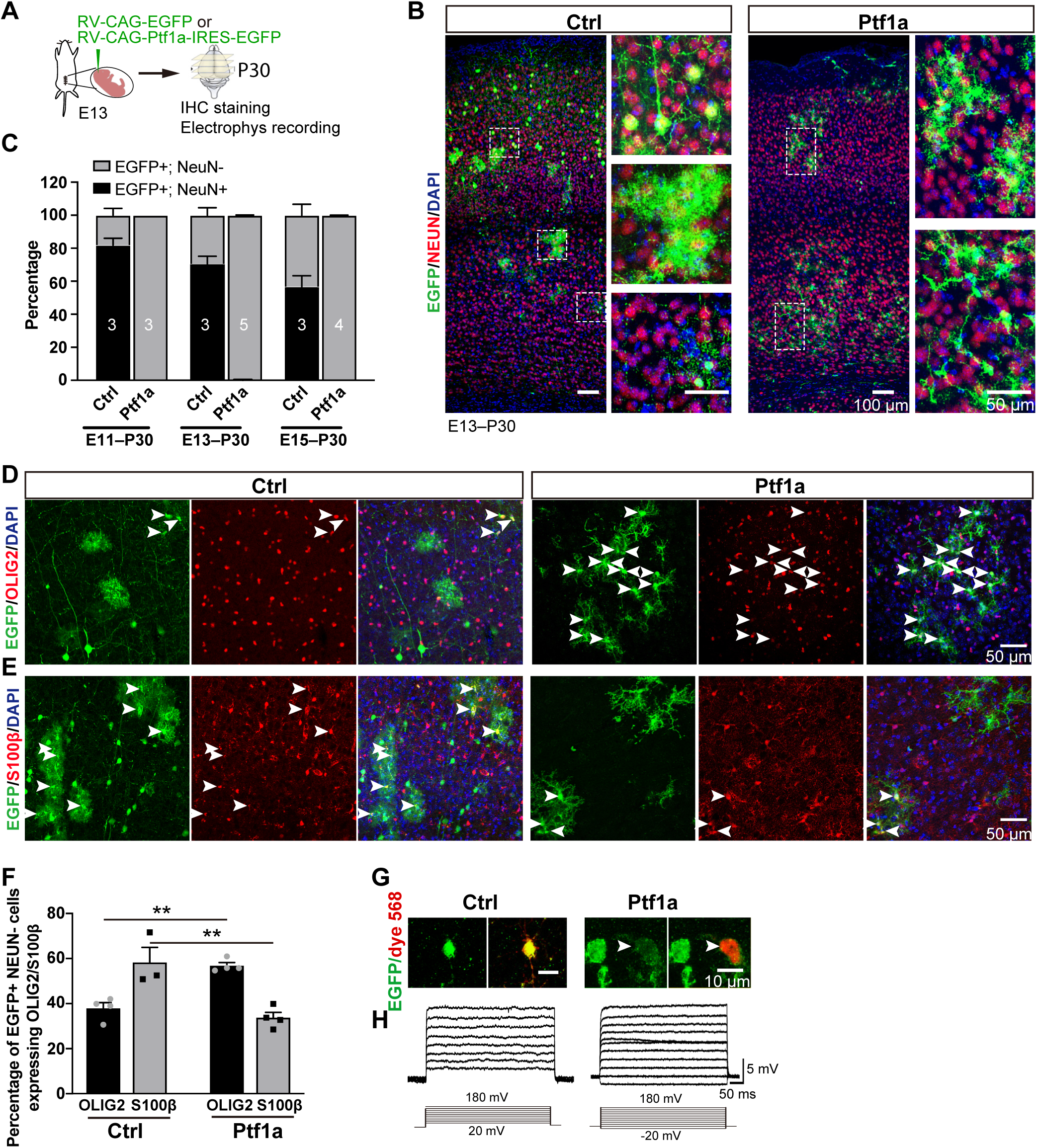
Viral expression of *Ptf1a* drives gliogenesis in the cortex. (A) Schematic of the experimental paradigm. (B) Representative images showing EGFP and NEUN immunostaining in cortices of P30 mice with retroviruses expressing EGFP (control) or *Ptf1a*-IRES-EGFP (*Ptf1a* mice) at E13. Right panel are magnified views of the boxed regions. (C) Quantification of the percentage of NEUN+ and NEUN- cells among total EGFP+ cells in the cortices of P30 control and *Ptf1a* mice that received *in utero* retrovirus injection at E11, E13 and E15, respectively. (D, E) Representative images showing colocalization of EGFP with OLIG2 (*D*) or S100β (*E*) in P30 cortices from control (Ctrl) or *Ptf1a* mice. Arrowheads indicate colocalized cells. (F) Quantification of the percentage of OLIG2+ or S100β+ cells among total EGFP+ cells in P30 control and *Ptf1a* cortices. **p < 0.01, two-way ANOVA with Bonferroni’s *post hoc* test. (G) Representative images of whole-cell patch clamp recordings of EGFP+ cells filled with Alexa Fluor 568 (arrowheads, red) in P30 cortices of control or *Ptf1a* mice. (H) Sample traces showing membrane potential changes in response to 20 pA incremental current injections (Ctrl: n = 8 cells; *Ptf1a*: n = 12 cells). No action potentials were observed. Data are presented as mean ± SEM.

To determine whether *Ptf1a*-expressing cells adopt glial identities, we stained P30 brain sections with markers for astrocytes (S100β, GFAP), oligodendrocyte lineage cells (OLIG2), and microglia (IBA-1). In the cortex, the majority of *Ptf1a*-expressing cells were OLIG2+ oligodendrocytes (56.9%), followed by S100β+ astrocytes (33.7%) and GFAP+ reactive astrocytes (17.8%) (Fig. 1D–F; Suppl. Fig. 4A, B). No EGFP⁺ cells were IBA-1+ microglia or GABAergic interneurons (Suppl. Fig. 4C, D). Electrophysiological recordings confirmed that these cells lacked action potentials, consistent with a glial identity (Fig. 1G, H). Notably, while astrocytes are typically the predominant glial subtype in the cortex (Fig. 1D, F; Suppl. Fig. 2H), *Ptf1a*-expressing RGPs were biased toward oligodendrocyte differentiation in both embryonic and adult stages (Fig. 1D, F; Suppl. Fig. 2A, C, G, H).

To determine whether this oligodendrocyte bias results from differential proliferation, we injected BrdU 1 hour before sacrifice at P5 to label proliferating cells. *Ptf1a*-expressing S100β+ astrocytes and OLIG2+ oligodendrocytes showed similar BrdU incorporation compared to neighboring non-EGFP cells (Suppl. Fig. 5A–D), suggesting that *Ptf1a* promotes fate specification rather than selectively enhancing OPC proliferation.

Previous studies using in utero electroporation of *Ptf1a* into dorsal RGPs reported induction of GABAergic neurons (Hoshino *et al*., 2005; Russ *et al*, 2015). To address this discrepancy, we compared the effects of *Ptf1a* expression using viral delivery versus electroporation. We electroporated CAG-*Ptf1a*-IRES-EGFP plasmids into dorsal telencephalon at E13. At P30, most *Ptf1a*-expressing cells were NEUN+ but negative for GABA, OLIG2, or S100β (Suppl. Fig. 6), suggesting a neuronal identity. Since electroporated plasmids dilute during cell division, this result supports the notion that stable and high *Ptf1a* expression is necessary for gliogenesis. To test this, we expressed *Ptf1a* using a weaker ubiquitin (Ubi) promoter, resulting in much lower PTF1A expression compared to those driven by the stronger CAG promoter (Suppl. Fig. 7A, B). While most Ubi-*Ptf1a*-expressing cells still became glia, the fraction of glial cells decreased as development progressed, and a notable number of neurons were observed (Suppl. Fig. 7C, D), further supporting a dose-dependent effect of *Ptf1a* on glial fate commitment.

We next asked whether *Ptf1a* could induce gliogenesis in adult hippocampal neural stem cells, which typically produce granule cells. Retroviruses expressing *Ptf1a* were stereotaxically injected into the subgranular zone of the dentate gyrus in ∼P56 mice (Suppl. Fig. 8A). Remarkably, adult hippocampal stem cells began generating glial cells instead of neurons, including ∼54.8% OLIG2+ oligodendrocytes and ∼25.6% S100β⁺ astrocytes (Suppl. Fig. 8B–E).

Together, these results demonstrate that PTF1A is a potent gliogenic factor capable of redirecting both embryonic and adult neural stem cells toward glial fates, with a preferential bias toward oligodendrocyte specification under stable, high-expression conditions.

### PTF1A-expressing cells differentiate into oligodendrocyte lineages

Oligodendrocyte lineage cells include oligodendrocyte progenitors, immature oligodendrocytes, and mature oligodendrocytes, all of which express OLIG2 and SOX10 (Fig. 2A) (Zhou *et al*., 2000). To determine the identity of the OLIG2-positive cells derived from *Ptf1a*-expressing RGPs, we analyzed P30 brain sections using markers of different oligodendrocyte developmental stages. Approximately 23.5% of *Ptf1a*-expressing cells in the cortex were positive for PDGFRα, 12.5% for O4, and 27.4% for CC1 (Fig. 2B–E). Notably, the fraction of OLIG2+ cells expressing CC1, a marker of mature oligodendrocytes, was comparable between *Ptf1a*-expressing and neighboring control (GFP-negative) glial cells (Fig. 2F, G). Surprisingly, immunohistochemistry revealed colocalization of OLIG2 and SOX10 in *Ptf1a*- expressing cells specifically within the fimbria, but not in other brain regions such as the cortex or hippocampus (Fig. 2H–J). Since SOX10 is critical for the differentiation of myelinating oligodendrocytes (Stolt *et al*., 2002), these results suggest that most oligodendrocytes derived from *Ptf1a*-expressed RGPs progress to SOX10-negative, non-myelinating oligodendrocytes.

**Fig. 2.**
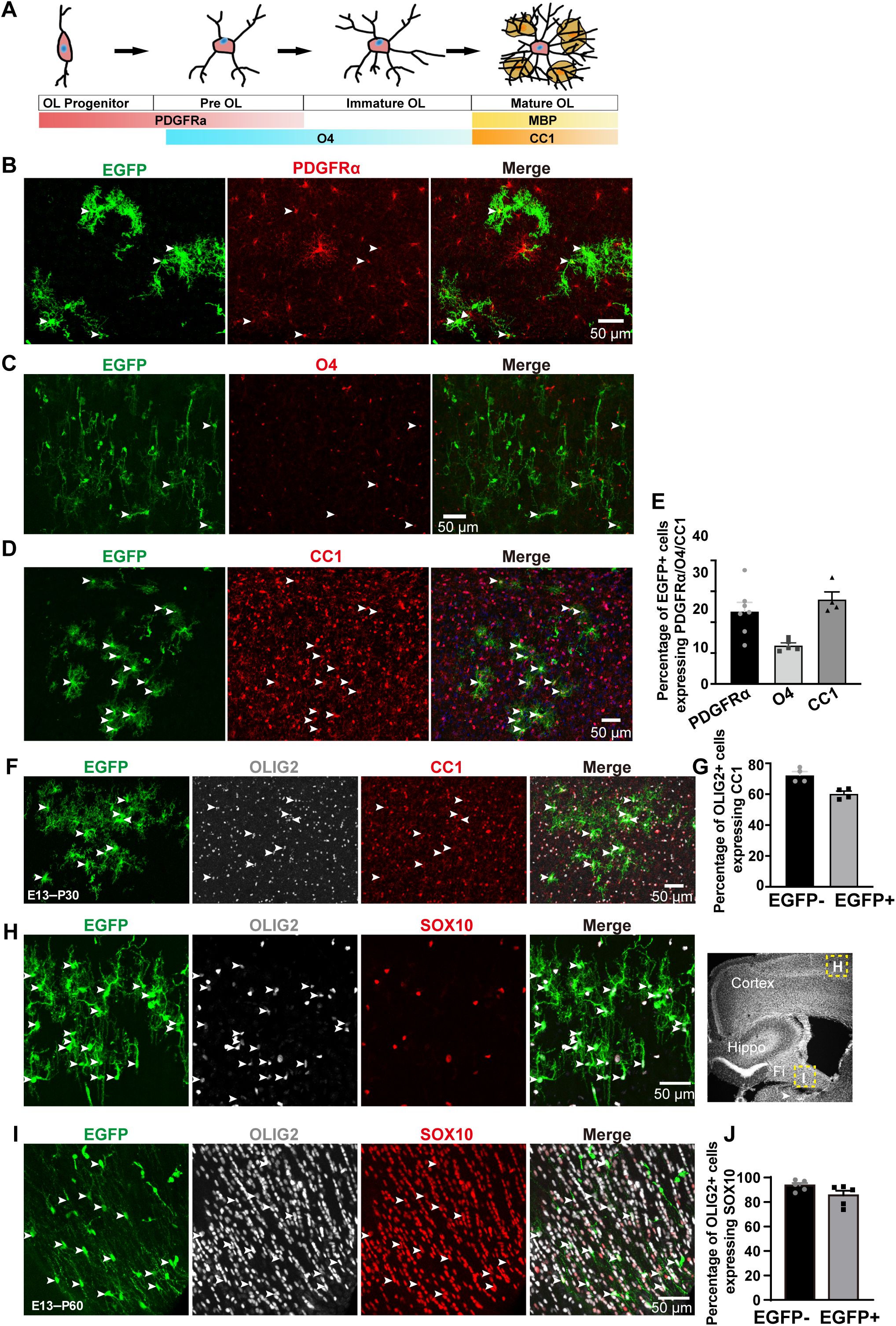
PTF1A-expressing RGPs are capable of generating oligodendrocyte lineages. (A) Schematic showing marker expression across oligodendrocyte developmental stages. (B-D) Representative images showing EGFP colocalization with PDGFRα (*B*), O4 (*C*), and CC1 (*D*) in P30 *Ptf1a* cortices. Arrowheads indicate colocalized cells. (E) Quantification of the percentage of EGFP+ cells expressing PDGFRα, O4 or CC1 cells. (F) Representative images showing colocalization of EGFP, CC1 and OLIG2 in a P30 *Ptf1a* cortex. Arrowheads indicate triple-labeled. (G) Quantification of the percentage of OLIG2+ cells co-expressing CC1 in EGFP- or EGFP+ populations. Data are represented as mean ± SEM. (H, I) Representative images showing EGFP+ OLIG2+ cells (arrowheads) negative for SOX10 in the cortex (*H*), but positive for SOX10 (arrowheads) in the fimbria (*I*). Right panel in (*H*) showing corresponding regions of images in (*H*) and (*I*). (J) Quantification of the percentage of OLIG2+ cells co-expressing SOX10 for EGFP- or EGFP+ cells in the P60 *Ptf1a* fimbria. Data are presented as mean ± SEM.

### *Ptf1a-*expressing RGPs exhibit normal progenitor behaviors during fate transition

During development, RGPs can directly give rise to oligodendrocyte progenitors that migrate widely throughout the brain, but most eventually retract their ventricular endfeet and differentiate into astrocytes at the end of embryogenesis in mammals (Kriegstein & Alvarez-Buylla, 2009; Laywell *et al*, 2000; Shen *et al*, 2021). We investigated whether *Ptf1a*-expressing RGPs follow this behavior. In contrast to the majority (>80%) of control RGPs that retained ventricular attachment at E16.5, *Ptf1a*- expressing RGPs exhibited progressive detachment over time: 67.3%, 48.7%, and 25.4% retained endfeet at 24, 48, and 72 hours after in utero retroviral injection at E13.5, respectively (Fig. 3A, B). We observed a significant increase in the proportion of PAX6+ EGFP+ cells in *Ptf1a*-expressing cortices compared to controls (Fig. 3C, D), with many delocalized outside the ventricular zone (VZ), likely due to endfoot retraction (Fig. 3C, E). Consistent with this, more EGFP+ KI67+ cells were found in extra-VZ regions in *Ptf1a* mice compared to controls, where most proliferating cells remained in the VZ (Fig. 3F, G). No differences were observed in the proportion of PAX6+ EGFP+ cells incorporating BrdU or the fraction of EGFP+ BrdU+ cells lacking KI67 expression (Suppl. Fig. 9A–E), suggesting unchanged cell cycle length.

**Fig. 3.**
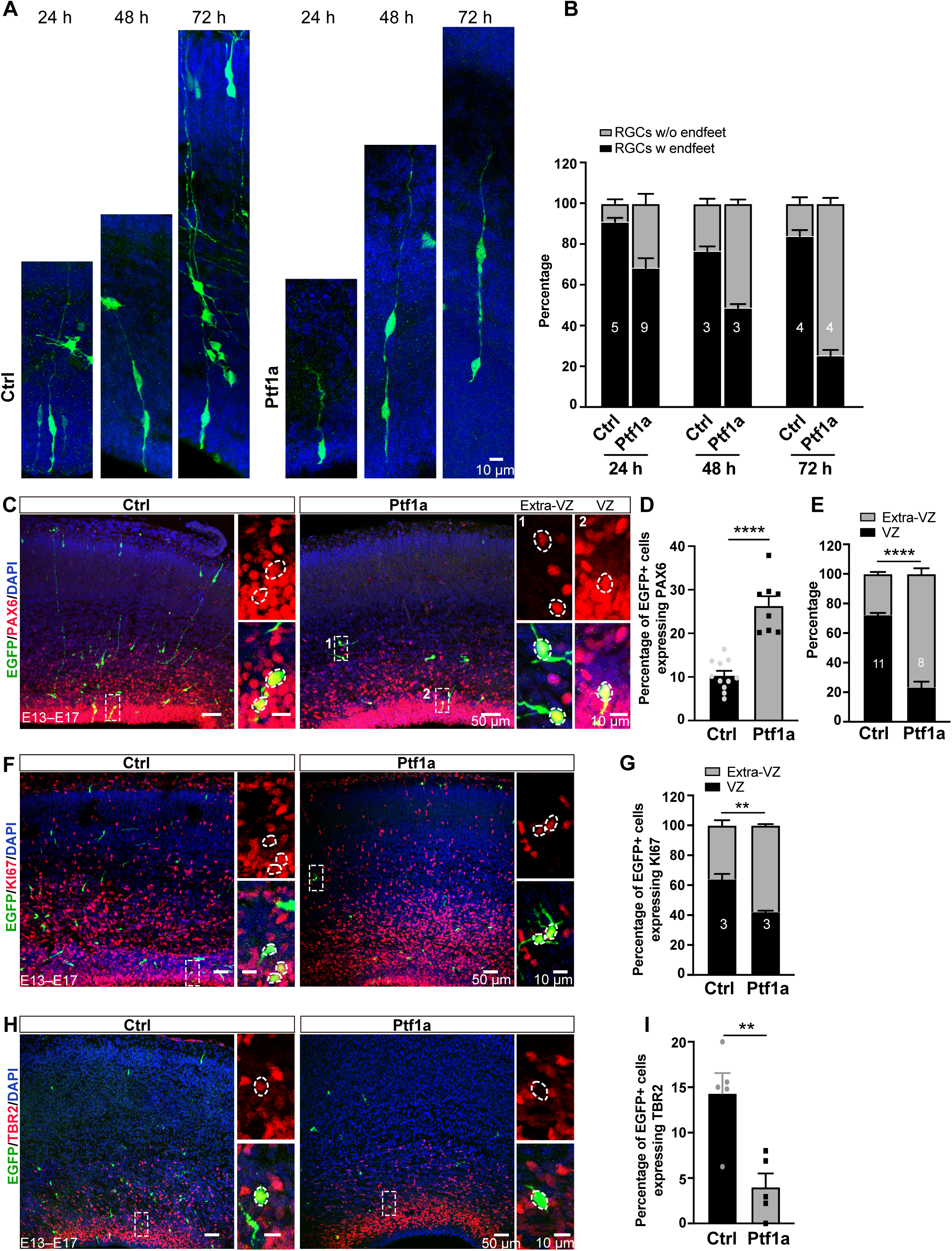
PTF1A-expressing RGPs recapitulate endfoot detachment normally from the basement membrane during embryonic development. (A) Representative images of EGFP-labeled RGP morphology at 24, 48, and 72 hours post-injection of retroviruses expressing EGFP or Ptf1a-IRES-EGFP at E13. (B) Quantification of the percentage of EGFP+ RGPs with or without endfoot attachments to the ventricular membrane. Note that RGPs in the cortices from *Ptf1a* embryos progressively retracted their endfeet from the ventricle membrane. (C-E) Representative images showing EGFP and PAX6 staining (*C*), quantification of the percentage of EGFP+ cells expressing PAX6 (*D*), and regional distribution of EGFP+ PAX6+ cells in the ventricular zone (VZ) versus extra-VZ regions (*E*) in E17 cortices. Right panels show magnified boxed areas. ****p < 0.0001, Student’s t-test. (F, G) Representative images of EGFP and KI67 staining (F), and quantification of EGFP+ KI67+ cells in the VZ and extra-VZ regions (G). **p < 0.01, Student’s t-test. (H, I) Representative images of EGFP and TBR2 staining (*H*) and quantification of EGFP+ cells expressing TBR2 (*I*). **p < 0.01, Student’s t-test. Data are presented as mean ± SEM.

We next assessed whether the initial response to *Ptf1a* expression biases RGPs toward glial fate. Brain sections were stained for TBR2, a marker of intermediate progenitor cells (IPCs), and doublecortin (DCX), an immature neuronal marker, at 1.5 and 4 days post retroviral infection, respectively. The percentage of TBR2+ cells, which typically give rise to neurons, was significantly reduced 1.5 days after *Ptf1a* retrovirus infection compared to controls (EGFP-only). Likewise, only 1.4% of *Ptf1a*-expressing cells were DCX+ at 4 days post-infection, compared to 82.7% in controls (Suppl. Fig. 9F, G). These results support that *Ptf1a* expression leads to early glial fate specification in RGPs.

### RNAseq reveals activation of gliogenic gene expression programs in *Ptf1a*- expressing RGPs

To uncover the molecular mechanisms underlying *Ptf1a*-mediated cell fate determination, we performed Smart-seq2-based transcriptomic profiling of individually isolated EGFP+ RGPs expressing either *Ptf1a* or EGFP only (control). 40 Cells were collected for a tube using a dual patch-clamp system, allowing simultaneous isolation and aspiration (Fig. 4A). Gene expression correlation analysis showed high intra-group consistency and low inter-group similarity, confirming robust data quality (Suppl. Fig. 9H). We identified 2,109 differentially expressed genes (DEGs) in *Ptf1a*-expressing RGPs relative to controls (FC ≥ 2, Padj < 0.01), with 929 upregulated and 1,180 downregulated genes (Fig. 4B, C; Suppl. Tables 1–2).

**Fig. 4.**
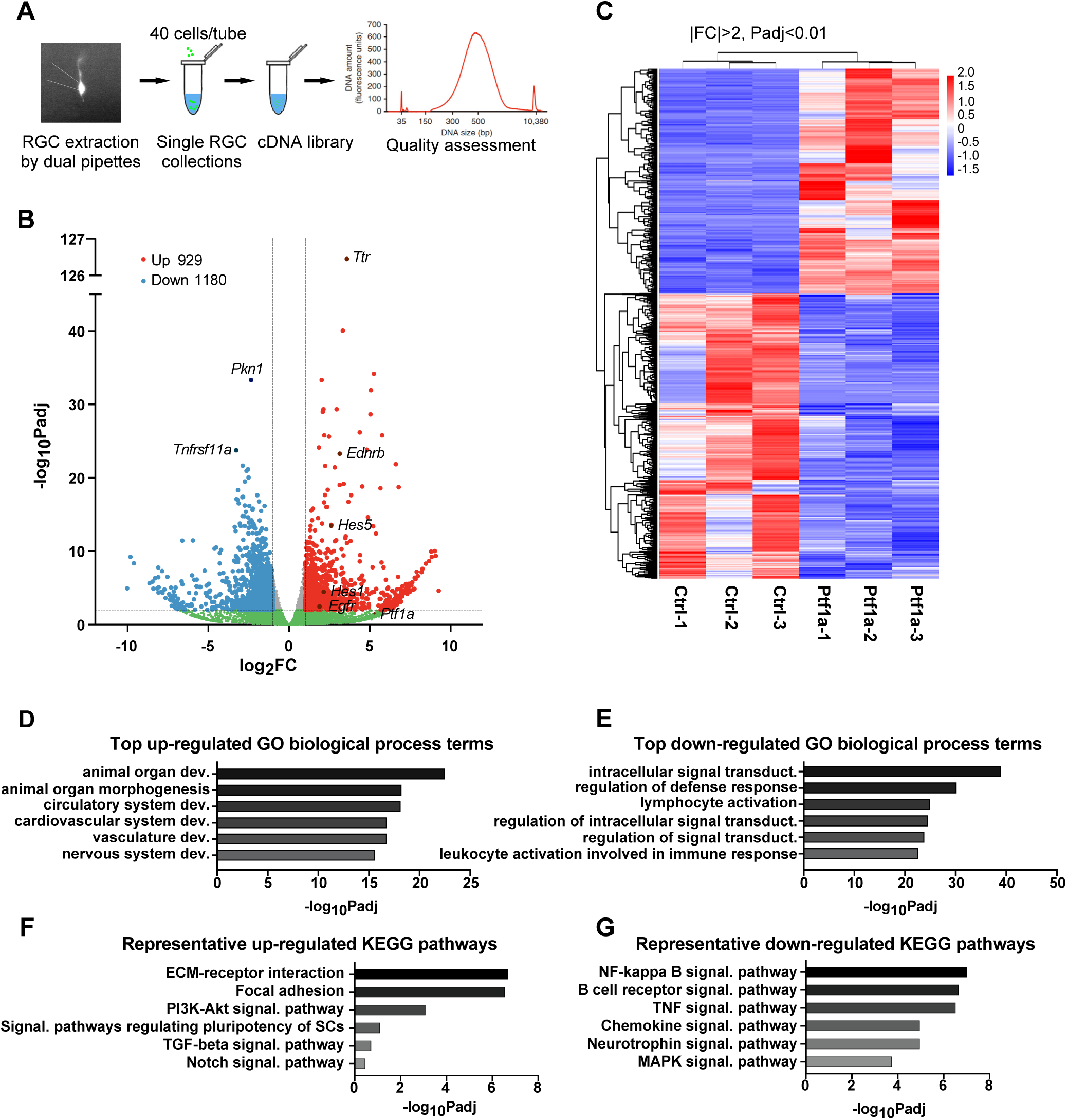
RNAseq reveals differentially expressed genes (DEGs) in *Ptf1a*-expressing RGPs primarily involved in neural development. (A) Schematic of dual-patch RGP acquisition and RNS sequencing. (B) Volcano plot comparing gene expression in PTF1A-expressing versus control RGPs. DEGs were defined by padj <0.01 and |log2 FC|>1. Red: upregulated; blue: downregulated; gray and green: non-significant. (C) Heatmap of highly significant DEGs (padj <0.01 and |log2 FC|>1). (D, E) Top gene ontology (GO) biological process (BP) terms enriched inr up-regulated (D) and down-regulated (*E*) DEGs. (F, G) Significantly enriched KEGG pathways among upregulated (*F*) and downregulated (*G*) DEGs.

Gene set enrichment analysis (GSEA) of upregulated DEGs revealed enrichment in developmental processes such as morphogenesis, organ development, nervous system development, and cell adhesion, as well as stem cell proliferation and gliogenesis (Fig. 4D, F; Suppl. Tables 3–4). Downregulated genes were associated with intracellular signaling, immune response, and neuronal differentiation/survival (Fig. 4E, G; Suppl. Tables 5–6). Many pro-glial genes were significantly upregulated, including core Notch signaling components and targets (*Notch1*, *Notch2*, *Hes1*, *Hes5*, *Hey1*). Specifically, genes associated with pro-astrocyte differentiation (*Bmp2*, *MT3*, *Hes1*), pro-oligodendrocyte identity (*Erbb2*, *Nkx2.1*, *Plp1*, *Olig1*, *Ednrb*, *Nog*), and general gliogenesis (*Sox9*, *Egfr*, *Sox8*, *Vtn*, *Gap43*, *Fgf43*, *Clcf1*) were all upregulated. In contrast, many pro-neuronal differentiation genes were downregulated (*Jade1, Bhlhe22, Gpc2, Brinp2, Foxa1, Adra2c, Lin28a, Rgs14, Actr3, Etv5, Adra2b, Psen1, Bcl6, Cebpb, Shc1*) (Fig. 5A, B; Suppl. Fig. 10A, B).

**Fig. 5.**
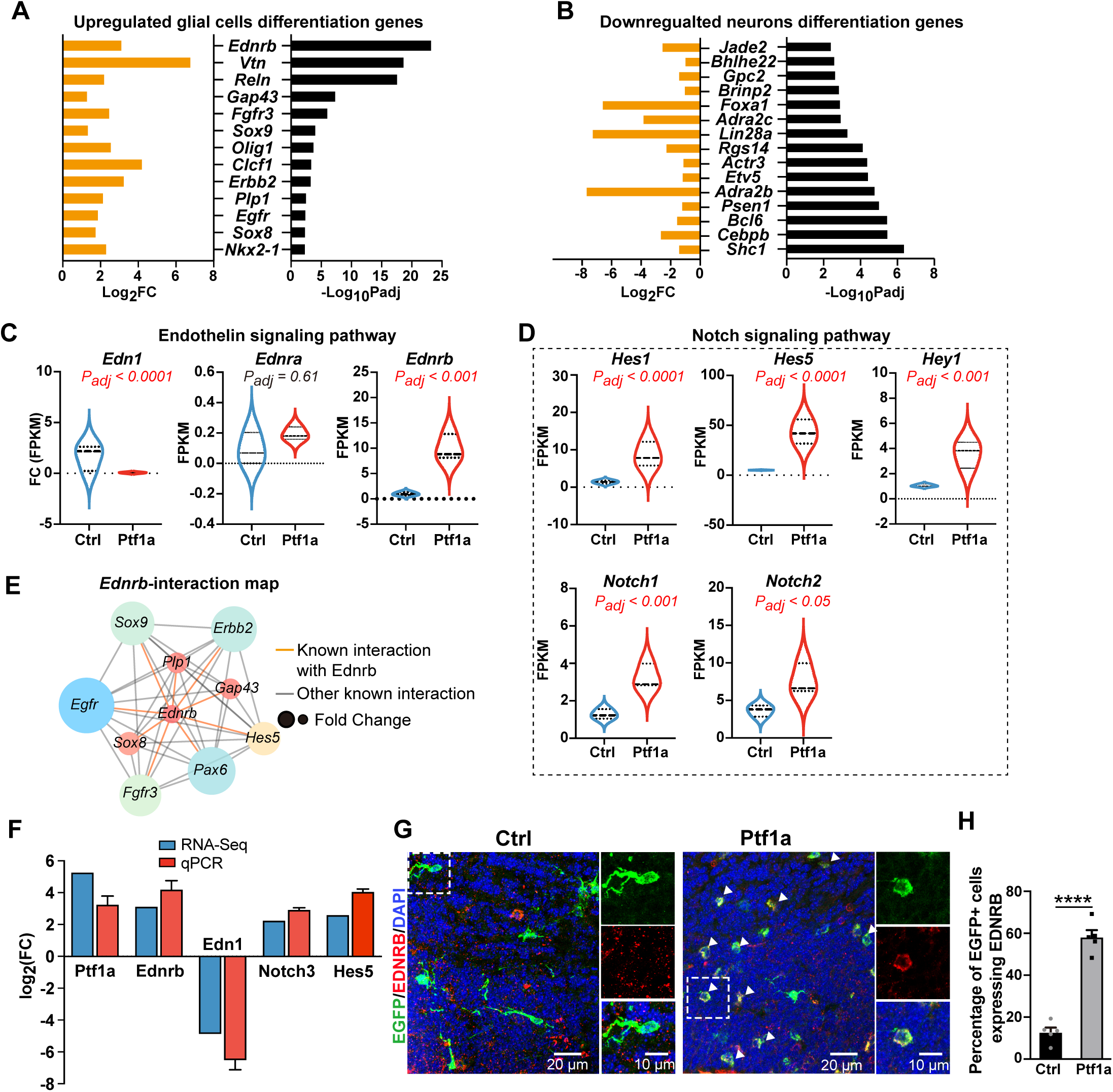
*Ednrb* is significantly upregulated in *Ptf1a-*expressing RGPs. (A, B) Bar graphs showing fold changes of upregulated glial cell differentiation genes (A) and downregulated neuronal differentiation genes (*B*) in *Ptf1a* versus control RGPs. (C) Violin plots showing expression levels of *Edn1*, *Ednra* and *Ednrb* in control and *Ptf1a* RGPs. Each plot shows transcript frequency distribution with max, average, and min values indicated. (D) Violin plots of DEGs involved in the Notch signaling pathway. (E) Interaction map of genes directly interacting with *Ednrb*. (F) RT-qPCR validation of five selected DEGs in E15 RGPs. Bar graph shows fold change normalized to control for RT-qPCR or RNA-seq. (G) Representative images showing EGFP and EDNRB immunostaining in E17 cortices. (H) Quantification of the percentage of EDNRB+ cells among EGFP⁺ cells. ****p < 0.0001, Student’s t test. Data are presented as mean ± SEM.

Importantly, *Ednrb* was upregulated more than eightfold in *Ptf1a*-expressing RGPs, while *Ednra* expression remained unchanged (Fig. 5C). *Ednrb* is known to mediate postnatal oligodendrocyte development and reactive astrogliosis (Adams *et al*, 2020; Gadea *et al*, 2008; Hammond *et al*, 2015; Schinelli, 2006; Swire *et al*, 2019). Of its ligands, *Edn1* expression was significantly reduced in *Ptf1a*-expressing cells, whereas *Edn3* was undetectable (Fig. 5F; Suppl. Table 1). Network analysis highlighted *Ednrb* as a central hub among pro-glial DEGs (Fig. 5E). qPCR validated several DEGs, including *Ptf1a, Ednrb, Edn1, Notch2*, and *Hes5* (Fig. 5F). Immunohistochemistry confirmed RNA-seq findings, showing elevated expression of *Ednrb* and *Sox9* in *Ptf1a*-expressing cells at E15, while *Nfia, Olig1,* and *Sox10* remained undetectable similar to control levels (Fig. 5G, H; Suppl. Fig. 10C–F).

### *Ednrb* knockdown attenuates the glial competency of *Ptf1a*-expressing RGPs

We next investigated whether *Ednrb* functions as a downstream target of *Ptf1a* in mediating the cell fate switch. To test this, we overexpressed (OE) EDNRB in RGPs via retroviral delivery at E13 and analyzed the fate of EGFP+ cells in the P30 cortex. We observed a significant increase in the proportion of EGFP+ NEUN- cells (∼60.47%) in the *Ednrb* OE group compared to control (∼17.6%) (Suppl. Fig 11A, B). Among these, approximately 56.09% were OLIG2+ oligodendrocytes and 43.2% were S100β+ astrocytes (Suppl. Fig. 11C-E).

To determine whether *Ptf1a* promotes gliogenesis via EDNRB signaling, we co-expressed a potent *Ednrb* short hairpin RNA (shRNA-1) with *Ptf1a* in RGPs using a retroviral vector (Suppl. Fig. 11F). In wild-type RGPs, knockdown of *Ednrb* had no significant effect on in wildtype RGPs with shRNA had no effect on the neurons-to-glia ratio (fraction of EGFP+ NEUN+ cells: scramble, ∼82.8%; shEdnrb, ∼81.0%; Suppl. Fig. 11G, H). However, knockdown of *Ednrb* in *Ptf1a*-expressing RGPs significantly increased the fraction of EGFP+ NEUN- cells compared to *Ptf1a* with scramble shRNA (Fig. 6A, B), indicating a partial rescue of neurogenesis.

**Fig. 6.**
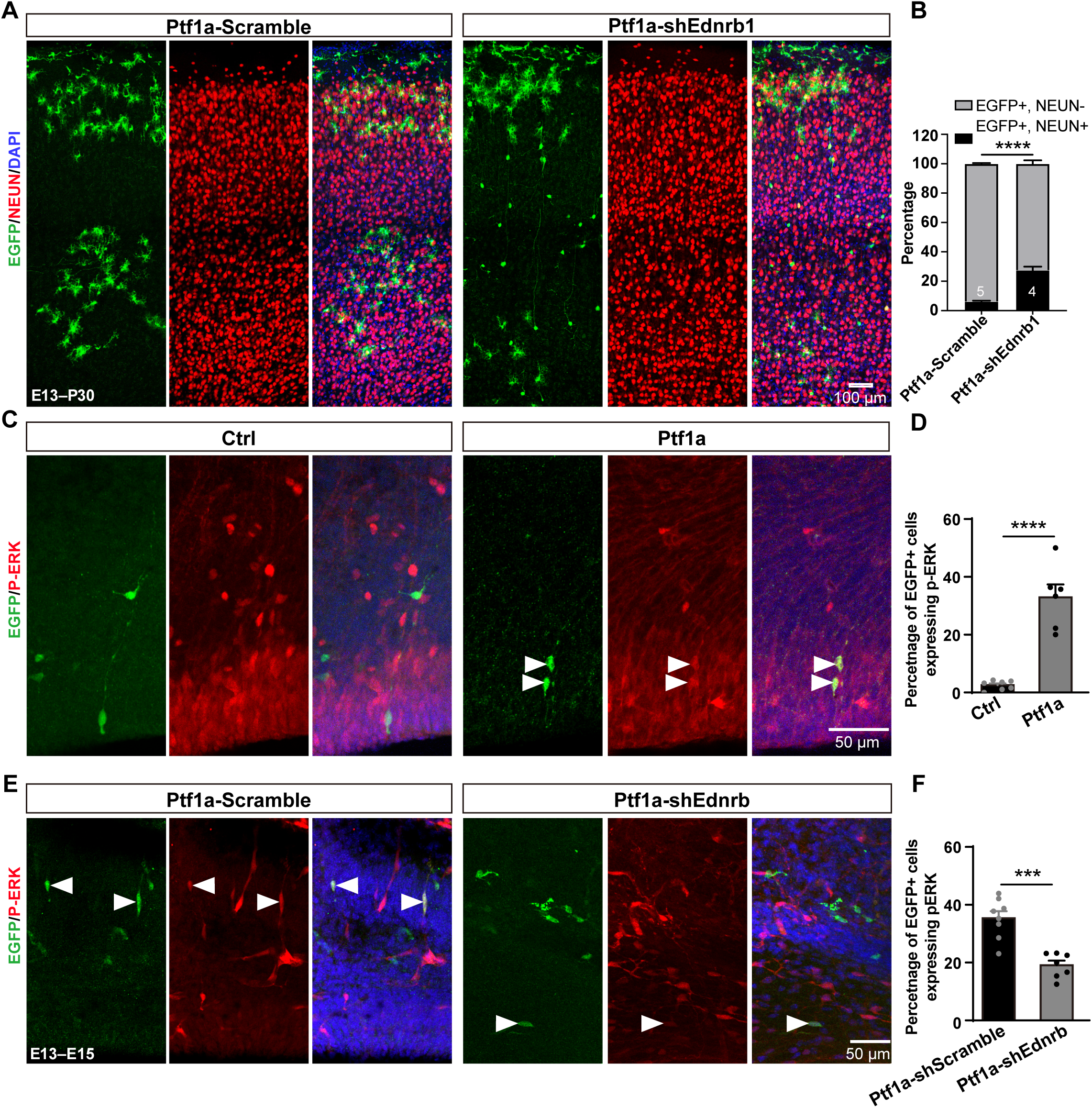
Knockdown of *Ednrb* partially rescues neurogenesis of *Ptf1a*-expressing RGPs. (A, B) Representative images of EGFP and NEUN staining in cortical *Ptf1a* cells expressing scramble or *Ednrb* shRNA. (B) Quantification of NEUN⁺ and NEUN⁻ cells among EGFP⁺ populations. (C, D) Representative images (*C*) and quantification (*D*) of EGFP and p-ERK staining in E15 control and *Ptf1a* cortices. Arrowheads indicate EGFP+ p-ERK+ cells. ****p < 0.0001, Student’s t-test. (E, F) Representative images (*E*) and quantification (*F*) of EGFP and p-ERK staining in E15 Ptf1a cells expressing scramble or Ednrb shRNA. N indicates the number of mice. ***p < 0.001, ****p < 0.0001, Student’s t-test. Data are presented as mean ± SEM.

Given that ERK signaling is essential for gliogenesis in the embryonic cortex and can be directly activated by EDNRB (Aquilla *et al*, 1996; Gao *et al*., 2025; Li *et al*., 2012), we examined whether EDNRB exerted its effect through ERK activation. Immunohistochemistry revealed approximately 33% of *Ptf1a*-expressing cells were positive for phosphorylated-ERK (p-ERK) at E15, compared to less than 5% in controls (Fig. 6C, D). Knockdown of *Ednrb* significantly reduced the proportion of *Ptf1a* cells with p-ERK (Fig. 6E, F), suggesting that EDNRB-ERK signaling pathway plays an important role in glial fate determination of *Ptf1a-*expressing RGPs.

## Discussion

### Dosage-dependent effect of PTF1A on the gliogenic switch

This study demonstrates for the first time that ectopic expression of the bHLH factor PTF1A, driven by a strong CAG promoter via retroviruses, efficiently induces gliogenesis in both the embryonic cortex and adult hippocampus. In contrast, retroviral expression driven by a weak Ubi promoter did not achieve the same efficiency, likely due to lower levels of PTF1A expression. Similarly, electroporation of *Ptf1a* plasmids had no impact on RGP fate, likely because these plasmids are not stably integrated into the genome and are thus diluted during RGP divisions. These results suggest that the gliogenic switch is driven by PTF1A in a dosage-dependent manner.

Unlike previous studies reporting GABAergic neurons following *Ptf1a*-expressing plasmid electroporation in the embryonic cortex, we found that progeny of electroporated RGPs differentiated into excitatory neurons in the P30 cortex. This raises the possibility that these cells may transiently express GABA during early development. The glial competency of *Ptf1a* appears specific to cortical RGPs in the embryonic forebrain, as *Ptf1a* is known to specify interneurons in other CNS regions such as the spinal cord and the cerebellum (Glasgow *et al*., 2005; Hoshino *et al*., 2005).

### *Ptf1a* directs RGPs to preferentially give rise to oligodendrocytes

Under normal conditions, cortical RGPs give rise to both astrocytes and oligodendrocytes near the end of embryonic development. Consistently, *Ptf1a*- expressing RGPs generated both glial subtypes, suggesting intact glial competency. Some *Ptf1a*-derived glia were not accounted for by OLIG2+ or S100β+ markers, possibly due to astrocyte heterogeneity (e.g., S100β+ vs. S100β- populations). Despite discrepancies in reported astrocyte-to-oligodendrocyte ratios across studies by employing different sample preparations and statistical methods (Carlo & Stevens, 2013; Herculano-Houzel, 2014; Rakic, 2008; Rockel *et al*, 1980), our quantification of EGFP-only glia showed that astrocytes outnumbered oligodendrocytes in controls. However, *Ptf1a*-expressing RGPs produced more oligodendrocytes than astrocytes, indicating a bias toward oligodendrocyte lineage. These oligodendrocytes spanned multiple developmental stages (OPC, immature, and mature CC1+ cells), similar to those derived from normal RGPs. Unexpectedly, most of them did not express SOX10, a key regulator of terminal oligodendrocyte differentiation, suggesting that they are arrested at a SOX10-negative, non-myelinating stage. While this study focused on *in vivo* embryonic gliogenesis, any future attempts to generate myelinating oligodendrocytes *in vitro* using *Ptf1a* will need to avoid its sustained expression.

### PTF1A-induced gliogenic switch is partially through EDNRB signaling

We observed that *Ptf1a*-expressing RGPs retracted their endfeet from the ventricular surface—behavior reminiscent of endogenous RGPs transitioning from neurogenesis to gliogenesis. However, since endfoot attachment is not essential for fate determination (Haubst *et al*, 2006), *Ptf1a* likely promotes gliogenesis through downstream gene expression programs rather than physical changes alone.

Consistent with this, we found that Notch pathway components—including *notch1*, *notch 2, hes1, hes5,* and *hey1*—were upregulated in *Ptf1a*-expressing RGPs. Although PTF1A can interact with RBPJ and HES1 (Ahnfelt-Ronne *et al*, 2012; Hori *et al*, 2008; Lelievre *et al*, 2011), disrupting the PTF1A-RBPJ-Notch tripartite complex by point mutation of PTF1A (W298A) did not impair its gliogenic function (Suppl. Fig. 12), indicating that PTF1A may activate the Notch pathway through a non-canonical, indirect mechanism.

Our transcriptomic, qPCR, and immunohistochemical analyses showed that *Ednrb*, encoding a GPCR known to enhance Notch signaling (Adams *et al*., 2020; Hammond *et al*, 2014), was upregulated by >8-fold in *Ptf1a*-expressing RGPs. Overexpression of *Ednrb* alone modestly promoted gliogenesis, while its knockdown partially suppressed *Ptf1a*-induced gliogenesis, suggesting that EDNRB contributes significantly to this cell fate switch. The incomplete suppression by EDNRB knockdown implies that additional pro-glial effectors are likely involved in downstream of *Ptf1a*. Likely, a cohort of the pro-glial molecules work together to fully mediate *Ptf1a*-induced gliogenesis.

The source of endothelin, the EDNRB ligand, remains a question. ET-1, but not ET-2 or ET-3, was enriched in embryonic RGPs, consistent with its established role in postnatal SVZ gliogenesis (Adams *et al*., 2020; Gadea *et al*., 2008). Although ET-1 was downregulated in *Ptf1a*-expressing cells, it may be secreted by adjacent non-*Ptf1a* RGPs or ependymal cells and act in a non-cell-autonomous manner, similar to the role of ET-1 in SVZ OPC proliferation and CT-1 in gliogenesis (Adams *et al*., 2020; Barnabe-Heider *et al*., 2005).

### Limitations of study

*Ptf1a* is not expressed in the normal embryonic cortex, and its knockdown had no effect on RGP fate (Suppl. Fig. 13), raising the possibility that ectopic Ptf1a activates non- physiological gene programs. Although EDNRB and Notch components were upregulated in *Ptf1a*-expressing RGPs, the mechanistic relationship among these factors remains unresolved.

## METHODS

### Animals

Animal care and use strictly followed institutional guidelines and governmental regulations. All animal studies and experimental procedures were approved by the institutional animal care and use committee (IACUC# 20250407001) of the animal facility at ShanghaiTech University. All ICR mice were purchased from Shanghai Jihui Laboratory Animal Care Co., Ltd. Animals were housed at controlled temperature (22- 25°C) on a 12-hours reverse light/dark cycle and were supplied with standard chow diet and ad libitum drinking water.

### Plasmids and viral production

*Ptf1a* and *Ednrb* cDNAs were cloned from a cDNA library of the adult mouse brain. For generation of retroviral plasmids, *Ptf1a* and *Ednrb* cDNA were subcloned into a CAG-EGFP vector (addgene #: 16664) between the two restriction enzyme sites BamHI and AgeI along with a fragment of IRES using In-Fusion cloning (Clontech). For generation of an Ubi-promoter-driven viral vector, *Ptf1a* cDNA was inserted into an Ubi-EGFP vector between the two restriction enzyme sites BamHI and AgeI along with a fragment of IRES using In-Fusion cloning (Clontech). For generation of the electroporation plasmid, *Ptf1a* cDNA was inserted into the vector of CAG-IRES-EGFP vector at the restriction enzyme site XhoI using In-Fusion cloning.

The target shRNA sequences used were as follow: shEdnrb-1, 5’-GCGGTATGCAGATTGCTTTGA-3’; shEdnrb-2, 5’-TCTAAGTATTGACAGATAT-3’; shEdnrb-3, 5’-AATAAATACAGCTCGTCTT-3’; scrambled shRNA, 5’-CCTAAGGTTAAGTCGCCCTCG-3’; shPtf1a-1, 5’-GGACGAGCAAGCAGAAGTAGA-3’; shPtf1a-2, 5’- GCTACACGAATACTGCTACCG-3’; shPtf1a-3, 5’- GGATTATGGTCTCCCTCCTCT-3’; shPtf1a-4, 5’- CAGTAAATCTTTCGACAACAT-3’. The sequences containing shRNA and miR30a were synthesized. shRNA-miR30a sequences were first inserted into the AAV-CAG-GFP vector (addgene #: 28014) at the restriction enzyme site EcoRI. Then Ptf1a-P2A was subcloned to the vector of AAV-CAG-GFP-shRNA-miR30a (28014-shRNA- miR30a) between the BamHI and AgeI sites using In-Fusion cloning. The segment containing Ptf1a-P2A-EGFP-shRNA-miR30a-WPRE was subcloned and inserted into the vector of CAG-EGFP between the BamHI and XhoI sites. All plasmids were confirmed by sequencing.

Viral particles of retroviruses were packaged in co-transfected HEK293T cells (at the density of >80%) with two helper plasmids Gag/Pol and VSVG using EZtrans transfection reagent (Life-iLab). Cell supernatants were collected 48 hours after the transfection, followed by filtration, and then concentrated through an ultracentrifuge (Beckman). The titer of retrovirus was determined via serial dilution of cell infection to achieve ∼10^9^ infectious units (IFU) per mL.

### In utero retrovirus injection and electroporation

Timed pregnant ICR mice at embryonic day (E) 11–15 were anesthetized with isoflurane and positioned on the heating pad for surgery as previously described (He *et al*, 2015; Xie *et al*, 2021). 1μL retroviral solution (∼10^9^ FU/mL) or DNA plasmid solution (2 μg/μL) mixed with fast green (Sigma) was injected into the lateral ventricle of the embryos with a sharpened glass micropipette (Drummond Scientific). For *in utero* electroporation, two 9-mm electrode paddles (BTX, ECM830) were used to deliver five 50 ms pulses of 50 mV with a 950 ms interval across the uterus to the head of the embryo. After completion of electroporation, the uterus was replaced back into the abdominal cavity and the wound was sutured. The surgery mice were placed into the warm ventilation chamber until they were recovered for showing physiological movements.

### Stereotaxic viral injection

For stereotaxic injection of retroviruses, ∼2-month-old mice were deeply anesthetized with isoflurane and were then fixed in a stereotaxic apparatus (stoelting Co.). CAG- Ptf1a-IRES-EGFP or CAG-EGFP retroviruses were injection into the subgranular zone of the mouse hippocampus at the coordinates of 2 mm posterior to the bregma, 1.6 mm lateral to the midline and 2.5 mm from the surface. Four weeks after retrovirus injection, mice were sacrificed for immunostaining.

### Brain sections, immunostaining and confocal imaging

Mice were deeply anesthetized with isoflurane and were transcardially perfused with 4% paraformaldehyde (PFA, Sigma) in ice-cold PBS (pH 7.4). Brains were removed and post-fixed with 4% PFA overnight at 4 °C, followed by serial coronal sections (60 μm) using a vibratome (Leica Microsystems). For cryosectioning, brains were dehydrated with 15% sucrose, followed by 30% sucrose, then embedded in OCT medium (Leica) and sectioned to 14 μm coronal sections using a cryostat (Leica CM1950).

For immunohistochemistry, brain sections were permeabilize with 0.3% triton solution in PBS for 10 mins at room temperature, followed by incubation in blocking solution (10% normal goat serum, 0.4% Triton X-100, 1% glycine and 3% BSA in 0.1% PBS) for 2 hours. Then brain sections were incubated for overnight at 4°C. The primary antibodies were used in this study as follows: chicken anti GFP (Aveslabs GFP-1020, 1:1000), rabbit anti NEUN (Abcam ab177487, 1:1000), mouse anti NEUN (Abcam ab104224, 1:1000), rabbit anti OLIG2 (Millipore ab9610, 1:500), rabbit anti S100β (Abcam ab41548, 1:500), rabbit anti GABA (Sigma A2052, 1:500), rabbit anti IBA-1 (Abcam ab5076, 1:500), rat anti BrdU (ABD Serotec OBT0030, 1:500), rabbit anti GFAP (Abcam ab7260, 1:500), Rat anti SOX10 (R&D systems MAB2864, 1:500), Rabbit anti SOX9 (Abcam ab185966, 1:500), Rabbit anti NFIA (Abcam, ab228897, 1:500), Rabbit anti OLIG1 (Cell Signaling 15848T, 1:500), rabbit anti PDGFRα (Cell signaling 3174s, 1:250), Rabbit anti p-ERK (Cell Signaling 4370T, 1:500), Mouse anti O4 (Millipore MAB345, 1:50), Rabbit anti CC1 (Millipore OP80, 1:500), rabbit anti PAX6 (Biolegend 901302, 1:500), Rabbit anti KI67 (Cell Signaling 9449T, 1:500), Rabbit anti EDNRB (Abcam ab117529, 1:500). The corresponding secondary antibodies were used as follows: goat anti-chicken Alexa-488 (Invitrogen A11039, 1:1000), goat anti-rat Alexa-546 (Invitrogen A11081, 1:1000), goat anti-rabbit Alexa- 546 (Invitrogen A11010, 1:1000), goat anti-mouse Alexa-546 (Invitrogen A11003, 1:1000), goat anti-rat Alexa-647 (Invitrogen A21247, 1:1000), goat anti-rabbit Alexa- 647 (Invitrogen A21244, 1:1000), goat anti-mouse Alexa-647 (Invitrogen A21235, 1:1000). DAPI (Roche) was added for counterstaining and sections were mounted with a coverslip.

Images were taken using the following laser scanning confocal microscopes: Leica SP8 STED, Zeiss LSM710, Zeiss LSM980 Airyscan2. Images were analyzed using ImageJ (NIH)-Fuji and Photoshop. For quantification of the fraction of NEUN+ or different glial subtypes in EGFP+ glial cells, cell number in the whole cortex of one hemisphere of coronal section was counted.

### BrdU labeling

For 5-bromo-2’-deoxyuridine (BrdU, Sigma) labeling, E15 pregnant mice were intraperitoneally injected with BrdU (50 mg per kg body weight) every 6 hours for three time or one pulse 1 hour prior to sacrifice. Animals were sacrificed 1 hour after the last injection and brains were fixed. For recovering of BrdU, brain slices were incubated with 2N HCl at 37°C for 25 mins to denature DNA, followed by incubation with primary antibodies and secondary antibodies.

### Western Blotting

Cultured cells were washed three times with ice-cold PBS and then lysed in RIPA buffer (CWBIO) containing fresh protease inhibitor cocktails (Targetmol). Crude protein was extracted by centrifugation at 12,000 rpm for 15 mins at 4°C and then the supernatants were transferred to new tubes containing 1× loading buffer, and heated for 5 mins at 95℃. Protein was separated by SDS-PAGE electrophoresis and then transferred to nitrocellulose membrane at 200 mA for 2 hours in 4℃. Membranes were blocked in 0.1% TBST containing 5% non-fat powdered milk for 2 hours at room temperature and then incubated with primary antibodies overnight at 4°C. The primary antibodies were used as follows: rabbit anti Ednrb (Abcam ab117529, 1∶5000), mouse anti β-actin (Proteintech 60008, 1∶5000) overnight at 4°C. Membranes were then incubated with HRP goat anti-rabbit (Proteintech SA00001-2, 1:5000) or HRP goat anti-mouse (Proteintech SA00001-1, 1:5000). Immunoblots were developed with Ultra-sensitive ECL chemiluminescent solution (iBio Co.) and visualized using scanned using a Gel imaging system (GE, AI600). Data were analyzed using ImageJ (NIH)-Fuji software.

### Electrophysiology

For preparation of acute brain slices, mice (∼P30) that received *in utero* retroviral injections were deeply anesthetized by isoflurane. Brains were quickly removed and sectioned to 350 μm brain slices in ice-cold artificial cerebral spinal fluid (ACSF) containing 126 mM NaCl, 4.9 mM KCl, 1.2 mM KH_2_PO_4_, 2.4 mM MgSO_4_, 2.5 mM CaCl_2_, 26 mM NaHCO_3_ and 10 mM glucose, bubbled with 95% O_2_/5% CO_2_ by a Vibratome (Leica VT1200S). Slices were recovered in the interface chamber at room temperature for at least one hour before being transferred to the recoding chamber. An infrared-differential interference contrast (DIC) microscope (Olympus) equipped with epifluorescence illumination was used to visualize and localize the recording electrodes to EGFP-expressing cells. Voltage and current signals were recorded by an Axon 700B amplifier (Axon). Glass pipettes (10-12 MΩ resistance) filled with a solution consisting of 136 mM K-gluconate, 6 mM KCl, 1 mM EGTA, 10 mM HEPES, 2.5 mM Na_2_ATP and 0.3% Alexa Fluor 568 hydrazide (Invitrogen) (pH 7.25 and 280 mOsm/kg) were used. Data were collected with a low-pass filter at 2 kHz using Multiclamp 700B and Digidata 1322A/D converter, and sampled at 10 kHz using pCLAMP software. To assess the current-voltage (I-V) curve, membrane voltages of EGFP-expressing cells were recorded in response to steps of current injections with an increment of 20 pA.

### mRNA sequencing (RNAseq) and analysis

Brain slices from E15 embryos were prepared in the same way as electrophysiological recordings 2 days after *in utero* injection of retroviruses (E13). For preparation of RNAseq, radial glial progenitors were collected using a dual-patch clamping system in which one pipette was used to isolate an EGFP-expressing radial glial cell (RGC) through positive air blow while the other pipette (tip diameter ∼6 µm) filled with intracellular solution containing RNase inhibitor was applied to extract the RGC via negative pressure. Then the extracted RGCs were quickly transferred to the cell lysate with RNase inhibitor on ice and ∼40 RGCs were collected into a tube. cDNA was prepared following the smart-seq2 method. The cDNA libraries were constructed using TruSeq Stranded mRNA LTSample Prep Kit (Illumina, San Diego, CA, USA) according to the manufacturer’s instructions. Then these libraries were sequenced on the Illumina sequencing platform (HiSeqTM 2500 or Illumina HiSeq X Ten) and 125bp/150bp paired-end reads were generated.

Raw data (raw reads) were processed using NGS QC Toolkit. The reads containing ploy-N and the low quality reads were removed to obtain the clean reads. Then the clean reads were mapped to reference genome using hisat2 (Kim *et al*, 2019). Differential expression gene (DEG) analysis was performed using DESeq2. Statistical values were FDR correction using the Benjamini-Hochberg method. FPKM was calculated using cufflinks, and the read counts were obtained by htseq-count. The threshold for DEGs was set as log2(foldchange) ≥1 and Padj ≤0.01 in expression of a gene between control RGCs and *Ptf1a*-expressing RGCs. GraphPad Prism was used to generate volcano plots of all genes according to fold change and significances. The normalized FPKM of DEGs were applied to generate a heatmap using R. GO Biological Process and KEGG pathways was performed using David (https://david.ncifcrf.gov) and visualized using GraphPad Prism and R. Protein interaction networks were generated using string (https://www.string-db.org) and optimized with Cytoscape. Violin plots of genes were generated using GraphPad Prism.

### Quantitative real-time PCR

Total RNA from extracted cortical EGFP-expressing RGCs was extracted using miRNeasy mini Kit (Qiagen) according to the manufacturer’s instructions. For cDNA synthesis, RNA was incubated with 2.5 mM OligodT primers (Invitrogen) and 10 mM dNTP (Thermofisher) for 5 mins at 65°C. cDNA was reversely transcribed using Moloney murine leukemia virus reverse transcriptase (M-MLV RT) (Invitrogen, 28025013) and RNaseOUT Recombinant RNase Inhibitor (Takara) according to the manufacturer’s instructions. A 1/10 aliquot of cDNA was added into ChamQ SYBR qPCR Master Mix (Vazyme) to undergo PCR amplification with primers as follows: Ptf1a, forward 5’-CCACGCTACCCTACGAAAAGC-3’, reverse 5’- CCTTCTGGGCCTGGTTACC-3’; Ednrb, forward 5’- TTCCCACCTGCCCAAGCCAC-3’, reverse 5’-AGGTGCGGAGGAACGCATCA-3’; Edn1, 5’-GGCCCAAAGTACCATGCAGA-3’, reverse 5’-GATGGCCTCCAACCTTCGTA-3’; Notch3, forward 5’- GGTTGCTGCTGATATCCGTGT-3’, reverse 5’-GTGGGGTGAAGCCATCAGG-3’; Hes5, forward 5’-AACTCCAAGCTGGAGAAGGC-3’, reverse 5’-GTCAGGAACTGTACCGCCTC-3’; GAPDH, forward 5’- AGGTCGGTGTGAACGGATTTG-3’, reverse 5’-TGTAGACCATGTAGTTGAGGTCA-3’. qPCRs reactions were performed using the ABI ViiA™ 7 Real-Time PCR System. The relative expression levels of target genes to an endogenous reference gene GAPDH that were calculated using the ΔΔCT method were used for normalization.

### Statistical analysis

Statistical analyses were performed blindly to the experimenter using DESeq2, GraphPad Prism, R, DAVID, STRING and Cytoscape. N values and experimental replicates are detailed in figure legends. Graphs display as means ± SEM unless otherwise stated. Statistical testing was based on unpaired student t-test, one-way

ANOVA or two-way ANOVA with *posthoc* Bonferroni’s test. Statistical significance was defined as *p < 0.05; **p<0.01; ***p<0.001; ****p<0.0001.

## KEY RESOURCE TABLE

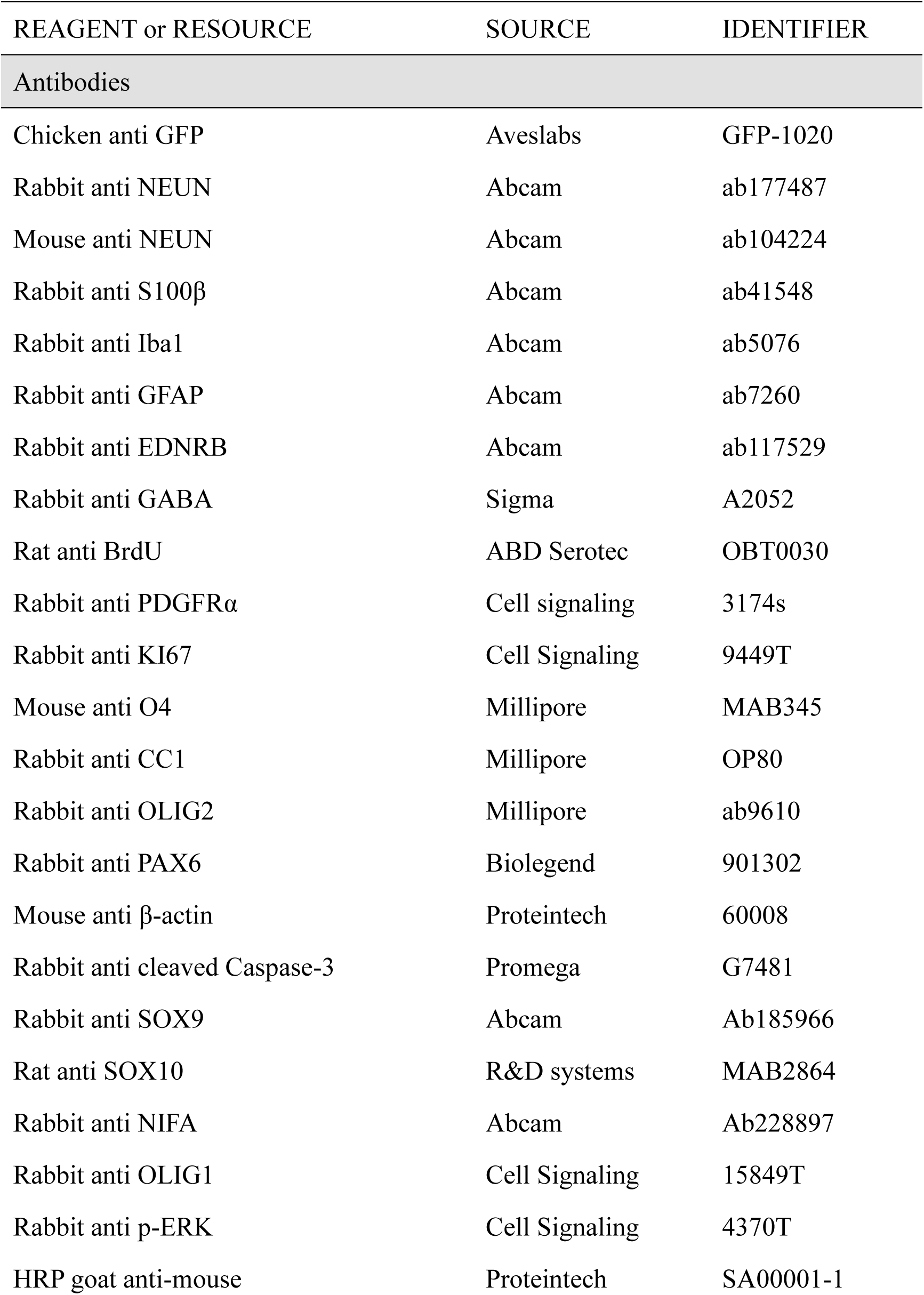
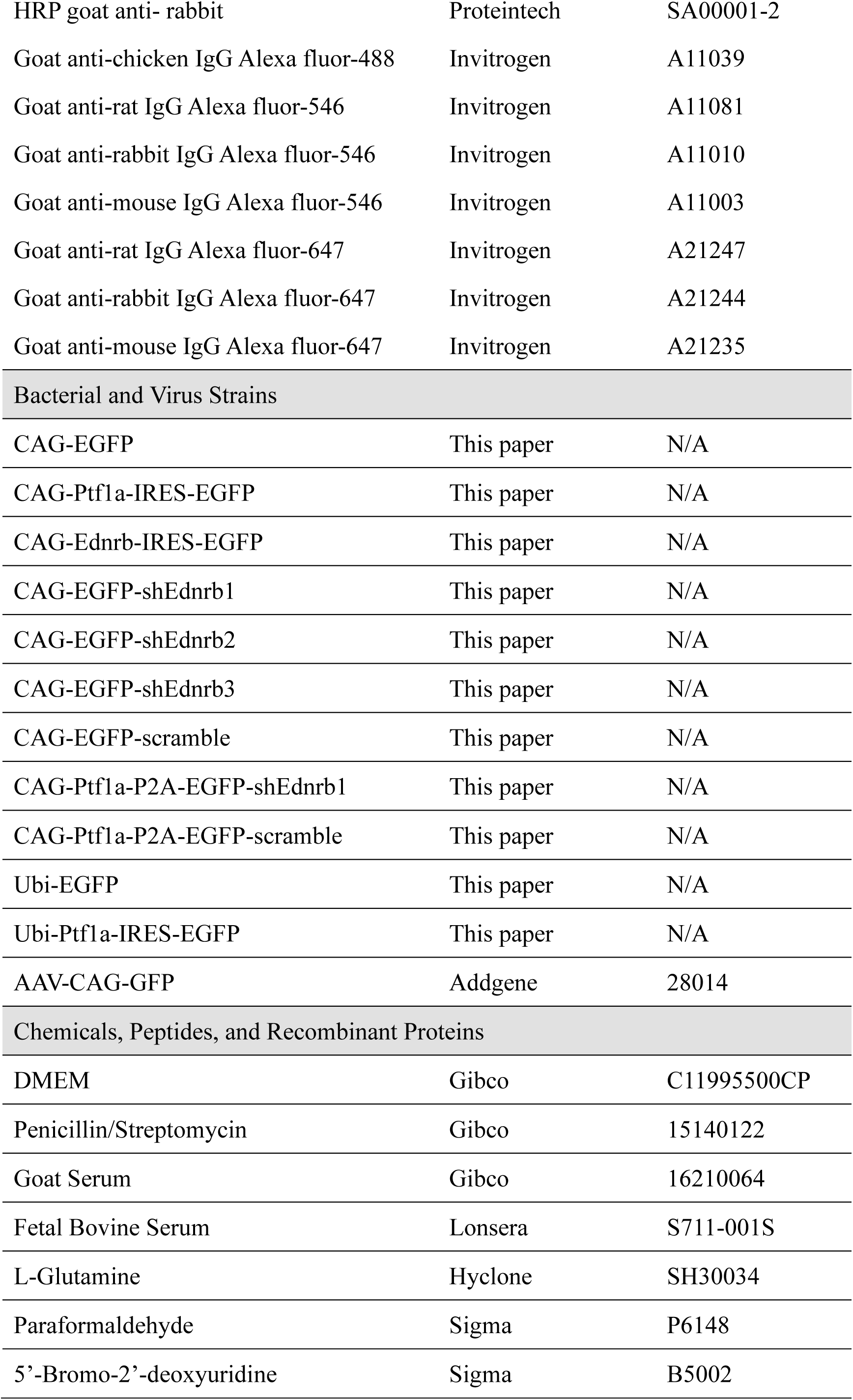
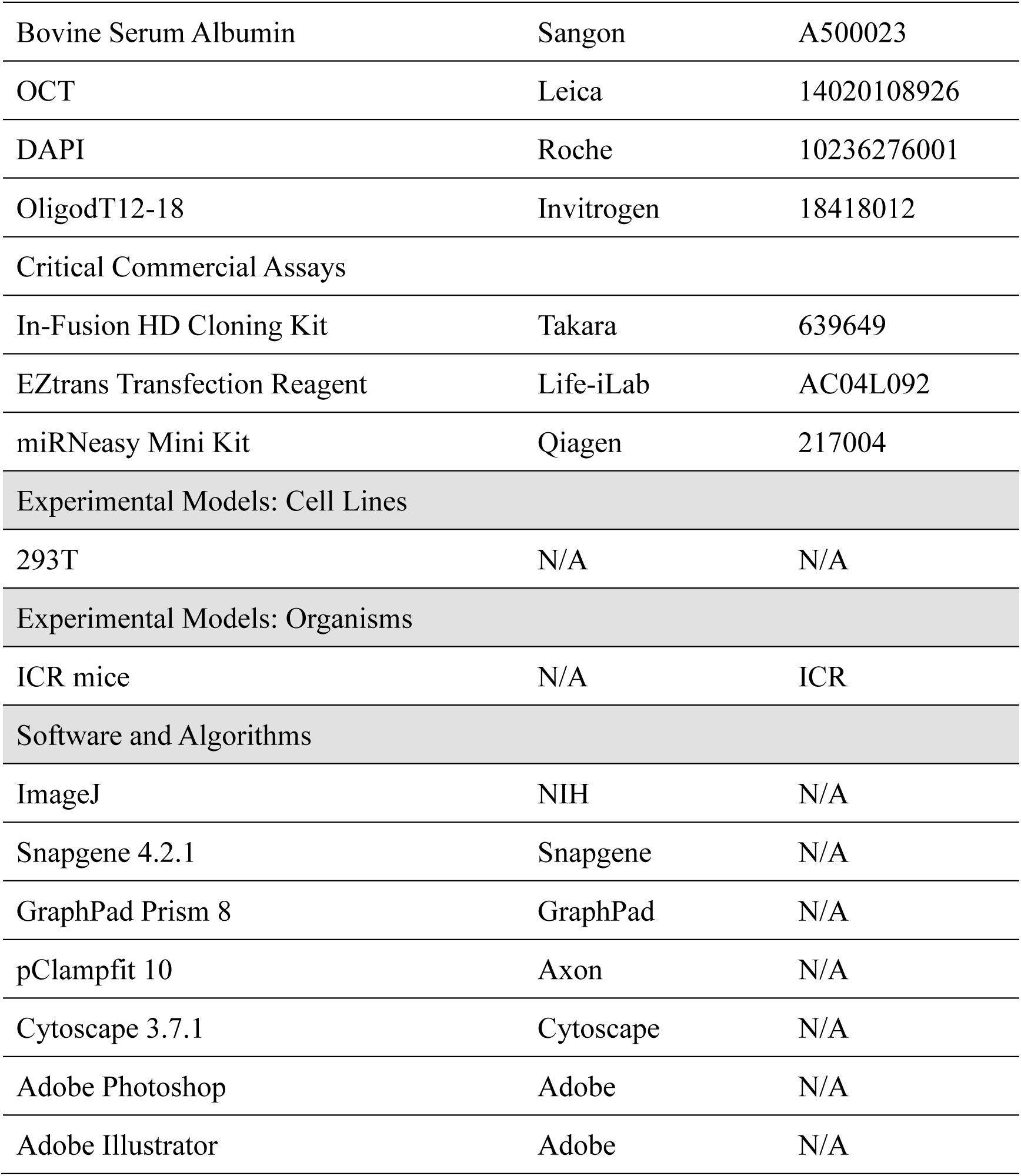

### RESOURCE AVAILABILITY

#### Lead contact

Further information and requests for resources and reagents should be directly addressed to the lead contact, Shuijin He.

#### Materials availability

All unique and stable reagents generated in this study of which sufficient quantities exists are available from the Lead Contact with a completed Materials Transfer Agreement.

#### Data and code availability

The accession number for the RNA-Seq raw data generated in this paper is PRJNA882943: https://www.ncbi.nlm.nih.gov/bioproject/PRJNA882943.

## ACKNOWLEDGEMENTS

We thank the facilities of Imaging Core of Life School of Science and Technology at ShanghaiTech University for technical supports. This study was supported by the National Key Research and Development Program of China (2021ZD0202500), National Natural Science Foundation of China (32270851, 81870734), Shanghai Natural Science Foundation (Grant No.22ZR1441500), and the Shanghai Municipal Government and ShanghaiTech University.

## AUTHOR CONTRIBUTIONS

Conceptualization, S.H.; Methodology, H.L., S.H., S.C. and Y.L.; Investigation, H.L., K.L. X.W., C.J., K.H., and S.X.; Writing, H.L. and S.H.; Funding Acquisition, S.H.; Supervision, S.H.

## DECLARATION OF INTERESTS

The authors declare no competing financial interests.

## Notes

https://www.ncbi.nlm.nih.gov/bioproject/PRJNA882943

### Competing Interest Statement

The authors have declared no competing interest.

