## Supplementary material for "*Ptf1a* robustly drives the gliogenic switch in the rodent embryonic cortex in a dosage-dependent manner by activating pro-glial gene expression programs": Suppl figures and Tables

Haixiang Li *et al.*

**This file includes:**

Figs. S1 to S13
Tables S1 to S6

**Fig. S1.**

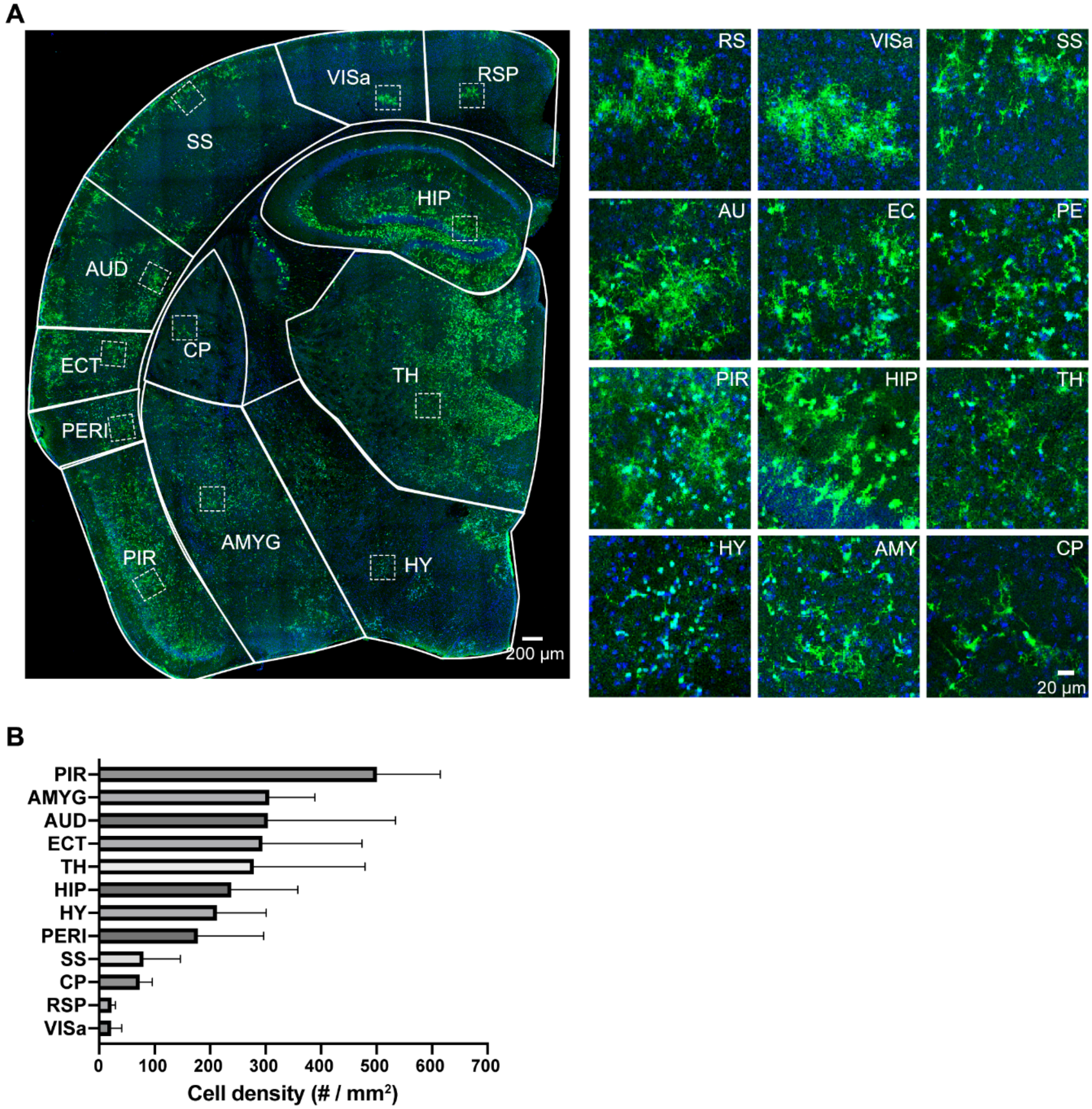


**EGFP-expressing glial cells are distributed across the brain of *Ptf1a* mice.**
Representative images of EGFP staining (*A*) and quantification of the density of EGFP+ cells (*B*) in different brain regions of P30 *Ptf1a* mice (n = 3 mice). Right panels in (*A*) are magnified from the corresponding boxed regions in the left.

Fig. S2.

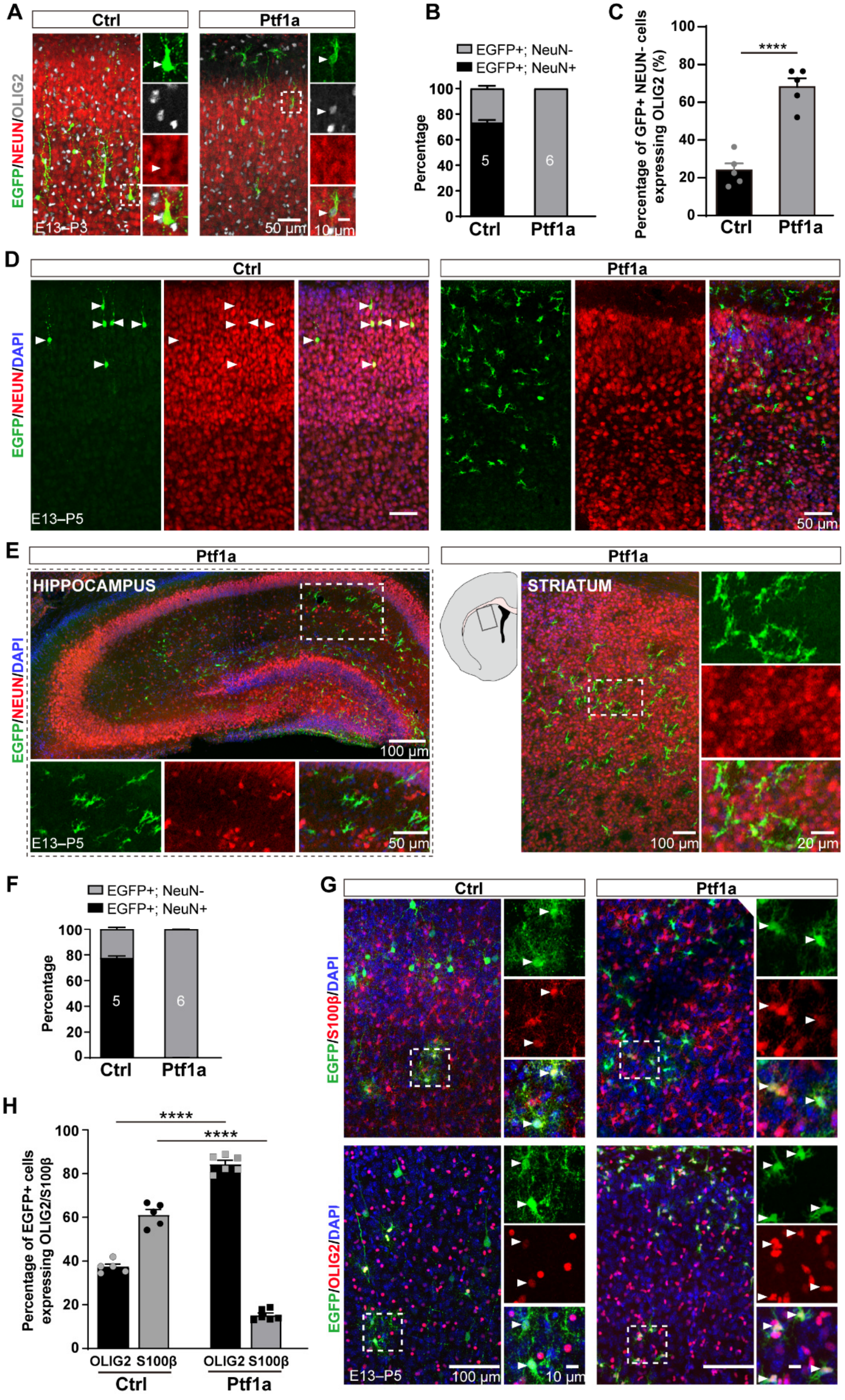
A
Ctrl
Ptf1a
EGFP/NEUN/OLIG2
E13–P3
50 µm
10 µm
B
EGFP+; NeuN-
EGFP+; NeuN+
Percentage
100
80
60
40
20
0
5
6
Ctrl
Ptf1a
C
Percentage of GFP+ NEUN- cells expressing OLIG2 (%)
100
80
60
40
20
0
****
Ctrl
Ptf1a
D
Ctrl
Ptf1a
EGFP/NEUN/DAPI
E13–P5
50 µm
E
Ptf1a
HIPPOCAMPUS
EGFP/NEUN/DAPI
100 µm
E13–P5
50 µm
Ptf1a
STRIATUM
100 µm
20 µm
F
EGFP+; NeuN-
EGFP+; NeuN+
100
80
60
40
20
0
Percentage
5
6
Ctrl
Ptf1a
G
Ctrl
Ptf1a
EGFP/S100β/DAPI
EGFP/OLIG2/DAPI
E13–P5
100 µm
10 µm
H
Percentage of EGFP+ cells expressing OLIG2/S100β
100
80
60
40
20
0
****
****
OLIG2
S100β
Ctrl
OLIG2
S100β
Ptf1a

**PTF1A-expressing cells are glia in the postnatal mouse brain.**

(A) Representative images showing NEUN and OLIG2 immunostaining in cortices of P3 mouse brains with EGFP alone (control; *A, left*) or with Ptf1a-IRES-EGFP (Ptf1a; *A, right*). Dashed boxes are enlarged in the right panels. Arrowheads indicate EGFP+ cells co-expressing NEUN or OLIG2.

(B) Quantification of the percentage of EGFP+ cells expressing NEUN.

(C) Quantification of the percentage of EGFP+ NEUN- cells that express OLIG2. ****p < 0.0001, Student's t-test.

(D, E) Representative images showing EGFP and NEUN staining in the cortex (*D*), hippocampus (*E, left*), and striatum (*E, right*) from P5 mouse brains. Arrowheads indicate EGFP+ NEUN+ cells. Dashed boxes are enlarged in the right panels.

(F) Quantification of the percentage of NEUN+ and NEUN- cells among total EGFP+ cells in the P5 cortices of control and *Ptf1a* mice.

(G) Representative images showing EGFP with S100β or OLIG2 staining in the P5 cortex of EGFP-only control mice (*left*) and *Ptf1a* mice (*right*). Arrowheads indicate EGFP+ cells co-expressing S100β or OLIG2. Dashed boxes are enlarged in the right panels.

(H) Quantification of the percentage of S100β+ and OLIG2+ cells among total EGFP+ cells in the P5 cortices of control and *Ptf1a* mice. ****p < 0.0001, Two-way ANOVA with *post hoc* Bonferroni's test.

**Fig. S3.**

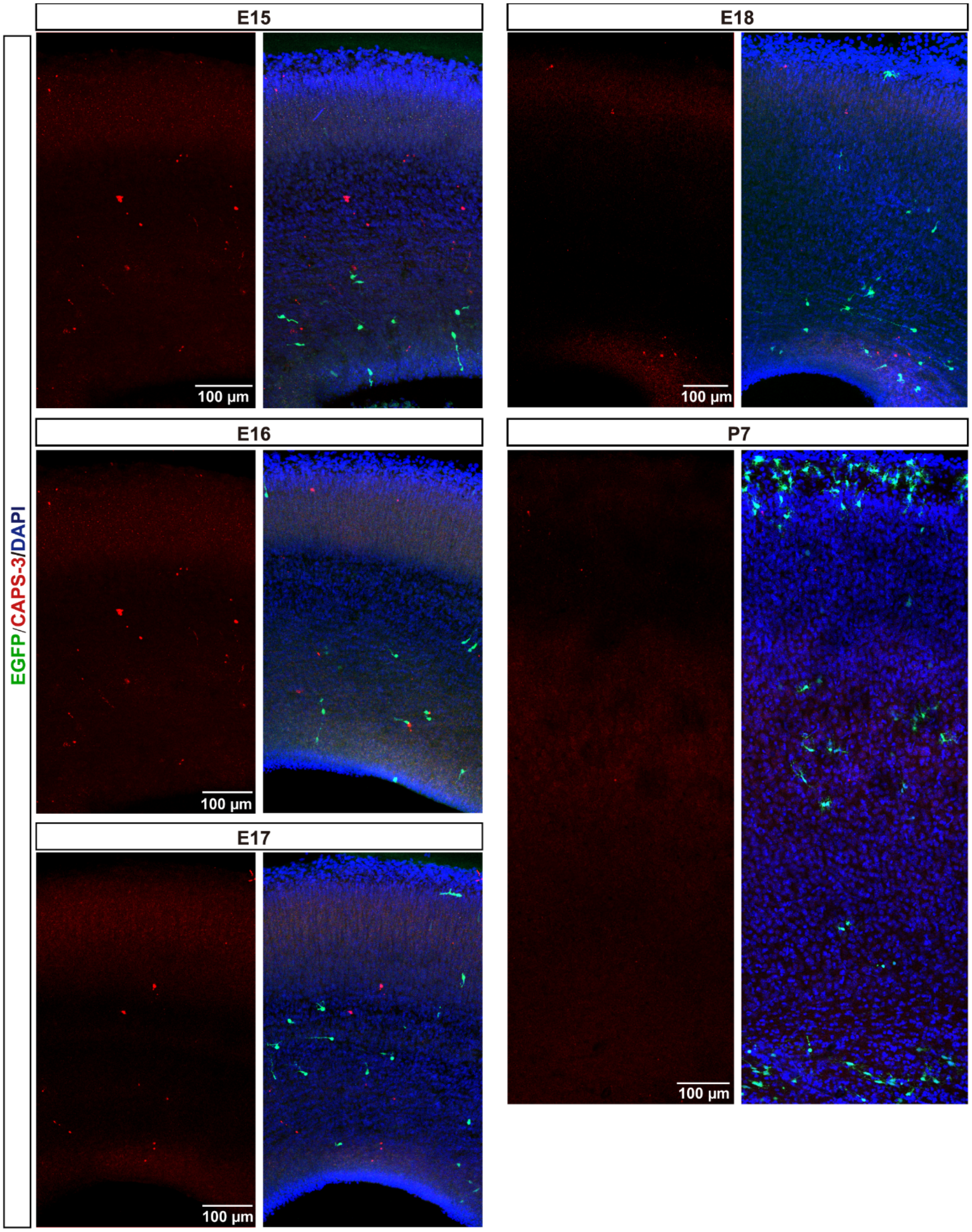


**PTF1A expression does not cause cell apoptosis.**

Representative images showing immunostaining for an apoptotic maker cleaved caspase-3 in E15, E16, E18, E17 and P7 mouse cortices. 4 mouse brains have been assessed for each age.

**Fig. S4.**

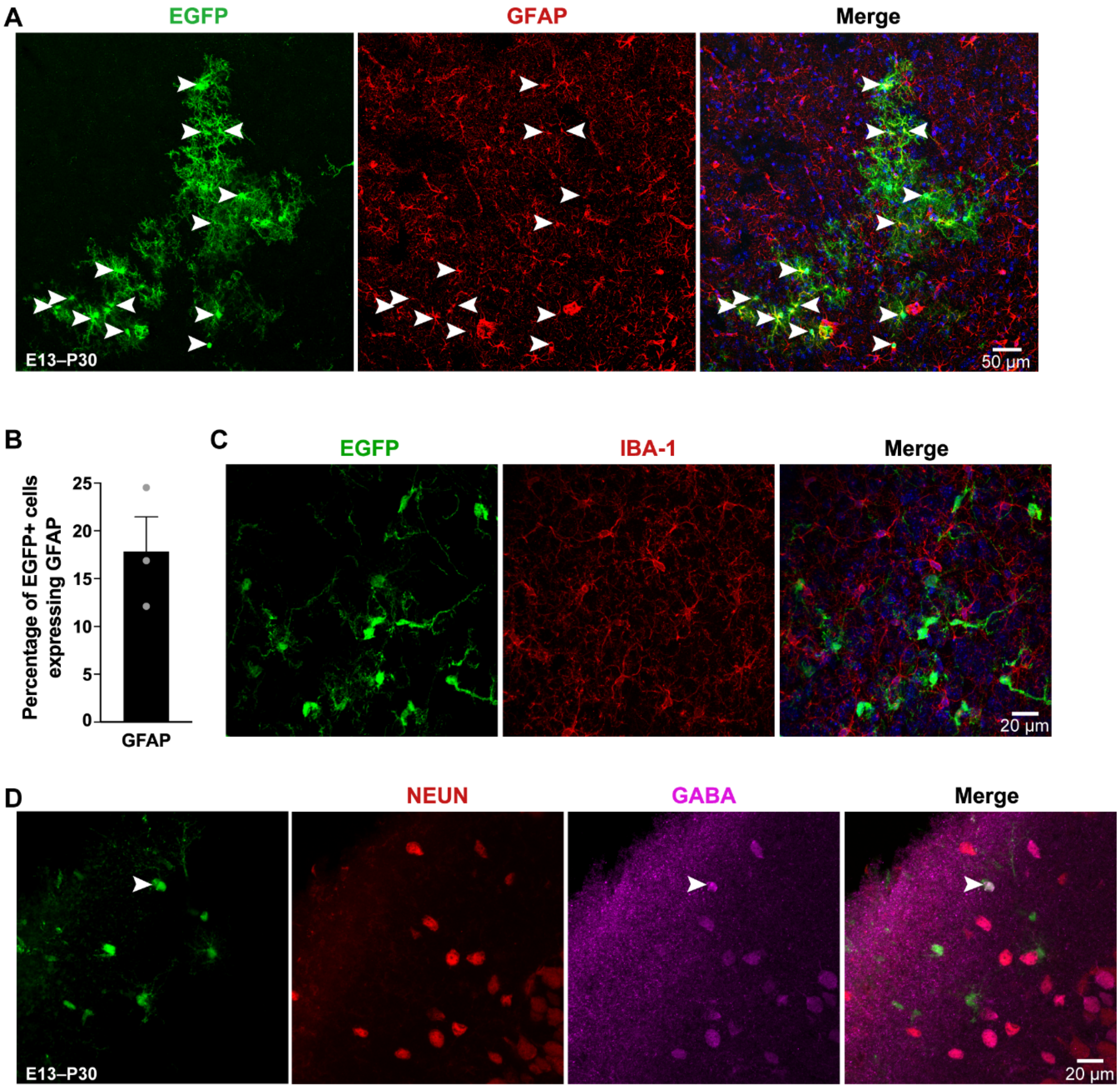


**PTF1A-expressing RGPs give rise to astrocytes and oligodendrocytes, but not microglia or interneurons.**

(A, B) Representative images showing EGFP and GFAP staining (*A*) and quantification of the percentage of GFAP+ cells among total EGFP+ cells (*B*) in P30 *Ptf1a* cortices. Arrowheads indicate colocalized cells.

(C) Representative images showing EGFP and Iba1 staining in a P30 *Ptf1a* cortex (n = 6 mice).

(D) Representative images showing GFP, NEUN and GABA staining of a P30 *Ptf1a* cortex (n = 7 mice). Arrowhead indicates an EGFP+ cell positive for GABA but negative for NEUN, indicative of an astrocyte expressing GABA. Data are presented as mean ± SEM.

**Fig. S5.**

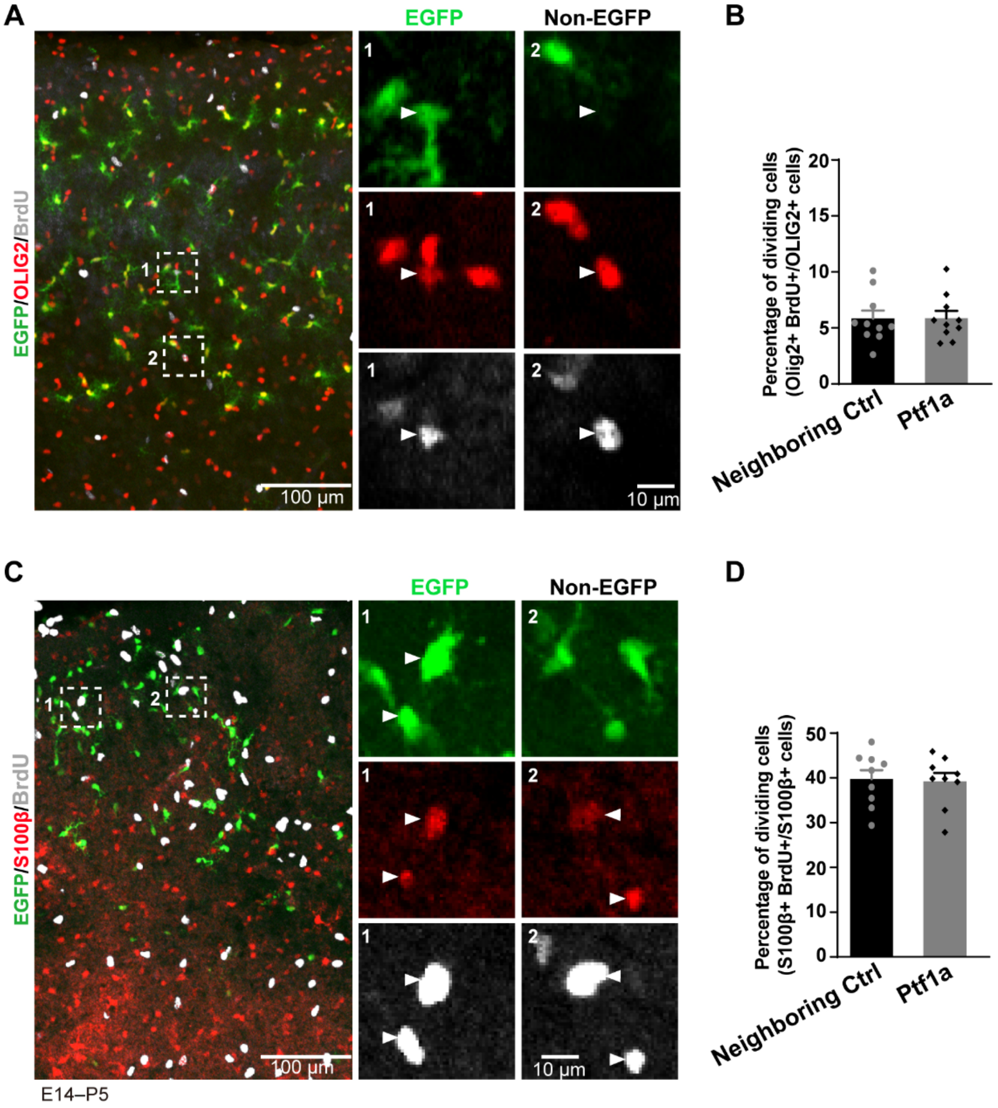


**PTF1A does not alter the proliferating rate of glial progenitor cells.**

(A, C) Representative images showing immunostaining of P5 *Ptf1a* cortices for EGFP, OLIG2 and BrdU (*A*) and for EGFP, S100β and BrdU (*C*). Dashed square areas are enlarged in the right panels. Arrowheads indicate cells co-labeled with BrdU and OLIG2 or with BrdU and S100β.

(B, D) Quantification of the percentage of BrdU+ cells among EGFP+ OLIG2+ cells or adjcent EGFP- OLIG2+ cells (*B*), and among EGFP+ S100β+ or adjacent EGFP- S100β+ cells (*D*). Data are presented as mean ± SEM.

**Fig. S6.**

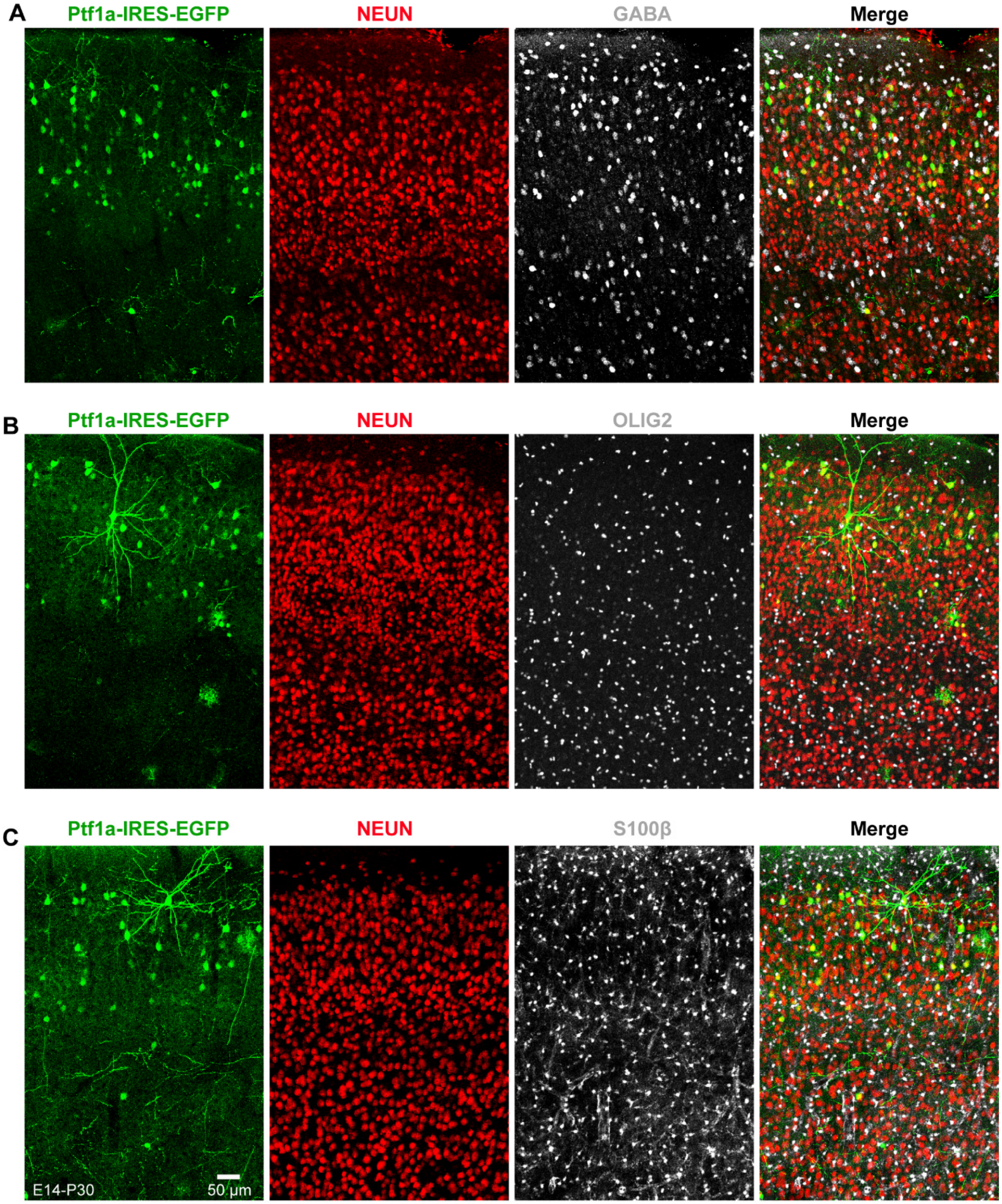


**Electroporation of *Ptf1a* has no effect on the cell fate of RGPs.**
(A-C) Representative images showing immunostaining of P3- cortices for EGFP, NEUN and GABA staining (*A*), EGFP, NEUN and OLIG2 staining (*B*), and EGFP, NEUN and S100β (*C*) Sample sizes: GABA (n = 6 mice), OLIG2 (n = 7 mice), S100β (n =7 mice).

**Fig. S7.**

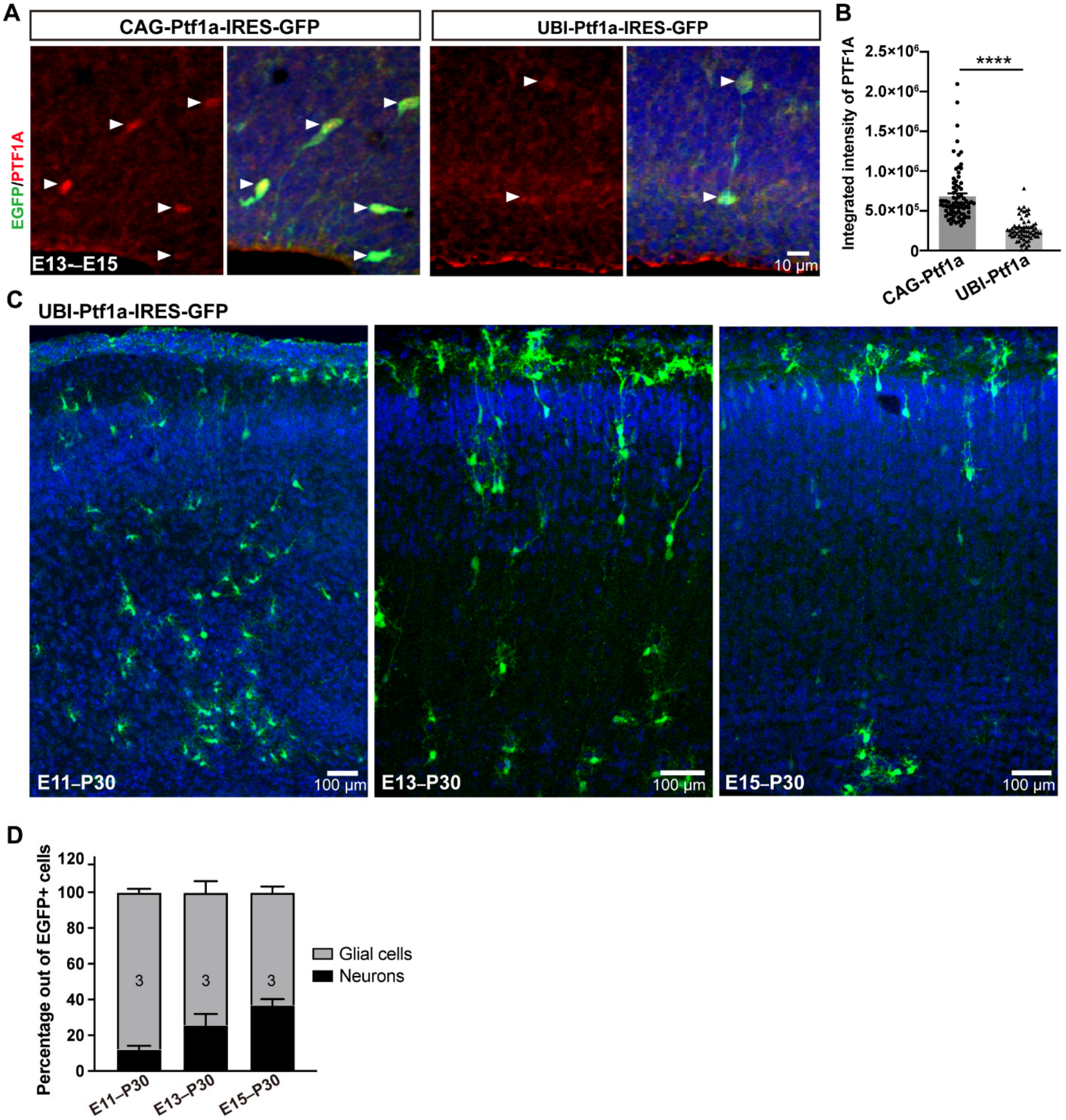


**Viral expression of *Ptf1a* driven by a Ubi promoter results in a mild phenotype of gliogenesis in the cortex.**

(A, B) Immunohistochemistry analysis showing that PTF1A expression levels driven by a strong CAG promoter are significantly higher than those driven by the Ubi promoter. Arrowheads indicates colocalized cells. N represents cell number analyzed. ****$p < 0.0001$, Student's t-test.

(C, D) Representative images of EGFP staining (*C*) and quantification of the percentage of glial cells and neurons among total EGFP+ cells (*D*) in P30 *Ubi-Ptf1a* cortices at different embryonic time points of virus injection. Neurons and glial cells are identified according to cell morphology. Data are presented as mean ± SEM.

**Fig. S8.**

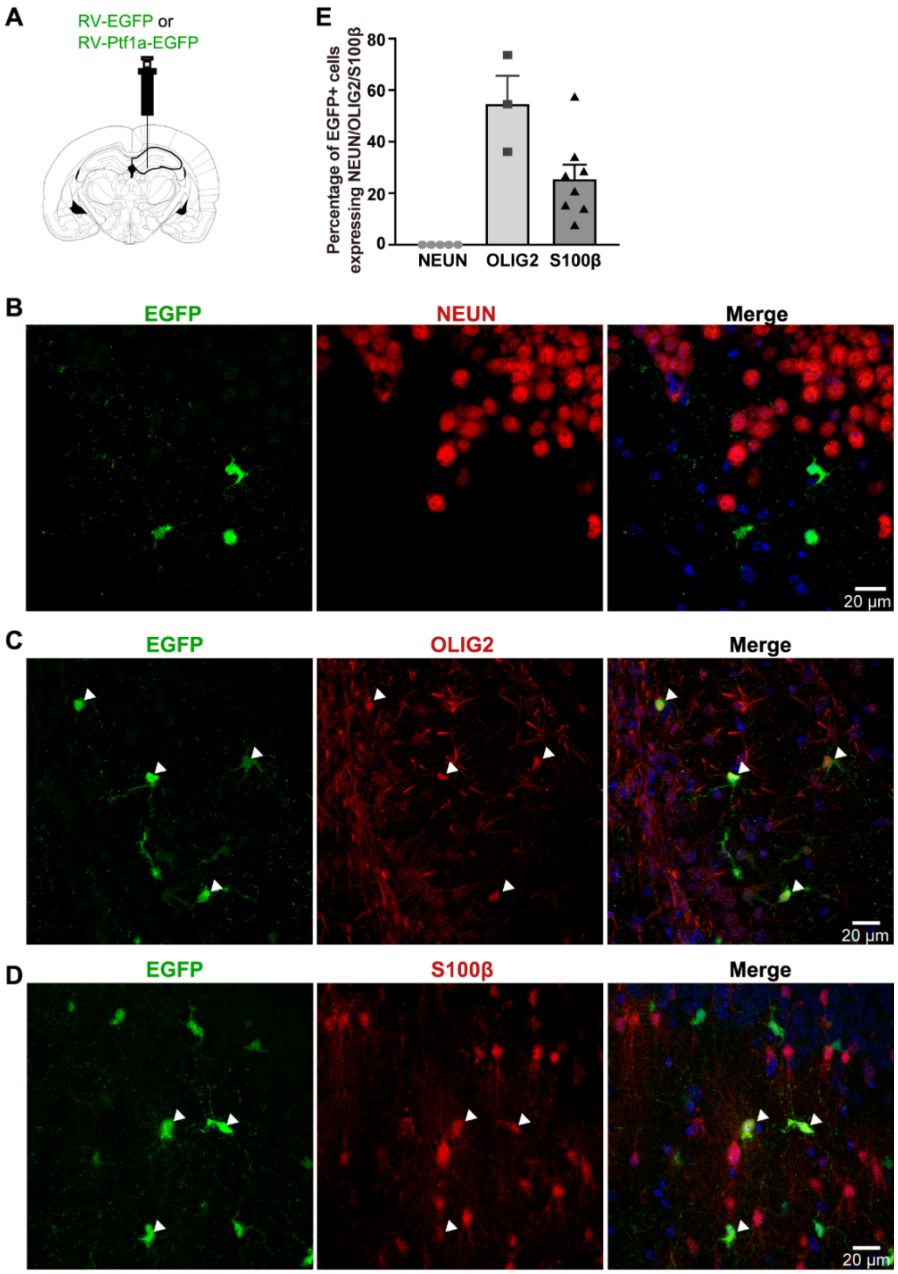


**Viral expression of *Ptf1a* strongly drives adult hippocampal neural stem cells to differentiate into glial cells.**

(A) Schematic of stereotaxic injection of retroviruses-expressing *Ptf1a* into the subgranular zone of P56 mouse hippocampus.

(B-D) Representative images showing staining of P80 *Ptf1a* mouse hippocampi for EGFP and NEUN (*B*), EGFP and OLIG2 (*C*), and EGFP and S100β (*D*).

Arrowheads indicate colocalization of EGFP and OLIG2 staining or colocalization of EGFP and S100β staining.

(E) Quantification of the percentage of NEUN+ cells, Olig2+ cells and S100β+ cells out of total EGFP+ cells. Data are presented as mean ± SEM.

Fig. S9.

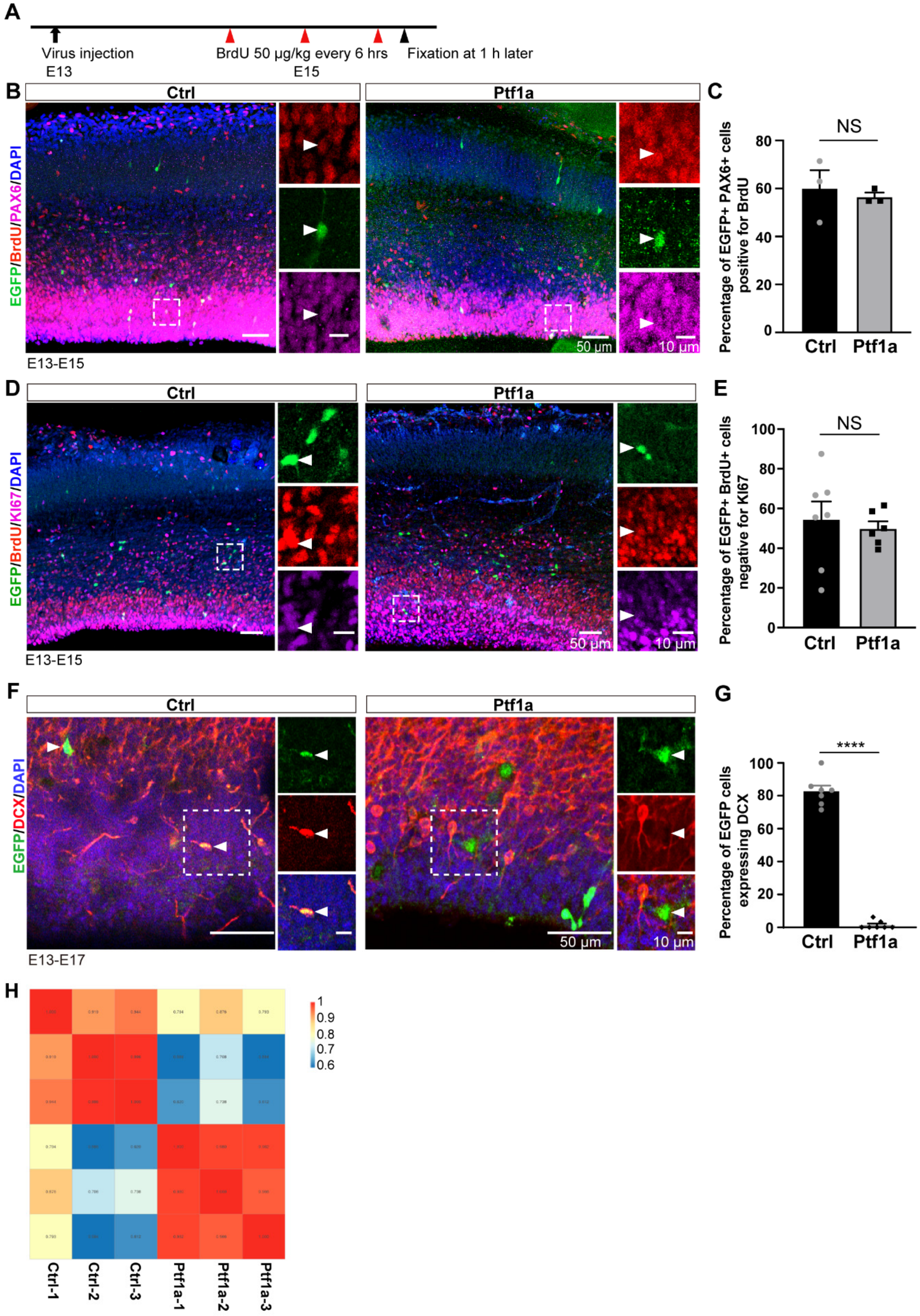

A
Virus injection
E13
BrdU 50 µg/kg every 6 hrs
E15
Fixation at 1 h later
B
Ctrl
Ptf1a
EGFP/BrdU/PAX6/DAPI
50 µm
10 µm
E13-E15
C
Percentage of EGFP+ PAX6+ cells positive for BrdU
NS
80
60
40
20
0
Ctrl
Ptf1a
D
Ctrl
Ptf1a
EGFP/BrdU/KI67/DAPI
50 µm
10 µm
E13-E15
E
Percentage of EGFP+ BrdU+ cells negative for KI67
NS
100
80
60
40
20
0
Ctrl
Ptf1a
F
Ctrl
Ptf1a
EGFP/DCX/DAPI
50 µm
10 µm
E13-E17
G
Percentage of EGFP cells expressing DCX
****
100
80
60
40
20
0
Ctrl
Ptf1a
H
1
0.9
0.8
0.7
0.6
Ctrl-1
Ctrl-2
Ctrl-3
Ptf1a-1
Ptf1a-2
Ptf1a-3

**PTF1A-expressing RGPs undergo normal cell divisions and cell cycle exit.**
(A) Schematic of experimental paradigm.
(B, C) Representative images showing EGFP, BrdU and PAX6 staining (*B*) and quantification of the percentage of BrdU+ PAX6+ cells among EGFP+ cells (*C*) in E15 control and *Ptf1a* cortices. Right panels in (*B*) are enlarged from the boxed regions in the left. Arrowheads indicate triple colocalized cells.
(D, E) Representative images showing EGFP, BrdU and KI67 staining (*D*) and quantification of the percentage of KI67- cells among EGFP+ BrdU+ cells (*E*) in E15 control and *Ptf1a* cortices. Right panels in (*D*) are enlarged from the boxed regions in the left. Arrowheads indicate a cell positive for EGFP, BrdU and KI67. Data are presented as mean ± SEM.
(F, G) Representative images showing EGFP and doublecortin (DCX) staining (*G*) and quantification of the percentage of DCX+ cells among EGFP+ cells (*H*) in E15 control and *Ptf1a* cortices. Right panels in (*B*) are enlarged from the boxed regions in the left. Arrowheads indicate an EGFP+ DCX+ cell in a control cortex and an EGFP+ DCX- cell in a *Ptf1a* cortex. ****p < 0.0001, Student's t test.
(H) Correlation between control and *Ptf1a* RGPs sequencing samples.

**Fig. S10.**

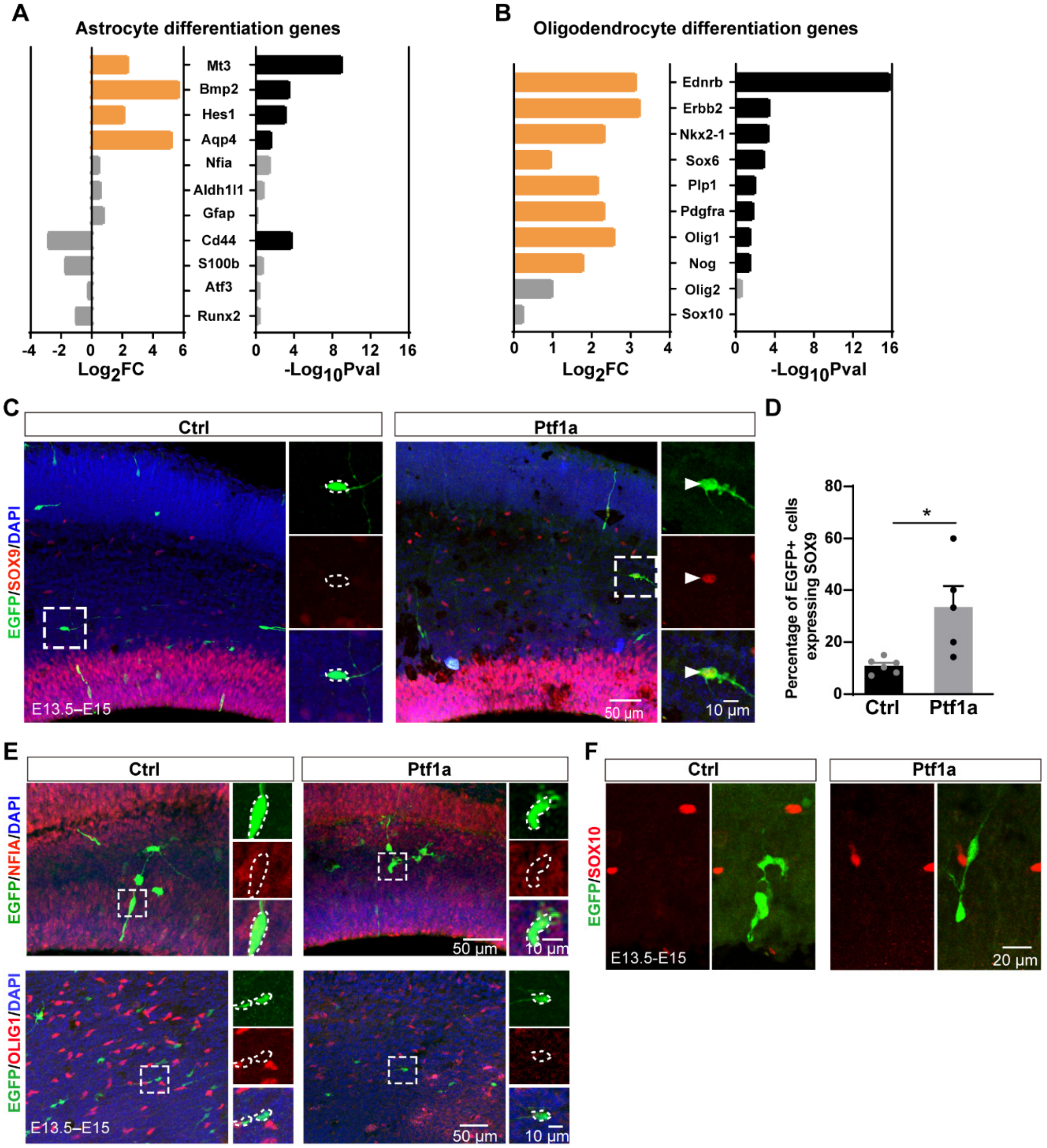


**Validation of known pro-glial genes from RNAseq**

(A, B) Bar graphs showing fold changes and statistical significance of pro-astrocyte (*A*) and pro-oligodendrocyte (*B*) differentiation genes. Yellow bars indicate upregulated genes; black bars denote genes with $p < 0.05$.

(C–F) Validation of the expression of SOX2 (*C, D*), NFIA (*E, top*), OLIG1 (*E, bottom*), and SOX10 (*F*) in EGFP+ cells from E15 control and *Ptf1a* cortices. Notably, only SOX9 expression is upregulated in *Ptf1a*-expressing cells. Right panels show magnified views of boxed regions. Arrowheads in (*C*) indicate EGFP+ cells co-expressing SOX9; dashed circles highlight EGFP+ cells without colocalization.

Fig. S11.

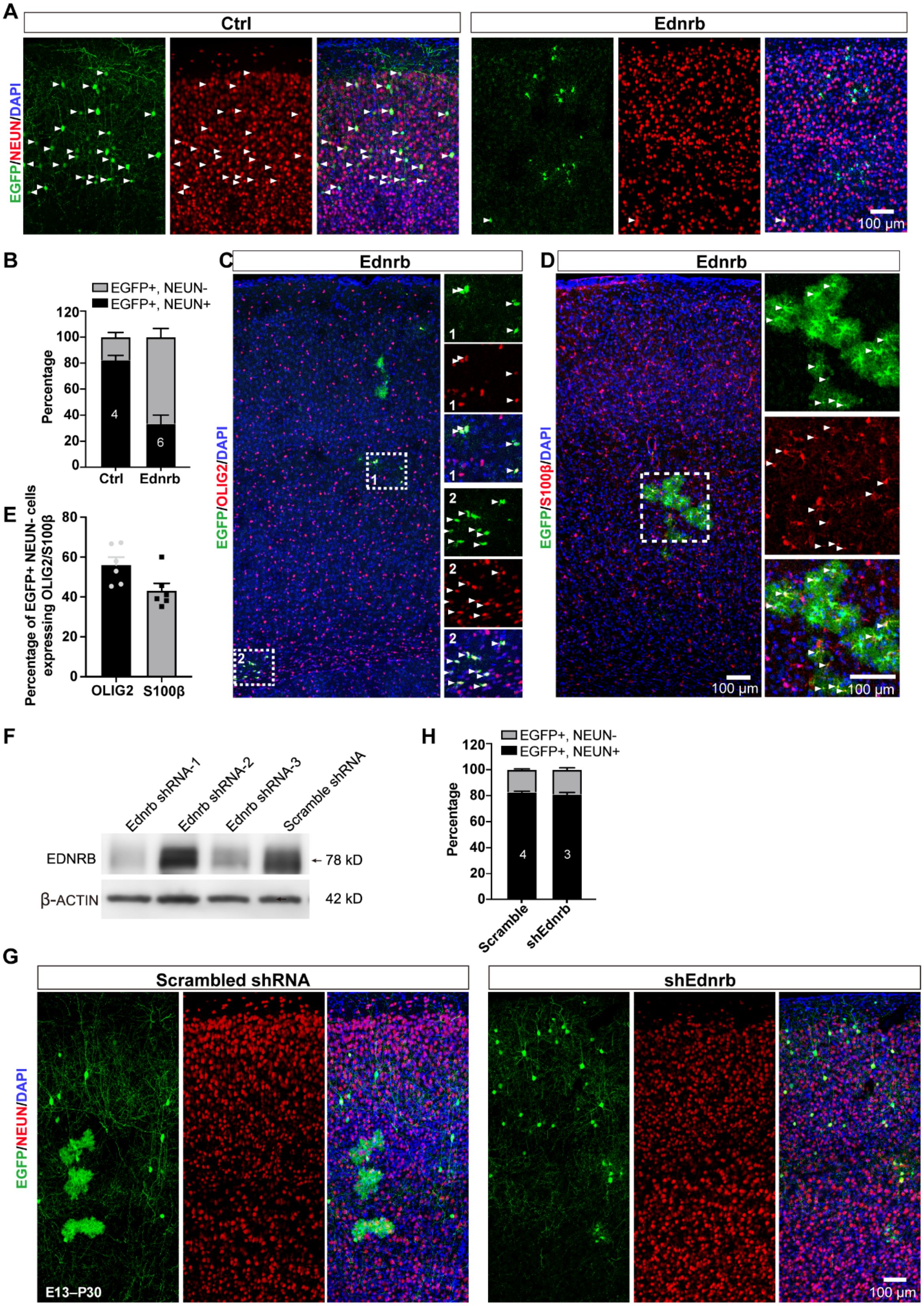

A
Ctrl
Ednrb
EGFP/NEUN/DAPI
100 µm
B
EGFP+, NEUN-
EGFP+, NEUN+
Percentage
120
100
80
60
40
20
0
4
6
Ctrl
Ednrb
C
Ednrb
EGFP/OLIG2/DAPI
D
Ednrb
EGFP/S100β/DAPI
100 µm
100 µm
E
Percentage of EGFP+ NEUN- cells expressing OLIG2/S100β
80
60
40
20
0
OLIG2
S100β
F
Ednrb shRNA-1
Ednrb shRNA-2
Ednrb shRNA-3
Scramble shRNA
EDNRB
← 78 kD
β-ACTIN
42 kD
H
EGFP+, NEUN-
EGFP+, NEUN+
Percentage
120
100
80
60
40
20
0
4
3
Scramble
shEdnrb
G
Scrambled shRNA
shEdnrb
EGFP/NEUN/DAPI
E13–P30
100 µm

**Overexpression of EDNRB promotes the cell fate switch of RGPs in the cortex**

(A, B) Representative images of EGFP and NEUN staining (*A*) and quantification of the percentage of NEUN+ and NEUN- cells among EGFP+ cells (*B*) in P30 control and *Ednrb*-expressing cortices. Arrowheads indicate colocalized cells.

(C-E) Representative images of EGFP and OLIG2 staining (*C*), EGFP and S100β staining (*D*) and quantification of the percentage of OLIG2+ or S100β+ cells among EGFP+ NeuN- cells (*E*) in P30 *Ednrb*-expressing cortices. Arrowheads indicate colocalization. Scale bars, 100 µm.

(F) Representative immunoblot for EDNRB expression (n = 3 replicates).

(G, H) Representative images of EGFP and NEUN staining (*G*) and quantification of the percentage of NEUN+ or NEUN- cells among total EGFP+ cells (*H*) in P30 cortices from mice injected with retroviruses expressing scramble shRNA control or *Ednrb* shRNA. Retroviruses expressing scramble shRNA or *Ednrb* shRNA were injected into RGPs. N indicates the number of mice analyzed. Data are presented as mean ± SEM.

**Fig. S12.**

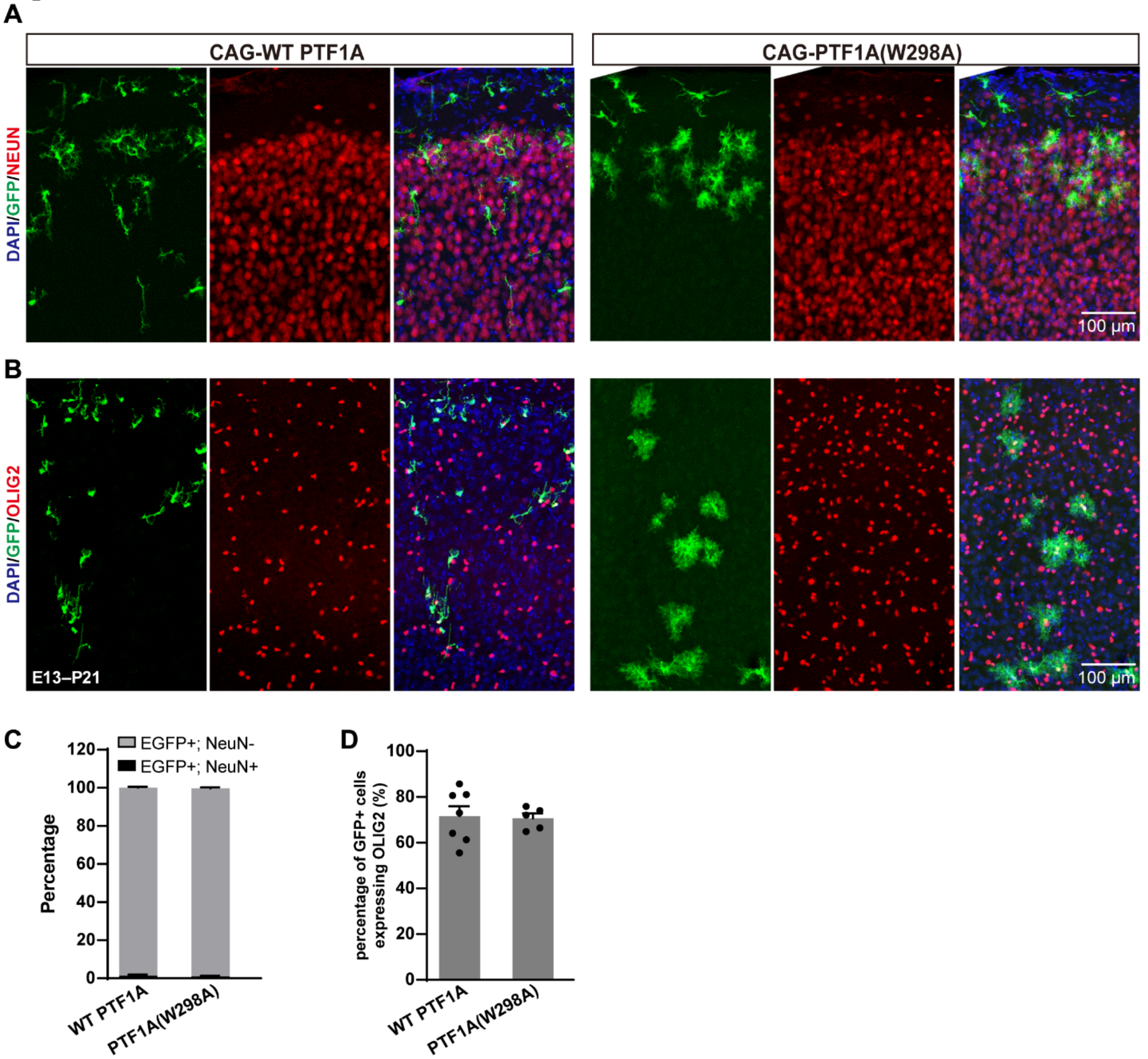


**Point mutation of PTF1A(W298A) has no effect on the glial competency of *Ptf1a*-expressing RGPs**

Representative images showing staining for NEUN (*A*) and OLIG2 (*B*) in cortices of mice injected with retroviruses expressing WT PTF1A or mutant PTF1A(W298A) and quantification of EGFP+ cells expressing NEUN (*C*) and the percentage of EGFP+ NEUN- cells expressing OLIG2 (*D*).

Fig. S13.

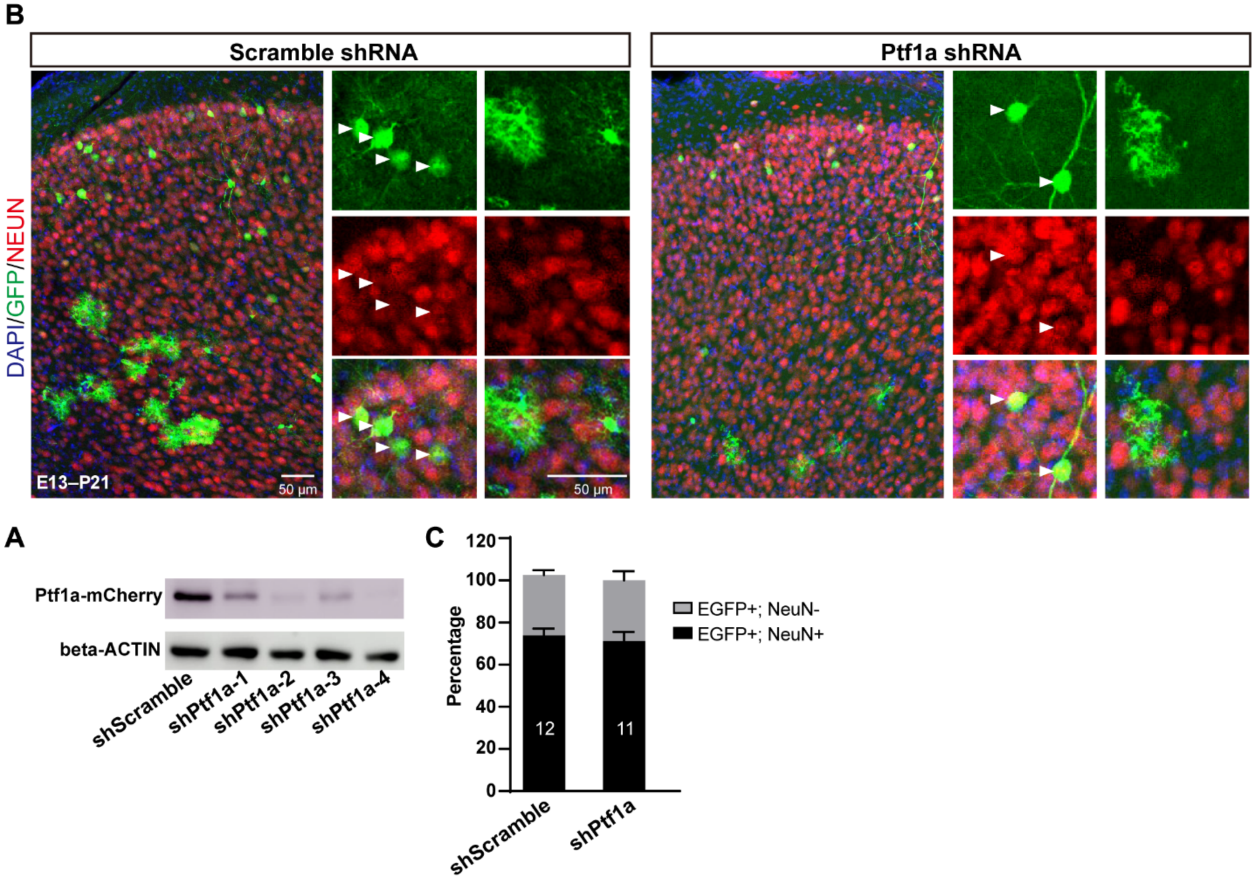


**Knockdown of *Ptf1a* has no effect on the neurogenic-to-gliogenic switch of RGPs during normal development**

(A) Representative immunoblot showing expression of PTF1A-mCherry ($n$ = 3 replicates). (B, C) Representative images of EGFP and NEUN staining (*B*) and quantification of the percentage of $NEUN^+$ and $NEUN^-$ cells among total $EGFP^+$ cells (*C*) in P21 cortices from mice injected with retroviruses expressing scramble shRNA or *Ptf1a* shRNA. *Ptf1a* shRNA-4 was used for knockdown. Retroviruses were introduced into E13 RGPs. $N$ represents the number of analyzed cortices from 4 mice. Data are presented as mean ± SEM.

### Appendix Tables

### Appendix Table S1. FC and padj in Ctrl and Ptf1a RGC.

| gene | log2FoldChange | padj |
|---|---|---|
| Ttr | 3.578625869 | 3.55E-127 |
| Rbp1 | 3.326787472 | 8.89E-41 |
| Angpt1 | 5.25150262 | 6.78E-35 |
| Nnat | 2.026623262 | 4.64E-34 |
| Pkn1 | -2.351431431 | 4.64E-34 |
| Col1a2 | 5.070778977 | 1.15E-32 |
| mt-Cytb | 2.952083894 | 4.59E-30 |
| mt-Nd1 | 2.137227822 | 4.59E-30 |
| Rpl26 | 2.094251697 | 1.02E-29 |
| Col1a1 | 5.040418552 | 2.24E-29 |
| Col3a1 | 4.372819165 | 6.54E-27 |
| Dcn | 5.766957561 | 1.58E-26 |
| Dbi | 2.177391583 | 1.58E-26 |
| Fabp7 | 2.472496588 | 2.41E-26 |
| mt-Atp6 | 1.852308273 | 7.37E-25 |
| Rgs5 | 4.829789623 | 1.46E-24 |
| Tnfrsf11a | -3.274653597 | 1.74E-24 |
| Ednrb | 3.137158676 | 5.00E-24 |
| Cldn11 | 6.607665308 | 1.41E-22 |
| Ptk2b | -2.870725013 | 2.16E-22 |
| Ddah1 | 2.235928201 | 2.29E-22 |
| Sparc | 2.833033025 | 3.63E-22 |
| Syk | -2.542443798 | 6.37E-22 |
| Plekho2 | -2.630970908 | 1.09E-21 |
| Rhoq | -2.510720826 | 8.83E-21 |
| Ikzf1 | -2.727086869 | 3.24E-20 |
| Pik3cg | -2.735409292 | 3.31E-20 |
| Atp1a2 | 3.391285591 | 6.72E-20 |
| Itm2a | 3.139180358 | 9.40E-20 |
| Apod | 4.537167166 | 1.50E-19 |
| Vtn | 6.792950812 | 1.89E-19 |
| Myl9 | 5.666463148 | 2.55E-19 |
| Mt3 | 2.393469977 | 4.16E-19 |
| Ass1 | -3.165768586 | 4.75E-19 |
| Reln | 2.228716056 | 2.11E-18 |
| A730017C20Rik | 3.845058974 | 2.26E-18 |
| Upf1 | -1.971386877 | 2.33E-18 |
| Plbd1 | -2.843632424 | 3.89E-18 |
| 2510039O18Rik | -2.885638555 | 4.17E-18 |
| Sirpb1c | -3.254070348 | 8.26E-18 |
| Adgre5 | -3.194271624 | 9.68E-18 |
| Sla | -1.856625565 | 1.67E-17 |
| Igf2 | 3.678032717 | 1.89E-17 |
| mt-Co3 | 2.166278708 | 2.22E-17 |
| Slc16a3 | -3.111903309 | 2.99E-17 |
| Ptprj | -2.747199239 | 5.92E-17 |
| Ptpn1 | -1.513728902 | 6.64E-17 |
| Myh9 | -2.464761153 | 6.75E-17 |
| Dusp26 | 2.365690929 | 9.31E-17 |
| Ccnd2 | 2.309311015 | 9.31E-17 |
| mt-Nd2 | 1.48045394 | 1.55E-16 |
| Apoc2 | -2.251150665 | 1.55E-16 |
| Rnf149 | -2.259025176 | 2.31E-16 |
| Skp1a | 1.377975889 | 2.88E-16 |
| Arhgef1 | -1.730653988 | 3.30E-16 |
| Stx11 | -2.674514877 | 4.28E-16 |
| Rassf5 | -2.876955151 | 1.63E-15 |
| Slc8b1 | -2.760671637 | 1.68E-15 |
| Pcdh11x | 4.885866765 | 2.30E-15 |
| Csf2rb | -2.535167109 | 2.49E-15 |
| mt-Co2 | 1.794662769 | 3.06E-15 |
| mt-Nd4 | 1.429582875 | 6.00E-15 |
| Tlr8 | -2.323072271 | 6.00E-15 |
| Akna | -1.820501527 | 6.52E-15 |
| Usp25 | -2.307858358 | 8.38E-15 |
| 6330416G13Rik | -2.486132328 | 1.20E-14 |
| Zfhx4 | 2.042319831 | 1.65E-14 |
| Emilin2 | -3.053147043 | 1.65E-14 |
| Prex1 | -2.207584861 | 1.73E-14 |
| Ptx3 | 2.618952642 | 2.50E-14 |
| Cpd | -2.151646459 | 3.17E-14 |
| Hes5 | 2.613293018 | 3.40E-14 |
| Ndn | 1.331409488 | 3.40E-14 |
| Irak3 | -2.745027509 | 3.56E-14 |
| Ptafr | -2.991149834 | 3.78E-14 |
| Serpinf1 | 5.209261661 | 3.96E-14 |
| Gsn | -2.379635986 | 4.33E-14 |
| Siglec1 | -3.366555663 | 4.75E-14 |
| Ppp6r1 | -2.141938185 | 6.26E-14 |
| Lacc1 | -2.495106819 | 6.26E-14 |
| Lrch1 | -1.675095062 | 7.63E-14 |
| Sema4c | -2.066500576 | 7.98E-14 |
| Tm9sf3 | -1.411474808 | 1.04E-13 |
| Atp5g1 | 1.304789367 | 1.29E-13 |
| H2afv | 1.601651234 | 1.40E-13 |
| Soat1 | -1.743169804 | 1.40E-13 |
| Gm5424 | -3.301628686 | 1.74E-13 |
| Msn | -1.866255508 | 3.25E-13 |
| Inpp5b | -2.252467021 | 3.38E-13 |
| Rras | -2.51362774 | 3.50E-13 |
| Pik3ap1 | -2.509473909 | 3.83E-13 |
| E2f1 | -2.002508292 | 3.97E-13 |
| Ecel1 | 5.379001034 | 4.00E-13 |
| Plce1 | 3.514450862 | 9.38E-13 |
| Snrpg | 1.325011782 | 1.25E-12 |
| Hivep3 | -1.631934554 | 1.53E-12 |
| Col4a1 | 3.488477069 | 1.63E-12 |
| Sel1l | -1.462511395 | 1.90E-12 |
| Sun2 | -1.735096402 | 2.37E-12 |
| Lpcat2 | -1.949987919 | 2.43E-12 |
| Tgfb1 | -2.835083184 | 2.70E-12 |
| AI593442 | 2.922937987 | 3.19E-12 |
| Aldoc | 2.050657853 | 3.39E-12 |
| Adgre4 | -5.956490171 | 3.44E-12 |
| Atp1a1 | -1.674242196 | 3.49E-12 |
| Ttyh3 | -1.65689265 | 3.61E-12 |
| C1rl | -6.613956606 | 3.61E-12 |
| S1pr3 | 3.816540201 | 3.97E-12 |
| Id1 | 2.843053988 | 4.30E-12 |
| Bex2 | 1.123591371 | 5.05E-12 |
| Zfp710 | -1.561966135 | 5.14E-12 |
| Efhd2 | -1.722625429 | 5.52E-12 |
| Cox6c | 1.14360851 | 5.58E-12 |
| Mdk | 1.317994705 | 6.46E-12 |
| Uqcrh | 1.099093189 | 6.93E-12 |

| gene | log2FoldChange | padj |
|---|---|---|
| S1pr4 | -3.376088903 | 0.026587512 |
| Dynlt1f | 0.698171385 | 0.02666422 |
| Lsm3 | 0.538136726 | 0.026669305 |
| Klhl40 | 5.517905657 | 0.026685616 |
| Ccdc113 | 4.057245062 | 0.026718183 |
| Szt2 | -0.934346488 | 0.026745733 |
| Ly6c1 | 4.703995154 | 0.026826862 |
| Zbtb38 | 0.54790336 | 0.026840697 |
| Olfr1351 | -6.413887298 | 0.026857366 |
| Fkbp1a | 0.374577742 | 0.026861499 |
| Kbtbd2 | -0.516770643 | 0.026861499 |
| Csf1 | -1.595738767 | 0.026861499 |
| Arhgap29 | 0.996879061 | 0.026910091 |
| Btbd2 | -0.561548496 | 0.026929854 |
| Il7r | -3.74432165 | 0.026929854 |
| Jrk | 1.525253989 | 0.026981288 |
| Ell | -1.153108227 | 0.027071369 |
| H2-Q6 | 2.690035392 | 0.027142719 |
| Cd47 | -0.596688679 | 0.027177931 |
| Fbxw8 | -0.723851843 | 0.027192841 |
| Rasa3 | -0.779228054 | 0.027192841 |
| Smyd4 | -0.864003825 | 0.027192841 |
| Pik3cb | -0.872652145 | 0.027192841 |
| Xpa | 0.843928055 | 0.027202776 |
| Smarcb1 | 0.469952325 | 0.027202776 |
| Mpzl2 | 5.354501404 | 0.027232497 |
| Pgls | 0.463127127 | 0.027273977 |
| Rbm33 | -0.559114222 | 0.027273977 |
| Ssbp1 | 0.441340239 | 0.027338787 |
| Mmaa | 0.931526458 | 0.027344768 |
| Ccdc30 | 1.41920869 | 0.027369574 |
| Zswim8 | -0.628982746 | 0.027376541 |
| Dnajc16 | -0.979931341 | 0.027414006 |
| Cnih3 | -1.055316522 | 0.027414006 |
| Anxa1 | -1.737044622 | 0.027420846 |
| Gabra4 | 1.688863695 | 0.027422224 |
| Prorsd1 | 1.049131702 | 0.027504384 |
| Mrps35 | 0.580259006 | 0.0275486 |
| Mib1 | 0.502805004 | 0.027644093 |
| Cept1 | -1.0573607 | 0.027644093 |
| BC052040 | 1.064369002 | 0.027743025 |
| Polr1d | 0.532457282 | 0.02784397 |
| Rad21 | 0.680794301 | 0.027874921 |
| Cks1b | 0.585153392 | 0.02789244 |
| Anxa4 | -1.089706707 | 0.02789244 |
| Mfap1a | 0.775272624 | 0.027900568 |
| Zfp335 | -0.888617691 | 0.027900568 |
| Alg2 | 0.523683394 | 0.028020189 |
| Wasf2 | -0.581062525 | 0.028028402 |
| Ssr1 | -0.463636474 | 0.028157499 |
| Atf1 | -0.717870843 | 0.028199253 |
| Yipf5 | 0.594946454 | 0.028282657 |
| Ptf1a | 5.285332223 | 0.028318764 |
| Slc41a3 | -1.225642529 | 0.028356893 |
| Tanc1 | 1.092326864 | 0.028378882 |
| Ehd2 | 2.144077523 | 0.028414419 |
| Cdkl5 | -0.840530323 | 0.028449738 |
| Acaa1a | -0.574279511 | 0.028491401 |
| Gramd1b | -0.825953821 | 0.028492097 |
| E130309D02Rik | -0.62043162 | 0.028493686 |
| Npas4 | 4.76788203 | 0.028591762 |
| 2010109I03Rik | -6.355561834 | 0.028629342 |
| Nudt13 | 1.205293086 | 0.028680631 |
| Scoc | 0.630807781 | 0.028778107 |
| Fkbp8 | -0.395007209 | 0.028957659 |
| Rsph4a | 3.398801052 | 0.028967039 |
| AU041133 | 1.159033815 | 0.028989542 |
| Ptgr1 | -0.933948266 | 0.029002683 |
| Thoc2 | 0.540756992 | 0.029007099 |
| Eif2ak3 | -1.130023468 | 0.029007099 |
| Kcns1 | 5.379304311 | 0.029029783 |
| 2210016L21Rik | 0.542827111 | 0.029043134 |
| Ier5 | -0.679510775 | 0.029043134 |
| Ube2v1 | -2.590858 | 0.029045241 |
| Mfsd6 | -0.901504443 | 0.02908018 |
| Aven | -1.086417185 | 0.02908018 |
| Ctdp1 | -0.852645925 | 0.029103457 |
| Col27a1 | 3.233917689 | 0.029105816 |
| 4930427A07Rik | 1.16899963 | 0.029105816 |
| Lage3 | 0.862539683 | 0.029105816 |
| Gatb | -1.054626128 | 0.029105816 |
| Abca8b | 3.733151816 | 0.029113802 |
| Lat2 | -1.107407462 | 0.029141129 |
| Praf2 | 0.795667602 | 0.029153769 |
| Commd2 | 0.506003603 | 0.029153769 |
| Tarsl2 | 1.017210243 | 0.029283963 |
| Cyb561d2 | 0.829367228 | 0.029353051 |
| Ankrd34b | 5.620568971 | 0.029367223 |
| Adarb1 | -0.954172898 | 0.029402215 |
| Limk2 | 0.562968599 | 0.029482088 |
| Ppp1r37 | -0.897866329 | 0.029506904 |
| Inpp5k | -0.602126354 | 0.029521084 |
| Dpagt1 | -0.635935412 | 0.029521084 |
| Abcc5 | -0.501432288 | 0.029582669 |
| Trak1 | -0.53940259 | 0.029583853 |
| B4galt6 | -0.780571693 | 0.02959327 |
| Kcns3 | 3.467351677 | 0.029597061 |
| Zrsr2 | 0.697889092 | 0.029597061 |
| Dip2b | -0.578180754 | 0.029597061 |
| Cdc42bpg | -2.39585218 | 0.029657433 |
| Cep72 | -1.094245589 | 0.029672289 |
| Psma1 | 0.452465791 | 0.029753816 |
| Itpk1 | -0.728559936 | 0.02976281 |
| Runx1t1 | 0.583186102 | 0.029778659 |
| Tom1 | -1.069586156 | 0.029778659 |
| Gltscr2 | 0.518357922 | 0.029846479 |
| Crim1 | -1.347707312 | 0.02985481 |
| 4930502E18Rik | -6.345106838 | 0.029864239 |
| Tsku | -1.000056684 | 0.030001531 |
| Mid1ip1 | 0.707993853 | 0.030088536 |
| Sbk1 | -0.403021379 | 0.030091258 |
| Uqcrfs1 | 0.4433287 | 0.030118721 |
| Zfp950 | 0.938743243 | 0.030126009 |
| Abcc3 | -1.377770391 | 0.030130481 |
| Zfp94 | -1.332289164 | 0.030140596 |

| gene | log2FoldChange | padj |
|---|---|---|
| Erp29 | -0.297825576 | 0.245582104 |
| Sccpdh | 0.274360279 | 0.246076293 |
| Unc5b | -1.130509052 | 0.246245781 |
| Zfp521 | 0.454069482 | 0.246474576 |
| Rhpn1 | 0.883417401 | 0.246524448 |
| Crb1 | 1.182672677 | 0.246536266 |
| Polr1e | 0.581985989 | 0.246541887 |
| Sec11c | -0.383728689 | 0.246541887 |
| Fam198a | -4.227578022 | 0.246997892 |
| Vsig2 | 4.169414227 | 0.247020998 |
| Ube2e3 | 0.265805987 | 0.247138333 |
| Ist1 | -0.273291 | 0.247138333 |
| Trim29 | -3.012425549 | 0.247165373 |
| Srpk3 | -1.07907097 | 0.247254294 |
| Rest | -0.377907221 | 0.247294542 |
| Ldlrad4 | -0.648342045 | 0.247310003 |
| Rccd1 | -0.805444069 | 0.247390194 |
| Otos | -4.964971287 | 0.247741345 |
| Tsr1 | 0.553626306 | 0.247914112 |
| Fam208a | 0.324401095 | 0.247914112 |
| Car5b | -3.08999384 | 0.247920604 |
| Magi2 | -0.424727701 | 0.248017287 |
| Zfp287 | -0.633707488 | 0.248017287 |
| Wfdc1 | -1.864917523 | 0.248017287 |
| Src | -0.466848225 | 0.248093089 |
| Pcgf5 | -0.474569069 | 0.248128109 |
| Ggt5 | -1.834897866 | 0.248161395 |
| Prmt3 | 0.377031183 | 0.248322478 |
| Usp4 | -0.257970305 | 0.248322478 |
| Ets2 | -0.572179547 | 0.24833729 |
| Tmem207 | 4.540457042 | 0.248466834 |
| Mzf1 | -0.957545709 | 0.248466862 |
| Tespa1 | -2.159943765 | 0.248494877 |
| Hspa4l | 0.342111791 | 0.248498763 |
| Nr1d1 | -0.690483843 | 0.248554087 |
| Zfp646 | -0.580329352 | 0.248596495 |
| Sdk1 | 1.614365224 | 0.248716949 |
| Lrba | 0.771552896 | 0.248933451 |
| Mtch2 | 0.241185185 | 0.249042049 |
| Mut | 0.420535747 | 0.249067307 |
| Hbs1l | 0.271351155 | 0.249067307 |
| Npr1 | 2.52473832 | 0.249177687 |
| Fam60a | -0.266187099 | 0.249248885 |
| Ddias | -0.597701785 | 0.249248885 |
| 1700113H08Rik | 4.120449708 | 0.249250474 |
| Hmgxb4 | 0.426653053 | 0.249282343 |
| Aldh7a1 | 0.477712305 | 0.249377801 |
| Cdc42bpa | 0.33119331 | 0.249377801 |
| Aurkaip1 | 0.285791883 | 0.249392906 |
| Nipal3 | 1.093248316 | 0.249555972 |
| Fam71e1 | 0.647986262 | 0.249555972 |
| Dnajb6 | 0.23777364 | 0.249555972 |
| Pdgfa | 0.875558464 | 0.249562006 |
| Rassf10 | 1.264549577 | 0.249738273 |
| Igf2r | -0.433915514 | 0.249738273 |
| Ddit4l | 2.317874982 | 0.249862701 |
| Gm10735 | 4.451013616 | 0.249968756 |
| Wfdc9 | 4.245733735 | 0.250048143 |
| Scamp5 | -0.357359488 | 0.250050223 |
| Eif2a | 0.326059757 | 0.250215538 |
| Tob2 | 0.68663707 | 0.250346233 |
| Asb10 | 2.485909256 | 0.250519378 |
| Foxred1 | -0.460328795 | 0.250519378 |
| Bsdc1 | -0.401092263 | 0.250639578 |
| Fam58b | 0.453848051 | 0.25070553 |
| Adat3 | -0.656423566 | 0.25070553 |
| Gtpbp6 | -0.699944595 | 0.250773035 |
| Rlbp1 | 0.525541915 | 0.250780476 |
| Cacng1 | -2.179957289 | 0.25088345 |
| Defb10 | 4.068829316 | 0.250991867 |
| Cplx2 | 0.360137589 | 0.250991867 |
| Cdc25c | 0.631482844 | 0.250996163 |
| Arfgef2 | -0.619805294 | 0.250999496 |
| Dynlrb2 | 1.428009234 | 0.251223571 |
| 9330182L06Rik | 0.597339526 | 0.251223571 |
| Actr10 | 0.232140593 | 0.251223571 |
| Carns1 | -0.991194684 | 0.251223571 |
| Cnnm2 | 0.577878263 | 0.251266182 |
| Enkd1 | 0.752684356 | 0.251296554 |
| Ccnc | 0.379193918 | 0.251308292 |
| Stkld1 | -3.252054449 | 0.2515005 |
| Tusc3 | 0.287996606 | 0.251599976 |
| Plekhg3 | -1.711073507 | 0.251693189 |
| Cyb561d1 | -0.629918819 | 0.251704922 |
| Msh5 | -1.034887796 | 0.251704922 |
| Ttc21a | 1.783986397 | 0.251852693 |
| Akap7 | 0.537937009 | 0.251852693 |
| Sav1 | -0.937654497 | 0.251876924 |
| Foxo3 | -0.311578645 | 0.251927192 |
| Riiad1 | 1.006939016 | 0.252123658 |
| Gm5814 | 4.042782423 | 0.252131191 |
| Slc5a1 | 4.180527782 | 0.252215176 |
| Rab34 | 0.407119598 | 0.252251249 |
| Cwc22 | -0.316039639 | 0.252304427 |
| Fam35a | -0.601369146 | 0.252419064 |
| Zdhhc7 | 0.440171224 | 0.252665991 |
| Wipi1 | 0.687880851 | 0.252812904 |
| Zfp87 | -0.897252537 | 0.252914139 |
| Kat8 | 0.523649438 | 0.253115555 |
| Npdc1 | 0.317377254 | 0.253115555 |
| Vmn2r56 | -4.935902247 | 0.25323829 |
| Rnf145 | -0.431751678 | 0.253247996 |
| Ankrd29 | 0.831598317 | 0.253275756 |
| Slc6a20a | 0.990510056 | 0.253451841 |
| Baz1b | -0.265245346 | 0.253464592 |
| 4930444P10Rik | 1.405725266 | 0.253475817 |
| Zbtb2 | -0.509061183 | 0.25374823 |
| Hmg20b | 0.438342766 | 0.253807228 |
| Map4 | -0.281423656 | 0.253807228 |
| Setd2 | -0.34164674 | 0.253807228 |
| Gm4131 | 3.795710663 | 0.254183572 |
| Kdm5b | -0.251121616 | 0.254280383 |
| Cpb2 | 4.102896805 | 0.254492295 |
| Spn | -2.375433771 | 0.254505719 |
| Stk11 | -0.321782059 | 0.254578911 |

| gene | log2FoldChang | padj |
|---|---|---|
| Dpp3 | -0.189201101 | 0.625888333 |
| Axin2 | 0.386892443 | 0.626008495 |
| Zfp81 | -0.160812555 | 0.626008495 |
| Dact1 | -0.253800674 | 0.626008495 |
| Lrrc61 | -0.318507322 | 0.626008495 |
| Dmrtc1a | -1.387207597 | 0.626071411 |
| Nosip | -0.150704897 | 0.626145762 |
| Emc3 | -0.147818312 | 0.626266176 |
| Fezf2 | -0.298456238 | 0.626266176 |
| Msc | 1.59570035 | 0.627118418 |
| Arhgef25 | 0.2442506 | 0.627118418 |
| Mettl17 | -0.261145501 | 0.627118418 |
| Dhx57 | -0.16295182 | 0.627404545 |
| Pex3 | 0.226033888 | 0.62742838 |
| Enox1 | 0.195814413 | 0.627627351 |
| Ube2l6 | -0.593898259 | 0.628031289 |
| A430105I19Ril | 0.737140106 | 0.628165964 |
| Jmjd6 | 0.15066025 | 0.628178206 |
| Ak9 | -1.702583418 | 0.628432219 |
| Mrpl43 | 0.131331262 | 0.628781983 |
| Farp2 | -0.272618606 | 0.628781983 |
| Fam46a | -0.315039157 | 0.628781983 |
| Omd | -0.668641135 | 0.628795203 |
| Zfp160 | 0.351844331 | 0.628862836 |
| Foxg1 | 0.157085246 | 0.628862836 |
| Ccnl2 | -0.156846261 | 0.628862836 |
| Atg16l2 | -0.208886824 | 0.628877729 |
| Rab11fip2 | -0.210070998 | 0.628976889 |
| BC017158 | 0.373413806 | 0.62950917 |
| B3galnt1 | 0.193004356 | 0.62950917 |
| Lztfl1 | -0.190446152 | 0.62950917 |
| Clspn | -0.328926397 | 0.62950917 |
| Nrg2 | -0.876846528 | 0.62950917 |
| Tacstd2 | -1.218785605 | 0.62950917 |
| Cma1 | 2.082070282 | 0.62965733 |
| Fam222a | 0.682919391 | 0.62965733 |
| Rhot1 | 0.152173477 | 0.62965733 |
| Ahcyl2 | -0.175224458 | 0.62965733 |
| Spryd4 | -0.233668585 | 0.63005885 |
| Myo16 | 0.284895567 | 0.630109452 |
| Myh6 | 1.057355461 | 0.630118374 |
| Zfp768 | 0.316072406 | 0.630240359 |
| Sh3d19 | 0.284704787 | 0.630443491 |
| Rnf185 | -0.234601788 | 0.630560257 |
| Nkx6-1 | 1.858166511 | 0.630968633 |
| Rab11a | 0.107583263 | 0.631166623 |
| Fmr1 | 0.169973609 | 0.631173232 |
| Gars | -0.118831285 | 0.631299914 |
| Gm3629 | 0.815636483 | 0.631307965 |
| Ranbp9 | 0.259696265 | 0.631307965 |
| Baz1a | 0.190259874 | 0.631307965 |
| Tmem39b | 0.182185331 | 0.631307965 |
| Nol6 | 0.142037163 | 0.631307965 |
| Ptrhd1 | -0.256815822 | 0.631307965 |
| Rasal2 | -0.151306669 | 0.631599018 |
| Prmt7 | 0.156048014 | 0.631678427 |
| Gnl3 | 0.195341909 | 0.631877423 |
| Yif1a | -0.174183139 | 0.631910195 |
| Mon2 | -0.23768999 | 0.632010128 |
| Hs2st1 | -0.19472623 | 0.632031988 |
| Cd1d2 | 1.567932523 | 0.632229364 |
| Sec22b | -0.117113443 | 0.632229364 |
| Hspa13 | -0.15637521 | 0.632229364 |
| Cyp2u1 | 0.31348042 | 0.632445508 |
| Hexdc | 0.283176721 | 0.632914807 |
| Garnl3 | 0.278697161 | 0.632914807 |
| Dohh | -0.158297202 | 0.632914807 |
| C2cd5 | -0.158651354 | 0.6332509 |
| Atp2b2 | -0.222792633 | 0.633379496 |
| Chst7 | -0.42529856 | 0.633379496 |
| Gm28372 | -1.630645387 | 0.633379496 |
| 1700023F06Ril | 1.441032588 | 0.63406238 |
| 1700093K21Ri | 1.609444965 | 0.634535894 |
| Mrpl44 | -0.181729262 | 0.634537531 |
| 1110037F02Ril | -0.142722433 | 0.635089611 |
| Rb1 | 0.202871723 | 0.635140304 |
| Zdhhc2 | 0.255934325 | 0.635219795 |
| Odf2 | -0.133251219 | 0.635219795 |
| Dnm3 | -0.23982136 | 0.635219795 |
| Hist1h2bq | 0.631220665 | 0.635237391 |
| Gm4799 | -0.733902308 | 0.635568769 |
| Kctd4 | 0.4025399 | 0.635578521 |
| Bahd1 | 0.369757947 | 0.635578521 |
| Cadm2 | 0.199806617 | 0.635578521 |
| Smim15 | 0.144493655 | 0.635578521 |
| Abcb6 | -0.202008379 | 0.635578521 |
| Brca1 | -0.221434052 | 0.635578521 |
| Gm17615 | -0.675537937 | 0.635578521 |
| Cabp7 | -1.036152759 | 0.635578521 |
| Phf19 | -0.655895454 | 0.635713375 |
| Mum1 | 0.161931691 | 0.635786653 |
| Timm8a1 | 0.212921153 | 0.635812854 |
| B3galnt2 | -0.362481344 | 0.635915244 |
| Gm14410 | 0.355912984 | 0.635965103 |
| Gucy1a2 | 0.327089899 | 0.63604795 |
| Atn1 | 0.164073619 | 0.636241483 |
| Nos3 | 0.618506555 | 0.636485664 |
| Adgrg3 | -0.844195909 | 0.636598703 |
| Rras2 | -0.247338831 | 0.636771531 |
| Zfp952 | -0.176789426 | 0.637455042 |
| Trpm4 | 0.375932188 | 0.63748559 |
| Ubl4a | -0.152362416 | 0.637528577 |
| Jazf1 | -0.256161212 | 0.63754868 |
| Zbtb12 | 0.521026326 | 0.637939154 |
| Kdm7a | -0.193089918 | 0.63821205 |
| Stk32a | -0.606260587 | 0.638590486 |
| Wdr74 | 0.154968905 | 0.639132665 |
| Myo1b | 0.129097225 | 0.639436353 |
| Rplp0 | 0.114525784 | 0.639436353 |
| Lrrn1 | 0.256561472 | 0.639583907 |
| Slfn9 | 0.250023217 | 0.639583907 |
| Mllt1 | -0.19571597 | 0.639583907 |
| Sh2b3 | -0.310645504 | 0.639583907 |
| Ccdc14 | -0.382322975 | 0.639583907 |
| Xkr5 | -0.588618556 | 0.639583907 |

| | | |
|---|---|---|
| Id4 | 3.592035052 | 7.51E-12 |
| Fkbp3 | 1.180242235 | 8.19E-12 |
| Gstm5 | 1.4751183 | 8.64E-12 |
| Pfkfb3 | -2.200891155 | 1.18E-11 |
| Nadk | -1.413326329 | 1.21E-11 |
| Ndrg2 | 1.537877679 | 1.26E-11 |
| Mfap2 | 1.618489712 | 1.68E-11 |
| Mapk14 | -1.815643948 | 1.74E-11 |
| Mest | 1.69452196 | 1.76E-11 |
| Arf3 | -1.465716487 | 1.84E-11 |
| Mpeg1 | -1.880458289 | 2.17E-11 |
| Gk | -2.095084635 | 2.31E-11 |
| Coro1b | -1.520904296 | 2.48E-11 |
| Map3k3 | -1.980737357 | 2.80E-11 |
| Scarf2 | -4.516552094 | 2.93E-11 |
| Tmem104 | -2.063718682 | 3.20E-11 |
| Mfsd12 | -3.111467841 | 3.35E-11 |
| Pygl | -2.597973786 | 3.35E-11 |
| Rpl23 | 1.071949781 | 3.42E-11 |
| mt-Nd6 | 1.24164689 | 3.72E-11 |
| Pik3r5 | -2.716520934 | 3.74E-11 |
| Hsp90aa1 | 1.15246506 | 3.87E-11 |
| Gna15 | -2.262361719 | 4.22E-11 |
| Lrrc8d | -1.670952713 | 4.26E-11 |
| Ltb4r1 | -4.614006114 | 4.52E-11 |
| Fign | 3.256083909 | 4.86E-11 |
| Nes | 2.391525073 | 4.96E-11 |
| 2700094K13Rik | 1.14711168 | 5.33E-11 |
| Sbno2 | -2.014224066 | 5.68E-11 |
| Anxa6 | -2.237604107 | 5.98E-11 |
| Piezo1 | -3.010503087 | 6.26E-11 |
| Usmg5 | 1.155512132 | 6.71E-11 |
| Rps27 | 1.407649859 | 6.80E-11 |
| Bex4 | 1.365608108 | 6.80E-11 |
| Trmt44 | -2.177746641 | 6.80E-11 |
| P2ry2 | -4.204027074 | 6.80E-11 |
| Nkain4 | 2.45278701 | 7.73E-11 |
| Aldh1a2 | 9.028747124 | 9.20E-11 |
| Gng11 | 2.778625706 | 9.55E-11 |
| Zdhhc5 | -1.857223477 | 1.02E-10 |
| Grn | -1.846653504 | 1.08E-10 |
| Kirrel2 | 8.808032712 | 1.12E-10 |
| Slc1a3 | 1.864500568 | 1.12E-10 |
| Camta2 | -1.785384618 | 1.13E-10 |
| Hist3h2ba | 1.470524261 | 1.21E-10 |
| Nr2e1 | 2.615746135 | 1.37E-10 |
| Nap1l5 | 1.908825593 | 1.38E-10 |
| Stxbp2 | -1.798111391 | 1.39E-10 |
| Pten | -1.191279055 | 1.44E-10 |
| Fosl2 | -2.453771998 | 1.48E-10 |
| Hist1h2ao | 1.947792798 | 1.56E-10 |
| Nlrc3 | -4.282756578 | 1.78E-10 |
| Fstl1 | 2.257810969 | 1.85E-10 |
| Scrn1 | 1.559301498 | 1.96E-10 |
| Fxyd6 | 1.407212441 | 2.06E-10 |
| Cyth4 | -2.177015518 | 2.20E-10 |
| Dnajc14 | -1.374500507 | 2.23E-10 |
| Wdr1 | -1.554053395 | 2.23E-10 |
| Tmsb10 | 1.150722419 | 2.77E-10 |
| Agps | -1.621770574 | 2.77E-10 |
| Trim8 | -1.658744021 | 2.77E-10 |
| Ero1l | -2.489666263 | 2.77E-10 |
| Itgal | -2.646394645 | 3.28E-10 |
| Ramp2 | 4.401014639 | 3.38E-10 |
| Sst | 3.372272558 | 3.77E-10 |
| Zc3h12a | -2.88276251 | 3.88E-10 |
| Pnmal2 | 1.372582061 | 3.90E-10 |
| mt-Co1 | 1.069107383 | 3.95E-10 |
| Uba7 | -2.578836788 | 4.33E-10 |
| Lum | 9.092301931 | 4.38E-10 |
| Il21r | -3.28308795 | 4.45E-10 |
| Btk | -1.994208424 | 4.51E-10 |
| Il4ra | -1.875222933 | 4.56E-10 |
| Aff1 | -2.384751518 | 4.59E-10 |
| Ptpn6 | -2.062520435 | 4.76E-10 |
| Bgn | 8.835698721 | 5.81E-10 |
| Slc11a1 | -1.976325533 | 5.81E-10 |
| Arhgap27 | -3.240612223 | 5.81E-10 |
| Ly6g | -9.821742139 | 5.81E-10 |
| Scube1 | -1.903238619 | 6.42E-10 |
| Clec12a | -1.674310298 | 6.79E-10 |
| Pax6 | 2.350973269 | 7.51E-10 |
| Acp5 | -2.299427353 | 8.62E-10 |
| Dnm2 | -1.335761087 | 8.98E-10 |
| Diaph2 | -1.688662488 | 9.33E-10 |
| Atp1b1 | 1.651422291 | 9.43E-10 |
| Tmem38b | -1.815917692 | 9.81E-10 |
| Ikbkb | -1.374833581 | 9.84E-10 |
| Mapkapk2 | -2.167119093 | 1.06E-09 |
| Pqbp1 | 1.53010605 | 1.09E-09 |
| Arl4a | 1.776361352 | 1.11E-09 |
| Tmem170b | -1.32086607 | 1.11E-09 |
| Car2 | 2.941433595 | 1.15E-09 |
| Cln3 | -1.703554523 | 1.19E-09 |
| Efna5 | 1.742009517 | 1.22E-09 |
| Zbtb20 | 1.17675401 | 1.22E-09 |
| Sec24d | -1.862484265 | 1.22E-09 |
| Pram1 | -3.152475434 | 1.22E-09 |
| Ralgds | -1.566512193 | 1.35E-09 |
| Slc7a8 | -1.873352931 | 1.42E-09 |
| Slc37a2 | -2.460933241 | 1.48E-09 |
| Vav1 | -2.008636283 | 1.53E-09 |
| Tpm1 | 1.407729713 | 1.57E-09 |
| Ppic | 1.981591325 | 1.74E-09 |
| Igsf6 | -2.099673193 | 1.91E-09 |
| Ap5b1 | -3.409600456 | 1.91E-09 |
| Abcc9 | 8.675406448 | 2.16E-09 |
| Raver2 | -2.318762542 | 2.17E-09 |
| Folr1 | 4.373993889 | 2.40E-09 |
| Tgm2 | -2.363173173 | 2.48E-09 |
| Scg2 | 2.705718018 | 2.55E-09 |
| Fuca2 | -1.954165287 | 2.78E-09 |
| Ppp1r3b | -3.160440268 | 2.89E-09 |
| Psd4 | -3.291168873 | 2.94E-09 |
| Dgkz | -1.79119541 | 3.02E-09 |
| Cox7c | 1.021361322 | 3.33E-09 |
| Vmp1 | -1.282088666 | 3.40E-09 |
| Tex15 | 5.725233864 | 3.40E-09 |
| Rac3 | 1.44144998 | 3.44E-09 |

| | | |
|---|---|---|
| Ptprm | -1.155800731 | 0.030156478 |
| Fxn | 0.641776548 | 0.030166062 |
| Tnxb | 2.903479309 | 0.030173702 |
| Meis3 | 0.581758946 | 0.030193068 |
| Kmt2e | 0.449402111 | 0.030201038 |
| Dnajc8 | 0.436128136 | 0.030201038 |
| Zfp777 | -1.175736479 | 0.030269851 |
| Hmgcll1 | 0.994377123 | 0.030316683 |
| Pradc1 | 0.880539705 | 0.030316683 |
| Trak2 | 0.814641224 | 0.030316683 |
| Mbnl2 | -0.722388356 | 0.030316683 |
| Ism1 | -4.398045848 | 0.030316683 |
| Ick | 0.583157199 | 0.030318819 |
| Atp6v0c | -0.905787474 | 0.030321878 |
| Ttc4 | 0.704597895 | 0.030346055 |
| Dmtn | 0.996241385 | 0.030426817 |
| Arrdc2 | -1.321825626 | 0.030448031 |
| Lmtk3 | -1.214502832 | 0.030478467 |
| Tstd2 | -0.792120611 | 0.030509764 |
| Zfp174 | 3.056391565 | 0.030562522 |
| Nat10 | -0.726729997 | 0.030568818 |
| Kcnq4 | -2.56230124 | 0.030611803 |
| Kif3c | -0.530925587 | 0.030646292 |
| Itgb7 | -2.037508044 | 0.030655013 |
| Susd1 | -2.294133745 | 0.030668058 |
| Itgb8 | 1.066743532 | 0.030690463 |
| Ppid | 0.471408649 | 0.030690463 |
| Hn1 | 0.407528753 | 0.030690463 |
| Rundc3b | -0.697650867 | 0.030690463 |
| Hnrnpa1 | 0.364828118 | 0.030760213 |
| S100a9 | -2.999523973 | 0.030760217 |
| R3hcc1l | -0.920029061 | 0.030826321 |
| Dpys | -1.980554951 | 0.030860426 |
| Lnx2 | -0.87259424 | 0.030941791 |
| Tlr6 | -1.391500604 | 0.03109943 |
| Clec4d | -2.66030781 | 0.031106195 |
| Tmem231 | 1.876017943 | 0.031157698 |
| Arhgef6 | -2.217443159 | 0.031157698 |
| Tmem151b | -0.546678448 | 0.031170818 |
| Znrd1 | 0.593926668 | 0.031276048 |
| Itgb2l | -5.997349945 | 0.031276048 |
| Tmem218 | 0.892597403 | 0.031287602 |
| Med13l | 0.679313507 | 0.03129842 |
| Crk | -0.428381774 | 0.031345168 |
| Elp2 | -0.444712264 | 0.031353537 |
| Nup88 | 0.491593375 | 0.031436015 |
| Hspb1 | 1.816362028 | 0.031467059 |
| Xylt1 | -1.539548741 | 0.031502595 |
| Whsc1 | -0.40312528 | 0.031557012 |
| C1qtnf6 | 3.874548395 | 0.031590242 |
| Metap2 | 0.387679735 | 0.031590242 |
| Atp6v1g2 | 0.803325221 | 0.031594058 |
| Gm5617 | 1.092769263 | 0.031686092 |
| Cldn9 | 5.280066171 | 0.0317009 |
| Thsd7a | 1.363714558 | 0.0317009 |
| Ncbp1 | 0.545757865 | 0.0317009 |
| Slc25a22 | -0.689196708 | 0.0317009 |
| Dopey2 | -0.688926001 | 0.031730707 |
| Cacna1g | 1.301107565 | 0.031788377 |
| Wnt8b | 2.740851437 | 0.031815337 |
| Timm9 | 0.731724691 | 0.031815337 |
| Hnrnpul1 | -0.478109092 | 0.031815337 |
| Irf9 | -1.200872966 | 0.031832679 |
| Prkar1a | -0.366430899 | 0.031890917 |
| Pus10 | -0.813249922 | 0.031947061 |
| AI429214 | -1.221382792 | 0.031955624 |
| Aars2 | -1.215732808 | 0.031980755 |
| Th | 5.005283957 | 0.032019479 |
| Cdc42ep5 | 2.879950738 | 0.032077129 |
| Pttg1ip | -0.650870672 | 0.032107864 |
| Tsen34 | 0.572657177 | 0.032227742 |
| Glud1 | -0.444439581 | 0.032317109 |
| Mt1 | 0.569167189 | 0.032359056 |
| Fam134a | -0.498182763 | 0.032360259 |
| Pcyt1a | -0.702444868 | 0.032367173 |
| Ndc1 | -0.931271084 | 0.032372495 |
| Recql5 | -0.942277095 | 0.032372495 |
| Bmpr1b | 2.241934347 | 0.03237786 |
| Pacsin3 | 1.756224344 | 0.032426212 |
| Aaed1 | -0.975710806 | 0.032426212 |
| Cops4 | 0.481974104 | 0.032472783 |
| Ms4a6c | -1.525649557 | 0.032472783 |
| Ugt8a | 5.418203081 | 0.032495293 |
| Pbdc1 | 0.545142994 | 0.032517835 |
| Ecscr | 2.630814386 | 0.03253025 |
| Ms4a4a | -0.97876369 | 0.032568546 |
| Mfsd7b | -1.088232334 | 0.032568546 |
| Cript | 0.468955253 | 0.03258993 |
| Prrx1 | 1.578837755 | 0.032648695 |
| Phf24 | -0.89358123 | 0.032648695 |
| Eapp | 0.555388168 | 0.032675021 |
| Csnk1g2 | -0.783111286 | 0.032678177 |
| Hist1h3e | -1.373660691 | 0.032739079 |
| Dnhd1 | -1.519271396 | 0.032779161 |
| Iqcg | 1.721228615 | 0.032844639 |
| Mob4 | 0.43680037 | 0.032844639 |
| Cep78 | 0.696464125 | 0.03287027 |
| Mtap7d3 | 5.239939952 | 0.032895636 |
| Ghr | 2.731717435 | 0.032927766 |
| Mapk7 | -0.676760333 | 0.032980501 |
| Tctn1 | -1.181507645 | 0.032989115 |
| Gm11733 | -3.664999776 | 0.032989115 |
| Imp3 | 0.647690763 | 0.032996519 |
| Smc4 | 0.609276394 | 0.033029452 |
| Slco1a4 | 4.770670061 | 0.033030776 |
| Zfp318 | 0.689115385 | 0.033051282 |
| Dpp9 | -0.622788556 | 0.033058149 |
| Calcr | 5.626240599 | 0.033074985 |
| Vwc2 | 3.940256562 | 0.033083725 |
| Aqp4 | 4.994810728 | 0.033086949 |
| Agpat2 | -1.527012867 | 0.033086949 |
| Ube2d1 | 0.459808831 | 0.033112632 |
| Chmp4b | -0.819887112 | 0.033298071 |
| Ap4s1 | 0.681997801 | 0.033318619 |
| Arhgap23 | -0.786430338 | 0.033424578 |
| Chad | 5.411054259 | 0.033584711 |
| Efr3a | -0.521557573 | 0.033603377 |
| Emc6 | 0.426594419 | 0.033603994 |
| Cd300a | -0.911779694 | 0.033610866 |

| | | |
|---|---|---|
| Tln2 | 0.502602159 | 0.254611615 |
| Lrrcc1 | 0.457846145 | 0.254713541 |
| Zfp449 | 0.934443563 | 0.254835821 |
| Zfp959 | 0.581917107 | 0.254835821 |
| Zfp91 | 0.362616864 | 0.254936828 |
| Llgl2 | -1.524873475 | 0.254962252 |
| Mypop | -1.036765756 | 0.25522493 |
| Gls2 | 2.88382709 | 0.25522572 |
| Pspn | 2.535397543 | 0.25522572 |
| Ndufaf3 | 0.293940068 | 0.25522572 |
| Batf2 | -1.974160775 | 0.255430404 |
| Lpo | 4.469431227 | 0.255436819 |
| Gstk1 | 0.672305394 | 0.255541276 |
| Sorbs2 | -0.270456787 | 0.255646602 |
| Zfp747 | -1.108067067 | 0.255646602 |
| Smarcc1 | -0.232049511 | 0.255716397 |
| Tnfrsf17 | 4.02163104 | 0.255833173 |
| Icam2 | 1.24426887 | 0.255833173 |
| Ids | -0.435353602 | 0.255833173 |
| Soga3 | 0.299457029 | 0.255976858 |
| Flcn | -0.336070996 | 0.256010161 |
| Sox14 | 4.013706523 | 0.256179505 |
| Anp32a | 0.31048064 | 0.256469613 |
| Acad11 | -0.829505893 | 0.256469613 |
| Agbl5 | -0.607699394 | 0.256528095 |
| Mrpl38 | 0.466816118 | 0.256538033 |
| Gm4951 | 2.058067601 | 0.25658021 |
| Lhfpl5 | -1.280655603 | 0.256617235 |
| Actr5 | -0.649968309 | 0.256656965 |
| Calcrl | 1.241554753 | 0.256787364 |
| Myb | 1.172550832 | 0.257119376 |
| 1700029I15Rik | -1.57879764 | 0.257119376 |
| Naprt | -0.96222707 | 0.257180406 |
| Fzd4 | 2.256217676 | 0.257563146 |
| Ehd3 | 0.503437485 | 0.257926301 |
| Acbd5 | -0.453441109 | 0.258089801 |
| Prkar2b | -0.359619378 | 0.258171637 |
| Grm3 | -0.880510288 | 0.258182477 |
| Obsl1 | 0.635507977 | 0.258188361 |
| Pcdhb4 | 3.420060768 | 0.258249692 |
| Arpp19 | 0.248540637 | 0.258256939 |
| Paics | 0.293548811 | 0.258302186 |
| Dync1i1 | -1.023628126 | 0.258315402 |
| Tmem181a | -0.46404313 | 0.258318098 |
| Chrm4 | -2.938135007 | 0.258684505 |
| Msl3l2 | 0.622021065 | 0.258789587 |
| Asap3 | 1.095523433 | 0.258791732 |
| Col24a1 | 3.70942208 | 0.258864588 |
| Rprm | 0.493795293 | 0.258864588 |
| Cic | -0.40330189 | 0.258864588 |
| Fam166a | -2.314810588 | 0.258979601 |
| Anks1 | -0.38667686 | 0.259077382 |
| Fam47e | 4.090363943 | 0.259172335 |
| Cyp26b1 | 4.073927339 | 0.259214111 |
| Gdpd1 | 0.290968107 | 0.259214111 |
| F8 | 2.544276118 | 0.259394939 |
| Tpm2 | -0.795926609 | 0.259410276 |
| Slc10a3 | -1.064841411 | 0.259502418 |
| Cfap58 | 4.492820154 | 0.259817333 |
| Nox3 | 4.492820154 | 0.259817333 |
| Mars | 0.305307287 | 0.259817333 |
| Tars2 | -0.351995644 | 0.259817333 |
| Ctla2b | -1.157156286 | 0.259817333 |
| 4930486L24Rik | -4.926550632 | 0.259817333 |
| Shisa8 | 1.943406025 | 0.260127218 |
| Trpc1 | -0.726954578 | 0.260127869 |
| Nod1 | -1.019828459 | 0.260296832 |
| Prss48 | -3.083664954 | 0.260359031 |
| Gpatch2 | 0.519161596 | 0.260366827 |
| Cyp4a12b | 4.041249541 | 0.260722649 |
| Cep126 | 1.495436362 | 0.260722649 |
| Psmd6 | 0.240520384 | 0.260780762 |
| E2f2 | -0.790372034 | 0.260822364 |
| Wdr3 | -0.389162175 | 0.260922393 |
| Tbl2 | -0.470971578 | 0.261046807 |
| Adam1b | -4.906253393 | 0.261046807 |
| Il1f6 | -4.906253393 | 0.261046807 |
| Il22ra1 | -4.906253393 | 0.261046807 |
| Tgtp1 | -4.906253393 | 0.261046807 |
| Kdr | 0.829698105 | 0.261082899 |
| Aqp9 | -2.176392506 | 0.261082899 |
| Matn2 | 0.999940622 | 0.261115907 |
| Sh2d2a | -3.273368161 | 0.261115907 |
| Grb7 | 4.030016678 | 0.261182761 |
| Cep290 | 0.425033777 | 0.261272044 |
| Cnksr2 | -0.514044453 | 0.261365467 |
| Stau1 | 0.28470043 | 0.261397986 |
| Ccdc22 | -0.369802811 | 0.261409033 |
| Slc35a2 | -0.500607281 | 0.261603264 |
| Med13 | 0.32315954 | 0.262063239 |
| Mrap2 | 4.120943704 | 0.262157549 |
| Dennd2a | -0.471573605 | 0.262157549 |
| Stard3nl | 0.251624107 | 0.26221923 |
| Aldh1l1 | 0.606469649 | 0.262367152 |
| Tprgl | -0.28902515 | 0.262706474 |
| Tmem30a | -0.321380121 | 0.262718934 |
| Rpl10-ps3 | 0.535430991 | 0.262806696 |
| Rabif | -0.296302534 | 0.263018125 |
| Tnpo3 | -0.326634691 | 0.263018125 |
| Stard4 | -0.598479245 | 0.263018125 |
| Oas1g | -1.682833998 | 0.263018125 |
| Elovl4 | 0.367534099 | 0.263023249 |
| BC100451 | 2.465709871 | 0.263036833 |
| Tsks | 3.693166703 | 0.26325395 |
| Etv1 | 0.58003869 | 0.263307526 |
| Kcna1 | -1.015981255 | 0.263307526 |
| P2ry13 | 1.282996812 | 0.263320632 |
| Tmem183a | 0.251656592 | 0.263360051 |
| Vps4b | -0.31036202 | 0.263375483 |
| Fkbp1b | 0.509679907 | 0.263401426 |
| Nudt18 | 0.482665887 | 0.263580407 |
| Klc1 | 0.248932272 | 0.263604963 |
| BC003331 | -0.336023315 | 0.263604963 |
| Srrt | 0.293397828 | 0.263695597 |
| Gdpd5 | -0.430598769 | 0.263695597 |
| Rspo3 | -0.584118409 | 0.263735041 |
| Trp53i11 | 0.331797986 | 0.263770798 |
| Gtf3c2 | 0.50994138 | 0.263851585 |
| Pigu | 0.489685524 | 0.263979248 |

| | | |
|---|---|---|
| Slc25a42 | 0.516446254 | 0.639963979 |
| Micu1 | -0.165952749 | 0.639963979 |
| Arsb | 0.220071677 | 0.639973271 |
| Ampd2 | -0.163046148 | 0.640000991 |
| Hr | 0.638848202 | 0.640203318 |
| Gabrr2 | -1.656934821 | 0.640203318 |
| Gypc | 0.309218537 | 0.640247253 |
| Kif15 | 0.166802726 | 0.640247253 |
| Flot2 | -0.147886726 | 0.640247253 |
| Mpst | 0.179710835 | 0.640280752 |
| R3hdm1 | 0.125096261 | 0.640280752 |
| Tmem167 | 0.130547339 | 0.640538599 |
| Atp4a | 1.332312133 | 0.640951543 |
| Frzb | 0.826925834 | 0.641073685 |
| 1700037C18Ri | 0.445231763 | 0.641073685 |
| Zfp472 | 0.363503464 | 0.641073685 |
| Mthfsl | 0.194624712 | 0.641073685 |
| Matk | -0.230323198 | 0.641073685 |
| Tmem74b | -0.269158962 | 0.641073685 |
| Ifit2 | -0.53157293 | 0.641073685 |
| Ptgis | -0.835397531 | 0.641073685 |
| 2810408A11Ri | 0.231786081 | 0.641097367 |
| Doc2b | 0.944020475 | 0.641116482 |
| Ryr2 | 0.619455103 | 0.641116482 |
| Gan | -0.195575767 | 0.641117214 |
| Gpr160 | 0.725232341 | 0.641214579 |
| Pcsk2 | 0.378677069 | 0.641509506 |
| Aar2 | -0.149879329 | 0.641523513 |
| Drd1 | -0.689401848 | 0.641523513 |
| Cc2d1a | 0.274833772 | 0.641956358 |
| Smarca4 | 0.114064937 | 0.64204392 |
| Fance | 0.219243542 | 0.642091684 |
| Prx | 0.713775881 | 0.642217249 |
| Uevld | 0.313745207 | 0.64233628 |
| Btbd6 | 0.23766779 | 0.64233628 |
| Vprbp | -0.238432586 | 0.64233628 |
| Pex5l | 0.582459088 | 0.642536681 |
| Il2ra | -1.596979165 | 0.642536681 |
| Plcd1 | 0.318994711 | 0.64299875 |
| Vim | -0.215357408 | 0.643238462 |
| Cdh1 | -0.627193663 | 0.643238462 |
| Lmntd2 | 1.935142084 | 0.643842255 |
| Cpxm2 | 0.780309858 | 0.643905227 |
| Slc7a4 | 0.297949649 | 0.644309846 |
| Rpl36-ps3 | 1.103423582 | 0.644348337 |
| Xlr3c | 0.909835738 | 0.644348337 |
| Pyroxd2 | -0.397558774 | 0.644382394 |
| Usp2 | 0.24709016 | 0.644901046 |
| Fam189a2 | -1.423731668 | 0.644901046 |
| Slc7a2 | -0.588044963 | 0.645206437 |
| Slc25a37 | -0.376120065 | 0.645526859 |
| Tmem9b | 0.174902764 | 0.64586016 |
| Rfc5 | -0.166637412 | 0.64586016 |
| Cep295 | -0.181578193 | 0.64586016 |
| Nucks1 | 0.117836067 | 0.646171833 |
| Slc1a6 | 0.494172852 | 0.646316661 |
| Ahdc1 | -0.242657026 | 0.646316661 |
| Mro | 0.76064676 | 0.646354028 |
| Cyth1 | -0.216199759 | 0.646354028 |
| 4921507P07Ril | -0.765898131 | 0.646354028 |
| Ggps1 | 0.211617891 | 0.646428775 |
| Ccdc50 | -0.131008232 | 0.646432825 |
| Ptpn4 | -0.253354019 | 0.646432825 |
| Rnf17 | -2.126549808 | 0.646432825 |
| Prrg4 | 2.26757648 | 0.646478822 |
| Rnd2 | -0.162572122 | 0.646478822 |
| Gab3 | -0.366100914 | 0.646611328 |
| Hsdl1 | -0.124225327 | 0.646845351 |
| Sycp1 | -1.294846649 | 0.646882687 |
| Fam228a | -0.996968962 | 0.64728867 |
| Skiv2l | -0.173636273 | 0.647332332 |
| Cep85 | 0.243694689 | 0.647349195 |
| Cuedc1 | -0.171072299 | 0.647363024 |
| Spr | 0.175234839 | 0.647369284 |
| Glipr2 | -0.248909468 | 0.647369284 |
| Zscan2 | -0.418747315 | 0.647369284 |
| Aspscr1 | 0.122137763 | 0.647673362 |
| Ndufs1 | -0.128129517 | 0.647904708 |
| Cidea | -1.058176172 | 0.647990934 |
| BC021785 | 2.206399184 | 0.648000055 |
| Foxl2 | 2.206399184 | 0.648000055 |
| Naa15 | 0.137524104 | 0.648048169 |
| Med4 | 0.141101043 | 0.648075914 |
| Fbxl19 | 0.255538069 | 0.648155214 |
| Eps15 | -0.184786433 | 0.64818524 |
| Cryz | 0.217296914 | 0.648361265 |
| Zak | -0.228867026 | 0.648361265 |
| Acss3 | 1.65513514 | 0.648693562 |
| Eef2kmt | 0.253172907 | 0.648693562 |
| Mynn | 0.202537715 | 0.648693562 |
| Clk4 | 0.132446287 | 0.648693562 |
| Gm9920 | 1.879023667 | 0.648961966 |
| Ccdc167 | 0.643019428 | 0.648961966 |
| Anapc2 | -0.15039261 | 0.649088417 |
| Gm5468 | -0.407072856 | 0.649088417 |
| Nfib | -0.169114974 | 0.649368224 |
| Cldn19 | -0.877065139 | 0.649458946 |
| Bean1 | -0.463721701 | 0.64951928 |
| Gm10837 | 0.899894149 | 0.65007893 |
| Tvp23b | -0.158595158 | 0.650521947 |
| Epha5 | 0.180017355 | 0.650524096 |
| Tcp1 | -0.105327474 | 0.650524096 |
| Thop1 | -0.165050351 | 0.650586696 |
| Gas6 | 0.255937042 | 0.650610764 |
| Faap100 | -0.14018412 | 0.650610764 |
| Edn3 | 0.668763012 | 0.650622796 |
| Tnnt1 | -0.32295572 | 0.650742171 |
| Nvl | -0.149934697 | 0.650879712 |
| Tmem208 | 0.13723886 | 0.650922999 |
| Klhl9 | 0.101986708 | 0.651241104 |
| Ewsr1 | -0.100256184 | 0.651241104 |
| Calhm2 | -0.683030637 | 0.651248995 |
| Gm21886 | -0.617969163 | 0.651770416 |
| Car3 | -0.898272828 | 0.651810295 |
| Rheb | 0.10794812 | 0.652038445 |
| Tle1 | -0.129903393 | 0.652060071 |
| Cd5 | -0.44944082 | 0.652617654 |
| Prdm10 | -0.18945264 | 0.652824815 |
| Myo5b | 0.625925722 | 0.652846827 |

| | | |
|---|---|---|
| Ptgds | 2.537059468 | 3.84E-09 |
| Blcap | 1.100585059 | 4.19E-09 |
| Tbc1d13 | -1.318713246 | 4.19E-09 |
| Itgb2 | -2.146091339 | 4.19E-09 |
| Pparg | -9.601147392 | 4.19E-09 |
| Actn4 | -1.360311664 | 4.28E-09 |
| Ttyh1 | 1.454788322 | 4.30E-09 |
| Prok2 | -4.286491412 | 4.47E-09 |
| Acp2 | -1.361852632 | 4.64E-09 |
| Arhgap26 | -2.216129027 | 4.66E-09 |
| Atpif1 | 1.013188806 | 4.69E-09 |
| Hmgcs2 | 8.477214522 | 4.71E-09 |
| Spag5 | 2.642445942 | 5.22E-09 |
| Stat6 | -2.215572355 | 5.29E-09 |
| Nckap1l | -2.031151377 | 6.09E-09 |
| Pcsk7 | -1.694818995 | 6.20E-09 |
| Cnksr3 | -2.632703473 | 6.39E-09 |
| Tor1aip1 | -1.735881486 | 6.56E-09 |
| Rinl | -2.589899481 | 6.75E-09 |
| Nefm | 2.202982805 | 6.96E-09 |
| C130050O18Rik | -2.631133305 | 7.02E-09 |
| Sox21 | 2.148289271 | 7.44E-09 |
| Ndufa4 | 1.013803808 | 7.54E-09 |
| Ehbp1l1 | -1.690388779 | 7.54E-09 |
| Ptn | 1.604386037 | 7.77E-09 |
| Polr2l | 1.381163887 | 8.04E-09 |
| Capza1 | -1.386190936 | 8.08E-09 |
| Pmp22 | 2.450324715 | 8.39E-09 |
| Chpf2 | -1.777242388 | 8.97E-09 |
| Adar | -1.465661889 | 1.00E-08 |
| Plxnb2 | -1.15159105 | 1.03E-08 |
| 15-Sep | 1.523743786 | 1.06E-08 |
| Snx18 | -1.767137732 | 1.08E-08 |
| Nova1 | 2.553287582 | 1.12E-08 |
| Mtmr3 | -1.24927398 | 1.16E-08 |
| Col4a2 | 2.228556859 | 1.18E-08 |
| 1600014C10Rik | -2.09531261 | 1.21E-08 |
| Tnfaip8 | -1.68638331 | 1.25E-08 |
| Adam17 | -1.527089121 | 1.35E-08 |
| Lamp1 | -1.331431319 | 1.42E-08 |
| Rcn3 | 2.026546988 | 1.44E-08 |
| Eno1b | -1.300074067 | 1.49E-08 |
| Plekha7 | 5.17868386 | 1.54E-08 |
| Lrrc59 | -1.480352556 | 1.58E-08 |
| Akap9 | 1.034973386 | 1.61E-08 |
| 1500015O10Rik | 5.855248051 | 1.61E-08 |
| Slc15a3 | -2.197869472 | 1.61E-08 |
| Esrra | -2.024082239 | 1.64E-08 |
| Flna | -1.313863917 | 1.71E-08 |
| Il18rap | -8.097252116 | 1.72E-08 |
| Cntn5 | 3.260876373 | 1.74E-08 |
| Spc25 | 1.4126405 | 1.81E-08 |
| Gm9844 | 1.684845105 | 1.85E-08 |
| Arpc1b | -1.594347804 | 1.86E-08 |
| Zfp869 | -1.584666899 | 1.90E-08 |
| Samhd1 | -2.095886423 | 1.92E-08 |
| Lsm5 | 1.443726354 | 1.95E-08 |
| Abhd8 | 1.251849068 | 2.11E-08 |
| Arid5a | -2.036010607 | 2.32E-08 |
| Actb | -1.177541045 | 2.41E-08 |
| Plbd2 | -1.483024706 | 2.41E-08 |
| 1110001J03Rik | 1.379309369 | 2.43E-08 |
| Clu | 1.795250118 | 2.43E-08 |
| Esco2 | 1.729584228 | 2.44E-08 |
| Phyhipl | 1.241086273 | 2.44E-08 |
| mt-Nd5 | 0.917464274 | 2.44E-08 |
| Slc6a6 | -1.13030181 | 2.44E-08 |
| Ikbke | -2.513959093 | 2.44E-08 |
| Lama4 | 8.262224858 | 2.54E-08 |
| Cmtm6 | -1.665918766 | 2.57E-08 |
| mt-Nd4l | 1.983440778 | 2.77E-08 |
| Frmd8 | -1.667049705 | 2.93E-08 |
| Sgk3 | -1.802291933 | 3.06E-08 |
| Myo1f | -1.516976535 | 3.12E-08 |
| Sepw1 | 0.844433492 | 3.22E-08 |
| Sash3 | -1.832439697 | 3.22E-08 |
| Med31 | 1.569697819 | 3.23E-08 |
| Gsr | -1.823512913 | 3.44E-08 |
| Actr3b | 2.208919363 | 3.49E-08 |
| Romo1 | 1.095180624 | 3.50E-08 |
| Jtb | 1.28824517 | 3.63E-08 |
| Irf5 | -1.875079744 | 3.66E-08 |
| Erp44 | -1.115244239 | 3.72E-08 |
| Gap43 | 1.315533958 | 3.84E-08 |
| Lin9 | -1.669641967 | 3.85E-08 |
| Gpx8 | 2.729921345 | 3.95E-08 |
| Pik3r4 | -1.630494176 | 4.05E-08 |
| Megf10 | 2.853453091 | 4.15E-08 |
| Cenpf | 1.672196269 | 4.15E-08 |
| Bpgm | 1.349079479 | 4.17E-08 |
| Tm9sf2 | -1.04109696 | 4.20E-08 |
| Lamp2 | -1.357605146 | 4.26E-08 |
| Arhgap17 | -1.382140762 | 4.26E-08 |
| Rhbdd1 | -2.907500074 | 4.35E-08 |
| Mcm10 | -1.958575995 | 4.35E-08 |
| Gm1673 | 1.090997268 | 4.41E-08 |
| Igfbp5 | 2.615204185 | 4.42E-08 |
| Rab13 | 2.091110857 | 4.42E-08 |
| Inpp5d | -1.563997494 | 4.42E-08 |
| Pirb | -2.555097687 | 4.42E-08 |
| Acvr1c | -5.219352491 | 4.42E-08 |
| Hiatl1 | -1.572609242 | 4.44E-08 |
| Glo1 | 1.138237344 | 4.60E-08 |
| Dok3 | -2.267448623 | 4.62E-08 |
| Klf3 | -1.393957745 | 4.77E-08 |
| Cyp4f18 | -3.125406866 | 4.91E-08 |
| Sptbn1 | 1.169120907 | 4.97E-08 |
| Rassf3 | -2.386137289 | 5.02E-08 |
| Pid1 | -1.487441146 | 5.07E-08 |
| Vdr | -7.860460601 | 5.11E-08 |
| S1pr1 | 2.469388837 | 5.18E-08 |
| Vac14 | -1.433640584 | 5.19E-08 |
| Ppp1r15b | -1.483233322 | 5.28E-08 |
| Sema4d | -1.809546084 | 5.31E-08 |
| Bst1 | -3.485418536 | 5.42E-08 |
| Chd3os | 1.347852168 | 5.60E-08 |
| Nme1 | 0.984432418 | 5.70E-08 |
| Msx1 | 7.908498871 | 6.15E-08 |
| Ttc7 | -2.33850854 | 6.27E-08 |

| | | |
|---|---|---|
| Slc29a4 | -0.540890713 | 0.03361719 |
| Cs | -0.447001157 | 0.033629509 |
| Gpatch11 | 0.719151436 | 0.033629838 |
| Psmb3 | 0.410514763 | 0.033635318 |
| Tom1l1 | 0.916101129 | 0.033656661 |
| Fcnb | 2.581212481 | 0.033708299 |
| Snw1 | 0.477744844 | 0.033713549 |
| Dnah5 | 5.460237846 | 0.033716478 |
| Nos1 | -0.746305781 | 0.033812416 |
| Matr3 | 0.407105315 | 0.033818208 |
| Il11ra1 | 0.881952466 | 0.033847515 |
| Try5 | 5.621720597 | 0.033856124 |
| Heatr1 | -0.806680407 | 0.033900981 |
| Snx20 | -0.961792596 | 0.033945304 |
| 2900011O08Rik | 0.582153659 | 0.033976628 |
| Hist1h2ae | 0.804224175 | 0.034006374 |
| Znhit1 | 0.628098903 | 0.034043213 |
| Rab3ip | 0.588338891 | 0.034067581 |
| Nfs1 | -0.694580287 | 0.034087028 |
| Shroom1 | 5.206159626 | 0.034120336 |
| Rnf152 | 0.986689275 | 0.03412106 |
| Mcoln3 | 5.630094997 | 0.034121252 |
| Dpyd | 1.464390727 | 0.034121252 |
| Slc13a2 | -4.258826946 | 0.034161632 |
| Slc2a5 | 1.639659279 | 0.03418929 |
| Pcdhgc3 | -0.826748841 | 0.03418929 |
| Chka | -0.584621182 | 0.034221342 |
| Slc30a7 | -0.742816627 | 0.034221342 |
| Pus1 | -0.973031619 | 0.034221342 |
| Dedd2 | -1.027692041 | 0.034221342 |
| Evpl | -5.95918493 | 0.034221342 |
| Ly6k | -5.95918493 | 0.034221342 |
| Rwdd2b | 1.230214562 | 0.034291178 |
| Camk4 | 0.86284252 | 0.034291361 |
| Gm14085 | -3.855330574 | 0.034297364 |
| Clip1 | -0.501669784 | 0.034343807 |
| Pxylp1 | 1.060260927 | 0.034422283 |
| Ccr9 | -4.55425939 | 0.034422283 |
| Sec16a | -0.465295542 | 0.034436057 |
| Ddr1 | 0.540162844 | 0.034545612 |
| Acvr1b | -0.851673625 | 0.034547966 |
| Acad12 | -2.798126807 | 0.034556173 |
| Rsph9 | 0.917363721 | 0.034582986 |
| Rgs18 | -2.148471014 | 0.034615808 |
| Ipo5 | -0.564793752 | 0.034618203 |
| Rab32 | -1.08159093 | 0.034618203 |
| Pou6f2 | -1.542930726 | 0.034648121 |
| Bbs2 | 1.30724302 | 0.03471552 |
| Cers4 | 0.738536406 | 0.03471552 |
| Slc38a2 | -0.45460877 | 0.03471552 |
| Lypla1 | -0.639538692 | 0.03471552 |
| Dhx34 | -1.091482068 | 0.03471552 |
| Vit | 2.048512358 | 0.034722213 |
| Nup37 | 1.051541314 | 0.034740646 |
| Inhba | -2.119844367 | 0.034748745 |
| Arih1 | -0.494852661 | 0.034910328 |
| Zscan18 | 1.13159606 | 0.034910494 |
| Tcof1 | -0.574597477 | 0.034910494 |
| Ubr2 | -0.613926119 | 0.034910494 |
| Tmprss11a | -5.604191308 | 0.034918445 |
| Agap3 | 0.694160277 | 0.034939461 |
| Kifc3 | -1.242446002 | 0.034939494 |
| Mnat1 | 0.681174493 | 0.034983906 |
| Hsbp1 | 0.506535347 | 0.035022422 |
| Zfp558 | -0.848354498 | 0.035104007 |
| Dnajb11 | -0.503094264 | 0.035210682 |
| Emp2 | 1.503246602 | 0.035258644 |
| Gmppb | -0.886730272 | 0.035258644 |
| Notch2 | 0.617353095 | 0.035295739 |
| Ctse | -1.683362095 | 0.035295739 |
| Lats2 | -0.952240845 | 0.035306196 |
| Klhl13 | 0.868951997 | 0.035330104 |
| Pcnxl3 | -0.673843779 | 0.035335112 |
| H3f3a | 0.362397822 | 0.035353678 |
| Uckl1 | -0.619190653 | 0.035353678 |
| Ppp2r2a | -0.455359074 | 0.035381181 |
| Jak1 | -0.627679515 | 0.035405403 |
| 9030624J02Rik | -0.452189494 | 0.035415439 |
| Parp9 | -1.292546948 | 0.035423898 |
| Dpm3 | 0.791430318 | 0.035530983 |
| Abhd4 | 0.676436337 | 0.035542476 |
| Efcab7 | 1.636327462 | 0.035654076 |
| Lpxn | -0.946480769 | 0.035679371 |
| Tmem98 | 0.720514212 | 0.035730348 |
| Grpel2 | -0.522689165 | 0.035730348 |
| Reep2 | 0.68500988 | 0.035764295 |
| Pnlip | 5.499331784 | 0.035779271 |
| Rab3a | 0.472675875 | 0.035808945 |
| Noct | -0.841667656 | 0.03583603 |
| Pdcd2l | 0.54284982 | 0.035930975 |
| Mia3 | -0.506574305 | 0.036026555 |
| Tshz1 | 0.996065031 | 0.036028381 |
| Slc25a2 | -2.672987496 | 0.036042 |
| Stxbp5 | -1.171048942 | 0.036057283 |
| Adnp2 | -0.858830857 | 0.03606469 |
| Scrn3 | 1.881690504 | 0.036084718 |
| Tmem59 | 0.396853017 | 0.036122887 |
| Ppp3r1 | -0.438976656 | 0.036122887 |
| Frmd5 | -0.666199122 | 0.036158146 |
| C1qtnf2 | 5.303678535 | 0.036186208 |
| Fbrs | -0.785449965 | 0.036288658 |
| Exoc1 | -0.481749704 | 0.036300369 |
| Fus | -0.508527239 | 0.036300369 |
| Adamts5 | 1.607264469 | 0.03637176 |
| Cd80 | -2.364175658 | 0.036373039 |
| Abcb7 | 0.834314207 | 0.036407591 |
| Tmem129 | -0.718320376 | 0.036407591 |
| B3gnt9 | 5.45558671 | 0.036434392 |
| Ptprb | 5.192799196 | 0.036434392 |
| Odf3b | -5.270016132 | 0.036434392 |
| Cacna1f | -1.526067994 | 0.036514207 |
| Mblac1 | 1.180671589 | 0.036580564 |
| Hnrnph3 | 0.487263359 | 0.036580564 |
| Trhde | 2.058780786 | 0.036601955 |
| Srsf1 | -0.402277972 | 0.036606142 |
| Ube3a | 0.543052483 | 0.036625879 |
| Naa11 | 5.58803611 | 0.036644014 |
| Ror1 | 4.689909226 | 0.036644014 |
| Zfp934 | 2.121345623 | 0.036644014 |

| | | |
|---|---|---|
| Trmt2a | -0.39790226 | 0.264043654 |
| Sertad1 | -0.536955645 | 0.264043654 |
| Grin3b | 4.075283292 | 0.264367625 |
| Rnf225 | 2.552822141 | 0.264432565 |
| Dnd1 | 4.086187654 | 0.264624623 |
| Ppp5c | -0.274018121 | 0.264743029 |
| Pcdha12 | -0.960331259 | 0.264836142 |
| Tssk4 | -2.996652138 | 0.265147471 |
| Tra2b | 0.271647674 | 0.265152227 |
| Nnt | -0.324330956 | 0.265177154 |
| Tmem39a | -0.384797451 | 0.265177154 |
| Exoc2 | -0.389375062 | 0.265552426 |
| 9930022D16Rik | -2.168675831 | 0.265701269 |
| Plscr5 | 4.353237835 | 0.266002511 |
| Pla2g15 | -0.500520196 | 0.266002511 |
| Ptger3 | -1.367460192 | 0.266053118 |
| Isg15 | -1.438888485 | 0.266245344 |
| Picalm | -0.357104957 | 0.266332111 |
| Zfp114 | 1.028586308 | 0.266522156 |
| Csrp2 | 0.342364758 | 0.266594981 |
| Samd9l | -2.033664831 | 0.266772092 |
| Stx6 | -0.263667203 | 0.266791692 |
| Evl | -0.417934294 | 0.26693108 |
| Ccdc59 | 0.321402023 | 0.266961547 |
| Uvssa | 0.372741509 | 0.267002327 |
| BC021891 | 4.117686615 | 0.267063443 |
| Pcgf6 | 0.425347257 | 0.267301638 |
| Smyd2 | 0.311249546 | 0.267301638 |
| Klhl17 | -0.45347832 | 0.267301638 |
| Fcho1 | 0.380707694 | 0.267374413 |
| Cav2 | 0.704805557 | 0.267493057 |
| Nos1ap | 0.653632421 | 0.267608847 |
| Gprasp1 | 0.307534557 | 0.267624221 |
| Arl6ip6 | 0.338471306 | 0.267658158 |
| Scaf4 | -0.331022595 | 0.268029677 |
| Mycbp2 | 0.269423431 | 0.268093005 |
| Nat6 | 0.718705016 | 0.26814786 |
| Dus4l | 0.703547924 | 0.268189941 |
| Nod2 | -1.065357831 | 0.268284781 |
| Ift57 | 0.35320985 | 0.268425422 |
| Rpl9-ps6 | 0.307500233 | 0.268449896 |
| Cenpl | -0.46115433 | 0.268532466 |
| Fndc1 | -2.526202747 | 0.268532466 |
| Klhdc8a | 1.528589374 | 0.268593481 |
| Camk1g | 0.55361596 | 0.268660877 |
| Mtrr | -0.6260202 | 0.268804259 |
| Stat1 | -0.99359805 | 0.268900966 |
| Mid2 | 0.94466714 | 0.26896057 |
| BC055324 | -0.64827316 | 0.269058269 |
| Prtg | 2.018160673 | 0.269382519 |
| Ccno | -1.046192793 | 0.269382519 |
| Camk2b | -0.474025444 | 0.269442993 |
| Hexb | -0.422687505 | 0.269480151 |
| Uqcc1 | 0.377655782 | 0.269647519 |
| Ncapg2 | -0.472553131 | 0.269780483 |
| Klhdc1 | 0.727969614 | 0.270130028 |
| Vps37a | -0.342284806 | 0.270130028 |
| Vangl1 | 0.74506219 | 0.270248569 |
| Prox2 | -1.169403957 | 0.270337884 |
| Cfap221 | 4.016681792 | 0.270405965 |
| Zfp770 | 0.500334047 | 0.270405965 |
| Kxd1 | 0.32905496 | 0.270405965 |
| C2 | -1.196736845 | 0.270405965 |
| Grhl2 | 4.441394353 | 0.270458476 |
| Zmym2 | -0.266828434 | 0.270705487 |
| Kcnh5 | -1.403555231 | 0.270911362 |
| Ly86 | 0.571273524 | 0.270970258 |
| Cacng8 | -0.729858618 | 0.27104398 |
| Stag3 | -0.835392981 | 0.271059083 |
| Rassf2 | -0.418374463 | 0.271271333 |
| Sorl1 | -0.52887156 | 0.271301134 |
| Polr3f | -0.357914429 | 0.271307979 |
| Mars2 | -0.661141045 | 0.27144916 |
| Mgst1 | 0.708332692 | 0.271513921 |
| Cacna1c | -0.490292766 | 0.271513921 |
| Tex14 | -2.363991809 | 0.271513921 |
| Mip | 3.425128101 | 0.271674375 |
| Rpl28 | 0.231899879 | 0.271834761 |
| Trpt1 | 0.729090822 | 0.27184552 |
| Dgat2l6 | -4.875950491 | 0.27186517 |
| Insl3 | -4.875950491 | 0.27186517 |
| Irf7 | -1.491601661 | 0.271943811 |
| Gramd2 | 1.579336035 | 0.272042516 |
| Clec2l | 0.837465168 | 0.272042516 |
| Ssr2 | 0.250892908 | 0.272042516 |
| Kctd20 | -0.258912407 | 0.272042516 |
| Man1b1 | -0.470470436 | 0.272042516 |
| Gm3252 | 1.13226281 | 0.272055747 |
| Akt1 | 0.352613482 | 0.272068061 |
| 1600012H06Rik | -0.407972392 | 0.272068061 |
| Crtc1 | -0.282293144 | 0.272071388 |
| Zfp931 | 0.644218649 | 0.272119557 |
| Ugdh | -0.365533745 | 0.272197604 |
| Lcmt2 | -0.465065921 | 0.272197604 |
| Timp1 | -1.20747888 | 0.272377005 |
| Zfp719 | -0.450497949 | 0.272408363 |
| Gm7879 | 1.221722069 | 0.272438545 |
| Tpbg | 0.691701469 | 0.272438545 |
| Mllt3 | 0.340443508 | 0.272467431 |
| Zfp821 | -0.29445404 | 0.272476148 |
| Gamt | 0.435399387 | 0.272589509 |
| Gm8108 | 3.98658075 | 0.272729168 |
| Nrbp2 | 0.751606717 | 0.272794718 |
| Ackr3 | 0.490477332 | 0.272832808 |
| Mrps27 | 0.479312086 | 0.272879795 |
| Pde11a | -2.009469946 | 0.272950342 |
| Mettl1 | 0.598195097 | 0.272971991 |
| Nudt9 | 0.33902421 | 0.273042632 |
| Neto1 | 0.732142538 | 0.27311395 |
| Socs4 | -0.287671053 | 0.27311395 |
| Zfp935 | -0.57597784 | 0.27311395 |
| Plb1 | -3.107751098 | 0.27311395 |
| Snap91 | -0.39058176 | 0.273158882 |
| Wdr62 | -0.968372178 | 0.273230111 |
| Nr2c2ap | 1.005832348 | 0.273301305 |
| Tpcn2 | -0.618299302 | 0.273363244 |
| Fam53c | -0.348912651 | 0.273489234 |
| Hdac11 | 0.578681256 | 0.273538211 |
| Mtf2 | 0.375638906 | 0.273538211 |

| | | |
|---|---|---|
| Cbx6 | 0.197979018 | 0.652846827 |
| Slk | 0.175413736 | 0.652846827 |
| Plxnb1 | 0.164492595 | 0.652846827 |
| Wasl | 0.140976549 | 0.652846827 |
| Creld2 | -0.158799905 | 0.652846827 |
| Zfp937 | -0.373285006 | 0.652846827 |
| Ebf2 | -0.466236829 | 0.652846827 |
| Slc5a5 | -1.382308047 | 0.652846827 |
| C8a | -1.389820932 | 0.652846827 |
| Cplx1 | 0.226743356 | 0.652900901 |
| Dolpp1 | -0.168683707 | 0.652900901 |
| Zfp93 | 0.522131924 | 0.652997595 |
| Agbl4 | 0.713249393 | 0.653296143 |
| Tkt | -0.109374054 | 0.653296143 |
| 1300017J02Rik | -0.822695307 | 0.653296143 |
| Fxr1 | 0.148864188 | 0.653352455 |
| Aacs | -0.269099358 | 0.653430424 |
| H2afy | 0.127290139 | 0.653555061 |
| Rhobtb3 | -0.151831086 | 0.653555061 |
| Etnk1 | 0.140897503 | 0.653565336 |
| Gbas | 0.116207514 | 0.654059268 |
| Mfsd2b | -0.520816523 | 0.654209838 |
| Kif26b | 0.362442846 | 0.654224073 |
| Greb1l | 0.666524223 | 0.654421411 |
| Stc2 | -0.593077669 | 0.654421471 |
| Haus2 | 0.188164285 | 0.654573539 |
| Tmem106b | 0.163872006 | 0.654573539 |
| Prob1 | -2.218408357 | 0.654600893 |
| Cubn | -1.050631371 | 0.654606622 |
| Spsb4 | 0.250444083 | 0.65461287 |
| Wdr77 | 0.184615335 | 0.654921565 |
| Zfp97 | -0.94109683 | 0.654921565 |
| Sec22c | 0.25498603 | 0.655052383 |
| Thpo | -1.29102252 | 0.655052383 |
| Oxnad1 | -0.297267573 | 0.655104909 |
| 1700025G04Ri | 0.249704563 | 0.655179848 |
| Copz1 | 0.131482573 | 0.655179848 |
| Coa7 | -0.155636299 | 0.655179848 |
| Dalrd3 | -0.263120741 | 0.655179848 |
| Bdnf | 0.918348275 | 0.655204232 |
| Pde12 | 0.219129255 | 0.655210044 |
| Diras1 | -0.221848835 | 0.655467321 |
| Galnt13 | 0.43882001 | 0.655565242 |
| Cetn4 | -0.620271278 | 0.655571131 |
| Amigo3 | -0.708362754 | 0.655571131 |
| Fam24a | -1.420000004 | 0.655571131 |
| Fam110b | 0.17118256 | 0.656066598 |
| Gstcd | -0.239729968 | 0.656202062 |
| B9d2 | 0.158696771 | 0.6568785 |
| Osbp | -0.178811521 | 0.6568785 |
| Pik3c2b | -0.162633842 | 0.657142292 |
| Dgat1 | 0.166721086 | 0.657164554 |
| Hgf | 0.663877807 | 0.657207688 |
| Prim2 | 0.198389469 | 0.657207688 |
| Zik1 | -0.155728202 | 0.657207688 |
| Brinp1 | -0.264050972 | 0.657207688 |
| Gabrb2 | -0.46554242 | 0.657207688 |
| Rasd2 | 0.888845765 | 0.65750523 |
| Htr5b | -1.490643284 | 0.65750523 |
| Tbpl1 | -0.126955217 | 0.657538302 |
| Gigyf1 | 0.1992241 | 0.657762188 |
| Pbrm1 | -0.103037292 | 0.657823592 |
| Aasdhppt | 0.18388515 | 0.657847042 |
| Slc39a8 | 0.555392179 | 0.657899256 |
| Vgll4 | 0.167604635 | 0.657899256 |
| Fam192a | 0.11501899 | 0.657907131 |
| Commd5 | -0.181770154 | 0.658102493 |
| Stard9 | 0.253375052 | 0.658138856 |
| St6galnac6 | -0.205128164 | 0.658225917 |
| RP23-466J17.3 | -1.2824513 | 0.658408295 |
| Pp2d1 | 1.000477068 | 0.658461949 |
| Fkbp14 | -0.314957301 | 0.658461949 |
| 6430550D23Ri | -0.610344302 | 0.658461949 |
| Nol4l | -0.173303383 | 0.659111972 |
| Ap4b1 | -0.915136634 | 0.659132869 |
| Ext2 | 0.12934101 | 0.659327202 |
| Rpl3 | -0.097678706 | 0.659327202 |
| Panx2 | 0.801703708 | 0.659456904 |
| Fam221a | 0.460767768 | 0.659456904 |
| Adamtsl1 | -0.388135015 | 0.659456904 |
| Pex2 | 0.124088365 | 0.659897321 |
| Lrrtm4 | -0.35366518 | 0.659897321 |
| Lrrc73 | 0.598401203 | 0.659965525 |
| Csl | -1.276660221 | 0.659965525 |
| Snx21 | 0.25111036 | 0.660096043 |
| Eif2b1 | 0.146191224 | 0.660096043 |
| Ubp1 | -0.152311861 | 0.660096043 |
| Lrrc24 | -0.268439307 | 0.660096043 |
| Nbeal1 | -0.166205917 | 0.660205276 |
| Tmem173 | 0.352205929 | 0.660225391 |
| Tpp1 | -0.191780369 | 0.660247309 |
| Oas1c | 0.653620375 | 0.660272745 |
| Gm28285 | 0.339558912 | 0.660371079 |
| Cabin1 | -0.173162799 | 0.660425015 |
| Zbtb25 | -0.26271299 | 0.660425015 |
| Gm16494 | 1.959521539 | 0.660577019 |
| Slc1a1 | 0.388329782 | 0.660696622 |
| Zbtb17 | 0.15409423 | 0.660868103 |
| Myadml2 | 1.285877748 | 0.660972914 |
| Ighmbp2 | 0.256192699 | 0.661129616 |
| Rnf217 | 0.384659619 | 0.661176255 |
| Inha | -0.316621741 | 0.661233554 |
| Cfh | 0.189357767 | 0.661406284 |
| Cyp2j6 | 0.513079519 | 0.661594851 |
| Slc22a23 | 0.265202014 | 0.661601924 |
| Rbbp8 | 0.187309479 | 0.661645883 |
| Zfp488 | 1.77748975 | 0.661662052 |
| Nmi | -0.315506977 | 0.661713015 |
| Mycn | 0.218687945 | 0.661722184 |
| Fam207a | 0.169627229 | 0.661749021 |
| Thumpd1 | -0.154445654 | 0.661749021 |
| Smek2 | -0.145998102 | 0.661941339 |
| Cep55 | -0.220276354 | 0.661941339 |
| Tubg1 | 0.139214947 | 0.662011318 |
| Cgn | -0.429896179 | 0.66205826 |
| Adra1b | 1.422045207 | 0.662160116 |
| Zfp784 | -1.695754378 | 0.66220558 |
| Sqle | -0.133730631 | 0.662327635 |
| Kcp | -0.470025481 | 0.662575777 |

| | | |
|---|---|---|
| Pitpna | -1.366292374 | 6.47E-08 |
| Ctsz | -1.662869788 | 6.73E-08 |
| Rpl36 | 1.094100083 | 6.99E-08 |
| Atp2a2 | -1.124601704 | 6.99E-08 |
| Gjb6 | 8.207795339 | 7.17E-08 |
| Edem1 | -1.771525612 | 7.37E-08 |
| Pom121 | -1.428632917 | 7.42E-08 |
| Sub1 | 0.921074046 | 7.66E-08 |
| Nbl1 | 3.358160213 | 7.87E-08 |
| Prkcb | -1.688030008 | 8.43E-08 |
| Myo1c | -1.791155791 | 8.43E-08 |
| Dtx2 | -1.884857853 | 8.44E-08 |
| Arap1 | -1.726440062 | 8.50E-08 |
| Slc2a1 | -2.041912801 | 8.50E-08 |
| Coro7 | -1.365101228 | 8.55E-08 |
| Cks2 | 1.62548888 | 8.96E-08 |
| Cox7a2 | 0.985528494 | 8.96E-08 |
| Thbs1 | -4.078814113 | 9.37E-08 |
| Sdhaf4 | 1.244941573 | 9.64E-08 |
| Timm8b | 0.970362549 | 9.64E-08 |
| Fgf13 | 1.383937181 | 9.83E-08 |
| Acot1 | 1.5395073 | 9.96E-08 |
| Ncstn | -1.099485206 | 1.05E-07 |
| AW551984 | 2.924555628 | 1.06E-07 |
| Hmgb3 | 0.851751164 | 1.08E-07 |
| Sgms2 | -4.284361838 | 1.10E-07 |
| Ube2c | 1.279172001 | 1.24E-07 |
| Atp5e | 1.120774914 | 1.32E-07 |
| Pfkl | -1.092204073 | 1.38E-07 |
| Nrarp | 2.299923557 | 1.39E-07 |
| Lrp10 | -1.420087857 | 1.40E-07 |
| Dkk3 | 3.2014166 | 1.42E-07 |
| Rhbdf2 | -2.343287699 | 1.46E-07 |
| Myo9b | -1.145165736 | 1.48E-07 |
| Hcls1 | -1.999944813 | 1.49E-07 |
| Cald1 | 1.648928926 | 1.49E-07 |
| Snrpd2 | 1.018542137 | 1.50E-07 |
| Wdr81 | -1.462292815 | 1.51E-07 |
| Zcchc12 | 1.746900363 | 1.61E-07 |
| Cpe | 1.055093306 | 1.66E-07 |
| Ndufb11 | 0.977067583 | 1.73E-07 |
| Lyn | -1.945471811 | 1.74E-07 |
| Pttg1 | 1.256315732 | 1.78E-07 |
| Ndufb4 | 1.253228419 | 1.85E-07 |
| Olfm4 | -5.996600165 | 1.99E-07 |
| Plin2 | -1.783722262 | 2.01E-07 |
| Mpo | 4.534933062 | 2.03E-07 |
| Tram1l1 | 1.444651865 | 2.07E-07 |
| C3ar1 | -1.950180124 | 2.14E-07 |
| Ms4a6d | -2.059332062 | 2.17E-07 |
| Sulf2 | 2.889138125 | 2.20E-07 |
| Ppm1b | -1.089521047 | 2.24E-07 |
| Plagl1 | 1.24930245 | 2.25E-07 |
| Ncf1 | -1.784887072 | 2.34E-07 |
| Gcsh | 1.172580664 | 2.35E-07 |
| B230118H07Rik | 1.566024375 | 2.38E-07 |
| Prdx5 | -1.113459241 | 2.40E-07 |
| Gltscr1 | -1.578527857 | 2.40E-07 |
| Plk2 | -0.994904095 | 2.43E-07 |
| Igfbp7 | 7.969606915 | 2.45E-07 |
| Parp8 | 2.507521683 | 2.47E-07 |
| Sgol1 | 2.106365868 | 2.50E-07 |
| Mlc1 | 2.328739686 | 2.52E-07 |
| Sphk2 | -1.474636248 | 2.53E-07 |
| Mvp | -2.023247352 | 2.53E-07 |
| Hist1h2ap | 1.722797278 | 2.60E-07 |
| Fgfbp3 | 2.313951144 | 2.64E-07 |
| Akap12 | 2.324885734 | 2.68E-07 |
| Cox6a1 | 1.12972677 | 2.68E-07 |
| Fxr2 | -1.306670823 | 2.71E-07 |
| Arl6 | 1.241210065 | 2.73E-07 |
| Dpy30 | 1.145052897 | 2.73E-07 |
| Marcks | 1.070271009 | 2.73E-07 |
| Fkbp15 | -1.133065302 | 2.73E-07 |
| Neu1 | -1.365818752 | 2.73E-07 |
| Bok | 1.236558443 | 2.74E-07 |
| Adrbk2 | -2.03725235 | 2.78E-07 |
| Ppp1r1a | 1.520922034 | 2.79E-07 |
| Nfam1 | -2.109899431 | 2.90E-07 |
| Tspan7 | 1.492503805 | 2.92E-07 |
| Pik3cd | -1.627904304 | 2.94E-07 |
| Cisd1 | 0.96005954 | 2.97E-07 |
| Rims3 | 2.187748768 | 2.98E-07 |
| Tmem2 | -1.069614816 | 3.07E-07 |
| Sall2 | 1.964353161 | 3.08E-07 |
| Pdcd6ip | -1.237387318 | 3.12E-07 |
| Cetn2 | 1.071227476 | 3.14E-07 |
| Mal2 | 6.751243808 | 3.14E-07 |
| Cd300lb | -7.529654792 | 3.16E-07 |
| Tmem123 | -1.286059242 | 3.22E-07 |
| Gm8186 | 1.484480484 | 3.26E-07 |
| Slc35d2 | -2.860101659 | 3.27E-07 |
| Bex1 | 1.20345344 | 3.39E-07 |
| Nup98 | -1.285528238 | 3.39E-07 |
| Snap23 | -1.36446719 | 3.39E-07 |
| Fam83f | -8.093249641 | 3.44E-07 |
| Sh2b2 | -1.528021093 | 3.53E-07 |
| Atxn2l | -1.3400849 | 3.53E-07 |
| Gpr107 | -1.134101552 | 3.63E-07 |
| Pilra | -2.280754766 | 3.72E-07 |
| Slco2b1 | 4.450627263 | 3.76E-07 |
| Dgcr6 | 0.986633525 | 3.83E-07 |
| Iqgap1 | -1.716288695 | 3.84E-07 |
| Slc2a6 | -2.224486214 | 3.84E-07 |
| Nr2f1 | 1.763093907 | 3.90E-07 |
| Col14a1 | -3.337625171 | 3.90E-07 |
| Tapbp | -1.712486795 | 3.91E-07 |
| Igf2bp3 | -1.13522515 | 3.92E-07 |
| B9d1 | 1.705897812 | 3.92E-07 |
| Shc1 | -1.46108176 | 4.12E-07 |
| Fen1 | -1.515735096 | 4.12E-07 |
| Pla2g7 | -1.699482727 | 4.17E-07 |
| Rel | -1.434287309 | 4.17E-07 |
| Sfrp2 | 2.029642289 | 4.24E-07 |
| Cttnbp2nl | -1.217523405 | 4.24E-07 |
| Lrrc3b | 7.783659845 | 4.24E-07 |
| Rps26 | 0.873395193 | 4.26E-07 |
| Psmb4 | 1.113862497 | 4.31E-07 |
| G0s2 | -3.858975574 | 4.31E-07 |

| | | |
|---|---|---|
| Chd2 | -0.442601743 | 0.036647827 |
| Depdc1a | 1.216235331 | 0.036663707 |
| Enox2 | -1.040033067 | 0.036674366 |
| Htr5a | 3.535429606 | 0.036703919 |
| Snx29 | -0.864519849 | 0.036703919 |
| Dnajc17 | 0.981520603 | 0.036789364 |
| Cep83 | -0.855559473 | 0.036834472 |
| Mrps7 | 0.520031209 | 0.03687104 |
| Rcor1 | 0.99056088 | 0.036873615 |
| Rpl35a | 0.526668922 | 0.036879392 |
| Cela1 | -1.376348976 | 0.036931576 |
| Psma2 | 0.458623078 | 0.036932235 |
| Arfgap1 | -0.559897825 | 0.037105057 |
| Nsun4 | -0.841363618 | 0.037137098 |
| Orc1 | -1.823844885 | 0.037157748 |
| Ifi47 | -3.199875115 | 0.037157748 |
| Dlg2 | -0.792567733 | 0.037198271 |
| Slc18a1 | -1.463413584 | 0.037252738 |
| Tmc8 | -1.472156459 | 0.037252738 |
| Abca5 | 1.011406354 | 0.03727482 |
| Rpl9 | 0.358393276 | 0.03727482 |
| Tox | 0.954337752 | 0.037310367 |
| Abat | 0.771040957 | 0.037317983 |
| Ost4 | 0.827728697 | 0.037321686 |
| Acly | -0.477125556 | 0.037338101 |
| Nudt17 | 2.027555567 | 0.037401756 |
| Spp1 | -2.196404277 | 0.037407847 |
| Pkp3 | -4.87760873 | 0.037407847 |
| Fam227b | -5.291447043 | 0.037407847 |
| Bcl10 | -0.5045831 | 0.037438103 |
| Arel1 | -0.639935717 | 0.037542447 |
| Nop58 | 0.420638268 | 0.03755634 |
| Rpl6 | 0.391979078 | 0.03755634 |
| Pdlim1 | 1.168739029 | 0.037582911 |
| Mre11a | -0.713196541 | 0.037615383 |
| Milr1 | -1.003242027 | 0.037615383 |
| 9530077C05Rik | 1.123476747 | 0.037646466 |
| Gpn3 | 0.939530186 | 0.037678758 |
| Lhcgr | 5.440995894 | 0.037706042 |
| Hopx | 1.254311893 | 0.037706042 |
| Taf12 | 0.580790515 | 0.037715027 |
| Ntsr2 | 4.089777132 | 0.037734513 |
| Il1rap | -1.214589488 | 0.037734513 |
| Zfp704 | 0.55603706 | 0.037737114 |
| Atp6ap2 | -0.460830752 | 0.037737114 |
| Echs1 | 0.511774168 | 0.037801865 |
| Rpp25l | 0.774159218 | 0.037874834 |
| Chmp7 | 0.456937111 | 0.037909373 |
| Gpr149 | 4.728691518 | 0.037910134 |
| Chl1 | 0.621579239 | 0.03794735 |
| Ly6c2 | -2.241719149 | 0.03794735 |
| Zim1 | 2.416632174 | 0.03796792 |
| Gm9843 | 0.497062977 | 0.03796792 |
| Abi1 | -0.549125886 | 0.038051404 |
| Orc2 | 0.66089761 | 0.038054304 |
| 1700001C02Rik | 5.194746529 | 0.038078234 |
| Ascl1 | 1.136059215 | 0.038109312 |
| Rqcd1 | -0.465032757 | 0.038153134 |
| Pfdn2 | 0.53196497 | 0.038263599 |
| Tmcc1 | 0.451455438 | 0.038263599 |
| Lgr5 | -3.213521107 | 0.038263599 |
| Atg10 | -0.753012264 | 0.038266809 |
| Cebpzos | 0.881520614 | 0.038382658 |
| Tm2d1 | 0.492952453 | 0.038437476 |
| Arpp21 | -1.740403224 | 0.038473558 |
| Gm5868 | 5.302156743 | 0.038491392 |
| 4921506M07Rik | 4.814395133 | 0.038508695 |
| Zfp653 | -0.952230873 | 0.038529522 |
| Arhgap9 | -0.886590817 | 0.038568095 |
| Fam32a | 0.418577956 | 0.03857331 |
| 3110035E14Rik | -0.865694698 | 0.038591628 |
| Mlxip | -0.531835901 | 0.038653907 |
| 4930590J08Rik | -2.000050982 | 0.038674387 |
| Sfn | -5.867972185 | 0.038674387 |
| Ankrd55 | 2.505946597 | 0.038738689 |
| Ankrd40 | -0.857390718 | 0.038745832 |
| Zfp54 | 3.265541427 | 0.038940864 |
| Poli | 1.044526741 | 0.038984335 |
| Gbp7 | -1.705946489 | 0.039026286 |
| Dnajc15 | 0.638859416 | 0.039102723 |
| Skil | -0.585125871 | 0.039102723 |
| Dcc | -0.925403024 | 0.039102723 |
| Hdhd2 | 0.670645694 | 0.039108975 |
| B230217C12Rik | 0.90023207 | 0.039139899 |
| Gnpat | 0.507380634 | 0.039139899 |
| Glb1l | -1.935110662 | 0.039150681 |
| Rab9b | 0.965053227 | 0.039224965 |
| Emc8 | 0.516348967 | 0.039224965 |
| Zfp358 | 0.468851254 | 0.039268982 |
| Radil | 1.853291011 | 0.039370556 |
| Fscn1 | -0.414850176 | 0.0393739 |
| Slc20a1 | -0.596795817 | 0.039391277 |
| Dnm1 | 0.98613981 | 0.039394212 |
| Usb1 | -0.630610174 | 0.039501234 |
| Stk32c | 1.303999698 | 0.039589886 |
| Prdm5 | 1.600146741 | 0.03971912 |
| Cyp7b1 | 2.366340453 | 0.039727762 |
| Dnajc3 | -0.556530725 | 0.039771145 |
| Stam2 | -0.450602923 | 0.03980018 |
| Ccdc151 | 2.39537141 | 0.039861745 |
| Zbtb48 | -1.377011353 | 0.039861745 |
| Hist1h2bc | 0.768488907 | 0.039866736 |
| 2700062C07Rik | 0.809505552 | 0.039870572 |
| Iqub | 5.326919913 | 0.039877187 |
| Pcdhb8 | 3.319813361 | 0.039880807 |
| Col22a1 | 1.990985033 | 0.039880807 |
| Nmt1 | -0.569815691 | 0.039880807 |
| Pex11a | 1.124976089 | 0.039887023 |
| 1700001O22Rik | 1.681654933 | 0.039889044 |
| Gcnt4 | 5.529607549 | 0.039932186 |
| Nme6 | 0.744543634 | 0.039932186 |
| 3830408C21Rik | 5.246637387 | 0.039940161 |
| Kcnk7 | -2.578673734 | 0.039996306 |
| Frem2 | 1.779123264 | 0.040049533 |
| Sumo3 | 0.503929257 | 0.040049533 |
| Sybu | -0.906991988 | 0.040088553 |
| Eif4e | 0.484526389 | 0.0401015 |
| Wipi2 | -0.473200405 | 0.0401015 |
| Ndfip1 | 0.4426793 | 0.040105159 |

| | | |
|---|---|---|
| Olfr1386 | 4.337624875 | 0.273595919 |
| Ppp1r2 | 0.268555748 | 0.273607689 |
| Ube4b | -0.337289174 | 0.273711438 |
| A930002H24Rik | 3.939710065 | 0.27380296 |
| Ugcg | -0.427472838 | 0.273861222 |
| 9430038I01Rik | -0.45354296 | 0.273861222 |
| Spin4 | 1.300275252 | 0.27390691 |
| 1700029J07Rik | 0.96746125 | 0.27390691 |
| Sf3b5 | 0.276285474 | 0.274073948 |
| Gm9803 | 0.737077347 | 0.274108705 |
| Atg2a | -0.450904304 | 0.274108705 |
| Shank3 | -0.686484441 | 0.274114057 |
| Agpat5 | -0.356530793 | 0.274239936 |
| Taldo1 | -0.228347121 | 0.274361908 |
| Tbc1d8 | -0.423959081 | 0.274598226 |
| Supt3 | 0.569886939 | 0.274920612 |
| Map2 | 0.301942827 | 0.275116558 |
| Zswim7 | -0.531484588 | 0.275116558 |
| Armcx4 | 0.379483426 | 0.275120266 |
| Agap1 | -0.302438121 | 0.275120266 |
| Zfp960 | 2.162853297 | 0.275243205 |
| Fam117a | 1.586594817 | 0.275243205 |
| Fuca1 | -0.311859347 | 0.275332142 |
| Dot1l | -0.364231735 | 0.275332142 |
| Ltbr | -0.482815855 | 0.275338996 |
| Kcng1 | -0.940176269 | 0.275396339 |
| Mpdz | 0.436609424 | 0.275532765 |
| Gm21967 | 1.111764547 | 0.275566387 |
| Rcan1 | -0.345914795 | 0.27576425 |
| Slc25a16 | 0.485282002 | 0.275782345 |
| 3-Sep | 0.313237807 | 0.275782345 |
| Tbx2 | 4.13322903 | 0.275937459 |
| Polr2m | 0.227566711 | 0.276006706 |
| Igsf9 | -0.462141913 | 0.276031032 |
| Scaf11 | 0.293025797 | 0.276252806 |
| Btbd11 | -0.814267177 | 0.276252806 |
| Borcs5 | 0.32891175 | 0.276545077 |
| Pcmtd1 | 0.348560427 | 0.276571489 |
| Csmd2 | 0.698356984 | 0.276665373 |
| 0610007P14Rik | 0.316294736 | 0.276665373 |
| Hcfc2 | -0.375437548 | 0.276665373 |
| Gm14403 | -1.28016111 | 0.276665373 |
| Tdp1 | 0.339905941 | 0.276769992 |
| Kdm4c | -0.378565143 | 0.276769992 |
| Map3k6 | -2.027899866 | 0.276796864 |
| Pex11g | 0.640896906 | 0.276920204 |
| Spice1 | 0.471996818 | 0.277037174 |
| Taz | 0.409557032 | 0.277037174 |
| Dtx1 | 0.40332266 | 0.277037174 |
| Pcdhga6 | -0.888763694 | 0.277037174 |
| Tm7sf3 | 0.387982158 | 0.277283262 |
| Gak | -0.262926219 | 0.277283262 |
| Smyd3 | -0.414270007 | 0.277283262 |
| Pde4a | -0.48734513 | 0.277283262 |
| Mpg | -0.542190983 | 0.277283262 |
| Kctd21 | 1.751034316 | 0.277411118 |
| Srp54b | -0.322885089 | 0.277559946 |
| Hexim1 | -0.405320885 | 0.277559946 |
| Fig4 | -0.433230445 | 0.277707984 |
| Trpv2 | -0.635662141 | 0.277724433 |
| Cdc7 | 0.330987379 | 0.277963915 |
| Pepd | -0.291803955 | 0.277963915 |
| Brox | -0.303699629 | 0.277963915 |
| Etnppl | 3.742619067 | 0.277981394 |
| Hbq1b | 3.336029102 | 0.277981394 |
| Trmt1l | -0.429731784 | 0.277981394 |
| Rabl3 | -0.499727316 | 0.277981394 |
| Haus3 | 0.455925962 | 0.278016362 |
| Kctd18 | 0.492917007 | 0.278129229 |
| Trim59 | 0.45159523 | 0.278129229 |
| Aff2 | -0.544936442 | 0.278129229 |
| Kpna3 | -0.246973122 | 0.278139285 |
| Samd14 | 0.303625521 | 0.278148373 |
| Hist2h2be | 0.519239865 | 0.278159253 |
| Kdm5a | -0.261324258 | 0.278190294 |
| Spdya | -0.839787561 | 0.278225155 |
| Nov | 1.824480642 | 0.278607579 |
| Kat6a | -0.312290609 | 0.278774092 |
| Slc26a1 | 1.781910638 | 0.278953596 |
| Nrg1 | -0.37618828 | 0.278953596 |
| Tmem132c | -0.997487881 | 0.278953596 |
| Ntmt1 | 0.271077681 | 0.279128371 |
| Smim8 | 0.345961793 | 0.27931483 |
| Ulk3 | -0.992802909 | 0.27931483 |
| Rad51ap1 | -0.558427844 | 0.279357339 |
| Stag1 | 0.28002119 | 0.279372259 |
| Sac3d1 | 0.367674115 | 0.279390679 |
| Smarcd3 | -0.423044063 | 0.279397463 |
| Yars2 | 0.419357531 | 0.279410692 |
| Shroom4 | -0.70321164 | 0.27942708 |
| Il1r1 | -1.680650392 | 0.27942708 |
| Slc22a29 | -1.609133732 | 0.279484373 |
| Tfrc | -0.297931401 | 0.279495919 |
| Tmem222 | 0.289334951 | 0.279627714 |
| Vamp5 | 0.923173308 | 0.279719477 |
| Siglece | 0.672714214 | 0.280163017 |
| Nebl | 0.867541618 | 0.280206985 |
| Mbd3 | 0.295920837 | 0.280252714 |
| Ankrd16 | -0.420794094 | 0.280252714 |
| Zfp874b | -0.998814763 | 0.280252714 |
| Trub2 | 0.442901598 | 0.280266044 |
| Eif4g3 | -0.256956999 | 0.280266044 |
| Odf3l1 | 4.058281137 | 0.280275682 |
| Pdgfrb | 0.82585175 | 0.280303646 |
| Mef2c | -0.348788905 | 0.280325527 |
| Tiparp | -0.329524677 | 0.280326389 |
| Smpdl3b | 1.61992728 | 0.280483107 |
| Mydgf | 0.302551741 | 0.280527083 |
| Lmnb2 | -0.438722533 | 0.280527083 |
| Fam199x | -0.534962532 | 0.280527083 |
| Mettl10 | 0.344254243 | 0.280611207 |
| Cltc | -0.246709882 | 0.280715772 |
| Zfp583 | -0.570443365 | 0.280736791 |
| Cacna1h | -0.578397329 | 0.280803913 |
| Qsox2 | -0.54365933 | 0.280835201 |
| Zfp956 | 0.475461878 | 0.28096586 |
| A130010J15Rik | 0.66246741 | 0.281028452 |
| Anapc15-ps | 1.703415307 | 0.281032822 |
| Tbx3 | 4.33011879 | 0.281070499 |

| | | |
|---|---|---|
| Gm2564 | 1.651002155 | 0.662576717 |
| Brcc3 | 0.126281742 | 0.662576717 |
| Gpd1l | -0.198110381 | 0.662576717 |
| Txlnb | -0.985644201 | 0.662576717 |
| Rph3al | 0.869150687 | 0.662642522 |
| Psg16 | 2.402408465 | 0.662678408 |
| Podxl2 | 0.124866419 | 0.663012353 |
| Kcna4 | -0.802335065 | 0.663042651 |
| Gm14685 | 1.733405314 | 0.663055203 |
| Gm10490 | -1.7157925 | 0.663326774 |
| Usp22 | -0.109464462 | 0.663354381 |
| Bloc1s1 | -0.712568823 | 0.663477191 |
| Stxbp1 | 0.120574997 | 0.663520431 |
| Pkib | 0.292474112 | 0.6635834 |
| Kdm1a | -0.125483893 | 0.6635834 |
| Apobec1 | -0.20310901 | 0.6635834 |
| Slc32a1 | -0.434742981 | 0.6635834 |
| Dpp10 | 0.445420721 | 0.663688338 |
| Phf7 | -0.261615264 | 0.66383809 |
| Il1a | -0.329916342 | 0.664182159 |
| Rsl1d1 | 0.114984983 | 0.664266407 |
| Arx | 0.211890134 | 0.664268035 |
| Ints9 | 0.16636941 | 0.664609113 |
| Gpr137c | 0.354967943 | 0.664691325 |
| Etfb | 0.329513038 | 0.664691325 |
| Sass6 | 0.167137259 | 0.66471646 |
| Zfp148 | -0.130492688 | 0.66471646 |
| Cry2 | 0.191359846 | 0.664748924 |
| Rabep2 | -0.225540225 | 0.664748924 |
| Paip1 | 0.119605353 | 0.665252239 |
| 1700056E22Ril | -0.770208216 | 0.665252239 |
| Hmgcr | 0.121719048 | 0.665273312 |
| Dapk2 | -0.965080957 | 0.665297724 |
| Btbd19 | 0.344473381 | 0.665422204 |
| Zfp532 | -0.155698735 | 0.665422204 |
| Them6 | -1.003601214 | 0.665422204 |
| Zfp870 | 0.495720273 | 0.665470044 |
| Map10 | -0.33569829 | 0.665470044 |
| Mtdh | -0.109203789 | 0.6654796 |
| Ppm1k | 0.197003529 | 0.665793884 |
| Sat1 | -0.138578048 | 0.666042431 |
| Dmrt2 | -1.5197526 | 0.666042431 |
| Upp2 | -0.416915428 | 0.666096599 |
| Meis1 | 0.232524486 | 0.666467634 |
| Gpr68 | -1.964697228 | 0.666467634 |
| Epm2a | 0.360856849 | 0.666549921 |
| Kcnb2 | 0.24470614 | 0.666564534 |
| Snx33 | -0.50290725 | 0.666655577 |
| Dpy19l2 | -2.248069321 | 0.666657426 |
| Trip13 | 0.235310293 | 0.666969866 |
| Focad | -0.235185728 | 0.666969866 |
| Crem | -0.272185227 | 0.666969866 |
| Dhx32 | -0.180969967 | 0.667243526 |
| Kcnab1 | 0.510597193 | 0.667262395 |
| Gtf2e1 | 0.160912276 | 0.667262395 |
| Tor1b | -0.166924368 | 0.667262395 |
| Tmtc4 | -0.199350265 | 0.667262395 |
| Zfp36l2 | -0.323015989 | 0.667262395 |
| H2-Oa | -0.830450444 | 0.667262395 |
| Sytl1 | 1.856776319 | 0.667753139 |
| Aff4 | -0.138382125 | 0.667832158 |
| Zfp458 | 0.407592614 | 0.667895068 |
| Hfe2 | 2.367816565 | 0.667976995 |
| Cdh17 | 1.070573255 | 0.667976995 |
| Ccdc64 | 0.272098867 | 0.667976995 |
| Sirt5 | 0.2387321 | 0.667976995 |
| Mink1 | -0.264029123 | 0.667976995 |
| Mettl2 | 0.149598969 | 0.668007819 |
| Pik3r2 | 0.169954768 | 0.668433878 |
| Cizl | -0.138451626 | 0.668887675 |
| Atp6v1g3 | 2.155543087 | 0.668899705 |
| Gm5616 | 0.624137904 | 0.669063952 |
| Vdac1 | 0.095752201 | 0.669063952 |
| Gm10320 | -0.213355293 | 0.669063952 |
| Slc7a14 | -0.402135201 | 0.669063952 |
| Pcdhb16 | -0.431753218 | 0.669063952 |
| Ttc14 | 0.161136005 | 0.669398766 |
| Phf11c | 1.222636302 | 0.669403016 |
| Ccdc130 | 0.258432519 | 0.669403016 |
| Ubac1 | 0.174974855 | 0.669403016 |
| Epha4 | 0.167572945 | 0.669403016 |
| Tdrkh | 0.149615347 | 0.669403016 |
| Cisd2 | -0.114258066 | 0.669403016 |
| Spire1 | -0.142390509 | 0.669472318 |
| Fam73a | -0.241195197 | 0.669753499 |
| Ufsp1 | 0.402065516 | 0.669818001 |
| Hid1 | 0.16581909 | 0.669818001 |
| Yars | 0.113099073 | 0.669818001 |
| Gnb3 | -0.539038348 | 0.669818001 |
| Meig1 | 0.565802326 | 0.669911802 |
| Fnip1 | 0.189532127 | 0.669911802 |
| Cdk15 | 2.159602584 | 0.670177995 |
| Ccdc6 | 0.117282491 | 0.670335666 |
| Ak8 | 0.488996619 | 0.670518773 |
| Paf1 | 0.155403363 | 0.670518773 |
| Atp6v0e | -0.123017713 | 0.670695533 |
| Ushbp1 | 1.502858576 | 0.670711235 |
| Bmper | 0.420009331 | 0.670711235 |
| Rbm18 | 0.109813375 | 0.670711235 |
| Cryl1 | -0.193598216 | 0.670711235 |
| Lonrf1 | -0.197788612 | 0.670711235 |
| Ankef1 | -1.4344771 | 0.670711235 |
| D5Ertd579e | -0.130100512 | 0.670778447 |
| Bin1 | 0.115442811 | 0.672126564 |
| Skiv2l2 | 0.151311878 | 0.672162633 |
| Gpr150 | 0.738758786 | 0.672264962 |
| Fam175b | 0.17732282 | 0.67241225 |
| Myo19 | -0.321641679 | 0.672565877 |
| Cd151 | 0.163389957 | 0.672643049 |
| Dcps | 0.170553033 | 0.67271239 |
| R3hdm4 | 0.120853691 | 0.672909394 |
| Gm3558 | 0.675094226 | 0.673040479 |
| Cxcr4 | -0.527847562 | 0.673200647 |
| Senp2 | 0.165237883 | 0.673290015 |
| Me1 | -0.253181481 | 0.673421777 |
| Gas2 | -0.277215639 | 0.673478014 |
| Col20a1 | 0.745928725 | 0.673484528 |
| Limch1 | -0.187216734 | 0.673484528 |
| Arhgef39 | 0.241287804 | 0.673550747 |

| | | |
|---|---|---|
| Snx1 | -0.989550415 | 4.42E-07 |
| Lrrk1 | -1.827620183 | 4.48E-07 |
| Cap1 | -1.173864629 | 4.51E-07 |
| Nrros | -1.791339808 | 4.56E-07 |
| Rasa4 | -2.023207381 | 4.56E-07 |
| Mfge8 | 2.254357255 | 4.59E-07 |
| Rgcc | 1.646443051 | 4.63E-07 |
| Tceb2 | 0.908955671 | 4.68E-07 |
| Alas1 | -1.261730196 | 4.83E-07 |
| Cx3cr1 | -1.918103372 | 4.83E-07 |
| Igsf1 | -8.308120008 | 4.85E-07 |
| Phf11b | -2.786932231 | 4.92E-07 |
| Samp | 1.000100605 | 4.96E-07 |
| Csdc2 | 1.440600483 | 5.12E-07 |
| Wbp5 | 0.923477807 | 5.15E-07 |
| Lrrfip1 | -1.219802938 | 5.16E-07 |
| Elf4 | -1.890626512 | 5.16E-07 |
| Trim21 | -2.950064782 | 5.16E-07 |
| Pigp | 0.985748513 | 5.17E-07 |
| Rps29 | 1.132508634 | 5.19E-07 |
| Tro | 1.370925043 | 5.24E-07 |
| Coro1a | -1.376264838 | 5.26E-07 |
| Tor3a | -1.704272028 | 5.36E-07 |
| Lcp1 | -1.72013357 | 5.38E-07 |
| Ndrg1 | -1.670069868 | 5.41E-07 |
| Sod2 | -1.641241444 | 5.50E-07 |
| P4hb | -1.107830446 | 5.60E-07 |
| Rps6ka1 | -1.678687174 | 5.66E-07 |
| Cav1 | 4.39357914 | 5.66E-07 |
| Clic6 | 4.349317321 | 5.73E-07 |
| Ldlr | -1.924088266 | 5.83E-07 |
| Smarcd2 | -1.555585274 | 5.86E-07 |
| Inf2 | -2.263709824 | 5.89E-07 |
| Smad6 | -2.946152038 | 5.89E-07 |
| Tcta | 1.891268463 | 5.99E-07 |
| Rspo2 | 3.374937038 | 6.01E-07 |
| Spryd7 | 1.339144762 | 6.15E-07 |
| Hck | -1.932438646 | 6.16E-07 |
| Tmem130 | 1.432734465 | 6.23E-07 |
| Stc1 | 7.839568031 | 6.30E-07 |
| Phf11a | -2.983591976 | 6.32E-07 |
| Slc16a10 | -1.664345658 | 6.50E-07 |
| Ap1g2 | -1.775195094 | 6.50E-07 |
| Bach1 | -1.054280692 | 6.66E-07 |
| Cacng4 | 1.324338712 | 6.88E-07 |
| 1700017B05Rik | -1.459972015 | 6.99E-07 |
| Ngfrap1 | 0.873945648 | 6.99E-07 |
| Ankfy1 | -1.036093721 | 7.02E-07 |
| Spaca6 | -1.569882069 | 7.05E-07 |
| Ccnf | -2.055699493 | 7.05E-07 |
| Ndufb3 | 0.947051083 | 7.08E-07 |
| Tmem258 | 0.925140553 | 7.21E-07 |
| Asprv1 | -5.718531708 | 7.30E-07 |
| Scrg1 | 2.630786777 | 7.44E-07 |
| Snhg11 | 2.417144342 | 7.50E-07 |
| Fgfr3 | 2.497639537 | 7.54E-07 |
| P2rx7 | -1.615692689 | 7.58E-07 |
| Wdfy3 | -1.269006697 | 7.65E-07 |
| Tlr1 | -2.235724424 | 7.65E-07 |
| Rbp4 | 5.99437429 | 7.92E-07 |
| Phf5a | 1.037385868 | 7.92E-07 |
| Tspan3 | 0.906739734 | 7.92E-07 |
| Adam15 | -1.410812117 | 7.92E-07 |
| Ulbp1 | -2.638673771 | 7.92E-07 |
| Il1r2 | -5.866766369 | 7.92E-07 |
| Tox3 | 1.405241932 | 7.98E-07 |
| Alpk1 | -3.247055769 | 8.14E-07 |
| Otud5 | -1.062406753 | 8.18E-07 |
| Ndufc1 | 0.929176563 | 8.37E-07 |
| Uqcrq | 0.868806454 | 8.45E-07 |
| Ogn | 3.83199574 | 8.45E-07 |
| Ndufaf4 | 1.169591234 | 8.50E-07 |
| Atp1b2 | 1.686961123 | 8.53E-07 |
| Acsl5 | -1.707855348 | 8.54E-07 |
| Sostdc1 | 3.366901716 | 8.62E-07 |
| Lyz2 | -1.98780147 | 8.63E-07 |
| Rbm43 | -1.877981943 | 8.89E-07 |
| Nfasc | -1.101437679 | 8.95E-07 |
| Ctsa | -1.212130036 | 8.98E-07 |
| Aldoa | -1.247190027 | 9.27E-07 |
| H1f0 | 1.134220974 | 9.49E-07 |
| Emb | -1.990951539 | 9.52E-07 |
| Gch1 | -2.04896478 | 9.84E-07 |
| Agpat3 | -1.100809813 | 9.88E-07 |
| Tlr4 | -3.282932043 | 1.01E-06 |
| Ntng1 | 3.453367205 | 1.03E-06 |
| Csf3r | -2.229883269 | 1.04E-06 |
| Col25a1 | 4.622650241 | 1.04E-06 |
| Lrrc47 | -1.598343693 | 1.04E-06 |
| Myd88 | -1.952812971 | 1.04E-06 |
| Sipa1 | -1.543612708 | 1.05E-06 |
| Tbca | 1.052498204 | 1.06E-06 |
| Tnfrsf14 | -4.11031272 | 1.08E-06 |
| Arid3b | -1.317740594 | 1.10E-06 |
| Ncf4 | -1.726773859 | 1.11E-06 |
| Atp5j | 1.073245965 | 1.12E-06 |
| M6pr | -1.458521013 | 1.14E-06 |
| Ankrd13a | -1.484704615 | 1.20E-06 |
| Lrp4 | 1.407996893 | 1.20E-06 |
| Mmp17 | -1.923146251 | 1.22E-06 |
| Adamts1 | 2.292914841 | 1.23E-06 |
| Ltf | -6.654995869 | 1.26E-06 |
| Pfdn1 | 0.886254277 | 1.27E-06 |
| Prdx2 | 0.835594451 | 1.27E-06 |
| Odc1 | -0.902085988 | 1.28E-06 |
| Spi1 | -1.587424663 | 1.30E-06 |
| H6pd | -2.731049669 | 1.31E-06 |
| Rab8b | -1.073800567 | 1.31E-06 |
| Hyou1 | -1.473408141 | 1.32E-06 |
| Crip2 | 0.9622242 | 1.34E-06 |
| Sppl2a | -1.104328563 | 1.35E-06 |
| Tgfbrap1 | -1.325935253 | 1.36E-06 |
| Gclc | -1.793772464 | 1.37E-06 |
| Slc38a8 | -8.457101652 | 1.37E-06 |
| Bnip3 | -1.273343505 | 1.42E-06 |
| Ablim3 | 2.039978239 | 1.43E-06 |
| Fmnl1 | -1.940304834 | 1.43E-06 |
| Map1lc3a | 0.848636537 | 1.43E-06 |
| Bcl3 | -2.389017599 | 1.43E-06 |

| | | |
|---|---|---|
| Stac | 4.767591738 | 0.040179054 |
| Bche | 5.51682811 | 0.040196091 |
| Plxdc2 | 1.109553382 | 0.040196091 |
| Pilrb1 | -1.819900607 | 0.040196091 |
| Il1rn | -2.451708252 | 0.040197458 |
| E130308A19Rik | 0.8716287 | 0.040230095 |
| Dbnl | -0.512293163 | 0.040237032 |
| Lca5l | 1.989369725 | 0.040280998 |
| Pet100 | 0.640155475 | 0.040301136 |
| Fut10 | 1.704017466 | 0.04038134 |
| Slc2a12 | 2.427803238 | 0.040386578 |
| Rpl14 | 0.424191224 | 0.040386578 |
| Tst | 1.056999398 | 0.040448721 |
| 5330417C22Rik | 1.00557174 | 0.040448721 |
| Sec63 | -0.621117177 | 0.040472898 |
| Fam234b | -0.689518003 | 0.040512828 |
| Nog | 1.784086261 | 0.040515931 |
| Samd4 | 1.539944868 | 0.04052049 |
| Caap1 | -0.704985383 | 0.040525559 |
| Fgf12 | 1.054651057 | 0.040544595 |
| Ctage5 | -0.53899018 | 0.040544595 |
| Wdr90 | -0.844948256 | 0.040544595 |
| Isl1 | 2.965939146 | 0.040565437 |
| Pidd1 | -1.159350594 | 0.040580502 |
| Etv3 | -0.950839265 | 0.040652827 |
| Hira | 1.133132348 | 0.040695764 |
| Mllt11 | 0.493775864 | 0.040713972 |
| Gcfc2 | -1.443719749 | 0.040771161 |
| Hist1h1e | 0.553872428 | 0.040813732 |
| Lamtor3 | 0.484680007 | 0.040815775 |
| Celsr1 | 0.860477 | 0.040878782 |
| Arrdc1 | -0.71435496 | 0.040911697 |
| Lrig2 | -0.696891499 | 0.040946698 |
| Trim5 | 5.169876025 | 0.040970374 |
| Fam195b | 0.467405453 | 0.041053558 |
| Smad5 | 0.608915394 | 0.0411134 |
| Morf4l1 | 0.363410472 | 0.041187877 |
| Cnp | -0.7357942 | 0.041209812 |
| Il17b | 5.228536567 | 0.041248758 |
| Hmcn1 | 2.123983026 | 0.041255774 |
| Me2 | -0.560880364 | 0.041264275 |
| Rnf168 | 0.649106548 | 0.041364378 |
| Tmem205 | 0.730136601 | 0.041380031 |
| Pi4ka | -0.639506529 | 0.041434288 |
| Bicc1 | 2.382829877 | 0.041508835 |
| Zmym3 | 0.485984903 | 0.04153638 |
| Bmpr1a | 0.679736788 | 0.041591724 |
| Zfp750 | -2.505984844 | 0.041591724 |
| Stom | -1.856748665 | 0.041622402 |
| Gja5 | 3.512377183 | 0.04165941 |
| 9-Mar | 1.393516472 | 0.041777612 |
| Cdpf1 | 0.824171005 | 0.041787397 |
| Tspo2 | 5.296218752 | 0.041824025 |
| Tspan2 | 1.891283929 | 0.041824025 |
| Wdyhv1 | 0.84916756 | 0.041824025 |
| Rec8 | -5.883206189 | 0.041840613 |
| Ptpro | -0.741109844 | 0.041882092 |
| Wee1 | -0.872999215 | 0.042023145 |
| 2810006K23Rik | 0.877994049 | 0.042154991 |
| Dhrs7 | -1.117236793 | 0.042304268 |
| AI467606 | -1.130880707 | 0.042338075 |
| Pias2 | -0.513467339 | 0.042338101 |
| Hnrnpm | 0.494974067 | 0.042375309 |
| Atg13 | -0.550167237 | 0.042375309 |
| Nyap1 | -0.633628334 | 0.042375309 |
| Mgl2 | 5.372592913 | 0.042388497 |
| Tsc22d1 | 0.459504662 | 0.042388497 |
| Ipo9 | -0.568323119 | 0.042388497 |
| Nfatc4 | 1.254134342 | 0.042454965 |
| Hsd17b7 | -0.748153769 | 0.042518592 |
| Spem1 | 5.213467449 | 0.042525915 |
| Cuta | 0.559090829 | 0.042579213 |
| Ampd1 | -5.844571506 | 0.042607009 |
| Tfpi | 0.994537069 | 0.042660008 |
| Pcdhga11 | -1.04933343 | 0.042679382 |
| Ubox5 | -0.953239376 | 0.042685933 |
| Lad1 | 4.520503154 | 0.042739504 |
| Lsm10 | 0.605963364 | 0.042739504 |
| Arf2 | 0.440703256 | 0.042739504 |
| 2-Mar | -0.63707684 | 0.042739504 |
| AB124611 | -0.87125379 | 0.042853729 |
| Krt8 | 4.517640834 | 0.042867645 |
| Cdip1 | 0.609522241 | 0.042867645 |
| Usp15 | -0.554043447 | 0.042867645 |
| Gsta4 | 0.951525593 | 0.04309984 |
| Elmo3 | -2.843107378 | 0.043175834 |
| Lpar1 | 0.893920362 | 0.043187034 |
| Dnajc28 | 1.61619094 | 0.04321161 |
| Hbegf | 1.939369339 | 0.043282062 |
| Ikbip | 0.916400094 | 0.043285281 |
| Npas3 | 0.877814346 | 0.043317815 |
| Faim | 0.530834671 | 0.043518712 |
| Med24 | -0.543666488 | 0.043548511 |
| Trim34a | -4.183788866 | 0.043625517 |
| Nusap1 | 0.734575257 | 0.043767231 |
| Kdm4a | -0.687175961 | 0.043872966 |
| Supt4a | 0.519751184 | 0.043893583 |
| Ccser1 | -0.772462125 | 0.0439634 |
| Polh | -1.565713937 | 0.044081321 |
| Cldn23 | 5.744556749 | 0.044118942 |
| 4930438A08Rik | -2.965413902 | 0.04412538 |
| Syngr2 | -0.80945397 | 0.044243017 |
| Zfpl1 | 0.685458526 | 0.044292233 |
| Unc5d | 0.668576247 | 0.044335237 |
| Nop9 | -0.562470211 | 0.044425772 |
| Cirh1a | -0.602305699 | 0.044456432 |
| Bmi1 | -0.556402535 | 0.044488848 |
| Casp9 | -0.752123974 | 0.044537172 |
| Dock1 | 0.915720059 | 0.044543441 |
| Gm5113 | 1.063044971 | 0.044583991 |
| Lilra6 | -2.187588451 | 0.044583991 |
| Zfp456 | -2.284890552 | 0.044655195 |
| Nckap1 | 0.603652008 | 0.044660567 |
| Asb6 | -0.810560182 | 0.044660567 |
| Tmem101 | 0.700206367 | 0.044719112 |
| Uqcc2 | 0.486044627 | 0.044780955 |
| Aggf1 | 0.497064288 | 0.044805468 |
| Rsu1 | -0.579577146 | 0.044921246 |
| Etnk2 | 2.953266225 | 0.044995292 |

| | | |
|---|---|---|
| Tbc1d16 | 0.303570922 | 0.281070499 |
| Gipc1 | -0.309003711 | 0.281070499 |
| Galnt10 | -0.794989018 | 0.281070499 |
| Zfp658 | -0.809760705 | 0.281070499 |
| Supt5 | -0.315213137 | 0.281095633 |
| Vash1 | -0.608800256 | 0.2812177 |
| Tmub1 | 0.422473284 | 0.281409116 |
| Mmp28 | 2.359331267 | 0.281463522 |
| Kcnh1 | -2.080403888 | 0.281556854 |
| Arfl | -0.236763079 | 0.281808028 |
| Atoh7 | 3.944923923 | 0.281929028 |
| Sephs1 | -0.317638532 | 0.281929028 |
| Mlx | -0.365580573 | 0.281929028 |
| Pex16 | 0.436584523 | 0.282276829 |
| 9330161L09Rik | -3.816755627 | 0.282376434 |
| Rnf220 | 0.355350632 | 0.282456447 |
| Orai2 | -0.443322612 | 0.282456447 |
| Guca2a | -4.819689817 | 0.282456447 |
| Mrpl10 | 0.33373395 | 0.282560144 |
| Hsd3b3 | 3.950158684 | 0.282609069 |
| Srd5a1 | 0.792944506 | 0.282609069 |
| Rnf150 | -0.692546327 | 0.282609069 |
| Gm9821 | -1.570435841 | 0.282609069 |
| Pex1 | 0.584530122 | 0.282677396 |
| Tlr7 | -0.652585854 | 0.282810059 |
| Cluh | -0.437322272 | 0.283114254 |
| Rab9 | 0.334290227 | 0.283267851 |
| Sec13 | 0.283433066 | 0.283295423 |
| Pold2 | -0.32170788 | 0.283439005 |
| Stfa1 | -2.284954799 | 0.283441318 |
| Bahcc1 | -0.456047776 | 0.283454391 |
| Mkrn3 | -0.470359613 | 0.283454391 |
| Mtg2 | -0.392139032 | 0.283689577 |
| Rnf41 | -0.319929333 | 0.283727175 |
| Rad21l | -3.125824459 | 0.283899895 |
| Gcat | 0.413836078 | 0.283949327 |
| Pofut2 | -0.337597093 | 0.283990598 |
| Jam3 | 0.522285086 | 0.284138056 |
| D430041D05Rik | -0.424789765 | 0.284216586 |
| Ddit4 | -0.603784734 | 0.28439353 |
| Shprh | -0.294959351 | 0.284564889 |
| Cfap70 | 3.999473658 | 0.284672994 |
| Tigd3 | 1.35829454 | 0.284672994 |
| Timeless | -0.393259424 | 0.284672994 |
| Gemin5 | 0.489509069 | 0.284891849 |
| Efcab5 | 1.627064461 | 0.284984162 |
| Ddhd2 | -0.319944506 | 0.284984528 |
| Cib2 | 0.418908077 | 0.285221653 |
| B4galnt4 | -0.489013445 | 0.285221653 |
| Maf | 0.505777807 | 0.285635264 |
| Gramd1a | -0.266526314 | 0.285635264 |
| Slc35a1 | 0.36829869 | 0.285700921 |
| Pcdhb21 | -0.920259135 | 0.285731648 |
| Cd2 | 4.271328587 | 0.286608239 |
| 1700020L24Rik | -4.844865372 | 0.286608239 |
| 1810024B03Rik | 1.001568385 | 0.286824236 |
| Gm684 | -1.351616312 | 0.287145927 |
| Fastkd5 | -1.08881609 | 0.287205282 |
| G3bp2 | 0.226730797 | 0.287211582 |
| Nrde2 | 0.61367682 | 0.287279689 |
| Acacb | -1.135434725 | 0.287279689 |
| D830039M14Rik | -1.308184748 | 0.287279689 |
| Lifr | 0.500739062 | 0.287321023 |
| Hadha | 0.277921162 | 0.287593913 |
| Gm15800 | -0.296836197 | 0.287593913 |
| Zbed5 | 0.357909836 | 0.287623552 |
| Iah1 | 0.417049758 | 0.287665557 |
| Frmpd3 | 0.833210531 | 0.287802546 |
| Gas2l3 | -0.513342823 | 0.287802546 |
| Ldb3 | 3.736807696 | 0.287912246 |
| Cdkl3 | 0.484369881 | 0.287944368 |
| Mastl | -0.461731606 | 0.287963033 |
| Caps2 | 4.321842133 | 0.288078879 |
| Krt17 | 4.321842133 | 0.288078879 |
| Ccdc162 | 3.982613121 | 0.288078879 |
| Vps45 | 0.386229229 | 0.288133824 |
| Chst11 | -0.576032356 | 0.288133824 |
| Smad9 | 4.274671786 | 0.288173403 |
| Lgi1 | -0.63824524 | 0.288173403 |
| Lgi2 | 1.388654183 | 0.2882599 |
| Gm17732 | -2.379178198 | 0.288497515 |
| Nek5 | 4.292275941 | 0.288552047 |
| Eps8l1 | 3.281030579 | 0.288615268 |
| Kiz | 0.450874486 | 0.288615268 |
| Mrpl48 | 0.304453991 | 0.288615268 |
| Racgap1 | 0.354415445 | 0.28865958 |
| Asap1 | -0.275921313 | 0.28865958 |
| Eif3d | -0.232572511 | 0.288910052 |
| Zfp101 | -0.506324496 | 0.28899794 |
| Lrrtm1 | -0.574861937 | 0.289139658 |
| Tuft1 | 0.746027122 | 0.289397784 |
| Akr1c12 | -4.832727394 | 0.289411617 |
| Ctfl | 1.031086674 | 0.289562495 |
| Pomgnt1 | -0.367006009 | 0.289570323 |
| Prickle3 | -0.932915172 | 0.289570323 |
| Cdr1 | 0.611669329 | 0.289869995 |
| Cenpb | 0.327138343 | 0.289873684 |
| Zic5 | 1.53788573 | 0.290189987 |
| Olig2 | 1.008726984 | 0.290189987 |
| Zfp58 | -1.266575547 | 0.290373906 |
| 4930555F03Rik | -1.534906804 | 0.290818205 |
| Sned1 | 1.22251961 | 0.29082161 |
| 5730409E04Rik | 0.725062444 | 0.29082161 |
| Rps26-ps1 | 0.404690572 | 0.29082161 |
| Thnsl2 | -1.300197784 | 0.29082161 |
| Syn2 | 0.627629566 | 0.29096815 |
| Ung | -0.490380703 | 0.290986545 |
| Thbs4 | 4.353611288 | 0.291324992 |
| Tex35 | -4.491135433 | 0.291347185 |
| Cramp1l | -0.337419968 | 0.29153229 |
| Cmklr1 | -0.951286436 | 0.291551547 |
| 2310061I04Rik | 0.544646073 | 0.291565088 |
| Topbp1 | -0.315935825 | 0.291659777 |
| Plekhg5 | -0.68556806 | 0.291698322 |
| Atg16l1 | 0.454803664 | 0.291847841 |
| Cd209f | 4.269197491 | 0.292102481 |
| Fam171a1 | 0.459392511 | 0.292252547 |
| Snrk | -0.288277218 | 0.292252547 |
| Lif | -2.211647354 | 0.292444671 |

| | | |
|---|---|---|
| Mfhas1 | -0.234651788 | 0.674090197 |
| Fam76b | -0.140464674 | 0.674160049 |
| Klhl10 | -1.100295249 | 0.674174227 |
| Npepps | 0.114786746 | 0.674231193 |
| Mphosph10 | 0.171621618 | 0.674939629 |
| Arl4c | -0.12827911 | 0.675116612 |
| Thsd1 | 1.658430659 | 0.675211105 |
| Nek8 | 0.513119049 | 0.675245065 |
| Ppp1r9a | 0.163561131 | 0.675245065 |
| Cdc23 | 0.131535813 | 0.675245065 |
| Mtf1 | -0.15825894 | 0.675245065 |
| Skp2 | -0.221185228 | 0.675245065 |
| Cacna2d4 | 1.410940161 | 0.675613041 |
| Pdik1l | -0.176952383 | 0.675731447 |
| Notum | 0.907914556 | 0.676104112 |
| Mtrf1l | 0.23958371 | 0.676104112 |
| Astn2 | 0.872730638 | 0.676338513 |
| Slc35e2 | -0.138023596 | 0.676338513 |
| Klhl7 | 0.12706792 | 0.676435296 |
| Mmgt1 | 0.122756385 | 0.676435296 |
| Pcdha4 | 2.132689627 | 0.676625752 |
| Fndc3c1 | -0.877522101 | 0.676798419 |
| Prkcq | 0.494986509 | 0.677013783 |
| Msantd4 | 0.156272999 | 0.677235704 |
| Pmel | -1.540921022 | 0.677526412 |
| Gsx1 | 0.791958135 | 0.678277736 |
| Grik5 | -0.203592689 | 0.678277736 |
| Lins1 | 0.282405194 | 0.67848248 |
| Zrsr1 | 0.260080902 | 0.678503392 |
| Sntg1 | 0.227618268 | 0.678503392 |
| Apeh | -0.153639288 | 0.678503392 |
| Kcnc3 | -0.340400962 | 0.678503392 |
| Kctd17 | 0.180977817 | 0.67862287 |
| Armcx2 | 0.138873034 | 0.67862287 |
| Cerkl | -0.862586791 | 0.67862287 |
| Comp | -1.224839817 | 0.678740381 |
| Gp6 | 2.086020034 | 0.67893742 |
| St8sia5 | 1.265409224 | 0.67893742 |
| Baalc | 0.546293568 | 0.67893742 |
| Cish | 0.270645852 | 0.67893742 |
| Babam1 | 0.126231367 | 0.67893742 |
| Pank2 | -0.185519475 | 0.67893742 |
| Elac1 | -0.432780225 | 0.67893742 |
| Mei4 | -1.55871513 | 0.67893742 |
| Paqr6 | 0.997157511 | 0.6791502 |
| Zfp40 | 0.254114449 | 0.6791502 |
| Reps2 | 0.247322203 | 0.679289327 |
| AA415398 | 0.547343999 | 0.679290108 |
| Zfp354c | 0.159402676 | 0.679290108 |
| Pdzd7 | -0.981207302 | 0.679377808 |
| Kcnc4 | 0.69731394 | 0.679392199 |
| Zfp423 | -0.259202749 | 0.679663338 |
| Gm10392 | -1.302142695 | 0.679663338 |
| Eif2s3y | 0.200272272 | 0.679671989 |
| Ifnb1 | -0.630322339 | 0.679671989 |
| Wdr5 | -0.14243441 | 0.679877426 |
| Glt28d2 | -0.858237058 | 0.679877426 |
| Bace1 | -0.134781628 | 0.679885022 |
| Lats1 | -0.136820594 | 0.679885022 |
| Adgrg5 | -0.375412854 | 0.680004314 |
| Dtwd1 | 0.224142401 | 0.680059396 |
| Fstl4 | 0.540519323 | 0.680280929 |
| Rrp9 | 0.1470554 | 0.680280929 |
| Runx2 | -1.053619721 | 0.680487672 |
| Tfcp2l1 | 0.889862823 | 0.680951952 |
| Atcay | -0.144802281 | 0.680951952 |
| Atp8b3 | -0.782074801 | 0.680951952 |
| Jmjd8 | -0.22047759 | 0.681020172 |
| Cct3 | 0.089950113 | 0.681068279 |
| F2 | 0.815973559 | 0.681313414 |
| Cpne1 | -1.484368858 | 0.681348458 |
| Zfp36 | -0.182635786 | 0.6813658 |
| Angptl7 | 0.922844229 | 0.681836196 |
| Tpr | 0.096134241 | 0.681836196 |
| Ccdc60 | -1.239912042 | 0.681836196 |
| Gnao1 | 0.13143605 | 0.681893956 |
| Timp2 | -0.143824056 | 0.681893956 |
| Alkbh2 | -0.384441025 | 0.681893956 |
| Tspyl5 | 0.179962646 | 0.681948447 |
| Uba2 | -0.109263788 | 0.682152449 |
| Serpina3i | 1.083364807 | 0.682482043 |
| Syn3 | 0.229125074 | 0.682482043 |
| Gas8 | 0.177112931 | 0.682482043 |
| Skida1 | 0.170276778 | 0.682482043 |
| Pip4k2b | -0.261224971 | 0.682482043 |
| 0610040J01Rik | -0.276167675 | 0.682482043 |
| Galnt9 | -0.392170992 | 0.682482043 |
| Dnajc22 | -0.868263383 | 0.682482043 |
| Atp6v0a4 | -1.173744624 | 0.682482043 |
| Fasn | -0.110842852 | 0.682500812 |
| Lrrc34 | -1.076571595 | 0.682500812 |
| 2700029M09R | -0.157557568 | 0.683048374 |
| Fam160a1 | -0.648719486 | 0.683059142 |
| Rab1a | 0.100758105 | 0.683201872 |
| Vwa3a | -0.671302562 | 0.683389238 |
| Sv2b | -0.346080384 | 0.683392812 |
| Rnf39 | 1.343682416 | 0.683434497 |
| Sec23ip | 0.140144687 | 0.683643104 |
| Rell1 | 0.324974991 | 0.683649965 |
| Thtpa | 0.134881492 | 0.683689126 |
| Saal1 | 0.157557086 | 0.683886189 |
| Lsg1 | -0.148411558 | 0.683886189 |
| Chtop | 0.111135321 | 0.68409445 |
| Lrfn3 | 0.366639587 | 0.684122598 |
| Foxj3 | 0.18576779 | 0.684172631 |
| Sidt2 | -0.159754287 | 0.684207905 |
| Armc3 | -0.822215317 | 0.684207905 |
| Otof | 0.749980461 | 0.684218444 |
| Prpf31 | 0.106189298 | 0.684218444 |
| Rimbp3 | 1.162428232 | 0.685128911 |
| Neb | 0.322197074 | 0.685160038 |
| Zfp593 | 0.825941414 | 0.685277898 |
| Atg4a-ps | 2.047884365 | 0.685439174 |
| Mettl16 | 0.146631108 | 0.685439174 |
| Npat | -0.215817931 | 0.685439174 |
| Gm14420 | -0.598699988 | 0.685439174 |
| Alkbh1 | 0.175557853 | 0.685469028 |
| Stx2 | -0.178404625 | 0.685469028 |
| Mdm1 | 0.176322223 | 0.685491098 |

| | | |
|---|---|---|
| Dennd4a | -1.658191669 | 1.45E-06 |
| 2010107E04Rik | 0.929939344 | 1.47E-06 |
| Dynlrb1 | 0.869272754 | 1.47E-06 |
| mt-Atp8 | 2.010772049 | 1.55E-06 |
| Stk10 | -1.824054472 | 1.56E-06 |
| Fgd3 | -2.007873903 | 1.58E-06 |
| Cd38 | -3.741264714 | 1.63E-06 |
| Kcnip3 | 1.658314296 | 1.63E-06 |
| Rpl18a | 0.790382617 | 1.64E-06 |
| Serf1 | 0.998103294 | 1.66E-06 |
| Mfsd1 | -1.260421057 | 1.66E-06 |
| Skap2 | -1.574647169 | 1.68E-06 |
| Adgrg6 | -3.907208496 | 1.69E-06 |
| Dut | 0.938663588 | 1.72E-06 |
| Cntn2 | -1.676661392 | 1.74E-06 |
| Hbb-y | 2.865607489 | 1.74E-06 |
| Cyp1b1 | 7.634677836 | 1.76E-06 |
| Rnf130 | -1.096542269 | 1.79E-06 |
| Kctd12 | -0.963795318 | 1.80E-06 |
| Crabp2 | 3.379853685 | 1.82E-06 |
| Rassf4 | -1.394866498 | 1.83E-06 |
| Isyna1 | -1.168771265 | 1.86E-06 |
| Eif2s2 | 0.91318471 | 1.88E-06 |
| Ptpn23 | -1.339668332 | 1.88E-06 |
| Pcsk1n | 1.232472372 | 1.88E-06 |
| Atxn7l3b | 0.834054638 | 1.91E-06 |
| Rnd3 | 1.17799964 | 1.95E-06 |
| Nab1 | -1.980144065 | 2.00E-06 |
| Dse | -1.629860264 | 2.00E-06 |
| Rpl13a | 0.90538857 | 2.00E-06 |
| Metrnl | -2.186762902 | 2.01E-06 |
| Prickle1 | -1.371178845 | 2.05E-06 |
| Atp5j2 | 0.862401099 | 2.10E-06 |
| St3gal1 | -2.029784533 | 2.14E-06 |
| Trmt112 | 1.236996579 | 2.17E-06 |
| Snrpe | 0.988415709 | 2.20E-06 |
| Bcas2 | 0.941037073 | 2.20E-06 |
| Flnc | 3.595826206 | 2.26E-06 |
| Ccr7 | -8.633455235 | 2.26E-06 |
| Crcp | 1.249846725 | 2.28E-06 |
| Tfeb | -1.937371502 | 2.28E-06 |
| Nup35 | 1.535614516 | 2.28E-06 |
| Npm1 | 0.921031686 | 2.32E-06 |
| Cdc42ep2 | -2.18975977 | 2.32E-06 |
| Ndufb9 | 0.869503001 | 2.39E-06 |
| Abcc8 | -2.415908076 | 2.45E-06 |
| Cxcl12 | 2.196663814 | 2.46E-06 |
| Trappc10 | -1.412720455 | 2.47E-06 |
| Serp1 | -1.510062804 | 2.47E-06 |
| Cox4i1 | 0.861348907 | 2.51E-06 |
| Tbk1 | -1.188342545 | 2.52E-06 |
| 4930452B06Rik | 4.086616012 | 2.59E-06 |
| Chst14 | -1.816813301 | 2.59E-06 |
| Ripk1 | -1.735831698 | 2.61E-06 |
| Inppl1 | -1.380973777 | 2.61E-06 |
| Flii | -1.200496078 | 2.62E-06 |
| Wfdc21 | -2.326865687 | 2.63E-06 |
| Muc5b | -4.96278754 | 2.68E-06 |
| Zfr | 0.873306622 | 2.68E-06 |
| Ibsp | 7.428768606 | 2.69E-06 |
| Mif | 0.94779806 | 2.76E-06 |
| Hmgb1 | 1.373996189 | 2.85E-06 |
| Cldn5 | 7.239904485 | 2.85E-06 |
| Paqr9 | 4.493576692 | 2.89E-06 |
| Atp5k | 0.992164809 | 2.91E-06 |
| Cybb | -1.956283957 | 2.92E-06 |
| Banf1 | 0.994892195 | 2.92E-06 |
| Rpl21 | 1.75109709 | 2.93E-06 |
| Cox20 | 1.004989165 | 2.93E-06 |
| Ldlrap1 | -1.916636405 | 2.99E-06 |
| Stmn2 | 0.916003479 | 3.06E-06 |
| D330045A20Rik | -2.829407146 | 3.10E-06 |
| Sik1 | -1.420538115 | 3.19E-06 |
| Ndufa11 | 0.869839699 | 3.25E-06 |
| Retnlg | -6.602144122 | 3.25E-06 |
| Surf4 | -0.898807111 | 3.27E-06 |
| Cebpb | -2.699337385 | 3.28E-06 |
| Bcl6 | -1.599934537 | 3.33E-06 |
| Rab11fip1 | -2.834632387 | 3.34E-06 |
| Sh3tc1 | -3.111049594 | 3.34E-06 |
| Slc13a4 | 7.488646219 | 3.35E-06 |
| Lipa | -1.283839928 | 3.37E-06 |
| Rap1b | -1.500194077 | 3.39E-06 |
| Rab27a | -3.203430464 | 3.44E-06 |
| Ago1 | -1.079386093 | 3.48E-06 |
| Ndufa1 | 1.003975786 | 3.57E-06 |
| Tmem189 | -1.955655221 | 3.58E-06 |
| Ndufa3 | 1.132715397 | 3.69E-06 |
| Ostf1 | -1.394337728 | 3.69E-06 |
| Atf6 | -1.443564573 | 3.69E-06 |
| Arhgap24 | -2.269156125 | 3.69E-06 |
| 1110004E09Rik | 1.319113689 | 3.73E-06 |
| Asah1 | -1.067387996 | 3.80E-06 |
| Megf9 | -1.081100532 | 3.85E-06 |
| Dmrtb1 | 2.377363945 | 3.92E-06 |
| Prkcd | -1.36537688 | 4.01E-06 |
| Egfl7 | 3.892324923 | 4.11E-06 |
| Dlk1 | 3.717094218 | 4.15E-06 |
| Fbln5 | 7.570405035 | 4.21E-06 |
| Uchl1 | 0.955574913 | 4.27E-06 |
| Rpl31 | 0.967655494 | 4.41E-06 |
| Gla | -2.158085024 | 4.41E-06 |
| Tnc | 1.928968885 | 4.44E-06 |
| Glrx | -1.927669989 | 4.50E-06 |
| Dusp16 | -1.552190591 | 4.64E-06 |
| Pdlim3 | 7.572089857 | 4.65E-06 |
| Ptger2 | -5.110854083 | 4.75E-06 |
| Ucp2 | -1.166635673 | 4.78E-06 |
| Il16 | -1.927972303 | 4.78E-06 |
| Prune2 | -1.965797338 | 4.78E-06 |
| Dock5 | -3.706651604 | 4.78E-06 |
| Dusp8 | 0.994627741 | 4.83E-06 |
| Tlr13 | -2.263275309 | 4.87E-06 |
| Hmha1 | -1.881664569 | 5.14E-06 |
| Hps6 | -3.67311374 | 5.16E-06 |
| Stt3a | -0.930040241 | 5.18E-06 |
| Plek | -1.818225666 | 5.18E-06 |
| Cndp2 | -1.363970178 | 5.23E-06 |
| Cdh18 | 7.312468084 | 5.32E-06 |

| | | |
|---|---|---|
| Elk4 | -0.625480736 | 0.044995292 |
| Khk | 1.008382484 | 0.044999785 |
| Stat4 | -3.657366448 | 0.045040921 |
| Ptprs | -0.471672724 | 0.045088818 |
| Pcsk5 | 1.657032102 | 0.045097859 |
| Arid2 | 0.544175643 | 0.045104861 |
| Mcf2 | 5.505350935 | 0.045250973 |
| Camkmt | 0.834697107 | 0.045250973 |
| Rc3h1 | -0.69494634 | 0.045250973 |
| Zfp14 | -1.135578851 | 0.045294821 |
| Gbp5 | -2.544807564 | 0.045381343 |
| Asb7 | -0.664025681 | 0.045429536 |
| Ilvbl | -0.736478695 | 0.045429536 |
| Tnf | -1.653511117 | 0.0454639 |
| Usp47 | -0.447601231 | 0.045481504 |
| Atp5g2 | 0.426891932 | 0.045498939 |
| Fbxo10 | -0.890242825 | 0.045538222 |
| Zfp654 | -0.680866754 | 0.045573885 |
| Hsp90ab1 | 0.366730883 | 0.045578092 |
| Arhgef26 | 0.92401872 | 0.045607849 |
| Zfp790 | -0.715658692 | 0.045607849 |
| Suco | -0.529353271 | 0.045621108 |
| C1qtnf1 | 5.239777465 | 0.045647847 |
| Sh3bgr | 3.887678806 | 0.045660243 |
| Usf3 | 1.108212808 | 0.045660243 |
| Coa4 | 0.885480972 | 0.045660243 |
| Btg1 | -0.443186867 | 0.045660243 |
| Myo15b | -2.065790958 | 0.045660243 |
| Arhgef3 | -2.507835487 | 0.045694082 |
| Sdr39u1 | 0.655027385 | 0.045704084 |
| Mrpl11 | 0.445148253 | 0.045734 |
| Eda | -2.450999442 | 0.045749654 |
| Ubap1 | -0.633992098 | 0.045873275 |
| Rufy3 | 0.463399711 | 0.045874522 |
| Cntfr | 0.894867 | 0.045952766 |
| Ier3ip1 | 0.580300628 | 0.045952766 |
| Fat1 | 0.866964785 | 0.045997391 |
| Mprip | 0.489791035 | 0.045997391 |
| BC031181 | 0.432242002 | 0.045997391 |
| Ppm1f | -0.956696253 | 0.045997391 |
| Aldh4a1 | 0.95351693 | 0.04600741 |
| Slco4c1 | -4.43458855 | 0.046025768 |
| Mtmr1 | -0.564352437 | 0.046040521 |
| Smox | -0.635061968 | 0.046040521 |
| Ube2w | 0.502546557 | 0.04609008 |
| Cds1 | -1.804648244 | 0.046111848 |
| Cmpk2 | -1.454240589 | 0.046131336 |
| Ccdc39 | 1.063256526 | 0.046180437 |
| Tigd5 | -1.845180108 | 0.046180437 |
| Pigs | -0.585744793 | 0.046275331 |
| Tle3 | -0.734557216 | 0.046275331 |
| Ano5 | 2.582141702 | 0.046325043 |
| Nim1k | 1.243714748 | 0.046343181 |
| BC089491 | 4.59091911 | 0.046400126 |
| Rpl23a-ps3 | 0.720312497 | 0.046539812 |
| Marco | -2.208002518 | 0.046539812 |
| Fgf5 | 5.139032464 | 0.046548775 |
| Ddx46 | 0.523681266 | 0.046551133 |
| Cnot4 | 0.496788305 | 0.046551223 |
| Cirbp | 0.427465646 | 0.046636841 |
| Crybb1 | 2.125151056 | 0.04672804 |
| Nme7 | 0.811493907 | 0.046818499 |
| Gm10260 | 0.401444886 | 0.046818499 |
| Kcnt1 | -1.494238018 | 0.046887755 |
| Ppig | 0.428637992 | 0.046890249 |
| Apex2 | -0.677314864 | 0.046912127 |
| Dus2 | 0.714145411 | 0.04696119 |
| Zbtb11 | -0.705719397 | 0.04696119 |
| Rnf6 | -0.846559622 | 0.04696119 |
| Malt1 | -1.046194353 | 0.047075854 |
| Rpa1 | -0.483074082 | 0.047099934 |
| Lman2 | -0.615591784 | 0.047170395 |
| Cygb | 0.981791077 | 0.047261453 |
| Uck2 | 0.566486199 | 0.047286056 |
| Wdr20 | -0.659856605 | 0.047286056 |
| Adcy2 | 3.260329132 | 0.047327032 |
| Kti12 | 0.747673022 | 0.047384799 |
| Pccb | 0.699682916 | 0.047384799 |
| Naca | 0.350296427 | 0.047384799 |
| Got1l1 | 5.815796177 | 0.047533107 |
| Drc1 | 2.561412001 | 0.04756788 |
| Ttc41 | 3.860336454 | 0.047647141 |
| Mrps23 | 0.639207578 | 0.048041997 |
| Atat1 | 0.498407668 | 0.048073264 |
| Ext1 | 0.760515239 | 0.048095147 |
| Mpzl1 | 0.437933709 | 0.048127737 |
| Cdk17 | -0.585748289 | 0.048150783 |
| Adamts18 | -0.961471864 | 0.04815137 |
| Bicd1 | 0.558721481 | 0.048162167 |
| Pnrc1 | -0.417353606 | 0.048162167 |
| Tpst2 | -0.578644417 | 0.048259073 |
| Mef2d | -0.584747496 | 0.048304563 |
| Pdzd11 | 0.470737341 | 0.048357115 |
| Slitrk1 | -1.309785147 | 0.048357115 |
| Enkur | 2.409410248 | 0.048361654 |
| Tlr5 | -6.105852154 | 0.048361654 |
| Mgat3 | 1.116946927 | 0.048441427 |
| Trim44 | 0.624606545 | 0.048677475 |
| Bcdin3d | 1.071433244 | 0.048728091 |
| Cops2 | 0.450160625 | 0.048915094 |
| Tmem159 | -1.123666499 | 0.048943571 |
| Camk2d | -0.693742156 | 0.048954651 |
| Rab21 | 1.253737601 | 0.049003436 |
| Sbsn | 1.355998048 | 0.049024659 |
| Prkrip1 | 0.58609522 | 0.049024659 |
| Bop1 | -0.625774519 | 0.049024659 |
| Mc4r | -1.368225699 | 0.049024659 |
| Chchd4 | 0.518720751 | 0.049029003 |
| Ly6h | 0.584023103 | 0.049088565 |
| Polr3b | 0.701551829 | 0.049126923 |
| Ufc1 | 0.44262628 | 0.049126923 |
| 1110059G10Rik | 0.675045506 | 0.049271392 |
| Cyp2a5 | 5.340117303 | 0.049491236 |
| Kif26a | -0.997380059 | 0.049491236 |
| Tnnt2 | -2.780868296 | 0.049631968 |
| Bola2 | 0.472103254 | 0.04972534 |
| Arpc1a | 0.408943451 | 0.04972534 |
| Srgn | -1.815191554 | 0.049784766 |
| Taok3 | -0.547548715 | 0.049792629 |

| | | |
|---|---|---|
| Akr1b3 | 0.380690456 | 0.292759727 |
| I830077J02Rik | -0.636661181 | 0.292759727 |
| Sema6b | -0.723661321 | 0.292821539 |
| Mier3 | 0.474823696 | 0.292827945 |
| Spsb2 | -0.501894215 | 0.292827945 |
| Mybphl | -2.069417545 | 0.292827945 |
| Polr3d | 0.437818894 | 0.292838061 |
| Dhrs1 | -0.321786834 | 0.292838061 |
| Ctdnep1 | -0.420734877 | 0.292838061 |
| 4930550C14Rik | 3.419974011 | 0.292862171 |
| Napg | 0.309230864 | 0.292862171 |
| Trdmt1 | -0.550675649 | 0.292866447 |
| Uncx | 4.281523195 | 0.293026455 |
| Lag3 | 1.370306171 | 0.293026455 |
| Bloc1s6 | 0.321362334 | 0.293026455 |
| Eefsec | -0.367732816 | 0.293026455 |
| Zbtb40 | -0.496856385 | 0.293026455 |
| Glra2 | 0.572282785 | 0.293200618 |
| Fgfr1op2 | -0.270440114 | 0.29326089 |
| P2ry1 | 1.221204344 | 0.293289761 |
| Cpped1 | 0.515813852 | 0.293364952 |
| Zfp628 | -0.768314068 | 0.293364952 |
| Cep170b | 0.560579253 | 0.29351587 |
| Gcg | -1.941313207 | 0.293651505 |
| Gpr87 | 4.33501704 | 0.293814584 |
| Tysnd1 | -0.422084647 | 0.293814584 |
| Sema6c | -0.460667452 | 0.293814584 |
| Tmem200b | 4.054078956 | 0.293918159 |
| Magt1 | -0.373818212 | 0.294050401 |
| Tspan17 | 0.493405011 | 0.294238904 |
| Wnt9b | 2.538693093 | 0.294245979 |
| Ptk2 | -0.324112897 | 0.294245979 |
| Arih2 | -0.361716414 | 0.294245979 |
| Zfp598 | -0.59072346 | 0.294245979 |
| Prr16 | 1.455997115 | 0.294390111 |
| Syncrip | 0.221499246 | 0.294390111 |
| Micall1 | -0.650244856 | 0.294390111 |
| Galnt15 | -1.751063895 | 0.294404636 |
| Bsg | 0.290595982 | 0.294764929 |
| Il17rd | 1.126040095 | 0.294922651 |
| 1700012B09Rik | 1.093271738 | 0.294922651 |
| Chmp1a | -0.233331622 | 0.294922651 |
| Zic1 | 1.236224055 | 0.295065199 |
| Chrna1 | 3.884189706 | 0.29511449 |
| Slc6a11 | 0.874842901 | 0.29511449 |
| Wwox | -0.48582433 | 0.29511449 |
| Cnih4 | 0.321267851 | 0.295194502 |
| Tac1 | 1.001585446 | 0.295343502 |
| Gemin6 | 0.44440094 | 0.295343502 |
| Cdh2 | -0.38885759 | 0.295660035 |
| Cds2 | -0.417162683 | 0.295700136 |
| Ptdss2 | 0.340496205 | 0.295786124 |
| Ntrk2 | 0.425227494 | 0.296132001 |
| Stradb | 0.271935968 | 0.296169542 |
| Mknk2 | -0.385899598 | 0.296287019 |
| Lrriq1 | 1.63535614 | 0.2964274 |
| A430033K04Rik | -0.460047487 | 0.296610141 |
| Bcl2l12 | 0.607163304 | 0.296883597 |
| E2f7 | -0.470771605 | 0.296883597 |
| Exo1 | -0.498386307 | 0.296908741 |
| Gpr153 | 0.924381183 | 0.296943817 |
| Eef1d | 0.219447439 | 0.297098521 |
| Usp8 | -0.380077276 | 0.297115226 |
| Ccr10 | -4.759909225 | 0.297146085 |
| Cd200r3 | -4.759909225 | 0.297146085 |
| Opn4 | -4.759909225 | 0.297146085 |
| Zfp541 | -2.237852676 | 0.297298908 |
| Syde2 | -1.026507303 | 0.29735985 |
| Med21 | 0.311191805 | 0.297425273 |
| Snrnp35 | 0.429117322 | 0.297511301 |
| Pgk1-rs7 | -0.872607339 | 0.297511301 |
| Zfp651 | 0.698900357 | 0.297786526 |
| Ddx27 | 0.309814132 | 0.298050527 |
| Gprin2 | 1.493648363 | 0.298123474 |
| Gp1bb | 1.414817244 | 0.298471137 |
| Aldh6a1 | -0.420510366 | 0.298625479 |
| Acot11 | -1.3677954 | 0.298625479 |
| Bbof1 | 1.014703897 | 0.298732143 |
| Lipo1 | 0.579644809 | 0.298732143 |
| Gclm | -0.289855571 | 0.298732143 |
| Tnfsf13 | -3.752870184 | 0.298732143 |
| Oprk1 | 3.293419841 | 0.298735402 |
| Sgce | 0.435513931 | 0.298847099 |
| Ppp6r3 | -0.289316688 | 0.298847099 |
| Sh2d4b | -1.253686381 | 0.298847099 |
| Sphkap | 0.617323099 | 0.2989034 |
| Zfp385b | -0.850092544 | 0.29905684 |
| Cdyl | 0.701655588 | 0.299110871 |
| Fam103a1 | 0.34915021 | 0.299129662 |
| Slit3 | 0.863265575 | 0.299150228 |
| Barx2 | 4.207662131 | 0.299221303 |
| Zfp639 | 0.405785001 | 0.299525731 |
| Sncg | -1.097166593 | 0.299850894 |
| Crhbp | 1.881797712 | 0.300099796 |
| Snph | 0.693632125 | 0.300320965 |
| Hnrnpdl | 0.202306048 | 0.300320965 |
| Ache | 0.921473135 | 0.300407698 |
| Tceal5 | 0.599924957 | 0.300407698 |
| Zfp128 | -0.808185649 | 0.300407698 |
| Tg | -2.071676792 | 0.300627974 |
| Bard1 | 0.692911385 | 0.300649082 |
| Ephb1 | -0.401542296 | 0.300681928 |
| AU040320 | -0.337659184 | 0.300752915 |
| Rnf44 | -0.281069938 | 0.300874699 |
| Ikzf5 | 0.748060097 | 0.300917367 |
| St8sia3 | 0.3435552 | 0.301087346 |
| Actn3 | 4.216912043 | 0.301289358 |
| Vps37c | -0.380724551 | 0.301429526 |
| Trappc5 | 0.317105251 | 0.301661542 |
| Dusp18 | 0.394678099 | 0.301690997 |
| Tert | -0.681539102 | 0.301852038 |
| Lypd6 | -0.564147521 | 0.301931036 |
| Ssx2ip | -0.353410456 | 0.302215336 |
| Zfp280b | 0.667879573 | 0.302270595 |
| Cep104 | -0.371113207 | 0.302270595 |
| Sncaip | -0.375950176 | 0.302352128 |
| Bcr | 0.429539348 | 0.302352277 |
| Scamp1 | -0.295803287 | 0.302352277 |
| Serinc2 | 0.327826055 | 0.302440347 |

| | | |
|---|---|---|
| Tgm4 | -1.229471442 | 0.685896304 |
| AI846148 | -0.192967276 | 0.686065166 |
| Rpia | -0.32450069 | 0.686257136 |
| Lzts1 | 0.145169157 | 0.686263326 |
| Gm10709 | -0.823353328 | 0.68646576 |
| Sln | 0.659145283 | 0.686702682 |
| Gabarap | 0.092588343 | 0.686702682 |
| Bcl9 | -0.13483141 | 0.686702682 |
| Gm7334 | 0.43141367 | 0.686964974 |
| Zcchc2 | -0.31109329 | 0.687321727 |
| Cdk10 | 0.161415468 | 0.687608057 |
| Ube3b | 0.154874076 | 0.687642087 |
| Mrps31 | -0.199154218 | 0.687708853 |
| Artn | 1.191137698 | 0.688057911 |
| Ddx17 | 0.106961766 | 0.688057911 |
| Odf2l | 0.201576087 | 0.688627491 |
| Fam184b | -0.505040495 | 0.688717095 |
| Tmem245 | 0.156268416 | 0.689080219 |
| Gpat2 | -1.471949147 | 0.689080219 |
| Ptms | -0.102754623 | 0.689173047 |
| Pcca | -0.176505705 | 0.689462913 |
| Zfp273 | 0.538915613 | 0.689602802 |
| BC025920 | -0.299894168 | 0.689618462 |
| Cbwd1 | -0.186979324 | 0.689739661 |
| Msto1 | 0.145514891 | 0.690052651 |
| Map3k9 | -0.201463659 | 0.690190345 |
| Nr4a3 | -0.279969642 | 0.690250669 |
| Cactin | -0.183763921 | 0.690387287 |
| Gnl2 | 0.143833223 | 0.690527768 |
| Vkorc1l1 | 0.166568048 | 0.690691045 |
| Gtf3c5 | 0.156330204 | 0.690691045 |
| Rab5b | -0.113226713 | 0.690691045 |
| Mus81 | -0.168824881 | 0.690691045 |
| Robo2 | -0.196874542 | 0.690691045 |
| Moxd1 | -0.88247663 | 0.690691045 |
| Slc25a51 | 0.112836118 | 0.691038014 |
| Yif1b | -0.148438295 | 0.69127291 |
| Serpinc1 | -0.719350017 | 0.691327815 |
| Gpm6a | 0.132153297 | 0.691383761 |
| Oxr1 | -0.127190843 | 0.691410528 |
| Ssu72 | 0.105344233 | 0.69153804 |
| Iscu | -0.116044333 | 0.69153804 |
| Gm17252 | 2.264750786 | 0.691989914 |
| Pgam5 | -0.130938929 | 0.69212836 |
| Tnfrsf23 | 0.644876074 | 0.692472874 |
| Dgkd | -0.162041135 | 0.692472874 |
| Iqcd | -0.797623259 | 0.692472874 |
| Psmd7 | 0.092844606 | 0.692630217 |
| Nptn | -0.125896202 | 0.692630217 |
| Bre | -0.133112105 | 0.692736637 |
| Dyrk3 | 0.17935059 | 0.692795772 |
| Parl | 0.120522557 | 0.692795772 |
| Dennd2d | -0.777235996 | 0.693176731 |
| BC037034 | -0.235819047 | 0.693261697 |
| Zfp712 | 0.602841053 | 0.693483782 |
| Zbtb45 | 0.242292937 | 0.693483782 |
| Afg3l1 | 0.165844761 | 0.693483782 |
| Obfc1 | -0.180533692 | 0.693483782 |
| Smarcal1 | -0.19471748 | 0.693483782 |
| Lsr | -0.244608506 | 0.693483782 |
| Ntn4 | -0.537766216 | 0.693483782 |
| Naaladl2 | -0.725493054 | 0.693483782 |
| Trim30c | -0.80018814 | 0.693483782 |
| Sco1 | 0.209364298 | 0.693588012 |
| Bcl7c | 0.145207629 | 0.693588012 |
| Pfkm | 0.166122738 | 0.693756432 |
| Abca3 | -0.175740221 | 0.693762192 |
| Mitf | -0.265296139 | 0.693957581 |
| Ptcd2 | 0.138115996 | 0.6940044 |
| Kcnh8 | 0.379883586 | 0.694308673 |
| Nsmf | 0.129977192 | 0.694308673 |
| Htr6 | -1.862510021 | 0.694308673 |
| Pla2g4e | 0.327712809 | 0.694515928 |
| Cxadr | -0.138951435 | 0.694980514 |
| Svil | -0.19695025 | 0.694980514 |
| Slc7a5 | -0.189955101 | 0.694980813 |
| Zbtb7c | -0.395274563 | 0.694980813 |
| Dbf4 | 0.135450088 | 0.694996454 |
| Crtc3 | 0.160186313 | 0.695361693 |
| Txndc9 | 0.10793333 | 0.695421665 |
| Mettl20 | 0.334220552 | 0.695659861 |
| Syt3 | 0.194058643 | 0.695659861 |
| 4930539E08Ril | -1.236368146 | 0.69580614 |
| Gm572 | 1.836638778 | 0.696178961 |
| Mrpl45 | -0.169078191 | 0.696348605 |
| Csrnp1 | 0.196596529 | 0.696411711 |
| Tbcc | 0.126394054 | 0.696505758 |
| Tmem221 | -0.835122702 | 0.696505758 |
| Fam189b | -0.279055679 | 0.696612962 |
| Ryr1 | -0.234649449 | 0.696704513 |
| Gm9970 | -1.869381585 | 0.696704513 |
| Tfb2m | 0.154864121 | 0.696817193 |
| Vps13b | -0.15243686 | 0.696833859 |
| Ifrd1 | -0.131794821 | 0.696842295 |
| Hes7 | 1.419741202 | 0.697018006 |
| Dcdc2a | 0.432129062 | 0.697098907 |
| Ier3 | 0.156570863 | 0.697169884 |
| Snapc5 | 0.135632317 | 0.697169884 |
| Lpar4 | -0.3688489 | 0.697169884 |
| Adgb | 1.170200229 | 0.697279627 |
| Pced1b | 0.612606005 | 0.697831109 |
| Cwc25 | 0.141910494 | 0.697889562 |
| Map3k14 | 0.462944837 | 0.698020483 |
| Cdkn2c | 0.328804929 | 0.698020483 |
| Trabd2b | 1.057379025 | 0.698101958 |
| Krt18 | 0.44286934 | 0.698101958 |
| Celf3 | 0.128592995 | 0.698101958 |
| C87436 | 0.190964518 | 0.69836465 |
| Snx22 | 0.469159196 | 0.698399022 |
| Lbx2 | 1.194293377 | 0.698506151 |
| Sft2d3 | -0.223367076 | 0.698576803 |
| Rap2b | 0.139402917 | 0.698587479 |
| Slc39a6 | 0.120899719 | 0.698587479 |
| Rtel1 | 0.199134185 | 0.698600863 |
| Fam227a | -0.304931917 | 0.698962909 |
| Letm2 | -0.270130155 | 0.699298764 |
| Trmt6 | 0.180540017 | 0.699368383 |
| Apln | 0.374158597 | 0.699378989 |
| Mog | -0.488853242 | 0.699443324 |

| | | |
|---|---|---|
| Dock2 | -2.091824622 | 5.33E-06 |
| Nfatc3 | -1.561304198 | 5.37E-06 |
| Csf2ra | -1.351838692 | 5.42E-06 |
| Mbnl1 | -1.495660528 | 5.42E-06 |
| Slc12a2 | 1.735741862 | 5.51E-06 |
| Spcs3 | -1.199704663 | 5.59E-06 |
| Saa3 | -2.940713045 | 5.59E-06 |
| Map3k11 | -1.756328586 | 5.71E-06 |
| Gda | -3.34133399 | 5.72E-06 |
| Ndnf | 7.029433869 | 5.78E-06 |
| Ccng2 | -0.94350901 | 5.81E-06 |
| Polr2i | 1.119534844 | 5.90E-06 |
| F5 | -5.734823247 | 6.06E-06 |
| Dnajc9 | 0.93756919 | 6.11E-06 |
| Oma1 | -1.250701855 | 6.11E-06 |
| Card9 | -1.986451633 | 6.17E-06 |
| Ltb | -2.06966483 | 6.18E-06 |
| Clec4a1 | -1.830026343 | 6.20E-06 |
| Mzt2 | 1.478970595 | 6.27E-06 |
| Rnf138rt1 | 7.36818327 | 6.27E-06 |
| Was | -1.556789004 | 6.36E-06 |
| Upf3a | 1.249240752 | 6.37E-06 |
| Immp1l | 1.106107109 | 6.47E-06 |
| Rfwd3 | -1.633170533 | 6.47E-06 |
| Mapk13 | -3.646297308 | 6.47E-06 |
| Haus7 | 1.123664276 | 6.50E-06 |
| Sipa1l2 | -1.329723293 | 6.55E-06 |
| Gm5150 | -2.633011221 | 6.55E-06 |
| Mmp14 | -1.467526061 | 6.56E-06 |
| Sh3kbp1 | -1.064709003 | 6.61E-06 |
| Rtn1 | 0.979587801 | 6.63E-06 |
| Gm10800 | 3.426959979 | 6.65E-06 |
| Spry2 | 1.641108714 | 6.70E-06 |
| Rragd | 1.193729598 | 6.74E-06 |
| Dock8 | -2.108676854 | 6.74E-06 |
| Naaa | -2.377103777 | 6.76E-06 |
| Slc43a2 | -1.203340982 | 6.86E-06 |
| Tmed3 | -1.363742486 | 6.97E-06 |
| Foxk1 | -1.492708127 | 7.00E-06 |
| Atp5o | 0.865291879 | 7.01E-06 |
| Wnt7b | -1.412391846 | 7.04E-06 |
| Nr4a2 | 2.091626375 | 7.14E-06 |
| Myo7a | -2.31832709 | 7.20E-06 |
| Prss46 | -5.486279521 | 7.29E-06 |
| Ip6k3 | -5.553699694 | 7.35E-06 |
| Snrpn | 0.791445077 | 7.42E-06 |
| Rac2 | -1.501740591 | 7.48E-06 |
| Thbs2 | 7.392851066 | 7.54E-06 |
| Serping1 | 1.855812622 | 7.55E-06 |
| Ccdc3 | 7.330039631 | 7.60E-06 |
| Atp6ap1 | -0.875485835 | 7.72E-06 |
| Rffl | -1.649155796 | 7.76E-06 |
| Gsap | -1.864136102 | 7.81E-06 |
| Mfsd7a | -8.074241609 | 7.83E-06 |
| Zc3hav1 | -1.943487606 | 7.90E-06 |
| C3 | -2.170621621 | 8.00E-06 |
| Anapc13 | 1.202051906 | 8.53E-06 |
| Sec31a | -1.031635096 | 8.67E-06 |
| Cetn3 | 0.87998839 | 8.74E-06 |
| Mocos | -7.026133183 | 8.79E-06 |
| 1500009L16Rik | 1.722356105 | 8.82E-06 |
| Chchd6 | 0.96254509 | 9.13E-06 |
| Slc25a4 | 0.799835555 | 9.13E-06 |
| Eva1c | 6.134245978 | 9.17E-06 |
| Coprs | 0.966874743 | 9.17E-06 |
| Pdcd1lg2 | -7.304926478 | 9.17E-06 |
| H2afx | 1.211107927 | 9.17E-06 |
| Psen1 | -1.251730732 | 9.17E-06 |
| Stxbp3 | -1.336037537 | 9.24E-06 |
| Ldha | -1.095725456 | 9.27E-06 |
| Eef1e1 | 0.986398105 | 9.57E-06 |
| Atg9a | -1.186195147 | 9.66E-06 |
| Cfdp1 | 0.748021174 | 9.70E-06 |
| Heyl | 7.309369831 | 9.78E-06 |
| Phgdh | 1.517936171 | 9.92E-06 |
| Bin2 | -1.73004989 | 9.97E-06 |
| Hnrnpl | -0.943438915 | 1.00E-05 |
| Rock2 | 1.381260111 | 1.02E-05 |
| Uqcrb | 0.846732721 | 1.02E-05 |
| Arid3a | -1.698936175 | 1.02E-05 |
| Helb | -1.70063508 | 1.02E-05 |
| Fbxl5 | -1.357487369 | 1.03E-05 |
| Hnrnpa0 | 1.231032777 | 1.03E-05 |
| Ttc3 | 0.802442479 | 1.03E-05 |
| S100a16 | 2.291375596 | 1.03E-05 |
| Numbl | -1.09446818 | 1.03E-05 |
| Arpc5 | -0.93023596 | 1.04E-05 |
| Unc93b1 | -1.549533951 | 1.04E-05 |
| Cd53 | -1.567122275 | 1.06E-05 |
| Pitpnm1 | -1.200343462 | 1.08E-05 |
| Kpna4 | -1.022724861 | 1.11E-05 |
| 9830107B12Rik | -8.167236533 | 1.12E-05 |
| Tshz2 | 2.117002982 | 1.13E-05 |
| Mrpl28 | 0.807327148 | 1.13E-05 |
| Kcng4 | -10.02447704 | 1.13E-05 |
| Snrpb2 | 0.953008387 | 1.14E-05 |
| Pdcd5 | 0.910368216 | 1.21E-05 |
| Cnpy2 | 1.003473527 | 1.21E-05 |
| Stat3 | -1.424854675 | 1.28E-05 |
| Cd3d | -8.395711683 | 1.28E-05 |
| Cd300lf | -2.143238948 | 1.28E-05 |
| Trim10 | 7.243847571 | 1.28E-05 |
| Mctp1 | -1.763540235 | 1.29E-05 |
| Zfyve1 | -1.097570115 | 1.30E-05 |
| Itgam | -1.676874821 | 1.31E-05 |
| Sh3bp2 | -1.162653165 | 1.32E-05 |
| Plpp6 | -1.99227249 | 1.32E-05 |
| Plcg2 | -1.858015774 | 1.32E-05 |
| Slc31a2 | -1.77981025 | 1.32E-05 |
| Alpl | 5.402047884 | 1.33E-05 |
| Socs3 | -1.263981621 | 1.33E-05 |
| Dmrt3 | 6.104596152 | 1.34E-05 |
| Kndc1 | 3.48434186 | 1.34E-05 |
| Plagl2 | -1.415546944 | 1.34E-05 |
| Prss12 | 7.768976883 | 1.36E-05 |
| Dgki | -1.466635953 | 1.37E-05 |
| Jund | -1.062573303 | 1.45E-05 |
| Hint1 | 0.775906026 | 1.45E-05 |
| Sh3bp4 | -1.228595495 | 1.45E-05 |

| | | |
|---|---|---|
| Morn4 | 0.463484242 | 0.049814972 |
| Sdf2 | 0.593793597 | 0.049833223 |
| Natd1 | 0.550262345 | 0.049833223 |
| Sqstm1 | -0.427212114 | 0.04983711 |
| Enpp2 | 0.669730319 | 0.049948502 |
| Arhgap33 | -0.598270528 | 0.05015543 |
| Gabarapl1 | 0.437420858 | 0.050189987 |
| Fam64a | 0.678253625 | 0.050254137 |
| Slc39a10 | 0.665455102 | 0.050289546 |
| Trim2 | 0.467141964 | 0.050378166 |
| Mss51 | 5.611074878 | 0.050467276 |
| F830016B08Rik | 5.672324793 | 0.050585833 |
| Htr2c | 2.66289825 | 0.050585833 |
| Zfand1 | 0.648890737 | 0.050585833 |
| Cdc16 | 0.410667992 | 0.050585833 |
| 4930407I10Rik | -5.786005544 | 0.050585833 |
| 1700011H14Rik | -5.014013064 | 0.050644936 |
| Kcnmb4 | 1.135951091 | 0.050657038 |
| Prdm2 | -0.485872959 | 0.050663744 |
| Asb14 | -3.120433445 | 0.050669961 |
| Slc24a1 | -4.966463913 | 0.050679108 |
| Bnip1 | 0.708059773 | 0.050685369 |
| Ncdn | -0.520539088 | 0.050715703 |
| Papd7 | 0.739604197 | 0.050730116 |
| Itga2b | 2.11489734 | 0.05090511 |
| Surf2 | 0.729230602 | 0.050921466 |
| Gid4 | 0.501185632 | 0.050957763 |
| Qrfpr | 5.223216269 | 0.05100042 |
| Palm3 | 5.136813908 | 0.051024686 |
| Dennd4c | -0.742553695 | 0.051024686 |
| Chmp4c | -2.368409692 | 0.051024686 |
| Col5a3 | 5.12085339 | 0.051055138 |
| Afap1l1 | 1.362406311 | 0.051055138 |
| Gprin3 | 1.574536035 | 0.051061585 |
| Trmt10c | 0.697684907 | 0.051061585 |
| Ppp1r15a | 0.549804313 | 0.051117533 |
| Aldh1a3 | 3.32429018 | 0.051150386 |
| Slmo2 | 0.526863994 | 0.051154133 |
| Clint1 | -0.575633555 | 0.051154133 |
| Dnmt3b | -1.397679922 | 0.05115657 |
| Gm21981 | -5.802120541 | 0.051165076 |
| Gm1661 | 5.223960499 | 0.051224522 |
| Akap6 | 0.554177035 | 0.051228873 |
| Lgr6 | 5.290228641 | 0.051297879 |
| Creg2 | -1.692711212 | 0.051329855 |
| Glce | 1.020368549 | 0.051357912 |
| Rps15a | 0.363928751 | 0.051378662 |
| Ctu1 | -0.765725453 | 0.051389437 |
| Zbtb44 | -0.639661068 | 0.051390299 |
| Npepl1 | -0.671665231 | 0.051537591 |
| Pax8 | -5.753405404 | 0.051537591 |
| Inpp5a | -1.126336776 | 0.051591877 |
| Tex10 | -0.629855329 | 0.051634222 |
| Naga | -0.724595423 | 0.051634222 |
| Hba-x | 1.748649133 | 0.051639556 |
| Ergic1 | 0.481613732 | 0.051641031 |
| Nedd4 | 0.555394275 | 0.051785556 |
| Psma3 | 0.403895519 | 0.051901995 |
| Lyve1 | 5.635999565 | 0.051930266 |
| Pi15 | 5.193482213 | 0.05199804 |
| Sct | 5.181034202 | 0.05199804 |
| Acadvl | -0.691649217 | 0.05199804 |
| Wnk4 | -2.032338649 | 0.05199804 |
| Gas7 | -0.539158419 | 0.051998621 |
| Abcc10 | -1.30037625 | 0.052063344 |
| D10Jhu81e | 0.592588266 | 0.052164622 |
| Dnaja1 | 0.35067909 | 0.052164622 |
| Rbm22 | 0.532138102 | 0.052186581 |
| BC107364 | -5.015115498 | 0.052223591 |
| Pcna-ps2 | 0.573276004 | 0.052243688 |
| Shb | 1.007244469 | 0.052320519 |
| Pmepa1 | 0.823479119 | 0.052320519 |
| F7 | -2.179133001 | 0.052320519 |
| Skor1 | 3.15727579 | 0.052368847 |
| Pcdhb3 | 1.820241094 | 0.052368847 |
| Tmem17 | 1.378814734 | 0.052403372 |
| Car12 | 1.260585524 | 0.052403372 |
| Rasgrf2 | 2.552046989 | 0.052411687 |
| Taf5l | -0.715014044 | 0.052441508 |
| Usp40 | 0.677497271 | 0.052477249 |
| AI314180 | -0.487576531 | 0.052477249 |
| BC022687 | 1.114056338 | 0.052533399 |
| Ppp2r2c | 0.790820474 | 0.052533399 |
| Dpp8 | -0.47742178 | 0.052533399 |
| Gpcpd1 | -0.484233804 | 0.052533399 |
| Wdr37 | -0.707529009 | 0.052533399 |
| Pcdhgb8 | -2.301538971 | 0.052533399 |
| Adgrf4 | -6.100408613 | 0.052567301 |
| Rnf146 | -0.428722556 | 0.05258196 |
| Rnf11 | -0.569509432 | 0.05258196 |
| Col6a2 | 1.350778703 | 0.052583669 |
| Dnah17 | -1.407614057 | 0.052618491 |
| Vdac3 | 0.37611652 | 0.052686115 |
| Gal | 2.61136601 | 0.052782166 |
| Rnft1 | -0.710802813 | 0.052782166 |
| Acvr2b | 0.930878131 | 0.052830973 |
| Cadps | 0.772572023 | 0.052850753 |
| Igfbp2 | 0.59434089 | 0.052851714 |
| Fahd2a | 0.805276068 | 0.052882912 |
| Wdr34 | 0.563511287 | 0.052932666 |
| Zfp949 | 1.058304475 | 0.0530342 |
| Slc1a5 | -1.024726884 | 0.053074029 |
| Eftud2 | -0.456145625 | 0.053119802 |
| Pcdh20 | 4.534121556 | 0.05316427 |
| Icosl | -1.526698955 | 0.053202149 |
| Rttn | -1.478828972 | 0.053204086 |
| Zc3h7b | -0.571400711 | 0.053207242 |
| Ogt | -0.424908691 | 0.053333686 |
| Acap1 | -2.992741954 | 0.053333686 |
| Rab44 | -2.772396192 | 0.053348663 |
| Mcu | 0.697037575 | 0.053401463 |
| Ampd3 | -0.922448328 | 0.053445851 |
| Pnma1 | 1.859364736 | 0.053469913 |
| Rab6b | 0.444083084 | 0.053469913 |
| Rac1 | -0.501070752 | 0.053473534 |
| Nol9 | -0.517367406 | 0.053476821 |
| Kif5b | 0.415972662 | 0.053513826 |
| Slc26a2 | -1.061957118 | 0.053513826 |
| Mtmr14 | -0.497734156 | 0.053520201 |

| | | |
|---|---|---|
| Epg5 | -0.563982121 | 0.302440347 |
| Crh | 3.8705278 | 0.30244254 |
| Capn3 | 0.971503423 | 0.30244254 |
| D11Wsu47e | -0.474207337 | 0.302472357 |
| Ap3d1 | -0.284197383 | 0.302785641 |
| Ppargc1a | 1.01161377 | 0.302882257 |
| Rasef | 4.258185678 | 0.303350148 |
| Myo3b | 4.19830935 | 0.303350148 |
| Vps29 | 0.232994161 | 0.303350148 |
| Gpam | -0.552228072 | 0.303663764 |
| Shpk | -2.270724267 | 0.303663764 |
| Tmem158 | 0.574352782 | 0.303690023 |
| Pgrmc2 | -0.443293576 | 0.303751448 |
| D630033O11Rik | 4.219387531 | 0.303836585 |
| Umps | -0.371547587 | 0.304284035 |
| Wdr35 | 0.418372205 | 0.304315762 |
| Yy1 | -0.332996232 | 0.304602181 |
| Gramd1c | 4.208845575 | 0.304631568 |
| Man1a | -0.509140952 | 0.304631568 |
| Met | -0.90439473 | 0.304631568 |
| Kctd14 | 3.870012135 | 0.304981782 |
| Plpp2 | 0.624953653 | 0.305278109 |
| Zfp426 | -0.429432501 | 0.305586131 |
| 1700020N01Rik | 3.933203551 | 0.305590355 |
| Zfp612 | 0.613969681 | 0.305590355 |
| Slc16a12 | 3.924581293 | 0.305687567 |
| Gm10322 | -4.726406654 | 0.306020213 |
| AW822073 | 3.896647111 | 0.306156755 |
| Pcnxl2 | -0.966280856 | 0.306156755 |
| 4930578G10Rik | 1.86847201 | 0.306235526 |
| Pir | 1.362893093 | 0.306235526 |
| Iqcb1 | 0.28605541 | 0.306235526 |
| Galk2 | 0.471450692 | 0.307017193 |
| Zdhhc13 | -0.420259613 | 0.307099756 |
| Cd69 | -0.597346205 | 0.307099756 |
| Camp | -1.603487207 | 0.307122496 |
| Ccdc125 | -1.258931059 | 0.307677936 |
| Zfp800 | 0.416013061 | 0.307994431 |
| Prom1 | 0.384718571 | 0.307994431 |
| Arrdc4 | -0.279873686 | 0.308093401 |
| Kpna1 | -0.246158657 | 0.308157098 |
| Aox1 | -4.728741345 | 0.308348695 |
| Sla2 | -4.728741345 | 0.308348695 |
| Ahcy | -0.450025345 | 0.308563047 |
| A830080D01Rik | 0.412527054 | 0.308879533 |
| Myo1h | 3.199785347 | 0.308963878 |
| Rasgrp2 | -0.494426077 | 0.309168616 |
| Gpatch2l | -0.480584666 | 0.309308665 |
| Rabgap1 | 0.280709429 | 0.309377217 |
| Ttl | -0.339848468 | 0.309499459 |
| Thsd4 | 0.671259951 | 0.309909759 |
| Prkar1b | 0.336626278 | 0.309959053 |
| Arhgef28 | 0.675448812 | 0.310003235 |
| Tgfbr3 | 1.10648173 | 0.310142534 |
| BC021614 | -3.497487918 | 0.310142534 |
| Slbp | -0.305475503 | 0.310405791 |
| Rpl8 | 0.191176713 | 0.31069588 |
| Nupr1l | 1.313931197 | 0.310920737 |
| Adprhl2 | 0.35326382 | 0.310920737 |
| Mtr | -0.486295191 | 0.311216111 |
| 2700049A03Rik | -0.374794441 | 0.311389364 |
| Mrpl41 | 0.250229162 | 0.311728903 |
| Grm5 | 0.584560319 | 0.311793759 |
| Ero1lb | 0.722452398 | 0.311999461 |
| Caprin2 | 0.717401436 | 0.312118843 |
| Tcaim | 1.197279704 | 0.312137056 |
| Ufd1l | 0.238007817 | 0.312178833 |
| Bag1 | -0.266990739 | 0.312178833 |
| Tepp | 1.57769218 | 0.312224095 |
| Nanos1 | -0.590161708 | 0.312260775 |
| Crygs | 3.974774212 | 0.312431422 |
| Wdr49 | 2.807883908 | 0.312431422 |
| Oxld1 | 0.618887458 | 0.312431422 |
| Srgap3 | 0.335828775 | 0.312431422 |
| Lrrc40 | -0.29475609 | 0.312431422 |
| Suv39h1 | -0.330028182 | 0.312431422 |
| Hist2h3b | -0.782569521 | 0.312431422 |
| Gm20696 | -1.111838418 | 0.312431422 |
| Adamts19 | 1.669525539 | 0.312575905 |
| Smim1 | 0.813089145 | 0.312575905 |
| Acin1 | 0.265263214 | 0.312575905 |
| Iqcc | -0.712119475 | 0.312575905 |
| Fnbp1l | 0.248088629 | 0.312736547 |
| Tango6 | 0.979144068 | 0.312906631 |
| Mrps10 | 0.312382822 | 0.312906631 |
| Ftl1 | -0.255594424 | 0.312906631 |
| Lta | 3.556405639 | 0.31291251 |
| Pak6 | 0.464278464 | 0.31291251 |
| Cyp51 | 0.30656628 | 0.31291251 |
| Pard6g | 0.293607587 | 0.31291251 |
| Fads6 | -2.96519078 | 0.313044688 |
| Pou2f2 | -0.455363937 | 0.313298335 |
| Dnajc30 | 0.434039654 | 0.313367006 |
| Aox4 | 1.823608399 | 0.313621625 |
| Gpr65 | -0.642242794 | 0.313621625 |
| Foxd1 | 3.819601643 | 0.313887164 |
| Zic3 | 0.794762403 | 0.313887164 |
| Sil1 | 0.507949103 | 0.313887164 |
| Wdr54 | 0.484738805 | 0.313887164 |
| Sez6l | 0.457924028 | 0.313887164 |
| Haus1 | 0.397725642 | 0.313887164 |
| Actg1 | -0.193845762 | 0.313887164 |
| Eya3 | -0.32654278 | 0.313887164 |
| Zmpste24 | -0.328412471 | 0.313887164 |
| Pofut1 | -0.354527654 | 0.313887164 |
| Mpp3 | -0.479817568 | 0.313887164 |
| Plxnd1 | -0.729793808 | 0.313887164 |
| Tnfrsf18 | -1.301175924 | 0.313887164 |
| Trcg1 | -1.644653268 | 0.313887164 |
| Rgmb | -0.420307258 | 0.313944081 |
| Pdzd4 | -0.323968632 | 0.314020218 |
| Rps6 | 0.207782332 | 0.314061017 |
| Asic2 | -0.814513576 | 0.314072305 |
| Baiap3 | 2.2804066 | 0.314314992 |
| Zfp961 | 0.633828191 | 0.314385793 |
| Polr3g | 0.76720996 | 0.314625708 |
| Mycbp | -0.393387117 | 0.314630871 |
| Slc7a11 | -0.56539224 | 0.314949446 |
| Bbs7 | 0.567198066 | 0.314962544 |

| | | |
|---|---|---|
| Slc25a35 | 1.394596174 | 0.699643703 |
| Clpx | 0.142838466 | 0.699665553 |
| Arhgef12 | 0.157448886 | 0.699722217 |
| Gm21994 | -1.030006383 | 0.699909188 |
| Galnt14 | 0.421875228 | 0.700662507 |
| Ahsa1 | -0.091747431 | 0.700662507 |
| Nostrin | -0.42309928 | 0.700662507 |
| Glod5 | -1.240259276 | 0.700662507 |
| Ccdc136 | 0.211219197 | 0.700714982 |
| Chd8 | -0.131408015 | 0.700806159 |
| Fam214a | -0.179961752 | 0.701041109 |
| Map4k5 | -0.200869146 | 0.701041109 |
| Arl13b | 0.256705405 | 0.701298406 |
| Fancd2 | -0.282030668 | 0.701298406 |
| 4931428F04Ril | 0.141864506 | 0.701462266 |
| Traf7 | -0.136508562 | 0.701634211 |
| Cdkn1a | 0.219212809 | 0.702033441 |
| Nfkbil1 | 0.218161565 | 0.702294411 |
| Rgs8 | 0.242930331 | 0.702495218 |
| Brk1 | 0.09664614 | 0.702495218 |
| Tinf2 | -0.230772713 | 0.70293285 |
| Adamts13 | -1.188920399 | 0.70361269 |
| Fam131b | -0.192333959 | 0.7038715 |
| 0610010K14Ri | 0.67676437 | 0.703918695 |
| Cacnb4 | 0.18939341 | 0.704043074 |
| Hcst | -0.37268376 | 0.704178988 |
| Cnnm1 | -0.45773868 | 0.704429503 |
| Ldoc1l | 0.163832935 | 0.704509157 |
| Arhgap42 | -0.201285455 | 0.704509157 |
| Gm5145 | 1.781526984 | 0.704825772 |
| Prdm8 | 0.281019654 | 0.704825772 |
| Prkd3 | 0.16385415 | 0.704825772 |
| Exosc8 | 0.125088147 | 0.704825772 |
| Gm10699 | -0.318425485 | 0.704825772 |
| Rnf34 | -0.129329921 | 0.704889863 |
| Prkce | 0.201303649 | 0.704946922 |
| Pus7 | -0.211895875 | 0.705130415 |
| Lsm14b | 0.142050226 | 0.705245951 |
| Phyhip | 0.463619402 | 0.705416562 |
| Fbxo2 | 0.618521617 | 0.705619199 |
| Gabpa | 0.161768829 | 0.705646529 |
| Vkorc1 | 0.615346954 | 0.705778686 |
| Ppm1g | -0.103194078 | 0.705778686 |
| Trmt5 | -0.203885727 | 0.705778686 |
| Kras | 0.104140243 | 0.705941712 |
| Rbms3 | -0.182210086 | 0.705941712 |
| Sult6b1 | -1.681478022 | 0.705941712 |
| Mab21l2 | -0.595383432 | 0.705960453 |
| Lysmd3 | -0.140350743 | 0.705981428 |
| Ube2a | 0.114942012 | 0.705983978 |
| Ccdc32 | 0.150423877 | 0.706157652 |
| Nefh | 1.174880949 | 0.706182041 |
| Dis3 | 0.153858976 | 0.706280552 |
| Appl2 | -0.196911982 | 0.706322702 |
| Ms4a4d | 1.58945319 | 0.706377985 |
| Scrt2 | -0.235830269 | 0.706475877 |
| Lrrc51 | 0.225303806 | 0.706784007 |
| Dnali1 | 0.915340864 | 0.706992194 |
| A330050F15Ri | 0.999654993 | 0.70706768 |
| Vps54 | -0.273358215 | 0.70737893 |
| Gm3636 | -0.617885915 | 0.70737893 |
| Tk2 | -0.153140959 | 0.707436075 |
| Gm10801 | 0.899629509 | 0.707453059 |
| Zfp867 | 0.281460596 | 0.707453059 |
| Nradd | 0.236676363 | 0.707453059 |
| Nsrp1 | 0.147564912 | 0.707460083 |
| Htr1a | -0.935738564 | 0.707576558 |
| Ece2 | -0.141766037 | 0.707822725 |
| Glt1d1 | -0.53247142 | 0.707947118 |
| 2810021J22Rik | 0.27909614 | 0.708666136 |
| 9030612E09Ril | -0.488629568 | 0.708873446 |
| Dgcr2 | 0.133267974 | 0.709088121 |
| Atxn10 | 0.083177332 | 0.709088121 |
| Npsr1 | 1.515881103 | 0.709196107 |
| Nlrp1b | -0.692894242 | 0.709196107 |
| Gm13150 | -1.964887721 | 0.709514708 |
| Gstt3 | -1.964887721 | 0.709514708 |
| Mtfmt | -0.284387392 | 0.709583711 |
| Tmtc3 | 0.153917059 | 0.709616451 |
| Wdr31 | 0.369987611 | 0.709621787 |
| Trim11 | -0.205599721 | 0.709755528 |
| Rad54b | -0.251177409 | 0.709755528 |
| Slc46a3 | 0.271321509 | 0.709763492 |
| Agk | -0.128966271 | 0.709763492 |
| Atoh8 | -0.851221615 | 0.709763492 |
| St7l | 0.128529739 | 0.709791481 |
| St14 | 0.924035282 | 0.710048382 |
| Fam135b | 0.340506165 | 0.710048382 |
| Vwa5a | 0.186120525 | 0.710048382 |
| Cdh24 | -0.277796312 | 0.710337497 |
| Gp5 | 1.264948369 | 0.71044611 |
| Dip2c | 0.162230248 | 0.710725813 |
| Pigv | 0.204685207 | 0.710872702 |
| Pcdha9 | -0.413057421 | 0.710969871 |
| Sacs | -0.168766575 | 0.711632323 |
| DXBay18 | 1.374907843 | 0.711765996 |
| Il15 | 0.554883854 | 0.711765996 |
| Gpr171 | 0.611191579 | 0.711936512 |
| Megf6 | 0.41836501 | 0.712030736 |
| Pdp2 | 0.169778581 | 0.71214403 |
| Wdr83 | 0.136650757 | 0.712146274 |
| Naaladl1 | 0.952448374 | 0.712191383 |
| Cd300e | 0.438288582 | 0.712542724 |
| Gm16181 | 1.288375008 | 0.712871856 |
| Rltpr | 0.256433448 | 0.713291288 |
| Mfsd2a | -0.522358039 | 0.71336597 |
| Fbn1 | 0.280858316 | 0.713499311 |
| Srsf10 | 0.087755693 | 0.71393997 |
| Rbm39 | -0.089390321 | 0.714219662 |
| Adamtsl5 | 0.963029107 | 0.714368496 |
| Wbscr22 | 0.136803857 | 0.714368496 |
| Sypl2 | 0.522828722 | 0.714840529 |
| Purg | 0.224676175 | 0.715087402 |
| Nudcd3 | 0.138415699 | 0.715087402 |
| Chd9 | 0.116492979 | 0.715087402 |
| Trip4 | -0.141647869 | 0.715087402 |
| Slc22a15 | -0.282928978 | 0.715087402 |
| Rnf144b | -0.425964743 | 0.715087402 |
| Adi1 | -0.149552323 | 0.715137215 |

| | | |
|---|---|---|
| Cdo1 | 1.077371483 | 1.48E-05 |
| Gatm | -1.306165352 | 1.48E-05 |
| Prdm13 | 6.70244909 | 1.48E-05 |
| Map1a | 1.04953423 | 1.48E-05 |
| Plpp3 | 1.803914253 | 1.49E-05 |
| Frem1 | 4.826771007 | 1.50E-05 |
| Cnpy3 | -1.128992846 | 1.50E-05 |
| Crabp1 | 2.337342989 | 1.52E-05 |
| Pcdh10 | 1.620975953 | 1.52E-05 |
| Svep1 | 7.295569089 | 1.52E-05 |
| 1810022K09Rik | 0.791329527 | 1.54E-05 |
| Clvs2 | 3.106723503 | 1.56E-05 |
| 5730508B09Rik | -2.180589627 | 1.57E-05 |
| Cd72 | -1.547246325 | 1.57E-05 |
| Pdpn | 2.103986042 | 1.57E-05 |
| Dzip1 | 1.338451649 | 1.57E-05 |
| Fgfbp1 | 6.89613899 | 1.58E-05 |
| Lrrc25 | -1.64765324 | 1.58E-05 |
| Adra2b | -7.714160846 | 1.59E-05 |
| Tspan6 | 0.927505766 | 1.61E-05 |
| Wdr11 | -1.122505338 | 1.61E-05 |
| Rhoj | 3.596179605 | 1.62E-05 |
| Rai2 | 2.393939032 | 1.65E-05 |
| Ms4a7 | -1.61511151 | 1.65E-05 |
| Cxcr2 | -3.128226656 | 1.65E-05 |
| Zkscan2 | 1.882597343 | 1.68E-05 |
| Ndst2 | -1.445983584 | 1.68E-05 |
| Rps27a | 0.906425362 | 1.70E-05 |
| Nrsn1 | 1.839156827 | 1.71E-05 |
| Rerg | 7.694403381 | 1.72E-05 |
| Kif5c | 0.855341661 | 1.73E-05 |
| Cox7b | 0.837403161 | 1.73E-05 |
| Tspan5 | -1.031265451 | 1.74E-05 |
| Rcan2 | 2.029184374 | 1.75E-05 |
| Ndufa5 | 0.925594015 | 1.76E-05 |
| Rfx4 | 2.275733632 | 1.76E-05 |
| Tbc1d10b | -1.397045054 | 1.78E-05 |
| Fam229b | 2.39994239 | 1.80E-05 |
| Lsm6 | 1.128954837 | 1.80E-05 |
| Ptrf | 3.414191241 | 1.83E-05 |
| Nop10 | 0.921489567 | 1.84E-05 |
| Gpr31b | -2.965927485 | 1.84E-05 |
| Slamf7 | -3.674167709 | 1.84E-05 |
| Minos1 | 0.966580361 | 1.85E-05 |
| D8Ertd738e | 0.922867872 | 1.87E-05 |
| Il17ra | -1.705916038 | 1.94E-05 |
| Per1 | -1.3570281 | 1.95E-05 |
| Nupr1 | -1.788469352 | 1.98E-05 |
| Rpl32 | 0.895148014 | 1.98E-05 |
| Ep300 | -1.095192128 | 2.00E-05 |
| Smcr8 | -1.353993457 | 2.00E-05 |
| Ccdc184 | 2.011103294 | 2.00E-05 |
| Kif13b | -1.671984237 | 2.00E-05 |
| Cox14 | 0.922290895 | 2.00E-05 |
| Rgs2 | 1.563892385 | 2.01E-05 |
| Fkbp9 | 2.935235168 | 2.02E-05 |
| Tmem184b | -1.25250085 | 2.02E-05 |
| Rpa3 | 1.048777225 | 2.04E-05 |
| Cox6b1 | 0.872904759 | 2.07E-05 |
| Cyb5r1 | -1.215996274 | 2.09E-05 |
| Ap3m1 | -0.888985376 | 2.10E-05 |
| Lepr | 7.474441338 | 2.10E-05 |
| Tmem107 | 1.308289406 | 2.11E-05 |
| Nfil3 | -1.016268767 | 2.12E-05 |
| Slc38a6 | -1.433606403 | 2.13E-05 |
| Vstm2a | 1.872146182 | 2.16E-05 |
| Tpcn1 | -1.346123195 | 2.18E-05 |
| Gm8730 | 0.890756581 | 2.19E-05 |
| Frmd4b | -1.002270581 | 2.19E-05 |
| 2-Sep | -0.798353414 | 2.20E-05 |
| Irak1bp1 | -0.931078455 | 2.23E-05 |
| Arf6 | -1.214899832 | 2.24E-05 |
| Lgals8 | -1.584326713 | 2.24E-05 |
| Edn1 | -4.889767863 | 2.24E-05 |
| Nek9 | -1.28201524 | 2.24E-05 |
| Ndufa7 | 0.842539095 | 2.24E-05 |
| Eef1g | 0.699059053 | 2.24E-05 |
| Map2k3 | -1.151460425 | 2.29E-05 |
| Vasp | -1.401455963 | 2.31E-05 |
| Eif4ebp2 | -1.112302373 | 2.31E-05 |
| Polr2f | 1.042173272 | 2.33E-05 |
| Tctex1d2 | 1.120566815 | 2.34E-05 |
| Mrpl34 | 0.880765361 | 2.34E-05 |
| Cfap36 | 0.974322863 | 2.35E-05 |
| Zeb1 | 1.123737036 | 2.39E-05 |
| Hspe1 | 0.761551095 | 2.39E-05 |
| Psen2 | -1.506599296 | 2.39E-05 |
| Elane | 9.268867325 | 2.40E-05 |
| Arhgap36 | 7.081974084 | 2.44E-05 |
| B630005N14Rik | -1.071122719 | 2.47E-05 |
| Alx3 | 7.198974 | 2.47E-05 |
| Csrnp3 | 0.793625893 | 2.50E-05 |
| Osm | -2.083196669 | 2.52E-05 |
| Ctsg | 6.397791928 | 2.55E-05 |
| Lsm4 | 0.907852948 | 2.55E-05 |
| Oas2 | -2.506702432 | 2.56E-05 |
| Cyth3 | -1.17991892 | 2.56E-05 |
| Cadm4 | -1.473345474 | 2.56E-05 |
| Susd6 | -1.306743404 | 2.62E-05 |
| Abca7 | -1.557968334 | 2.66E-05 |
| Satb2 | -1.337342596 | 2.68E-05 |
| Ppp2ca | -0.754718939 | 2.77E-05 |
| Basp1 | 0.814699206 | 2.78E-05 |
| Acox3 | -1.700512751 | 2.78E-05 |
| P2rx4 | -1.437367962 | 2.78E-05 |
| Slc44a5 | 2.275933529 | 2.95E-05 |
| Tpd52 | -1.416915969 | 2.96E-05 |
| Peak1 | -1.834579062 | 2.96E-05 |
| Erh | 0.756556864 | 2.97E-05 |
| Sec61g | 0.863002667 | 2.99E-05 |
| Zfp592 | -1.127338425 | 2.99E-05 |
| Tbc1d10c | -3.338121016 | 2.99E-05 |
| Adamtsl3 | -2.202209981 | 3.03E-05 |
| Olfml1 | 7.261060702 | 3.03E-05 |
| Plekhm1 | -2.087977133 | 3.03E-05 |
| Rps19 | 0.860150222 | 3.10E-05 |
| Mctp2 | -7.138059103 | 3.12E-05 |
| Rps6ka4 | -1.427162826 | 3.20E-05 |
| Cntnap2 | 1.301169171 | 3.23E-05 |

| | | |
|---|---|---|
| Olfr212 | 5.597947963 | 0.053537754 |
| Cgref1 | 1.224426094 | 0.053545865 |
| Syt1 | 0.61365474 | 0.05356365 |
| Rad54l | -0.90158639 | 0.053637485 |
| Nudt2 | 0.506902399 | 0.053662892 |
| Pdlim2 | -1.044659451 | 0.053681733 |
| Kifap3 | 0.475829882 | 0.053694695 |
| Adamts6 | 1.515287916 | 0.05375957 |
| Eif5 | 0.356123182 | 0.053904794 |
| Psmb9 | -0.863716324 | 0.053982712 |
| Fgf2 | 5.088423334 | 0.053994343 |
| N6amt2 | 0.639093466 | 0.0540316 |
| Abi3 | -1.481830921 | 0.0540316 |
| Itgb3 | -2.19036299 | 0.0540316 |
| Lcn2 | -2.535934448 | 0.0540316 |
| Il23a | -6.108496509 | 0.0540316 |
| Cchcr1 | -1.114652001 | 0.054082712 |
| Medag | -1.302058637 | 0.054144756 |
| Cdk9 | -0.6500856 | 0.054146466 |
| Slc16a6 | -0.900475268 | 0.054174436 |
| Homer2 | 1.363491992 | 0.054205296 |
| Wdr73 | 0.592681764 | 0.054209064 |
| Mamstr | 3.255096119 | 0.054362967 |
| F8a | 0.663915794 | 0.054398561 |
| Gm4070 | -2.542184921 | 0.054398561 |
| Dennd1c | -1.558248391 | 0.054427712 |
| Alg3 | 0.718484713 | 0.054451348 |
| Dnah11 | 5.073052594 | 0.05450261 |
| Zfp51 | -1.226807779 | 0.05450261 |
| Pop5 | 0.497551036 | 0.05452096 |
| Spats2l | 0.987284128 | 0.054574262 |
| Htr1b | 3.023058545 | 0.0545852 |
| Cnih2 | 0.405328106 | 0.054625589 |
| Tmem185b | -0.685075691 | 0.054625589 |
| Fut11 | 0.825814902 | 0.05463957 |
| Rhoh | -1.105187025 | 0.05463957 |
| Fsbp | 2.535983147 | 0.054751872 |
| Metap1d | -0.8022471 | 0.05475327 |
| Sfrp5 | 5.166010956 | 0.054873556 |
| Pde7a | 0.575079976 | 0.054902991 |
| Meaf6 | 0.446463881 | 0.054947238 |
| Pias1 | -0.62191414 | 0.055029106 |
| Adk | -0.624631568 | 0.055035751 |
| Hdgfrp3 | 0.484992009 | 0.055236403 |
| Erbb4 | -0.790659074 | 0.05530228 |
| Tbc1d22a | -0.746023424 | 0.055302949 |
| Nsf | -0.514034287 | 0.055319375 |
| Acy3 | 5.03329952 | 0.055392026 |
| Ahsa2 | 0.589433963 | 0.055433042 |
| Txn2 | 0.461239706 | 0.055456992 |
| Psma5 | 0.43364362 | 0.055475145 |
| Bbx | -0.500644574 | 0.055577648 |
| Mpzl3 | -2.782419625 | 0.055588533 |
| Cd79b | -1.136947376 | 0.055858588 |
| BC051019 | 5.139024578 | 0.055867596 |
| Pkia | 0.41674818 | 0.055927162 |
| Pls1 | 2.37534789 | 0.056002293 |
| Dll4 | -3.562384083 | 0.056029686 |
| Dnmt3a | 0.440850495 | 0.056056429 |
| Zfp706 | 0.36913649 | 0.056056429 |
| Mrps30 | 0.579500374 | 0.056152658 |
| Lrp1b | 1.471840719 | 0.056186486 |
| Fbln7 | 2.816636278 | 0.056214633 |
| Fam65b | -0.995704767 | 0.056329834 |
| Trappc1 | 0.520503806 | 0.056375809 |
| Arl5a | -0.486114634 | 0.056375809 |
| Rtn4rl2 | -1.275132737 | 0.056375809 |
| Csmd1 | 1.000412542 | 0.056410414 |
| Tbc1d23 | -0.55275777 | 0.056410414 |
| 0610009O20Rik | 0.593314714 | 0.056442302 |
| Pcolce2 | 2.160832027 | 0.05644596 |
| Tomm20 | 0.400563185 | 0.056466546 |
| Tchp | 0.779679626 | 0.056500675 |
| Fbrsl1 | -0.620743318 | 0.056653847 |
| Ccdc180 | -3.652982094 | 0.056659817 |
| Lap3 | -0.625971011 | 0.056678212 |
| Pnrc2 | 0.524409028 | 0.056714516 |
| Trap1 | -0.551738307 | 0.056714516 |
| Cab39l | -0.554046876 | 0.05673328 |
| Syvn1 | -0.675937173 | 0.056803765 |
| Atxn2 | 0.641462675 | 0.056842858 |
| Ly96 | -1.066400037 | 0.056907944 |
| Msh6 | -0.550373128 | 0.057096556 |
| Tmem234 | 0.44346313 | 0.057323077 |
| Serf2 | 0.39373912 | 0.057329571 |
| Hadh | 0.674081353 | 0.057401506 |
| Zdhhc15 | 2.362948553 | 0.057413684 |
| Lpar6 | -1.009632332 | 0.057666203 |
| Zgrf1 | -0.740338374 | 0.057699052 |
| Rrp12 | -0.784281256 | 0.057699052 |
| Fam185a | -1.027462381 | 0.057719973 |
| Trmu | 0.765212918 | 0.057724293 |
| Slc23a2 | -0.86398693 | 0.057724293 |
| Mpp1 | -0.810912879 | 0.057767822 |
| Nudt1 | 1.021246975 | 0.057786173 |
| Casp4 | -1.351275399 | 0.057786173 |
| 1700015E13Rik | -4.409139815 | 0.057839653 |
| Fbln2 | 2.178348877 | 0.057852508 |
| Tm2d3 | 0.586677007 | 0.057877201 |
| Gimap4 | 5.208410027 | 0.05789142 |
| 9130401M01Rik | 0.792756801 | 0.057896666 |
| Amer1 | 0.525021763 | 0.057898599 |
| Tmem119 | -1.002483482 | 0.057950011 |
| Prdx4 | 0.457750961 | 0.058061199 |
| Wdr93 | 5.085830083 | 0.058181136 |
| Oasl1 | -2.849388552 | 0.058247652 |
| Klf11 | -0.745921097 | 0.05838114 |
| Serpine1 | 3.830419795 | 0.058431744 |
| Ccni | -0.410909532 | 0.058459804 |
| Lhfpl3 | 1.463579941 | 0.058673713 |
| BC035044 | -1.337138729 | 0.058673713 |
| Cmah | 2.556590185 | 0.058706971 |
| Ispd | 1.245784926 | 0.058761337 |
| Lipg | -2.238062837 | 0.058761337 |
| Sec14l1 | -0.441093912 | 0.058766867 |
| Tmem94 | -0.645211362 | 0.058766867 |
| Mybpc3 | -2.5315498 | 0.058766867 |
| Tmem219 | -0.537599415 | 0.058786465 |
| Rubcn | -0.637206324 | 0.058786465 |

| | | |
|---|---|---|
| Fam222b | -0.362974511 | 0.315032434 |
| Ppm1d | -0.319111452 | 0.315070784 |
| Tenm2 | -0.425157001 | 0.315148832 |
| Hpgds | -0.337142764 | 0.315456531 |
| Ddit3 | 0.366605147 | 0.315527721 |
| Atxn7 | -0.504261574 | 0.315931258 |
| Fgd1 | -0.450368079 | 0.316014258 |
| U2surp | 0.245052337 | 0.316066644 |
| Gm14391 | 1.514320158 | 0.316341574 |
| Cntln | 0.461459717 | 0.316341574 |
| Fez2 | -0.259684153 | 0.316374927 |
| Hrh1 | -4.686625743 | 0.316374927 |
| Olfr652 | -4.686625743 | 0.316374927 |
| Wbp2 | -0.256339476 | 0.31649133 |
| Nmb | 1.088415628 | 0.316550224 |
| Prdx3 | -0.295047266 | 0.316986379 |
| Patz1 | -0.308990716 | 0.316986379 |
| Cnn1 | 3.250575736 | 0.317018635 |
| Armc6 | 0.489682117 | 0.317018635 |
| Chst9 | 3.910822237 | 0.317188049 |
| Mtrf1 | 0.573494679 | 0.317210491 |
| Zmynd12 | -2.392575538 | 0.317420749 |
| 9130008F23Rik | -4.69209702 | 0.317508631 |
| Il19 | -4.69209702 | 0.317508631 |
| Phf2 | 0.440892826 | 0.31754306 |
| Tnfrsf9 | -3.331050685 | 0.317775956 |
| Eif2s3x | -0.383267101 | 0.317847781 |
| Casq2 | 4.240190061 | 0.318038595 |
| Pla2g2c | 3.861658408 | 0.31838103 |
| Il3ra | -1.324398449 | 0.318398533 |
| Ms4a6b | -0.536790086 | 0.318546891 |
| Gm13157 | 0.758462132 | 0.318743639 |
| Ak2 | -0.254239181 | 0.318743639 |
| Mtmr11 | 0.595707647 | 0.318776786 |
| Ankrd33b | -1.476917761 | 0.318776786 |
| Wipf3 | 0.821762052 | 0.318870052 |
| Scd2 | 0.235068842 | 0.319158898 |
| Rpl10 | 0.211976669 | 0.319158898 |
| Tuba1c | -0.205961162 | 0.319158898 |
| Rab43 | -1.303882909 | 0.319158898 |
| Tph1 | 2.792000577 | 0.319275683 |
| Mogat2 | 4.145856057 | 0.319491846 |
| Creld1 | 0.373628114 | 0.319491846 |
| Kctd9 | 0.634279775 | 0.31963286 |
| Prps1 | 0.269080257 | 0.319643069 |
| Erc1 | 0.337444315 | 0.319974093 |
| Ddx50 | 0.284288172 | 0.319974093 |
| Mks1 | -0.670451873 | 0.319974093 |
| Mtmr12 | -0.677583074 | 0.319974093 |
| Dusp12 | 0.370922056 | 0.320052112 |
| Myot | 4.191590477 | 0.320179454 |
| Efcab10 | 1.089686914 | 0.320179454 |
| Slc38a1 | -0.272634965 | 0.320179454 |
| Mettl13 | 0.463954783 | 0.320563384 |
| Saxo1 | 4.225836784 | 0.320679029 |
| Rtn4rl1 | 0.674422373 | 0.320679029 |
| 1700016K19Rik | 1.2853593 | 0.320717592 |
| Casc4 | 0.339897912 | 0.321218325 |
| Thap1 | 0.437766326 | 0.321234561 |
| Sdccag3 | -0.229419195 | 0.321396092 |
| Rsg1 | 1.002559902 | 0.321567579 |
| Parp6 | 0.231545916 | 0.321570218 |
| Acoxl | -3.027336308 | 0.32183703 |
| Gorasp1 | 0.566923197 | 0.321917411 |
| Armcx1 | 0.425144987 | 0.321917411 |
| Pycr2 | -0.352533724 | 0.321957836 |
| 1700040L02Rik | 4.220160179 | 0.322159636 |
| Hspa8 | -0.198320053 | 0.322414341 |
| Tbc1d25 | -0.453709861 | 0.322414341 |
| Tagln | 1.575553119 | 0.322418968 |
| Htr1d | -3.910673359 | 0.32252229 |
| Phospho2 | 0.343111072 | 0.322533557 |
| Thrap3 | -0.202971817 | 0.323069501 |
| Fah | -0.540006823 | 0.323180641 |
| 9230109A22Rik | 3.852752615 | 0.323214844 |
| Eaf2 | 1.803124559 | 0.323214844 |
| Bbc3 | -0.366514257 | 0.323320674 |
| Furin | -0.455275276 | 0.323320674 |
| Ifi27 | -0.545784216 | 0.323320674 |
| Car7 | 1.919064462 | 0.323324258 |
| Smim24 | 1.244273996 | 0.323324258 |
| Git1 | 0.399876509 | 0.323324258 |
| Smim7 | 0.237911486 | 0.323324258 |
| Dennd6a | -0.372898623 | 0.323443115 |
| Cox11 | -0.481694256 | 0.32356814 |
| Slc35g3 | -2.60254456 | 0.323601941 |
| Dyx1c1 | 2.130399392 | 0.32361121 |
| Gnmt | -2.424459328 | 0.323668704 |
| BC034090 | -1.071485618 | 0.323718307 |
| Fzr1 | 0.283779766 | 0.324019539 |
| Cdca2 | 0.439781329 | 0.324093214 |
| Fam53b | -0.404563056 | 0.324168173 |
| Vps16 | -0.253120502 | 0.324270342 |
| Pomt1 | -0.425889205 | 0.324901326 |
| Foxj2 | -0.482570677 | 0.324901326 |
| Mpl | 4.214856438 | 0.32498064 |
| Clstn3 | 0.35077792 | 0.32498064 |
| Adam3 | -1.069617968 | 0.324986786 |
| Slc46a2 | -3.9152866 | 0.325304628 |
| Unc80 | -0.891735277 | 0.325519389 |
| Msx3 | 4.166495556 | 0.325597882 |
| Lrrc16b | -0.452216106 | 0.325597882 |
| Gorasp2 | 0.280234145 | 0.325672825 |
| Spic | -1.702893708 | 0.325672825 |
| Psma6 | 0.231492329 | 0.325811894 |
| Cdh10 | 0.630786357 | 0.325826154 |
| Vezf1 | 0.295588955 | 0.325827085 |
| Lmo2 | 0.36558844 | 0.325978616 |
| Bpnt1 | -0.516297579 | 0.325984828 |
| Ccpg1 | -0.500840408 | 0.326010689 |
| Rnase1 | 3.880475934 | 0.326108611 |
| Raph1 | -0.333699992 | 0.326162813 |
| Sox17 | 4.133863747 | 0.326463324 |
| Ctsw | 1.943383672 | 0.326570302 |
| Nthl1 | 0.667277986 | 0.3266208 |
| Bcar3 | 0.969790839 | 0.326680209 |
| Cpox | -0.626579547 | 0.326683703 |
| 4930447C04Rik | 3.799059411 | 0.32709957 |
| Rit1 | 0.308081911 | 0.32709957 |

| | | |
|---|---|---|
| 1-Sep | 0.328339224 | 0.715138669 |
| Pianp | 0.181539907 | 0.715138669 |
| Pde10a | 0.169758598 | 0.715163999 |
| Bnipl | 1.062789497 | 0.715386364 |
| Gm10767 | 0.430000148 | 0.715386364 |
| Dcun1d1 | 0.116874559 | 0.715386364 |
| Atp2b3 | -0.339071713 | 0.715386364 |
| Serpinb9 | -0.460508923 | 0.715386364 |
| 5530401A14Ri | -0.72211742 | 0.715439876 |
| Speg | 0.208677081 | 0.715505069 |
| Erlin2 | -0.140096357 | 0.715659314 |
| Ptchd1 | 0.512393304 | 0.715807266 |
| Zfp422 | 0.093666788 | 0.715807266 |
| Mtmr7 | -0.190103058 | 0.715807266 |
| Tbc1d31 | -0.228736317 | 0.715935289 |
| Lin52 | -0.170035935 | 0.71594785 |
| Fbxl21 | -0.46238244 | 0.71622219 |
| Chrnb4 | -0.541325307 | 0.716295711 |
| Pcdh18 | 0.225905687 | 0.717000769 |
| Olfr731 | 1.492868483 | 0.717133697 |
| Tmem80 | -0.139455099 | 0.717206612 |
| Cebpz | 0.134198933 | 0.717291187 |
| Trim56 | -0.441742016 | 0.717307909 |
| Ccdc61 | -0.182272063 | 0.717322723 |
| Gpr37 | -1.441783028 | 0.717511641 |
| 4933406P04Ril | -0.571069599 | 0.717702495 |
| Serinc3 | -0.159968053 | 0.717709337 |
| Tiprl | -0.092141442 | 0.717871653 |
| Nsl1 | -0.232960474 | 0.717871653 |
| Zcchc4 | 0.218841229 | 0.71803352 |
| Ppib | 0.112327634 | 0.718325622 |
| Slc30a3 | 0.926403937 | 0.718597324 |
| Zmiz2 | 0.100738143 | 0.718706187 |
| Fbxl7 | -0.440490997 | 0.718706187 |
| Sgtb | 0.14218777 | 0.718726352 |
| Sfxn3 | 0.125863276 | 0.718758122 |
| Rsbn1l | -0.163653697 | 0.71878628 |
| Olfr804 | 1.318632683 | 0.718970055 |
| Hist1h2af | 0.801898955 | 0.719014337 |
| Ccnk | -0.173257737 | 0.719014337 |
| Tiam1 | -0.187418102 | 0.719014337 |
| Ctif | -0.231233587 | 0.719014337 |
| Tex9 | 0.1904973 | 0.719163889 |
| Palld | 0.197705878 | 0.719201044 |
| Gm21992 | 0.929124736 | 0.719301146 |
| Glb1l2 | 0.526150302 | 0.719301146 |
| Lrrc28 | 0.147069219 | 0.719301146 |
| Casp2 | 0.130573204 | 0.719301146 |
| Atf4 | 0.121038429 | 0.719301146 |
| H13 | -0.10187497 | 0.719301146 |
| Smpd1 | -0.119404865 | 0.719301146 |
| Fpgs | -0.234340616 | 0.719301146 |
| Cops7a | -0.12921773 | 0.719343586 |
| Trp53bp2 | -0.163291281 | 0.719343586 |
| Slc25a43 | -0.823350838 | 0.719343586 |
| Armc2 | 0.384478977 | 0.719556615 |
| Fam63b | 0.197919807 | 0.719556615 |
| Habp4 | 0.176356711 | 0.719556615 |
| Snx12 | 0.102332815 | 0.719556615 |
| Tm9sf1 | -0.142034548 | 0.719556615 |
| Bcl7a | -0.16891365 | 0.719556615 |
| Trf | -0.511393563 | 0.719556615 |
| Akip1 | -0.149423259 | 0.719745793 |
| Znrd1as | 0.422524975 | 0.719894643 |
| Ryk | 0.191141931 | 0.719894643 |
| Srpk1 | -0.090631435 | 0.719894643 |
| Ckm | -0.619051781 | 0.719894643 |
| Setd4 | 0.269236389 | 0.719922329 |
| Pcdhb20 | 0.374590155 | 0.720338383 |
| Nubp2 | -0.102059435 | 0.720338383 |
| Lipo2 | 1.256641986 | 0.720533101 |
| Dennd5a | 0.111074297 | 0.720533101 |
| Hist1h2bf | 0.823112436 | 0.720687028 |
| Slc27a3 | 0.209096589 | 0.720687028 |
| Vps13c | 0.190990233 | 0.720687028 |
| Sfxn5 | 0.117811424 | 0.720687028 |
| Rab2a | -0.092136096 | 0.720687028 |
| Dhx30 | -0.092255033 | 0.720687028 |
| Hist2h3c2 | -0.196342141 | 0.720687028 |
| Rbl1 | 0.254816322 | 0.720942588 |
| Mepce | -0.238368961 | 0.720942588 |
| Itgb1bp1 | -0.132928432 | 0.720949452 |
| Wnt16 | 1.210926113 | 0.721119205 |
| Rnf170 | 0.235451546 | 0.721274207 |
| Pnp2 | 1.727942502 | 0.721717954 |
| Gm7120 | -1.387462205 | 0.721717954 |
| Rmdn3 | -0.192751082 | 0.721819847 |
| 4930402H24Ri | -0.114762446 | 0.721859448 |
| Pcdhb9 | -0.335766853 | 0.721859448 |
| G6b | -0.836567147 | 0.721859448 |
| Trim23 | 0.160106699 | 0.721925244 |
| Lbp | -0.491888481 | 0.721935009 |
| Prokr1 | 0.478621896 | 0.722261241 |
| Cdyl2 | -0.211993164 | 0.722414958 |
| Rnf207 | 1.893557046 | 0.722452966 |
| Smim10l2a | 0.235552055 | 0.722452966 |
| Dusp10 | 0.167699459 | 0.722452966 |
| Elp6 | 0.14946955 | 0.722452966 |
| Taf11 | 0.127565753 | 0.722452966 |
| Hps4 | -0.147317103 | 0.722452966 |
| Otx2 | -0.29599869 | 0.722452966 |
| Gm9992 | -1.008334647 | 0.722452966 |
| Scn5a | 0.794221078 | 0.722490566 |
| Nmnat2 | 0.130668501 | 0.722490566 |
| Gtpbp2 | -0.191428838 | 0.722490566 |
| Nxph2 | -0.520700865 | 0.722490566 |
| Hnrnpr | 0.105924854 | 0.722762589 |
| Tmem25 | 0.437828713 | 0.723178731 |
| Hsd17b12 | -0.094188099 | 0.723178731 |
| Unc79 | -0.160432634 | 0.723178731 |
| Slc6a9 | -0.57486675 | 0.723178731 |
| Myo5a | 0.115685935 | 0.723605247 |
| 3110021N24Ri | 0.995341293 | 0.723610266 |
| Acadm | 0.131937268 | 0.723863102 |
| Plekha8 | -0.192422326 | 0.723869099 |
| Gzf1 | -0.155757882 | 0.723898182 |
| 5-Sep | -0.157525419 | 0.723898182 |
| Osmr | -0.947222178 | 0.723898182 |
| Phb | 0.088215857 | 0.723958045 |

| | | |
|---|---|---|
| Zdhhc6 | -1.040637906 | 3.23E-05 |
| Slc35b3 | -1.880179535 | 3.23E-05 |
| Cnih1 | 0.86496776 | 3.23E-05 |
| Eno1 | -0.913283357 | 3.23E-05 |
| Stxbp6 | 1.613689791 | 3.24E-05 |
| Ntng2 | -1.366489846 | 3.30E-05 |
| Fam19a1 | 2.973657795 | 3.30E-05 |
| Hif3a | -1.930535995 | 3.30E-05 |
| Sik2 | -1.54038099 | 3.30E-05 |
| Olig3 | 7.482583391 | 3.30E-05 |
| Agfg1 | -0.949708017 | 3.33E-05 |
| Tcte2 | 2.49817969 | 3.33E-05 |
| Acsl1 | -1.552302106 | 3.34E-05 |
| Sephs2 | -0.996707652 | 3.34E-05 |
| Abcc1 | -1.528333045 | 3.34E-05 |
| Itga6 | -1.60536824 | 3.35E-05 |
| Raver1 | -1.10280408 | 3.37E-05 |
| Mmp8 | -4.94990531 | 3.37E-05 |
| Nlrp12 | -7.919817656 | 3.38E-05 |
| Opcml | -1.536821513 | 3.40E-05 |
| Cbln1 | 4.809021354 | 3.42E-05 |
| Cd248 | 2.787414278 | 3.45E-05 |
| Morc3 | -0.965121621 | 3.45E-05 |
| 6720489N17Rik | -1.653013352 | 3.45E-05 |
| Acot7 | 0.86202481 | 3.45E-05 |
| Hes1 | 2.151356582 | 3.46E-05 |
| Adgre1 | -1.706617193 | 3.46E-05 |
| Pign | -1.256470159 | 3.47E-05 |
| Hspa5 | -0.973676558 | 3.50E-05 |
| Ndp | 5.26238686 | 3.54E-05 |
| Sirpa | -1.410188785 | 3.54E-05 |
| Cytip | -1.922384149 | 3.54E-05 |
| Rps21 | 1.095421416 | 3.56E-05 |
| Kif6 | 6.96090843 | 3.57E-05 |
| Gas1 | 1.44596024 | 3.60E-05 |
| Tmbim6 | -0.91232596 | 3.60E-05 |
| Tmed1 | 1.243057694 | 3.66E-05 |
| Pkm | -0.90520511 | 3.67E-05 |
| Kpna6 | -1.229812965 | 3.68E-05 |
| Etv5 | -1.214276505 | 3.69E-05 |
| Stard3 | -0.971820985 | 3.73E-05 |
| Tor4a | -2.280990493 | 3.73E-05 |
| Nxph1 | 1.304031024 | 3.84E-05 |
| Qk | 0.813999234 | 3.86E-05 |
| Junb | -0.915174475 | 3.86E-05 |
| Galnt6 | -3.257517001 | 3.90E-05 |
| Col6a1 | 5.148913893 | 3.90E-05 |
| Unc13c | 3.286616326 | 3.93E-05 |
| Gng3 | 0.849451616 | 3.93E-05 |
| Cd82 | -1.283298303 | 3.96E-05 |
| Polr2h | 0.973358654 | 3.99E-05 |
| Uqcr10 | 0.947423591 | 4.00E-05 |
| Gprasp2 | 1.48434445 | 4.04E-05 |
| Dok1 | -1.480049762 | 4.05E-05 |
| Fam49b | -1.161114078 | 4.06E-05 |
| Actr3 | -1.162699644 | 4.06E-05 |
| Man2b1 | -1.225580872 | 4.06E-05 |
| Rab3b | 1.759921618 | 4.11E-05 |
| 1-Mar | -1.440393421 | 4.13E-05 |
| Car9 | -3.512892396 | 4.14E-05 |
| Nos2 | -3.45526741 | 4.19E-05 |
| Alyref | -1.107267514 | 4.21E-05 |
| Kcnj8 | 5.807618976 | 4.23E-05 |
| Fchsd1 | -1.879282226 | 4.24E-05 |
| Gmnc | -8.079327233 | 4.24E-05 |
| Tbc1d1 | -1.11563814 | 4.25E-05 |
| Pde1b | 1.12500807 | 4.26E-05 |
| Vwc2l | 6.529806638 | 4.27E-05 |
| Sox1 | 1.565399705 | 4.44E-05 |
| Atraid | 1.149644213 | 4.44E-05 |
| Angptl3 | -2.037305535 | 4.46E-05 |
| Magoh | 0.974478713 | 4.47E-05 |
| Nme5 | 2.431042279 | 4.48E-05 |
| Aldh3b1 | -1.986477214 | 4.48E-05 |
| Dhrs3 | -1.253994215 | 4.49E-05 |
| Aebp1 | 3.088087213 | 4.55E-05 |
| Grik1 | 4.039101188 | 4.59E-05 |
| Wdr72 | 5.716786543 | 4.59E-05 |
| Grb2 | -0.98464026 | 4.59E-05 |
| Acsbg1 | 2.418703174 | 4.61E-05 |
| Nap1l3 | 1.794403403 | 4.62E-05 |
| Prkcdbp | 1.848420462 | 4.67E-05 |
| Krt80 | -6.189717791 | 4.68E-05 |
| Tdo2 | -7.831603445 | 4.72E-05 |
| Map1s | -1.158035986 | 4.72E-05 |
| Mark2 | -1.089427747 | 4.79E-05 |
| Smc3 | 0.740244653 | 4.79E-05 |
| Tnip1 | -1.724809763 | 4.79E-05 |
| Slc10a4 | 2.906300938 | 4.82E-05 |
| Adcy7 | -1.889172802 | 4.83E-05 |
| Kcne1l | 1.574522173 | 4.84E-05 |
| Fli1 | -1.569997365 | 4.94E-05 |
| Tmem41a | 1.520209361 | 4.96E-05 |
| B4galt1 | -1.328162402 | 4.98E-05 |
| Ptpn14 | 7.145222043 | 4.99E-05 |
| Kmt2d | -0.795466389 | 5.03E-05 |
| Rpl36a | 0.735053526 | 5.07E-05 |
| Ctsl | -1.272216859 | 5.08E-05 |
| Gpr132 | -5.164879673 | 5.10E-05 |
| Scpep1 | -1.216451205 | 5.13E-05 |
| Med19 | 1.21039134 | 5.24E-05 |
| Map3k8 | -1.440681592 | 5.24E-05 |
| Stmn4 | 0.917505597 | 5.27E-05 |
| Pdzrn4 | 2.243365839 | 5.28E-05 |
| Ccl27b | 3.831256958 | 5.31E-05 |
| Cnr1 | -1.302992199 | 5.31E-05 |
| Clec2i | -5.760776835 | 5.32E-05 |
| Ptprc | -1.428919478 | 5.33E-05 |
| Rnh1 | -1.123515419 | 5.38E-05 |
| Ppa2 | 0.906578454 | 5.38E-05 |
| Cox7a2l | 0.918770377 | 5.41E-05 |
| Gm10282 | 0.886293298 | 5.44E-05 |
| Arpc2 | -0.876781665 | 5.45E-05 |
| Xpo6 | -0.957383856 | 5.45E-05 |
| Dusp1 | -0.831451605 | 5.47E-05 |
| Tiam2 | -1.336853301 | 5.51E-05 |
| Ano8 | -1.305742576 | 5.56E-05 |
| Ak4 | -1.972916868 | 5.59E-05 |
| Man2c1 | -1.007372977 | 5.81E-05 |

| | | |
|---|---|---|
| Adgrv1 | 1.060370316 | 0.058797198 |
| Oasl2 | -2.411059282 | 0.058797198 |
| Cmtm4 | -0.611237857 | 0.058849593 |
| Tc2n | 5.339177346 | 0.058925186 |
| Cyfip2 | -0.432778425 | 0.058938162 |
| Cmkl1 | 0.605906008 | 0.059162755 |
| Rbm8a | 0.46510103 | 0.059229365 |
| Wrb | 0.493849404 | 0.059268904 |
| Slc25a23 | -0.498541653 | 0.059268904 |
| Edem2 | -0.613501245 | 0.059268904 |
| Igtp | -1.985884937 | 0.059268904 |
| Tmem52 | 1.881646504 | 0.05936569 |
| Hist1h2ad | 1.421227129 | 0.059391302 |
| Map3k10 | -0.719135007 | 0.059391302 |
| Psrc1 | 1.085031881 | 0.059433451 |
| Cdv3 | -0.511361954 | 0.059447369 |
| Csnk1g3 | -0.531188872 | 0.059447369 |
| Als2 | -0.765631024 | 0.059447369 |
| Ano3 | 4.465271683 | 0.059519301 |
| Dctd | 1.031559908 | 0.059519301 |
| Elof1 | 0.497275071 | 0.059519301 |
| Gpr108 | -0.704346017 | 0.059519301 |
| Col4a5 | 1.651847862 | 0.059542062 |
| Adam9 | -0.609842394 | 0.059690077 |
| Wwc1 | 0.816102494 | 0.05971232 |
| E330009J07Rik | -0.659427398 | 0.05971232 |
| Ctbs | -0.983972032 | 0.05971232 |
| Kansl1l | -0.699140619 | 0.059742057 |
| Endov | 0.792736487 | 0.059780758 |
| Wasf1 | 0.598275326 | 0.059780758 |
| Glis3 | -2.203195471 | 0.059780758 |
| Tpi1 | -0.519121867 | 0.059814783 |
| Mettl5 | 0.62649486 | 0.059816177 |
| Ptges3 | 0.5357451 | 0.059816177 |
| Dlx5 | 0.757304245 | 0.059823629 |
| Vps8 | -0.660963016 | 0.05985591 |
| Usp18 | -1.876329082 | 0.05985591 |
| Ets1 | -1.401794145 | 0.059869781 |
| Slc25a40 | -1.074071352 | 0.059870434 |
| Actr1b | 0.490700191 | 0.059905467 |
| Tnfaip8l2 | -0.773812969 | 0.059924376 |
| Smim3 | -1.065805528 | 0.059991174 |
| Tmppe | -1.2300529 | 0.059991174 |
| Nrep | 0.413208396 | 0.060187273 |
| P2rx6 | 5.499014939 | 0.060443377 |
| Ranbp3l | 2.212523338 | 0.060443377 |
| Ginm1 | 0.692684419 | 0.060443377 |
| Pik3r3 | 0.486209032 | 0.060443377 |
| Ndufs8 | 0.41724633 | 0.060443377 |
| Ogg1 | 1.683224202 | 0.060541812 |
| Sntb2 | -0.873389877 | 0.060874338 |
| Pirt | 5.108356548 | 0.060964027 |
| Pctp | -1.755723356 | 0.060967741 |
| Pnkd | 0.519669074 | 0.060980661 |
| Sema4a | -2.018808971 | 0.060990972 |
| Chit1 | -5.995855657 | 0.060990972 |
| Fpr2 | -1.145129757 | 0.061079519 |
| Ergic3 | 0.415938419 | 0.061099416 |
| Nppc | 2.727489346 | 0.061147558 |
| Rhobtb1 | 1.336125111 | 0.061222076 |
| Ccl25 | 0.718300962 | 0.061222076 |
| Rab28 | 0.420496594 | 0.061222076 |
| Vnn1 | -5.27240215 | 0.061225053 |
| Kcne4 | 5.582481902 | 0.061243838 |
| Sepp1 | 0.57453882 | 0.061243838 |
| Acaa2 | 0.503850847 | 0.061338221 |
| Ccnh | 0.508947659 | 0.061411885 |
| Tsc1 | -0.573828119 | 0.061493131 |
| Tlr12 | -4.186489531 | 0.061494531 |
| Aplp2 | 0.480412838 | 0.061556124 |
| Dnah7c | -2.916578695 | 0.061558729 |
| D15Ertd621e | -0.680612055 | 0.061569034 |
| 1700024G13Rik | 5.034467174 | 0.061575818 |
| Nckap5l | -0.882418055 | 0.061679626 |
| Slc13a3 | -2.316118241 | 0.061679626 |
| Pde4d | -0.509585639 | 0.061826603 |
| Mrpl57 | 0.465402117 | 0.061874389 |
| Rora | 1.200839833 | 0.0618752 |
| Smim10l1 | 0.456703168 | 0.061963607 |
| Samsn1 | -1.833630417 | 0.061963607 |
| Klf13 | -0.484912984 | 0.062005536 |
| Greb1 | 3.102846442 | 0.062068305 |
| Tmem72 | 2.70986347 | 0.062102381 |
| Zbtb3 | 1.922609715 | 0.06210262 |
| Plec | -0.806076443 | 0.062159135 |
| Fam92b | 5.129323122 | 0.062189684 |
| Msr1 | -2.49913974 | 0.062195352 |
| Ppm1l | -0.644445928 | 0.062276571 |
| Gm8251 | -5.656073175 | 0.062276571 |
| Sh3bgrl | -0.404831631 | 0.062412055 |
| Slc25a18 | -1.212488049 | 0.062429238 |
| Ncs1 | 0.479714497 | 0.062453198 |
| P2rx5 | 5.058696008 | 0.062544693 |
| Rorc | 4.514937298 | 0.062544693 |
| Ift20 | 0.560623851 | 0.062544693 |
| Prrc2a | -0.523161455 | 0.062544693 |
| Erf | -0.784618552 | 0.062544693 |
| Gm17484 | -1.562582556 | 0.062544693 |
| Zap70 | 4.318664918 | 0.062557824 |
| Srsf12 | 1.014850791 | 0.062668942 |
| Tub | -0.711587426 | 0.062668942 |
| Abcf2 | -0.439495308 | 0.06267021 |
| Rbm17 | 0.389071725 | 0.062856903 |
| Tmem154 | -2.365707028 | 0.063021358 |
| Dnajc24 | 0.558097104 | 0.063028996 |
| Eps15l1 | -0.571925652 | 0.063033892 |
| Gimap6 | 2.807363171 | 0.063072821 |
| Zfp942 | -0.964753815 | 0.06310057 |
| Slc26a11 | 1.514145126 | 0.063102774 |
| Map7d2 | 0.768173243 | 0.063154318 |
| Nfia | 0.518106789 | 0.063154318 |
| Bscl2 | 0.557617148 | 0.063177928 |
| Hrk | 1.939697977 | 0.06318222 |
| Ffar2 | -4.554511753 | 0.063234323 |
| Sox18 | 2.840044824 | 0.063300433 |
| Cmtr2 | 1.538235574 | 0.063300635 |
| Chpf | -0.628946538 | 0.063341411 |
| Maml1 | -1.222527607 | 0.063341411 |
| Ngdn | 0.517320735 | 0.063421807 |

| | | |
|---|---|---|
| Zcchc11 | 0.304981656 | 0.32709957 |
| Baiap2l2 | -1.703867088 | 0.32709957 |
| Myo15 | -1.573136567 | 0.327203396 |
| Endod1 | 0.687353461 | 0.327938993 |
| Arhgap6 | -2.461490399 | 0.327938993 |
| Gm20654 | 3.772868968 | 0.328189519 |
| Cryab | 0.563846538 | 0.328189519 |
| Tyms | 0.335760924 | 0.328189519 |
| Xpo4 | -0.477813688 | 0.328189519 |
| Trnp1 | -0.959332419 | 0.328189519 |
| Pde2a | -0.484907222 | 0.32860165 |
| Depdc7 | -0.708479933 | 0.32860165 |
| Best1 | -0.844031973 | 0.32860165 |
| Cbr2 | -2.278463628 | 0.328814648 |
| Ccdc120 | 0.528887086 | 0.328831723 |
| Camsap3 | -0.394908184 | 0.329034513 |
| Treh | 3.843770551 | 0.329125262 |
| Kctd2 | -0.521548808 | 0.329190679 |
| Cep135 | 0.466853615 | 0.329246338 |
| Olfr689 | 3.849125965 | 0.329326385 |
| Ago3 | 0.355019499 | 0.329326385 |
| Neurl1a | -0.685007447 | 0.329374151 |
| Nop2 | -0.298953631 | 0.329417533 |
| Ubr7 | -0.448512799 | 0.329936582 |
| Lama5 | 0.90141232 | 0.330214874 |
| Mthfd2 | -0.29074879 | 0.33037292 |
| Tada1 | -0.325461475 | 0.330827146 |
| Plcl1 | 0.515734063 | 0.330964384 |
| Mamdc4 | -0.796114798 | 0.331058867 |
| Sin3a | -0.298981978 | 0.331152861 |
| Ap5s1 | -0.418848823 | 0.331152861 |
| Zfp106 | 0.252097173 | 0.33123125 |
| Dcaf6 | -0.443043213 | 0.33123125 |
| Gm8394 | 0.427653576 | 0.331252558 |
| Trpv6 | -1.381968686 | 0.331397617 |
| Morn5 | 1.494212281 | 0.331621438 |
| Tspan18 | -0.51321479 | 0.331785627 |
| Stard7 | -0.263560012 | 0.331883638 |
| Echdc3 | 4.092436619 | 0.331957303 |
| Rhod | 0.950403965 | 0.33214652 |
| H2-Q7 | 0.889555463 | 0.33214652 |
| Pkd2l2 | -0.924071401 | 0.33214652 |
| Eif3f | 0.218123837 | 0.332213822 |
| Coq6 | 0.34267845 | 0.332257203 |
| Zfp955b | 0.524501352 | 0.33242954 |
| Adgrd1 | 3.815753798 | 0.332471047 |
| Eid2b | 0.367343321 | 0.332471047 |
| Cpsf3 | -0.357359921 | 0.332716435 |
| Zfp644 | -0.22892151 | 0.333153289 |
| Nras | -0.201136248 | 0.333181891 |
| Cul7 | -0.335230403 | 0.333181891 |
| Ogfod2 | 0.330233128 | 0.333495611 |
| Gpank1 | -0.458830198 | 0.333591687 |
| Trmt2b | -0.691169882 | 0.33368198 |
| Mipep | 0.336376827 | 0.333881859 |
| Ssbp3 | -0.276786082 | 0.334069536 |
| Slc33a1 | -0.504712584 | 0.334080022 |
| Crtac1 | 0.596818459 | 0.334308543 |
| Fdx1l | 0.377901216 | 0.334384253 |
| Parg | -0.276843164 | 0.334462715 |
| Slc35f2 | -1.067251403 | 0.334462715 |
| Rps4x | 0.195963165 | 0.334765949 |
| Gm2178 | -3.232392897 | 0.334813177 |
| Fancm | 0.504109781 | 0.335168227 |
| Homez | 0.666374745 | 0.335263348 |
| Fyn | -0.236362545 | 0.335263348 |
| Enoph1 | -0.410420984 | 0.335325243 |
| Gm14322 | 1.196115186 | 0.335326571 |
| Phc3 | -0.276066763 | 0.335326571 |
| Slc25a15 | 0.778080392 | 0.335366204 |
| Bcl2l2 | 0.639196656 | 0.335667533 |
| Seh1l | -0.313440271 | 0.335667533 |
| Dnttip1 | -0.323133795 | 0.335667533 |
| Ankrd34a | 0.797008401 | 0.335703288 |
| Hcn3 | -0.332887926 | 0.335703288 |
| Arf4 | 0.199423734 | 0.335718242 |
| Gm8237 | 2.584009298 | 0.33574067 |
| Bub1 | -0.296953498 | 0.33574067 |
| Npffr1 | 4.112930979 | 0.335831624 |
| Pcmt1 | 0.220946225 | 0.336146998 |
| Cdc42se2 | -0.220468398 | 0.336146998 |
| Ogfr | -0.302499047 | 0.336146998 |
| Zfp68 | -0.323599166 | 0.336146998 |
| Svop | -0.344957643 | 0.336146998 |
| Vrk2 | -0.498725107 | 0.336146998 |
| Gm3550 | 0.603174989 | 0.336225885 |
| Casp8 | -0.978928411 | 0.336473804 |
| Epha1 | 2.111246392 | 0.336542511 |
| Rhbdd2 | -0.247627602 | 0.336573493 |
| Riok1 | -0.39226875 | 0.337367327 |
| Insc | 3.834692833 | 0.337395117 |
| Yes1 | 0.467614208 | 0.337395117 |
| Iqce | 0.392177133 | 0.337395117 |
| Kcmf1 | 0.2965745 | 0.337395117 |
| Stac2 | -0.791138362 | 0.337395117 |
| Igfbp1 | 4.159106367 | 0.337581789 |
| 8030423J24Rik | 4.067612951 | 0.337602281 |
| Gm4950 | 0.578533001 | 0.33768366 |
| Dmap1 | 0.311074561 | 0.33768366 |
| Arid4a | 0.297888762 | 0.33768366 |
| Smg9 | -0.3952188 | 0.33768366 |
| Plod3 | -0.443856425 | 0.33768366 |
| Scml2 | 0.942548127 | 0.33788282 |
| Teddm1b | -3.207677377 | 0.33788282 |
| Smco4 | 0.522389852 | 0.338051106 |
| Eif4a3 | -0.236594324 | 0.338094192 |
| Mettl21c | 2.657292176 | 0.338230801 |
| Tmem35 | 0.301412363 | 0.338286487 |
| Zfp354a | 0.915601993 | 0.338545772 |
| Abhd2 | 0.271559566 | 0.338754516 |
| Clptm1 | -0.340970615 | 0.338919039 |
| Bace2 | 3.873359122 | 0.338952308 |
| Msh2 | -0.316947916 | 0.338982896 |
| Myl12b | 0.226144539 | 0.339247659 |
| Dnajc27 | 0.675610685 | 0.339357204 |
| 5830454E08Rik | 1.56120953 | 0.339448101 |
| Man2a2 | -0.382867345 | 0.339448101 |
| Sf3b2 | 0.204725873 | 0.339613575 |
| Pkdcc | 0.909423872 | 0.33999985 |

| | | |
|---|---|---|
| Ticrr | -0.238363145 | 0.724099478 |
| Rtbdn | 1.869762087 | 0.724440345 |
| Gng8 | -0.434699967 | 0.72445382 |
| Ppfibp1 | -0.180546939 | 0.724500738 |
| Lrp8 | 0.110691468 | 0.72458915 |
| Xrcc6 | 0.130303628 | 0.724648477 |
| Sema4b | -0.139218729 | 0.724648477 |
| Commd9 | 0.129413303 | 0.724844438 |
| Lrrc42 | 0.124712629 | 0.725171193 |
| Ccl12 | -0.411100303 | 0.725745841 |
| Stra8 | 1.875877499 | 0.725825003 |
| Twistnb | 0.114552522 | 0.72583123 |
| Rab18 | 0.102226566 | 0.72583123 |
| Tnpo1 | -0.1008488 | 0.72583123 |
| Grid1 | 0.503621053 | 0.726092822 |
| Psmc2 | -0.088831089 | 0.726405599 |
| Rabepk | 0.182141174 | 0.72642688 |
| Rims4 | 0.30554639 | 0.726441588 |
| H2-T22 | 0.210559938 | 0.726441588 |
| Slc25a53 | 0.180076135 | 0.726486098 |
| Dio3 | -1.040176545 | 0.726564944 |
| Msrb3 | 0.622937914 | 0.726698531 |
| N4bp2l2 | 0.098545294 | 0.72672356 |
| Rprd1b | -0.118560174 | 0.72672356 |
| Gosr1 | -0.142612638 | 0.72672356 |
| Klhl2 | -0.195214863 | 0.72672356 |
| Ctxn3 | -0.812721098 | 0.72672356 |
| Anapc10 | 0.117118127 | 0.726966641 |
| Nell2 | -0.146103401 | 0.727179259 |
| Gm38393 | 0.19940133 | 0.727615127 |
| Fam172a | 0.108909859 | 0.727615127 |
| Tbc1d24 | -0.184856537 | 0.727615127 |
| Klhl36 | -0.192909629 | 0.727615127 |
| Kctd7 | -0.321978955 | 0.727831189 |
| B430203G13Ri | -1.206080054 | 0.727831189 |
| 2610042L04Ri | 0.42625247 | 0.727874838 |
| Gm14295 | -0.488545861 | 0.727891105 |
| Cbx8 | 0.222694928 | 0.727928729 |
| Mkx | -0.638563173 | 0.727928729 |
| Card6 | -0.573520331 | 0.728151466 |
| Kif27 | -0.626263691 | 0.728151466 |
| Mtum | -0.122980552 | 0.728173826 |
| Foxb1 | 0.90519033 | 0.72830319 |
| Pter | -0.276726058 | 0.728409408 |
| Exo5 | 0.136867048 | 0.728417494 |
| Sclt1 | 0.162124639 | 0.728514492 |
| Fam188b | -0.327537696 | 0.728514492 |
| Otub1 | -0.084639401 | 0.729033587 |
| Idh3b | 0.099238369 | 0.7290487 |
| Dnah9 | 0.646915034 | 0.729084364 |
| Tmem87a | -0.129981627 | 0.729362402 |
| Samd11 | 0.585792245 | 0.729563876 |
| Nrip2 | 0.4249153 | 0.729949787 |
| Tns3 | 0.163823716 | 0.730060156 |
| Espnl | 1.855231377 | 0.730627371 |
| Tmem138 | 0.159191329 | 0.730627371 |
| Gm10941 | 0.685508531 | 0.730777052 |
| Ndufb2 | 0.290408115 | 0.730777052 |
| Efs | 0.190036685 | 0.730777052 |
| Kmt2a | -0.124827381 | 0.730777052 |
| Armcx6 | 0.245400222 | 0.730833793 |
| Elk3 | -0.197646582 | 0.730833793 |
| Engase | -0.297462241 | 0.730833793 |
| Nexn | -0.43024902 | 0.730833793 |
| Neurod4 | 0.609509469 | 0.730856109 |
| Cecr2 | 0.181032952 | 0.730856109 |
| Gpr26 | 0.232893738 | 0.730879166 |
| Pcdhac2 | -0.339249351 | 0.730879166 |
| Pemt | 0.50214977 | 0.731195551 |
| Fam63a | 0.14822011 | 0.731195551 |
| Cd9 | -0.181739606 | 0.731195551 |
| Sra1 | 0.101424582 | 0.731208815 |
| Ephx4 | 0.27448993 | 0.731481487 |
| Sstr2 | 0.200690095 | 0.731481487 |
| Nsun3 | -0.190582046 | 0.731481487 |
| 1700067K01Ri | 1.039938372 | 0.731529435 |
| Fbxl16 | -0.159596099 | 0.731767822 |
| Eif4h | -0.079034673 | 0.731787213 |
| 4933416C03Ri | -1.137950172 | 0.731787213 |
| Mfrp | 1.478429435 | 0.732090015 |
| Obscn | 0.707960121 | 0.732293746 |
| Ptpn9 | -0.145119714 | 0.732293746 |
| Zwilch | 0.322254493 | 0.732349436 |
| Atp6ap1l | 0.625886392 | 0.732467896 |
| Msantd1 | 0.405979491 | 0.732467896 |
| Gm10184 | 0.293651686 | 0.732467896 |
| Arv1 | 0.220512842 | 0.732467896 |
| Tpra1 | 0.165537037 | 0.732467896 |
| Bysl | 0.14714899 | 0.732467896 |
| Psmf1 | 0.123318528 | 0.732467896 |
| Vamp4 | 0.111272041 | 0.732467896 |
| Dmxl1 | -0.130489225 | 0.732467896 |
| Gmppa | -0.137463183 | 0.732467896 |
| Pdk1 | -0.146949203 | 0.732467896 |
| Piga | -0.215574108 | 0.732467896 |
| Mgst2 | -0.738282774 | 0.7325131 |
| Kcnj14 | 0.741792901 | 0.732854401 |
| Ctdsp1 | -0.204830208 | 0.732854401 |
| Msl3 | 0.158698267 | 0.733056386 |
| Mroh2a | -0.355702618 | 0.733179677 |
| Ddx59 | 0.219843325 | 0.733397835 |
| Ncor1 | -0.085002708 | 0.733598836 |
| Arf5 | 0.092761673 | 0.733914164 |
| Yipf1 | 0.128482977 | 0.733980149 |
| 2810403A07Ri | 0.100058659 | 0.733980149 |
| Mgea5 | -0.095924212 | 0.734000051 |
| Tenm4 | -0.15707487 | 0.734000051 |
| Tesk2 | -0.586013002 | 0.73400869 |
| Tubgcp5 | -0.11828822 | 0.734370814 |
| Erich2 | -0.531213553 | 0.734375318 |
| Rogdi | 0.154507242 | 0.734457481 |
| Edil3 | -0.176434692 | 0.734457481 |
| Zdhhc14 | -0.20531107 | 0.734457481 |
| Slc37a1 | -0.295618415 | 0.73450828 |
| Ctnnbip1 | 0.111399992 | 0.734614264 |
| Klhl11 | -0.192910053 | 0.734790226 |
| Ankrd22 | -0.934706014 | 0.734885912 |
| Kcng2 | -0.678028429 | 0.734896258 |
| Cngb1 | -0.565419096 | 0.735032955 |

| | | |
|---|---|---|
| Lipe | -1.109582971 | 5.81E-05 |
| Fmnl3 | -1.719108943 | 5.81E-05 |
| Tmem63a | -2.062047004 | 5.81E-05 |
| Rab3c | 1.110680386 | 5.81E-05 |
| Mtmr6 | -0.956018679 | 5.82E-05 |
| Zbtb1 | -1.152328401 | 5.84E-05 |
| Cryba4 | 2.115812519 | 5.87E-05 |
| Tnfrsf21 | -0.96454812 | 5.87E-05 |
| Il6ra | -1.501568771 | 6.00E-05 |
| Pqlc3 | -1.750419116 | 6.00E-05 |
| Col26a1 | 3.574610071 | 6.01E-05 |
| Tnfrsf1a | -1.435488515 | 6.01E-05 |
| Slc39a11 | -1.181900213 | 6.13E-05 |
| Smc6 | 0.864373634 | 6.20E-05 |
| Psmg4 | 1.060467803 | 6.20E-05 |
| Mob3a | -1.048512803 | 6.22E-05 |
| Med23 | -1.251251119 | 6.24E-05 |
| Blvra | -0.995306163 | 6.26E-05 |
| Gna13 | -1.143379792 | 6.27E-05 |
| Cflar | -1.553412635 | 6.38E-05 |
| 2210013O21Rik | 0.982568757 | 6.41E-05 |
| Zfp871 | -0.836319151 | 6.41E-05 |
| Zfp518a | 1.9076515 | 6.49E-05 |
| Dnajc2 | 0.87777042 | 6.49E-05 |
| Higd2a | 0.845350291 | 6.50E-05 |
| Psma4 | 0.743156262 | 6.50E-05 |
| Frrs1 | -1.766175366 | 6.62E-05 |
| Psat1 | 1.224751727 | 6.71E-05 |
| Tgfbr1 | -0.852416165 | 6.71E-05 |
| Crlf3 | -1.410599103 | 6.71E-05 |
| Npy | 1.721995222 | 6.72E-05 |
| Pcdh19 | 1.29165485 | 6.72E-05 |
| Slc35f6 | -1.477564411 | 6.73E-05 |
| Wwtr1 | 1.432785883 | 6.74E-05 |
| Ehd1 | -1.26129177 | 6.74E-05 |
| H2-Ab1 | 7.321241098 | 6.77E-05 |
| Rpl18 | 0.772364248 | 6.79E-05 |
| Mrps6 | 1.222900834 | 6.79E-05 |
| Cass4 | -4.252235268 | 6.79E-05 |
| 1110008F13Rik | -0.971577919 | 6.83E-05 |
| Lrrc4c | 2.583182157 | 6.88E-05 |
| Rgs14 | -2.321886876 | 6.92E-05 |
| Cyp4f13 | -1.543908198 | 6.93E-05 |
| Ripply3 | 4.262249777 | 7.09E-05 |
| Ift27 | 1.206202109 | 7.13E-05 |
| Hivep2 | -1.313972501 | 7.13E-05 |
| Cd6 | -7.414842855 | 7.15E-05 |
| Pim2 | -1.316399883 | 7.16E-05 |
| Tmem147 | 0.791897243 | 7.20E-05 |
| Tfap2b | 4.961613739 | 7.22E-05 |
| Gm4353 | 2.085398072 | 7.23E-05 |
| Tmem261 | 0.941599361 | 7.25E-05 |
| Synj1 | -0.840060195 | 7.33E-05 |
| Gopc | -1.415552473 | 7.35E-05 |
| Lsm2 | 0.895270627 | 7.38E-05 |
| Tagap | -1.202166756 | 7.41E-05 |
| Dynll1 | 0.691318674 | 7.41E-05 |
| Map4k1 | -2.426832059 | 7.44E-05 |
| Cttnbp2 | -1.069720824 | 7.45E-05 |
| Esf1 | 1.15094329 | 7.46E-05 |
| Rcn1 | 1.174984689 | 7.47E-05 |
| Fmn1 | -1.869437429 | 7.50E-05 |
| Copz2 | 2.239715335 | 7.52E-05 |
| Rps14 | 0.705032182 | 7.54E-05 |
| Vcl | -1.370710102 | 7.57E-05 |
| Dock10 | -1.413666024 | 7.57E-05 |
| Nudt21 | 0.845919451 | 7.58E-05 |
| Srp14 | 0.709845419 | 7.58E-05 |
| Lsm7 | 0.967533716 | 7.62E-05 |
| Ly6a | 3.965027034 | 7.64E-05 |
| Bcan | 1.93522389 | 7.66E-05 |
| Sox9 | 1.364741521 | 7.69E-05 |
| Mrps11 | 1.159485448 | 7.69E-05 |
| Mrpl13 | 0.875756438 | 7.76E-05 |
| Gns | -1.150756457 | 7.76E-05 |
| Derl1 | -1.165350863 | 7.92E-05 |
| Adgra1 | 2.930388374 | 7.94E-05 |
| Slirp | 0.954620092 | 7.94E-05 |
| Dctn3 | 0.796082589 | 7.95E-05 |
| Pecam1 | -2.527600411 | 7.95E-05 |
| Rdx | 0.829502118 | 7.97E-05 |
| Cd34 | 2.32685241 | 8.06E-05 |
| Anxa5 | -1.357571165 | 8.10E-05 |
| Vps33a | -1.024781361 | 8.12E-05 |
| Higd1b | 4.557579603 | 8.14E-05 |
| Zc3h7a | -1.040286838 | 8.18E-05 |
| 8-Sep | -1.09486137 | 8.22E-05 |
| Kcnmb2 | 2.586839929 | 8.25E-05 |
| Ptma | 0.69029394 | 8.25E-05 |
| Cc2d1b | -1.340252407 | 8.26E-05 |
| Ccr1 | -2.318514967 | 8.28E-05 |
| Cntn6 | 6.057906431 | 8.48E-05 |
| Tmem14a | 1.817964113 | 8.48E-05 |
| Zc3h14 | -0.73571051 | 8.57E-05 |
| Laptm4b | 1.491743302 | 8.61E-05 |
| Gtf2h5 | 0.841866159 | 8.63E-05 |
| Slc25a27 | 0.993825783 | 8.66E-05 |
| Tpm3 | -0.777766849 | 8.76E-05 |
| Elmsan1 | -1.451186487 | 8.82E-05 |
| Lmbrd1 | -1.301801463 | 8.84E-05 |
| Rnf7 | 0.76813244 | 8.86E-05 |
| Mmel1 | -4.313120599 | 8.89E-05 |
| Ddx25 | 1.201094736 | 8.91E-05 |
| Chrna4 | 1.9802831 | 8.95E-05 |
| 2410004B18Rik | 0.97618552 | 8.97E-05 |
| Fam160b1 | -1.111146803 | 9.00E-05 |
| Adamts9 | 6.915216194 | 9.09E-05 |
| Kcne2 | 6.61017429 | 9.09E-05 |
| Arl3 | 0.822662912 | 9.29E-05 |
| Klhdc10 | -1.013303644 | 9.29E-05 |
| Pgap2 | -1.471226999 | 9.33E-05 |
| Pcm1 | 0.794518692 | 9.41E-05 |
| Gpx7 | 2.024624691 | 9.41E-05 |
| Fam118b | 1.032175231 | 9.45E-05 |
| Rps28 | 0.82685468 | 9.46E-05 |
| Chchd1 | 0.895323399 | 9.84E-05 |
| Tlr2 | -1.71127689 | 9.88E-05 |
| Ifnar1 | -0.883609642 | 9.91E-05 |
| Rit2 | 2.50418197 | 9.92E-05 |

| | | |
|---|---|---|
| Dda1 | 0.370632657 | 0.063421807 |
| Mab21l3 | -2.624446352 | 0.063421807 |
| Hspb8 | 2.255019502 | 0.063454686 |
| Ppme1 | -0.382373406 | 0.063454686 |
| F2rl1 | -5.979660418 | 0.063454686 |
| Ranbp2 | -0.41229047 | 0.063491551 |
| Pgf | 2.149939678 | 0.063534005 |
| Vcpkmt | 0.63005869 | 0.063567858 |
| Nek11 | 1.910395948 | 0.063608698 |
| Syt13 | 2.473726537 | 0.063662797 |
| Kctd1 | 1.353532777 | 0.063703058 |
| Zranb2 | 0.369734443 | 0.063724649 |
| Zfp280c | 0.673651644 | 0.063765481 |
| Cmtm7 | -0.710421472 | 0.063915091 |
| Usp46 | 0.647740083 | 0.063958528 |
| Crebzf | -0.749779064 | 0.06401257 |
| Tnfsf10 | -1.867695646 | 0.064071461 |
| Anp32b | 0.383807946 | 0.064077684 |
| Rbfa | -0.491825385 | 0.064398108 |
| Bola3 | 0.573763526 | 0.064521301 |
| Ube2m | -0.731316093 | 0.064645254 |
| Emcn | 5.113509393 | 0.064755272 |
| Zfp941 | 1.36816255 | 0.064855018 |
| Prss16 | -5.588411038 | 0.064881135 |
| Ly6g6f | -2.289818707 | 0.064933041 |
| Gfm2 | -0.727714407 | 0.064964177 |
| Ptprc | -1.372199342 | 0.064973443 |
| F13a1 | -1.973702227 | 0.064981504 |
| Cenpp | 0.971207692 | 0.065172083 |
| Psmb6 | 0.396534039 | 0.065206407 |
| Pla2g4c | 2.884577096 | 0.065207068 |
| Gm5784 | -1.555906009 | 0.06521267 |
| Paip2 | 0.322801093 | 0.06521769 |
| 4833427G06Rik | 4.916777925 | 0.06522882 |
| Dbil5 | -5.548551608 | 0.06522882 |
| Polr2a | -0.419541699 | 0.065261929 |
| Trib2 | 0.663292525 | 0.065322961 |
| Dolk | 0.632622124 | 0.065374799 |
| Mybbp1a | -0.514425737 | 0.065389439 |
| Nupl1 | -0.527155067 | 0.065389439 |
| Zbtb21 | -0.58035914 | 0.065389439 |
| Rgp1 | -0.83090392 | 0.065458832 |
| Lrrc17 | 5.126860263 | 0.065524389 |
| 1700088E04Rik | 2.200251594 | 0.065524389 |
| Pmf1 | 0.581886864 | 0.065524389 |
| Fiz1 | -0.547107149 | 0.065560714 |
| Gpd1 | 2.929059714 | 0.065628205 |
| Agtpbp1 | -0.582759755 | 0.065679361 |
| Camta1 | 0.460715628 | 0.065798772 |
| Arl5c | -1.035296831 | 0.065982363 |
| 6030458C11Rik | -0.754937549 | 0.066077366 |
| Sh3pxd2a | 0.764583077 | 0.066122984 |
| H2-Q10 | -4.260591596 | 0.06615247 |
| Ezh2 | 0.448609535 | 0.066238701 |
| Cybrd1 | 5.171410602 | 0.066246806 |
| Ywhab | -0.352407585 | 0.066260748 |
| Pla2g6 | 0.907938034 | 0.066260778 |
| Scnm1 | 0.525239378 | 0.066263441 |
| BC029214 | 0.570211662 | 0.066338043 |
| Cenpw | 0.766589523 | 0.066352709 |
| Hnf1a | -4.087741411 | 0.06637422 |
| Gm21954 | -5.536946621 | 0.066506845 |
| Map3k4 | -0.660406104 | 0.066609266 |
| Cbfa2t2 | -0.461385943 | 0.066627322 |
| Prdm4 | -0.626157979 | 0.0666886 |
| Kif2a | 0.562867348 | 0.066885988 |
| Emilin1 | -1.098794781 | 0.066933361 |
| Cxx1b | 0.808433207 | 0.06700905 |
| Creb3l2 | 1.317038764 | 0.06701245 |
| Jmjd7 | -2.96564222 | 0.067091491 |
| 2610524H06Rik | 0.460575608 | 0.067104374 |
| Clk3 | -0.405907856 | 0.067166838 |
| Myc | -0.579806434 | 0.067172244 |
| Syt4 | 0.619551073 | 0.067210905 |
| Traf3 | -0.574279468 | 0.067336391 |
| Ggcx | -0.862104423 | 0.067383965 |
| Serpinb8 | -1.748765222 | 0.067383965 |
| Pak1 | 0.531742504 | 0.067421225 |
| Gm10718 | 2.489360456 | 0.067499402 |
| Slpi | -2.845709578 | 0.067643323 |
| Tekt4 | 3.303208055 | 0.06769878 |
| Perm1 | -2.315977591 | 0.067732478 |
| Fbxl15 | 0.823411713 | 0.067738151 |
| Kdm2b | -0.577995703 | 0.067738151 |
| Ahnak | 0.889607067 | 0.067745154 |
| Snap29 | -0.543633576 | 0.067745154 |
| Trim7 | 5.044089363 | 0.067812015 |
| Tmem260 | -0.98823631 | 0.067812015 |
| Keap1 | -0.520814888 | 0.067837604 |
| Gjb4 | 5.120967111 | 0.067869151 |
| Rasgrf1 | -2.116679665 | 0.067940062 |
| Cpne3 | 0.531091291 | 0.067961095 |
| Dtymk | 0.497849252 | 0.068181827 |
| Ctss | -0.737904931 | 0.068222594 |
| Snx10 | -0.427299837 | 0.068263928 |
| 2010300C02Rik | 4.911693809 | 0.06827389 |
| Myo18a | -0.480931398 | 0.068327988 |
| Gpr61 | 4.766252166 | 0.068355568 |
| Unc13b | -0.598066912 | 0.068419053 |
| B3gnt5 | 0.704284332 | 0.068465098 |
| Fbln1 | 0.71812966 | 0.068468583 |
| Adamtsl4 | -1.421817853 | 0.0684687 |
| Lgals9 | -0.762499311 | 0.068484295 |
| Tcea2 | -0.763228194 | 0.068512654 |
| Nrip1 | -0.531920191 | 0.068546975 |
| Tgif2 | 0.670560715 | 0.068750556 |
| Bend4 | -1.178173997 | 0.06879084 |
| Atp8a1 | -0.603961668 | 0.068822483 |
| Prdm16 | 1.319663 | 0.06895214 |
| Ahi1 | 0.580507358 | 0.06895214 |
| Aph1a | -0.44254113 | 0.068965125 |
| Mbnl3 | -1.384587543 | 0.069047486 |
| Fam20a | 3.195412348 | 0.069073343 |
| Tmem50a | 0.397929151 | 0.069081037 |
| F2rl2 | 3.91161275 | 0.069139395 |
| Mff | 0.397488686 | 0.069246423 |
| Grm4 | 5.524575869 | 0.069274019 |
| Extl2 | 0.879032272 | 0.069350186 |
| Atp6v0e2 | 0.428450823 | 0.069406605 |

| | | |
|---|---|---|
| Sumf2 | -0.400937368 | 0.340060016 |
| Astn1 | 0.326757432 | 0.340212358 |
| Zbtb4 | 0.821607941 | 0.340229721 |
| Cyp39a1 | 0.458056938 | 0.340229721 |
| Nup50 | -0.273800455 | 0.340504339 |
| Uchl4 | 0.727738058 | 0.340626427 |
| Sdad1 | -0.343381131 | 0.340657676 |
| Fer1l5 | 1.142885447 | 0.341502771 |
| Lrwd1 | 0.349388905 | 0.341508344 |
| Lnx1 | 1.086885898 | 0.341576015 |
| Hist1h3c | 0.956969059 | 0.341576015 |
| Nxt2 | 0.473315533 | 0.341576015 |
| Usp16 | 0.269255096 | 0.341576015 |
| Cpxm1 | 0.807470111 | 0.341634137 |
| Tnfrsf12a | 0.717293392 | 0.341634137 |
| Elfn1 | 0.857343154 | 0.342001519 |
| 4933427G17Rik | 3.800853222 | 0.342013731 |
| Gp1ba | 2.13139442 | 0.342375073 |
| Smu1 | 0.237554717 | 0.342382019 |
| Il15ra | -1.134166593 | 0.342382019 |
| Vwa7 | -2.950191002 | 0.342457311 |
| Taf2 | 0.27840524 | 0.342532124 |
| Trnau1ap | 0.289946363 | 0.342547068 |
| Rps5 | 0.18957213 | 0.342612865 |
| Gng7 | 0.625603191 | 0.343039988 |
| Cep57 | 0.243252838 | 0.34356645 |
| Nphp1 | 0.605437876 | 0.343631523 |
| Cox16 | 0.591469762 | 0.343631523 |
| AU022252 | 0.326946468 | 0.343631523 |
| Pkhd1l1 | 3.710930539 | 0.34375406 |
| Tmem198b | 0.642221009 | 0.34375406 |
| Amz2 | 0.285307864 | 0.344013568 |
| Dlx4 | 4.140136531 | 0.344064791 |
| Ggnbp2 | -0.246889258 | 0.344130334 |
| Dnah8 | 0.797999419 | 0.34433139 |
| Rxrb | -0.31594117 | 0.34433139 |
| 1700026L06Rik | -0.598485101 | 0.34433139 |
| Fam214b | -0.363026828 | 0.344404223 |
| Ccdc174 | 0.43006484 | 0.345337498 |
| Rap1gap | 0.585585842 | 0.345420456 |
| Asah2 | 2.49335181 | 0.345542276 |
| Rassf6 | 2.558837707 | 0.345964246 |
| 1700049G17Rik | 0.686661314 | 0.345965899 |
| Dcst1 | -0.927179603 | 0.346116218 |
| Slc12a5 | 0.767074303 | 0.346179139 |
| Olfr330 | 4.061117891 | 0.346187214 |
| Slc22a18 | 4.4300326 | 0.346431543 |
| Unc45a | -0.281386502 | 0.346431543 |
| Calr3 | -1.497517099 | 0.346488882 |
| Trmt13 | -0.585662451 | 0.346498825 |
| Mlkl | -1.42047519 | 0.346580983 |
| Dpysl3 | 0.181400752 | 0.34675086 |
| 9-Sep | -0.244373694 | 0.34675086 |
| Nars | -0.221089046 | 0.346774739 |
| Spag9 | -0.218621835 | 0.346829103 |
| Chsy1 | 0.392096864 | 0.346864702 |
| Ccdc146 | -0.991474592 | 0.346864702 |
| Sap130 | -0.259267377 | 0.346936269 |
| Rec114 | -1.207117707 | 0.346936269 |
| Prph | 4.052561837 | 0.347029423 |
| Cdh4 | 0.430362864 | 0.347029423 |
| Nfrkb | -0.466188788 | 0.347517081 |
| Tmc2 | -4.290802324 | 0.347517081 |
| A1cf | 3.761688418 | 0.347545697 |
| Vat1l | 0.465402606 | 0.347648777 |
| Plekho1 | 0.3392459 | 0.347710192 |
| Gm16253 | -2.637199395 | 0.347739607 |
| Dpf3 | 0.535859687 | 0.347976378 |
| Chadl | 0.49959967 | 0.347976378 |
| Gstm4 | -0.516872211 | 0.347976378 |
| Rsph1 | 0.688021518 | 0.348197988 |
| Rnf112 | -0.679696675 | 0.348211973 |
| Gpkow | 0.355834277 | 0.348244682 |
| Mfsd7c | -1.852782582 | 0.348244682 |
| Cystm1 | 0.405747677 | 0.348410218 |
| Cars | -0.309446567 | 0.348481597 |
| Pms2 | -0.441018803 | 0.348481597 |
| Snx30 | -0.338909614 | 0.348505933 |
| 2010109A12Rik | 1.670173631 | 0.348694734 |
| Gorab | 0.507154282 | 0.348694734 |
| Ankrd53 | 3.748546183 | 0.348850608 |
| Spast | 0.359865646 | 0.348850608 |
| Ube3c | -0.291957117 | 0.348850608 |
| Nwd2 | -1.853946442 | 0.348850608 |
| Slfn14 | -4.576616186 | 0.348850608 |
| Foxc1 | 1.934483604 | 0.348930751 |
| Gm3893 | -3.212615061 | 0.349008336 |
| 6330408A02Rik | 0.424033691 | 0.349087789 |
| Cpne8 | -0.579193435 | 0.349185982 |
| Fam13b | -0.296119154 | 0.349466558 |
| Aox3 | 4.115687132 | 0.349555106 |
| Olfr1112 | 4.115687132 | 0.349555106 |
| Hinfp | -0.533628937 | 0.349619556 |
| B4galnt1 | -0.35204728 | 0.349745682 |
| Msl2 | -0.343602087 | 0.34986035 |
| 4930426L09Rik | -3.198459704 | 0.34986035 |
| Fras1 | 0.761672043 | 0.349939945 |
| Alg13 | -0.371835103 | 0.349939945 |
| Exosc5 | 0.31267791 | 0.350387199 |
| B3galt2 | 0.460757235 | 0.350526738 |
| Gm1527 | 3.771227763 | 0.350535845 |
| Lactb2 | -0.651453127 | 0.350535845 |
| Sec14l3 | 3.825466623 | 0.35056744 |
| Calu | 0.25683112 | 0.35056744 |
| Gucy2e | 4.054774148 | 0.350580969 |
| Nr2c2 | -0.324510578 | 0.350765982 |
| Alg5 | 0.316984704 | 0.351406013 |
| Pdzd8 | -0.308195658 | 0.351502187 |
| Sod3 | 3.353000768 | 0.351577045 |
| Shoc2 | -0.257139084 | 0.351669647 |
| Gm3739 | 0.491991341 | 0.351796316 |
| Chd3 | 0.359710568 | 0.352238674 |
| Slc35d3 | 3.351192781 | 0.352642487 |
| Ttf1 | 0.502047641 | 0.352642487 |
| Trim6 | -1.538030899 | 0.352748841 |
| Max | -0.243594775 | 0.352806972 |
| Epb41l4a | -0.414089109 | 0.35287851 |
| Tbrg1 | 0.337742659 | 0.353103087 |
| Kbtbd4 | -0.254864389 | 0.353103087 |

| | | |
|---|---|---|
| Drap1 | 0.094460124 | 0.735110659 |
| Nt5c2 | -0.111878686 | 0.735130233 |
| Cldn10 | 0.436765392 | 0.735131116 |
| 1700039E15Ril | 1.141470034 | 0.735232007 |
| Il18 | 0.177010745 | 0.735232007 |
| Mrpl24 | -0.148669661 | 0.735251346 |
| Bik | -0.563662013 | 0.735333197 |
| Ncapd3 | 0.12286479 | 0.735414535 |
| Exosc3 | 0.122589961 | 0.735414535 |
| Dnajc25 | 0.28197876 | 0.735436821 |
| Pitx2 | -1.220090342 | 0.735494376 |
| Prpf38b | -0.107802538 | 0.735904692 |
| N4bp2 | 0.123918969 | 0.73601447 |
| Begain | -0.225136096 | 0.73601447 |
| Polr1b | -0.187473794 | 0.736021467 |
| Zkscan4 | -0.43920381 | 0.736021467 |
| Egr4 | -1.167293212 | 0.73602604 |
| Ccdc173 | 0.258198704 | 0.736187008 |
| Cux1 | -0.108190976 | 0.736370949 |
| Lss | -0.14917626 | 0.736370949 |
| Fam228b | 0.495205242 | 0.736465048 |
| B3gat2 | 0.367658062 | 0.736787644 |
| Akr1a1 | -0.080341227 | 0.736787644 |
| Lman2l | -0.138344834 | 0.736787644 |
| Il11ra2 | 1.116945996 | 0.73685094 |
| Tulp1 | 0.645035825 | 0.737149579 |
| Mfsd9 | -0.482112166 | 0.737149579 |
| Bcl2 | -0.197960375 | 0.737427464 |
| Cdk5r2 | -0.299938693 | 0.737441805 |
| A830010M20R | -0.154321664 | 0.73768632 |
| Mina | -0.241048307 | 0.737860874 |
| Dnaja2 | -0.083169233 | 0.737996504 |
| Tmem88 | 0.380525892 | 0.738005224 |
| Faap20 | 0.117049545 | 0.738005224 |
| Primpol | 0.256892554 | 0.738198288 |
| Ccdc68 | -0.74063949 | 0.738198288 |
| Prpf19 | -0.108704062 | 0.738390357 |
| Doc2g | 0.875909859 | 0.738501403 |
| Pole4 | 0.115034912 | 0.73856311 |
| Svopl | 0.382881915 | 0.738566745 |
| Dffa | 0.17057834 | 0.73856994 |
| Gnaq | -0.089332689 | 0.73856994 |
| Ephx1 | -0.409316767 | 0.738588823 |
| Ehmt2 | 0.103101039 | 0.738598077 |
| Slx4ip | 0.186797024 | 0.738762096 |
| Rnf2 | 0.084187441 | 0.738762096 |
| Rdh12 | -1.025331569 | 0.738762096 |
| Mcm7 | -0.113668676 | 0.738946716 |
| Pacs1 | -0.11685422 | 0.738956161 |
| Zcchc14 | -0.110783808 | 0.739015659 |
| Notch4 | -0.319163803 | 0.739040521 |
| Ercc3 | -0.195471724 | 0.73943153 |
| Ptges3l | 0.558471092 | 0.739435423 |
| Gm10340 | 0.886897855 | 0.739492097 |
| Prrt1 | 0.417474345 | 0.739492097 |
| Ube2d2b | -0.844760919 | 0.739492097 |
| E2f3 | -0.184213271 | 0.739503371 |
| Rft1 | 0.189414167 | 0.7396018 |
| Kcnj11 | -0.50340054 | 0.7396018 |
| Atp6v1d | 0.093134432 | 0.739656479 |
| Abca13 | -0.828049431 | 0.739788923 |
| Abcb10 | -0.169717781 | 0.739902443 |
| Pigb | -0.129611746 | 0.739949803 |
| Spata21 | -1.303698824 | 0.740488211 |
| Fancc | -0.240356388 | 0.740673874 |
| Zcchc10 | -0.174281973 | 0.740677621 |
| Irgc1 | -1.382813953 | 0.740677621 |
| Commd8 | 0.10585482 | 0.741042443 |
| Lrif1 | -0.222800035 | 0.741231859 |
| Ccdc38 | -0.436549948 | 0.74123349 |
| Peli1 | 0.097072772 | 0.741497995 |
| Nup210 | -0.140278165 | 0.741497995 |
| Pkp1 | 1.807048594 | 0.741551473 |
| Prss35 | 0.712433292 | 0.741551473 |
| H2-Q4 | 0.417182751 | 0.741551473 |
| Thrb | 0.307428477 | 0.741551473 |
| Vwa8 | 0.173163782 | 0.741551473 |
| Miip | 0.157346694 | 0.741551473 |
| Dnajc11 | -0.105116929 | 0.741551473 |
| Nomo1 | -0.149664619 | 0.741551473 |
| Enthd2 | -0.182996994 | 0.741551473 |
| Hspb6 | 0.405916164 | 0.741698849 |
| Kbtbd6 | -0.707389353 | 0.741698849 |
| Pcdha2 | -0.414786276 | 0.741755463 |
| Tardbp | 0.080304355 | 0.741776826 |
| Rasgef1c | -0.290346543 | 0.741846477 |
| Hoga1 | -0.721403431 | 0.741846477 |
| Slc3a1 | -1.222757812 | 0.742031805 |
| Catsperg1 | 1.807044775 | 0.742936064 |
| Bcorl1 | -0.18968147 | 0.743076955 |
| Gm11744 | -0.689665853 | 0.743076955 |
| Ifi205 | -0.601176069 | 0.74318022 |
| Hist1h4b | 1.23699497 | 0.743306344 |
| Pex13 | -0.105606697 | 0.743306344 |
| Ago4 | -0.165727091 | 0.743390295 |
| Fech | 0.138090251 | 0.743593002 |
| Atf7ip | -0.097725011 | 0.743593002 |
| Aebp2 | -0.185055966 | 0.743593002 |
| Tcf19 | -0.187971624 | 0.743593002 |
| Rpap2 | 0.187349605 | 0.743715847 |
| Cnot6 | 0.093135383 | 0.743715847 |
| Neurog1 | 0.680959253 | 0.743984581 |
| Cfap53 | 1.989770222 | 0.74433707 |
| Wrn | 0.21439908 | 0.744720386 |
| Flt3l | -0.515758499 | 0.744813624 |
| Oxct2b | -1.066197338 | 0.744840355 |
| Zfp212 | -0.135307614 | 0.745008512 |
| Ephb3 | -0.254641072 | 0.74508455 |
| Tubgcp3 | -0.122028503 | 0.745130683 |
| Ms4a4c | -0.45185973 | 0.745224157 |
| Tspan9 | 0.182902944 | 0.74543713 |
| Hilpda | 0.134192882 | 0.74543713 |
| Neurl4 | -0.203492025 | 0.74543713 |
| Snta1 | -0.214282308 | 0.74543713 |
| Crocc2 | -1.554307255 | 0.74543713 |
| Pycrl | 0.101551296 | 0.745446086 |
| Glt8d1 | 0.108925351 | 0.745489553 |
| Usp3 | -0.085841236 | 0.745518199 |
| Efcab14 | -0.099640548 | 0.745518199 |

| | | |
|---|---|---|
| Ptprg | 1.59552361 | 0.000100312 |
| Rpl13 | 0.757615487 | 0.000100737 |
| Tmem135 | -1.023218822 | 0.000100963 |
| Uhrf1 | -0.999618522 | 0.000103606 |
| Osr1 | 5.922896562 | 0.000104169 |
| Rps15 | 0.848848644 | 0.000104193 |
| Ccnd3 | -0.927280287 | 0.000104964 |
| Wdfy2 | -1.3369364 | 0.000105064 |
| Gpi1 | -0.950659363 | 0.000105138 |
| Tatdn2 | -1.609304952 | 0.000105191 |
| Ogfrl1 | -1.2494591 | 0.000105626 |
| Rpl41 | 0.671600316 | 0.000105727 |
| Med17 | -1.034404302 | 0.000106 |
| A630033H20Rik | -2.484721821 | 0.000106512 |
| Taf13 | 1.306939345 | 0.000108368 |
| Rcor2 | -1.006317756 | 0.000108368 |
| Lsm1 | 0.841372667 | 0.00010863 |
| Pde4b | -1.35641407 | 0.00010863 |
| Rab29 | -1.089605592 | 0.000109114 |
| Celf4 | 0.807161631 | 0.000109362 |
| Trim16 | 6.926626203 | 0.000110129 |
| Tomm7 | 0.753502527 | 0.000110406 |
| A130051J06Rik | -1.590253683 | 0.000110406 |
| Ntsr1 | -2.454631388 | 0.000110406 |
| Ccl27a | 1.042238801 | 0.000111345 |
| Mast4 | 2.310524217 | 0.000112773 |
| Rbm46 | 6.741538676 | 0.000113474 |
| Evc | 2.869977907 | 0.00011383 |
| Cast | -1.720552545 | 0.000114691 |
| Adam10 | -0.891123051 | 0.000114778 |
| Gpx1 | 1.190871927 | 0.000115999 |
| Syngr3 | 1.029373287 | 0.000117455 |
| Bbs5 | -1.720129941 | 0.00011822 |
| Cd177 | -7.358780514 | 0.000118322 |
| Tcirg1 | -1.867613989 | 0.000118553 |
| Tceal6 | 1.753660862 | 0.000120427 |
| Fbxo45 | 1.702662045 | 0.000120427 |
| Ceacam10 | -7.064499357 | 0.000120515 |
| Atp5h | 0.748318644 | 0.000121792 |
| B3gnt3 | -7.20844217 | 0.000121792 |
| Polr2e | 0.71522816 | 0.000122425 |
| Stk11ip | -1.245529205 | 0.000122912 |
| Adgrl2 | -1.053800724 | 0.000123148 |
| Ndufs5 | 0.713776497 | 0.000125206 |
| Tmem50b | -0.970768897 | 0.000125206 |
| Dtna | 2.159314418 | 0.000125824 |
| Nedd8 | 0.701605713 | 0.000125824 |
| Calb1 | 2.00958304 | 0.00012762 |
| Cyba | -1.317320453 | 0.00012772 |
| Cbx1 | 0.74686292 | 0.00012914 |
| Col4a6 | 6.713503865 | 0.000129667 |
| Schip1 | 1.094546438 | 0.00013041 |
| Cyp11a1 | -5.069472987 | 0.000130454 |
| Hcn1 | -3.240432628 | 0.000130774 |
| Trem3 | -2.88086925 | 0.000131471 |
| Mcmbp | -0.753961723 | 0.000131936 |
| Ddr2 | 2.741546228 | 0.000132238 |
| mt-Nd3 | 1.462710741 | 0.000132807 |
| Sod1 | 0.91886797 | 0.000133125 |
| Rap1gds1 | -0.966181662 | 0.000133187 |
| Gm6377 | -2.172168239 | 0.000133727 |
| Rab31 | -0.834251056 | 0.000134962 |
| Crhr1 | 3.85513367 | 0.000135013 |
| Ifnar2 | -0.995351613 | 0.000135131 |
| Spag6l | 2.75663329 | 0.000135511 |
| Irak2 | -2.074140837 | 0.000135589 |
| Pbx4 | 1.452324178 | 0.000135947 |
| Lyst | -1.760532203 | 0.000137538 |
| Lhpp | 1.372668216 | 0.000137859 |
| Anapc11 | 0.928982939 | 0.000138253 |
| Top1 | 0.771746743 | 0.00013932 |
| Neurl3 | -1.900957065 | 0.00013932 |
| Mlf1 | 1.382091378 | 0.000139427 |
| Sdsl | 6.975208412 | 0.000139531 |
| Pstpip1 | -2.327236766 | 0.000140234 |
| Cdc26 | 0.791403395 | 0.000140405 |
| 1500011B03Rik | 0.828235133 | 0.000140521 |
| Usp13 | 2.628494192 | 0.000140855 |
| Csf2rb2 | -2.143289753 | 0.000142561 |
| Mettl9 | 0.73805617 | 0.000142699 |
| Xpnpep2 | 7.001992737 | 0.00014292 |
| Rps27rt | 1.112225434 | 0.000143634 |
| Osgin1 | -2.870909818 | 0.000144926 |
| Serpinh1 | 1.88340716 | 0.000145359 |
| Olig1 | 2.592287143 | 0.000145552 |
| Tdrd7 | -1.574554261 | 0.000145777 |
| Plek2 | 6.221576128 | 0.000145809 |
| Pxmp2 | 3.795729648 | 0.00014747 |
| Pira2 | -4.534541961 | 0.00014747 |
| Ssh1 | -1.047302021 | 0.000148949 |
| H2afz | 0.692926636 | 0.000149486 |
| Ajuba | 2.097246 | 0.000150363 |
| Dusp5 | -1.752332806 | 0.000150363 |
| 2810428I15Rik | 0.833795522 | 0.000151397 |
| Rbm47 | -1.375291974 | 0.000151404 |
| Fam181b | 2.291863414 | 0.000151788 |
| Ggta1 | -1.698775288 | 0.000151971 |
| Nts | 3.791120406 | 0.000153495 |
| Slfn3 | -4.541808207 | 0.000153495 |
| Nbeal2 | -3.285062553 | 0.000154828 |
| Dapp1 | -1.757110416 | 0.000156111 |
| Ppp1r14c | 2.000195416 | 0.000156237 |
| Pnp | -1.107041139 | 0.000157433 |
| Shisa4 | 1.114115849 | 0.000158666 |
| Mxd1 | -1.890980813 | 0.000159021 |
| Rbm20 | 3.83496345 | 0.000159263 |
| Tceal8 | 0.874507197 | 0.000159263 |
| Cherp | -1.067944026 | 0.000159263 |
| Ccdc80 | 2.394240834 | 0.000160156 |
| Arhgap21 | 0.86154658 | 0.000160156 |
| Afap1l2 | 6.763126603 | 0.000162379 |
| Pdap1 | 0.80742958 | 0.000162438 |
| Krtcap2 | 0.787543047 | 0.000162719 |
| Rpl5 | 0.666646113 | 0.000162719 |
| Rps10 | 0.737322098 | 0.000162759 |
| Sdcbp | -0.897030614 | 0.000162759 |
| Pip4k2a | -1.382469217 | 0.000163237 |
| Zbbx | 6.683778389 | 0.000163271 |
| Ppp1r12c | -1.076535987 | 0.000163936 |

| | | |
|---|---|---|
| Ptpn7 | -1.061142703 | 0.069406605 |
| Insm1 | 0.720037155 | 0.069463645 |
| Upp1 | -2.900523175 | 0.069463645 |
| Gm12216 | -3.074264217 | 0.069463645 |
| Ak1 | 0.443499821 | 0.069482882 |
| Dlat | -0.467934101 | 0.069482882 |
| Cstf2 | -0.443869865 | 0.069574441 |
| Cep85l | -0.744621163 | 0.069580884 |
| Bbip1 | 0.441834063 | 0.06962886 |
| Haus6 | -0.615455376 | 0.069730214 |
| Scn2b | 1.333593164 | 0.069743936 |
| Wfdc2 | 1.733971332 | 0.06984013 |
| Gyltl1b | -4.370753602 | 0.069840284 |
| Lrpap1 | -0.394978415 | 0.069868702 |
| Slc2a4 | 2.871878611 | 0.069872744 |
| Mpped1 | -0.66632929 | 0.069968483 |
| Rpl27-ps3 | 0.720283009 | 0.069990586 |
| Ppil6 | 3.093151753 | 0.069991527 |
| Gm10076 | 0.647433843 | 0.070002756 |
| Hdac8 | 0.848962589 | 0.07004683 |
| Rab7b | -1.315388749 | 0.070125834 |
| St13 | 0.362233489 | 0.070158842 |
| Bet1l | 0.674741736 | 0.070171715 |
| Cox5b | 0.404673045 | 0.070180514 |
| Rab5c | -0.475410442 | 0.070180514 |
| Pus7l | 1.682469688 | 0.070457997 |
| Tmem176a | -1.592414679 | 0.070457997 |
| Psmg1 | 0.516456279 | 0.070563541 |
| Olfr543 | -4.304223135 | 0.070668318 |
| Vopp1 | 0.431058759 | 0.070730376 |
| Spib | -1.892304248 | 0.070730376 |
| Plcb2 | -1.067649313 | 0.07073051 |
| Tmem151a | 1.251499936 | 0.070757283 |
| Mogat1 | -2.820696858 | 0.0708202 |
| Ralgapb | -0.665967032 | 0.070912927 |
| Psme2 | -0.557014923 | 0.070973438 |
| Fmo1 | 3.047483301 | 0.070992888 |
| Tubb2a | 0.369594646 | 0.070992888 |
| Pgam1 | -0.394839156 | 0.070992888 |
| Fam189a1 | -1.722300001 | 0.070992888 |
| Zfp105 | 1.035418628 | 0.071034578 |
| Scamp4 | -2.568083531 | 0.071034578 |
| Bcas1 | 2.9083783 | 0.071290636 |
| Gm3752 | 4.876600504 | 0.07136765 |
| Noc2l | -0.56928675 | 0.071450757 |
| Slc25a48 | 5.409645472 | 0.071474997 |
| Dleu7 | 0.84939356 | 0.071474997 |
| Zfp455 | -3.305381353 | 0.07153997 |
| Nemf | 0.424906518 | 0.07159241 |
| Nucb1 | -0.521507838 | 0.07159241 |
| Dpy19l4 | -0.813475596 | 0.071625584 |
| Morn1 | -0.997679804 | 0.071625584 |
| Mmd | 0.435084498 | 0.07185367 |
| Il7 | 5.015622872 | 0.071892971 |
| Alx1 | 5.009392356 | 0.071892971 |
| Fam104a | 0.413974044 | 0.071981003 |
| Cited1 | 2.323434143 | 0.071997035 |
| Idi2 | -5.57895046 | 0.071997035 |
| Rnf122 | 0.67319734 | 0.07204007 |
| Kalrn | -0.505512071 | 0.072156849 |
| Acsl4 | -0.752654229 | 0.072165921 |
| 1110017D15Rik | 1.253693817 | 0.072325224 |
| Gad1 | 0.812696035 | 0.07233263 |
| Kdsr | 0.716198537 | 0.07233263 |
| Ubb | 0.30565445 | 0.07233263 |
| Mdn1 | -0.588740956 | 0.072351261 |
| Msrb1 | -1.064003954 | 0.072421025 |
| Penk | 1.397367387 | 0.07243043 |
| She | 3.656055661 | 0.072444698 |
| Prkag2 | 0.590982129 | 0.072444698 |
| Mast1 | -0.780025683 | 0.072444698 |
| Cfap126 | 0.932768301 | 0.072528008 |
| Gmeb2 | -0.8313117 | 0.072586804 |
| Ibtk | 0.510158404 | 0.072616072 |
| Zfp503 | 1.382364318 | 0.072622413 |
| Gmcl1 | -0.506407985 | 0.072622413 |
| Igfbp3 | -1.219087104 | 0.072659303 |
| Krt1 | 1.09081621 | 0.072671142 |
| Lyzl | -5.88011854 | 0.072705409 |
| Zfp180 | 0.819276101 | 0.072803642 |
| Ppie | 0.533483364 | 0.072803642 |
| Ppef2 | -5.52665285 | 0.072868821 |
| Slc10a1 | 4.916460723 | 0.072943677 |
| Tnfsf13b | -1.095520121 | 0.073131014 |
| Tktl2 | -5.890872655 | 0.073131014 |
| Rtkn2 | 1.045973531 | 0.073185792 |
| N4bp3 | -2.328058636 | 0.073467879 |
| Naa16 | -0.733722129 | 0.07354611 |
| Pzp | 5.363387372 | 0.073624642 |
| Kbtbd12 | 5.173269354 | 0.073624642 |
| Rbm12 | 0.682482182 | 0.073704308 |
| Myo1g | -1.941071121 | 0.073783717 |
| Tnfaip1 | 0.410815811 | 0.073830785 |
| Ilf2 | 0.387482435 | 0.073835787 |
| Ints1 | -0.617955117 | 0.073835787 |
| Fam135a | 0.580964252 | 0.073864483 |
| Prokr2 | 1.018819206 | 0.073916618 |
| Lhfpl4 | 0.489774029 | 0.073916618 |
| Zfp493 | 4.977748311 | 0.073923053 |
| Fahd1 | 1.2938166 | 0.073923053 |
| Clip4 | 1.518879534 | 0.07394489 |
| D2hgdh | -1.126439685 | 0.07394489 |
| Hibch | 0.728919333 | 0.073991791 |
| Sharpin | -0.555834456 | 0.073991791 |
| Nmnat3 | -1.999844074 | 0.073991791 |
| Lrrc7 | -0.636630486 | 0.074159988 |
| Mrps22 | 0.482425477 | 0.074228759 |
| Capn15 | -0.788736994 | 0.074355358 |
| Adamts10 | 0.825709224 | 0.074361539 |
| Sars2 | -0.677968087 | 0.074362586 |
| Zfp689 | 0.859325617 | 0.074458551 |
| Txnl4a | 0.432209623 | 0.074458551 |
| Lix1l | 0.7700453 | 0.074480832 |
| Dscc1 | -0.765165015 | 0.074480832 |
| Gjb2 | 5.003168381 | 0.074516623 |
| Sgms1 | -1.066802293 | 0.074566763 |
| Vps72 | 0.349844967 | 0.074609533 |
| Entpd7 | -0.78827534 | 0.074652951 |
| 4930503L19Rik | -0.680968522 | 0.074658437 |

| | | |
|---|---|---|
| Vgll3 | 4.101993351 | 0.353114376 |
| Lmln | -0.679887686 | 0.353114376 |
| Lrrc46 | 1.733035101 | 0.353145021 |
| Fam50a | 0.264737309 | 0.353145021 |
| Ereg | -1.364195184 | 0.353305416 |
| Flg2 | -4.559388174 | 0.353305416 |
| Olfr78 | -4.559388174 | 0.353305416 |
| S100a7a | -4.559388174 | 0.353305416 |
| Tmem89 | -4.559388174 | 0.353305416 |
| Prelid1 | 0.266049155 | 0.353321116 |
| Rif1 | 0.246265929 | 0.353419472 |
| Tns2 | 0.917165665 | 0.353555414 |
| Foxj1 | 0.857486293 | 0.353555414 |
| Mbtps1 | -0.33166815 | 0.353555414 |
| Htr3a | 0.994150128 | 0.353646836 |
| Anxa2 | -0.897221046 | 0.354178323 |
| Xlr3b | 0.47397425 | 0.354180047 |
| Zzz3 | 0.274231144 | 0.354428523 |
| Psmd3 | -0.226174608 | 0.354661198 |
| Ppp6r2 | -0.334758774 | 0.354717138 |
| Mccc1 | -0.295107528 | 0.354856202 |
| Ngf | 3.114437935 | 0.355313381 |
| Zfp703 | -0.49309579 | 0.355719678 |
| Pcdhac1 | -1.600099533 | 0.35581891 |
| Cecr6 | 0.836295874 | 0.355838612 |
| Slc16a4 | -0.827380369 | 0.355838612 |
| Ppm1n | -1.133852885 | 0.355930173 |
| 4930518I15Rik | 0.815153554 | 0.356036324 |
| Incenp | -0.357023855 | 0.356144726 |
| Rab15 | 0.466034238 | 0.356939452 |
| Prdm12 | 3.762040337 | 0.357264183 |
| Cabp2 | 4.032615384 | 0.35733085 |
| Wdr18 | -0.217255485 | 0.357372138 |
| Zfp575 | 0.654940128 | 0.357477339 |
| Rab14 | 0.209207702 | 0.357477339 |
| Crip1 | 0.675069803 | 0.357575357 |
| Pla2g4d | 4.048401939 | 0.357655376 |
| Ak5 | 1.414958195 | 0.357668189 |
| Nkx2-2 | 4.018100511 | 0.357901336 |
| 9330151L19Rik | -0.566284445 | 0.357947665 |
| D630036H23Rik | -1.815775917 | 0.357947665 |
| Rpl13-ps3 | 1.323136984 | 0.357955916 |
| Dyrk1a | 0.318796252 | 0.357955916 |
| Ttc30a1 | 0.706804248 | 0.358222252 |
| Fam179a | 4.04701501 | 0.358403875 |
| Trpm1 | 4.04701501 | 0.358403875 |
| Trim46 | 0.373104749 | 0.358403875 |
| Set | 0.22126653 | 0.358604129 |
| Arrb2 | -0.450066655 | 0.35895065 |
| Slc22a5 | -0.658842538 | 0.35895065 |
| Nufip1 | -0.302957875 | 0.35916391 |
| Klk10 | 3.75431513 | 0.359224931 |
| Rnf167 | -0.292413207 | 0.359773224 |
| Galk1 | 0.273913712 | 0.359944446 |
| Cacul1 | 0.424811914 | 0.359984017 |
| Rpap3 | 0.338750329 | 0.359984017 |
| Qtrtd1 | 0.516758716 | 0.360002516 |
| 4933408J17Rik | -4.245403233 | 0.360002516 |
| Ihh | -4.245403233 | 0.360002516 |
| Mfsd11 | -0.410008269 | 0.360042937 |
| C1qa | -0.432293236 | 0.360042937 |
| Nolc1 | 0.258233771 | 0.360435717 |
| Ovol1 | -2.040851879 | 0.360435717 |
| Fam98c | -0.546380546 | 0.360469182 |
| Napepld | 0.857333452 | 0.360638327 |
| Ppp2r3a | 0.294213868 | 0.360678744 |
| Irgm2 | -1.158744469 | 0.361065133 |
| Acss1 | -0.414791974 | 0.361451508 |
| C2cd4d | 1.951104767 | 0.361599003 |
| Haghl | 0.294244603 | 0.361741278 |
| Palm | 0.259493184 | 0.361741278 |
| Gpc4 | 0.620827266 | 0.361834094 |
| Txn1 | 0.267546412 | 0.361897977 |
| Cox6b2 | 1.030659477 | 0.362060248 |
| Pml | -0.753209478 | 0.362075234 |
| Ube2j1 | -0.415720733 | 0.362113822 |
| Krtcap3 | -1.070064673 | 0.362290678 |
| Col6a4 | 4.365872275 | 0.362442089 |
| Fgf11 | 0.578364239 | 0.362442089 |
| Ivl | -4.520855086 | 0.362442089 |
| Mrgprh | -4.520855086 | 0.362442089 |
| RP24-254M13.1 | -4.520855086 | 0.362442089 |
| Ntrk1 | -3.299911077 | 0.362644542 |
| Ect2 | 0.408373788 | 0.362645363 |
| 1700019G17Rik | -1.092325292 | 0.3626767 |
| Kansl3 | 0.305994716 | 0.362748609 |
| Wrap73 | 0.33343079 | 0.363032963 |
| Top3b | -0.317580772 | 0.363078173 |
| Fgd6 | -0.460821633 | 0.363078173 |
| Wbscr27 | 0.940456752 | 0.363350453 |
| Sun1 | -0.291994927 | 0.363350453 |
| Cdr2 | 0.904442353 | 0.363432272 |
| Dap3 | 0.249586669 | 0.36351997 |
| Muc2 | 3.699324499 | 0.363542065 |
| Myo5c | 3.699324499 | 0.363542065 |
| Vrk1 | 0.28237029 | 0.363563067 |
| Gnptab | -0.473793812 | 0.363632889 |
| Arhgap15 | -0.814882912 | 0.363682177 |
| Nphp4 | -0.975627035 | 0.363719272 |
| Pde4c | -3.767370785 | 0.363719272 |
| Mttp | -0.773901462 | 0.363932548 |
| Ccdc142 | -0.8670559 | 0.363941164 |
| Golt1b | -0.255945968 | 0.364116861 |
| Pde6b | -3.395602644 | 0.364188844 |
| Spop | -0.203747542 | 0.364353208 |
| Mettl21a | 0.550380624 | 0.364410254 |
| Tmem67 | -0.453865167 | 0.364410254 |
| Pfkfb2 | 0.530634715 | 0.364449607 |
| Zfp382 | -0.603305332 | 0.364550005 |
| Kcnc1 | -0.427708368 | 0.364587396 |
| Agbl3 | -0.640415985 | 0.364587396 |
| Glyctk | -0.866959296 | 0.364587396 |
| Ppp1r9b | -0.36367007 | 0.364828626 |
| Chn2 | 0.333823696 | 0.365212058 |
| Tmbim1 | -0.585030562 | 0.365225191 |
| Aatk | -0.63280327 | 0.365225191 |
| Cdc27 | -0.254864741 | 0.365359552 |
| Syt12 | 2.27788685 | 0.365535769 |
| Klhdc7a | 3.995524926 | 0.365556534 |

| | | |
|---|---|---|
| Ints4 | -0.126358639 | 0.745518199 |
| Slc6a15 | -0.182006341 | 0.745518199 |
| Dnase2a | -0.191290186 | 0.745518199 |
| Pigl | -0.276044979 | 0.745518199 |
| Chil1 | -0.357582306 | 0.745607623 |
| P4ha2 | -0.265238638 | 0.74569208 |
| Triqk | -0.23702513 | 0.745839052 |
| Naa35 | -0.104163021 | 0.745851865 |
| Zfpm1 | -0.547055802 | 0.746191294 |
| Decr1 | 0.194306705 | 0.746215992 |
| Gabpb1 | 0.111299401 | 0.746543226 |
| Pcyox1l | -0.168439814 | 0.746543226 |
| Gm10479 | -0.731489174 | 0.746543226 |
| Orm1 | -0.876995116 | 0.746543226 |
| Gspt2 | -0.208730981 | 0.746547455 |
| Apoo-ps | 0.228388489 | 0.746789119 |
| Ercc5 | -0.137727104 | 0.74685442 |
| Rpl7a-ps3 | -0.720392951 | 0.747099268 |
| Sycp2 | 0.459605447 | 0.747212214 |
| Csrp1 | 0.155026312 | 0.747212214 |
| Tmem251 | -0.139676503 | 0.747299848 |
| Ccdc17 | 0.28874484 | 0.747301443 |
| Gm17545 | -0.994656128 | 0.747301443 |
| Thy1 | 0.324222919 | 0.747541694 |
| Ercc4 | -0.148049147 | 0.747541694 |
| Rnf128 | -0.3913526 | 0.747541694 |
| Dnaja4 | -0.51196615 | 0.747541694 |
| Gm43951 | -0.926279302 | 0.747541694 |
| 3110062M04Ri | -0.25930346 | 0.74756559 |
| Brms1 | -0.124099356 | 0.747611957 |
| Esco1 | -0.160506785 | 0.747611957 |
| Ube2e2 | 0.124617922 | 0.747781861 |
| Chid1 | -0.144922148 | 0.747808526 |
| Rnf25 | -0.185325356 | 0.747855672 |
| Leprot | -0.108346326 | 0.747949777 |
| Marveld2 | 0.843781924 | 0.748510551 |
| Ldah | 0.090876403 | 0.748604155 |
| Smim20 | -0.098956313 | 0.748604155 |
| Traip | 0.183416964 | 0.7489004 |
| Efhb | 0.59611298 | 0.748915688 |
| Swsap1 | 0.163976534 | 0.748970903 |
| Nup155 | -0.117456466 | 0.749070961 |
| Plp2 | -0.190673118 | 0.74908374 |
| Adcy1 | 0.185615494 | 0.749377294 |
| Lrrc14 | -0.129513468 | 0.749377294 |
| Gm29609 | -0.508367176 | 0.749377294 |
| Aldh1a7 | 0.512848287 | 0.749474143 |
| Vcpip1 | -0.098451509 | 0.749556705 |
| Antxr1 | 0.235841093 | 0.749577359 |
| C2cd2l | -0.157273365 | 0.749577359 |
| Hrh3 | -0.41928715 | 0.749577359 |
| Rpp40 | -0.197268907 | 0.749809405 |
| 3110052M02Ri | -0.24813 | 0.749856171 |
| Klrk1 | 0.87394045 | 0.749865218 |
| Vdac2 | -0.087626103 | 0.749865218 |
| Cib1 | -0.107618296 | 0.750068356 |
| Ppm1j | 0.480960725 | 0.750284596 |
| Lig4 | 0.332283737 | 0.750341379 |
| Ikbkap | 0.104816433 | 0.750341379 |
| Acsf2 | 0.389790318 | 0.750565647 |
| Nsdhl | 0.113025153 | 0.750571058 |
| Rsl1 | -0.23478925 | 0.750571058 |
| Toe1 | 0.227478218 | 0.750733553 |
| Zfp820 | 0.466990522 | 0.750903352 |
| Orc6 | 0.117943252 | 0.750903352 |
| Alox8 | -1.479366874 | 0.750903352 |
| Setbp1 | -0.11532918 | 0.751230074 |
| Xrcc2 | -0.303811229 | 0.751230074 |
| Ptges2 | -0.110879899 | 0.751489359 |
| Secisbp2 | -0.118126226 | 0.751994462 |
| Dock6 | 0.275730785 | 0.752132851 |
| Slc2a2 | 1.089713788 | 0.75217007 |
| Add3 | 0.12903363 | 0.75217007 |
| Dnajc10 | 0.10396472 | 0.75217007 |
| Ecsit | 0.150260316 | 0.752343198 |
| Zfp944 | -0.178518326 | 0.752471415 |
| Rprml | 0.417401643 | 0.75259294 |
| Gm13199 | 1.117901123 | 0.752642779 |
| Cfap73 | 1.26500596 | 0.752643896 |
| Dmpk | 0.337796361 | 0.752643896 |
| Pcmtd2 | 0.138747747 | 0.752643896 |
| Zhx3 | -0.145320386 | 0.752643896 |
| Pdcl3 | -0.102976067 | 0.752760276 |
| Encl | -0.105512074 | 0.753127195 |
| Tapbpl | 0.426355082 | 0.753147897 |
| Ypel4 | -0.161000951 | 0.753797463 |
| Ucp1 | 1.006618021 | 0.753844364 |
| Aqr | -0.107776312 | 0.753844364 |
| Atp13a3 | -0.148669933 | 0.753844364 |
| Kcnh3 | -0.331212641 | 0.753890615 |
| Pgr | -0.702201073 | 0.753944165 |
| Gm10309 | -0.578390812 | 0.7540273 |
| Klf10 | -0.18164277 | 0.754462758 |
| Cspg5 | -0.16801293 | 0.754509641 |
| Polm | -0.22078646 | 0.754552579 |
| Rictor | 0.168759964 | 0.754555261 |
| Cldn1 | 0.551703276 | 0.75461531 |
| Rps6kb2 | 0.140183936 | 0.75461531 |
| Uprt | -0.234693082 | 0.75461531 |
| BC106179 | -0.377534506 | 0.75461531 |
| Plekhm2 | 0.206913459 | 0.754814645 |
| Adgrl1 | 0.143287214 | 0.754814645 |
| Zfp72 | 0.576362716 | 0.754873759 |
| Anpep | 0.566977489 | 0.754924699 |
| Car5a | -0.86844098 | 0.754924699 |
| Slc24a5 | 0.33434826 | 0.755132233 |
| Rad51d | 0.186093622 | 0.755132233 |
| Grk5 | -0.325723069 | 0.755143502 |
| Hus1b | 1.510766232 | 0.755654929 |
| Olfm1 | -0.102394424 | 0.755654929 |
| Eif1ad | 0.096040475 | 0.755699166 |
| Kdm6b | 0.090682544 | 0.756060411 |
| Cxcl10 | 0.214713831 | 0.756071991 |
| A430005L14Ri | 0.14451515 | 0.756615345 |
| Muc13 | -1.622916399 | 0.756615345 |
| Aff3 | -0.123784075 | 0.756620429 |
| Kctd6 | 0.161331293 | 0.756849662 |
| Usp51 | -0.364814474 | 0.757057662 |
| Apobec2 | -0.720839106 | 0.757083005 |

| | | |
|---|---|---|
| Cyfip1 | -0.935280888 | 0.000164508 |
| Wbp1 | 0.723710221 | 0.000164702 |
| Ppa1 | 0.741399081 | 0.000165486 |
| Snx9 | -1.453869229 | 0.000165532 |
| Esrrg | 2.645491006 | 0.000167288 |
| Cobl | 2.528039822 | 0.000167966 |
| Tbc1d7 | 0.883796361 | 0.000168243 |
| Pin4 | 0.963298872 | 0.000168334 |
| Bmp5 | 6.722368964 | 0.00016891 |
| Mkl1 | -1.067964186 | 0.000170934 |
| Sltm | 0.766849239 | 0.000171044 |
| N4bp1 | -1.42610938 | 0.000171145 |
| Sydel | 6.24509617 | 0.000171885 |
| Slc31a1 | -1.272152073 | 0.000172381 |
| Ndufa2 | 0.715661609 | 0.000173725 |
| Mrpl32 | 0.845602831 | 0.000173979 |
| Arhgef2 | -0.813747331 | 0.000174033 |
| Ndufab1 | 0.674268354 | 0.00017491 |
| Nenf | 0.913803702 | 0.000175414 |
| Dapk1 | -0.894730958 | 0.000175852 |
| D8Ertd82e | -1.049307948 | 0.000176371 |
| Itih5 | 4.024552071 | 0.000177071 |
| Zfp395 | -1.568843676 | 0.000177395 |
| Ly9 | -1.695121915 | 0.000178131 |
| Nploc4 | -1.193513244 | 0.000178861 |
| Pxk | -1.196693375 | 0.000179014 |
| Borcs7 | 1.321732584 | 0.000180194 |
| Mrpl42 | 0.767236158 | 0.000180255 |
| Ppdpf | 0.905837347 | 0.000180492 |
| Zscan21 | 0.947167181 | 0.000181126 |
| Sox2 | 1.099932585 | 0.000181273 |
| Gtf2ird1 | -0.93807999 | 0.000182188 |
| Cnn2 | -1.390376076 | 0.000184219 |
| Nin | -1.474425208 | 0.000185381 |
| Rplp1 | 0.939548443 | 0.000186719 |
| Gsdmd | -1.435094573 | 0.000187089 |
| 0610011F06Rik | 0.954435734 | 0.000187921 |
| Hes6 | 0.687829078 | 0.000188668 |
| P4ha3 | 2.586213169 | 0.00018884 |
| Cdc45 | -1.28997003 | 0.00018884 |
| Pbxip1 | -1.313724258 | 0.000190085 |
| Mast3 | -1.27401289 | 0.000190747 |
| Bmp4 | 6.706423555 | 0.000192153 |
| Stbd1 | 4.534705762 | 0.000193001 |
| Ubl5 | 0.849603441 | 0.000193038 |
| Mphosph8 | 1.164915272 | 0.000193163 |
| Ccdc88a | 0.749907705 | 0.000193309 |
| Sgk1 | -1.575883013 | 0.000196172 |
| Atp5g3 | 0.709729388 | 0.000196646 |
| Polr3h | 1.10299113 | 0.000196954 |
| Pih1d2 | 2.830967413 | 0.000197789 |
| Sash1 | -1.745251743 | 0.000197971 |
| Col5a1 | 6.61506981 | 0.000198711 |
| Rpl39 | 0.694982186 | 0.000198711 |
| Hk3 | -1.546693941 | 0.000198711 |
| Oard1 | 0.921695567 | 0.000199343 |
| Angptl2 | 3.187011308 | 0.000201129 |
| Serpina3g | -2.478568427 | 0.000201367 |
| Prkrir | 1.014185207 | 0.000202314 |
| Hmgn2 | 0.775901725 | 0.000202314 |
| Epn1 | -1.168553207 | 0.000203488 |
| Slamf6 | -3.846224589 | 0.000203488 |
| Siva1 | 0.973507881 | 0.000204066 |
| Nap1l1 | 0.642376259 | 0.000204773 |
| Mafb | -1.266438279 | 0.000208703 |
| Wbp1l | -1.018356449 | 0.000208808 |
| Egr1 | 0.705473518 | 0.000210244 |
| Pycard | -1.055243285 | 0.000210301 |
| Ndufaf2 | 1.26697158 | 0.000212357 |
| Hps5 | -1.454739307 | 0.0002136 |
| Gab1 | -1.121468709 | 0.000213603 |
| Uchl3 | 0.834549211 | 0.000215119 |
| Adora2a | -2.488558809 | 0.000215418 |
| Fam107b | -0.977905997 | 0.000216968 |
| Sgsh | -2.238994655 | 0.000217262 |
| Stk24 | -1.491300643 | 0.000220735 |
| Brpf1 | -1.01465078 | 0.000220891 |
| Rab11fip5 | -1.142543693 | 0.000220891 |
| Rpp21 | 1.229133343 | 0.000221499 |
| Eif3k | 0.669831319 | 0.000221507 |
| Col23a1 | 1.728427071 | 0.000221517 |
| Pcbd1 | 1.954097587 | 0.000221649 |
| Ceacam18 | -4.890026804 | 0.000223251 |
| Mrps21 | 0.843676509 | 0.00022525 |
| Wnt5a | 1.435078532 | 0.000225644 |
| 4932438H23Rik | 6.880062174 | 0.000226666 |
| Lingo2 | 2.678017964 | 0.000226666 |
| Spint1 | -7.643750124 | 0.000226666 |
| Ctnnb1 | 0.687706237 | 0.000227467 |
| Mtpn | -0.660755312 | 0.000228371 |
| U2af2 | -0.834394462 | 0.000228371 |
| Specc1l | -0.861707469 | 0.000228526 |
| Ltbp3 | 2.456636105 | 0.000229953 |
| Eif1ax | 0.735269983 | 0.00023495 |
| Bola1 | 1.071137132 | 0.000238354 |
| Tspan14 | -0.956952421 | 0.000238354 |
| Cant1 | -0.906600449 | 0.000239038 |
| Timm13 | 0.796304459 | 0.000239528 |
| Slitrk6 | 6.795736849 | 0.000241301 |
| Nlrp3 | -2.268374228 | 0.000241641 |
| Cxcl1 | 2.737013296 | 0.000242184 |
| Upf3b | 1.103752302 | 0.000242184 |
| Hnmpf | -0.768779333 | 0.000242532 |
| Map4k2 | -1.371491496 | 0.000242532 |
| Zc3h12d | -2.631017641 | 0.000244189 |
| Sgpp1 | -1.274979133 | 0.000244473 |
| Gm2000 | 0.827709415 | 0.000245444 |
| Cysltr1 | -2.008869836 | 0.000245815 |
| B3gat1 | -1.567376764 | 0.000246011 |
| Trpc4ap | -0.756042843 | 0.000247567 |
| Tm6sf1 | -1.569788168 | 0.000248279 |
| Ccr2a | -7.175345068 | 0.000248518 |
| Tln1 | -1.042197481 | 0.000248861 |
| Pck2 | 1.576955693 | 0.000250953 |
| Slc35a3 | -1.296763233 | 0.000250953 |
| Lnpep | -0.823721355 | 0.000251557 |
| Ccdc85b | 1.136955515 | 0.000252389 |
| Zfp329 | 0.9829862 | 0.000252389 |
| Telo2 | -1.568818569 | 0.000254032 |

| | | |
|---|---|---|
| Scg3 | 0.474218316 | 0.074670211 |
| Nsmce2 | 0.508639859 | 0.074683172 |
| Tmpo | -0.371316023 | 0.074720044 |
| Armc5 | -0.638473366 | 0.074720044 |
| Slc26a7 | 2.254052672 | 0.07479313 |
| Sbspon | -4.150896145 | 0.074835271 |
| Serpina10 | 5.15499678 | 0.074840633 |
| Atp10b | 3.703914101 | 0.074840633 |
| Ppfibp2 | -1.044011306 | 0.074927711 |
| Tek | 5.055444212 | 0.07496344 |
| Tmem71 | -3.381782293 | 0.07496344 |
| Tbc1d20 | -0.523401552 | 0.074989304 |
| 3100002H09Rik | -5.490986103 | 0.075041366 |
| Desi1 | 0.459081978 | 0.075076267 |
| Tdrd6 | -2.965019963 | 0.075163623 |
| Sucla2 | 0.392046307 | 0.075184588 |
| Gpr157 | -1.390066703 | 0.075209364 |
| Caprin1 | -0.390442227 | 0.075336732 |
| Sec61a1 | -0.420599708 | 0.0754638 |
| Zc3h6 | 1.137941217 | 0.075480119 |
| Spag1 | 2.605410194 | 0.075492546 |
| Dram1 | -2.10204386 | 0.075520596 |
| Dok7 | -1.859628688 | 0.075521347 |
| Emp3 | -0.777422966 | 0.075550629 |
| I7Rn6 | 0.39783187 | 0.07557342 |
| Zbp1 | -2.127896638 | 0.075636109 |
| E130311K13Rik | 1.383553177 | 0.075645006 |
| Pold4 | -0.676650944 | 0.075648012 |
| Plcxd2 | 0.818041889 | 0.075776099 |
| Zbtb24 | 0.671508692 | 0.075776099 |
| Cyp2d22 | 4.28187491 | 0.075782928 |
| Patl1 | -0.797154619 | 0.075794711 |
| Gm5475 | -5.570519409 | 0.075794711 |
| Tnfsf15 | -5.570519409 | 0.075794711 |
| Mamdc2 | 4.950117592 | 0.075806287 |
| Ubxn2a | 0.643804374 | 0.075806287 |
| Gmps | -0.388298793 | 0.075855921 |
| Fbxl22 | 2.004534462 | 0.076029857 |
| Ttc19 | -0.514474219 | 0.07612202 |
| Gpt | -1.312605854 | 0.076137602 |
| Itga2 | 3.598017064 | 0.076190087 |
| Zscan10 | -5.600540164 | 0.076190087 |
| Frmpd4 | -2.012154964 | 0.076206908 |
| Actc1 | 5.077826562 | 0.076295658 |
| Ipo11 | 0.52805034 | 0.076369202 |
| Fam132b | -4.812317938 | 0.07637625 |
| Nedd9 | -0.791225585 | 0.076456775 |
| Adm | -1.279596221 | 0.076456775 |
| Crybg3 | -1.265730677 | 0.076465928 |
| Lyrm9 | 0.752850256 | 0.076634484 |
| Ripply1 | -4.833115916 | 0.076634484 |
| Dbx2 | 5.080738237 | 0.076679872 |
| Pf4 | 1.084562202 | 0.076809084 |
| Ebf1 | 1.031855018 | 0.076833416 |
| Bckdk | -0.40539638 | 0.076835249 |
| Adamts16 | 3.482777074 | 0.076935334 |
| Plk1 | 0.635909326 | 0.076935334 |
| Vdac3-ps1 | 0.480596915 | 0.076955878 |
| Asic4 | -2.528899294 | 0.076955878 |
| Zfp605 | 0.845155161 | 0.076965927 |
| Med10 | 0.413147375 | 0.076977295 |
| Serpinb2 | -5.8506706 | 0.0769984 |
| Gabrq | 3.118999941 | 0.077032228 |
| Rasal3 | -0.795700203 | 0.077036193 |
| Rgn | -5.53266571 | 0.077118583 |
| Nkiras1 | 0.695590373 | 0.077251922 |
| Ankrd26 | 0.724260783 | 0.077252838 |
| Psmd4 | 0.377118548 | 0.077263998 |
| Arl11 | -1.118462217 | 0.077279555 |
| Arhgap44 | 0.682066862 | 0.077305281 |
| Syp | 0.628789483 | 0.077305281 |
| Zbed3 | 0.597244503 | 0.077305281 |
| Mrgbp | 0.555524072 | 0.077305281 |
| Col2a1 | 1.160342035 | 0.077372457 |
| Tmem51 | -1.314990008 | 0.077372457 |
| Glipr1 | -1.734303143 | 0.077386798 |
| Dnajb7 | -5.486453115 | 0.077388799 |
| Mrpl15 | 0.396717932 | 0.077444138 |
| Ccdc67 | 5.281390963 | 0.07749436 |
| Gm10083 | 4.986618135 | 0.077528653 |
| Pex19 | 0.525256341 | 0.077736287 |
| Tmem150a | 1.131505892 | 0.077739232 |
| Plxna3 | -0.576767943 | 0.077783 |
| Asf1b | -0.635476421 | 0.077783 |
| Smg7 | -0.434723705 | 0.077807457 |
| Ttn | 1.487834082 | 0.077820072 |
| Atg4b | 0.455027894 | 0.077906039 |
| Ik | 0.376730776 | 0.077913354 |
| Top2a | 0.465326583 | 0.077915752 |
| Nprl3 | 0.568327255 | 0.078060624 |
| Ccdc134 | -0.756467266 | 0.078123247 |
| Stxbp5l | 1.051654367 | 0.078138492 |
| Cep97 | 0.785568239 | 0.078228625 |
| Tagap1 | -0.67510007 | 0.078228625 |
| Cd180 | -0.803187534 | 0.078228625 |
| Kri1 | -0.895268944 | 0.078228625 |
| Rnpepl1 | -0.614661229 | 0.07823885 |
| Pex12 | -0.786806749 | 0.078437497 |
| Atp6v1g1 | 0.347091504 | 0.078442954 |
| Capg | -2.390837973 | 0.078496603 |
| Gm4922 | 5.367766878 | 0.078513072 |
| Shox2 | 2.674937463 | 0.078513072 |
| Tra2a | 0.380795173 | 0.078514087 |
| Usp36 | -0.756370442 | 0.078514087 |
| Cdon | 0.634896031 | 0.078563429 |
| Mavs | -0.921366946 | 0.078563429 |
| F11r | -0.931313234 | 0.078563429 |
| Gm20604 | 0.894791249 | 0.078604165 |
| Stfa2l1 | -2.589058968 | 0.0786309 |
| Ap3m2 | -0.63454962 | 0.07865412 |
| Marcksl1 | 0.358500421 | 0.078682011 |
| H2-D1 | -0.70459456 | 0.078682011 |
| Plaur | -2.17017723 | 0.078682011 |
| Slc36a3 | -5.8349129 | 0.078682011 |
| Hfe | -0.869520793 | 0.078727649 |
| Fam204a | 0.564183716 | 0.078730433 |
| Tas1r1 | 5.133825832 | 0.078987254 |
| Brd1 | 0.591722999 | 0.07904074 |
| Hif1an | -0.768336257 | 0.07904074 |

| | | |
|---|---|---|
| Son | 0.191020568 | 0.365666075 |
| Zbtb33 | -0.375433976 | 0.365796236 |
| Gm6710 | 0.76962762 | 0.365836194 |
| 2-Mar | 0.264684657 | 0.365961385 |
| Drg1 | 0.22631312 | 0.366221991 |
| Vax2 | 3.984508907 | 0.36651652 |
| Wnt11 | 3.984508907 | 0.36651652 |
| Rrp1 | 0.192753764 | 0.366559726 |
| Wdr13 | -0.330833962 | 0.366559726 |
| Ttc7b | 0.498870487 | 0.366598786 |
| Krcc1 | -0.333323294 | 0.366678714 |
| Akt1s1 | 0.249917616 | 0.366705116 |
| Rpn2 | -0.236704931 | 0.367088857 |
| Stab2 | 2.571919051 | 0.367100874 |
| Cntn3 | 1.372833477 | 0.367100874 |
| Cpa6 | 3.708609796 | 0.367147098 |
| Fhit | 0.784560657 | 0.367147098 |
| Sowaha | 1.880156506 | 0.367720191 |
| Pld2 | 0.617866566 | 0.36813543 |
| Tmem97 | 0.412158401 | 0.368273347 |
| Zcchc5 | 3.689374088 | 0.368311453 |
| Neurod1 | 0.454640961 | 0.368357412 |
| Tctel | 2.06440085 | 0.368403487 |
| Osbpl1a | 0.362972611 | 0.368490673 |
| Ssxb5 | -3.787610435 | 0.368490673 |
| Nid2 | 0.985385266 | 0.368509356 |
| Spata13 | -0.382381129 | 0.368685429 |
| Prpf6 | 0.339378056 | 0.368810026 |
| Ubap2 | -0.269914678 | 0.368867635 |
| Tmem240 | -0.807675732 | 0.368867635 |
| Ccl4 | -0.78654132 | 0.368968257 |
| Zbtb14 | 0.411234866 | 0.369113497 |
| Sema4f | -0.663020878 | 0.369113497 |
| Akr1e1 | 0.382312911 | 0.36920236 |
| Irgq | 0.336966981 | 0.36920236 |
| Gm21975 | -0.663922739 | 0.36920236 |
| Mbip | 0.302369381 | 0.36921806 |
| Snx15 | 0.352549451 | 0.369527026 |
| Zfp410 | 0.295704101 | 0.369527026 |
| Ofd1 | 0.454601888 | 0.369720051 |
| Pla2g3 | 3.695539891 | 0.369726004 |
| Xxylt1 | 0.4978576 | 0.369726004 |
| FRMPD2 | -4.523559944 | 0.369726004 |
| Rab26 | 1.074574569 | 0.369741131 |
| Eif2b3 | 0.27927234 | 0.369989632 |
| Diexf | -0.417931332 | 0.369989632 |
| Dusp2 | -0.61283384 | 0.369989632 |
| Plekha6 | 0.522811135 | 0.370283071 |
| Lrrc41 | -0.262707397 | 0.370587437 |
| Gucd1 | -0.462333391 | 0.370587437 |
| Plgrkt | -0.253066869 | 0.371002731 |
| Kdelc2 | 0.559333388 | 0.371309599 |
| Wdr6 | -0.255290808 | 0.371383771 |
| Slc4a8 | 0.395655573 | 0.371396755 |
| Cdk7 | -0.316673158 | 0.371672035 |
| Cbx4 | -0.418169656 | 0.371767639 |
| Nme4 | 0.315392308 | 0.372000871 |
| Raly | 0.20607138 | 0.372000871 |
| Ubxn2b | -0.362260516 | 0.372083263 |
| Txnrd3 | -0.479037064 | 0.372083263 |
| Olfr370 | 3.744576213 | 0.372207853 |
| Cldn7 | -4.216720729 | 0.372509116 |
| Gimap5 | 3.627797528 | 0.372514896 |
| Gm10563 | -2.928654607 | 0.372514896 |
| 9930021J03Rik | 0.456192316 | 0.372571646 |
| Sobp | -0.404103394 | 0.372571646 |
| Morc2a | -0.280426055 | 0.372775119 |
| Ap4m1 | 0.511673981 | 0.37287819 |
| Gm9780 | 3.650181354 | 0.372956032 |
| Nphs1 | 0.746678017 | 0.372969714 |
| Fhl2 | 0.692860215 | 0.37299269 |
| Cmss1 | 0.422347183 | 0.37327984 |
| Eif2b5 | -0.23004213 | 0.373575293 |
| Stx3 | -0.677913481 | 0.373575293 |
| Fbxo36 | 0.658219271 | 0.373861817 |
| Gcnt2 | -0.671048648 | 0.373876197 |
| Gnai3 | -0.202588752 | 0.374177734 |
| Hoxa9 | -4.481290539 | 0.374577307 |
| Tmc1 | -4.481290539 | 0.374577307 |
| Dusp22 | 0.425181127 | 0.374609158 |
| Dzank1 | 1.03315841 | 0.374680015 |
| Nr0b2 | -4.474950216 | 0.374722559 |
| Sord | -0.460606675 | 0.374878062 |
| Ankrd13c | -0.278256912 | 0.374997787 |
| Gga3 | -0.404011672 | 0.374997787 |
| Slc35f1 | 0.321893246 | 0.375034748 |
| Gid8 | -0.329023379 | 0.375212897 |
| Errfi1 | 0.385816732 | 0.375401732 |
| Urgcp | -0.366551909 | 0.375401732 |
| Tmem102 | 4.311243083 | 0.375499067 |
| Gm3264 | 0.763693782 | 0.375532747 |
| Scn1b | -0.581739129 | 0.375532747 |
| Klhdc3 | -0.240779991 | 0.375547584 |
| Pnkp | -0.391224581 | 0.375824494 |
| Uros | 0.380029748 | 0.375983818 |
| Nubpl | 0.477020868 | 0.3762719 |
| Vangl2 | -0.325491538 | 0.3762719 |
| Gm9903 | -1.630398346 | 0.3762719 |
| Anln | -0.457136274 | 0.376345681 |
| Ctnnd1 | -0.467595038 | 0.376396968 |
| Apitd1 | 0.336849418 | 0.37681037 |
| Tnfrsf4 | 3.672493061 | 0.37702611 |
| Tia1 | 0.219602504 | 0.377120123 |
| Ogfod3 | 0.515487352 | 0.377150552 |
| Hgfac | -2.356593193 | 0.377383548 |
| Prrc1 | 0.308708207 | 0.377509987 |
| Duxf3 | 3.616535418 | 0.377834564 |
| Rnf123 | -0.397476323 | 0.377834564 |
| Fbxl14 | -0.55820009 | 0.377834564 |
| Gm10273 | 0.78050867 | 0.377857698 |
| Abcb9 | 0.774928405 | 0.377857698 |
| 1810011O10Rik | -0.592089015 | 0.377857698 |
| Ak7 | 3.660480535 | 0.378159456 |
| Usp39 | -0.264133083 | 0.378283841 |
| Dnase1l1 | -0.598764025 | 0.378659937 |
| Loxl2 | -0.823543837 | 0.378659937 |
| Gm17324 | -2.122677057 | 0.378778284 |
| Ncoa5 | -0.256658613 | 0.378996073 |
| Rbm12b2 | 0.544772886 | 0.379146415 |

| | | |
|---|---|---|
| Bambi | -0.176332782 | 0.757205031 |
| Epcam | 0.397210977 | 0.757380094 |
| Adck1 | 0.151680766 | 0.757540437 |
| Cd164 | 0.099673402 | 0.757730596 |
| Ttc1 | 0.09839557 | 0.757740787 |
| Hacd3 | -0.102260489 | 0.757740787 |
| Mtap | 0.111417535 | 0.757837857 |
| Tmem120b | -0.240891801 | 0.757963022 |
| Boc | 0.247747935 | 0.758126819 |
| Wdr45 | -0.173620829 | 0.758126819 |
| Rundc3a | 0.103257506 | 0.758196871 |
| Prr18 | -0.217752212 | 0.758196871 |
| Rims1 | 0.198138831 | 0.758220105 |
| Lalba | -1.015677225 | 0.758299871 |
| Ezr | 0.124782675 | 0.758317102 |
| Prepl | -0.118416734 | 0.758389435 |
| Mcph1 | -0.154428506 | 0.758640361 |
| Alas2 | 0.226956055 | 0.758645398 |
| Pawr | 0.380421686 | 0.758756654 |
| Rbm48 | 0.195628363 | 0.758756654 |
| Sgsm3 | 0.109741526 | 0.758756654 |
| Angel2 | 0.099137063 | 0.758756654 |
| Atp1b3 | -0.146408597 | 0.758756654 |
| Cys1 | 0.569409797 | 0.758804735 |
| Ly6g6d | -1.406150812 | 0.758999541 |
| Msantd2 | 0.205323444 | 0.759029343 |
| Apc2 | -0.110965218 | 0.759029343 |
| Fbf1 | 0.139295253 | 0.759055717 |
| Dcaf5 | -0.125553152 | 0.759055717 |
| Whrn | -0.232520554 | 0.759490098 |
| Scn8a | 0.134970988 | 0.759514355 |
| Nlk | -0.11771806 | 0.759514355 |
| Zfp947 | -0.886315012 | 0.759514355 |
| Sptlc1 | -0.138191198 | 0.759968152 |
| Iars2 | -0.172645264 | 0.759968152 |
| Nudt12 | -0.782718791 | 0.760120528 |
| Zfp26 | -0.11728755 | 0.760158051 |
| Rhbdl1 | -1.614560474 | 0.760297912 |
| Zfp638 | 0.122958827 | 0.76045632 |
| Rpgrip1l | -0.22150217 | 0.760747373 |
| Sec14l2 | -0.797187469 | 0.760747373 |
| Cbln2 | 0.547837607 | 0.760774302 |
| Zpr1 | -0.094510932 | 0.760774302 |
| Rufy1 | -0.109737158 | 0.760774302 |
| Larp4 | -0.110359469 | 0.760774302 |
| Tpm3-rs7 | -0.122128289 | 0.760774302 |
| Grasp | -0.401388012 | 0.760774302 |
| Pcbd2 | -0.15496937 | 0.76081559 |
| Lars | 0.096768052 | 0.760954052 |
| Ppp3ca | 0.093609893 | 0.760954052 |
| Rpl17 | 0.076363617 | 0.760954052 |
| Trpc4 | -0.295542204 | 0.760954052 |
| Creb3l4 | 0.412445474 | 0.7610441 |
| Rpusd3 | 0.197332801 | 0.761126092 |
| Fam101a | -0.47340723 | 0.761424503 |
| Cacna2d3 | 0.198710885 | 0.761518833 |
| Tbxas1 | -0.215913974 | 0.761682937 |
| Zscan30 | 0.734085365 | 0.761777481 |
| Gm26897 | 0.312360717 | 0.761777481 |
| Otub2 | 0.191444037 | 0.761777481 |
| Nufip2 | 0.12527333 | 0.761777481 |
| Luc7l3 | 0.082293372 | 0.761777481 |
| Bzw1 | -0.070597015 | 0.761777481 |
| Kif5a | -0.105245114 | 0.761777481 |
| Zfp534 | -1.390953121 | 0.761777481 |
| Psme3 | 0.12494903 | 0.761901071 |
| Cldn3 | -0.400476373 | 0.761901071 |
| Slc25a47 | 0.428723807 | 0.762068332 |
| Nfl | -0.115079328 | 0.76220562 |
| Neurog2 | 0.218280037 | 0.762253033 |
| D430019H16Ri | -0.109010647 | 0.762309575 |
| Qpct | -0.233460983 | 0.762549546 |
| Phlda3 | 0.276153416 | 0.76262829 |
| Trip12 | 0.080656444 | 0.76262829 |
| Ccdc177 | -0.157718258 | 0.76262829 |
| Egln3 | -0.239609819 | 0.76262829 |
| Dync1li1 | -0.082614094 | 0.762632966 |
| Vars | -0.104416734 | 0.762657708 |
| Rnf182 | 0.136286421 | 0.76300443 |
| Ebf4 | -0.37611057 | 0.763055322 |
| Bcat1 | 0.128860449 | 0.763102784 |
| Mapk6 | 0.105853506 | 0.763119014 |
| Slitrk2 | 0.177306632 | 0.763348985 |
| Clec3b | 0.711530036 | 0.76344477 |
| Rnf183 | 0.813968047 | 0.763445104 |
| Arhgap1 | 0.089147796 | 0.763973515 |
| Ccdc106 | -0.149797003 | 0.764486157 |
| Prkaa1 | -0.219581396 | 0.764662824 |
| Bbs4 | -0.10627223 | 0.765261823 |
| Ice2 | -0.140327548 | 0.76527591 |
| Podxl | 0.200405524 | 0.765355325 |
| Ddb2 | 0.263096611 | 0.76574229 |
| Gins4 | -0.127392781 | 0.76574229 |
| Siglecf | -0.374587054 | 0.76574229 |
| Pif1 | 0.372831519 | 0.765744014 |
| Ptgs1 | 0.173197258 | 0.765744014 |
| Glb1 | -0.145566913 | 0.765744014 |
| Kifc5b | -0.291555724 | 0.765744014 |
| Rhov | -0.20556616 | 0.765785595 |
| Dhx36 | 0.102600308 | 0.765865225 |
| Calcoco1 | -0.147560978 | 0.765865225 |
| Gm38394 | -0.526427287 | 0.765865225 |
| Sytl2 | 0.406792347 | 0.765922949 |
| Slc18a2 | -0.182092712 | 0.765922949 |
| AI464131 | 0.295402154 | 0.766251906 |
| Tex261 | 0.095918363 | 0.76646028 |
| Rer1 | -0.084349 | 0.766462743 |
| Ing2 | -0.151056219 | 0.766543558 |
| Slc16a9 | 1.089314271 | 0.766547757 |
| 1110065P20Ri | 0.206167345 | 0.766547757 |
| Vta1 | 0.088421902 | 0.766547757 |
| Ap2a2 | -0.087204981 | 0.766547757 |
| Slu7 | -0.096314231 | 0.766547757 |
| Hyal2 | 0.120466537 | 0.766588491 |
| Uba3 | -0.08364513 | 0.766598244 |
| Lin37 | 0.141524349 | 0.766667896 |
| Dlgap3 | -0.255634769 | 0.766667896 |
| Pvrl1 | -0.262757943 | 0.766703556 |
| Sgsm1 | 0.226622416 | 0.766785217 |

| | | |
|---|---|---|
| Prrg1 | 3.39403997 | 0.000255629 |
| Prpf4b | 0.992836796 | 0.000257885 |
| Pgk1 | -0.773321802 | 0.000257885 |
| Elmo1 | -1.100230897 | 0.000258581 |
| Kif1c | -1.57452884 | 0.000258606 |
| Hey1 | 1.505911659 | 0.000260514 |
| Kdm2a | -0.823896606 | 0.0002637 |
| Slc22a8 | 6.607196497 | 0.000264179 |
| Ybx3 | 1.248759411 | 0.000264179 |
| Palm2 | 4.818356271 | 0.000264767 |
| Avp | 3.841094618 | 0.000264767 |
| Actn1 | -1.635753173 | 0.000264767 |
| Pnoc | 3.220358327 | 0.000265911 |
| S1pr2 | -1.317533492 | 0.000266972 |
| Ybx1 | 0.989372456 | 0.000270449 |
| Slc38a3 | 2.683581234 | 0.00027068 |
| Synrg | -0.870685261 | 0.00027068 |
| Relb | -1.821099147 | 0.00027068 |
| Mocs1 | -1.557596633 | 0.000272801 |
| Siah2 | -1.358692556 | 0.000274552 |
| Twist2 | 6.591084004 | 0.000274872 |
| Ankrd49 | 1.364897105 | 0.000275832 |
| Csgalnact1 | 6.554147524 | 0.000276082 |
| 2310057M21Rik | -1.420052069 | 0.000278113 |
| Tet2 | -0.990879669 | 0.000278815 |
| Ltc4s | 6.625984432 | 0.000279228 |
| Apopt1 | 0.999287509 | 0.000279228 |
| Xylt2 | -1.954469536 | 0.000279228 |
| Fpr1 | -2.4272157 | 0.000279228 |
| Fermt2 | 1.264731411 | 0.000281093 |
| Rab5a | -0.781601336 | 0.000281093 |
| Plcd4 | 4.745433052 | 0.00028138 |
| Mrpl20 | 0.677933802 | 0.00028138 |
| Cers6 | -0.885713174 | 0.00028138 |
| Gm5431 | -2.328718517 | 0.00028138 |
| Mrps24 | 0.790991689 | 0.000281607 |
| Snapc2 | 1.244257429 | 0.000281714 |
| Cdkn3 | 1.923598854 | 0.000282132 |
| Lhx5 | 2.337688887 | 0.000283151 |
| Asb4 | 2.845017406 | 0.000283505 |
| Olfml2b | -1.606193657 | 0.000284941 |
| Prr3 | 1.100203043 | 0.000286054 |
| Polr1c | 0.955957203 | 0.000286443 |
| Tnfsf14 | -6.881890601 | 0.000286443 |
| Mrpl30 | 0.888782018 | 0.000287407 |
| Guk1 | 0.800896168 | 0.000287748 |
| Klf6 | -1.014378775 | 0.000287748 |
| Trip6 | 2.182142167 | 0.000290283 |
| Sft2d2 | -1.553355012 | 0.000290921 |
| Stab1 | -1.284275585 | 0.000291812 |
| Ppp3cc | -1.540474116 | 0.000293578 |
| Pebp1 | 0.612237476 | 0.000295563 |
| Slc7a7 | -1.655359944 | 0.000295726 |
| Cby1 | 0.881740765 | 0.00029676 |
| Prox1 | 1.590388747 | 0.000297625 |
| Mien1 | 0.685120599 | 0.000298836 |
| Fam155a | 1.736093046 | 0.000301848 |
| Angel1 | -1.289271374 | 0.000303628 |
| Bmp2 | 5.614154916 | 0.000304983 |
| Trim14 | -2.004024635 | 0.000304983 |
| Adam8 | -3.767690869 | 0.000304983 |
| Spa17 | 1.827896676 | 0.000306895 |
| Ccdc34 | 0.982540742 | 0.000306895 |
| Cd244 | -1.932523246 | 0.000306895 |
| Nutf2 | 0.745141887 | 0.000307266 |
| Zswim4 | -0.879818658 | 0.000309664 |
| Ttc9 | -1.617908931 | 0.000310498 |
| Slc44a2 | -1.121899138 | 0.000310801 |
| Grap | -1.569866353 | 0.000310801 |
| Chodl | 4.861043977 | 0.000313543 |
| Cd2ap | -1.306167063 | 0.000314256 |
| Gabrg1 | 3.095915815 | 0.000315726 |
| Il17d | 5.668913216 | 0.000317549 |
| Dhx58 | -2.430522197 | 0.000319727 |
| Hnrnpc | 0.584318611 | 0.000323305 |
| Abhd14a | 1.128474031 | 0.000324382 |
| Rangap1 | 0.775715262 | 0.000324709 |
| Slc22a6 | 3.718530765 | 0.00032644 |
| Itga11 | 3.541029856 | 0.00032644 |
| Ywhaq | 0.617828958 | 0.00032644 |
| Ctsb | -1.103726348 | 0.000326711 |
| Atrx | 0.700663058 | 0.00032683 |
| Creg1 | -1.09537272 | 0.000327323 |
| Trim25 | -1.714259283 | 0.000328342 |
| Tma7 | 0.8353481 | 0.000329867 |
| Znrf1 | -0.904985743 | 0.000332223 |
| 2310009B15Rik | 1.233368438 | 0.000333668 |
| Optn | -1.40507348 | 0.000333668 |
| Ppia | 0.632497581 | 0.000333821 |
| Lancl1 | 0.991576544 | 0.000335682 |
| Ilkap | 0.717743664 | 0.000335904 |
| Mafg | -1.071492777 | 0.000335904 |
| Uap1l1 | -1.403741509 | 0.000337674 |
| Rita1 | 1.478141685 | 0.000338608 |
| Clcf1 | 4.222496998 | 0.000340444 |
| Ranbp1 | 0.700213912 | 0.000340444 |
| Aars | -0.709858317 | 0.000340444 |
| Lrrk2 | -2.918929402 | 0.000341768 |
| Mrpl53 | 0.839099488 | 0.000342402 |
| Rabgef1 | -1.166699821 | 0.000342449 |
| Itgb5 | -1.14962747 | 0.000343438 |
| Mcl1 | -1.206189173 | 0.000345126 |
| Extl3 | -1.046463669 | 0.000346537 |
| C4b | -5.890429561 | 0.000346816 |
| A230046K03Rik | -0.92722198 | 0.000348343 |
| Xab2 | -0.876634862 | 0.000349235 |
| Eef1b2 | 0.724295337 | 0.000349491 |
| Mertk | -2.014956222 | 0.000349491 |
| 1500011K16Rik | 1.583858081 | 0.000350961 |
| Abr | -0.866465702 | 0.000351039 |
| Cbs | 3.391758819 | 0.000352421 |
| Slc6a13 | 2.622453526 | 0.000354368 |
| Aif1l | 1.08841621 | 0.000354368 |
| Pqlc2 | -1.553062853 | 0.000354368 |
| Ccdc71l | -2.396426972 | 0.000359202 |
| Nr2f2 | 1.557507278 | 0.000364572 |
| Ica1 | 1.118366159 | 0.000364572 |
| Slc35c1 | -2.225606724 | 0.000364572 |
| Akap10 | -1.137139174 | 0.000364925 |

| | | |
|---|---|---|
| Fcgr4 | -1.620258204 | 0.07904074 |
| Prr12 | 0.693128569 | 0.079041187 |
| Card10 | 2.673504471 | 0.079239852 |
| Naif1 | -0.918775789 | 0.079239852 |
| Hdac10 | -1.087228772 | 0.079239852 |
| Tcf7l2 | 1.814259974 | 0.079242517 |
| Kifc1 | -0.874695652 | 0.079263897 |
| Gcc2 | 0.539213699 | 0.079325499 |
| Ccdc163 | 0.958233805 | 0.079348827 |
| Stau2 | 0.43792554 | 0.079484945 |
| Scfd1 | -0.426738286 | 0.079484945 |
| Fkbp4 | 0.347905482 | 0.079532453 |
| Slc38a10 | -0.497212378 | 0.079532453 |
| Nipsnap1 | 0.425047156 | 0.079751598 |
| Bag4 | 0.707385708 | 0.079820455 |
| Chrna2 | -3.345553443 | 0.079820455 |
| Mvd | -0.571188287 | 0.079843019 |
| Hunk | -0.688768939 | 0.07991536 |
| Exd1 | -3.137880004 | 0.07991536 |
| Trpm3 | 0.856093382 | 0.079916752 |
| Eva1b | 1.585856428 | 0.079920759 |
| Dennd4b | -0.70598389 | 0.079920759 |
| 1700066M21Rik | -0.843593629 | 0.079957987 |
| Mospd3 | 0.481902944 | 0.080045516 |
| Syn1 | 0.530859703 | 0.080058177 |
| Mkrn1 | 0.328759087 | 0.080095106 |
| Iffo1 | 0.792055968 | 0.080129642 |
| Glg1 | -0.391630113 | 0.080129642 |
| Card11 | -2.387226276 | 0.080137629 |
| Cd200 | 0.481157202 | 0.080151616 |
| Rsph14 | 5.574045495 | 0.08017399 |
| Txndc17 | 0.488087035 | 0.080266775 |
| Zfp688 | 0.820595651 | 0.080272573 |
| Ccnt1 | -0.430755249 | 0.080272573 |
| Got2 | -0.338275813 | 0.080321238 |
| Xpc | -0.668897148 | 0.080321238 |
| Tmem132d | 3.775016221 | 0.080506589 |
| Rilp | 4.999186539 | 0.080550113 |
| Eif3a | 0.373958152 | 0.080689049 |
| Bcl2l1 | -0.475215954 | 0.080689049 |
| Pcdhgb7 | -1.109960443 | 0.080785069 |
| Insm2 | -2.646756436 | 0.080872173 |
| Supt20 | -0.481102913 | 0.080935501 |
| Ubash3a | -5.507327089 | 0.080935501 |
| Cpsf7 | 0.551594358 | 0.081017103 |
| Hs3st3b1 | -2.426961637 | 0.081034205 |
| Snrnp48 | 0.763057698 | 0.081034861 |
| Jak2 | -0.72028183 | 0.081077903 |
| Pacs2 | -0.649407707 | 0.081181099 |
| Eif3j1 | 0.502312288 | 0.081239817 |
| Adora2b | -2.803726911 | 0.08137514 |
| Stfa3 | -2.651339103 | 0.081380998 |
| 1110008P14Rik | 0.484995804 | 0.081448059 |
| Tas2r108 | 5.363123206 | 0.081489143 |
| Tctn3 | 0.818394339 | 0.081618138 |
| Trim17 | 1.548444376 | 0.081635744 |
| Nr1i2 | -5.519224063 | 0.081651956 |
| Gli2 | 0.988441015 | 0.081707009 |
| Rtn3 | 0.327283866 | 0.08171087 |
| Gadd45b | -1.134385441 | 0.08171087 |
| Ppp1r3d | -1.259735101 | 0.081788992 |
| Aspg | -5.820239178 | 0.081935374 |
| Deptor | 2.345199015 | 0.081985292 |
| BC094916 | -2.249173417 | 0.082116184 |
| Fam102b | 0.817538558 | 0.08215334 |
| Htt | -0.569411556 | 0.082233844 |
| Sart3 | -0.489443077 | 0.082359869 |
| Ssbp4 | -0.494781854 | 0.082383422 |
| Rab12 | -0.547142981 | 0.082383422 |
| Mfap1b | 0.881867617 | 0.082432221 |
| Izumo1r | -5.814027896 | 0.082461721 |
| Rhob | 0.538000502 | 0.082475437 |
| Limd1 | -0.684235629 | 0.08255548 |
| Clec2g | -2.770258839 | 0.082564997 |
| Myh14 | 1.861315589 | 0.082576311 |
| Zfp13 | 0.646618767 | 0.082592422 |
| Sms | 0.493375747 | 0.082609748 |
| Traf4 | 0.40945977 | 0.082617494 |
| Mcm4 | -0.476854911 | 0.082617494 |
| Ints8 | -0.558707807 | 0.082617494 |
| Lin54 | -0.63593586 | 0.082617494 |
| Gm28043 | -2.121673076 | 0.082617494 |
| Lingo4 | -2.403362433 | 0.082617494 |
| Rara | -0.80436964 | 0.082680148 |
| 3110009E18Rik | 1.10963546 | 0.082711196 |
| Alox15 | -4.752492495 | 0.082711196 |
| Abcb8 | -0.466316303 | 0.082717758 |
| Ccdc150 | -2.538622722 | 0.082860399 |
| Dctn1 | -0.380238238 | 0.082936845 |
| Akr7a5 | 0.537109237 | 0.083116286 |
| Alg10b | -0.741532945 | 0.083139183 |
| Gprin1 | -0.666754735 | 0.083214356 |
| Wdr61 | 0.446314372 | 0.083287225 |
| Rai14 | -0.660684517 | 0.083287225 |
| Lrp1 | -0.354699724 | 0.083333997 |
| Adrbk1 | -0.504377933 | 0.083471729 |
| Itgb4 | 2.618334678 | 0.083474977 |
| Clcc1 | 0.538401779 | 0.083486864 |
| Gng5 | 0.45478146 | 0.083534258 |
| R3hcc1 | 0.434914977 | 0.083534258 |
| Apba3 | -0.889456101 | 0.083534258 |
| Magel2 | 1.26434386 | 0.083574764 |
| Cenpn | 1.0733739 | 0.08359917 |
| Hn1l | -0.484444034 | 0.08359917 |
| Unc13d | -1.506409162 | 0.08359917 |
| Npy2r | 4.996878228 | 0.083641049 |
| Kir3dl2 | 5.335175967 | 0.083649277 |
| Col9a3 | 1.167173326 | 0.08369821 |
| Ccpg1os | 0.886264418 | 0.083699927 |
| Col4a3bp | -0.678771876 | 0.083887858 |
| 2610528A11Rik | 4.472225562 | 0.083921655 |
| Brca2 | 0.789818096 | 0.083921655 |
| Ubfd1 | -0.460665457 | 0.083921655 |
| Tlk1 | 0.540937887 | 0.084055037 |
| Cfap69 | 1.396975319 | 0.084064193 |
| Taf4b | -1.793678128 | 0.084158356 |
| Cr1l | -0.469408514 | 0.084213827 |
| Cfap45 | 1.711470907 | 0.084455445 |
| Hnrnpul2 | -0.526633921 | 0.084607953 |

| | | |
|---|---|---|
| Igsf23 | 3.64194826 | 0.379219486 |
| Sspo | -1.554115563 | 0.379437946 |
| Tecpr2 | -0.493035189 | 0.37957588 |
| Ccdc102a | -0.500747595 | 0.379987125 |
| Ooep | 1.7730275 | 0.38000941 |
| Atp10a | -1.42411072 | 0.380013262 |
| Gnpda2 | 0.344631438 | 0.380298784 |
| Gm10451 | 3.643836433 | 0.380336819 |
| Il10ra | -0.562652381 | 0.380423464 |
| Oca2 | 2.959280047 | 0.380652404 |
| Eomes | 0.337436498 | 0.380652404 |
| Gzmk | 3.962826161 | 0.380775533 |
| Hsd11b2 | -1.231867855 | 0.380885347 |
| Ccl6 | 0.56649767 | 0.380993421 |
| Frrs1l | 0.53225518 | 0.380993421 |
| Rfk | 0.236060541 | 0.381036055 |
| Acmsd | 0.720231487 | 0.381094482 |
| Ifit3 | -1.686432994 | 0.381131161 |
| Lamp5 | -1.264826451 | 0.381204204 |
| Dazap1 | -0.282794247 | 0.381600194 |
| Psd2 | 0.477442127 | 0.381718153 |
| Myh15 | 3.677033212 | 0.381900219 |
| Vav3 | -0.696618792 | 0.382030884 |
| Gm17571 | 2.178225264 | 0.382089193 |
| Prpf40a | 0.201201157 | 0.382095454 |
| Nde1 | 0.422901155 | 0.382106415 |
| Ndufaf7 | 0.33430768 | 0.382106415 |
| Gm11808 | 0.31015659 | 0.382157999 |
| Tbccd1 | 0.454853649 | 0.382278025 |
| Ar | 3.669185917 | 0.382437767 |
| Amigo2 | -0.954436048 | 0.382640579 |
| Swt1 | 0.319322922 | 0.382646603 |
| Gdap1l1 | 0.257371294 | 0.382646603 |
| Zfp563 | 0.425398777 | 0.382733501 |
| Hbp1 | -0.219689994 | 0.382908853 |
| Pnpla2 | -0.299948061 | 0.382994715 |
| Ftsj2 | -0.439983692 | 0.382994715 |
| Mgat5b | -0.344743692 | 0.383109478 |
| Olfr1392 | -4.449998796 | 0.383109478 |
| Brsk1 | -0.696690877 | 0.383118755 |
| E4f1 | 0.338933074 | 0.383153232 |
| Hsd17b14 | -1.563687004 | 0.383194755 |
| Tceal7 | 2.086879951 | 0.383338648 |
| Fbxo24 | 2.92897903 | 0.383574251 |
| Klhl22 | -0.268168099 | 0.383574251 |
| Srsf5 | -0.186813699 | 0.383579694 |
| Khnyn | -0.627096262 | 0.383579694 |
| Glrx5 | 0.249074989 | 0.383951023 |
| Gabre | 3.648373164 | 0.384404393 |
| 2610044O15Rik8 | 0.588382638 | 0.384404393 |
| Lrrc8a | 0.232436104 | 0.384404393 |
| Pdf | -3.01183226 | 0.384545396 |
| Lpcat1 | -0.256004582 | 0.384699326 |
| Cry1 | 0.416569975 | 0.38477215 |
| Smarcad1 | 0.256623978 | 0.384871357 |
| Comtd1 | -0.397913298 | 0.384975465 |
| Pth2r | 3.923886657 | 0.384976149 |
| Fam83c | 2.712893174 | 0.385147089 |
| Gon4l | 0.378257878 | 0.385147089 |
| Synpo | -1.158208769 | 0.385156729 |
| Tssc4 | 0.341798373 | 0.385163003 |
| Msh3 | 0.394924347 | 0.38518227 |
| Ostc | 0.214005405 | 0.385540875 |
| Taco1 | -0.513800241 | 0.385704153 |
| Proca1 | -0.827704228 | 0.385744166 |
| Inadl | 0.634610912 | 0.385826091 |
| Vmac | 0.328629821 | 0.386210011 |
| E2f6 | -0.325104525 | 0.38623017 |
| Mppe1 | -0.657269761 | 0.386283467 |
| Pno1 | 0.316297136 | 0.386316188 |
| Gyg | -0.280846502 | 0.386316188 |
| Wdr76 | -0.3831438 | 0.386316188 |
| Epb41l4b | -2.350498326 | 0.386316188 |
| Ubl3 | -0.240663221 | 0.38656416 |
| Katnal2 | -0.497664618 | 0.38656416 |
| Celf5 | -0.343456013 | 0.386572516 |
| Itga10 | -1.659947944 | 0.386663549 |
| Zxdb | 1.249433838 | 0.386765825 |
| Zgpat | -0.300952623 | 0.386765825 |
| H2-M5 | 1.353823422 | 0.387078538 |
| Zfp341 | 0.822591959 | 0.387297027 |
| Mag | -0.811922861 | 0.387297027 |
| Fhad1 | -0.818646692 | 0.387297027 |
| Fancf | 0.840614722 | 0.38738464 |
| Tbxa2r | 1.825529623 | 0.388091326 |
| Dnlz | 0.243284899 | 0.388091326 |
| 1700061G19Rik | -1.305783322 | 0.388296531 |
| Ccdc13 | -1.407840417 | 0.388309381 |
| Hist1h3i | -0.634586112 | 0.388341552 |
| Srgap2 | -0.231774031 | 0.388341931 |
| Mapk15 | -2.678381108 | 0.388396917 |
| Abracl | 0.218366349 | 0.388551346 |
| Hiat1 | -0.382373877 | 0.388551346 |
| Slc45a1 | -0.472000335 | 0.388551346 |
| Hspd1 | 0.186951712 | 0.388600713 |
| Btg2 | 0.308926198 | 0.388681997 |
| Fgfrl1 | -1.098129174 | 0.388829654 |
| Rbm15 | -0.204073611 | 0.389427702 |
| Rnf151 | 3.958703041 | 0.389557518 |
| Rbfox2 | -0.212156213 | 0.389557518 |
| Tfb1m | -0.54480942 | 0.389557518 |
| Brms1l | 0.414992316 | 0.389664798 |
| Adarb2 | -0.661022821 | 0.389664798 |
| Zswim1 | -0.452509945 | 0.389967853 |
| Rgs22 | -1.728861278 | 0.390104462 |
| Oacyl | -1.928343641 | 0.390191424 |
| Kif19a | -1.149069163 | 0.390373014 |
| A530016L24Rik | 3.902845338 | 0.390435576 |
| Ccnb1ip1 | 3.902845338 | 0.390435576 |
| Sbk3 | 3.902845338 | 0.390435576 |
| Gpr173 | -0.492052775 | 0.390435576 |
| Gdap1 | 0.293891446 | 0.390521912 |
| Nln | 0.276532466 | 0.390521912 |
| Hk1 | -0.243261772 | 0.390521912 |
| 1700030J22Rik | 0.818996709 | 0.390660041 |
| Stat5b | -0.549311241 | 0.390660041 |
| Alg1 | -0.404445042 | 0.390765262 |
| Pip5k1b | 1.190326517 | 0.390840463 |
| Fam13c | 0.448085964 | 0.391001755 |

| | | |
|---|---|---|
| Ror2 | -0.325150317 | 0.766822146 |
| Morn3 | 0.384548316 | 0.767136854 |
| 1810055G02Ri | 0.148105784 | 0.767279946 |
| BC049715 | 0.46314802 | 0.767884175 |
| Irf2bp2 | 0.106904334 | 0.767884175 |
| Exosc4 | 0.088862012 | 0.767884175 |
| Gm16314 | -0.307762037 | 0.767931998 |
| Trub1 | -0.255395893 | 0.768399183 |
| Ifitm7 | -0.423881568 | 0.768721831 |
| 6820408C15Ri | -0.58549918 | 0.768721831 |
| Ift46 | 0.103740501 | 0.768745561 |
| Elac2 | -0.206616198 | 0.768970387 |
| Rab35 | 0.158018154 | 0.769068093 |
| Gtf2h1 | -0.095963507 | 0.769068093 |
| Cntnap5b | -0.385738949 | 0.769472055 |
| Mapkap1 | 0.102128544 | 0.769750692 |
| Lonp2 | 0.086169569 | 0.769750692 |
| Ccdc82 | -0.133356792 | 0.769750692 |
| A830018L16Ri | -0.386170769 | 0.769775103 |
| Gm17359 | 0.718625834 | 0.769802282 |
| Cep44 | 0.127171032 | 0.769802282 |
| Dazap2 | 0.091471643 | 0.769888771 |
| Tec | 0.239070673 | 0.770111148 |
| Arg2 | 0.232134874 | 0.770111148 |
| Supt6 | -0.093198751 | 0.770111148 |
| Aasdh | -0.140479193 | 0.770111148 |
| Gm3944 | 0.38623115 | 0.770386807 |
| Krit1 | 0.093669198 | 0.770386807 |
| 5730522E02Ril | 0.155660301 | 0.770472071 |
| Clcn1 | -0.41572344 | 0.770472071 |
| Grifin | -0.954665523 | 0.770472071 |
| Slc39a5 | 0.688093193 | 0.770549312 |
| Zfp1 | 0.15593906 | 0.770549312 |
| Phtf1 | 0.129132515 | 0.770549312 |
| Srrm1 | 0.067971227 | 0.770549312 |
| Gsto1 | 0.105111259 | 0.770709124 |
| Zfp738 | -0.211135361 | 0.770709124 |
| Asb3 | -0.150139668 | 0.770971163 |
| Fitm2 | 0.236602066 | 0.771032327 |
| Clec18a | 0.506241136 | 0.771238906 |
| Gm2974 | 0.322422701 | 0.771238906 |
| Anapc1 | 0.085154502 | 0.771238906 |
| Dmrta1 | 1.026340707 | 0.77127644 |
| Ttc30a2 | -0.521664259 | 0.771695669 |
| Rhot2 | -0.216908523 | 0.771747005 |
| Zkscan3 | -0.081963701 | 0.771848365 |
| 4930595D18Ri | -0.342541404 | 0.771848365 |
| Crygn | 1.219238254 | 0.771890221 |
| Xpnpep1 | -0.088969321 | 0.771914468 |
| Stra6l | -0.373292017 | 0.772068802 |
| Slc40a1 | 0.262629058 | 0.772187263 |
| Coq5 | 0.092460921 | 0.772187263 |
| Uchl5 | 0.08723162 | 0.772187263 |
| Gm4737 | -0.111993715 | 0.772187263 |
| Tsga10 | -0.114785198 | 0.77236531 |
| Tmx3 | -0.097130647 | 0.772737203 |
| Bcat2 | -0.318456154 | 0.772776696 |
| Strn3 | 0.093118864 | 0.772912857 |
| Rab42 | -0.895683359 | 0.772949017 |
| Tirap | -0.217190657 | 0.772953536 |
| Dars2 | 0.148599369 | 0.773075886 |
| B3gat3 | -0.122775386 | 0.773075886 |
| Igf2bp2 | -0.157461859 | 0.773105304 |
| Oip5 | 0.229643842 | 0.7731174 |
| Chmp2b | 0.133911719 | 0.77333313 |
| Dock4 | 0.122926061 | 0.77333313 |
| Tmem8b | 0.161982428 | 0.773412272 |
| Cpsf3l | -0.097893408 | 0.773412272 |
| Hyal3 | -0.672477184 | 0.773412272 |
| Zc3hc1 | 0.123718735 | 0.773552539 |
| Actl6a | 0.087302748 | 0.773552539 |
| Tsc2 | -0.12348411 | 0.773552539 |
| Vps39 | -0.138997857 | 0.773552539 |
| Jag1 | -0.241577066 | 0.773552539 |
| Zbtb6 | -0.122320761 | 0.773653316 |
| H2-K1 | -0.146071638 | 0.773794967 |
| Egln1 | 0.174499172 | 0.773952256 |
| Bcam | 0.462342577 | 0.774083654 |
| Zfp59 | 0.258487311 | 0.774085519 |
| Ch25h | -0.3884859 | 0.774245356 |
| Acrbp | -0.247707007 | 0.774457506 |
| Kif17 | -0.678432628 | 0.774457506 |
| Ybey | 0.237979243 | 0.774608854 |
| Hells | 0.137798138 | 0.774608854 |
| Kif23 | 0.134267448 | 0.774608854 |
| Whamm | 0.26783279 | 0.774631566 |
| Fzd1 | -0.149588896 | 0.774646896 |
| Gm9949 | -0.417451482 | 0.774646896 |
| Slc39a1 | 0.119362098 | 0.774700133 |
| Unc119 | 0.117235033 | 0.774700133 |
| Rfc2 | 0.107132324 | 0.774700133 |
| Rab40c | -0.134954763 | 0.774700133 |
| Pbld2 | -0.470272996 | 0.774700133 |
| E030037K01Ri | 0.392179213 | 0.774790728 |
| Gm17430 | 0.950330808 | 0.774868872 |
| Nup93 | 0.109770443 | 0.775074193 |
| Fam163a | 0.277191717 | 0.775116759 |
| Prkab2 | -0.156405152 | 0.775163702 |
| Dlgap5 | 0.154225461 | 0.775178925 |
| Zmynd11 | 0.081088829 | 0.77522276 |
| 4933434E20Ril | -0.074957786 | 0.77522276 |
| Ninl | 0.23451743 | 0.775272913 |
| Aip | -0.094463249 | 0.775272913 |
| Dgcr8 | -0.132466146 | 0.775272913 |
| Ccdc40 | -0.243350102 | 0.775272913 |
| Tmem202 | -0.896532122 | 0.775272913 |
| Med8 | -0.077433578 | 0.775650662 |
| Rasl10a | -1.041598613 | 0.775650662 |
| Clec1a | 1.267111917 | 0.775844231 |
| Hist1h2bn | 0.558981498 | 0.775864744 |
| Ppp1r3g | -0.558709644 | 0.776165986 |
| Pcdhga4 | 0.217252397 | 0.776278492 |
| Bmx | -1.480449602 | 0.776439565 |
| Tnni2 | 0.250364808 | 0.776445965 |
| Kdm1b | 0.145305631 | 0.776445965 |
| Cdca4 | -0.089033326 | 0.776445965 |
| Lsmem1 | -0.683269386 | 0.776445965 |
| Naalad2 | 0.674201005 | 0.776458893 |
| Papola | -0.076354676 | 0.777025567 |

| | | |
|---|---|---|
| Rragc | -1.360258191 | 0.000368926 |
| Trip11 | 1.043864707 | 0.00037162 |
| Lgmn | -1.087488112 | 0.000371905 |
| Rpl37a | 0.650945233 | 0.000374489 |
| Tbx18 | 6.118337878 | 0.000374513 |
| Il13ra1 | -1.860398581 | 0.000376226 |
| Fam169b | -7.360710097 | 0.000379023 |
| Pacsin1 | 1.425406274 | 0.000379814 |
| Vps11 | -0.928543255 | 0.00038033 |
| Rpsa | 0.72709002 | 0.000380709 |
| Akap5 | 1.308407037 | 0.000382396 |
| Cenpa | 1.036334265 | 0.000382396 |
| Gm9733 | -3.161678326 | 0.000384679 |
| Tmem8 | -2.748639198 | 0.000387606 |
| Pprc1 | -1.511865643 | 0.000387625 |
| Camk2n1 | 1.317824048 | 0.000392236 |
| Renbp | -1.101467431 | 0.000394113 |
| Prkra | 0.816604505 | 0.000395191 |
| Tial1 | 0.987561141 | 0.000395381 |
| Zfp11 | -1.924815257 | 0.000395632 |
| Mageh1 | 0.833627728 | 0.000396928 |
| Ict1 | 0.773706511 | 0.000396928 |
| Slitrk5 | -1.427175445 | 0.00039698 |
| Pdcd4 | 0.900571135 | 0.000402756 |
| Otulin | -1.010892944 | 0.000405267 |
| Tfap2c | 2.239140435 | 0.000405403 |
| Clstn2 | -1.568730442 | 0.000405427 |
| Ddx3x | 0.849810068 | 0.000410233 |
| Cdc42ep1 | 2.381287288 | 0.000410766 |
| Cep70 | 1.253824956 | 0.000411126 |
| Pced1a | -1.078061028 | 0.000411308 |
| Apbb1ip | -1.235750455 | 0.000412688 |
| Rassf9 | 6.50693449 | 0.000415329 |
| Ythdf1 | -1.170120491 | 0.000415734 |
| Igfbp4 | 0.816236865 | 0.000423139 |
| Cfap43 | -1.412440255 | 0.000425468 |
| Kitl | 1.678293372 | 0.000427063 |
| Fbxo44 | 1.27458378 | 0.000429305 |
| Tekt1 | 3.213736712 | 0.000429796 |
| Gpr18 | -2.095623471 | 0.000432453 |
| Rps27l | 0.793817386 | 0.000433217 |
| Eci2 | 0.713268235 | 0.000433334 |
| Hmox1 | -2.878683386 | 0.000433557 |
| Apold1 | 6.744466364 | 0.00043495 |
| Elavl4 | 1.027055393 | 0.000440206 |
| Pomp | 0.68604624 | 0.000441469 |
| Sumo2 | 0.576314237 | 0.000444602 |
| Piezo2 | 6.044256258 | 0.000444716 |
| Asf1a | 0.72340539 | 0.000444716 |
| Atp5l | 0.706992572 | 0.000448042 |
| Ndufb10 | 0.63222632 | 0.000448866 |
| Grm8 | 3.763308611 | 0.0004491 |
| Slc44a1 | -1.27757015 | 0.0004491 |
| Trpm2 | -2.22042816 | 0.000450192 |
| Apoo | 0.748229707 | 0.000451241 |
| Hapln3 | 6.499805617 | 0.000451434 |
| Coa5 | -0.818488382 | 0.000452455 |
| Slc22a4 | -3.557110891 | 0.000452455 |
| Erbb2 | 3.261895074 | 0.000452929 |
| Vcam1 | 1.723590909 | 0.000454467 |
| Manba | -2.366813946 | 0.000455384 |
| Carkd | 0.743255906 | 0.000456072 |
| Mical2 | -1.393406004 | 0.000456088 |
| Copa | -0.667814546 | 0.0004569 |
| Selk | 0.595044786 | 0.000461026 |
| Atp2b4 | 1.374424595 | 0.000461964 |
| Wdtc1 | -1.054714118 | 0.000465497 |
| Nasp | 0.679100565 | 0.000467175 |
| Nefl | 2.098508993 | 0.000468394 |
| Mrpl27 | 0.745249674 | 0.000472459 |
| Tmem106a | -1.865968594 | 0.000477361 |
| Lin28a | -7.309065229 | 0.000477361 |
| Zfp819 | -3.662968853 | 0.000480847 |
| Ptpra | -0.69365418 | 0.000481809 |
| Selplg | -1.667663264 | 0.000485468 |
| Arg1 | -3.861492599 | 0.000486796 |
| Gm20939 | -7.211355556 | 0.000486796 |
| Gmip | -1.231996625 | 0.000489843 |
| Ell2 | -2.02667933 | 0.000490657 |
| Iqsec3 | 1.899686622 | 0.000494659 |
| Layn | 6.464217496 | 0.000497767 |
| Slc11a2 | -1.104994324 | 0.000498212 |
| Rab7 | -0.714525109 | 0.000502756 |
| Pacsin2 | -1.037867373 | 0.000503687 |
| Fermt3 | -1.411096211 | 0.000506261 |
| Sall3 | 1.893924092 | 0.000507805 |
| Arxes2 | 1.090860367 | 0.000509026 |
| Adrb2 | -1.281038603 | 0.000509026 |
| Sirpb1b | -2.957397466 | 0.000509664 |
| Alkbh6 | 0.817357511 | 0.000509868 |
| Lcp2 | -1.35784457 | 0.000509868 |
| Notch1 | 0.970018592 | 0.000510119 |
| Oaz2 | 0.77677717 | 0.000510119 |
| Galntl6 | 3.296862183 | 0.000512394 |
| Ifnlr1 | -6.907549738 | 0.000513239 |
| Nol8 | 1.25087064 | 0.000514331 |
| Usp28 | -1.016376478 | 0.000514502 |
| Mgrn1 | -0.705978595 | 0.000515133 |
| Cab39 | -0.858788429 | 0.000515133 |
| Rnf103 | -1.327913379 | 0.00051656 |
| Ehd4 | -1.161295695 | 0.000516988 |
| Cdk13 | -0.940840334 | 0.000531516 |
| Zfand5 | -0.78095838 | 0.000534409 |
| Tmem256 | 0.768367125 | 0.000538853 |
| Rem2 | 0.993655131 | 0.000542413 |
| Cpa3 | -7.182886962 | 0.000544494 |
| Gpr35 | -1.905185135 | 0.000544778 |
| Pax3 | 6.597058911 | 0.000551363 |
| Gstp1 | 0.684571571 | 0.000552232 |
| Myo6 | 1.393370441 | 0.000552732 |
| Gm11627 | 3.04040338 | 0.000554322 |
| Srsf2 | -0.690377376 | 0.000554856 |
| Sstr1 | 6.166282488 | 0.000555355 |
| Lgals1 | -1.111956559 | 0.000555684 |
| Hnrnpab | -0.731632172 | 0.000558933 |
| Dph3 | 0.858367043 | 0.000560028 |
| Amdhd2 | -1.178238941 | 0.000561155 |
| Ephx2 | 6.484206328 | 0.000562824 |
| Lzic | 1.002767271 | 0.000564118 |

| | | |
|---|---|---|
| Zmynd19 | -0.578961343 | 0.084947005 |
| Casp12 | 4.32521912 | 0.084964898 |
| Trim36 | -0.659500922 | 0.085081087 |
| Zfp687 | -0.576610152 | 0.085086778 |
| Pop4 | 0.827596794 | 0.085159906 |
| Crym | 0.781426722 | 0.085203473 |
| Ghdc | -1.126072537 | 0.085203473 |
| Mau2 | -0.411609151 | 0.085225554 |
| Trps1 | -0.686273427 | 0.085541082 |
| Uhrf1bp1 | -0.952241375 | 0.085555281 |
| Ostm1 | -0.519494462 | 0.085570532 |
| Gsx2 | 1.679177971 | 0.085660054 |
| Mterf1a | 1.353585023 | 0.085660054 |
| C030039L03Rik | 1.021764256 | 0.085660054 |
| Marf1 | 0.390607558 | 0.085710956 |
| Daam1 | 0.443965484 | 0.085748276 |
| Zbtb26 | -0.850610458 | 0.085812947 |
| Hist1h4i | -1.044374649 | 0.085822188 |
| Rab39b | 0.627662709 | 0.085859586 |
| Agfg2 | -0.746150015 | 0.085864359 |
| Fam57b | 0.707847101 | 0.085919272 |
| Nxpe4 | 3.099994437 | 0.0859334 |
| Plxnb3 | -1.461318699 | 0.0859334 |
| Ganc | -0.83613505 | 0.085943835 |
| Mrps18c | 0.472531764 | 0.085955059 |
| Cd33 | -1.670897104 | 0.086016355 |
| Sult4a1 | 0.550172537 | 0.086036837 |
| Mns1 | 1.607518626 | 0.086087263 |
| Tulp3 | 0.922714916 | 0.086092279 |
| Ranbp17 | -0.833899211 | 0.086167864 |
| Zfyve28 | -1.253777518 | 0.086167864 |
| Slc25a19 | -0.601603991 | 0.086242956 |
| Tacc1 | -0.466926883 | 0.086259004 |
| Pdxdc1 | -0.518265856 | 0.086259004 |
| Zfp292 | 0.400289332 | 0.086302567 |
| 9130230L23Rik | -2.052876234 | 0.086302567 |
| Api5 | -0.36458272 | 0.086420857 |
| L1cam | -0.714126996 | 0.086420857 |
| Snapin | 0.591733456 | 0.086429477 |
| Alkbh3 | -0.60436459 | 0.086429477 |
| Gm12166 | 0.794307049 | 0.086460679 |
| F3 | 1.348233464 | 0.08651751 |
| Gm10094 | 0.540026292 | 0.086535766 |
| Lrriq3 | 4.883892231 | 0.086570004 |
| Zyg11b | -0.435647507 | 0.086645437 |
| Nudt19 | 0.537941053 | 0.086691093 |
| 1700019D03Rik | 2.285818329 | 0.086744107 |
| Usf2 | 0.644181358 | 0.087051854 |
| Gbp10 | -3.196411556 | 0.087081515 |
| Ddx21 | 0.495739071 | 0.087183383 |
| Tonsl | 0.339405954 | 0.087204727 |
| Fbxo16 | 0.815191313 | 0.087236921 |
| Plpp4 | 3.131775392 | 0.087374006 |
| Arhgef40 | 0.658836951 | 0.08751367 |
| Cbx2 | -0.71926411 | 0.087571525 |
| Usp21 | 0.456207348 | 0.087647215 |
| Smg5 | -0.454208789 | 0.087647215 |
| Tmem114 | -5.786558815 | 0.087692947 |
| Ptgir | -4.126106281 | 0.08776634 |
| Hdlbp | -0.453491189 | 0.087790635 |
| Acd | 0.422811135 | 0.087899189 |
| Aph1b | -0.772798873 | 0.087899189 |
| Dab1 | -0.679713107 | 0.087905316 |
| Rnf180 | 0.73206009 | 0.087983915 |
| Prss57 | 5.393091224 | 0.08800694 |
| Kcne3 | 4.960711572 | 0.08812877 |
| Sppl2b | -0.561767986 | 0.088131505 |
| Pnpt1 | -0.688729079 | 0.088131505 |
| Tdrd12 | 4.848252143 | 0.08814463 |
| Dok4 | 0.476610158 | 0.08814463 |
| Fam3b | -5.776651272 | 0.088147295 |
| D130040H23Rik | 1.750674687 | 0.088158712 |
| Flad1 | -0.599150044 | 0.088158712 |
| Zyx | -0.669901617 | 0.088158712 |
| Nr1h3 | -1.660259925 | 0.088199044 |
| Bmp7 | 1.39264441 | 0.088233036 |
| Pomt2 | -0.883373783 | 0.088242561 |
| Acbd7 | 4.836890561 | 0.088312177 |
| Rnmtl1 | 0.913139155 | 0.088312177 |
| 1700037H04Rik | 0.435546109 | 0.088322625 |
| Sort1 | -0.600472014 | 0.088398905 |
| Ggh | -0.59765212 | 0.088578105 |
| Clasp1 | -0.403925721 | 0.088649401 |
| Cd84 | -1.0712693 | 0.088649401 |
| Mapk4 | -0.952904882 | 0.088868279 |
| 5-Mar | -0.400110538 | 0.088942736 |
| Impact | -0.454646516 | 0.088942921 |
| Gbp2 | -1.425306723 | 0.0890103 |
| Inpp1 | 0.831732233 | 0.08903461 |
| Pink1 | 0.385478973 | 0.089036194 |
| Ubxn8 | 0.916512927 | 0.089138377 |
| Edc4 | -0.715783083 | 0.089157238 |
| Pdzph1 | 1.581213947 | 0.089178729 |
| Ascc3 | -0.600388395 | 0.089285254 |
| Hrct1 | 5.111929035 | 0.089352582 |
| Spats1 | 4.154184406 | 0.089421187 |
| Jagn1 | 0.538974749 | 0.089421187 |
| Vav2 | -0.573034755 | 0.089421187 |
| Erich3 | 3.96592794 | 0.089422729 |
| Selp | -5.760051413 | 0.089422729 |
| Gtf2h3 | 0.618897763 | 0.089559184 |
| Mrrf | 0.613826657 | 0.089564467 |
| Nt5dc2 | 0.590933898 | 0.089564467 |
| Fdxr | 0.764304559 | 0.089566524 |
| Sec24a | -0.631965964 | 0.089566524 |
| Dach1 | 1.113852256 | 0.089650121 |
| Alg14 | 0.795802902 | 0.089697523 |
| Bmp10 | 5.034066916 | 0.089741221 |
| Mgat4c | -1.084906332 | 0.089744015 |
| Epdr1 | 1.070340047 | 0.089818405 |
| Kcnj13 | 2.666790815 | 0.089878973 |
| Uba1 | -0.308947003 | 0.0900386 |
| Cnep1r1 | 0.604506288 | 0.09027422 |
| Copg2 | 0.390506087 | 0.090378123 |
| Casd1 | 0.60910344 | 0.090644677 |
| Clcn6 | -0.618730323 | 0.090644677 |
| Abhd5 | -0.671457383 | 0.090644677 |
| Snx13 | -0.636258618 | 0.090678015 |
| Top3a | -1.133909092 | 0.090719302 |

| | | |
|---|---|---|
| Myl2 | -1.997968919 | 0.391014513 |
| Ppp1ca | -0.227732283 | 0.391082031 |
| Ppp2r3d | -0.383951276 | 0.391139944 |
| Cyp4f37 | 3.212981953 | 0.39116819 |
| Zfp365 | 0.993935115 | 0.39116819 |
| Agpat4 | 0.354317965 | 0.39116819 |
| Igfbpl1 | 0.260154701 | 0.39116819 |
| Nkain3 | -0.400232829 | 0.39116819 |
| Gm10073 | 0.627375077 | 0.391328228 |
| Stambp | 0.302004393 | 0.391392543 |
| D430042O09Rik | -0.389628233 | 0.391500777 |
| Lyrm1 | -0.387161117 | 0.391549942 |
| A930018P22Rik | -4.440514289 | 0.391549942 |
| Fcmr | -4.440514289 | 0.391549942 |
| Rfx8 | -4.440514289 | 0.391549942 |
| Fam179b | -0.363348659 | 0.391796858 |
| Epc1 | 0.258689451 | 0.391857825 |
| Pde6h | -2.076782856 | 0.391970901 |
| Mxra7 | 0.999541962 | 0.392224728 |
| Slc39a3 | -0.336081954 | 0.392584279 |
| Cpsf4 | -0.259860187 | 0.392618645 |
| Egfl6 | 0.939831304 | 0.392624026 |
| Ptprq | -2.599797375 | 0.392756116 |
| Sh3gl2 | -0.398729545 | 0.392820187 |
| Npc2 | 0.222819609 | 0.392900807 |
| Wdr47 | -0.287227951 | 0.392900807 |
| Poc1b | -0.32274726 | 0.392900807 |
| Fastk | 0.222675928 | 0.39292293 |
| Blzf1 | -0.350776001 | 0.39292293 |
| Ppp2r5a | -0.438130981 | 0.392973088 |
| Adra1a | 0.87625108 | 0.393101719 |
| Mgll | 0.517953981 | 0.393106325 |
| Mrps9 | 0.287797006 | 0.393106325 |
| Zfp454 | 0.52231873 | 0.393398005 |
| Zfp52 | -0.813592936 | 0.393404693 |
| Rab39 | -1.017714289 | 0.393643028 |
| Ap3b1 | -0.238938559 | 0.393768828 |
| Cxxc5 | -0.365546097 | 0.393768828 |
| Pard6b | -0.732104569 | 0.393768828 |
| Tceanc2 | 0.274919341 | 0.394141474 |
| Adamtsl2 | 0.896950651 | 0.394290113 |
| Thap4 | -0.248355754 | 0.394309082 |
| Lysmd4 | -0.29113629 | 0.394309082 |
| Gpr75 | 1.249447908 | 0.394315461 |
| Auts2 | -0.268441564 | 0.394402367 |
| Grp | 1.698175946 | 0.394499144 |
| Lrp12 | -0.396285632 | 0.394499144 |
| Cfap100 | 1.023859566 | 0.39455943 |
| Ggt7 | -0.377774283 | 0.394678491 |
| Npnt | 0.812767182 | 0.394718994 |
| Echdc1 | -0.495567917 | 0.394743165 |
| Gm9312 | 1.817365308 | 0.395018676 |
| Gprc5c | 1.223027689 | 0.395074932 |
| Neurl1b | 0.727003182 | 0.395074932 |
| Letmd1 | 0.300113483 | 0.39542958 |
| Tmem120a | 0.227536833 | 0.39542958 |
| Gm8281 | 1.385273645 | 0.395567675 |
| Dhx9 | 0.25293009 | 0.395567675 |
| Cmip | 0.241468327 | 0.395567675 |
| Ubxn6 | 0.235357607 | 0.395567675 |
| Gm8444 | -4.424487639 | 0.395567675 |
| Nodal | -4.424487639 | 0.395567675 |
| Creb3l3 | 1.67261321 | 0.395867886 |
| Dnah7b | 1.179574983 | 0.395867886 |
| Agbl2 | 2.965690445 | 0.396419478 |
| Polr3a | -0.304706279 | 0.396419478 |
| Glrb | -0.479591747 | 0.396452799 |
| Tbx6 | 2.49667702 | 0.396471016 |
| Chkb | 0.321033816 | 0.396626028 |
| Tbkbp1 | 0.526168094 | 0.396742676 |
| Cartpt | 2.908103794 | 0.396802477 |
| Iqgap3 | -0.703766158 | 0.39701996 |
| Clec11a | 0.752398733 | 0.397175622 |
| Cep164 | 0.356097873 | 0.397185357 |
| Kcnq3 | -0.407145506 | 0.397213779 |
| Cyb5d2 | -0.630804966 | 0.397329309 |
| 4930507D05Rik | 1.96381901 | 0.397469724 |
| Sp4 | 0.456886197 | 0.39750972 |
| Ccdc85c | -1.466496201 | 0.397738924 |
| Ciart | 0.515670564 | 0.397887968 |
| Inhbb | -0.512414184 | 0.397967241 |
| Pex5 | -0.271054774 | 0.398069911 |
| Sox4 | -0.235516601 | 0.398228542 |
| Cd1d1 | 0.468476202 | 0.398338888 |
| Rab8a | -0.213496228 | 0.398338888 |
| Dcbld2 | -0.514571546 | 0.398474496 |
| Crbn | -0.283118245 | 0.399082277 |
| Fut2 | 3.534521206 | 0.399294599 |
| Zfp202 | 0.748884067 | 0.399302177 |
| Kmt2c | -0.271483735 | 0.399302177 |
| Atp2c1 | -0.362284399 | 0.399302177 |
| Tspan32 | -0.548428607 | 0.399314167 |
| Dnaaf2 | -0.327971702 | 0.39962166 |
| Rbks | -0.677012471 | 0.400393373 |
| Lrp11 | -0.322265173 | 0.400520409 |
| Slc4a10 | 0.718813832 | 0.400553266 |
| Vgf | -0.718658527 | 0.400553266 |
| Cryzl1 | 0.247714256 | 0.400685019 |
| Prr14l | -0.284460177 | 0.40074386 |
| Zbtb37 | -0.516201278 | 0.40074386 |
| Tbc1d19 | 0.270576934 | 0.400836757 |
| Wdr46 | -0.309088498 | 0.40084884 |
| Pon3 | -0.680337862 | 0.40085263 |
| Tbc1d12 | 0.561173969 | 0.400975514 |
| Fgfr2 | -0.355590036 | 0.401126996 |
| Chd1l | -0.521409993 | 0.401126996 |
| 1600014C23Rik | 3.601498226 | 0.401276443 |
| Zfp868 | -0.306485888 | 0.401276443 |
| Ndufa9 | 0.21057608 | 0.401514472 |
| Mpnd | -0.259533534 | 0.401712301 |
| Dbn1 | 0.204563855 | 0.401820083 |
| Baiap2 | -0.407185866 | 0.401933111 |
| Gm10116 | -0.253495027 | 0.402014312 |
| Ceacam2 | -1.308971396 | 0.402065144 |
| Inip | 0.270725537 | 0.40224603 |
| Dexi | -0.395212522 | 0.40224603 |
| Soat2 | -2.396908702 | 0.40224603 |
| Ubxn10 | 1.340514761 | 0.402454318 |
| Cacnb2 | 0.646983556 | 0.402454318 |

| | | |
|---|---|---|
| Cep162 | 0.155117717 | 0.777152111 |
| Hnrnpa3 | 0.113547174 | 0.777152111 |
| Hist2h4 | -0.396498078 | 0.777152111 |
| 2210418O10Ri | -0.549887204 | 0.777152111 |
| Anapc7 | 0.109673965 | 0.777156746 |
| 1700001L19Ri | 0.291931876 | 0.777193114 |
| Clic4 | 0.083506689 | 0.777193114 |
| Snrpa | 0.0719755 | 0.777193114 |
| Ccs | 0.115351986 | 0.777246901 |
| Eef1a2 | 0.221846609 | 0.777351975 |
| Hpca | -0.151962729 | 0.777351975 |
| Poln | 0.411377143 | 0.777357564 |
| Gm3336 | 1.219872756 | 0.777403468 |
| Prcp | -0.169434399 | 0.777613287 |
| 4933428G20Ri | -0.677017059 | 0.777988684 |
| Iqsec2 | -0.276612234 | 0.778082776 |
| Phf23 | -0.092421677 | 0.778313529 |
| Fam159b | 0.777639169 | 0.778623031 |
| Sugp1 | 0.114003656 | 0.778623031 |
| Bdh1 | -0.128352508 | 0.778623031 |
| Kctd5 | 0.090313298 | 0.779597606 |
| Mthfd2l | -0.103507824 | 0.779597606 |
| Cenpj | -0.143062594 | 0.779597606 |
| Dock9 | -0.261985019 | 0.77979061 |
| Cep152 | -0.217296077 | 0.779811595 |
| Rsph3b | 0.147108772 | 0.779858204 |
| Bves | 1.043574613 | 0.779937111 |
| Mettl25 | 0.152980663 | 0.779937111 |
| Clec1b | -0.540908003 | 0.779937111 |
| Zan | -1.074216156 | 0.780047693 |
| Wbscr17 | 0.167861966 | 0.780171977 |
| Dusp28 | -0.159718341 | 0.78032113 |
| Ccdc89 | 0.638797114 | 0.7803757 |
| Hist1h2bj | 0.539678837 | 0.7803757 |
| Mpp4 | 0.493354559 | 0.7803757 |
| Trim39 | 0.125102902 | 0.7803757 |
| Adam23 | -0.16796965 | 0.780454509 |
| Slc7a6 | 0.116598668 | 0.780617374 |
| Ddrgk1 | 0.088183247 | 0.780617374 |
| Triap1 | -0.09150362 | 0.780617374 |
| Prr15 | -0.405514088 | 0.780617374 |
| Abca4 | 0.9913619 | 0.78069089 |
| Hist1h2be | 0.306014716 | 0.78069089 |
| Ttc21b | 0.174308431 | 0.78069089 |
| Abhd11 | 0.097113917 | 0.78069089 |
| Gskip | -0.103843056 | 0.78069089 |
| Pin1rt1 | -0.979906505 | 0.78069089 |
| Ski | -0.092361746 | 0.780717476 |
| Capn2 | -0.1458817 | 0.781439362 |
| Fkrp | 0.239965619 | 0.781456446 |
| Ppp1r42 | 0.554298486 | 0.781467836 |
| Capn5 | 0.226574779 | 0.781467836 |
| Btbd17 | 0.195666785 | 0.781467836 |
| Cacna1i | -0.348838503 | 0.781467836 |
| Efhc1 | 0.612829176 | 0.781518192 |
| Anxa11 | -0.363544338 | 0.781518192 |
| Zfp715 | 0.216345019 | 0.781754449 |
| Tmbim4 | 0.086609229 | 0.781754449 |
| Cmas | 0.091624696 | 0.781835971 |
| Cfap74 | -0.220490325 | 0.782572399 |
| Dscaml1 | 0.133517298 | 0.782933546 |
| Zfp65 | -0.22984347 | 0.782933546 |
| C130026I21Ri | -0.360303314 | 0.783102461 |
| Filip1 | -0.403521435 | 0.783241454 |
| C1ql2 | 0.781162259 | 0.783928994 |
| Adcyap1 | 0.271023288 | 0.783928994 |
| Gria4 | 0.131699146 | 0.78419258 |
| Alkbh4 | -0.15344043 | 0.784217393 |
| Kidins220 | -0.100575765 | 0.784377219 |
| Glp1r | 0.481686435 | 0.784483541 |
| Alkbh8 | 0.100919404 | 0.784483541 |
| Eea1 | -0.149561433 | 0.784483541 |
| Enpp3 | 0.537098504 | 0.784579866 |
| Rapgefl1 | 0.257953649 | 0.784579866 |
| Vps4a | 0.084313578 | 0.784579866 |
| 4833420G17Ri | 0.082619015 | 0.784579866 |
| Rai1 | -0.109434738 | 0.784579866 |
| Dact2 | -0.472896168 | 0.784584368 |
| Gm8923 | -0.728546801 | 0.784584368 |
| Elovl7 | 0.400222403 | 0.784792992 |
| Rspry1 | -0.088774472 | 0.785070662 |
| Eri1 | -0.11316869 | 0.78522823 |
| Fundc2 | -0.088829269 | 0.785604844 |
| Gpr17 | -0.819986271 | 0.785654966 |
| Rnf219 | 0.128611448 | 0.785676694 |
| Gm960 | -0.226305674 | 0.785676694 |
| Lyplal1 | 0.134525137 | 0.785955932 |
| Cd55 | 0.304738673 | 0.786010053 |
| Tor2a | -0.109592042 | 0.786010053 |
| Ank1 | -0.277434066 | 0.786010053 |
| Gulp1 | 0.232190183 | 0.786494628 |
| Nfic | -0.108880394 | 0.786494628 |
| Paxbp1 | 0.155786835 | 0.78688072 |
| Lekr1 | -0.178578404 | 0.78688072 |
| 1810032O08Ri | 0.27951887 | 0.786960073 |
| Ints10 | -0.109736712 | 0.786960073 |
| Slc25a32 | -0.198190271 | 0.787168211 |
| Pak2 | 0.070527145 | 0.787254046 |
| Pias4 | 0.134929583 | 0.787564864 |
| Klhl1 | 0.360209878 | 0.787771473 |
| Hecw2 | 0.211382376 | 0.788342762 |
| Zfp963 | 0.352677895 | 0.788524103 |
| Rd3 | -0.396101209 | 0.788525453 |
| Chml | 0.233060682 | 0.788681273 |
| Zmym6 | -0.139979824 | 0.788681273 |
| Cyct | -1.159484711 | 0.788681273 |
| Musk | 0.67603551 | 0.788685509 |
| A230052G05Ri | -0.318242544 | 0.788685509 |
| 1700092M07Ri | -1.006543148 | 0.788685509 |
| Tmem14c | 0.105781135 | 0.788784952 |
| Mtfr2 | -0.242578487 | 0.788790303 |
| Cers5 | -0.088316802 | 0.788992334 |
| Hmga1 | 0.092892631 | 0.789027522 |
| Itga4 | -0.286423091 | 0.789481935 |
| Lpgat1 | 0.11426523 | 0.789738387 |
| Gm6169 | 0.252350705 | 0.789773973 |
| Ldb1 | 0.101048761 | 0.789773973 |
| Phf20l1 | 0.077225318 | 0.789773973 |
| Ap3s2 | 0.090265739 | 0.789830669 |

| | | |
|---|---|---|
| Pigg | -1.54540373 | 0.000569295 |
| Mtfp1 | 3.62111716 | 0.000571426 |
| Antxr2 | -2.108166034 | 0.000573445 |
| Ccdc97 | -0.80713396 | 0.000574331 |
| Lsm14a | -0.929657583 | 0.000586288 |
| Map9 | 1.090322195 | 0.000586307 |
| Gse1 | -1.335102514 | 0.000586307 |
| Fam105a | -1.199888299 | 0.000588947 |
| Sh2d5 | 6.506498137 | 0.000592347 |
| Dact3 | 3.151223474 | 0.000592347 |
| Ndufa6 | 0.664205422 | 0.000592347 |
| Plekhf2 | -0.90667699 | 0.000592347 |
| Trappc8 | -1.026592913 | 0.000592347 |
| Kcnab2 | -1.450357056 | 0.000592347 |
| Nudt16l1 | 1.003277311 | 0.000592544 |
| Hmgb2 | 0.848898102 | 0.000592544 |
| Ercc6l2 | 1.350372841 | 0.000595162 |
| Snx11 | -1.05866149 | 0.000595486 |
| Cxxc4 | 1.522466362 | 0.000596908 |
| Rgs4 | 2.68435467 | 0.000598127 |
| Bcl2l11 | -1.428228804 | 0.000598127 |
| Nfkbie | -1.693841792 | 0.000599275 |
| Ints7 | -0.731027865 | 0.000601299 |
| Stat5a | -2.421947814 | 0.000601299 |
| Zfp92 | 6.503972074 | 0.000604449 |
| Zfp580 | 1.217329161 | 0.00060712 |
| Rnf31 | -1.15068716 | 0.000607251 |
| Pgd | -0.897883313 | 0.000609037 |
| Clec10a | 6.085477505 | 0.000612239 |
| Rps3 | 0.702806604 | 0.000613333 |
| Mcemp1 | -3.339638134 | 0.000613387 |
| Gstm2 | 3.284653742 | 0.000613841 |
| Ripk3 | -1.762870261 | 0.000613841 |
| Six3 | 2.86976296 | 0.00061512 |
| Elovl1 | -1.008077754 | 0.000625733 |
| Rpl10a | 0.790899357 | 0.000627812 |
| Oaf | -1.574378137 | 0.000627909 |
| Col18a1 | 1.573875014 | 0.000628361 |
| Fas | -3.074411043 | 0.000629235 |
| Pcdhga1 | -2.65028522 | 0.00062939 |
| Ccdc90b | 1.022434484 | 0.000638991 |
| Sowahc | -1.008122293 | 0.000638991 |
| Smad2 | -0.697543882 | 0.00063979 |
| Gm14597 | 1.438782432 | 0.000640225 |
| Tgfbi | -2.423083715 | 0.000642524 |
| Arhgap25 | -1.638362185 | 0.000643649 |
| Runx3 | -2.179244359 | 0.00064642 |
| Tspan12 | 1.034932336 | 0.000648798 |
| Mrgpre | 2.414013356 | 0.000652125 |
| Olfml3 | -1.381070523 | 0.000652276 |
| Hacd2 | -0.915230519 | 0.00065479 |
| Adcy3 | -1.628293755 | 0.00065479 |
| Mefv | -3.566513267 | 0.000667103 |
| Cenpq | 1.402437662 | 0.000669649 |
| Gbx2 | 2.781684392 | 0.000670911 |
| Htra4 | -4.139732612 | 0.0006711 |
| Ms4a3 | 6.936600236 | 0.000676235 |
| Adgrl4 | 6.743295278 | 0.00067826 |
| Bphl | 1.000860821 | 0.000681809 |
| Prep | -0.695962751 | 0.000683258 |
| Snrpd3 | 0.771235845 | 0.000688893 |
| Ran | 0.627359008 | 0.000690269 |
| Pgm1 | -0.993146549 | 0.000690269 |
| Eml2 | -1.256757993 | 0.000696623 |
| Zcrb1 | 0.730096539 | 0.000698728 |
| Nhej1 | 2.578712062 | 0.000699462 |
| Abtb2 | -1.86596055 | 0.000699704 |
| Plau | -2.116716709 | 0.000699771 |
| St6galnac5 | 1.672705222 | 0.00070407 |
| Cnn3 | 0.650592503 | 0.000706432 |
| Fam46b | -6.965367489 | 0.000709342 |
| Syt17 | 2.368859921 | 0.000713921 |
| Xpo5 | -1.093562013 | 0.000713921 |
| Cyb561a3 | -1.181826628 | 0.000714015 |
| D630003M21Rik | 6.402219586 | 0.00071642 |
| Myadm | -1.483212284 | 0.00071642 |
| Atf7 | -1.006849778 | 0.000716512 |
| Dnajc19 | 0.817825298 | 0.000716691 |
| Ndufa12 | 0.660831893 | 0.000716691 |
| Sf1 | -0.729773037 | 0.000716691 |
| Gpr137b | -1.260360438 | 0.000716691 |
| Proser3 | -1.591514348 | 0.000716691 |
| Por | -0.955164819 | 0.000718207 |
| Naglu | -1.502873922 | 0.000720832 |
| Dnajc13 | -0.814378439 | 0.000722526 |
| Tm9sf4 | -0.725195708 | 0.000725705 |
| Yeats4 | 0.703934973 | 0.000728748 |
| Selt | -0.696328778 | 0.00072883 |
| Lpcat3 | -0.932685422 | 0.00072883 |
| Xrcc5 | 1.212070271 | 0.000731183 |
| Zdhhc9 | -0.917380524 | 0.000731183 |
| Cd37 | -1.351284775 | 0.000732013 |
| Acvrl1 | -1.674549144 | 0.000735211 |
| Crb3 | 6.466377902 | 0.000740321 |
| Blvrb | -1.424986186 | 0.000741875 |
| Cacybp | 0.660361229 | 0.000742211 |
| Tmem243 | -0.987208041 | 0.000748993 |
| Jam2 | 1.321845709 | 0.000750195 |
| Pvrl3 | 1.689390001 | 0.000750221 |
| Gnat2 | -7.027890387 | 0.000750381 |
| Trit1 | 1.653905872 | 0.000755111 |
| Rhoa | -0.699331172 | 0.000755146 |
| Slc5a6 | 1.683389733 | 0.000757863 |
| Snrpd1 | 0.587319434 | 0.000757863 |
| Tmem192 | -0.93583491 | 0.00075954 |
| Zfp697 | 2.030285546 | 0.000761906 |
| Wbp4 | 0.798597973 | 0.000763525 |
| Nup160 | -0.973915459 | 0.000763525 |
| Efnb3 | 1.021097037 | 0.000766039 |
| Hcrtr2 | 6.489202267 | 0.000766198 |
| Zcchc17 | 0.724707882 | 0.000768303 |
| Atp13a2 | -0.87238615 | 0.000770002 |
| Sarm1 | -0.850636093 | 0.000772257 |
| Slc12a9 | -1.22134628 | 0.000772994 |
| Ptger4 | -1.79333638 | 0.000774464 |
| Frg1 | 0.73535478 | 0.00077623 |
| Tmem110 | -1.208544193 | 0.000776969 |
| Paqr8 | 1.676932333 | 0.000777683 |
| Tcn2 | -1.443348886 | 0.000777683 |

| | | |
|---|---|---|
| Zcchc24 | 1.197377266 | 0.090868066 |
| Sec24c | -0.450218002 | 0.090868066 |
| Rpn1 | 0.396750898 | 0.090885859 |
| Kdm4d | 5.009474486 | 0.091080977 |
| Cfl1 | 0.409590071 | 0.091080977 |
| Gnptg | 0.509907845 | 0.091084422 |
| Fzd10 | 2.009275378 | 0.091110128 |
| Psme4 | 0.544298304 | 0.091110128 |
| Traf6 | -0.52289885 | 0.091110128 |
| Pnmal1 | 0.602257197 | 0.091126672 |
| Pfkp | -0.876015582 | 0.091164325 |
| Aco1 | -0.480559792 | 0.091417095 |
| Actrt3 | -5.743274763 | 0.091438759 |
| Tdrd5 | -5.743274763 | 0.091438759 |
| Tyrobp | -0.528360205 | 0.091676186 |
| Arsa | -0.636623429 | 0.091676186 |
| Fam65a | -0.558349708 | 0.091684432 |
| Eno2 | -0.698575574 | 0.091747818 |
| Fmod | 2.029175513 | 0.091806918 |
| Adam12 | 1.585858714 | 0.091806918 |
| Cd276 | 0.69920716 | 0.091806918 |
| Mrpl4 | 0.461226303 | 0.091806918 |
| 9330159F19Rik | 0.424687665 | 0.091806918 |
| Colec12 | -1.124065173 | 0.091806918 |
| Mrps14 | 0.368992443 | 0.091986417 |
| Lpar2 | -0.900599896 | 0.092102246 |
| Gtf3c6 | 0.428517948 | 0.0921044 |
| Spata2 | -0.621062881 | 0.0921044 |
| Slc25a24 | -1.487586304 | 0.092223181 |
| Mcts1 | 0.383272733 | 0.09227163 |
| Pdzrn3 | -0.600284761 | 0.092274465 |
| Ift172 | -0.666667905 | 0.09249543 |
| Mob1b | -0.613366817 | 0.092498593 |
| Abca2 | -0.718934712 | 0.092498593 |
| Ccl9 | -1.752943831 | 0.09256153 |
| Snx2 | -0.555857716 | 0.092571527 |
| Rps6ka6 | 0.55856535 | 0.092741172 |
| Mtcp1 | -1.156200333 | 0.092741172 |
| Rpl27a | 0.379249249 | 0.092743536 |
| Parp10 | -1.621644323 | 0.092743536 |
| Donson | -0.872911672 | 0.092752197 |
| Atp6v1f | 0.376335227 | 0.092839905 |
| Pdlim7 | 0.487347982 | 0.092915897 |
| Ppp1r18 | -0.630760343 | 0.092915897 |
| Klhl33 | 4.855586086 | 0.092960414 |
| Gk5 | 1.413385005 | 0.092960414 |
| Mis18a | 0.581647352 | 0.093028233 |
| Epb42 | 5.072768314 | 0.093039459 |
| Dag1 | -0.430691833 | 0.09305157 |
| Pelo | 0.595974028 | 0.093075636 |
| Rps8 | 0.315230682 | 0.093075636 |
| Cfap20 | 0.449915747 | 0.09309759 |
| Mllt4 | 0.400326815 | 0.093144143 |
| Ttc34 | 2.807963681 | 0.093307459 |
| Rgs19 | -0.609195688 | 0.093326674 |
| Tcf3 | -0.499271344 | 0.09337483 |
| Hmgxb3 | -0.629027258 | 0.093408411 |
| Irx1 | -3.823769845 | 0.093411259 |
| Uggt2 | 0.765189083 | 0.093428358 |
| Gldn | 4.994620492 | 0.093455445 |
| Gtdc1 | -0.699370024 | 0.093459942 |
| Casq1 | 1.671837742 | 0.093554519 |
| Rad1 | 0.725171661 | 0.093554519 |
| Tagln2 | -0.707882129 | 0.093565045 |
| Stxbp4 | 0.618378695 | 0.093593881 |
| Xlr4c | -4.036298574 | 0.093593881 |
| Kif4 | 0.68214023 | 0.09372152 |
| Pyhin1 | -2.592197533 | 0.093779918 |
| Tbc1d14 | -0.443439591 | 0.093785955 |
| Prpf3 | -0.688864978 | 0.093807566 |
| Ccnj | -0.573206782 | 0.093889608 |
| Hras | -0.598950482 | 0.093889608 |
| Rpl7a-ps5 | 0.682857344 | 0.093899356 |
| Mical1 | -0.809342034 | 0.093921962 |
| Meis2 | 0.636373194 | 0.093934716 |
| Stx5a | -0.380670542 | 0.093997788 |
| Llph | 0.419963535 | 0.094027162 |
| Il33 | 5.348300998 | 0.094070971 |
| Acbd3 | -0.578787397 | 0.094070971 |
| Olfr920 | 5.288362915 | 0.094075396 |
| Patl2 | 4.96327002 | 0.094075848 |
| Gsk3a | -0.72345135 | 0.094091161 |
| Dnase1 | -1.798246885 | 0.094112554 |
| Pkmyt1 | -0.707936537 | 0.094129109 |
| Zc3h4 | -0.469549247 | 0.094213052 |
| Crls1 | 1.13215876 | 0.094241737 |
| Mr1 | 2.294509068 | 0.094245868 |
| Pigz | 2.228937924 | 0.09428671 |
| Ralgapa2 | -0.557500494 | 0.0943014 |
| As3mt | -0.82501217 | 0.094317846 |
| Nxph4 | 0.963499023 | 0.094441051 |
| Gtse1 | 0.950008155 | 0.094445044 |
| Chp2 | 5.266600613 | 0.094565718 |
| Zfp213 | 1.101672673 | 0.094659927 |
| Ttc39b | -0.623656199 | 0.094720174 |
| Wfikkn1 | -2.079601131 | 0.094909841 |
| Angptl8 | -3.801114596 | 0.095074371 |
| Rab6a | 0.410978633 | 0.095134764 |
| Mdp1 | 0.496123535 | 0.0951797 |
| Hspb2 | 4.876146221 | 0.095251938 |
| Dynlt1b | 0.488726293 | 0.095256485 |
| Xrcc1 | 0.449110018 | 0.095300251 |
| Zfp866 | -0.736065069 | 0.095352245 |
| Nisch | -0.364304471 | 0.095356671 |
| Bcl2a1b | -0.99213873 | 0.095394688 |
| Kiss1r | 3.428551294 | 0.095397379 |
| Loxl1 | 1.263977697 | 0.095397379 |
| Rps6ka5 | -0.655418025 | 0.095657751 |
| Pnn | 0.389242311 | 0.095691324 |
| Mga | -0.468269628 | 0.095941654 |
| Gpr88 | 2.154564522 | 0.096167019 |
| Acad9 | 0.445895681 | 0.096171507 |
| Nek2 | -0.73180097 | 0.096171507 |
| 4931440F15Rik | -4.335235869 | 0.096196013 |
| Pdss2 | 0.802798732 | 0.096286479 |
| Trim26 | -0.656728721 | 0.096307267 |
| Gm21738 | 2.930855167 | 0.096360203 |
| Fam193a | 0.466688246 | 0.096360203 |
| Zfp457 | 4.786407084 | 0.096399513 |

| | | |
|---|---|---|
| Mpp5 | 0.339690427 | 0.402464518 |
| Rbm25 | 0.173552643 | 0.402464518 |
| Ccdc74a | 0.841491964 | 0.402478261 |
| Rrp8 | 0.4016859 | 0.402478261 |
| Ccnt2 | -0.274106472 | 0.402478261 |
| Pelp1 | -0.294173521 | 0.402486086 |
| Hs1bp3 | -0.787202826 | 0.402843214 |
| Dlec1 | -1.838670796 | 0.402876669 |
| Cacna1a | -0.542973111 | 0.403048538 |
| Adamts8 | 2.051429685 | 0.403162953 |
| Eps8l3 | 0.659136295 | 0.403421734 |
| Zfp239 | -0.640614082 | 0.403612835 |
| Sp6 | 3.873423124 | 0.403704657 |
| Utp14b | -0.480846153 | 0.4038118 |
| Spata33 | 1.067593694 | 0.40438257 |
| Dars | -0.205237577 | 0.404484838 |
| Tfdp2 | -0.268985206 | 0.405446319 |
| Asic3 | 3.83548438 | 0.405656143 |
| Olfr874 | 3.83548438 | 0.405656143 |
| Vegfb | 0.311191341 | 0.405656143 |
| Scgb1b30 | -2.333428045 | 0.405656143 |
| Sf3b1 | 0.198446755 | 0.405877357 |
| Fam26f | -0.954570406 | 0.405877357 |
| Smarce1 | 0.203092566 | 0.405938616 |
| Hist1h4h | 0.704109796 | 0.406244018 |
| Erlec1 | 0.258240697 | 0.406253262 |
| Psmb10 | -0.351339027 | 0.406253262 |
| Ston1 | -1.180211698 | 0.406253262 |
| Epha2 | 3.898293046 | 0.40631046 |
| Lrrc75b | 0.4338389 | 0.40631046 |
| Amer3 | -0.46096861 | 0.40644596 |
| Cyyr1 | 1.268912548 | 0.406510144 |
| Lmo1 | -0.275761291 | 0.406588104 |
| Spink13 | 3.892333719 | 0.406601106 |
| Gpatch1 | 0.403216991 | 0.40663421 |
| Trim28 | -0.253379277 | 0.40663421 |
| Bsn | -0.279649468 | 0.40663421 |
| M1ap | -1.009069828 | 0.40663421 |
| Tmprss9 | -2.59132857 | 0.406818151 |
| Mcf2l | 0.280364335 | 0.406873713 |
| Irgm1 | -1.007154221 | 0.406896495 |
| Ptrh1 | -0.843867157 | 0.40699977 |
| Ddx31 | 0.508624406 | 0.407213313 |
| Serhl | -0.6412563 | 0.40754704 |
| Tsen15 | 0.36624016 | 0.407863384 |
| Tada2b | 0.686558844 | 0.407896604 |
| Psme1 | 0.24565006 | 0.407896604 |
| Ipo8 | 0.300732713 | 0.407941953 |
| Erbb2ip | 0.310402125 | 0.407966449 |
| Ddx58 | -1.236246333 | 0.408038067 |
| Mical3 | -0.257548478 | 0.408047502 |
| Gm17641 | 3.885191703 | 0.408410703 |
| Klk1b3 | 3.889535061 | 0.408418643 |
| Nek10 | 3.889535061 | 0.408418643 |
| Rassf7 | -0.458116782 | 0.409153759 |
| Cad | 0.381870295 | 0.409162231 |
| Btbd10 | -0.23917427 | 0.409227874 |
| Hmbs | 0.316914244 | 0.409666903 |
| Ttpal | -0.281043207 | 0.409666903 |
| Eno3 | -0.684293795 | 0.409666903 |
| Rmdn1 | 0.326549169 | 0.409902374 |
| Flt4 | 2.15171539 | 0.409922861 |
| Pou5f1 | 3.882277302 | 0.410082407 |
| Zic4 | 1.265238541 | 0.410120689 |
| Helz | -0.293297153 | 0.410120689 |
| Ufm1 | 0.223157652 | 0.41028693 |
| Fam229a | 2.89984532 | 0.410546318 |
| Pydc4 | 2.458770127 | 0.410617872 |
| Gm12184 | 0.62274737 | 0.410617872 |
| Dido1 | -0.259693351 | 0.410617872 |
| Cfap157 | -1.405984008 | 0.410617872 |
| Rad9b | 0.56853547 | 0.410728433 |
| Il25 | 3.826329854 | 0.410787124 |
| Gm5160 | 1.90578365 | 0.410787124 |
| Bckdhb | 0.358769821 | 0.410787124 |
| Dram2 | -0.304299731 | 0.410787124 |
| Rprd1a | -0.328166071 | 0.410787124 |
| Cnot6l | -0.382048135 | 0.410787124 |
| Nceh1 | -0.424315685 | 0.410787124 |
| Arhgap31 | -0.442034731 | 0.410787124 |
| Tmem241 | -0.4391975 | 0.410903825 |
| Slc9a7 | -0.718320669 | 0.411145282 |
| Sh2d3c | -0.317931497 | 0.411155741 |
| Ccdc92 | 0.338308226 | 0.411158573 |
| Rptor | -0.291462681 | 0.411158573 |
| Plekhb1 | 1.198027257 | 0.411569817 |
| Rxra | -0.622989277 | 0.411569817 |
| 1700028P14Rik | 1.789112626 | 0.411633899 |
| Ferl1 | -0.972850842 | 0.411693123 |
| L3mbtl2 | -0.354495956 | 0.411733853 |
| Golga1 | 0.336875202 | 0.411756773 |
| Slc14a1 | -1.039718145 | 0.411852911 |
| Pald1 | -0.440784541 | 0.411880515 |
| Ddx1 | 0.210505728 | 0.411907013 |
| Esd | -0.242958497 | 0.411952694 |
| Dmrta2 | 0.631691579 | 0.411954838 |
| Atrn | -0.259402206 | 0.411954838 |
| Npas1 | -0.568792378 | 0.411954838 |
| Spry4 | 1.101859957 | 0.411988048 |
| Prune | 0.370612077 | 0.411988048 |
| Sertm1 | 1.017449563 | 0.412368739 |
| Igsf21 | 0.405137858 | 0.412368739 |
| Ciao1 | -0.341202428 | 0.412483476 |
| Parm1 | 0.897283726 | 0.412567086 |
| Swap70 | -0.441270955 | 0.412567086 |
| Gstz1 | 0.346962178 | 0.412987606 |
| Lst1 | -0.385295093 | 0.412987606 |
| Fdxacb1 | 0.609360146 | 0.41301698 |
| Mms19 | -0.266543049 | 0.41301698 |
| Pitpnb | 0.208923591 | 0.413080832 |
| Vipr2 | 1.518382238 | 0.413092665 |
| AF529169 | 0.838531556 | 0.413092665 |
| L3hypdh | 0.511597967 | 0.413092665 |
| Psd | 0.417035829 | 0.413092665 |
| Armcx5 | 0.406708022 | 0.413092665 |
| Scai | 0.296544621 | 0.413179558 |
| 4930442H23Rik | -4.336475286 | 0.41348938 |
| Thbd | 1.168141113 | 0.41420328 |
| Wnt3 | 1.121443377 | 0.41420328 |

| | | |
|---|---|---|
| Arid1b | 0.085655418 | 0.789830669 |
| Suclg1 | 0.07334559 | 0.789830669 |
| Pggt1b | -0.153595965 | 0.789830669 |
| 1600002H07Ri | -0.190573301 | 0.79003943 |
| Amigo1 | 0.211125442 | 0.790700622 |
| Hpcal1 | -0.1444371 | 0.790700622 |
| Akirin2 | -0.107358896 | 0.790838432 |
| Papln | -0.462058671 | 0.791172542 |
| Zfp383 | 0.140303718 | 0.791596312 |
| Foxn4 | -0.853368847 | 0.791613092 |
| Dnajb4 | 0.110674627 | 0.791797282 |
| Ppp1r10 | -0.103173389 | 0.791806099 |
| Wdr44 | 0.129321439 | 0.79192331 |
| Tbp | -0.110065285 | 0.792004132 |
| Ankrd66 | 0.550039299 | 0.792249183 |
| Fam151b | -0.662583313 | 0.792249183 |
| Dld | -0.071713043 | 0.792265937 |
| Ctsf | 0.121897355 | 0.792271805 |
| Sigmar1 | 0.119783199 | 0.792330687 |
| Scn3a | 0.100282585 | 0.792330687 |
| Gm10110 | 0.526755408 | 0.792580344 |
| Glt8d2 | -0.294109093 | 0.792990332 |
| Bhlhe41 | 0.100619119 | 0.793149108 |
| A230050P20Ri | 0.242331502 | 0.793269606 |
| Gm4724 | 0.895416209 | 0.793270069 |
| 4933430I17Rik | 0.418425714 | 0.793270069 |
| Taf4a | -0.191692353 | 0.793270069 |
| Med20 | -0.101924319 | 0.793431322 |
| Car4 | 0.584423835 | 0.793696487 |
| Spata2l | 0.280293314 | 0.793914408 |
| Asxl2 | 0.13554721 | 0.793949067 |
| Bad | -0.09531753 | 0.793949067 |
| Fam159a | 0.911081712 | 0.794417921 |
| Trpv4 | 0.538874044 | 0.794417921 |
| Satb1 | 0.099915477 | 0.794417921 |
| Ammecr1 | -0.146224174 | 0.794478094 |
| Park2 | 0.21036208 | 0.794515866 |
| Gm2956 | 1.118391354 | 0.794543908 |
| Tmed8 | -0.177582572 | 0.794543908 |
| Hsf2bp | -0.734162541 | 0.794543908 |
| Scml4 | 0.232995271 | 0.794606014 |
| Chgb | 0.132894232 | 0.794606014 |
| Rbfox3 | 0.112928276 | 0.794606014 |
| Gstp2 | 1.172275316 | 0.79487431 |
| Mdc1 | -0.115872034 | 0.79487431 |
| Ckb | 0.071809124 | 0.794880405 |
| Fabp3-ps1 | 0.478190554 | 0.79518063 |
| Myl4 | -0.53680182 | 0.795189937 |
| Rpl34 | 0.103964394 | 0.79520766 |
| Ndufaf1 | -0.098548021 | 0.795280146 |
| Wars2 | 0.16669981 | 0.795750377 |
| Zfp182 | 0.139934462 | 0.795750377 |
| Terf1 | -0.111181885 | 0.795750377 |
| Pnpla3 | -0.246097565 | 0.795750377 |
| 4930562C15Ri | -0.373880997 | 0.795750377 |
| Cox15 | 0.151339993 | 0.795884266 |
| Tmem186 | 0.148983657 | 0.795884266 |
| Rab1b | 0.068201722 | 0.795884266 |
| Atg9b | -0.757128157 | 0.795884266 |
| Cdk12 | -0.095661495 | 0.795929118 |
| Trp53inp1 | 0.100192188 | 0.795966465 |
| Ift80 | -0.138901041 | 0.796440663 |
| Gmnn | -0.113827532 | 0.796771503 |
| Gm8206 | 1.041411092 | 0.796816265 |
| D3Ertd751e | 0.258827859 | 0.796873725 |
| Mmp2 | 0.197069871 | 0.796912235 |
| Snx3 | 0.071728783 | 0.79697841 |
| Psmd13 | 0.068265436 | 0.797052176 |
| Gm13139 | 0.292547725 | 0.797367743 |
| Gtf2f2 | 0.103229465 | 0.797367743 |
| Ergic2 | -0.077672587 | 0.797367743 |
| Kcnrg | 0.456489032 | 0.79739089 |
| Casp7 | 0.176667755 | 0.79739089 |
| Otx1 | 0.12250446 | 0.79739089 |
| Zfp512b | 0.104620265 | 0.79739089 |
| Atxn7l3 | -0.119505342 | 0.79739089 |
| Ttc37 | -0.124868691 | 0.79739089 |
| Plcd3 | -0.314470385 | 0.79739089 |
| Pik3c2a | 0.118694215 | 0.797487985 |
| Prkx | -0.086406303 | 0.797487985 |
| Gpr22 | 0.264156275 | 0.797499386 |
| Gm10093 | -0.116359481 | 0.797499386 |
| Tdg | 0.094155563 | 0.797596756 |
| Gcnt7 | 0.452122674 | 0.797599254 |
| 4930522L14Ri | 0.325186477 | 0.797599254 |
| Coq4 | 0.148988819 | 0.797641358 |
| Dhx15 | 0.077096365 | 0.797641358 |
| Srpr | -0.096886513 | 0.797641358 |
| 4933427D14Ri | 0.159635083 | 0.797788204 |
| BC048507 | -0.860853757 | 0.797788204 |
| Lysmd1 | 0.172476152 | 0.797841733 |
| Pum1 | -0.084125496 | 0.797841733 |
| Pcdhb14 | -0.243389607 | 0.797841733 |
| Eif2s1 | 0.075340477 | 0.798003481 |
| 9930012K11Ri | -0.254813461 | 0.798003481 |
| Wipf1 | -0.209732143 | 0.798022905 |
| Jmjd4 | 0.180383568 | 0.798097987 |
| Klhl5 | 0.094043323 | 0.79826739 |
| Tubb2b | 0.080742189 | 0.79826739 |
| Adck4 | -0.116954504 | 0.79826739 |
| Usp6nl | 0.114476952 | 0.798367815 |
| Ccdc171 | -0.273384968 | 0.798413762 |
| Ifit3b | -0.631233885 | 0.798413762 |
| Tmem143 | 0.336458332 | 0.798617449 |
| Acot6 | -0.489031641 | 0.798617449 |
| D16Ertd472e | 0.21858981 | 0.79886296 |
| Stx8 | 0.097683621 | 0.79886296 |
| B3gnt2 | -0.521014635 | 0.799299166 |
| Sfxn4 | 0.158478817 | 0.799514813 |
| Papss1 | 0.07439827 | 0.799514813 |
| RP23-198F11. | 0.958909139 | 0.79983534 |
| Lin7c | 0.079251565 | 0.799846663 |
| BC068281 | -0.256526603 | 0.799846663 |
| Gm3604 | 0.536548273 | 0.800104654 |
| Ints3 | -0.130787464 | 0.800252958 |
| A930018M24R | 0.499385058 | 0.800410374 |
| Wdr25 | 0.293511045 | 0.800523631 |
| Plekha5 | 0.092305052 | 0.800523631 |
| Foxo4 | 0.331223571 | 0.800771113 |

| | | |
|---|---|---|
| Mtm1 | -1.484945413 | 0.000778524 |
| Zfp263 | -1.094189116 | 0.000779217 |
| Arxes1 | 1.530709252 | 0.000780905 |
| Ppil1 | 0.836208024 | 0.00078225 |
| Fam96a | -0.831410226 | 0.000782368 |
| Hba-a1 | 1.407264852 | 0.000784416 |
| Rpl24 | 0.578917492 | 0.000788576 |
| Uggt1 | -0.753298664 | 0.000788811 |
| Kbtbd3 | 1.789914179 | 0.000789821 |
| Zc3h15 | 0.766014977 | 0.000794763 |
| Trip10 | -1.60574421 | 0.000796197 |
| Mrpl47 | 1.053431226 | 0.000797968 |
| Pafah1b3 | 0.65814555 | 0.0007987 |
| Icam1 | -1.530975768 | 0.0007987 |
| Camkk1 | -1.019287295 | 0.000799346 |
| Slco4a1 | -2.012875603 | 0.000800545 |
| Npff | -6.95897848 | 0.00080112 |
| Pik3ca | 1.025982327 | 0.000807561 |
| Acyp2 | 2.209625818 | 0.00080823 |
| Grm1 | -2.684084905 | 0.000812728 |
| Fabp3 | 1.159214855 | 0.000813075 |
| Pglyrp1 | -2.242552717 | 0.000814533 |
| Zfp428 | 0.713904445 | 0.000820953 |
| Sult1a1 | 6.902083498 | 0.000825488 |
| Smpdl3a | -1.248142549 | 0.000825984 |
| Tspo | -1.32128466 | 0.000825984 |
| Pik3r6 | -1.694755362 | 0.00082878 |
| Prss27 | -4.623786535 | 0.000831847 |
| Fcrl5 | -4.77395675 | 0.000831847 |
| Fut4 | -4.675963685 | 0.000832405 |
| Adora1 | 1.786163244 | 0.0008389 |
| Siglech | 1.8961193 | 0.000850584 |
| Flot1 | -0.776104935 | 0.000851263 |
| Slc2a13 | -2.237191207 | 0.000851263 |
| Calm2 | 0.756620285 | 0.000851432 |
| Sema3d | 5.966632798 | 0.000856111 |
| Fopnl | 0.683663619 | 0.000856239 |
| Ndufb7 | 0.658334143 | 0.000874574 |
| Psma7 | 0.538009711 | 0.000874574 |
| Ajap1 | 4.010606072 | 0.000876101 |
| Tmeff2 | 1.293792148 | 0.000876101 |
| Map2k1 | -0.866462669 | 0.00087997 |
| Rala | 0.621768655 | 0.000881657 |
| Stmn3 | 0.710779026 | 0.000882166 |
| Plxnc1 | 1.633809045 | 0.000887405 |
| Camk2a | 1.511515176 | 0.000888454 |
| Fxyd1 | 3.191577935 | 0.000891761 |
| Gm10146 | 0.967919469 | 0.00089191 |
| Ptdss1 | 0.790894153 | 0.00089191 |
| Kcna3 | -1.793541188 | 0.000894615 |
| Cthrc1 | 2.40083832 | 0.000897499 |
| Adgrf5 | 6.719358066 | 0.000902417 |
| Kif21a | 0.642764061 | 0.000908221 |
| Deb1 | 0.806526454 | 0.000908244 |
| Clec4a3 | -1.326640221 | 0.000910657 |
| Fam171a2 | -1.279931351 | 0.000915843 |
| Rplp2 | 0.721656028 | 0.000916534 |
| Scg5 | 0.743794097 | 0.000918483 |
| Xlr3a | 3.874697985 | 0.000920891 |
| Vti1b | 0.697291103 | 0.000927189 |
| Ice1 | -0.835645843 | 0.000931387 |
| Ncf2 | -1.031181537 | 0.000933037 |
| Rnaseh2c | 0.855308514 | 0.000935567 |
| Vti1a | -1.115069508 | 0.000936772 |
| Elf1 | -1.213184201 | 0.000938066 |
| Traf3ip3 | -1.715895897 | 0.000938066 |
| Csnk1d | -0.757340665 | 0.000938168 |
| Pi4kb | -0.842482816 | 0.000938168 |
| Slc24a3 | 2.441283767 | 0.000940495 |
| Mis18bp1 | 1.013609377 | 0.000944373 |
| Pigyl | 0.890342411 | 0.000945649 |
| Gnai2 | -0.784150596 | 0.000946794 |
| Chst1 | 1.522353306 | 0.000947058 |
| Fam20c | -0.883746929 | 0.000948319 |
| Sugt1 | 0.791383598 | 0.000953155 |
| Cd68 | -1.279688514 | 0.000953701 |
| Hey2 | 6.282051408 | 0.000955909 |
| Ckap2 | 1.650658668 | 0.000960666 |
| Adipor1 | -0.812005823 | 0.000966486 |
| Prss23 | 2.557323992 | 0.000967255 |
| Lrmp | -1.662357465 | 0.000968924 |
| Hmgn3 | 0.986101958 | 0.000970523 |
| Sgf29 | 1.01133259 | 0.000972016 |
| Pcdhga8 | -2.723015206 | 0.000973612 |
| Srf | -0.941932272 | 0.000978847 |
| Nrsn2 | 2.21671958 | 0.000979308 |
| Pgs1 | -0.947673674 | 0.000979308 |
| Fsd1 | 0.84058443 | 0.000983173 |
| Fbxl8 | -6.568234497 | 0.000983173 |
| Abcg3 | 6.341119463 | 0.000987197 |
| Thoc7 | 0.750008164 | 0.000992786 |
| Rpl11 | 0.57683764 | 0.000993826 |
| Rwdd1 | 0.773317408 | 0.000997108 |
| Ccar1 | 0.662440989 | 0.000997807 |
| E130114P18Rik | 1.24636564 | 0.001001489 |
| Zbtb41 | 0.923082723 | 0.001008038 |
| Rnf187 | 0.669341683 | 0.001008219 |
| Ifngr2 | -0.603650148 | 0.001010546 |
| Stap1 | -1.837206153 | 0.001010546 |
| Nr3c1 | -1.34376564 | 0.00101428 |
| Ptpn3 | 6.116203024 | 0.001017901 |
| Iqca | 6.46352207 | 0.001022359 |
| Lilr4b | -3.957619734 | 0.001024772 |
| Gabra3 | 1.656980966 | 0.001024821 |
| Steap2 | 2.059900374 | 0.001025165 |
| Uqcr11 | 0.643948945 | 0.001029712 |
| Fcer1g | -1.075053376 | 0.001029887 |
| 1810043G02Rik | 1.018597982 | 0.001032162 |
| Alox5ap | -1.291416789 | 0.001035546 |
| Dusp27 | 6.632043858 | 0.001036329 |
| Pdgfra | 2.330076949 | 0.001039907 |
| Atf6b | -0.728553714 | 0.001044132 |
| Colgalt1 | -0.881988255 | 0.001048852 |
| Fkbp11 | 3.437992077 | 0.001050467 |
| Filip1l | -1.636243912 | 0.001053116 |
| Shisa5 | -1.336018574 | 0.001058835 |
| Fhl4 | 4.182411826 | 0.001062759 |
| Ctla2a | 4.48762496 | 0.001062857 |
| Necap1 | -0.665802366 | 0.001062857 |

| | | |
|---|---|---|
| Krt7 | -5.707570941 | 0.096429133 |
| Ifitm10 | 3.603086592 | 0.096538191 |
| Lysmd2 | 0.695486277 | 0.096538191 |
| D230025D16Rik | -0.623974972 | 0.09656112 |
| 2700097O09Rik | 0.690474687 | 0.096566101 |
| Exoc3l4 | 4.749436986 | 0.096635409 |
| Ttll1 | 0.552206163 | 0.096650438 |
| Aplnr | 5.198300562 | 0.096720465 |
| Zfp677 | 1.657504393 | 0.096750933 |
| Stat2 | -1.125289834 | 0.096828686 |
| 2510009E07Rik | -0.591512283 | 0.096959094 |
| Ccm2l | -2.836146997 | 0.097023032 |
| Tpbpb | -5.709136413 | 0.097029109 |
| Gpr27 | 0.826045682 | 0.097038575 |
| Vamp8 | -0.564954863 | 0.097038575 |
| Liph | -1.914864527 | 0.097038575 |
| Helz2 | -2.946506486 | 0.097041438 |
| Eif3i | 0.372805905 | 0.097069316 |
| Foxo1 | 0.887077423 | 0.097100075 |
| Slc7a1 | -0.448457491 | 0.097100075 |
| Spopl | -0.568178804 | 0.097100075 |
| Aim1 | -1.915405225 | 0.097100075 |
| Ptbp2 | 0.39137391 | 0.097224608 |
| Arhgap5 | 0.488755861 | 0.097232478 |
| Acot9 | -0.645141143 | 0.097521908 |
| Reep6 | -1.789599034 | 0.097693896 |
| Zufsp | -0.618166892 | 0.097784588 |
| Slc16a7 | -0.937464242 | 0.097784588 |
| Trp53tg5 | -4.716415777 | 0.097784588 |
| Ifngr1 | -0.658833195 | 0.09779683 |
| 8030462N17Rik | -0.974984726 | 0.097918544 |
| Zfp597 | 0.730411851 | 0.09809892 |
| Krt222 | 2.576180744 | 0.098161101 |
| Vipas39 | -0.545045957 | 0.098161101 |
| Nop14 | 0.498258731 | 0.098257392 |
| Fam178b | -2.231360333 | 0.098257392 |
| Ift43 | 0.577112765 | 0.098284207 |
| Gsg1l | -1.017980407 | 0.098284207 |
| Styk1 | 4.812663233 | 0.09836166 |
| Syndig1l | 1.794991964 | 0.098363184 |
| Cdc123 | 0.430615708 | 0.098363184 |
| Efemp2 | 0.571141738 | 0.098391819 |
| Man1c1 | -0.641560977 | 0.098524834 |
| Cbfb | -0.605875458 | 0.09870417 |
| Tapt1 | -0.870473988 | 0.098770915 |
| Fbp1 | -1.473940249 | 0.098770915 |
| Tmem100 | 1.896307778 | 0.098811449 |
| Fam118a | -0.605310899 | 0.098811449 |
| Pkd2l1 | 4.693119405 | 0.098867053 |
| Armc7 | -1.163726633 | 0.098867053 |
| Paox | -0.763881472 | 0.098917558 |
| Slc43a1 | 1.290277799 | 0.098929091 |
| Prr11 | 0.995737837 | 0.099209689 |
| Arnt2 | -0.40160191 | 0.099230657 |
| Mgarp | -1.223935753 | 0.099286411 |
| Pnldc1 | -1.556353363 | 0.099364135 |
| Gm15155 | -5.376624847 | 0.099364135 |
| Slc38a11 | 4.878400926 | 0.099406876 |
| Med6 | 0.58365635 | 0.099427488 |
| Rcn2 | 0.374617019 | 0.099562695 |
| Gpt2 | 0.597621175 | 0.099594438 |
| Tbc1d15 | -0.535959736 | 0.099594438 |
| Hectd2 | 0.720739869 | 0.099643139 |
| Gm7173 | 2.203741803 | 0.099706075 |
| Cpa2 | 2.131607438 | 0.09972461 |
| Actr6 | 0.621037933 | 0.09972461 |
| Med22 | 0.394758034 | 0.09972461 |
| Pmpcb | 0.37053006 | 0.09972461 |
| Kcnu1 | 1.643230633 | 0.099748171 |
| Hbb-bs | 1.141972362 | 0.099748171 |
| Fam160b2 | -0.81137588 | 0.099821624 |
| Pllp | 4.048974671 | 0.099830698 |
| Pik3c3 | -0.559367256 | 0.099830698 |
| Wdr70 | 0.623933892 | 0.099923079 |
| Rcsd1 | -0.959453206 | 0.099928486 |
| Ifi202b | -1.867012107 | 0.099998714 |
| Agmo | -1.220673696 | 0.100043182 |
| BC051142 | 2.187870427 | 0.100046752 |
| Gtpbp1 | -0.613100854 | 0.100210534 |
| Insig2 | 0.597591139 | 0.100259546 |
| Ttc13 | -0.453587031 | 0.100279 |
| Apol9b | 4.675461683 | 0.100328553 |
| Rpl12 | 0.4165033 | 0.100422719 |
| Crtc2 | -0.663089737 | 0.100651459 |
| Sel1l3 | 0.88860268 | 0.100913356 |
| Pcnx | -0.710515447 | 0.100975593 |
| Plekhd1 | 5.23105175 | 0.10097771 |
| Appl1 | 0.451097557 | 0.10097771 |
| Dgkh | -0.799916413 | 0.101120416 |
| Tmem206 | -0.591628364 | 0.10133376 |
| Gm9936 | 5.219691124 | 0.101436229 |
| Impa2 | 1.053156486 | 0.101436229 |
| Erlin1 | -0.714491796 | 0.101436229 |
| AI481877 | -2.747270572 | 0.1016117 |
| Naa25 | -0.438809323 | 0.101630689 |
| Apoa1bp | 0.390412915 | 0.101660797 |
| Pcdha5 | 2.396319703 | 0.101718399 |
| Alg8 | -1.003470072 | 0.101718399 |
| Lrrc72 | -4.709881896 | 0.101853547 |
| Gria2 | 0.48534242 | 0.101878391 |
| Slc20a2 | -0.675404122 | 0.101891469 |
| Tmem126a | 0.414484139 | 0.101951927 |
| Lfng | -0.906442122 | 0.10201642 |
| Ppp6c | -0.37730279 | 0.102084038 |
| Usp20 | -0.591057353 | 0.102186198 |
| St3gal5 | -0.615924258 | 0.102186198 |
| Sap30l | -0.788146685 | 0.102435055 |
| Zbtb43 | -0.715742821 | 0.102436626 |
| Pydc3 | -1.892548036 | 0.102453687 |
| Eri3 | 0.349293672 | 0.102456845 |
| Rock1 | -0.382332283 | 0.102502609 |
| 2410004P03Rik | 1.946931685 | 0.102717952 |
| 11-Mar | 1.2850009 | 0.102826895 |
| Bmp2k | -0.658428536 | 0.102826895 |
| Ppip5k1 | 0.912401922 | 0.102838852 |
| Tet1 | 0.564156086 | 0.102838852 |
| Ebi3 | -0.762147031 | 0.10283954 |
| P3h3 | 1.086517976 | 0.10295332 |
| Apobr | -1.795076391 | 0.10295332 |

| | | |
|---|---|---|
| Sh2d6 | -1.434327718 | 0.414218962 |
| Ifitm1 | -1.086774017 | 0.414253472 |
| Rgs12 | -0.302135307 | 0.414565737 |
| Gm42715 | 0.443920898 | 0.414648278 |
| Zfp24 | 0.244444308 | 0.414677072 |
| Klf7 | 0.224889172 | 0.414693346 |
| Tnrc6c | -0.259609876 | 0.414738715 |
| Sh3bgrl2 | -0.294059434 | 0.414823991 |
| Zfp758 | 0.524839455 | 0.414863229 |
| Tubd1 | 0.623621847 | 0.415298717 |
| Eogt | -0.781342005 | 0.415298717 |
| Homer3 | -0.33662225 | 0.41538958 |
| Suv420h2 | -0.259193821 | 0.415652563 |
| Atg14 | 0.336311797 | 0.41585745 |
| Smim17 | 1.579324028 | 0.415907052 |
| Arhgap12 | -0.268494294 | 0.41617128 |
| Dcaf13 | 0.273953471 | 0.416190886 |
| AU019823 | 0.302930922 | 0.416597519 |
| Orc3 | -0.270307672 | 0.416721774 |
| Tmem65 | -0.273015712 | 0.416721774 |
| Smchd1 | 0.214126571 | 0.417059172 |
| C77080 | 0.737381619 | 0.417232426 |
| Srgap1 | -0.302362072 | 0.417458104 |
| Gem | -0.396043089 | 0.417475894 |
| Npas2 | -1.078966668 | 0.417650639 |
| Fam26e | 3.502964722 | 0.417685458 |
| Ncoa7 | -0.342647376 | 0.417849971 |
| Zbtb10 | -0.445494418 | 0.418251703 |
| Txndc12 | 0.289247299 | 0.418407735 |
| Ric8b | 0.251326338 | 0.418466289 |
| Mgat4d | 3.513441998 | 0.418528049 |
| Mtfr1 | -0.24339195 | 0.418681032 |
| Fzd5 | 1.336836541 | 0.418813777 |
| Tmem132e | 2.369661249 | 0.418943999 |
| Acot4 | 1.701354933 | 0.418943999 |
| Cnot1 | -0.195891737 | 0.418943999 |
| Acaca | -0.30495104 | 0.418943999 |
| Coq10b | 0.260143218 | 0.419022949 |
| Ngef | 0.719050846 | 0.419118351 |
| Rundc1 | 0.369651883 | 0.419195112 |
| Fars2 | 0.256104198 | 0.419195112 |
| Impg2 | -1.445634644 | 0.419195112 |
| Usp54 | -0.382057325 | 0.419211391 |
| Zfp386 | 0.28540705 | 0.419329709 |
| Tcf7 | 0.923374471 | 0.419383732 |
| Ephb4 | 0.564254031 | 0.41958387 |
| Wscd2 | -0.656529974 | 0.41958387 |
| Gjc1 | 0.34536869 | 0.420135164 |
| Ifi203 | -0.969289009 | 0.42055359 |
| Aifm3 | 1.171542127 | 0.420599066 |
| Mxd4 | -0.270842237 | 0.420599066 |
| Pknox1 | 0.255815833 | 0.42062403 |
| Gabrg2 | 0.326289826 | 0.420657457 |
| Polr1a | -0.44226273 | 0.420657457 |
| Syne3 | -1.357961679 | 0.421249261 |
| Mylpf | 0.774141571 | 0.421281464 |
| Etfa | 0.205500835 | 0.421419845 |
| Gjb1 | 3.781572765 | 0.421584166 |
| 1700066B19Rik | 2.858428683 | 0.421584166 |
| Cops6 | 0.193754782 | 0.421694734 |
| Rell2 | 0.595753197 | 0.421709586 |
| Chordc1 | 0.237711297 | 0.421709586 |
| Pdia4 | -0.244300161 | 0.42179195 |
| Gm13420 | -4.326420454 | 0.42179195 |
| Bmp15 | 3.802837307 | 0.422157648 |
| Prr23a1 | 3.802837307 | 0.422157648 |
| Ccl3 | -0.389347412 | 0.422157648 |
| Xndc1 | -0.514968894 | 0.422157648 |
| Hsh2d | -2.217859644 | 0.422157648 |
| Galr2 | 3.105267046 | 0.422288644 |
| Star | -0.684014058 | 0.422288644 |
| Otud4 | -0.240781154 | 0.422385938 |
| Mok | 0.482982358 | 0.422617128 |
| Tstd3 | -0.483363582 | 0.422776911 |
| Smim6 | 1.366906631 | 0.423041617 |
| Frmd7 | 1.666950689 | 0.423055119 |
| Wdhd1 | -0.42664211 | 0.423088048 |
| Rgs9 | 0.497486362 | 0.423415433 |
| Pcdhgb5 | 0.954738885 | 0.423463714 |
| Tspan33 | -0.521401262 | 0.423513056 |
| Nudt8 | -0.342426722 | 0.423735623 |
| Zc3hav1l | -0.405659955 | 0.423826741 |
| Gli3 | 0.314478718 | 0.423861662 |
| Idua | -0.534180557 | 0.423883 |
| Zfp791 | -1.102704195 | 0.423883 |
| Hist1h3f | -1.714275664 | 0.423905132 |
| Cdc37 | 0.197614163 | 0.424052268 |
| Pxdc1 | 0.478035658 | 0.424101206 |
| Cutc | 0.422623992 | 0.424307963 |
| Bcor | -0.271484319 | 0.424307963 |
| Cmc4 | 0.726743999 | 0.424450621 |
| Fam162a | 0.254433015 | 0.425013079 |
| Cnbd2 | -0.912990101 | 0.425013079 |
| Ddn | -1.445236805 | 0.425013079 |
| Clec2d | -1.011015091 | 0.425049269 |
| Mtmr4 | -0.228046074 | 0.425114398 |
| Jdp2 | -1.970734492 | 0.425131844 |
| Nkx6-2 | 1.733212861 | 0.42519432 |
| D030056L22Rik | 0.251537873 | 0.425198816 |
| Asxl3 | 0.316450982 | 0.425203582 |
| Slc25a46 | 0.21440758 | 0.425203582 |
| Hba-a2 | -0.598958325 | 0.425304806 |
| Hist1h2bp | 1.261104734 | 0.425540971 |
| Gria1 | -0.218621114 | 0.425540971 |
| Pdxk | -0.343395143 | 0.425546579 |
| Ppat | -0.335172839 | 0.425594464 |
| Entpd5 | -0.369363108 | 0.425656258 |
| Chuk | -0.298196813 | 0.425728254 |
| Adap1 | -0.478931569 | 0.425792266 |
| Slc9a3r1 | 0.371277528 | 0.425862806 |
| Atp5sl | -0.300745847 | 0.425862806 |
| Gngt2 | -0.426189195 | 0.425862806 |
| Tep1 | -0.478832855 | 0.425862806 |
| Aptx | -0.234239632 | 0.425864576 |
| Mfsd10 | -0.276151415 | 0.425864576 |
| Slc16a13 | -0.435803665 | 0.425864576 |
| Lsp1 | -0.508084129 | 0.425864576 |
| Slco5a1 | 0.364262386 | 0.425953137 |
| Trh | 3.815261402 | 0.42612006 |

| | | |
|---|---|---|
| 1810026J23Rik | -0.085078488 | 0.800833514 |
| Ptchd2 | -0.202061682 | 0.800833514 |
| Tcf4 | 0.089024254 | 0.800891772 |
| Lcorl | 0.087919193 | 0.800931926 |
| Rexo2 | 0.084286765 | 0.800931926 |
| Irf1 | -0.154316717 | 0.80117281 |
| Il2 | -0.734200769 | 0.80117281 |
| Gm5751 | 0.433015364 | 0.801239829 |
| Ubr3 | -0.085744833 | 0.801379538 |
| Gpr12 | -0.220671785 | 0.801410332 |
| Gm10020 | 0.308025924 | 0.801580368 |
| Phf1 | 0.193299178 | 0.801580368 |
| Atp5a1 | -0.062373499 | 0.801580368 |
| Slx4 | -0.124894665 | 0.801580368 |
| Gm10604 | 0.605543609 | 0.801601115 |
| Gtf2h2 | 0.119258084 | 0.801782706 |
| Gm42641 | 0.519565649 | 0.80189715 |
| Cdh9 | 0.394644611 | 0.80195852 |
| Gm6483 | 0.118438463 | 0.802171778 |
| Cd40 | -0.442255782 | 0.802184942 |
| Zscan22 | 0.15482431 | 0.802190413 |
| Cotl1 | 0.097833757 | 0.802333854 |
| Gins2 | 0.096786159 | 0.802473112 |
| Nacad | -0.097525288 | 0.802933149 |
| Khdrbs2 | -0.105865198 | 0.803048277 |
| Man2a1 | -0.188084793 | 0.803243639 |
| Tnni3 | 1.132310338 | 0.803254618 |
| Ccdc117 | -0.169408296 | 0.803264242 |
| Rcan3 | -0.122187112 | 0.803531346 |
| Fbxl12 | 0.15054901 | 0.803595785 |
| Pola1 | 0.088674914 | 0.803595785 |
| Kif1bp | -0.084653115 | 0.803595785 |
| Grid2ip | 0.60670439 | 0.803601646 |
| Mapk1ip1l | 0.085478293 | 0.803601646 |
| AI182371 | -0.844766184 | 0.803603564 |
| Adgrb3 | 0.126881048 | 0.803842854 |
| Sptan1 | 0.084615711 | 0.803907631 |
| Sos2 | 0.128053827 | 0.804041809 |
| Fn1 | 0.112066223 | 0.804041809 |
| Blnk | 0.168991527 | 0.804096603 |
| Ddx56 | -0.0706027 | 0.804096603 |
| Prkch | 0.233249555 | 0.804487941 |
| Ccdc108 | -0.507039829 | 0.804545973 |
| Gm3095 | 0.729133539 | 0.804780908 |
| Sgip1 | 0.123531993 | 0.804833973 |
| Epor | 0.452851946 | 0.805068486 |
| Pank1 | 0.196657159 | 0.805068486 |
| Raet1e | 0.589464022 | 0.805198659 |
| Vstm2b | 0.179952009 | 0.805621276 |
| Galt | 0.159134617 | 0.805621276 |
| Sptbn4 | -0.324039761 | 0.80562944 |
| Golga5 | -0.094294137 | 0.805749514 |
| Chst8 | -0.211463401 | 0.806038732 |
| Aup1 | -0.11838929 | 0.806408152 |
| Prrg2 | -0.278597009 | 0.806408152 |
| Zswim6 | 0.10980162 | 0.806502976 |
| Camk1 | -0.074258922 | 0.806502976 |
| Phf21a | -0.080362745 | 0.806502976 |
| 5031439G07Ri | -0.134643617 | 0.806583064 |
| Dimt1 | 0.134228017 | 0.806618615 |
| Dnajc5 | -0.072579978 | 0.806910238 |
| Wif1 | 0.322308331 | 0.807129416 |
| Lrp2 | 0.264509077 | 0.807390708 |
| Gm4787 | 0.72759449 | 0.807419675 |
| Gm3642 | 0.631459671 | 0.807419675 |
| Tmem30c | 0.628551142 | 0.807419675 |
| Pcsk6 | 0.596413993 | 0.807419675 |
| Cd93 | 0.180284719 | 0.807419675 |
| Slc37a4 | 0.112647939 | 0.807419675 |
| Scarb2 | 0.066800048 | 0.807419675 |
| Gpaa1 | -0.080302244 | 0.807419675 |
| Dcp2 | -0.086458945 | 0.807419675 |
| Far1 | -0.08879543 | 0.807419675 |
| 4430402I18Rik | -0.316153614 | 0.808294407 |
| Zfp786 | -0.232042631 | 0.808605358 |
| Faap24 | 0.127589051 | 0.808815084 |
| Glis1 | -0.60814682 | 0.808815084 |
| Clic3 | 0.603713304 | 0.808825869 |
| Larp4b | 0.09453921 | 0.808825869 |
| Ppp2r5e | -0.081678478 | 0.808825869 |
| Samd8 | -0.084129132 | 0.808825869 |
| Tns1 | -0.169889644 | 0.808955009 |
| Pyroxd1 | 0.150648691 | 0.808974548 |
| Zfp459 | 0.650908491 | 0.809055827 |
| Gpr135 | 0.327025619 | 0.809055827 |
| Slc35a4 | 0.098997221 | 0.809372196 |
| Rbm15b | -0.089907927 | 0.809372196 |
| Htr4 | -0.576266298 | 0.809731457 |
| Tmem203 | -0.156329495 | 0.809869958 |
| Fam163b | -0.425690804 | 0.810278499 |
| Pla2r1 | 0.901114646 | 0.810395608 |
| Saraf | -0.097431314 | 0.810395608 |
| Ctr9 | 0.0745135 | 0.810991653 |
| Alg6 | -0.152174498 | 0.811040693 |
| Hltf | -0.105134091 | 0.811086553 |
| Asnsd1 | -0.175450149 | 0.811086553 |
| Tacr3 | -0.626372368 | 0.811086553 |
| Hook1 | 0.134721058 | 0.811153878 |
| Simc1 | -0.128050734 | 0.811153878 |
| Zfp940 | -0.138674717 | 0.811153878 |
| 4933408B17Ri | -0.583586579 | 0.811153878 |
| Wdr27 | 0.972175864 | 0.811230488 |
| Nle1 | 0.169305007 | 0.811476282 |
| Calm3 | -0.06016933 | 0.811476282 |
| Snx24 | 0.141155418 | 0.811506496 |
| Smurf2 | 0.104826468 | 0.811506496 |
| Rbpj | -0.08260173 | 0.811506496 |
| Gfm1 | -0.097738915 | 0.811506496 |
| Cfap44 | 0.346234727 | 0.811826664 |
| Hsd3b7 | 0.864210768 | 0.811873877 |
| Trim45 | 0.161039546 | 0.812049997 |
| Cyb5d1 | -0.1646628 | 0.812250002 |
| Aida | 0.136512616 | 0.812320452 |
| Gpr158 | 0.354852821 | 0.81236308 |
| Sp5 | -0.395312162 | 0.812377317 |
| Zfp808 | 0.340649403 | 0.812407398 |
| Xk | -0.230792927 | 0.812703947 |
| Ccne2 | -0.143458992 | 0.813099774 |
| Ccdc96 | 0.347102386 | 0.813244172 |

| | | |
|---|---|---|
| Pcgf1 | 1.293261024 | 0.001070413 |
| Exoc4 | -0.825047334 | 0.001074884 |
| Rreb1 | -1.078017227 | 0.001074884 |
| 3830403N18Rik | 6.599252797 | 0.001076314 |
| Prr32 | 4.623807272 | 0.001076314 |
| Mrps33 | 0.757455525 | 0.001079406 |
| Ttc9b | 0.807874531 | 0.001082329 |
| Prrc2c | 0.616671932 | 0.001082329 |
| Zeb2 | -0.698225964 | 0.001082329 |
| Map3k5 | -2.064881488 | 0.001082329 |
| Zdhhc12 | -1.11516067 | 0.001085131 |
| Pld4 | -1.082648823 | 0.00108613 |
| Sirpb1a | -4.160802587 | 0.001088686 |
| Pfdn4 | 1.09814338 | 0.001097225 |
| Itsn2 | -1.216751315 | 0.001099355 |
| Herpud2 | -1.07765152 | 0.001100876 |
| Tab3 | -1.196537259 | 0.001105451 |
| Aard | 6.538131882 | 0.001110101 |
| Sp8 | 1.351887847 | 0.001110101 |
| Higd1a | 0.831729303 | 0.001110101 |
| Ndufv3 | 0.735430543 | 0.001110101 |
| Chp1 | -0.718400542 | 0.001110101 |
| Adra2c | -3.871353425 | 0.001110101 |
| Cyhr1 | 1.227948657 | 0.001118439 |
| Tmem216 | 1.224834092 | 0.001118977 |
| Morf4l2 | 0.655574924 | 0.001119896 |
| Nrip3 | -1.890759865 | 0.001124689 |
| Traf5 | 4.499263589 | 0.001127987 |
| Themis2 | -1.776806839 | 0.001136326 |
| Fam149a | 2.361876342 | 0.001138904 |
| Dmxl2 | -0.880587737 | 0.001138904 |
| Il1rl2 | -3.529279743 | 0.001141153 |
| Tmem38a | 1.505705338 | 0.001142831 |
| Aoc2 | -2.190075698 | 0.001151642 |
| Coro2a | -1.835145564 | 0.00115582 |
| Rbx1 | 0.576427213 | 0.001157494 |
| Serp2 | 0.922227871 | 0.0011592 |
| Htatsf1 | 0.641795042 | 0.001160654 |
| Chtf18 | -1.705843059 | 0.001160654 |
| Phf14 | 0.867211594 | 0.001160674 |
| Rabac1 | 0.648629491 | 0.001160785 |
| Atp13a1 | -1.085124377 | 0.001167401 |
| Znfx1 | -1.845954489 | 0.001168331 |
| Triobp | -1.053620456 | 0.001170817 |
| Siae | -1.426034663 | 0.001176041 |
| 2410015M20Rik | 0.779129955 | 0.001176096 |
| Acot13 | 1.016062487 | 0.001184627 |
| Osbpl9 | -0.837453877 | 0.001185499 |
| Ndufv2 | 0.68956813 | 0.001186123 |
| D630045J12Rik | -0.963760602 | 0.001186123 |
| Foxf2 | 3.355478882 | 0.00118665 |
| 1700048O20Rik | 1.733987683 | 0.001186992 |
| Mrpl21 | 0.691020386 | 0.001186992 |
| Sdf2l1 | -0.862525046 | 0.001186992 |
| P2ry6 | -1.4143151 | 0.001186992 |
| Tnks | -0.902868015 | 0.001200562 |
| Plcb3 | -1.383662751 | 0.001204313 |
| Rpl38 | 0.634023212 | 0.001205368 |
| Tmem194 | -1.873176238 | 0.00120614 |
| Tnks2 | -0.792286017 | 0.001210921 |
| Foxa1 | -6.635006067 | 0.001210921 |
| Pfn4 | 2.107689684 | 0.001219051 |
| Aftph | -0.806365661 | 0.001221335 |
| Lamc3 | 6.465101874 | 0.001223732 |
| Zfp566 | 2.092612592 | 0.001223732 |
| Gm973 | 2.674458204 | 0.001227449 |
| Dctpp1 | 0.815871785 | 0.001227449 |
| Csgalnact2 | -1.324176136 | 0.001227449 |
| Fhl3 | -1.879169308 | 0.001228039 |
| Ccdc166 | 1.725737895 | 0.001230582 |
| Mylk | 2.81425601 | 0.001235823 |
| Diras2 | 1.762135957 | 0.001238304 |
| Hipk3 | -0.912953837 | 0.001238954 |
| Cbln4 | 6.24682822 | 0.001239387 |
| Naip5 | -1.692166615 | 0.001247632 |
| Fth1 | -1.073723753 | 0.00126162 |
| Tmem43 | -0.95670808 | 0.001271597 |
| Cdkl4 | 3.503152057 | 0.001276424 |
| Xrn1 | 0.915497013 | 0.001277204 |
| Nkap | 0.892488064 | 0.001294228 |
| Ppp1cc | -0.815474905 | 0.00129584 |
| Uqcc3 | 0.815023709 | 0.001298751 |
| Ccrl2 | -1.402286888 | 0.001298751 |
| Mcm5 | -0.903231583 | 0.001301475 |
| Etf1 | -0.734784226 | 0.001304009 |
| Aoah | -2.068959218 | 0.001304055 |
| Srek1ip1 | 1.078082101 | 0.001307638 |
| Psph | 0.833618855 | 0.001308524 |
| Xdh | -2.462203033 | 0.001314578 |
| Atp6v1a | -0.671875552 | 0.001316213 |
| AA467197 | -3.016804716 | 0.001317467 |
| Mpc2 | 0.66205826 | 0.001319724 |
| Hdc | -4.079652642 | 0.001325277 |
| Ly6i | -3.542489107 | 0.001331357 |
| Mfap4 | 1.298663977 | 0.001331666 |
| Zfp661 | -2.21537699 | 0.001336455 |
| Fam20b | -0.898276543 | 0.001339637 |
| Mex3a | 0.725938092 | 0.001347756 |
| Psmb5 | 0.673594927 | 0.001348531 |
| Sumo1 | 0.547436438 | 0.001352322 |
| Aldh2 | -0.706263086 | 0.001352765 |
| Mb21d2 | 2.339068428 | 0.001356033 |
| Ccdc93 | -1.252084882 | 0.001356033 |
| Mlec | -0.84708892 | 0.00136551 |
| Exoc5 | -0.617285754 | 0.001365831 |
| Dach2 | 3.270008334 | 0.001366839 |
| Rps7 | 0.589532543 | 0.001368627 |
| Nid1 | 2.534015081 | 0.001369835 |
| Mov10 | -1.299286908 | 0.001375862 |
| Fam84b | -1.617581722 | 0.001377023 |
| Brinp2 | -1.065980659 | 0.001377874 |
| Il10 | -2.194902556 | 0.001388192 |
| Snrnp27 | 0.916423915 | 0.001389926 |
| Slc9a3r2 | 1.384022759 | 0.001393553 |
| Cdc6 | -2.04005408 | 0.001393553 |
| Mxra8 | 2.871592394 | 0.001398444 |
| Trappc2l | 0.702478342 | 0.001402142 |
| Rab10 | -0.66346392 | 0.001404891 |
| Nfkb2 | -1.764013693 | 0.001405491 |

| | | |
|---|---|---|
| Eif2ak4 | -0.739585367 | 0.102953682 |
| Fxyd7 | 0.872869806 | 0.102975147 |
| Lnp | 0.470455367 | 0.10300392 |
| Pde3b | -0.553915671 | 0.103025708 |
| Tomm22 | 0.312307958 | 0.103057853 |
| Lims2 | 4.963550429 | 0.103098323 |
| Map6d1 | 3.009286691 | 0.103098323 |
| Armc9 | 0.675359387 | 0.103253506 |
| Zkscan5 | -1.066858908 | 0.103332452 |
| Hist1h2bg | -1.291795587 | 0.103332452 |
| Thap3 | 0.704159237 | 0.103570392 |
| Tmem191c | 1.811385786 | 0.103734856 |
| Rwdd3 | 0.881153406 | 0.10386714 |
| Neto2 | -0.659859175 | 0.103994469 |
| Anapc16 | 0.384980701 | 0.104067797 |
| Kcnh4 | 1.666071278 | 0.104165204 |
| Prmt1 | 0.412560742 | 0.104165204 |
| Sgpl1 | -0.593759522 | 0.104165204 |
| Scimp | -1.693686892 | 0.10422084 |
| Mycl | -0.499410121 | 0.104390402 |
| Snrpb | 0.408992009 | 0.104433631 |
| Clec4a2 | -0.783538155 | 0.104683351 |
| Gale | -0.779876475 | 0.104706429 |
| Ube2v2 | 0.520092874 | 0.104850955 |
| Cln6 | -0.818225223 | 0.104871132 |
| Cpeb2 | -0.660380275 | 0.105009311 |
| Gm3696 | 1.820340954 | 0.10523919 |
| Srcap | -0.585886232 | 0.105378266 |
| Cenpt | 0.503331625 | 0.105492851 |
| Mdh1 | 0.317744381 | 0.105492851 |
| Chmp6 | 0.516629254 | 0.105522165 |
| Pdia6 | -0.435519931 | 0.105558519 |
| Nifk | 0.489281606 | 0.105647479 |
| Zfp420 | 0.97408006 | 0.105648336 |
| Tmem87b | -0.700024961 | 0.105648336 |
| Ifitm6 | -2.669245553 | 0.105666557 |
| Luc7l2 | -0.320808495 | 0.105705713 |
| BC100530 | -2.154005021 | 0.105877413 |
| Rbbp5 | 0.600846188 | 0.105947557 |
| Tha1 | -1.496402969 | 0.106100585 |
| Pigf | 0.703065549 | 0.106142474 |
| Fbxo17 | 1.043481301 | 0.106224728 |
| Nrcam | -0.603216311 | 0.106224728 |
| Slc35e4 | 0.957064644 | 0.106248337 |
| BC016579 | 5.293173062 | 0.106255538 |
| Wdr89 | 0.595372831 | 0.106256475 |
| Sctr | 4.078673916 | 0.10629793 |
| G6pdx | -0.477808658 | 0.106297931 |
| Myt1l | -0.424664036 | 0.106348525 |
| Mpv17l | 0.835954934 | 0.106476219 |
| Arhgdia | -0.393075732 | 0.106476219 |
| Gtf2i | 0.393225536 | 0.1066086 |
| Nanp | 1.92080487 | 0.106631536 |
| Ncaph | -0.50442272 | 0.106631536 |
| Plekhj1 | 0.402636775 | 0.106647919 |
| Grin3a | 1.017478238 | 0.106861007 |
| Gabra1 | 1.332695934 | 0.106930038 |
| Col19a1 | -2.726371036 | 0.106930038 |
| 3110040N11Rik | 0.497178482 | 0.106960518 |
| Srp19 | 0.378226033 | 0.106967841 |
| Elp5 | 0.406559834 | 0.107121765 |
| Bod1l | 0.448606957 | 0.107129473 |
| Dguok | 0.471391979 | 0.107209517 |
| Psme2b | -0.59318577 | 0.10728859 |
| Foxo6 | -0.797887105 | 0.107364572 |
| Rhebl1 | 0.649903019 | 0.107421028 |
| Sec22a | 0.509625843 | 0.107421028 |
| Mc5r | 4.681957322 | 0.107470599 |
| Wscd1 | 0.728929241 | 0.107470599 |
| Csf3 | -3.922133469 | 0.107470599 |
| Lrrc18 | 4.90314124 | 0.107875816 |
| Mrs2 | -0.711131186 | 0.107875816 |
| Strbp | -0.350944742 | 0.108117127 |
| Tap1 | -0.989063002 | 0.108227514 |
| Fam101b | 1.764081276 | 0.10841302 |
| Rxfp1 | -5.282300604 | 0.108460563 |
| Slc25a11 | -0.40542538 | 0.108494325 |
| Rexo4 | -0.452391563 | 0.108907896 |
| Zfp729b | 0.709500718 | 0.109034125 |
| Atxn7l1 | -0.58226623 | 0.109036429 |
| Hipk2 | 0.495201166 | 0.109135215 |
| Dyrk1b | -0.873296574 | 0.109135215 |
| Snrnp25 | 0.449438079 | 0.109211288 |
| Ano4 | 1.249336087 | 0.10932511 |
| Mcur1 | -0.650101162 | 0.10932511 |
| Ccdc77 | 0.874990576 | 0.109332262 |
| Ftsj1 | 0.494906455 | 0.109559061 |
| Tsr2 | 0.674304493 | 0.109586887 |
| Vill | 2.422560421 | 0.109660925 |
| Tcf12 | 0.34638493 | 0.109723096 |
| Med28 | 0.344484054 | 0.109861083 |
| Pcyt2 | 0.398040216 | 0.109970368 |
| Zfp276 | -0.899081611 | 0.110018739 |
| Mark3 | -0.443269707 | 0.110131255 |
| Vasn | -1.339894526 | 0.110171783 |
| Slc25a13 | 0.883318858 | 0.110273855 |
| Amotl2 | 0.532695976 | 0.110287747 |
| Cxcr3 | -2.337238586 | 0.110513476 |
| Rpgr | 0.554778378 | 0.110618033 |
| Nck2 | -0.60750722 | 0.110718345 |
| Bzrap1 | -0.670011679 | 0.110718345 |
| 2810474O19Rik | -0.629930648 | 0.110718985 |
| Mrpl51 | 0.402545331 | 0.110731419 |
| Lhx9 | 1.827417381 | 0.110914853 |
| Phf10 | 0.846414399 | 0.110936537 |
| Mboat7 | -0.608209704 | 0.110936537 |
| Qtrt1 | 0.563507229 | 0.11112664 |
| Pip5k1c | -0.417021088 | 0.11112664 |
| Irs2 | 0.572825294 | 0.111128608 |
| C1qbp | 0.320938194 | 0.111128608 |
| Fbxl20 | 0.56165507 | 0.111309358 |
| Mafk | -0.854251081 | 0.111309358 |
| Nppa | -1.915195827 | 0.111309358 |
| Slc6a17 | -0.748846184 | 0.111374231 |
| Cpm | 1.538537008 | 0.111426966 |
| Cdc42ep3 | 0.595736241 | 0.111426966 |
| Rab3il1 | -0.693875406 | 0.11146573 |
| Map6 | 0.576447082 | 0.11149425 |
| Zkscan6 | -0.717343808 | 0.111498024 |

| | | |
|---|---|---|
| Tmf1 | 0.235931784 | 0.42612006 |
| Cadm1 | 0.284171799 | 0.426461125 |
| Dvl2 | -0.323436583 | 0.426476552 |
| Erich6 | 2.367010827 | 0.426616085 |
| Brat1 | -0.459727484 | 0.426647102 |
| Eml6 | -0.565029204 | 0.426647102 |
| Has3 | -1.101627034 | 0.426825129 |
| Dnah1 | 0.721207466 | 0.427250037 |
| Drosha | -0.233508855 | 0.427326985 |
| Tceb3 | -0.297388513 | 0.427367656 |
| Tufm | 0.222359701 | 0.427530678 |
| Farsa | -0.185951731 | 0.427811326 |
| Gm1604b | 2.831222983 | 0.427966042 |
| Ccnb2 | 0.303569584 | 0.427966042 |
| Galnt2 | -4.301892016 | 0.427966042 |
| Impad1 | 0.275078135 | 0.428038554 |
| Ccdc47 | -0.231310444 | 0.428076918 |
| Fkbp6 | 3.807727853 | 0.428118473 |
| Capns1 | 0.328271519 | 0.428137227 |
| Eif2ak1 | -0.248645635 | 0.428137227 |
| Myof | -0.975468853 | 0.428137227 |
| Murc | -1.001064252 | 0.428173319 |
| Cttn | 0.193050062 | 0.428361775 |
| Atp13a4 | -3.284077568 | 0.428434077 |
| Aloxe3 | 1.833790565 | 0.428767706 |
| Aass | 1.210778851 | 0.429075163 |
| Ap3s1 | 0.203193547 | 0.429075163 |
| Zmiz1 | 0.211043564 | 0.429398316 |
| Lipc | -1.203088566 | 0.429398316 |
| Mamld1 | -0.446123828 | 0.429416332 |
| Hook2 | -0.435199625 | 0.429425281 |
| B4galt3 | -0.284015349 | 0.429476127 |
| Ldhc | 2.89535298 | 0.429677131 |
| Epc2 | 0.309852454 | 0.429677131 |
| Kcnh6 | -1.144786087 | 0.42987921 |
| Entpd6 | -0.304395555 | 0.429898803 |
| Olfr1233 | -3.520381776 | 0.430006451 |
| Rpl29 | 0.345796779 | 0.430381194 |
| Gm15440 | -1.100910766 | 0.430381194 |
| Lrch4 | -1.899996873 | 0.430381194 |
| Hdx | 0.476891969 | 0.430452822 |
| Eno4 | -0.74919258 | 0.430759994 |
| Dscr3 | -0.253957698 | 0.430760404 |
| Mapre3 | 0.287233653 | 0.43077692 |
| Ptcd1 | -0.32336781 | 0.43077692 |
| Zfand2a | -0.337936618 | 0.43077692 |
| Hydin | 1.241981913 | 0.43081587 |
| Qrich1 | -0.190285956 | 0.43083088 |
| Kctd12b | -0.933826667 | 0.4309319 |
| Zbtb32 | -1.94963207 | 0.431114505 |
| Clmp | -0.313695684 | 0.431347303 |
| Tnfaip6 | 0.860834774 | 0.431684365 |
| Zfp930 | 0.623594241 | 0.431684365 |
| Gm16286 | -0.216561802 | 0.431684365 |
| Srpx2 | 3.734701827 | 0.43179191 |
| Tmed10 | -0.204755141 | 0.431901599 |
| Cyp4f17 | 1.191120849 | 0.432206206 |
| Vmn2r91 | 3.795307843 | 0.432501574 |
| Fam174b | -0.437153349 | 0.432720676 |
| Ceacam19 | -1.476623916 | 0.432720676 |
| Fer1l6 | 2.911477562 | 0.432738364 |
| Atp8a2 | -0.568743354 | 0.432738364 |
| Spin1 | 0.196451036 | 0.432756546 |
| Abhd17c | 0.328183736 | 0.433004021 |
| Zkscan17 | -0.315573721 | 0.43324071 |
| Tuba1b | 0.188778943 | 0.433271985 |
| Scaf1 | -0.282796491 | 0.43339468 |
| Camkk2 | -0.236024379 | 0.433512712 |
| Cnot3 | -0.276574145 | 0.433512712 |
| Gtf2f1 | -0.224637928 | 0.433793875 |
| Fundc1 | 0.220919727 | 0.433962545 |
| Phactr3 | 0.38783733 | 0.43404728 |
| Klhl41 | 2.053173995 | 0.434252419 |
| Gins3 | 0.94039711 | 0.434252419 |
| Baz2a | 0.296620951 | 0.434252419 |
| BC005561 | 0.204990569 | 0.434252419 |
| Mecp2 | -0.225167741 | 0.434252419 |
| Ccdc141 | 0.8253853 | 0.434261179 |
| Wdsub1 | 0.628881147 | 0.434327679 |
| Gm9869 | -2.337039589 | 0.434354426 |
| Mob3b | -0.493564354 | 0.434364369 |
| Vmn1r4 | 4.07622504 | 0.434780129 |
| Wwp1 | 0.29008888 | 0.434869052 |
| Tspyl1 | 0.187202272 | 0.434869052 |
| Taf6 | -0.270720415 | 0.434988464 |
| Atg4a | 0.453170517 | 0.435217561 |
| Rfx5 | -0.628662297 | 0.435217561 |
| Kif3b | -0.257041187 | 0.435460731 |
| Kat7 | -0.272268566 | 0.435485877 |
| Apcdd1 | -0.574274168 | 0.435485877 |
| Shcbp1 | 0.305311647 | 0.435687342 |
| Wwc2 | -0.391628247 | 0.435952552 |
| Nt5dc1 | -0.503197968 | 0.435987367 |
| Igsf10 | 0.597529182 | 0.435996051 |
| 9130023H24Rik | 0.514229739 | 0.437430946 |
| Usp42 | 0.283607967 | 0.437439482 |
| 9930111J21Rik2 | -0.911385101 | 0.437606575 |
| 9430069I07Rik | -2.36975654 | 0.437606575 |
| Grin2d | 1.131167744 | 0.437880028 |
| Dpf2 | -0.233102952 | 0.437883634 |
| Lpl | -0.416020068 | 0.437914283 |
| Hist1h2bh | 2.246934171 | 0.437936401 |
| Disp1 | 0.670455157 | 0.437936401 |
| Frs2 | -0.327829617 | 0.438164076 |
| Ptprk | -0.410421609 | 0.438412429 |
| Leprotl1 | 0.214498979 | 0.438561169 |
| Ubiad1 | -0.288445035 | 0.438662779 |
| Elmod2 | 0.304742236 | 0.438757124 |
| Vsx1 | 3.735759443 | 0.438857056 |
| Gm14308 | 2.748624469 | 0.438857056 |
| Gm10031 | 0.267869631 | 0.438857056 |
| Zfp579 | 0.263732245 | 0.438857056 |
| Map1lc3b | 0.160234574 | 0.438857056 |
| Ubac2 | -0.277006353 | 0.438857056 |
| Gpd2 | -0.311242526 | 0.438857056 |
| Bmp1 | -0.34905314 | 0.438857056 |
| Ddx11 | -0.662790536 | 0.438857056 |
| Sdccag8 | 0.266485561 | 0.438858002 |
| Zfp398 | -0.294315094 | 0.439005186 |

| | | |
|---|---|---|
| Lilra5 | 0.496972686 | 0.81344656 |
| Exosc9 | 0.111978938 | 0.813760756 |
| Chac1 | -0.242384216 | 0.814000185 |
| Tacr1 | 0.861411562 | 0.814251443 |
| Bglap | 0.654271656 | 0.814332927 |
| Sntg2 | 0.649939202 | 0.814332927 |
| Ppil2 | -0.063515176 | 0.814332927 |
| Gm16380 | -0.580899875 | 0.814332927 |
| Topors | 0.081523637 | 0.814389738 |
| Sepn1 | -0.145939141 | 0.814439375 |
| Anks6 | -0.236974229 | 0.814993183 |
| Gm29666 | -0.265742951 | 0.815068011 |
| Ddx19a | -0.109391283 | 0.815176815 |
| Slc35e1 | 0.184693416 | 0.815183212 |
| Xylb | 0.169932364 | 0.815183212 |
| Mettl14 | 0.110365713 | 0.815211022 |
| Lmbrd2 | -0.108176479 | 0.815465766 |
| Cdkn2d | 0.080367595 | 0.815809955 |
| Gnas | 0.170396228 | 0.816026348 |
| Clgn | -0.246090044 | 0.816026348 |
| Chd7 | -0.09665008 | 0.816238726 |
| Sema3f | -0.203846898 | 0.816238726 |
| Dhodh | 0.145422394 | 0.816444707 |
| Hhat | 0.270434928 | 0.816832257 |
| Sdc4 | -0.257341976 | 0.816852113 |
| 3110001I22Rik | 0.174324685 | 0.816885915 |
| Cped1 | -0.264177527 | 0.817656583 |
| Acvr1 | 0.156916482 | 0.81767168 |
| Hist1h2bb | -0.380489051 | 0.81794698 |
| Rom1 | 0.432738498 | 0.818190563 |
| Ppp2r1a | -0.056813411 | 0.818190563 |
| Gpr161 | -0.148409153 | 0.818190563 |
| Iqck | 0.373780077 | 0.818413435 |
| Fh1 | 0.070969771 | 0.818594508 |
| Ulk4 | 0.182800613 | 0.818625325 |
| Spns3 | -0.477216384 | 0.81876466 |
| Mad2l1 | 0.114186287 | 0.818830571 |
| Vps41 | 0.068316569 | 0.818872502 |
| A730015C16Ri | -0.729407373 | 0.818872502 |
| Snapc1 | 0.109893047 | 0.818926739 |
| Dhps | 0.077735082 | 0.818926739 |
| Tcf20 | -0.079819819 | 0.818926739 |
| Spata4 | 1.176905817 | 0.819144906 |
| AA986860 | 0.636966548 | 0.819144906 |
| Asl | -0.102914668 | 0.819144906 |
| B3glct | -0.117652219 | 0.819144906 |
| Ccdc57 | -0.190006085 | 0.819144906 |
| Rnf113a2 | -0.085323929 | 0.819519652 |
| Snx17 | -0.065743015 | 0.819590064 |
| Pisd | -0.102272477 | 0.819590064 |
| Slc22a21 | 0.220788316 | 0.819825986 |
| Dpf1 | -0.088297154 | 0.819825986 |
| Dfnb59 | -0.481006981 | 0.819825986 |
| Ap1s2 | 0.084359147 | 0.820121748 |
| Shisa2 | -0.179827541 | 0.820121748 |
| Ccnd1 | -0.086033085 | 0.820445748 |
| Txndc11 | -0.105066992 | 0.820546638 |
| Dis3l | 0.097542338 | 0.820611037 |
| Gm21092 | 0.090912207 | 0.820716682 |
| Nhlrc3 | -0.146318509 | 0.820781092 |
| Rnf8 | -0.188737769 | 0.820781092 |
| Zfr2 | 0.134987557 | 0.821418874 |
| Atg101 | 0.079050793 | 0.821601793 |
| Rps2 | 0.301810651 | 0.821983762 |
| Naa10 | 0.092642546 | 0.8229232 |
| Nrxn2 | 0.077021028 | 0.823004624 |
| Apba1 | -0.110837268 | 0.823004624 |
| Fbxo5 | 0.13144823 | 0.823153268 |
| 4931406C07Ri | 0.110427704 | 0.823320708 |
| Olfr120 | 1.036871929 | 0.823342776 |
| Efhc2 | 0.414717899 | 0.823342776 |
| Ppp1r26 | 0.165452747 | 0.823342776 |
| Hist1h1d | 0.15724833 | 0.823342776 |
| Mta1 | 0.132720747 | 0.823342776 |
| Fgfr1 | 0.104077515 | 0.823342776 |
| Mrpl37 | 0.076169049 | 0.823342776 |
| Gmfb | 0.069892538 | 0.823342776 |
| Dnpep | 0.061784073 | 0.823342776 |
| Ap1m1 | -0.063379046 | 0.823342776 |
| Plaa | -0.075463268 | 0.823342776 |
| Nfyc | -0.077135551 | 0.823342776 |
| Wipf2 | -0.136284583 | 0.823342776 |
| Kcnd3 | -0.167781607 | 0.823342776 |
| Trpc7 | -0.362943457 | 0.823342776 |
| Gm6563 | 0.062839681 | 0.823463581 |
| Fuk | 0.281776874 | 0.823564979 |
| Ebag9 | 0.119345673 | 0.823564979 |
| Sh3gl3 | 0.111334093 | 0.823629353 |
| Uba5 | 0.062088274 | 0.823857579 |
| Aurka | 0.110238318 | 0.824448636 |
| Arl6ip5 | -0.06530789 | 0.824448636 |
| Ccl8 | -0.522257584 | 0.824448636 |
| 4930558K02Ri | -0.740877621 | 0.824448636 |
| Fibcd1 | 0.214703293 | 0.824515761 |
| Rsad1 | -0.161571469 | 0.824515761 |
| Tjap1 | -0.1532576 | 0.824585309 |
| Ilf3 | 0.062563163 | 0.824692205 |
| Fam234a | 0.118403473 | 0.825184714 |
| Avl9 | -0.070318689 | 0.825184714 |
| Zswim5 | -0.117779653 | 0.825184714 |
| Cpne6 | 0.321823105 | 0.825208907 |
| Exog | -0.125059025 | 0.825351661 |
| Rab4b | -0.440246736 | 0.825422465 |
| Vpreb1 | 0.65988416 | 0.825447654 |
| Senp6 | -0.071744674 | 0.825989068 |
| Fam188a | 0.087597647 | 0.826045435 |
| Sp1 | -0.090588919 | 0.826153537 |
| Gm10654 | -0.552733574 | 0.82637367 |
| Cd3eap | 0.094591685 | 0.826623049 |
| Gnl1 | 0.105799267 | 0.826893102 |
| Utp3 | 0.07322893 | 0.826893102 |
| Ccdc159 | -0.230187534 | 0.826893102 |
| Gm5819 | 1.024564099 | 0.82695968 |
| Prrt4 | 0.280151418 | 0.827113275 |
| Slc25a25 | 0.108208235 | 0.827113275 |
| Zbtb46 | 0.191823732 | 0.827291709 |
| Ash1l | -0.064190948 | 0.827291709 |
| Rad54l2 | -0.09089891 | 0.827291709 |
| B230307C23Ri | -0.151915692 | 0.827291709 |

| | | |
|---|---|---|
| Fcfl | 0.72052906 | 0.001407489 |
| Rpl35 | 0.71020506 | 0.001408788 |
| Zfp946 | 1.1699722 | 0.001413469 |
| Tfcp2 | 1.140985547 | 0.001413469 |
| Mpped2 | 0.736251122 | 0.001413469 |
| Eif4enif1 | -0.956190338 | 0.001413469 |
| Zbtb49 | -1.43621466 | 0.001413469 |
| Pou3f1 | -1.454394032 | 0.001413469 |
| Krt10 | 1.154911007 | 0.001414675 |
| Pnck | 0.793057976 | 0.001414854 |
| Adamts3 | -1.786102725 | 0.001418175 |
| Cobll1 | 3.647580312 | 0.001418186 |
| Lsm8 | 0.644602106 | 0.001418186 |
| Rps25 | 0.753215591 | 0.001421479 |
| E430025E21Rik | -0.64547299 | 0.001421479 |
| Cdk2 | -0.958148787 | 0.001421479 |
| Klhl8 | -1.094227334 | 0.001421479 |
| Jak3 | -1.87388356 | 0.001421479 |
| Ptgfrn | 1.751623642 | 0.001423087 |
| Coq9 | 1.015897722 | 0.001423785 |
| Ap2a1 | -0.899561007 | 0.001423785 |
| Mgat5 | -0.854378139 | 0.001425252 |
| Rapgef1 | -0.831073762 | 0.001429622 |
| Cuedc2 | 0.614925713 | 0.001432092 |
| Hmgn5 | 1.084524555 | 0.001437518 |
| Tac2 | 3.033397288 | 0.001439918 |
| Ripk2 | -1.410933913 | 0.001439918 |
| Polr2j | 0.800158141 | 0.001452871 |
| Ndufs6 | 0.621227648 | 0.001461872 |
| Eml3 | -1.198706967 | 0.001461872 |
| Chst12 | -1.033242904 | 0.001462423 |
| B430306N03Rik | -2.590487831 | 0.001462423 |
| Itga5 | -2.950389933 | 0.001462868 |
| Fgr | -3.036784932 | 0.001465721 |
| Rabggtb | 1.086899592 | 0.001466718 |
| Pthlh | 6.342488074 | 0.001467674 |
| Nicn1 | 0.896344309 | 0.001467724 |
| Dhdh | -2.292637018 | 0.001467902 |
| Dcaf17 | 1.243815862 | 0.001474246 |
| AK157302 | 2.120043436 | 0.001482315 |
| Fstl5 | 1.116718792 | 0.001482315 |
| Frmd4a | -0.855425938 | 0.001485798 |
| Pstpip2 | -2.527267832 | 0.00149338 |
| Hist1h1b | 0.910994571 | 0.001497447 |
| Asb5 | -5.158111753 | 0.001497447 |
| Tifa | -0.955361887 | 0.001504544 |
| Pbk | 0.93916596 | 0.001505788 |
| Nfkb1 | -1.121762821 | 0.001508936 |
| Arl16 | 0.977061962 | 0.00151046 |
| Kpnb1 | 0.835672992 | 0.001526296 |
| Polr2d | 0.801587819 | 0.001526296 |
| Nacc1 | -1.21411856 | 0.001526309 |
| Hint2 | 0.863970999 | 0.001527188 |
| AW209491 | 1.249649721 | 0.00154635 |
| Col15a1 | 4.943481408 | 0.001552372 |
| Zfp882 | 1.772251733 | 0.001560751 |
| Smc1a | 0.630317149 | 0.001562242 |
| Gjd2 | 2.939865383 | 0.001563716 |
| Cnot11 | -1.245014152 | 0.001564031 |
| Zfp277 | 1.121913901 | 0.001566886 |
| Zer1 | -0.881123267 | 0.001566886 |
| Sox6 | 0.962768191 | 0.001567649 |
| Tbcb | 0.563508614 | 0.001569295 |
| Mpc1 | 0.711527466 | 0.001575622 |
| Ctxn1 | 0.564507487 | 0.001577444 |
| 9430020K01Rik | 2.287043891 | 0.001586192 |
| Actr2 | -0.74159202 | 0.001588186 |
| Ndufs4 | 0.672124653 | 0.001589512 |
| Zfp30 | 1.154475083 | 0.001593491 |
| Cd300ld | -3.279480531 | 0.001593491 |
| Lhfp | 1.629244986 | 0.001594716 |
| Gpbp1 | 0.599806295 | 0.001594716 |
| Sbf2 | -0.778604616 | 0.001605176 |
| Casc3 | -0.819371665 | 0.001605176 |
| Crat | -0.749924961 | 0.001607965 |
| Sez6l2 | -0.804035658 | 0.00161631 |
| Cd274 | -3.145474523 | 0.00161631 |
| Sdc1 | 1.495828796 | 0.001617237 |
| Efna1 | 2.404137576 | 0.001618245 |
| Psmb2 | 0.62032545 | 0.001618661 |
| Pcdh17 | 1.294717839 | 0.001619234 |
| Ccne1 | -1.402347067 | 0.001619418 |
| Sf3b6 | 0.638437964 | 0.001620239 |
| Spock1 | -1.0957113 | 0.001620239 |
| Cbln3 | -6.763965919 | 0.001620412 |
| Arhgef11 | -1.114066556 | 0.001628268 |
| Smim11 | 0.874044934 | 0.001643188 |
| Calca | 6.398172451 | 0.001645147 |
| Ptprn2 | -1.3640251 | 0.001647346 |
| Mrps28 | 0.923595891 | 0.001656181 |
| 3110043O21Rik | -0.784718348 | 0.001661697 |
| Mmp11 | 1.875655506 | 0.001664585 |
| Zmym5 | 0.890517657 | 0.001668022 |
| Litaf | -0.885971952 | 0.001680824 |
| Rpl22l1 | 0.619796324 | 0.001693888 |
| Nuak2 | -1.196292502 | 0.00169498 |
| Rpp25 | 1.975905672 | 0.001698909 |
| Prmt8 | 1.765679432 | 0.001701618 |
| E030030I06Rik | 6.128037218 | 0.001702291 |
| Lefty2 | 4.222487683 | 0.001702291 |
| Commd1 | 0.941797595 | 0.001704658 |
| Arid4b | 0.715687963 | 0.00171183 |
| Tpd52l1 | 1.752080929 | 0.001718335 |
| Bbs10 | -3.079485874 | 0.001719272 |
| Uhrf2 | -0.624683252 | 0.001719731 |
| Piwil4 | 6.269643154 | 0.001724886 |
| Cdca8 | 0.871400121 | 0.001727579 |
| Ctnnbl1 | 0.768528105 | 0.001727675 |
| Mrgpra2b | -6.047046214 | 0.001728361 |
| Vwa1 | 2.779649328 | 0.001733649 |
| Cdk16 | 0.599482489 | 0.00173742 |
| Dhx8 | -0.76317464 | 0.001739175 |
| Relt | -1.951227317 | 0.001739175 |
| Eid2 | 1.976829053 | 0.001741609 |
| Mfsd8 | -1.01230954 | 0.001741609 |
| Lrp5 | -1.361134484 | 0.001742955 |
| Cdk4 | 0.637254276 | 0.001743833 |
| Kcnq2 | -1.009939612 | 0.00176692 |
| Spsb1 | -1.755255167 | 0.001777641 |

| | | |
|---|---|---|
| Nedd1 | 0.649956634 | 0.111521868 |
| Angptl1 | -1.8858362 | 0.111521868 |
| Cnr2 | -4.432835516 | 0.111521868 |
| Kif11 | 0.577255271 | 0.111570786 |
| Adgrf3 | 3.285303395 | 0.111701427 |
| Dstn | 0.316981102 | 0.11181179 |
| Xaf1 | -1.045369381 | 0.111816523 |
| Arrb1 | -0.717162775 | 0.111820464 |
| Eif5b | 0.366414954 | 0.111843469 |
| Sap30bp | 0.454979261 | 0.111882677 |
| Syt9 | -1.038114176 | 0.111882677 |
| Narfl | -0.473511322 | 0.111906536 |
| Exosc2 | 0.518091468 | 0.111953492 |
| Nptxr | 1.102878634 | 0.112125431 |
| Acadl | 0.481310825 | 0.11218655 |
| Cpne9 | -1.576515709 | 0.11218655 |
| Acat2 | 0.375301225 | 0.112240478 |
| Rhof | -0.968841576 | 0.112308472 |
| Josd1 | -0.527144009 | 0.112338835 |
| Wfikkn2 | 2.316149786 | 0.112354958 |
| Rapgef6 | 0.429891823 | 0.112354958 |
| Adgrb1 | 1.052741927 | 0.11236557 |
| S100a6 | -1.800429963 | 0.11236557 |
| Pafah2 | 1.357729757 | 0.11250864 |
| Zfp516 | -0.616802552 | 0.11250864 |
| Tox4 | -0.369692255 | 0.112516572 |
| Socs5 | 0.708587324 | 0.112528364 |
| Myl12a | -0.48617981 | 0.112528364 |
| Il4 | 2.406076539 | 0.112549208 |
| Kank2 | -1.246170675 | 0.112549208 |
| Map2k2 | 0.329214956 | 0.112606409 |
| Msmo1 | -0.420780268 | 0.112661791 |
| Ascl2 | -1.253529478 | 0.112661791 |
| Smc5 | 0.448524632 | 0.112676709 |
| Usp34 | 0.441655466 | 0.112728005 |
| Tspan15 | 1.171791736 | 0.112728097 |
| Brip1 | -0.986194049 | 0.112744135 |
| Zc4h2 | 0.506231704 | 0.112789393 |
| Rap2c | -0.652763179 | 0.112789393 |
| Efcc1 | 2.956187748 | 0.11283209 |
| Eif4g1 | 0.348072525 | 0.11303318 |
| Araf | 0.389285006 | 0.113119048 |
| Als2cl | -2.671580197 | 0.113119048 |
| Hal | -3.458451617 | 0.113189124 |
| Cyp3a13 | -4.087104405 | 0.113196801 |
| Amica1 | -1.765750263 | 0.113274722 |
| Tcp11 | -2.350585498 | 0.113345212 |
| Ptpn11 | 0.389321506 | 0.113507986 |
| Rbbp9 | 0.703080553 | 0.113556908 |
| Il11 | 4.624135696 | 0.113614624 |
| Sdhd | 0.414231047 | 0.113614624 |
| Fkbp2 | 0.411957006 | 0.113614624 |
| Chac2 | 0.893000346 | 0.113719005 |
| Cdk1 | 0.503444407 | 0.113779378 |
| Tbc1d8b | 1.577871803 | 0.113784002 |
| Hipk1 | 0.688091323 | 0.113788114 |
| Rbmx2 | 0.543730484 | 0.113929186 |
| Nkd2 | 1.265161299 | 0.113957735 |
| Atf7ip2 | -3.654955883 | 0.113959955 |
| Itpka | 4.860915459 | 0.113960917 |
| Ptpdc1 | -0.881386675 | 0.114003624 |
| Zdhhc23 | -1.993537698 | 0.114185788 |
| Mmab | 0.800657749 | 0.11432862 |
| Mrvi1 | 4.830388703 | 0.114530746 |
| Egfem1 | 1.06723698 | 0.114530746 |
| Slc7a10 | 2.220671971 | 0.114606197 |
| Tma16 | -0.616824622 | 0.11470073 |
| Ptch2 | 1.889547344 | 0.114742766 |
| Kif18a | 0.858658853 | 0.114742766 |
| Irx2 | -1.634755244 | 0.114742766 |
| Lama1 | 1.378256014 | 0.114763003 |
| Camk2n2 | 0.999534699 | 0.114763003 |
| Styx | 0.571744412 | 0.114763003 |
| Smndc1 | 0.448913016 | 0.114763003 |
| Tmem74 | -0.992908338 | 0.114763003 |
| Wasf3 | 0.494923221 | 0.114853171 |
| Zfyve16 | 0.659510052 | 0.114951978 |
| Grpel1 | 0.401092174 | 0.114973948 |
| Gdf15 | 3.981068164 | 0.11501082 |
| Ppan | 0.682218429 | 0.115050508 |
| Cap2 | -0.903981262 | 0.115105519 |
| Col11a1 | 0.691090293 | 0.115131761 |
| Stk39 | 0.513817921 | 0.115185035 |
| Nudcd2 | 0.405007562 | 0.115185976 |
| Parvg | -0.680493822 | 0.115185976 |
| Vps35 | -0.359331864 | 0.115456431 |
| Fjx1 | 0.652233744 | 0.115521358 |
| Dtx3 | 0.389195296 | 0.115946927 |
| Atp6v1b2 | -0.400903759 | 0.115946927 |
| Zfp37 | 0.722219256 | 0.115954959 |
| Gm21762 | -5.584303075 | 0.116088107 |
| Cspg4 | 2.439645078 | 0.116107152 |
| Plekhh2 | 1.845576865 | 0.116108795 |
| Tmx4 | 0.387273038 | 0.11626824 |
| Oser1 | -0.520593612 | 0.116289207 |
| Lrrc48 | 1.928568657 | 0.116424309 |
| Hdac5 | -0.405171994 | 0.116521275 |
| Rhbdl3 | 0.778267204 | 0.116587165 |
| Cops5 | 0.365398753 | 0.116602752 |
| Gm11127 | -2.245596749 | 0.116602752 |
| Ppp2cb | 0.79658893 | 0.116651864 |
| Hist1h1c | -0.698256496 | 0.116651864 |
| Proser2 | 2.743545417 | 0.11674815 |
| Folh1 | 3.198575767 | 0.116784635 |
| Prpf40b | 0.600789892 | 0.116937722 |
| Pigt | -0.325073658 | 0.116937722 |
| Slc9a6 | 0.513565683 | 0.117008689 |
| 1110004F10Rik | 0.322828017 | 0.117014238 |
| Lrrc63 | -5.590241484 | 0.117328565 |
| Brd9 | 0.438559863 | 0.117386929 |
| Trem2 | 0.539624224 | 0.117415441 |
| Cdkn1c | 0.461767795 | 0.11804444 |
| Atxn3 | -0.735006458 | 0.11804444 |
| Nit2 | 0.566848226 | 0.11809748 |
| Ubr5 | 0.400086069 | 0.11809748 |
| Hsf5 | -5.584266508 | 0.11809748 |
| Stx18 | -0.477757041 | 0.118123866 |
| Il1f9 | -2.543339394 | 0.118123866 |
| Chrdl1 | 3.787352015 | 0.118146901 |

| | | |
|---|---|---|
| Gal3st3 | 0.519118563 | 0.439387835 |
| Rbm10 | 0.241841792 | 0.439387835 |
| Flt3 | 3.070044914 | 0.439434577 |
| Ccdc148 | 0.751200193 | 0.43975935 |
| Gpalpp1 | 0.252916089 | 0.439829687 |
| Ino80 | 0.460218681 | 0.439992032 |
| Prr13 | -0.2331574 | 0.439992032 |
| Troap | 0.552428887 | 0.440214974 |
| 2010106E10Rik | -4.255740799 | 0.440214974 |
| Sbk2 | -4.255740799 | 0.440214974 |
| Lrrc4 | -0.59378677 | 0.440224196 |
| Nudt10 | 0.269690564 | 0.440286549 |
| Rxfp3 | 2.320230192 | 0.440854474 |
| Smtnl1 | -4.234247935 | 0.440987464 |
| Tcp11l2 | -0.312843222 | 0.441052108 |
| Rgl3 | 2.787008655 | 0.441173182 |
| Abcd3 | 0.292188016 | 0.441173182 |
| Gm10645 | 3.702960298 | 0.441337436 |
| Hebp2 | 0.656838049 | 0.441356688 |
| Syngap1 | -0.507199332 | 0.441356688 |
| Thsd7b | -0.561444164 | 0.441356688 |
| Slc9b2 | -2.323734522 | 0.441356688 |
| Myom1 | 1.040630169 | 0.441392807 |
| Rrp7a | -0.217241703 | 0.441406783 |
| Tbc1d9b | -0.256919026 | 0.441492131 |
| Spsb3 | -1.016149871 | 0.441492131 |
| Hist1h4a | 2.756213324 | 0.441843414 |
| Tsta3 | 0.207929186 | 0.441910339 |
| Ctns | -0.468592129 | 0.441910339 |
| Asna1 | 0.245565014 | 0.44198262 |
| Scn11a | 1.801132744 | 0.442044463 |
| Fam175a | 0.556111547 | 0.442044463 |
| Coq3 | 0.334439698 | 0.442153379 |
| Adamdec1 | -1.865564531 | 0.442153379 |
| Actn2 | -1.929282837 | 0.442307259 |
| Apon | 3.689542307 | 0.442315735 |
| Il6 | -2.106011491 | 0.442562946 |
| Eepd1 | 0.46246896 | 0.442574557 |
| Aim2 | -0.440037993 | 0.442574557 |
| Gm14327 | 1.190839311 | 0.44271586 |
| Cep112 | 0.603020266 | 0.44271586 |
| Gm28551 | -3.256161424 | 0.44271586 |
| Gm10100 | -4.284287712 | 0.443069163 |
| Sh3rf2 | -4.284287712 | 0.443069163 |
| Mettl8 | 0.44297127 | 0.443175287 |
| Atg4d | 0.353992381 | 0.443704705 |
| Nrl | 3.008409252 | 0.444027485 |
| Setd8 | -0.334368535 | 0.44406427 |
| Nell1 | 0.920330066 | 0.444066809 |
| Tmem196 | 0.952096581 | 0.444205315 |
| Golgb1 | 0.233740548 | 0.444205315 |
| Tmem57 | -0.246525253 | 0.444205315 |
| Caskin1 | 0.43443227 | 0.445005204 |
| Gm3055 | 2.8649455 | 0.44512944 |
| Slc2a10 | 1.100777131 | 0.44512944 |
| Ccdc114 | 1.049418451 | 0.44512944 |
| Edrf1 | -0.390776553 | 0.445308212 |
| Zfp113 | -0.46006118 | 0.445357691 |
| Ptpru | 0.753839482 | 0.445445352 |
| Pcdh1 | -0.503458796 | 0.445445352 |
| Chst2 | 0.425659538 | 0.445519575 |
| Calr | 0.212123093 | 0.445519575 |
| Dyrk2 | -0.353599687 | 0.445553498 |
| Clpb | -0.278172901 | 0.445635906 |
| Zfp143 | -0.312064893 | 0.445696302 |
| P2ry14 | 0.597230316 | 0.445764829 |
| Zfp109 | -0.691533057 | 0.445802462 |
| Nol3 | 1.154537639 | 0.445959575 |
| Gm12353 | -1.068953399 | 0.445973088 |
| Tinagl1 | 1.045462201 | 0.4460884 |
| Ska2 | 0.1996541 | 0.446132196 |
| Olfr1420 | 3.436879008 | 0.446219418 |
| Ccdc175 | 3.409875531 | 0.446219418 |
| H2-Eb1 | 1.289255817 | 0.446432491 |
| Tspan4 | 0.391544269 | 0.446432491 |
| Mmp15 | 0.402822812 | 0.446526953 |
| Slc9c1 | -3.480521757 | 0.44656055 |
| Mark1 | 0.37943701 | 0.446692311 |
| Fbxo15 | 2.553950818 | 0.446901106 |
| Hnrnpu | 0.237870744 | 0.447091914 |
| Ppp2r3c | 0.242618122 | 0.447191905 |
| Cct8 | 0.169992796 | 0.447191905 |
| Usp9x | -0.17027905 | 0.447191905 |
| Neu3 | -1.365632494 | 0.447191905 |
| Mlip | -0.724137721 | 0.447252122 |
| Spata9 | -0.78067937 | 0.447252122 |
| Saxo2 | 0.585912229 | 0.447835383 |
| L2hgdh | 0.579338679 | 0.447835383 |
| Pam | 0.264700762 | 0.447835383 |
| Arap3 | -0.7306037 | 0.447835383 |
| Gm11084 | -3.480488319 | 0.447835383 |
| 4930538K18Rik | 3.410756959 | 0.448011885 |
| Kdf1 | 3.410756959 | 0.448011885 |
| Pld5 | 1.461001606 | 0.448011885 |
| Otud6b | 0.343129757 | 0.448011885 |
| Orc5 | 0.39127052 | 0.448156564 |
| Ermard | 0.296947725 | 0.448156564 |
| Csnk1e | 0.191662109 | 0.448156564 |
| Gpr19 | 0.263035494 | 0.448245966 |
| Cadps2 | -0.918795024 | 0.448245966 |
| Rnpc3 | 0.334058808 | 0.448328077 |
| Ttc9c | -0.309396237 | 0.448551567 |
| Oprl1 | 0.353319754 | 0.448572765 |
| Fnbp4 | -0.20075755 | 0.448572765 |
| Cggbp1 | -0.206705305 | 0.448688571 |
| Afg3l2 | -0.282302087 | 0.448786715 |
| Hhipl1 | 1.299103373 | 0.448864226 |
| Arhgap35 | -0.190891916 | 0.448864226 |
| Srek1 | 0.240625657 | 0.448875932 |
| Gtsf1l | 3.725865703 | 0.449031921 |
| Lrrc45 | -0.365513379 | 0.449178705 |
| Ackr2 | 2.801445531 | 0.449305113 |
| Slc12a1 | 3.670172504 | 0.449395777 |
| Fam198b | 1.865379797 | 0.449395777 |
| Hist1h2ai | 0.983479003 | 0.449395777 |
| H1fx | 0.538988426 | 0.449395777 |
| Pkp2 | 1.190378472 | 0.449421747 |
| Sirt2 | 0.212389548 | 0.449421747 |
| Enpp4 | 0.578940015 | 0.449613464 |

| | | |
|---|---|---|
| Clybl | 0.302600949 | 0.827318718 |
| Ctnna3 | -0.678109715 | 0.827318718 |
| Slc25a28 | 0.169158435 | 0.827331585 |
| Cd36 | 0.165689584 | 0.827331585 |
| 4930579G24Ri | -0.132801426 | 0.827505304 |
| Rnf14 | 0.059514041 | 0.827887069 |
| Aldh5a1 | 0.146927326 | 0.828113092 |
| Casp8ap2 | 0.070319769 | 0.828283385 |
| Hacd4 | -0.111274211 | 0.828323062 |
| Trim33 | -0.067723727 | 0.828366496 |
| Dtx4 | -0.107842282 | 0.828366496 |
| WDR97 | -0.749642631 | 0.828444133 |
| Ypel3 | -0.064270108 | 0.828484748 |
| Vapa | -0.059158451 | 0.828576966 |
| Erdr1 | 0.172061772 | 0.828853005 |
| Fbxl4 | 0.210615266 | 0.829337454 |
| Ntan1 | 0.060667407 | 0.829392527 |
| Clcn2 | -0.18903359 | 0.82951474 |
| Slc25a39 | -0.081920619 | 0.82960332 |
| Snx32 | 0.126554971 | 0.82977685 |
| Ppp1r14b | 0.068781704 | 0.82992064 |
| Reep1 | -0.079123944 | 0.830099398 |
| C330011M18R | 0.254282268 | 0.830112514 |
| Myl6 | -0.063835707 | 0.830112514 |
| 6330409D20Ri | -0.398393042 | 0.830112514 |
| Ttpa | -0.532545021 | 0.830112514 |
| Bhlhe23 | 0.566348386 | 0.830270982 |
| Rnaseh1 | 0.146067691 | 0.830281461 |
| Cachd1 | -0.109830732 | 0.830281461 |
| Clmn | -0.244631264 | 0.830281461 |
| Vhl | -0.07487464 | 0.830536988 |
| Fhod1 | 0.282601881 | 0.830578951 |
| Stx16 | 0.101766208 | 0.830578951 |
| Smpd3 | -0.09700626 | 0.830578951 |
| AI413582 | 0.087882877 | 0.830643182 |
| 1700020D05Ri | -1.081864404 | 0.83087891 |
| Zfp235 | -0.125449826 | 0.831004948 |
| Ahctf1 | -0.080488875 | 0.831040842 |
| Phldb1 | -0.110026075 | 0.831040842 |
| Dcaf7 | 0.061178947 | 0.831224441 |
| Rchy1 | 0.068335097 | 0.831271355 |
| Zfp825 | 0.112992619 | 0.831290221 |
| Tbata | -0.327911577 | 0.831290221 |
| 9430015G10Ri | 0.085191106 | 0.831385774 |
| Syf2 | 0.075573201 | 0.831568619 |
| Dqx1 | -0.520275357 | 0.831568619 |
| Ska3 | 0.120232167 | 0.831646552 |
| Gm11146 | -0.232275894 | 0.831646552 |
| Fgf10 | 0.497134531 | 0.831800096 |
| Enpp5 | -0.090081816 | 0.831806146 |
| Rev3l | 0.071961204 | 0.831871105 |
| St8sia1 | -0.11242846 | 0.831871105 |
| Zfp119a | -0.182910843 | 0.831871105 |
| Kcnj12 | -0.539089927 | 0.831871105 |
| Ido2 | -0.900790101 | 0.831871105 |
| Mpv17l2 | -0.079459749 | 0.831929379 |
| Tmcc2 | -0.138477958 | 0.831929379 |
| Cxcl5 | -0.70437489 | 0.831980829 |
| Hrc | 0.65454301 | 0.832065484 |
| Mgat1 | -0.092869701 | 0.832065484 |
| Fcgr3 | -0.118552751 | 0.832065484 |
| Fam219a | -0.129053848 | 0.832065484 |
| Pabpc5 | 0.492816451 | 0.832113126 |
| 1190005I06Rik | -0.759377358 | 0.832113126 |
| Pitpnc1 | -0.068363051 | 0.832234046 |
| Usp49 | 0.159268851 | 0.832261567 |
| Anapc5 | -0.0494246 | 0.832379704 |
| Mmp27 | -0.637138184 | 0.832523988 |
| Mesdc2 | 0.079885432 | 0.832592675 |
| Cntnap4 | 0.149752957 | 0.832614693 |
| Khdrbs1 | -0.056041441 | 0.832732632 |
| Fam72a | 0.221934238 | 0.832915107 |
| Snap25 | 0.086159776 | 0.832915107 |
| Grhl1 | -0.349925059 | 0.832915107 |
| Proser1 | -0.087359516 | 0.833023939 |
| Mllt10 | 0.084252834 | 0.833103334 |
| Hip1r | -0.123437113 | 0.833103334 |
| Mgme1 | 0.144030983 | 0.83377282 |
| Gm9797 | 0.247517228 | 0.833890499 |
| Tbc1d32 | -0.19957086 | 0.834119907 |
| P2rx1 | -0.207965933 | 0.834119907 |
| Ctf2 | 0.328754042 | 0.834179258 |
| Ipo4 | 0.087212489 | 0.834179258 |
| Phka1 | -0.334489716 | 0.834179258 |
| Cxcl3 | -0.713763283 | 0.834216827 |
| C2cd4b | 0.370671604 | 0.834310975 |
| Fam169a | 0.119140832 | 0.834310975 |
| Usp11 | -0.073117786 | 0.834310975 |
| Rpgrip1 | -0.230021021 | 0.834310975 |
| Angpt4 | -0.744974158 | 0.834310975 |
| 2210016F16Ril | -0.09091783 | 0.834388759 |
| Tmem145 | -0.145063361 | 0.834406136 |
| Yrdc | -0.074081514 | 0.834495003 |
| Npr2 | -0.180378336 | 0.834495003 |
| Med27 | -0.09766336 | 0.83462235 |
| Lemd3 | 0.16639231 | 0.834678688 |
| Rngtt | -0.066647745 | 0.834770133 |
| Rnmt | 0.079036712 | 0.83478915 |
| Pde9a | -0.125067873 | 0.83478915 |
| Zfp568 | -0.178530715 | 0.83478915 |
| Lamp3 | -0.778766711 | 0.83478915 |
| Proz | 0.499953911 | 0.83486674 |
| Pou3f2 | 0.09121194 | 0.834918527 |
| Canx | 0.055386638 | 0.835844557 |
| Aste1 | 0.135347894 | 0.836339728 |
| Rbm3 | 0.056719494 | 0.836339728 |
| Ssr3 | -0.051595452 | 0.836361571 |
| Zfp667 | -0.113046159 | 0.836361571 |
| Zfp146 | 0.082991101 | 0.836697627 |
| Tmem194b | -0.123483616 | 0.836697627 |
| Ppox | 0.107080501 | 0.836987094 |
| Myo1d | 0.300917042 | 0.837382809 |
| Ifit1bl1 | -0.355890704 | 0.837766796 |
| Pld1 | -0.127510139 | 0.837937335 |
| Nox1 | 0.577438209 | 0.838079561 |
| Cercam | -0.134564955 | 0.838079561 |
| Parp11 | -0.095655846 | 0.838186953 |
| Iigp1 | 0.3592353 | 0.838626802 |
| Hist3h2a | 0.10874293 | 0.838626802 |

| | | |
|---|---|---|
| 1110032A03Rik | 1.139675699 | 0.00178185 |
| Plscr1 | -1.328119547 | 0.00178185 |
| Setd6 | 1.176827443 | 0.001782471 |
| Smad7 | -1.174457089 | 0.001782471 |
| Tmem204 | 6.454900144 | 0.001785768 |
| Psmg3 | 1.287827608 | 0.001785768 |
| Chchd7 | 0.859742958 | 0.001793641 |
| Hspa4 | -0.640236677 | 0.001793641 |
| Kansl1 | -0.643781323 | 0.001793641 |
| Cited2 | -0.775303291 | 0.00179373 |
| Cdc5l | 0.851585116 | 0.001803367 |
| Gadd45g | 1.160940235 | 0.001806611 |
| Nfkbia | -1.183630995 | 0.001813979 |
| Ndufa13 | 0.607972603 | 0.001815345 |
| Rbm6 | -0.65332953 | 0.001818755 |
| Nxf1 | 0.589582344 | 0.001844013 |
| Ngrn | 0.957971156 | 0.001844212 |
| Hacd1 | 1.033827774 | 0.001845947 |
| Capn10 | 0.834228256 | 0.001851008 |
| Maged2 | 0.58377637 | 0.001857622 |
| Shfm1 | 0.606369657 | 0.001859467 |
| Mrpl39 | 0.83520211 | 0.00185972 |
| Mrpl36 | 0.786766727 | 0.00185972 |
| Trappc4 | 0.619707529 | 0.001863242 |
| Smad4 | 0.962816021 | 0.001863605 |
| Rnf19b | -1.202908328 | 0.001865391 |
| Rtn4 | 0.698441767 | 0.001865511 |
| Fbxo7 | 0.94901152 | 0.001869862 |
| Nav2 | -0.803563447 | 0.001869862 |
| Cd44 | -2.861046501 | 0.001869862 |
| Pnliprp2 | -7.072009069 | 0.001869862 |
| Trank1 | 2.22911042 | 0.001872565 |
| Gad2 | 1.092541456 | 0.001874547 |
| Zcchc18 | 0.672440408 | 0.001874547 |
| Rpl15 | 0.590285082 | 0.001874547 |
| Ogdh | -0.671126815 | 0.001882927 |
| Derl2 | -0.747950656 | 0.001885572 |
| Pfdn5 | 0.583524937 | 0.001891885 |
| Rnaset2a | -0.922898786 | 0.001896679 |
| Col6a3 | 2.823189952 | 0.001896807 |
| Zfp760 | 1.260024544 | 0.001904067 |
| Ckap5 | 0.745207737 | 0.001904067 |
| Sp140 | -1.252332913 | 0.001907418 |
| Zc3h18 | -0.744345716 | 0.001907465 |
| Ndufb6 | 0.769357835 | 0.001915145 |
| Dek | 0.69988645 | 0.001915772 |
| Kcnk2 | 1.299640643 | 0.001922386 |
| Micu3 | 0.698218716 | 0.001922386 |
| Tefm | 1.477520117 | 0.001923387 |
| Dnah7a | 3.133926391 | 0.001926206 |
| Paxip1 | -0.846729853 | 0.001941373 |
| Mterf1b | 4.548582174 | 0.001951715 |
| Slc2a3 | -1.540780455 | 0.00195208 |
| Map3k7 | -0.858653756 | 0.001953213 |
| Lamb1 | 1.443994153 | 0.001957261 |
| Irf8 | -0.899342842 | 0.001963211 |
| Snx7 | -1.87446876 | 0.001964051 |
| Herc4 | -0.821056588 | 0.001969352 |
| Gemin7 | 0.852864032 | 0.001972831 |
| Rgs17 | 1.469212196 | 0.001975031 |
| Tmem156 | -3.148097211 | 0.001991979 |
| Npm3 | 0.794534938 | 0.00200076 |
| Emp1 | 1.723034404 | 0.002006648 |
| Plekha2 | -1.439674476 | 0.002009059 |
| Smc2 | 0.826742502 | 0.002015506 |
| Ears2 | 1.693673495 | 0.002021708 |
| Twf2 | -0.949292906 | 0.002030662 |
| Zfp626 | -1.579106585 | 0.00203331 |
| 10-Sep | 1.329530862 | 0.002033565 |
| Pfkfb4 | -1.005768932 | 0.002039394 |
| Wdr45b | 0.864834814 | 0.002044123 |
| Frs3 | 1.733319862 | 0.00205266 |
| Ccdc126 | -1.520323264 | 0.00205266 |
| Pih1d1 | 0.745286175 | 0.002059576 |
| Slmap | -0.805581168 | 0.002064971 |
| Poldip3 | -0.607881281 | 0.002071706 |
| Gdf9 | -6.793439848 | 0.002071706 |
| Zfp36l1 | 0.900881177 | 0.002075824 |
| Cdc34 | -0.996882681 | 0.002084496 |
| Myl7 | 6.668925151 | 0.002088886 |
| Fbxl18 | -0.932612788 | 0.002091604 |
| Nedd4l | -0.841402315 | 0.002101651 |
| Rhpn2 | 6.092904523 | 0.002101987 |
| Pkd1 | -1.262805947 | 0.002106402 |
| C1d | 0.619724918 | 0.0021108 |
| Zfp964 | 4.588733823 | 0.002130013 |
| Rpl7 | 0.50490187 | 0.002132513 |
| Prss50 | -3.525342395 | 0.002132513 |
| Zmynd15 | -2.533477707 | 0.002137455 |
| Fam184a | 1.903701402 | 0.00213755 |
| Sh3glb1 | -0.735106238 | 0.00213755 |
| Ifi35 | -1.662375222 | 0.00213755 |
| Tnfaip3 | -1.377458944 | 0.002145394 |
| Pomgnt2 | 1.565865298 | 0.00215867 |
| Rdm1 | 1.34492087 | 0.002164378 |
| Clpp | 0.689358533 | 0.002166114 |
| Ppef1 | -6.320573544 | 0.002166114 |
| Magi1 | 0.839874757 | 0.002177997 |
| Slc38a7 | -1.099784243 | 0.002181456 |
| Tox2 | 1.115092074 | 0.002183863 |
| Erg | 6.080050301 | 0.002207023 |
| Malsu1 | 0.823675276 | 0.002210394 |
| Plin3 | -0.930020785 | 0.002210394 |
| Dennd2c | -3.293019419 | 0.002210394 |
| Maob | 5.840169257 | 0.002213313 |
| Tgfbr2 | -1.557516225 | 0.002213472 |
| Nkain1 | -1.0906147 | 0.002215567 |
| Gpc2 | -1.451761674 | 0.002216513 |
| Ccdc28b | 1.022549326 | 0.002221108 |
| Pag1 | -0.904631474 | 0.002235383 |
| 2310030G06Rik | 6.45647142 | 0.002236638 |
| Ssscal | 0.876835992 | 0.002242875 |
| Mcm6 | -0.839157767 | 0.00225181 |
| Mtmr9 | -1.013282552 | 0.002257267 |
| Ermp1 | -1.191718001 | 0.002257267 |
| Tti1 | -1.107014964 | 0.002267889 |
| Msantd3 | 0.967612271 | 0.002279203 |
| Yap1 | 2.132800615 | 0.002290635 |
| 1500009C09Rik | 1.003427569 | 0.002297176 |
| Esyt3 | 2.593937178 | 0.118190018 |
| Myzap | 2.937492171 | 0.118334063 |
| Mrpl49 | 0.559274333 | 0.118373535 |
| Emd | 0.447480619 | 0.118373535 |
| Ankrd11 | 0.37431579 | 0.118545423 |
| Timp3 | 1.101021933 | 0.118678777 |
| Pgrmc1 | 0.388164687 | 0.118714737 |
| Fam195a | 1.02363133 | 0.118785113 |
| Sirt3 | 0.800034098 | 0.118785113 |
| Cdca7l | -0.862310231 | 0.11887769 |
| Apc | 0.369438431 | 0.118936987 |
| Cwc15 | 0.335131427 | 0.118936987 |
| Mcm9 | -0.91395867 | 0.118936987 |
| Gm13154 | -1.985733156 | 0.118969717 |
| Rtkn | -1.413553866 | 0.119072905 |
| Bag6 | -0.479338549 | 0.119137459 |
| Vps51 | -0.433357834 | 0.119301933 |
| Vps18 | -0.540742132 | 0.119330407 |
| Fam69b | 0.551841867 | 0.119346317 |
| Pdhb | 0.32845841 | 0.119353652 |
| Inpp5j | -1.754460749 | 0.119370632 |
| Bcl2a1d | -0.683746922 | 0.119391461 |
| Rbpms2 | 2.332381766 | 0.119702048 |
| Vps53 | -0.476682554 | 0.11976821 |
| Gckr | 4.620907883 | 0.119795183 |
| Fbxo34 | 0.722950008 | 0.119796102 |
| Epm2aip1 | -0.45056308 | 0.119894002 |
| Kcnk4 | 4.7457243 | 0.119899792 |
| St6galnac2 | -1.063327162 | 0.119998693 |
| Cfl2 | 0.396949697 | 0.120007555 |
| Grhl3 | -3.982747851 | 0.120056163 |
| Gm5483 | -2.349033091 | 0.120095171 |
| BC003965 | 0.459444427 | 0.120357972 |
| Abl1 | -0.602101617 | 0.1204751 |
| Sgpp2 | 4.048657611 | 0.120487488 |
| Zfp185 | 2.968406578 | 0.120608234 |
| 1700007G11Rik | 2.66144039 | 0.120613985 |
| Ankrd35 | 4.79066873 | 0.120765084 |
| Sfr1 | 0.403690844 | 0.120765084 |
| Glra3 | 4.885771852 | 0.120804989 |
| AU022751 | -5.248913814 | 0.120883006 |
| Myh3 | -5.248913814 | 0.120883006 |
| Taok1 | 0.350899203 | 0.121042015 |
| 4931422A03Rik | -0.981249705 | 0.121081468 |
| Lrrn3 | 0.618284594 | 0.121119737 |
| Plscr3 | -0.607455223 | 0.121167887 |
| Heg1 | -0.434831556 | 0.121168349 |
| Spx | -3.872644339 | 0.121296602 |
| Aak1 | -0.434170628 | 0.12131935 |
| Rrs1 | -0.510735922 | 0.121596697 |
| Foxc2 | 5.039033948 | 0.122043691 |
| Dgke | -0.523935398 | 0.122241258 |
| Slc7a6os | 0.559590687 | 0.122341535 |
| Coa6 | 0.658411006 | 0.122341612 |
| D2Wsu81e | 0.611096586 | 0.122341612 |
| Pcdhgc5 | -1.170998463 | 0.122341612 |
| Il18r1 | -2.91562009 | 0.122341612 |
| Gm10643 | 1.38423917 | 0.122354682 |
| Atxn1 | 0.415331974 | 0.122552287 |
| Ppm1m | -0.632400811 | 0.122606086 |
| Gm13152 | -5.578753508 | 0.122752326 |
| Tac4 | -5.578753508 | 0.122752326 |
| Rad50 | -0.506902489 | 0.122808111 |
| Nuf2 | 0.591770427 | 0.122814378 |
| Stk3 | -0.575023505 | 0.122814378 |
| A530064D06Rik | -2.629342512 | 0.122814378 |
| Gm10961 | -2.915164304 | 0.122814378 |
| Grsf1 | -0.336445248 | 0.122834752 |
| Dnajc21 | 0.863207974 | 0.122863768 |
| Cyb5a | 0.317913497 | 0.122931433 |
| Zfhx2 | -0.525893623 | 0.123107706 |
| Tmem45a2 | -5.571339431 | 0.123107706 |
| Rgs7 | 0.830650529 | 0.123210529 |
| Lrp3 | 0.708826483 | 0.123210529 |
| Ccdc94 | 0.69541726 | 0.123210529 |
| Gnb2 | 0.340880323 | 0.123210529 |
| Emc10 | 0.314736011 | 0.123210529 |
| Dlgap2 | -0.738730723 | 0.123210529 |
| Gpr146 | -0.887663617 | 0.123210529 |
| Gdpd3 | -1.834507961 | 0.123210529 |
| Cd209g | -5.559683333 | 0.123231303 |
| Setd3 | -0.403727165 | 0.123532025 |
| Brd7 | -0.430973692 | 0.123561679 |
| Snip1 | -0.785626036 | 0.123616779 |
| Plcxd3 | 0.764599859 | 0.123656275 |
| Col9a1 | 5.158506845 | 0.123802274 |
| Gm28051 | -5.597645368 | 0.123832762 |
| Cenph | 0.778315943 | 0.123891383 |
| Rpl6l | 0.424067172 | 0.124242301 |
| Abi2 | 0.391119781 | 0.124264911 |
| Ctps | -0.481751938 | 0.124264911 |
| Fam212a | -0.614676407 | 0.124264911 |
| Elmod3 | 0.937781612 | 0.124483604 |
| Slc18b1 | 0.852283502 | 0.124483604 |
| 2310039H08Rik | 0.510190513 | 0.124483604 |
| Shkbp1 | -0.508463918 | 0.124483604 |
| Pgam2 | 3.592098939 | 0.124529765 |
| Dok6 | -0.998232271 | 0.124529765 |
| Spock2 | 0.597174703 | 0.124619192 |
| Rpain | 0.515960046 | 0.124656191 |
| Tgoln1 | -0.477338317 | 0.124771115 |
| Cyb561 | 1.228785772 | 0.124788623 |
| Med12 | -0.762759394 | 0.124825186 |
| Bub1b | -0.458498603 | 0.124852712 |
| Ercc2 | -0.680837734 | 0.124911652 |
| C1qb | -0.622199134 | 0.124936189 |
| Chchd5 | 0.627347276 | 0.125005971 |
| Zfp324 | -0.999357783 | 0.125005971 |
| Gata1 | 4.711958708 | 0.125014942 |
| Cltb | 0.390770492 | 0.125055551 |
| Pdk2 | 0.559624225 | 0.12518679 |
| Akap8 | -0.420961859 | 0.125220294 |
| Ap1b1 | -0.476723914 | 0.125298028 |
| Prr7 | 0.539738185 | 0.125355154 |
| Khdc3 | -5.221844324 | 0.125355154 |
| Pde1c | 0.939234261 | 0.125613343 |
| Hook3 | 0.355141603 | 0.125631714 |
| Golim4 | 0.623817158 | 0.125694194 |
| Srprb | 0.540047949 | 0.125701675 |
| Bod1 | -0.215384675 | 0.449613464 |
| Tbck | -0.18380986 | 0.449672582 |
| Sdcbp2 | 3.014797984 | 0.449905221 |
| Dusp14 | 0.485013555 | 0.449956589 |
| Repin1 | 0.400384464 | 0.450149295 |
| Trim41 | 0.292001866 | 0.450149295 |
| Oosp3 | -4.197104022 | 0.450149295 |
| Tigd4 | -4.197104022 | 0.450149295 |
| Hpn | -1.13356771 | 0.450428723 |
| Apobec4 | 1.811294751 | 0.450490161 |
| Aagab | 0.249348612 | 0.450490161 |
| Tmem91 | -0.679902443 | 0.450490161 |
| Hcn4 | -0.560091616 | 0.450633141 |
| Eml5 | 0.288604114 | 0.450696134 |
| Ing5 | -0.372766997 | 0.450716627 |
| BC048403 | -1.225636828 | 0.450716627 |
| Gm10509 | -1.204864466 | 0.451044023 |
| F2rl3 | 2.989557043 | 0.451298367 |
| Zscan26 | 0.257295125 | 0.45156445 |
| Cnot10 | -0.356053251 | 0.451659209 |
| Lyl1 | -0.529239178 | 0.451659209 |
| Zfp142 | -0.390035797 | 0.45171031 |
| Cdc25b | 0.451473547 | 0.45175301 |
| Rbbp4 | -0.173871871 | 0.452084303 |
| Teddm2 | 1.512241491 | 0.452495523 |
| Slc35a5 | -0.403481655 | 0.452560649 |
| Fat2 | -1.282483308 | 0.452705944 |
| Pls3 | 0.322004605 | 0.452778049 |
| Gbp3 | -1.137094751 | 0.452881013 |
| Cx3cl1 | 0.449328412 | 0.452933475 |
| Hdhd3 | 0.581315532 | 0.453023882 |
| Dhx29 | 0.273277028 | 0.453023882 |
| 4921536K21Rik | 3.391028189 | 0.453026061 |
| Stk25 | 0.236052022 | 0.453026061 |
| Zfp300 | -0.795552123 | 0.453046534 |
| Cdc42ep4 | 0.268272197 | 0.453119555 |
| Hivep1 | -0.30958089 | 0.453340482 |
| Wdr78 | 0.4904084 | 0.453485574 |
| Ecm2 | 1.019083356 | 0.453598235 |
| Mapkbp1 | -0.318927227 | 0.453672396 |
| Cacna1e | -0.414693886 | 0.453859027 |
| Car11 | 0.426577879 | 0.45411621 |
| Adck5 | 0.389193301 | 0.45411621 |
| Fam150b | 3.659915765 | 0.454460866 |
| Klrb1b | 3.659915765 | 0.454460866 |
| Dph6 | 0.275418442 | 0.454460866 |
| Ttll3 | -0.779540247 | 0.454460866 |
| Tmem56 | 0.906113866 | 0.454551325 |
| Barhl2 | -0.772981672 | 0.454551325 |
| Nptx1 | 0.58224963 | 0.454611008 |
| Apol11b | 3.993723478 | 0.455118998 |
| Tex40 | 3.676076724 | 0.455118998 |
| Celf6 | 0.477750509 | 0.455118998 |
| Afap1 | -0.182268155 | 0.455118998 |
| Ankrd2 | 3.704508614 | 0.455408044 |
| Sgca | 3.704508614 | 0.455408044 |
| Gm13306 | 1.640787515 | 0.455408044 |
| D630044L22Rik | 1.34526855 | 0.455408044 |
| Ifih1 | -0.485802727 | 0.455408044 |
| Jade3 | -0.749092898 | 0.455408044 |
| Gm10775 | 3.702643002 | 0.455498371 |
| Snap47 | 0.189245912 | 0.455542228 |
| Ddo | -2.042855825 | 0.455576014 |
| Ttll5 | 0.307224542 | 0.455868201 |
| Fam174a | -0.246851871 | 0.455868201 |
| Tnfrsf25 | -1.227179717 | 0.456124693 |
| Txnl1 | 0.196428384 | 0.456204658 |
| Ddx42 | 0.188941053 | 0.456717439 |
| Olfr433 | -1.620910755 | 0.456866948 |
| Utp15 | -0.250044658 | 0.457138527 |
| Cipc | -0.231478765 | 0.457301986 |
| Athl1 | 0.522004556 | 0.457457661 |
| Sf3a3 | 0.195041436 | 0.457715213 |
| Itsn1 | -0.314982854 | 0.457715213 |
| Gm1110 | 3.637972176 | 0.457727558 |
| Acot8 | -0.318952463 | 0.457842072 |
| Gm6792 | 3.388668997 | 0.457921148 |
| Loxhd1 | 3.388668997 | 0.457921148 |
| Dcst2 | 3.694496741 | 0.45805187 |
| Mcpt8 | 3.694496741 | 0.45805187 |
| Tas2r137 | 3.694496741 | 0.45805187 |
| Kdm5d | 0.357870972 | 0.45805187 |
| Anks1b | 0.286111499 | 0.45805187 |
| Gm21190 | -4.166884506 | 0.45805187 |
| Bcap31 | -0.233215489 | 0.458464046 |
| Klf15 | 0.484073859 | 0.458469567 |
| Gfpt2 | 0.717307703 | 0.458474882 |
| Ddost | -0.193891761 | 0.45847708 |
| Islr2 | -0.333193323 | 0.458502545 |
| Mcrs1 | 0.223493591 | 0.458817229 |
| Hus1 | -0.550982993 | 0.458873861 |
| Toporsos | 0.945705846 | 0.459035477 |
| Lef1 | 0.541805964 | 0.459035477 |
| Vat1 | 0.3123245 | 0.459035477 |
| Fbxo31 | 0.261867217 | 0.459035477 |
| Slc6a8 | -0.371616891 | 0.459035477 |
| Dcstamp | 2.984953932 | 0.459040857 |
| Prrx2 | 1.494164795 | 0.459040857 |
| Fam126b | 0.411270212 | 0.459040857 |
| Slc17a5 | 0.347237658 | 0.459040857 |
| Ppp2r5b | 0.270992206 | 0.459040857 |
| Vsig10 | 2.324494483 | 0.45908429 |
| Zcchc9 | 0.309733166 | 0.45908429 |
| Hdac7 | 0.284926286 | 0.45908429 |
| Mllt6 | -0.21687947 | 0.459551668 |
| C1galt1c1 | -0.350333517 | 0.459751579 |
| St8sia2 | 0.227753891 | 0.459841715 |
| Grin2b | -0.273622759 | 0.459841715 |
| Phf11d | -0.649051873 | 0.459841715 |
| Upb1 | 3.684419834 | 0.459869637 |
| Aurkb | 0.343479193 | 0.459909406 |
| Zfp830 | 0.332372212 | 0.459909406 |
| Idh2 | 0.174055912 | 0.460027597 |
| Hsd3b6 | 3.37868148 | 0.460096306 |
| Dus3l | 0.275255691 | 0.460096306 |
| Kcnk9 | 3.000124377 | 0.460120793 |
| Zcchc8 | -0.200542861 | 0.460531176 |
| Mill2 | 3.649562557 | 0.460632117 |
| Rho | 3.649562557 | 0.460632117 |
| Gm1604a | 0.756847516 | 0.838826366 |
| Wdr82 | 0.073831253 | 0.838983379 |
| B3galt5 | -0.560604559 | 0.838983379 |
| Rpap1 | -0.103259767 | 0.839023624 |
| Gnrh1 | 0.412731277 | 0.839040847 |
| Id2 | -0.054501962 | 0.839040847 |
| Afp | -0.266763229 | 0.839040847 |
| Usp10 | -0.068810611 | 0.839112434 |
| Thnsl1 | -0.106946042 | 0.839213042 |
| Ccdc129 | 0.456774549 | 0.839226807 |
| Strap | -0.050768858 | 0.839296754 |
| 3830406C13Ri | -0.07789628 | 0.839296754 |
| Ift122 | -0.106456969 | 0.839500868 |
| Abcg5 | 0.345876328 | 0.839730873 |
| Fntb | -0.095501069 | 0.84028312 |
| Gemin2 | -0.133433643 | 0.840316397 |
| Pabpn1 | 0.146092853 | 0.840544862 |
| Acp6 | 0.112850608 | 0.840544862 |
| Rbl2 | -0.088966335 | 0.840544862 |
| Rgs1 | -0.146935296 | 0.840691035 |
| Dpy19l3 | 0.129082873 | 0.840715478 |
| Cklf | 0.100398875 | 0.840715478 |
| Med9 | 0.085895866 | 0.840715478 |
| Ddhd1 | -0.121440619 | 0.841035901 |
| Tatdn3 | 0.148727415 | 0.841107366 |
| Isoc1 | 0.081872437 | 0.841127093 |
| Gpr3 | -0.306514081 | 0.841226178 |
| Decr2 | -0.104790138 | 0.841290295 |
| Gm10311 | 0.470819598 | 0.841317152 |
| Zdhhc8 | -0.132863956 | 0.841317152 |
| Daam2 | 0.3077355 | 0.841347504 |
| Sertad4 | 0.120274575 | 0.841347504 |
| Gss | -0.105944975 | 0.841347504 |
| Zbtb8b | -0.26868449 | 0.841347504 |
| Cpne2 | -0.148442729 | 0.841417344 |
| Aspdh | 0.517410059 | 0.841487368 |
| Rbm12b1 | 0.197059237 | 0.841487368 |
| BC029722 | 0.101714942 | 0.841487368 |
| Mtmr2 | -0.064548788 | 0.841528345 |
| Ficd | -0.152483032 | 0.841580932 |
| Man1a2 | 0.074074832 | 0.841640069 |
| Kcnt2 | 0.344054045 | 0.841712068 |
| Ndnl2 | -0.068128523 | 0.841740737 |
| Klhl15 | -0.115383545 | 0.841740737 |
| Ptprh | 0.766806713 | 0.842179411 |
| Zfp397 | -0.087712855 | 0.842179411 |
| Polr3e | -0.095911718 | 0.842179411 |
| Epb41l2 | -0.082456504 | 0.842262051 |
| Dusp4 | -0.089947845 | 0.842357931 |
| Six5 | -0.259943377 | 0.842712889 |
| Cacng2 | 0.137985876 | 0.843300394 |
| Pogz | 0.085305217 | 0.843300394 |
| Ccdc73 | 0.239710992 | 0.843378733 |
| AI987944 | 0.178664539 | 0.843682523 |
| A430078G23Ri | -0.802849328 | 0.843682523 |
| Dcakd | -0.057425905 | 0.843950389 |
| Chek2 | -0.128880087 | 0.844118044 |
| Grcc10 | 0.154817279 | 0.844271039 |
| Cenpv | -0.094768728 | 0.844271039 |
| Dclre1a | -0.129568968 | 0.844405853 |
| Casp6 | 0.090334265 | 0.844453927 |
| Srp68 | 0.067376791 | 0.844453927 |
| Phf8 | -0.07056481 | 0.844453927 |
| Nup153 | -0.079231901 | 0.844453927 |
| Ldhal6b | 0.944207748 | 0.844709181 |
| Ecd | -0.08085827 | 0.84503671 |
| Hemk1 | 0.088621395 | 0.845068604 |
| Alg11 | 0.072250499 | 0.845068604 |
| Dcp1b | -0.093849558 | 0.845068604 |
| BC028528 | -0.122431355 | 0.845075278 |
| Gm9996 | -0.120430319 | 0.845213053 |
| Sytl3 | -0.618994278 | 0.845227045 |
| Gtf2h4 | -0.1020612 | 0.845294909 |
| Lrrc9 | -0.210679709 | 0.845373491 |
| Gm7694 | 0.480353138 | 0.845844514 |
| Prickle2 | 0.114426084 | 0.845909356 |
| Tm7sf2 | 0.078025779 | 0.845909356 |
| Tmem116 | -0.239573524 | 0.845909356 |
| Acer3 | 0.138980142 | 0.845920572 |
| Spns2 | 0.242651261 | 0.845953563 |
| Prps1l3 | -0.111734991 | 0.845953563 |
| Lrrtm3 | -0.166318124 | 0.845953563 |
| Dnah10 | -0.272632324 | 0.845953563 |
| Anapc15 | -0.114999857 | 0.845966728 |
| Unc13a | -0.105904847 | 0.846024657 |
| Hsph1 | 0.078578752 | 0.8462223 |
| Ubqln2 | 0.067356112 | 0.846262796 |
| Sft2d1 | 0.062640456 | 0.846395736 |
| Pnpla8 | -0.064130404 | 0.846395736 |
| Spag8 | 0.283769945 | 0.846460119 |
| Abhd17a | 0.08218958 | 0.846638656 |
| Vrk3 | -0.07843448 | 0.846957038 |
| A830005F24Ri | -0.467823134 | 0.847216997 |
| Kcnma1 | 0.105584783 | 0.847247422 |
| Flt1 | 0.226850676 | 0.847331093 |
| Fastkd2 | -0.103661189 | 0.847709986 |
| Zfp599 | 0.197067891 | 0.847826226 |
| Scn2a1 | -0.101258253 | 0.847826226 |
| Shf | -0.217399269 | 0.847826226 |
| Ift88 | 0.120360031 | 0.847957508 |
| Isoc2b | 0.457739715 | 0.84824866 |
| Clec16a | -0.088306464 | 0.84824866 |
| Rad51c | 0.134349291 | 0.848401819 |
| Mib2 | 0.090297715 | 0.848401819 |
| Pus3 | 0.6322422 | 0.848812025 |
| Plpp5 | 0.094358575 | 0.848812025 |
| Ipp | -0.095972251 | 0.848812025 |
| Il12a | -0.310321703 | 0.848812025 |
| Zfp385c | -0.617913351 | 0.848812025 |
| Gldc | -0.126955161 | 0.849704952 |
| Zfp62 | 0.090505759 | 0.850398122 |
| Gm17333 | -0.813668124 | 0.850753393 |
| Ppp2r4 | -0.057188672 | 0.851332833 |
| Nxn | -0.150228309 | 0.851332833 |
| Gpn1 | -0.06150049 | 0.851725177 |
| Aldh1l2 | 0.098884861 | 0.851923293 |
| Med7 | -0.077902869 | 0.851923293 |
| Tcerg1 | -0.052535831 | 0.851930849 |
| Dynll2 | -0.059271563 | 0.851930849 |

| | | |
|---|---|---|
| Nol7 | 0.727015726 | 0.002300964 |
| Cdc40 | 0.770039461 | 0.002319586 |
| Amfr | -0.818631376 | 0.002319761 |
| Gpm6b | 0.77930017 | 0.002321645 |
| Wdr60 | 1.278091042 | 0.002329833 |
| Grik2 | 0.827064123 | 0.002329833 |
| Eva1a | 5.971158438 | 0.002346174 |
| Tmx1 | -0.592047754 | 0.00234896 |
| Nadsyn1 | -1.49120216 | 0.002366669 |
| Rin3 | -1.411315083 | 0.002372704 |
| Sh3pxd2b | -0.983806527 | 0.00237475 |
| Spta1 | 6.11630541 | 0.00237572 |
| Id3 | 1.199857753 | 0.002377053 |
| Tradd | -0.994088146 | 0.002379002 |
| Ube2cbp | 2.675720564 | 0.002379544 |
| Qsox1 | -1.173773361 | 0.00240045 |
| Scd3 | 6.067442961 | 0.002415191 |
| 4921511H03Rik | 6.22590767 | 0.002423769 |
| Gng12 | -0.766946068 | 0.002423769 |
| AU021092 | 6.141848692 | 0.002432065 |
| Cenpm | 1.261676518 | 0.002433429 |
| Smad1 | -0.864618826 | 0.002433429 |
| Rasd1 | 4.950345123 | 0.002438495 |
| Acat1 | 0.687493781 | 0.002438495 |
| Klra2 | -2.172272991 | 0.002449199 |
| Parp12 | -1.446243397 | 0.002468033 |
| Cdca3 | 1.022746477 | 0.002474034 |
| P2ry12 | 0.98221576 | 0.002477183 |
| Prkaca | -0.888648556 | 0.002477183 |
| Csf1r | -0.929925861 | 0.002478127 |
| Bhlhe22 | -1.033942704 | 0.002478127 |
| Lrg1 | -6.006267469 | 0.002478127 |
| Hp1bp3 | 0.564080951 | 0.00248432 |
| Rapgef5 | -1.195149869 | 0.002491319 |
| Sdhc | 0.739839102 | 0.002505621 |
| 1700112E06Rik | -1.861723746 | 0.002518303 |
| Adprh | 0.595835381 | 0.002529695 |
| Hap1 | 1.511225639 | 0.002533173 |
| Nfatc2 | -1.587231923 | 0.002533173 |
| Yae1d1 | 0.88109697 | 0.00253535 |
| Klhl4 | 5.533654736 | 0.002535508 |
| Fcrla | -6.173168951 | 0.002552347 |
| Slc15a4 | -0.950873502 | 0.002554863 |
| Cinp | 0.775759604 | 0.002572255 |
| Rnf115 | -0.830543746 | 0.002572255 |
| Rab20 | -1.385112382 | 0.002572255 |
| Gpnmb | -2.485774308 | 0.002572255 |
| Gpr84 | -2.91905732 | 0.002579712 |
| Tmed7 | -0.831046655 | 0.002598897 |
| Prpsap1 | 0.602735008 | 0.002604929 |
| Crip3 | 6.02867523 | 0.002614059 |
| Fat4 | -1.006422857 | 0.002622 |
| Mybl1 | 2.85745757 | 0.002627955 |
| Mgam | -7.261016957 | 0.002641722 |
| Plcl2 | -0.971983333 | 0.00264286 |
| Ecm1 | -2.164118209 | 0.002649691 |
| Pikfyve | -0.912131416 | 0.002653155 |
| Eda2r | 6.019529847 | 0.002663309 |
| Pja1 | 0.566865933 | 0.002664239 |
| Spint2 | 1.492940222 | 0.002667182 |
| Tspyl4 | 0.686673656 | 0.002668174 |
| Ahrr | -4.272776725 | 0.002684015 |
| Gm10250 | 0.806790149 | 0.002689896 |
| Eif3b | -0.62749378 | 0.002701925 |
| Plp1 | 2.168482529 | 0.002705002 |
| Tmem59l | 1.215938678 | 0.002705002 |
| Cxx1c | 1.071515262 | 0.002705002 |
| Khdrbs3 | 0.911373752 | 0.002705002 |
| Cd200r1 | -2.940480124 | 0.002705002 |
| Sapcd2 | 1.703546087 | 0.002705418 |
| Fam133b | 0.874191589 | 0.002715399 |
| Dnase1l3 | 6.530749602 | 0.002718541 |
| 2610318N02Rik | 1.299700804 | 0.002724756 |
| Fgl2 | -1.809350686 | 0.002734373 |
| Pdyn | 6.449093319 | 0.002745212 |
| Sdc3 | -1.125998102 | 0.002745212 |
| Mta2 | -0.667930056 | 0.002761055 |
| Tamm41 | 1.049203625 | 0.002773069 |
| Capza2 | -0.828652424 | 0.002773069 |
| Cfap46 | 6.268150756 | 0.002774035 |
| Swi5 | 0.568029987 | 0.002777725 |
| Leng8 | -0.76263053 | 0.002793802 |
| Pcdhb19 | -1.391061968 | 0.002804177 |
| Chd5 | 1.21818124 | 0.002814784 |
| Scaper | 0.870649174 | 0.002837196 |
| Cml1 | 2.372526461 | 0.00284114 |
| Mboat1 | -1.530146607 | 0.002849318 |
| Slc12a7 | -1.335588447 | 0.002850152 |
| Myrf | -4.482105151 | 0.002850152 |
| Sdc2 | 2.046310666 | 0.002859325 |
| Arpc5l | 0.581927287 | 0.002864117 |
| Mcoln2 | -2.761705296 | 0.002867757 |
| Rabggta | -0.8787415 | 0.002873907 |
| Tbrg4 | -0.841169413 | 0.002877786 |
| Vps52 | -0.658559008 | 0.002877799 |
| Plppr4 | 1.60044158 | 0.002879285 |
| Ankrd37 | 1.653195394 | 0.002880653 |
| Taf10 | -0.841784263 | 0.002881643 |
| Gdf10 | 5.760829411 | 0.002899746 |
| Mcfd2 | -0.744181351 | 0.002911001 |
| Fam49a | -0.980415622 | 0.002915369 |
| Zxdc | -1.0451467 | 0.002915837 |
| Pomc | 2.840998464 | 0.002919204 |
| Rftn1 | -1.448499485 | 0.002923119 |
| Ndufs7 | 0.575656945 | 0.002927934 |
| Zfp811 | 0.94578413 | 0.002937441 |
| Slco1c1 | 5.671639445 | 0.002938299 |
| Tlr9 | -1.963642425 | 0.002947627 |
| Spaca4 | -6.730870123 | 0.002947627 |
| Fzd2 | 1.617188731 | 0.002961864 |
| Fbxw11 | -0.613223041 | 0.002964592 |
| Dnaaf5 | -1.375880036 | 0.002967627 |
| Zwint | 0.58927616 | 0.002980312 |
| Fbxo11 | 0.708487032 | 0.002982589 |
| Txndc5 | -0.853278812 | 0.002982589 |
| Dbx1 | -3.50104406 | 0.002982589 |
| Uaca | 1.772538753 | 0.002994498 |
| Park7 | 0.57814928 | 0.002998725 |
| Cmbl | 6.278231738 | 0.003001166 |

| | | |
|---|---|---|
| Dnajb5 | 0.516106396 | 0.125718527 |
| Cmtm8 | 1.192448397 | 0.125751149 |
| Coq2 | -0.471323113 | 0.125751149 |
| Ddx28 | -0.591756231 | 0.125751149 |
| Atp11a | -0.75698129 | 0.125751149 |
| Qprt | -1.676001742 | 0.125825237 |
| Myh7 | 1.233428099 | 0.125878269 |
| Setmar | 2.386951729 | 0.125979347 |
| Cul5 | 0.511509133 | 0.125979347 |
| Rps9 | 0.314231712 | 0.125979347 |
| 2610507B11Rik | -0.376572898 | 0.125979347 |
| Grap2 | 4.644397778 | 0.126024048 |
| Gm12689 | 4.616035219 | 0.126024048 |
| Ankrd9 | 1.774726411 | 0.126098165 |
| Pdzd2 | 1.046554441 | 0.126108489 |
| Rasgrp3 | 1.07208279 | 0.126131033 |
| Adam4 | 1.794290232 | 0.126435125 |
| Def8 | -0.492763101 | 0.126454148 |
| Epn2 | 0.555949373 | 0.126536963 |
| Ssfa2 | -0.587664574 | 0.12667587 |
| Lgi4 | 4.557477101 | 0.126678955 |
| Stub1 | 0.416843721 | 0.12668988 |
| Dcun1d5 | -0.361312711 | 0.12668988 |
| Slc48a1 | -0.380638793 | 0.126713273 |
| Cga | 4.650997148 | 0.126723461 |
| Pura | 0.695434505 | 0.12686012 |
| Sectm1a | 4.641657244 | 0.126948951 |
| Nipa2 | -0.520202428 | 0.12695052 |
| Wdr48 | -0.446949799 | 0.12704828 |
| Ltbp2 | 3.78903791 | 0.127114396 |
| Nxt1 | 0.418108807 | 0.127145386 |
| Rad23b | -0.405628908 | 0.127208441 |
| Slc35g1 | -1.081069404 | 0.127363516 |
| Rfx1 | 0.939487127 | 0.12744903 |
| Cgrrf1 | 0.657409417 | 0.12744903 |
| Oxa1l | 0.417380762 | 0.12744903 |
| Lmo7 | -1.054316817 | 0.12744903 |
| Tbc1d4 | 0.681085628 | 0.127458585 |
| Tnpo2 | -0.554925858 | 0.127536433 |
| Mrpl18 | 0.418276541 | 0.127682707 |
| Fads1 | 0.360072509 | 0.127699582 |
| Traf3ip2 | -0.941659376 | 0.127699582 |
| Slc41a1 | 0.599602106 | 0.127829409 |
| Zfp110 | 0.820452164 | 0.127940702 |
| Atp6v0a1 | -0.54189419 | 0.127984986 |
| Impdh1 | 0.474831714 | 0.128276727 |
| Grina | -0.522356399 | 0.128373492 |
| Fgf1 | 4.573034223 | 0.128374077 |
| Npc1 | -0.520339103 | 0.128374077 |
| Psapl1 | 4.722359087 | 0.128456401 |
| 0610037L13Rik | 0.388883507 | 0.128582345 |
| Dcp1a | -0.537723175 | 0.12866095 |
| Ppp1r32 | 2.555436636 | 0.128718887 |
| Pddc1 | 0.40954616 | 0.128767276 |
| Glul | -0.404511807 | 0.128821578 |
| Cyp2r1 | -3.791555806 | 0.128852833 |
| Miefl | -0.564905974 | 0.12892671 |
| 1700086D15Rik | -4.510203247 | 0.1289621 |
| Fzd7 | -0.756830983 | 0.129059887 |
| Fam181a | 1.724994035 | 0.129062443 |
| Madd | 0.458700378 | 0.129062443 |
| Dsc2 | 4.781147788 | 0.129089581 |
| Sorbs3 | 0.86259999 | 0.129117926 |
| Mfap5 | 4.672934405 | 0.129226239 |
| Ptar1 | -0.550348351 | 0.129443728 |
| Dock7 | -0.399183025 | 0.129546706 |
| Insrr | 2.506536834 | 0.129574223 |
| Kcnk1 | 1.025765669 | 0.129693301 |
| Sec31b | 4.685409033 | 0.129719565 |
| Flnb | 0.55169141 | 0.129780393 |
| Fhdc1 | 4.536484816 | 0.129844557 |
| Kif16b | -0.831109746 | 0.12988751 |
| Tubb4a | -0.39708432 | 0.129891028 |
| Usp19 | -0.52670769 | 0.12995224 |
| Thrsp | 1.446225599 | 0.129996714 |
| Ldb2 | 0.941439945 | 0.129996714 |
| Drg2 | 0.41363105 | 0.129996714 |
| Rpl22 | 0.320584408 | 0.130005386 |
| Pcyox1 | -0.422171131 | 0.130005386 |
| Hapln1 | 4.744065311 | 0.130052037 |
| Six4 | 1.893552022 | 0.130100306 |
| Arl14 | 5.094842092 | 0.13014736 |
| Dpp6 | -0.501462256 | 0.13014736 |
| Dlc1 | 1.157824663 | 0.130196534 |
| Gm9776 | 3.389100722 | 0.130244558 |
| Abhd17b | -0.472030533 | 0.130388699 |
| Alad | 0.363397912 | 0.130405977 |
| Terf2ip | 0.512898495 | 0.130483379 |
| Ormdl3 | 0.530910679 | 0.130672478 |
| Ahcyl1 | -0.357036838 | 0.130709414 |
| Rgs7bp | 0.809261629 | 0.130712649 |
| Npr3 | -2.17451334 | 0.130712649 |
| Edf1 | 0.294997052 | 0.130841559 |
| Zfp771 | 0.423402609 | 0.130866125 |
| Gmpr2 | 0.451164746 | 0.130872498 |
| Pabpc4 | 0.68710402 | 0.130985403 |
| Lypla2 | 0.56160784 | 0.130985403 |
| Prr14 | -0.537534794 | 0.130985403 |
| BC004004 | 0.417346026 | 0.130994489 |
| Herpud1 | -0.449274297 | 0.130994489 |
| Recql | -0.625196689 | 0.130994489 |
| Ube2d3 | -0.349888159 | 0.131009004 |
| Plvap | 1.743967362 | 0.131122606 |
| Fbxw10 | 3.493873347 | 0.131255625 |
| Igsf3 | -0.418886235 | 0.13126229 |
| Apof | -5.501096692 | 0.131440333 |
| Nsa2 | 0.502103622 | 0.13145297 |
| Abhd6 | 0.85321107 | 0.131494551 |
| Slc6a12 | -1.616862624 | 0.131531282 |
| Wdr86 | 1.932908032 | 0.131590212 |
| Kat2a | 0.543451563 | 0.131725615 |
| Eif4e2 | 0.432696108 | 0.131725615 |
| Celf1 | 0.338919413 | 0.131780097 |
| Ralgps1 | -0.488242988 | 0.131816634 |
| Tfr2 | 2.012982479 | 0.132024784 |
| Ankrd50 | -0.666130515 | 0.132062094 |
| Cep95 | -0.809949368 | 0.132062094 |
| Ddi2 | -0.437535497 | 0.132293858 |
| Zdhhc17 | 0.594974104 | 0.132387702 |

| | | |
|---|---|---|
| Ccl24 | 1.120119379 | 0.460632117 |
| Spag4 | 0.644542872 | 0.460732796 |
| Sdr42e1 | -1.12532972 | 0.460832542 |
| Rnase6 | 1.269836129 | 0.461037265 |
| Ric3 | 0.363126702 | 0.461037265 |
| D1Ertd622e | -0.198997443 | 0.461037265 |
| Hps1 | -0.352523532 | 0.461037265 |
| Mtus2 | 0.394266998 | 0.461057254 |
| Scn9a | 0.757393881 | 0.461059808 |
| Tbl1x | 0.223133709 | 0.461315453 |
| Mapk8 | -0.181798295 | 0.461315453 |
| Myh10 | 0.207225262 | 0.461429187 |
| Tsnax | 0.195788188 | 0.461429187 |
| Rbm26 | 0.179088659 | 0.461429187 |
| Hic2 | -0.359614272 | 0.461429187 |
| Hsf2 | 0.188651088 | 0.461505058 |
| Rps18-ps3 | 0.336008168 | 0.461677207 |
| Atl2 | -0.338554708 | 0.461748814 |
| Mmgt2 | 0.321105531 | 0.462257996 |
| Rgs13 | 1.33739601 | 0.462258103 |
| Kif22 | 0.301198726 | 0.462464067 |
| Tgs1 | 0.255385425 | 0.462652058 |
| Rrm2b | 0.396947831 | 0.462830384 |
| Zfp518b | 0.272570358 | 0.462879502 |
| Cacnb3 | 0.243749481 | 0.462879502 |
| Adcy9 | -0.922816584 | 0.462879502 |
| Taok2 | -0.230503938 | 0.463634358 |
| Lrrc55 | 0.678773797 | 0.46370639 |
| Sp3 | -0.212290096 | 0.46370639 |
| Ppp1r13b | -0.29843989 | 0.46370639 |
| Dcun1d3 | -0.433482576 | 0.46370639 |
| S100a13 | -0.373043249 | 0.464029554 |
| Lclat1 | -0.415304559 | 0.464227457 |
| Ccdc43 | 0.274740384 | 0.46425836 |
| Elp4 | 0.263667206 | 0.46425836 |
| Lman1 | -0.291319485 | 0.46425836 |
| Polg | -0.316163515 | 0.464379632 |
| 3010026O09Rik | 0.543391554 | 0.464440496 |
| Gxylt1 | 0.388951393 | 0.464775144 |
| Tcfl5 | 3.949390318 | 0.465001138 |
| Gm10912 | 2.718809455 | 0.465277268 |
| Lor | 1.078697334 | 0.465277268 |
| Dna2 | 0.462096338 | 0.465277268 |
| Pik3r1 | 0.239927593 | 0.465277268 |
| Chd6 | 0.223313962 | 0.465277268 |
| Tcf25 | 0.175996294 | 0.465277268 |
| Copg1 | -0.190815191 | 0.465277268 |
| Dcaf12 | -0.19108701 | 0.465277268 |
| Robo1 | -0.286355099 | 0.465277268 |
| Kcna5 | -1.386831504 | 0.465277268 |
| Ipo7 | -0.191610244 | 0.465422943 |
| Gatad1 | 0.260076022 | 0.465827303 |
| Adrb1 | -0.736212706 | 0.465923597 |
| Dcdc2b | 1.262164316 | 0.465927743 |
| Tmem200a | 0.553216115 | 0.46625463 |
| Gdf5 | 3.357639328 | 0.466365706 |
| 5430427O19Rik | -0.880858042 | 0.466365706 |
| Ssr4 | 0.214140799 | 0.466458099 |
| Bloc1s3 | -0.332232725 | 0.466458099 |
| Fcrls | -0.623436474 | 0.466458099 |
| Amn | 1.752367012 | 0.466653262 |
| Klhl28 | 0.361132425 | 0.466678824 |
| Phka2 | -0.328173354 | 0.466678824 |
| Pip5kl1 | 0.886575002 | 0.466705616 |
| Dhcr7 | -0.28326993 | 0.466899406 |
| 2200002D01Rik | 3.332383741 | 0.467058094 |
| Gps1 | 0.222698176 | 0.467081217 |
| Cdk14 | -0.212216208 | 0.467089002 |
| Smim5 | 0.996477223 | 0.467110536 |
| Tle6 | -0.878159769 | 0.467443821 |
| Pbx1 | 0.184562265 | 0.467448963 |
| Cdk5rap2 | 0.241256392 | 0.46748544 |
| Evi5 | -0.279463216 | 0.467596003 |
| Kcnab3 | 0.625658094 | 0.467606295 |
| Cep131 | -0.339070025 | 0.467606295 |
| Ube2d2a | 0.181275829 | 0.467813209 |
| Irx5 | -0.888215385 | 0.467813209 |
| Aatf | -0.218870424 | 0.46822832 |
| Neil3 | -0.363025276 | 0.46822832 |
| Dhfr | 0.301538671 | 0.468414356 |
| Atp11c | -0.273283237 | 0.468414356 |
| Gm16432 | 1.260849038 | 0.468718365 |
| Ccna2 | 0.287886242 | 0.468718365 |
| Pear1 | 0.91381049 | 0.468722421 |
| Ccnb1 | 0.268269455 | 0.468722421 |
| Ccnl1 | 0.255647742 | 0.468722421 |
| Cxxc1 | -0.251739821 | 0.468722421 |
| Ttll12 | -0.266591052 | 0.468722421 |
| Robo3 | -1.339539149 | 0.468722421 |
| Tlr3 | 1.779847631 | 0.468966558 |
| Yod1 | -0.206792874 | 0.468966558 |
| E130309D14Rik | -0.54303138 | 0.468966558 |
| B3gnt8 | -0.596852786 | 0.468966558 |
| Zfp330 | 0.213538141 | 0.469331851 |
| Vwa9 | -0.245024208 | 0.469331851 |
| Ddx5 | -0.14344741 | 0.46941173 |
| Bcar1 | -0.257067361 | 0.469741587 |
| Usf1 | -0.225604926 | 0.46977183 |
| Ept1 | -0.241676054 | 0.469817673 |
| Mcidas | -1.304222504 | 0.469817673 |
| Gm2237 | 0.68816055 | 0.469937266 |
| Ippk | 0.511576139 | 0.469990526 |
| Ppp1r21 | -0.320088354 | 0.470028193 |
| BC030867 | -0.628485583 | 0.470028193 |
| Plat | 0.441791254 | 0.470031554 |
| Tmem163 | 0.529444953 | 0.470049535 |
| Lancl2 | 0.313610498 | 0.470049535 |
| Fam114a2 | 0.226444288 | 0.470049535 |
| Zfp334 | -0.361904546 | 0.470049535 |
| Gm3278 | -3.389791206 | 0.470049535 |
| Wfdc3 | 3.639071497 | 0.470432928 |
| Cox7a1 | -0.919003649 | 0.470432928 |
| Porcn | 0.293798842 | 0.470463576 |
| Vars2 | -0.558095816 | 0.470537139 |
| Eif4b | 0.199124374 | 0.470776419 |
| Kbtbd7 | -0.525967841 | 0.470806412 |
| Gng10 | 1.372498803 | 0.470959928 |
| Copb1 | -0.166068131 | 0.471109068 |
| 9030617O03Rik | 1.096920533 | 0.471123652 |

| | | |
|---|---|---|
| Eme1 | -0.169523176 | 0.851930849 |
| Acad10 | -0.312263085 | 0.851930849 |
| Hist1h3h | 0.299998726 | 0.852052738 |
| Recql4 | 0.394031388 | 0.852360815 |
| Atg5 | -0.06198051 | 0.852503463 |
| Fhod3 | 0.105474671 | 0.852612295 |
| Galr1 | -0.512346198 | 0.852663711 |
| Tmem18 | 0.092458124 | 0.852677534 |
| Hspa14 | 0.055337395 | 0.852677534 |
| Zfp46 | 0.100730071 | 0.85269297 |
| Gm9242 | 0.794042544 | 0.852795671 |
| Adgrb2 | 0.131955557 | 0.853118189 |
| Cxcr5 | -0.330941662 | 0.853170348 |
| Retn | -0.881999332 | 0.853189167 |
| Mak | 0.648985759 | 0.853219479 |
| Rpe | 0.091854394 | 0.853284252 |
| Ftsj3 | 0.082959491 | 0.853337518 |
| Ythdc1 | 0.063045239 | 0.853500136 |
| Nr2f6 | -0.169788181 | 0.853526171 |
| Cd27 | -0.835742452 | 0.853565201 |
| Ccdc36 | 0.384368854 | 0.853792263 |
| Pusl1 | 0.083358885 | 0.853797181 |
| Dusp7 | 0.191121783 | 0.853808677 |
| Bcl7b | 0.070046923 | 0.85381273 |
| 2810417H13Ri | -0.080804302 | 0.85388898 |
| Cluap1 | -0.138095412 | 0.854031261 |
| Mchr1 | -0.475611095 | 0.854226527 |
| Tcaf1 | -0.048486504 | 0.854249646 |
| Clcn4 | -0.062133205 | 0.854287064 |
| Pon1 | -0.446370206 | 0.854349577 |
| Hlf | 0.203893084 | 0.854711833 |
| Jun | 0.059947111 | 0.854711833 |
| Nbr1 | 0.061182844 | 0.854899825 |
| Lgalsl | -0.069982546 | 0.854964817 |
| Nqo1 | -0.319037251 | 0.855065408 |
| Prodh | -0.228669315 | 0.855234324 |
| 3110039M20Ri | -0.107349956 | 0.855278257 |
| Itch | 0.061892834 | 0.85539972 |
| Cebpg | -0.073925714 | 0.85539972 |
| Tmem254c | 0.449063906 | 0.855467796 |
| Hfm1 | -0.300065902 | 0.855704997 |
| Katnal1 | -0.101845568 | 0.855795657 |
| Zfp764 | -0.222557083 | 0.855824784 |
| Ager | -0.418686122 | 0.856009231 |
| Abcc2 | 0.434296454 | 0.856097997 |
| Nr1d2 | -0.124965176 | 0.856137547 |
| Cul4b | -0.09561183 | 0.856150157 |
| Prrt2 | -0.339366372 | 0.856222072 |
| Slc4a2 | -0.088729651 | 0.856278487 |
| Samd1 | 0.142659629 | 0.856301949 |
| Josd2 | 0.077124804 | 0.856327019 |
| Tulp4 | 0.058118365 | 0.856327019 |
| Il18bp | -0.142221682 | 0.856343362 |
| Tef | -0.077550227 | 0.856656844 |
| Nol11 | 0.063104353 | 0.85672753 |
| Ifi27l2a | -0.101886179 | 0.857053639 |
| Adrm1 | 0.0654083 | 0.857192804 |
| Rgag4 | -0.137590838 | 0.857242845 |
| Mbp | -0.10655109 | 0.857534574 |
| Trmt12 | 0.149982727 | 0.857565707 |
| Uqcrc2 | -0.044993453 | 0.857565707 |
| Mmachc | 0.088880508 | 0.857671763 |
| Taf3 | 0.075106545 | 0.857681969 |
| Arl1 | -0.045863939 | 0.858087089 |
| Chchd10 | 0.11250326 | 0.858353466 |
| Samm50 | -0.053917647 | 0.858353466 |
| 4932438A13Ri | -0.06161297 | 0.858534149 |
| 1810041L15Ri | 0.17356949 | 0.858726528 |
| Ube2s | 0.069656963 | 0.858726528 |
| Clta | 0.050699486 | 0.858726528 |
| Kcnk12 | -0.177554036 | 0.858726528 |
| Tmed9 | -0.055914277 | 0.858989988 |
| Cep350 | 0.081427812 | 0.85907103 |
| 1700001P01Ri | 0.607449442 | 0.859210722 |
| 6-Mar | 0.090918891 | 0.859210722 |
| Shank1 | -0.087929789 | 0.859210722 |
| Gpr89 | -0.101057352 | 0.859210722 |
| Pcdhb15 | 0.208141441 | 0.859211022 |
| Cck | -0.11097318 | 0.859211022 |
| Slx1b | -0.116311803 | 0.859211022 |
| Cryba1 | -0.459378537 | 0.859211022 |
| Dnah12 | 0.36493831 | 0.859326008 |
| Mterf4 | 0.08480246 | 0.859326008 |
| Cxcl13 | -0.278862027 | 0.859354833 |
| Zbtb5 | -0.085700028 | 0.859473197 |
| Tbr1 | -0.100872931 | 0.859486088 |
| Tmem19 | -0.100947985 | 0.859792918 |
| Rbm11 | 0.439114411 | 0.859903236 |
| 1700028J19Rik | 0.437028893 | 0.859903236 |
| Gm13251 | 0.102572748 | 0.859903236 |
| Phtf2 | 0.081548833 | 0.859903236 |
| Fgfr1op | 0.068495718 | 0.859903236 |
| Amn1 | -0.06472394 | 0.859903236 |
| Arhgap10 | -0.17607687 | 0.859903236 |
| BC024139 | -0.227712887 | 0.859903236 |
| Rbm42 | 0.045712918 | 0.859953356 |
| Pklr | 0.857262963 | 0.859980956 |
| Mpp7 | 0.107768037 | 0.860021528 |
| Zdhhc24 | 0.155461174 | 0.860201145 |
| Fbxo4 | 0.105529875 | 0.860201145 |
| Mn1 | -0.070827632 | 0.860201145 |
| Dnajc1 | -0.077884134 | 0.860201145 |
| Apip | 0.077786591 | 0.860482572 |
| Foxp1 | 0.06466143 | 0.860621399 |
| Thumpd2 | -0.121499641 | 0.860621399 |
| Gm3173 | -0.450407899 | 0.860621399 |
| Ppp2r5c | 0.061910175 | 0.86091016 |
| Lasp1 | -0.064291122 | 0.86091016 |
| Zfp692 | -0.112312374 | 0.86091016 |
| Tmem200c | -0.330773792 | 0.86091016 |
| Nudt16 | 0.087687801 | 0.861090895 |
| Psma8 | -0.270721988 | 0.861104116 |
| Klhdc9 | 0.265689673 | 0.861143509 |
| Cracr2b | 0.194645769 | 0.861178963 |
| Krt73 | 0.440087009 | 0.861188467 |
| Specc1 | 0.090169375 | 0.861188467 |
| Bach2 | 0.07644505 | 0.861188467 |
| Bcl2l15 | -0.420682722 | 0.861294275 |
| Zfp53 | -0.098963907 | 0.861570802 |

| | | |
|---|---|---|
| Fhl1 | 0.85376699 | 0.0030059 |
| Nlrp1a | -3.350249035 | 0.003009091 |
| Vsir | -0.974197624 | 0.003018046 |
| Ccl22 | -6.78219075 | 0.003034127 |
| Nadk2 | -1.03278692 | 0.003039855 |
| Gm13212 | -1.993666534 | 0.003044984 |
| Dnal4 | 0.737194584 | 0.003059382 |
| Atp5b | 0.506782861 | 0.003063958 |
| Tmem246 | 0.891332782 | 0.003069488 |
| Spata5 | -0.867172099 | 0.003078734 |
| Rpl27 | 0.623598521 | 0.003087225 |
| Alox5 | -1.980180747 | 0.003087225 |
| Cdh12 | -1.926883425 | 0.003102277 |
| Rgl1 | -1.32840057 | 0.003106388 |
| Deaf1 | 0.696073603 | 0.003108328 |
| Gucy1b2 | 2.809197034 | 0.003112395 |
| Usp7 | 0.836994151 | 0.003120453 |
| Srebf2 | -0.653737359 | 0.003128467 |
| Tpm4 | -0.664861557 | 0.003128467 |
| Sat2 | 1.515842728 | 0.003134173 |
| Lgr4 | 1.208212009 | 0.003138942 |
| Nfe2l2 | -1.093834122 | 0.003141543 |
| Hpgd | 1.727500614 | 0.003141624 |
| Capns2 | -6.649281809 | 0.003163872 |
| Thbs3 | 1.289491409 | 0.003182057 |
| 2810004N23Rik | 0.870196939 | 0.003183508 |
| Spef1 | 1.335402771 | 0.003208101 |
| Dclre1c | -0.992916409 | 0.003208101 |
| Itpripl2 | -2.047442531 | 0.003208101 |
| Hp | -3.126130695 | 0.003208101 |
| Mthfr | -1.436216911 | 0.003215674 |
| Mrpl14 | 0.833262823 | 0.003219697 |
| Pgm5 | 6.102719036 | 0.003226656 |
| Mrps16 | 0.751759987 | 0.003228947 |
| Dap | -0.897327442 | 0.003230373 |
| Fam69c | 5.961036948 | 0.003243952 |
| Ugt1a7c | -2.340309843 | 0.003258483 |
| Itga1 | 2.910140815 | 0.003262377 |
| Mob1a | -0.996887878 | 0.003264102 |
| Cir1 | 0.934733324 | 0.003273415 |
| Ypel1 | 0.788547652 | 0.003287475 |
| Pea15a | 0.757166785 | 0.003287475 |
| Plekhm3 | -1.180916118 | 0.003287475 |
| Slc5a11 | -6.674373278 | 0.003287475 |
| Ss18 | -0.756240914 | 0.003298928 |
| Srm | 0.775496792 | 0.003305259 |
| Egfr | 1.892357417 | 0.00331858 |
| Eif2b2 | 0.820529324 | 0.003318747 |
| Cul3 | 0.700577327 | 0.003318747 |
| Mogs | -1.250649481 | 0.003318747 |
| Ubxn4 | 0.637554515 | 0.003321502 |
| Mfsd4 | -1.119457372 | 0.003330643 |
| Il10rb | -1.286949034 | 0.003330643 |
| Fibp | 0.678502891 | 0.003335789 |
| Kynu | 2.390257265 | 0.003345423 |
| Rtl1 | -2.192453924 | 0.003345423 |
| Rarb | 2.385143407 | 0.003360225 |
| Chchd2 | 0.495975973 | 0.003376189 |
| Kif3a | 0.745713644 | 0.003385249 |
| Anxa7 | -1.365098391 | 0.003385302 |
| Gpr155 | -1.380717771 | 0.003394326 |
| Btbd3 | 0.983557779 | 0.003401727 |
| Eif3m | 0.568289567 | 0.00341829 |
| Rcc2 | 0.687025028 | 0.003418654 |
| Zfp12 | -1.036761746 | 0.00341893 |
| Gm17455 | 4.662369611 | 0.003422371 |
| Zfyve26 | -1.12275372 | 0.00342299 |
| Nmral1 | 0.78311953 | 0.003423762 |
| H2afy2 | 0.678716376 | 0.003448888 |
| Ssh2 | -0.783862522 | 0.003448888 |
| Cfp | -2.195934806 | 0.003448888 |
| F10 | -5.435532218 | 0.003448888 |
| Spty2d1 | -0.956269396 | 0.003455184 |
| Ccdc149 | -1.59341993 | 0.00345737 |
| Nhp2l1 | 0.61731714 | 0.003458389 |
| Gm9930 | 6.032865927 | 0.003463059 |
| Apbb2 | 0.855223781 | 0.003467805 |
| Tomm5 | 0.836222939 | 0.003486504 |
| Gper1 | 5.773729995 | 0.003522272 |
| Cd52 | -0.951585809 | 0.003542692 |
| Ifitm2 | -1.073218061 | 0.003551428 |
| Os9 | -0.62205849 | 0.003555875 |
| 2010107G23Rik | 0.954157597 | 0.003560271 |
| Rps17 | 0.515173696 | 0.003560271 |
| Btf3l4 | 0.576222095 | 0.003560685 |
| 1110038F14Rik | 0.893138208 | 0.003573351 |
| Lyar | 0.779193834 | 0.003573351 |
| Col13a1 | 5.995866795 | 0.003594767 |
| Mkl2 | -0.859600089 | 0.003598659 |
| Alg12 | 2.000041807 | 0.003600929 |
| Rims2 | 1.303003341 | 0.003612992 |
| Mxd3 | 1.477694657 | 0.003613826 |
| Taf6l | -0.662875787 | 0.003620213 |
| Irg1 | -3.22574535 | 0.003627937 |
| Bmp6 | 6.03186083 | 0.003635513 |
| Pam16 | 0.738849266 | 0.003646075 |
| Atg7 | -0.80949378 | 0.003646075 |
| Myo1e | -1.681393189 | 0.003646075 |
| Zfp74 | 1.392356045 | 0.003656987 |
| Gstm1 | 0.924883122 | 0.003656987 |
| Il1b | -3.08319597 | 0.003656987 |
| Lmtk2 | -0.937344474 | 0.003663199 |
| Ip6k2 | 0.681758221 | 0.003664361 |
| Bcs1l | 0.975474731 | 0.003676254 |
| Mphosph6 | 0.914298942 | 0.003676254 |
| Sox8 | 1.775934347 | 0.00368013 |
| L3mbtl3 | 0.889201592 | 0.003680262 |
| Col8a1 | 5.99347342 | 0.003680514 |
| Sdha | -0.600131463 | 0.003704613 |
| Dnajb2 | 0.955773953 | 0.003710122 |
| Pwp1 | -0.832154637 | 0.00371462 |
| Ctsk | 5.946524591 | 0.003718348 |
| Slc14a2 | 2.192434196 | 0.003718348 |
| Nacc2 | 1.128575649 | 0.00372442 |
| Zbtb34 | -1.301824579 | 0.003763126 |
| Psip1 | 0.638810136 | 0.003764944 |
| Azi2 | -0.590830162 | 0.003764944 |
| Mrgpra2a | -3.344336721 | 0.003764944 |
| Fancl | 1.259796364 | 0.00376527 |

| | | |
|---|---|---|
| P3h2 | 5.10097407 | 0.132521021 |
| Acsl3 | 0.496347984 | 0.132554708 |
| Esyt2 | -0.660042333 | 0.132554708 |
| Plet1 | 4.973919314 | 0.1326481 |
| Trappc13 | 0.443896666 | 0.132720651 |
| Hmox2 | -0.441678673 | 0.132838918 |
| Mmp12 | 4.696659191 | 0.132958549 |
| Slc9a8 | -0.530950773 | 0.132964827 |
| Trpm5 | 3.396725284 | 0.133120137 |
| Birc3 | -1.639448892 | 0.133120137 |
| Rtp4 | -1.683525804 | 0.133120137 |
| Rab17 | -5.499279066 | 0.133120137 |
| Sstr3 | -5.499279066 | 0.133120137 |
| Slfn1 | -2.553608562 | 0.133126513 |
| Gmds | 0.566641327 | 0.133245413 |
| Gm10277 | 2.189863114 | 0.133247307 |
| Nek1 | 0.615768305 | 0.133266839 |
| Golph3 | -0.349646106 | 0.133288352 |
| Crispld2 | -1.784471786 | 0.133305024 |
| Zfp523 | 0.56468218 | 0.133350636 |
| Psmg2 | 0.476350474 | 0.133505053 |
| Rraga | 0.370359725 | 0.133505053 |
| Trdn | 4.612298373 | 0.133509207 |
| Ap1ar | -0.559163478 | 0.133509207 |
| Csnk2b | 0.360085359 | 0.133603816 |
| Senp3 | -0.441792134 | 0.133603816 |
| Mthfd1l | -0.794126439 | 0.133603816 |
| Tlk2 | -0.434738304 | 0.133622386 |
| Dhx16 | -0.460488683 | 0.133767927 |
| Ndor1 | -0.569770857 | 0.1340199 |
| Gapdhs | 0.937482482 | 0.134038478 |
| Pik3c2g | 4.677818806 | 0.134084502 |
| Med26 | -0.617220463 | 0.134084502 |
| Upk1a | 3.590042985 | 0.13417046 |
| Timmdc1 | 0.496809657 | 0.134294699 |
| Btaf1 | -0.433794922 | 0.134316801 |
| Abhd18 | -0.73270869 | 0.134348416 |
| Atr | -0.512874466 | 0.134436023 |
| Colgalt2 | 2.373791567 | 0.13445575 |
| Ovgp1 | -1.803109868 | 0.13464978 |
| Ankrd6 | 0.484941191 | 0.134854799 |
| Dgkk | 4.714763907 | 0.134882183 |
| Naa40 | -0.413561205 | 0.134962956 |
| Noxa1 | 4.593590311 | 0.134997373 |
| Hbb-bt | 1.157348114 | 0.135008325 |
| Gaa | -0.490455011 | 0.135102334 |
| Gabra5 | 0.885018407 | 0.135111252 |
| Dtx3l | -0.76017254 | 0.13511643 |
| Slc22a14 | 4.791269754 | 0.135125508 |
| Pptc7 | -0.910707455 | 0.135319024 |
| C2cd2 | -0.823828392 | 0.135331431 |
| Jrkl | 0.871234723 | 0.135413502 |
| Hibadh | -0.390798751 | 0.135416602 |
| Mlxipl | 4.042860411 | 0.135521573 |
| Hist2h2ab | 4.022534104 | 0.135521573 |
| Eif3h | 0.325491191 | 0.135521573 |
| Stk17b | -0.579665402 | 0.135521573 |
| C130032M10Rik | -5.149359358 | 0.135521573 |
| Arpc4 | -0.433820465 | 0.135587021 |
| Zfp804a | 1.188426261 | 0.13563112 |
| Fgf22 | -4.735845188 | 0.135859979 |
| Ly6g6e | 1.278970416 | 0.135863407 |
| Trrap | -0.37083183 | 0.135878441 |
| Umad1 | 0.510169089 | 0.136037754 |
| Nrbp1 | 0.388069336 | 0.136073692 |
| Sfswap | -0.453261036 | 0.136077878 |
| Suv39h2 | -0.626958256 | 0.136142843 |
| Rsrp1 | 0.449281272 | 0.136193936 |
| Naa20 | 0.350293918 | 0.136193936 |
| Nfat5 | -0.570606963 | 0.136193936 |
| Pgpep1l | 4.650727725 | 0.13627211 |
| Smpd2 | 0.741273411 | 0.13627211 |
| Spidr | -0.66742835 | 0.13627211 |
| Col11a2 | 3.072308425 | 0.136359713 |
| Idh3g | 0.319149429 | 0.136359713 |
| Atp1a3 | -0.536845592 | 0.136359713 |
| Emc9 | 0.651398971 | 0.136416069 |
| Nsg1 | 0.390382368 | 0.136497771 |
| Acsm3 | -5.1243137 | 0.136566767 |
| Kif13a | 0.910430772 | 0.136625233 |
| Pim1 | -0.59519742 | 0.136625233 |
| Gm1043 | -1.987959953 | 0.13670248 |
| 1700019A02Rik | 4.676607198 | 0.136951419 |
| Steap4 | -3.87951675 | 0.137158326 |
| Btbd7 | -0.465451148 | 0.137245335 |
| Gm11110 | 1.228053888 | 0.137343843 |
| Egln2 | -0.43149771 | 0.137431636 |
| Dcaf12l1 | 1.097798875 | 0.137453718 |
| Mef2a | -0.366685209 | 0.137453718 |
| Cyp4x1 | 1.434932345 | 0.137561816 |
| St3gal6 | -0.619576091 | 0.137561816 |
| Lgals3bp | -0.743579759 | 0.137561816 |
| Axl | -0.649058201 | 0.137589448 |
| App | -0.348755045 | 0.137602941 |
| Thoc6 | -0.617955713 | 0.137676481 |
| Bzw2 | 0.409320246 | 0.137690802 |
| Tmem53 | 1.488957105 | 0.137765887 |
| Atad2b | -0.574077195 | 0.137843773 |
| Fgf17 | 3.503094295 | 0.137958508 |
| Cldn20 | -5.462127593 | 0.137958508 |
| Sirt1 | 1.034230269 | 0.138022348 |
| Parp3 | 4.11285365 | 0.138082729 |
| Scp2 | 0.361131535 | 0.138194364 |
| Zfp444 | 0.355004826 | 0.138220901 |
| Jph1 | -1.349976025 | 0.138220901 |
| Tm2d2 | 0.319732267 | 0.138340858 |
| Klhl29 | -0.590713688 | 0.138487274 |
| Hddc2 | 0.437468497 | 0.138653377 |
| Hnrnph2 | -0.310270603 | 0.138653377 |
| Cxcl17 | 3.991397755 | 0.138798477 |
| Rfx7 | 0.437965772 | 0.13881137 |
| Lmcd1 | 1.237076332 | 0.138824957 |
| Nelfe | 0.438194996 | 0.139044053 |
| Nutf2-ps1 | 0.641210507 | 0.139109365 |
| Eid1 | 0.420228719 | 0.139165089 |
| Rabl2 | 0.753807925 | 0.139168374 |
| Plekhg4 | -1.985441221 | 0.139510826 |
| Slc6a7 | 2.101731439 | 0.139917973 |
| Rwdd2a | 0.898053627 | 0.13995898 |

| | | |
|---|---|---|
| Ralbp1 | 0.220918766 | 0.471360695 |
| Kpna2 | 0.229046471 | 0.471474668 |
| Pcp2 | -3.389840517 | 0.471585088 |
| Trim35 | -0.154791866 | 0.471726357 |
| Txnrd2 | -0.320621178 | 0.471943398 |
| Tnfrsf10b | 1.085328995 | 0.471992766 |
| Zfp322a | 0.22583746 | 0.472250867 |
| Slc16a14 | 0.396390961 | 0.472302957 |
| Rhbdd3 | 0.305318809 | 0.472323024 |
| Isca1 | 0.211737651 | 0.472400709 |
| Naip1 | -2.34240649 | 0.472653639 |
| Poc1a | 0.433925224 | 0.472683496 |
| Bend6 | 0.324946024 | 0.472900752 |
| Rbsn | 0.214708565 | 0.472900752 |
| Lck | 0.77704653 | 0.472926387 |
| 9430016H08Rik | 0.299828193 | 0.472926387 |
| Capn7 | -0.212512608 | 0.472926387 |
| Riok3 | -0.268718352 | 0.472926387 |
| Klhl26 | -0.482464369 | 0.472926387 |
| 3110070M22Rik | 0.815348869 | 0.473019771 |
| 5430403G16Rik | -1.222444189 | 0.473025763 |
| Mtif2 | 0.267152865 | 0.473084109 |
| Osbpl6 | -0.375616238 | 0.473180935 |
| Vipr1 | -1.313731096 | 0.473180935 |
| H2-DMb1 | 1.399736327 | 0.473359702 |
| Cebpa | 0.674438616 | 0.473417572 |
| Hectd1 | 0.174038677 | 0.473417572 |
| Tars | -0.23601567 | 0.473417572 |
| Rnf121 | -0.249382529 | 0.473417572 |
| Ephb6 | -0.279667968 | 0.473417572 |
| Pdpr | -0.33537534 | 0.473417572 |
| Cfap52 | -1.64964658 | 0.473417572 |
| 2010315B03Rik | 0.535449793 | 0.473460062 |
| Maged1 | 0.157878677 | 0.473460062 |
| H2-Ke6 | 0.230809044 | 0.473606488 |
| Trpm7 | -0.196466301 | 0.473606488 |
| Evc2 | -0.746190688 | 0.473749163 |
| B130006D01Rik | -0.811460554 | 0.474107001 |
| Gen1 | -0.529774802 | 0.474126839 |
| Ascl4 | -1.531532299 | 0.474356036 |
| Abca8a | 2.361673053 | 0.47440022 |
| Ube2e1 | 0.233923279 | 0.474569352 |
| Faim2 | 0.569275925 | 0.474699949 |
| Dnm1l | -0.204821325 | 0.474699949 |
| Atf3 | -0.273014327 | 0.474699949 |
| Gm7008 | 1.029757222 | 0.474784968 |
| Pask | 0.456876635 | 0.474784968 |
| Srrm3 | 0.35146738 | 0.474784968 |
| Syt14 | -0.377819696 | 0.474784968 |
| Gabrg3 | -0.805903816 | 0.474784968 |
| Prima1 | -2.070275746 | 0.474784968 |
| Gsto2 | -0.997769022 | 0.474806674 |
| Rfng | 0.279113885 | 0.475212754 |
| Rae1 | 0.18954396 | 0.475212754 |
| Fbxo21 | -0.211888464 | 0.475212754 |
| Xpr1 | -0.260239695 | 0.475261829 |
| Rce1 | -0.246529094 | 0.475338716 |
| Ebna1bp2 | 0.215674673 | 0.475393241 |
| Psg23 | 3.906219802 | 0.4754981 |
| Vmn2r18 | 3.906219802 | 0.4754981 |
| Grip1 | 0.305283244 | 0.475604502 |
| Arhgap11a | 0.3303148 | 0.475863938 |
| Fads2 | 0.235710852 | 0.475863938 |
| Xiap | -0.228598572 | 0.475863938 |
| Trappc11 | -0.329450101 | 0.475863938 |
| Shisa9 | -0.557345058 | 0.475863938 |
| Trmt1 | 0.250835484 | 0.475874131 |
| Slc50a1 | -0.27401512 | 0.475881243 |
| Sertad3 | 0.438311474 | 0.475895814 |
| Clk1 | -0.161677302 | 0.475895814 |
| Pias3 | -0.318851331 | 0.475895814 |
| Ireb2 | 0.251721594 | 0.476199561 |
| Slc17a6 | 0.330353376 | 0.47622838 |
| Fam196a | 0.379679091 | 0.476232217 |
| Tle2 | -0.372681303 | 0.476232217 |
| Ino80b | 0.427405428 | 0.47638695 |
| Scamp2 | -0.230677128 | 0.476475537 |
| Magi3 | 0.340828141 | 0.47677101 |
| Pou2af1 | 3.568902282 | 0.476794887 |
| Klhdc8b | 0.353704748 | 0.476794887 |
| Lbh | -0.26824224 | 0.476794887 |
| Apoa2 | 1.707195893 | 0.477287313 |
| Zfp943 | 0.403590001 | 0.477779598 |
| Noxred1 | -0.925484613 | 0.477791784 |
| Sh3bgrl3 | -0.205695027 | 0.47791272 |
| Zfp281 | -0.214728039 | 0.47791272 |
| Slc38a9 | -0.31466809 | 0.47791272 |
| Cmtr1 | -0.535011549 | 0.47791272 |
| Scube3 | -0.640366444 | 0.47791272 |
| Capn6 | -0.770440167 | 0.47791272 |
| Cblc | -1.565188551 | 0.477936412 |
| Lrrc39 | -1.584132553 | 0.477936412 |
| Cdh22 | -0.888270184 | 0.477977261 |
| Puf60 | 0.169281325 | 0.478027907 |
| Crlf1 | 0.733803746 | 0.478237318 |
| Rep15 | 3.25019604 | 0.478844478 |
| Manea | 0.429635956 | 0.478844478 |
| Ing4 | 0.202872846 | 0.478926414 |
| Etohi1 | 0.677593378 | 0.479222911 |
| Lamtor1 | -0.212417315 | 0.479347246 |
| Csn1s2a | 3.603676875 | 0.479443612 |
| Yme1l1 | -0.165730748 | 0.479464337 |
| 1810013L24Rik | -0.305525695 | 0.479488339 |
| Lrfn1 | -0.501457574 | 0.479488339 |
| 4933407L21Rik | 1.342317478 | 0.479542876 |
| Gm6583 | -1.313895683 | 0.479542876 |
| Otud1 | -0.327160369 | 0.479883009 |
| Rasgrp1 | -0.869256058 | 0.480135209 |
| Abtb1 | -0.290884885 | 0.480684874 |
| Hist1h2an | 1.185490567 | 0.480714019 |
| Ifitm3 | -0.940919379 | 0.480714019 |
| Klhl12 | 0.228165575 | 0.480765116 |
| Pigk | 0.272478503 | 0.480809098 |
| Ogfod1 | -0.440446336 | 0.481280552 |
| Zfp647 | 0.41950695 | 0.481529538 |
| 4932414N04Rik | 3.596900876 | 0.481606576 |
| Gnat3 | 3.596900876 | 0.481606576 |
| Hsd17b2 | 3.596900876 | 0.481606576 |
| Pdc | 3.596900876 | 0.481606576 |

| | | |
|---|---|---|
| Slc19a2 | 0.151473427 | 0.861655267 |
| Nsmce4a | -0.074949125 | 0.861655267 |
| Padi4 | -0.373799394 | 0.861655267 |
| Mto1 | -0.077625902 | 0.861669344 |
| Pcnxl4 | 0.091091054 | 0.861919536 |
| Smtnl2 | 0.323262372 | 0.862339723 |
| Pdia3 | 0.050355407 | 0.862339723 |
| Crot | -0.116298465 | 0.862339723 |
| Hist1h2ak | -0.233127113 | 0.862339723 |
| Trim67 | 0.077989865 | 0.862539046 |
| Tfg | -0.051869718 | 0.862574628 |
| Elk1 | -0.07236342 | 0.862608862 |
| Cdk5rap3 | -0.053955542 | 0.862791249 |
| P2rx3 | 0.633635648 | 0.862995263 |
| Kdm8 | -0.338922707 | 0.862995263 |
| Nyap2 | -0.098680266 | 0.863042596 |
| Parvb | -0.174216171 | 0.863042596 |
| Fam213b | -0.067078281 | 0.863179339 |
| Fadd | -0.090611363 | 0.86333382 |
| Cenpc1 | -0.068896368 | 0.863674785 |
| Lpin2 | 0.062356678 | 0.863934708 |
| Tmem5 | 0.087150404 | 0.863943219 |
| Lhx6 | 0.111966376 | 0.863950969 |
| Nop16 | 0.062773095 | 0.864111801 |
| Ap2m1 | 0.039408763 | 0.864221176 |
| Rasal1 | 0.303311202 | 0.864263909 |
| Zfp438 | 0.255677328 | 0.864485584 |
| Dusp19 | 0.151900844 | 0.864485584 |
| Nbea | 0.071311904 | 0.864485584 |
| Dtl | -0.06103134 | 0.864485584 |
| Cdr2l | -0.074073854 | 0.864485584 |
| Stx1b | -0.085823249 | 0.864485584 |
| Atg4c | -0.104070278 | 0.864485584 |
| Zfp446 | -0.154094626 | 0.864485584 |
| Pcdhgb4 | -0.187995342 | 0.864485584 |
| Zfp28 | -0.115962398 | 0.864572587 |
| Setdb2 | -0.148626384 | 0.864687725 |
| Kcnv1 | -0.177524197 | 0.864834078 |
| 2310022B05Ri | 0.080154221 | 0.864840457 |
| Ddx10 | 0.06852889 | 0.864840457 |
| Stk38 | -0.053412922 | 0.864840457 |
| Cd300lg | 0.416977202 | 0.865202804 |
| Dnah3 | -0.614142652 | 0.865246622 |
| Cdkal1 | 0.07103065 | 0.865269658 |
| Ctdspl | 0.120700014 | 0.8653242 |
| Msra | 0.069805409 | 0.8653242 |
| Slc17a3 | 0.741090685 | 0.86543491 |
| Fank1 | -0.191282879 | 0.865790714 |
| Nsfl1c | 0.044704182 | 0.866252276 |
| Vimp | -0.064380942 | 0.866396707 |
| Mex3d | 0.091784058 | 0.866523204 |
| Adat2 | -0.117692167 | 0.866823937 |
| Cep57l1 | 0.068970053 | 0.866911539 |
| Lrrc19 | 0.242459845 | 0.866990149 |
| Acsbg2 | 0.680906018 | 0.867005576 |
| Pcsk1 | 0.406387971 | 0.867163141 |
| Tti2 | -0.090586865 | 0.867163141 |
| Pitpnm3 | -0.206037843 | 0.867163141 |
| Ndst3 | 0.085715793 | 0.867212355 |
| Surf6 | -0.08289689 | 0.867212355 |
| Ppm1a | -0.066834305 | 0.867502156 |
| Psors1c2 | -0.588939618 | 0.867672372 |
| Cnpy4 | 0.095043491 | 0.867767189 |
| Dr1 | -0.07124691 | 0.867767189 |
| Nom1 | -0.080754032 | 0.867918941 |
| Gatsl2 | 0.068325288 | 0.868213206 |
| 2410002F23Ril | 0.048269842 | 0.868213206 |
| Nlrc5 | -0.266348791 | 0.868213206 |
| Ada | 0.109271937 | 0.868328618 |
| Myef2 | 0.05081721 | 0.868528447 |
| Bloc1s4 | 0.071600312 | 0.868620539 |
| Zar1 | 0.687642264 | 0.868721005 |
| Ypel5 | 0.061167997 | 0.868721005 |
| Zdhhc3 | -0.055241056 | 0.868821888 |
| Bri3bp | -0.066100486 | 0.869029482 |
| Zfp189 | -0.11300793 | 0.869029482 |
| Vstm2l | -0.315520711 | 0.869029482 |
| 6030419C18Ri | 0.19910816 | 0.869071479 |
| Fbxl3 | 0.085261755 | 0.869147774 |
| Sh2d1b1 | 0.130859015 | 0.869178367 |
| Ncmap | 0.812810483 | 0.869217471 |
| Gpld1 | 0.278028187 | 0.869305727 |
| Zfp236 | -0.086641455 | 0.869305727 |
| Tpk1 | -0.144466751 | 0.869305727 |
| 4931423N10Ri | -0.561502339 | 0.869305727 |
| Ndufaf6 | 0.074597215 | 0.869372136 |
| Agl | -0.079019327 | 0.869584744 |
| Osbpl7 | 0.160039774 | 0.869593802 |
| Gucy2g | 0.672489349 | 0.869645662 |
| Slc27a5 | 0.324625721 | 0.869645662 |
| Mpp2 | -0.120600892 | 0.869645662 |
| Zbtb16 | -0.127480362 | 0.869899248 |
| Cd59a | 0.131353141 | 0.869904018 |
| Cox17 | 0.203114501 | 0.869920172 |
| Strip2 | 0.201429123 | 0.870295035 |
| Zfp408 | -0.084060721 | 0.870295035 |
| Eps8 | -0.104843852 | 0.870295035 |
| Hs3st1 | 0.094523943 | 0.870598443 |
| Psmd1 | -0.04684676 | 0.870598443 |
| Bet1 | 0.059490658 | 0.870710486 |
| Arntl | 0.094171025 | 0.870831439 |
| Ccser2 | 0.065612915 | 0.87084058 |
| Ccdc138 | 0.125473253 | 0.871196454 |
| Fam83d | -0.087504653 | 0.871496235 |
| Dtwd2 | -0.180806302 | 0.871689908 |
| Olfr494 | 0.308799878 | 0.872107281 |
| Utrn | -0.095244588 | 0.872107281 |
| Wls | 0.064818492 | 0.872130137 |
| Daw1 | 0.429212774 | 0.872475809 |
| Cwf19l1 | 0.067462455 | 0.872475809 |
| Stx17 | 0.077858855 | 0.872508675 |
| Ipo13 | 0.074758533 | 0.872581138 |
| Tbl3 | 0.05576643 | 0.872602726 |
| Nono | -0.039936995 | 0.872602726 |
| Commd7 | 0.055802817 | 0.872921192 |
| Dclk1 | -0.054301287 | 0.87302816 |
| Bbs9 | -0.093822676 | 0.87302816 |
| Dlg5 | 0.080581387 | 0.873277362 |
| Rph3a | 0.107432775 | 0.873341189 |

| | | |
|---|---|---|
| Rnf139 | -0.778835557 | 0.003780779 |
| Adpgk | -1.103347857 | 0.003780779 |
| Exosc7 | 0.733872608 | 0.003784011 |
| Eif4a2 | 0.628475461 | 0.003784011 |
| Hsd11b1 | -1.613551147 | 0.003786047 |
| Plekhb2 | -0.763181816 | 0.003786714 |
| Prim1 | 0.745542179 | 0.003791106 |
| Ncoa6 | -0.711189347 | 0.003794431 |
| Jade2 | -2.577544431 | 0.003794431 |
| Nat8l | 1.598546219 | 0.003802747 |
| Cyp4f16 | -1.097033612 | 0.003802811 |
| Stard5 | -0.99861402 | 0.003818708 |
| Kcnq5 | -1.990426501 | 0.003823958 |
| Chrna3 | -2.153148832 | 0.003823958 |
| Snrpa1 | 0.605721356 | 0.003829458 |
| Polb | 0.66256975 | 0.003829815 |
| Srd5a3 | -0.812870386 | 0.003829815 |
| Sox7 | 5.928162958 | 0.003837832 |
| Smim13 | -0.735573462 | 0.003837832 |
| Msl1 | 0.84228791 | 0.003839233 |
| Psmb8 | -1.164318032 | 0.003843234 |
| Foxs1 | 6.067482729 | 0.003845014 |
| Nfkbiz | -1.143191108 | 0.003845014 |
| Otor | 6.040287406 | 0.003850445 |
| Rapgef2 | -0.992316785 | 0.003850445 |
| Areg | -3.875781357 | 0.003858115 |
| Ubn1 | -0.708411593 | 0.00385897 |
| Cox5a | 0.578949534 | 0.003870462 |
| Rpl19 | 0.528977428 | 0.003871268 |
| Slc8a1 | -0.847131473 | 0.003871806 |
| Anp32e | 0.560419814 | 0.003877903 |
| Itpripl1 | -1.111568369 | 0.003893733 |
| Dab2 | -1.265709045 | 0.003898691 |
| Hddc3 | 0.764846008 | 0.003903286 |
| Usp12 | -1.034797685 | 0.003903286 |
| Prc1 | 1.218063397 | 0.003927319 |
| Gmfg | -1.050282884 | 0.003951864 |
| Nhs | 1.968026829 | 0.003990207 |
| Isca2 | 0.693948321 | 0.004003039 |
| Tceal1 | 1.163998794 | 0.004019842 |
| Unc5c | 1.687789198 | 0.004039864 |
| Ktn1 | 0.788935046 | 0.004039864 |
| Mocs3 | -2.258834181 | 0.004042642 |
| Dnaic1 | 4.183144082 | 0.004049685 |
| Cmc1 | 0.799132885 | 0.004049685 |
| Atp6v0a2 | -0.990967003 | 0.004049685 |
| Lbr | -0.889099445 | 0.004057946 |
| Adamts12 | 6.011770999 | 0.004063918 |
| Tcaf2 | 6.25356391 | 0.004070562 |
| Dnal1 | 1.877661629 | 0.004070562 |
| Tpt1 | 0.566700385 | 0.00408216 |
| Fam210b | 1.135701253 | 0.004088501 |
| Epas1 | 2.74831126 | 0.004089615 |
| Ptbp3 | -0.818035182 | 0.004105576 |
| Gatad2a | -0.860951023 | 0.004105576 |
| Nkx2-1 | 2.3375457 | 0.00411939 |
| Larp7 | 0.784528909 | 0.00411939 |
| Zfp740 | -0.729535231 | 0.004129489 |
| Cog3 | -1.288962897 | 0.004129489 |
| Ccdc25 | 0.79783085 | 0.004134924 |
| Vegfa | -1.078506225 | 0.004134924 |
| Tceal3 | 0.913403042 | 0.004137666 |
| Peg10 | 2.32171758 | 0.00414476 |
| Borcs8 | 0.769672887 | 0.00414476 |
| Slc12a4 | -1.643134431 | 0.00414476 |
| Pip4k2c | -0.957000068 | 0.004145761 |
| Dpm1 | -0.877706297 | 0.004146899 |
| Hvcn1 | -2.174717666 | 0.004146899 |
| Itpa | 0.742738604 | 0.004153966 |
| Spryd3 | -0.958578573 | 0.004157644 |
| Acpp | -6.770346946 | 0.004168951 |
| Plxdc1 | 4.936486149 | 0.004173197 |
| Eif1b | 0.80503647 | 0.004188514 |
| Il2rg | -2.086743645 | 0.004188514 |
| Slc30a5 | -0.947075432 | 0.004189695 |
| Smpd5 | -4.70730583 | 0.004197882 |
| Atl3 | -1.032761813 | 0.004206875 |
| Fgf9 | 2.376081225 | 0.0042088 |
| Slc39a13 | -1.253047328 | 0.004224759 |
| Dvl3 | -0.873991287 | 0.004229851 |
| Epha10 | 2.875118185 | 0.004243068 |
| Apaf1 | -1.145145733 | 0.004249086 |
| Hrsp12 | 0.924237262 | 0.004269698 |
| Emc7 | 0.753262828 | 0.00427737 |
| Mcee | 1.006435373 | 0.004285979 |
| Stra13 | 0.745409887 | 0.004286113 |
| Ppara | 6.007518607 | 0.004292938 |
| Ralb | -0.913815722 | 0.004311515 |
| Plod1 | -1.138709092 | 0.004315255 |
| Snx19 | -1.014045014 | 0.004332184 |
| Emid1 | 2.383842767 | 0.004359385 |
| Cdk5 | 0.647618078 | 0.004370407 |
| Fem1a | -0.928491792 | 0.004370407 |
| Galnt18 | 2.747520449 | 0.004388203 |
| Rhou | 1.556550972 | 0.004388203 |
| Fam213a | 0.857647497 | 0.004389115 |
| Car10 | 2.628705285 | 0.004397094 |
| Gnaz | 1.092495868 | 0.00441293 |
| Tbcd | -0.700619438 | 0.00441438 |
| Comt | 0.643747935 | 0.004452292 |
| Esm1 | 6.114510052 | 0.004458157 |
| Ubxn7 | -0.791420783 | 0.00447118 |
| S100a10 | 1.24766067 | 0.004480371 |
| Eif2b4 | 0.840466383 | 0.004480371 |
| Tsn | 0.540942308 | 0.0044924 |
| Fbxo28 | -0.876846329 | 0.004495186 |
| Tmtc2 | 1.610805124 | 0.004507817 |
| Ndufb5 | 0.535194111 | 0.004520195 |
| Galnt11 | -0.79704682 | 0.004525595 |
| Tmem132b | 1.753903881 | 0.004527351 |
| Arhgdib | -1.101922938 | 0.004528202 |
| Postn | 3.459550254 | 0.004530365 |
| Grid2 | 2.953259375 | 0.00453223 |
| Usp48 | -0.588279827 | 0.00453223 |
| Wdr41 | -0.987162686 | 0.00453223 |
| Gm10033 | -6.506689986 | 0.004550692 |
| Fam136a | 0.571532157 | 0.004555881 |
| Scyl3 | -0.89867675 | 0.004566011 |
| Rpl30 | 0.814337732 | 0.004566798 |

| | | |
|---|---|---|
| Cdhr1 | 4.977968963 | 0.139971256 |
| Exoc3 | 0.32794921 | 0.139971256 |
| Adck2 | -0.67055838 | 0.139991419 |
| Chga | 0.495514255 | 0.140000621 |
| Cacng3 | 1.060062098 | 0.140145832 |
| Pdp1 | 0.494910704 | 0.140239627 |
| Misp | -4.324717845 | 0.140344004 |
| Zfp551 | -0.940261253 | 0.14039249 |
| Zfp707 | 0.855741014 | 0.140475268 |
| Invs | -0.654154068 | 0.140485564 |
| Stag2 | 0.40630493 | 0.140515047 |
| 3110082I17Rik | -0.548774885 | 0.140522181 |
| Scel | 3.524978263 | 0.14058085 |
| Nova2 | 0.444534474 | 0.14058085 |
| Gm9493 | 0.572621802 | 0.140663969 |
| Enpp1 | -1.081552299 | 0.14086469 |
| Lgi3 | 4.913091483 | 0.140958723 |
| Scap | -0.404876048 | 0.141050906 |
| Map4k4 | 0.353084593 | 0.141087216 |
| Pcdh15 | 1.10806245 | 0.141096866 |
| Txnip | -0.532316453 | 0.141096866 |
| Tuba1a | 0.311820831 | 0.141212318 |
| Zfp61 | 0.64945388 | 0.141236155 |
| Synpr | -0.742207442 | 0.141241337 |
| Prkca | 0.588213821 | 0.141248551 |
| Cenpe | 0.721523359 | 0.141305553 |
| Tcp11l1 | 0.483594253 | 0.141392792 |
| Coro1c | -0.386117416 | 0.141485863 |
| Ppp4r1 | -0.475821983 | 0.141485863 |
| Cables2 | -0.752833739 | 0.141497968 |
| Cyc1 | 0.292670403 | 0.141548525 |
| Nagpa | -0.626161181 | 0.141548525 |
| Tymp | -1.806876676 | 0.141548525 |
| Lypd3 | -5.449504737 | 0.141548525 |
| Ube2z | -0.511905182 | 0.141745618 |
| Elfn2 | 1.363330756 | 0.14187548 |
| Hsbp1l1 | 4.555554665 | 0.141877849 |
| Tes | -0.630372511 | 0.141880783 |
| Zfp69 | -0.923078714 | 0.141880783 |
| Slfn5 | -2.069782002 | 0.141880783 |
| Avil | -2.876638339 | 0.141880783 |
| Pcdha3 | -1.534517088 | 0.141926431 |
| Tnik | -0.559203013 | 0.142119998 |
| 3110082J24Rik | 1.567385002 | 0.142291393 |
| Foxm1 | -0.606129238 | 0.142477115 |
| Ptprd | -0.510242948 | 0.142631728 |
| Tmcc3 | -0.845872762 | 0.142632657 |
| Gabbr1 | -0.413089055 | 0.142940598 |
| Sp110 | -0.93531169 | 0.142940598 |
| Cxcl2 | -1.298309689 | 0.142940598 |
| Fam167b | 2.384691593 | 0.143019452 |
| Oscar | -5.088999965 | 0.143074391 |
| Dydc2 | 2.054600007 | 0.1432448 |
| Zkscan7 | 1.329839194 | 0.1432448 |
| Zfp746 | -0.529143395 | 0.1432448 |
| Jade1 | -0.643593089 | 0.143267202 |
| Srxn1 | -0.975524847 | 0.14344262 |
| Snx5 | 0.409713314 | 0.143617717 |
| Pds5a | -0.360656441 | 0.143658786 |
| Zcchc7 | 0.503251905 | 0.143778977 |
| Taf7 | 0.428723674 | 0.143867816 |
| Tubgcp6 | -0.998468248 | 0.143867816 |
| Mob2 | 0.542235943 | 0.143888113 |
| Rapsn | 1.077632895 | 0.143954594 |
| Phkb | 0.50916958 | 0.143954594 |
| Rps6ka3 | 0.579339367 | 0.144170715 |
| Col8a2 | 3.448453597 | 0.14436153 |
| Rgs16 | -0.724618708 | 0.144367362 |
| Ddah2 | 0.336872958 | 0.144475973 |
| A930003A15Rik | -1.856702632 | 0.144475973 |
| Sidt1 | 2.17331666 | 0.14448334 |
| Gm5611 | 1.905096593 | 0.144739708 |
| Fam111a | -0.565720546 | 0.144865874 |
| Carf | -0.596423083 | 0.145005926 |
| Il17rb | -2.187906899 | 0.145005926 |
| Tpmt | 0.597402539 | 0.145162808 |
| Gtpbp4 | 0.332308873 | 0.145162808 |
| Pi16 | -1.371933559 | 0.145198873 |
| Gm5900 | 0.462927258 | 0.145229679 |
| Ttc24 | -5.44798755 | 0.145301138 |
| Serinc5 | -0.902447277 | 0.14535745 |
| Zcchc16 | 3.826751742 | 0.14536497 |
| B4galt4 | 0.725527118 | 0.14536497 |
| Phf6 | 0.357041471 | 0.145530146 |
| Rad17 | 0.628003714 | 0.145859548 |
| Susd4 | 0.800284537 | 0.145963336 |
| Srebf1 | -0.742873973 | 0.145963336 |
| Hmces | 0.439908881 | 0.146097771 |
| Nckipsd | -0.846973405 | 0.146097771 |
| Lmbr1l | -0.565834138 | 0.146182724 |
| Rps12-ps3 | 0.927477981 | 0.146301504 |
| Tceb1 | 0.327716721 | 0.146682521 |
| Rnf215 | -0.570757842 | 0.146768979 |
| Gstt2 | -0.990102431 | 0.14685261 |
| Sbno1 | -0.359986206 | 0.146856964 |
| Ccng1 | -0.515586307 | 0.146856964 |
| Nlrp4c | -5.427133593 | 0.146856964 |
| Olfr374 | -5.427133593 | 0.146856964 |
| Dpysl4 | 0.429884683 | 0.146857559 |
| Mmadhc | 0.359447554 | 0.1468995 |
| Net1 | -0.814782622 | 0.1468995 |
| Ttc39c | 0.667769441 | 0.14690166 |
| Gsg2 | -0.862080369 | 0.14690166 |
| Tceanc | 0.842743726 | 0.146920779 |
| Pan2 | -0.624045134 | 0.147131636 |
| Rtn4r | -1.071698454 | 0.147131636 |
| Grm6 | -5.108753455 | 0.147138095 |
| Gm4944 | -1.0864345 | 0.147156674 |
| Mrpl55 | 0.461491075 | 0.147206821 |
| Eml1 | -0.722628902 | 0.147333663 |
| Mtor | -0.45075249 | 0.14741215 |
| Gm5464 | -1.403739328 | 0.147706725 |
| Stam | 0.531699476 | 0.147789703 |
| Wdr7 | -0.426232628 | 0.147895692 |
| Ppm1h | -0.641936925 | 0.147895692 |
| Hgh1 | 0.95536592 | 0.147984113 |
| Abhd14b | -1.979145656 | 0.148164261 |
| Gltscr1l | -0.741930849 | 0.148170233 |
| C530008M17Rik | -0.41538522 | 0.148175266 |

| | | |
|---|---|---|
| Fam53a | 0.242969683 | 0.481650751 |
| Nrgn | -0.616152682 | 0.481707842 |
| Sdhaf3 | 0.323096302 | 0.481792839 |
| Slc26a9 | -3.318670656 | 0.481792839 |
| Cwf19l2 | 0.296519341 | 0.481848989 |
| Kcnq1 | -1.47361944 | 0.481848989 |
| Gm14418 | -0.747734548 | 0.481849825 |
| Bag3 | -0.420733972 | 0.4820855 |
| Sys1 | 0.221141766 | 0.482096877 |
| Dmbx1 | 3.87542307 | 0.482111619 |
| Arc | -1.303904839 | 0.482111619 |
| Pde6g | -2.392673533 | 0.482111619 |
| Zfpm2 | 0.514691656 | 0.482349769 |
| Ccdc115 | 0.202920305 | 0.482636409 |
| Vps50 | 0.260468456 | 0.483051583 |
| Mvb12a | -0.201866754 | 0.483207287 |
| Trp53bp1 | -0.23245468 | 0.483285029 |
| Arhgap20 | -0.37893175 | 0.483285029 |
| Rrp36 | 0.258121882 | 0.48353337 |
| Trim43c | 2.63950813 | 0.483691677 |
| Fem1c | -0.204363531 | 0.483691677 |
| Mlh1 | 0.317632191 | 0.483843266 |
| Ccdc12 | 0.177701603 | 0.483843266 |
| Cstf2t | -0.212964271 | 0.483843266 |
| Psmd10 | -0.21344968 | 0.483843266 |
| Hmx1 | 3.529062443 | 0.483860061 |
| Actr1a | 0.168703549 | 0.483898679 |
| Gm12169 | 3.586116113 | 0.484189689 |
| Tmem45b | 3.586116113 | 0.484189689 |
| Pan3 | 0.194809254 | 0.484247318 |
| Dis3l2 | -0.298034155 | 0.484423933 |
| Jph3 | -0.266418881 | 0.484441091 |
| Ush1c | -1.228799445 | 0.484449663 |
| Prdm9 | -0.443051433 | 0.484505473 |
| Sprn | 0.590162679 | 0.484699978 |
| Cox4i2 | 1.752260355 | 0.484999165 |
| Ccl2 | -0.901236697 | 0.485110622 |
| Ppm1e | 0.252605293 | 0.485438348 |
| Htra3 | -0.741382183 | 0.485438348 |
| Gli1 | 1.049776821 | 0.485724947 |
| Kif20b | 0.266847064 | 0.485736007 |
| AY358078 | 1.883636316 | 0.485744975 |
| Gm6665 | 2.597071359 | 0.485788277 |
| Dlg4 | -0.377399662 | 0.48589848 |
| Prg4 | -0.84710282 | 0.486204699 |
| Sh3tc2 | -2.660752611 | 0.486204699 |
| Smim14 | 0.189248514 | 0.48626498 |
| Lima1 | -0.234727452 | 0.486314554 |
| Syne1 | 0.555367242 | 0.486580313 |
| Maats1 | -1.384385895 | 0.486596691 |
| Tex13 | 3.5259157 | 0.487284349 |
| 9830147E19Rik | 1.031454565 | 0.487284349 |
| C730034F03Rik | 0.660527697 | 0.487284349 |
| Ctnna1 | 0.233515092 | 0.487284349 |
| Usp29 | 0.221111592 | 0.487284349 |
| Lrrc71 | -1.682394891 | 0.487284349 |
| Got1 | 0.199113643 | 0.487684236 |
| Cd226 | -1.730663709 | 0.487684236 |
| Necab1 | 0.416387077 | 0.487858747 |
| Ing3 | 0.216219267 | 0.488081464 |
| Usp24 | -0.240729165 | 0.488088162 |
| Tssc1 | -0.246485661 | 0.488088162 |
| Trove2 | -0.248204761 | 0.488088162 |
| Wsb1 | -0.235817171 | 0.488156349 |
| C2cd4c | 2.586312664 | 0.488196422 |
| Prr36 | -0.234465303 | 0.488196422 |
| Gnl3l | -0.1755136 | 0.488428261 |
| Thap7 | -0.240479135 | 0.488486257 |
| Atm | -0.388046492 | 0.488525492 |
| Strn4 | -0.213884241 | 0.488535215 |
| Nrbf2 | 0.29690555 | 0.488653584 |
| Cpeb4 | 0.24236702 | 0.488834224 |
| Zmat5 | 0.339982847 | 0.488972722 |
| Scyl2 | -0.246754732 | 0.489193072 |
| Asxl1 | 0.249777154 | 0.489372953 |
| Trim30b | -1.035962673 | 0.489372953 |
| Kars | -0.186382097 | 0.489420989 |
| Fubp1 | 0.148915302 | 0.489470413 |
| Carnmt1 | 0.380536687 | 0.489623 |
| R3hdm2 | 0.23811637 | 0.489679369 |
| Crebbp | 0.172168623 | 0.489679369 |
| Gfpt1 | 0.234892709 | 0.489722089 |
| Tmem214 | -0.271291277 | 0.489787368 |
| Ptrh2 | 0.223597486 | 0.489818135 |
| Birc6 | -0.164382989 | 0.489818951 |
| Fam3c | 0.279149669 | 0.489912095 |
| Plod2 | 0.418650638 | 0.489916475 |
| Dnajc18 | 0.266329863 | 0.489916475 |
| Kat6b | -0.227068486 | 0.489916475 |
| Trappc9 | -0.23199977 | 0.489916475 |
| Slc4a7 | -0.290405796 | 0.489916475 |
| Zfp41 | -0.30070061 | 0.489916475 |
| Mthfs | -0.370591018 | 0.489916475 |
| Sh3gl1 | -0.193649456 | 0.489916847 |
| Nav1 | 0.20678115 | 0.489936307 |
| Cse1l | -0.181456788 | 0.489936307 |
| Mitd1 | -0.420729643 | 0.48999742 |
| Timm23 | 0.249830101 | 0.490028227 |
| Ankhd1 | -0.232012937 | 0.490055249 |
| Dll1 | 0.341960827 | 0.490058962 |
| Klhl6 | -0.441361409 | 0.490058962 |
| Snrpf | -1.837784265 | 0.490252881 |
| Gnb1l | 0.433071741 | 0.490404652 |
| Egf | -1.456769233 | 0.490404652 |
| Zfp874a | 0.466253839 | 0.4905863 |
| Trim27 | -0.227416419 | 0.490615629 |
| Gdf3 | 3.555724494 | 0.490652004 |
| Ccl28 | 3.505339689 | 0.490943479 |
| Syt5 | 0.223886021 | 0.490943479 |
| Gm11007 | 1.186152666 | 0.491139764 |
| Rfxank | 0.467899601 | 0.491139764 |
| Zfp260 | 0.206882932 | 0.491139764 |
| Dchs1 | -0.242387705 | 0.491139764 |
| Kif2c | -0.288920531 | 0.491139764 |
| Gm14305 | 0.772892211 | 0.491311122 |
| Scrt1 | 0.56646341 | 0.491335119 |
| Hsd17b11 | -0.62801306 | 0.491375477 |
| Cgnl1 | 0.800087992 | 0.491700993 |
| Disc1 | -0.650238937 | 0.491700993 |

| | | |
|---|---|---|
| Noc4l | 0.07804161 | 0.87385027 |
| Ypel2 | 0.101573947 | 0.874136541 |
| Noc3l | 0.082297445 | 0.87441955 |
| Zfp119b | -0.112279539 | 0.874720474 |
| Gm8225 | 0.796865442 | 0.874885706 |
| Pou5f2 | 0.286090412 | 0.875136493 |
| Ddc | 0.264887394 | 0.875136493 |
| Yjefn3 | 0.207333835 | 0.875136493 |
| Fam57a | 0.109505823 | 0.875136493 |
| Slc38a4 | -0.252335109 | 0.875136493 |
| Slc4a11 | 0.302830882 | 0.875548136 |
| Ep400 | 0.049220256 | 0.875654206 |
| Mroh7 | 0.629173574 | 0.875749766 |
| G6pc3 | -0.090371698 | 0.875749766 |
| Adam11 | -0.125035987 | 0.875749766 |
| Bend3 | -0.109161229 | 0.875924984 |
| Ntf3 | 0.300094828 | 0.875932274 |
| Trp73 | -0.169835207 | 0.875932274 |
| Abhd15 | 0.227995251 | 0.875972937 |
| Gm3839 | -0.46433746 | 0.875972937 |
| Tmprss4 | -0.662084156 | 0.875972937 |
| Zfp600 | 0.518180766 | 0.87607505 |
| Cnga3 | 0.613390107 | 0.876234531 |
| Vmn2r29 | 0.464067787 | 0.876234531 |
| Amz1 | 0.141946572 | 0.876234531 |
| Brf1 | -0.072306971 | 0.876234531 |
| Rdh14 | -0.089415303 | 0.876234531 |
| Stil | -0.107174445 | 0.876234531 |
| Trim68 | -0.140946415 | 0.876234531 |
| Cdc25a | -0.085224974 | 0.876242442 |
| Nt5dc3 | 0.131652298 | 0.876496384 |
| Grhpr | 0.08526195 | 0.876496384 |
| Taf7l | -0.49343472 | 0.876496384 |
| Dctn4 | 0.039438559 | 0.876835301 |
| Snx6 | 0.043381829 | 0.877072017 |
| Zfp607 | -0.36695547 | 0.877426397 |
| Nxnl2 | 0.388560119 | 0.877761723 |
| Stk38l | 0.092125985 | 0.877761723 |
| Trim9 | 0.072988452 | 0.877761723 |
| Acvr2a | 0.066455108 | 0.877821345 |
| Idi1 | -0.072496612 | 0.877821345 |
| Rb1cc1 | 0.047546367 | 0.877831114 |
| Rdh13 | 0.055710883 | 0.877973875 |
| Ppcdc | -0.090435903 | 0.877973875 |
| Lrrc3 | 0.146514124 | 0.878131912 |
| Pomk | 0.086599331 | 0.878131912 |
| Pole2 | 0.139221018 | 0.878212623 |
| Ndrg4 | 0.055749712 | 0.878241308 |
| Abca9 | -0.106295508 | 0.878241308 |
| Gm14296 | -0.26611188 | 0.878263794 |
| Ksr2 | 0.114365558 | 0.878733725 |
| Amd2 | 0.218985267 | 0.878778787 |
| Rimklb | -0.051525561 | 0.878778787 |
| Tmem136 | -0.10637002 | 0.878778787 |
| Vax1 | -0.241222719 | 0.878778787 |
| Pcdhga2 | -0.354583384 | 0.878778787 |
| Gm12394 | 0.275317616 | 0.878846983 |
| Akr1c19 | 0.176146535 | 0.878846983 |
| Ankle1 | 0.170499623 | 0.878846983 |
| Pop1 | 0.119217729 | 0.878846983 |
| Xrcc4 | 0.082160517 | 0.878846983 |
| Mast2 | 0.066874921 | 0.878846983 |
| Rnf38 | -0.055546393 | 0.878846983 |
| Tmem175 | -0.063990281 | 0.878873675 |
| Ssrp1 | 0.044973597 | 0.879076185 |
| Prnp | -0.071721915 | 0.879552662 |
| Ctnnd2 | 0.063033294 | 0.879663379 |
| Rfwd2 | 0.070731113 | 0.879672332 |
| Dydc1 | 0.747148226 | 0.879723268 |
| Evx2 | 0.72785585 | 0.879723268 |
| Pcdh7 | 0.077095275 | 0.879723268 |
| Asb1 | 0.076073961 | 0.879723268 |
| Bag5 | -0.049022157 | 0.879723268 |
| Bcap29 | -0.060253086 | 0.879723268 |
| Nck1 | -0.082892778 | 0.879723268 |
| Tk1 | -0.086935491 | 0.879723268 |
| Zfp3 | -0.089934819 | 0.879723268 |
| Ptprn | -0.144474527 | 0.879723268 |
| Gm996 | -0.184349174 | 0.879723268 |
| Mmrn2 | 0.279286549 | 0.879783701 |
| 5730559C18Ri | 0.115560769 | 0.879783701 |
| Pogk | 0.054087867 | 0.879783701 |
| Tmem150b | -0.412932843 | 0.879783701 |
| Tmem28 | -0.298261847 | 0.879822283 |
| Reep4 | -0.099119317 | 0.880095086 |
| Sh2d1b2 | -0.542559066 | 0.880166896 |
| Nkapl | -0.190942833 | 0.880166921 |
| Vsnl1 | 0.191157685 | 0.880176637 |
| Adamts17 | -0.117174246 | 0.880176637 |
| R3hdml | -0.572844966 | 0.880176637 |
| Fancb | 0.15016175 | 0.880187417 |
| Ccsap | -0.128905999 | 0.880187417 |
| Acsf3 | 0.111436727 | 0.880250728 |
| Mtus1 | -0.049377772 | 0.880317143 |
| Foxq1 | 0.423119129 | 0.880530901 |
| Coro2b | 0.070000196 | 0.880530901 |
| Mthfsd | 0.067468453 | 0.880530901 |
| Nbas | -0.076037124 | 0.880530901 |
| Cps1 | 0.447225358 | 0.881043885 |
| Gabrd | 0.268293303 | 0.881043885 |
| Pex26 | 0.106960572 | 0.881043885 |
| Pigc | 0.069287148 | 0.881043885 |
| Melk | -0.082852668 | 0.881043885 |
| Gfod1 | -0.09917471 | 0.881043885 |
| Gm19684 | 0.754249199 | 0.881186383 |
| Zfp282 | 0.06999756 | 0.881186383 |
| Rab2b | 0.049190291 | 0.881186383 |
| Rbbp6 | -0.048676455 | 0.881186383 |
| Fuom | -0.10293999 | 0.881186383 |
| Gm766 | -0.364606335 | 0.881544241 |
| Cand1 | -0.049787533 | 0.882083739 |
| Gpr137 | -0.08182107 | 0.882083739 |
| Cnnm3 | -0.082564832 | 0.882152702 |
| Phxr4 | 0.109690999 | 0.882190498 |
| Gm14325 | 0.086606152 | 0.882190498 |
| Fam69a | 0.079298055 | 0.882190498 |
| Mcoln1 | -0.063142883 | 0.882190498 |
| Slc22a12 | -0.467204548 | 0.882190498 |
| Gm3591 | 0.227079067 | 0.88243733 |

| | | |
|---|---|---|
| Lamtor5 | 0.523659259 | 0.004567419 |
| Tusc2 | 0.809545808 | 0.004576265 |
| Olr1 | -2.979783194 | 0.004576265 |
| Frmd6 | 2.419883202 | 0.004577902 |
| Sipa1l3 | -1.158327891 | 0.004580718 |
| Kcnip4 | 2.359686797 | 0.00459076 |
| Pcdhb7 | 1.737457486 | 0.004607775 |
| Efcab2 | 1.330663561 | 0.00462383 |
| Tchh | 4.016913735 | 0.004644974 |
| Cblb | -0.76327994 | 0.004651876 |
| Fosb | 0.740112084 | 0.00466112 |
| Fsd1l | 0.817773825 | 0.004661714 |
| Brap | -0.828127281 | 0.004668625 |
| Irak4 | -1.15092015 | 0.004668625 |
| Mmp9 | -4.755262531 | 0.004669129 |
| Spock3 | 2.244608749 | 0.004684887 |
| Aspn | 5.913541484 | 0.004690316 |
| Fam129b | -1.319971357 | 0.004695331 |
| Fem1b | -0.686216873 | 0.004715129 |
| Krba1 | -0.856158796 | 0.004726242 |
| Wdr83os | 0.966857926 | 0.00472675 |
| Sf3b3 | -0.643731166 | 0.00472675 |
| AA474408 | -2.149189468 | 0.00472675 |
| Egflam | 4.730941208 | 0.004762093 |
| Cacna2d2 | 1.661804275 | 0.004762093 |
| 1110051M20Rik | 0.634699305 | 0.004762093 |
| Lias | 0.646681486 | 0.004769734 |
| Ube2b | 0.563816464 | 0.004770837 |
| Prkg2 | 5.902122279 | 0.004777087 |
| Ndel1 | -0.595532844 | 0.004777087 |
| Mrps15 | 0.627058716 | 0.004780475 |
| Zfp641 | 2.679718823 | 0.004782934 |
| Kcnj2 | 6.457162933 | 0.004829961 |
| Mtpap | 0.903279353 | 0.004850827 |
| Igbp1 | 0.892225332 | 0.004859844 |
| Denr | 0.581042305 | 0.004867982 |
| Wdfy4 | -1.573261842 | 0.00487827 |
| Col5a2 | 1.389696573 | 0.00488288 |
| Gpr34 | 1.599020395 | 0.004886836 |
| Senp5 | -0.900212568 | 0.004889064 |
| Vwa5b1 | 4.724448618 | 0.004899374 |
| Rfx2 | 3.073707129 | 0.004899374 |
| Ceacam1 | -1.774545098 | 0.004904252 |
| Cbr1 | 0.64725805 | 0.004925464 |
| Zfp384 | -0.926781195 | 0.004925464 |
| Prkar2a | -0.840172898 | 0.004933843 |
| Gpr62 | -6.514826625 | 0.004948006 |
| Fam216a | 0.837712089 | 0.004962595 |
| Ppard | -1.740696831 | 0.004966212 |
| Ckap4 | -0.891231848 | 0.004967382 |
| Ndufa8 | 0.645202359 | 0.004971802 |
| Rilpl2 | -0.770314691 | 0.004980384 |
| Sall1 | 1.173618106 | 0.004983583 |
| Diablo | 0.788972237 | 0.004983583 |
| Mfsd5 | -0.745233075 | 0.004983583 |
| E2f4 | -0.79458378 | 0.004983583 |
| Bcl2a1a | -1.355026432 | 0.004983756 |
| Pvr | -1.69630505 | 0.004993407 |
| Gm21188 | -2.930424569 | 0.004993407 |
| 1700109H08Rik | -1.538249669 | 0.005026975 |
| Stk40 | -1.143178512 | 0.005030376 |
| Txndc15 | 0.771925885 | 0.005038204 |
| Cops8 | 0.521066935 | 0.005038204 |
| Car14 | 2.448812856 | 0.00504074 |
| Nupl2 | 1.467856942 | 0.005049988 |
| Morn2 | 1.283381687 | 0.005049988 |
| Gabra2 | 1.146775263 | 0.005049988 |
| Phax | 0.636561203 | 0.005058067 |
| K230010J24Rik | 6.014261782 | 0.005065434 |
| Lxn | 1.108577722 | 0.00507367 |
| Rps19bp1 | 0.800908956 | 0.00507367 |
| Traf2 | -1.106398087 | 0.005074245 |
| Ppp3cb | 0.703430726 | 0.005083228 |
| Plxna4 | -0.781357002 | 0.005099308 |
| Trp53i13 | 1.335785992 | 0.005103874 |
| Ambp | -7.037393045 | 0.005117688 |
| Tmem168 | -0.885816254 | 0.005134205 |
| Tmc6 | -1.629356683 | 0.005138479 |
| Hscb | 1.097634078 | 0.005146167 |
| Whsc1l1 | -0.530115029 | 0.005148551 |
| Kremen1 | -1.444285774 | 0.005156611 |
| Socs2 | 0.991963627 | 0.005164975 |
| Ntn1 | 2.791257295 | 0.005187808 |
| Cdc42se1 | -0.671814477 | 0.005187808 |
| Mbd2 | -1.753666578 | 0.005187808 |
| Optc | 2.839296022 | 0.005193033 |
| Unk | -0.812851755 | 0.005195924 |
| Mdfic | -1.322040233 | 0.005227011 |
| Tnfrsf26 | -3.411240069 | 0.005236161 |
| 9130011E15Rik | -1.267351105 | 0.005248967 |
| Defb11 | 6.029453425 | 0.00525834 |
| Kcnh7 | 1.044834319 | 0.00525834 |
| Spc24 | 0.892414927 | 0.005259108 |
| Cd302 | -1.252009427 | 0.005275763 |
| Tgm1 | -2.732510348 | 0.005275763 |
| Cdc14b | -1.75367637 | 0.00528834 |
| Slc9a5 | -1.547143225 | 0.005358324 |
| Hhex | -1.231614992 | 0.00536941 |
| Zfp850 | 3.033321993 | 0.005370505 |
| Btnl9 | -6.567084354 | 0.005414275 |
| Drc7 | 4.739760847 | 0.005417279 |
| Ccdc88b | -1.520553456 | 0.005422808 |
| H2-T24 | -1.715688339 | 0.005422808 |
| Lonrf2 | 0.921906774 | 0.00543586 |
| Timm22 | 0.636881244 | 0.005455703 |
| Tmod2 | 0.573388032 | 0.005458012 |
| Galnt1 | -0.794754473 | 0.005475131 |
| Phrf1 | -0.751572183 | 0.005478498 |
| Cog2 | -0.755443237 | 0.005478498 |
| Iqsec1 | -0.87789274 | 0.005478498 |
| Fam109b | -4.556062244 | 0.005493699 |
| Slc3a2 | -0.632306658 | 0.005497404 |
| Smdt1 | 0.606426129 | 0.005499148 |
| Rhog | -0.899846681 | 0.005502508 |
| Gm561 | 0.897107268 | 0.005509509 |
| Gabrb1 | 1.604028895 | 0.00552348 |
| Rsf1 | 0.586056003 | 0.005524492 |
| Mrps17 | 0.663198708 | 0.005529406 |
| Atp8b4 | -1.953892407 | 0.005529769 |

| | | |
|---|---|---|
| Ppp1r35 | 0.510817421 | 0.148388294 |
| Ppif | 0.411069681 | 0.148570678 |
| Ncoa2 | -0.500132043 | 0.148608488 |
| Bora | 0.671815007 | 0.148693996 |
| Cxx1a | 0.809934903 | 0.148704787 |
| Thyn1 | 0.700020811 | 0.148877089 |
| Rnf216 | -0.407793285 | 0.148958223 |
| Ap2s1 | 0.340175595 | 0.148963538 |
| Glod4 | 0.359760407 | 0.149206252 |
| Rab30 | 0.594815233 | 0.149376235 |
| Zfp846 | 0.484170087 | 0.149486544 |
| Spag17 | 3.981060065 | 0.149504153 |
| Sparcl1 | 0.708554986 | 0.149504153 |
| Nmt2 | -0.346064807 | 0.149532696 |
| Plekha1 | -0.451847168 | 0.149532696 |
| Slc34a3 | 5.00378339 | 0.149755278 |
| Sntb1 | 4.692604273 | 0.149755278 |
| Sptssb | 4.692604273 | 0.149755278 |
| Chsy3 | 1.860561517 | 0.149755278 |
| Ppwd1 | 0.963185669 | 0.149755278 |
| Celsr3 | -0.499642828 | 0.149865667 |
| Hgsnat | -0.602759313 | 0.149875527 |
| Nbn | 0.46750878 | 0.149921524 |
| Nfkbib | -0.437072947 | 0.149921524 |
| Trpc3 | -0.975003583 | 0.149958051 |
| Gnai1 | 0.491962279 | 0.150135211 |
| Gm8113 | 2.767025338 | 0.150274273 |
| Il1rapl2 | 4.967831127 | 0.15051587 |
| Gm14443 | -4.665582275 | 0.150796798 |
| Bag2 | 0.66392514 | 0.1510943 |
| Ttc23 | 1.066315242 | 0.151151462 |
| Pnpla7 | -0.629166567 | 0.151188736 |
| Rab19 | 2.229281011 | 0.151325935 |
| Eif1a | -0.386633903 | 0.151708513 |
| Tap2 | -0.890880425 | 0.151716621 |
| Pde1a | 1.525020466 | 0.151784424 |
| Prpf38a | 0.332350822 | 0.151885753 |
| Pcdhga7 | -0.894670949 | 0.151885753 |
| Ring1 | -0.823997013 | 0.152001602 |
| Pcgf3 | 0.478519263 | 0.152005203 |
| Pick1 | 0.563516621 | 0.152133978 |
| Zfp513 | -0.866471695 | 0.152198809 |
| Naip6 | -1.435896525 | 0.152590151 |
| Trim71 | 4.464424576 | 0.152599377 |
| Wnt5b | 1.394132634 | 0.152599377 |
| Brwd3 | 0.947674872 | 0.152599377 |
| Dnph1 | 0.637358077 | 0.152599377 |
| Nudc | 0.308945225 | 0.152619091 |
| Tmco6 | -0.594555602 | 0.152771111 |
| Naa50 | -0.314981027 | 0.15296367 |
| Rnf141 | -0.539543786 | 0.15303147 |
| Tuba4a | -0.915525255 | 0.153068711 |
| Pla2g4a | -1.062708118 | 0.153068711 |
| Cdhr4 | -5.078933817 | 0.153068711 |
| Gabrb3 | -0.503203714 | 0.153226498 |
| Zpld1 | -3.922452287 | 0.153269312 |
| Grm7 | 1.562991698 | 0.153287175 |
| Spink10 | 1.394405909 | 0.153287175 |
| Ormdl2 | -0.536463456 | 0.153287175 |
| Slc25a45 | -0.732115635 | 0.153393803 |
| Smtn | 0.98989353 | 0.153769155 |
| Msi1 | 0.509973911 | 0.153842865 |
| Alms1 | -0.522163084 | 0.153859075 |
| Ugp2 | -0.467994033 | 0.153936918 |
| Tfe3 | -0.608121539 | 0.153964245 |
| Ccdc18 | 0.793128582 | 0.15399905 |
| Arfgap3 | -0.580567631 | 0.154069208 |
| Mospd1 | 0.485260187 | 0.154084349 |
| Tcea3 | 4.568916419 | 0.154090809 |
| AF251705 | -0.634475302 | 0.154090809 |
| Dcaf10 | -0.57695688 | 0.154118516 |
| Xkr7 | -1.386172002 | 0.154210449 |
| Atox1 | 0.33531052 | 0.154260118 |
| Dnaaf3 | 2.286171505 | 0.154322506 |
| Sez6 | -0.665594204 | 0.154400867 |
| Lhx8 | 1.793851847 | 0.154486946 |
| C330007P06Rik | 0.417547432 | 0.154543929 |
| Cntrob | -0.578041186 | 0.154667549 |
| Oaz3 | 1.811910432 | 0.154719497 |
| Dkk2 | 1.635733064 | 0.154734262 |
| Dlx6 | 0.793758721 | 0.154805261 |
| Usp38 | -0.505301348 | 0.154818137 |
| Gstm6 | 1.984040965 | 0.154824687 |
| Wnt4 | 1.684684316 | 0.154824687 |
| Mcm8 | -0.688850858 | 0.154861085 |
| Kctd3 | -0.474969172 | 0.1548813 |
| L3mbtl1 | -2.138338207 | 0.1548813 |
| Adamts14 | 3.340282072 | 0.154924071 |
| Mlf2 | 0.289464485 | 0.154966358 |
| Hif1a | -0.367834723 | 0.154966358 |
| Edc3 | -0.430279186 | 0.15500055 |
| Srsf9 | -0.419653879 | 0.155280945 |
| Pafah1b1 | -0.36660728 | 0.155302425 |
| BC049352 | -1.223269964 | 0.155452514 |
| Mb21d1 | -1.06809527 | 0.155540239 |
| Atp2a3 | -1.933532124 | 0.155566104 |
| Gm10263 | 0.479044953 | 0.15570191 |
| Trmo | 1.202879377 | 0.155762234 |
| Cpne5 | 0.862340616 | 0.155779327 |
| Dhx37 | 0.517655729 | 0.155832746 |
| Scrib | 0.689256224 | 0.155927708 |
| Oxsm | -0.844925006 | 0.155929018 |
| Npbwr1 | 3.990699769 | 0.155933793 |
| Bst2 | -1.487604105 | 0.155933793 |
| Twf1 | 0.379426346 | 0.156000998 |
| Rab33a | 0.529713862 | 0.156038797 |
| A3galt2 | -2.732007782 | 0.156073949 |
| Crtam | -2.323099246 | 0.156085762 |
| Nfatc1 | -0.740721685 | 0.156130739 |
| 2410131K14Rik | -0.520630903 | 0.156614364 |
| Gpr139 | -1.334866513 | 0.156614364 |
| Zfp663 | 1.656413413 | 0.156688562 |
| Rbms2 | 0.642242835 | 0.156688562 |
| Ern1 | -0.902679413 | 0.156688562 |
| Hspa1a | -1.837158798 | 0.156717454 |
| Myct1 | 4.539429279 | 0.156828439 |
| Zscan29 | -0.702149634 | 0.156896204 |
| Fgd2 | -0.77191546 | 0.156896204 |
| Med25 | 0.33600138 | 0.157371295 |

| | | |
|---|---|---|
| St3gal4 | -0.387892753 | 0.491748239 |
| Gpr162 | 0.488117565 | 0.491889959 |
| Accs | 0.325557949 | 0.491889959 |
| Rrnad1 | -0.350849243 | 0.491889959 |
| Taf1a | 0.396921226 | 0.49192607 |
| Agrp | -1.525515937 | 0.49192607 |
| Ccl7 | -0.880749781 | 0.492032158 |
| Ccdc91 | 0.266274797 | 0.49215437 |
| C1s1 | 3.255324026 | 0.492186169 |
| Glis2 | 0.333904015 | 0.492484439 |
| Shroom3 | 0.449826593 | 0.492504416 |
| Ercc1 | 0.255348059 | 0.492522236 |
| Arhgap18 | 0.343290031 | 0.492596633 |
| B3galt1 | -0.361954215 | 0.492720325 |
| Rcbtb2 | 0.179967377 | 0.492757423 |
| Snx27 | 0.209471285 | 0.492867376 |
| Las1l | -0.18846326 | 0.492958519 |
| Dnmbp | -0.511229679 | 0.493307661 |
| Sergef | 0.283875457 | 0.493351913 |
| Gm128 | -2.439875968 | 0.493362774 |
| Arsg | 0.45123244 | 0.4933917 |
| Plppr2 | 0.243260397 | 0.4933917 |
| Mcmdc2 | 0.513114135 | 0.493412331 |
| Fastkd3 | -0.506236597 | 0.493490147 |
| Klc3 | -1.017363901 | 0.493490147 |
| Rasl11b | -0.321394686 | 0.493615772 |
| Tmem42 | 0.337840203 | 0.493773905 |
| Gm2897 | 0.631124725 | 0.493961319 |
| Abcd4 | -0.351692821 | 0.493961319 |
| Nsun5 | 0.35037785 | 0.494017293 |
| Gm3468 | 3.200589976 | 0.494066804 |
| Wdr26 | -0.196484967 | 0.494066804 |
| Ehhadh | -0.886158754 | 0.494308593 |
| Tmem180 | -0.490755665 | 0.494489478 |
| Tmie | 2.13428995 | 0.494726664 |
| Trim62 | 0.599524019 | 0.494726664 |
| Shisa7 | 0.486431397 | 0.494726664 |
| Itgax | -1.460346073 | 0.494726664 |
| Ly75 | -1.124452816 | 0.494843662 |
| Dnajb14 | -0.309127961 | 0.494929509 |
| Rpl23a | 0.44745269 | 0.495034362 |
| Asphd1 | 0.54938986 | 0.495166107 |
| Fanca | -0.487463911 | 0.495251951 |
| Nipa1 | -0.349512004 | 0.495334144 |
| Adad2 | 3.813047964 | 0.495732545 |
| Myo7b | 3.813047964 | 0.495732545 |
| Cpq | 0.62647471 | 0.495732545 |
| Zfp948 | -0.31071338 | 0.495764273 |
| Ankrd13b | 0.270448894 | 0.496032987 |
| Ruvbl1 | 0.205053408 | 0.496032987 |
| Spdl1 | 0.401842331 | 0.496099567 |
| Gm5108 | 3.506724496 | 0.496589537 |
| Mmp20 | 3.506724496 | 0.496589537 |
| Sptbn5 | 2.127192923 | 0.496589537 |
| Isy1 | 0.316985749 | 0.496638343 |
| Leng1 | 0.398773666 | 0.49680104 |
| Abt1 | -0.237752305 | 0.497157579 |
| Abhd13 | -0.26600119 | 0.497252519 |
| Sfpq | 0.159048658 | 0.497252591 |
| Hist2h2ac | 0.6100706 | 0.4973339 |
| Mapk3 | 0.237434644 | 0.497507627 |
| Scfd2 | -0.282833734 | 0.49750989 |
| E130012A19Rik | -0.359462834 | 0.49750989 |
| Mknk1 | -0.25932671 | 0.497529066 |
| Adtrp | 3.797489348 | 0.497878957 |
| Npy1r | 0.624931579 | 0.498214473 |
| Klhl23 | 0.237441674 | 0.498229879 |
| Zfp609 | -0.255160173 | 0.498229879 |
| Dxo | 0.250993512 | 0.4983952 |
| 1700030K09Rik | 0.284584401 | 0.49850837 |
| Ndc80 | 0.425056406 | 0.498550666 |
| Arid5b | 0.443284742 | 0.498982987 |
| Dbp | -0.729769838 | 0.498982987 |
| Lrguk | -1.103146977 | 0.498982987 |
| Ascc1 | 0.22094783 | 0.499089553 |
| Cpb1 | 3.787653446 | 0.499208983 |
| Krtap16-1 | 3.787653446 | 0.499208983 |
| Xcr1 | 1.028640906 | 0.499224186 |
| Wdr66 | -0.727422519 | 0.499227623 |
| A230065H16Rik | 3.186582849 | 0.499229504 |
| Ckmt1 | 0.390589242 | 0.499229504 |
| Adrb3 | 3.253662183 | 0.499414035 |
| Elf5 | 3.253662183 | 0.499414035 |
| Exoc3l2 | 3.253662183 | 0.499414035 |
| Zc3h11a | 0.52646227 | 0.499414035 |
| Adcyap1r1 | -0.297360539 | 0.499414035 |
| Gvin1 | -1.319853061 | 0.499414035 |
| Armc8 | -0.234682265 | 0.499492086 |
| Mpi | 0.229141228 | 0.499744405 |
| Noa1 | -0.29781708 | 0.499744405 |
| Tnnt3 | -1.608453534 | 0.500548837 |
| Mzt1 | -0.185321428 | 0.500641866 |
| AW549877 | 0.2567785 | 0.500646777 |
| Manbal | -0.252719935 | 0.500646777 |
| Nars2 | -0.367310829 | 0.500646777 |
| Nhsl2 | -0.398629468 | 0.500646777 |
| Fos | 0.239105747 | 0.500862979 |
| Vsig8 | 3.463314018 | 0.500948722 |
| Rapgef4 | 0.617725602 | 0.501104626 |
| Lmna | -0.239253422 | 0.501271969 |
| Naa60 | 0.228541819 | 0.501274056 |
| Ccdc170 | 3.471755231 | 0.5013213 |
| 3300002I08Rik | 0.61077366 | 0.501590976 |
| Rabl6 | -0.188017359 | 0.501590976 |
| Nol4 | -0.210766464 | 0.501590976 |
| Abhd10 | -0.288200132 | 0.501590976 |
| Ncam2 | 0.472775833 | 0.501809062 |
| Strc | -3.236940061 | 0.50202499 |
| Anapc4 | -0.173515052 | 0.502238153 |
| Vsig10l | -0.58901333 | 0.502494361 |
| Nfix | 0.296354556 | 0.502584739 |
| Gcn1l1 | -0.209623538 | 0.502887563 |
| Stard6 | 1.963923386 | 0.503052788 |
| Gcdh | -0.277284905 | 0.503052788 |
| Tgtp2 | -1.66608279 | 0.503052788 |
| Itpr3 | 0.938845261 | 0.50314798 |
| Ggct | -0.500207063 | 0.503156972 |
| Slc35b2 | -0.186102019 | 0.503224109 |
| Sorcs2 | 0.496829836 | 0.503400975 |

| | | |
|---|---|---|
| Inpp5e | -0.079062055 | 0.882713572 |
| 2410089E03Ril | -0.059670986 | 0.882742222 |
| Ifit1bl2 | -0.346836582 | 0.882742222 |
| Mroh6 | -0.565045842 | 0.882930542 |
| Gdpgp1 | -0.133592864 | 0.883301662 |
| Gm5580 | -0.490486819 | 0.883301662 |
| Mustn1 | -0.429383189 | 0.883449536 |
| Spred3 | 0.124214579 | 0.883571686 |
| Ccl17 | 0.53442533 | 0.883670479 |
| Ggact | 0.132741195 | 0.883670479 |
| Tmem86b | -0.175542516 | 0.883670479 |
| Phpt1 | 0.302434892 | 0.883970659 |
| Arfgap2 | -0.06295218 | 0.884040845 |
| Cabp1 | -0.126746454 | 0.884116448 |
| Dffb | 0.15723303 | 0.884134625 |
| Brf2 | 0.114933981 | 0.884134625 |
| Rorb | -0.075562781 | 0.884134625 |
| Nudt7 | -0.212674593 | 0.884134625 |
| Dusp13 | -0.611668608 | 0.884245558 |
| Mettl3 | 0.071823903 | 0.884358661 |
| Prdx6 | -0.055030834 | 0.884481874 |
| Ppfia2 | 0.069803846 | 0.884967198 |
| Rbm34 | 0.061578335 | 0.884967198 |
| Wrnip1 | -0.062785471 | 0.884967198 |
| Srcin1 | -0.076219411 | 0.884967198 |
| Zfp780b | -0.081606848 | 0.884967198 |
| Mfsd3 | -0.138560926 | 0.884967198 |
| Cda | 0.538195089 | 0.88521662 |
| Nub1 | 0.049916006 | 0.885750636 |
| E030018B13Ri | 0.350689865 | 0.885797416 |
| Golga3 | -0.056439203 | 0.886061172 |
| Arnt | -0.070046434 | 0.886061172 |
| Tstd1 | -0.257662007 | 0.886281698 |
| Prkacb | 0.046677784 | 0.8862893 |
| Nr3c2 | -0.20319673 | 0.8862893 |
| Stip1 | -0.040539627 | 0.886305112 |
| Plac8 | -0.257193557 | 0.886312973 |
| Zfp951 | -0.178402997 | 0.886322993 |
| Cnot7 | 0.045504832 | 0.88647919 |
| Blmh | -0.040577306 | 0.88648025 |
| Ctcf | 0.042358834 | 0.886680054 |
| Fcho2 | 0.06317018 | 0.886719025 |
| Tsc22d3 | 0.050037474 | 0.886719025 |
| Dhrs11 | -0.134825467 | 0.886719025 |
| Cfap61 | 0.368509278 | 0.886853189 |
| Psmd11 | 0.037164597 | 0.886853189 |
| Cog4 | -0.055385967 | 0.886960991 |
| Med11 | -0.077892664 | 0.886961237 |
| Nptx2 | 0.229874409 | 0.887057932 |
| Suox | 0.153739999 | 0.887057932 |
| Wdr12 | 0.045982588 | 0.887057932 |
| Bcl11b | -0.071831212 | 0.887057932 |
| Alyref2 | -0.077411516 | 0.887074461 |
| Rwdd4a | 0.056083468 | 0.887245532 |
| Bcl11a | 0.06614015 | 0.887311262 |
| Hccs | 0.074266444 | 0.887332638 |
| Dsp | -0.647277794 | 0.887655956 |
| Zfp799 | 0.067941988 | 0.887970474 |
| Exoc3l | -0.127498542 | 0.887970474 |
| Folr2 | 0.200546081 | 0.888095517 |
| Kdm5c | 0.0449271 | 0.888168106 |
| Nepro | -0.146780736 | 0.888168106 |
| Pja2 | 0.048814322 | 0.88851854 |
| Uba52 | -0.127598903 | 0.888998055 |
| Syt2 | 0.20127514 | 0.889301902 |
| Omg | -0.235761609 | 0.889301902 |
| Adprm | 0.070661497 | 0.88944376 |
| Dsel | -0.081084005 | 0.88944376 |
| Osbpl11 | -0.070096472 | 0.88951533 |
| Shisa6 | 0.103936786 | 0.889927723 |
| Olfr1414 | -0.672067315 | 0.890181858 |
| Ttll7 | 0.088369891 | 0.890224816 |
| Gtf2a1 | 0.046270463 | 0.890224816 |
| Srsf4 | -0.043140039 | 0.890224816 |
| Fam206a | -0.074991683 | 0.890235975 |
| Ankrd42 | -0.085329609 | 0.890235975 |
| Kbtbd13 | -0.64113466 | 0.890235975 |
| Irx3 | 0.259674667 | 0.890303054 |
| Mrpl19 | 0.057083217 | 0.890303054 |
| Armc1 | 0.043446028 | 0.890303054 |
| Apbb1 | 0.038853498 | 0.890303054 |
| Gm2026 | -0.431359548 | 0.890303054 |
| Ddx51 | 0.107637046 | 0.890385964 |
| Hdac9 | 0.100699681 | 0.890555583 |
| Sufu | 0.068892714 | 0.89061081 |
| Rnf114 | -0.041044953 | 0.89061081 |
| Utp23 | -0.065550215 | 0.89061081 |
| Zdhhc1 | -0.102738695 | 0.89061081 |
| Tead3 | 0.216537858 | 0.890661308 |
| Clcn5 | 0.099298077 | 0.890661308 |
| Hsd17b4 | -0.057391575 | 0.890661308 |
| Utp11l | 0.059860297 | 0.891242762 |
| Zfp672 | -0.07279736 | 0.891242762 |
| Gatc | 0.054112641 | 0.89140832 |
| Eif4e3 | -0.110685862 | 0.89140832 |
| Rab11fip4 | 0.059709954 | 0.89142592 |
| Etfdh | 0.071508909 | 0.891567405 |
| Smo | -0.094857237 | 0.891937371 |
| Pdk3 | 0.056193242 | 0.892077349 |
| Pcyt1b | 0.075769044 | 0.892087847 |
| Chd4 | 0.040316639 | 0.892087847 |
| Wdr53 | 0.080899512 | 0.89233934 |
| Hspbap1 | 0.103990178 | 0.892361191 |
| Cep170 | 0.042890978 | 0.892361191 |
| Fosl1 | 0.27325486 | 0.89255108 |
| Ptpn18 | 0.071171045 | 0.89255108 |
| Hist2h2aa2 | -0.107377019 | 0.89255108 |
| Fcer2a | -0.275045511 | 0.89255108 |
| Rpl7l1 | 0.038784806 | 0.892709227 |
| Zc3h12b | 0.159652094 | 0.892856013 |
| Plxna1 | 0.076108572 | 0.89298896 |
| Zfp526 | -0.155827395 | 0.89298896 |
| Trnt1 | 0.050120352 | 0.893089023 |
| Arhgef9 | 0.064683246 | 0.893321677 |
| Lonp1 | 0.060959425 | 0.893321677 |
| Hs6st3 | -0.293703978 | 0.893510016 |
| Dok5 | 0.073796512 | 0.893511837 |
| Orai3 | 0.109166089 | 0.893661264 |
| Fam84a | -0.084987002 | 0.893661264 |

| | | |
|---|---|---|
| U2af1l4 | 0.824745191 | 0.005546196 |
| Fcrlb | -2.954628444 | 0.005549556 |
| Glrx2 | 0.651506358 | 0.005559063 |
| Dvl1 | 1.263105297 | 0.005574149 |
| P4ha1 | -0.794655804 | 0.0055813 |
| Rps12 | 0.846795899 | 0.005598308 |
| Scara5 | 6.159904543 | 0.005598527 |
| Cd320 | 0.884342293 | 0.005605768 |
| Flywch2 | 1.336638258 | 0.005614051 |
| Lrrc49 | 0.70653499 | 0.005625436 |
| Kdm4b | -0.77026536 | 0.0056499 |
| Gm15056 | -4.10142745 | 0.005662149 |
| Etv6 | -0.942361962 | 0.005664578 |
| Ndufc2 | 0.72245098 | 0.00566936 |
| Fancg | -1.046339185 | 0.005669372 |
| Sulf1 | 1.239318706 | 0.005671406 |
| Notch3 | 2.259596448 | 0.005678254 |
| Rhbdf1 | 2.750121165 | 0.005685338 |
| Col12a1 | 1.767102076 | 0.005702005 |
| Brinp3 | 1.517066379 | 0.005702005 |
| Paqr7 | -1.305763084 | 0.005702005 |
| Prpf8 | -0.535883115 | 0.005711778 |
| Arhgap39 | -0.967129707 | 0.005721418 |
| Tmem81 | 6.058374696 | 0.005737179 |
| Trafd1 | -0.759833977 | 0.00574555 |
| Qrsl1 | -1.096100106 | 0.005761512 |
| Kcnf1 | -2.490947723 | 0.00576291 |
| Ncald | 1.42525725 | 0.005763116 |
| Rnf5 | 0.642419859 | 0.005772718 |
| Sap18 | 0.594382192 | 0.005772718 |
| Uty | -0.771143388 | 0.005774645 |
| Prdm6 | 6.021423204 | 0.00578095 |
| Ppp1cb | 0.60459199 | 0.005816985 |
| Stim2 | -0.856660528 | 0.005816985 |
| Cln5 | -1.308284863 | 0.005816985 |
| Tifab | -1.440986245 | 0.005816985 |
| Zc3h3 | -1.249614869 | 0.005826491 |
| Dync2li1 | 0.914457138 | 0.00583054 |
| H2-Aa | 6.280009018 | 0.005839674 |
| Spon2 | 4.427863988 | 0.005840125 |
| Hsd17b10 | 0.595053764 | 0.005841247 |
| Slc45a3 | -2.599554252 | 0.005860911 |
| Hrasls | 6.00003813 | 0.005862613 |
| Fignl1 | -1.508882629 | 0.005876825 |
| Elavl1 | -0.564411925 | 0.00588302 |
| Rhcg | 5.979137528 | 0.005914689 |
| Caly | 1.086309397 | 0.005914689 |
| Gm15448 | -3.008457203 | 0.005918854 |
| Eef1a1 | 0.509422442 | 0.005929519 |
| E2f8 | -1.284925095 | 0.005929912 |
| Tasp1 | 1.06725692 | 0.00598348 |
| Coa3 | 0.574225425 | 0.005996166 |
| Cdk6 | 0.882972931 | 0.005997917 |
| Ift74 | 1.103293657 | 0.006005916 |
| Fbxw5 | 0.664977761 | 0.006036367 |
| Hmgn1 | 0.474090212 | 0.006036367 |
| Smoc2 | 4.488227905 | 0.006038928 |
| Frmpd1 | 6.161671058 | 0.006045883 |
| Mrpl12 | 0.643242571 | 0.00605405 |
| Ddx6 | -0.503059485 | 0.006082538 |
| Mrpl9 | 0.700048968 | 0.00609603 |
| Tceal | 0.851753537 | 0.006110545 |
| Dnajc12 | 1.253917389 | 0.00612233 |
| Zfc3h1 | -0.776820029 | 0.006127261 |
| Rnf126 | -0.825654961 | 0.006127261 |
| Rbm7 | -0.647254547 | 0.006141853 |
| Dync2h1 | 0.870460831 | 0.006156891 |
| Slc2a9 | -2.045704969 | 0.006168695 |
| Zzef1 | -0.685545654 | 0.006170458 |
| Fap | -4.566011202 | 0.006184469 |
| Acyp1 | 0.953064495 | 0.006202861 |
| Siah1b | 0.94703222 | 0.006223721 |
| Socs1 | -1.840030689 | 0.00622534 |
| Zfp652 | -0.753533548 | 0.006252415 |
| Birc5 | 0.811986659 | 0.006272083 |
| Itgb1 | -0.608391749 | 0.006282702 |
| Fbxo30 | -0.6526003 | 0.006282702 |
| Atp7a | -0.759184973 | 0.006282702 |
| Pcdhgb6 | -1.748300075 | 0.006300121 |
| Mrc2 | 2.571686024 | 0.0063128 |
| Cdk8 | -0.745167793 | 0.006313287 |
| Mex3b | 0.968329498 | 0.006324495 |
| Smurf1 | -0.976166532 | 0.006325902 |
| Laptm5 | -1.012085721 | 0.006334784 |
| Rnf113a1 | 1.795942025 | 0.00634707 |
| 7-Sep | 0.533593021 | 0.006357659 |
| Serpind1 | 3.790374181 | 0.006357994 |
| Arhgef10l | -1.014857069 | 0.006362836 |
| Pi4k2a | -1.011294271 | 0.006370918 |
| Pde6d | 0.633528695 | 0.006395419 |
| C1ra | 6.27165175 | 0.006411164 |
| Stk19 | 1.028688903 | 0.006424265 |
| Arl8a | -0.835692166 | 0.006425248 |
| Dmwd | 0.883041053 | 0.006435426 |
| Rnpep | -0.761420174 | 0.006452718 |
| Tbx15 | 6.047739325 | 0.006458627 |
| Atp9b | -0.865191892 | 0.00645983 |
| Clec4e | -2.633117218 | 0.00645983 |
| Zfp473 | 6.068404456 | 0.00648558 |
| Fkbp10 | 2.457046145 | 0.006488691 |
| Tmem179 | 1.286691669 | 0.00651811 |
| Sh3bp1 | -0.729767081 | 0.006565838 |
| Abca1 | -0.90722106 | 0.006565838 |
| Slc4a1 | 3.73882532 | 0.0065759 |
| Glcci1 | 0.894558173 | 0.0065759 |
| Fyb | -0.938983572 | 0.006636949 |
| 1810037I17Rik | 0.661651115 | 0.00663695 |
| Ipcef1 | -1.523394415 | 0.00663695 |
| Snca | 0.785685347 | 0.006654958 |
| Sptlc2 | -0.931139185 | 0.006663727 |
| Fbxl17 | 0.97691727 | 0.006680442 |
| Mcm3 | -0.7374956 | 0.006706258 |
| Vps36 | 0.657268133 | 0.00673182 |
| Pcbp4 | 0.655308836 | 0.006736694 |
| Pcna | 0.648606841 | 0.006736694 |
| Myeov2 | 0.655347102 | 0.006766091 |
| Midn | -0.710648998 | 0.006766091 |
| Pltp | -1.12013181 | 0.006766091 |
| Mrpl22 | 0.76473052 | 0.006777216 |

| | | |
|---|---|---|
| Apex1 | 0.522780098 | 0.157437382 |
| Nkpd1 | 4.592928866 | 0.157621985 |
| Osgin2 | -0.485297912 | 0.157621985 |
| A630001G21Rik | -0.825121881 | 0.157621985 |
| Fam134b | -1.003239269 | 0.157621985 |
| Gm26992 | -1.131050493 | 0.157716855 |
| Ptp4a3 | -0.381973769 | 0.157790971 |
| Pcnt | 0.523360473 | 0.157817794 |
| S100pbp | 0.819042162 | 0.157867105 |
| Itpr1 | -0.485464354 | 0.157883015 |
| Glmp | -0.549018569 | 0.157883015 |
| Tmem141 | 0.625397208 | 0.158011818 |
| Sgcb | 0.495141764 | 0.158011818 |
| Gfer | 0.445913542 | 0.158011818 |
| Pafah1b2 | -0.268400098 | 0.158134525 |
| Lrrc8b | -0.553655484 | 0.158249651 |
| Ccdc183 | 3.204137032 | 0.158258703 |
| Shroom2 | 0.676764847 | 0.15826546 |
| Zfp366 | -2.419986103 | 0.15826546 |
| Rdh5 | -1.472705863 | 0.15826688 |
| Xntrpc | 3.747555752 | 0.158268309 |
| Uvrag | -0.524465624 | 0.158268309 |
| Lrrc6 | -2.122588852 | 0.158268309 |
| Syt6 | -0.56513284 | 0.158414136 |
| Exosc1 | 0.450640911 | 0.158519774 |
| Ccbe1 | 1.362900368 | 0.15857558 |
| Mrps18b | 0.652546493 | 0.158772339 |
| Gm6904 | -2.174441892 | 0.159006146 |
| Slc25a17 | 0.357677119 | 0.15931191 |
| Phyh | -0.456563779 | 0.159522874 |
| Eif2d | 0.425591065 | 0.159549825 |
| Snrnp40 | 0.400017001 | 0.159549825 |
| Btf3 | 0.268291037 | 0.159549825 |
| Mnt | -0.519983194 | 0.159577586 |
| Rad51 | -0.520924157 | 0.159577586 |
| Slc36a1 | -1.020866312 | 0.15960782 |
| Fam19a2 | 1.20431812 | 0.15972649 |
| Slc39a7 | -0.37157752 | 0.159748394 |
| Ifi30 | -0.625969183 | 0.159752993 |
| Gdap2 | -0.46887239 | 0.159830873 |
| Nkd1 | 0.971748772 | 0.16001253 |
| Kcnd2 | 1.002365645 | 0.160021713 |
| Naip2 | -0.894228855 | 0.160021713 |
| Prpf39 | 0.479329854 | 0.160088814 |
| Smn1 | 0.480661143 | 0.160241092 |
| Fam71f2 | 2.658216252 | 0.160250259 |
| Snf8 | -0.398579159 | 0.160250259 |
| Tango2 | 0.525411707 | 0.160275137 |
| Sptbn2 | -0.535397525 | 0.160302649 |
| Sugp2 | -0.585975827 | 0.160302649 |
| Kansl2 | 0.348370119 | 0.160343831 |
| Isg20 | -1.503125805 | 0.160343831 |
| Dgat2 | -0.806908245 | 0.160812517 |
| Emx1 | 0.61455114 | 0.160924471 |
| Mrpl50 | 0.403328807 | 0.160924471 |
| Ptpn21 | 1.242644429 | 0.160999791 |
| Svip | 0.621676762 | 0.161006629 |
| Fcna | 1.982532328 | 0.161013558 |
| Ttll8 | 3.786921575 | 0.161091494 |
| Eri2 | 0.799367452 | 0.161091494 |
| Celsr2 | 0.505619739 | 0.16112257 |
| Hyls1 | 0.692300842 | 0.16139733 |
| S100b | -1.757971429 | 0.161493646 |
| Cmtm5 | 4.389439924 | 0.16152402 |
| Tm4sf1 | 1.746937894 | 0.16152402 |
| Ubl7 | 0.339221202 | 0.161754586 |
| Sae1 | 0.2781619 | 0.161754586 |
| Mbl1 | -5.342342403 | 0.161781437 |
| Sstr4 | 4.511416112 | 0.161803857 |
| Serpinb5 | 4.841141219 | 0.161841789 |
| S100a8 | -2.004566705 | 0.161855234 |
| Cebpe | -2.475585076 | 0.161855234 |
| Pds5b | 0.347191408 | 0.161929638 |
| Arsi | 4.568271372 | 0.161939598 |
| Baz2b | 0.394280633 | 0.162059871 |
| Pcdhb11 | -1.109353176 | 0.162089937 |
| Spaca1 | 4.559367621 | 0.162124202 |
| Lamc1 | -0.757097913 | 0.162163803 |
| Mfn2 | -0.522151705 | 0.162234039 |
| Fbxo42 | -0.544098829 | 0.162312999 |
| Ss18l1 | -0.483572633 | 0.162383149 |
| Hdgf | 0.339867442 | 0.162452451 |
| Vps9d1 | 0.457296725 | 0.162454365 |
| Pgm2l1 | 0.708409166 | 0.162511497 |
| Ctbp1 | -0.350477842 | 0.162689933 |
| Itga8 | -1.288671652 | 0.162851034 |
| Chmp3 | 0.367116127 | 0.162851367 |
| Ebf3 | -0.890345353 | 0.162851367 |
| Fam168b | -0.324558893 | 0.162913686 |
| Pmvk | 0.339220746 | 0.162969217 |
| Pold3 | -0.363014229 | 0.163086248 |
| Rnf135 | -0.783399039 | 0.163146432 |
| 11-Sep | 0.270430293 | 0.163400432 |
| Mfap3 | 0.360902947 | 0.163478596 |
| Rusc2 | -0.32605158 | 0.163573298 |
| Bdp1 | 0.382694398 | 0.163607042 |
| Btd | 0.829433414 | 0.163615573 |
| Pvalb | 4.923983934 | 0.163723114 |
| Stard10 | 1.211594212 | 0.164005893 |
| Igsf11 | 1.589903577 | 0.16401144 |
| Ykt6 | -0.363833697 | 0.164027435 |
| Batf3 | -1.006384379 | 0.164113746 |
| Mapk8ip2 | 0.431282972 | 0.164211198 |
| Fam162b | 4.873893025 | 0.164266546 |
| Rhoc | 0.65804917 | 0.164273268 |
| Atxn1l | -0.722330958 | 0.164273268 |
| Mcam | 1.767406355 | 0.164575843 |
| Myo10 | 0.563614745 | 0.16460031 |
| Dusp23 | 1.311801184 | 0.164622974 |
| Eprs | 0.294507884 | 0.164656176 |
| Krt19 | 4.456772167 | 0.164662152 |
| Gmeb1 | 0.450297547 | 0.164766034 |
| Ddb1 | -0.310105777 | 0.164766034 |
| Gpr156 | 0.984718729 | 0.164769512 |
| Hnrnph1 | 0.315298816 | 0.164860732 |
| Csnk2a2 | 0.69981908 | 0.164933125 |
| Ribc1 | -1.170013046 | 0.164933125 |
| Stoml2 | 0.304659983 | 0.164994491 |
| Slc12a8 | 3.916119244 | 0.165006769 |

| | | |
|---|---|---|
| Ppip5k2 | -0.200698948 | 0.503400975 |
| Rnf125 | -1.050641004 | 0.503585999 |
| Akap8l | 0.207779346 | 0.503595828 |
| Zfp407 | -0.322049369 | 0.503595828 |
| Tmem252 | 1.094255284 | 0.503656482 |
| Oxsr1 | 0.296267889 | 0.503656482 |
| Akt2 | -0.228510198 | 0.503656482 |
| Zfp90 | 0.275804054 | 0.503716341 |
| Fpgt | 0.30896238 | 0.503889272 |
| Casc5 | 0.274753407 | 0.503889272 |
| Clip3 | -0.190946332 | 0.503889272 |
| Ankrd45 | -0.352136418 | 0.504933748 |
| Fam122b | -0.441355907 | 0.504933748 |
| Slc13a5 | 1.116668751 | 0.505084536 |
| A130057D12Rik | 1.705098926 | 0.505240352 |
| Tmem144 | -1.463359212 | 0.505263855 |
| Arhgef10 | 0.77940938 | 0.505585566 |
| Ccdc65 | 0.916781892 | 0.505781167 |
| Krtdap | 3.497418898 | 0.505893635 |
| Hdac3 | 0.152181093 | 0.50594092 |
| Nxnl1 | 3.471034437 | 0.505957482 |
| Abcb1a | 2.63098131 | 0.505957482 |
| Hist1h2ah | 2.010567747 | 0.505957482 |
| Doc2a | 1.704917303 | 0.505957482 |
| Plekhf1 | 1.482497188 | 0.505957482 |
| Mme | 0.708688056 | 0.505957482 |
| Nans | 0.171678078 | 0.505957482 |
| Csnk1a1 | 0.144653312 | 0.505957482 |
| Gucy1a3 | -0.280626761 | 0.505957482 |
| Mmp24 | -0.432186634 | 0.505957482 |
| Bms1 | 0.283415439 | 0.505987953 |
| Oaz1 | 0.152990358 | 0.506015386 |
| Scn4b | 3.172415496 | 0.506111144 |
| 6430531B16Rik | 1.876424445 | 0.506405956 |
| Sh3glb2 | 0.202376006 | 0.506405956 |
| Cstf3 | -0.263883435 | 0.506534866 |
| Abcg2 | 0.33417896 | 0.506543515 |
| Csde1 | -0.150550899 | 0.506702816 |
| Pot1a | -0.323230675 | 0.506836895 |
| Gcc1 | -0.555217471 | 0.506943455 |
| Insl6 | -0.882514881 | 0.506969437 |
| 2310003L06Rik | 3.464708336 | 0.507088179 |
| Fndc3a | 0.213136529 | 0.507125309 |
| Aox2 | 2.815377857 | 0.507125557 |
| Dab2ip | -0.260440435 | 0.507152543 |
| Asb13 | 0.279428399 | 0.507381524 |
| Lrrfip2 | 0.228274564 | 0.507381524 |
| Emsy | 0.301929241 | 0.507524391 |
| Gm3411 | 0.824868484 | 0.507623872 |
| Ino80e | 0.278976557 | 0.507623872 |
| Camkv | -0.372395041 | 0.507719873 |
| Dpep2 | -0.815419997 | 0.507748465 |
| Trim43b | 3.175064474 | 0.507976475 |
| Trpc6 | 3.175064474 | 0.507976475 |
| Phactr1 | -0.243792499 | 0.508076081 |
| Pon2 | -0.230795671 | 0.508167816 |
| Thap11 | 0.208268719 | 0.508255366 |
| Wash1 | -0.291733874 | 0.508460917 |
| Mkrn2 | 0.249033307 | 0.508588053 |
| Cd163l1 | 3.713342179 | 0.508677268 |
| Slc27a6 | 3.713342179 | 0.508677268 |
| Dlg1 | -0.280621272 | 0.508677268 |
| Wtap | -0.193361294 | 0.508719857 |
| Ahnak2 | 1.093280278 | 0.508780515 |
| Gtf3a | 0.405398931 | 0.508780515 |
| Tpst1 | 0.242875828 | 0.50919504 |
| A930004D18Rik | 0.611979898 | 0.509210994 |
| Dscam | 0.441376627 | 0.509287283 |
| Zfp954 | -0.345332049 | 0.509335966 |
| Usp27x | 0.483639239 | 0.509404461 |
| Tspan11 | 0.450932779 | 0.509876014 |
| Tecr | 0.152696275 | 0.509876014 |
| Lin7a | 0.317583599 | 0.509879397 |
| Mier2 | -0.300945794 | 0.509968024 |
| Ogdhl | -0.294095213 | 0.510056217 |
| Slc25a20 | 0.371037252 | 0.510192374 |
| Ptpn5 | 0.290840049 | 0.510349482 |
| Klf8 | 0.504104628 | 0.510386053 |
| Igsf8 | 0.2547465 | 0.510386053 |
| Ccdc105 | 3.476410878 | 0.510605009 |
| Ferd3l | 3.476410878 | 0.510605009 |
| Rassf8 | 0.719901077 | 0.510854468 |
| Lonrf3 | 0.741255437 | 0.511116442 |
| Cntd1 | 3.694533357 | 0.511244451 |
| Krt79 | 3.694533357 | 0.511244451 |
| Cd2bp2 | -0.196180237 | 0.511409042 |
| Wdr5b | -0.675251578 | 0.511437753 |
| Gm26558 | -1.272182249 | 0.511691598 |
| 1700003F12Rik | -0.97653918 | 0.511731916 |
| Elovl2 | -0.266132485 | 0.511739725 |
| Tdrd9 | 3.44742736 | 0.511741216 |
| Reps1 | -0.31884954 | 0.511791738 |
| Tex26 | 1.326715282 | 0.51196116 |
| Mkln1 | -0.184255083 | 0.512068767 |
| Gpatch4 | 0.272051481 | 0.512117591 |
| Ank2 | -0.202145553 | 0.512117591 |
| Npm2 | 3.151377946 | 0.5125377 |
| Xrra1 | 1.990420147 | 0.5125377 |
| Eral1 | 0.393719735 | 0.5125377 |
| Bfar | 0.261246149 | 0.5125377 |
| Acss2 | -0.334884131 | 0.5125377 |
| Ptprt | -0.339567437 | 0.5125377 |
| Cand2 | -0.357904951 | 0.5125377 |
| A330017A19Rik | -0.645777139 | 0.5125377 |
| Olfr1000 | 3.468936733 | 0.51265802 |
| Plrg1 | -0.200722621 | 0.513073754 |
| Pkn3 | -0.743101842 | 0.513078902 |
| Ambra1 | -0.263316494 | 0.513179981 |
| Loxl3 | -0.502394895 | 0.513503112 |
| Trim13 | -0.296498843 | 0.513549193 |
| Ovca2 | 0.570532374 | 0.513806458 |
| Fam109a | 0.296330716 | 0.513897602 |
| Fnbp1 | -0.220984478 | 0.513897602 |
| Tnfrsf8 | -1.069348456 | 0.513897602 |
| Poldip2 | 0.183509339 | 0.51390046 |
| Carhsp1 | -0.212373631 | 0.51390046 |
| Tslp | 2.775233235 | 0.513916301 |
| Col16a1 | -0.827122884 | 0.513928501 |
| Prkcsh | 0.208358532 | 0.514130584 |

| | | |
|---|---|---|
| Ankdd1a | -0.50052525 | 0.893740136 |
| Hint3 | -0.085730445 | 0.893755278 |
| Thra | -0.051511371 | 0.894069618 |
| Csk | -0.048502926 | 0.894104029 |
| Zmat1 | 0.100079484 | 0.894118249 |
| Myrip | 0.470667784 | 0.894255726 |
| 4930430F08Ril | 0.108494125 | 0.894429538 |
| Zfp655 | 0.042331282 | 0.894429538 |
| Cbll1 | -0.054414025 | 0.894429538 |
| Dirc2 | -0.063414157 | 0.894429538 |
| Zfp629 | -0.083539923 | 0.894429538 |
| Nfatc2ip | -0.093093422 | 0.894429538 |
| Hist1h4f | -0.372474065 | 0.894506117 |
| Rbm38 | 0.051896246 | 0.894590622 |
| Chd1 | 0.043831011 | 0.894831905 |
| Tnr | -0.085419023 | 0.895097842 |
| Cst7 | 0.54887073 | 0.895705175 |
| Ptger1 | -0.113487363 | 0.895791747 |
| 1700003E16Ril | 0.247395628 | 0.895840008 |
| Fndc5 | 0.212880007 | 0.895840008 |
| Gm7293 | -0.596582905 | 0.896086415 |
| 2610021A01Ri | -0.088881318 | 0.896632043 |
| Osbpl10 | 0.286454728 | 0.896645591 |
| C1ql1 | 0.254693667 | 0.896645591 |
| Zfp608 | -0.044946855 | 0.896645591 |
| Becn1 | -0.049159751 | 0.896645591 |
| Slc23a1 | -0.168281186 | 0.896645591 |
| Pcdhb10 | -0.21102921 | 0.896645591 |
| Fcgrt | 0.089416128 | 0.89672527 |
| Tprn | 0.089587017 | 0.896928058 |
| Adat1 | -0.115077346 | 0.896928058 |
| Atp7b | -0.145389971 | 0.897180326 |
| P2rx2 | -0.130931937 | 0.897199161 |
| Dusp11 | -0.043370027 | 0.897323927 |
| Ints6 | -0.09559803 | 0.897323927 |
| Drd4 | -0.526956917 | 0.897413167 |
| Gm7535 | -0.222235769 | 0.897619345 |
| Micalcl | 0.351763113 | 0.898083811 |
| Taf1c | -0.101436436 | 0.898083811 |
| B2m | -0.061063566 | 0.898132123 |
| Btg3 | 0.07324399 | 0.89844668 |
| Ranbp3 | 0.055498516 | 0.898557794 |
| Dzip1l | 0.102926986 | 0.898970635 |
| Rbm24 | -0.153798344 | 0.898970635 |
| Xlr4b | -0.260475884 | 0.898970635 |
| Dsn1 | 0.077940228 | 0.899023568 |
| Rnf20 | 0.043558226 | 0.899410661 |
| Kirrel | -0.122051862 | 0.899410661 |
| Spg7 | 0.068963237 | 0.89945324 |
| Parn | -0.059246612 | 0.899500706 |
| Cyp27b1 | -0.475935478 | 0.899628247 |
| BC023829 | -0.071747012 | 0.899662983 |
| Masp2 | 0.373027105 | 0.899924279 |
| Neil2 | 0.336592688 | 0.899924279 |
| Srrm4 | -0.056219518 | 0.899924279 |
| Gm11710 | -0.472767233 | 0.899924279 |
| Serpina3n | 0.367881189 | 0.900032606 |
| Lrrn2 | 0.058089332 | 0.900032606 |
| Aph1c | -0.134343632 | 0.900032606 |
| Atp6v0b | -0.033077725 | 0.900063378 |
| Xpnpep3 | 0.156440457 | 0.90017645 |
| Nudt5 | 0.044232793 | 0.900224065 |
| Dusp15 | -0.12172745 | 0.900260209 |
| Nus1 | -0.056044341 | 0.900267416 |
| Vcan | -0.069992949 | 0.900267416 |
| Rgl2 | -0.079999821 | 0.900267416 |
| Nme2 | -0.270740166 | 0.900267416 |
| Rmnd1 | -0.050999711 | 0.900534292 |
| Ehf | -0.363687131 | 0.900605155 |
| Gm3164 | -0.45432943 | 0.900633462 |
| Ppp2r5d | -0.051046982 | 0.900874588 |
| Ttyh2 | 0.055659442 | 0.90115192 |
| Znhit2 | 0.054074987 | 0.90115192 |
| Sar1a | 0.034980443 | 0.90115192 |
| Gpr176 | -0.169210367 | 0.90115192 |
| A830031A19Ri | -0.441874169 | 0.90115192 |
| Nrxn3 | 0.061364519 | 0.901175087 |
| Plppr3 | -0.042105498 | 0.901175087 |
| Ngb | -0.238721015 | 0.901175087 |
| Rfx3 | 0.054677849 | 0.901197977 |
| Apoc1 | 0.066198036 | 0.901319901 |
| Tmem40 | -0.235153933 | 0.901319901 |
| Ccm2 | -0.052238124 | 0.901447449 |
| Masp1 | -0.068845838 | 0.901447449 |
| Fan1 | 0.12114433 | 0.901703954 |
| Zfp251 | -0.05605729 | 0.901703954 |
| Ublcp1 | -0.067527326 | 0.901703954 |
| Klc4 | -0.046179502 | 0.901798058 |
| Saa2 | -0.246258637 | 0.901927771 |
| Ugt1a6b | -0.299901659 | 0.901927771 |
| Prss52 | 0.323842475 | 0.90200345 |
| Entpd1 | 0.098624276 | 0.90200345 |
| Fbxo25 | 0.061843575 | 0.902613939 |
| Ndst1 | 0.057309304 | 0.902786188 |
| Shmt1 | -0.06428994 | 0.903323737 |
| Sorcs1 | -0.113516598 | 0.903416907 |
| Pde8b | 0.152368649 | 0.903517194 |
| Heatr5b | 0.067057017 | 0.903517194 |
| Fgd4 | -0.045347458 | 0.903517194 |
| Serpina3f | 0.233189433 | 0.903645619 |
| Plin5 | 0.244098026 | 0.903810147 |
| Zfyve9 | -0.05484327 | 0.903850029 |
| Atg2b | -0.060295158 | 0.903956341 |
| Ttbk1 | -0.120237189 | 0.904129941 |
| Dnajb13 | 0.114464196 | 0.904415635 |
| Raf1 | -0.039973208 | 0.904415635 |
| Fbll1 | -0.129072354 | 0.904415635 |
| Tal1 | -0.253853827 | 0.904486843 |
| Ppcs | -0.083472932 | 0.904541119 |
| Lta4h | -0.045234517 | 0.904675777 |
| Gm17296 | 0.116941591 | 0.90476756 |
| Ube2i | 0.032077265 | 0.90476756 |
| Cntrl | -0.048473097 | 0.90476756 |
| Elmod1 | -0.07863448 | 0.90476756 |
| Aco2 | -0.037388544 | 0.90483043 |
| Zfp362 | -0.051223709 | 0.904897607 |
| Slc35f3 | -0.269688561 | 0.904897607 |
| Gna11 | -0.041002706 | 0.904907405 |
| Ccdc88c | 0.069669681 | 0.904919636 |

| | | |
|---|---|---|
| Dkkl1 | 5.853761085 | 0.006817225 |
| Cdh7 | 1.935575587 | 0.006843656 |
| Jkamp | 0.583900777 | 0.006846971 |
| Map7d1 | -0.547535873 | 0.006846971 |
| Tnnc1 | 5.902641862 | 0.006855031 |
| Clcn7 | -0.839714273 | 0.006861989 |
| Pvrl4 | -2.491314447 | 0.00688611 |
| Dctn6 | 0.651963974 | 0.006888062 |
| Fbxo38 | -0.617169792 | 0.00690796 |
| Galnt4 | -1.631770342 | 0.006942507 |
| Rfc4 | 0.727455355 | 0.006968296 |
| Cdk19 | -0.985400147 | 0.00696909 |
| Atp11b | -0.877501376 | 0.007001992 |
| Alkbh7 | 0.973956455 | 0.007004217 |
| Rfesd | 0.966973132 | 0.007004217 |
| Zfp326 | 0.604315191 | 0.007004217 |
| Bub3 | 0.580148771 | 0.007004217 |
| Crlf2 | -1.549037016 | 0.007004217 |
| Gm12942 | 1.308157787 | 0.007004324 |
| Reep5 | -0.537861254 | 0.007021446 |
| Hcar2 | -3.533964168 | 0.007030475 |
| Asphd2 | 1.176516974 | 0.007071278 |
| Rabep1 | 0.596024369 | 0.007095965 |
| Ccdc124 | 0.685671998 | 0.007111895 |
| Sema5a | 1.016444635 | 0.007114793 |
| Wars | -0.69078188 | 0.007114793 |
| Me3 | 3.007804067 | 0.007121614 |
| Hebp1 | -1.079425574 | 0.007121614 |
| Hspa1b | -2.50492207 | 0.007152466 |
| Eya4 | 5.846882884 | 0.007162759 |
| Tgfb1i1 | 1.90209418 | 0.007175792 |
| Eif3e | 0.531610783 | 0.007200555 |
| Arhgap32 | 0.870074997 | 0.007220464 |
| Nfe2l1 | -0.784344179 | 0.007225664 |
| Zfhx3 | 0.917770473 | 0.007227479 |
| Ppih | 0.745753159 | 0.007230694 |
| Kcnn4 | -2.416172857 | 0.007242902 |
| Dus1l | 0.822590415 | 0.007258703 |
| Ncoa1 | 0.926079054 | 0.007279825 |
| Necap2 | -0.838542523 | 0.00734198 |
| Gkap1 | 0.996440918 | 0.007384356 |
| Reep3 | -0.615703903 | 0.007384356 |
| Gpx4 | 0.523056734 | 0.007391239 |
| Nfkbid | -1.449970078 | 0.007419727 |
| Aqp3 | -6.9168404 | 0.007419727 |
| Zfp831 | -6.47686039 | 0.007434923 |
| Gstm7 | 0.784207481 | 0.007439418 |
| Slc39a12 | 3.78897366 | 0.007454551 |
| Hars | 0.609035617 | 0.007496619 |
| Tmem63c | -1.926915044 | 0.007527003 |
| Ing1 | 0.879975489 | 0.00752764 |
| Ncln | -0.835835813 | 0.00752764 |
| Cwc27 | 0.94883016 | 0.007531145 |
| Acy1 | 1.032197571 | 0.00754447 |
| Nabp1 | -1.958586228 | 0.00754447 |
| Ppp1r16a | 0.695641671 | 0.007572353 |
| Zfp57 | -0.683472434 | 0.007608266 |
| Slc25a26 | -1.545818097 | 0.00763616 |
| Dclk3 | 3.081938126 | 0.007638642 |
| Coch | -2.05535798 | 0.007650522 |
| Gng2 | 0.521974093 | 0.007676771 |
| Psmb1 | 0.493273644 | 0.007676771 |
| Plin4 | -3.529825113 | 0.007676771 |
| Cst3 | 0.625343117 | 0.007679553 |
| Ccdc116 | -3.810791332 | 0.007679553 |
| Nsmce1 | 0.55990433 | 0.007688318 |
| Fam178a | -0.625010034 | 0.007734941 |
| C77370 | 0.792144408 | 0.00775231 |
| Mettl22 | 0.975055386 | 0.007800347 |
| Serpine2 | 0.99943141 | 0.007825301 |
| Them5 | -6.558155032 | 0.007825301 |
| Ripply2 | 2.852847444 | 0.007835264 |
| Gbp4 | -2.768015996 | 0.007835264 |
| Scarf1 | 5.736687928 | 0.007835597 |
| Wnk3 | 0.898026333 | 0.007853139 |
| Nfe2l3 | -1.721354534 | 0.007854033 |
| Rsph3a | 0.843163633 | 0.007863556 |
| Tbc1d2b | -1.270960807 | 0.007879026 |
| Timm17b | 0.692516829 | 0.007880126 |
| H60b | -2.916658532 | 0.007880126 |
| Prkdc | -0.813716677 | 0.007889715 |
| Cyp4v3 | -1.619817126 | 0.007889715 |
| Pcdhga9 | -1.451814657 | 0.007925667 |
| Dennd1a | -0.846229676 | 0.007927336 |
| Tmem128 | 0.660787094 | 0.00793069 |
| Rp2 | -0.950122281 | 0.00793176 |
| Map3k1 | -1.215660773 | 0.00793176 |
| Gphn | 0.940911963 | 0.007952783 |
| Gas2l1 | -0.895281796 | 0.00795316 |
| Sardh | 5.745990867 | 0.007960744 |
| Trib3 | -3.746270711 | 0.007970898 |
| Zfp35 | 0.87978485 | 0.00797626 |
| Tnip3 | -2.68468903 | 0.007984979 |
| Urb2 | -1.184574079 | 0.007989804 |
| Atg12 | 0.628615365 | 0.008028519 |
| Kazald1 | 5.73708524 | 0.008046943 |
| Gys1 | -0.915713503 | 0.008078374 |
| Pcdha11 | -2.096326176 | 0.008078374 |
| 1700013F07Rik | 4.550624731 | 0.008097995 |
| Mrps12 | 0.624767086 | 0.008098799 |
| Kcnn2 | -0.782469976 | 0.008104867 |
| BC117090 | -2.438958543 | 0.008169595 |
| Sec24b | -0.625605884 | 0.008182222 |
| Dpcd | 0.750207207 | 0.008202053 |
| Wdr91 | -0.988188924 | 0.008215893 |
| Clk2 | -0.991222661 | 0.008259951 |
| Tmed2 | -0.569342838 | 0.008300461 |
| Smim18 | 1.119067716 | 0.008313582 |
| Spcs1 | 0.491473548 | 0.00833135 |
| St3gal3 | -0.85705933 | 0.008338676 |
| Sox11 | 0.603769189 | 0.00834435 |
| Tsg101 | 0.476624942 | 0.008357622 |
| Gbx1 | 5.761794264 | 0.008389495 |
| Atp6v1h | -0.491326002 | 0.008407243 |
| 8-Mar | -0.647409727 | 0.008411992 |
| Cfap206 | 5.729403384 | 0.008422851 |
| Egr3 | -6.526864721 | 0.008422851 |
| Prr5l | 2.486230834 | 0.008458791 |
| Vstm5 | 2.914444874 | 0.008474343 |

| | | |
|---|---|---|
| Ano6 | -0.551534267 | 0.165027925 |
| Pcdha6 | 2.075789339 | 0.165032032 |
| Mysm1 | -0.368014882 | 0.165060391 |
| 8430408G22Rik | -2.233678985 | 0.165195847 |
| Nup62cl | -5.328641809 | 0.165343273 |
| Ube2u | -5.345643621 | 0.165343273 |
| Rhd | 4.433140811 | 0.165358517 |
| Rbm5 | -0.312075237 | 0.165541084 |
| Snx16 | 0.501680818 | 0.165555505 |
| Atp9a | 0.445074818 | 0.165555505 |
| Lrtm1 | 1.184968622 | 0.165568955 |
| Oscp1 | 0.543236908 | 0.165568955 |
| Hnmpll | -0.302005631 | 0.165611476 |
| Zbtb7a | -0.496531058 | 0.165611476 |
| Ltbp1 | 0.756629939 | 0.165632826 |
| Cacna1d | -0.653697844 | 0.165632826 |
| Chia1 | -4.59734786 | 0.165632826 |
| Efemp1 | 2.192492539 | 0.1657029 |
| Klhl21 | -0.954132828 | 0.1657029 |
| Foxk2 | 0.855755345 | 0.165726729 |
| Ruvbl2 | 0.373078846 | 0.165726729 |
| Aprt | -0.391957535 | 0.165837293 |
| Bank1 | -1.529910757 | 0.165846062 |
| St6galnac1 | -5.328349702 | 0.165846062 |
| Gdpd2 | 0.953526166 | 0.166003042 |
| 0610009L18Rik | 1.012145291 | 0.16608429 |
| Il27 | -4.152789135 | 0.16608429 |
| Heatr9 | 4.857227238 | 0.166103123 |
| Matn4 | 4.406885113 | 0.166257407 |
| Hmgcl | -0.416985564 | 0.166257518 |
| Bmp8a | -4.332189504 | 0.166257518 |
| Ctsh | -0.564450798 | 0.166264693 |
| Cpt1a | 0.585950744 | 0.166320069 |
| Hs6st2 | -0.517936236 | 0.166767425 |
| Ism2 | 4.233757623 | 0.166826654 |
| Bdkrb1 | 3.896362458 | 0.166829137 |
| Gart | -0.39496569 | 0.166906654 |
| Dkc1 | 0.362852733 | 0.167085978 |
| Prdx1 | -0.31555104 | 0.167152459 |
| Gm12355 | 0.587567053 | 0.167309434 |
| Tmem248 | -0.441724078 | 0.167512462 |
| Rad23a | -0.403510374 | 0.16774775 |
| Dnajc4 | 0.761692261 | 0.167893596 |
| 2700081O15Rik | 0.42026383 | 0.167920912 |
| Hectd3 | -0.509812881 | 0.167920912 |
| Herc1 | -0.342180583 | 0.167987235 |
| Adra2a | 1.826007039 | 0.168007788 |
| Klf12 | 0.623231138 | 0.168020424 |
| Hat1 | -0.347918209 | 0.168020424 |
| Tmem47 | 0.550232483 | 0.168387013 |
| Tmc4 | -2.322308038 | 0.168547776 |
| Clhc1 | 4.365448004 | 0.16858061 |
| Hist2h2bb | 1.364231401 | 0.168801051 |
| 4931409K22Rik | 4.44847047 | 0.168882915 |
| Oxct1 | 0.346018595 | 0.168954483 |
| Gnb2l1 | 0.333518698 | 0.168968769 |
| Adamts2 | -1.133609319 | 0.169042644 |
| Gm10226 | -1.75806125 | 0.169095173 |
| Tfec | -1.287904832 | 0.169127565 |
| Eif4g2 | -0.287706112 | 0.169144565 |
| Vmn2r86 | 4.487543961 | 0.169163836 |
| Crb2 | 0.907663915 | 0.169181602 |
| Aim1l | -1.930469644 | 0.169248625 |
| Ercc8 | 0.736286026 | 0.169288185 |
| Osgep | 0.395339029 | 0.169319262 |
| Fam129a | -1.493553772 | 0.169319262 |
| Chil3 | -1.637533952 | 0.169646823 |
| Exph5 | -2.283777392 | 0.169734493 |
| Mob3c | -1.18245901 | 0.169814214 |
| Fam8a1 | -0.538323028 | 0.169834043 |
| Phldb3 | -3.77281245 | 0.169838095 |
| Rlf | -0.400269543 | 0.16989983 |
| Klrb1c | 4.488554394 | 0.169969412 |
| Pacrgl | 0.681379176 | 0.170013043 |
| B4gat1 | -0.43787785 | 0.170029152 |
| Bivm | -0.491049249 | 0.170061685 |
| Cd63 | 0.390270659 | 0.170063693 |
| Stard13 | 2.5083004 | 0.170201502 |
| Trappc6b | 0.391639355 | 0.170214573 |
| Fam122a | -0.509614759 | 0.170214573 |
| Tspear | -5.314780683 | 0.170214573 |
| Dio2 | 3.161806238 | 0.170240408 |
| Arhgef7 | -0.374752678 | 0.170240408 |
| Pacrg | 1.464636472 | 0.170359584 |
| Dlx1 | 0.431619907 | 0.170361877 |
| Akr1b8 | 1.881139812 | 0.170370792 |
| Ube2f | -0.413592296 | 0.170435658 |
| Abcc12 | 4.64641568 | 0.170568585 |
| Nudt11 | 0.611242856 | 0.170568585 |
| Adgra2 | 1.955429794 | 0.170616693 |
| Gm21093 | 0.724049926 | 0.170700518 |
| Mlycd | -0.700288728 | 0.170774071 |
| Pfkfb1 | -1.054301389 | 0.170789778 |
| Cox6a2 | -2.884944642 | 0.170878883 |
| Dmc1 | 3.965516052 | 0.171015339 |
| Serpini1 | -0.502648154 | 0.171015339 |
| Oc90 | 4.882365834 | 0.17102043 |
| Fxyd5 | -1.179317961 | 0.171139781 |
| Sfmbt2 | -1.159082196 | 0.171207391 |
| Zbtb7b | -1.461777055 | 0.171239438 |
| Gtf2e2 | 0.31908792 | 0.171355849 |
| Ciapin1 | 0.352712703 | 0.171445049 |
| Eif2ak2 | -0.503385886 | 0.171445049 |
| C130060K24Rik | 3.940434667 | 0.171655072 |
| Sh3yl1 | 0.504298299 | 0.171754639 |
| Figf | -5.310939723 | 0.171937292 |
| Pphln1 | -0.328799493 | 0.172055462 |
| Commd4 | 0.308974521 | 0.172072428 |
| Kcnj9 | 2.387295925 | 0.172081592 |
| Hepacam | 2.091321609 | 0.172124604 |
| Arfgef3 | -0.59056264 | 0.172124604 |
| 9130019O22Rik | -1.562613474 | 0.172203627 |
| Med18 | -0.783573876 | 0.17230664 |
| Acta2 | 1.766360333 | 0.172329351 |
| Gm10053 | 0.413903593 | 0.172329351 |
| Map2k6 | -0.459402959 | 0.172387035 |
| Trim37 | 0.358836405 | 0.172390421 |
| Brix1 | 0.384020664 | 0.172573833 |
| Gm340 | -0.654790194 | 0.172573833 |

| | | |
|---|---|---|
| Rab33b | -0.27341606 | 0.514130584 |
| Twist1 | 1.28626154 | 0.51414114 |
| Fgf7 | 1.29923308 | 0.514473108 |
| Ddx47 | -0.216597953 | 0.514473108 |
| Rint1 | -0.250526355 | 0.514473108 |
| Gps2 | 0.222252096 | 0.514726746 |
| Neurod2 | -0.286939179 | 0.514750418 |
| Ppp1r12b | -0.262284213 | 0.515235457 |
| Mnd1 | -0.674352149 | 0.51538054 |
| St6galnac4 | -0.232796959 | 0.515497429 |
| Zfat | -0.492694643 | 0.515710288 |
| Spata45 | 3.412705864 | 0.515872537 |
| Wdr17 | -0.399491271 | 0.515872537 |
| Cers1 | 0.420642022 | 0.516681309 |
| Spag6 | 3.163137578 | 0.51681303 |
| Olfm2 | -0.310827284 | 0.517099687 |
| Pdgfb | -0.764124931 | 0.517099687 |
| Icam5 | 0.865378152 | 0.517167631 |
| Ermap | 0.799320238 | 0.517167631 |
| Lpcat4 | 0.601189825 | 0.517252883 |
| Rgag1 | 3.158038135 | 0.51734683 |
| Ablim2 | 0.612251568 | 0.51734683 |
| Alk | -1.026630477 | 0.51734683 |
| Frk | 3.448031374 | 0.517500655 |
| Magohb | 0.266189107 | 0.517525773 |
| Gm10964 | -1.855853242 | 0.517525773 |
| Slc25a14 | 0.314628627 | 0.517564984 |
| Bloc1s5 | -0.376747406 | 0.517564984 |
| Mdfi | 0.877900988 | 0.517689132 |
| Iglon5 | -0.624513138 | 0.51782821 |
| Cyp2j9 | 1.354536374 | 0.517922173 |
| Tjp3 | 0.82855984 | 0.517922173 |
| Thoc1 | 0.185395974 | 0.517922173 |
| Polk | 0.71048458 | 0.518413427 |
| Rmdn2 | -0.61023922 | 0.518413427 |
| Phlda1 | 0.464751951 | 0.518600087 |
| Slc30a1 | -0.438188956 | 0.518600087 |
| Tomt | -0.76206797 | 0.518600087 |
| Desi2 | 0.273714946 | 0.518734109 |
| Lurap1 | -0.572030345 | 0.518734109 |
| Ifi204 | -1.02628892 | 0.518734109 |
| Ubr4 | -0.172509424 | 0.51874951 |
| Rasip1 | 0.404035207 | 0.519008023 |
| Rpa2 | 0.302648663 | 0.519008023 |
| Srrm2 | 0.134875408 | 0.519035857 |
| Gja6 | 0.869458544 | 0.519489944 |
| Arhgap19 | 0.299208112 | 0.519489944 |
| Tab2 | 0.197577805 | 0.51962687 |
| Gm3373 | 1.101605844 | 0.51963083 |
| Narf | -0.207608935 | 0.519790037 |
| Tcf15 | -1.3064446 | 0.519874098 |
| Zfp9 | -0.554158674 | 0.520059243 |
| Usp30 | -0.354862758 | 0.520147947 |
| Lrrc43 | -2.305251131 | 0.520227732 |
| Clnk | 3.146330397 | 0.520292945 |
| Depdc1b | 0.658998256 | 0.520295039 |
| Tbc1d30 | -0.419245604 | 0.520295039 |
| Ankrd28 | -0.317078673 | 0.520571638 |
| Ankrd46 | 0.203202001 | 0.520597331 |
| Ankdd1b | -0.617922596 | 0.520650208 |
| Cdc37l1 | 0.260498281 | 0.520741944 |
| Btrc | -0.215536843 | 0.520741944 |
| Fam208b | 0.272035968 | 0.520858843 |
| Mrpl1 | 0.296563438 | 0.520889426 |
| Ace | 1.372470876 | 0.521398592 |
| Rasl10b | -0.462362658 | 0.52141706 |
| Gm266 | -1.885706344 | 0.521507484 |
| Pnliprp1 | 2.461531541 | 0.521563085 |
| Vldlr | 0.412333891 | 0.521563085 |
| Dgkb | 0.489244232 | 0.521876074 |
| Mlst8 | 0.283836795 | 0.521876074 |
| 6430571L13Rik | 0.927717668 | 0.521992781 |
| Typ23a | -0.390734464 | 0.521992781 |
| Cnst | -0.290528373 | 0.522014167 |
| Dnah14 | 1.162786118 | 0.522022367 |
| Slc9b1 | -2.817343331 | 0.522190764 |
| Camk2g | 0.177157413 | 0.522333212 |
| Yipf3 | 0.186441034 | 0.522350914 |
| Ubr1 | -0.241137715 | 0.522350914 |
| Gm10130 | -0.912298908 | 0.522362415 |
| Snmp70 | 0.162727071 | 0.522396676 |
| Rybp | 0.382571766 | 0.522451466 |
| Adcy5 | 0.494440611 | 0.522694047 |
| Hoxc6 | 3.606230313 | 0.52281645 |
| Srsf6 | -0.145809845 | 0.52281645 |
| Havcr2 | -0.461374101 | 0.52281645 |
| Kif18b | 0.507830155 | 0.523001813 |
| Pum3 | -0.210238986 | 0.523094359 |
| Wdr36 | -0.21440344 | 0.523221323 |
| Fam13a | 0.529539813 | 0.523344886 |
| Taf1 | 0.20823058 | 0.523344886 |
| Tle4 | -0.269375981 | 0.523344886 |
| Slc4a3 | 0.272756244 | 0.523443291 |
| Zfp286 | 0.32623648 | 0.523452662 |
| Prkcz | 0.226636139 | 0.523510788 |
| Zdhhc18 | -0.448297555 | 0.523510788 |
| Zfp763 | -0.334494346 | 0.523792804 |
| Zfp553 | 0.201729898 | 0.52389243 |
| Eya1 | 0.657656964 | 0.523907305 |
| Elp3 | 0.262158145 | 0.523913177 |
| Golga7b | 0.193488502 | 0.523913177 |
| Gsc2 | 3.594985812 | 0.524142972 |
| Rhox7a | 3.594985812 | 0.524142972 |
| Vmn2r85 | 3.594985812 | 0.524142972 |
| Akr1b10 | -0.406011094 | 0.524206925 |
| Stx7 | -0.157589498 | 0.524394053 |
| Rimbp2 | 0.670420717 | 0.524424105 |
| Epha8 | 1.26486309 | 0.524672981 |
| Lman1l | 3.1037786 | 0.524746788 |
| Procr | 1.336779864 | 0.524746788 |
| Tor1aip2 | -0.23208918 | 0.524984462 |
| Lrrn4 | 3.392776526 | 0.525204151 |
| Hsf1 | -0.221665131 | 0.525537783 |
| Timm10b | 0.346523081 | 0.525544987 |
| Gm6588 | -2.118820942 | 0.525567826 |
| Zfp512 | 0.214349278 | 0.525591336 |
| Kifc2 | 0.556746717 | 0.525893394 |
| Senp8 | 0.51297489 | 0.525934665 |
| Armc10 | 0.198173329 | 0.525941565 |

| | | |
|---|---|---|
| Mtss1 | 0.045454385 | 0.905075392 |
| Senp7 | -0.048999849 | 0.905145392 |
| Dhtkd1 | -0.136885463 | 0.905145392 |
| Ola1 | 0.037864236 | 0.905172612 |
| Pyurf | 0.058869981 | 0.905296326 |
| Yipf4 | 0.040496287 | 0.905393248 |
| Srp54a | 0.187768101 | 0.905677338 |
| Gjd3 | -0.438595053 | 0.905874349 |
| Hist1h4k | 0.362668921 | 0.906033756 |
| Peo1 | -0.074244843 | 0.906033756 |
| Gm10778 | -0.290466582 | 0.906247478 |
| Nrf1 | 0.048532901 | 0.906403343 |
| Atp2a1 | 0.753936124 | 0.906405732 |
| Wdr38 | 0.311321319 | 0.906405732 |
| Chaf1b | -0.063119176 | 0.906405732 |
| Kbtbd8 | -0.296390398 | 0.906405732 |
| Gpr183 | 0.105356838 | 0.906701651 |
| Gm13225 | -0.162810882 | 0.90675989 |
| Col28a1 | -0.339922462 | 0.906801111 |
| Dedd | 0.16064546 | 0.907110649 |
| Zfp367 | -0.055821413 | 0.907110649 |
| Paqr3 | 0.06459833 | 0.907355927 |
| Immp2l | -0.098267823 | 0.907355927 |
| Mettl7a1 | 0.129668284 | 0.907452334 |
| Gfra4 | 0.069629151 | 0.907505495 |
| Tmem255a | -0.217150947 | 0.907505495 |
| Sap30 | -0.037955807 | 0.907511715 |
| Gtf3c4 | 0.056528465 | 0.907514549 |
| Zswim3 | 0.113274865 | 0.908258861 |
| Adamts20 | 0.162687289 | 0.908338439 |
| Tmem259 | 0.086971899 | 0.908338439 |
| Parp1 | 0.030780775 | 0.908338439 |
| Klf1 | -0.287791524 | 0.908338439 |
| Rin2 | -0.061107113 | 0.908432707 |
| Onecut3 | 0.287510347 | 0.908503682 |
| Krr1 | 0.057242672 | 0.9085231 |
| Pak7 | -0.049546478 | 0.9085231 |
| Hhip | 0.255477095 | 0.908587623 |
| Ammecr1l | 0.044577873 | 0.908731795 |
| Metrn | 0.071620117 | 0.908842188 |
| Ttc30b | 0.076279029 | 0.908903579 |
| Rbm41 | -0.105095935 | 0.908903579 |
| Dagla | -0.063764112 | 0.909018335 |
| Emc2 | -0.043722611 | 0.90913975 |
| Gm10638 | 0.402792096 | 0.909142017 |
| Plscr4 | 0.361468562 | 0.909142017 |
| Plppr5 | 0.133881105 | 0.909142017 |
| Mrm1 | 0.091430033 | 0.909142017 |
| Cdipt | 0.039300854 | 0.909142017 |
| Auh | -0.035389879 | 0.909142017 |
| Stk32b | -0.073381379 | 0.909142017 |
| Lbhd1 | -0.103041234 | 0.909142017 |
| Mecom | -0.129656264 | 0.909142017 |
| Cdh6 | -0.175551786 | 0.909142017 |
| Rbm8a2 | -0.370905146 | 0.909201081 |
| Adss | 0.05404013 | 0.909204837 |
| Hnmpk | -0.025651346 | 0.909204837 |
| Scyl1 | -0.049316109 | 0.909204837 |
| AW554918 | -0.085337569 | 0.909204837 |
| Fignl2 | 0.590971718 | 0.909420097 |
| Calml4 | -0.079637783 | 0.909504241 |
| Map7 | -0.214746339 | 0.909535492 |
| Zmym1 | 0.053451058 | 0.909686905 |
| Pard3b | 0.104381069 | 0.909866129 |
| Csnk1g1 | -0.043715651 | 0.909974804 |
| Coasy | 0.048428164 | 0.91013468 |
| Tmem126b | -0.050188505 | 0.91018463 |
| Ccdc24 | -0.543013063 | 0.910206183 |
| Abi3bp | 0.242739815 | 0.910390596 |
| Psmc6 | 0.027609004 | 0.910390596 |
| Zfp574 | -0.041759876 | 0.910390596 |
| Uxs1 | 0.064104167 | 0.910421345 |
| Sfmbt1 | 0.051100402 | 0.910475199 |
| Dgkq | -0.13996104 | 0.910475199 |
| Mccc2 | 0.058029396 | 0.910592625 |
| Ddx39b | 0.029920546 | 0.910592625 |
| Izumo4 | -0.071256533 | 0.910721872 |
| 3632451O06Ri | 0.070598928 | 0.911489016 |
| Slc25a1 | -0.03611357 | 0.911489016 |
| Alx4 | 0.225247592 | 0.911526577 |
| Gm1818 | 0.112126669 | 0.911615059 |
| Tpd52l2 | 0.03458527 | 0.911620373 |
| Lix1 | 0.040322592 | 0.911940718 |
| Nudt6 | -0.230609143 | 0.911940718 |
| Slc43a3 | -0.099457572 | 0.911945955 |
| Txnl4b | -0.04884065 | 0.912147391 |
| Ccdc28a | 0.095393503 | 0.912376468 |
| Eif4a1 | -0.025970006 | 0.912376468 |
| Coil | 0.050449448 | 0.912410653 |
| 2610001J05Rik | -0.037583805 | 0.912410653 |
| Otud7a | 0.126485585 | 0.912442987 |
| Prps1l1 | 0.717723511 | 0.912635785 |
| Trp53inp2 | -0.040408183 | 0.913002724 |
| Bhlhb9 | -0.041020465 | 0.913002724 |
| Aldh9a1 | -0.062235426 | 0.9130486 |
| Hist1h4c | 0.132201106 | 0.913311026 |
| Gosr2 | -0.036153189 | 0.913311026 |
| Prrg3 | -0.070477362 | 0.913311026 |
| Erich5 | -0.533003854 | 0.913311026 |
| Pcgf2 | 0.053432798 | 0.913338494 |
| Zfp759 | -0.149697759 | 0.913338494 |
| Thegl | 0.436725642 | 0.913394104 |
| Fam220a | -0.035395439 | 0.913394104 |
| Cpt2 | -0.065361739 | 0.913394104 |
| S1pr5 | -0.212525843 | 0.913732249 |
| Zfp418 | -0.112652015 | 0.913866044 |
| Gba2 | -0.043167484 | 0.913929148 |
| Trappc6a | -0.048902282 | 0.913929148 |
| Rtfdc1 | 0.035632164 | 0.913981498 |
| Lrrc4b | 0.0779137 | 0.914047561 |
| Cdkn2aipnl | -0.035594795 | 0.914047561 |
| Wnt1 | -0.332724515 | 0.914047561 |
| Mon1b | 0.05877492 | 0.91413836 |
| Lrrc29 | 0.158924077 | 0.914584711 |
| Ikbkg | 0.054970568 | 0.914584711 |
| Ndrg3 | -0.035991605 | 0.914584711 |
| Huwe1 | -0.038185606 | 0.914584711 |
| Lurap1l | -0.077808063 | 0.914584711 |
| Dym | 0.034852516 | 0.914948554 |

| | | |
|---|---|---|
| Itih3 | 5.96789797 | 0.008511471 |
| Mrps26 | 0.728688315 | 0.008512165 |
| Mrpl33 | 0.714197382 | 0.008544431 |
| Il20rb | -4.753478371 | 0.008544431 |
| Prkcg | -2.198900469 | 0.008545764 |
| Zfos1 | 0.912117985 | 0.008551838 |
| Lmo3 | 1.137360895 | 0.008586894 |
| Gipr | -1.831007095 | 0.008614894 |
| Daxx | -0.98531746 | 0.008625729 |
| Atp5d | 0.482973597 | 0.008649283 |
| Vwf | 5.7668548 | 0.008660823 |
| Kank4 | 5.785151079 | 0.008661987 |
| Dixdc1 | 0.991651025 | 0.00866389 |
| Hbq1a | 5.689566159 | 0.008666772 |
| Kbtbd11 | 0.90665381 | 0.008669707 |
| Stfa2 | -3.558596836 | 0.008681112 |
| Dynlt1c | 0.814841883 | 0.008683008 |
| Tcerg1l | 1.749558298 | 0.008745879 |
| Ankib1 | -0.721086427 | 0.008753158 |
| Akap17b | 1.331144355 | 0.008764967 |
| Tnip2 | -1.677296345 | 0.008778842 |
| Rln1 | 3.17488302 | 0.008795233 |
| Nt5m | 0.763970867 | 0.008795233 |
| Smarca2 | 0.746958026 | 0.008795233 |
| Eny2 | 0.546964438 | 0.008821281 |
| Sumf1 | -0.943101636 | 0.008834071 |
| 5031414D18Rik | -2.853452027 | 0.008856613 |
| Slc39a14 | -1.186848709 | 0.008868761 |
| Ift81 | 0.765013132 | 0.008909841 |
| Slc39a4 | -1.790868181 | 0.008916909 |
| Rps6ka2 | -0.954426823 | 0.008918973 |
| Pfdn6 | 0.735558556 | 0.008944022 |
| Depdc5 | -0.834413118 | 0.008953346 |
| Arid1a | 0.524587811 | 0.008973507 |
| Tppp3 | 1.453344465 | 0.008993843 |
| Arglu1 | 0.759928858 | 0.008993843 |
| 2700060E02Rik | 0.520012745 | 0.009005829 |
| Spred2 | -0.850641085 | 0.009027811 |
| C2cd3 | -0.683838201 | 0.009030784 |
| Tmem108 | -1.128476534 | 0.009055953 |
| Tspyl3 | -1.049679806 | 0.009066371 |
| Prmt10 | -0.745328917 | 0.009120661 |
| Pkd1l2 | 5.900956802 | 0.009145801 |
| Klra17 | -5.816052971 | 0.009149276 |
| Nprl2 | 0.854641025 | 0.009172395 |
| Dgkg | -1.7077088 | 0.009187945 |
| Pars2 | 2.377051089 | 0.009200994 |
| Pdgfrl | 1.462848366 | 0.009200994 |
| Per2 | -1.023174648 | 0.009200994 |
| Spocd1 | -6.497250933 | 0.009200994 |
| Ankrd52 | -0.831088454 | 0.009206615 |
| Ms4a4b | -3.069058523 | 0.009207153 |
| Tubgcp2 | -0.544585512 | 0.009219291 |
| Hlx | -1.696368309 | 0.009219291 |
| Stim1 | -0.972813383 | 0.00922324 |
| Eci1 | 1.009233383 | 0.00922795 |
| Tmem164 | -0.829201915 | 0.00922795 |
| Pabpc4l | 1.938546655 | 0.009258761 |
| Parp4 | -1.89377748 | 0.009260434 |
| Ctsc | -2.030757954 | 0.009265882 |
| Adnp | 5.675002594 | 0.009271594 |
| Etv4 | 2.772573551 | 0.009275277 |
| Lgals3 | -2.943572306 | 0.009282185 |
| Gga1 | -0.740710007 | 0.009304287 |
| Polr2k | 0.700072621 | 0.009322621 |
| Trib1 | -0.998601083 | 0.009360777 |
| Cntnap5c | -2.287552654 | 0.009478854 |
| Dpp7 | -1.002043115 | 0.009483711 |
| Snrpc | 0.684637998 | 0.009495245 |
| Fam21 | -0.579498892 | 0.00960148 |
| Aamdc | 0.994508278 | 0.009625047 |
| Shisa3 | -5.05928846 | 0.009654546 |
| Prtn3 | 1.366319777 | 0.009681338 |
| Bmyc | 0.714720529 | 0.009681338 |
| Pvrl2 | -0.734946631 | 0.009681338 |
| Hspbp1 | 0.634237644 | 0.00968424 |
| Tsr3 | 0.735677571 | 0.009691999 |
| Kif1b | 0.526515761 | 0.009694052 |
| Rpl37 | 0.469283162 | 0.009707333 |
| Tmem127 | -0.864679891 | 0.009718225 |
| Mea1 | 0.604261965 | 0.009736706 |
| Amotl1 | 0.852626364 | 0.009761965 |
| Trp53 | 0.569283864 | 0.009769434 |
| Adam19 | -1.001705217 | 0.009771146 |
| Arhgap4 | -1.467953083 | 0.009772789 |
| Gpc3 | 1.9120264 | 0.00979 |
| Wdr75 | 0.997178062 | 0.00981519 |
| Ctnnal1 | 1.093289129 | 0.009815548 |
| Pilrb2 | -2.871101916 | 0.009821747 |
| Pcdhgb2 | 5.70277003 | 0.009829206 |
| Smek1 | -0.643005429 | 0.009829206 |
| Cpa4 | 5.906490927 | 0.009831332 |
| Cpsf1 | -0.622686235 | 0.009831332 |
| Mroh1 | -0.991418807 | 0.009831332 |
| Sema7a | -1.341676433 | 0.009831332 |
| Nup62 | -0.685746867 | 0.009852496 |
| Scn1a | 1.41159629 | 0.009899794 |
| Cfi | -6.470773113 | 0.009899794 |
| Bfsp1 | 4.106127317 | 0.009902817 |
| Arfrp1 | 0.607863182 | 0.009902817 |
| Sema3e | 3.637834516 | 0.00990364 |
| Rab3gap2 | -0.863330419 | 0.009909311 |
| Frat2 | -1.13531557 | 0.009915683 |
| 1190002N15Rik | 1.29499865 | 0.00993074 |
| Zfyve27 | -0.70772929 | 0.00995053 |
| Trim24 | 0.793922253 | 0.009980012 |
| Ndufa4l2 | 2.839869999 | 0.009982289 |
| Arhgap22 | -2.545564856 | 0.009982289 |
| C5ar1 | -2.763686112 | 0.010035595 |
| Osr2 | 5.684176119 | 0.010051571 |
| Cenpo | -1.266625231 | 0.010065122 |
| Sh3rf1 | 0.696206252 | 0.010085446 |
| Nae1 | 0.520472238 | 0.010101779 |
| Spata17 | -6.477454419 | 0.010110793 |
| Tgif1 | -1.18612538 | 0.010113224 |
| Prmt2 | 0.57699021 | 0.010115143 |
| Pold1 | -0.804416154 | 0.010121591 |
| Pxn | -1.074225563 | 0.010127506 |
| Fam120a | 0.76520747 | 0.010144525 |

| | | |
|---|---|---|
| Zbtb42 | 2.861934694 | 0.172653657 |
| Slc15a2 | 0.856401767 | 0.172653657 |
| Sertad2 | 0.403468934 | 0.172653657 |
| Ly6e | -0.530702275 | 0.172653657 |
| Siglecg | -5.326925292 | 0.172771521 |
| 2610305D13Rik | -1.191013193 | 0.17279918 |
| Kdelr2 | 0.344239945 | 0.172826394 |
| Them4 | 0.458481436 | 0.172920369 |
| Efnb1 | 0.463725828 | 0.172982249 |
| Gtf3c1 | -0.481927842 | 0.17302034 |
| Aanat | -5.304203271 | 0.173035095 |
| Dhrs9 | -5.304203271 | 0.173035095 |
| Mmp25 | -1.894543995 | 0.173067001 |
| Amd1 | 0.423532157 | 0.173092264 |
| St6gal1 | 0.643813163 | 0.173139057 |
| 1700021F05Rik | 0.398680345 | 0.173139057 |
| Amelx | 4.747293287 | 0.17325288 |
| Rnf214 | 0.367839252 | 0.173326689 |
| Zfp108 | -2.358677701 | 0.173326689 |
| Htra1 | 0.852044781 | 0.173402738 |
| Mul1 | -0.474767436 | 0.173422659 |
| Ptch1 | 0.64526882 | 0.173432706 |
| Zbtb39 | 0.766746464 | 0.17383588 |
| Tatdn1 | 0.645070879 | 0.173836884 |
| Trpm6 | 2.083675416 | 0.173933 |
| Gck | 1.396756742 | 0.173933 |
| Zfp595 | -2.456578018 | 0.173933 |
| Prelid2 | -0.894081503 | 0.173980655 |
| Ascc2 | -0.540044219 | 0.174404755 |
| Epn3 | 2.317146771 | 0.174447701 |
| Stox1 | 1.196277559 | 0.174515103 |
| Vps13d | -0.460551876 | 0.174573454 |
| Lipt2 | 0.730115087 | 0.174633712 |
| Hsf4 | 4.482197173 | 0.17474128 |
| Gm4924 | 1.652657708 | 0.17474128 |
| Mum1l1 | 1.066158179 | 0.17474128 |
| Gtpbp8 | 0.580072786 | 0.17474128 |
| 4-Mar | -0.682063673 | 0.17474128 |
| Fcgr2b | -1.718119346 | 0.17474128 |
| Ldlrad3 | 0.483380104 | 0.17487817 |
| Opn3 | 1.501585714 | 0.174957095 |
| Vamp1 | -0.864363565 | 0.175085546 |
| Fn3krp | 0.559660175 | 0.175119401 |
| Zbtb22 | 0.762160815 | 0.175128308 |
| Otud3 | -0.98549076 | 0.175316346 |
| D17Wsu92e | -0.45629741 | 0.175618554 |
| Apol9a | -3.793989786 | 0.175618554 |
| Rasgef1a | 1.89343373 | 0.175725681 |
| Ccz1 | -0.440655658 | 0.175725681 |
| Akt3 | -0.512514871 | 0.175754975 |
| Tnfaip8l1 | 0.951885501 | 0.175883525 |
| Mri1 | 0.710221782 | 0.175883525 |
| Fanci | -1.465089573 | 0.176055776 |
| Fam126a | -0.406076185 | 0.176081942 |
| Arrdc3 | -0.35286545 | 0.176348963 |
| Bicd2 | -0.427930122 | 0.176359985 |
| Clec14a | 4.781272633 | 0.17647295 |
| 2510002D24Rik | 0.545763307 | 0.17647884 |
| Nox4 | 1.104020348 | 0.176533101 |
| Caskin2 | 0.826355248 | 0.176635558 |
| Pla2g4b | -3.423821558 | 0.176838742 |
| Chchd3 | 0.363909783 | 0.176860339 |
| Zbtb8a | 0.71233581 | 0.176884357 |
| Zfp938 | 0.57987648 | 0.176946345 |
| Hist1h2al | -0.477602933 | 0.1771146 |
| Rtcb | -0.354305543 | 0.177333895 |
| Edn2 | -5.34945321 | 0.177374873 |
| Hepacam2 | 1.430456853 | 0.177509218 |
| Ndufaf5 | 0.508289213 | 0.177509218 |
| Cend1 | 0.539377841 | 0.17753039 |
| Hjurp | -0.424173562 | 0.177772714 |
| A730049H05Rik | 4.313034433 | 0.177790818 |
| Pcsk4 | 1.601150273 | 0.177790818 |
| Cox18 | 0.450919734 | 0.177790818 |
| Nol12 | 0.446409925 | 0.177790818 |
| Ube2k | 0.279243602 | 0.177790818 |
| Serinc1 | -0.362814898 | 0.177790818 |
| Numb | -0.517723052 | 0.177790818 |
| Aldh3a2 | -0.557098126 | 0.177790818 |
| Nup205 | -0.567395872 | 0.177790818 |
| Ppargc1b | -1.598917361 | 0.177790818 |
| Lca5 | 0.53741556 | 0.177812389 |
| Nagk | -0.332708457 | 0.177812389 |
| Ankrd17 | 0.359435119 | 0.177966154 |
| Hspa12b | 1.441106461 | 0.177976264 |
| Ppp1r12a | 0.349774528 | 0.177976264 |
| Lrp6 | -0.498946403 | 0.178100129 |
| Mtmr10 | -0.836638282 | 0.178117265 |
| Nek7 | -0.627753981 | 0.178241788 |
| Napsa | -1.316238987 | 0.17831728 |
| Fam19a3 | -3.263805221 | 0.17831728 |
| Mrps25 | 0.380086689 | 0.178413935 |
| Spin2c | 0.729924651 | 0.178494938 |
| Phf3 | -0.330669114 | 0.178501617 |
| Fcor | 1.010999547 | 0.178542955 |
| Rad9a | -0.610751809 | 0.178550524 |
| Plscr2 | 4.89817342 | 0.178575503 |
| Gpc6 | 0.713153367 | 0.178575503 |
| Qars | 0.497995065 | 0.178575992 |
| Cit | 0.479493227 | 0.178575992 |
| Sepsecs | -0.680059681 | 0.178575992 |
| Efna2 | 0.84883277 | 0.178596764 |
| Rcbtb1 | -0.313185569 | 0.178596764 |
| Cul2 | -0.338562781 | 0.178596764 |
| Ccdc9 | -0.612990501 | 0.178596764 |
| Leng9 | -1.124701788 | 0.178596764 |
| Lipt1 | 1.52053464 | 0.178635466 |
| Zmynd10 | 1.131393756 | 0.178635466 |
| Fbxw17 | 0.570382112 | 0.178635466 |
| Slc16a2 | 0.421626785 | 0.178635466 |
| Rars | -0.295272595 | 0.178635466 |
| Slc27a4 | -0.513681621 | 0.178635466 |
| Avpr1a | 4.470014579 | 0.178772309 |
| Tmem109 | -0.481163862 | 0.178824868 |
| Fam78b | 0.826730222 | 0.178867516 |
| Zhx2 | 0.741736074 | 0.178867516 |
| Pcdhb22 | 1.347033207 | 0.178951188 |
| Senp1 | 0.49388668 | 0.179067514 |
| Tsc22d2 | -0.356818959 | 0.179076222 |

| | | |
|---|---|---|
| 4930563M21Rik | 3.088800306 | 0.526050976 |
| Tmem160 | 0.189803873 | 0.526050976 |
| Rnf43 | 2.260435522 | 0.526158313 |
| Tigar | 0.334732467 | 0.526158313 |
| Tmc7 | 0.398525301 | 0.526311146 |
| Nipsnap3b | -0.153981002 | 0.526311146 |
| Apoa1 | 1.234360657 | 0.526381229 |
| Fbxw9 | 0.223671938 | 0.526381229 |
| Htr2a | 1.312807947 | 0.526526082 |
| Numa1 | -0.181767808 | 0.526580941 |
| Heatr5a | -0.380562635 | 0.526612189 |
| Treml4 | -0.702672777 | 0.526612189 |
| Fdps | 0.499016634 | 0.5269519 |
| Fgf15 | 0.979420817 | 0.527060039 |
| Gm20498 | 0.915200354 | 0.527362661 |
| Ret | 1.106775975 | 0.527708717 |
| Efr3b | 0.225339618 | 0.527755843 |
| Ift140 | -0.351541951 | 0.527755843 |
| Pim3 | -0.186263774 | 0.527760539 |
| Krt24 | 3.118638158 | 0.527803186 |
| Ankrd39 | -0.600500009 | 0.527809188 |
| Cdh13 | -0.591132882 | 0.527931485 |
| Ankrd54 | 0.216312313 | 0.52813757 |
| Klhl20 | -0.316319244 | 0.528321962 |
| AW011738 | 0.667911286 | 0.528464271 |
| Adhfe1 | -0.608458657 | 0.528632932 |
| Msi2 | 0.215331957 | 0.52875731 |
| 1700011E24Rik | -1.837070686 | 0.52894974 |
| Myg1 | -0.212112276 | 0.529016725 |
| Brsk2 | -0.25342879 | 0.529246928 |
| Cc2d2a | 0.420665162 | 0.529271972 |
| Lemd1 | 0.351384577 | 0.529271972 |
| Ddx24 | 0.173070192 | 0.529271972 |
| Golga7 | 0.162482602 | 0.529271972 |
| Fam98a | -0.204799906 | 0.529271972 |
| Map3k2 | 0.293115275 | 0.529386164 |
| Cops7b | 0.251492158 | 0.529423683 |
| 4930432K21Rik | -0.842677204 | 0.529423683 |
| Ildr2 | 0.765114482 | 0.530025709 |
| Cdk5r1 | -0.249625903 | 0.530133276 |
| Wdr63 | 1.59354336 | 0.530213106 |
| Foxp4 | -0.269070495 | 0.530213106 |
| Smg6 | -0.19809137 | 0.530553325 |
| AW146154 | 0.55460829 | 0.530865519 |
| Ncapd2 | -0.284120308 | 0.530865519 |
| Samd12 | 0.470510607 | 0.530871228 |
| Nat2 | -1.195413579 | 0.530871228 |
| Slc24a2 | 0.793942436 | 0.531340955 |
| Gls | 0.236721379 | 0.531345929 |
| Kptn | 0.228985147 | 0.531345929 |
| Sec23a | -0.180350847 | 0.531471396 |
| Lpp | -0.251036836 | 0.531471396 |
| Kctd19 | -1.068650062 | 0.531471396 |
| Abcb4 | -2.143596373 | 0.531471396 |
| Commd10 | 0.192840671 | 0.531753376 |
| Iws1 | 0.170482541 | 0.531788252 |
| Med12l | 0.332897655 | 0.531831612 |
| Abhd3 | 3.534975214 | 0.53204209 |
| Erp27 | 3.534975214 | 0.53204209 |
| Gfra3 | 3.534975214 | 0.53204209 |
| Zpbp | 1.089751306 | 0.53204209 |
| 1110008L16Rik | 0.453589831 | 0.53204209 |
| Rgs3 | 0.340382508 | 0.53204209 |
| Prosc | 0.277738204 | 0.53204209 |
| Rgs10 | 0.216205808 | 0.53204209 |
| Tmem63b | -0.235224495 | 0.53204209 |
| Diaph1 | -0.269667041 | 0.53204209 |
| Uck1 | -0.253945535 | 0.532137175 |
| 4930524J08Rik | 1.050380827 | 0.532256268 |
| Pkd2 | 0.324037044 | 0.532256268 |
| Ocln | -0.953330875 | 0.532256268 |
| Kat5 | -0.253126216 | 0.532443799 |
| Prrt3 | -0.94396215 | 0.532678752 |
| Ttc38 | 0.646382915 | 0.532682799 |
| Nt5c3 | 0.193627177 | 0.532682799 |
| Zfp606 | 0.227057892 | 0.532832627 |
| Lrpprc | 0.198927912 | 0.532840241 |
| Scn3b | -0.243578106 | 0.532938729 |
| Rbmx | 0.159420003 | 0.533124343 |
| Dnaja3 | 0.311563963 | 0.533135942 |
| Sf3a1 | -0.164426227 | 0.533308312 |
| H2-T23 | -0.557514152 | 0.533397037 |
| Robo4 | 3.382708289 | 0.533487341 |
| Ethe1 | -0.213552213 | 0.533487341 |
| Piwil2 | -2.24190172 | 0.533487341 |
| Irs1 | 0.29925706 | 0.533555064 |
| Mansc1 | -0.688365832 | 0.533558851 |
| Fam161b | -0.29143401 | 0.533657525 |
| Krtap17-1 | 3.354918321 | 0.533865974 |
| Trim55 | 3.354918321 | 0.533865974 |
| Lrrd1 | 3.335352673 | 0.533865974 |
| Slc22a17 | 0.201687013 | 0.534094329 |
| Heatr4 | -1.422140543 | 0.534282634 |
| Rhbg | 3.091628183 | 0.534412435 |
| Sirt4 | -0.345962268 | 0.534412435 |
| Cd46 | -0.724306914 | 0.534412435 |
| Megf8 | -0.242748393 | 0.534427873 |
| Ltbp4 | 0.610632718 | 0.534434162 |
| Pycr1 | -0.231312794 | 0.534434162 |
| Rnls | 0.34698289 | 0.534548744 |
| Dennd6b | 0.37045079 | 0.534561724 |
| Dph1 | 0.511223801 | 0.534697289 |
| Ube2j2 | -0.168588381 | 0.535137709 |
| Gm14434 | 1.5756324 | 0.535289346 |
| Ccl1 | 3.347277773 | 0.535342014 |
| Leo1 | 0.220925327 | 0.535466021 |
| Ccdc62 | 1.282446085 | 0.535643423 |
| Kcnj5 | 1.424845706 | 0.536055351 |
| Prkg1 | 0.856768849 | 0.536900725 |
| Chpt1 | 0.418509297 | 0.536900725 |
| Spata7 | -0.251932791 | 0.536900725 |
| Klk8 | -0.648497792 | 0.536993518 |
| Yaf2 | 0.215561791 | 0.537189629 |
| Maz | 0.183404404 | 0.537189629 |
| Grk1 | 3.10660876 | 0.537520442 |
| Arfip2 | 0.195303115 | 0.537520442 |
| Ddx26b | -0.252467996 | 0.537520442 |
| Tfap4 | -0.581356903 | 0.537520442 |
| Gm9047 | 1.54936345 | 0.537520653 |

| | | |
|---|---|---|
| Trim3 | 0.054948029 | 0.915085195 |
| Bmpr2 | -0.037747941 | 0.915085195 |
| Dynlt3 | -0.038625721 | 0.915085195 |
| Fam171b | -0.053051866 | 0.915170799 |
| Il12b | -0.297327631 | 0.915450568 |
| Rnft2 | 0.076131285 | 0.915611939 |
| Ppp2r2d | 0.039124546 | 0.915611939 |
| Nhlh2 | -0.088449404 | 0.915611939 |
| Gm9887 | -0.477397438 | 0.915611939 |
| Mybpc2 | -0.492866477 | 0.915611939 |
| Ropn1l | 0.105743018 | 0.915955058 |
| Ckap2l | -0.048063933 | 0.916021162 |
| Gm20517 | 0.219176299 | 0.91606084 |
| Ivns1abp | -0.033355812 | 0.91606084 |
| Pdcd11 | 0.062247198 | 0.916090688 |
| Eif5a2 | -0.099487434 | 0.916090688 |
| Pank4 | -0.069225231 | 0.916346987 |
| Cpeb1 | 0.263317867 | 0.916368108 |
| Hdac2 | 0.027553866 | 0.916368108 |
| Arpc3 | -0.028736413 | 0.916383842 |
| Dlg3 | -0.073403537 | 0.916383842 |
| Txndc16 | 0.042565076 | 0.916529066 |
| Gufl | -0.046497769 | 0.91657682 |
| Dnah2 | -0.136961937 | 0.91657682 |
| Gm8797 | 0.031257267 | 0.916778298 |
| 4931414P19Ril | 0.116262615 | 0.916878276 |
| Atad3a | 0.038534024 | 0.916878276 |
| Ccdc51 | 0.107828168 | 0.917084256 |
| Btnl2 | -0.209033722 | 0.917084256 |
| Rab37 | -0.234147462 | 0.917084256 |
| Slc52a3 | -0.27679972 | 0.917177377 |
| Prkaa2 | -0.056962934 | 0.917190977 |
| Mief2 | -0.097947303 | 0.917226462 |
| Trp53rkb | 0.065610102 | 0.917259352 |
| Ciita | 0.315203043 | 0.917328971 |
| Ablim1 | 0.059430479 | 0.917392583 |
| Unc93a | 0.539452741 | 0.917855273 |
| AI837181 | 0.062543627 | 0.917855273 |
| Gm10642 | 0.364446816 | 0.917962277 |
| Smad3 | 0.076485145 | 0.917962462 |
| Tgfa | 0.10912118 | 0.918009191 |
| Itgav | -0.064816762 | 0.918113312 |
| Selenbp2 | 0.241567101 | 0.918769423 |
| Gm19345 | 0.236175706 | 0.918769423 |
| Rps6kl1 | -0.076481294 | 0.918769423 |
| Zdhhc16 | -0.057126256 | 0.918797659 |
| Slc35f4 | 0.132695663 | 0.919016054 |
| Jag2 | -0.092540933 | 0.920007182 |
| S100g | -0.347199388 | 0.920197191 |
| Meioc | 0.500561442 | 0.920285943 |
| Otud7b | 0.070172233 | 0.920800042 |
| Kcnj6 | 0.224881207 | 0.920858037 |
| Ddx23 | -0.053890496 | 0.920858037 |
| Setd5 | -0.039167753 | 0.920965985 |
| Mrpl16 | 0.042677332 | 0.920967047 |
| Sart1 | -0.04658513 | 0.921104022 |
| Pcdhga5 | -0.132224556 | 0.921124177 |
| 2900055J20Rik | 0.097617254 | 0.92117219 |
| Fbn2 | 0.076769814 | 0.921644641 |
| Eng | 0.082602295 | 0.92170675 |
| Orm2 | 0.572414328 | 0.921798264 |
| Dcxr | -0.048887886 | 0.921798264 |
| Pcx | -0.081003689 | 0.921798264 |
| Syngr1 | 0.042310822 | 0.921891533 |
| Pak1ip1 | -0.032750649 | 0.921891533 |
| Gm6741 | 0.125378691 | 0.922357126 |
| Rbp7 | 0.508590334 | 0.922569697 |
| Mbd6 | 0.068069958 | 0.922569697 |
| Mrpl3 | 0.032760829 | 0.922580879 |
| Gm11437 | -0.355001924 | 0.922665473 |
| Rpp38 | 0.085068668 | 0.923037657 |
| B3galt4 | 0.177743655 | 0.923202279 |
| Pgm3 | -0.045614059 | 0.923202279 |
| Fxyd4 | -0.108844751 | 0.923202279 |
| Ildr1 | 0.313648262 | 0.923246316 |
| Ap3b2 | 0.046443813 | 0.923246316 |
| Aamp | 0.033107116 | 0.923246316 |
| Ccl19 | 0.351295158 | 0.923297024 |
| Kcng3 | 0.267947417 | 0.923297024 |
| Tomm40l | 0.075861525 | 0.923297024 |
| Ralyl | 0.044520452 | 0.923297024 |
| Map4k3 | -0.036260864 | 0.923297024 |
| Rad51b | -0.236240684 | 0.923297024 |
| Sfxn2 | 0.064101223 | 0.923390571 |
| Wdr24 | 0.051014998 | 0.923404404 |
| Aimp2 | -0.054806071 | 0.923516732 |
| Ttc12 | 0.168129185 | 0.923540314 |
| Pkn2 | -0.038566508 | 0.923865313 |
| Slc26a6 | -0.133981085 | 0.923867606 |
| Zkscan16 | 0.252204425 | 0.923876165 |
| Prkag1 | -0.036662015 | 0.923876165 |
| Fndc9 | 0.117613039 | 0.923919075 |
| Psmd12 | 0.025557842 | 0.924006163 |
| Saa1 | 0.136409954 | 0.924256405 |
| Panx1 | -0.039704134 | 0.924256405 |
| Kcnd1 | -0.108862466 | 0.924256405 |
| Ywhaz | -0.023387713 | 0.92433574 |
| Hnrnpd | -0.027904556 | 0.9245242 |
| Ttc39d | -0.437401273 | 0.9245242 |
| Lace1 | 0.129786412 | 0.924714241 |
| Yipf2 | 0.059000272 | 0.924714241 |
| Bud13 | 0.054847843 | 0.924714241 |
| Mtx2 | 0.030820657 | 0.924714241 |
| Gfra2 | -0.106051039 | 0.924714241 |
| Hist1h2br | -0.278377414 | 0.924714241 |
| Synj2 | 0.163690031 | 0.924725767 |
| Actr8 | 0.035130035 | 0.924856692 |
| Ddx4 | 0.311716643 | 0.925127427 |
| Mon1a | 0.052517731 | 0.925251622 |
| B4galt7 | 0.043697711 | 0.926384596 |
| Pcdhga12 | -0.081050371 | 0.926384596 |
| Gpr4 | -0.134971528 | 0.926384596 |
| Tex29 | 0.484911332 | 0.926395372 |
| Arhgap40 | 0.235388563 | 0.926395372 |
| Klhl34 | 0.214527474 | 0.926395372 |
| Wnt9a | 0.198325903 | 0.926395372 |
| Igdcc3 | 0.085309738 | 0.926395372 |
| Ccbl2 | 0.078194857 | 0.926395372 |
| Ttf2 | 0.054575608 | 0.926395372 |

| | | |
|---|---|---|
| Sh3bp5 | -0.592942729 | 0.010148461 |
| Ubald1 | -0.56768494 | 0.010157514 |
| Prelp | 2.701289525 | 0.010164088 |
| Rilpl1 | 1.342295803 | 0.010164088 |
| Trmt10b | 1.008342351 | 0.010164088 |
| Ngp | -3.902846583 | 0.010164088 |
| Atf5 | 1.889282638 | 0.010170373 |
| BC017643 | -0.809253274 | 0.010177967 |
| Ttc26 | 1.633722577 | 0.01019299 |
| Mettl23 | 0.754951552 | 0.01019299 |
| Ankrd61 | -1.818108226 | 0.01020261 |
| Pskh1 | -1.024045268 | 0.010207646 |
| Slc25a36 | -0.53532953 | 0.010254843 |
| Nrk | 5.648336281 | 0.010255696 |
| Degs1 | -0.790164138 | 0.010264939 |
| Lrfn5 | -1.194540278 | 0.010264939 |
| Cox19 | 0.687042528 | 0.010272544 |
| Agap2 | -1.280995581 | 0.010292218 |
| Mgst3 | 0.600328047 | 0.010299032 |
| Ssc5d | 5.644199462 | 0.010305394 |
| Rps24 | 0.473248361 | 0.010310629 |
| Insr | -0.807376924 | 0.010331935 |
| Ptges | -3.278249842 | 0.01034675 |
| Caml | 0.618812921 | 0.010355205 |
| Nup85 | 0.748227971 | 0.010366934 |
| Pmpca | -0.572640338 | 0.010368952 |
| Selm | 1.071278996 | 0.010404735 |
| Gm10269 | 0.820876052 | 0.010404735 |
| Gfra1 | 1.746497055 | 0.010423542 |
| Luzp2 | 1.214322799 | 0.0104294 |
| Sri | -0.624826764 | 0.010457576 |
| Zfp955a | 0.935947525 | 0.010462131 |
| AK010878 | 0.860181773 | 0.01047273 |
| Commd6 | 0.564978017 | 0.01047273 |
| Hmcn2 | -2.08009907 | 0.010479404 |
| Dtd2 | 1.01694329 | 0.010482787 |
| Gm4955 | 5.051550176 | 0.010484356 |
| Asns | -0.851540507 | 0.010496673 |
| Tpgs1 | 0.81383318 | 0.010497742 |
| Zfp319 | -0.734780455 | 0.010499293 |
| Fam161a | 2.28723541 | 0.010500623 |
| Rnf213 | -1.368184552 | 0.010513202 |
| Sec23b | -0.637314483 | 0.010516625 |
| Mmp19 | -2.380375483 | 0.010516625 |
| Cyth2 | 0.487029546 | 0.010525134 |
| Sqrdl | -0.873074307 | 0.010525134 |
| Siah3 | 2.077704211 | 0.010526414 |
| Stk33 | 2.186315054 | 0.0105428 |
| Nsun2 | -0.633307677 | 0.010563743 |
| Ctdsp2 | 0.74596628 | 0.010599829 |
| Dera | -0.722650063 | 0.010602037 |
| Brs3 | -6.440187407 | 0.010625669 |
| Rgs20 | 1.633820657 | 0.01062695 |
| Resp18 | 1.552327738 | 0.01062695 |
| Zfp560 | 0.738838462 | 0.01062695 |
| Hcfc1r1 | 0.636157418 | 0.01062695 |
| Mgat4a | -0.715891342 | 0.01062695 |
| Nuak1 | -0.784381024 | 0.01062695 |
| Akap13 | -0.854864595 | 0.01062695 |
| Ubap2l | -0.45096422 | 0.010634333 |
| Mrpl52 | 0.590382313 | 0.010662903 |
| Ldhb | 0.625549898 | 0.010665522 |
| Papolb | 6.219828331 | 0.010680843 |
| Dhrs13 | 1.137958231 | 0.010722052 |
| Nipal1 | -6.263115851 | 0.010754462 |
| Klrb1f | -6.770613985 | 0.010754462 |
| Rpp30 | 0.697555943 | 0.010779734 |
| Lhx2 | 0.789568643 | 0.010790618 |
| Btbd18 | 5.634228036 | 0.010803887 |
| Rps23 | 0.49019024 | 0.010803887 |
| Ppp1r11 | 0.660629487 | 0.010812665 |
| Pde8a | -2.147888775 | 0.010812665 |
| Pde3a | 1.790920022 | 0.010829265 |
| A2m | 2.543240743 | 0.010831845 |
| Ccr5 | -1.124560323 | 0.010850456 |
| Fam134c | -0.665606988 | 0.010854314 |
| Ift22 | 0.768256367 | 0.010881875 |
| Pard6a | 0.821842728 | 0.010882408 |
| Kdelr3 | 2.075318733 | 0.01088566 |
| Add1 | -0.463558751 | 0.010897549 |
| Zdhhc4 | 0.818441725 | 0.010902285 |
| Pramef8 | -1.050464734 | 0.010930508 |
| Has2 | 3.18658603 | 0.010959693 |
| Mocs2 | 0.577919259 | 0.010959693 |
| Hspb3 | 5.789288956 | 0.010960081 |
| Cdk11b | 0.601212009 | 0.010960081 |
| Trem1 | -4.43021017 | 0.010960081 |
| Tmod3 | -0.614274612 | 0.010964892 |
| Cln8 | -0.698120787 | 0.010969193 |
| Ticam2 | -1.941382611 | 0.010969193 |
| Rap1gap2 | -1.411068466 | 0.011011582 |
| Svbp | 0.566110391 | 0.011031607 |
| Ctgf | 2.772480369 | 0.011045774 |
| Echdc2 | 2.207911165 | 0.011049221 |
| Dhrs4 | 0.807676236 | 0.011049221 |
| Tsfm | 0.723632328 | 0.011049221 |
| Fam43b | -1.960876432 | 0.011049221 |
| Pax2 | 6.008553889 | 0.011051027 |
| Slc9a1 | -0.999584306 | 0.011096038 |
| Gm11273 | 0.682110797 | 0.011096924 |
| Pmm1 | 0.762108975 | 0.011102941 |
| Stmn1 | 0.538437797 | 0.011106234 |
| Ddx60 | -2.398439597 | 0.011119471 |
| Agbl1 | -6.766995508 | 0.011122924 |
| Mrfap1 | 0.483818286 | 0.011143755 |
| Ankrd44 | -0.67508794 | 0.011143755 |
| Jakmip2 | 0.787068554 | 0.011157496 |
| Nat14 | 1.012656296 | 0.011165849 |
| Htatip2 | -1.08918958 | 0.011167737 |
| Sp9 | 1.045558714 | 0.011180784 |
| Dcaf4 | 0.936957338 | 0.01118465 |
| Tob1 | 0.980274732 | 0.011224739 |
| Ptgr2 | -0.868966619 | 0.011224739 |
| Gpr152 | -6.739300984 | 0.011252873 |
| Rps11 | 0.486532133 | 0.011254965 |
| Dyrk4 | 6.285291531 | 0.011264901 |
| Mfap3l | 2.555379091 | 0.011264901 |
| Ubqln1 | -0.509514801 | 0.01127374 |
| Urod | 0.640919056 | 0.011282909 |

| | | |
|---|---|---|
| Timm17a | 0.320563566 | 0.179125642 |
| Slfn8 | -1.415935365 | 0.179125642 |
| Aldh3b2 | -5.257837615 | 0.179125642 |
| Mzb1 | -5.257837615 | 0.179125642 |
| Anxa3 | -1.111712313 | 0.179435024 |
| Trim66 | -2.626919483 | 0.179435024 |
| Scamp3 | -0.452845161 | 0.179527489 |
| Lrfn4 | -0.519960164 | 0.179583629 |
| Ccdc191 | 0.757059779 | 0.179636682 |
| Xrcc6bp1 | 0.890997658 | 0.179693802 |
| Tfap2d | 1.169794154 | 0.179827381 |
| Gm6576 | 1.82687968 | 0.179886358 |
| Papd5 | 0.522355045 | 0.180135786 |
| Timp4 | 1.018193022 | 0.180239086 |
| Yipf6 | -0.496615521 | 0.180534061 |
| Rnf10 | -0.293135152 | 0.180539952 |
| Ush2a | 3.831518057 | 0.180559078 |
| Nlrx1 | -0.579109416 | 0.18059993 |
| Lrcol1 | -5.009547151 | 0.180723131 |
| Gga2 | -0.403789532 | 0.180825983 |
| Ak6 | 0.388017091 | 0.181056518 |
| Hsp90b1 | -0.284934759 | 0.181088374 |
| Ccdc58 | 0.466749835 | 0.181151145 |
| Dand5 | 1.784178316 | 0.181227867 |
| Stoml1 | 0.4257377 | 0.18133217 |
| Aimp1 | 0.356715212 | 0.181384127 |
| Hspa12a | -0.633238162 | 0.181384127 |
| Slc35g2 | 1.293046889 | 0.181475234 |
| Nfxl1 | -0.574085709 | 0.181478386 |
| Dhx38 | -0.496412691 | 0.181491797 |
| Shq1 | -0.531737535 | 0.181524457 |
| Abce1 | 0.332702597 | 0.182187257 |
| Sacm1l | -0.503311569 | 0.182189806 |
| Catip | -2.249075353 | 0.182189806 |
| Rsrc1 | 0.373914944 | 0.182198136 |
| Tmeff1 | -0.43222411 | 0.182223865 |
| Gnb1 | -0.372673713 | 0.182254603 |
| Rab3gap1 | -0.457222798 | 0.18232024 |
| Orc4 | -0.351225495 | 0.182398463 |
| Arvcf | 0.430258686 | 0.182417956 |
| Ubtd1 | -0.643569435 | 0.182419309 |
| Rmi1 | -0.609738906 | 0.182683018 |
| Aif1 | -0.553900446 | 0.182776747 |
| Rab24 | 0.337747942 | 0.182822628 |
| Olfr878 | 4.715914582 | 0.18282724 |
| Rufy2 | 0.431173373 | 0.182880076 |
| Grin1 | -0.710651852 | 0.18322044 |
| Fam167a | 0.798296056 | 0.183631397 |
| Pcdhb18 | -1.558685369 | 0.18383033 |
| S100a11 | 0.856474725 | 0.183868869 |
| Tril | 1.324904427 | 0.184021899 |
| Med29 | 0.357930233 | 0.184021899 |
| Kmt2b | -0.489361168 | 0.184021899 |
| Sema3b | 3.440954488 | 0.184091436 |
| Ticam1 | -0.721306308 | 0.184091436 |
| Shmt2 | -0.326272409 | 0.184106385 |
| Bcas3 | -0.476609344 | 0.184155079 |
| Ppp1r17 | 4.740175535 | 0.184313157 |
| Prlr | 4.316707933 | 0.184313157 |
| Amt | 0.63206699 | 0.184322434 |
| Slain1 | -0.582887417 | 0.184588383 |
| Tmem86a | -0.455841179 | 0.184623075 |
| Fzd6 | 2.335780115 | 0.184686483 |
| Rfc3 | 0.41160437 | 0.184686483 |
| Ccdc85a | 1.03608541 | 0.184801142 |
| Gpr83 | 2.558659767 | 0.184849437 |
| Gna14 | 2.258536483 | 0.184849437 |
| Dkk1 | -5.234113579 | 0.184849437 |
| Atmin | -0.687354344 | 0.184876485 |
| Slc28a2 | -3.671445914 | 0.184876485 |
| Nup43 | -0.445197671 | 0.184896765 |
| Mex3c | 0.669218314 | 0.184919832 |
| Srfbp1 | -0.606375161 | 0.184950608 |
| Mdga2 | -0.758661462 | 0.184950608 |
| Rgs6 | -0.924263715 | 0.184973677 |
| Dstyk | -0.494481269 | 0.185076652 |
| Sypl | -0.367483353 | 0.185218256 |
| Gpr21 | -0.595648987 | 0.185218256 |
| Camsap1 | 0.386312626 | 0.185258699 |
| Fam81b | 4.722941677 | 0.185301503 |
| Aes | -0.314802189 | 0.185324803 |
| Cnbp | 0.279565387 | 0.185459465 |
| Tdp2 | 0.413166095 | 0.185570288 |
| Psmd9 | 0.356245135 | 0.185570288 |
| Mbd4 | 0.785525901 | 0.185781165 |
| Cbx7 | -3.636406244 | 0.185984342 |
| Dcun1d4 | -0.457590566 | 0.186045371 |
| Sema5b | 0.513213753 | 0.18613616 |
| Lrrc1 | 1.02856415 | 0.186253289 |
| Dlk2 | 1.629965948 | 0.186388003 |
| Ptprr | 1.14265204 | 0.186388003 |
| Gprc5d | -5.226498909 | 0.186388003 |
| Haus8 | -0.538093091 | 0.1865012 |
| Trmt10a | 0.902925684 | 0.186569188 |
| Dkk4 | -5.25569876 | 0.186569188 |
| Oas1a | -1.415450336 | 0.186694859 |
| Tdrd3 | 0.346277938 | 0.186810733 |
| Spata24 | 0.986570258 | 0.186924831 |
| Cdc14a | -0.598191927 | 0.187004215 |
| Pygb | -0.430066464 | 0.18704357 |
| C1qtnf7 | 4.852080453 | 0.187057461 |
| Mapk12 | -1.285611769 | 0.187096825 |
| Slc36a4 | -0.386305898 | 0.187316158 |
| Tagln3 | 0.349401036 | 0.187384295 |
| Cnnm4 | -0.41342245 | 0.187454076 |
| Lypd1 | 1.264229244 | 0.187513096 |
| Gm15446 | 0.585948625 | 0.187567258 |
| Smoc1 | 0.693986889 | 0.187635966 |
| Trim69 | 4.795840402 | 0.187676048 |
| Tead2 | 0.473299342 | 0.187782333 |
| Ntrk3 | -0.68248388 | 0.187855222 |
| Polg2 | -0.711940231 | 0.188091789 |
| Nmrk1 | -0.915884629 | 0.188096337 |
| Bend7 | 1.803651102 | 0.188107916 |
| Chrna7 | -0.755357469 | 0.188322094 |
| Gpr85 | -0.358658217 | 0.188495794 |
| Bcl9l | -0.597348478 | 0.188495794 |
| Clns1a | 0.321458507 | 0.188590992 |
| Polr2g | 0.313202616 | 0.188590992 |

| | | |
|---|---|---|
| Cpne4 | 0.621559593 | 0.537520653 |
| Ramp3 | 0.594948013 | 0.537520653 |
| Wfs1 | -0.555325997 | 0.537520653 |
| Unc119b | 0.245719086 | 0.53754567 |
| Tubgcp4 | -0.24732117 | 0.53754567 |
| Paqr5 | -0.845670562 | 0.53754567 |
| Ldhd | -1.427001561 | 0.53754567 |
| Nav3 | -0.233796368 | 0.537666456 |
| Clca4a | 3.490524496 | 0.537667804 |
| Gm10447 | 3.490524496 | 0.537667804 |
| Gsdmc2 | 3.490524496 | 0.537667804 |
| Tmem64 | 0.298107225 | 0.537667804 |
| Ulk2 | 0.260736583 | 0.537667804 |
| Prps2 | -0.259566967 | 0.537667804 |
| Polr3c | 0.212559535 | 0.537687255 |
| Kcnj4 | 3.488056824 | 0.538061918 |
| Bbs1 | 0.408450456 | 0.538187351 |
| Mmp16 | -0.292493624 | 0.538187351 |
| Ubqln4 | -0.178225884 | 0.538293987 |
| Anxa9 | -1.357897371 | 0.538293987 |
| Tll1 | 0.997759451 | 0.538464712 |
| Dcun1d2 | -0.419691821 | 0.538783985 |
| Fut8 | 0.345274209 | 0.538789349 |
| Bnip3l | 0.136585046 | 0.538789349 |
| Dnajc7 | 0.145709544 | 0.538820539 |
| Lyrm5 | -0.279231498 | 0.53904435 |
| Sely | -0.489395199 | 0.539510005 |
| Stac3 | -2.010749113 | 0.539510005 |
| Sox3 | 0.97263936 | 0.539540711 |
| Ptcd3 | -0.181527926 | 0.539766727 |
| Wiz | -0.257533841 | 0.539849601 |
| Zfp280d | -0.172955519 | 0.539861636 |
| D130043K22Rik | 1.09192627 | 0.539934063 |
| AI854703 | 0.463608771 | 0.539946885 |
| Ehbp1 | 0.255518638 | 0.539949886 |
| Muc1 | -0.861005451 | 0.54005522 |
| Fzd8 | -0.576496731 | 0.540320886 |
| Phb2 | 0.161369386 | 0.540474112 |
| Sgcz | -0.906447475 | 0.540474112 |
| Cask | 0.199786056 | 0.540520047 |
| Gm7102 | 2.43053456 | 0.540621271 |
| Cfap57 | 1.25339234 | 0.540621271 |
| Cep41 | 0.35989006 | 0.540621271 |
| Gnpnat1 | 0.321356739 | 0.540621271 |
| Arl5b | 0.183626441 | 0.540621271 |
| Elavl3 | 0.160108138 | 0.540621271 |
| Htra2 | -0.275019633 | 0.540621271 |
| Shc2 | -0.42145541 | 0.540621271 |
| Jakmip3 | -0.860284991 | 0.540621271 |
| Pygo2 | -0.260807906 | 0.540754951 |
| Gm20379 | -0.666175183 | 0.540854242 |
| Flrt2 | -0.277788166 | 0.540981094 |
| Syt7 | 0.348898886 | 0.541106357 |
| Fkbpl | -0.345322408 | 0.541106357 |
| Cckar | 1.398405521 | 0.54125757 |
| Pitrm1 | 0.227140687 | 0.541342142 |
| Il1rl1 | 2.058323551 | 0.541460523 |
| Slc10a7 | -0.336781647 | 0.541473313 |
| C1ql3 | 1.22805053 | 0.54151249 |
| Lgals7 | 0.840407615 | 0.54151249 |
| 2610020H08Rik | 0.409813761 | 0.54151249 |
| Sfi1 | 0.317833394 | 0.54151249 |
| Pxmp4 | 0.268431576 | 0.54151249 |
| Irf3 | 0.259548144 | 0.54151249 |
| Zmym4 | 0.201828445 | 0.54151249 |
| Nelfb | -0.199312379 | 0.54151249 |
| Tmem132a | -0.201623797 | 0.54151249 |
| Vps25 | 1.455626475 | 0.541607593 |
| Usp32 | 0.266368134 | 0.541665117 |
| Dcx | -0.168950997 | 0.541882409 |
| Mterf3 | -0.234326076 | 0.541882409 |
| Zkscan1 | 0.204277554 | 0.542152231 |
| Morc4 | 0.626086009 | 0.542161652 |
| Mboat4 | 3.342092972 | 0.542368327 |
| Slco1a5 | 3.342092972 | 0.542368327 |
| Arhgef33 | 1.49573834 | 0.542828116 |
| Gm29096 | 3.34140145 | 0.543203333 |
| Lhx4 | 3.34140145 | 0.543203333 |
| Pih1h3b | 3.34140145 | 0.543203333 |
| Mgp | 1.048019496 | 0.543203333 |
| Sorcs3 | 0.505785729 | 0.543203333 |
| Ccdc69 | -0.842919588 | 0.543391135 |
| Mrpl17 | 0.168127656 | 0.543427314 |
| Adsl | 0.202175017 | 0.543707038 |
| Abca12 | -2.09796598 | 0.543707038 |
| Gm6793 | 0.203073757 | 0.543788074 |
| Use1 | 0.198286033 | 0.543861904 |
| Ptprcap | 1.937821765 | 0.544306346 |
| Gimap9 | -1.046755874 | 0.54437578 |
| Pou3f4 | 0.372955923 | 0.544382339 |
| Ralgps2 | 0.179455198 | 0.544382339 |
| Serbp1 | 0.129350352 | 0.544531952 |
| Dnajb9 | -0.234697562 | 0.544742603 |
| Ppp1r8 | 0.209260768 | 0.544823804 |
| Pelo | 0.600981597 | 0.544883036 |
| Nlgn3 | -0.199555395 | 0.54504166 |
| Ube2r2 | -0.247396195 | 0.545247906 |
| Kin | 0.344818932 | 0.545288276 |
| Sde2 | -0.1941973 | 0.545368546 |
| Zfp839 | -0.627307283 | 0.545524575 |
| Myh7b | 1.19470674 | 0.545558125 |
| Ccdc155 | 1.66379094 | 0.545566127 |
| Ndufa10 | 0.158254035 | 0.545566127 |
| Epha6 | 0.820800967 | 0.545732166 |
| Tns4 | -0.866471767 | 0.545732166 |
| Edem3 | 0.292542572 | 0.546019957 |
| Ufsp2 | -0.221830872 | 0.546019957 |
| Dynlt1a | 0.258489774 | 0.546056487 |
| Ube2o | 0.289860044 | 0.546109627 |
| Ptprf | 0.222987647 | 0.546715222 |
| Mapk11 | 0.307411565 | 0.546801623 |
| Ptgdr | 1.584466292 | 0.546831321 |
| Cbx3 | 0.169977438 | 0.546831321 |
| Ttc5 | 0.177603208 | 0.546866901 |
| Letm1 | -0.184249545 | 0.546923084 |
| Ncam1 | 0.171372621 | 0.546934742 |
| Pcid2 | -0.322492161 | 0.546934742 |
| Upk3a | 2.387442745 | 0.546956286 |
| Znrf2 | 0.251723676 | 0.547133472 |

| | | |
|---|---|---|
| Rbfox1 | 0.046492481 | 0.926395372 |
| Mcat | 0.045708066 | 0.926395372 |
| Spg21 | -0.026809719 | 0.926395372 |
| Tbc1d22b | -0.044252407 | 0.926395372 |
| Mcm2 | -0.044281848 | 0.926395372 |
| Smim12 | -0.050025223 | 0.926395372 |
| Zc3h10 | -0.053270351 | 0.926395372 |
| Tmem62 | 0.089394032 | 0.926447785 |
| Cyp4f15 | 0.366709153 | 0.926563834 |
| Zfp933 | 0.093155523 | 0.926563834 |
| Gm4707 | 0.078353838 | 0.926672936 |
| Snrnp200 | 0.033276806 | 0.9267002 |
| Fam173a | 0.035001351 | 0.926795177 |
| Cd74 | 0.072138961 | 0.926928979 |
| Cldnd1 | 0.03154902 | 0.927119597 |
| Rap2a | -0.050155657 | 0.927251772 |
| Aaas | -0.043582262 | 0.927723543 |
| Rsl24d1 | -0.032484843 | 0.927757295 |
| Ribc2 | -0.184306521 | 0.927757295 |
| Selenbp1 | 0.05532969 | 0.928011607 |
| Azin1 | -0.035280712 | 0.928030096 |
| Gfod2 | -0.044703863 | 0.928030096 |
| Osbpl3 | -0.064910039 | 0.928030096 |
| Ggnbp1 | -0.121973226 | 0.928030096 |
| Ccr2 | -0.164467787 | 0.928030096 |
| Gm14419 | -0.241562779 | 0.928030096 |
| Dusp9 | 0.229151121 | 0.928074491 |
| Mtif3 | 0.054432922 | 0.928074491 |
| Rnf166 | 0.049560076 | 0.928074491 |
| Ddx3y | -0.036033224 | 0.928074491 |
| Eftud1 | -0.059139662 | 0.928074491 |
| Wnt2b | 0.158310688 | 0.928098987 |
| Gm6356 | 0.081435645 | 0.928098987 |
| Asap2 | 0.036963648 | 0.928098987 |
| Fam193b | -0.04577864 | 0.928098987 |
| Fn3k | -0.114235524 | 0.928098987 |
| Erich1 | 0.053511599 | 0.928495407 |
| Hirip3 | 0.045083612 | 0.928495407 |
| Crybb3 | 0.387989337 | 0.928685609 |
| Diaph3 | 0.07620628 | 0.928915165 |
| Rimkla | -0.051195422 | 0.929102101 |
| Srl | -0.06075326 | 0.929122727 |
| Ankrd13d | -0.036682441 | 0.929248408 |
| Pole | -0.054432463 | 0.92928798 |
| S100a1 | 0.049465148 | 0.929351614 |
| Usp31 | -0.063355324 | 0.92942897 |
| Inca1 | 0.224936714 | 0.92975832 |
| Fbxo46 | -0.056011995 | 0.930018675 |
| Tnfrsf13b | 0.066855369 | 0.930030176 |
| Mras | -0.038730954 | 0.930210724 |
| Gm21860 | -0.436211596 | 0.930210724 |
| Tusc1 | 0.086548277 | 0.930273551 |
| Pex7 | -0.044230298 | 0.930273551 |
| Vps37b | -0.042552992 | 0.930399057 |
| Styxl1 | -0.274131718 | 0.930403637 |
| Ap4e1 | -0.045894541 | 0.931009032 |
| Gm9833 | -0.049759985 | 0.931009032 |
| Gm16372 | 0.141354725 | 0.931081352 |
| Fbxo22 | -0.038135591 | 0.931204857 |
| Pou2f1 | -0.042160211 | 0.931204857 |
| Synpo2 | -0.175193575 | 0.931297421 |
| Lyg2 | -0.40312334 | 0.931297421 |
| Ccdc7b | 0.385545272 | 0.931494265 |
| Atl1 | -0.048677944 | 0.931512451 |
| 1110034G24Ri | -0.044082953 | 0.931601697 |
| Socs6 | -0.037322736 | 0.931752625 |
| Nob1 | 0.03939791 | 0.931853058 |
| H2-DMb2 | 0.144898618 | 0.932006731 |
| Taf5 | 0.05589915 | 0.932006731 |
| Ctps2 | 0.034863358 | 0.932096273 |
| Lingo3 | 0.103756509 | 0.932117904 |
| Nelfa | -0.047026457 | 0.932216589 |
| Atf2 | 0.031896058 | 0.932231184 |
| Mixl1 | 0.230377089 | 0.932344532 |
| Ptp4a1 | -0.029089126 | 0.932371399 |
| Fastkd1 | -0.061214132 | 0.93254602 |
| Necab3 | -0.07158185 | 0.93254602 |
| Tff3 | 0.344447233 | 0.932920613 |
| Gins1 | 0.044038382 | 0.932920613 |
| Tfdp1 | -0.028383133 | 0.932920613 |
| Sox12 | -0.048581984 | 0.932920613 |
| Gm17728 | -0.253923315 | 0.93309847 |
| Fam187b | -0.388305498 | 0.93309847 |
| Strip1 | -0.029732272 | 0.933108922 |
| Ppfia3 | -0.052856945 | 0.933333773 |
| Ceacam16 | -0.280926433 | 0.933359168 |
| Zfx | -0.026956025 | 0.933815627 |
| Guca1a | -0.099899062 | 0.933993967 |
| Zfp618 | 0.06892539 | 0.934026375 |
| F2r | 0.049794029 | 0.934026375 |
| Lamb3 | 0.161458017 | 0.934036167 |
| Mthfd1 | -0.047313707 | 0.934036167 |
| Selo | -0.048193972 | 0.934290025 |
| Iars | 0.034607607 | 0.934576996 |
| Atp6v0d1 | 0.02390226 | 0.934750193 |
| Tbc1d10a | -0.0376104 | 0.934750193 |
| Csrp2bp | -0.062536509 | 0.934796868 |
| Ino80d | -0.043541742 | 0.935362044 |
| Scx | 0.152828903 | 0.935670255 |
| Nanos3 | 0.077975246 | 0.935670255 |
| Fam132a | -0.065444302 | 0.935701957 |
| Cd99l2 | -0.037712995 | 0.935767904 |
| Crebrf | 0.037418795 | 0.935993861 |
| Fam177a | -0.165471419 | 0.935993861 |
| Tesc | 0.163259596 | 0.936051835 |
| Aarsd1 | -0.107777877 | 0.936259938 |
| Pex14 | 0.036310691 | 0.936432523 |
| Syt10 | 0.202538188 | 0.936541347 |
| Alpk3 | 0.123412705 | 0.936579244 |
| Mettl18 | 0.077176325 | 0.936579244 |
| Cars2 | 0.038361847 | 0.936579244 |
| Glyr1 | 0.022396198 | 0.936579244 |
| Cpsf6 | -0.021448075 | 0.936579244 |
| Gm17669 | -0.370173675 | 0.936579244 |
| Cul1 | -0.027905784 | 0.936614157 |
| Hist1h2bm | -0.099064254 | 0.936614157 |
| Rps18 | 0.038613147 | 0.936695597 |
| Ccdc33 | 0.227960951 | 0.936844137 |
| Rnf144a | -0.039833373 | 0.937323808 |

| | | |
|---|---|---|
| Tmem9 | 0.618271506 | 0.011309584 |
| Mrpl35 | 0.666052176 | 0.011320522 |
| Ifi44 | 3.211024221 | 0.01132234 |
| Pes1 | -0.626286774 | 0.011346274 |
| Acta1 | 2.216285902 | 0.011352168 |
| Ube2n | 0.481722669 | 0.011352168 |
| Dad1 | 0.509899259 | 0.011352905 |
| Churc1 | 0.801631281 | 0.011355661 |
| Hbb-bh1 | 5.815181163 | 0.011371072 |
| Sdhb | 0.530343858 | 0.011375449 |
| Adssl1 | -1.03239231 | 0.01137989 |
| Tldc1 | -1.198864317 | 0.011382068 |
| Tmem161a | -0.632616762 | 0.011387994 |
| Steap3 | -2.778103884 | 0.01139588 |
| Setd7 | -0.977088124 | 0.011498991 |
| Hgs | -0.58021562 | 0.0115148 |
| Aktip | 0.532293913 | 0.011557042 |
| Tnfsf9 | -1.449719252 | 0.011557042 |
| Nlrc4 | -1.839888236 | 0.01160327 |
| Tpx2 | 0.744882992 | 0.011605079 |
| Tnfrsf19 | 0.948093896 | 0.011608196 |
| 4833439L19Rik | 0.508501352 | 0.01161938 |
| Ssb | 0.523379883 | 0.011620723 |
| Unc5a | -1.171032621 | 0.011622298 |
| Rnf13 | -0.666720754 | 0.01163223 |
| Rsrc2 | 0.565034956 | 0.011666893 |
| Eif4ebp1 | -0.919873118 | 0.011674311 |
| Itpkb | -1.3211852 | 0.011689602 |
| Traf1 | -2.22029218 | 0.011689602 |
| Mrps36 | 0.718146706 | 0.011692332 |
| Tead1 | 0.821842944 | 0.01169385 |
| Pin1 | 0.530994538 | 0.011697164 |
| Pi4k2b | -1.256415052 | 0.011697164 |
| Abcd1 | -1.086194633 | 0.011704005 |
| Jakmip1 | -1.062722181 | 0.01170411 |
| 4921524J17Rik | 0.695291081 | 0.011710006 |
| Upk3b | -3.351821964 | 0.011710006 |
| Fbxl2 | 1.344843898 | 0.01171307 |
| Itgb1bp2 | -2.707541838 | 0.011715279 |
| Thap2 | 0.999302459 | 0.01172473 |
| Bnip2 | -0.595702355 | 0.011727221 |
| Vstm4 | 4.057183369 | 0.011730682 |
| Smarca1 | 0.760363331 | 0.01173695 |
| Ssna1 | 0.602841689 | 0.01173695 |
| Ppp2r1b | -0.645985521 | 0.01173695 |
| Etaa1 | 1.33194263 | 0.011750097 |
| Brd3 | 0.603283033 | 0.011763473 |
| Chic1 | 1.09589613 | 0.01183177 |
| Ssh3 | -1.362085062 | 0.011845542 |
| Pcdh12 | 5.940489248 | 0.011858539 |
| Pak3 | 0.681575418 | 0.011859784 |
| Ust | -1.09437238 | 0.011860099 |
| Gpsm3 | -1.455413352 | 0.011860099 |
| Stra6 | 2.184570077 | 0.01186159 |
| Lrrc75a | -1.163774728 | 0.011862941 |
| Pola2 | -0.945135743 | 0.011877962 |
| Poglut1 | 0.720938196 | 0.011907882 |
| Nrg3 | -1.492169508 | 0.011965564 |
| Tmem165 | -0.793641922 | 0.011969353 |
| Ninj1 | -0.944624071 | 0.011970274 |
| Ghitm | -0.478537891 | 0.01197282 |
| Thap8 | 6.009303831 | 0.01198882 |
| U2af1 | 0.546097436 | 0.012006996 |
| 4932443I19Rik | 5.877727688 | 0.012024276 |
| 9230110C19Rik | 0.924501861 | 0.012028979 |
| Bccip | 0.586706168 | 0.012044225 |
| Gtf2a2 | 0.517495432 | 0.012044225 |
| 2310033P09Rik | 1.045071101 | 0.012073013 |
| Sec61b | 0.533247079 | 0.012107148 |
| Wnk1 | -0.573907501 | 0.012113748 |
| Bnc2 | 5.401369714 | 0.012113815 |
| Rabgap1l | -0.832388866 | 0.012132568 |
| Aldh16a1 | -0.84868661 | 0.01217875 |
| Nlrp6 | 5.902217151 | 0.012195631 |
| Chmp2a | 0.538443383 | 0.012197808 |
| Rnaseh2b | 0.835331535 | 0.012204501 |
| Plekhn1 | -3.168394792 | 0.012204501 |
| Ap5z1 | -1.360186804 | 0.012219392 |
| Micall2 | -1.963569137 | 0.012219392 |
| Grb14 | 1.293125622 | 0.012221612 |
| Ppp4r4 | -2.589797273 | 0.01224604 |
| Cdh8 | 0.72844615 | 0.01224768 |
| Tgfb2 | 1.19698332 | 0.012252132 |
| Kcnk3 | 4.575863477 | 0.012293067 |
| Ebp | -0.814176848 | 0.012309022 |
| Tbl1xr1 | -0.907510261 | 0.012309022 |
| Fez1 | 0.683817655 | 0.012310306 |
| Snapc4 | -0.966857341 | 0.012310306 |
| Art5 | -4.013793635 | 0.01232052 |
| Cep19 | 1.041626743 | 0.012368853 |
| Ywhae | 0.472086738 | 0.012368853 |
| Lrrc8c | -1.629891506 | 0.012392459 |
| Tktl1 | -2.57694844 | 0.012422624 |
| Card19 | -1.046250272 | 0.01244338 |
| Soga1 | -1.165236404 | 0.01244338 |
| Pfn1 | -0.692306007 | 0.012458597 |
| Lect1 | 3.422946462 | 0.012467151 |
| Agpat1 | -0.545009667 | 0.012475851 |
| Ube2h | -0.646990479 | 0.012475851 |
| Mxi1 | -0.682928285 | 0.012483947 |
| Aifm2 | -1.67345932 | 0.012499316 |
| Mpv17 | 0.690195114 | 0.012503809 |
| Tmco3 | -0.721103237 | 0.012507449 |
| Sell | -3.491555274 | 0.012512452 |
| Ndufb8 | 0.595020015 | 0.012538129 |
| Stox2 | 0.714031124 | 0.012563069 |
| Sptssa | 0.554023281 | 0.012569471 |
| Slc17a7 | -1.80245946 | 0.012580425 |
| Pcdha8 | 5.91257889 | 0.012600841 |
| Gbp2b | -6.703996398 | 0.012634089 |
| Cnpy1 | 5.625897062 | 0.012650966 |
| Foxp2 | 1.1573904 | 0.012659912 |
| Mtcl1 | -1.036174346 | 0.012716454 |
| Xkr6 | 1.642147016 | 0.012747328 |
| Capzb | -0.538139396 | 0.012756293 |
| Rnaset2b | -0.950119195 | 0.012796728 |
| Sirt6 | 0.637555305 | 0.012799995 |
| Dck | -0.643040634 | 0.012803041 |
| Pfn2 | 0.493413929 | 0.012842662 |

| | | |
|---|---|---|
| Tmem26 | 1.466583974 | 0.188694139 |
| Cxcl16 | -1.205455318 | 0.188732683 |
| Tarbp2 | 0.392030563 | 0.188840017 |
| Smug1 | 0.736168865 | 0.188853105 |
| Ston2 | -0.793170708 | 0.188853105 |
| Meiob | -5.211681534 | 0.188853105 |
| Wisp3 | 3.678775133 | 0.189047478 |
| Carm1 | -0.616629566 | 0.189047478 |
| Mnda | -1.127280606 | 0.189160375 |
| 4933415A04Rik | -0.878334337 | 0.189196486 |
| Fndc3b | -0.651920695 | 0.189233037 |
| Cntnap5a | 1.256577378 | 0.18927988 |
| Gm9726 | 1.271442631 | 0.18932538 |
| Pole3 | 0.359821889 | 0.18932538 |
| Zfand6 | 0.284306808 | 0.189485799 |
| Cradd | 0.761601322 | 0.189561777 |
| Smpd4 | 0.367978113 | 0.189584142 |
| Mrc1 | 1.181775207 | 0.189678922 |
| Kif1a | 0.421033646 | 0.18996602 |
| Ino80c | -0.499487878 | 0.18996602 |
| Mmrn1 | 4.701351495 | 0.189972618 |
| Myoz1 | 4.701351495 | 0.189972618 |
| Atp8b5 | 1.757024852 | 0.189972618 |
| Clvs1 | -0.484904098 | 0.189972618 |
| Fdx1 | 0.552481142 | 0.19012362 |
| 1700007K13Rik | 1.870153449 | 0.190125597 |
| Pou4f1 | -2.298367105 | 0.190125597 |
| Tmem79 | 2.289817768 | 0.190162385 |
| Slc16a1 | -0.716641002 | 0.190396571 |
| Tmem117 | -1.88474356 | 0.190501849 |
| Kcnn1 | 0.96762455 | 0.190562802 |
| Stk26 | -1.64953299 | 0.190721266 |
| Ubash3b | -0.576790424 | 0.190794655 |
| Tgfb3 | 1.351276059 | 0.191074612 |
| Cbr3 | -1.753441592 | 0.191104659 |
| Gzmm | 2.657377758 | 0.1911391 |
| Ydjc | -0.645307484 | 0.191262443 |
| Nfe2 | -1.857495361 | 0.191262443 |
| Phkg2 | -0.493659139 | 0.191376776 |
| Tgm3 | -4.464864686 | 0.191597061 |
| Fnta | 0.311213993 | 0.191634277 |
| Atp6v1c1 | -0.310953621 | 0.191668624 |
| Eps8l2 | 4.032654276 | 0.191771214 |
| Olfm3 | 1.456796587 | 0.191799328 |
| Uba6 | 0.469075742 | 0.191799328 |
| Cnppd1 | -0.384943445 | 0.191799328 |
| Rnf4 | -0.314048288 | 0.191987884 |
| Psmc4 | 0.292440822 | 0.192119662 |
| Xrn2 | -0.257957768 | 0.192119662 |
| Ttll6 | 4.656652178 | 0.192178665 |
| Elmo2 | -0.437584051 | 0.192229685 |
| Tsnaxip1 | 4.31591704 | 0.192237916 |
| Nipbl | 0.329009974 | 0.192496925 |
| Ltn1 | -0.351843258 | 0.192496925 |
| Nynrin | 0.748351353 | 0.192562098 |
| Tnrc18 | -0.370720034 | 0.192628673 |
| Fbxo6 | -0.497163475 | 0.192691114 |
| Ctsd | -0.503620294 | 0.192691114 |
| Lrr1 | 0.812399423 | 0.192698127 |
| Fes | -0.696015667 | 0.193015738 |
| Gm10036 | 0.474436862 | 0.193065851 |
| Tmem184c | 0.320007716 | 0.193065851 |
| Zfp445 | -0.382744146 | 0.193065851 |
| Ppp1r1b | -0.733044159 | 0.193065851 |
| Cdca7 | 0.49201291 | 0.193130201 |
| Gpx3 | 0.440207725 | 0.193130201 |
| Lrsam1 | -0.794802527 | 0.193130201 |
| Hpse | -2.744894979 | 0.193130201 |
| Supt7l | 0.473943856 | 0.193170933 |
| Mapk8ip3 | -0.326546049 | 0.193269797 |
| Edar | 4.330146378 | 0.193382731 |
| Lynx1 | 1.466013797 | 0.193382731 |
| Samd5 | 1.067797828 | 0.193382731 |
| Dbndd1 | 0.370371647 | 0.193382731 |
| Sv2c | 0.894578555 | 0.193802439 |
| Trim34b | -4.14307697 | 0.193836215 |
| Agpat9 | -1.566240973 | 0.19401967 |
| Ankrd63 | 3.748639133 | 0.194058703 |
| Adck3 | 1.405793026 | 0.194329894 |
| Plch1 | 0.866680157 | 0.194329894 |
| Atp2b1 | -0.277637798 | 0.194418106 |
| Ddx55 | 0.564957603 | 0.194424336 |
| Tmem29 | 0.334090601 | 0.194424336 |
| Zadh2 | -0.80293256 | 0.194577105 |
| 2300009A05Rik | 0.529388065 | 0.194714385 |
| Snx4 | -0.388389937 | 0.194793286 |
| Ccdc84 | -0.908122133 | 0.194793286 |
| Spry1 | 0.894430925 | 0.194794257 |
| Tmem150c | 0.735539019 | 0.194817456 |
| 7-Mar | -0.421852051 | 0.194817456 |
| St6galnac3 | -0.679796052 | 0.194817456 |
| Naa30 | -0.442776944 | 0.194876559 |
| Safb2 | -0.353800508 | 0.194980608 |
| Ctc1 | -0.400184384 | 0.194980608 |
| Rnf26 | -0.401434094 | 0.194980608 |
| Lefty1 | 4.674990927 | 0.194999709 |
| Lamb2 | 1.12951895 | 0.194999709 |
| Napb | 0.430045219 | 0.194999709 |
| Nuggc | 4.809140695 | 0.195114213 |
| Ddx39 | 0.373454933 | 0.195114213 |
| Cers2 | -0.37932282 | 0.195114213 |
| Cpsf4l | -1.891469866 | 0.195161321 |
| Tpo | -2.729447907 | 0.195304782 |
| Gle1 | -0.366733112 | 0.195342399 |
| Ankra2 | 0.428336799 | 0.195413685 |
| Slmo1 | 0.435111217 | 0.195439434 |
| Usp5 | -0.33715514 | 0.195481932 |
| Smc1b | -2.224326504 | 0.195481932 |
| Mei1 | 5.116271668 | 0.195605721 |
| Herc2 | -0.343180077 | 0.195659698 |
| Ncbp2 | 0.306506182 | 0.196170302 |
| Wnt7a | 0.757261252 | 0.196201673 |
| Insig1 | -0.298327314 | 0.196201673 |
| Trim30a | -1.37830726 | 0.196334254 |
| Prdm15 | -0.649937552 | 0.1963871 |
| Phf13 | -0.743748947 | 0.196410513 |
| Gtpbp10 | 0.584608984 | 0.196500978 |
| Ttll9 | 1.804252641 | 0.196583598 |
| Gng4 | 0.827381071 | 0.196702256 |

| | | |
|---|---|---|
| Ccdc160 | -0.89044239 | 0.547133472 |
| Spg11 | -0.417650266 | 0.547268357 |
| Dnajb12 | 0.330990275 | 0.548015969 |
| Wapl | -0.184751655 | 0.548277336 |
| Ppp1r16b | -0.317227659 | 0.5482883 |
| Tomm34 | -0.171025721 | 0.548315431 |
| Tubb3 | 0.129164782 | 0.548425884 |
| Zfp507 | -0.369234051 | 0.54885554 |
| Col4a4 | 0.761053798 | 0.548991194 |
| 2310035C23Rik | -0.186631541 | 0.548991194 |
| Rnf24 | -0.205386095 | 0.548991194 |
| Dclk2 | -0.220871959 | 0.548992448 |
| 0610010F05Rik | 0.159848448 | 0.54920293 |
| Amh | -3.063193094 | 0.549396408 |
| Arhgef19 | 1.706619052 | 0.549415858 |
| Man2b2 | -0.325713695 | 0.549606742 |
| Mapk1ip1 | -0.291629686 | 0.549653919 |
| Mat2b | -0.150941199 | 0.549689088 |
| Ddx18 | 0.241486625 | 0.549818284 |
| Fdft1 | 0.154357554 | 0.549882075 |
| Hipk4 | -1.313195363 | 0.549882075 |
| Fbxw4 | -0.573285374 | 0.549953664 |
| Slc5a3 | 0.268191655 | 0.550119205 |
| Acadsb | -0.163316392 | 0.550232984 |
| Nckap5 | -0.285437496 | 0.550232984 |
| Srrd | -0.338690186 | 0.550476663 |
| Nmd3 | -0.194561521 | 0.550619053 |
| Trappc2 | -0.214343968 | 0.550865669 |
| Susd3 | 0.264353612 | 0.550905804 |
| Rcl1 | 0.237099706 | 0.550976529 |
| Clcn3 | -0.173802757 | 0.551130445 |
| 4931429L15Rik | 3.269928566 | 0.551291315 |
| Lsamp | 0.208412649 | 0.551291315 |
| Tacc3 | 0.174751358 | 0.551291315 |
| Gm14399 | -1.34905258 | 0.551291315 |
| Vmn2r1 | -1.177075567 | 0.551627263 |
| Dlgap1 | 0.261671586 | 0.551894736 |
| Cntn1 | -0.419912508 | 0.551894736 |
| Mtbp | -0.49907161 | 0.551894736 |
| Map2k4 | 0.20580287 | 0.55213615 |
| Ccna1 | 0.559379177 | 0.55218762 |
| Kcnip1 | 0.242723735 | 0.55218762 |
| Cdk5rap1 | 0.505644864 | 0.552316107 |
| Gm12258 | 0.41681072 | 0.552316107 |
| Dll3 | 0.276754953 | 0.552316107 |
| Gatad2b | -0.169256344 | 0.552705933 |
| Rpusd2 | 0.467020371 | 0.552775091 |
| Fgf3 | 1.084232659 | 0.552930346 |
| Brdt | 0.780345556 | 0.553091029 |
| Hdac4 | 0.18647028 | 0.553127907 |
| Gm10762 | -0.368191959 | 0.553214258 |
| Fam129c | -1.049871184 | 0.553221656 |
| Mettl15 | 0.424193527 | 0.553278122 |
| Gm11168 | 3.372563511 | 0.553357068 |
| Gp9 | 3.372563511 | 0.553357068 |
| Iqgap2 | 0.481150444 | 0.553357068 |
| Wtip | -0.419347417 | 0.553357068 |
| Knop1 | 0.191957405 | 0.553444434 |
| Zmynd8 | 0.220056509 | 0.553731164 |
| Sesn1 | -0.278873045 | 0.553908144 |
| Pld6 | -0.950618251 | 0.554144366 |
| Slc1a4 | -0.36398951 | 0.55419158 |
| 4921513D11Rik | 3.364724965 | 0.554394083 |
| Clrn1 | 3.364724965 | 0.554394083 |
| Loxl4 | 3.364724965 | 0.554394083 |
| Ppbp | 3.364724965 | 0.554394083 |
| Lyrm7 | 0.848252184 | 0.554394083 |
| G3bp1 | -0.142803485 | 0.554394083 |
| Gemin8 | 0.34803429 | 0.554400226 |
| Ocel1 | -0.384746648 | 0.554400226 |
| Gm3002 | 2.353666902 | 0.554424298 |
| Magee2 | 0.556245929 | 0.554424298 |
| Plch2 | 0.528833916 | 0.554502886 |
| Tcf24 | 1.758273915 | 0.55467929 |
| Ap1s1 | -0.195475594 | 0.554683469 |
| Gtf2b | 0.206482933 | 0.554869957 |
| Mrps2 | 0.214322961 | 0.554892612 |
| Zfp748 | 0.365443291 | 0.555028743 |
| Rc3h2 | -0.195876883 | 0.555057317 |
| Ern2 | 3.269051162 | 0.555146811 |
| Ros1 | 3.269051162 | 0.555146811 |
| Sh2b1 | 0.22114122 | 0.555146811 |
| Gm42791 | -1.962340525 | 0.555234127 |
| Arntl2 | 1.315394396 | 0.555314424 |
| Adal | 0.250735461 | 0.555314424 |
| Xkr4 | -0.42695149 | 0.555654003 |
| Cacng5 | 0.821995207 | 0.555941478 |
| Neurod6 | -0.25316273 | 0.555941478 |
| Tmco4 | 0.709075049 | 0.55621457 |
| Hist1h2ac | 0.694378629 | 0.55621457 |
| Zfp219 | 0.489142248 | 0.55621457 |
| Fam3a | 0.286999865 | 0.55621457 |
| Cul4a | -0.159622354 | 0.55621457 |
| Hmg20a | -0.161539081 | 0.55621457 |
| Htr2b | -0.932223432 | 0.55621457 |
| Ube2g1 | -0.176502981 | 0.556348674 |
| Cacna1b | -0.221843595 | 0.556348674 |
| Mfn1 | -0.231620665 | 0.556348674 |
| Gm10406 | 0.813504364 | 0.556421867 |
| Car15 | -2.70318641 | 0.556421867 |
| Nkain2 | -0.253171733 | 0.556509913 |
| Borcs6 | 0.315111107 | 0.556943984 |
| Trpc5 | 1.342619217 | 0.557111768 |
| Gm6685 | 1.30478789 | 0.557273516 |
| Ctu2 | -0.218517861 | 0.557896258 |
| Zfp451 | 0.191269675 | 0.557977075 |
| Ica1l | 0.302462895 | 0.558018536 |
| Nop56 | 0.178434664 | 0.558719691 |
| Gar1 | -0.264610291 | 0.558767982 |
| Mrpl54 | 0.173346155 | 0.559159417 |
| Cfap77 | -3.029914547 | 0.559159417 |
| 8030474K03Rik | 3.328999779 | 0.559185738 |
| B230104I21Rik | 3.328999779 | 0.559185738 |
| Zic2 | 0.819472717 | 0.559185738 |
| Usp1 | 0.202628575 | 0.559185738 |
| Wwp2 | -0.189049211 | 0.559185738 |
| Lactb | -0.336206201 | 0.559185738 |
| Magee1 | -0.209885478 | 0.559216981 |
| Gspt1 | -0.168431427 | 0.559366417 |

| | | |
|---|---|---|
| Pla2g5 | 0.297661983 | 0.937632709 |
| Sox10 | 0.26247606 | 0.937632709 |
| Pcdha1 | 0.247141038 | 0.937632709 |
| Fbxo27 | 0.145057955 | 0.937632709 |
| Tmem230 | 0.03415157 | 0.937632709 |
| Fbxo8 | -0.045684597 | 0.937632709 |
| Ccdc189 | 0.060851655 | 0.937749358 |
| Larp1b | 0.103342361 | 0.937756402 |
| Klhl42 | 0.051570748 | 0.937756402 |
| Chrnb2 | 0.031236996 | 0.937756402 |
| Itm2c | -0.028513267 | 0.937756402 |
| Osbpl2 | -0.036694731 | 0.937756402 |
| AA474331 | -0.097538048 | 0.937768774 |
| Dnaic2 | 0.163393387 | 0.937789615 |
| Disp2 | -0.048534733 | 0.937835635 |
| Tsacc | -0.11085377 | 0.937971847 |
| Jmy | 0.031789417 | 0.938798834 |
| Pqlc1 | -0.03536556 | 0.938798834 |
| Mpdu1 | 0.026600346 | 0.938932603 |
| Fam149b | -0.036314349 | 0.938932603 |
| Tmem37 | -0.04905717 | 0.938932603 |
| Arap2 | -0.117776737 | 0.938932603 |
| Adam33 | -0.233821808 | 0.938932603 |
| Gm4969 | 0.341804078 | 0.939157504 |
| Galnt7 | 0.047049394 | 0.939157504 |
| Klhl3 | 0.084303129 | 0.939198153 |
| Gm9923 | 0.057684357 | 0.939337887 |
| Casp1 | -0.056499292 | 0.939337887 |
| Zscan25 | -0.073158757 | 0.939337887 |
| Nr6a1 | -0.113964973 | 0.939337887 |
| Ahr | 0.106341345 | 0.939677032 |
| Snx25 | -0.085994746 | 0.940096327 |
| Fam76a | 0.034097514 | 0.940137615 |
| Atxn7l2 | -0.044207028 | 0.940137615 |
| Zfp619 | -0.148428038 | 0.940137615 |
| Hnmt | 0.121638607 | 0.940332422 |
| Pigo | 0.056327621 | 0.940332422 |
| Glmn | 0.045285566 | 0.940332422 |
| Srsf3 | 0.021122579 | 0.940332422 |
| Myom2 | -0.288314114 | 0.940332422 |
| Alkbh5 | -0.025920929 | 0.940469551 |
| Pglyrp2 | 0.315577401 | 0.940531958 |
| Hykk | 0.078582297 | 0.940531958 |
| Nlgn2 | 0.036237132 | 0.940531958 |
| Zfp414 | 0.026234212 | 0.940531958 |
| Srp72 | -0.02339394 | 0.940531958 |
| Tada3 | -0.030981214 | 0.940531958 |
| Palb2 | -0.086859833 | 0.940531958 |
| Pigq | -0.026567326 | 0.940638319 |
| Adam22 | -0.036019213 | 0.940790279 |
| Gm17509 | 0.351661029 | 0.940803459 |
| Gm3239 | 0.274791567 | 0.940849132 |
| Mark4 | 0.04312143 | 0.940849132 |
| Ntf5 | -0.328691848 | 0.940849132 |
| Xlr | 0.061440887 | 0.94088081 |
| Trp53rka | -0.031263204 | 0.94088081 |
| Aqp11 | -0.065139074 | 0.941105405 |
| Prkd1 | -0.079936753 | 0.941105405 |
| Dnttip2 | 0.024081429 | 0.941239516 |
| Itga3 | 0.068691865 | 0.941267803 |
| Vegfc | 0.177322634 | 0.941304329 |
| Grik4 | -0.077910476 | 0.941304329 |
| Kctd13 | -0.026327365 | 0.941385317 |
| Sec11a | -0.021202392 | 0.941517156 |
| Mphosph9 | -0.040698185 | 0.941567695 |
| Fam186a | -0.251018459 | 0.941567695 |
| Macrod2 | -0.040809267 | 0.941746219 |
| 4930451G09Ri | 0.201218735 | 0.941750597 |
| Tex2 | 0.036275607 | 0.941750597 |
| Pank3 | 0.027977274 | 0.941750597 |
| BC005624 | 0.023889257 | 0.941750597 |
| Dhdds | -0.027344888 | 0.941750597 |
| Katna1 | -0.030038047 | 0.941750597 |
| Nudcd1 | -0.033501971 | 0.941774337 |
| Srbd1 | -0.034948692 | 0.941774337 |
| Btbd9 | -0.021036472 | 0.941836185 |
| Zfp266 | 0.024572033 | 0.941937816 |
| Vezt | -0.030415609 | 0.941937816 |
| 5730455P16Ril | -0.033011238 | 0.941937816 |
| Zfp873 | -0.0401255 | 0.941937816 |
| Peli3 | -0.071784913 | 0.941937816 |
| Polrmt | 0.040946008 | 0.942209684 |
| Zfp809 | 0.067706601 | 0.942547323 |
| Zfyve19 | -0.030181269 | 0.943033528 |
| Fbxo33 | 0.056221378 | 0.943147256 |
| Mgmt | 0.054896678 | 0.943147256 |
| Prss36 | 0.049511779 | 0.943147256 |
| Gabpb2 | 0.038209113 | 0.943147256 |
| Draxin | 0.034324936 | 0.943147256 |
| Mecr | 0.028356581 | 0.943147256 |
| Safb | -0.024504203 | 0.943147256 |
| Clock | -0.027028619 | 0.943147256 |
| Klf2 | -0.052474035 | 0.943147256 |
| Adam1a | -0.111163907 | 0.943147256 |
| Bcl6b | -0.174468545 | 0.943147256 |
| Zhx1 | -0.029377136 | 0.943484273 |
| Kntc1 | -0.0558164 | 0.943649894 |
| Mms22l | 0.041292442 | 0.94383135 |
| Tbc1d17 | -0.028835483 | 0.94383135 |
| Sfrp4 | -0.101440647 | 0.94383153 |
| Capn12 | -0.207194709 | 0.943932788 |
| Rbbp7 | -0.023780869 | 0.944255175 |
| Gbp6 | -0.099726427 | 0.944314889 |
| Plk4 | 0.038623433 | 0.94446251 |
| Rfxap | 0.032054767 | 0.94446251 |
| Btbd8 | -0.073594811 | 0.94446251 |
| Abcg4 | 0.056296557 | 0.944585131 |
| Samd4b | 0.038096159 | 0.944585131 |
| Slc25a5 | 0.019699228 | 0.944585131 |
| Lin7b | -0.070737338 | 0.944585131 |
| Gm13090 | -0.172811806 | 0.944585131 |
| Amot | 0.048527244 | 0.945246356 |
| Itfg2 | 0.027307385 | 0.945246356 |
| Osbpl8 | -0.027303831 | 0.945246356 |
| Bbs12 | -0.048422654 | 0.945452113 |
| Aen | 0.029259632 | 0.945512319 |
| Plcb1 | -0.029017479 | 0.945512319 |
| Gimap1 | 0.191723497 | 0.945773552 |
| Igfbp6 | 0.094242509 | 0.945773552 |

| | | |
|---|---|---|
| Cabp4 | -2.612523714 | 0.012854472 |
| Tmprss6 | -6.226650864 | 0.012861227 |
| Chrd | 3.21004408 | 0.012892894 |
| Bin3 | -0.802320122 | 0.012904905 |
| Epb41l5 | 0.885276216 | 0.012906515 |
| Pcdhgb1 | -1.153356467 | 0.012906515 |
| Esyt1 | -1.268639732 | 0.012917273 |
| Nt5e | -5.074003152 | 0.012930409 |
| Kank1 | 5.629757997 | 0.012972457 |
| Gstt1 | 1.667206653 | 0.012972457 |
| Fezf1 | 5.648933592 | 0.012997404 |
| Dcbld1 | 0.877939338 | 0.012997404 |
| Cox8a | 0.531382468 | 0.01309033 |
| Sp100 | -1.479704207 | 0.013130215 |
| Hsdl2 | 0.971308018 | 0.01313092 |
| Tmem41b | -0.640665374 | 0.01313092 |
| Onecut2 | 1.157370594 | 0.013136997 |
| Apmap | -1.111336156 | 0.013149796 |
| Srpx | 5.704753228 | 0.01316125 |
| Rsbn1 | 0.909644889 | 0.013180914 |
| Ttbk2 | 1.023508018 | 0.013184364 |
| Saysd1 | 0.813964929 | 0.0131865 |
| Gfi1b | 5.984589375 | 0.013193088 |
| Vps26a | -0.57731903 | 0.013193088 |
| Stt3b | -0.642797174 | 0.01320504 |
| Nhp2 | 0.672002764 | 0.013249661 |
| Neu2 | -5.301190816 | 0.013249661 |
| Mgat2 | -0.578480119 | 0.013286953 |
| Larp6 | 1.492684687 | 0.013292205 |
| Cct2 | 0.498413424 | 0.013293082 |
| Tm4sf5 | 5.566465865 | 0.013298899 |
| Ube2q2 | -0.787090613 | 0.013298899 |
| 4931429I11Rik | 5.095361 | 0.013304632 |
| Npl | 1.012649339 | 0.013306658 |
| Sdpr | 2.280982558 | 0.013352423 |
| Rrp15 | 0.781780342 | 0.013359235 |
| Phactr2 | 0.79115015 | 0.013379408 |
| Slc38a5 | 2.095404757 | 0.013387962 |
| Ywhag | -0.596248627 | 0.013387962 |
| Pygo1 | 0.761324425 | 0.01342229 |
| Clic1 | -0.732941038 | 0.013426219 |
| Fam98b | 0.736259461 | 0.013427017 |
| Grk4 | 2.995774906 | 0.013443621 |
| 2310011J03Rik | -0.778651921 | 0.013443621 |
| Ermn | 5.714181266 | 0.013445993 |
| Kl | 5.658347241 | 0.013446255 |
| Med30 | 0.748737252 | 0.013446255 |
| Nabp2 | 0.630014589 | 0.013451175 |
| Pitpnm2 | -1.034689664 | 0.013465842 |
| Maneal | 1.909100749 | 0.013476459 |
| Clasrp | -0.865499249 | 0.01347646 |
| Emx2 | 1.535183917 | 0.013510868 |
| 9530053A07Rik | 4.997054132 | 0.013531059 |
| Slc8a3 | -1.32760482 | 0.013546504 |
| Uhmk1 | -0.893918633 | 0.013551818 |
| Trappc12 | -0.729791722 | 0.013559904 |
| Ank | -0.858398781 | 0.013575408 |
| Rbms1 | -0.98156942 | 0.013578773 |
| Dpysl5 | -0.619208525 | 0.013593115 |
| Rab11fip3 | -0.849718376 | 0.013619912 |
| Foxred2 | -1.062285928 | 0.013619912 |
| Macf1 | 0.536512704 | 0.013626928 |
| Acbd4 | -1.362083722 | 0.013666573 |
| Rpl36al | 0.489134856 | 0.013742913 |
| Cdc42bpb | -0.797496068 | 0.013751176 |
| Tmod4 | 5.53598196 | 0.01376434 |
| Preb | -0.679162066 | 0.013774 |
| Cntn4 | 2.400265156 | 0.013785081 |
| Gm5148 | 0.716581916 | 0.013787814 |
| Adap2 | -0.995455392 | 0.013792048 |
| Zdhhc21 | -0.757004754 | 0.013814166 |
| Fnip2 | -1.041671732 | 0.013832515 |
| Map2k5 | 0.654317748 | 0.013852571 |
| Rps20 | 0.555221325 | 0.013876151 |
| Tmem55a | 0.814128178 | 0.013941874 |
| Lmx1a | 2.607011544 | 0.014000454 |
| Dlst | -0.582927201 | 0.014000454 |
| Fam124a | 1.110812113 | 0.014016055 |
| Phip | 0.559603227 | 0.014016055 |
| Eed | -0.622524911 | 0.014031063 |
| Pmm2 | -0.640668401 | 0.014049015 |
| Oraov1 | -0.782491539 | 0.01408882 |
| Dpp4 | 5.703179204 | 0.014213076 |
| Nrxn1 | 0.742575248 | 0.014231209 |
| Pcdhb12 | 5.685905319 | 0.014259364 |
| Tenm1 | 1.30729406 | 0.014261673 |
| Mafl | 0.549071539 | 0.014280806 |
| Slit1 | -1.120142646 | 0.01430677 |
| Ncl | 0.481525392 | 0.014339643 |
| Scaf8 | -0.60678112 | 0.014344649 |
| Cldn6 | -6.318359941 | 0.01436041 |
| 0610012G03Rik | 0.556933901 | 0.014399219 |
| Rasl11a | 5.685370159 | 0.014441184 |
| Vamp2 | 0.610293805 | 0.014450327 |
| Rev1 | -0.675361382 | 0.014450327 |
| Mbd1 | -0.938744092 | 0.014450327 |
| Zfp709 | -1.231832735 | 0.014450327 |
| Bhlhe40 | -1.390401522 | 0.014450327 |
| Mapkapk3 | -2.024663816 | 0.014450327 |
| Wnk2 | -0.681549603 | 0.014463132 |
| Xlr4a | -5.978820408 | 0.014463132 |
| Psmc3 | 0.580809069 | 0.014468721 |
| Nup133 | -0.652442495 | 0.014468721 |
| Gm2a | -0.787953602 | 0.014482059 |
| Rps16 | 0.558524859 | 0.014482776 |
| Rab4a | 0.630661825 | 0.014489154 |
| Ppil3 | 0.985469666 | 0.014492751 |
| Tmem178b | 0.683747114 | 0.014524811 |
| Rftn2 | 0.976075004 | 0.014575127 |
| Calb2 | 2.1863708 | 0.014599458 |
| Enpep | 5.872896697 | 0.014601581 |
| Nup107 | -0.908822876 | 0.01461308 |
| 2900026A02Rik | -1.1030722 | 0.01461308 |
| Plk3 | -1.222500274 | 0.014627448 |
| Ncoa3 | -0.787817413 | 0.01463935 |
| Cpne7 | 5.506428387 | 0.014642256 |
| Gxylt2 | 2.554846278 | 0.014642256 |
| Etl4 | 1.862836343 | 0.014642256 |
| Znhit6 | 0.939576295 | 0.014642256 |

| | | |
|---|---|---|
| Eif3j2 | 0.524891503 | 0.196702256 |
| Gcnt1 | -1.010313107 | 0.196702256 |
| Matn3 | -2.869588465 | 0.196702256 |
| 1700018B08Rik | -5.190669229 | 0.196702256 |
| Slfl | 0.420795324 | 0.196936465 |
| Slamf9 | -1.534400309 | 0.196963008 |
| Slc37a3 | 0.474751112 | 0.197134032 |
| Acot12 | -5.181077504 | 0.197134032 |
| Scara3 | 1.022236393 | 0.197230755 |
| Rhobtb2 | -0.418060659 | 0.197230755 |
| Kirrel3 | -0.689158316 | 0.197730819 |
| Plag1 | -0.794549569 | 0.197730819 |
| Tmem45a | 4.791520337 | 0.197778106 |
| Pnisr | 0.3259699 | 0.197778106 |
| Rab22a | -0.392835226 | 0.197778106 |
| Btbd1 | -0.39509547 | 0.197778106 |
| Pigm | -0.424169317 | 0.197778106 |
| Prpf4 | -0.460931487 | 0.197806425 |
| Gypa | 1.341466558 | 0.19785152 |
| Rrbp1 | -0.429151627 | 0.19785152 |
| Cdan1 | -0.799117617 | 0.197865465 |
| Ythdf3 | -0.36148874 | 0.197942153 |
| Utp18 | 0.321627586 | 0.197981807 |
| Clasp2 | 0.310036913 | 0.198003161 |
| Kcnn3 | -0.57042055 | 0.198003161 |
| Lmnb1 | -0.295470328 | 0.198136977 |
| Klk4 | 4.659477699 | 0.198175835 |
| Champ1 | 0.454060907 | 0.198284053 |
| Bri3 | -0.403921611 | 0.198284053 |
| Cep68 | -0.446641949 | 0.198284053 |
| Fam102a | -0.711642999 | 0.198284053 |
| Trmt61a | -0.642691555 | 0.198350911 |
| Heph | 2.082546733 | 0.19838202 |
| Zfp433 | 1.640918524 | 0.198488705 |
| Gpr182 | 4.265008205 | 0.198630867 |
| Fip1l1 | -0.274158692 | 0.198630867 |
| Kdm6a | -0.401452217 | 0.198630867 |
| Mad2l1bp | 0.493295727 | 0.198782506 |
| H2afj | 0.370994193 | 0.198782506 |
| Phykpl | -0.421497254 | 0.19878801 |
| Pwwp2b | -1.019448028 | 0.19878801 |
| 1700125H20Rik | -5.171455855 | 0.198958742 |
| Cyr61 | 0.812295467 | 0.199033701 |
| Sv2a | 0.559359602 | 0.199037728 |
| Blm | 0.546083101 | 0.199037728 |
| Tpp2 | -0.373930187 | 0.199040297 |
| Dcaf8 | -0.306562911 | 0.19932455 |
| Eif5a | 0.270821322 | 0.199474532 |
| Tor1a | -0.317259532 | 0.199474532 |
| Hist1h3d | -1.373038571 | 0.199474532 |
| Pef1 | 0.39772539 | 0.19952102 |
| Setx | 0.418306588 | 0.199611183 |
| Mfng | 0.510813994 | 0.19981907 |
| 2610301B20Rik | 0.378823212 | 0.199961642 |
| Gm9830 | 4.253234638 | 0.200053662 |
| Col9a2 | 1.709174981 | 0.200055322 |
| Cdk2ap1 | 0.359750142 | 0.200088553 |
| Caln1 | 0.733241089 | 0.200133113 |
| Tfip11 | -0.489475444 | 0.200263444 |
| Prkd2 | 0.95875635 | 0.200309526 |
| Atad1 | 0.363451518 | 0.200746657 |
| H2-M3 | -0.975943589 | 0.200746657 |
| Adora3 | -1.164959137 | 0.200771915 |
| Rbak | -0.743519987 | 0.200807546 |
| Dbh | -3.405721538 | 0.200823271 |
| Rab36 | 0.650159152 | 0.200919175 |
| Itpr2 | -0.756566926 | 0.20094684 |
| Fbxw2 | -0.291999145 | 0.200962523 |
| Setd1a | -0.51712748 | 0.201119158 |
| Ccnjl | -0.748717795 | 0.201179251 |
| Mapk1 | -0.269957423 | 0.201186906 |
| Sppl3 | 0.45653749 | 0.201528413 |
| Prex2 | 1.26136356 | 0.201648885 |
| Psmd2 | 0.236596569 | 0.20168087 |
| Gdi2 | -0.309519557 | 0.201819636 |
| Kit | -0.764615896 | 0.201993937 |
| Rtf1 | 0.331436576 | 0.20238586 |
| Arhgef37 | -2.598427895 | 0.20238586 |
| D6Wsu163e | -0.510516183 | 0.202389674 |
| Pak4 | -0.54271934 | 0.20244107 |
| Mettl6 | -0.369790814 | 0.202498846 |
| Pdzk1 | 4.474621384 | 0.20270049 |
| Ppp1r36 | 4.474621384 | 0.20270049 |
| Csnk2a1 | -0.281995158 | 0.20270049 |
| Acap3 | -0.411837674 | 0.20271541 |
| C8b | -5.161791103 | 0.20271541 |
| Gm438 | -4.421127927 | 0.202773491 |
| Zfp60 | 0.523380174 | 0.203282038 |
| Creb3 | -0.32435529 | 0.203282038 |
| Cnot2 | -0.347167841 | 0.203282038 |
| Snn | -0.3919418 | 0.203282038 |
| Nipal2 | 3.673757741 | 0.203399421 |
| Rsad2 | -2.523476323 | 0.203469787 |
| Rfc1 | 0.394687275 | 0.203504343 |
| Efcab6 | 1.320600572 | 0.203514692 |
| Cep128 | -0.778511841 | 0.203561154 |
| Sgta | 0.29311139 | 0.203731211 |
| Gadl1 | 0.446218757 | 0.203901916 |
| Cd4 | -2.58517746 | 0.204051464 |
| Cdkn2b | 4.662494755 | 0.204100656 |
| Ttk | 0.747089976 | 0.20411541 |
| Arhgap8 | -3.069381938 | 0.20411541 |
| Olfr70 | 4.235029181 | 0.204275024 |
| Rnf40 | -0.451381993 | 0.204482469 |
| Xrcc3 | -0.772208066 | 0.204482469 |
| Spata3 | 4.610210153 | 0.204647573 |
| Speer4b | 2.208917303 | 0.204647573 |
| Fam92a | 0.351658554 | 0.204647573 |
| Kcna2 | -1.09295281 | 0.204647573 |
| Gripap1 | -0.330869396 | 0.204693911 |
| Maea | -0.46773999 | 0.20479313 |
| Ociad1 | -0.255188093 | 0.204802898 |
| Itk | -5.169818835 | 0.204802898 |
| Ankrd10 | 0.329585091 | 0.204820115 |
| Eml4 | -0.378885452 | 0.204820115 |
| Golph3l | -0.395665 | 0.205011364 |
| Stap2 | 1.904425635 | 0.205061867 |
| Slc39a9 | -0.579169234 | 0.205061867 |
| Ccdc53 | 0.382640151 | 0.205178327 |

| | | |
|---|---|---|
| Mipol1 | -0.376819433 | 0.559368125 |
| Dtnbp1 | 0.214543613 | 0.559710918 |
| Dhx40 | -0.177004586 | 0.559710918 |
| Mettl24 | 2.623149927 | 0.559855091 |
| Ppp1r3e | 0.640540544 | 0.559855091 |
| Efna4 | 0.449422398 | 0.559855091 |
| Apool | -0.304335529 | 0.559855091 |
| Itga9 | -0.422342063 | 0.559855091 |
| Gm5141 | -0.366512041 | 0.560153868 |
| Gm11149 | 0.691503987 | 0.560256026 |
| Osgepl1 | 0.360345968 | 0.560256026 |
| Myl6b | -0.712781844 | 0.560256026 |
| Cth | 0.736074734 | 0.560257499 |
| Gdi1 | 0.15140719 | 0.560257499 |
| Txlna | -0.233693785 | 0.56026668 |
| Slc24a4 | 1.417646964 | 0.560490857 |
| Gpr151 | 0.972505241 | 0.560490857 |
| Dnajb3 | 0.415565648 | 0.560490857 |
| 2610008E11Rik | 0.249116604 | 0.560490857 |
| Ccdc122 | -1.098677006 | 0.560490857 |
| Tubb5 | 0.127388774 | 0.560757416 |
| Slc6a20b | -0.71420663 | 0.560782943 |
| Dhx35 | 0.294224528 | 0.560847241 |
| Tmem106c | -0.178099895 | 0.560847241 |
| Pot1b | -0.359891142 | 0.560847241 |
| Tmem177 | 0.359072426 | 0.560895759 |
| Slitrk3 | -0.380427937 | 0.561008748 |
| Sox13 | -0.676119348 | 0.561043072 |
| Kcnip2 | 0.476184673 | 0.56118106 |
| B3gnt7 | -0.822898808 | 0.56118106 |
| Oplah | -0.342695908 | 0.561423884 |
| Vnn3 | -0.940675624 | 0.561423884 |
| Clstn1 | 0.203683819 | 0.561626971 |
| Ano10 | 0.250467088 | 0.562202031 |
| Trpv1 | 3.220366042 | 0.562716516 |
| Gpr101 | 2.850360991 | 0.563121916 |
| Cby3 | -1.852426259 | 0.563121916 |
| Zcchc3 | 0.239571627 | 0.563215469 |
| Tram2 | -0.335933139 | 0.563215469 |
| Crebl2 | -0.542625234 | 0.563215469 |
| Arr3 | 3.258083219 | 0.563286028 |
| Ly6g5c | 3.258083219 | 0.563286028 |
| Sycp3 | 2.543079812 | 0.563372467 |
| Gpr45 | 0.263983426 | 0.563518991 |
| Vwa5b2 | -0.682398574 | 0.56381724 |
| Smim22 | -1.446059211 | 0.563820478 |
| B4galt2 | -0.211240566 | 0.563997929 |
| Dnajb1 | 0.205779536 | 0.564240484 |
| Aplf | -0.392582594 | 0.564240484 |
| Zkscan14 | -0.31522284 | 0.564348319 |
| Rbmxl1 | 0.166804087 | 0.564626217 |
| Tmem169 | -0.225683686 | 0.56470549 |
| Nhlrc1 | -0.511538095 | 0.56470549 |
| Rnase10 | 1.68125436 | 0.564833158 |
| Gm6658 | -1.112298533 | 0.564833158 |
| Sphk1 | 1.480676407 | 0.56501989 |
| Fry | -0.199358783 | 0.56501989 |
| Cacnb1 | -0.265726551 | 0.565079827 |
| Pcdhb2 | -0.839062785 | 0.565143438 |
| Tmem82 | 1.894052746 | 0.56530045 |
| Ralgapa1 | -0.177192022 | 0.56530045 |
| Ang | 0.530018976 | 0.565360999 |
| Cep192 | -0.215623444 | 0.565360999 |
| Zbtb18 | -0.276041697 | 0.565379189 |
| Tomm40 | 0.15542525 | 0.565427159 |
| BC049730 | 2.8258228 | 0.565468269 |
| Sar1b | 0.152761442 | 0.565712521 |
| Laptm4a | 0.1347232 | 0.565894353 |
| Eif3l | -0.138934961 | 0.56608913 |
| Shc3 | -0.549599891 | 0.56608913 |
| Ptk7 | -0.256902047 | 0.566589597 |
| Zfand3 | 0.218999769 | 0.567252014 |
| Sorbs1 | 0.194311054 | 0.567564187 |
| Fbxo41 | -0.46360201 | 0.567570185 |
| C8g | -0.691888708 | 0.567570185 |
| Kctd15 | 0.313326228 | 0.567749544 |
| Lypd6b | 0.724555714 | 0.567811481 |
| Nnmt | 1.164150711 | 0.567954539 |
| Phf12 | -0.209650999 | 0.568143003 |
| 1700123O20Rik | 0.228673861 | 0.568201603 |
| Tmed4 | 0.167295071 | 0.568201603 |
| Immt | 0.143056774 | 0.568201603 |
| 9630041A04Rik | -1.045617703 | 0.568201603 |
| Hspa1l | 1.22238372 | 0.568247154 |
| Mboat2 | 0.183830567 | 0.568437114 |
| Tnrc6b | 0.174175533 | 0.568625816 |
| Ythdc2 | 0.21956207 | 0.568832612 |
| Mbtps2 | 0.239989797 | 0.569176942 |
| Il17rc | -0.927300294 | 0.569176942 |
| Helt | -1.111992361 | 0.569176942 |
| Pdss1 | 0.301093448 | 0.569389136 |
| Gm5595 | 0.533172375 | 0.569486473 |
| Gm7030 | -0.506003553 | 0.569486473 |
| Kcnk13 | -0.69040493 | 0.569587623 |
| Tmprss5 | 1.214924007 | 0.569788171 |
| Haus4 | -0.267008764 | 0.56980635 |
| Skap1 | 0.878705429 | 0.569965027 |
| Amacr | 0.392131293 | 0.569965027 |
| Gm9774 | 0.38371094 | 0.569965027 |
| Lsm11 | 0.245963568 | 0.569984676 |
| Fam170a | 3.247013202 | 0.570390249 |
| Vmn2r49 | 3.247013202 | 0.570390249 |
| Zfp872 | 3.247013202 | 0.570390249 |
| Ftcd | 3.227746392 | 0.570390249 |
| Stk36 | 0.318590702 | 0.570390249 |
| Ngly1 | -0.243748172 | 0.570390249 |
| Lmf2 | -0.287770998 | 0.570390249 |
| Lmf1 | -0.296288507 | 0.570390249 |
| Kif7 | -0.514369815 | 0.570605645 |
| Mbtd1 | 0.187668869 | 0.570931509 |
| Chn1 | -0.215043569 | 0.570931509 |
| Fam114a1 | -0.415115825 | 0.570969585 |
| Snai1 | 1.047405602 | 0.571065746 |
| Brpf3 | -0.472226746 | 0.571240518 |
| Ier5l | 0.590792932 | 0.571326902 |
| Mtx3 | -0.368073371 | 0.571401204 |
| Fam166b | -1.114537847 | 0.571401204 |
| Cldn18 | 3.204052947 | 0.571422547 |
| Tssk2 | 3.204052947 | 0.571422547 |

| | | |
|---|---|---|
| Aldh1b1 | 0.044781057 | 0.945773552 |
| Jph4 | 0.027759769 | 0.945773552 |
| Kcnb1 | -0.057829042 | 0.945773552 |
| Capsl | -0.108315511 | 0.945773552 |
| Gsg1 | -0.18836378 | 0.945773552 |
| Galnt12 | 0.088765066 | 0.945851393 |
| Tmem178 | 0.040469817 | 0.945851393 |
| Nap1l2 | 0.030310319 | 0.945851393 |
| Fam173b | -0.038525892 | 0.945851393 |
| Nrtn | -0.121366091 | 0.945851393 |
| Wfdc18 | -0.348198299 | 0.945851393 |
| Tbce | 0.035404146 | 0.946083984 |
| Gbf1 | -0.024231327 | 0.946083984 |
| Add2 | -0.02677015 | 0.94639398 |
| Pdcd7 | -0.038840778 | 0.94639398 |
| Zfp691 | 0.045455362 | 0.946483157 |
| Zfyve21 | 0.029861047 | 0.946483157 |
| Zfp207 | -0.017202975 | 0.946483157 |
| Retsat | -0.045701797 | 0.946525967 |
| Haao | -0.088659589 | 0.946525967 |
| Grem1 | 0.137152994 | 0.946551423 |
| Ncan | 0.02232444 | 0.946656637 |
| Pde7b | -0.065309574 | 0.946658181 |
| Cstad | 0.130876263 | 0.946732529 |
| Epb41l3 | 0.035486524 | 0.946737932 |
| Psd3 | -0.036320469 | 0.946737932 |
| Golga4 | 0.022301309 | 0.94680531 |
| BC005537 | -0.02048171 | 0.94680531 |
| Mab21l1 | -0.10493431 | 0.94680531 |
| Calm4 | -0.226248588 | 0.9468183 |
| Napa | 0.01798817 | 0.946818474 |
| Nqo2 | -0.045972438 | 0.946818474 |
| Unc50 | -0.021872799 | 0.94687728 |
| Usp35 | 0.041738248 | 0.947051283 |
| Cdadc1 | -0.028430534 | 0.947170413 |
| Spon1 | -0.079703671 | 0.947545636 |
| Chdh | -0.289891236 | 0.947584753 |
| Pxdn | 0.040288058 | 0.947714463 |
| Uri1 | 0.023411032 | 0.947714463 |
| Mapre1 | -0.016225073 | 0.947714463 |
| Unkl | -0.07160826 | 0.947714463 |
| Cadm3 | 0.029349816 | 0.947723438 |
| Armcx3 | 0.028242902 | 0.947777306 |
| Cox10 | 0.039298707 | 0.94789226 |
| Gbe1 | -0.044491248 | 0.94789226 |
| Slc30a6 | -0.045987577 | 0.94789226 |
| Acaa1b | -0.076953172 | 0.94789226 |
| Arl2bp | 0.0221376 | 0.947900358 |
| Fmo5 | -0.137112054 | 0.947900358 |
| Cspp1 | -0.032329781 | 0.948166189 |
| Peli2 | 0.043109647 | 0.948296456 |
| Crmp1 | -0.020093511 | 0.948296456 |
| Cdc73 | -0.028385512 | 0.948367467 |
| Lztr1 | -0.021633968 | 0.948804318 |
| Grtp1 | 0.033107893 | 0.94951517 |
| Ift52 | -0.030471143 | 0.949658748 |
| Igf1r | -0.021038061 | 0.949670313 |
| Def6 | -0.039177786 | 0.949670313 |
| Cd86 | 0.035420753 | 0.949889323 |
| Fam210a | 0.027924134 | 0.950016717 |
| Akr1c13 | -0.157095185 | 0.950016717 |
| Zfp64 | -0.027550378 | 0.950265397 |
| Tcf7l1 | -0.05334049 | 0.950338768 |
| Mst1 | -0.186878908 | 0.950341845 |
| Arl4d | -0.04596844 | 0.95072012 |
| Slc35e3 | -0.041038121 | 0.950774914 |
| Podnl1 | 0.2499541 | 0.950918189 |
| Pecr | 0.04620697 | 0.950918189 |
| Tmem198 | 0.028285895 | 0.950918189 |
| Pcbp3 | 0.030878909 | 0.951067719 |
| Fam131a | -0.033765141 | 0.951185348 |
| Wdr33 | -0.018484487 | 0.951224135 |
| Ppp1r3f | -0.068530231 | 0.951260155 |
| Trmt11 | -0.041910301 | 0.951281231 |
| Ppt2 | -0.037479006 | 0.951339014 |
| Dph2 | -0.045472725 | 0.951873146 |
| Ash2l | -0.023230215 | 0.951875195 |
| Trim47 | -0.047796528 | 0.952012571 |
| Nap1l4 | -0.017239739 | 0.952030529 |
| Nsd1 | 0.017597641 | 0.952309548 |
| Ado | -0.019790271 | 0.952309548 |
| Sugct | 0.08401961 | 0.952609067 |
| Scd1 | -0.078242982 | 0.952609067 |
| Rab25 | 0.201640461 | 0.952618866 |
| Nubp1 | -0.038025879 | 0.952618866 |
| Klhl25 | 0.037420013 | 0.952786796 |
| Gm14288 | -0.123888621 | 0.952786796 |
| Phf20 | -0.019082003 | 0.952986439 |
| Cdkl2 | -0.05203557 | 0.953167659 |
| Gapt | 0.101975946 | 0.953179554 |
| Neurl2 | 0.051444627 | 0.953179554 |
| Gigyf2 | 0.025690454 | 0.953179554 |
| Ier2 | -0.020581969 | 0.953179554 |
| 0610030E20Ril | -0.027481888 | 0.953179554 |
| Gm527 | 0.104872474 | 0.95325408 |
| Vapb | -0.024698387 | 0.953692596 |
| Zbtb8os | 0.033815257 | 0.953992896 |
| Ube2l3 | 0.016831893 | 0.954055555 |
| Tubb4b | 0.01926744 | 0.954062279 |
| Ccdc78 | -0.06397245 | 0.954062279 |
| Mutyh | 0.074478996 | 0.954110117 |
| Plpp7 | -0.040046317 | 0.954110117 |
| Hist2h3c1 | -0.046993543 | 0.954260682 |
| 2610002M06Ri | 0.023403928 | 0.954353727 |
| Ywhah | 0.015370189 | 0.954353727 |
| Adamts4 | 0.09376897 | 0.954425824 |
| Zfp184 | 0.035996648 | 0.954425824 |
| Sf3b4 | -0.01941482 | 0.954425824 |
| Acad8 | -0.029809199 | 0.954425824 |
| Gin1 | -0.046923233 | 0.954425824 |
| Supv3l1 | 0.029535456 | 0.954492148 |
| Polr2b | 0.025050294 | 0.954624773 |
| Dgcr14 | 0.024281945 | 0.954624773 |
| Amph | -0.021938981 | 0.954624773 |
| Hecw1 | -0.029028061 | 0.954624773 |
| Tbc1d9 | 0.030563169 | 0.954733441 |
| Cyb5rl | 0.050596501 | 0.954739351 |
| Uspl1 | 0.022356282 | 0.955046525 |
| 4921509C19Ri | 0.219496793 | 0.955206535 |

| | | |
|---|---|---|
| Car13 | -1.752715232 | 0.014642256 |
| Zfp429 | -2.71619702 | 0.014642256 |
| Trhr | -3.49903367 | 0.014668554 |
| Srpk2 | 0.465258189 | 0.014678099 |
| Rmi2 | -1.502844294 | 0.014703873 |
| Crocc | -1.401933711 | 0.014742604 |
| Aga | -1.086591404 | 0.014750502 |
| Pmaip1 | -0.988754658 | 0.014769919 |
| Atp5c1 | 0.527054981 | 0.014805359 |
| AA386476 | -5.0635689 | 0.014805359 |
| Uxt | 0.727144807 | 0.014809361 |
| Rasgef1b | -0.711894949 | 0.014826341 |
| Gabarapl2 | 0.466826519 | 0.014842209 |
| Plxna2 | -0.740079652 | 0.014857675 |
| Mad2l2 | 0.554250666 | 0.014857997 |
| Asph | -0.645782073 | 0.014868291 |
| Tipin | 0.614361743 | 0.014905087 |
| Kcns2 | -3.387303129 | 0.014911568 |
| Stx19 | -3.501172692 | 0.014918983 |
| Timm44 | 0.991081801 | 0.014938118 |
| Islr | 1.75379805 | 0.014958224 |
| Batf | -1.155963027 | 0.014962672 |
| Cyp2b19 | 3.621182052 | 0.014978505 |
| Arl10 | -0.754942956 | 0.014999091 |
| Ndfip2 | -0.667316854 | 0.015053688 |
| Mbd3l2 | -6.3123336 | 0.015065831 |
| Synj2bp | -0.79967208 | 0.015077244 |
| Gatsl3 | -2.214901082 | 0.015077244 |
| Scrn2 | 1.730048368 | 0.015178072 |
| Ercc6 | 1.202205987 | 0.015227386 |
| Knstrn | 1.023239819 | 0.015227386 |
| Cd300lh | -4.236616568 | 0.015227386 |
| Gja4 | 5.551527935 | 0.015229066 |
| Dip2a | -0.831376645 | 0.015233795 |
| Mycbpap | 2.593158663 | 0.01529333 |
| Veph1 | 1.497901198 | 0.01529333 |
| Impa1 | -0.58612923 | 0.015328124 |
| D930015E06Rik | -0.616275154 | 0.015338378 |
| Prss30 | -2.636291078 | 0.015345114 |
| Grk6 | -0.79176683 | 0.015346269 |
| Ccdc181 | 1.043005566 | 0.015365025 |
| Igf2bp1 | -0.58996547 | 0.015365025 |
| Mt2 | 1.012348333 | 0.015399661 |
| Pcp4 | 1.465867059 | 0.015441821 |
| Lat | -4.244710392 | 0.015481792 |
| Ddx19b | -1.071350379 | 0.015494432 |
| Git2 | -0.608245553 | 0.015545073 |
| Map3k12 | -0.772953085 | 0.015562521 |
| Pts | 0.629783492 | 0.015572931 |
| Arpin | 0.943388736 | 0.015596825 |
| Znhit3 | 0.653142723 | 0.015626271 |
| Cd14 | -2.647241479 | 0.015626271 |
| Rnaseh2a | 0.668796204 | 0.015666873 |
| Phactr4 | 0.944573923 | 0.015701763 |
| Hexa | -0.784276758 | 0.015701932 |
| Itgb3bp | 0.902422157 | 0.015730691 |
| Grb10 | 0.986149176 | 0.015756587 |
| Psap | -0.520204931 | 0.015756587 |
| Lemd2 | -0.879891696 | 0.015769891 |
| Cfap161 | 5.550723959 | 0.0157843 |
| Scarb1 | -0.697698687 | 0.015837353 |
| Mesdc1 | -0.957390773 | 0.015914598 |
| Pgm2 | -0.903667552 | 0.015924274 |
| Parp14 | -1.930313319 | 0.015924274 |
| Chtf8 | -0.636089187 | 0.015935909 |
| Slc35b4 | -0.86187855 | 0.015935909 |
| Spp2 | 5.904475365 | 0.015961508 |
| Hnrnpa2b1 | 0.424155678 | 0.015961508 |
| Zc3h13 | 0.662205513 | 0.015969575 |
| Ccdc157 | 1.270549009 | 0.015970629 |
| Lhfpl2 | -1.094808413 | 0.015970629 |
| Cntnap3 | 5.732246849 | 0.016086145 |
| Nlgn1 | -1.037335998 | 0.016116948 |
| Enho | 0.672785427 | 0.016133083 |
| Hist1h1a | 2.3625653 | 0.016205021 |
| Nudt4 | -0.702346679 | 0.016233894 |
| Zfp623 | 0.857090921 | 0.016278651 |
| Fbxo18 | -0.618465208 | 0.016286418 |
| Lims1 | -0.607995083 | 0.016347601 |
| Ppil4 | 0.524286093 | 0.016367941 |
| Spg20 | -1.050751767 | 0.016367941 |
| Tut1 | -0.87980033 | 0.016380556 |
| Tmem229b | -0.824649409 | 0.016385939 |
| Cdc20 | 0.879710934 | 0.016391003 |
| Cyp46a1 | 2.172530929 | 0.016392772 |
| Sec62 | 0.590516546 | 0.016418782 |
| Zfp511 | 0.694193293 | 0.016442691 |
| Arl8b | -0.500538202 | 0.016442691 |
| Mdga1 | -0.758944231 | 0.016442691 |
| 4930523C07Rik | -2.070147635 | 0.016442691 |
| Ngfr | 2.709343591 | 0.016444839 |
| Cdk18 | -2.545191187 | 0.016444839 |
| Sesn3 | 0.811930048 | 0.016447669 |
| Timm10 | 0.777351729 | 0.016447669 |
| Papd4 | -0.630813788 | 0.016453692 |
| Prss41 | 5.673596908 | 0.016454543 |
| Serac1 | -1.526625606 | 0.016481927 |
| Evi5l | 0.608504641 | 0.016524754 |
| Ubtd2 | -0.699157492 | 0.016524754 |
| Cbl | -0.700943034 | 0.016524754 |
| Mlh3 | -1.473859475 | 0.016527598 |
| Nfyb | 0.579455481 | 0.016535943 |
| Sf3a2 | -0.741644844 | 0.016577729 |
| Bsx | 5.528736223 | 0.016591298 |
| Gata2 | -3.781080936 | 0.016594195 |
| Mtss1l | 1.045123361 | 0.016601163 |
| Cd101 | -2.203162415 | 0.016625268 |
| Orai1 | -0.884713128 | 0.016681997 |
| Ei24 | -0.555850494 | 0.016684936 |
| Rcor3 | 0.996800977 | 0.016736233 |
| Tyw5 | 1.014124254 | 0.016774082 |
| Ntpcr | -1.067764494 | 0.016793612 |
| Rlim | -0.604226075 | 0.016818195 |
| Dbt | 1.134706947 | 0.016854872 |
| Rpl3l | -6.168256747 | 0.016920001 |
| Aunip | -1.475495699 | 0.016923912 |
| Dennd3 | -2.307583944 | 0.016935847 |
| Kmo | -4.492157878 | 0.016945255 |
| Mgat4b | -0.857822757 | 0.016991251 |
| Slain2 | -0.414793673 | 0.205226815 |
| Thada | 0.674435946 | 0.205261146 |
| Dbndd2 | 1.419880426 | 0.205279827 |
| Gdf11 | 0.503466902 | 0.205279827 |
| Pspc1 | 0.352076857 | 0.205279827 |
| Rnf111 | -0.327773437 | 0.205279827 |
| C1qc | -0.507051784 | 0.205279827 |
| Icmt | -0.883970906 | 0.205279827 |
| Tmem171 | 4.755231285 | 0.205408864 |
| Gm19426 | 1.304159354 | 0.205408864 |
| Abcf3 | 0.378640306 | 0.205560831 |
| Tmem263 | 0.326909384 | 0.205685417 |
| Rxrg | 1.222602798 | 0.205833607 |
| Pdha1 | -0.351980513 | 0.205975177 |
| Ankrd24 | -0.630947294 | 0.205975177 |
| Pla2g16 | 0.631757807 | 0.206006724 |
| Mndal | -1.147958575 | 0.206099425 |
| Slc6a19 | -5.144518684 | 0.206248572 |
| Lair1 | -0.513963647 | 0.2062804 |
| Irs4 | 3.69795537 | 0.206453659 |
| Krt28 | 4.360515579 | 0.20653173 |
| Nr0b1 | 4.360515579 | 0.20653173 |
| 2410016O06Rik | 0.674679849 | 0.20653173 |
| H2-DMa | 0.993359476 | 0.206620955 |
| Cmc2 | 0.515254584 | 0.206633331 |
| Efcab1 | 0.808071185 | 0.206685886 |
| Pgbd5 | 0.751446562 | 0.206685886 |
| Ccdc112 | 0.709137416 | 0.206982906 |
| Akap11 | 0.361414345 | 0.207084825 |
| Tmem201 | -0.509404821 | 0.207084825 |
| Stk35 | -0.572120494 | 0.207418209 |
| Syap1 | -0.364757177 | 0.207613347 |
| Cct6b | 4.315824742 | 0.20769434 |
| Thoc5 | -0.381039498 | 0.20785367 |
| Fscn2 | 2.425597857 | 0.207961124 |
| Agm | 0.342065548 | 0.207961124 |
| Ldoc1 | 1.523947078 | 0.208071327 |
| Epb41l1 | -0.371420062 | 0.208079676 |
| Fmn2 | 0.586933108 | 0.20811431 |
| Hic1 | 3.579777674 | 0.20827433 |
| Ryr3 | -0.759194988 | 0.208401763 |
| Med1 | -0.289144069 | 0.208410151 |
| Zfp229 | -0.714030552 | 0.208411227 |
| Srsf7 | 0.250027769 | 0.208415406 |
| Cabyr | -1.812174115 | 0.208459841 |
| Dclre1b | -0.760742215 | 0.208730536 |
| Pcdhb6 | -1.052909385 | 0.208862433 |
| Rps6kb1 | -0.325618017 | 0.208867101 |
| Far2 | 0.91766831 | 0.208904338 |
| Zfp131 | 0.410122373 | 0.208904338 |
| Dnmt1 | -0.390854604 | 0.208904338 |
| Idnk | 0.383348705 | 0.208911941 |
| Oas3 | -2.343170423 | 0.208911941 |
| Slc26a10 | 4.215707424 | 0.208933708 |
| Nr1h2 | -0.420627698 | 0.20897555 |
| Slc19a1 | -0.625207163 | 0.208998981 |
| Dopey1 | -0.492037499 | 0.209129823 |
| Crkl | -0.354853123 | 0.209224512 |
| Mrgprf | 4.226909062 | 0.209293679 |
| Sall4 | 4.226909062 | 0.209293679 |
| Fam46c | 1.009887333 | 0.209293679 |
| Cdkn1b | 0.373207565 | 0.209358992 |
| Adcy8 | -1.838167332 | 0.209358992 |
| Rpf2 | 0.415972798 | 0.20946835 |
| Plekhg2 | -0.53144898 | 0.209535001 |
| Ccdc109b | -0.844561811 | 0.209595555 |
| Bax | 0.281845857 | 0.20985856 |
| Poc5 | 0.419379677 | 0.209902944 |
| Phlpp1 | 0.64503781 | 0.2099278 |
| Chst15 | -0.613058044 | 0.209957356 |
| Efna3 | 0.674549117 | 0.209970873 |
| Pcif1 | 0.305462696 | 0.210064071 |
| Dnase2b | -5.118833882 | 0.210069986 |
| Lig1 | 0.416054069 | 0.210072352 |
| Slc46a1 | -0.551793564 | 0.210139297 |
| Osbp2 | -0.915668332 | 0.210139297 |
| Csrnp2 | 0.496607392 | 0.210258502 |
| Adam32 | 4.201817302 | 0.21033671 |
| Dennd1b | -0.761994832 | 0.21059936 |
| Vwa2 | 4.430326784 | 0.210612743 |
| Epyc | 4.200447233 | 0.210612743 |
| Ppp1r14a | 1.243468121 | 0.210685002 |
| Mid1 | -0.461205608 | 0.210915777 |
| Dmd | 0.511046444 | 0.211027305 |
| Smim19 | 0.617030512 | 0.211863955 |
| Uhrf1bp1l | -0.331569258 | 0.211896203 |
| Nup188 | -0.449942722 | 0.211923481 |
| Scand1 | 0.403568415 | 0.211992894 |
| G2e3 | 0.40034132 | 0.211992894 |
| Kdm3a | -0.42307992 | 0.211992894 |
| Ercc6l | -0.764778053 | 0.212083244 |
| Ppp1r3c | 1.912401719 | 0.212245374 |
| Rtca | 0.316297539 | 0.212410895 |
| Tbcel | -1.036362296 | 0.212410895 |
| Emc1 | -0.392653931 | 0.21241144 |
| Jup | 0.404955892 | 0.212461102 |
| Akap1 | -0.646020115 | 0.212483986 |
| Eif3c | 0.288008168 | 0.212585217 |
| Rtn2 | -0.445526609 | 0.212978239 |
| Vamp3 | -0.295781995 | 0.213008478 |
| Ankzf1 | -0.749110638 | 0.213008478 |
| Psmd14 | -0.267079197 | 0.213154864 |
| A730071L15Rik | 4.35923685 | 0.213228493 |
| Gm14698 | 0.735851933 | 0.213311346 |
| Fggy | -0.709368645 | 0.213339487 |
| Vcp | -0.242456671 | 0.213347489 |
| Arl15 | -0.591709921 | 0.213347489 |
| Tmem176b | -0.583555552 | 0.213422229 |
| Arl2 | 0.403724332 | 0.21343201 |
| Psmd5 | 0.314525041 | 0.21343201 |
| Creb1 | 0.420487026 | 0.213570863 |
| Map1b | 0.320460439 | 0.213625165 |
| Atp5f1 | 0.242243574 | 0.213625165 |
| Pld3 | -0.291536623 | 0.213625165 |
| Agtrap | -0.776297585 | 0.213625165 |
| Dok2 | -1.642190937 | 0.213783923 |
| Rpl39l | 3.921142819 | 0.21384345 |
| Amer2 | -0.361771743 | 0.21384345 |
| Polr3gl | 0.46861635 | 0.214063615 |
| Chrac1 | 0.222362413 | 0.571558691 |
| Zfp467 | 0.235054493 | 0.571615837 |
| Zfp120 | -0.351656847 | 0.571615837 |
| Gnpda1 | 0.235971885 | 0.572257223 |
| Rspo1 | 0.507458179 | 0.572270606 |
| Ccdc71 | 0.22443212 | 0.572270606 |
| Fam110a | 0.165110886 | 0.572485425 |
| Fam45a | -0.174194511 | 0.572485425 |
| Nphp3 | 0.46042524 | 0.572653308 |
| Phyhd1 | -0.842171689 | 0.572653308 |
| Bdkrb2 | 3.226898691 | 0.572711874 |
| Gm28557 | 3.226898691 | 0.572711874 |
| Gm5741 | 3.226898691 | 0.572711874 |
| Gm6970 | 3.226898691 | 0.572711874 |
| Lrrc2 | 3.226898691 | 0.572711874 |
| Otop2 | 3.226898691 | 0.572711874 |
| Prr19 | 3.226898691 | 0.572711874 |
| Psg26 | 3.226898691 | 0.572711874 |
| Vmn1r20 | 3.226898691 | 0.572711874 |
| N4bp2l1 | 0.325027418 | 0.572711874 |
| Ctdspl2 | 0.193596819 | 0.572711874 |
| Pcbp2 | 0.170782259 | 0.572711874 |
| Asun | -0.162199211 | 0.572711874 |
| Tgds | -0.270788111 | 0.572711874 |
| Trim43a | 1.220495699 | 0.572748604 |
| C330018D20Rik | 0.337800373 | 0.57284803 |
| Tacc2 | -0.229550125 | 0.573046686 |
| Mtg1 | 0.194080491 | 0.573072608 |
| Iba57 | -0.350607812 | 0.573072608 |
| Pappa | 3.215536699 | 0.573119142 |
| Gm10577 | 1.129341843 | 0.573308845 |
| Cenpu | -0.392971713 | 0.573354921 |
| Cdkl1 | -1.043299888 | 0.573354921 |
| Rab23 | -0.304617757 | 0.573377549 |
| Efcab9 | -0.719338371 | 0.573406356 |
| Slc27a1 | 0.206910273 | 0.573879322 |
| Gpbp1l1 | 0.296473893 | 0.57409195 |
| Prickle4 | -1.04024649 | 0.57409195 |
| Lrrc20 | -0.334751567 | 0.57450947 |
| Ocrl | -0.214840576 | 0.574518526 |
| Slc18a3 | 3.213818678 | 0.574531907 |
| Zfp524 | 0.396401353 | 0.574553232 |
| Arhgef15 | 0.574027529 | 0.574559087 |
| Pibf1 | 0.266119732 | 0.574582921 |
| D17H6S53E | -0.307650782 | 0.574582921 |
| Pnpo | 0.261034163 | 0.574655671 |
| Tspyl2 | -0.257665401 | 0.574749855 |
| Asb18 | -1.815155833 | 0.575424852 |
| Tollip | -0.178580734 | 0.575427042 |
| Gpr180 | 0.26694436 | 0.575571887 |
| Metap1 | -0.181724004 | 0.575707638 |
| Uimc1 | -0.265459282 | 0.575707638 |
| Heatr3 | -0.226959811 | 0.575824313 |
| St8sia4 | 0.24287823 | 0.575903948 |
| H2-Q1 | -0.804205668 | 0.575903948 |
| 2610028H24Rik | 3.201492553 | 0.576149095 |
| Gm5458 | 3.201492553 | 0.576149095 |
| Lipo4 | 3.201492553 | 0.576149095 |
| Tpsb2 | 3.201492553 | 0.576149095 |
| Hk2 | -0.295732659 | 0.576241516 |
| Slc29a2 | -0.456385001 | 0.576241516 |
| Mios | 0.18315484 | 0.576328516 |
| Apobec3 | 0.298303227 | 0.576790043 |
| Tnfsf12 | 1.113509952 | 0.57682731 |
| Atg3 | 0.147953637 | 0.57682731 |
| Fbl | 0.147131469 | 0.576846582 |
| Pm20d2 | -0.603951722 | 0.576846582 |
| Spns1 | -0.205114319 | 0.576948954 |
| Itga7 | -0.87029755 | 0.576948954 |
| Zdhhc22 | -0.870730057 | 0.577008511 |
| Kdm3b | -0.195852824 | 0.577111545 |
| Limk1 | -0.203652696 | 0.577430833 |
| Rpusd4 | -0.351085578 | 0.577771618 |
| Strn | -0.221641985 | 0.578477113 |
| Gpsm2 | -0.22872176 | 0.578477113 |
| Slc6a1 | 0.199436718 | 0.578592469 |
| Rbm4b | -0.155869887 | 0.578772767 |
| Sytl5 | -1.13608579 | 0.578772767 |
| Aspa | 1.037662248 | 0.578808443 |
| Nit1 | 0.195709329 | 0.578808443 |
| Sin3b | 0.153389465 | 0.579149503 |
| Zfp775 | 0.471777487 | 0.579370455 |
| Pnma3 | 1.099506673 | 0.579383996 |
| Ppp4r2 | 0.238278604 | 0.579517517 |
| Psmc5 | 0.1274019 | 0.579558509 |
| Wdr4 | -0.190436683 | 0.579719847 |
| Zfp668 | -0.200623992 | 0.579719847 |
| Lsm12 | 0.142974861 | 0.579807686 |
| Hist1h3g | -0.747183573 | 0.579807686 |
| Gm17606 | -1.476573478 | 0.580202589 |
| Fto | -0.20080692 | 0.580478766 |
| Olfr350 | 1.430261114 | 0.58053452 |
| Bfsp2 | 1.01841438 | 0.58053452 |
| Pth1r | 0.549554558 | 0.58053452 |
| 1110059E24Rik | 0.227764037 | 0.58053452 |
| Imp4 | 0.204787811 | 0.58053452 |
| Brd8 | 0.159550556 | 0.58053452 |
| Galnt16 | 0.399270518 | 0.580549617 |
| Ddx41 | -0.181241411 | 0.580549617 |
| Shtn1 | -0.211904849 | 0.580549617 |
| Slco3a1 | -0.498812248 | 0.580549617 |
| Pwp2 | 0.283182323 | 0.580666516 |
| P3h4 | 0.278291647 | 0.580666516 |
| Casc1 | -1.133303758 | 0.580725293 |
| Ttc28 | -0.209932155 | 0.58080778 |
| Gtpbp3 | -0.405108141 | 0.58080778 |
| Taf8 | -0.264775721 | 0.580897891 |
| Slc29a3 | -0.249096835 | 0.581237964 |
| Top1mt | 0.397805882 | 0.581302474 |
| Cldn12 | 0.609622027 | 0.58130355 |
| Qdpr | 0.15824798 | 0.58130355 |
| Rpf1 | 0.208886538 | 0.581554292 |
| 5430421N21Rik | -1.894055584 | 0.581554292 |
| Lrrc36 | 3.149357598 | 0.582011138 |
| 1700006E09Rik | 1.602432761 | 0.582039873 |
| Mansc4 | 1.550265163 | 0.582039873 |
| Rnf208 | 0.234020854 | 0.582039873 |
| Casz1 | -0.574974071 | 0.582340644 |
| Fgfr4 | 1.003225502 | 0.582504131 |
| Nxpe2 | 0.179215772 | 0.955625421 |
| Phex | -0.249096576 | 0.955928395 |
| Klc2 | 0.045055189 | 0.956108395 |
| Ptprz1 | -0.027102236 | 0.956119342 |
| Nhlrc2 | -0.028797447 | 0.956186905 |
| Mx1 | 0.18510728 | 0.956411592 |
| Slc39a2 | 0.131025911 | 0.956411592 |
| 1190007I07Rik | 0.049020258 | 0.956411592 |
| Rad18 | 0.031725766 | 0.956411592 |
| Slc25a38 | 0.028329848 | 0.956411592 |
| Itfg1 | -0.017017237 | 0.956411592 |
| Upf2 | -0.025998251 | 0.956411592 |
| Slc52a2 | -0.035791685 | 0.956411592 |
| Igdcc4 | -0.043693185 | 0.956411592 |
| Pabpc1l | -0.095740003 | 0.956411592 |
| Arhgef5 | 0.179679219 | 0.956412278 |
| Map3k13 | -0.044328384 | 0.956412278 |
| Gm9755 | -0.051211202 | 0.956412278 |
| Edaradd | -0.053672168 | 0.956535838 |
| Rnf138 | -0.016019919 | 0.956653144 |
| Dgka | 0.050858153 | 0.956693132 |
| Gm8909 | 0.18737852 | 0.956837609 |
| Cmtm3 | -0.023825891 | 0.956837609 |
| Lrrc16a | -0.025781363 | 0.956863162 |
| Prdm11 | 0.049193626 | 0.956882633 |
| Gtf2ird2 | 0.039647853 | 0.956882633 |
| Alcam | 0.030929064 | 0.956882633 |
| Ttc27 | 0.030613179 | 0.956882633 |
| Nkiras2 | -0.018441757 | 0.956882633 |
| Wnt6 | -0.066468159 | 0.956882633 |
| Mst1r | -0.163535575 | 0.956882633 |
| Foxn3 | 0.021104847 | 0.957196812 |
| Tmprss7 | 0.1408951 | 0.957199853 |
| Lpin1 | 0.04566042 | 0.957327544 |
| Elovl5 | -0.021085217 | 0.957327544 |
| Cisd3 | 0.034349755 | 0.957374681 |
| Creb3l1 | 0.056466119 | 0.957419211 |
| Tmem115 | 0.027045909 | 0.957419211 |
| Maff | -0.054042432 | 0.957419211 |
| Sgcd | -0.061137107 | 0.957419211 |
| Iqcj | -0.160042898 | 0.957419211 |
| Btla | 0.140067977 | 0.957580974 |
| Zfp317 | 0.023658795 | 0.957580974 |
| Cope | -0.016079971 | 0.957580974 |
| Cyb5r4 | -0.018335301 | 0.957580974 |
| Nol10 | -0.025542073 | 0.957580974 |
| Omp | 0.312830012 | 0.957632268 |
| Gm5434 | 0.160536193 | 0.958178161 |
| Epha7 | -0.024124687 | 0.958178161 |
| Ccdc107 | 0.021197516 | 0.958205177 |
| Actl6b | 0.018505732 | 0.958402494 |
| Mylip | 0.052291899 | 0.958436794 |
| Fam73b | -0.027772111 | 0.958909549 |
| Prkci | -0.026297536 | 0.959057132 |
| Gpr63 | 0.085467309 | 0.959199278 |
| Tmem161b | 0.043452199 | 0.959365961 |
| Rprd2 | 0.018914507 | 0.959365961 |
| Idh3a | 0.013879698 | 0.959365961 |
| Ube4a | -0.018527613 | 0.959365961 |
| Naf1 | -0.036014304 | 0.959365961 |
| Sprtn | -0.050645389 | 0.959365961 |
| Tssk1 | -0.25387761 | 0.959365961 |
| Sestd1 | 0.015669834 | 0.959388138 |
| Zfp827 | 0.027137997 | 0.959884242 |
| Klhl24 | 0.014587163 | 0.96052563 |
| Galc | -0.034664987 | 0.960577308 |
| Asgr2 | 0.310015745 | 0.960736817 |
| Dicer1 | -0.021445307 | 0.960736817 |
| Wdpcp | 0.0440837 | 0.960844063 |
| Tpbgl | -0.063646508 | 0.960844063 |
| Rcc1 | -0.019691233 | 0.960938155 |
| Camsap2 | -0.015907236 | 0.961255888 |
| Rps2-ps6 | -0.118759356 | 0.961802133 |
| Frmd3 | 0.034483718 | 0.961869304 |
| Lrrc56 | 0.037540993 | 0.961902604 |
| Sema4g | 0.02745616 | 0.96200225 |
| Ehmt1 | 0.017884974 | 0.962346047 |
| Ttll13 | 0.164274426 | 0.962389803 |
| Zmat3 | 0.027366756 | 0.962389803 |
| Ubtf | 0.014723493 | 0.962389803 |
| Uqcrc1 | 0.012546121 | 0.962389803 |
| Tmem70 | 0.018061962 | 0.962651074 |
| Ylpm1 | -0.015516119 | 0.962714441 |
| B930041F14Ri | 0.054963436 | 0.962961542 |
| Adgrg2 | 0.115496986 | 0.963201635 |
| Efcab12 | 0.079072938 | 0.963201635 |
| Sncb | 0.05279624 | 0.963334936 |
| Cables1 | 0.053519304 | 0.963445322 |
| Nsg2 | -0.016629475 | 0.963455728 |
| Micu2 | -0.018445241 | 0.963698799 |
| Dlx2 | -0.018757627 | 0.963751906 |
| Thg1l | -0.032664171 | 0.963751906 |
| G430095P16Ri | -0.088797297 | 0.963751906 |
| Odf4 | -0.231469679 | 0.963864336 |
| Neil1 | -0.03511161 | 0.963914448 |
| Pcf11 | 0.017541153 | 0.96400799 |
| Cpsf2 | -0.014656514 | 0.964179016 |
| Gm21887 | 0.092127634 | 0.964253359 |
| Klf5 | -0.053589748 | 0.964253359 |
| Nup54 | 0.017695415 | 0.964312691 |
| Taf1b | -0.019488431 | 0.964392435 |
| Llgl1 | -0.023004578 | 0.964392435 |
| Nanos2 | 0.123126026 | 0.964783781 |
| Zfp958 | -0.044528761 | 0.964783781 |
| Zfp474 | -0.226711979 | 0.964783781 |
| Nrp2 | 0.027439983 | 0.964794561 |
| Eif1 | -0.011214637 | 0.964794561 |
| Pcdha7 | -0.062787476 | 0.964975416 |
| Ccer2 | 0.152560669 | 0.965108823 |
| Tctex1d4 | 0.10541979 | 0.965108823 |
| Pik3ip1 | -0.02635188 | 0.965108823 |
| Duox1 | -0.129875228 | 0.965108823 |
| Angpt2 | -0.035305873 | 0.965138001 |
| Fam117b | 0.020203531 | 0.965223029 |
| Wdr95 | -0.107748571 | 0.96548393 |
| Gm7808 | 0.079616828 | 0.965530623 |
| Ttc32 | 0.027711803 | 0.965530623 |
| Gpatch8 | -0.014264907 | 0.96559577 |
| Pla1a | -0.102325452 | 0.96559577 |

| | | |
|---|---|---|
| Rps13 | 0.445802631 | 0.017042941 |
| Lilrb4a | -2.992015714 | 0.017054909 |
| Ccdc66 | 0.873536459 | 0.017087646 |
| Nudt14 | 0.674772597 | 0.017112686 |
| Polq | -1.222057527 | 0.017174377 |
| Ccdc86 | -0.994703903 | 0.017186508 |
| Pbx3 | 0.840786543 | 0.017222377 |
| Lyrm4 | 0.547591601 | 0.017242873 |
| Ndufs2 | 0.504522525 | 0.017242873 |
| Fam219b | 0.6401267 | 0.017246077 |
| Poll | 1.302659986 | 0.017271339 |
| Pkig | 0.664895866 | 0.017287285 |
| Mybl2 | -1.190755455 | 0.017287285 |
| Slc30a9 | 0.614022138 | 0.017295112 |
| Adgrl3 | 0.868841744 | 0.017344846 |
| Tada2a | 0.870209632 | 0.017389227 |
| Lyrm2 | 0.612525289 | 0.017389227 |
| Nup214 | -0.616056033 | 0.017406348 |
| Ccny | -0.564510006 | 0.017553234 |
| Lox | 2.167176353 | 0.017586938 |
| Fau | 0.456006069 | 0.017591538 |
| Ivd | 0.923451215 | 0.01761133 |
| Mrpl23 | 0.603468241 | 0.01761133 |
| Ankrd12 | 0.63704575 | 0.01762394 |
| Lin28b | 2.736652578 | 0.017643487 |
| Ttc25 | 4.866076735 | 0.01765339 |
| Cd7 | -6.565452746 | 0.017669104 |
| Zdbf2 | 0.78527981 | 0.017692701 |
| 1700028K03Rik | -2.902240621 | 0.017736456 |
| Mad1l1 | -0.765739324 | 0.017736876 |
| Klf4 | 1.212884019 | 0.017767848 |
| Spred1 | 0.911408941 | 0.017795322 |
| Qrich2 | -6.160197598 | 0.017816383 |
| Abcg1 | -0.611682165 | 0.017821599 |
| Gbgt1 | -2.469618107 | 0.017827601 |
| Asb8 | 0.937875392 | 0.017876211 |
| Nt5c3b | 0.588659766 | 0.017876211 |
| Sfxn1 | 0.553399134 | 0.017876211 |
| Xpo7 | -0.664070973 | 0.017876211 |
| 6430548M08Rik | -1.052685864 | 0.017937413 |
| Tmem199 | 0.697543889 | 0.017950246 |
| Pigh | 1.230931493 | 0.017974391 |
| BC030336 | 0.957107502 | 0.017981869 |
| Cog7 | 0.940260752 | 0.018006722 |
| Mapt | 0.559913174 | 0.018006722 |
| B3galt6 | 0.825031402 | 0.018026352 |
| Parpbp | -1.209470033 | 0.018064034 |
| RP23-457J22.1 | -6.245923789 | 0.01806805 |
| Rnf32 | 1.353815384 | 0.018101977 |
| Psenen | 0.543980961 | 0.018134852 |
| Rpl7a | 0.626953002 | 0.018150207 |
| Fgf8 | 5.82678944 | 0.018266455 |
| Xbp1 | -0.645102755 | 0.018332888 |
| Cdh23 | -1.484621208 | 0.018378286 |
| Syt11 | 0.435560438 | 0.018413226 |
| Fv1 | 1.816254513 | 0.018430549 |
| B230219D22Rik | 0.492109555 | 0.018436712 |
| Gnb5 | 1.097301724 | 0.018558183 |
| Gadd45a | -1.209365946 | 0.018558183 |
| Capn1 | -2.063325807 | 0.018592459 |
| Csmd3 | 1.24352557 | 0.018625128 |
| Aspm | 1.12158251 | 0.018706028 |
| Bptf | 0.584912728 | 0.018706028 |
| St18 | 0.839932503 | 0.018729356 |
| Nek4 | 0.930839766 | 0.018732091 |
| Pabpc1 | 0.617890785 | 0.018745173 |
| Ddx54 | -0.571557981 | 0.01879778 |
| Pard3 | 0.726131821 | 0.018802645 |
| Accsl | -4.932934081 | 0.018848676 |
| Derl3 | -2.20591717 | 0.018862456 |
| Kctd11 | -3.264234459 | 0.018862456 |
| Polr2c | 0.683061187 | 0.01892508 |
| Igfals | -6.560029926 | 0.01892978 |
| Treml2 | -1.76947404 | 0.018950331 |
| Acp1 | 0.511761099 | 0.018951107 |
| Slfn4 | -4.249505481 | 0.018971608 |
| Serpinb12 | -6.530082277 | 0.018971608 |
| Ncaph2 | 0.605058798 | 0.019008011 |
| Psmc1 | 0.546348459 | 0.019035419 |
| Adcy10 | -2.374950632 | 0.019039639 |
| Dpm2 | 0.559772015 | 0.019147349 |
| Fbxo3 | -0.609561692 | 0.019240895 |
| Abcd2 | -0.686693507 | 0.019298252 |
| Pdia5 | -1.337214969 | 0.019310492 |
| Itm2b | -0.565542409 | 0.01931803 |
| Elf2 | -0.516000403 | 0.01935121 |
| Exoc6 | -1.117677655 | 0.01935121 |
| Sfrp1 | 0.991097852 | 0.019358023 |
| Parp16 | 2.859529475 | 0.019374003 |
| Pdrg1 | 0.598903382 | 0.019374483 |
| Pcdhb17 | 1.525271376 | 0.019413506 |
| Peg12 | 1.097687082 | 0.019446433 |
| 1810030O07Rik | -0.67453199 | 0.019449753 |
| Garem | 1.347956754 | 0.019484544 |
| Crtap | 1.248354067 | 0.019485271 |
| Cycs | 0.447023037 | 0.019506921 |
| Usp37 | -0.520403615 | 0.019523676 |
| Pms1 | 1.15825912 | 0.019552676 |
| Fis1 | 0.464860207 | 0.019587655 |
| Hspg2 | 1.531708847 | 0.019593455 |
| Tarm1 | -2.494088521 | 0.019602028 |
| Ttc39a | -2.001030741 | 0.019605648 |
| Utp20 | -0.895835722 | 0.019615044 |
| Rnf165 | 0.636406728 | 0.019724243 |
| Glrx3 | 0.426339379 | 0.019763492 |
| Rsph10b | -3.816823157 | 0.019763492 |
| Tspan13 | 0.428832526 | 0.019812797 |
| Adgra3 | -1.039098066 | 0.019812797 |
| Pros1 | 1.011464027 | 0.019857551 |
| Prmt5 | 0.575665026 | 0.019862786 |
| Foxn2 | -0.69269506 | 0.01994049 |
| Rps3a1 | 0.450285648 | 0.019958705 |
| Arl6ip1 | 0.481474334 | 0.020031639 |
| Med14 | -0.67880105 | 0.020036308 |
| C1qtnf4 | 1.059331837 | 0.020076002 |
| Jmjd1c | -0.576126541 | 0.020077113 |
| Ggt1 | -3.514077085 | 0.020086744 |
| Fkbp7 | 1.307249683 | 0.020092952 |
| Hspb11 | 0.859960822 | 0.020094257 |

| | | |
|---|---|---|
| Sema3c | -0.49725517 | 0.214063615 |
| Fblim1 | 0.675243142 | 0.214204333 |
| Chil5 | 4.243134903 | 0.214384111 |
| Mreg | 2.349514646 | 0.214388076 |
| Plppr1 | -0.56698749 | 0.214675071 |
| Ackr1 | 0.688941836 | 0.214784206 |
| Exosc10 | -0.317570343 | 0.21491974 |
| Abhd12 | -0.316351626 | 0.214958757 |
| Slc27a2 | 1.709008667 | 0.215021747 |
| Slco2a1 | -1.979246501 | 0.215171261 |
| Tssk3 | 4.243686623 | 0.215201542 |
| Serpinb6a | -0.506589317 | 0.215322646 |
| Mc1r | 3.566417332 | 0.215337173 |
| Rapgef3 | 1.265555871 | 0.215353933 |
| Mtfr1l | 0.259612087 | 0.215353933 |
| Bap1 | 0.344624365 | 0.215395609 |
| Nrap | 3.41326631 | 0.215437942 |
| Oat | 0.369301502 | 0.215437942 |
| Iqcf5 | 4.704255144 | 0.215674849 |
| Mapre2 | -0.292557727 | 0.215674849 |
| Tmem229a | -1.468951557 | 0.215674849 |
| Egr2 | -1.21977208 | 0.215945806 |
| Msrb2 | 0.980663787 | 0.216033867 |
| Smim4 | 0.50922188 | 0.216033867 |
| Coq7 | 0.411416659 | 0.21608725 |
| Gm5416 | -1.954281381 | 0.216110063 |
| Ranbp10 | -0.452735156 | 0.216143861 |
| Herc6 | -0.928081674 | 0.216179575 |
| Txnrd1 | -0.248913445 | 0.216242872 |
| Tlcd2 | -0.753280384 | 0.216242872 |
| Tnfrsf22 | 2.428376655 | 0.21625098 |
| Cbx5 | 0.235715713 | 0.216715965 |
| Zfp442 | 1.490884069 | 0.216752333 |
| Rasl12 | 1.382050788 | 0.216752333 |
| Spata6 | -0.882512925 | 0.216853931 |
| Rab40b | 1.144509404 | 0.216914145 |
| Smarca5 | -0.34782631 | 0.217067305 |
| Gprc5b | 0.66887891 | 0.217189276 |
| Mapk9 | -0.341986826 | 0.217189276 |
| Kif21b | -0.34399671 | 0.21726474 |
| Xkr8 | -1.473252398 | 0.21726474 |
| Slc5a10 | -4.798944457 | 0.21726474 |
| Ints5 | -0.470268134 | 0.21732925 |
| Cel5 | -0.718461994 | 0.217380363 |
| Ndst4 | 2.183162459 | 0.217620322 |
| Wbp11 | -0.310301074 | 0.217638653 |
| 2810459M11Rik | 0.736598564 | 0.217713347 |
| Lrrc27 | 1.189677149 | 0.217940504 |
| Dst | 0.380113791 | 0.217940504 |
| Aig1 | -0.411097924 | 0.21806794 |
| Evi2a | -0.543691813 | 0.218162045 |
| Bmp3 | 1.408426508 | 0.218183893 |
| Sdhaf2 | 0.414308785 | 0.218306111 |
| Zc2hc1a | 0.337760087 | 0.218306111 |
| Isoc2a | -1.63665251 | 0.218306111 |
| Cenpk | 0.684464545 | 0.218435402 |
| Suz12 | 0.340468097 | 0.218479157 |
| Tm4sf19 | -2.648142777 | 0.218531037 |
| Pdlim5 | -0.714291592 | 0.218569872 |
| Col17a1 | -3.429979038 | 0.218577411 |
| Tubg2 | 0.646103431 | 0.218705744 |
| Inpp4b | -1.045040546 | 0.218705744 |
| Fam216b | -4.348425885 | 0.218787396 |
| 3-Mar | 1.566992578 | 0.218844044 |
| Fzd3 | -0.353631458 | 0.218844044 |
| Tmem68 | -0.36994387 | 0.218844044 |
| Cage1 | 1.454639849 | 0.218881313 |
| Adcy6 | 0.817618178 | 0.218881313 |
| Smg8 | -0.435071228 | 0.218915498 |
| Il1rapl1 | 0.78403296 | 0.218916972 |
| Ppfia1 | 0.43872654 | 0.21894993 |
| Rarres2 | 2.599012029 | 0.219112073 |
| Brd2 | -0.256129898 | 0.219180343 |
| Dnase1l2 | -5.084316829 | 0.219187326 |
| Mmp7 | -5.084316829 | 0.219187326 |
| Mis12 | -0.691083941 | 0.219292972 |
| Gm21969 | -5.100487603 | 0.2194182 |
| Abhd16a | -0.401196971 | 0.219539566 |
| C330021F23Rik | 0.555300526 | 0.219626856 |
| Fam65c | 4.598552824 | 0.219630799 |
| Ap2b1 | -0.324944489 | 0.219630799 |
| Ate1 | 0.334763427 | 0.21970413 |
| Tnfaip2 | -1.221328272 | 0.21970413 |
| Sp2 | -0.494285587 | 0.219707986 |
| Amy1 | 2.06900418 | 0.219782261 |
| Kif9 | 1.078518983 | 0.219825648 |
| Bspry | 2.807076269 | 0.220100873 |
| Slc26a8 | -2.248848696 | 0.220104305 |
| Entpd2 | 2.620001504 | 0.220187601 |
| Gm6525 | 1.032064499 | 0.220187601 |
| Slc30a4 | -0.870964983 | 0.220187601 |
| Slc25a44 | -0.365956178 | 0.220201712 |
| Ap5m1 | -0.460224634 | 0.220201712 |
| Timm21 | 0.494510677 | 0.220328088 |
| BC048546 | 1.828392936 | 0.220398902 |
| Wdr92 | 0.497277763 | 0.220409267 |
| Cstf1 | 0.459022702 | 0.220434458 |
| Stambpl1 | -0.676476837 | 0.220434458 |
| Glp2r | -3.406217248 | 0.220434458 |
| 4930453N24Rik | 0.363567881 | 0.220460786 |
| Endog | -0.94829203 | 0.220549547 |
| Dcaf15 | 0.795642785 | 0.220669723 |
| Dync1i2 | 0.301410446 | 0.220898688 |
| Snd1 | -0.328583735 | 0.220923531 |
| Susd5 | -5.067972794 | 0.221174255 |
| Arhgdig | 0.565406046 | 0.221206254 |
| 1700034I23Rik | 3.001900431 | 0.221272872 |
| Csad | 0.539587719 | 0.221341567 |
| Fam196b | 1.344797436 | 0.221406002 |
| Taf9b | 0.884676243 | 0.221436719 |
| Ddx49 | 0.571749176 | 0.221483933 |
| Syt16 | 0.473293509 | 0.221818124 |
| Ric1 | -0.362639572 | 0.221914012 |
| Dusp6 | -0.3789826 | 0.221914012 |
| Zfp773 | -0.831772846 | 0.221926355 |
| Art3 | -2.640585567 | 0.221926355 |
| Cmya5 | 3.996955139 | 0.222015669 |
| Zfp622 | -0.313179293 | 0.222015669 |
| Mtl5 | 4.138476284 | 0.222102461 |

| | | |
|---|---|---|
| Kif14 | 0.406900256 | 0.582504131 |
| Pex11b | 0.205085047 | 0.582504131 |
| 4931406P16Rik | -0.179022264 | 0.582658504 |
| Tfpt | 0.205758293 | 0.582790334 |
| Pgp | 0.200350663 | 0.582790334 |
| Slc35c2 | -0.162961492 | 0.582790334 |
| Gata3 | -0.759660926 | 0.582790334 |
| Gm5493 | -0.845995217 | 0.582814944 |
| Zfp637 | 0.188054921 | 0.583132292 |
| Papolg | -0.262103742 | 0.58315257 |
| Cog8 | -0.211750177 | 0.5831647 |
| Stx12 | 0.141099951 | 0.583233473 |
| Ssbp2 | -0.266231277 | 0.583233473 |
| Eya2 | -0.765773021 | 0.583233473 |
| Zscan12 | 0.346722686 | 0.583368214 |
| Nfya | -0.255541326 | 0.583605253 |
| Cux2 | -0.289884598 | 0.583766611 |
| Gca | 0.285489913 | 0.583860949 |
| Luzp1 | 0.237920183 | 0.583959961 |
| Zfp865 | -0.456114944 | 0.583959961 |
| Marveld3 | -0.858820105 | 0.583959961 |
| C1s2 | -2.289537536 | 0.584322223 |
| Gtsf1 | -2.133132078 | 0.584493717 |
| Vps13a | 0.265238972 | 0.585333585 |
| Tsen2 | 0.201004319 | 0.585333585 |
| Tuba3a | 2.514654198 | 0.585380326 |
| Gpr50 | 3.133489095 | 0.58542389 |
| Cers3 | 1.446006805 | 0.585423988 |
| Hist4h4 | -0.6238471 | 0.585423988 |
| Gm17660 | 3.133698906 | 0.585484826 |
| Med15 | -0.208205648 | 0.585484826 |
| Mybpc1 | -2.586689924 | 0.585484826 |
| Adipor2 | -0.177413644 | 0.585964228 |
| Akirin1 | 0.136284547 | 0.586108401 |
| Marveld1 | 1.109081955 | 0.586156958 |
| Zc2hc1c | -0.66681149 | 0.586196438 |
| Szrd1 | -0.179481229 | 0.586388419 |
| Zfp78 | -0.788408245 | 0.58659543 |
| Ankfn1 | 0.617497864 | 0.586913408 |
| Gde1 | 0.138729727 | 0.587178059 |
| Slc16a11 | 0.941674494 | 0.587188966 |
| A930009A15Rik | 3.121640605 | 0.587301457 |
| Aadat | 3.121640605 | 0.587301457 |
| Arl9 | 3.121640605 | 0.587301457 |
| Tigit | -1.035602476 | 0.58746161 |
| AI606181 | 0.456776762 | 0.587480521 |
| Tmem266 | 0.597629424 | 0.587498811 |
| Nat9 | 0.298918075 | 0.587498811 |
| Fads3 | -0.340590507 | 0.587498811 |
| Slc41a2 | -0.554129907 | 0.587498811 |
| Zfp648 | 3.118681433 | 0.587555967 |
| Pdcd10 | -0.168586865 | 0.587722028 |
| Atic | -0.209941266 | 0.5879139 |
| Gm17567 | 3.115898902 | 0.587977641 |
| D630023F18Rik | -1.042475981 | 0.588209156 |
| Hax1 | -0.154200458 | 0.588612418 |
| Haus5 | -0.284686996 | 0.588612418 |
| Cd5l | 3.1094863 | 0.588945448 |
| Gpr15 | 3.1094863 | 0.588945448 |
| Rgsl1 | 3.1094863 | 0.588945448 |
| Tomm20l | 3.1094863 | 0.588945448 |
| Rbm45 | 0.240182808 | 0.589189711 |
| Prr33 | 1.131613846 | 0.58929657 |
| Kazn | 0.335271205 | 0.589622233 |
| Hexim2 | 0.306042584 | 0.589622233 |
| Gna12 | 0.255468205 | 0.589664801 |
| D10Wsu102e | 0.261872263 | 0.589692713 |
| Agr3 | 3.103730524 | 0.589727038 |
| Hapln4 | 3.103730524 | 0.589727038 |
| Palmd | 0.34967509 | 0.590135921 |
| Ksr1 | -0.462189168 | 0.590677074 |
| Epha3 | -0.185890051 | 0.590688624 |
| Ppp1r13l | 0.598711316 | 0.590919322 |
| Lama2 | 0.52840008 | 0.590919322 |
| Neo1 | -0.185205267 | 0.591222707 |
| Gm6096 | 3.094406773 | 0.591298141 |
| Katnbl1 | -0.223324483 | 0.591346002 |
| Krt12 | 2.450287692 | 0.591466793 |
| Sars | -0.11784321 | 0.59167447 |
| Dlgap4 | -0.1260081 | 0.5917043 |
| Copb2 | -0.143534569 | 0.5917043 |
| A530032D15Rik | -0.612636836 | 0.5917043 |
| Adgrg1 | 0.178782078 | 0.591708642 |
| Cd79a | 1.678752328 | 0.591748214 |
| Mier1 | 0.166586112 | 0.591768511 |
| 4933402P03Rik | 3.088635255 | 0.591828467 |
| Ctsr | 3.088635255 | 0.591828467 |
| Zfp953 | 3.088635255 | 0.591828467 |
| Dbpht2 | 0.54518253 | 0.591828467 |
| Aifm1 | -0.176081772 | 0.592033637 |
| Hacl1 | 0.447650106 | 0.592209765 |
| Rnase4 | 0.282963061 | 0.592209765 |
| Lrrc23 | -0.716458222 | 0.592337726 |
| Hs6st1 | -0.278185524 | 0.592414324 |
| Perp | -0.422090218 | 0.592414324 |
| Atrnl1 | -0.352135471 | 0.592677316 |
| Sdhaf1 | 0.315541959 | 0.592678272 |
| Galm | -0.498043127 | 0.592678272 |
| Slc25a3 | 0.122359257 | 0.592843946 |
| Dnah6 | -0.85231831 | 0.592843946 |
| Tsx | 2.035437947 | 0.592957398 |
| Paqr4 | 0.246733839 | 0.592957398 |
| Emg1 | 0.140881368 | 0.592957398 |
| Exoc8 | -0.243488654 | 0.593022623 |
| Elovl6 | 0.197653189 | 0.593040855 |
| Epb41 | -0.183453152 | 0.593088379 |
| Slc35d1 | -0.470666805 | 0.593088379 |
| Slc4a4 | -0.364099658 | 0.593089251 |
| Faah | 0.442970861 | 0.593344017 |
| Ncapg | 0.26873406 | 0.593344017 |
| Ucma | -1.158463119 | 0.593479493 |
| Ifna15 | 3.074503447 | 0.593579107 |
| Acads | -0.250657495 | 0.593602439 |
| Tie1 | 0.935700099 | 0.593699867 |
| Wsb2 | -0.190565226 | 0.593718035 |
| Zfp84 | -0.286263976 | 0.59373241 |
| Adamts15 | 2.419557483 | 0.594051729 |
| Sigirr | 0.818792021 | 0.594051729 |
| Lrtm2 | 0.650656311 | 0.594051729 |

| | | |
|---|---|---|
| Sdf4 | -0.014940982 | 0.965621388 |
| Tab1 | 0.026664055 | 0.965837981 |
| Cep76 | 0.024390661 | 0.965935409 |
| Rarres1 | -0.123508898 | 0.965935409 |
| Ech1 | -0.017229908 | 0.966091945 |
| Gtf3c3 | -0.014458941 | 0.966182388 |
| Creb5 | -0.023714505 | 0.966209676 |
| Il4i1 | 0.116415277 | 0.966236011 |
| Bak1 | -0.051999392 | 0.966381752 |
| Mmp23 | 0.108110869 | 0.966409297 |
| Stx4a | 0.01969294 | 0.966779703 |
| Duxbl1 | 0.19989099 | 0.966824505 |
| Angptl6 | 0.035627876 | 0.966824505 |
| Twsg1 | 0.015398454 | 0.966824505 |
| Ppp4c | 0.013014997 | 0.966824505 |
| Rexo1 | -0.01382548 | 0.966824505 |
| Mettl4 | -0.020425273 | 0.966824505 |
| Armt1 | -0.028316674 | 0.966824505 |
| Gsta3 | 0.120653426 | 0.96709537 |
| Chrm2 | 0.096521218 | 0.96709537 |
| Ssc4d | -0.075705913 | 0.96709537 |
| Trio | -0.021098934 | 0.967668024 |
| Nucb2 | 0.020191638 | 0.967812482 |
| Hspa2 | 0.054854751 | 0.968226634 |
| Zscan20 | 0.042539561 | 0.968226634 |
| Plekhg1 | 0.035115883 | 0.968226634 |
| Lcor | 0.020582444 | 0.968226634 |
| Supt16 | -0.013786206 | 0.968226634 |
| Dnajc6 | -0.018567748 | 0.968226634 |
| Coq10a | -0.025515018 | 0.968226634 |
| Sgsm2 | -0.026517781 | 0.968226634 |
| Gnal | -0.026616521 | 0.968226634 |
| Spire2 | -0.02836495 | 0.968226634 |
| Jchain | -0.194212228 | 0.968226634 |
| Gm10644 | -0.219125737 | 0.968226634 |
| Gpr165 | 0.056037379 | 0.968243993 |
| Banp | 0.016267469 | 0.968243993 |
| Rarg | -0.052222445 | 0.968337433 |
| Bdh2 | 0.052851447 | 0.968472851 |
| Erc2 | -0.01498982 | 0.968542476 |
| Hist1h4n | -0.045189575 | 0.968552466 |
| Dner | 0.016155495 | 0.968610269 |
| Gm5415 | -0.048982064 | 0.968621003 |
| Hist1h3a | -0.057661197 | 0.968721248 |
| Tsen54 | 0.02503957 | 0.968868256 |
| Smap2 | 0.016243958 | 0.968868256 |
| Zfp2 | -0.021765642 | 0.969118739 |
| Txk | -0.150348602 | 0.969153439 |
| Atrip | 0.054526008 | 0.969169513 |
| Slc9a9 | 0.037842071 | 0.969169513 |
| Ric8 | 0.014842308 | 0.969169513 |
| Golm1 | 0.012706245 | 0.969169513 |
| Suv420h1 | 0.011387453 | 0.969169513 |
| Sos1 | -0.019025444 | 0.969169513 |
| Kif24 | -0.031497412 | 0.969169513 |
| Cops3 | -0.012154269 | 0.9691767 |
| Wrap53 | 0.020950048 | 0.969442067 |
| Nif3l1 | -0.014200323 | 0.969442067 |
| Bid | -0.019268242 | 0.969442067 |
| Bckdha | -0.031265427 | 0.969442067 |
| Cst6 | 0.119163373 | 0.969575675 |
| Hagh | 0.013744729 | 0.969623722 |
| Cd83 | -0.020503664 | 0.969623722 |
| Arhgef17 | -0.020982968 | 0.969623722 |
| Ccdc190 | 0.046068787 | 0.969711409 |
| Cog6 | -0.018367043 | 0.969711409 |
| Tmem170 | 0.118458348 | 0.96994874 |
| Cilp2 | 0.042823721 | 0.96994874 |
| Katnb1 | 0.01758965 | 0.96994874 |
| Shank2 | -0.022983669 | 0.96994874 |
| Rpusd1 | -0.023294403 | 0.96994874 |
| Nxpe3 | -0.03202788 | 0.96994874 |
| Akap14 | -0.118192163 | 0.96994874 |
| Ces1d | 0.121535986 | 0.969995994 |
| Ndufv1 | 0.010762108 | 0.970119487 |
| Heca | 0.031081335 | 0.970493738 |
| Rrh | -0.139392303 | 0.970493738 |
| 5830473C10Ri | -0.151189607 | 0.970493738 |
| Slc29a1 | -0.018225682 | 0.97049566 |
| Ifitm5 | 0.183231652 | 0.970751987 |
| Cntnap1 | -0.123718692 | 0.970751987 |
| Igflr1 | 0.053370504 | 0.971050313 |
| Ube2g2 | 0.013308498 | 0.971050313 |
| Helq | -0.022850115 | 0.971260094 |
| Ifrd2 | 0.03154407 | 0.971321183 |
| Awat2 | 0.092606878 | 0.971472478 |
| BC024978 | 0.021228423 | 0.971475772 |
| Fam209 | 0.146034182 | 0.971602549 |
| Klhl35 | 0.042196307 | 0.971602549 |
| Vma21 | 0.015497608 | 0.971602549 |
| Gbp9 | -0.039626852 | 0.971602549 |
| Cdk20 | 0.020184107 | 0.971604339 |
| Dfna5 | -0.012349086 | 0.971604339 |
| Abl2 | -0.016755986 | 0.971720793 |
| Spats2 | 0.014074465 | 0.971772485 |
| Cdkn2aip | -0.020470215 | 0.971772485 |
| Papss2 | 0.038547869 | 0.971891106 |
| Gria3 | 0.019818961 | 0.971891106 |
| Gne | 0.018711023 | 0.971891106 |
| Farp1 | 0.017541344 | 0.971891106 |
| Manf | 0.012534903 | 0.971891106 |
| Ccp110 | -0.013114463 | 0.971891106 |
| Gm11992 | -0.196221062 | 0.971891106 |
| Slc30a10 | 0.025974128 | 0.972029272 |
| Atp5s | 0.019981409 | 0.972193711 |
| Igsf5 | -0.113991121 | 0.972193711 |
| Scnn1b | 0.159528475 | 0.972503453 |
| Zfp788 | 0.021524463 | 0.972542871 |
| Rrn3 | 0.011226543 | 0.972694121 |
| Gm3453 | 0.119136564 | 0.973040237 |
| Ccdc15 | -0.022368218 | 0.973040237 |
| Ccdc8 | -0.029689767 | 0.973040237 |
| Intu | 0.029183334 | 0.97304931 |
| Ano1 | 0.046913196 | 0.973150379 |
| Zfp810 | -0.016484608 | 0.973150379 |
| Zfp369 | -0.019942712 | 0.973150379 |
| Fyco1 | -0.021148881 | 0.973150379 |
| Ubxn11 | -0.031306526 | 0.973150379 |
| Cdt1 | -0.016118712 | 0.973182084 |

| | | |
|---|---|---|
| Tpgs2 | 0.596459498 | 0.020096399 |
| Setd1b | -0.702408852 | 0.020096964 |
| Dock3 | -0.886995777 | 0.020106512 |
| Gchfr | 2.240217338 | 0.020112552 |
| Abcc4 | -1.40841112 | 0.020173773 |
| Ctxn2 | 2.108455752 | 0.020176266 |
| Mki67 | 0.794877841 | 0.020176266 |
| Tex264 | -0.617938251 | 0.020176266 |
| Ntm | 0.803326735 | 0.020205984 |
| Kctd10 | -0.737913649 | 0.020212148 |
| Parva | 0.987003535 | 0.020269747 |
| Cp | -1.213723351 | 0.020280049 |
| Tmem242 | 0.689269744 | 0.020294246 |
| Cerk | -0.996410666 | 0.020294246 |
| Gipc2 | 2.590943254 | 0.020307937 |
| Eafl | -0.970630949 | 0.020307937 |
| Galns | -1.261458753 | 0.020334722 |
| Brd4 | -0.535928487 | 0.020343773 |
| Fam120c | 0.942883647 | 0.020394623 |
| Naa38 | 0.959895905 | 0.020397822 |
| Tdrp | 2.687498217 | 0.020426784 |
| Atp8b2 | 1.319792673 | 0.020426784 |
| Spen | -0.600200275 | 0.020426784 |
| Kcnk6 | -1.429527572 | 0.020426784 |
| BC147527 | -2.409504176 | 0.020426784 |
| Lrig3 | -2.761215358 | 0.020448485 |
| Stk4 | -0.692067045 | 0.020458017 |
| Aqp1 | 3.936697023 | 0.020459792 |
| Ephb2 | 0.838923652 | 0.020507006 |
| Rbm27 | 0.589172436 | 0.020508001 |
| Maml2 | -0.933648777 | 0.020525566 |
| Fzd9 | 1.855638413 | 0.020540827 |
| Rnps1 | 0.57781862 | 0.020540827 |
| Brwd1 | 0.489777624 | 0.020545443 |
| Otp | 5.419421684 | 0.020546405 |
| Erap1 | -1.407796608 | 0.020555006 |
| Tprkb | 0.784899048 | 0.020555104 |
| Nip7 | 0.577921748 | 0.020555104 |
| Rpl4 | 0.433985905 | 0.020555104 |
| Tyro3 | 1.264138366 | 0.020577195 |
| Opn1sw | -6.172944106 | 0.020591721 |
| Msx2 | 5.474842894 | 0.020597331 |
| Ppfia4 | 1.02373148 | 0.020602562 |
| Tmtc1 | -1.705864365 | 0.020602562 |
| Dhx33 | -0.808373444 | 0.020653903 |
| Hyal1 | 3.142593412 | 0.020658633 |
| Sbf1 | -0.654641015 | 0.020681144 |
| C330027C09Rik | 0.889466664 | 0.02069439 |
| Dync1li2 | 0.624280082 | 0.02070826 |
| Tjp2 | 1.285799227 | 0.02076237 |
| Mta3 | -0.990403741 | 0.020764076 |
| Tjp1 | 0.856759337 | 0.020777992 |
| Commd3 | 0.495240605 | 0.020795559 |
| Rela | -0.767427936 | 0.020847007 |
| Epsti1 | -1.479721191 | 0.02087135 |
| Pigx | 0.731740236 | 0.020899261 |
| Qser1 | 0.610899844 | 0.020899261 |
| Bud31 | 0.662776307 | 0.02090628 |
| Ctso | -0.882734306 | 0.02090628 |
| Lamtor2 | 0.498076121 | 0.020919343 |
| Smagp | -1.112201123 | 0.020944513 |
| Eef2 | -0.494317896 | 0.020975782 |
| Nsmaf | -0.686177194 | 0.021087491 |
| Spatc1 | -3.411924891 | 0.021164644 |
| Serpinb1b | 5.753503937 | 0.021167827 |
| Hmmr | 1.072481919 | 0.021219741 |
| Atad2 | -0.735797921 | 0.021226245 |
| Hmga2 | 2.528809486 | 0.021242752 |
| Flywch1 | 0.540697485 | 0.021242752 |
| Rasgrp4 | -3.583401697 | 0.021243782 |
| Rpp14 | 0.816919925 | 0.021276461 |
| Rragb | 1.282303324 | 0.021312562 |
| Arhgap30 | -1.910869551 | 0.021312562 |
| Tfap2a | 5.57351068 | 0.021363633 |
| Znrf3 | 1.123360212 | 0.02136503 |
| Gpc1 | 0.620344771 | 0.021405254 |
| Phkg1 | 3.83248699 | 0.021408521 |
| E2f5 | 0.89930665 | 0.021411673 |
| Samd3 | 5.738890613 | 0.021431812 |
| Shd | 0.679405622 | 0.021443113 |
| Fndc4 | 0.600144741 | 0.021463172 |
| 2010111I01Rik | -0.931134594 | 0.021490774 |
| Gm14548 | -3.799598305 | 0.021524789 |
| Fam83g | -4.000085284 | 0.021602692 |
| Ptchd4 | 2.066948456 | 0.02161368 |
| Samd10 | 0.754628296 | 0.021651763 |
| Fkbp5 | -0.711605826 | 0.021708428 |
| 0610009B22Rik | 0.822595397 | 0.02171624 |
| Arl13a | -6.028028945 | 0.021842394 |
| Tnfrsf1b | -2.19665131 | 0.02184591 |
| Prdm1 | -1.530989117 | 0.021885958 |
| Cd109 | 3.370062328 | 0.021913462 |
| Rps6kc1 | -0.847474459 | 0.021949763 |
| Mblac2 | 1.158649446 | 0.021969971 |
| Pnma2 | 0.956980053 | 0.021975066 |
| Epgn | 5.719729009 | 0.022001337 |
| Mdm2 | -0.621711804 | 0.02203876 |
| Chst5 | 5.820947511 | 0.022056459 |
| Pdcl | -0.615684242 | 0.0220866 |
| Gm10134 | -2.131118119 | 0.02211286 |
| Fam19a5 | 1.451138948 | 0.022123171 |
| Arl14ep | 0.518037543 | 0.022140728 |
| Ndufs3 | 0.482204589 | 0.022145902 |
| Sympk | -0.653011897 | 0.022163278 |
| CK137956 | 5.367013507 | 0.022205985 |
| Mif4gd | -0.97874354 | 0.022252389 |
| Smg1 | -0.51583252 | 0.022252408 |
| Cep89 | 0.858268158 | 0.022303137 |
| Fafl | -0.509420904 | 0.022332561 |
| Gmpr | -0.877910154 | 0.022332561 |
| Gm13889 | 1.254681843 | 0.022337255 |
| Tmem237 | 0.594686568 | 0.022376434 |
| Kcnj10 | 1.446856465 | 0.022382421 |
| Mrps5 | 0.645048616 | 0.022427294 |
| Kank3 | 1.219090683 | 0.022432933 |
| Gab2 | -0.566562737 | 0.022452046 |
| Lrch2 | 0.740049363 | 0.022465871 |
| 1810043H04Rik | 0.654842535 | 0.022486381 |
| Daglb | -1.181407993 | 0.022486381 |

| | | |
|---|---|---|
| Mdm4 | -0.340783364 | 0.222102461 |
| Golga2 | -0.418457039 | 0.222102461 |
| A930011G23Rik | -0.458449768 | 0.222102461 |
| Chmp5 | 0.245378592 | 0.222163979 |
| Chst10 | 0.487098423 | 0.22219568 |
| Fat3 | -0.475474667 | 0.22219568 |
| Asb2 | -1.173590859 | 0.222298472 |
| Ppp2r2b | -0.422021234 | 0.222308506 |
| Fam221b | 0.381463018 | 0.222388982 |
| Kdelr1 | 0.345558853 | 0.222451618 |
| Fgd5 | 1.212274561 | 0.222465692 |
| Fbxo9 | 0.372337141 | 0.222465692 |
| 4930415F15Rik | -4.811268934 | 0.222465692 |
| Nfu1 | 0.314021849 | 0.222559168 |
| Ipmk | -0.528379464 | 0.222559168 |
| Bcl2l13 | -0.458437724 | 0.222593955 |
| Fchsd2 | 0.336435708 | 0.222679358 |
| Nwd1 | 1.690048205 | 0.222874076 |
| 4930579F01Rik | -5.066110138 | 0.222874076 |
| Gm43738 | 4.207722895 | 0.222981805 |
| Nxph3 | -1.166270983 | 0.223569251 |
| 1700016H13Rik | 4.253253642 | 0.223673015 |
| Cdh3 | 4.227311138 | 0.223673015 |
| Ptpn12 | 0.523643675 | 0.223673015 |
| Snapc3 | 0.304718358 | 0.223673015 |
| Plcg1 | -0.343799425 | 0.223673015 |
| Cd70 | 4.546552779 | 0.223820786 |
| Kat2b | -0.686621877 | 0.223879035 |
| Zfp772 | 0.908300514 | 0.224131799 |
| Ythdf2 | -0.260131818 | 0.224281141 |
| Purb | 0.356500131 | 0.22450759 |
| Fam217b | -0.697822772 | 0.224515023 |
| Dspp | -5.116513887 | 0.224563232 |
| Hmga1-rs1 | 0.492161663 | 0.224626497 |
| Azin2 | -0.791127383 | 0.224626497 |
| Btn1a1 | -3.498628877 | 0.224692343 |
| Aqp7 | -5.051436333 | 0.224692343 |
| Olfr1509 | -5.051436333 | 0.224692343 |
| Atpaf1 | -0.467731525 | 0.224773022 |
| Mpp6 | -0.400713524 | 0.224798274 |
| Get4 | 0.46033361 | 0.22484796 |
| Cdh11 | 0.820859982 | 0.224879253 |
| Fabp5 | 0.276513114 | 0.22492002 |
| BC026585 | -0.56279816 | 0.22492002 |
| Yeats2 | -0.416753328 | 0.225112746 |
| Zc3h12c | -0.47244071 | 0.22537954 |
| Gpr174 | -3.000314621 | 0.22537954 |
| Slc45a4 | 0.883384452 | 0.22545829 |
| Tmed5 | -0.417496652 | 0.22572342 |
| Klf16 | -0.581875444 | 0.225761733 |
| 4930415O20Rik | 4.132438335 | 0.226003671 |
| Klhdc4 | -0.463008816 | 0.226003671 |
| Pcdhb13 | 3.732646499 | 0.226011122 |
| Opalin | 4.651413433 | 0.226073029 |
| Synpo2l | 4.651413433 | 0.226073029 |
| Plekhh3 | 1.250957986 | 0.226257796 |
| Tmsb4x | 0.382167774 | 0.226257796 |
| Ophn1 | -0.33225408 | 0.226257796 |
| Mbd5 | -0.51368242 | 0.226257796 |
| Apol8 | -4.747003539 | 0.226470975 |
| Nudt22 | -0.503294431 | 0.226592592 |
| Tm6sf2 | -2.952806644 | 0.226641927 |
| Lama3 | -1.837504099 | 0.226962862 |
| Lrrc57 | 0.482454865 | 0.226993015 |
| Fam120b | -0.327988997 | 0.22704292 |
| Limd2 | -0.349926704 | 0.22704292 |
| Tshr | 4.215893853 | 0.22738331 |
| Tax1bp1 | 0.354308913 | 0.22738331 |
| Sike1 | -0.352794453 | 0.22738331 |
| Cep120 | 0.403589974 | 0.227409731 |
| Clec5a | -1.808086121 | 0.227554068 |
| Ptov1 | -0.329898952 | 0.227665879 |
| Clca3a1 | 4.52076317 | 0.227747834 |
| Phf21b | -0.370298064 | 0.227802985 |
| Ms4a14 | -1.749672685 | 0.227802985 |
| Fryl | -0.385404934 | 0.227843381 |
| Dpysl2 | -0.257635871 | 0.227922988 |
| D3Ertd254e | -0.633932629 | 0.227925321 |
| Eif3g | 0.245685455 | 0.228225258 |
| Appbp2 | 0.310326447 | 0.228250221 |
| Khsrp | -0.34862678 | 0.228250221 |
| 1700011I03Rik | 3.040920009 | 0.228272061 |
| Klf9 | 0.976141429 | 0.228272061 |
| Mvk | 0.493851163 | 0.228272061 |
| Mmd2 | 0.394085912 | 0.228272061 |
| Cul9 | -0.6656656 | 0.228272061 |
| Birc2 | -0.322824763 | 0.22836243 |
| Xpot | -0.475898244 | 0.228561361 |
| Umodl1 | 3.302389617 | 0.228648472 |
| 2310036O22Rik | 0.324663251 | 0.228714344 |
| Fibin | 1.339258468 | 0.22880729 |
| Rgma | 0.446256283 | 0.228894786 |
| Ska1 | 0.77238053 | 0.229117139 |
| Pstk | 0.624440784 | 0.229186886 |
| Acot2 | 0.435895235 | 0.229272322 |
| Gja1 | 1.490772255 | 0.229498257 |
| Ociad2 | 0.707284388 | 0.229498257 |
| Mtnr1a | 4.58662542 | 0.229520783 |
| Ttc8 | 0.554860363 | 0.229588501 |
| Igsf9b | -0.987773713 | 0.229588501 |
| Nek6 | -0.394689994 | 0.229654017 |
| Dmkn | -4.741890923 | 0.229680207 |
| Nrn1 | 0.493655277 | 0.229769193 |
| Trim32 | 0.336532984 | 0.229843583 |
| Ankmy2 | 0.356961 | 0.229915114 |
| Prpsap2 | 0.341866437 | 0.230114541 |
| Zcwpw1 | 1.281316049 | 0.230180711 |
| Fgf14 | 1.087104862 | 0.230218699 |
| Ccdc186 | 0.377001065 | 0.230218699 |
| Nt5c | 0.313552933 | 0.230218699 |
| Grik3 | -0.536231892 | 0.230218699 |
| Gltp | -0.463792483 | 0.230261432 |
| Afmid | 1.769104367 | 0.230653787 |
| Olfr111 | 4.171911713 | 0.230663648 |
| Tlcd1 | 1.461758856 | 0.231058452 |
| Tmem140 | -1.628334855 | 0.231058452 |
| Rab11b | 0.247183812 | 0.231200467 |
| 1600002K03Rik | 0.910307121 | 0.231296336 |
| Gusb | -0.458471446 | 0.231413357 |

| | | |
|---|---|---|
| Klhl32 | -0.268084338 | 0.594051729 |
| Slc25a10 | -0.319166603 | 0.594051729 |
| Tmem238 | 0.587906194 | 0.594214304 |
| Nkx2-3 | 0.670586803 | 0.594292285 |
| Otoa | 3.067686166 | 0.594377858 |
| Thoc3 | -0.142699545 | 0.594536093 |
| Acr | 0.702959635 | 0.594569424 |
| Zfp664 | 0.165023585 | 0.595196811 |
| Espn | 3.06193022 | 0.595228807 |
| Rhag | 3.06193022 | 0.595228807 |
| Pkp4 | 0.250947679 | 0.595228807 |
| Ankrd27 | -0.197261564 | 0.595228807 |
| Hs3st4 | -0.355196744 | 0.595228807 |
| 2310022A10Rik | -0.18491664 | 0.595342182 |
| Rassf1 | 0.212247393 | 0.595894635 |
| Mrpl46 | 0.20395706 | 0.595960104 |
| Vwa3b | -1.308075444 | 0.596258052 |
| Mospd2 | -0.287181624 | 0.596504963 |
| Rbpms | -0.524673592 | 0.596528557 |
| 1110032F04Rik | -0.694679116 | 0.596775142 |
| Pgpep1 | 0.252436332 | 0.596814456 |
| Kdelc1 | -0.325051172 | 0.59697023 |
| Gadd45gip1 | 0.447952229 | 0.597041424 |
| Prr5 | 0.356858406 | 0.597041424 |
| Cep63 | 0.190951147 | 0.597041424 |
| Ccndbp1 | 0.175764976 | 0.597041424 |
| Cct7 | 0.113985703 | 0.597041424 |
| Zfp27 | -0.350041686 | 0.597041424 |
| Syce1l | -1.607163575 | 0.597041424 |
| Pcnp | 0.17177326 | 0.597180179 |
| Cdsn | 3.046320563 | 0.597375412 |
| Fam19a4 | 3.046320563 | 0.597375412 |
| Slc26a5 | 3.046320563 | 0.597375412 |
| Tssk6 | 1.189431605 | 0.59737586 |
| Cfap97 | 0.258572517 | 0.59737586 |
| Ensa | 0.122969099 | 0.59737586 |
| Mterf2 | -0.282948239 | 0.59737586 |
| Tyk2 | -0.3067715 | 0.59737586 |
| Osbpl5 | 0.253124987 | 0.597702254 |
| Ubxn1 | 0.134980462 | 0.59775923 |
| Clp1 | -0.249259072 | 0.59775923 |
| Cyb5r3 | -0.131804261 | 0.597796839 |
| Zw10 | -0.206356207 | 0.598004774 |
| Scmh1 | -0.173130472 | 0.598143097 |
| Zfp617 | 0.306149212 | 0.598154465 |
| Cemip | -1.144442616 | 0.598247084 |
| Ybx2 | -0.68312331 | 0.598306477 |
| Clec4a4 | 1.044093366 | 0.598334647 |
| Leap2 | -1.751844113 | 0.598739764 |
| Zfp536 | 0.198839629 | 0.598942174 |
| Itgbl1 | -0.746918929 | 0.599013664 |
| Gm3512 | 1.794308904 | 0.599092249 |
| Oit3 | -1.604003008 | 0.599372668 |
| Hist1h4m | -1.099443321 | 0.599622825 |
| Gfap | 0.800339286 | 0.599971612 |
| Rrm1 | -0.190256153 | 0.60017418 |
| Fgf18 | 1.029962943 | 0.600439649 |
| 1110012L19Rik | 0.219984298 | 0.600439649 |
| St7 | -0.1622728 | 0.600439649 |
| Pbx2 | 0.337902157 | 0.600454847 |
| Usp33 | 0.163371242 | 0.600454847 |
| 1700047I17Rik2 | 0.782774209 | 0.600567618 |
| Gpat4 | -0.194762009 | 0.600738858 |
| Ptpn2 | 0.198332656 | 0.601130827 |
| Nudt3 | 0.126154829 | 0.601130827 |
| Nab2 | -0.457744441 | 0.601130827 |
| Rbm28 | 0.18500107 | 0.601213935 |
| Cacfd1 | 0.162136496 | 0.601213935 |
| Tmem33 | -0.142376946 | 0.601245376 |
| 9230104M06Rik | -1.304278764 | 0.601420658 |
| Rnf157 | 0.250171234 | 0.601436728 |
| Tmco1 | 0.139690844 | 0.601470716 |
| Pum2 | 0.11911087 | 0.601470716 |
| Esr1 | -2.271781897 | 0.601470716 |
| Macrod1 | -0.509137089 | 0.601905362 |
| Lrig1 | 0.505757449 | 0.602118039 |
| Mapkapk5 | -0.80532902 | 0.602292339 |
| Zfp932 | 0.336821827 | 0.602596657 |
| Irak1 | -0.207040331 | 0.602596657 |
| Casp3 | -0.1495141 | 0.602717633 |
| Nsun6 | 0.32289212 | 0.603035606 |
| Ifit1 | -1.006119236 | 0.603035606 |
| Ankrd23 | -0.969873808 | 0.60318865 |
| Hars2 | -0.204198306 | 0.603288 |
| Lancl3 | -0.334058593 | 0.604059443 |
| Strada | -0.203056063 | 0.604340031 |
| Tex12 | -1.069316036 | 0.604340031 |
| Hs3st2 | -0.534228534 | 0.604496786 |
| Opa3 | 0.203318392 | 0.604540463 |
| Hist1h4d | 0.188643355 | 0.604540463 |
| Samd15 | -1.015018625 | 0.604606563 |
| Rp9 | 0.172471428 | 0.60500781 |
| Nrd1 | -0.138670518 | 0.60500781 |
| Prcc | 0.317167734 | 0.605079806 |
| Minpp1 | 0.170369631 | 0.605290628 |
| Rgs11 | -1.372161828 | 0.60553259 |
| Slc4a1ap | 0.199885262 | 0.605721994 |
| Pigw | -0.55194284 | 0.605721994 |
| Gimap8 | 2.394666199 | 0.605740792 |
| Ubn2 | 0.153709893 | 0.605740792 |
| Nrg4 | -0.639488751 | 0.605740792 |
| Det1 | 0.330505692 | 0.605767339 |
| Ncoa4 | -0.229335947 | 0.605767339 |
| Ak3 | 0.157427778 | 0.606131627 |
| Rin1 | 1.063582266 | 0.606281731 |
| Cdh5 | 0.834767693 | 0.606281731 |
| Ltv1 | 0.191851528 | 0.606304324 |
| Zfp275 | -0.223678106 | 0.607040815 |
| Gpn2 | 0.279857277 | 0.607074716 |
| Kcnh2 | -0.423068157 | 0.607107558 |
| Pln | -0.889995703 | 0.607419805 |
| Klhdc2 | 0.116497338 | 0.607736574 |
| Gabbr2 | 0.365867083 | 0.607796499 |
| Sdk2 | 0.337956403 | 0.607796499 |
| Smarcd1 | -0.148540024 | 0.607796499 |
| Hist1h2bl | 1.263474939 | 0.608095 |
| Nkrf | 0.248003819 | 0.60842642 |
| Maoa | -0.237436995 | 0.60842642 |
| Nelfcd | 0.158341209 | 0.608666535 |

| | | |
|---|---|---|
| Pcdhga3 | -0.046313642 | 0.973182084 |
| Gm2174 | -0.085090641 | 0.973182084 |
| Myo1a | -0.113494526 | 0.973182084 |
| Gm7221 | -0.094871124 | 0.973242594 |
| B3gntl1 | 0.027565389 | 0.973605635 |
| Wdr59 | 0.02350938 | 0.973605635 |
| Slc25a12 | -0.010402707 | 0.973605635 |
| Tyw1 | -0.017769795 | 0.973819207 |
| Pkd1l3 | -0.080553749 | 0.973819207 |
| Bend5 | 0.014923045 | 0.974080442 |
| Rmnd5b | 0.011813858 | 0.974080442 |
| Arsk | -0.031582971 | 0.974080442 |
| Tmem209 | 0.017672568 | 0.974354155 |
| Sync | 0.09263142 | 0.974391528 |
| Fubp3 | 0.010158913 | 0.974438469 |
| Tmx2 | 0.009366703 | 0.974438469 |
| Mapk10 | -0.01057664 | 0.974438469 |
| Hadhb | -0.011926477 | 0.974438469 |
| Sc5d | -0.012634949 | 0.974438469 |
| Dpy19l1 | -0.01967005 | 0.974438469 |
| Plekha4 | -0.117188389 | 0.974438469 |
| Gm21953 | -0.125469923 | 0.974438469 |
| Slc25a30 | -0.031739208 | 0.974798237 |
| Rnf181 | 0.009803039 | 0.974866696 |
| Sgol2a | 0.018528714 | 0.974907587 |
| Zfp462 | -0.014970655 | 0.975054081 |
| Il34 | 0.071172507 | 0.975223263 |
| Tecpr1 | 0.012877544 | 0.975223263 |
| Egfl8 | 0.09884495 | 0.975346036 |
| Lrrc10b | 0.129120702 | 0.975403663 |
| Setdb1 | 0.013284469 | 0.975403949 |
| Parp2 | 0.011567441 | 0.975937612 |
| Hs3st6 | 0.090986907 | 0.976107891 |
| Dph5 | 0.012443241 | 0.976107891 |
| Slit2 | -0.019357036 | 0.976107891 |
| Aldh18a1 | -0.013063767 | 0.976174058 |
| Fsip1 | 0.031166573 | 0.976237472 |
| Smap1 | 0.015665417 | 0.976397695 |
| Shc4 | 0.061122984 | 0.976613124 |
| Usp14 | -0.008269949 | 0.976613124 |
| Rmnd5a | 0.015000231 | 0.976706478 |
| Ten1 | -0.011870176 | 0.976795413 |
| Ccdc137 | -0.020511989 | 0.976893756 |
| Ctnna2 | -0.011348204 | 0.977122984 |
| Kcna6 | -0.039209575 | 0.977122984 |
| Rab27b | -0.042306201 | 0.977122984 |
| Tyw3 | -0.025034589 | 0.97741383 |
| Zranb3 | 0.023436352 | 0.977440399 |
| Adamts7 | 0.016816855 | 0.977556649 |
| Shcbp1l | -0.050983559 | 0.977556649 |
| Exd2 | 0.013678714 | 0.977704074 |
| Srr | 0.012780784 | 0.97812455 |
| Kif20a | -0.013997916 | 0.97812455 |
| Qpctl | -0.013987042 | 0.978521373 |
| Gm37240 | 0.166624225 | 0.978681428 |
| Raet1d | -0.041820764 | 0.978681428 |
| Tppp | 0.034150536 | 0.978907044 |
| Nsun7 | -0.05245552 | 0.978937055 |
| Cd164l2 | 0.099459337 | 0.979122198 |
| Rnasek | 0.081646144 | 0.979122198 |
| Terf2 | -0.014119562 | 0.979122198 |
| Slc8a2 | -0.044405054 | 0.979122198 |
| Eef2k | -0.013237527 | 0.979132752 |
| Ankle2 | 0.009466737 | 0.979238865 |
| Cacna2d1 | -0.010165328 | 0.979238865 |
| St6gal2 | 0.010392119 | 0.97957812 |
| Pgap3 | 0.041977349 | 0.979697173 |
| Hps3 | -0.013503111 | 0.979720372 |
| Hmgcs1 | 0.007539486 | 0.979916525 |
| Mkks | 0.012308599 | 0.979929388 |
| Gm9938 | 0.042044564 | 0.980231675 |
| Gm6594 | 0.079966041 | 0.980391009 |
| Cxcl14 | 0.029193003 | 0.980722169 |
| Atpaf2 | 0.010665836 | 0.980722169 |
| Spcs2 | 0.006692153 | 0.980871336 |
| Slc4a5 | -0.069108281 | 0.980871336 |
| Hist2h2aa1 | 0.015894912 | 0.98131399 |
| Tekt2 | 0.018862236 | 0.981388929 |
| Zfp787 | 0.013407208 | 0.981388929 |
| Prmt6 | 0.010587986 | 0.981435725 |
| Ip6k1 | -0.008748216 | 0.981592984 |
| Abcb1b | 0.030849805 | 0.981767379 |
| Impdh2 | -0.007425864 | 0.982014197 |
| Cyp20a1 | 0.010952938 | 0.982710044 |
| Hypk | 0.038744932 | 0.982712607 |
| Cd81 | 0.007400226 | 0.982712607 |
| Rnf169 | -0.014296496 | 0.982712607 |
| Pde4dip | 0.006821209 | 0.983058526 |
| Zmat4 | 0.059767736 | 0.983108032 |
| Lrfn2 | 0.025863421 | 0.983108032 |
| 1700010I14Rik | 0.024708844 | 0.983108032 |
| Hlcs | 0.023385583 | 0.983108032 |
| Rbm14 | 0.009459558 | 0.983108032 |
| Fam43a | 0.020985723 | 0.983213387 |
| Magix | 0.076085094 | 0.983247539 |
| Onecut1 | 0.049250999 | 0.983249263 |
| Ccl21a | -0.055739934 | 0.983249263 |
| Padi2 | -0.041611444 | 0.983649974 |
| Mcts2 | 0.011631815 | 0.984149187 |
| Rbck1 | 0.007487612 | 0.984149187 |
| Fbxo32 | 0.017226434 | 0.984160598 |
| Chaf1a | 0.009256395 | 0.98425663 |
| Rusc1 | -0.010089917 | 0.98425663 |
| Chm | -0.011092829 | 0.984425814 |
| Il17c | -0.046653197 | 0.985444136 |
| Mrps34 | -0.014131442 | 0.985564452 |
| Ints12 | 0.007513711 | 0.98563475 |
| Chrnb1 | 0.041405832 | 0.985722982 |
| Nek3 | -0.01239003 | 0.985910189 |
| Tomm70a | -0.005731074 | 0.986012372 |
| Jarid2 | -0.006416841 | 0.986012372 |
| Apoe | 0.016662387 | 0.98601748 |
| Lrch3 | 0.007477856 | 0.98601748 |
| Slc2a8 | -0.012092816 | 0.986049978 |
| Slc25a29 | -0.012773336 | 0.986049978 |
| Lgals2 | -0.059198461 | 0.986049978 |
| Kctd8 | 0.022176471 | 0.98620437 |
| Gm28062 | -0.022153783 | 0.98620437 |
| Rad52 | -0.012927789 | 0.986763822 |

| | | |
|---|---|---|
| Pgbd1 | 2.340258666 | 0.022509552 |
| Lamtor4 | 0.6028012 | 0.022520157 |
| Bloc1s2 | 0.475642453 | 0.022520157 |
| Hs3st5 | -1.556319667 | 0.022520157 |
| Ppt1 | -0.657264638 | 0.022527972 |
| Mvb12b | -0.659795905 | 0.022573578 |
| Sipa1l1 | -0.872260759 | 0.022619159 |
| Zcchc6 | -0.572488022 | 0.022687637 |
| Lhx1 | 2.315954195 | 0.0226883 |
| Eme2 | -1.944948237 | 0.022715599 |
| Fcgr1 | -1.065376279 | 0.022729957 |
| Timm50 | 0.47682705 | 0.022793149 |
| St3gal2 | -0.98741689 | 0.022820329 |
| Plk5 | 5.596373295 | 0.022911481 |
| Flrt1 | -0.876198356 | 0.022911481 |
| Zfp385a | -0.876569967 | 0.022911481 |
| Grip2 | -1.897858375 | 0.022911481 |
| Clptm1l | -0.630377591 | 0.022917833 |
| H3f3b | 0.448269239 | 0.022923947 |
| Cacna1s | 5.440729488 | 0.022930981 |
| Nalcn | 1.62979472 | 0.022945669 |
| Zfp711 | 0.709732126 | 0.022945669 |
| Tmem11 | 0.622812599 | 0.022945669 |
| Ncor2 | -0.496944928 | 0.022945669 |
| Irf2bpl | -0.49091842 | 0.022995615 |
| Lrp2bp | -2.320940353 | 0.023022255 |
| AI607873 | -1.408083651 | 0.023099946 |
| Spag7 | 0.544486003 | 0.023113107 |
| Fbxo47 | -3.826026976 | 0.023113107 |
| Slc7a15 | -5.328899618 | 0.023134466 |
| 1810011H11Rik | 2.137798 | 0.023156732 |
| Tmem44 | -0.853046142 | 0.023205296 |
| Lpar5 | -2.533596025 | 0.023235051 |
| Kcnk10 | 0.925055055 | 0.023286209 |
| Ptbp1 | -0.573217517 | 0.023288261 |
| Cdk2ap2 | -0.600424693 | 0.023288261 |
| Mat2a | -0.46137793 | 0.02337977 |
| Npy5r | 5.570858787 | 0.023408643 |
| Zmat2 | 0.614469784 | 0.023452643 |
| Asrgl1 | 0.980726986 | 0.023483262 |
| 9530068E07Rik | -0.541909518 | 0.023485461 |
| Synm | 2.640284129 | 0.023650255 |
| Tubb6 | -0.997307038 | 0.0237095 |
| Asgr1 | 4.78827708 | 0.023742956 |
| Mrpl40 | 0.672233413 | 0.023753207 |
| Ece1 | -0.759974294 | 0.023821509 |
| Esam | 2.619714422 | 0.023849485 |
| Ccar2 | -0.595862831 | 0.023980734 |
| Pdcd6 | 0.505081495 | 0.023994495 |
| Tekt5 | 5.571242234 | 0.023997087 |
| Snx14 | -0.817823498 | 0.023998911 |
| Gba | -0.704550436 | 0.024007155 |
| 2210408I21Rik | 1.107531554 | 0.02404153 |
| Gsk3b | 0.371171899 | 0.02404153 |
| Tank | -0.86163666 | 0.02404153 |
| Srp9 | 0.542033947 | 0.024080655 |
| 6430573F11Rik | 1.912999305 | 0.024096001 |
| Camk1d | -0.797119895 | 0.024105539 |
| Trim12c | -0.938129593 | 0.024105539 |
| Tram1 | -0.595785176 | 0.024125671 |
| Tax1bp3 | 0.611515177 | 0.024166939 |
| Lingo1 | -0.93602352 | 0.02418409 |
| 4933424G06Rik | -3.039399019 | 0.02418409 |
| Apba2 | 0.493181467 | 0.024205036 |
| Rdh10 | 1.186191467 | 0.024205051 |
| Suds3 | -0.822817797 | 0.024218508 |
| Idh1 | 0.398425409 | 0.024230732 |
| Avpi1 | 1.526248961 | 0.024293257 |
| Tmem69 | 1.266990701 | 0.024333086 |
| Aldh1a1 | 5.71188877 | 0.024361448 |
| Wfdc17 | -2.639804462 | 0.024362152 |
| Irf2 | -0.926011026 | 0.024370013 |
| Alg9 | -0.777846029 | 0.024463076 |
| Iffo2 | 1.802399487 | 0.024477918 |
| Pknox2 | 1.152138369 | 0.02457888 |
| Rrp1b | -0.738084679 | 0.024587777 |
| Lrrc58 | -0.44801818 | 0.024596704 |
| Surf1 | 0.648696218 | 0.024632885 |
| Agtr2 | 5.715909421 | 0.024666778 |
| C1galt1 | 1.920368707 | 0.024666778 |
| Cdh20 | 4.038992175 | 0.024692663 |
| Atad5 | 0.724798219 | 0.024692663 |
| Hcfc1 | -0.597579384 | 0.024692663 |
| Cd48 | -1.232404045 | 0.024692663 |
| Cfb | -2.694543134 | 0.024692663 |
| Pde5a | 1.03064147 | 0.024722202 |
| Wac | -0.53459112 | 0.024722202 |
| Dhrs7b | 0.75145317 | 0.024748655 |
| Zfp111 | 1.969457852 | 0.024758602 |
| Cnrip1 | 0.669334802 | 0.024758602 |
| Mrps18a | 0.583154395 | 0.024758602 |
| Steap1 | 4.953665917 | 0.02476396 |
| Trappc3 | 0.540031997 | 0.02476396 |
| Siah1a | -0.573557402 | 0.02476396 |
| Dhcr24 | -1.045538473 | 0.02476396 |
| Slfn2 | -1.880910835 | 0.02476396 |
| Gm9847 | 2.702912503 | 0.024799517 |
| Ago2 | -0.588662392 | 0.024799517 |
| Ccnyl1 | 1.885323729 | 0.024841823 |
| Tbc1d5 | -1.014364809 | 0.024869383 |
| Fam96b | 0.510410032 | 0.024895672 |
| 6330403A02Rik | 1.954528337 | 0.024956366 |
| Cd247 | 5.489480211 | 0.025062988 |
| Fbxo40 | -4.522341946 | 0.02508903 |
| Fbxl6 | 0.686712379 | 0.025136651 |
| Aplp1 | -0.596649757 | 0.025167102 |
| Irf2bp1 | -0.712165525 | 0.025177915 |
| Ulk1 | -0.596527849 | 0.025178382 |
| Grm2 | 1.570037412 | 0.025193934 |
| Cyp27a1 | 1.967541378 | 0.025232868 |
| Tube1 | 1.136801569 | 0.025262406 |
| Cog1 | -0.581739571 | 0.025286387 |
| Serpinb1a | 2.543019895 | 0.025300166 |
| Chic2 | -0.569181452 | 0.025329639 |
| Arhgef18 | -0.950759556 | 0.025346503 |
| Lcmt1 | 0.607787919 | 0.025351411 |
| Taf1d | 0.901069134 | 0.02539074 |
| Hist1h2ag | 2.767965926 | 0.025406737 |
| Braf | -0.601398672 | 0.025452936 |

| | | |
|---|---|---|
| Vbp1 | 0.30155175 | 0.231426508 |
| Gucy1b3 | 0.632890088 | 0.231616247 |
| Cfap99 | 4.204973274 | 0.231656508 |
| Fam183b | 0.772941392 | 0.231741315 |
| Flrt3 | -0.372647644 | 0.231741315 |
| 9930111J21Rik1 | -2.3799203 | 0.231741315 |
| Cog5 | -1.076907192 | 0.231886852 |
| 4-Sep | 0.713558658 | 0.23188727 |
| Acbd6 | 0.372538792 | 0.231891426 |
| Hspa9 | -0.26961386 | 0.231923258 |
| Inpp4a | -0.340680924 | 0.231923258 |
| Utp14a | -0.443748594 | 0.231923258 |
| Sik3 | -0.569341482 | 0.231923258 |
| Pdpk1 | -0.312131302 | 0.232077744 |
| Nampt | -0.449346056 | 0.232077744 |
| Pla2g12a | 0.498436087 | 0.232099898 |
| Secisbp2l | 0.344862882 | 0.232107404 |
| P4htm | 0.737229057 | 0.232207505 |
| BC030500 | 0.862801187 | 0.232295437 |
| Tanc2 | -0.393755622 | 0.232295437 |
| Trim30d | -1.742623028 | 0.232308161 |
| Tmem131 | -0.340796154 | 0.232532255 |
| Klhl18 | -0.456896354 | 0.232532255 |
| Sh3rf3 | -0.481121481 | 0.232532255 |
| Tkfc | -0.641664582 | 0.232532255 |
| Mtch1 | -0.342200175 | 0.232678193 |
| Lamc2 | -1.349242732 | 0.232678193 |
| Cyb5r2 | -5.038963842 | 0.232678193 |
| Mdh2 | 0.231840317 | 0.232747084 |
| Pou6f1 | -0.485661495 | 0.232747084 |
| Pifo | 1.368182198 | 0.232775005 |
| Ganab | 0.361408609 | 0.232826073 |
| Susd2 | -1.524574408 | 0.232885855 |
| Megf11 | -0.647966999 | 0.233437084 |
| Gm13248 | -1.331276247 | 0.233437084 |
| A930033H14Rik | -2.880369591 | 0.233437084 |
| Per3 | 0.993624584 | 0.233524369 |
| Rasa1 | -0.429517028 | 0.233524369 |
| Zfp248 | -0.604851169 | 0.233593227 |
| Phc1 | 0.402703364 | 0.233761862 |
| Mtx1 | 0.339947176 | 0.233941859 |
| Gm17546 | -1.912259292 | 0.234018179 |
| Utp6 | 0.377716657 | 0.234078284 |
| Tmem121 | 0.458437425 | 0.234194948 |
| Uap1 | -0.426485932 | 0.234199885 |
| Tshz3 | -0.693618359 | 0.234199885 |
| Sox5 | -0.939839867 | 0.234199885 |
| Cd300c | 4.479764676 | 0.234226359 |
| Tnni3k | 4.110378033 | 0.234288353 |
| Efnb2 | 0.420109685 | 0.234617408 |
| Catsperd | -1.704574147 | 0.234834121 |
| Nyx | -2.109859587 | 0.23512775 |
| 4930519P11Rik | -5.017051731 | 0.23512775 |
| Degs2 | 1.639111737 | 0.235190679 |
| Slc1a2 | 0.547054137 | 0.235190679 |
| Heatr6 | -0.333432341 | 0.235374508 |
| Aipl1 | 4.309924869 | 0.235610566 |
| Ikzf4 | 0.646951036 | 0.235831472 |
| Fer | -0.355815961 | 0.235999838 |
| Nf2 | -0.358738549 | 0.235999838 |
| Zfp141 | -0.557517745 | 0.235999838 |
| Zfp217 | -0.61755039 | 0.236034806 |
| E430018J23Rik | -1.119688201 | 0.236034806 |
| Fam71a | -4.71391599 | 0.236034806 |
| Pate2 | -4.71391599 | 0.236034806 |
| Cfap54 | 1.133700632 | 0.236102904 |
| Spata1 | 1.122127364 | 0.236155081 |
| Cdc42 | -0.265563193 | 0.236238426 |
| Ttc33 | 0.399944226 | 0.236238673 |
| Pinx1 | 0.622901788 | 0.236260457 |
| Map2k7 | -0.517391023 | 0.236260457 |
| Cyld | -0.398354983 | 0.236400121 |
| Pfas | 0.286366402 | 0.236600123 |
| Ramp1 | 1.032962809 | 0.236711101 |
| Pex6 | -0.42399122 | 0.236711101 |
| Rbm4 | 0.862505385 | 0.23673002 |
| Pdzk1ip1 | 4.50686048 | 0.236919 |
| Lars2 | -0.417380451 | 0.236922429 |
| Fbxo48 | 4.456251935 | 0.236923981 |
| Muc6 | -1.184944964 | 0.2369982 |
| Tnrc6a | 0.411346659 | 0.237000183 |
| Acot3 | -1.668031149 | 0.237007588 |
| Prrc2b | -0.2970744 | 0.237066799 |
| Hdgfrp2 | 0.376090885 | 0.237198205 |
| Cacng7 | -0.796120758 | 0.237198205 |
| Ccdc103 | 1.292272286 | 0.237586549 |
| Ccbl1 | 0.748983558 | 0.237586549 |
| Stx1a | 0.584567635 | 0.237586549 |
| Cdca5 | 0.525185493 | 0.238064958 |
| Ddx20 | -0.36792862 | 0.238064958 |
| Dennd5b | 0.388292661 | 0.238077647 |
| Ranbp6 | -0.359814358 | 0.238164031 |
| Ankub1 | 4.291769929 | 0.238379427 |
| Aoc1 | 4.291769929 | 0.238379427 |
| Mrpl2 | 0.352027891 | 0.238379427 |
| Thumpd3 | -0.350270045 | 0.238658287 |
| Rrad | -0.944161545 | 0.238924235 |
| Cbr4 | -0.541032783 | 0.23900037 |
| Ap1g1 | -0.355955114 | 0.239296354 |
| Nr4a1 | -0.691951941 | 0.239609677 |
| Vash2 | 0.354554397 | 0.239710942 |
| Mylk3 | -4.248987958 | 0.239710942 |
| Sh3bp5l | 0.466726619 | 0.239774018 |
| Pcdh8 | -0.457578014 | 0.239774018 |
| Pappa2 | 1.090860341 | 0.239963114 |
| Necab2 | -1.61530952 | 0.239963114 |
| Isg20l2 | -0.434560666 | 0.240129992 |
| Dock11 | -0.305003631 | 0.240258702 |
| Ly6g5b | 4.465458734 | 0.240354797 |
| Gpr82 | 1.261384705 | 0.240354797 |
| Ufl1 | 0.407487668 | 0.240509428 |
| Eln | 0.9553392 | 0.241055278 |
| Ddx52 | 0.359763894 | 0.241099126 |
| Tsc22d4 | -0.321011638 | 0.241309548 |
| Slc35f5 | -0.478487871 | 0.241508581 |
| Grrp1 | 1.806349866 | 0.241765674 |
| Pdgfc | 1.151204607 | 0.241938921 |
| Fsd2 | 4.101465665 | 0.241958914 |
| Maml3 | 0.458150655 | 0.241965179 |

| | | |
|---|---|---|
| Hip1 | 0.220783186 | 0.608684235 |
| Mcm3ap | -0.171545702 | 0.608814529 |
| Wbscr16 | 0.227144005 | 0.608822104 |
| Spag16 | -0.695771917 | 0.608988935 |
| Lrrn4cl | 1.929953872 | 0.609162259 |
| Asic1 | -0.231383613 | 0.609162259 |
| St8sia6 | -0.899789098 | 0.609197012 |
| Stard8 | 0.510552944 | 0.609285168 |
| Fra10ac1 | 0.19944589 | 0.609299575 |
| Top2b | 0.12437197 | 0.609299575 |
| Faxc | -0.218276188 | 0.609299575 |
| Zfp39 | 0.344812664 | 0.60972242 |
| Zfp7 | 0.427049956 | 0.609760267 |
| Fxyd2 | 0.702950968 | 0.609792333 |
| Exoc7 | -0.152649365 | 0.609792333 |
| Igf1 | 0.472745513 | 0.610029072 |
| Ednra | 0.786545044 | 0.610142684 |
| Exoc6b | -0.191962778 | 0.610142684 |
| Fam83h | 1.121882443 | 0.610274516 |
| Pdlim4 | 0.160289751 | 0.610697182 |
| Adh5 | 0.124441589 | 0.610697182 |
| B4galt5 | -0.355749208 | 0.610697182 |
| Wdr19 | -0.390341502 | 0.61084986 |
| Pym1 | 0.282777952 | 0.611138821 |
| Slc25a33 | 0.235243351 | 0.611138821 |
| Tmem55b | -0.160366026 | 0.611191325 |
| Bmf | 0.431730353 | 0.611683704 |
| Ube2q1 | 0.231319552 | 0.611698406 |
| Myt1 | 0.207369199 | 0.611711343 |
| Pgap1 | -0.179370985 | 0.611789285 |
| Nup210l | 0.905416482 | 0.612218818 |
| Cyp2t4 | 2.136198949 | 0.612592661 |
| Cldn34c1 | 0.330150974 | 0.612754372 |
| Ccdc127 | 0.186808123 | 0.612765088 |
| Gm5422 | 0.429672444 | 0.612942254 |
| P3h1 | 0.418724777 | 0.613239211 |
| Ide | -0.18581354 | 0.613413953 |
| Chek1 | -0.269009558 | 0.613477735 |
| Dusp3 | 0.202679566 | 0.613617409 |
| Ccdc87 | 0.952388994 | 0.613828669 |
| Pdhx | 0.192993899 | 0.614005027 |
| Cct5 | 0.104863882 | 0.614005027 |
| Plekha3 | -0.475765187 | 0.614005027 |
| Plekhh1 | 0.779886789 | 0.614008362 |
| Rhno1 | 0.170145593 | 0.614008362 |
| Lrrtm2 | 0.382517264 | 0.614020772 |
| Plpp1 | 0.173466141 | 0.614020772 |
| Pop7 | 0.140881271 | 0.614026963 |
| Gm10382 | 1.269297539 | 0.614485543 |
| Mak16 | 0.193999954 | 0.614571826 |
| Xpo1 | 0.138567426 | 0.614571826 |
| Rs1 | -0.999719504 | 0.61462494 |
| Gapdh | -0.17184086 | 0.614953172 |
| Ttll4 | -0.206492925 | 0.614953172 |
| Mcc | -0.379592734 | 0.614953342 |
| Fam160a2 | -0.19128653 | 0.615074829 |
| Hpcal4 | -0.481746913 | 0.615297211 |
| Nudt15 | 0.617213616 | 0.615438129 |
| Arhgef4 | -0.349330512 | 0.615438129 |
| Tmub2 | -0.153988983 | 0.615656578 |
| Gm5089 | -0.599860507 | 0.615656578 |
| Traf3ip1 | 0.250330236 | 0.615736202 |
| Pcdh9 | 0.232988446 | 0.615965875 |
| Spdef | 0.939981327 | 0.616085998 |
| Dph7 | 0.196840378 | 0.616286947 |
| Gm6768 | 0.283465198 | 0.616361023 |
| Gm5786 | 1.104380926 | 0.616400037 |
| Lig3 | 0.275210627 | 0.616482535 |
| Lzts3 | 0.342214541 | 0.616693346 |
| Nhlh1 | 0.305036572 | 0.616693346 |
| Prkab1 | 0.167638358 | 0.616711532 |
| Vps26b | -0.167987662 | 0.616711532 |
| Ctbp2 | -0.173337751 | 0.616711532 |
| Sag | 1.142943795 | 0.616836638 |
| Plcb4 | 0.270650338 | 0.616881037 |
| Ptp4a2 | -0.151054059 | 0.616910579 |
| Mplkip | 0.165034424 | 0.617064864 |
| Uso1 | 0.214675862 | 0.617726779 |
| Cyb5b | -0.160619264 | 0.617773895 |
| Trim65 | -0.460931317 | 0.617773895 |
| Tmem220 | -1.14522712 | 0.617773895 |
| Celf2 | -0.159551877 | 0.618100538 |
| Pygm | 0.605904865 | 0.618189172 |
| Rtn4ip1 | 0.297721062 | 0.618189172 |
| Rbm19 | -0.304349551 | 0.618189172 |
| Hist1h2ab | 0.357451764 | 0.618244777 |
| Cenpi | 0.346732317 | 0.618244777 |
| Bco2 | -0.645019858 | 0.618244777 |
| Oprm1 | -0.677810775 | 0.618244777 |
| Ano7 | 1.900102166 | 0.618258981 |
| Ppp1r7 | -0.179311382 | 0.618258981 |
| Dtd1 | 0.193348026 | 0.618471186 |
| Hist1h2bk | 0.671033761 | 0.618971845 |
| Tet3 | -0.140101457 | 0.618971845 |
| Cdc20b | 1.180060326 | 0.618985806 |
| Il6st | 0.260530391 | 0.619265766 |
| Psmd8 | 0.116082119 | 0.619265766 |
| Fam168a | -0.157152958 | 0.619265766 |
| Wdr43 | -0.168617664 | 0.619265766 |
| Clip2 | -0.301309465 | 0.619265766 |
| Slc7a3 | 1.849685895 | 0.619368496 |
| Memo1 | -0.206453414 | 0.619553667 |
| Ints2 | -0.212034919 | 0.619823937 |
| Nmnat1 | 0.331158531 | 0.619872807 |
| Ormdl1 | 0.193472672 | 0.619911989 |
| Sptb | 0.582057323 | 0.619978935 |
| Hmbox1 | -0.239930457 | 0.620165714 |
| Mrto4 | 0.153619279 | 0.620240153 |
| Tom1l2 | 0.208151695 | 0.620445332 |
| Calm1 | 0.118262049 | 0.620938481 |
| Prr22 | 1.104043942 | 0.621070687 |
| Cecr5 | 0.272588007 | 0.621070687 |
| Pithd1 | 0.191719243 | 0.621452787 |
| Gareml | -0.240089393 | 0.621452787 |
| Inafm1 | 0.231603529 | 0.621518189 |
| Ddt | 0.371938998 | 0.621575065 |
| Pou3f3 | 0.172344845 | 0.621599285 |
| Ikzf2 | -0.419967757 | 0.621630641 |
| Gbp8 | -0.936223684 | 0.621630641 |

| | | |
|---|---|---|
| Nccrp1 | 0.047538222 | 0.986823709 |
| Reck | 0.021321604 | 0.986823709 |
| Gm9958 | 0.01742102 | 0.986823709 |
| Paip2b | 0.005760092 | 0.986823709 |
| Urm1 | -0.012082099 | 0.986823709 |
| Cd24a | -0.006079607 | 0.98692364 |
| Zfp354b | -0.030485864 | 0.98692364 |
| Sec61a2 | 0.00719229 | 0.987034217 |
| Ptpn22 | -0.023932303 | 0.987034217 |
| Cdh15 | -0.026051876 | 0.987034217 |
| Faf2 | -0.006295462 | 0.987123356 |
| Akap2 | 0.054310625 | 0.987251112 |
| Cct4 | -0.003995282 | 0.987272036 |
| Zdhhc20 | -0.005461941 | 0.987594198 |
| Usp50 | -0.008651041 | 0.987594198 |
| Serpine3 | -0.06117821 | 0.988019705 |
| Olfr57 | -0.018776136 | 0.988091284 |
| Olfml2a | -0.048902575 | 0.988091284 |
| Tnk2 | -0.010959492 | 0.988297724 |
| Tenm3 | 0.006912283 | 0.988387807 |
| Pwwp2a | 0.006488559 | 0.988387807 |
| Usp45 | -0.008875661 | 0.988387807 |
| Drp2 | -0.011100639 | 0.988396785 |
| Hist1h3b | 0.027738457 | 0.988412549 |
| Barhl1 | -0.031554914 | 0.988646004 |
| Smyd5 | 0.007201477 | 0.988802303 |
| Usp53 | -0.009386034 | 0.989150456 |
| Zfp112 | -0.018449139 | 0.989150456 |
| Grwd1 | 0.006538763 | 0.989176166 |
| Smarcc2 | -0.004483292 | 0.989176166 |
| Suclg2 | 0.008579811 | 0.989241049 |
| Men1 | 0.004654794 | 0.989729211 |
| Kctd16 | -0.025504319 | 0.989859504 |
| Nrm | 0.007124604 | 0.989934847 |
| Catsper2 | 0.011111621 | 0.990462096 |
| Noxo1 | -0.020254276 | 0.990462096 |
| Rab38 | -0.020791304 | 0.990462096 |
| Orm3 | -0.031952408 | 0.990462096 |
| Gm5239 | 0.034002216 | 0.990517412 |
| Hcn2 | 0.039028596 | 0.990613492 |
| Tbc1d2 | 0.024609275 | 0.990613492 |
| Pcdhb5 | 0.018532683 | 0.990613492 |
| Ina | 0.004595972 | 0.990613492 |
| Eif6 | 0.004049491 | 0.990613492 |
| Rdh11 | -0.006112119 | 0.990613492 |
| Pex10 | -0.007162896 | 0.990613492 |
| Ube2t | -0.007238781 | 0.990613492 |
| Arfip1 | -0.014397801 | 0.990613492 |
| Fam89a | -0.024700685 | 0.990613492 |
| Ncbp3 | 0.00521443 | 0.990725747 |
| Sema6d | -0.004134469 | 0.990725747 |
| Rars2 | -0.004266795 | 0.990725747 |
| Ube2ql1 | -0.007061046 | 0.990725747 |
| Phldb2 | -0.008917801 | 0.990725747 |
| 2210404O09Ri | -0.010398268 | 0.990725747 |
| Gramd3 | -0.013001892 | 0.990725747 |
| Abhd12b | -0.045445372 | 0.990725747 |
| Tex30 | -0.005474184 | 0.990891888 |
| Phlpp2 | -0.00665883 | 0.990891888 |
| Gm14326 | -0.010799567 | 0.990891888 |
| Htr7 | 0.030648843 | 0.990965044 |
| Slc17a9 | 0.026789491 | 0.991332563 |
| Tmem167b | 0.004501553 | 0.991332563 |
| Ttll11 | -0.012281332 | 0.991332563 |
| Wdr55 | 0.007908412 | 0.991404257 |
| Atp6v1e1 | 0.003311268 | 0.991762214 |
| Gapvd1 | -0.004054964 | 0.991762214 |
| Slc16a8 | -0.023287863 | 0.991802854 |
| Nktr | -0.003927125 | 0.991880463 |
| Lhb | 0.034461847 | 0.992101699 |
| Fbxw7 | 0.003631234 | 0.992101699 |
| Cat | 0.003119729 | 0.992101699 |
| Farsb | 0.003027456 | 0.992101699 |
| Myo9a | -0.003447949 | 0.992101699 |
| Fktn | -0.005348241 | 0.992101699 |
| Pnpla6 | -0.00800802 | 0.992101699 |
| Grin2a | -0.015137972 | 0.992101699 |
| Rnasel | -0.004624566 | 0.992122118 |
| Nfx1 | -0.003653349 | 0.992430957 |
| Zfp157 | -0.005005955 | 0.992430957 |
| Pcp4l1 | 0.007691915 | 0.992431659 |
| Ttc17 | -0.004466257 | 0.992431659 |
| Slc12a6 | -0.004740492 | 0.992692709 |
| Itgae | -0.019744245 | 0.993177262 |
| Zfp945 | -0.003138079 | 0.993214357 |
| Hprt | -0.004332453 | 0.993223868 |
| 6-Sep | -0.002975065 | 0.993638237 |
| Vps37d | 0.005016627 | 0.993688883 |
| Socs7 | 0.003686252 | 0.993935961 |
| Med16 | -0.002601775 | 0.993935961 |
| Anks3 | -0.005207463 | 0.993935961 |
| Lactbl1 | 0.037801821 | 0.993958935 |
| Abcf1 | -0.002156556 | 0.994110676 |
| Zfp85 | 0.011821738 | 0.994202128 |
| Pcbp1 | -0.002150226 | 0.994334923 |
| Dct | -0.023523683 | 0.994334923 |
| Ptpn13 | 0.005494065 | 0.994476353 |
| Zfp879 | -0.015225964 | 0.994552251 |
| Zfp169 | -0.005230262 | 0.994629812 |
| Htr1f | 0.01610097 | 0.995392135 |
| Zfp729a | 0.005534901 | 0.995392135 |
| Sema6a | 0.002545318 | 0.995392135 |
| Tfam | 0.00237726 | 0.995392135 |
| Acox1 | 0.002147563 | 0.995392135 |
| Gpsm1 | 0.002103946 | 0.995392135 |
| Slc35b1 | 0.001797511 | 0.995392135 |
| Fmnl2 | -0.002253737 | 0.995392135 |
| Dtnb | -0.003014735 | 0.995392135 |
| Arfgef1 | 0.002820666 | 0.995459495 |
| Madcam1 | -0.012640714 | 0.995459495 |
| Tnks1bp1 | -0.004551467 | 0.995484625 |
| Cyp2s1 | 0.010690329 | 0.995622124 |
| Large | 0.003375732 | 0.995798448 |
| Nrp1 | 0.002355536 | 0.995798448 |
| Slamf8 | 0.005133685 | 0.995800859 |
| Fut9 | -0.002872821 | 0.996067621 |
| Cstb | -0.002056216 | 0.996135736 |
| Vps33b | -0.002097432 | 0.996135736 |
| Serpinb6b | 0.006404864 | 0.996540026 |

| | | |
|---|---|---|
| Fst | 2.068914809 | 0.025475772 |
| Fuz | 1.09259296 | 0.025481855 |
| Acer2 | 1.515161623 | 0.025512796 |
| Itpkc | -1.472699521 | 0.025554375 |
| Syne2 | 0.716937774 | 0.025576641 |
| Dapk3 | -0.746980931 | 0.025578402 |
| St5 | -1.0925513 | 0.025578402 |
| Btbd16 | -3.785464534 | 0.025593229 |
| Cep250 | -0.893770037 | 0.025630231 |
| Cptp | -1.075526485 | 0.025633143 |
| Cbfa2t3 | 0.762624827 | 0.025724742 |
| Phc2 | 0.631990791 | 0.025756942 |
| Emc4 | 0.505329128 | 0.025756942 |
| Polr3k | 0.58994186 | 0.025764949 |
| Pcolce | 2.821218639 | 0.025782344 |
| Psmc3ip | 0.847358812 | 0.025782344 |
| Cd3g | -5.439998211 | 0.025829246 |
| Zkscan8 | 1.493765063 | 0.025838557 |
| Syce2 | 0.677448431 | 0.025838557 |
| Dcaf11 | 0.541087345 | 0.025846202 |
| Tesk1 | -1.688316743 | 0.025857642 |
| Crispld1 | 2.280239027 | 0.025867981 |
| Usp43 | -3.107378318 | 0.02600033 |
| Rrm2 | -0.817766795 | 0.026048283 |
| Opa1 | 0.538975143 | 0.026093287 |
| Stk16 | 0.565049893 | 0.026167019 |
| Gpatch3 | -0.97232161 | 0.026201752 |
| Wdfy1 | 0.664103252 | 0.026206509 |
| 3110002H16Rik | -0.650093844 | 0.026206509 |
| Psmb7 | 0.683944597 | 0.026212165 |
| Runx1 | -1.2997218 | 0.026214079 |
| Rab3d | -0.747615565 | 0.026260058 |
| Rnf19a | -0.69148954 | 0.026359002 |
| Ubc | 0.806415394 | 0.026455762 |
| Sirt7 | -0.619726162 | 0.026455762 |
| Ebpl | 0.674071778 | 0.026466702 |
| Taf15 | -0.521221622 | 0.02646946 |
| Negr1 | 0.966703893 | 0.026492588 |
| Bclaf1 | 0.43352029 | 0.026492588 |
| Hace1 | -0.971903145 | 0.026548441 |
| Rap1a | -0.515717061 | 0.026564069 |
| Fam78a | -1.791979975 | 0.026587512 |

| | | |
|---|---|---|
| Gramd4 | 0.535690312 | 0.241991388 |
| Cpeb3 | -0.723907485 | 0.241991388 |
| Ms4a12 | 4.502196021 | 0.242008216 |
| Arhgap28 | 1.287146838 | 0.242008216 |
| Dmtf1 | 0.334187788 | 0.242231007 |
| Herc3 | -0.51012104 | 0.242407704 |
| Syndig1 | 0.647225141 | 0.242481366 |
| Spef2 | 1.306529033 | 0.242506519 |
| Dzip3 | 0.282590868 | 0.242747266 |
| Arcn1 | -0.330051961 | 0.24283855 |
| Chfr | -0.310688095 | 0.242855062 |
| Cutal | 1.95873206 | 0.242983092 |
| Tal2 | 2.300502007 | 0.243110629 |
| Reg3g | 4.279599371 | 0.243452094 |
| Angptl4 | 1.461249054 | 0.243540211 |
| Tmem60 | 0.314244714 | 0.243540211 |
| Zfp346 | 0.577541151 | 0.243632348 |
| Dctn5 | 0.251962626 | 0.244050505 |
| N6amt1 | 0.512098945 | 0.244616785 |
| Mapk8ip1 | -0.353820017 | 0.244616785 |
| 1700008O03Rik | -2.023616647 | 0.244616785 |
| Gemin4 | 3.397376604 | 0.244871378 |
| Zfand4 | 0.795270319 | 0.244871378 |
| Gnb4 | 0.462272312 | 0.244871378 |
| Sbds | 0.362636946 | 0.244871378 |
| Homer1 | -0.503602198 | 0.244871378 |
| Gm10722 | 3.552644012 | 0.24494878 |
| AI597479 | -0.407194736 | 0.244984685 |
| Cmpk1 | -0.296149067 | 0.245061425 |
| Rnd1 | -0.519067276 | 0.245073817 |
| Snupn | 0.50385129 | 0.245198252 |
| Ptgs2 | -1.852357631 | 0.245240128 |
| Chst3 | 1.82526922 | 0.245244616 |
| Car8 | 1.252508918 | 0.245244616 |
| Gpr141 | -2.310282148 | 0.245244616 |
| Axin1 | 0.622893935 | 0.245325889 |
| Urb1 | -0.778651581 | 0.245437271 |
| Zbed4 | -0.500781122 | 0.245477727 |
| Nr2c1 | -0.561852953 | 0.245502914 |
| Peg3 | 0.379362863 | 0.245512837 |
| Des | 4.432674904 | 0.245569171 |
| Lzts2 | 0.497934557 | 0.245582104 |

| | | |
|---|---|---|
| Sema3a | 0.336951312 | 0.6219636 |
| Rere | -0.161361926 | 0.622016572 |
| Zranb1 | 0.165571552 | 0.622265562 |
| Tmc3 | -0.754897145 | 0.622440916 |
| Hdac6 | 0.140578865 | 0.622540366 |
| Fam81a | -0.268081282 | 0.62254234 |
| Fyttd1 | 0.14656029 | 0.622583715 |
| Scube2 | -1.375100667 | 0.622583715 |
| Gm4245 | -1.302292392 | 0.622691641 |
| Dctn2 | 0.12549305 | 0.622949833 |
| Srsf11 | 0.130576846 | 0.622974057 |
| Clec4n | -0.744830118 | 0.622997355 |
| Pipox | 0.573116607 | 0.622999055 |
| Cpt1c | 0.216123184 | 0.623191637 |
| Gm5796 | 0.560920521 | 0.623228071 |
| Gm3667 | 0.432727001 | 0.623228071 |
| Chrm1 | -0.522722822 | 0.623228071 |
| Chrm3 | 0.796235333 | 0.623278146 |
| Myo18b | 0.842279245 | 0.623292111 |
| Gm17305 | 1.064650132 | 0.623745098 |
| Inpp5f | 0.147570186 | 0.62377514 |
| Tigd2 | -0.208806332 | 0.62377514 |
| Icos | 0.785006924 | 0.623808382 |
| Hdac1 | -0.147452334 | 0.623808382 |
| Sesn2 | -0.294766602 | 0.623808382 |
| Luc7l | -0.13804922 | 0.623824229 |
| Vmn2r87 | -1.324412703 | 0.623824229 |
| Ubald2 | 0.202949458 | 0.624114265 |
| Larp1 | 0.160450966 | 0.624114914 |
| Acap2 | -0.183244543 | 0.624114914 |
| Podn | -0.832664603 | 0.624114914 |
| Cited4 | 0.720004572 | 0.624248842 |
| Txlng | -0.207375175 | 0.624248842 |
| Cnot8 | 0.138025662 | 0.624454632 |
| Tspan31 | 0.183835433 | 0.624496241 |
| Ank3 | 0.180578932 | 0.624519876 |
| Fam212b | 0.867788752 | 0.624668266 |
| Ezh1 | 0.211448614 | 0.624668266 |
| Lmbr1 | -0.268913279 | 0.624668266 |
| Dync1h1 | 0.13482122 | 0.624845155 |
| Cort | -0.812346925 | 0.624846176 |
| Kcnj3 | -0.426808 | 0.625547606 |

| | | |
|---|---|---|
| Pip5k1a | 0.002313896 | 0.996540026 |
| Elavl2 | 0.001722168 | 0.996540026 |
| Efcab11 | -0.003783634 | 0.996540026 |
| BC067074 | -0.006249352 | 0.996746367 |
| Gm26620 | 0.024484112 | 0.996957059 |
| Espl1 | 0.002756531 | 0.996957059 |
| 9030624G23Ri | 0.01334925 | 0.997021326 |
| B630019K06R | 0.001948693 | 0.997371694 |
| Ggn | 0.008572604 | 0.997541889 |
| Col7a1 | 0.009873232 | 0.997591432 |
| Kcnc2 | 0.002527688 | 0.997591432 |
| Slitrk4 | -0.002939979 | 0.997591432 |
| Baiap2l1 | -0.004654849 | 0.997591432 |
| Wnt3a | -0.007351148 | 0.997663493 |
| Arl6ip4 | -0.00117142 | 0.997772696 |
| Lmo4 | -0.000850599 | 0.997873611 |
| Harbi1 | 0.001745955 | 0.997948197 |
| Acsl6 | 0.001283668 | 0.998007778 |
| Trabd | 0.001120771 | 0.998091728 |
| Cbarp | -0.001254682 | 0.998139497 |
| Acat3 | -0.004681709 | 0.998248569 |
| Fam131c | 0.001641815 | 0.998458577 |
| Rasa2 | -0.001105836 | 0.998458577 |
| Plcxd1 | -0.004848326 | 0.998458577 |
| Zfp316 | 0.000792062 | 0.998549989 |
| Prpf18 | 0.000767974 | 0.998549989 |
| Pa2g4 | 0.000414414 | 0.998700331 |
| C130074G19Ri | -0.003223673 | 0.998902372 |
| Zfand2b | -0.000626389 | 0.998907821 |
| Gm21370 | 0.004591785 | 0.998964301 |
| Enah | 0.000408827 | 0.998977887 |
| Gm17349 | -0.002873669 | 0.998977887 |
| Zc3h8 | -0.000819664 | 0.999079877 |
| Nhsl1 | 0.000464103 | 0.999161851 |
| Dbr1 | 0.000549145 | 0.999195569 |
| 2010107G12Ri | 0.001746058 | 0.999512328 |
| Pcdhga10 | 0.000839685 | 0.999550036 |
| Tgfbr3l | -0.000654501 | 0.999550036 |
| Gm21685 | -0.001335936 | 0.999550036 |
| Fndc7 | -0.000186423 | 0.999819311 |
| Snx8 | 7.03E-05 | 0.999829881 |

**Appendix Table S2.** FPKM in Ctrl and Ptf1a RGC.

| gene | fpkm_Gtrl1 | fpkm_Gtrl2 | fpkm_Gtrl3 | fpkm_Ptfla1 | fpkm_Ptfla2 | fpkm_Ptfla3 |
|---|---|---|---|---|---|---|
| Elane | 0 | 0 | 0 | 26.8913 | 1.78083 | 13.3779 |
| Lum | 0 | 0 | 0 | 2.93772 | 1.35067 | 5.40425 |
| Aldh1a2 | 0 | 0 | 0 | 2.95136 | 3.96369 | 3.74589 |
| Bgn | 0.00927537 | 0 | 0 | 0.317808 | 0.54265 | 0.767978 |
| Kirrel2 | 0 | 0 | 0 | 4.14407 | 3.46757 | 2.67621 |
| Abcc9 | 0 | 0 | 0 | 0.631981 | 0.491438 | 0.310925 |
| Hmgcs2 | 0 | 0 | 0 | 1.72868 | 1.29639 | 1.13643 |
| Lama4 | 0 | 0 | 0 | 0.254795 | 0.117448 | 0.271583 |
| Gjb6 | 0 | 0 | 0 | 1.26897 | 0.860833 | 2.55475 |
| Igfbp7 | 0.0946539 | 0 | 0 | 13.1752 | 11.6769 | 2.33866 |
| Msx1 | 0 | 0.0403419 | 0 | 2.38133 | 2.56498 | 5.48718 |
| Stc1 | 0 | 0 | 0 | 0.862666 | 0.317088 | 0.997203 |
| Lrrc3b | 0 | 0 | 0 | 1.88873 | 2.15452 | 1.28698 |
| Prss12 | 0 | 0 | 0 | 3.54759 | 0.106962 | 0.534975 |
| Rerg | 0 | 0 | 0 | 2.51718 | 0.15038 | 0.637578 |
| Cyp1b1 | 0 | 0 | 0 | 0.488432 | 0.947022 | 0.390835 |
| Pdlim3 | 0 | 0 | 0 | 2.41618 | 0.643762 | 3.55562 |
| Fbln5 | 0 | 0 | 0 | 0.145869 | 0.534913 | 0.383063 |
| Slc13a4 | 0 | 0 | 0.0140166 | 0.359497 | 0.388234 | 1.95698 |
| Olig3 | 0 | 0 | 0 | 0.218667 | 3.05447 | 2.32711 |
| Lepr | 0 | 0 | 0 | 0.0819855 | 0.110972 | 0.562904 |
| Ibsp | 0 | 0 | 0.0314601 | 4.06384 | 0.829705 | 1.22081 |
| Thbs2 | 0 | 0 | 0 | 0.557478 | 0.337553 | 0.190155 |
| Rnf138rt1 | 0 | 0 | 0 | 2.63834 | 5.20762 | 2.46374 |
| Ccdc3 | 0 | 0 | 0 | 1.01766 | 1.28988 | 0.586353 |
| H2-Ab1 | 0 | 0 | 0 | 0.08431 | 0.780037 | 1.54715 |
| Cdh18 | 0 | 0 | 0 | 0.417417 | 0.241381 | 0.36174 |
| Heyl | 0 | 0 | 0 | 0.249422 | 0.309997 | 0.713789 |
| Svep1 | 0 | 0 | 0 | 0.0419757 | 0.126219 | 0.136776 |
| Olfml1 | 0 | 0 | 0 | 2.06509 | 0.381464 | 1.53476 |
| Trim10 | 0 | 0 | 0 | 1.16431 | 1.8248 | 0.724999 |
| Cldn5 | 0 | 0.102286 | 0 | 11.2271 | 6.1995 | 3.06634 |
| Alx3 | 0 | 0 | 0 | 2.63418 | 1.56675 | 0.749943 |
| Ptpn14 | 0 | 0 | 0 | 0.300356 | 0.0860131 | 0.0557486 |
| Arhgap36 | 0 | 0 | 0 | 0.940581 | 0.416719 | 0.527537 |
| Ndnf | 0 | 0.00569584 | 0 | 0.145126 | 0.176618 | 0.422614 |
| Xpnpep2 | 0 | 0 | 0 | 0.0248306 | 0.360603 | 0.275806 |
| Sdsl | 0 | 0 | 0 | 1.27557 | 0.771649 | 4.27849 |
| Kif6 | 0 | 0 | 0 | 0.172001 | 0.134009 | 0.114086 |
| Ms4a3 | 0 | 0 | 0 | 0.694047 | 0 | 0.381705 |
| Trim16 | 0 | 0 | 0 | 0.371334 | 0.573479 | 0.133084 |
| Adamts9 | 0 | 0 | 0.00292574 | 0.0550896 | 0.0116907 | 0.119219 |
| Sult1a1 | 0 | 0 | 0 | 0.951132 | 0.585322 | 0 |
| Fgfbp1 | 0 | 0 | 0.0310959 | 0.646003 | 1.27507 | 2.20772 |
| 4932438H23Rik | 0 | 0 | 0 | 0.150932 | 0.386687 | 1.09147 |
| Slitrk6 | 0 | 0.0139241 | 0 | 0.0531668 | 1.30035 | 0.225774 |
| Vtn | 0.0415109 | 0.0903201 | 0.0395195 | 9.35838 | 4.43129 | 5.12552 |
| Afap1l2 | 0 | 0 | 0 | 0.30795 | 0.337017 | 0.141389 |
| Mal2 | 0 | 0.0218953 | 0.0191767 | 2.20745 | 1.43366 | 0.927361 |
| Apold1 | 0 | 0 | 0 | 0.625242 | 0.0754194 | 0.594184 |
| Adgrl4 | 0 | 0 | 0 | 0.556073 | 0.121178 | 0.0474339 |
| Rbm46 | 0 | 0 | 0 | 0.266482 | 0.343633 | 0.482194 |
| Bmp5 | 0 | 0 | 0 | 0.334166 | 0.161866 | 0.405188 |
| Adgrf5 | 0 | 0 | 0 | 0.257866 | 0.145963 | 0.0138764 |
| Col4a6 | 0 | 0 | 0 | 0.177655 | 0.200913 | 0.15176 |
| Bmp4 | 0 | 0 | 0 | 0.547351 | 0.655949 | 1.11767 |
| Prdm13 | 0 | 0 | 0.0212374 | 0.678087 | 0.62489 | 0.751767 |
| Zbbx | 0 | 0 | 0 | 0.315518 | 0.212196 | 0.309774 |
| Myl7 | 0 | 0 | 0 | 10.8622 | 0.54201 | 0.222825 |
| Dusp27 | 0 | 0 | 0 | 0.0364485 | 0.719597 | 0.571263 |
| Ltc4s | 0 | 0 | 0 | 5.01256 | 1.76273 | 4.10452 |
| Col5a1 | 0 | 0 | 0 | 0.179041 | 0.205523 | 0.158426 |
| Kcne2 | 0.0457749 | 0 | 0 | 1.27282 | 0.483577 | 3.1587 |
| Cldn11 | 0.207495 | 0.111748 | 0.196879 | 9.39014 | 15.0276 | 33.4838 |
| Slc22a8 | 0 | 0 | 0 | 0.431727 | 0.23477 | 0.194511 |
| 3830403N18Rik | 0 | 0 | 0 | 1.31391 | 1.90744 | 0.161704 |
| Pax3 | 0 | 0 | 0 | 0.283918 | 0.0561362 | 0.193808 |
| Twist2 | 0 | 0 | 0 | 0.39116 | 0.619324 | 0.289845 |
| Csgalnact1 | 0 | 0 | 0 | 0.336325 | 0.214742 | 0.202872 |
| Aard | 0 | 0 | 0 | 4.0244 | 0.411173 | 1.36856 |
| Dnase1l3 | 0 | 0 | 0 | 0.13452 | 0.355907 | 0.0052218 |
| Vwc2l | 0 | 0 | 0 | 0.286994 | 0.107982 | 0.22771 |
| Rassf9 | 0 | 0 | 0 | 0.645253 | 0.620447 | 0.395288 |
| Sh2d5 | 0 | 0 | 0 | 0.622393 | 0.18482 | 0.271192 |
| Zfp92 | 0 | 0 | 0 | 0.193687 | 0.204296 | 0.0579863 |
| Hapln3 | 0 | 0 | 0 | 0.316992 | 0.0810578 | 0.277096 |
| Hcrtr2 | 0 | 0 | 0 | 0.271062 | 0.0664885 | 0.111226 |
| Ephx2 | 0 | 0 | 0 | 0.879357 | 0.669001 | 0.227528 |
| Crb3 | 0 | 0 | 0 | 0.763995 | 0.339954 | 1.73965 |
| Lamc3 | 0 | 0 | 0 | 0.282892 | 0.049439 | 0.357065 |
| Layn | 0 | 0 | 0 | 0.780627 | 0.427949 | 0.945012 |
| Iqca | 0 | 0 | 0 | 0.138814 | 0.0569018 | 0.582716 |
| Kcnj2 | 0 | 0 | 0 | 0.00607105 | 0.494516 | 0.105283 |
| 2310030G06Rik | 0 | 0 | 0 | 0.449851 | 2.58763 | 0.194824 |
| Tmem204 | 0 | 0 | 0 | 0.132549 | 0.200728 | 1.18821 |
| Pdyn | 0 | 0 | 0 | 1.13265 | 0.152286 | 0.0658474 |
| D630003M21Rik | 0 | 0 | 0 | 0.369847 | 0.137976 | 0.307565 |
| Calca | 0 | 0 | 0 | 0.0964905 | 0.306509 | 0.979932 |
| Ctsg | 0.192149 | 0 | 0 | 13.562 | 1.55691 | 3.57475 |
| Pthlh | 0 | 0 | 0 | 2.0926 | 0.471522 | 0.623208 |
| Abcg3 | 0 | 0 | 0 | 0.282732 | 0.205268 | 0.0965336 |
| Hey2 | 0 | 0 | 0 | 0.419727 | 0.458453 | 0.232517 |
| H2-Aa | 0 | 0 | 0 | 0.100576 | 0.0427676 | 1.31276 |
| Cmbl | 0 | 0 | 0 | 0.105406 | 1.37926 | 0.895286 |
| C1ra | 0 | 0 | 0 | 0.467101 | 0.666738 | 0.0111697 |
| Piwil4 | 0 | 0 | 0 | 0.248714 | 0.0779766 | 0.0316648 |
| Cfap46 | 0 | 0 | 0 | 0.0836982 | 0.0040372 | 0.0516578 |
| Tcaf2 | 0 | 0 | 0 | 0.103773 | 0.0293426 | 0.459574 |
| Cbln4 | 0 | 0 | 0 | 0.586383 | 0.136997 | 0.401523 |
| Syde1 | 0 | 0 | 0.0256156 | 1.18558 | 0.504389 | 0.900534 |
| 4921511H03Rik | 0 | 0 | 0 | 0.212752 | 0.232684 | 0.051034 |
| Plek2 | 0 | 0 | 0 | 1.2249 | 0.945182 | 1.22666 |

| gene | fpkm_Gtrl1 | fpkm_Gtrl2 | fpkm_Gtrl3 | fpkm_Ptfla1 | fpkm_Ptfla2 | fpkm_Ptfla3 |
|---|---|---|---|---|---|---|
| Itpripl1 | 14.7298 | 6.52567 | 14.7117 | 8.50632 | 5.51985 | 5.61012 |
| Lgals1 | 378.857 | 355.162 | 548.264 | 198.972 | 350.705 | 167.403 |
| Eif4ebp2 | 26.0976 | 19.8675 | 22.8595 | 10.6501 | 11.4265 | 15.9297 |
| Prdx5 | 32.3022 | 35.5301 | 45.9507 | 26.5644 | 31.6555 | 25.4497 |
| Arhgef11 | 1.83274 | 1.30205 | 1.67022 | 0.967369 | 0.795715 | 1.03616 |
| Vti1a | 9.36679 | 4.8969 | 7.44685 | 5.00944 | 3.8519 | 2.6641 |
| Zdhhc12 | 6.01465 | 7.10104 | 9.39034 | 3.91572 | 4.28638 | 4.49351 |
| Erp44 | 7.89451 | 8.12007 | 10.3975 | 5.37066 | 4.22307 | 4.64281 |
| Tbc1d1 | 2.76354 | 3.49723 | 3.88123 | 2.23845 | 1.75116 | 1.76543 |
| Mfsd4 | 1.539 | 1.25769 | 1.16209 | 0.551884 | 0.597559 | 0.873969 |
| Pltp | 8.24527 | 9.86689 | 13.1948 | 11.8691 | 8.33077 | 4.24118 |
| Gab1 | 5.88691 | 6.01635 | 7.29146 | 3.44182 | 2.36179 | 4.16716 |
| Nfkb1 | 0.550678 | 0.631062 | 0.830754 | 0.394488 | 0.298805 | 0.247695 |
| Slc44a2 | 4.68956 | 6.86165 | 6.55373 | 2.75991 | 3.6755 | 4.02747 |
| Wdr11 | 1.39643 | 1.15295 | 1.64188 | 0.62008 | 0.829024 | 0.834935 |
| Zfyve26 | 1.11817 | 1.19905 | 1.22794 | 0.453886 | 1.01592 | 0.589361 |
| Rnh1 | 33.6359 | 50.2701 | 61.6874 | 30.4996 | 29.4935 | 21.9024 |
| Atp2a2 | 9.11757 | 11.5039 | 9.52253 | 5.71654 | 5.12386 | 5.50199 |
| Sdc3 | 6.54762 | 11.5398 | 11.1524 | 4.65517 | 9.25601 | 4.57763 |
| Zfp592 | 2.44098 | 2.44087 | 3.03677 | 1.36427 | 1.24715 | 1.71366 |
| Tmem108 | 8.03427 | 8.92503 | 2.76532 | 3.00353 | 3.74239 | 4.0518 |
| Cnpy3 | 14.7248 | 10.2518 | 15.1302 | 8.45621 | 7.68213 | 7.53056 |
| Slc6a6 | 3.64601 | 4.05825 | 4.28825 | 2.08858 | 2.39207 | 2.27132 |
| Fkbp15 | 3.32986 | 3.67904 | 3.74487 | 1.96741 | 1.96222 | 1.93702 |
| Gpr107 | 1.35766 | 1.37118 | 1.2388 | 0.836194 | 0.793269 | 0.718522 |
| Igf2bp3 | 2.95951 | 2.41942 | 2.98734 | 1.72286 | 1.60551 | 1.55604 |
| Frat2 | 13.9695 | 6.41861 | 12.1645 | 7.22575 | 4.84575 | 5.75336 |
| Akap10 | 0.991117 | 0.97714 | 1.31134 | 0.482382 | 0.589839 | 0.563736 |
| Plod1 | 2.73468 | 1.10591 | 2.75378 | 1.27661 | 1.9062 | 1.3175 |
| Rab11fip5 | 2.82238 | 2.24716 | 3.03915 | 1.4185 | 1.6945 | 1.64633 |
| Stk40 | 2.92904 | 5.19382 | 4.86956 | 2.58124 | 3.23497 | 1.41263 |
| Nfkbiz | 22.4706 | 27.2457 | 35.7445 | 11.7775 | 23.2567 | 8.22761 |
| Gna13 | 10.223 | 10.5529 | 14.3365 | 6.47958 | 6.73168 | 4.33407 |
| Apaf1 | 1.56611 | 1.14921 | 1.36681 | 0.51125 | 0.563894 | 1.19304 |
| Myo9b | 3.79798 | 3.30747 | 3.99302 | 2.33624 | 2.78409 | 2.05269 |
| Itgb5 | 10.1148 | 8.63929 | 12.2593 | 4.80897 | 7.91361 | 4.53921 |
| Rnf31 | 1.6968 | 1.52186 | 2.38594 | 1.4181 | 1.20343 | 0.897156 |
| Gns | 16.9424 | 18.3477 | 23.0732 | 9.95475 | 14.2852 | 7.59384 |
| Irak4 | 2.97598 | 2.19341 | 3.75416 | 1.82862 | 1.75701 | 0.912247 |
| Map2k3 | 4.85674 | 4.1821 | 6.42053 | 3.26086 | 4.27626 | 3.3847 |
| Plxnb2 | 17.2178 | 13.8675 | 19.5607 | 8.82025 | 9.95573 | 9.5364 |
| Zbtb1 | 1.29562 | 1.08734 | 1.11933 | 0.585403 | 0.478474 | 0.668364 |
| Map1s | 10.549 | 11.3668 | 14.3348 | 8.75297 | 6.02292 | 7.53801 |
| Sipa1l3 | 0.35277 | 0.441207 | 0.338581 | 0.164671 | 0.316664 | 0.235708 |
| Fam49b | 47.448 | 30.8404 | 51.2956 | 22.0405 | 23.7687 | 16.5898 |
| Ehd4 | 7.51977 | 6.2841 | 8.25805 | 3.26572 | 6.06718 | 3.39499 |
| Sh3bp2 | 5.66216 | 4.98773 | 7.2157 | 3.7399 | 3.26828 | 4.62724 |
| Actr3 | 53.7871 | 34.9045 | 59.1843 | 24.9389 | 30.8978 | 20.2735 |
| Psmb8 | 6.32712 | 7.28614 | 9.18826 | 4.17154 | 5.4187 | 1.98875 |
| Derl1 | 27.0108 | 27.3016 | 39.4631 | 11.753 | 20.5326 | 14.0733 |
| Ucp2 | 16.8631 | 11.889 | 20.3133 | 9.49556 | 10.0085 | 7.48921 |
| Rabgef1 | 4.49654 | 4.77349 | 5.86492 | 2.73446 | 3.57677 | 2.16005 |
| Epn1 | 5.38569 | 3.86456 | 6.29657 | 3.401 | 4.70638 | 3.33432 |
| Isyna1 | 30.7349 | 26.2237 | 31.8716 | 13.0062 | 17.7595 | 19.1703 |
| Ythdf1 | 7.63563 | 9.6353 | 9.22001 | 3.98363 | 6.23585 | 3.51593 |
| Qsox1 | 1.53956 | 2.93502 | 2.71346 | 1.40125 | 1.6082 | 1.74357 |
| Cap1 | 20.9077 | 16.6779 | 22.6968 | 11.0094 | 13.1442 | 9.57501 |
| Smad7 | 2.79519 | 2.97264 | 3.77111 | 1.53505 | 2.18616 | 1.64424 |
| Actb | 3406.26 | 2972.73 | 4201.38 | 2337.77 | 2270.2 | 1695.73 |
| Amdhd2 | 2.93103 | 2.86721 | 4.61206 | 2.39094 | 1.62056 | 1.89966 |
| Cyth3 | 2.04344 | 3.08183 | 2.35193 | 1.10074 | 1.22795 | 1.31338 |
| Plekhm3 | 0.536342 | 0.832771 | 0.88957 | 0.41109 | 0.363093 | 0.530333 |
| Cyb561a3 | 11.0345 | 5.20359 | 11.0243 | 4.69311 | 4.2307 | 4.23768 |
| Slc39a11 | 7.13981 | 6.73246 | 8.03595 | 4.9635 | 3.2297 | 4.55867 |
| Nfkbia | 134.421 | 399.177 | 396.622 | 169.809 | 191.29 | 139.319 |
| Atg9a | 3.86269 | 5.01966 | 4.0769 | 1.85853 | 2.7139 | 2.09553 |
| Slc39a14 | 0.402662 | 0.538817 | 0.655962 | 0.239867 | 0.387628 | 0.278551 |
| Tbk1 | 7.05866 | 9.56093 | 9.13872 | 3.7575 | 4.48127 | 4.41483 |
| Pten | 5.74487 | 6.21887 | 5.59936 | 2.73213 | 2.36179 | 3.0789 |
| Ermp1 | 0.80133 | 0.748677 | 1.0865 | 0.667709 | 0.32258 | 0.420909 |
| Nploc4 | 3.85215 | 5.29333 | 6.09039 | 2.47541 | 1.92069 | 3.65767 |
| Rapgef5 | 5.27612 | 2.78784 | 4.89963 | 1.84803 | 3.37466 | 1.41252 |
| Nuak2 | 2.28774 | 3.43905 | 4.03424 | 1.19491 | 2.10078 | 2.03926 |
| Tab3 | 1.02066 | 1.54576 | 1.60758 | 0.748383 | 0.538691 | 0.782239 |
| Pxk | 9.21179 | 11.2221 | 12.8305 | 7.90324 | 5.43056 | 4.37099 |
| Eml3 | 3.21356 | 2.25616 | 3.85526 | 0.939628 | 1.52882 | 1.82395 |
| Spcs3 | 12.8559 | 10.2768 | 14.799 | 4.34617 | 6.44245 | 7.08446 |
| Fam105a | 14.923 | 8.4814 | 14.7344 | 8.5649 | 6.14987 | 4.58014 |
| Pitpnm1 | 5.43312 | 3.71669 | 5.21279 | 1.97595 | 3.10242 | 2.42718 |
| Flii | 5.69072 | 5.50393 | 7.75283 | 3.53566 | 3.78818 | 2.91077 |
| Tagap | 6.48622 | 4.62901 | 6.89471 | 3.12734 | 3.49785 | 3.17639 |
| Rnf19b | 4.93826 | 10.9372 | 11.1614 | 3.7127 | 5.7271 | 5.17178 |
| Slc43a2 | 1.0247 | 1.37568 | 1.38611 | 0.706407 | 0.681994 | 0.706982 |
| Mcl1 | 24.9863 | 44.6929 | 46.49 | 22.3374 | 20.1476 | 12.8234 |
| Tmem110 | 2.08863 | 1.39781 | 2.29305 | 1.12593 | 0.974331 | 0.926366 |
| Ctsa | 14.6784 | 18.3705 | 25.3586 | 13.4578 | 14.5316 | 11.272 |
| Elf1 | 4.10446 | 3.42567 | 4.45858 | 1.72985 | 2.62144 | 1.40623 |
| Nacc1 | 3.81022 | 6.56728 | 6.52915 | 2.67021 | 2.15684 | 4.18117 |
| Etv5 | 5.11434 | 4.69316 | 6.91428 | 3.47517 | 2.09654 | 3.14196 |
| Arf6 | 23.8165 | 20.148 | 30.523 | 13.2452 | 12.1095 | 8.31043 |
| Map3k1 | 6.66964 | 3.98764 | 7.45252 | 3.70083 | 4.0149 | 1.34746 |
| Cyb5r1 | 2.15946 | 2.77485 | 3.68476 | 1.87337 | 1.4437 | 1.5691 |
| Scpep1 | 19.4583 | 30.3726 | 35.0107 | 13.0565 | 16.9049 | 12.1649 |
| Itsn2 | 2.75049 | 3.58438 | 5.19406 | 2.13338 | 2.14632 | 1.23593 |
| Cttnbp2nl | 7.29421 | 5.50822 | 8.36105 | 2.6567 | 2.82843 | 3.61477 |
| Lrrfip1 | 2.94042 | 2.88656 | 3.96514 | 1.81693 | 2.0614 | 1.0993 |
| Slc12a9 | 3.41349 | 2.79375 | 4.70048 | 2.14532 | 2.10145 | 1.43319 |
| Man2b1 | 14.41 | 16.5206 | 20.3879 | 10.8854 | 9.4352 | 5.67723 |
| Sh3bp4 | 3.52426 | 4.63206 | 3.9218 | 1.83058 | 2.34851 | 2.26867 |
| Kpna6 | 1.64684 | 2.89659 | 2.42174 | 1.26268 | 1.14082 | 1.23847 |
| Hhex | 4.47832 | 3.74203 | 6.88003 | 3.56054 | 2.32854 | 2.33935 |
| Gmip | 6.76753 | 3.44757 | 5.90361 | 3.04956 | 3.8144 | 2.08694 |

| | | | | | | |
|---|---|---|---|---|---|---|
| Sstr1 | 0 | 0.0560199 | 0 | 2.37875 | 0.419795 | 0.942569 |
| Frmpd1 | 0 | 0 | 0 | 0.223078 | 0.304539 | 0.00757046 |
| Scara5 | 0 | 0 | 0 | 0.215493 | 0.0131009 | 0.252788 |
| AU021092 | 0 | 0 | 0 | 1.66363 | 0.348314 | 0.645271 |
| Eva1c | 0 | 0.0447882 | 0 | 1.31965 | 0.612773 | 0.777744 |
| E030030I06Rik | 0 | 0 | 0 | 0.219537 | 0.259016 | 0.310637 |
| Tbx18 | 0 | 0.016461 | 0 | 0.736562 | 0.257392 | 0.263257 |
| Spta1 | 0 | 0 | 0 | 0.0369512 | 0.130853 | 0.0500176 |
| Ptpn3 | 0 | 0 | 0 | 0.114434 | 0.453731 | 0.0178054 |
| Esm1 | 0 | 0 | 0 | 0.645006 | 0.050269 | 0.603736 |
| Dmrt3 | 0 | 0.0845557 | 0 | 2.48625 | 1.04653 | 2.80833 |
| Pgm5 | 0 | 0 | 0 | 0.538352 | 0.163561 | 0.152053 |
| Rhpn2 | 0 | 0 | 0 | 0.117613 | 0.0435259 | 0.175362 |
| Clec10a | 0.0264714 | 0 | 0 | 0.25809 | 0.533813 | 0.208884 |
| Erg | 0 | 0 | 0 | 0.241167 | 0.233106 | 0.168097 |
| Zfp473 | 0 | 0 | 0 | 0.129414 | 0.00607424 | 0.0646662 |
| Foxs1 | 0 | 0 | 0 | 2.13055 | 1.75774 | 0.393564 |
| Scd3 | 0 | 0 | 0 | 0.155552 | 0.284772 | 0.0869527 |
| Tmem81 | 0 | 0 | 0 | 0.622202 | 0.559872 | 0.0526944 |
| Cntn6 | 0.00898415 | 0 | 0.00830143 | 0.513843 | 0.426396 | 0.0526063 |
| Tbx15 | 0 | 0 | 0 | 0.110373 | 0.0389308 | 0.484811 |
| Piezo2 | 0.00442657 | 0 | 0 | 0.110227 | 0.0465512 | 0.084076 |
| Otor | 0 | 0 | 0 | 1.05067 | 1.4675 | 0.248792 |
| Gm9930 | 0 | 0 | 0 | 0.341372 | 0.158864 | 0.0786349 |
| Bmp6 | 0 | 0 | 0 | 0.395844 | 0.819955 | 0.187644 |
| Defb11 | 0 | 0 | 0 | 2.88424 | 2.17861 | 8.05285 |
| Crip3 | 0 | 0 | 0 | 0.313402 | 0.336313 | 0.306501 |
| Prdm6 | 0 | 0 | 0 | 0.080733 | 0.452193 | 0.86551 |
| Eda2r | 0 | 0 | 0 | 0.0578855 | 0.0644107 | 0.117562 |
| K230010J24Rik | 0 | 0 | 0 | 0.221172 | 0.288591 | 0.0278542 |
| Adamts12 | 0 | 0 | 0 | 0.0891782 | 0.06319 | 0.221708 |
| Ppara | 0 | 0 | 0 | 0.530157 | 0.0726062 | 0.147782 |
| Hrasls | 0 | 0 | 0 | 0.379146 | 0.146133 | 0.0474916 |
| Col13a1 | 0 | 0 | 0 | 0.279976 | 0.142375 | 0.0393999 |
| Rbp4 | 0.269964 | 0 | 0 | 8.81859 | 2.85629 | 8.82741 |
| Col8a1 | 0 | 0 | 0 | 0.221655 | 0.0750542 | 0.180244 |
| Rhcg | 0 | 0 | 0 | 0.191588 | 0.250955 | 0.873777 |
| Eva1a | 0 | 0.0277005 | 0 | 0.834871 | 0.0468145 | 0.980383 |
| Itih3 | 0 | 0 | 0 | 0.0744817 | 0.0227619 | 0.392927 |
| Sema3d | 0 | 0.00636979 | 0 | 0.105046 | 0.0150741 | 0.141321 |
| Fam69c | 0 | 0 | 0 | 0.366556 | 0.232886 | 0.268586 |
| Ctsk | 0 | 0 | 0 | 0.311959 | 0.580569 | 0.56551 |
| Sox7 | 0 | 0 | 0 | 0.292498 | 0.13892 | 0.253677 |
| Osr1 | 0.0507204 | 0 | 0.0477425 | 1.8648 | 0.655745 | 4.87833 |
| Aspn | 0 | 0 | 0 | 0.276766 | 0.137379 | 0.110044 |
| Cpa4 | 0 | 0 | 0 | 0.574897 | 0.178693 | 0.0518007 |
| Tnnc1 | 0 | 0 | 0 | 0.839631 | 0.956886 | 0 |
| Prkg2 | 0 | 0 | 0 | 0.100339 | 0.119319 | 0.209775 |
| Pkd1l2 | 0 | 0 | 0 | 0.0106584 | 0.148714 | 0.0332419 |
| 1500015O10Rik | 0.21956 | 0 | 0.208815 | 12.0664 | 4.27533 | 9.2995 |
| Dkkl1 | 0 | 0 | 0 | 2.42575 | 0.959786 | 0.424548 |
| Eya4 | 0 | 0 | 0 | 0.171605 | 0.014325 | 0.164167 |
| Maob | 0 | 0 | 0 | 0.39464 | 0.843581 | 0.0651843 |
| Kcnj8 | 0.0330512 | 0 | 0.0316191 | 2.12951 | 1.02741 | 1.31502 |
| Kank4 | 0 | 0 | 0 | 0.109269 | 0.272744 | 0.0712688 |
| Gper1 | 0 | 0 | 0 | 0.789941 | 0.0827928 | 1.10567 |
| Dcn | 0.032803 | 0.318935 | 0.3412 | 16.9825 | 7.3892 | 11.4074 |
| Vwf | 0 | 0 | 0 | 0.0436267 | 0.0129174 | 0.011051 |
| Gbx1 | 0 | 0 | 0 | 0.211524 | 0.191618 | 0.586241 |
| Gdf10 | 0.0341993 | 0 | 0 | 0.706207 | 1.60228 | 0.148669 |
| Sardh | 0 | 0 | 0 | 0.0303529 | 0.0489072 | 0.11455 |
| Kazald1 | 0 | 0 | 0 | 0.893446 | 0.696352 | 0.56733 |
| Scarf1 | 0 | 0 | 0 | 0.334337 | 0.200748 | 0.420205 |
| Cfap206 | 0 | 0 | 0 | 0.183756 | 0.161758 | 0.255553 |
| Tex15 | 0.00535182 | 0 | 0 | 0.31108 | 0.3524 | 0.418075 |
| Wdr72 | 0 | 0 | 0 | 0 | 0.00827173 | 0 |
| Pcdhgb2 | 0.0212113 | 5.28E-17 | 0.0206068 | 0.252216 | 0.263337 | 0.529206 |
| Hbq1a | 0 | 0 | 0 | 1.44158 | 1.66362 | 1.88816 |
| Adnp | 6.06E-07 | 5.25E-07 | 4.87E-07 | 0.16271 | 0.122352 | 0.226764 |
| Slco1c1 | 0 | 0 | 0 | 0.265404 | 0.084843 | 0.0364633 |
| Il17d | 0.111729 | 0 | 0.107176 | 7.79656 | 3.65875 | 1.0654 |
| Myl9 | 0.145501 | 0.323177 | 0.21101 | 14.808 | 10.2912 | 24.84 |
| Bmp2 | 0 | 0 | 0.0445384 | 0.76399 | 1.26779 | 0.358771 |
| Klhl4 | 0 | 0 | 0 | 0.340267 | 0.135256 | 0.169753 |
| Alpl | 0.0198239 | 0 | 0.0376777 | 0.684071 | 0.48501 | 1.21141 |
| Ecel1 | 0.161274 | 0 | 0.0766795 | 6.17609 | 1.71358 | 6.43524 |
| Ndp | 0 | 0.122481 | 0 | 1.89651 | 1.0256 | 2.01084 |
| Angpt1 | 0.07639 | 0.0800009 | 0.12751 | 3.46108 | 4.00015 | 4.08608 |
| Serpinf1 | 0.181109 | 0.119153 | 0 | 5.01962 | 4.37481 | 3.45819 |
| Plekha7 | 0 | 0 | 0.0160487 | 0.262541 | 0.338862 | 0.468607 |
| Col6a1 | 0 | 0.0178457 | 0.0155641 | 0.356501 | 0.798027 | 0.836592 |
| Col1a2 | 0.0326788 | 0.0579188 | 0.035502 | 1.24139 | 1.04731 | 1.40878 |
| Col1a1 | 0.233159 | 0.100731 | 0.0443909 | 5.0982 | 4.63499 | 5.61554 |
| Tfap2b | 0 | 0 | 0.0122529 | 0.39455 | 0.219037 | 0.340658 |
| Rasd1 | 0 | 0.073833 | 0 | 2.25034 | 0.671454 | 2.13706 |
| Col15a1 | 0.0126222 | 0 | 0.0119963 | 0.204661 | 0.219273 | 0.21032 |
| Plxdc1 | 0.0162163 | 0 | 0 | 0.202518 | 0.588364 | 0.121438 |
| Pcdh11x | 0.0298526 | 0 | 0.0168837 | 0.324814 | 0.694695 | 0.672756 |
| Chodl | 0 | 0 | 0 | 0.263323 | 0.212726 | 0.709232 |
| Rgs5 | 0.142036 | 0.176529 | 0.0593909 | 5.11101 | 3.6486 | 2.29786 |
| Frem1 | 0.00829457 | 0.00883668 | 0 | 0.163623 | 0.158322 | 0.096486 |
| Palm2 | 3.06E-06 | 0.0268381 | 0.0699883 | 1.10885 | 1.03974 | 0.592036 |
| Cbln1 | 0.0966584 | 0 | 0.0920985 | 1.6945 | 3.17849 | 1.64219 |
| Plcd4 | 5.48E-12 | 2.08E-09 | 6.61E-09 | 0.199189 | 0.125214 | 0.133229 |
| Drc7 | 0.0543538 | 0 | 0 | 0.364879 | 0.317457 | 0.53047 |
| Egflam | 0.0108355 | 0 | 0.0101741 | 0.215353 | 0.0719194 | 0.203158 |
| Vwa5b1 | 0 | 0 | 0 | 0.075274 | 0.123439 | 0.149577 |
| Gm17455 | 0.120781 | 0 | 0.234576 | 3.46981 | 2.99658 | 0.265786 |
| Prr32 | 0.193 | 0 | 0 | 1.48632 | 1.60817 | 2.20665 |
| Col25a1 | 0.0196717 | 0 | 0.00912225 | 0.38392 | 0.103828 | 0.21381 |
| Zfp964 | 0.0214734 | 0 | 0 | 0.155377 | 0.253651 | 0.0418031 |
| Higd1b | 0.434107 | 0.11961 | 0 | 6.12129 | 3.13498 | 1.96012 |
| 1700013F07Rik | 0 | 0 | 0 | 0.773134 | 0.497412 | 0.408119 |
| Mterf1b | 0 | 0 | 0 | 0.602438 | 2.26054 | 1.72239 |
| Apod | 0.0348154 | 0.0378705 | 0.0333172 | 1.14855 | 1.87615 | 1.4542 |
| Mpo | 0.125337 | 0.0818853 | 0.0238708 | 5.49478 | 0.740925 | 2.16038 |

| | | | | | | |
|---|---|---|---|---|---|---|
| Apbb1ip | 3.35979 | 2.18304 | 3.84399 | 1.27179 | 1.85483 | 0.980659 |
| Pdcd6ip | 8.23593 | 6.26061 | 9.20276 | 3.93109 | 4.36259 | 3.32784 |
| Cnot11 | 6.76806 | 5.27589 | 7.59636 | 3.91628 | 1.80364 | 4.54288 |
| Stk11ip | 1.24094 | 1.15884 | 1.83108 | 0.806978 | 0.708657 | 0.875945 |
| Aldoa | 46.376 | 56.4835 | 72.0334 | 37.0776 | 38.5965 | 25.5566 |
| Smpdl3a | 7.47855 | 16.9792 | 19.1596 | 7.15386 | 7.92937 | 6.47312 |
| Mtmr3 | 3.98244 | 4.45476 | 4.86974 | 2.11266 | 2.36544 | 2.6385 |
| Ogfrl1 | 1.77079 | 1.89985 | 2.17802 | 1.04551 | 1.03871 | 0.738749 |
| Zc3h3 | 1.44725 | 1.28351 | 1.77304 | 0.759389 | 0.759386 | 1.09931 |
| Mogs | 4.12933 | 4.60665 | 6.3556 | 4.09078 | 2.58187 | 2.05311 |
| Oma1 | 8.20219 | 7.39837 | 10.3474 | 3.91499 | 5.16476 | 3.87349 |
| Med23 | 2.03058 | 3.72428 | 3.23759 | 1.54072 | 1.64612 | 1.04166 |
| Psen1 | 10.6719 | 13.2262 | 14.9791 | 7.43453 | 7.71469 | 3.74234 |
| Cd302 | 45.5824 | 29.9226 | 58.3497 | 14.6196 | 36.223 | 14.8722 |
| Ccdc93 | 0.693227 | 0.794049 | 0.711546 | 0.238845 | 0.31663 | 0.624797 |
| Sp140 | 1.91722 | 4.07369 | 3.96606 | 1.30999 | 1.94242 | 1.33725 |
| Tmem184b | 4.04561 | 4.74459 | 6.26693 | 2.63682 | 3.29644 | 2.47657 |
| Slc39a13 | 1.82788 | 1.40514 | 1.87079 | 0.626088 | 1.1332 | 1.10715 |
| Dhrs3 | 22.0122 | 18.7592 | 29.4416 | 12.7205 | 14.4628 | 8.11065 |
| Pign | 0.46199 | 0.408151 | 0.453958 | 0.214236 | 0.348522 | 0.204322 |
| Eml2 | 2.49565 | 4.09906 | 4.31144 | 2.25489 | 1.59001 | 2.55063 |
| Gpr137b | 4.15581 | 5.71776 | 5.92945 | 2.04813 | 3.25083 | 1.69113 |
| Mfsd1 | 14.942 | 19.6754 | 25.6483 | 10.4362 | 10.7361 | 8.03778 |
| Ehd1 | 34.0634 | 31.7421 | 45.1548 | 20.3194 | 27.9991 | 12.7319 |
| Alas1 | 4.68221 | 4.6689 | 5.94128 | 2.88356 | 2.54981 | 2.03782 |
| Pkd1 | 1.45363 | 0.698977 | 1.2516 | 0.633493 | 0.53955 | 1.10576 |
| Socs3 | 27.8475 | 44.0208 | 52.5543 | 21.7064 | 26.7522 | 17.7817 |
| Dab2 | 2.33345 | 6.40069 | 6.33828 | 1.6036 | 3.05552 | 1.8908 |
| Mafb | 19.6887 | 12.6496 | 19.5431 | 10.8084 | 5.52039 | 9.15926 |
| 9130011E15Rik | 3.24157 | 5.26029 | 4.67465 | 1.03755 | 2.17177 | 3.4046 |
| Wdfy3 | 5.29442 | 2.98657 | 5.11 | 2.11782 | 2.58026 | 1.94259 |
| Tbc1d2b | 0.688332 | 0.461636 | 0.661706 | 0.251209 | 0.394075 | 0.348125 |
| Slc31a1 | 5.15868 | 11.501 | 11.7017 | 4.22596 | 4.74669 | 4.30659 |
| Ctsl | 55.5412 | 111.442 | 123.456 | 46.1349 | 48.5108 | 38.2658 |
| Bnip3 | 15.6916 | 29.1196 | 29.6831 | 11.0231 | 13.027 | 11.1403 |
| Mast3 | 4.33137 | 2.70053 | 3.65382 | 1.57345 | 2.2709 | 1.69888 |
| Sgpp1 | 5.85531 | 3.58409 | 5.09564 | 2.32583 | 2.28943 | 1.71088 |
| Slc44a1 | 2.04279 | 2.8253 | 2.64938 | 0.758568 | 1.54312 | 1.29731 |
| Cd68 | 37.919 | 74.1369 | 82.7081 | 31.5844 | 47.485 | 21.8019 |
| Fam171a2 | 10.3281 | 11.9803 | 9.08173 | 8.8207 | 5.18333 | 5.22734 |
| Adrb2 | 3.84849 | 5.54952 | 5.99061 | 2.22822 | 2.37519 | 2.73213 |
| Nek9 | 4.01876 | 4.35551 | 4.98079 | 1.93305 | 1.44349 | 3.22778 |
| Vmp1 | 16.7236 | 15.6195 | 19.6018 | 7.96117 | 7.03803 | 6.61427 |
| Cd82 | 7.5234 | 6.22224 | 9.36322 | 4.29771 | 5.37653 | 3.59865 |
| Lipa | 14.3306 | 24.0235 | 23.9595 | 9.75916 | 10.1581 | 8.47555 |
| Stab1 | 3.55387 | 2.7775 | 4.40072 | 1.91165 | 2.66479 | 1.06725 |
| E2f8 | 1.84475 | 0.792795 | 1.71749 | 0.832251 | 0.512085 | 0.87779 |
| Nup98 | 5.49308 | 4.97374 | 4.90096 | 2.2719 | 2.4508 | 2.45759 |
| Tmem123 | 11.897 | 11.158 | 15.7876 | 5.86107 | 6.12947 | 5.08145 |
| Il10rb | 4.52082 | 6.91123 | 9.79915 | 3.25223 | 4.94306 | 2.27933 |
| Cog3 | 2.70779 | 2.24035 | 2.87848 | 2.25619 | 0.403918 | 1.08133 |
| Angel1 | 1.91602 | 2.2986 | 2.10869 | 0.822384 | 1.09331 | 1.32721 |
| Cdc45 | 9.68677 | 8.09989 | 12.2423 | 7.55231 | 4.13093 | 4.09236 |
| Alox5ap | 22.4809 | 39.8651 | 46.5843 | 25.1488 | 26.5775 | 11.8786 |
| Slc35a3 | 0.557685 | 0.729862 | 0.788896 | 0.299682 | 0.233369 | 0.417614 |
| Mov10 | 1.55808 | 1.48055 | 1.82249 | 0.510482 | 1.05013 | 0.608352 |
| Eno1b | 90.9714 | 132.058 | 153.995 | 60.9706 | 66.3914 | 60.9953 |
| Lmbrd1 | 2.28028 | 2.49696 | 2.73006 | 1.66161 | 0.672947 | 1.48206 |
| Zbtb34 | 0.775633 | 0.882211 | 1.31847 | 0.477933 | 0.344085 | 0.468249 |
| Cnr1 | 23.014 | 17.0391 | 10.3966 | 5.93137 | 6.96024 | 9.55743 |
| Ano8 | 5.63111 | 3.82576 | 4.30681 | 2.20852 | 2.57681 | 2.42628 |
| Paqr7 | 1.97026 | 3.05982 | 2.85822 | 1.91609 | 0.804592 | 1.2181 |
| Gatm | 127.883 | 105.141 | 160.922 | 70.0781 | 70.8733 | 38.2247 |
| Cd2ap | 4.94421 | 3.74589 | 5.67496 | 2.0133 | 2.84053 | 1.20552 |
| Fxr2 | 4.23021 | 5.08844 | 4.14732 | 3.06989 | 2.28247 | 2.0017 |
| Susd6 | 3.63949 | 3.64533 | 4.96664 | 1.84329 | 2.57637 | 1.73654 |
| Cln5 | 4.14245 | 12.1303 | 11.5472 | 4.0424 | 5.2432 | 3.23758 |
| Pbxip1 | 5.25095 | 4.94858 | 7.27572 | 4.18076 | 2.38482 | 2.3456 |
| Flna | 8.40643 | 8.25907 | 10.9588 | 4.05324 | 5.43058 | 4.23515 |
| Hivep2 | 3.21863 | 3.42343 | 2.20586 | 0.868977 | 1.29325 | 2.02709 |
| Pim2 | 10.6775 | 5.65969 | 10.3972 | 4.23564 | 5.25921 | 4.82171 |
| S1pr2 | 7.09093 | 5.38072 | 8.44649 | 4.39662 | 3.62252 | 2.45204 |
| Arid3b | 1.54556 | 1.75588 | 1.69958 | 0.902623 | 0.867779 | 0.699508 |
| Tbc1d13 | 6.92445 | 5.37035 | 7.50816 | 3.24397 | 4.23013 | 3.80112 |
| Fam129b | 1.00235 | 2.40219 | 1.48434 | 1.18113 | 0.702932 | 1.09002 |
| Tmem170b | 11.8548 | 11.0496 | 12.8549 | 3.97708 | 4.68998 | 6.40483 |
| Tspo | 75.4813 | 165.704 | 189.416 | 70.4936 | 100.537 | 44.9603 |
| Mdfic | 0.418082 | 1.11893 | 1.08395 | 0.252016 | 0.516464 | 0.528016 |
| Csgalnact2 | 1.98142 | 1.25463 | 1.8455 | 0.984194 | 0.700285 | 0.489915 |
| Tgfbrap1 | 1.39967 | 1.70273 | 1.38047 | 0.667129 | 0.797024 | 0.623768 |
| Clec4a3 | 72.4264 | 58.2425 | 100.881 | 38.7841 | 46.5559 | 16.3629 |
| Rnf103 | 2.93546 | 3.24146 | 2.67237 | 2.09179 | 0.847448 | 0.971573 |
| Plscr1 | 9.2019 | 13.4107 | 15.443 | 4.28728 | 8.68683 | 3.63254 |
| B4galt1 | 18.6331 | 20.2405 | 30.4392 | 13.2036 | 12.8731 | 6.87311 |
| Rgl1 | 2.36301 | 4.15647 | 4.4985 | 1.04028 | 2.62073 | 1.20473 |
| Sipa1l2 | 3.39082 | 2.62943 | 3.11903 | 1.47017 | 1.88892 | 1.37134 |
| Lamp1 | 150.071 | 203.682 | 240.47 | 106.944 | 93.3655 | 77.221 |
| Slc12a7 | 2.76863 | 1.58145 | 3.29436 | 1.78482 | 0.861958 | 0.89845 |
| Dnm2 | 10.4519 | 8.81037 | 11.9714 | 5.06054 | 5.99905 | 4.91412 |
| Shisa5 | 8.24455 | 15.4682 | 20.1452 | 9.32973 | 12.9637 | 5.43465 |
| Stxbp3 | 1.18215 | 1.10018 | 1.51992 | 0.557544 | 0.601574 | 0.539508 |
| Tiam2 | 13.3524 | 9.05618 | 9.30353 | 5.386 | 4.63152 | 8.74614 |
| Wdfy2 | 8.40764 | 7.03909 | 9.10958 | 2.64585 | 2.75746 | 5.47278 |
| Satb2 | 8.54752 | 6.20764 | 6.27796 | 3.37592 | 1.94886 | 4.72623 |
| Ptpn23 | 3.91077 | 4.2274 | 4.94978 | 1.59624 | 2.6018 | 2.27804 |
| Atxn2l | 13.0996 | 14.9066 | 15.1561 | 5.27888 | 7.19365 | 9.29909 |
| Cc2d1b | 1.86239 | 1.90291 | 1.91298 | 1.41339 | 1.09446 | 0.745971 |
| Sema7a | 4.21296 | 2.50856 | 2.49762 | 0.817729 | 2.34761 | 1.83204 |
| Nr3c1 | 1.31331 | 2.49637 | 2.57355 | 1.1909 | 0.879921 | 0.609284 |
| Tpcn1 | 1.28736 | 1.06201 | 1.30166 | 0.617047 | 0.479925 | 0.670504 |
| Cd37 | 14.8397 | 21.5177 | 27.9909 | 12.6228 | 11.1758 | 6.5312 |
| Csf2ra | 15.9502 | 22.1865 | 26.4823 | 12.633 | 10.163 | 8.70392 |
| Smcr8 | 1.78386 | 2.7147 | 2.64342 | 0.847458 | 1.25087 | 1.17441 |
| Bcl2a1a | 47.6703 | 52.047 | 69.1133 | 26.3638 | 47.5481 | 14.3488 |
| Pde4b | 3.18568 | 4.89152 | 5.52697 | 2.08837 | 2.66721 | 1.36025 |
| Per1 | 4.60263 | 5.85706 | 6.12508 | 1.98421 | 3.69883 | 2.54458 |

| | | | | | | |
|---|---|---|---|---|---|---|
| Stbd1 | 0.0314322 | 0 | 0.0592946 | 0.28682 | 2.02464 | 1.74802 |
| Traf5 | 0.0267548 | 0 | 0 | 0.308992 | 0.209379 | 0.185269 |
| Paqr9 | 0.0436181 | 0.00927594 | 0.00815084 | 0.648228 | 0.471726 | 0.302619 |
| Smoc2 | 0 | 0.0243811 | 0.0424402 | 0.157012 | 0.181766 | 1.17113 |
| Ctla2a | 0.0484832 | 0.104385 | 0 | 0.961847 | 1.04147 | 1.51109 |
| Slco2b1 | 0.0305268 | 0.0434732 | 0.00948971 | 0.329206 | 1.38822 | 0.703927 |
| Spon2 | 0.0267921 | 0 | 0 | 0.080184 | 0.36216 | 0.819126 |
| Ramp2 | 12.1209 | 9.05102 | 12.7869 | 17.9921 | 18.3774 | 15.0935 |
| Cav1 | 0 | 0.107428 | 0.0564868 | 1.02675 | 0.869994 | 1.2587 |
| Folr1 | 0 | 0.13934 | 0.0913744 | 1.85581 | 2.14087 | 2.67984 |
| Col3a1 | 0.107039 | 0.0311095 | 0.0760613 | 1.27462 | 1.69009 | 1.71713 |
| Clic6 | 0.0887411 | 0 | 0.105794 | 1.015 | 1.53168 | 1.92399 |
| Ripply3 | 0 | 0.164162 | 0.0960154 | 1.78831 | 1.32584 | 3.14273 |
| Clcf1 | 0 | 0.0686529 | 0.0401314 | 0.78431 | 0.43607 | 1.10233 |
| Lefty2 | 0.0543969 | 0 | 0.0510936 | 0.434896 | 1.25527 | 0.513874 |
| Dnaic1 | 0 | 0.0274029 | 0 | 0.834727 | 0.243286 | 0.337951 |
| Fhl4 | 0.0199532 | 0.0210311 | 0.0366065 | 0.800845 | 0.295871 | 0.341425 |
| Bfsp1 | 0 | 0 | 0.0773528 | 1.18691 | 0.325342 | 0.552006 |
| 4930452B06Rik | 0 | 0.159126 | 0.139099 | 2.65081 | 2.10013 | 1.09429 |
| Grik1 | 0 | 0 | 0 | 1.19844 | 0.109811 | 0.560904 |
| Itih5 | 0.0316874 | 0 | 0.00741079 | 0.359391 | 0.511634 | 0.117063 |
| Tchh | 0 | 0.0540895 | 0.0474719 | 0.76092 | 1.0098 | 0.228118 |
| Ajap1 | 0.065639 | 0 | 0.0924786 | 2.23397 | 0.686709 | 0.747735 |
| Ly6a | 0.054789 | 0.291329 | 0.153095 | 0.999617 | 2.79809 | 5.39973 |
| Egfl7 | 0.0419042 | 0.0224875 | 0 | 1.10038 | 0.435609 | 0.66385 |
| Xlr3a | 0.0463719 | 0.0330687 | 0.0434219 | 0.332267 | 0.230312 | 0.384522 |
| Crhr1 | 0.0340271 | 0 | 0 | 1.54536 | 1.59319 | 0.239565 |
| A730017C20Rik | 0.207044 | 0.107297 | 0.112875 | 2.21592 | 4.45991 | 2.60726 |
| Avp | 1.69341 | 0.743814 | 0.649949 | 16.563 | 3.38333 | 35.1966 |
| Rbm20 | 0 | 0.0115916 | 0 | 0.580171 | 0.13126 | 0.900706 |
| Ogn | 0.023919 | 0.20402 | 0.155734 | 0.940148 | 0.95274 | 3.1973 |
| Ccl27b | 0.602669 | 0 | 0 | 13.8136 | 8.3131 | 0.640032 |
| S1pr3 | 0.0737642 | 0.266814 | 0.0961482 | 3.12349 | 2.3962 | 1.79944 |
| Pxmp2 | 0.355723 | 0.193356 | 0.0850599 | 1.67424 | 3.19127 | 1.04137 |
| Nts | 0 | 0.185969 | 0.435093 | 3.60817 | 0.951629 | 4.63141 |
| Serpind1 | 0.0597878 | 0 | 0.0563667 | 0.605085 | 1.15822 | 0.233006 |
| Slc39a12 | 0 | 0 | 0.05517 | 0.701351 | 0.17574 | 0.188069 |
| Grm8 | 0 | 0.0354346 | 0.0186087 | 0.187735 | 0.327689 | 0.175693 |
| Slc4a1 | 0 | 0 | 0.0268084 | 0.455072 | 0.108144 | 0.310945 |
| Slc22a6 | 0.0507303 | 0 | 0.0793913 | 0.792988 | 0.483771 | 0.619861 |
| Dlk1 | 0 | 0.031197 | 0.0685554 | 0.57942 | 0.397254 | 0.683374 |
| Igf2 | 0.14097 | 0.100823 | 0.102978 | 2.37802 | 2.59115 | 3.25372 |
| Cobll1 | 0 | 0 | 0 | 0.0827528 | 0.177384 | 0.0322472 |
| Sema3e | 0 | 0.0184166 | 0.00536661 | 0.307818 | 0.0149125 | 0.225102 |
| Mtfp1 | 0.180812 | 0 | 0.127383 | 2.97113 | 0.973402 | 1.09059 |
| Rhoj | 0 | 0.0635185 | 0.0461506 | 0.622959 | 0.590922 | 0.384374 |
| Flnc | 0.0662912 | 0.00901502 | 0.0627376 | 0.805004 | 0.385634 | 0.739992 |
| Id4 | 1.96051 | 2.00949 | 3.26953 | 14.0654 | 52.2855 | 16.9494 |
| Ttr | 45.953 | 44.64 | 49.5643 | 667.658 | 594.203 | 712.468 |
| Col26a1 | 0 | 0.0439402 | 0 | 0.493787 | 0.37533 | 0.348707 |
| Itga11 | 0 | 0 | 0 | 0.573051 | 0.361995 | 0.394712 |
| Plce1 | 0.0506828 | 0.0249696 | 0.032713 | 0.652509 | 0.439893 | 0.687957 |
| Cdkl4 | 0 | 0 | 0 | 1.54678 | 0.511934 | 0.489988 |
| Col4a1 | 0.10038 | 0.131942 | 0.167627 | 2.85245 | 1.6496 | 1.20864 |
| Kndc1 | 0.0809963 | 0.0332739 | 0.0192964 | 0.389781 | 0.684363 | 0.76644 |
| Postn | 0.0130881 | 0.00691109 | 0.0240795 | 0.137079 | 0.0406508 | 0.181485 |
| Ntng1 | 0.222669 | 0.0337791 | 0.169599 | 1.07059 | 2.95772 | 0.738686 |
| Fkbp11 | 0 | 0.301264 | 0.789526 | 2.52073 | 7.20275 | 6.35117 |
| Gm10800 | 0.219009 | 0.230999 | 0.410239 | 9.5057 | 1.62885 | 1.39708 |
| Ptrf | 0.185304 | 0.0347019 | 0.244205 | 2.96998 | 1.34379 | 2.38681 |
| Prrg1 | 0.0222009 | 0 | 0.0402681 | 0.189029 | 0.334357 | 0.130545 |
| Cbs | 0.139866 | 0.0764372 | 0.0222087 | 0.569897 | 0.910449 | 0.666307 |
| Atp1a2 | 0.242161 | 0.274099 | 0.153341 | 2.35335 | 2.64924 | 3.10819 |
| Crabp2 | 0.218723 | 0.481795 | 0.701979 | 3.22444 | 4.82818 | 11.3046 |
| Rspo2 | 0.0452688 | 0.292321 | 0.447158 | 2.68547 | 4.55475 | 2.18275 |
| Sst | 11.3911 | 10.7513 | 7.6823 | 260.617 | 51.1103 | 89.9334 |
| Sostdc1 | 0.115182 | 0.563743 | 0.552344 | 4.185 | 4.19007 | 6.60879 |
| Nbl1 | 0.133712 | 0.826553 | 0.597927 | 10.3348 | 7.06799 | 4.24999 |
| Foxf2 | 0.140765 | 0.202184 | 0.0444272 | 1.79 | 1.4642 | 1.28325 |
| Rbp1 | 1.98076 | 2.32534 | 2.32268 | 29.9082 | 19.9562 | 26.883 |
| Galntl6 | 0.0711512 | 0 | 0.0650215 | 0.190404 | 0.477648 | 0.699701 |
| Unc13c | 0.0417224 | 0.0527167 | 0.0327175 | 0.73083 | 0.222064 | 0.328501 |
| Gstm2 | 0.040202 | 0.130668 | 0.190072 | 0.962967 | 0.440345 | 1.9679 |
| Dach2 | 0 | 0.0432944 | 0.0189202 | 0.190571 | 0.258097 | 0.0994299 |
| Erbb2 | 0.05507 | 0.0892765 | 0.0259348 | 0.263401 | 0.565972 | 0.893525 |
| Cntn5 | 0.0154963 | 0.0674193 | 0.0443314 | 0.97706 | 0.579886 | 0.509928 |
| Fign | 0.1643 | 0.136922 | 0.12 | 1.8023 | 1.70391 | 0.66062 |
| Pnoc | 0.0769631 | 0.126155 | 0.293463 | 1.63663 | 1.19526 | 4.64912 |
| Tekt1 | 0.334027 | 0.211892 | 0 | 3.45972 | 0.774756 | 2.36353 |
| Dkk3 | 0.747007 | 0.160852 | 0.140393 | 4.7402 | 4.06195 | 3.58047 |
| Fxyd1 | 0 | 0 | 0 | 0.0194588 | 0 | 0 |
| Angptl2 | 0.0457654 | 0.0393198 | 0 | 0.334884 | 0.323381 | 0.678933 |
| Rln1 | 0.0207511 | 0.0220741 | 0.0194193 | 0.346473 | 0.4509 | 0.183021 |
| Dact3 | 0.197834 | 0.215462 | 0.047427 | 3.30525 | 2.05814 | 1.41975 |
| Itm2a | 2.80938 | 1.84552 | 1.667 | 24.9091 | 16.1737 | 19.9011 |
| Ednrb | 0.93394 | 0.704036 | 1.37213 | 8.81277 | 8.10968 | 12.7809 |
| Dnah7a | 0.00489459 | 0.00532589 | 0.0185233 | 0.0568616 | 0.168332 | 0.17877 |
| Clvs2 | 0.0167701 | 0.158432 | 0.0770465 | 0.851116 | 0.714708 | 0.515053 |
| Gabrg1 | 0.0258478 | 0 | 0.0174537 | 0.143919 | 0.144201 | 0.214481 |
| Aebp1 | 0.180951 | 0.0796564 | 0.155222 | 1.31451 | 0.620494 | 2.16366 |
| Dclk3 | 0.0260613 | 0.115819 | 0.101155 | 0.311106 | 1.97306 | 0.531121 |
| Rfx2 | 0 | 0 | 0.0206954 | 0.177683 | 0.910173 | 0.840197 |
| Gm11627 | 0.158183 | 0.0866462 | 0.230068 | 1.96507 | 1.3434 | 1.56774 |
| Tac2 | 1.05707 | 1.17443 | 0.389104 | 5.20704 | 2.44352 | 25.9647 |
| Zfp850 | 0.0616107 | 0 | 0.0282189 | 0.205572 | 0.134859 | 0.425384 |
| Me3 | 0 | 0.00794161 | 0.0138459 | 0.126731 | 0.00409278 | 0.0500748 |
| Fam19a1 | 0.0261187 | 0.28719 | 0.167229 | 1.19155 | 1.80441 | 0.898442 |
| Grid2 | 0.0145458 | 0.00382297 | 0.00668238 | 0.057344 | 0.091264 | 0.0804384 |
| mt-Cytb | 130.253 | 151.343 | 150.36 | 1715.88 | 1103.05 | 894.116 |
| Car2 | 0.59093 | 0.385823 | 0.645974 | 6.41548 | 5.68723 | 2.7447 |
| Gjd2 | 0.473583 | 0 | 0.29448 | 3.89029 | 1.51259 | 1.2661 |
| Fkbp9 | 0.345478 | 0.790011 | 0.122112 | 1.40807 | 4.63933 | 4.68614 |
| Adgra1 | 0.0311569 | 0.033238 | 0.0383106 | 0.302066 | 0.585835 | 0.528531 |
| AW551984 | 0.0631772 | 0.190703 | 0.108506 | 0.984515 | 1.60348 | 0.879148 |
| AI593442 | 0.80629 | 0.848224 | 0.715005 | 9.50937 | 6.72355 | 3.06922 |
| Vstm5 | 0.110284 | 0.238553 | 0.069449 | 2.83237 | 0.599523 | 0.899281 |

| | | | | | | |
|---|---|---|---|---|---|---|
| Fbxl5 | 3.29508 | 5.16813 | 5.49493 | 2.45641 | 2.03747 | 2.06809 |
| Anxa5 | 23.0118 | 49.9618 | 55.9007 | 19.5943 | 23.9956 | 17.2052 |
| Lamp2 | 8.39785 | 11.5761 | 13.5219 | 4.64721 | 5.10632 | 4.10514 |
| Lcp2 | 7.98615 | 5.91191 | 9.65286 | 3.84584 | 5.24417 | 1.94642 |
| Siah2 | 8.66954 | 4.43478 | 8.65245 | 3.24924 | 4.68695 | 2.78906 |
| Rragc | 10.2174 | 14.3872 | 16.7045 | 8.0104 | 7.3281 | 3.49176 |
| Actn4 | 5.25809 | 7.25993 | 7.39722 | 4.27393 | 3.2232 | 3.23477 |
| Lrp5 | 1.03025 | 0.862037 | 1.00963 | 0.208047 | 0.354432 | 0.74146 |
| Acp2 | 6.57888 | 6.98051 | 8.24469 | 3.05014 | 4.96902 | 3.38439 |
| Tmed3 | 23.9223 | 13.5922 | 28.572 | 10.8488 | 8.06736 | 11.274 |
| Cndp2 | 37.0249 | 70.7364 | 73.4457 | 27.7031 | 35.7136 | 25.9439 |
| Ptprn2 | 0.743147 | 0.456134 | 0.476196 | 0.190628 | 0.319667 | 0.307195 |
| Snap23 | 4.04896 | 6.81082 | 7.3593 | 2.49397 | 2.6918 | 2.7708 |
| Anxa7 | 2.66819 | 7.3703 | 5.6143 | 3.34176 | 1.83805 | 1.65584 |
| Coro7 | 3.07331 | 2.94787 | 3.39947 | 1.65304 | 1.61472 | 1.26771 |
| Prkcd | 3.95634 | 6.02648 | 6.13264 | 3.0252 | 3.6457 | 2.18497 |
| Neu1 | 7.03068 | 9.951 | 9.97132 | 3.94186 | 4.88374 | 4.17375 |
| Pitpna | 16.2748 | 18.2747 | 21.5057 | 9.07674 | 11.1805 | 7.1788 |
| Ntng2 | 5.00874 | 4.24622 | 4.18844 | 1.73854 | 2.60881 | 2.56078 |
| Vcl | 4.50302 | 2.83854 | 5.3177 | 2.04299 | 1.72047 | 1.43711 |
| Prickle1 | 5.23169 | 3.69269 | 5.57585 | 2.01839 | 2.7081 | 2.3716 |
| Map4k2 | 0.657258 | 0.382341 | 0.589606 | 0.436458 | 0.433551 | 0.444989 |
| Dnajc14 | 9.95869 | 9.58275 | 11.2356 | 4.43824 | 5.45382 | 5.79156 |
| Ikbkb | 1.15519 | 0.972598 | 0.99864 | 0.593112 | 0.408969 | 0.603454 |
| Rbm47 | 2.18105 | 1.45964 | 2.32398 | 1.13497 | 1.02536 | 0.656534 |
| Dnaaf5 | 1.94223 | 2.28793 | 2.19079 | 0.552359 | 1.77456 | 0.725083 |
| Coro1a | 53.4587 | 49.3473 | 72.6628 | 30.4584 | 36.8996 | 20.4104 |
| Tnfaip3 | 4.13319 | 10.387 | 9.94156 | 3.18873 | 5.76515 | 1.99729 |
| Gpr155 | 0.704992 | 0.663116 | 0.668736 | 0.123692 | 0.22776 | 0.397828 |
| Inppl1 | 3.52796 | 4.7784 | 5.53861 | 2.24918 | 2.04769 | 2.38118 |
| Olfml3 | 8.57699 | 3.56302 | 7.11986 | 3.62479 | 3.31884 | 2.06946 |
| Arhgap17 | 3.16017 | 3.73762 | 4.33002 | 1.39271 | 2.09512 | 1.55118 |
| Pip4k2a | 13.4029 | 7.14329 | 13.4948 | 5.86538 | 5.91614 | 3.33722 |
| Plcb3 | 2.08632 | 2.99365 | 3.47657 | 1.10197 | 2.41962 | 1.13804 |
| Rab20 | 13.3103 | 31.306 | 34.349 | 16.2918 | 14.4525 | 6.60604 |
| Capza1 | 14.8759 | 15.5589 | 21.0406 | 7.79803 | 7.20128 | 5.66491 |
| Cnn2 | 65.7454 | 26.7252 | 66.6623 | 29.8804 | 27.9003 | 17.9339 |
| Pcdhb19 | 1.72146 | 1.42247 | 0.887772 | 0.570919 | 0.318569 | 0.78644 |
| Mical2 | 0.465347 | 0.656831 | 0.572169 | 0.201405 | 0.277219 | 0.324486 |
| Klf3 | 1.63833 | 2.41214 | 2.67408 | 1.14351 | 0.988375 | 1.04959 |
| Ostf1 | 10.6102 | 8.53584 | 13.8353 | 4.73595 | 6.31395 | 3.1973 |
| Rassf4 | 4.07807 | 2.84034 | 4.83522 | 2.12932 | 2.53383 | 1.39488 |
| Tbc1d10b | 3.74756 | 4.67992 | 5.1372 | 3.15384 | 1.90055 | 1.86487 |
| Vasp | 9.67912 | 8.60064 | 12.7161 | 4.87644 | 7.17964 | 4.06585 |
| Ccrl2 | 33.2721 | 51.2359 | 61.5189 | 19.1833 | 33.4254 | 10.0258 |
| Ccne1 | 5.7127 | 2.20536 | 6.01103 | 2.68485 | 2.12834 | 1.73871 |
| Uap1l1 | 2.41561 | 3.73148 | 4.8491 | 2.04942 | 1.81984 | 1.12266 |
| Optn | 3.07644 | 2.34411 | 3.57338 | 1.29702 | 2.12045 | 0.990756 |
| Sirpa | 14.6002 | 16.5553 | 19.4674 | 10.2648 | 10.5085 | 4.27468 |
| Crlf3 | 3.20449 | 3.28462 | 4.52544 | 1.91143 | 1.00561 | 1.53324 |
| Adam15 | 3.29317 | 2.9607 | 4.08095 | 2.51822 | 1.73055 | 1.4405 |
| Ripk2 | 3.5409 | 5.35119 | 5.72237 | 2.27491 | 2.27135 | 0.974508 |
| Fermt3 | 56.3002 | 42.6015 | 71.5175 | 28.5771 | 44.7299 | 12.387 |
| Rin3 | 0.417511 | 0.400076 | 0.807854 | 0.219323 | 0.406255 | 0.239841 |
| Tm9sf3 | 13.8087 | 12.5102 | 14.734 | 5.25954 | 5.76182 | 5.26108 |
| Wnt7b | 23.0401 | 16.8632 | 13.8546 | 8.00099 | 7.31617 | 11.3803 |
| Cfap43 | 0.836006 | 0.895268 | 0.901894 | 0.426919 | 0.290952 | 0.588613 |
| Trappc10 | 3.84792 | 2.63408 | 3.47441 | 1.27837 | 1.72631 | 1.39545 |
| Nadk | 43.9472 | 34.8957 | 53.8692 | 21.8205 | 21.5875 | 18.3273 |
| Dock10 | 0.885674 | 0.547263 | 1.06626 | 0.345407 | 0.350519 | 0.267694 |
| P2ry6 | 14.2148 | 15.3256 | 21.6496 | 8.31992 | 13.4174 | 3.54331 |
| Plagl2 | 5.875 | 4.10905 | 6.94359 | 1.8172 | 3.12844 | 2.91668 |
| Gopc | 2.29816 | 1.70031 | 2.08464 | 0.775528 | 0.836364 | 1.06527 |
| Tpd52 | 10.2223 | 15.9738 | 18.9733 | 6.57301 | 7.71904 | 3.7286 |
| 2310057M21Rik | 0.286843 | 0.397428 | 0.299785 | 0.133026 | 0.115063 | 0.0960956 |
| Lrp10 | 9.17723 | 14.4555 | 13.9175 | 6.99641 | 4.95517 | 5.98872 |
| Sik1 | 6.62886 | 4.6976 | 7.10497 | 2.89721 | 2.40647 | 3.6633 |
| Stat3 | 3.16716 | 3.42473 | 4.58185 | 1.46041 | 2.41432 | 2.15115 |
| Blvrb | 21.0665 | 8.45527 | 22.0447 | 7.35174 | 11.6791 | 4.47444 |
| Siae | 1.72773 | 1.31123 | 1.63599 | 0.819865 | 0.55753 | 0.608642 |
| N4bp1 | 3.51108 | 8.09194 | 8.08307 | 2.86364 | 3.59647 | 2.24427 |
| Rps6ka4 | 12.0815 | 7.1148 | 13.0012 | 5.76282 | 5.4845 | 4.39851 |
| Slitrk5 | 3.79975 | 3.25829 | 1.90896 | 1.27941 | 1.23508 | 1.35136 |
| Bcl2l11 | 1.56073 | 1.34018 | 2.2863 | 0.762944 | 0.976036 | 0.500821 |
| Pom121 | 5.66362 | 7.55517 | 6.44792 | 4.07192 | 2.95542 | 3.37477 |
| Ptpre | 2.48243 | 2.41394 | 3.66179 | 1.25966 | 1.84304 | 1.04854 |
| Slc38a6 | 0.528504 | 0.422477 | 0.538815 | 0.37975 | 0.156045 | 0.330747 |
| Vac14 | 16.0731 | 11.9635 | 18.5172 | 7.80548 | 6.24369 | 7.75362 |
| Rel | 3.98226 | 4.1077 | 5.96153 | 2.05981 | 1.78597 | 1.55817 |
| Gsdmd | 21.617 | 24.8012 | 32.3027 | 12.6065 | 16.7264 | 6.06359 |
| Tnfrsf1a | 10.285 | 9.98222 | 14.3849 | 6.45326 | 6.91368 | 3.235 |
| Zbtb49 | 0.541397 | 0.897747 | 0.531066 | 0.270335 | 0.180894 | 0.294412 |
| Mthfr | 1.59542 | 1.72213 | 1.99091 | 0.392565 | 1.07016 | 1.15875 |
| P2rx4 | 3.3976 | 4.75812 | 5.91885 | 2.29274 | 2.54678 | 1.20938 |
| Plekha2 | 3.53636 | 2.08959 | 3.9349 | 1.26934 | 1.91385 | 0.700198 |
| 1-Mar | 6.32154 | 4.84968 | 7.45269 | 2.01399 | 4.00285 | 1.93079 |
| Map3k8 | 2.96939 | 4.90708 | 6.10769 | 2.07801 | 2.13032 | 1.68333 |
| Tifab | 7.82955 | 4.05379 | 8.91476 | 3.21651 | 4.06158 | 1.14116 |
| Tcn2 | 8.39404 | 20.1957 | 21.0431 | 10.7131 | 6.02665 | 5.45618 |
| Atf6 | 1.4698 | 1.21576 | 1.54276 | 0.688533 | 0.456984 | 0.73269 |
| Kremen1 | 0.565383 | 0.962343 | 0.991483 | 0.214724 | 0.405673 | 0.400392 |
| Ndst2 | 4.20505 | 6.38287 | 5.67683 | 2.82614 | 2.95233 | 1.85645 |
| Parp12 | 2.52527 | 6.23633 | 6.51809 | 1.44328 | 3.07819 | 1.82024 |
| Rftn1 | 2.59106 | 3.45887 | 3.87254 | 0.574757 | 2.36565 | 1.68751 |
| Nfkbid | 1.22499 | 2.02355 | 2.61865 | 0.904085 | 1.28387 | 1.16938 |
| Kcnab2 | 2.41219 | 2.39046 | 3.25853 | 1.57929 | 1.81983 | 0.414155 |
| Elmsan1 | 1.78933 | 1.58059 | 2.35107 | 0.627577 | 1.01737 | 0.914592 |
| Gpc2 | 12.8761 | 11.9237 | 7.61048 | 6.60096 | 6.28534 | 7.86951 |
| Pcdhga9 | 1.81206 | 1.01169 | 1.48342 | 0.676761 | 1.13654 | 0.678242 |
| Snx9 | 5.11396 | 4.57936 | 4.50067 | 1.71168 | 2.76609 | 1.78226 |
| Pou3f1 | 14.1936 | 5.9067 | 8.57918 | 5.35473 | 3.70474 | 4.74115 |
| Hps5 | 5.21313 | 2.84035 | 5.71963 | 1.06674 | 2.8928 | 1.32303 |
| M6pr | 13.0232 | 17.6401 | 20.8353 | 7.65787 | 9.10843 | 4.96163 |
| 1700017B05Rik | 4.43263 | 3.1506 | 4.99704 | 1.91581 | 1.93143 | 2.05554 |
| Shc1 | 3.52247 | 3.05414 | 4.74659 | 1.35918 | 2.18885 | 1.89979 |
| Wdr81 | 3.88334 | 4.17681 | 4.55544 | 1.93487 | 1.56558 | 2.42105 |

| | | | | | | |
|---|---|---|---|---|---|---|
| Itga1 | 0 | 0.146427 | 0.0456913 | 0.530133 | 0.465997 | 0.332961 |
| Slc10a4 | 0.730649 | 0.440765 | 0.516534 | 4.12754 | 2.06896 | 9.58327 |
| Sulf2 | 0.22759 | 0.55349 | 0.342124 | 1.6972 | 5.19181 | 2.18112 |
| Epha10 | 0 | 0.0237051 | 0 | 0.374867 | 0.481635 | 0.220058 |
| Mxra8 | 0.132187 | 0 | 0.124078 | 0.515842 | 1.0362 | 1.28502 |
| Evc | 0.00804294 | 0.0172844 | 0.00754313 | 0.409818 | 0.244817 | 0.474167 |
| Six3 | 0.0329421 | 0.336015 | 0.295356 | 3.75632 | 0.99424 | 1.21292 |
| Hbb-y | 0.543188 | 2.65711 | 3.75513 | 21.6405 | 17.2684 | 17.4205 |
| Mybl1 | 0.00994986 | 0.0103916 | 0.0272226 | 0.228461 | 0.433725 | 0.130617 |
| Megf10 | 0.10267 | 0.217184 | 0.0875047 | 0.748021 | 1.24295 | 1.10661 |
| Ripply2 | 0.423237 | 0.238221 | 1.07077 | 3.09492 | 2.76213 | 4.77021 |
| Asb4 | 0.0261291 | 0.108075 | 0.0941793 | 0.896767 | 0.452632 | 0.828859 |
| Id1 | 2.30283 | 2.55265 | 2.92942 | 22.9811 | 14.6162 | 22.1772 |
| Pomc | 0.388391 | 0.43056 | 0.88531 | 8.31003 | 4.60485 | 3.48475 |
| Ndufa4l2 | 0.12913 | 0.568033 | 0.624886 | 5.23422 | 2.44029 | 3.53343 |
| Optc | 0.0273114 | 0.238389 | 0.0517594 | 1.38978 | 0.431797 | 0.670545 |
| Sparc | 1.19592 | 1.633 | 2.11829 | 19.1774 | 13.0654 | 16.6751 |
| Pih1d2 | 0 | 0.0236777 | 0 | 0.574702 | 0.442839 | 0.369106 |
| Col6a3 | 0.0311559 | 0.00676718 | 0.0414796 | 0.221937 | 0.174264 | 0.252758 |
| Mylk | 0.0290038 | 0.00787656 | 0.0137732 | 0.241564 | 0.492106 | 0.387933 |
| Gucy1b2 | 0.0530494 | 0.0876048 | 0.0507634 | 0.959986 | 0.603649 | 0.424604 |
| Ntn1 | 0.0141435 | 0.0920919 | 0.053705 | 0.377971 | 0.323352 | 2.0863 |
| Cd248 | 0.273866 | 0.445877 | 0.194314 | 2.83737 | 1.9099 | 3.47529 |
| Gbx2 | 0.559752 | 1.11264 | 0.823782 | 2.9238 | 18.9124 | 2.36404 |
| Vwa1 | 0.0417993 | 0.202077 | 0.236638 | 1.67283 | 1.64588 | 0.504774 |
| Gng11 | 2.49773 | 3.96263 | 3.49213 | 30.8767 | 28.9171 | 19.5775 |
| Etv4 | 0 | 0 | 0 | 1.05902 | 0.512047 | 0.422976 |
| Spag6l | 0.0242298 | 0.050956 | 0.0332976 | 0.376108 | 0.821029 | 0.364242 |
| Rhbdf1 | 0 | 0.0328642 | 0.0577887 | 0.48663 | 1.24457 | 0.197066 |
| Epas1 | 0.0336824 | 0.0238284 | 0.010276 | 0.581486 | 0.279087 | 0.219191 |
| Galnt18 | 0.208779 | 0.0326613 | 0.199154 | 0.770236 | 2.7616 | 1.29911 |
| Ddr2 | 0.0124136 | 0 | 0.0115514 | 0.190221 | 0.0689778 | 0.282149 |
| Cxcl1 | 0.165469 | 0.730975 | 0.641926 | 1.5609 | 5.91341 | 1.28968 |
| Gpx8 | 1.30902 | 2.36139 | 2.22711 | 22.0729 | 8.43877 | 22.7444 |
| Scg2 | 0.602948 | 0.744765 | 0.949077 | 8.00129 | 5.6465 | 3.42077 |
| Rgs4 | 0.465146 | 0 | 0.566877 | 3.06205 | 1.46423 | 2.44345 |
| Slc38a3 | 0.150534 | 0.148544 | 0.0470638 | 0.428728 | 1.09662 | 0.309445 |
| Zfp641 | 0.0463487 | 0 | 0.014493 | 0.600475 | 0.446056 | 0.292341 |
| Lingo2 | 0.0260383 | 0.0545874 | 0.0535364 | 0.365154 | 0.540617 | 0.258002 |
| Ube2cbp | 0.0514516 | 0.397153 | 0.649194 | 1.65524 | 2.83919 | 3.17882 |
| Gm973 | 0.0744747 | 0.0332778 | 0 | 0.250848 | 0.355279 | 0.394091 |
| Esrrg | 0.131852 | 0.13996 | 0.236032 | 2.35968 | 0.437908 | 0.675111 |
| Spag5 | 0.0937536 | 0.151967 | 0.177328 | 1.11861 | 1.09148 | 0.764721 |
| Scrg1 | 2.1676 | 1.28398 | 2.61667 | 16.3906 | 12.8219 | 16.7155 |
| Car10 | 0.0335157 | 0.0880284 | 0.106903 | 0.176904 | 1.06913 | 0.267993 |
| Usp13 | 0.0176868 | 0.0283211 | 0.0413776 | 0.394424 | 0.421326 | 0.233496 |
| Slc6a13 | 0.05315 | 0.628066 | 0.426421 | 3.48648 | 1.21469 | 2.12449 |
| Ptx3 | 1.07609 | 1.23743 | 1.54869 | 6.95099 | 9.84106 | 8.5963 |
| Nr2e1 | 0.578774 | 0.684955 | 0.842474 | 5.60137 | 3.13924 | 6.23508 |
| Igfbp5 | 0.136079 | 0.335602 | 0.433741 | 2.2537 | 2.00047 | 2.27555 |
| Hes5 | 5.3274 | 5.00057 | 4.9729 | 42.0839 | 31.6931 | 55.9588 |
| Olig1 | 1.40486 | 0.880554 | 0.976702 | 16.503 | 3.55139 | 6.93903 |
| Kcnmb2 | 0.316407 | 0.062797 | 0.238449 | 1.30557 | 1.38973 | 1.39495 |
| P4ha3 | 0.12891 | 0.179598 | 0.0699539 | 1.21172 | 0.395999 | 1.07256 |
| Lrrc4c | 0.117108 | 0.345674 | 0.150457 | 1.4124 | 1.93377 | 0.833489 |
| Nhej1 | 0.149782 | 0.9065 | 0.652232 | 5.02991 | 3.09653 | 3.94993 |
| Mrc2 | 0 | 0.0229959 | 0.0501093 | 0.514739 | 0.461669 | 0.716427 |
| Prss23 | 0.0343963 | 0.182461 | 0.239566 | 0.933348 | 0.797165 | 1.30335 |
| Nova1 | 0.278495 | 0.325555 | 0.239979 | 1.0344 | 2.59707 | 1.58172 |
| Ptgds | 0.537881 | 0.584519 | 0.74398 | 4.01593 | 6.58587 | 2.54498 |
| Nid1 | 0.160694 | 0.182833 | 0.0187578 | 1.27232 | 0.753393 | 0.464078 |
| Cobl | 0.115309 | 0.0698811 | 0.0539874 | 0.758442 | 0.437157 | 0.250516 |
| Parp8 | 0.1763 | 0.262749 | 0.394931 | 1.96527 | 2.53639 | 2.04956 |
| Rit2 | 0.184242 | 0.312701 | 0.353484 | 0.817242 | 3.27646 | 1.284 |
| Tcte2 | 0.0155011 | 0.0912757 | 0.0363178 | 0.921151 | 0.541902 | 0.442524 |
| Fgfr3 | 0.235418 | 0.296182 | 0.20954 | 1.54356 | 1.09262 | 2.67247 |
| Prr5l | 0.156227 | 0 | 0.0605211 | 0.350308 | 0.807031 | 0.538092 |
| Fabp7 | 39.2077 | 53.1142 | 54.6305 | 365.237 | 225.26 | 269.606 |
| S1pr1 | 0.845799 | 0.348354 | 0.704536 | 4.61499 | 3.31636 | 4.2903 |
| Fkbp10 | 0.0256586 | 0.608241 | 0.410503 | 2.40584 | 1.92862 | 2.45781 |
| Ltbp3 | 0.154935 | 0.319228 | 0.204499 | 1.2158 | 1.36815 | 1.28206 |
| Nkain4 | 0.717552 | 0.700431 | 0.614938 | 8.03637 | 6.00906 | 7.77672 |
| Pmp22 | 0.700778 | 1.16313 | 0.984844 | 7.17328 | 3.9297 | 6.95339 |
| Car14 | 0.0254532 | 0.191162 | 0.190853 | 1.21009 | 0.563865 | 0.706515 |
| Slc24a3 | 0.164498 | 0.357706 | 0.104216 | 1.32997 | 1.60288 | 0.689582 |
| Nme5 | 0.376754 | 0.616651 | 0.542817 | 3.39525 | 2.5499 | 2.53213 |
| Frmd6 | 0 | 0.0302608 | 0 | 0.816572 | 0.557082 | 0.760278 |
| Acsbg1 | 0.269465 | 0.197292 | 0.258224 | 2.22598 | 1.17739 | 2.47301 |
| Snhg11 | 0.564431 | 1.2749 | 1.29235 | 6.41305 | 9.11865 | 2.32242 |
| Mrgpre | 0.201492 | 0.197336 | 0.184842 | 0.494402 | 1.88501 | 1.57656 |
| Efna1 | 0.280217 | 0.393661 | 0.227982 | 2.41806 | 1.38964 | 1.96339 |
| Cthrc1 | 0.16453 | 1.18481 | 1.19559 | 6.14655 | 5.25192 | 3.42255 |
| Fam229b | 0 | 0.0165568 | 0.0434011 | 0.217494 | 0.137738 | 0.302178 |
| Ccdc80 | 0.913483 | 0.11613 | 0.522733 | 4.69337 | 2.38519 | 3.5976 |
| Rai2 | 0.481726 | 0.790936 | 0.731278 | 5.95254 | 3.05807 | 4.76718 |
| Mt3 | 40.9957 | 31.8331 | 40.8295 | 320.085 | 180.591 | 244.106 |
| Nes | 1.05027 | 1.65382 | 1.73262 | 9.18051 | 6.51862 | 14.9974 |
| Kynu | 0.0596865 | 0.0104914 | 0.0829753 | 0.186953 | 0.194284 | 0.15862 |
| Rarb | 0.0907351 | 0.269814 | 0.0640873 | 2.77891 | 0.545189 | 1.88181 |
| Emid1 | 0 | 0.428002 | 0.321509 | 2.24887 | 1.46071 | 2.44827 |
| Cdc42ep1 | 0.282917 | 0.269733 | 0.304308 | 2.05519 | 2.00599 | 1.67988 |
| Dmrtb1 | 0.944441 | 2.12989 | 1.57582 | 6.28753 | 15.5302 | 8.13367 |
| Pars2 | 0 | 0.0258117 | 0.0676244 | 0.522741 | 0.74739 | 0.904208 |
| Fgf9 | 0.138682 | 0.0368855 | 0.17821 | 0.594851 | 0.754313 | 0.809661 |
| Cml1 | 0.383997 | 0.724166 | 0.826292 | 4.9705 | 4.6677 | 1.6571 |
| Syt17 | 0.0879234 | 0.0944992 | 0.0828925 | 0.634818 | 0.940525 | 0.573204 |
| Dusp26 | 1.73158 | 2.31779 | 2.77889 | 13.923 | 20.7645 | 13.6178 |
| Fam149a | 0.1472 | 0.0727768 | 0.265188 | 0.803211 | 1.58147 | 0.47607 |
| Kcnip4 | 0.174289 | 0.186272 | 0.0162609 | 0.532823 | 1.22203 | 0.724352 |
| Pax6 | 0.255217 | 0.579681 | 0.801824 | 3.28636 | 2.57547 | 3.43108 |
| Mb21d2 | 0.0842393 | 0.791252 | 0.455284 | 2.6907 | 2.16696 | 2.95507 |
| Lhx5 | 0.307688 | 0.74011 | 0.820987 | 2.27951 | 7.17705 | 2.94739 |
| Nkx2-1 | 0.104745 | 0.929882 | 0.827941 | 2.82611 | 3.2081 | 5.89552 |
| Crabp1 | 2.02072 | 5.25706 | 2.49307 | 25.1532 | 14.354 | 23.5386 |
| Pdgfra | 0.0203375 | 0.116482 | 0.108023 | 0.28984 | 0.803941 | 0.355402 |
| Mlc1 | 0.876966 | 1.31992 | 0.524493 | 6.8285 | 3.8056 | 5.96739 |

| | | | | | | |
|---|---|---|---|---|---|---|
| Sell1 | 4.73252 | 4.73801 | 5.49711 | 2.77984 | 2.24593 | 2.24711 |
| Adar | 4.84912 | 6.4353 | 6.01106 | 2.11048 | 2.77986 | 2.98079 |
| Arf3 | 19.1777 | 17.7243 | 20.8426 | 9.18165 | 8.69189 | 6.44498 |
| Dgki | 1.64601 | 1.32986 | 0.972537 | 0.699406 | 0.529569 | 0.736982 |
| Mmp14 | 7.31608 | 16.2167 | 16.1718 | 5.49292 | 6.6259 | 7.82465 |
| Arhgap4 | 2.70093 | 2.07679 | 3.0921 | 2.74262 | 2.69093 | 1.81323 |
| Pgap2 | 16.371 | 7.26523 | 16.1821 | 8.54747 | 5.18818 | 4.11134 |
| Cadm4 | 20.3932 | 13.4635 | 10.7792 | 6.88977 | 5.61183 | 7.81947 |
| Hyou1 | 1.30084 | 1.88129 | 2.08845 | 0.628574 | 0.681373 | 0.857129 |
| Nin | 1.63856 | 1.61059 | 1.02654 | 0.613556 | 1.00338 | 0.883799 |
| Sphk2 | 4.75699 | 5.91833 | 6.98345 | 3.13599 | 3.23332 | 2.48264 |
| Slc35f6 | 1.36131 | 2.13649 | 2.01638 | 0.680725 | 1.1737 | 0.64918 |
| Dok1 | 8.08948 | 9.09973 | 15.2837 | 6.24705 | 5.45363 | 3.54349 |
| Lrrc59 | 34.7769 | 45.6008 | 51.5425 | 24.1844 | 17.4878 | 14.5907 |
| Plbd2 | 2.74825 | 3.75731 | 4.28842 | 1.50254 | 1.63285 | 1.66759 |
| Myadm | 3.23874 | 11.2663 | 9.97129 | 2.92448 | 2.99993 | 3.95021 |
| Ppp1r15b | 5.22676 | 6.90684 | 8.5438 | 3.07441 | 2.42427 | 2.87596 |
| Ankrd13a | 4.99052 | 6.01645 | 6.61435 | 3.06896 | 3.1149 | 1.78874 |
| Mtm1 | 0.200809 | 0.378686 | 0.250624 | 0.105909 | 0.10953 | 0.169483 |
| Pid1 | 6.7117 | 8.64618 | 10.8214 | 3.21399 | 4.22828 | 2.99625 |
| Nadsyn1 | 0.715668 | 0.757462 | 0.934113 | 0.305752 | 0.287772 | 0.416545 |
| Stk24 | 5.48293 | 6.4236 | 7.99919 | 1.78774 | 1.74821 | 4.21088 |
| Foxk1 | 1.62057 | 2.67873 | 1.97048 | 0.893665 | 1.0343 | 0.996973 |
| Mbnl1 | 2.74788 | 1.43465 | 2.68424 | 0.997857 | 0.889987 | 0.620172 |
| Rap1b | 107.067 | 116.698 | 154.697 | 46.9302 | 57.4108 | 28.1044 |
| Il6ra | 4.20029 | 5.04609 | 6.11061 | 2.75001 | 2.51734 | 1.08067 |
| Rac2 | 130.493 | 107.271 | 177.925 | 58.8553 | 81.9733 | 33.9047 |
| Naglu | 5.29018 | 5.72038 | 7.54771 | 3.20864 | 1.4706 | 3.58768 |
| Psen2 | 2.39482 | 3.68394 | 4.65698 | 1.78611 | 1.96954 | 1.50783 |
| Fignl1 | 5.21178 | 1.79296 | 5.51692 | 2.48927 | 1.416 | 0.84979 |
| Serp1 | 176.72 | 130.563 | 208.686 | 52.4654 | 91.8577 | 47.0727 |
| Pprc1 | 1.35784 | 2.53009 | 2.28847 | 0.547469 | 0.637779 | 1.36029 |
| Ptpn1 | 23.5441 | 23.6388 | 31.9868 | 11.8303 | 12.2373 | 12.4066 |
| Fen1 | 36.2338 | 21.449 | 39.2531 | 11.8913 | 14.2179 | 16.1935 |
| Myo1f | 8.49852 | 8.81014 | 10.7061 | 4.09653 | 4.83985 | 2.81293 |
| Ccdc126 | 1.43145 | 2.21114 | 2.10243 | 0.506912 | 0.744601 | 0.752163 |
| Ccdc88b | 3.10868 | 2.07644 | 3.55163 | 2.46353 | 1.49557 | 0.608492 |
| Coro1b | 37.9797 | 31.278 | 47.5897 | 21.7945 | 19.5948 | 15.6568 |
| Ipcef1 | 0.392992 | 0.287925 | 0.256814 | 0.110578 | 0.0816888 | 0.107264 |
| Adam17 | 2.30165 | 2.06347 | 3.42347 | 1.1938 | 1.12987 | 0.894193 |
| Sh2b2 | 2.4877 | 2.30623 | 1.91547 | 1.12522 | 0.916998 | 0.915951 |
| Abcc1 | 0.437382 | 0.857382 | 0.775378 | 0.378581 | 0.394663 | 0.294254 |
| Mboat1 | 1.18712 | 2.28103 | 2.0528 | 1.06151 | 0.696631 | 0.503099 |
| Icam1 | 9.10508 | 10.6706 | 14.9738 | 5.92748 | 7.06713 | 1.83749 |
| Opcml | 3.60086 | 3.92786 | 1.82376 | 0.975894 | 1.08375 | 1.77296 |
| 1700109H08Rik | 0.739027 | 2.06095 | 2.21856 | 0.888238 | 0.392597 | 0.497446 |
| Sik2 | 1.16622 | 0.799663 | 0.823756 | 0.496381 | 0.191194 | 0.562587 |
| Ppp3cc | 4.02226 | 2.57488 | 4.27374 | 1.25914 | 1.54111 | 1.40019 |
| Slc2a3 | 0.681376 | 2.24105 | 2.0035 | 0.50621 | 0.879926 | 0.766158 |
| Sipa1 | 11.6736 | 10.158 | 13.885 | 6.66948 | 6.38437 | 3.47728 |
| Cyp4f13 | 2.07377 | 1.19401 | 2.20562 | 0.704102 | 0.840342 | 0.634274 |
| Pigg | 1.57709 | 1.59677 | 1.14261 | 0.599777 | 0.318215 | 0.637977 |
| Slc25a26 | 0.733897 | 0.251153 | 0.673544 | 0.152247 | 0.662856 | 0.171843 |
| Hk3 | 8.67597 | 7.17737 | 11.866 | 3.54086 | 7.48082 | 2.3739 |
| Slc9a5 | 0.835019 | 0.963367 | 0.651103 | 0.47636 | 0.189449 | 0.683462 |
| Cd72 | 59.0619 | 31.6017 | 63.2428 | 24.189 | 31.0418 | 14.8271 |
| Crlf2 | 3.14697 | 2.7876 | 4.71194 | 2.17778 | 1.5316 | 2.3272 |
| Unc93b1 | 67.6286 | 40.43 | 80.4405 | 36.5959 | 32.4045 | 15.693 |
| Dusp16 | 1.75533 | 2.84297 | 3.15741 | 0.760075 | 1.36973 | 1.03985 |
| Acsl1 | 0.493181 | 0.530972 | 0.911277 | 0.247046 | 0.415246 | 0.226676 |
| Pqlc2 | 1.35159 | 2.55475 | 2.39578 | 0.802942 | 0.847529 | 1.33349 |
| Sft2d2 | 1.42417 | 1.40229 | 2.09592 | 0.390561 | 0.663572 | 0.664574 |
| Cflar | 1.42279 | 2.98215 | 3.02468 | 0.591028 | 1.08649 | 0.824707 |
| Wdr1 | 16.2906 | 12.8006 | 18.6449 | 7.31705 | 9.2979 | 5.20842 |
| Smarcd2 | 5.47921 | 4.25086 | 5.88388 | 3.11427 | 3.13794 | 2.12486 |
| Was | 8.06518 | 7.46086 | 9.24918 | 4.14698 | 3.81784 | 2.34133 |
| Tgfbr2 | 9.75002 | 6.13468 | 10.2865 | 2.03571 | 6.71134 | 1.76515 |
| Mocs1 | 1.36208 | 1.45366 | 1.96096 | 0.646471 | 0.616583 | 0.866556 |
| Abca7 | 0.913112 | 1.45493 | 1.38507 | 0.392184 | 0.558262 | 0.344091 |
| Nfatc3 | 1.37522 | 1.76036 | 1.57125 | 0.634264 | 0.417249 | 0.67034 |
| Zfp710 | 6.73765 | 6.07417 | 7.2547 | 2.99509 | 3.13007 | 2.40591 |
| Inpp5d | 2.75578 | 2.54701 | 4.04956 | 1.51796 | 1.34289 | 1.30855 |
| Ralgds | 6.04535 | 11.0582 | 9.91975 | 4.51786 | 4.06804 | 4.57017 |
| Cd53 | 117.857 | 115.83 | 166.729 | 48.1894 | 69.5954 | 27.41 |
| B3gat1 | 11.697 | 12.152 | 4.27025 | 3.06075 | 4.0652 | 5.45744 |
| Clstn2 | 5.1708 | 4.23942 | 1.73135 | 1.32731 | 1.69487 | 1.97722 |
| Telo2 | 1.26714 | 1.34729 | 0.75401 | 0.378784 | 0.326287 | 0.699558 |
| Zfp395 | 1.50523 | 2.11335 | 2.54149 | 0.957847 | 0.811479 | 0.705747 |
| Tm6sf1 | 3.58434 | 2.27251 | 3.89623 | 1.26937 | 1.49774 | 0.579444 |
| Grap | 3.81101 | 5.35379 | 7.51674 | 2.34276 | 3.39328 | 1.72473 |
| Spaca6 | 3.54785 | 4.83927 | 2.89124 | 1.07323 | 1.29658 | 1.79465 |
| Fli1 | 5.22764 | 4.27385 | 7.16209 | 2.98694 | 1.71355 | 1.60994 |
| Hiatl1 | 7.80596 | 12.9462 | 13.6263 | 3.23874 | 3.60237 | 5.19457 |
| Wdfy4 | 0.711179 | 0.570113 | 0.959309 | 0.391619 | 0.47423 | 0.127987 |
| Oaf | 2.25669 | 4.57166 | 5.3735 | 1.51875 | 1.78763 | 2.11323 |
| Kif1c | 0.769222 | 1.75325 | 1.62629 | 0.723362 | 0.560074 | 0.42702 |
| Tdrd7 | 3.09866 | 4.6967 | 4.29163 | 1.24936 | 2.28038 | 1.05794 |
| Skap2 | 5.0226 | 4.15022 | 5.97012 | 1.93269 | 2.12951 | 1.11869 |
| Sgk1 | 6.05362 | 16.8529 | 16.2272 | 6.24984 | 5.94647 | 3.15118 |
| Gltscr1 | 6.29272 | 5.42322 | 5.42578 | 1.77048 | 2.67685 | 3.32954 |
| Zfp626 | 1.11912 | 1.13127 | 1.01017 | 0.244811 | 0.694444 | 0.341449 |
| Lgals8 | 2.7668 | 5.05842 | 5.18598 | 1.49667 | 2.48981 | 1.15581 |
| Zfp869 | 9.15509 | 9.40379 | 8.83273 | 2.37817 | 2.8964 | 3.83925 |
| Nfatc2 | 1.03053 | 0.325627 | 0.991562 | 0.351295 | 0.435498 | 0.335805 |
| Spi1 | 68.799 | 61.1158 | 97.0251 | 33.5659 | 45.3779 | 19.7704 |
| A130051J06Rik | 2.59443 | 3.97448 | 2.92949 | 1.11697 | 1.02895 | 1.55478 |
| Proser3 | 1.21946 | 0.850532 | 1.0272 | 0.199909 | 0.533711 | 0.438687 |
| Ccdc149 | 1.09656 | 1.63937 | 1.25241 | 0.472001 | 0.389022 | 0.656696 |
| Arpc1b | 26.9276 | 23.2608 | 33.9693 | 11.5402 | 16.0439 | 7.93413 |
| Lrrc47 | 6.24367 | 6.43828 | 6.49621 | 2.90191 | 3.79703 | 2.85264 |
| Bcl6 | 1.49678 | 1.64247 | 2.31953 | 0.784973 | 0.493945 | 0.528402 |
| Itga6 | 6.70066 | 2.61292 | 5.82912 | 1.54742 | 2.716 | 1.72748 |
| Trip10 | 2.42322 | 6.38104 | 7.21308 | 2.21799 | 2.15992 | 2.33593 |
| Olfml2b | 2.11499 | 1.61611 | 1.58655 | 0.782977 | 0.578103 | 0.814962 |
| Tatdn2 | 2.03877 | 3.19453 | 3.24951 | 0.752513 | 1.23839 | 1.17854 |
| Hsd11b1 | 2.1375 | 1.73487 | 2.70719 | 0.721288 | 1.3338 | 0.321969 |

| | | | | | | |
|---|---|---|---|---|---|---|
| Cd34 | 0.219834 | 0.0662258 | 0.144885 | 1.10324 | 0.467807 | 0.564946 |
| Akap12 | 0.283481 | 1.22506 | 0.940477 | 4.79876 | 3.78498 | 4.56017 |
| Peg10 | 0.0298551 | 0.213297 | 0.213646 | 0.423565 | 0.862521 | 1.43116 |
| Fgfbp3 | 1.89343 | 1.45969 | 2.15685 | 14.2507 | 8.33133 | 8.3333 |
| Mast4 | 0.0499133 | 0.068798 | 0.0692336 | 0.207571 | 0.975424 | 0.224073 |
| Ccnd2 | 3.83035 | 8.10335 | 7.71886 | 28.7163 | 26.9593 | 31.9075 |
| Nrarp | 1.06039 | 1.43041 | 1.34817 | 8.05316 | 5.08384 | 9.95196 |
| Adamts1 | 0.295294 | 0.565118 | 0.339914 | 2.27032 | 1.48187 | 3.54614 |
| Fam181b | 2.85282 | 0.677948 | 2.12539 | 17.3204 | 7.79747 | 10.7838 |
| S100a16 | 1.12158 | 1.40791 | 1.06186 | 10.556 | 9.29572 | 3.75166 |
| 9430020K01Rik | 0.164272 | 0.0555165 | 0.177525 | 0.430008 | 0.551858 | 1.27833 |
| Slc44a5 | 0.0762403 | 0.0536227 | 0.0312677 | 0.36162 | 0.342104 | 0.496978 |
| Rfx4 | 0.0358888 | 0.139513 | 0.0992893 | 1.23455 | 0.365975 | 1.01167 |
| Notch3 | 0.0294927 | 0.147152 | 0.091866 | 0.451695 | 0.439667 | 0.830806 |
| Fstl1 | 0.76297 | 2.09173 | 2.3586 | 11.4578 | 7.85293 | 9.08351 |
| Mfge8 | 1.06199 | 0.854453 | 1.09504 | 6.64411 | 2.44366 | 7.43296 |
| Spock3 | 0.0143819 | 0.176483 | 0.0897182 | 0.703471 | 0.414105 | 0.230592 |
| Pdzrn4 | 0.643005 | 0.265432 | 0.78861 | 2.81523 | 5.19728 | 2.18466 |
| Copz2 | 0.0696991 | 0.185566 | 0.292816 | 0.951174 | 1.38375 | 0.903337 |
| Tfap2c | 1.17857 | 0.192731 | 0.735596 | 3.48876 | 2.72003 | 4.83549 |
| Ddah1 | 6.32496 | 5.17522 | 6.34921 | 37.8708 | 22.6643 | 30.1294 |
| Trank1 | 0.0777585 | 0.0603474 | 0.0684203 | 0.555455 | 0.496473 | 0.121371 |
| Reln | 0.123346 | 0.229425 | 0.171948 | 1.03355 | 0.94518 | 1.13987 |
| Col4a2 | 0.293216 | 0.542685 | 0.340689 | 3.12158 | 4.55394 | 2.97849 |
| Nrsn2 | 0.273271 | 1.29699 | 1.80751 | 5.00976 | 7.84135 | 7.47104 |
| Acyp2 | 0.633204 | 0.798724 | 0.901005 | 6.86954 | 4.61941 | 4.96747 |
| Actr3b | 0.90428 | 1.1582 | 0.833168 | 7.00138 | 3.02453 | 6.20014 |
| Nefm | 1.01404 | 2.22801 | 2.09708 | 10.0022 | 14.1395 | 6.60264 |
| Cxcl12 | 0.524688 | 0.224389 | 0.574177 | 2.73166 | 2.59933 | 1.39555 |
| Slc14a2 | 0.0911279 | 0.565225 | 0.492408 | 1.426 | 1.54619 | 2.80036 |
| Rims3 | 0.12009 | 0.218816 | 0.103912 | 1.1062 | 1.25289 | 1.15404 |
| Trip6 | 0.174624 | 1.39511 | 1.05605 | 5.39443 | 5.51714 | 4.77782 |
| Dbi | 9.45108 | 7.50262 | 9.19491 | 60.378 | 45.903 | 44.4512 |
| Plp1 | 0.0972955 | 0.115976 | 0.0790909 | 0.416023 | 0.616505 | 0.311571 |
| mt-Co3 | 20.0203 | 25.4046 | 24.5436 | 159.192 | 107.933 | 94.1843 |
| Dtna | 0.124127 | 0.0712889 | 0.0802741 | 0.543868 | 1.02265 | 0.564844 |
| Hes1 | 0.957868 | 1.8231 | 1.42091 | 7.81393 | 5.78292 | 12.1549 |
| Sox21 | 1.23388 | 1.16058 | 1.17947 | 6.17537 | 5.11406 | 7.62831 |
| mt-Nd1 | 236.106 | 284.416 | 296.57 | 1497.23 | 1198.18 | 1195.71 |
| Yap1 | 0.0143907 | 0.197499 | 0.133414 | 0.404371 | 0.579154 | 1.03824 |
| AK157302 | 1.08008 | 0.672633 | 0.946468 | 3.51748 | 2.19623 | 4.67099 |
| Tshz2 | 0.245298 | 0.191565 | 0.157845 | 1.7195 | 1.03233 | 0.901762 |
| Cryba4 | 0.896082 | 2.53121 | 1.24026 | 10.2402 | 13.6201 | 10.3519 |
| Pfn4 | 0.524445 | 1.28635 | 0.929091 | 4.30782 | 3.4238 | 5.46487 |
| Sgol1 | 0.17755 | 0.282722 | 0.316001 | 1.9639 | 1.55657 | 2.0305 |
| Pdpn | 0.995144 | 4.04365 | 4.10573 | 13.3166 | 13.5707 | 18.6265 |
| Nefl | 1.61405 | 4.06919 | 3.02418 | 7.85251 | 31.2443 | 7.04897 |
| Ajuba | 0.433543 | 0.144869 | 0.763985 | 3.01544 | 2.01783 | 2.9844 |
| Rpl26 | 5.15604 | 4.64767 | 4.8972 | 42.7713 | 40.6182 | 37.1806 |
| Zfp566 | 1.05475 | 0.533915 | 0.375344 | 2.98862 | 3.91943 | 2.94009 |
| Nr4a2 | 0.368612 | 0.81998 | 0.429856 | 3.03466 | 2.58937 | 2.32158 |
| Rab13 | 2.28251 | 1.33448 | 2.41901 | 11.3537 | 6.8876 | 12.2451 |
| Gm4353 | 0.888496 | 0.769422 | 0.670185 | 1.99585 | 2.68567 | 2.56337 |
| Steap2 | 0.0742412 | 0.034222 | 0.0783139 | 0.335665 | 0.178909 | 0.373498 |
| Aldoc | 3.03323 | 3.34913 | 5.11265 | 18.9053 | 13.1452 | 26.4668 |
| Sdc2 | 0.101703 | 0.541244 | 0.608797 | 1.18471 | 1.73441 | 2.69409 |
| Zfhx4 | 0.393004 | 0.413754 | 0.452857 | 1.99305 | 2.54796 | 1.85949 |
| Ablim3 | 0.343464 | 0.418705 | 0.377951 | 2.6629 | 3.72685 | 1.99516 |
| Zfp697 | 0.295358 | 0.183633 | 0.122594 | 1.21477 | 0.67328 | 0.796913 |
| Sfrp2 | 1.39248 | 2.15929 | 2.26118 | 12.5835 | 7.52343 | 9.25269 |
| Rcan2 | 0.613697 | 1.60899 | 1.12204 | 3.50232 | 8.06915 | 4.06776 |
| Nnat | 194.009 | 233.312 | 248.224 | 1213.94 | 1226.59 | 1043.99 |
| Rcn3 | 2.64772 | 5.20625 | 3.65189 | 26.3705 | 20.0784 | 23.1556 |
| Gpx7 | 1.69033 | 4.6307 | 3.70453 | 24.3515 | 13.817 | 11.8512 |
| Ccdc184 | 0.843442 | 2.35969 | 2.20002 | 7.9882 | 11.6635 | 7.17471 |
| mt-Atp8 | 272.749 | 411.179 | 356.944 | 1572.69 | 1201.88 | 1338.51 |
| Calb1 | 0.385591 | 0.907764 | 0.509769 | 2.11316 | 3.83876 | 1.17804 |
| Ppp1r14c | 2.37942 | 0.72623 | 0.724126 | 7.23873 | 4.69777 | 5.33076 |
| Alg12 | 0.0258173 | 0.361456 | 0.170147 | 1.70122 | 1.6069 | 0.937454 |
| mt-Nd4l | 20.2966 | 28.3 | 24.1957 | 104.356 | 82.8184 | 84.8104 |
| Ppic | 4.07285 | 6.124 | 7.49581 | 27.8098 | 21.1215 | 33.8806 |
| Chrna4 | 0.159344 | 0.344949 | 0.400442 | 1.72477 | 2.03482 | 1.18042 |
| Eid2 | 1.66265 | 2.09987 | 0.981721 | 6.75934 | 8.10625 | 7.44307 |
| Rpp25 | 0.861949 | 1.47095 | 2.18138 | 12.5926 | 4.50784 | 6.14908 |
| Nhs | 0.0590502 | 0.0952148 | 0.0967596 | 0.232502 | 0.658908 | 0.47233 |
| Sall2 | 0.673686 | 1.09738 | 1.17431 | 5.89569 | 5.35581 | 3.22338 |
| Pcbd1 | 2.03866 | 4.20145 | 3.67826 | 11.8604 | 24.1054 | 10.8698 |
| Hist1h2ao | 67.5584 | 87.5312 | 103.907 | 685.885 | 494.505 | 369.392 |
| Pabpc4l | 0.160638 | 0.0416819 | 0.136972 | 0.50231 | 0.299994 | 0.497347 |
| Cdh7 | 0.155729 | 0.0783948 | 0.0146774 | 0.223744 | 0.356465 | 0.490074 |
| Bcan | 0.750926 | 0.8408 | 0.449143 | 2.83275 | 2.15881 | 4.46958 |
| Tnc | 0.493658 | 0.299721 | 0.267349 | 2.05247 | 1.69081 | 1.03655 |
| Cdkn3 | 3.68621 | 1.59873 | 2.04816 | 16.671 | 13.9944 | 6.85446 |
| Gpc3 | 0.349696 | 1.2421 | 0.751981 | 2.64455 | 1.2239 | 7.37663 |
| Nap1l5 | 4.48723 | 3.25351 | 3.83995 | 14.8589 | 18.439 | 14.5293 |
| Zfp518a | 0.126999 | 0.251808 | 0.306398 | 0.718703 | 0.760647 | 0.95409 |
| Fam184a | 0.118127 | 0.242328 | 0.113436 | 0.773761 | 0.819012 | 0.386438 |
| Tgfb1i1 | 0.0459811 | 0.184693 | 0.23683 | 0.336559 | 1.07568 | 0.690382 |
| Iqsec3 | 0.146017 | 0.113449 | 0.0708571 | 0.783204 | 0.297705 | 0.504766 |
| Siglech | 0.131901 | 0.246181 | 0.168614 | 1.41651 | 0.771288 | 1.07941 |
| Sall3 | 0.214103 | 0.508329 | 0.380917 | 2.33587 | 1.13153 | 1.19402 |
| Egfr | 0.0268386 | 0.0743829 | 0.0645126 | 0.314644 | 0.232219 | 0.318021 |
| Tcta | 1.88413 | 2.02823 | 2.79164 | 13.7374 | 10.4539 | 7.851 |
| Serpinh1 | 0.540182 | 1.94514 | 1.96383 | 7.49256 | 6.37904 | 7.64811 |
| Zkscan2 | 0.295272 | 0.286229 | 0.13995 | 1.28402 | 0.843858 | 1.01547 |
| Dnal1 | 0.146577 | 0.170992 | 0.0338077 | 0.431889 | 0.341779 | 0.674726 |
| Mmp11 | 0.608389 | 0.361191 | 0.459376 | 2.9646 | 1.97871 | 1.36321 |
| Vstm2a | 0.325634 | 0.24711 | 0.389527 | 2.03655 | 1.35158 | 1.17237 |
| Slc1a3 | 1.06269 | 1.36108 | 0.829834 | 6.79829 | 4.1689 | 5.79684 |
| Serping1 | 1.16082 | 1.6258 | 1.21264 | 5.0597 | 9.21645 | 6.92956 |
| mt-Atp6 | 68.0601 | 73.5405 | 77.3297 | 279.392 | 247.288 | 226.019 |
| Prkcdbp | 2.87335 | 4.38668 | 3.5698 | 21.8704 | 14.7599 | 18.8334 |
| Nrsn1 | 1.83114 | 1.09405 | 1.69565 | 6.39031 | 7.87476 | 4.17821 |
| Spa17 | 4.40501 | 3.38011 | 3.3802 | 14.1628 | 12.9567 | 13.3716 |
| Tmem14a | 1.60881 | 0.897076 | 1.56814 | 6.72145 | 5.37354 | 5.4966 |
| Plpp3 | 2.09693 | 1.86374 | 2.46372 | 8.069 | 4.97985 | 14.5692 |

| | | | | | | |
|---|---|---|---|---|---|---|
| Ms4a7 | 6.23772 | 5.7217 | 8.90709 | 2.04796 | 3.53531 | 1.56164 |
| P2rx7 | 1.9947 | 2.47062 | 2.88719 | 0.958871 | 1.10943 | 0.454666 |
| Fam84b | 1.38755 | 2.22675 | 1.13264 | 0.420434 | 0.527416 | 0.900013 |
| Ttc9 | 3.68653 | 2.67498 | 3.25305 | 0.950464 | 1.25953 | 1.59001 |
| Cyp4v3 | 0.510052 | 2.30666 | 1.82266 | 0.887419 | 0.327671 | 0.497695 |
| Agps | 2.06056 | 2.8469 | 2.85097 | 0.827991 | 1.1709 | 0.910625 |
| Pik3cd | 3.0747 | 2.60787 | 2.66763 | 1.41788 | 1.28598 | 0.866171 |
| Adcy3 | 0.65719 | 0.45669 | 0.390489 | 0.384507 | 0.348533 | 0.41275 |
| Tmc6 | 0.580805 | 0.480601 | 0.622646 | 0.341044 | 0.151825 | 0.136285 |
| Dse | 6.39897 | 4.2543 | 6.50813 | 2.66813 | 2.14715 | 1.55156 |
| Pik3r4 | 0.989276 | 0.917885 | 1.01648 | 0.420001 | 0.293497 | 0.416618 |
| Galnt4 | 0.830565 | 0.811355 | 0.727695 | 0.212035 | 0.314335 | 0.260465 |
| Hivep3 | 2.09487 | 1.68345 | 2.1771 | 0.89767 | 0.712228 | 0.832059 |
| Rfwd3 | 6.78788 | 4.12853 | 8.49577 | 1.81271 | 1.71265 | 3.41211 |
| Actn1 | 1.72639 | 2.52449 | 2.9949 | 0.698931 | 0.885049 | 1.34616 |
| Filip1l | 3.13091 | 2.18587 | 3.26286 | 1.20951 | 1.44968 | 0.426192 |
| Arhgap25 | 3.58908 | 1.845 | 4.03729 | 0.965458 | 2.07019 | 0.71795 |
| Sod2 | 57.1495 | 103.674 | 116.405 | 24.3326 | 42.4653 | 29.2366 |
| Slc12a4 | 1.20283 | 1.20044 | 1.29896 | 0.779288 | 0.137349 | 0.44244 |
| Lrrc25 | 47.1735 | 64.8978 | 82.7199 | 31.28 | 38.0097 | 13.6961 |
| Rftl | 1.28562 | 2.13849 | 2.57733 | 1.82241 | 0.979881 | 0.92869 |
| 6720489N17Rik | 1.48104 | 2.32841 | 1.92835 | 0.632602 | 0.51173 | 0.839117 |
| Slc7a7 | 9.6649 | 5.42784 | 12.4016 | 4.86301 | 4.64551 | 2.21543 |
| Ttyh3 | 11.4544 | 11.5982 | 11.4988 | 4.57692 | 4.85267 | 6.56278 |
| Dennd4a | 1.69499 | 1.59762 | 1.84369 | 0.761504 | 0.751498 | 0.424671 |
| Trim8 | 34.6314 | 51.8888 | 44.2533 | 22.9869 | 14.111 | 15.565 |
| Lrmp | 3.96153 | 2.02824 | 4.41382 | 1.83352 | 1.55418 | 0.522226 |
| Ifi35 | 3.87836 | 8.76577 | 9.7946 | 2.07205 | 4.90421 | 1.42398 |
| Ctsz | 301.551 | 433.808 | 556.37 | 181.781 | 232.563 | 112.049 |
| Slc16a10 | 3.51536 | 3.41135 | 4.19745 | 1.2144 | 1.50138 | 0.852977 |
| Cmtm6 | 8.31258 | 12.8449 | 15.616 | 3.90312 | 4.63013 | 4.73301 |
| Frmd8 | 3.68709 | 3.02015 | 4.63984 | 1.2195 | 1.69045 | 1.65513 |
| Selplg | 6.97284 | 7.15232 | 10.8307 | 4.23227 | 4.05506 | 1.15955 |
| Lin9 | 0.860307 | 1.08627 | 1.12667 | 0.402345 | 0.399564 | 0.441679 |
| Ndrg1 | 4.20905 | 5.51789 | 4.78316 | 1.14959 | 1.60849 | 2.78803 |
| Lrrc8d | 2.87973 | 4.08143 | 3.0868 | 1.26237 | 1.20797 | 1.20063 |
| Kif13b | 2.60651 | 1.43486 | 2.68271 | 0.415487 | 0.9051 | 0.930325 |
| Atp1a1 | 18.0718 | 24.6806 | 28.9183 | 9.66854 | 9.12301 | 6.97105 |
| Clec12a | 2.3729 | 2.18254 | 3.30602 | 0.946684 | 0.94551 | 0.77816 |
| Acvrl1 | 1.83728 | 2.15576 | 2.89037 | 1.08148 | 1.25544 | 0.359279 |
| Lrch1 | 4.5785 | 4.62437 | 5.4951 | 2.04877 | 1.85477 | 1.85075 |
| Cntn2 | 23.1775 | 18.913 | 16.6755 | 5.17618 | 6.32098 | 12.1978 |
| Itgam | 18.7251 | 15.9385 | 23.6911 | 7.59741 | 11.4253 | 3.51002 |
| Tnip2 | 3.5061 | 6.40718 | 6.26802 | 0.891272 | 4.30079 | 0.471416 |
| Rps6ka1 | 17.1056 | 8.80277 | 18.1442 | 6.17852 | 7.67558 | 4.47661 |
| Myo1e | 5.11933 | 5.73334 | 7.50755 | 0.782008 | 4.67802 | 1.54925 |
| Tnfaip8 | 4.93612 | 2.94049 | 5.88056 | 1.61959 | 1.75485 | 1.40053 |
| Prkcb | 13.9834 | 7.2337 | 15.2181 | 4.608 | 4.78447 | 3.34231 |
| Diaph2 | 1.03619 | 1.27846 | 1.32163 | 0.324102 | 0.296656 | 0.468979 |
| Ehbp1l1 | 7.01955 | 4.81145 | 8.16739 | 2.81043 | 3.51478 | 2.13156 |
| Naip5 | 0.571578 | 0.699223 | 0.837396 | 0.235706 | 0.256582 | 0.213359 |
| Nfkbie | 2.31094 | 4.62418 | 5.17415 | 1.46559 | 2.00457 | 1.24351 |
| Pik3r6 | 2.09768 | 1.25923 | 2.47538 | 1.03227 | 0.411027 | 1.03542 |
| Pcsk7 | 3.67607 | 4.26849 | 4.06715 | 2.02413 | 1.66213 | 1.20305 |
| Ly9 | 1.6868 | 5.33608 | 4.8611 | 1.2332 | 1.75032 | 1.58785 |
| Pvr | 0.315206 | 0.812353 | 0.792029 | 0.185187 | 0.400097 | 0.264133 |
| Hlx | 3.06188 | 2.9949 | 5.27514 | 1.55742 | 1.48101 | 0.964114 |
| Ggta1 | 2.57106 | 1.79244 | 2.8461 | 0.770103 | 1.31965 | 0.688344 |
| Arid3a | 4.6041 | 5.5103 | 6.03056 | 2.05086 | 2.65677 | 0.955736 |
| Pla2g7 | 31.4824 | 44.0728 | 57.0791 | 14.2816 | 23.6316 | 10.773 |
| Acox3 | 1.24377 | 0.897096 | 1.11234 | 0.238131 | 0.399054 | 0.53259 |
| Helb | 1.95041 | 1.47125 | 1.9017 | 0.506504 | 0.414772 | 0.507551 |
| Cln3 | 0.831131 | 1.35867 | 1.2339 | 0.650063 | 0.628451 | 0.46099 |
| Tor3a | 6.44767 | 6.21844 | 9.11758 | 2.36945 | 3.69131 | 1.96184 |
| Chtf18 | 1.15805 | 1.04717 | 1.11591 | 0.226598 | 0.574343 | 0.460422 |
| Il17ra | 2.40588 | 3.87028 | 4.68331 | 1.89952 | 1.49314 | 0.670018 |
| Adgre1 | 112.366 | 51.6662 | 126.326 | 35.2998 | 51.3959 | 18.4875 |
| Dgkg | 0.664492 | 0.436444 | 0.601426 | 0.280792 | 0.0778881 | 0.319794 |
| Acsl5 | 38.9789 | 32.7857 | 51.1093 | 14.9496 | 20.6747 | 8.29881 |
| Tlr2 | 35.9649 | 46.1515 | 59.7083 | 19.7591 | 24.7817 | 6.48989 |
| Tapbp | 13.8932 | 24.1898 | 24.1001 | 10.2843 | 11.5578 | 5.72095 |
| Trim25 | 0.836929 | 1.56683 | 1.7487 | 0.737327 | 0.693758 | 0.297937 |
| H2-T24 | 1.49847 | 8.41959 | 6.88739 | 1.61958 | 2.48233 | 1.23071 |
| Traf3ip3 | 1.38618 | 0.676851 | 1.52018 | 0.607198 | 0.402482 | 0.227195 |
| Iqgap1 | 2.63838 | 5.19149 | 5.36858 | 2.09057 | 2.08202 | 1.22872 |
| Fmnl3 | 3.07624 | 1.61939 | 2.99993 | 1.52004 | 0.874806 | 0.761633 |
| Bbs5 | 2.08381 | 0.924724 | 1.18834 | 0.675615 | 0.316539 | 0.278501 |
| Lcp1 | 17.1405 | 15.3286 | 21.5535 | 6.67876 | 8.78086 | 3.47592 |
| Cast | 2.32109 | 4.57141 | 5.56728 | 1.0979 | 1.38232 | 1.48338 |
| Nfe2l3 | 5.95849 | 5.23671 | 1.23587 | 1.72077 | 0.78442 | 2.59594 |
| Efhd2 | 201.147 | 168.221 | 258.251 | 83.7474 | 97.7174 | 59.5415 |
| Tnip1 | 11.1223 | 13.9893 | 18.0346 | 4.64223 | 10.261 | 3.2578 |
| Arap1 | 3.26573 | 3.87756 | 5.00895 | 1.5935 | 2.68037 | 1.3739 |
| Ncf4 | 25.0159 | 31.4905 | 43.1092 | 15.7694 | 19.0513 | 7.62793 |
| Bin2 | 46.9726 | 23.5193 | 47.3803 | 17.1436 | 18.7613 | 7.47435 |
| Arhgef1 | 4.44987 | 3.78539 | 5.14313 | 1.86157 | 2.16561 | 1.67663 |
| Sun2 | 5.81734 | 7.47931 | 8.10857 | 2.80868 | 2.44138 | 3.47207 |
| Ripk1 | 1.72024 | 3.00192 | 3.30373 | 0.61585 | 1.15056 | 0.62982 |
| Tor1aip1 | 4.84333 | 5.84792 | 6.93107 | 2.59801 | 2.1608 | 1.42773 |
| Ppard | 0.498285 | 0.301878 | 0.474117 | 0.263401 | 0.184989 | 0.4246 |
| Soat1 | 4.0211 | 4.66666 | 5.77333 | 1.90421 | 1.80664 | 1.63522 |
| Sash1 | 1.71946 | 6.676 | 5.69562 | 2.06838 | 1.49676 | 1.19388 |
| Pcdhgb6 | 1.00167 | 3.2284 | 1.9177 | 1.18273 | 1.12468 | 1.43485 |
| Pqlc3 | 0.910943 | 0.902378 | 0.945598 | 0.350146 | 0.314899 | 0.229985 |
| Dusp5 | 5.53252 | 13.7779 | 14.4985 | 3.28301 | 5.87445 | 3.16084 |
| Mbd2 | 0.566292 | 0.988517 | 1.68697 | 0.549032 | 0.56238 | 0.231025 |
| Cdc14b | 0.524787 | 0.214803 | 0.407678 | 0.246913 | 0.0866628 | 0.217741 |
| Spsb1 | 3.42529 | 1.14012 | 2.78548 | 0.466448 | 1.11581 | 1.00286 |
| Map3k11 | 3.53811 | 7.17146 | 8.41713 | 2.42321 | 3.05837 | 2.02264 |
| Dapp1 | 2.93485 | 1.50452 | 3.01674 | 1.18576 | 0.938743 | 0.463086 |
| Lyst | 0.565731 | 2.11411 | 1.75495 | 0.419405 | 0.425189 | 0.604341 |
| Ripk3 | 6.63868 | 6.90217 | 9.48587 | 4.61833 | 2.65528 | 1.33755 |
| Mctp1 | 0.662258 | 0.86975 | 0.835562 | 0.194756 | 0.195148 | 0.41208 |
| Nfkb2 | 2.42378 | 7.40648 | 8.3738 | 2.00441 | 2.79413 | 0.885228 |
| Frrs1 | 0.798755 | 2.0191 | 2.09785 | 0.565034 | 0.54027 | 0.895004 |
| Snx18 | 9.05881 | 15.5106 | 16.6466 | 5.85979 | 4.32249 | 3.9639 |

| | | | | | | |
|---|---|---|---|---|---|---|
| Rnf113a1 | 0.91331 | 1.92156 | 2.39043 | 11.5684 | 4.9625 | 7.63817 |
| Clu | 1.96288 | 1.81689 | 1.47084 | 7.52422 | 5.8155 | 9.25891 |
| mt-Co2 | 31.2882 | 38.1559 | 40.2044 | 159.207 | 126.573 | 117.04 |
| Nap1l3 | 0.827537 | 1.00085 | 1.31139 | 4.69188 | 2.54676 | 4.91597 |
| Kbtbd3 | 0.182083 | 0.441322 | 0.361915 | 2.40524 | 1.37524 | 1.28773 |
| Adora1 | 0.357784 | 0.323845 | 0.152356 | 1.73351 | 0.905205 | 1.56796 |
| Arl4a | 0.879603 | 1.53792 | 1.52626 | 4.39897 | 5.41281 | 4.99034 |
| Sox8 | 0.391629 | 0.545092 | 0.740232 | 1.87447 | 1.25378 | 3.25813 |
| Uaca | 0.789947 | 0.145101 | 0.861143 | 3.17561 | 2.82394 | 2.11482 |
| Zfp882 | 0.227692 | 0.451739 | 0.529295 | 1.5538 | 0.848706 | 2.36904 |
| Col12a1 | 0.101932 | 0.0699581 | 0.0419523 | 0.181261 | 0.207418 | 0.416077 |
| Prmt8 | 0.609069 | 1.3033 | 1.16782 | 4.13705 | 6.30702 | 2.14949 |
| Nr2f1 | 4.72406 | 1.99314 | 3.49507 | 17.3507 | 12.5689 | 10.6601 |
| Diras2 | 0.377101 | 0.220487 | 0.22642 | 1.0053 | 1.18548 | 0.859087 |
| Rab3b | 0.406257 | 0.694065 | 0.554253 | 2.81547 | 1.98337 | 1.63799 |
| Tmem132b | 0.22588 | 0.0415036 | 0.231759 | 0.966048 | 0.435068 | 0.958071 |
| Tceal6 | 1.95719 | 2.65103 | 4.02793 | 11.8253 | 11.3415 | 11.9886 |
| Tpd52l1 | 1.76003 | 1.87419 | 0.818039 | 7.23116 | 6.68317 | 3.70618 |
| Ptgfrn | 0.169002 | 0.180557 | 0.231124 | 0.768267 | 1.03569 | 0.396552 |
| Rpl21 | 1.4294 | 0.954876 | 1.58568 | 4.42854 | 1.90881 | 4.36553 |
| Tcerg1l | 0.353185 | 0.341041 | 0.438682 | 0.778068 | 2.78474 | 0.920292 |
| Zcchc12 | 4.89121 | 12.6231 | 10.8171 | 38.6542 | 39.9523 | 37.3826 |
| Efna5 | 2.40292 | 1.96352 | 1.42782 | 7.83335 | 8.55692 | 8.64573 |
| Pcdhb7 | 0.896425 | 0.671085 | 0.276052 | 2.10497 | 1.50358 | 3.48326 |
| Fam155a | 0.726392 | 0.465829 | 0.588401 | 1.37397 | 3.7342 | 2.12271 |
| Slc12a2 | 0.200144 | 0.591483 | 0.411835 | 1.71337 | 1.67207 | 2.44575 |
| 1700048O20Rik | 1.03519 | 0.739277 | 0.324852 | 4.74152 | 2.46476 | 2.59731 |
| Frs3 | 0.528714 | 0.302569 | 0.29065 | 2.11496 | 1.64033 | 1.49099 |
| Esco2 | 0.782601 | 0.896261 | 1.03813 | 3.80608 | 3.04873 | 3.3658 |
| Col23a1 | 0.209311 | 0.345286 | 0.208186 | 1.45652 | 1.27327 | 1.66843 |
| Hpgd | 0.418286 | 2.07755 | 1.34128 | 7.07924 | 3.19297 | 5.5211 |
| Ccdc166 | 0.444705 | 1.21337 | 0.912898 | 3.22257 | 3.57596 | 2.9957 |
| Vcam1 | 0.482117 | 0.14124 | 0.456326 | 0.85368 | 1.18008 | 1.63227 |
| Emp1 | 0.144559 | 0.461597 | 0.33696 | 1.19918 | 1.40411 | 1.00067 |
| Hist1h2ap | 21.5154 | 30.2683 | 28.8819 | 255.631 | 115.994 | 147.44 |
| 1500009L16Rik | 3.52281 | 4.95988 | 2.25825 | 14.2749 | 13.2739 | 17.6235 |
| Npy | 30.7131 | 48.328 | 34.711 | 268.973 | 90.4647 | 113.007 |
| B9d1 | 0.730755 | 1.34716 | 0.798714 | 4.43732 | 4.72337 | 4.70743 |
| Sapcd2 | 0.818241 | 0.366299 | 0.771378 | 2.39714 | 1.58779 | 3.29813 |
| Fbxo45 | 0.296648 | 0.645987 | 0.581456 | 2.12221 | 1.93724 | 1.98637 |
| Mest | 2.90786 | 5.25675 | 4.62645 | 20.8096 | 14.9132 | 15.5575 |
| Ears2 | 0.142373 | 0.180326 | 0.230065 | 0.545721 | 0.755689 | 1.02029 |
| Pvrl3 | 0.126337 | 0.266971 | 0.112498 | 0.440271 | 0.734059 | 0.890706 |
| Unc5c | 0.183298 | 0.0373586 | 0.165347 | 0.402599 | 0.433458 | 0.461513 |
| Atp1b2 | 0.89612 | 0.855397 | 0.616872 | 3.61148 | 3.67035 | 4.08953 |
| Gm9844 | 90.9568 | 80.9882 | 78.6614 | 278.946 | 395.225 | 232.088 |
| Slc5a6 | 0.315554 | 0.119362 | 0.331416 | 1.51534 | 1.49129 | 1.94068 |
| Kitl | 1.98764 | 2.04516 | 0.545539 | 4.78824 | 8.37216 | 4.81407 |
| Paqr8 | 0.388771 | 0.297011 | 0.362737 | 0.879969 | 1.3808 | 1.95411 |
| St6galnac5 | 0.207816 | 0.2516 | 0.240511 | 0.537943 | 1.07803 | 1.16644 |
| Cenpf | 0.451874 | 0.78987 | 0.766622 | 2.4367 | 1.45181 | 2.93321 |
| Cacna2d2 | 0.0804261 | 0.0856256 | 0.119182 | 0.346993 | 0.400367 | 0.474304 |
| Kcnip3 | 0.812447 | 0.793083 | 0.829706 | 3.41441 | 3.40294 | 3.40536 |
| Gabra3 | 0.81846 | 0.468384 | 0.527009 | 1.71199 | 3.50159 | 1.34632 |
| Trit1 | 1.1425 | 0.999211 | 0.729719 | 6.62039 | 5.09714 | 2.17097 |
| Ankrd37 | 0.803888 | 1.59834 | 1.40058 | 3.87413 | 4.9478 | 3.02323 |
| Atp1b1 | 3.02961 | 3.42368 | 3.88432 | 12.8411 | 12.7666 | 14.7082 |
| Ckap2 | 1.41859 | 0.506016 | 0.998811 | 4.4024 | 2.23242 | 6.28601 |
| Cald1 | 0.292044 | 0.201204 | 0.35187 | 0.827903 | 0.797191 | 1.20043 |
| Rgcc | 9.39017 | 18.4624 | 19.1314 | 84.255 | 49.8053 | 69.5595 |
| Spry2 | 4.82387 | 10.7173 | 8.09876 | 34.7913 | 28.6461 | 20.0489 |
| Plxnc1 | 0.177141 | 0.323035 | 0.392035 | 0.787021 | 1.30869 | 1.47771 |
| Lhfp | 0.647829 | 0.56878 | 0.215239 | 1.45179 | 1.32983 | 1.82133 |
| Cks2 | 24.9843 | 31.101 | 37.1373 | 147.312 | 95.9183 | 83.7663 |
| Pcdh10 | 0.138735 | 0.178545 | 0.242653 | 0.73835 | 0.548918 | 0.541442 |
| Mfap2 | 15.2377 | 21.0303 | 18.3938 | 90.5425 | 64.4654 | 86.2086 |
| Fzd2 | 1.24742 | 0.985588 | 1.15998 | 3.96412 | 2.3943 | 6.42357 |
| Stxbp6 | 0.306592 | 0.373025 | 0.455234 | 1.44603 | 1.46493 | 1.3045 |
| Tmtc2 | 0.138112 | 0.265703 | 0.119665 | 0.421302 | 0.682773 | 0.39225 |
| Ptn | 9.48611 | 14.7307 | 15.0866 | 44.6648 | 50.5628 | 34.2765 |
| Gabrb1 | 0.247414 | 0.0966195 | 0.188858 | 0.286245 | 0.721301 | 0.603732 |
| H2afv | 41.8328 | 51.6449 | 54.2785 | 205.191 | 146.085 | 153.68 |
| Plppr4 | 0.270338 | 0.198859 | 0.392638 | 0.864723 | 0.57811 | 1.66363 |
| Gpr34 | 0.494535 | 0.89756 | 1.59368 | 4.64979 | 2.95985 | 2.2139 |
| Nat8l | 0.239986 | 0.165256 | 0.240848 | 0.957586 | 0.813748 | 0.652794 |
| Ptprg | 0.294444 | 0.248521 | 0.236116 | 0.662048 | 0.656127 | 1.35244 |
| Prox1 | 0.513033 | 0.42558 | 0.789991 | 1.33918 | 3.06496 | 1.95263 |
| 1500011K16Rik | 3.95278 | 5.10553 | 8.76903 | 18.8372 | 20.5107 | 19.2717 |
| Pck2 | 0.857371 | 0.345566 | 0.860913 | 2.87641 | 3.12569 | 3.0511 |
| Kcne11 | 2.88358 | 3.58124 | 5.09647 | 15.3145 | 12.2958 | 16.9829 |
| Col18a1 | 0.152153 | 0.278071 | 0.218762 | 1.41142 | 1.3006 | 1.54923 |
| Med31 | 2.70323 | 2.36015 | 2.96582 | 10.5282 | 7.52491 | 7.85154 |
| B230118H07Rik | 0.0609496 | 0.103775 | 0.124643 | 0.561215 | 0.462501 | 0.566223 |
| Pomgnt2 | 1.41722 | 0.693757 | 0.890284 | 4.76516 | 3.25003 | 4.30328 |
| Sox1 | 1.15896 | 1.11875 | 1.28088 | 4.76776 | 3.88584 | 3.301 |
| Rgs2 | 2.73823 | 1.26461 | 2.76603 | 12.7688 | 5.14671 | 6.83313 |
| Scrn1 | 1.39912 | 1.5376 | 1.24387 | 5.10898 | 4.34534 | 6.25155 |
| Nr2f2 | 1.91218 | 1.29037 | 1.66791 | 9.61094 | 4.80496 | 2.78122 |
| Acot1 | 5.22627 | 5.1471 | 4.75976 | 15.955 | 19.3294 | 18.0255 |
| Ndrg2 | 8.65575 | 9.97442 | 9.6693 | 34.8579 | 26.4928 | 38.1833 |
| Nup35 | 0.879371 | 0.865281 | 0.87547 | 3.13024 | 3.66904 | 2.47114 |
| Arxes1 | 1.86368 | 5.09521 | 3.18025 | 11.0282 | 7.98589 | 12.9405 |
| Pqbp1 | 10.1201 | 9.49175 | 12.7452 | 42.4744 | 40.1873 | 30.7686 |
| 15-Sep | 4.66645 | 4.57582 | 5.5269 | 19.4536 | 14.2403 | 11.3702 |
| Cxxc4 | 0.138974 | 0.332346 | 0.382603 | 0.815994 | 0.904483 | 0.817453 |
| Chst1 | 1.35687 | 2.64151 | 1.74105 | 5.70685 | 9.58226 | 5.21757 |
| Ppp1r1a | 9.57721 | 15.1189 | 14.8057 | 58.2254 | 48.3667 | 34.6964 |
| Tmem41a | 3.86796 | 4.46 | 3.4599 | 15.619 | 13.151 | 11.7629 |
| Phgdh | 4.84968 | 11.2013 | 10.7238 | 35.6667 | 22.1059 | 41.2141 |
| Brinp3 | 0.18577 | 0.224535 | 0.171638 | 0.809736 | 0.454348 | 0.681111 |
| Sat2 | 0.820671 | 1.90099 | 1.55091 | 4.83943 | 4.37821 | 5.46988 |
| Camk2a | 0.664813 | 0.199088 | 0.782415 | 1.40868 | 1.92583 | 1.84766 |
| Hap1 | 0.684231 | 3.12809 | 1.62079 | 5.3783 | 6.60398 | 7.60952 |
| Hey1 | 1.16022 | 0.989247 | 1.01274 | 3.82846 | 2.43873 | 4.50686 |
| Tmem38a | 0.961215 | 0.889215 | 0.685156 | 3.97019 | 2.48635 | 4.35944 |
| Sdc1 | 0.744599 | 0.473041 | 0.85208 | 2.24986 | 1.89045 | 3.18317 |

| | | | | | | |
|---|---|---|---|---|---|---|
| Edem1 | 5.70179 | 7.60855 | 7.81971 | 1.86117 | 2.99877 | 1.76452 |
| Ceacam1 | 0.582474 | 0.471118 | 0.580873 | 0.151806 | 0.178765 | 0.0785489 |
| Ap1g2 | 1.77808 | 0.99458 | 1.44892 | 0.464088 | 0.750571 | 0.447465 |
| Themis2 | 6.84706 | 9.89648 | 12.1969 | 3.43426 | 6.4274 | 1.14428 |
| Chpf2 | 5.09972 | 5.52059 | 6.07004 | 1.96537 | 1.56776 | 2.73675 |
| Slc31a2 | 12.6489 | 26.9089 | 30.1391 | 9.64011 | 9.53143 | 4.21885 |
| Plin2 | 29.1831 | 45.718 | 51.9971 | 13.9252 | 22.6859 | 9.39778 |
| Ncf1 | 17.019 | 20.5768 | 25.2674 | 6.7165 | 10.8882 | 4.92285 |
| Camta2 | 3.76482 | 3.12226 | 3.51004 | 1.35792 | 1.11335 | 0.752269 |
| Adamts3 | 3.95412 | 4.67529 | 1.31442 | 0.700367 | 0.626093 | 2.02443 |
| Nupr1 | 2.76142 | 7.47609 | 6.82745 | 2.31569 | 2.209 | 1.8308 |
| Slc39a4 | 1.30971 | 0.966344 | 1.64706 | 0.21081 | 0.542779 | 0.242692 |
| Myo1c | 1.09663 | 1.60751 | 1.75901 | 0.657834 | 0.607351 | 0.422997 |
| Dgkz | 5.19799 | 5.90301 | 7.51618 | 3.34519 | 3.02026 | 1.59067 |
| Nrros | 8.38694 | 10.5892 | 13.575 | 3.27807 | 5.6257 | 2.55346 |
| Ptger4 | 3.16624 | 2.06919 | 3.99121 | 1.2574 | 1.26157 | 0.33009 |
| Kcna3 | 15.929 | 13.5527 | 19.2672 | 9.74246 | 5.13686 | 1.93828 |
| Gclc | 3.44664 | 3.90099 | 3.69221 | 1.37668 | 1.69744 | 0.834771 |
| Stxbp2 | 5.33493 | 6.04873 | 7.78837 | 1.71629 | 3.27 | 1.91911 |
| Sgk3 | 0.608704 | 0.573536 | 0.673928 | 0.259128 | 0.309653 | 0.146902 |
| Fgl2 | 0.953121 | 1.6711 | 1.68928 | 0.281636 | 0.674427 | 0.307826 |
| Sema4d | 5.77601 | 7.27104 | 8.75891 | 3.11385 | 3.47025 | 1.54882 |
| Mapk14 | 9.2829 | 6.31168 | 11.0312 | 2.55855 | 3.63967 | 3.36666 |
| Tmem38b | 2.91082 | 2.77878 | 4.62101 | 1.1001 | 1.20559 | 1.1283 |
| Chst14 | 5.52903 | 4.75901 | 6.31467 | 1.78441 | 2.38413 | 1.56655 |
| Plek | 15.3601 | 20.6591 | 25.5876 | 5.99253 | 8.92006 | 3.1888 |
| Akna | 5.47289 | 6.795 | 7.6288 | 2.31882 | 2.59543 | 2.17969 |
| Relb | 2.22095 | 4.69792 | 5.2952 | 1.24794 | 1.94482 | 0.763072 |
| Gsr | 16.0975 | 36.0274 | 33.6691 | 7.82914 | 12.0134 | 9.16209 |
| Stk10 | 3.84259 | 3.84744 | 5.18363 | 1.58478 | 2.00165 | 0.780433 |
| Lrrk1 | 0.86208 | 1.07345 | 1.02991 | 0.257647 | 0.411563 | 0.293247 |
| Clec4a1 | 94.3919 | 65.0081 | 119.878 | 32.4019 | 41.0691 | 13.4062 |
| Gipr | 2.08865 | 1.5944 | 0.355358 | 0.610853 | 0.549655 | 0.37453 |
| Sash3 | 17.0809 | 11.8885 | 19.2131 | 6.05635 | 7.55131 | 3.47166 |
| Peak1 | 0.728304 | 1.86328 | 1.56078 | 0.288022 | 0.475039 | 0.545983 |
| Coro2a | 1.38001 | 1.29579 | 1.7682 | 0.496901 | 0.923563 | 0.147709 |
| Stap1 | 1.18535 | 0.802706 | 1.45696 | 0.188819 | 0.591135 | 0.242254 |
| Socs1 | 3.88362 | 12.9888 | 10.0232 | 2.59702 | 3.49534 | 3.88386 |
| Znfx1 | 0.821749 | 3.04379 | 2.84266 | 0.292374 | 1.3963 | 0.398206 |
| Grn | 67.5239 | 67.5626 | 96.8372 | 32.7679 | 37.0062 | 17.9473 |
| Sla | 8.48829 | 5.73887 | 7.5671 | 2.13634 | 2.2762 | 2.74806 |
| Zdhhc5 | 5.25198 | 5.06367 | 6.0033 | 1.35198 | 1.71152 | 2.00118 |
| Plcg2 | 11.1055 | 5.74052 | 11.4512 | 4.63571 | 3.76504 | 1.6926 |
| Il13ra1 | 2.25822 | 9.05989 | 7.24203 | 1.99756 | 2.50763 | 1.05714 |
| 1700112E06Rik | 0.364538 | 0.224161 | 0.269161 | 0.502708 | 0.228397 | 0.0401262 |
| Sec24d | 0.835785 | 0.935857 | 0.973328 | 0.306266 | 0.343727 | 0.339848 |
| Gsap | 1.18373 | 1.34966 | 1.58388 | 0.181915 | 0.683314 | 0.387994 |
| Abtb2 | 0.634084 | 1.12923 | 1.28443 | 0.231586 | 0.638268 | 0.281666 |
| Tmem106a | 4.21836 | 7.6623 | 7.17927 | 2.25746 | 3.78504 | 0.887697 |
| Msn | 25.5548 | 28.7901 | 37.4264 | 11.135 | 10.3518 | 6.89788 |
| Tcirg1 | 1.85859 | 4.98744 | 5.41234 | 1.85076 | 2.50594 | 0.832527 |
| Fmn1 | 0.605061 | 0.293627 | 0.549545 | 0.13749 | 0.177708 | 0.085455 |
| Tmem194 | 0.537014 | 0.446762 | 0.534987 | 0.212084 | 0.104404 | 0.165366 |
| Slc7a8 | 5.1151 | 8.35922 | 9.18164 | 1.92347 | 3.09732 | 2.81043 |
| Jak3 | 6.42764 | 2.73116 | 7.03981 | 3.47259 | 1.85204 | 0.59414 |
| Snx7 | 0.446291 | 1.09043 | 1.02311 | 0.487903 | 0.173261 | 0.204223 |
| Irf5 | 14.7696 | 14.6235 | 22.7296 | 6.33954 | 8.42508 | 3.89824 |
| Il4ra | 6.77767 | 7.53016 | 10.4549 | 3.14454 | 3.85187 | 2.1649 |
| Rbm43 | 2.61744 | 3.06708 | 4.26374 | 1.21074 | 0.697884 | 1.1415 |
| Fhl3 | 4.20811 | 5.73585 | 7.92473 | 1.78371 | 3.23967 | 0.929366 |
| Fchsd1 | 1.41086 | 1.0465 | 1.26266 | 0.350212 | 0.79177 | 0.883586 |
| Slc35b3 | 5.37257 | 4.97279 | 5.45479 | 1.67849 | 0.484432 | 2.03774 |
| Mpeg1 | 22.256 | 24.9666 | 33.28 | 8.68289 | 10.3875 | 5.85493 |
| Hmha1 | 4.06313 | 2.44444 | 5.05609 | 1.89424 | 1.94017 | 0.972082 |
| Dtx2 | 2.65741 | 5.36766 | 5.75035 | 1.61343 | 1.53404 | 1.11783 |
| Adcy7 | 4.2417 | 1.22915 | 4.38104 | 0.975144 | 1.43488 | 0.855854 |
| Elf4 | 3.23375 | 1.90374 | 3.75342 | 0.776295 | 1.26039 | 0.929206 |
| Nrip3 | 0.697843 | 3.17892 | 2.81138 | 0.652324 | 0.854594 | 0.592249 |
| Mxd1 | 1.37141 | 5.74991 | 5.32146 | 1.71826 | 1.00661 | 1.49962 |
| Parp4 | 0.867681 | 1.03108 | 0.984279 | 0.080742 | 0.581436 | 0.211554 |
| Neurl3 | 6.84656 | 3.72949 | 7.30929 | 2.61917 | 2.30546 | 0.800269 |
| Scube1 | 1.91747 | 3.07506 | 2.39444 | 1.07148 | 0.645855 | 0.996721 |
| Gpr35 | 0.617131 | 1.44697 | 1.65384 | 0.364685 | 0.268821 | 0.566092 |
| Ldlrap1 | 1.8039 | 1.65996 | 2.0378 | 0.364716 | 0.896859 | 0.569275 |
| Cx3cr1 | 88.0921 | 32.307 | 87.9355 | 21.8199 | 27.6436 | 15.9353 |
| Cytip | 0.500686 | 0.750072 | 0.827736 | 0.19448 | 0.213321 | 0.186725 |
| Mmp17 | 1.82907 | 1.73339 | 1.05017 | 0.606725 | 0.422237 | 0.624547 |
| Ldlr | 3.18007 | 6.69411 | 7.07906 | 1.73923 | 2.29459 | 1.23126 |
| Zfp11 | 0.796179 | 0.553109 | 0.960304 | 0.210976 | 0.0694184 | 0.298548 |
| Cdh12 | 0.867641 | 0.346559 | 0.737304 | 0.338332 | 0.0561693 | 0.114584 |
| Tmem63c | 0.777801 | 0.863227 | 0.608655 | 0.413524 | 0.054 | 0.45781 |
| Glrx | 55.4509 | 199.021 | 186.496 | 41.4967 | 58.8597 | 33.4647 |
| Il16 | 4.36882 | 1.78346 | 4.4278 | 1.115 | 1.16899 | 0.62049 |
| Hif3a | 0.75845 | 0.663002 | 1.08571 | 0.32214 | 0.281581 | 0.21824 |
| Hck | 99.1469 | 66.8499 | 118.525 | 28.6545 | 51.8895 | 16.3895 |
| Cd244 | 0.796306 | 0.684838 | 1.05746 | 0.247713 | 0.509497 | 0.170462 |
| Tfeb | 0.969873 | 1.42306 | 1.43688 | 0.591356 | 0.421506 | 0.49623 |
| Fmnl1 | 2.21495 | 2.60518 | 2.64344 | 0.705612 | 1.35756 | 0.512487 |
| Zc3hav1 | 5.20487 | 6.26139 | 8.1748 | 1.82104 | 3.32033 | 0.974257 |
| Lyn | 24.395 | 20.5512 | 31.7192 | 7.51325 | 10.6755 | 3.56001 |
| Lpcat2 | 13.642 | 12.7607 | 17.2229 | 5.3272 | 4.80333 | 3.01448 |
| C3ar1 | 47.77 | 52.883 | 71.8217 | 16.0135 | 23.3669 | 7.82953 |
| Relt | 2.39907 | 1.12217 | 2.28988 | 1.00348 | 0.239483 | 0.674292 |
| Myd88 | 12.1871 | 16.6145 | 21.2196 | 5.64639 | 7.53379 | 2.67603 |
| Atp8b4 | 0.239177 | 0.95915 | 0.849743 | 0.296879 | 0.11201 | 0.183861 |
| Fuca2 | 4.52115 | 3.9212 | 5.81586 | 1.62489 | 1.48194 | 0.964785 |
| Xylt2 | 4.13976 | 3.82614 | 5.88857 | 0.711064 | 3.01948 | 0.905088 |
| Tmem189 | 9.87142 | 15.1663 | 15.7629 | 4.92494 | 4.35826 | 1.73628 |
| Cybb | 11.4031 | 13.2362 | 18.2388 | 3.91869 | 6.67023 | 1.87274 |
| Mcm10 | 1.67055 | 1.01397 | 1.51327 | 0.545539 | 0.38637 | 0.515499 |
| Nabp1 | 0.321547 | 0.175607 | 0.218587 | 0.0267674 | 0.0556471 | 0.109011 |
| Tlr9 | 3.28613 | 2.5622 | 3.10651 | 1.71118 | 0.879063 | 0.218245 |
| Prune2 | 1.23475 | 0.531719 | 1.36937 | 0.296821 | 0.314726 | 0.227703 |
| Upf1 | 11.314 | 8.84494 | 13.0109 | 4.40221 | 3.52789 | 4.55049 |
| Ak4 | 0.687453 | 1.89732 | 1.86918 | 0.370313 | 0.373917 | 0.53698 |
| Slc1a1 | 21.9769 | 20.1873 | 30.7676 | 8.30655 | 13.3922 | 5.80175 |

| | | | | | | |
|---|---|---|---|---|---|---|
| Spint2 | 0.417472 | 1.24001 | 1.02992 | 2.38811 | 2.41557 | 4.44576 |
| Tspan7 | 2.44419 | 2.3794 | 2.55676 | 9.56766 | 4.52299 | 9.45689 |
| Laptm4b | 2.50462 | 3.4159 | 2.21468 | 13.4955 | 6.4725 | 10.1862 |
| Gm8186 | 28.7974 | 34.1358 | 42.5399 | 118.468 | 128.127 | 109.93 |
| Gprasp2 | 1.54472 | 2.43099 | 1.60438 | 4.9668 | 8.03756 | 4.8519 |
| mt-Nd2 | 97.729 | 124.546 | 126.45 | 353.988 | 337.895 | 348.962 |
| Mzt2 | 3.26048 | 3.00985 | 3.02058 | 12.0137 | 11.7044 | 10.5982 |
| Rita1 | 1.01885 | 0.88804 | 1.0556 | 5.18792 | 4.31463 | 2.86214 |
| Mxd3 | 0.948127 | 0.937243 | 1.39449 | 5.60962 | 2.13578 | 4.21204 |
| Tefm | 1.29092 | 0.623748 | 1.59763 | 4.08978 | 4.55416 | 3.8155 |
| Gstm5 | 4.51863 | 5.59946 | 4.76868 | 18.4163 | 18.5022 | 16.4236 |
| Hist3h2ba | 213.25 | 141.048 | 193.155 | 734.632 | 630.52 | 606.444 |
| Rgs17 | 0.216574 | 0.5133 | 0.267586 | 1.01536 | 0.564164 | 1.22738 |
| Nupl2 | 0.423279 | 0.266285 | 0.291147 | 0.71142 | 1.21036 | 2.11144 |
| Pdgfrl | 0.589976 | 0.353822 | 0.452286 | 2.28529 | 0.885855 | 2.30967 |
| mt-Nd3 | 25.3414 | 40.3731 | 43.8435 | 108.878 | 122.012 | 76.3915 |
| Ttyh1 | 1.15112 | 1.11384 | 1.10966 | 4.33151 | 3.5231 | 4.88089 |
| Tppp3 | 1.00989 | 3.5546 | 2.67877 | 8.70722 | 10.5607 | 4.76535 |
| Pbx4 | 0.71985 | 0.500166 | 0.242453 | 1.23395 | 1.23226 | 1.45409 |
| Gas1 | 2.39414 | 3.83682 | 4.66033 | 13.8571 | 9.7266 | 10.9867 |
| Tram1l1 | 2.37029 | 2.12133 | 1.91084 | 6.15296 | 6.66962 | 5.46262 |
| Lamb1 | 0.27427 | 0.429484 | 0.181633 | 0.755446 | 1.02379 | 0.968565 |
| Lsm5 | 4.96261 | 4.36778 | 3.95911 | 13.6311 | 11.6651 | 12.5473 |
| Rac3 | 18.835 | 20.3281 | 16.3604 | 72.6906 | 70.6923 | 49.6571 |
| Csdc2 | 2.45759 | 2.5952 | 2.42547 | 11.2607 | 6.7217 | 7.27856 |
| Wnt5a | 0.431296 | 0.317505 | 0.254135 | 1.02624 | 1.12421 | 1.30197 |
| Wwtr1 | 1.15729 | 0.828778 | 1.21564 | 2.92158 | 2.21093 | 4.17 |
| Tmem130 | 3.23649 | 5.41478 | 4.54932 | 12.9213 | 16.7272 | 16.0111 |
| mt-Nd4 | 88.9855 | 110.964 | 109.756 | 329.842 | 276.238 | 290.586 |
| Pacsin1 | 0.744812 | 0.908943 | 0.557269 | 2.237 | 2.57802 | 2.47405 |
| Ncald | 0.52723 | 1.46793 | 1.10841 | 2.08696 | 5.30785 | 1.60321 |
| Spc25 | 3.97392 | 2.47642 | 4.37533 | 10.8379 | 9.27969 | 11.2768 |
| Scn1a | 0.0763518 | 0.0860233 | 0.0890053 | 0.581293 | 0.244854 | 0.276628 |
| Lrp4 | 0.777054 | 1.32088 | 1.31422 | 4.02206 | 3.25577 | 3.41004 |
| Tpm1 | 1.30607 | 1.2714 | 1.53472 | 4.54347 | 3.97369 | 4.09445 |
| Rps27 | 0.287697 | 0.730134 | 0.184692 | 0.856996 | 0.969749 | 0.546009 |
| Hba-a1 | 186.421 | 625.306 | 514.294 | 1483.8 | 1560.33 | 894.4 |
| Fxyd6 | 45.4758 | 45.8848 | 36.4257 | 166.79 | 132.351 | 125.791 |
| Tox3 | 2.05578 | 1.80534 | 2.32974 | 7.02809 | 6.16933 | 6.55134 |
| Cenpq | 0.880066 | 1.30148 | 1.96907 | 6.06385 | 3.33185 | 5.34467 |
| Myo6 | 0.297793 | 0.199586 | 0.170219 | 0.744136 | 0.636956 | 0.640652 |
| Zfp74 | 0.337121 | 0.419536 | 0.314606 | 0.918057 | 1.01753 | 2.07293 |
| Col5a2 | 0.092614 | 0.250327 | 0.212856 | 0.664754 | 0.42993 | 0.829158 |
| Slc9a3r2 | 1.38647 | 2.16238 | 1.29948 | 6.38026 | 5.17851 | 4.87576 |
| Fgf13 | 4.91986 | 4.72606 | 4.78531 | 16.6555 | 15.0923 | 11.7476 |
| Mlf1 | 1.32816 | 1.94808 | 2.6293 | 5.40807 | 6.60472 | 4.84973 |
| Rock2 | 0.797179 | 0.363586 | 0.837865 | 2.37531 | 2.07251 | 1.91169 |
| Polr2l | 4.95919 | 4.66441 | 5.58296 | 15.5131 | 15.484 | 18.2253 |
| 1110001J03Rik | 29.3954 | 37.4504 | 39.3522 | 123.264 | 121.739 | 100.06 |
| Skp1a | 28.2931 | 30.6329 | 30.2647 | 103.533 | 83.1889 | 83.0031 |
| Atp2b4 | 0.189628 | 0.168184 | 0.156881 | 0.653336 | 0.969733 | 0.806864 |
| Hmgb1 | 7.16091 | 5.28095 | 6.52967 | 17.3366 | 12.1943 | 14.7148 |
| Lhpp | 1.62434 | 1.47829 | 2.05399 | 7.06256 | 5.04514 | 5.27048 |
| Pnmal2 | 5.51252 | 5.66861 | 5.79599 | 15.8216 | 20.039 | 18.8468 |
| Tro | 1.44223 | 1.15793 | 1.31568 | 4.59186 | 4.71181 | 3.30668 |
| Prtn3 | 1.24751 | 0.962906 | 1.16024 | 4.79131 | 2.04568 | 3.11687 |
| Bex4 | 29.0108 | 26.5235 | 29.1168 | 91.6532 | 81.6623 | 86.4874 |
| Ankrd49 | 2.5801 | 2.9725 | 2.65193 | 11.2187 | 6.09116 | 7.04604 |
| Sox9 | 3.79169 | 5.88501 | 5.32578 | 16.0634 | 8.81214 | 19.5087 |
| Sp8 | 0.984683 | 1.10109 | 1.47378 | 4.21529 | 2.66596 | 3.33312 |
| Ercc6l2 | 0.266454 | 0.275946 | 0.256321 | 0.93359 | 0.69206 | 0.936998 |
| Bpgm | 2.59805 | 3.48406 | 3.99388 | 9.73221 | 9.33323 | 8.51598 |
| Chd3os | 11.8294 | 15.207 | 12.3935 | 36.5326 | 41.3206 | 31.6349 |
| Rdm1 | 2.07905 | 0.886771 | 2.41305 | 7.00011 | 6.38942 | 6.76115 |
| Spryd7 | 1.46846 | 1.11963 | 1.60605 | 3.52062 | 3.64569 | 3.65585 |
| Dzip1 | 1.81841 | 1.81816 | 1.43827 | 4.28674 | 5.97974 | 4.63982 |
| Flywch2 | 1.5411 | 3.51069 | 3.7309 | 8.48959 | 11.6675 | 6.35014 |
| Trp53i13 | 0.913915 | 1.16376 | 1.3626 | 7.01304 | 4.29285 | 4.58345 |
| Spef1 | 0.968325 | 1.15245 | 1.36044 | 3.26549 | 4.32548 | 3.13424 |
| Ndn | 23.501 | 25.8573 | 21.332 | 61.0139 | 64.3323 | 70.2178 |
| Akap17b | 0.270495 | 0.409394 | 0.398609 | 1.02161 | 1.31988 | 0.805344 |
| Efcab2 | 0.662908 | 0.348654 | 0.49955 | 1.31188 | 0.87013 | 2.30385 |
| 10-Sep | 0.919052 | 1.80256 | 2.15583 | 5.47262 | 4.31464 | 4.21432 |
| Snrpg | 1.24488 | 1.22003 | 0.73577 | 2.77973 | 2.63154 | 3.64682 |
| Cacng4 | 3.86412 | 3.93597 | 3.06978 | 11.7479 | 9.87829 | 13.0099 |
| Jam2 | 0.262481 | 0.416815 | 0.362543 | 1.02597 | 1.17525 | 0.973997 |
| Borcs7 | 3.90175 | 2.84287 | 4.72298 | 10.2945 | 12.9884 | 7.13135 |
| 1110004E09Rik | 7.7249 | 10.2115 | 8.2768 | 34.16 | 27.6052 | 24.232 |
| Mdk | 41.5815 | 41.4648 | 53.4067 | 168.526 | 141.399 | 142.759 |
| Camk2n1 | 1.83431 | 2.20776 | 1.78084 | 4.66877 | 7.29432 | 4.73544 |
| Gap43 | 93.2604 | 95.9178 | 81.9903 | 239.493 | 348.709 | 222.563 |
| Akap5 | 0.487496 | 0.686432 | 0.471157 | 1.54673 | 1.39863 | 1.39638 |
| Tmem107 | 1.56988 | 1.9285 | 1.02284 | 4.59288 | 3.94579 | 3.57994 |
| Gm12942 | 1.02392 | 1.1375 | 1.36635 | 4.80196 | 3.34026 | 3.45279 |
| Taf13 | 5.24843 | 2.73046 | 3.67752 | 11.1381 | 13.2326 | 8.32932 |
| Atp5g1 | 5.69316 | 5.53017 | 5.83365 | 17.4201 | 15.3185 | 17.1681 |
| Nxph1 | 2.30688 | 1.39255 | 2.50972 | 6.00302 | 5.97507 | 5.49314 |
| Rims2 | 0.693947 | 0.867618 | 0.831259 | 2.24717 | 3.35775 | 1.45 |
| Cntnap2 | 0.72515 | 1.14573 | 1.23516 | 2.31019 | 3.29032 | 2.87472 |
| 2610318N02Rik | 1.49984 | 2.65014 | 2.32128 | 6.7171 | 9.46452 | 6.21917 |
| Kcnk2 | 1.15468 | 0.641911 | 0.967337 | 2.2307 | 3.58991 | 2.31475 |
| Mfap4 | 3.9958 | 2.87881 | 3.41061 | 12.4099 | 6.60526 | 13.8132 |
| 1190002N15Rik | 0.664466 | 0.924228 | 0.564504 | 1.40584 | 1.64775 | 2.65083 |
| Pcdh17 | 0.68041 | 0.699358 | 0.584824 | 1.09001 | 2.48991 | 1.71481 |
| Tmeff2 | 0.368026 | 0.505126 | 0.467594 | 1.15403 | 1.37993 | 0.867605 |
| Pcgf1 | 0.386079 | 0.838543 | 0.522469 | 1.19896 | 1.21162 | 1.51334 |
| Pcdh19 | 0.528091 | 0.621488 | 0.646795 | 1.83515 | 1.25583 | 1.69263 |
| Thbs3 | 0.400904 | 0.297803 | 0.42763 | 1.13417 | 0.999378 | 1.27036 |
| Jtb | 6.38247 | 8.09364 | 7.06461 | 19.2258 | 18.5766 | 23.8582 |
| Psmg3 | 1.23256 | 0.97053 | 1.42784 | 3.79879 | 2.60394 | 3.22012 |
| Tmem179 | 2.18775 | 1.95328 | 0.858385 | 5.4936 | 4.43611 | 6.27662 |
| Morn2 | 9.81216 | 7.36864 | 8.50809 | 21.6128 | 25.8469 | 22.515 |
| Ube2c | 20.0731 | 14.9119 | 22.4971 | 66.4581 | 46.3221 | 52.5429 |
| Wdr60 | 0.527387 | 0.419299 | 0.850143 | 2.32068 | 1.55914 | 2.66593 |
| Fbxo44 | 0.896167 | 1.40681 | 1.55282 | 3.59644 | 3.92983 | 3.81424 |
| Ndufaf2 | 10.0645 | 11.3354 | 6.33548 | 30.5279 | 29.0899 | 28.8255 |

| | | | | | | |
|---|---|---|---|---|---|---|
| Nab1 | 1.46624 | 3.65726 | 2.31565 | 0.635377 | 0.801138 | 0.699152 |
| Alox5 | 0.719268 | 1.98263 | 1.7756 | 0.726049 | 0.557766 | 0.128604 |
| Map3k3 | 3.7933 | 3.94696 | 5.00846 | 1.06333 | 1.29337 | 0.675913 |
| Card9 | 5.08794 | 4.64388 | 7.11769 | 2.49839 | 0.89026 | 1.57115 |
| Aldh3b1 | 22.6181 | 12.128 | 28.7365 | 5.5764 | 11.972 | 3.47736 |
| Lyz2 | 1220.33 | 1075.7 | 1712.78 | 420.11 | 622.064 | 167.513 |
| Kcnq5 | 0.504293 | 0.724703 | 0.153731 | 0.125586 | 0.210229 | 0.236779 |
| Emb | 12.3435 | 29.8115 | 31.4052 | 5.44541 | 9.64579 | 6.20472 |
| Plpp6 | 3.95288 | 5.40803 | 5.97077 | 1.32907 | 0.740596 | 2.3323 |
| Gm13212 | 0.785295 | 0.561032 | 0.987066 | 0.174572 | 0.21567 | 0.0196593 |
| Btk | 4.03704 | 3.46897 | 4.99076 | 1.63999 | 0.881411 | 1.36015 |
| Hcls1 | 60.4449 | 45.4453 | 77.0058 | 17.5114 | 30.732 | 10.7282 |
| E2f1 | 9.26376 | 8.63831 | 11.3757 | 3.65402 | 2.26065 | 2.90344 |
| Trim14 | 1.51201 | 0.947312 | 1.88555 | 0.256988 | 0.670037 | 0.327419 |
| Fgd3 | 2.02541 | 1.25157 | 2.92899 | 0.585411 | 0.830203 | 0.397018 |
| Vav1 | 34.7948 | 27.0776 | 43.6917 | 13.1082 | 13.4119 | 5.94021 |
| Cysltr1 | 2.40663 | 1.70541 | 2.61785 | 1.06231 | 0.424303 | 0.241737 |
| Slco4a1 | 5.72608 | 1.4282 | 4.75729 | 1.20257 | 2.23255 | 0.496772 |
| Sbno2 | 4.597 | 4.97997 | 7.43544 | 2.11261 | 1.52196 | 2.32285 |
| Mertk | 1.73856 | 2.3412 | 2.90738 | 0.849895 | 1.08086 | 0.236453 |
| Rasa4 | 1.54059 | 2.4191 | 2.78433 | 0.94809 | 1.02442 | 0.28085 |
| Mvp | 5.48747 | 5.87012 | 7.74762 | 1.17811 | 2.79755 | 1.03367 |
| Esrra | 5.5022 | 6.58813 | 5.40925 | 2.93584 | 1.6785 | 1.82209 |
| Ell2 | 1.49003 | 3.85604 | 4.27512 | 0.558345 | 1.60815 | 0.601684 |
| St3gal1 | 3.25179 | 3.31302 | 1.9655 | 0.632705 | 0.589815 | 1.28152 |
| Ctsc | 168.329 | 113.502 | 204.607 | 35.4821 | 73.6675 | 15.5532 |
| Nckap1l | 16.4285 | 14.3695 | 22.1433 | 5.69858 | 6.65473 | 2.65086 |
| Arid5a | 1.11304 | 0.971475 | 1.64406 | 0.215372 | 0.394431 | 0.423128 |
| Adrbk2 | 1.21549 | 0.905224 | 1.29126 | 0.331742 | 0.465885 | 0.27214 |
| Angptl3 | 0.787742 | 0.792257 | 0.948687 | 0.316915 | 0.170713 | 0.123528 |
| Cdc6 | 1.70026 | 0.489312 | 1.65548 | 0.611314 | 0.253607 | 0.350774 |
| Slc2a1 | 6.78376 | 20.7278 | 17.4622 | 5.77573 | 4.18038 | 4.00538 |
| Slc2a9 | 0.160829 | 0.217256 | 0.328761 | 0.336982 | 0.146869 | 0.0210238 |
| Itpripl2 | 1.38515 | 0.613549 | 1.62948 | 0.362315 | 0.447224 | 0.103949 |
| Gch1 | 4.86503 | 9.64828 | 11.9234 | 2.2665 | 2.95923 | 1.68134 |
| Coch | 0.85793 | 1.04004 | 0.378311 | 0.175031 | 0 | 0.122391 |
| Ccnf | 4.64355 | 3.45759 | 5.00536 | 1.55331 | 0.628846 | 1.74121 |
| Ms4a6d | 64.062 | 96.9739 | 119.298 | 18.3267 | 43.0793 | 15.6956 |
| Tmem63a | 3.5867 | 2.15269 | 3.60172 | 1.28693 | 1.31959 | 0.439783 |
| Ptpn6 | 50.3245 | 41.4767 | 69.3398 | 18.9132 | 20.6361 | 8.95414 |
| Tmem104 | 4.437 | 6.59364 | 9.29903 | 2.63357 | 2.19781 | 1.92114 |
| Map3k5 | 0.547836 | 0.542412 | 0.733044 | 0.0135302 | 0.265682 | 0.137103 |
| Sema4c | 3.635 | 3.12124 | 3.72642 | 1.16629 | 1.73155 | 0.878993 |
| Aoah | 0.903625 | 1.73367 | 1.92719 | 0.454657 | 0.631324 | 0.145775 |
| Ltb | 31.1348 | 10.1934 | 31.4236 | 6.92616 | 10.2937 | 4.71187 |
| Irak2 | 1.01931 | 3.5234 | 3.86804 | 1.01144 | 1.15352 | 0.413684 |
| Osm | 2.0053 | 3.51042 | 4.723 | 1.12986 | 0.747196 | 1.0347 |
| Il2rg | 2.55496 | 5.34273 | 6.39062 | 0.685335 | 2.19612 | 0.61705 |
| Plekhm1 | 7.18395 | 2.17007 | 7.12844 | 2.11464 | 1.90839 | 0.698896 |
| Dock2 | 1.5883 | 1.76566 | 2.80432 | 0.627729 | 0.902565 | 0.265031 |
| Gk | 3.81354 | 3.1538 | 4.7131 | 0.729748 | 1.47789 | 0.746032 |
| 1600014C10Rik | 5.26985 | 12.1456 | 14.5108 | 2.96959 | 4.27997 | 3.48247 |
| Gpr18 | 19.1263 | 16.6946 | 25.6633 | 4.12042 | 12.1264 | 1.49729 |
| Samhd1 | 2.21271 | 3.06593 | 3.91427 | 0.555008 | 1.49152 | 0.697052 |
| Pcdha11 | 1.92104 | 0.966552 | 1.22872 | 0.632544 | 0.0968359 | 0.537352 |
| Igsf6 | 40.1514 | 32.2196 | 53.5171 | 15.35 | 12.2735 | 5.9056 |
| Antxr2 | 0.523851 | 2.63735 | 2.09891 | 0.263541 | 0.7019 | 0.501895 |
| Dock8 | 6.88381 | 5.02172 | 8.12941 | 1.33668 | 3.32579 | 0.813052 |
| Nfam1 | 11.4724 | 11.9684 | 16.4215 | 4.04954 | 5.89386 | 1.80328 |
| Plau | 4.86704 | 3.78891 | 6.50632 | 1.21643 | 2.09237 | 0.361356 |
| Ppp6r1 | 2.29183 | 1.97595 | 1.98752 | 0.64189 | 0.696168 | 0.499298 |
| Cd300lf | 2.283 | 6.69095 | 6.77788 | 1.88825 | 1.83397 | 0.593477 |
| Csf2rb2 | 0.960977 | 2.75053 | 2.16765 | 0.336482 | 0.947013 | 0.423639 |
| Itgb2 | 14.7701 | 22.8786 | 27.4854 | 8.15707 | 8.05045 | 3.36807 |
| AA474408 | 0.974018 | 0.662729 | 0.710351 | 0.307758 | 0.0913868 | 0.119592 |
| Cpd | 8.20849 | 12.4554 | 14.0271 | 3.25153 | 3.48523 | 1.94031 |
| Chrna3 | 0.763519 | 3.34406 | 3.04914 | 0.414248 | 0.988743 | 0.30934 |
| Gla | 1.50568 | 4.78704 | 4.71692 | 1.21335 | 1.39878 | 0.477585 |
| Ecm1 | 0.902777 | 4.46104 | 3.49634 | 1.60594 | 1.35899 | 0.303107 |
| Mapkapk2 | 14.1672 | 36.4108 | 35.8946 | 6.22965 | 9.72175 | 7.37026 |
| C3 | 2.08474 | 4.90489 | 4.71857 | 1.50567 | 1.38518 | 0.449219 |
| Gm6377 | 6.11662 | 5.27463 | 8.90804 | 0.676625 | 2.79463 | 0.879255 |
| Klra2 | 2.84202 | 0.897196 | 3.14154 | 0.877606 | 0.91617 | 0.0732624 |
| Hvcn1 | 2.50544 | 1.84974 | 2.92608 | 0.777441 | 0.741501 | 0.733322 |
| Cyth4 | 74.2151 | 40.4897 | 79.9677 | 17.6242 | 25.042 | 11.2876 |
| Trmt44 | 2.13993 | 1.89366 | 2.11506 | 0.312269 | 0.643287 | 0.635565 |
| Runx3 | 0.800965 | 0.620806 | 1.06506 | 0.155845 | 0.345856 | 0.233142 |
| 5730508B09Rik | 2.53411 | 6.106 | 5.76309 | 1.17225 | 1.52916 | 0.757378 |
| Metrnl | 14.5616 | 35.7677 | 39.333 | 7.04549 | 14.5269 | 5.32675 |
| Cdc42ep2 | 10.617 | 27.8135 | 25.3634 | 3.92331 | 8.63233 | 4.56085 |
| Aoc2 | 0.726208 | 0.719224 | 0.684156 | 0.156066 | 0.0929451 | 0.35695 |
| Rtl1 | 0.646873 | 0.393893 | 0.413766 | 0.0409258 | 0.241991 | 0.10768 |
| Il10 | 2.73208 | 3.88328 | 4.64586 | 0.105452 | 1.71332 | 0.522498 |
| Cfp | 36.8591 | 66.6827 | 83.2994 | 11.8794 | 26.978 | 6.09208 |
| Slc15a3 | 3.00366 | 6.45403 | 5.93866 | 1.56616 | 2.12311 | 1.36454 |
| Prkcg | 0.876683 | 0.951661 | 0.309507 | 0.241427 | 0.0484967 | 0.190921 |
| Pfkfb3 | 4.0538 | 6.1121 | 6.89253 | 2.20732 | 1.04401 | 1.31284 |
| Adamtsl3 | 1.01149 | 0.926625 | 0.748648 | 0.38503 | 0.0601652 | 0.234017 |
| Prex1 | 12.9965 | 6.0875 | 11.967 | 3.31531 | 3.1902 | 2.94174 |
| Zfp661 | 0.506477 | 1.00094 | 0.653631 | 0.290653 | 0.0682072 | 0.186552 |
| Stat6 | 5.22677 | 6.49054 | 8.51843 | 1.38287 | 2.85773 | 1.16481 |
| Arhgap26 | 0.84623 | 1.0378 | 1.45576 | 0.217051 | 0.337041 | 0.169016 |
| Trpm2 | 0.309308 | 0.175047 | 0.221715 | 0.0759188 | 0.0388532 | 0.0634169 |
| Slc2a6 | 2.37117 | 3.80857 | 3.89447 | 0.678159 | 1.42581 | 0.850032 |
| Slc35c1 | 1.61283 | 2.01885 | 1.99705 | 0.89269 | 0.313811 | 0.28753 |
| Csf3r | 7.05009 | 18.1069 | 19.4207 | 5.89182 | 4.84241 | 1.86615 |
| Tlr1 | 6.28084 | 7.87248 | 9.17658 | 0.967646 | 3.52095 | 1.45644 |
| Slc2a13 | 0.337643 | 0.490317 | 0.185004 | 0.0574532 | 0.029924 | 0.0313234 |
| Anxa6 | 58.442 | 50.6882 | 80.0041 | 13.2398 | 26.9332 | 10.7196 |
| Sgsh | 1.10226 | 0.686144 | 1.90138 | 0.265963 | 0.83633 | 0.393935 |
| Pglyrp1 | 1.20098 | 8.49529 | 6.9237 | 1.3496 | 1.14816 | 1.3761 |
| Apoc2 | 25.391 | 24.3767 | 37.1316 | 6.66562 | 9.49842 | 7.33491 |
| Inpp5b | 4.99805 | 3.46761 | 5.13738 | 1.59454 | 0.942549 | 1.21357 |
| Mocs3 | 4.4293 | 2.94815 | 2.68687 | 0.319683 | 1.22051 | 1.22713 |
| Rnf149 | 26.0068 | 28.7502 | 36.7483 | 9.59488 | 6.76616 | 5.34029 |
| Gna15 | 9.177 | 6.51285 | 8.61916 | 2.93398 | 1.77802 | 2.3003 |

| | | | | | | |
|---|---|---|---|---|---|---|
| Fermt2 | 1.55068 | 1.17214 | 0.822041 | 3.61328 | 3.05451 | 4.01916 |
| Dvl1 | 0.699484 | 0.772585 | 0.629758 | 1.97585 | 1.61405 | 2.68672 |
| Cenpm | 5.39091 | 3.7619 | 5.56569 | 21.9889 | 10.0185 | 16.8743 |
| Zfp760 | 0.289028 | 0.615645 | 0.586379 | 1.28011 | 1.60288 | 1.54147 |
| Fancl | 0.968848 | 1.50094 | 1.66724 | 3.5459 | 2.62503 | 2.43958 |
| Pttg1 | 2.61373 | 2.04207 | 3.72357 | 10.0199 | 7.85952 | 7.41961 |
| Dnajc12 | 0.701463 | 1.09556 | 1.13384 | 2.1559 | 2.9319 | 2.21366 |
| Cep70 | 0.578358 | 0.389885 | 0.372184 | 2.26428 | 1.3631 | 0.823472 |
| Ndufb4 | 8.38532 | 6.43739 | 8.95145 | 23.8478 | 20.9392 | 18.4229 |
| Abhd8 | 12.3981 | 17.3811 | 17.0133 | 52.2327 | 50.9104 | 48.6108 |
| Nol8 | 0.838713 | 0.971768 | 0.645055 | 2.07245 | 2.19818 | 2.54207 |
| Crcp | 5.94772 | 3.97758 | 5.1151 | 18.7247 | 12.9723 | 15.0377 |
| AW209491 | 2.67461 | 1.61687 | 2.65768 | 6.56728 | 5.48283 | 7.29155 |
| Plagl1 | 1.2099 | 1.49822 | 1.24504 | 2.94383 | 3.13267 | 3.63166 |
| Upf3a | 8.2448 | 6.98554 | 9.45636 | 24.2982 | 22.7733 | 29.9926 |
| Ybx3 | 1.09228 | 1.03155 | 1.16638 | 3.93008 | 4.14285 | 4.329 |
| S100a10 | 1.6735 | 5.9012 | 4.5282 | 10.168 | 13.721 | 13.0816 |
| E130114P18Rik | 2.50484 | 1.46235 | 1.42567 | 6.031 | 3.47262 | 5.94193 |
| Sdhaf4 | 26.8702 | 20.1779 | 18.7962 | 58.4538 | 56.9151 | 59 |
| Snapc2 | 3.56154 | 4.02588 | 5.407 | 16.8495 | 10.3169 | 13.3919 |
| Dcaf17 | 0.239483 | 0.288528 | 0.209433 | 0.86573 | 0.501036 | 0.647116 |
| Tmed1 | 6.88548 | 11.2591 | 10.843 | 25.7613 | 30.7383 | 26.3261 |
| mt-Nd6 | 151.448 | 151.56 | 175.394 | 374.056 | 395.386 | 415.226 |
| Arl6 | 2.23463 | 2.53396 | 2.09752 | 5.71153 | 6.21509 | 7.32819 |
| Phyhipl | 0.873447 | 0.885329 | 0.727976 | 2.42735 | 2.11391 | 2.11449 |
| Sulf1 | 0.27619 | 0.29617 | 0.284029 | 0.672755 | 0.475905 | 1.11693 |
| Trmt112 | 5.66697 | 11.0524 | 11.0093 | 20.1521 | 15.5546 | 15.7343 |
| Bok | 6.19512 | 6.27374 | 5.61908 | 21.6907 | 21.4548 | 20.7136 |
| 2310009B15Rik | 2.03881 | 1.37908 | 1.96882 | 3.44407 | 6.02483 | 4.71724 |
| Pcsk1n | 14.1898 | 14.3162 | 10.5778 | 32.3918 | 41.7952 | 37.1267 |
| Hnrnpa0 | 15.894 | 21.7023 | 21.8776 | 65.5266 | 46.5994 | 42.2432 |
| Rpp21 | 1.68996 | 1.60504 | 1.89962 | 5.33997 | 6.38351 | 3.19037 |
| Cyhr1 | 0.79494 | 1.34562 | 0.771622 | 3.25913 | 2.49956 | 2.84649 |
| Tmem216 | 1.22829 | 2.06144 | 1.56578 | 4.09394 | 4.52162 | 6.29459 |
| Psat1 | 4.05334 | 5.58335 | 6.46802 | 12.7142 | 11.7246 | 21.3711 |
| Mrps6 | 10.8264 | 9.15649 | 10.4454 | 39.3108 | 25.4075 | 30.3426 |
| Chd5 | 0.485175 | 0.801751 | 0.369352 | 1.80477 | 1.36017 | 1.70709 |
| Prcl | 0.580778 | 0.375578 | 0.75875 | 2.1582 | 1.17006 | 1.52581 |
| Zfp580 | 5.78118 | 9.9888 | 11.5551 | 30.2869 | 25.9593 | 24.279 |
| Tmem59l | 4.92447 | 5.53119 | 3.16636 | 10.1382 | 16.7033 | 13.0842 |
| Xrcc5 | 2.98785 | 2.57867 | 1.94574 | 6.98508 | 5.66339 | 7.2197 |
| H2afx | 61.1128 | 57.009 | 68.876 | 253.842 | 141.893 | 173.616 |
| Med19 | 8.44178 | 7.05392 | 9.25597 | 18.217 | 16.419 | 24.9859 |
| Lgr4 | 0.946462 | 0.610176 | 1.20801 | 3.35305 | 2.21754 | 2.0359 |
| Ift27 | 11.382 | 17.0332 | 19.7097 | 51.5837 | 48.403 | 35.8251 |
| Bex1 | 20.886 | 22.3924 | 17.8974 | 55.9512 | 62.0591 | 57.3758 |
| Anapc13 | 3.93616 | 3.59004 | 3.59261 | 11.3587 | 8.94157 | 9.67486 |
| Ddx25 | 1.93599 | 1.17005 | 1.46715 | 4.52844 | 3.19389 | 4.29671 |
| Id3 | 5.55626 | 13.3823 | 13.1991 | 23.8348 | 29.4329 | 32.0364 |
| Rragd | 1.26166 | 1.32781 | 1.25762 | 3.34037 | 3.27887 | 3.62701 |
| Gpx1 | 49.8678 | 47.495 | 66.8665 | 187.066 | 152.972 | 87.734 |
| Fkbp3 | 94.9884 | 107.003 | 105.242 | 273.73 | 239.367 | 264.941 |
| Rnd3 | 3.57125 | 5.3656 | 3.7923 | 8.97382 | 10.8455 | 10.5534 |
| Setd6 | 1.83338 | 1.72341 | 1.84521 | 5.46155 | 4.65354 | 7.58202 |
| Zbtb20 | 1.89342 | 1.64339 | 1.85842 | 4.34253 | 3.83193 | 5.55168 |
| Asphd2 | 0.790888 | 0.856875 | 0.790879 | 1.78198 | 2.81813 | 2.63987 |
| Rcn1 | 2.52217 | 4.98411 | 4.2739 | 10.4447 | 9.2915 | 10.8511 |
| Sall1 | 0.750278 | 0.815086 | 0.816575 | 2.1955 | 1.76484 | 2.90022 |
| Gcsh | 7.64984 | 8.44098 | 8.52716 | 24.5478 | 19.28 | 27.9854 |
| Zfp946 | 2.12256 | 0.849334 | 1.30751 | 3.23706 | 2.24079 | 4.0414 |
| Ndufaf4 | 2.42629 | 1.92038 | 2.17237 | 5.50686 | 4.84483 | 5.09067 |
| Sptbn1 | 1.47564 | 1.5789 | 1.22253 | 4.14865 | 3.56049 | 3.60186 |
| Mphosph8 | 2.82957 | 1.82643 | 1.47278 | 4.05175 | 3.74039 | 6.10945 |
| Tceal1 | 1.69228 | 1.15421 | 1.30867 | 2.31037 | 3.20892 | 4.04001 |
| Gadd45g | 10.7535 | 21.0175 | 18.6006 | 59.6857 | 37.3233 | 62.6757 |
| Mrps11 | 2.14347 | 1.8939 | 2.15698 | 5.7835 | 4.53456 | 4.41008 |
| Fabp3 | 28.04 | 21.1012 | 16.7833 | 55.7156 | 70.8524 | 47.9179 |
| Usmg5 | 382.934 | 360.015 | 427.658 | 1052.73 | 1062.09 | 1018.61 |
| Krt10 | 3.67605 | 1.62552 | 3.92446 | 9.83462 | 10.1837 | 8.96057 |
| Zfp30 | 1.00718 | 0.6337 | 0.851466 | 2.2552 | 2.51347 | 1.96409 |
| Hsp90aa1 | 26.6934 | 29.0045 | 29.2485 | 74.4116 | 64.993 | 61.7933 |
| Esf1 | 1.92042 | 2.1921 | 1.74466 | 4.64472 | 4.06547 | 6.374 |
| Tmsb10 | 211.383 | 197.926 | 210.692 | 631.763 | 613.12 | 509.444 |
| Atraid | 5.33999 | 3.25051 | 4.69138 | 12.6657 | 11.5007 | 8.58124 |
| 2700094K13Rik | 16.527 | 14.8403 | 16.588 | 43.8386 | 37.3203 | 41.1666 |
| Gabra2 | 1.09867 | 0.645741 | 0.72578 | 1.1856 | 2.0486 | 2.59374 |
| Dpy30 | 38.7986 | 39.6879 | 48.1154 | 120.809 | 114.351 | 92.4047 |
| Cox6c | 4.85268 | 4.47235 | 4.11484 | 12.5142 | 10.6857 | 11.023 |
| Tfcp2 | 1.47414 | 1.43585 | 1.09814 | 4.00732 | 3.42741 | 3.24362 |
| 1110032A03Rik | 0.660172 | 0.671843 | 0.501927 | 1.42372 | 2.10533 | 1.28813 |
| Glo1 | 29.5671 | 38.1039 | 40.4702 | 97.645 | 94.3261 | 82.8322 |
| Lmo3 | 2.7523 | 3.64395 | 1.99139 | 11.248 | 4.49482 | 5.60932 |
| Ccdc85b | 17.8848 | 17.2226 | 22.161 | 50.2874 | 41.3794 | 44.2237 |
| Fam210b | 5.84075 | 4.53267 | 6.38981 | 17.249 | 6.5532 | 20.065 |
| H1f0 | 69.032 | 68.0763 | 79.0684 | 243.695 | 153.565 | 171.511 |
| Ndufa3 | 3.24131 | 3.48539 | 2.05053 | 3.60082 | 3.94971 | 2.93318 |
| Rps29 | 58.1207 | 44.8632 | 57.7382 | 135.206 | 137.479 | 101.26 |
| Cox6a1 | 180.781 | 162.902 | 201.313 | 595.592 | 508.49 | 400.685 |
| Lsm6 | 5.40672 | 3.47234 | 5.78605 | 13.3108 | 9.06279 | 12.4622 |
| Nacc2 | 0.523453 | 0.410562 | 0.641559 | 1.16472 | 1.62915 | 1.40799 |
| Abhd14a | 1.74012 | 2.21366 | 1.70075 | 5.95709 | 6.12585 | 5.63528 |
| Pde1b | 3.06219 | 3.53284 | 4.11081 | 9.23111 | 10.5231 | 11.2247 |
| Zeb1 | 0.886076 | 1.20201 | 1.23592 | 2.66049 | 2.37386 | 3.4607 |
| Haus7 | 6.86293 | 9.18116 | 10.1507 | 23.8928 | 24.1281 | 20.9554 |
| Bex2 | 152.786 | 175.155 | 158.452 | 448.699 | 436.662 | 437.862 |
| Zfp277 | 1.76443 | 4.47241 | 3.20778 | 7.66564 | 8.27302 | 7.33557 |
| Atp5e | 14.4967 | 11.1802 | 12.1015 | 34.6607 | 26.4719 | 24.6509 |
| Tctex1d2 | 1.8356 | 2.04347 | 1.74397 | 4.30431 | 4.72198 | 4.07599 |
| Polr2i | 1.43881 | 0.765557 | 1.26416 | 2.14255 | 1.68503 | 2.4692 |
| Smim18 | 11.4371 | 10.5273 | 7.56695 | 26.1686 | 22.998 | 25.2483 |
| Ica1 | 0.804581 | 0.852226 | 1.19772 | 3.52334 | 2.93025 | 2.89118 |
| Fstl5 | 0.348269 | 0.616876 | 0.533935 | 1.51398 | 0.97715 | 1.14098 |
| Tox2 | 2.63598 | 2.82971 | 3.44937 | 8.57775 | 11.5197 | 6.2257 |
| Shisa4 | 3.91119 | 4.09956 | 4.98876 | 15.7843 | 11.0113 | 11.7823 |
| Psmb4 | 4.52098 | 4.79821 | 4.63925 | 12.7706 | 12.2704 | 8.76179 |
| Rps27rt | 14.5171 | 13.832 | 14.1502 | 48.7432 | 38.6686 | 34.6974 |

| | | | | | | |
|---|---|---|---|---|---|---|
| Tlr13 | 4.5886 | 6.10765 | 6.99993 | 1.99704 | 1.53871 | 0.44812 |
| Inf2 | 1.54986 | 1.33019 | 1.8458 | 0.534943 | 0.447157 | 0.285354 |
| Dok3 | 53.5448 | 35.1396 | 66.0516 | 13.0493 | 23.8838 | 7.30935 |
| Nlrp3 | 1.19485 | 6.81215 | 4.92987 | 1.04092 | 1.58709 | 0.753565 |
| Arhgap24 | 0.493121 | 1.03651 | 1.13298 | 0.21468 | 0.264973 | 0.12464 |
| Pilra | 6.68952 | 9.6897 | 11.2128 | 2.07246 | 4.00217 | 1.37504 |
| Tor4a | 6.03606 | 4.79806 | 7.17449 | 2.67793 | 1.3518 | 0.508808 |
| Cntnap5c | 0.265081 | 0.0319365 | 0.360254 | 0.0248817 | 0.0352012 | 0.00762666 |
| Dhdh | 1.63331 | 0.903233 | 1.45779 | 0.608418 | 0.246033 | 0.169907 |
| Acp5 | 123.08 | 48.2971 | 135.123 | 27.9698 | 33.3202 | 17.6587 |
| Usp25 | 7.82804 | 10.9137 | 12.5981 | 2.96562 | 2.14312 | 1.67797 |
| Myo7a | 1.47569 | 2.17095 | 2.63472 | 0.199534 | 0.828046 | 0.534021 |
| Ccr1 | 1.82641 | 10.2447 | 8.00021 | 1.98733 | 1.4036 | 0.889005 |
| Raver2 | 6.93364 | 2.31914 | 6.16954 | 1.48056 | 1.21565 | 1.10952 |
| Rgs14 | 1.77527 | 3.45118 | 3.94336 | 1.15551 | 0.504032 | 0.773676 |
| Tlr8 | 3.06952 | 3.75913 | 4.14272 | 0.688053 | 0.439232 | 0.544644 |
| Wfdc21 | 46.003 | 273.769 | 232.096 | 33.4898 | 49.7291 | 45.816 |
| Pstpip1 | 0.413396 | 1.1196 | 0.975425 | 0.235893 | 0.115536 | 0.061356 |
| Gm5431 | 1.08125 | 1.09755 | 2.20354 | 0.330437 | 0.499694 | 0.0684469 |
| Ttc7 | 0.79271 | 0.816423 | 1.08022 | 0.219712 | 0.234959 | 0.176824 |
| Ugt1a7c | 1.63519 | 1.55439 | 2.29399 | 0.689613 | 0.504861 | 0.443935 |
| Rhbdf2 | 1.06195 | 2.98634 | 2.92703 | 0.467248 | 0.72905 | 0.656475 |
| Pkn1 | 5.93437 | 5.72205 | 6.55126 | 1.51867 | 1.64353 | 1.64419 |
| Tgm2 | 2.50702 | 2.93642 | 3.71932 | 0.609809 | 1.15529 | 0.760096 |
| Manba | 0.842781 | 0.818313 | 1.16132 | 0.398689 | 0.28053 | 0.0462245 |
| Naaa | 9.83339 | 1.95517 | 8.85874 | 2.03942 | 1.18303 | 1.2551 |
| Gsn | 22.398 | 28.2238 | 33.1717 | 5.10064 | 10.8234 | 5.65799 |
| Aff1 | 1.39361 | 2.13893 | 2.76985 | 0.391523 | 0.709044 | 0.340858 |
| Rassf3 | 2.31514 | 3.63137 | 4.15583 | 0.997312 | 0.697657 | 0.599006 |
| Bcl3 | 4.07835 | 4.0437 | 6.80385 | 0.976011 | 1.95199 | 0.660748 |
| Ccdc71l | 1.7669 | 4.76332 | 3.85639 | 0.198399 | 1.26614 | 0.5276 |
| Abcc8 | 2.95467 | 2.13761 | 1.12706 | 0.2117 | 0.282101 | 0.690292 |
| Kcnn4 | 0.044407 | 0.539389 | 0.513338 | 0.00857281 | 0.164689 | 0.0778829 |
| Stat5a | 0.493593 | 0.268072 | 0.777905 | 0.0337957 | 0.203029 | 0.126924 |
| Tgfbi | 151.41 | 110.833 | 197.917 | 30.6795 | 62.4544 | 14.4556 |
| Map4k1 | 2.06643 | 1.08998 | 3.55129 | 0.621387 | 0.405373 | 0.198226 |
| Fpr1 | 138.559 | 195.381 | 254.274 | 36.158 | 67.8005 | 19.0948 |
| Dhx58 | 1.64152 | 2.81732 | 4.5336 | 0.92512 | 1.24839 | 0.138554 |
| BC117090 | 3.06128 | 25.8401 | 19.398 | 5.01504 | 3.71849 | 0.463818 |
| Fosl2 | 0.862282 | 1.49505 | 1.4835 | 0.232962 | 0.277465 | 0.285922 |
| Ntsr1 | 4.93206 | 5.65192 | 1.04084 | 0.439957 | 1.11818 | 1.17791 |
| Slc37a2 | 1.62355 | 2.30934 | 3.51418 | 0.360724 | 0.751072 | 0.425216 |
| Xdh | 1.58162 | 2.72564 | 2.4581 | 0.28391 | 1.05123 | 0.0718837 |
| Myh9 | 8.51177 | 5.05707 | 9.17768 | 1.69151 | 2.51494 | 1.37184 |
| Serpina3g | 2.37481 | 7.50892 | 7.1086 | 0.276192 | 2.19153 | 1.05402 |
| A630033H20Rik | 3.93614 | 2.29202 | 4.08981 | 0.175398 | 1.33454 | 0.316358 |
| Gpnmb | 0.298823 | 1.23465 | 1.10497 | 0.390898 | 0.0483584 | 0.0852342 |
| 6330416G13Rik | 4.53668 | 7.42669 | 8.66276 | 1.51809 | 1.86705 | 1.30444 |
| Adora2a | 1.00092 | 11.4582 | 8.66953 | 2.20353 | 1.93786 | 1.06845 |
| Ero1l | 5.83839 | 18.4555 | 17.719 | 2.85995 | 2.3106 | 2.6297 |
| Kcnf1 | 0.766883 | 0.168678 | 0.710034 | 0.175083 | 0.0690482 | 0.11628 |
| Pvrl4 | 0.590949 | 1.03506 | 0.643831 | 0.0763958 | 0.371637 | 0.0397642 |
| Lacc1 | 13.879 | 10.8363 | 17.8091 | 2.36495 | 4.12429 | 2.15628 |
| Hspa1b | 0.266528 | 3.36436 | 2.36482 | 0.538867 | 0.51445 | 0.256748 |
| Oas2 | 1.55329 | 3.69467 | 4.00503 | 0.480581 | 1.36051 | 0.323153 |
| Pik3ap1 | 27.8011 | 28.3474 | 38.0017 | 8.33999 | 9.4937 | 3.81074 |
| Rhoq | 5.81855 | 6.64986 | 7.28933 | 1.38599 | 1.11602 | 1.31822 |
| Rras | 21.5822 | 32.6525 | 45.1561 | 6.30563 | 8.15056 | 7.04137 |
| Ikbke | 1.43948 | 2.08306 | 2.49234 | 0.359715 | 0.680123 | 0.189958 |
| Pstpip2 | 1.40357 | 5.01845 | 4.07138 | 0.599972 | 1.24455 | 0.200301 |
| Pecam1 | 2.6889 | 1.07957 | 2.65144 | 0.804475 | 0.494699 | 0.0848608 |
| Zmynd15 | 0.729283 | 0.748085 | 1.39263 | 0.195216 | 0.298867 | 0.0469383 |
| Csf2rb | 6.59495 | 7.2997 | 10.836 | 2.189 | 2.11159 | 1.19959 |
| Syk | 4.78278 | 3.88997 | 5.96647 | 1.14106 | 1.18134 | 0.801658 |
| Arhgap22 | 0.746678 | 0.877011 | 1.43246 | 0 | 0.548608 | 0.193582 |
| Pirb | 1.22506 | 1.83289 | 2.27493 | 0.235166 | 0.493272 | 0.205551 |
| Jade2 | 0.369117 | 0.122509 | 0.429096 | 0.128203 | 0.0456442 | 0 |
| Uba7 | 0.797738 | 0.938173 | 1.2431 | 0.393923 | 0.0904746 | 0.142093 |
| Rinl | 2.54825 | 2.30427 | 3.01614 | 0.905003 | 0.310673 | 0.453327 |
| B430306N03Rik | 0.178031 | 1.47459 | 0.917672 | 0.175101 | 0.306334 | 0.0611051 |
| Pygl | 4.8614 | 11.5777 | 11.7731 | 2.79012 | 1.99059 | 1.27375 |
| Slc45a3 | 0.66388 | 0.803567 | 0.24212 | 0.163565 | 0.0540383 | 0.12513 |
| Plekho2 | 9.51396 | 8.95238 | 13.0627 | 1.53675 | 1.982 | 2.08612 |
| Zc3h12d | 1.76639 | 2.16754 | 2.99642 | 0.440866 | 0.785695 | 0.0198623 |
| C130050O18Rik | 2.14131 | 2.34607 | 3.46511 | 0.561989 | 0.469234 | 0.495027 |
| Cnksr3 | 7.58513 | 2.11888 | 5.30396 | 0.978374 | 0.678103 | 0.956213 |
| Gm5150 | 3.48405 | 2.38381 | 5.37849 | 0.587265 | 1.74088 | 0.32663 |
| Clec4e | 1.28368 | 7.67389 | 7.22045 | 0.796173 | 1.34719 | 0.673272 |
| Ulbp1 | 2.45057 | 1.62258 | 3.09981 | 0.170634 | 0.739688 | 0.442766 |
| Itgal | 7.25789 | 5.57793 | 8.9601 | 1.12012 | 2.6703 | 0.772667 |
| Pcdhga1 | 0.987226 | 0.73756 | 0.524155 | 0.253746 | 0.0551278 | 0.231052 |
| Stx11 | 7.12322 | 11.8714 | 13.8649 | 1.92625 | 2.00319 | 2.05522 |
| Grm1 | 0.56314 | 0.294262 | 0.18792 | 0.0402749 | 0.0744924 | 0.0469124 |
| Tnip3 | 0.25404 | 1.99385 | 1.86093 | 0.222857 | 0.258924 | 0.2164 |
| Cebpb | 18.0564 | 92.4362 | 79.752 | 8.41904 | 11.6523 | 17.4076 |
| Pik3r5 | 7.57127 | 7.123 | 11.5937 | 2.00137 | 2.41507 | 0.722902 |
| Pcdhga8 | 1.62593 | 0.670721 | 1.37023 | 0.219429 | 0.161588 | 0.549364 |
| Ikzf1 | 6.25353 | 3.82393 | 6.73155 | 1.13497 | 1.01662 | 0.913301 |
| H6pd | 1.22488 | 3.05014 | 2.816 | 0.858664 | 0.25699 | 0.320899 |
| Tgm1 | 0.781728 | 0.817319 | 1.05687 | 0.327261 | 0.111509 | 0.0264122 |
| Pik3cg | 8.78268 | 5.04816 | 9.95201 | 1.39117 | 1.65344 | 1.21168 |
| Irak3 | 31.2373 | 22.3791 | 38.2585 | 5.87636 | 7.90973 | 2.65003 |
| Ptprj | 6.15061 | 4.90662 | 7.73874 | 0.903223 | 1.38368 | 0.626322 |
| Tmem8 | 0.422584 | 3.65508 | 3.07965 | 0.701188 | 0.394166 | 0.203615 |
| Slc8b1 | 3.19419 | 3.53197 | 4.80932 | 0.654887 | 1.03335 | 0.454654 |
| Mcoln2 | 0.259271 | 1.15399 | 1.8388 | 0.408714 | 0.0560075 | 0.231528 |
| Gbp4 | 0.0461228 | 0.928669 | 0.837902 | 0.05554 | 0.191097 | 0.0113748 |
| Phf11b | 9.25468 | 19.4223 | 22.4996 | 1.33088 | 5.41363 | 1.22352 |
| D330045A20Rik | 5.69472 | 3.69136 | 5.92499 | 0.272178 | 1.61011 | 0.394118 |
| Rab11fip1 | 0.515888 | 1.61934 | 1.47793 | 0.150441 | 0.320986 | 0.115991 |
| Tgfb1 | 1.62574 | 1.09018 | 1.82091 | 0.320615 | 0.257496 | 0.260701 |
| Plbd1 | 18.5381 | 22.7531 | 30.2066 | 4.40064 | 5.25262 | 2.15812 |
| 5031414D18Rik | 0.275059 | 0.386617 | 0.182136 | 0.203474 | 0 | 0 |
| Slc35d2 | 1.19431 | 3.34689 | 2.50193 | 0.319442 | 0.36506 | 0.381255 |
| Cd44 | 3.42509 | 9.36293 | 8.72245 | 0.757022 | 2.03042 | 0.427887 |
| Ptk2b | 2.55405 | 2.91289 | 3.45401 | 0.500269 | 0.606419 | 0.352763 |

| | | | | | | |
|---|---|---|---|---|---|---|
| Rab3c | 2.51799 | 2.38503 | 1.81005 | 4.66372 | 6.63218 | 5.41142 |
| Lxn | 4.20913 | 5.8681 | 7.18041 | 15.2094 | 13.8955 | 13.1611 |
| Immp1l | 20.9796 | 19.8281 | 20.5137 | 58.4766 | 46.6808 | 42.1162 |
| Upf3b | 3.01749 | 3.30266 | 2.31374 | 7.93578 | 8.03111 | 5.57439 |
| Ift74 | 1.93851 | 3.26692 | 2.47137 | 4.38929 | 4.00897 | 8.76763 |
| Polr3h | 5.96921 | 5.3273 | 6.00466 | 20.3716 | 11.0083 | 18.3511 |
| Blcap | 9.86799 | 11.0941 | 11.1013 | 29.9657 | 29.5733 | 28.3023 |
| Prr3 | 2.02057 | 3.36047 | 2.26193 | 7.66642 | 6.31428 | 7.44572 |
| Sox2 | 8.35591 | 16.1163 | 13.0357 | 30.6994 | 32.9629 | 34.3096 |
| Uqcrh | 41.6997 | 36.1298 | 43.2829 | 93.2902 | 97.9357 | 101.983 |
| Pfdn4 | 4.72798 | 3.15634 | 3.01652 | 9.42807 | 10.2055 | 6.81202 |
| Hscb | 3.75673 | 2.93634 | 3.83545 | 10.4957 | 8.30841 | 6.00955 |
| Rps21 | 0.417875 | 0.452949 | 0.799544 | 0.322897 | 0.602933 | 0.3708 |
| Romo1 | 72.2268 | 79.5468 | 94.041 | 276.661 | 266.297 | 225.595 |
| Schip1 | 16.1917 | 14.5363 | 11.9709 | 48.8839 | 31.3953 | 39.9761 |
| Rpl36 | 327.097 | 325.285 | 367.61 | 983.023 | 1021.18 | 734.811 |
| Ctnnal1 | 0.313234 | 0.184794 | 0.234599 | 0.567868 | 0.629058 | 0.710749 |
| Gad2 | 1.96795 | 1.95568 | 1.65218 | 7.30699 | 2.89128 | 4.18562 |
| Gnaz | 2.36431 | 2.91274 | 1.78812 | 5.5114 | 7.63323 | 5.73651 |
| Gm1673 | 60.4383 | 70.0716 | 62.4383 | 197.747 | 161.341 | 185.182 |
| Arxes2 | 12.6229 | 15.2817 | 8.59926 | 29.3048 | 22.3573 | 31.1161 |
| Map9 | 1.74082 | 1.44887 | 0.995665 | 3.29397 | 3.52591 | 4.25276 |
| Aif1l | 1.41187 | 1.90771 | 1.5006 | 3.94217 | 3.90687 | 5.57261 |
| Rabggtb | 2.37561 | 1.16739 | 2.01392 | 5.83784 | 4.38228 | 5.26972 |
| Caly | 4.64276 | 4.32165 | 2.34554 | 11.1113 | 11.3058 | 10.3615 |
| Hmgn5 | 3.94168 | 4.57888 | 4.89086 | 11.4188 | 6.8269 | 12.6987 |
| Srek1ip1 | 0.76762 | 0.964664 | 0.971222 | 1.84076 | 1.1815 | 2.7148 |
| Cdo1 | 9.60424 | 9.22625 | 11.7871 | 26.7126 | 22.1294 | 25.0228 |
| Atp5j | 3.34355 | 2.38389 | 2.93635 | 7.85567 | 7.5281 | 5.96224 |
| Rpl23 | 23.8193 | 24.2472 | 25.4598 | 57.4071 | 62.2289 | 60.3875 |
| Cxx1c | 15.1942 | 11.751 | 22.653 | 46.2665 | 61.9425 | 32.223 |
| Cetn2 | 12.5044 | 14.1747 | 13.5401 | 31.5565 | 30.9933 | 32.6381 |
| Bola1 | 17.1151 | 13.8657 | 16.4958 | 53.1687 | 39.5073 | 44.5195 |
| Marcks | 56.4546 | 57.9757 | 61.7627 | 145.819 | 128.661 | 110.863 |
| mt-Co1 | 442.195 | 459.447 | 532.257 | 1187.56 | 926.085 | 1135.75 |
| Tasp1 | 0.634108 | 0.517314 | 0.419945 | 1.52513 | 1.80847 | 1.55054 |
| Psmg4 | 46.0609 | 53.2233 | 54.7772 | 123.5 | 162.757 | 108.649 |
| Cpe | 39.2312 | 46.9667 | 40.3589 | 121.278 | 95.7427 | 97.6476 |
| Mrpl47 | 1.52868 | 1.12142 | 1.39175 | 2.93406 | 2.51963 | 2.44423 |
| Tbca | 140.527 | 144.084 | 174.861 | 441.232 | 355.937 | 327.256 |
| Map1a | 0.943265 | 1.05338 | 1.08566 | 2.59547 | 2.68454 | 2.47055 |
| Tamm41 | 0.32571 | 0.231351 | 0.225348 | 0.841713 | 0.293363 | 0.615061 |
| Rpa3 | 5.23596 | 4.43378 | 6.29964 | 11.3694 | 7.8184 | 12.3567 |
| Kcnh7 | 0.295953 | 0.252729 | 0.170279 | 0.516592 | 0.426208 | 0.527351 |
| Trip11 | 0.527584 | 0.680785 | 0.56782 | 1.75683 | 1.55852 | 2.01853 |
| Ccl27a | 1.07204 | 1.62076 | 1.79148 | 4.96588 | 4.6299 | 4.78634 |
| Polr2f | 53.7488 | 49.5975 | 63.804 | 180.807 | 141.859 | 134.954 |
| Phf5a | 30.538 | 30.7296 | 34.7271 | 99.1398 | 83.0073 | 77.8731 |
| Cenpa | 3.07038 | 2.53199 | 3.59913 | 9.51461 | 5.18048 | 5.51018 |
| Akap9 | 1.82771 | 1.71441 | 1.78384 | 4.46024 | 3.77337 | 4.2221 |
| Tspan12 | 1.90619 | 2.03457 | 2.1934 | 5.47065 | 3.41334 | 5.07765 |
| Hacd1 | 2.48448 | 2.00897 | 2.44349 | 6.06454 | 4.25001 | 6.32282 |
| Acy1 | 0.66344 | 0.866045 | 0.869721 | 1.82654 | 1.05546 | 1.79203 |
| Fam118b | 1.55239 | 1.3438 | 1.69633 | 4.0542 | 3.76025 | 3.37726 |
| Syngr3 | 10.1648 | 12.6457 | 8.43647 | 29.1384 | 27.2571 | 26.542 |
| Stk19 | 0.867874 | 1.17208 | 0.978616 | 3.01011 | 2.92446 | 2.21175 |
| Elavl4 | 4.9609 | 4.60381 | 5.59964 | 16.6981 | 10.5483 | 8.67775 |
| Pik3ca | 0.925711 | 0.614477 | 0.496747 | 1.67124 | 1.36701 | 2.02516 |
| Cdca3 | 3.59795 | 4.23554 | 4.94623 | 13.9694 | 8.21976 | 8.72677 |
| Ccdc28b | 0.287909 | 0.494745 | 0.565591 | 1.66429 | 1.17505 | 1.00962 |
| Ccdc90b | 7.18629 | 10.8416 | 10.0591 | 19.6408 | 19.31 | 21.9157 |
| Cox7c | 12.5688 | 16.5767 | 13.4398 | 32.0303 | 31.3889 | 32.4802 |
| Efnb3 | 4.21468 | 2.87097 | 3.21805 | 8.33152 | 8.19346 | 10.386 |
| 1810043G02Rik | 0.563202 | 0.451492 | 0.849615 | 1.42669 | 1.76954 | 1.40635 |
| Snrpd2 | 11.185 | 10.1142 | 11.7497 | 29.0822 | 22.6158 | 26.5351 |
| Sema5a | 0.91393 | 0.576449 | 0.491954 | 1.66561 | 1.62879 | 1.58892 |
| Acot13 | 21.7373 | 13.8681 | 24.615 | 45.4007 | 46.9421 | 50.4025 |
| Coq9 | 1.6528 | 1.86031 | 1.87392 | 5.82818 | 5.77947 | 3.66212 |
| Prkrir | 2.41998 | 2.50358 | 2.25472 | 6.3744 | 4.57165 | 5.71311 |
| Ndufa4 | 32.2167 | 26.1974 | 30.4966 | 74.4347 | 64.8767 | 79.9514 |
| Mis18bp1 | 0.958972 | 0.848335 | 1.02641 | 2.21039 | 1.39082 | 1.49018 |
| Atpif1 | 44.9297 | 41.0457 | 37.871 | 110.185 | 108.866 | 90.6563 |
| Sgf29 | 0.430281 | 0.384659 | 0.633331 | 1.58174 | 1.80043 | 1.38433 |
| Eci1 | 5.68661 | 10.8087 | 9.23532 | 16.5273 | 26.4578 | 16.9078 |
| Mcee | 5.61705 | 5.79568 | 8.96115 | 15.9465 | 18.0016 | 13.6136 |
| Cox20 | 2.69089 | 2.01545 | 2.12446 | 4.95726 | 4.83898 | 5.59283 |
| Ndufa1 | 152.484 | 173.26 | 223.731 | 457.139 | 457.394 | 399.566 |
| Cnpy2 | 26.1094 | 35.9615 | 31.4909 | 88.1229 | 72.948 | 74.2407 |
| 1500009C09Rik | 6.96966 | 7.2072 | 4.75778 | 14.5665 | 12.5996 | 18.7106 |
| Nudt16l1 | 18.026 | 14.131 | 22.7348 | 58.7042 | 38.3347 | 46.329 |
| Lzic | 5.96969 | 7.24637 | 7.27299 | 20.8135 | 14.09 | 15.3681 |
| Bphl | 8.51351 | 7.84698 | 9.92766 | 17.2965 | 20.2269 | 23.1977 |
| Sarnp | 37.1973 | 30.0925 | 29.6799 | 91.1459 | 74.849 | 86.4385 |
| Adam19 | 1.07319 | 1.20722 | 0.852594 | 0.492674 | 0.656571 | 0.869391 |
| Dpp7 | 1.89154 | 2.15553 | 2.27554 | 0.771467 | 1.844 | 0.701 |
| Frmd4b | 3.22842 | 2.24378 | 2.30162 | 1.34415 | 1.67691 | 1.68083 |
| Pfkfb4 | 2.77024 | 2.10685 | 2.24681 | 1.19357 | 1.78875 | 1.4459 |
| Rcor2 | 6.55809 | 4.74564 | 6.13149 | 5.47953 | 4.70983 | 4.92845 |
| Fat4 | 3.0503 | 3.32244 | 1.62302 | 1.3477 | 1.60848 | 1.9055 |
| Atf7 | 1.57176 | 1.65109 | 1.2557 | 0.913131 | 0.830274 | 0.788126 |
| Man2c1 | 1.07034 | 1.30058 | 1.3236 | 0.907034 | 0.64309 | 1.10676 |
| Elovl1 | 5.68945 | 7.17367 | 9.71869 | 4.34779 | 5.05825 | 4.02046 |
| Sowahc | 3.46553 | 5.04369 | 5.43018 | 2.63877 | 2.22901 | 2.75957 |
| Kcnq2 | 1.18685 | 1.14132 | 0.632908 | 0.649661 | 0.654565 | 0.71779 |
| Otulin | 17.753 | 16.9227 | 23.9592 | 15.3036 | 10.8003 | 9.24327 |
| Pi4k2a | 2.8916 | 5.39308 | 4.94558 | 2.84805 | 2.85707 | 2.5875 |
| Laptm5 | 39.1529 | 22.4187 | 46.7314 | 27.6616 | 26.6253 | 13.1152 |
| Mfsd8 | 1.10067 | 0.972091 | 0.978554 | 0.561085 | 0.490579 | 0.894884 |
| Mtmr9 | 8.61451 | 8.077 | 9.18912 | 3.49297 | 4.39974 | 6.81895 |
| Klhdc10 | 3.02072 | 2.81598 | 2.21578 | 1.42799 | 1.64148 | 2.12717 |
| Snx19 | 3.28995 | 1.99755 | 2.22801 | 1.6094 | 1.54588 | 1.57084 |
| Klf6 | 60.1096 | 50.0308 | 77.6714 | 26.3222 | 41.2049 | 29.0286 |
| Brpf1 | 3.42667 | 3.53525 | 4.56901 | 1.89622 | 2.56597 | 2.00877 |
| Arhgef10l | 4.24001 | 4.56435 | 6.15599 | 2.624 | 5.62249 | 2.23594 |
| Nfil3 | 28.6445 | 30.6043 | 33.1125 | 19.2211 | 21.537 | 16.0348 |
| Usp28 | 2.66812 | 2.49151 | 2.17039 | 2.43832 | 1.71979 | 1.81086 |

| | | | | | | |
|---|---|---|---|---|---|---|
| Osgin1 | 1.93232 | 6.17014 | 5.32337 | 1.29673 | 0.954607 | 0.0987208 |
| Pilrb2 | 0.0665674 | 1.98224 | 2.00872 | 0.203408 | 0.175822 | 0.0341357 |
| Rassf5 | 12.2151 | 6.41449 | 12.0825 | 1.92653 | 1.74176 | 1.05487 |
| Hmox1 | 25.9033 | 158.865 | 141.901 | 14.926 | 21.7373 | 18.2579 |
| Trem3 | 1.95015 | 25.4594 | 20.4518 | 2.83773 | 3.74532 | 1.3336 |
| Zc3h12a | 3.32412 | 2.64687 | 3.62921 | 0.441259 | 0.896914 | 0.404288 |
| 2510039O18Rik | 9.40864 | 15.8671 | 17.6148 | 2.86463 | 2.68407 | 1.9566 |
| Rhbdd1 | 0.865398 | 1.27005 | 1.02904 | 0.227985 | 0.0620176 | 0.131257 |
| H60b | 0.0582625 | 0.822352 | 0.603393 | 0.0324459 | 0 | 0.0600376 |
| Lrrk2 | 0.0764868 | 0.310673 | 0.363852 | 0.00864197 | 0.00907786 | 0.016418 |
| Gpr84 | 37.2125 | 107.384 | 111.702 | 9.7694 | 27.6212 | 4.5115 |
| Gm21188 | 1.62436 | 0 | 2.08275 | 0.144286 | 0.15517 | 0.128044 |
| Cd200r1 | 1.00879 | 1.02877 | 1.89947 | 0.396401 | 0.0572362 | 0.0388668 |
| Saa3 | 7133.01 | 16147.5 | 17366.4 | 1606.51 | 3611.47 | 1515.67 |
| Lgals3 | 7.104 | 89.9247 | 76.5265 | 10.6533 | 11.815 | 5.43798 |
| Smad6 | 2.10793 | 1.68481 | 2.61385 | 0.392698 | 0.208549 | 0.464622 |
| Trim21 | 1.34564 | 1.60063 | 2.09009 | 0.449914 | 0.0822452 | 0.162221 |
| Itga5 | 3.24237 | 12.0062 | 12.1308 | 0.98133 | 2.31747 | 0.52235 |
| Fcrlb | 0.695884 | 3.32145 | 2.4145 | 0.399346 | 0.676574 | 0.0494197 |
| Sirpb1b | 1.83556 | 2.45313 | 3.57882 | 0.209712 | 0.446084 | 0.185998 |
| Gpr31b | 9.69873 | 11.5436 | 12.0915 | 1.08856 | 2.20602 | 0.108955 |
| Olr1 | 0.117007 | 0.939762 | 0.660713 | 0 | 0.0999642 | 0 |
| Phf11a | 1.81065 | 5.67654 | 4.766 | 0.49769 | 1.30178 | 0.562787 |
| Ptafr | 7.27627 | 16.7367 | 17.9178 | 1.91937 | 2.85724 | 1.48282 |
| Gm15448 | 0.214299 | 0.435402 | 0.535137 | 0.024369 | 0.116269 | 0.0444426 |
| AA467197 | 0.853311 | 10.3579 | 7.87138 | 0.474637 | 1.45542 | 0.541866 |
| Fgr | 6.25286 | 8.93364 | 11.8482 | 1.54674 | 2.16652 | 0.286161 |
| Emilin2 | 23.8989 | 40.2733 | 45.3824 | 5.08485 | 9.27176 | 2.61322 |
| Ms4a4b | 0.8662 | 2.0922 | 2.24913 | 0.337406 | 0.220464 | 0.0204013 |
| Fas | 0.3892 | 3.91769 | 3.28714 | 0.366469 | 0.589751 | 0.0801518 |
| Bbs10 | 1.10143 | 0.65775 | 0.694268 | 0.248213 | 0.0413513 | 0.0493158 |
| Il1b | 10.9236 | 46.7707 | 40.2746 | 3.82461 | 8.62418 | 1.14665 |
| Sh3tc1 | 0.443271 | 0.621653 | 0.541535 | 0.192709 | 0.0519423 | 0.0496982 |
| Mfsd12 | 2.30328 | 4.31833 | 5.30721 | 0.453942 | 1.04815 | 0.64614 |
| Slc16a3 | 9.17352 | 20.9771 | 23.0333 | 2.29889 | 3.5847 | 1.82141 |
| Cyp4f18 | 5.87658 | 20.7301 | 18.9677 | 3.46849 | 1.71074 | 1.13482 |
| Hp | 52.2431 | 391.013 | 346.98 | 51.1297 | 48.46 | 14.7778 |
| Cxcr2 | 0.373866 | 3.98444 | 3.25602 | 0.30773 | 0.245792 | 0.467923 |
| Cd274 | 3.25175 | 15.0886 | 11.466 | 1.23903 | 1.9097 | 0.406049 |
| Tmem156 | 0.231864 | 0.746829 | 0.70966 | 0.0441988 | 0.0153448 | 0.214021 |
| Pram1 | 1.53082 | 3.3527 | 2.29775 | 0.154289 | 0.186297 | 0.103371 |
| Ppp1r3b | 0.87691 | 2.69344 | 2.76966 | 0.283595 | 0.320377 | 0.236883 |
| Gm9733 | 4.01634 | 6.35777 | 9.7067 | 0.0818037 | 1.72406 | 1.53089 |
| Ass1 | 11.1692 | 18.3203 | 27.4421 | 2.45402 | 3.82801 | 1.77456 |
| Adgre5 | 2.60277 | 3.10683 | 4.42011 | 0.557319 | 0.885426 | 0.377531 |
| Rab27a | 1.32792 | 0.818274 | 1.78069 | 0.251532 | 0.0759317 | 0.0916685 |
| Irg1 | 24.144 | 193.065 | 159.927 | 16.0277 | 25.5154 | 5.66998 |
| Hcn1 | 0.064777 | 0.0766046 | 0.0776008 | 0.00560305 | 0.00722288 | 0.0111082 |
| Arhgap27 | 3.55908 | 3.36291 | 4.90664 | 0.295842 | 1.21233 | 0.211187 |
| Alpk1 | 0.297255 | 0.839278 | 0.758544 | 0.0422462 | 0.0593716 | 0.0926236 |
| Sirpb1c | 3.65907 | 2.52128 | 4.9459 | 0.269498 | 0.497432 | 0.168167 |
| Galnt6 | 0.300115 | 2.95514 | 1.79549 | 0.266415 | 0.311903 | 0.0972934 |
| Tnfrsf11a | 2.02896 | 1.88775 | 2.61299 | 0.335445 | 0.214893 | 0.24639 |
| Cd300ld | 0.901152 | 0.831292 | 1.27592 | 0.0791348 | 0.323669 | 0.020913 |
| Tlr4 | 1.18039 | 0.951777 | 1.45849 | 0.0350301 | 0.140316 | 0.00487008 |
| Il21r | 2.42228 | 1.34264 | 3.0549 | 0.181938 | 0.290883 | 0.397382 |
| Nbeal2 | 0.68729 | 0.193735 | 0.55835 | 0.0056524 | 0.0671394 | 0.104084 |
| Psd4 | 0.563764 | 1.61136 | 1.34363 | 0.0812401 | 0.274809 | 0.0936609 |
| Dennd2c | 0.187016 | 0.110013 | 0.245139 | 0.00498931 | 0.00525648 | 0.0141129 |
| Gm5424 | 12.4414 | 19.4473 | 18.0498 | 1.20906 | 3.68745 | 1.50488 |
| Col14a1 | 0.550211 | 0.68708 | 1.00729 | 0.113778 | 0.136388 | 0.0181504 |
| Tbc1d10c | 0.888766 | 3.05892 | 2.27916 | 0.314482 | 0.0564064 | 0.25435 |
| Mcemp1 | 5.90857 | 21.5589 | 19.9821 | 3.44242 | 1.27391 | 0.531883 |
| Gda | 0.204227 | 2.78993 | 2.06908 | 0.12431 | 0.0925499 | 0.334419 |
| Mrgpra2a | 0.409684 | 6.72209 | 4.14061 | 0.812006 | 0.427067 | 0.029329 |
| Nlrp1a | 1.36334 | 0.0771431 | 0.761041 | 0.0403803 | 0.146055 | 0.00739266 |
| Siglec1 | 4.72917 | 5.49422 | 7.29819 | 0.66087 | 1.21262 | 0.289629 |
| Ap5b1 | 2.69941 | 7.0452 | 6.05499 | 0.267779 | 0.974189 | 0.709501 |
| Tnfrsf26 | 0.162295 | 0.76041 | 0.681509 | 0 | 0.0293208 | 0.195663 |
| Nos2 | 0.195815 | 4.399 | 3.30279 | 0.108221 | 0.290434 | 0.424763 |
| Bst1 | 1.31927 | 4.70639 | 4.50045 | 0.081085 | 0.540818 | 0.31991 |
| Dbx1 | 0.241428 | 3.84742 | 3.6211 | 0.116056 | 0.0623955 | 0.651742 |
| Car9 | 1.55895 | 0.805953 | 2.31602 | 0.0993366 | 0 | 0.379098 |
| Prss50 | 0.921399 | 7.07409 | 6.1287 | 0.076579 | 1.31287 | 0.0335629 |
| Il1rl2 | 0.0684236 | 0.689231 | 0.457388 | 0.0159673 | 0.00562559 | 0.144532 |
| Plin4 | 0.232744 | 0.218691 | 0.0448976 | 0.0232314 | 0 | 0.0568589 |
| Hcar2 | 3.29742 | 25.9581 | 22.1917 | 3.83954 | 0.990465 | 0.475831 |
| Ly6i | 0.875051 | 3.58867 | 2.70372 | 0.130909 | 0.626865 | 0.0301197 |
| Slc22a4 | 0.142407 | 1.38431 | 1.15609 | 0.0166384 | 0.0886141 | 0.0752493 |
| Stfa2 | 28.3546 | 391.581 | 340.746 | 25.0723 | 39.8255 | 6.1375 |
| Mefv | 0.148605 | 1.42049 | 1.12577 | 0.0543418 | 0.0231786 | 0.245055 |
| Mapk13 | 1.8167 | 6.82862 | 6.46302 | 1.20498 | 0.162069 | 0.132528 |
| Zfp819 | 2.53171 | 0.278402 | 2.65751 | 0.361046 | 0.0175245 | 0.148291 |
| Hps6 | 1.38154 | 2.63697 | 2.58858 | 0.300912 | 0.0232206 | 0.340525 |
| Slamf7 | 0.162854 | 0.893835 | 0.689601 | 0 | 0 | 0.0208568 |
| Dock5 | 0.207437 | 1.46063 | 1.2662 | 0.018464 | 0.192132 | 0.071114 |
| Cd38 | 6.86861 | 15.5664 | 14.801 | 0.697428 | 1.68088 | 0.566438 |
| Trib3 | 0.181467 | 1.9809 | 1.9551 | 0 | 0 | 0.274522 |
| Adam8 | 0.227937 | 4.91016 | 3.39692 | 0.358393 | 0.347628 | 0.320699 |
| Ccdc116 | 0.235245 | 0.281393 | 0.179301 | 0.0109058 | 1.43E-09 | 0.0633626 |
| Slamf6 | 0.21699 | 0.10935 | 0.25904 | 0 | 0.00792825 | 0.0557985 |
| G0s2 | 3.36627 | 45.6916 | 36.425 | 2.05862 | 4.25009 | 1.21613 |
| Arg1 | 21.443 | 130.91 | 121.392 | 14.4583 | 5.13628 | 2.29591 |
| Adra2c | 1.1142 | 0.756457 | 1.0478 | 0.169917 | 0 | 0.0977443 |
| Areg | 0.073961 | 2.1298 | 2.44482 | 0.0876961 | 0.186296 | 0.197704 |
| Adgrg6 | 0.249761 | 0.0881054 | 0.188128 | 0 | 0.00276239 | 0.00510039 |
| Lilr4b | 0.724322 | 8.17476 | 5.7284 | 0.461248 | 0.783167 | 0.166558 |
| Thbs1 | 6.51274 | 34.3629 | 28.6545 | 1.11866 | 1.63506 | 2.03703 |
| Hdc | 0.335397 | 4.97499 | 3.78413 | 0.235023 | 0.505188 | 0.206177 |
| Gm15056 | 0 | 3.36532 | 2.01072 | 0.558483 | 0.296453 | 0 |
| Tnfrsf14 | 0.455666 | 0.140327 | 0.611135 | 0.0389117 | 0.0417833 | 0 |
| Htra4 | 0.0555161 | 2.68407 | 2.08131 | 0 | 0 | 0.060395 |
| Sirpb1a | 1.47276 | 1.00076 | 1.72418 | 0.191762 | 0.240587 | 0.0613633 |
| P2ry2 | 1.28105 | 3.19495 | 3.15668 | 0.0577368 | 0.246846 | 0.167373 |
| Cass4 | 0.0382489 | 0.618945 | 0.515956 | 0.0145824 | 0 | 0 |
| Ahrr | 0.169489 | 0.162131 | 0.0784961 | 0 | 0.0284198 | 0 |

| | | | | | | |
|---|---|---|---|---|---|---|
| Wbp1l | 6.54581 | 5.36433 | 6.46071 | 4.6486 | 2.77597 | 3.96483 |
| Camkk1 | 2.62511 | 2.58 | 2.95019 | 1.91512 | 1.72839 | 1.75369 |
| Kpna4 | 2.46248 | 2.5623 | 2.88239 | 1.20372 | 1.41206 | 1.53557 |
| Per2 | 0.838584 | 0.657252 | 0.646465 | 0.321871 | 0.327617 | 0.56465 |
| Tmem135 | 2.60459 | 2.72277 | 2.83733 | 0.979758 | 1.59036 | 1.26812 |
| Vps33a | 7.27996 | 4.53706 | 6.05165 | 4.06579 | 3.96053 | 4.1333 |
| Trappc8 | 3.17103 | 2.61963 | 2.8475 | 1.56464 | 1.85378 | 1.29393 |
| Ncf2 | 2.65222 | 2.60663 | 4.01995 | 1.90778 | 2.44469 | 1.49585 |
| Tspan5 | 6.90942 | 6.65326 | 5.24323 | 3.74547 | 3.68968 | 3.93803 |
| Sec31a | 3.33066 | 4.33172 | 5.23107 | 2.44925 | 2.26457 | 2.51576 |
| Atl3 | 2.64143 | 1.33053 | 2.43814 | 1.05277 | 0.992014 | 1.24364 |
| Nadk2 | 1.728 | 1.72806 | 2.09736 | 1.18858 | 0.75772 | 1.26015 |
| Chst12 | 10.3404 | 9.51725 | 12.362 | 7.66874 | 7.31408 | 4.83476 |
| Bhlhe22 | 12.1599 | 9.6797 | 7.18811 | 4.63301 | 5.30928 | 5.54083 |
| Med17 | 4.27244 | 5.17218 | 6.3992 | 2.87519 | 2.78598 | 3.2532 |
| Usp12 | 2.08713 | 1.80356 | 2.52157 | 1.55861 | 1.26858 | 1.08873 |
| Ankfy1 | 3.10351 | 4.04175 | 4.39195 | 2.27992 | 2.11239 | 2.03863 |
| Zfp12 | 2.2923 | 1.63297 | 1.84677 | 0.936695 | 1.17836 | 1.59764 |
| Pacsin2 | 5.8958 | 5.10582 | 8.48186 | 3.2141 | 4.65065 | 3.65154 |
| Zc3h7a | 1.92217 | 1.61589 | 1.67382 | 0.918251 | 1.33794 | 0.865138 |
| Zdhhc6 | 30.4144 | 23.5415 | 37.882 | 18.3387 | 17.9699 | 14.8992 |
| Tm9sf2 | 33.2448 | 27.7325 | 35.1379 | 16.1054 | 17.1271 | 16.3978 |
| Tln1 | 10.556 | 7.0015 | 12.4532 | 6.64923 | 7.55062 | 5.52486 |
| Zxdc | 1.52158 | 1.23658 | 1.48616 | 0.875314 | 0.761717 | 0.498951 |
| Fancg | 1.54262 | 1.35363 | 1.41859 | 0.728478 | 1.23564 | 0.882642 |
| Extl3 | 8.77338 | 5.72465 | 7.62538 | 3.23907 | 4.76579 | 4.96607 |
| Ssh1 | 1.61111 | 1.79314 | 1.74582 | 1.05092 | 1.1356 | 1.03171 |
| Mob3a | 9.31474 | 11.2968 | 13.6316 | 6.01405 | 6.7185 | 6.55548 |
| D8Ertd82e | 2.20052 | 1.98302 | 2.28613 | 1.24582 | 1.43599 | 1.05492 |
| Tspyl3 | 5.90583 | 3.62501 | 4.71441 | 2.68872 | 1.71547 | 3.59346 |
| Gmfg | 4.5746 | 2.41439 | 4.21049 | 1.13156 | 1.35164 | 0.616038 |
| Triobp | 2.31732 | 3.05658 | 2.8629 | 1.16059 | 1.60686 | 2.21708 |
| Adgrl2 | 3.24234 | 2.14454 | 1.8852 | 1.07722 | 1.46467 | 1.53886 |
| Bach1 | 6.05237 | 5.1183 | 6.99048 | 3.20564 | 3.17709 | 3.37032 |
| Wdtc1 | 2.86542 | 2.80479 | 3.57038 | 1.98325 | 1.99694 | 1.55925 |
| Pycard | 32.1108 | 22.7707 | 34.3939 | 18.874 | 17.5288 | 11.5271 |
| Snx11 | 3.48538 | 3.84793 | 3.96777 | 2.66028 | 2.90631 | 1.69027 |
| Otud5 | 4.07131 | 4.6486 | 4.35564 | 3.11292 | 2.87747 | 3.57151 |
| Jund | 91.9053 | 101.821 | 124.036 | 68.615 | 70.4077 | 47.8088 |
| Sh3kbp1 | 6.06532 | 3.94706 | 5.41997 | 2.79836 | 2.77955 | 2.73745 |
| Brinp2 | 4.39666 | 3.2609 | 2.62162 | 2.50516 | 1.49775 | 1.75396 |
| Asah1 | 12.3955 | 12.2352 | 15.8537 | 6.80575 | 7.18846 | 6.1043 |
| Cherp | 7.69473 | 7.29488 | 8.88819 | 6.62301 | 4.60105 | 6.2634 |
| Mkl1 | 1.8056 | 1.34409 | 1.6431 | 1.11598 | 1.12007 | 1.00651 |
| Tmem2 | 7.78494 | 10.3849 | 10.3193 | 4.81434 | 5.59156 | 5.31829 |
| Cttnbp2 | 10.1303 | 7.90662 | 5.91518 | 3.86817 | 4.49704 | 4.63362 |
| B630005N14Rik | 3.74993 | 3.32806 | 3.143 | 1.56132 | 1.90921 | 1.66012 |
| Mafg | 5.98366 | 7.38444 | 7.07507 | 5.19281 | 3.8681 | 3.08037 |
| Ifitm2 | 197.411 | 477.715 | 522.124 | 215.079 | 288.145 | 183.217 |
| Fth1 | 3607.49 | 6764.98 | 7610.58 | 3493.17 | 4360.76 | 2403.37 |
| Rab8b | 9.45727 | 8.99413 | 12.0006 | 5.55931 | 5.24331 | 5.06382 |
| Fcer1g | 115.75 | 124.055 | 183.334 | 97.6378 | 115.263 | 48.2023 |
| Ppp1r12c | 2.29472 | 2.62372 | 2.65775 | 1.9848 | 1.92861 | 1.5658 |
| Herpud2 | 11.14 | 12.3694 | 12.4665 | 8.23661 | 3.5642 | 8.11423 |
| Rreb1 | 0.690064 | 0.842994 | 0.946922 | 0.430524 | 0.513436 | 0.417171 |
| Pced1a | 2.07188 | 1.98357 | 1.54154 | 1.16889 | 1.33223 | 1.17732 |
| Vegfa | 2.8923 | 2.35062 | 2.63513 | 0.9314 | 1.09911 | 1.81989 |
| Ago1 | 8.68367 | 5.69164 | 7.58252 | 4.67132 | 4.24426 | 5.04389 |
| Hebp1 | 13.2439 | 6.46832 | 13.8717 | 7.18787 | 7.70694 | 3.69737 |
| Megf9 | 6.06242 | 5.56803 | 4.76033 | 2.3567 | 2.44204 | 3.44876 |
| Pld4 | 95.983 | 59.8151 | 117.637 | 57.617 | 72.8727 | 37.5797 |
| Atp13a1 | 6.90549 | 5.37485 | 5.50624 | 2.92954 | 2.6986 | 5.30322 |
| Lgmn | 147.549 | 114.824 | 186.9 | 99.5546 | 113.953 | 57.4838 |
| Mark2 | 2.91796 | 2.28391 | 2.61077 | 2.11124 | 1.78677 | 1.56215 |
| Ppm1b | 13.7795 | 13.1846 | 13.3391 | 5.47192 | 6.77655 | 8.19477 |
| Rab29 | 8.51861 | 7.61321 | 9.12258 | 4.96701 | 5.16074 | 4.79202 |
| Nkain1 | 4.68113 | 5.09157 | 4.25229 | 3.56518 | 4.89971 | 2.79564 |
| Pfkl | 15.9101 | 18.9079 | 21.6229 | 10.9218 | 12.1652 | 10.6412 |
| Xpo5 | 6.06252 | 7.70954 | 7.87395 | 2.27637 | 3.2548 | 4.89386 |
| Nfe2l2 | 9.41892 | 9.97078 | 14.4829 | 7.60439 | 6.19048 | 3.38704 |
| Zfp263 | 7.57586 | 13.7383 | 12.0622 | 6.83385 | 7.78712 | 4.83943 |
| Klhl8 | 5.93264 | 4.90404 | 5.42593 | 4.13528 | 2.77501 | 2.44616 |
| Numbl | 2.12484 | 1.85884 | 1.79595 | 1.08395 | 1.02923 | 1.18968 |
| 8-Sep | 2.47378 | 2.09812 | 2.60106 | 1.32942 | 1.36839 | 1.37842 |
| Ep300 | 1.83146 | 2.50879 | 2.43008 | 1.38299 | 0.935471 | 1.40358 |
| Creg1 | 16.3003 | 17.5465 | 26.5194 | 10.1169 | 11.9774 | 8.09326 |
| Spock1 | 3.51558 | 4.10872 | 2.08338 | 1.71968 | 1.55697 | 2.34734 |
| Ldha | 127.273 | 192.917 | 220.224 | 107.634 | 111.451 | 86.1365 |
| Qrsl1 | 1.82935 | 1.84606 | 1.64517 | 1.18584 | 0.665656 | 0.837508 |
| Rnf130 | 41.5959 | 42.6802 | 54.0194 | 27.1726 | 23.7696 | 19.1898 |
| Cyp4f16 | 1.71322 | 1.16046 | 1.83086 | 1.0953 | 0.815359 | 1.25616 |
| Zfyve1 | 5.35775 | 4.79426 | 5.75696 | 3.17116 | 2.67457 | 3.22446 |
| Nestn | 19.0814 | 21.274 | 26.074 | 10.6253 | 11.1344 | 14.1181 |
| Slc38a7 | 3.28859 | 5.68003 | 6.41951 | 3.92428 | 3.68305 | 2.00704 |
| Elmo1 | 4.40667 | 2.57019 | 4.17143 | 1.84448 | 2.42662 | 2.04773 |
| Agpat3 | 5.50545 | 4.75536 | 5.78351 | 3.20812 | 2.8005 | 3.72388 |
| Nfasc | 7.79859 | 6.09988 | 5.5217 | 3.10652 | 3.75119 | 3.26012 |
| Renbp | 0.971178 | 1.71477 | 2.2402 | 0.827857 | 1.15487 | 0.740159 |
| Arhgdib | 184.427 | 105.906 | 217.668 | 120.282 | 123.945 | 54.1623 |
| Raver1 | 15.2929 | 16.9701 | 19.1881 | 10.3386 | 7.1989 | 11.9151 |
| Adpgk | 5.35847 | 9.89801 | 8.11509 | 5.23431 | 2.70671 | 4.61383 |
| Ctsb | 71.272 | 139.455 | 144.009 | 67.6094 | 74.158 | 51.724 |
| Sppl2a | 3.68723 | 5.0445 | 5.09454 | 2.43803 | 2.3406 | 2.41343 |
| Slc11a2 | 1.58421 | 2.3354 | 1.72948 | 1.11334 | 1.52174 | 1.17688 |
| Traf2 | 1.29994 | 1.15369 | 2.12347 | 0.680507 | 1.05955 | 1.14093 |
| Tti1 | 4.90008 | 4.44976 | 5.62347 | 2.23743 | 1.95126 | 3.9304 |
| Pnp | 13.0422 | 16.3014 | 20.2884 | 7.85985 | 11.4571 | 7.55227 |
| Alyref | 11.3178 | 11.6113 | 12.4125 | 7.86356 | 5.55727 | 4.8093 |
| P4hb | 100.627 | 153.64 | 165.805 | 74.502 | 78.5867 | 85.427 |
| Lipe | 2.34016 | 2.69769 | 2.82008 | 1.87744 | 0.971898 | 1.447 |
| Fam160b1 | 3.2037 | 3.03663 | 2.80167 | 1.42156 | 1.8427 | 1.43167 |

| | | | | | | |
|---|---|---|---|---|---|---|
| Nlrc3 | 0.502078 | 0.27691 | 0.504148 | 0.0276399 | 0.0196392 | 0.00420998 |
| Sgms2 | 0.153836 | 2.70282 | 1.98429 | 0.093813 | 0.0659577 | 0.112806 |
| Prok2 | 1.05761 | 12.8939 | 10.7178 | 0.590475 | 0.505479 | 0.479565 |
| Mmel1 | 0.0867044 | 0.308162 | 0.12416 | 0.013474 | 0.0289528 | 0 |
| Myrf | 0.151213 | 0.0324408 | 0.134838 | 0.00442397 | 0 | 0 |
| Scarf2 | 8.59157 | 1.55081 | 8.53239 | 0.439071 | 0.358361 | 0.249844 |
| Pira2 | 0.0952953 | 0.863441 | 0.874224 | 0.0146919 | 0.178012 | 0.0274223 |
| Slfn3 | 0.681363 | 0.452247 | 1.11011 | 0 | 0 | 0 |
| Fam109b | 0.271445 | 0.368047 | 0.465203 | 0.041912 | 0 | 0 |
| Fap | 0 | 0.109747 | 0.105476 | 0 | 0 | 0.015066 |
| Ltb4r1 | 2.64925 | 25.1366 | 20.4889 | 1.16936 | 0.88157 | 0.470576 |
| Prss27 | 0.468272 | 0.218537 | 0.0634222 | 0 | 0 | 0.109787 |
| Fut4 | 0.411715 | 0.496045 | 0.828444 | 0.0817554 | 0 | 0 |
| Smpd5 | 0.353227 | 0.400515 | 0.246141 | 9.00E-08 | 0.0290329 | 0.115083 |
| Il20rb | 0.264789 | 0.0311 | 0.245748 | 0.00811912 | 0 | 0.00749403 |
| Mmp9 | 0.905032 | 16.721 | 12.587 | 1.03135 | 0.27572 | 0.0257394 |
| Fcrl5 | 3.09071 | 1.19238 | 4.13443 | 0 | 0.340655 | 0.0188596 |
| Edn1 | 0.235509 | 2.6127 | 2.16528 | 0.128858 | 0 | 0.068875 |
| Ceacam18 | 0.839049 | 0.201971 | 0.879107 | 0 | 0 | 0.0487854 |
| Mmp8 | 3.45088 | 55.7734 | 44.1427 | 1.25923 | 1.80278 | 0.644382 |
| Muc5b | 0.0608545 | 0.0269155 | 0.0898008 | 0 | 0 | 0 |
| Shisa3 | 0 | 0.117839 | 0.192982 | 0 | 0 | 0 |
| Cyp11a1 | 0.0126721 | 0.587387 | 0.454247 | 0 | 0 | 0 |
| Ptger2 | 0.0794073 | 0.923766 | 0.941175 | 0.0101381 | 0.0423808 | 0.0195456 |
| Asb5 | 0.314169 | 0.291314 | 0.339242 | 0 | 0 | 0 |
| Gpr132 | 0.184563 | 1.49508 | 1.45033 | 0.0364108 | 0.0585149 | 0 |
| Acvr1c | 0.424194 | 0.539234 | 0.149704 | 0.00306762 | 0.00318952 | 0.0269457 |
| F10 | 0.445938 | 8.10839 | 5.63996 | 0.0167924 | 0.360063 | 0 |
| Prss46 | 0.124637 | 4.86583 | 3.36891 | 0.146429 | 0 | 0.0661933 |
| Ip6k3 | 1.3494 | 0.604836 | 1.45197 | 0 | 0 | 0.0741193 |
| Asprv1 | 4.72787 | 95.467 | 76.7815 | 1.40498 | 1.16696 | 1.11448 |
| F5 | 0.0402541 | 0.488596 | 0.461854 | 0 | 0.00947404 | 0 |
| Clec2i | 0.535721 | 0.180543 | 0.489146 | 0 | 0 | 0 |
| Klra17 | 0.0993537 | 0.527191 | 0.32105 | 0 | 0 | 0 |
| Il1r2 | 2.60759 | 33.996 | 29.5651 | 0.662855 | 0.0426191 | 0.477725 |
| C4b | 0.127309 | 0.111118 | 0.254053 | 0.00777748 | 0 | 0.00691456 |
| Adgre4 | 0.283534 | 2.18298 | 1.92302 | 0.0145367 | 0.022572 | 0.00713715 |
| Olfm4 | 7.76504 | 117.258 | 88.9264 | 0.800994 | 2.05623 | 0.826975 |
| Lrg1 | 3.96864 | 70.4856 | 59.6053 | 1.38321 | 1.50178 | 0 |
| Mrgpra2b | 1.3521 | 16.6314 | 12.5466 | 0 | 0.492044 | 0.0819472 |
| Fcrla | 0.118977 | 0.596782 | 0.709305 | 0 | 0 | 0.0204761 |
| Krt80 | 0.0992878 | 2.11024 | 1.73628 | 0 | 0 | 0 |
| Ppef1 | 0.0440121 | 0.453238 | 0.290574 | 0 | 0 | 0 |
| Cfi | 0.0342519 | 0.372346 | 0.292876 | 0 | 0 | 0 |
| Zfp831 | 0.0124964 | 0.0599497 | 0.0290143 | 0 | 0 | 0 |
| Spocd1 | 0.31433 | 0.0433723 | 0.302315 | 0 | 0 | 0 |
| Gpr62 | 0.657874 | 0.260368 | 0.456815 | 0 | 0 | 0 |
| Egr3 | 0.104893 | 0.817231 | 0.61212 | 0 | 0 | 0 |
| Them5 | 0.697939 | 0 | 0.236721 | 0 | 0 | 0 |
| Btnl9 | 0.0234179 | 0.121831 | 0.0424697 | 0 | 0 | 0 |
| Fbxl8 | 0 | 0.932743 | 1.22372 | 0 | 0 | 0 |
| Retnlg | 40.691 | 734.605 | 598.098 | 5.43815 | 9.5393 | 0.745886 |
| C1rl | 1.45358 | 1.77169 | 2.43242 | 0 | 0.0320116 | 0 |
| Foxa1 | 0.0273978 | 0.821604 | 0.53733 | 0 | 0.0159098 | 0 |
| Capns2 | 0 | 0.958947 | 0.561143 | 0 | 0 | 0 |
| Ltf | 2.23489 | 31.6682 | 25.3798 | 0.118018 | 0.566151 | 0.139742 |
| Slc5a11 | 0.150947 | 0 | 0.165872 | 0 | 0 | 0 |
| Spaca4 | 1.80778 | 0.79254 | 0.522928 | 0 | 0 | 0 |
| Cbln3 | 0.121847 | 0.159109 | 0.10154 | 0 | 0 | 0 |
| Acpp | 0 | 0.0447924 | 0.0548833 | 0 | 0 | 0 |
| Ccl22 | 0.444287 | 0.0589757 | 0.206854 | 0 | 0 | 0 |
| Gdf9 | 0.604896 | 0.0765073 | 0.266986 | 0 | 0 | 0 |
| Tnfsf14 | 0.0438235 | 1.53861 | 0.838599 | 0.0270081 | 0 | 0 |
| Ifnlr1 | 0 | 0.280823 | 0.47454 | 0 | 0 | 0 |
| Aqp3 | 0 | 0.584336 | 0.341353 | 0 | 0 | 0 |
| Npff | 1.35566 | 1.4195 | 0.267315 | 5.85E-09 | 3.43E-09 | 2.19E-08 |
| Fam46b | 0.452097 | 0.28453 | 0.42713 | 0 | 0 | 0 |
| Mocos | 0.082943 | 1.74918 | 1.2632 | 0 | 0 | 0 |
| Gnat2 | 0.121226 | 0.184167 | 0 | 0 | 0 | 0 |
| Ambp | 0 | 0.737535 | 0.408274 | 0 | 0 | 0 |
| Ceacam10 | 0.0957599 | 1.4042 | 0.905015 | 0.0272569 | 0 | 0 |
| Pnliprp2 | 1.05852 | 0.438755 | 0.0478468 | 0 | 0 | 0 |
| Mctp2 | 0.0273762 | 0.0912194 | 0.105142 | 0 | 0 | 0.00220364 |
| Cracr2a | 0.748297 | 0.420739 | 0.607086 | 0 | 0 | 0 |
| Cpa3 | 0 | 0.729738 | 0.36485 | 0 | 0 | 0 |
| B3gnt3 | 0.11228 | 2.02487 | 1.63023 | 0 | 0 | 0 |
| Gm20939 | 0.026732 | 0.282298 | 0.317243 | 0 | 0 | 0 |
| Mgam | 0 | 0.0259091 | 0.07303 | 0 | 0 | 0 |
| Pdcd1lg2 | 1.0457 | 0.626538 | 1.34913 | 0 | 0 | 0.0277533 |
| Lin28a | 0.0376242 | 0.381587 | 0.300365 | 0 | 0 | 0 |
| Cd177 | 1.13369 | 16.6621 | 12.6073 | 0 | 0.213544 | 0.0224462 |
| Fam169b | 0.00931746 | 0.148237 | 0.156034 | 0 | 0 | 0 |
| Cd6 | 0.129516 | 0.234549 | 0.32768 | 0 | 0 | 0 |
| Cd300lb | 0.168564 | 1.57122 | 1.52729 | 0.012105 | 0.0128785 | 0 |
| Spint1 | 0.029793 | 0.550838 | 0.452333 | 0 | 0 | 0 |
| Adra2b | 0.240053 | 0.473135 | 0.499108 | 0 | 0 | 0 |
| Tdo2 | 0.278527 | 0.195046 | 0.036525 | 0 | 0 | 0 |
| Vdr | 0.925349 | 0.118956 | 0.932038 | 0.0127185 | 0 | 0 |
| Nlrp12 | 0.0158889 | 0.23831 | 0.125613 | 0 | 0 | 0 |
| Mfsd7a | 0.112547 | 0.633158 | 0.740049 | 0 | 0 | 0 |
| Gmnc | 0.010866 | 0.211867 | 0.252312 | 0 | 0 | 0 |
| Fam83f | 0.191491 | 1.07284 | 0.963997 | 0 | 0 | 0 |
| Il18rap | 0.0381797 | 0.584709 | 0.624037 | 0 | 0.00843019 | 0 |
| 9830107B12Rik | 0.0484774 | 0.674045 | 0.362658 | 0 | 0 | 0 |
| Igsf1 | 0.248817 | 0.180374 | 0.04627 | 0 | 0 | 0 |
| Cd3d | 0.0581766 | 2.89042 | 1.94329 | 0 | 0 | 0 |
| Slc38a8 | 0.0229477 | 0.741794 | 0.779131 | 0 | 0 | 0 |
| Ccr7 | 0.085661 | 2.36893 | 1.78326 | 0 | 0 | 0 |
| Pparg | 0.146579 | 2.06014 | 1.5411 | 0 | 0 | 0 |
| Ly6g | 0.584378 | 6.44698 | 6.33622 | 0 | 0 | 0 |
| Kcng4 | 0.180441 | 3.13994 | 1.90781 | 0 | 0 | 0 |

**Appendix Table S3.** Enriched GO terms in up-regulated genes in Ptf1a RGC.

| Term | Description | PValue | Fold Enrichment | FDR |
|---|---|---|---|---|
| GO:0035556 | intracellular signal transduction | 2.53E-43 | 2.142857143 | 1.12E-39 |
| GO:0031347 | regulation of defense response | 3.09E-34 | 3.171066908 | 6.85E-31 |
| GO:0046649 | lymphocyte activation | 7.79E-29 | 2.647308588 | 1.15E-25 |
| GO:1902531 | regulation of intracellular signal transduction | 2.52E-28 | 2.136779752 | 2.79E-25 |
| GO:0009966 | regulation of signal transduction | 1.70E-27 | 1.801984768 | 1.51E-24 |
| GO:0002366 | leukocyte activation involved in immune response | 3.24E-26 | 4.123781263 | 2.39E-23 |
| GO:0016477 | cell migration | 1.01E-25 | 2.19521985 | 6.39E-23 |
| GO:0002274 | myeloid leukocyte activation | 3.63E-25 | 4.520064205 | 2.01E-22 |
| GO:0002694 | regulation of leukocyte activation | 1.21E-24 | 2.732858879 | 5.96E-22 |
| GO:0050900 | leukocyte migration | 1.37E-24 | 3.731778426 | 6.05E-22 |
| GO:0009967 | positive regulation of signal transduction | 6.30E-24 | 2.087950895 | 2.54E-21 |
| GO:0007159 | leukocyte cell-cell adhesion | 1.64E-23 | 3.396482567 | 6.07E-21 |
| GO:0007166 | cell surface receptor signaling pathway | 2.46E-23 | 1.772496097 | 8.39E-21 |
| GO:1902533 | positive regulation of intracellular signal transduction | 6.78E-23 | 2.355828221 | 2.14E-20 |
| GO:0031349 | positive regulation of defense response | 1.15E-22 | 3.361050328 | 3.40E-20 |
| GO:0007249 | I-kappaB kinase/NF-kappaB signaling | 1.57E-22 | 4.591218306 | 4.36E-20 |
| GO:0050727 | regulation of inflammatory response | 4.80E-22 | 3.447664507 | 1.25E-19 |
| GO:0001819 | positive regulation of cytokine production | 1.17E-21 | 3.064456722 | 2.87E-19 |
| GO:0045088 | regulation of innate immune response | 2.98E-21 | 3.523809524 | 6.62E-19 |
| GO:0019221 | cytokine-mediated signaling pathway | 3.04E-21 | 3.317136378 | 6.62E-19 |
| GO:0002521 | leukocyte differentiation | 3.14E-21 | 2.764119601 | 6.62E-19 |
| GO:1903555 | regulation of tumor necrosis factor superfamily cytokine production | 1.34E-20 | 4.758017493 | 2.71E-18 |
| GO:0050778 | positive regulation of immune response | 3.59E-20 | 2.435433562 | 6.91E-18 |
| GO:0048534 | hematopoietic or lymphoid organ development | 5.40E-20 | 2.259301587 | 9.96E-18 |
| GO:0032103 | positive regulation of response to external stimulus | 5.70E-20 | 3.462095238 | 1.01E-17 |
| GO:0016310 | phosphorylation | 1.01E-19 | 1.781175346 | 1.72E-17 |
| GO:0002520 | immune system development | 1.42E-19 | 2.202912505 | 2.33E-17 |
| GO:0006897 | endocytosis | 1.77E-19 | 2.432597842 | 2.80E-17 |
| GO:0019220 | regulation of phosphate metabolic process | 1.95E-19 | 1.903744947 | 2.98E-17 |
| GO:0001818 | negative regulation of cytokine production | 2.21E-19 | 3.766492644 | 3.26E-17 |
| GO:0032680 | regulation of tumor necrosis factor production | 2.47E-19 | 4.642487047 | 3.53E-17 |
| GO:0051174 | regulation of phosphorus metabolic process | 2.62E-19 | 1.898676403 | 3.62E-17 |
| GO:0030097 | hemopoiesis | 3.49E-19 | 2.262325581 | 4.68E-17 |
| GO:0032640 | tumor necrosis factor production | 5.87E-19 | 4.64399093 | 7.65E-17 |
| GO:0010647 | positive regulation of cell communication | 1.13E-18 | 1.832128961 | 1.43E-16 |
| GO:0032675 | regulation of interleukin-6 production | 1.50E-18 | 4.840336134 | 1.84E-16 |
| GO:0031325 | positive regulation of cellular metabolic process | 4.74E-18 | 1.562323391 | 5.68E-16 |
| GO:0030334 | regulation of cell migration | 1.05E-17 | 2.249433107 | 1.22E-15 |
| GO:0010942 | positive regulation of cell death | 1.08E-17 | 2.484734263 | 1.23E-15 |
| GO:1903037 | regulation of leukocyte cell-cell adhesion | 1.26E-17 | 3.112462006 | 1.39E-15 |
| GO:0050866 | negative regulation of cell activation | 1.30E-17 | 4.090225564 | 1.41E-15 |
| GO:0002695 | negative regulation of leukocyte activation | 1.92E-17 | 4.281533101 | 2.02E-15 |
| GO:2000145 | regulation of cell motility | 2.51E-17 | 2.189496191 | 2.59E-15 |
| GO:0002285 | lymphocyte activation involved in immune response | 4.49E-17 | 4.04551201 | 4.52E-15 |
| GO:0006915 | apoptotic process | 5.56E-17 | 1.795027976 | 5.48E-15 |
| GO:0097529 | myeloid leukocyte migration | 6.22E-17 | 3.881049563 | 5.99E-15 |
| GO:0002685 | regulation of leukocyte migration | 1.01E-16 | 3.901972504 | 9.54E-15 |
| GO:0050764 | regulation of phagocytosis | 1.11E-16 | 5.285714286 | 1.02E-14 |
| GO:0043122 | regulation of I-kappaB kinase/NF-kappaB signaling | 2.04E-16 | 4.044766294 | 1.85E-14 |
| GO:0071345 | cellular response to cytokine stimulus | 2.13E-16 | 2.167450611 | 1.88E-14 |
| GO:0000165 | MAPK cascade | 2.58E-16 | 2.355146676 | 2.20E-14 |
| GO:0023014 | signal transduction by protein phosphorylation | 2.58E-16 | 2.355146676 | 2.20E-14 |
| GO:0043068 | positive regulation of programmed cell death | 3.10E-16 | 2.505622657 | 2.59E-14 |
| GO:0050863 | regulation of T cell activation | 3.36E-16 | 3.134693878 | 2.76E-14 |
| GO:0002699 | positive regulation of immune effector process | 4.44E-16 | 3.334780721 | 3.57E-14 |
| GO:0002286 | T cell activation involved in immune response | 5.32E-16 | 5.087003222 | 4.21E-14 |
| GO:0043065 | positive regulation of apoptotic process | 5.84E-16 | 2.49716803 | 4.54E-14 |
| GO:0060627 | regulation of vesicle-mediated transport | 6.87E-16 | 2.550828482 | 5.25E-14 |
| GO:0010562 | positive regulation of phosphorus metabolic process | 2.26E-15 | 2.012303486 | 1.67E-13 |
| GO:0045937 | positive regulation of phosphate metabolic process | 2.26E-15 | 2.012303486 | 1.67E-13 |
| GO:0030335 | positive regulation of cell migration | 3.83E-15 | 2.56427379 | 2.78E-13 |
| GO:0070663 | regulation of leukocyte proliferation | 5.73E-15 | 3.46122449 | 4.09E-13 |
| GO:0051249 | regulation of lymphocyte activation | 6.32E-15 | 2.401937046 | 4.44E-13 |
| GO:0045089 | positive regulation of innate immune response | 6.77E-15 | 3.352380952 | 4.69E-13 |
| GO:0050867 | positive regulation of cell activation | 7.11E-15 | 2.519101644 | 4.85E-13 |
| GO:0043408 | regulation of MAPK cascade | 7.97E-15 | 2.314943369 | 5.35E-13 |
| GO:0046631 | alpha-beta T cell activation | 1.02E-14 | 4.053019146 | 6.71E-13 |
| GO:2000377 | regulation of reactive oxygen species metabolic process | 1.26E-14 | 3.736645963 | 8.24E-13 |
| GO:0022409 | positive regulation of cell-cell adhesion | 1.31E-14 | 3.216931217 | 8.30E-13 |
| GO:0030217 | T cell differentiation | 1.31E-14 | 3.216931217 | 8.30E-13 |
| GO:0051246 | regulation of protein metabolic process | 2.33E-14 | 1.555473098 | 1.45E-12 |
| GO:0051250 | negative regulation of lymphocyte activation | 2.76E-14 | 4.203612479 | 1.70E-12 |

| | | | | |
|---|---|---|---|---|
| GO:2000147 | positive regulation of cell motility | 4.63E-14 | 2.453674121 | 2.81E-12 |
| GO:1903039 | positive regulation of leukocyte cell-cell adhesion | 5.33E-14 | 3.378881988 | 3.19E-12 |
| GO:0030036 | actin cytoskeleton organization | 5.54E-14 | 2.330035615 | 3.27E-12 |
| GO:0030100 | regulation of endocytosis | 6.43E-14 | 3.141104294 | 3.75E-12 |
| GO:0043067 | regulation of programmed cell death | 7.19E-14 | 1.776326531 | 4.14E-12 |
| GO:0031399 | regulation of protein modification process | 7.74E-14 | 1.712009106 | 4.40E-12 |
| GO:0051272 | positive regulation of cellular component movement | 9.01E-14 | 2.409745293 | 5.05E-12 |
| GO:0046651 | lymphocyte proliferation | 1.04E-13 | 2.996825397 | 5.76E-12 |
| GO:0043410 | positive regulation of MAPK cascade | 1.56E-13 | 2.518783542 | 8.44E-12 |
| GO:1901652 | response to peptide | 1.56E-13 | 2.518783542 | 8.44E-12 |
| GO:1903557 | positive regulation of tumor necrosis factor superfamily cytokine production | 2.14E-13 | 4.876190476 | 1.14E-11 |
| GO:0050729 | positive regulation of inflammatory response | 2.26E-13 | 4.136054422 | 1.19E-11 |
| GO:0032496 | response to lipopolysaccharide | 2.99E-13 | 2.650660264 | 1.54E-11 |
| GO:0002696 | positive regulation of leukocyte activation | 3.00E-13 | 2.452583071 | 1.54E-11 |
| GO:0032268 | regulation of cellular protein metabolic process | 3.07E-13 | 1.551783859 | 1.56E-11 |
| GO:0042981 | regulation of apoptotic process | 3.59E-13 | 1.75864923 | 1.81E-11 |
| GO:0032270 | positive regulation of cellular protein metabolic process | 4.30E-13 | 1.754582989 | 2.14E-11 |
| GO:0002718 | regulation of cytokine production involved in immune response | 4.68E-13 | 4.606324973 | 2.30E-11 |
| GO:0060326 | cell chemotaxis | 5.78E-13 | 3.057054401 | 2.82E-11 |
| GO:0010648 | negative regulation of cell communication | 6.03E-13 | 1.832336504 | 2.90E-11 |
| GO:0030595 | leukocyte chemotaxis | 6.28E-13 | 3.486682809 | 2.99E-11 |
| GO:0032760 | positive regulation of tumor necrosis factor production | 8.20E-13 | 4.803874092 | 3.86E-11 |
| GO:0050670 | regulation of lymphocyte proliferation | 9.94E-13 | 3.331118494 | 4.63E-11 |
| GO:0051247 | positive regulation of protein metabolic process | 1.10E-12 | 1.718053375 | 5.07E-11 |
| GO:0043549 | regulation of kinase activity | 1.22E-12 | 2.186841149 | 5.58E-11 |
| GO:0032944 | regulation of mononuclear cell proliferation | 2.01E-12 | 3.267789245 | 9.09E-11 |
| GO:0002292 | T cell differentiation involved in immune response | 2.45E-12 | 5.786618445 | 1.09E-10 |
| GO:0061082 | myeloid leukocyte cytokine production | 2.46E-12 | 6.372294372 | 1.09E-10 |
| GO:0050766 | positive regulation of phagocytosis | 2.56E-12 | 5.30875576 | 1.12E-10 |
| GO:0031098 | stress-activated protein kinase signaling cascade | 2.77E-12 | 3.099273608 | 1.20E-10 |
| GO:0002221 | pattern recognition receptor signaling pathway | 2.97E-12 | 3.817896389 | 1.28E-10 |
| GO:0002703 | regulation of leukocyte mediated immunity | 3.11E-12 | 3.047619048 | 1.30E-10 |
| GO:0072358 | cardiovascular system development | 3.12E-12 | 2.13037448 | 1.30E-10 |
| GO:0001944 | vasculature development | 3.12E-12 | 2.13037448 | 1.30E-10 |
| GO:0002275 | myeloid cell activation involved in immune response | 3.28E-12 | 4.571428571 | 1.36E-10 |
| GO:0002758 | innate immune response-activating signal transduction | 4.24E-12 | 3.695040711 | 1.74E-10 |
| GO:0042119 | neutrophil activation | 4.34E-12 | 7.896103896 | 1.76E-10 |
| GO:0001568 | blood vessel development | 4.44E-12 | 2.159545204 | 1.79E-10 |
| GO:0050870 | positive regulation of T cell activation | 4.91E-12 | 3.291428571 | 1.95E-10 |
| GO:0002446 | neutrophil mediated immunity | 4.94E-12 | 8.43956044 | 1.95E-10 |
| GO:0051092 | positive regulation of NF-kappaB transcription factor activity | 5.12E-12 | 4.025157233 | 2.01E-10 |
| GO:0032755 | positive regulation of interleukin-6 production | 5.82E-12 | 4.777348777 | 2.26E-10 |
| GO:2000379 | positive regulation of reactive oxygen species metabolic process | 6.32E-12 | 4.46344207 | 2.42E-10 |
| GO:0002253 | activation of immune response | 6.34E-12 | 2.268424465 | 2.42E-10 |
| GO:0002757 | immune response-activating signal transduction | 7.58E-12 | 2.382293763 | 2.87E-10 |
| GO:0002764 | immune response-regulating signaling pathway | 8.51E-12 | 2.360462257 | 3.19E-10 |
| GO:0002687 | positive regulation of leukocyte migration | 9.39E-12 | 3.84962406 | 3.49E-10 |
| GO:0010604 | positive regulation of macromolecule metabolic process | 1.28E-11 | 1.443072265 | 4.71E-10 |
| GO:0006644 | phospholipid metabolic process | 1.74E-11 | 2.657283603 | 6.38E-10 |
| GO:0051403 | stress-activated MAPK cascade | 1.91E-11 | 3.069387755 | 6.95E-10 |
| GO:0009617 | response to bacterium | 1.98E-11 | 1.868389567 | 7.12E-10 |
| GO:0043123 | positive regulation of I-kappaB kinase/NF-kappaB signaling | 2.31E-11 | 4.022857143 | 8.24E-10 |
| GO:0006464 | cellular protein modification process | 3.58E-11 | 1.405879581 | 1.27E-09 |
| GO:0036211 | protein modification process | 3.71E-11 | 1.405166479 | 1.30E-09 |
| GO:0032940 | secretion by cell | 4.76E-11 | 1.962553442 | 1.66E-09 |
| GO:0001525 | angiogenesis | 5.38E-11 | 2.317907445 | 1.86E-09 |
| GO:0031401 | positive regulation of protein modification process | 5.50E-11 | 1.781684982 | 1.88E-09 |
| GO:0042093 | T-helper cell differentiation | 5.52E-11 | 5.830227743 | 1.88E-09 |
| GO:0002218 | activation of innate immune response | 5.81E-11 | 3.27386151 | 1.97E-09 |
| GO:1903530 | regulation of secretion by cell | 6.49E-11 | 2.142259414 | 2.18E-09 |
| GO:0006909 | phagocytosis | 6.55E-11 | 2.523594053 | 2.18E-09 |
| GO:0002294 | CD4-positive, alpha-beta T cell differentiation involved in immune response | 7.49E-11 | 5.746938776 | 2.48E-09 |
| GO:0070304 | positive regulation of stress-activated protein kinase signaling cascade | 8.15E-11 | 3.577639752 | 2.67E-09 |
| GO:0051046 | regulation of secretion | 8.66E-11 | 2.054320988 | 2.82E-09 |
| GO:0002532 | production of molecular mediator involved in inflammatory response | 8.81E-11 | 4.968944099 | 2.85E-09 |
| GO:0032655 | regulation of interleukin-12 production | 9.27E-11 | 6 | 2.97E-09 |
| GO:0009968 | negative regulation of signal transduction | 9.82E-11 | 1.790796808 | 3.13E-09 |
| GO:0002293 | alpha-beta T cell differentiation involved in immune response | 1.01E-10 | 5.665995976 | 3.19E-09 |
| GO:0002287 | alpha-beta T cell activation involved in immune response | 1.35E-10 | 5.587301587 | 4.25E-09 |
| GO:0002819 | regulation of adaptive immune response | 1.52E-10 | 3.12195122 | 4.73E-09 |
| GO:0007162 | negative regulation of cell adhesion | 1.64E-10 | 2.808777429 | 5.09E-09 |
| GO:0051130 | positive regulation of cellular component organization | 1.66E-10 | 1.69829153 | 5.12E-09 |
| GO:0031348 | negative regulation of defense response | 1.71E-10 | 3.059477487 | 5.23E-09 |
| GO:0042098 | T cell proliferation | 2.51E-10 | 3.180124224 | 7.62E-09 |
| GO:0001776 | leukocyte homeostasis | 2.66E-10 | 4.110741971 | 8.03E-09 |
| GO:1901653 | cellular response to peptide | 2.91E-10 | 2.605352431 | 8.72E-09 |

| | | | | |
|---|---|---|---|---|
| GO:0002700 | regulation of production of molecular mediator of immune response | 3.12E-10 | 3.277628032 | 9.27E-09 |
| GO:0045597 | positive regulation of cell differentiation | 3.29E-10 | 1.793007641 | 9.71E-09 |
| GO:1902532 | negative regulation of intracellular signal transduction | 3.57E-10 | 2.253521127 | 1.04E-08 |
| GO:0022408 | negative regulation of cell-cell adhesion | 3.58E-10 | 3.262240107 | 1.04E-08 |
| GO:0032102 | negative regulation of response to external stimulus | 3.61E-10 | 2.648627139 | 1.04E-08 |
| GO:1902105 | regulation of leukocyte differentiation | 4.86E-10 | 2.62667719 | 1.40E-08 |
| GO:0051347 | positive regulation of transferase activity | 6.85E-10 | 2.190634657 | 1.96E-08 |
| GO:0043434 | response to peptide hormone | 7.26E-10 | 2.356994357 | 2.06E-08 |
| GO:0046903 | secretion | 7.39E-10 | 1.80638772 | 2.09E-08 |
| GO:0010934 | macrophage cytokine production | 7.67E-10 | 6.453781513 | 2.15E-08 |
| GO:0055082 | cellular chemical homeostasis | 7.73E-10 | 1.919604205 | 2.15E-08 |
| GO:0045619 | regulation of lymphocyte differentiation | 8.07E-10 | 3.114160948 | 2.23E-08 |
| GO:0051345 | positive regulation of hydrolase activity | 8.32E-10 | 2.166975881 | 2.29E-08 |
| GO:0050728 | negative regulation of inflammatory response | 8.47E-10 | 3.704948646 | 2.32E-08 |
| GO:1901342 | regulation of vasculature development | 8.54E-10 | 2.557105164 | 2.32E-08 |
| GO:1904018 | positive regulation of vasculature development | 9.16E-10 | 3.158441558 | 2.47E-08 |
| GO:1990266 | neutrophil migration | 9.65E-10 | 3.899159664 | 2.59E-08 |
| GO:0031329 | regulation of cellular catabolic process | 1.22E-09 | 2.072555852 | 3.25E-08 |
| GO:1900015 | regulation of cytokine production involved in inflammatory response | 1.25E-09 | 5.888619855 | 3.32E-08 |
| GO:0097190 | apoptotic signaling pathway | 1.43E-09 | 2.127583109 | 3.77E-08 |
| GO:0045807 | positive regulation of endocytosis | 1.72E-09 | 3.211149826 | 4.52E-08 |
| GO:0045055 | regulated exocytosis | 1.95E-09 | 2.752030578 | 5.07E-08 |
| GO:0050777 | negative regulation of immune response | 2.26E-09 | 3.117840685 | 5.85E-08 |
| GO:0001666 | response to hypoxia | 2.40E-09 | 2.597101449 | 6.19E-08 |
| GO:1903038 | negative regulation of leukocyte cell-cell adhesion | 2.67E-09 | 3.632923368 | 6.84E-08 |
| GO:0043300 | regulation of leukocyte degranulation | 3.06E-09 | 5.603686636 | 7.78E-08 |
| GO:0002822 | regulation of adaptive immune response based on somatic recombination of immune receptors built from immunoglobulin superfamily domains | 3.24E-09 | 3.021118012 | 8.21E-08 |
| GO:0050801 | ion homeostasis | 3.32E-09 | 1.888854003 | 8.37E-08 |
| GO:0030162 | regulation of proteolysis | 3.40E-09 | 1.974293059 | 8.52E-08 |
| GO:0031331 | positive regulation of cellular catabolic process | 3.50E-09 | 2.432090077 | 8.72E-08 |
| GO:1903706 | regulation of hemopoiesis | 3.87E-09 | 2.27638484 | 9.58E-08 |
| GO:0051051 | negative regulation of transport | 4.39E-09 | 2.168394437 | 1.08E-07 |
| GO:0051050 | positive regulation of transport | 4.46E-09 | 1.779591837 | 1.09E-07 |
| GO:0038061 | NIK/NF-kappaB signaling | 4.48E-09 | 4.155844156 | 1.09E-07 |
| GO:0006873 | cellular ion homeostasis | 4.70E-09 | 2.001172562 | 1.14E-07 |
| GO:0050868 | negative regulation of T cell activation | 5.23E-09 | 3.737226277 | 1.26E-07 |
| GO:0070302 | regulation of stress-activated protein kinase signaling cascade | 5.33E-09 | 2.868347339 | 1.28E-07 |
| GO:0045621 | positive regulation of lymphocyte differentiation | 6.07E-09 | 3.607385811 | 1.44E-07 |
| GO:0030099 | myeloid cell differentiation | 6.20E-09 | 2.305590062 | 1.47E-07 |
| GO:1903426 | regulation of reactive oxygen species biosynthetic process | 8.38E-09 | 3.896955504 | 1.96E-07 |
| GO:0032652 | regulation of interleukin-1 production | 8.38E-09 | 3.896955504 | 1.96E-07 |
| GO:0002283 | neutrophil activation involved in immune response | 9.06E-09 | 9.540372671 | 2.11E-07 |
| GO:0045766 | positive regulation of angiogenesis | 1.06E-08 | 3.124192391 | 2.45E-07 |
| GO:0010506 | regulation of autophagy | 1.21E-08 | 2.629318394 | 2.80E-07 |
| GO:0032872 | regulation of stress-activated MAPK cascade | 1.25E-08 | 2.829931973 | 2.87E-07 |
| GO:0045862 | positive regulation of proteolysis | 1.27E-08 | 2.468319559 | 2.90E-07 |
| GO:0071375 | cellular response to peptide hormone stimulus | 1.32E-08 | 2.587601078 | 3.00E-07 |
| GO:0032695 | negative regulation of interleukin-12 production | 1.34E-08 | 10.58646617 | 3.02E-07 |
| GO:0007015 | actin filament organization | 1.85E-08 | 2.256029685 | 4.16E-07 |
| GO:0071219 | cellular response to molecule of bacterial origin | 2.25E-08 | 2.5108742 | 5.04E-07 |
| GO:0043269 | regulation of ion transport | 2.62E-08 | 1.897142857 | 5.83E-07 |
| GO:0044267 | cellular protein metabolic process | 2.68E-08 | 1.27647748 | 5.93E-07 |
| GO:0048513 | animal organ development | 3.07E-08 | 1.327837932 | 6.76E-07 |
| GO:0010563 | negative regulation of phosphorus metabolic process | 3.31E-08 | 2.073637703 | 7.22E-07 |
| GO:0045936 | negative regulation of phosphate metabolic process | 3.31E-08 | 2.073637703 | 7.22E-07 |
| GO:0009896 | positive regulation of catabolic process | 3.55E-08 | 2.195800059 | 7.70E-07 |
| GO:0008154 | actin polymerization or depolymerization | 3.68E-08 | 2.862111801 | 7.96E-07 |
| GO:0002720 | positive regulation of cytokine production involved in immune response | 4.15E-08 | 4.363636364 | 8.93E-07 |
| GO:0043254 | regulation of protein complex assembly | 4.19E-08 | 2.119092123 | 8.97E-07 |
| GO:0045765 | regulation of angiogenesis | 4.21E-08 | 2.459482038 | 8.97E-07 |
| GO:0051251 | positive regulation of lymphocyte activation | 4.33E-08 | 2.132750615 | 9.18E-07 |
| GO:0007169 | transmembrane receptor protein tyrosine kinase signaling pathway | 4.55E-08 | 2.05597469 | 9.61E-07 |
| GO:0007596 | blood coagulation | 4.95E-08 | 3.131807419 | 1.04E-06 |
| GO:0051248 | negative regulation of protein metabolic process | 5.54E-08 | 1.663641457 | 1.16E-06 |
| GO:0002886 | regulation of myeloid leukocyte mediated immunity | 6.44E-08 | 4.694980695 | 1.34E-06 |
| GO:0032956 | regulation of actin cytoskeleton organization | 6.96E-08 | 2.339425587 | 1.44E-06 |
| GO:0071417 | cellular response to organonitrogen compound | 6.98E-08 | 1.928872815 | 1.44E-06 |
| GO:0002702 | positive regulation of production of molecular mediator of immune response | 7.59E-08 | 3.404926108 | 1.56E-06 |
| GO:0071222 | cellular response to lipopolysaccharide | 7.82E-08 | 2.468010517 | 1.60E-06 |
| GO:0070665 | positive regulation of leukocyte proliferation | 8.22E-08 | 3.213852814 | 1.67E-06 |
| GO:0032611 | interleukin-1 beta production | 8.28E-08 | 4.022857143 | 1.67E-06 |
| GO:0010935 | regulation of macrophage cytokine production | 8.98E-08 | 5.962732919 | 1.81E-06 |
| GO:0043087 | regulation of GTPase activity | 1.02E-07 | 2.361067504 | 2.03E-06 |
| GO:0033003 | regulation of mast cell activation | 1.11E-07 | 5.096018735 | 2.22E-06 |
| GO:0042554 | superoxide anion generation | 1.21E-07 | 5.835866261 | 2.41E-06 |
| GO:1902563 | regulation of neutrophil activation | 1.30E-07 | 10.15873016 | 2.58E-06 |

| | | | | |
|---|---|---|---|---|
| GO:0006887 | exocytosis | 1.38E-07 | 2.221628838 | 2.72E-06 |
| GO:1902107 | positive regulation of leukocyte differentiation | 1.42E-07 | 2.813186813 | 2.78E-06 |
| GO:0008284 | positive regulation of cell proliferation | 1.57E-07 | 1.692208959 | 3.07E-06 |
| GO:0016485 | protein processing | 1.66E-07 | 2.499047619 | 3.22E-06 |
| GO:1901222 | regulation of NIK/NF-kappaB signaling | 1.69E-07 | 3.721871049 | 3.27E-06 |
| GO:0098542 | defense response to other organism | 1.92E-07 | 1.781474964 | 3.70E-06 |
| GO:0001959 | regulation of cytokine-mediated signaling pathway | 1.94E-07 | 3.348088531 | 3.72E-06 |
| GO:0070664 | negative regulation of leukocyte proliferation | 1.99E-07 | 3.831292517 | 3.81E-06 |
| GO:0042542 | response to hydrogen peroxide | 2.04E-07 | 3.160493827 | 3.88E-06 |
| GO:0032269 | negative regulation of cellular protein metabolic process | 2.25E-07 | 1.651319683 | 4.26E-06 |
| GO:0002456 | T cell mediated immunity | 2.30E-07 | 3.226890756 | 4.34E-06 |
| GO:1903556 | negative regulation of tumor necrosis factor superfamily cytokine production | 2.37E-07 | 4.571428571 | 4.45E-06 |
| GO:0030183 | B cell differentiation | 2.59E-07 | 3.047619048 | 4.84E-06 |
| GO:0050672 | negative regulation of lymphocyte proliferation | 2.74E-07 | 3.918367347 | 5.11E-06 |
| GO:0002705 | positive regulation of leukocyte mediated immunity | 2.83E-07 | 2.840499307 | 5.24E-06 |
| GO:0072359 | circulatory system development | 3.25E-07 | 1.620616246 | 6.00E-06 |
| GO:0032945 | negative regulation of mononuclear cell proliferation | 3.27E-07 | 3.878787879 | 6.00E-06 |
| GO:0001667 | ameboidal-type cell migration | 3.47E-07 | 2.118945432 | 6.35E-06 |
| GO:0032651 | regulation of interleukin-1 beta production | 3.87E-07 | 3.84 | 7.06E-06 |
| GO:0043030 | regulation of macrophage activation | 4.31E-07 | 4.958837772 | 7.83E-06 |
| GO:0045428 | regulation of nitric oxide biosynthetic process | 4.44E-07 | 4.388571429 | 8.03E-06 |
| GO:0046486 | glycerolipid metabolic process | 4.52E-07 | 2.243938829 | 8.12E-06 |
| GO:2000401 | regulation of lymphocyte migration | 4.53E-07 | 4.639658849 | 8.12E-06 |
| GO:0070555 | response to interleukin-1 | 4.70E-07 | 3.401993355 | 8.39E-06 |
| GO:0031663 | lipopolysaccharide-mediated signaling pathway | 4.87E-07 | 5.274725275 | 8.66E-06 |
| GO:0006809 | nitric oxide biosynthetic process | 5.02E-07 | 4.136054422 | 8.89E-06 |
| GO:0051129 | negative regulation of cellular component organization | 5.21E-07 | 1.754272815 | 9.20E-06 |
| GO:0002688 | regulation of leukocyte chemotaxis | 5.42E-07 | 3.375824176 | 9.52E-06 |
| GO:2001233 | regulation of apoptotic signaling pathway | 5.85E-07 | 2.138925295 | 1.02E-05 |
| GO:0031400 | negative regulation of protein modification process | 5.99E-07 | 1.905354366 | 1.04E-05 |
| GO:0051604 | protein maturation | 6.23E-07 | 2.292226292 | 1.08E-05 |
| GO:0043124 | negative regulation of I-kappaB kinase/NF-kappaB signaling | 6.29E-07 | 5.175202156 | 1.09E-05 |
| GO:1903825 | organic acid transmembrane transport | 6.38E-07 | 3.72815534 | 1.10E-05 |
| GO:0002526 | acute inflammatory response | 6.49E-07 | 3.148533586 | 1.11E-05 |
| GO:0008064 | regulation of actin polymerization or depolymerization | 6.50E-07 | 2.792399719 | 1.11E-05 |
| GO:0080171 | lytic vacuole organization | 6.62E-07 | 4.27458256 | 1.13E-05 |
| GO:0007252 | I-kappaB phosphorylation | 6.78E-07 | 8.707482993 | 1.15E-05 |
| GO:1903305 | regulation of regulated secretory pathway | 7.24E-07 | 2.778711485 | 1.22E-05 |
| GO:0043069 | negative regulation of programmed cell death | 7.82E-07 | 1.654740141 | 1.32E-05 |
| GO:0032715 | negative regulation of interleukin-6 production | 8.06E-07 | 5.079365079 | 1.35E-05 |
| GO:0072538 | T-helper 17 type immune response | 8.88E-07 | 5.446808511 | 1.48E-05 |
| GO:0030832 | regulation of actin filament length | 8.93E-07 | 2.751733703 | 1.49E-05 |
| GO:0030833 | regulation of actin filament polymerization | 9.35E-07 | 2.866409266 | 1.55E-05 |
| GO:0016236 | macroautophagy | 9.58E-07 | 2.549800797 | 1.58E-05 |
| GO:0010631 | epithelial cell migration | 1.07E-06 | 2.299401198 | 1.77E-05 |
| GO:0071621 | granulocyte chemotaxis | 1.08E-06 | 3.250793651 | 1.77E-05 |
| GO:0006508 | proteolysis | 1.25E-06 | 1.472876712 | 2.04E-05 |
| GO:0032653 | regulation of interleukin-10 production | 1.33E-06 | 4.571428571 | 2.17E-05 |
| GO:0015031 | protein transport | 1.39E-06 | 1.480666559 | 2.26E-05 |
| GO:0032479 | regulation of type I interferon production | 1.40E-06 | 3.555555556 | 2.26E-05 |
| GO:1903708 | positive regulation of hemopoiesis | 1.45E-06 | 2.465489567 | 2.33E-05 |
| GO:0002698 | negative regulation of immune effector process | 1.48E-06 | 2.93877551 | 2.37E-05 |
| GO:0042886 | amide transport | 1.52E-06 | 1.464036232 | 2.43E-05 |
| GO:0034341 | response to interferon-gamma | 1.54E-06 | 3.009041591 | 2.46E-05 |
| GO:0034614 | cellular response to reactive oxygen species | 1.56E-06 | 2.860335196 | 2.47E-05 |
| GO:0032649 | regulation of interferon-gamma production | 1.58E-06 | 3.285714286 | 2.50E-05 |
| GO:0060548 | negative regulation of cell death | 1.70E-06 | 1.590708479 | 2.68E-05 |
| GO:0061081 | positive regulation of myeloid leukocyte cytokine production involved in immune response | 2.11E-06 | 5.528239203 | 3.30E-05 |
| GO:0032733 | positive regulation of interleukin-10 production | 2.11E-06 | 5.528239203 | 3.30E-05 |
| GO:0031334 | positive regulation of protein complex assembly | 2.16E-06 | 2.292879534 | 3.37E-05 |
| GO:0032271 | regulation of protein polymerization | 2.25E-06 | 2.496844521 | 3.49E-05 |
| GO:0080135 | regulation of cellular response to stress | 2.25E-06 | 1.732100034 | 3.49E-05 |
| GO:0045453 | bone resorption | 2.28E-06 | 4.144761905 | 3.51E-05 |
| GO:0009620 | response to fungus | 2.28E-06 | 4.144761905 | 3.51E-05 |
| GO:0015833 | peptide transport | 2.34E-06 | 1.459657893 | 3.59E-05 |
| GO:0042088 | T-helper 1 type immune response | 2.45E-06 | 5.019607843 | 3.74E-05 |
| GO:0050854 | regulation of antigen receptor-mediated signaling pathway | 2.48E-06 | 4.36673774 | 3.77E-05 |
| GO:0072539 | T-helper 17 cell differentiation | 2.50E-06 | 6.704761905 | 3.78E-05 |
| GO:0043032 | positive regulation of macrophage activation | 2.50E-06 | 6.704761905 | 3.78E-05 |
| GO:0032868 | response to insulin | 2.51E-06 | 2.33958634 | 3.78E-05 |
| GO:0051493 | regulation of cytoskeleton organization | 2.57E-06 | 1.880123194 | 3.86E-05 |
| GO:0000902 | cell morphogenesis | 2.63E-06 | 1.605418287 | 3.93E-05 |
| GO:0007229 | integrin-mediated signaling pathway | 2.64E-06 | 3.297423888 | 3.93E-05 |
| GO:1905954 | positive regulation of lipid localization | 2.64E-06 | 3.297423888 | 3.93E-05 |
| GO:0006461 | protein complex assembly | 2.67E-06 | 1.587407584 | 3.95E-05 |
| GO:0001774 | microglial cell activation | 2.75E-06 | 5.402597403 | 4.07E-05 |

| | | | | |
|---|---|---|---|---|
| GO:0017157 | regulation of exocytosis | 2.86E-06 | 2.393766234 | 4.21E-05 |
| GO:0006820 | anion transport | 3.02E-06 | 1.848792598 | 4.44E-05 |
| GO:0006812 | cation transport | 3.56E-06 | 1.572481572 | 5.20E-05 |
| GO:1903034 | regulation of response to wounding | 3.57E-06 | 2.576623377 | 5.20E-05 |
| GO:0009891 | positive regulation of biosynthetic process | 3.97E-06 | 1.405941904 | 5.77E-05 |
| GO:0015711 | organic anion transport | 4.07E-06 | 1.969802555 | 5.90E-05 |
| GO:0032720 | negative regulation of tumor necrosis factor production | 4.43E-06 | 4.179591837 | 6.40E-05 |
| GO:0006650 | glycerophospholipid metabolic process | 4.69E-06 | 2.342653787 | 6.74E-05 |
| GO:0002313 | mature B cell differentiation involved in immune response | 4.86E-06 | 6.285714286 | 6.97E-05 |
| GO:0034220 | ion transmembrane transport | 5.03E-06 | 1.629420085 | 7.19E-05 |
| GO:0043066 | negative regulation of apoptotic process | 5.81E-06 | 1.601917188 | 8.27E-05 |
| GO:1902622 | regulation of neutrophil migration | 6.09E-06 | 4.654545455 | 8.65E-05 |
| GO:0030258 | lipid modification | 6.38E-06 | 2.344322344 | 9.03E-05 |
| GO:0097191 | extrinsic apoptotic signaling pathway | 6.46E-06 | 2.458583433 | 9.11E-05 |
| GO:0071396 | cellular response to lipid | 6.49E-06 | 1.726852707 | 9.13E-05 |
| GO:0032608 | interferon-beta production | 7.13E-06 | 4.285714286 | 9.99E-05 |
| GO:0002833 | positive regulation of response to biotic stimulus | 7.66E-06 | 4.007827789 | 1.07E-04 |
| GO:0050920 | regulation of chemotaxis | 7.67E-06 | 2.438095238 | 1.07E-04 |
| GO:0050671 | positive regulation of lymphocyte proliferation | 7.69E-06 | 2.906338694 | 1.07E-04 |
| GO:0044070 | regulation of anion transport | 8.12E-06 | 2.982776089 | 1.12E-04 |
| GO:0072676 | lymphocyte migration | 8.50E-06 | 3.294723295 | 1.17E-04 |
| GO:0009306 | protein secretion | 8.86E-06 | 2.031746032 | 1.22E-04 |
| GO:1901224 | positive regulation of NIK/NF-kappaB signaling | 9.14E-06 | 3.745266781 | 1.25E-04 |
| GO:0002534 | cytokine production involved in inflammatory response | 9.29E-06 | 4.49122807 | 1.27E-04 |
| GO:2000404 | regulation of T cell migration | 9.35E-06 | 4.851311953 | 1.27E-04 |
| GO:0009615 | response to virus | 1.02E-05 | 2.080120937 | 1.39E-04 |
| GO:0032731 | positive regulation of interleukin-1 beta production | 1.04E-05 | 4.155844156 | 1.41E-04 |
| GO:0032946 | positive regulation of mononuclear cell proliferation | 1.07E-05 | 2.849721707 | 1.45E-04 |
| GO:0002262 | myeloid cell homeostasis | 1.08E-05 | 2.585858586 | 1.45E-04 |
| GO:0090322 | regulation of superoxide metabolic process | 1.10E-05 | 5.224489796 | 1.48E-04 |
| GO:1902903 | regulation of supramolecular fiber organization | 1.12E-05 | 2.031746032 | 1.49E-04 |
| GO:0032480 | negative regulation of type I interferon production | 1.13E-05 | 6.530612245 | 1.50E-04 |
| GO:0007009 | plasma membrane organization | 1.13E-05 | 2.012303486 | 1.50E-04 |
| GO:0007167 | enzyme linked receptor protein signaling pathway | 1.14E-05 | 1.616209074 | 1.52E-04 |
| GO:0010508 | positive regulation of autophagy | 1.35E-05 | 2.737382378 | 1.78E-04 |
| GO:2001236 | regulation of extrinsic apoptotic signaling pathway | 1.35E-05 | 2.737382378 | 1.78E-04 |
| GO:0030168 | platelet activation | 1.37E-05 | 3.464661654 | 1.80E-04 |
| GO:0033619 | membrane protein proteolysis | 1.39E-05 | 4.338983051 | 1.82E-04 |
| GO:1902624 | positive regulation of neutrophil migration | 1.40E-05 | 5.102990033 | 1.84E-04 |
| GO:0030838 | positive regulation of actin filament polymerization | 1.43E-05 | 3.308843537 | 1.86E-04 |
| GO:0032648 | regulation of interferon-beta production | 1.50E-05 | 4.033613445 | 1.94E-04 |
| GO:2000191 | regulation of fatty acid transport | 1.57E-05 | 5.587301587 | 2.02E-04 |
| GO:0002719 | negative regulation of cytokine production involved in immune response | 1.57E-05 | 5.587301587 | 2.02E-04 |
| GO:0002706 | regulation of lymphocyte mediated immunity | 1.59E-05 | 2.391802291 | 2.05E-04 |
| GO:1903793 | positive regulation of anion transport | 1.78E-05 | 3.750915751 | 2.29E-04 |
| GO:0002888 | positive regulation of myeloid leukocyte mediated immunity | 1.80E-05 | 4.571428571 | 2.30E-04 |
| GO:0043313 | regulation of neutrophil degranulation | 1.82E-05 | 10.66666667 | 2.33E-04 |
| GO:0002821 | positive regulation of adaptive immune response | 1.83E-05 | 2.68907563 | 2.33E-04 |
| GO:0032732 | positive regulation of interleukin-1 production | 2.09E-05 | 3.703435805 | 2.65E-04 |
| GO:0032735 | positive regulation of interleukin-12 production | 2.24E-05 | 4.876190476 | 2.83E-04 |
| GO:0032660 | regulation of interleukin-17 production | 2.24E-05 | 4.876190476 | 2.83E-04 |
| GO:0061041 | regulation of wound healing | 2.41E-05 | 2.583850932 | 3.03E-04 |
| GO:0008285 | negative regulation of cell proliferation | 2.48E-05 | 1.666546112 | 3.11E-04 |
| GO:0032692 | negative regulation of interleukin-1 production | 2.64E-05 | 5.293233083 | 3.29E-04 |
| GO:0050869 | negative regulation of B cell activation | 2.64E-05 | 5.293233083 | 3.29E-04 |
| GO:0050708 | regulation of protein secretion | 2.78E-05 | 2.105627706 | 3.46E-04 |
| GO:0045123 | cellular extravasation | 2.85E-05 | 3.611992945 | 3.54E-04 |
| GO:0007264 | small GTPase mediated signal transduction | 2.89E-05 | 1.938117182 | 3.58E-04 |
| GO:0045806 | negative regulation of endocytosis | 2.96E-05 | 3.80952381 | 3.65E-04 |
| GO:0010634 | positive regulation of epithelial cell migration | 3.02E-05 | 2.675958188 | 3.72E-04 |
| GO:0048660 | regulation of smooth muscle cell proliferation | 3.05E-05 | 2.493506494 | 3.73E-04 |
| GO:0032890 | regulation of organic acid transport | 3.05E-05 | 3.416012559 | 3.73E-04 |
| GO:0031328 | positive regulation of cellular biosynthetic process | 3.06E-05 | 1.368613386 | 3.73E-04 |
| GO:1902905 | positive regulation of supramolecular fiber organization | 3.23E-05 | 2.388674389 | 3.93E-04 |
| GO:0002690 | positive regulation of leukocyte chemotaxis | 3.57E-05 | 3.226890756 | 4.33E-04 |
| GO:0051047 | positive regulation of secretion | 3.62E-05 | 1.920417482 | 4.38E-04 |
| GO:2000193 | positive regulation of fatty acid transport | 3.63E-05 | 6.582857143 | 4.38E-04 |
| GO:0043112 | receptor metabolic process | 3.94E-05 | 2.568218299 | 4.75E-04 |
| GO:0051223 | regulation of protein transport | 3.97E-05 | 1.738736699 | 4.76E-04 |
| GO:0052547 | regulation of peptidase activity | 3.97E-05 | 1.897193314 | 4.76E-04 |
| GO:0070486 | leukocyte aggregation | 4.02E-05 | 7.69924812 | 4.80E-04 |
| GO:0032273 | positive regulation of protein polymerization | 4.08E-05 | 2.774384236 | 4.86E-04 |
| GO:0008643 | carbohydrate transport | 4.10E-05 | 2.695970696 | 4.87E-04 |
| GO:0043302 | positive regulation of leukocyte degranulation | 4.28E-05 | 5.028571429 | 5.06E-04 |
| GO:0045622 | regulation of T-helper cell differentiation | 4.28E-05 | 5.028571429 | 5.06E-04 |
| GO:1900077 | negative regulation of cellular response to insulin stimulus | 4.29E-05 | 4.571428571 | 5.06E-04 |

| | | | | |
|---|---|---|---|---|
| GO:0022604 | regulation of cell morphogenesis | 4.41E-05 | 1.795324675 | 5.18E-04 |
| GO:0060284 | regulation of cell development | 4.45E-05 | 1.509544073 | 5.21E-04 |
| GO:0051495 | positive regulation of cytoskeleton organization | 4.47E-05 | 2.267428571 | 5.23E-04 |
| GO:0032870 | cellular response to hormone stimulus | 4.53E-05 | 1.750117869 | 5.29E-04 |
| GO:0009395 | phospholipid catabolic process | 4.79E-05 | 4.170426065 | 5.56E-04 |
| GO:0010628 | positive regulation of gene expression | 4.79E-05 | 1.347195482 | 5.56E-04 |
| GO:0043409 | negative regulation of MAPK cascade | 4.95E-05 | 2.476190476 | 5.73E-04 |
| GO:0032722 | positive regulation of chemokine production | 5.13E-05 | 3.442016807 | 5.92E-04 |
| GO:0030593 | neutrophil chemotaxis | 5.23E-05 | 3.134693878 | 6.01E-04 |
| GO:1903531 | negative regulation of secretion by cell | 5.40E-05 | 2.463360474 | 6.20E-04 |
| GO:0046942 | carboxylic acid transport | 5.68E-05 | 1.992993383 | 6.47E-04 |
| GO:1902106 | negative regulation of leukocyte differentiation | 5.68E-05 | 2.995073892 | 6.47E-04 |
| GO:0010720 | positive regulation of cell development | 5.68E-05 | 1.674044266 | 6.47E-04 |
| GO:0043373 | CD4-positive, alpha-beta T cell lineage commitment | 5.89E-05 | 7.314285714 | 6.69E-04 |
| GO:0071346 | cellular response to interferon-gamma | 5.97E-05 | 2.879640045 | 6.76E-04 |
| GO:0010543 | regulation of platelet activation | 6.42E-05 | 4.388571429 | 7.25E-04 |
| GO:0033006 | regulation of mast cell activation involved in immune response | 6.73E-05 | 4.789115646 | 7.59E-04 |
| GO:0015849 | organic acid transport | 6.77E-05 | 1.976833977 | 7.60E-04 |
| GO:1903532 | positive regulation of secretion by cell | 6.77E-05 | 1.976833977 | 7.60E-04 |
| GO:0046434 | organophosphate catabolic process | 7.31E-05 | 2.596119929 | 8.16E-04 |
| GO:0008360 | regulation of cell shape | 7.31E-05 | 2.596119929 | 8.16E-04 |
| GO:0002790 | peptide secretion | 7.60E-05 | 1.832388905 | 8.46E-04 |
| GO:0090407 | organophosphate biosynthetic process | 7.76E-05 | 1.768508863 | 8.61E-04 |
| GO:0048661 | positive regulation of smooth muscle cell proliferation | 7.99E-05 | 2.919567827 | 8.85E-04 |
| GO:0002824 | positive regulation of adaptive immune response based on somatic recombination of immune receptors built from immunoglobulin superfamily domains | 8.03E-05 | 2.580192813 | 8.87E-04 |
| GO:0050832 | defense response to fungus | 8.14E-05 | 3.961904762 | 8.97E-04 |
| GO:1900026 | positive regulation of substrate adhesion-dependent cell spreading | 8.34E-05 | 4.677740864 | 9.15E-04 |
| GO:0046627 | negative regulation of insulin receptor signaling pathway | 8.34E-05 | 4.677740864 | 9.15E-04 |
| GO:0070498 | interleukin-1-mediated signaling pathway | 8.42E-05 | 6.965986395 | 9.21E-04 |
| GO:0032688 | negative regulation of interferon-beta production | 8.56E-05 | 8.533333333 | 9.32E-04 |
| GO:0032494 | response to peptidoglycan | 8.56E-05 | 8.533333333 | 9.32E-04 |
| GO:0002228 | natural killer cell mediated immunity | 8.88E-05 | 3.287319422 | 9.64E-04 |
| GO:0070371 | ERK1 and ERK2 cascade | 9.09E-05 | 2.013605442 | 9.85E-04 |
| GO:0010811 | positive regulation of cell-substrate adhesion | 9.19E-05 | 2.704225352 | 9.93E-04 |
| GO:0070301 | cellular response to hydrogen peroxide | 9.52E-05 | 2.992207792 | 0.0010242 |
| GO:2001235 | positive regulation of apoptotic signaling pathway | 9.53E-05 | 2.431610942 | 0.0010242 |
| GO:0032757 | positive regulation of interleukin-8 production | 9.64E-05 | 3.896955504 | 0.0010323 |
| GO:1902565 | positive regulation of neutrophil activation | 9.65E-05 | 10.97142857 | 0.0010323 |
| GO:0032088 | negative regulation of NF-kappaB transcription factor activity | 9.94E-05 | 3.428571429 | 0.00106137 |
| GO:0034762 | regulation of transmembrane transport | 1.01E-04 | 1.760846561 | 0.00107506 |
| GO:0051048 | negative regulation of secretion | 1.05E-04 | 2.275555556 | 0.00111402 |
| GO:1903428 | positive regulation of reactive oxygen species biosynthetic process | 1.08E-04 | 3.605633803 | 0.00114823 |
| GO:0009914 | hormone transport | 1.09E-04 | 1.896296296 | 0.00115516 |
| GO:1903317 | regulation of protein maturation | 1.12E-04 | 3.077793494 | 0.00118449 |
| GO:0042267 | natural killer cell mediated cytotoxicity | 1.14E-04 | 3.386243386 | 0.0012022 |
| GO:0046879 | hormone secretion | 1.25E-04 | 1.902828136 | 0.00130775 |
| GO:2000021 | regulation of ion homeostasis | 1.28E-04 | 2.20952381 | 0.00134531 |
| GO:0001771 | immunological synapse formation | 1.31E-04 | 8 | 0.00136242 |
| GO:1903978 | regulation of microglial cell activation | 1.31E-04 | 8 | 0.00136242 |
| GO:0008654 | phospholipid biosynthetic process | 1.33E-04 | 2.380952381 | 0.00137879 |
| GO:0032642 | regulation of chemokine production | 1.34E-04 | 2.912768647 | 0.00138788 |
| GO:1904407 | positive regulation of nitric oxide metabolic process | 1.35E-04 | 4.063492063 | 0.00139351 |
| GO:0090087 | regulation of peptide transport | 1.41E-04 | 1.652486772 | 0.00145101 |
| GO:0070542 | response to fatty acid | 1.42E-04 | 3.018030513 | 0.00146659 |
| GO:0051173 | positive regulation of nitrogen compound metabolic process | 1.46E-04 | 1.321957791 | 0.00149623 |
| GO:0010595 | positive regulation of endothelial cell migration | 1.49E-04 | 2.887218045 | 0.00153165 |
| GO:0032892 | positive regulation of organic acid transport | 1.60E-04 | 3.98961039 | 0.00163778 |
| GO:0002791 | regulation of peptide secretion | 1.61E-04 | 1.917050691 | 0.00164325 |
| GO:0070201 | regulation of establishment of protein localization | 1.62E-04 | 1.65232358 | 0.00164862 |
| GO:0002675 | positive regulation of acute inflammatory response | 1.62E-04 | 4.812030075 | 0.00164862 |
| GO:0034765 | regulation of ion transmembrane transport | 1.76E-04 | 1.744820065 | 0.00178269 |
| GO:0006865 | amino acid transport | 1.78E-04 | 2.445182724 | 0.00180143 |
| GO:0010959 | regulation of metal ion transport | 1.79E-04 | 1.793481572 | 0.00180916 |
| GO:0050820 | positive regulation of coagulation | 1.85E-04 | 4.279635258 | 0.00185989 |
| GO:0019751 | polyol metabolic process | 1.85E-04 | 2.837438424 | 0.00186199 |
| GO:0002295 | T-helper cell lineage commitment | 1.93E-04 | 7.529411765 | 0.00193124 |
| GO:0019216 | regulation of lipid metabolic process | 1.95E-04 | 1.846153846 | 0.00195321 |
| GO:0051056 | regulation of small GTPase mediated signal transduction | 1.96E-04 | 2.155632985 | 0.0019588 |
| GO:0048535 | lymph node development | 1.98E-04 | 5.30875576 | 0.00196668 |
| GO:0034446 | substrate adhesion-dependent cell spreading | 2.01E-04 | 2.932614555 | 0.00199008 |
| GO:0043304 | regulation of mast cell degranulation | 2.01E-04 | 4.688644689 | 0.00199008 |
| GO:0002768 | immune response-regulating cell surface receptor signaling pathway | 2.06E-04 | 1.841726619 | 0.00203304 |
| GO:0031324 | negative regulation of cellular metabolic process | 2.08E-04 | 1.267201267 | 0.00205204 |
| GO:1903707 | negative regulation of hemopoiesis | 2.11E-04 | 2.417077176 | 0.00207337 |
| GO:0030163 | protein catabolic process | 2.14E-04 | 1.495618306 | 0.00209762 |
| GO:0045063 | T-helper 1 cell differentiation | 2.17E-04 | 6.095238095 | 0.00212582 |

| | | | | |
|---|---|---|---|---|
| GO:0002279 | mast cell activation involved in immune response | 2.21E-04 | 3.368421053 | 0.0021596 |
| GO:0010557 | positive regulation of macromolecule biosynthetic process | 2.22E-04 | 1.335265141 | 0.0021596 |
| GO:0002429 | immune response-activating cell surface receptor signaling pathway | 2.22E-04 | 1.851146384 | 0.0021596 |
| GO:0097193 | intrinsic apoptotic signaling pathway | 2.24E-04 | 1.992673993 | 0.00217836 |
| GO:0042742 | defense response to bacterium | 2.28E-04 | 1.685031692 | 0.00221487 |
| GO:0042832 | defense response to protozoan | 2.47E-04 | 4.571428571 | 0.0023859 |
| GO:0045893 | positive regulation of transcription, DNA-templated | 2.58E-04 | 1.364120782 | 0.00248854 |
| GO:0002701 | negative regulation of production of molecular mediator of immune response | 2.65E-04 | 4.104956268 | 0.0025512 |
| GO:0050856 | regulation of T cell receptor signaling pathway | 2.65E-04 | 4.104956268 | 0.0025512 |
| GO:0070163 | regulation of adiponectin secretion | 2.71E-04 | 13.06122449 | 0.00259736 |
| GO:0070162 | adiponectin secretion | 2.71E-04 | 13.06122449 | 0.00259736 |
| GO:0070423 | nucleotide-binding oligomerization domain containing signaling pathway | 2.76E-04 | 7.111111111 | 0.00263242 |
| GO:0002448 | mast cell mediated immunity | 2.83E-04 | 3.495798319 | 0.00269853 |
| GO:0010507 | negative regulation of autophagy | 2.89E-04 | 3.282051282 | 0.00274408 |
| GO:0010632 | regulation of epithelial cell migration | 2.93E-04 | 2.10430839 | 0.00278159 |
| GO:0032370 | positive regulation of lipid transport | 3.01E-04 | 2.955266955 | 0.00284215 |
| GO:0030193 | regulation of blood coagulation | 3.01E-04 | 2.955266955 | 0.00284215 |
| GO:0032368 | regulation of lipid transport | 3.06E-04 | 2.477419355 | 0.0028839 |
| GO:1900024 | regulation of substrate adhesion-dependent cell spreading | 3.07E-04 | 3.719128329 | 0.0028839 |
| GO:1902680 | positive regulation of RNA biosynthetic process | 3.11E-04 | 1.356890459 | 0.00291541 |
| GO:1905521 | regulation of macrophage migration | 3.16E-04 | 4.022857143 | 0.0029578 |
| GO:0031344 | regulation of cell projection organization | 3.18E-04 | 1.540077953 | 0.00297389 |
| GO:0051607 | defense response to virus | 3.20E-04 | 2.003913894 | 0.00298626 |
| GO:0034612 | response to tumor necrosis factor | 3.26E-04 | 2.124481328 | 0.00302854 |
| GO:0032663 | regulation of interleukin-2 production | 3.26E-04 | 3.445134576 | 0.00302854 |
| GO:0032677 | regulation of interleukin-8 production | 3.29E-04 | 3.240506329 | 0.00304457 |
| GO:0015908 | fatty acid transport | 3.44E-04 | 2.800514801 | 0.00317828 |
| GO:0010803 | regulation of tumor necrosis factor-mediated signaling pathway | 3.65E-04 | 4.353741497 | 0.00335788 |
| GO:2000403 | positive regulation of lymphocyte migration | 3.65E-04 | 4.353741497 | 0.00335788 |
| GO:0051254 | positive regulation of RNA metabolic process | 3.67E-04 | 1.343482093 | 0.00336966 |
| GO:0002753 | cytoplasmic pattern recognition receptor signaling pathway | 3.74E-04 | 3.943977591 | 0.00341392 |
| GO:0006023 | aminoglycan biosynthetic process | 3.74E-04 | 3.395918367 | 0.00341392 |
| GO:0032303 | regulation of icosanoid secretion | 3.75E-04 | 5.626373626 | 0.00341392 |
| GO:0032930 | positive regulation of superoxide anion generation | 3.75E-04 | 5.626373626 | 0.00341392 |
| GO:0032963 | collagen metabolic process | 3.78E-04 | 2.675958188 | 0.00343395 |
| GO:0035872 | nucleotide-binding domain, leucine rich repeat containing receptor signaling pathway | 3.84E-04 | 6.736842105 | 0.00348987 |
| GO:0050921 | positive regulation of chemotaxis | 3.95E-04 | 2.430379747 | 0.00357791 |
| GO:0045620 | negative regulation of lymphocyte differentiation | 4.15E-04 | 3.597189696 | 0.00374796 |
| GO:0000768 | syncytium formation by plasma membrane fusion | 4.28E-04 | 3.348088531 | 0.00386445 |
| GO:0010940 | positive regulation of necrotic cell death | 4.29E-04 | 8.43956044 | 0.00386445 |
| GO:0050851 | antigen receptor-mediated signaling pathway | 4.38E-04 | 1.858712716 | 0.00393528 |
| GO:0045429 | positive regulation of nitric oxide biosynthetic process | 4.41E-04 | 3.868131868 | 0.0039503 |
| GO:0046883 | regulation of hormone secretion | 4.72E-04 | 1.912967033 | 0.00422535 |
| GO:1902582 | single-organism intracellular transport | 4.74E-04 | 1.516688919 | 0.00423441 |
| GO:0002673 | regulation of acute inflammatory response | 4.77E-04 | 3.12195122 | 0.00425471 |
| GO:0006691 | leukotriene metabolic process | 4.82E-04 | 5.417989418 | 0.00428228 |
| GO:2001234 | negative regulation of apoptotic signaling pathway | 4.84E-04 | 2.072874494 | 0.00428228 |
| GO:0043270 | positive regulation of ion transport | 4.84E-04 | 1.848555816 | 0.00428228 |
| GO:0051271 | negative regulation of cellular component movement | 4.84E-04 | 1.848555816 | 0.00428228 |
| GO:2000406 | positive regulation of T cell migration | 4.85E-04 | 4.702040816 | 0.00428228 |
| GO:1901214 | regulation of neuron death | 4.89E-04 | 1.797121797 | 0.00430545 |
| GO:0008630 | intrinsic apoptotic signaling pathway in response to DNA damage | 5.15E-04 | 2.70310559 | 0.00452395 |
| GO:0032495 | response to muramyl dipeptide | 5.24E-04 | 6.4 | 0.00458723 |
| GO:0030220 | platelet formation | 5.24E-04 | 6.4 | 0.00458723 |
| GO:0051208 | sequestering of calcium ion | 5.51E-04 | 2.59167604 | 0.00481004 |
| GO:0046834 | lipid phosphorylation | 5.53E-04 | 3.482993197 | 0.00481004 |
| GO:0030865 | cortical cytoskeleton organization | 5.53E-04 | 3.482993197 | 0.00481004 |
| GO:0006949 | syncytium formation | 5.57E-04 | 3.256360078 | 0.00482466 |
| GO:0034121 | regulation of toll-like receptor signaling pathway | 5.57E-04 | 3.256360078 | 0.00482466 |
| GO:0071674 | mononuclear cell migration | 5.65E-04 | 2.917933131 | 0.00488848 |
| GO:0002708 | positive regulation of lymphocyte mediated immunity | 5.93E-04 | 2.355828221 | 0.00511778 |
| GO:0032740 | positive regulation of interleukin-17 production | 6.13E-04 | 5.224489796 | 0.00527972 |
| GO:0006869 | lipid transport | 6.17E-04 | 1.759892689 | 0.00530763 |
| GO:0043316 | cytotoxic T cell degranulation | 6.24E-04 | 18.28571429 | 0.00534875 |
| GO:0002369 | T cell cytokine production | 6.25E-04 | 4.063492063 | 0.00534875 |
| GO:0001562 | response to protozoan | 6.25E-04 | 4.063492063 | 0.00534875 |
| GO:0098792 | xenophagy | 6.38E-04 | 7.836734694 | 0.00544316 |
| GO:0050767 | regulation of neurogenesis | 6.50E-04 | 1.454953691 | 0.00553945 |
| GO:0021782 | glial cell development | 6.60E-04 | 2.551495017 | 0.0056161 |
| GO:1903725 | regulation of phospholipid metabolic process | 6.80E-04 | 3.011764706 | 0.00576815 |
| GO:0032305 | positive regulation of icosanoid secretion | 7.00E-04 | 6.095238095 | 0.00591685 |
| GO:0060099 | regulation of phagocytosis, engulfment | 7.00E-04 | 6.095238095 | 0.00591685 |
| GO:0045124 | regulation of bone resorption | 7.02E-04 | 3.657142857 | 0.00592472 |
| GO:0033209 | tumor necrosis factor-mediated signaling pathway | 7.15E-04 | 3.16952381 | 0.00601113 |
| GO:0044744 | protein targeting to nucleus | 7.18E-04 | 2.126245847 | 0.00601113 |
| GO:0006606 | protein import into nucleus | 7.18E-04 | 2.126245847 | 0.00601113 |

| | | | | |
|---|---|---|---|---|
| GO:1902593 | single-organism nuclear import | 7.18E-04 | 2.126245847 | 0.00601113 |
| GO:1903364 | positive regulation of cellular protein catabolic process | 7.43E-04 | 2.44668008 | 0.00620737 |
| GO:0051767 | nitric-oxide synthase biosynthetic process | 7.70E-04 | 5.044334975 | 0.00641022 |
| GO:0051769 | regulation of nitric-oxide synthase biosynthetic process | 7.70E-04 | 5.044334975 | 0.00641022 |
| GO:0061515 | myeloid cell development | 7.79E-04 | 2.827687776 | 0.00647507 |
| GO:0035296 | regulation of tube diameter | 7.94E-04 | 2.247406225 | 0.0065723 |
| GO:0050880 | regulation of blood vessel size | 7.94E-04 | 2.247406225 | 0.0065723 |
| GO:0050830 | defense response to Gram-positive bacterium | 8.29E-04 | 2.19047619 | 0.00684944 |
| GO:0043303 | mast cell degranulation | 8.30E-04 | 3.324675325 | 0.00684944 |
| GO:0009100 | glycoprotein metabolic process | 8.43E-04 | 1.760846561 | 0.00694487 |
| GO:0045017 | glycerolipid biosynthetic process | 8.53E-04 | 2.234920635 | 0.00701433 |
| GO:0016482 | cytosolic transport | 8.71E-04 | 2.285714286 | 0.00710019 |
| GO:0090505 | epiboly involved in wound healing | 8.72E-04 | 4.330827068 | 0.00710019 |
| GO:0030866 | cortical actin cytoskeleton organization | 8.72E-04 | 4.330827068 | 0.00710019 |
| GO:0044319 | wound healing, spreading of cells | 8.72E-04 | 4.330827068 | 0.00710019 |
| GO:0071248 | cellular response to metal ion | 8.74E-04 | 2.096985583 | 0.00710019 |
| GO:0097696 | STAT cascade | 8.74E-04 | 2.096985583 | 0.00710019 |
| GO:0044724 | single-organism carbohydrate catabolic process | 8.76E-04 | 2.412698413 | 0.00711143 |
| GO:0032763 | regulation of mast cell cytokine production | 8.95E-04 | 10.15873016 | 0.00720506 |
| GO:1904415 | regulation of xenophagy | 8.95E-04 | 10.15873016 | 0.00720506 |
| GO:1904417 | positive regulation of xenophagy | 8.95E-04 | 10.15873016 | 0.00720506 |
| GO:0002291 | T cell activation via T cell receptor contact with antigen bound to MHC molecule on antigen presenting cell | 8.95E-04 | 10.15873016 | 0.00720506 |
| GO:1905153 | regulation of membrane invagination | 9.18E-04 | 5.818181818 | 0.00735557 |
| GO:0045624 | positive regulation of T-helper cell differentiation | 9.18E-04 | 5.818181818 | 0.00735557 |
| GO:0036344 | platelet morphogenesis | 9.18E-04 | 5.818181818 | 0.00735557 |
| GO:0090257 | regulation of muscle system process | 9.34E-04 | 1.956773853 | 0.00746916 |
| GO:0019362 | pyridine nucleotide metabolic process | 9.38E-04 | 2.272189349 | 0.00748869 |
| GO:0002009 | morphogenesis of an epithelium | 9.67E-04 | 1.591367582 | 0.00770104 |
| GO:0043623 | cellular protein complex assembly | 9.71E-04 | 1.552813219 | 0.00771895 |
| GO:0007265 | Ras protein signal transduction | 0.00104103 | 1.807909605 | 0.0082648 |
| GO:0010939 | regulation of necrotic cell death | 0.00104499 | 4.21978022 | 0.00828138 |
| GO:0016311 | dephosphorylation | 0.00105147 | 1.788819876 | 0.00831787 |
| GO:1901216 | positive regulation of neuron death | 0.00110859 | 2.438095238 | 0.00875412 |
| GO:0030203 | glycosaminoglycan metabolic process | 0.00113377 | 2.612244898 | 0.00893704 |
| GO:0051155 | positive regulation of striated muscle cell differentiation | 0.00114809 | 3.009041591 | 0.00903384 |
| GO:2001237 | negative regulation of extrinsic apoptotic signaling pathway | 0.0011677 | 2.715700141 | 0.00917184 |
| GO:0002507 | tolerance induction | 0.00117884 | 4.718894009 | 0.00924293 |
| GO:1900016 | negative regulation of cytokine production involved in inflammatory response | 0.00118538 | 5.565217391 | 0.00926142 |
| GO:0061760 | antifungal innate immune response | 0.00118538 | 5.565217391 | 0.00926142 |
| GO:0051346 | negative regulation of hydrolase activity | 0.00121285 | 1.741496599 | 0.00944875 |
| GO:0036294 | cellular response to decreased oxygen levels | 0.00121362 | 2.174517375 | 0.00944875 |
| GO:0051348 | negative regulation of transferase activity | 0.00123633 | 1.921325052 | 0.00958849 |
| GO:0071675 | regulation of mononuclear cell migration | 0.00123806 | 3.409200969 | 0.00958849 |
| GO:0043647 | inositol phosphate metabolic process | 0.00123806 | 3.409200969 | 0.00958849 |
| GO:0006937 | regulation of muscle contraction | 0.00134652 | 2.206896552 | 0.01041024 |
| GO:0048732 | gland development | 0.00138257 | 1.6 | 0.01067038 |
| GO:0051282 | regulation of sequestering of calcium ion | 0.00140154 | 2.467120181 | 0.01079792 |
| GO:0098739 | import across plasma membrane | 0.00140837 | 2.316190476 | 0.01083172 |
| GO:0032667 | regulation of interleukin-23 production | 0.00142657 | 9.142857143 | 0.01091489 |
| GO:0032762 | mast cell cytokine production | 0.00142657 | 9.142857143 | 0.01091489 |
| GO:0002604 | regulation of dendritic cell antigen processing and presentation | 0.00142657 | 9.142857143 | 0.01091489 |
| GO:0051960 | regulation of nervous system development | 0.00143274 | 1.393044784 | 0.01092181 |
| GO:0001885 | endothelial cell development | 0.00143487 | 2.934744268 | 0.01092181 |
| GO:0034333 | adherens junction assembly | 0.00143487 | 2.934744268 | 0.01092181 |
| GO:0002360 | T cell lineage commitment | 0.00143866 | 4.571428571 | 0.01093186 |
| GO:0002312 | B cell activation involved in immune response | 0.00144495 | 2.782608696 | 0.01096086 |
| GO:1904149 | regulation of microglial cell mediated cytotoxicity | 0.00149644 | 14.62857143 | 0.01133199 |
| GO:0031579 | membrane raft organization | 0.0015082 | 5.333333333 | 0.01140161 |
| GO:0046474 | glycerophospholipid biosynthetic process | 0.00153297 | 2.367934224 | 0.01156907 |
| GO:0042982 | amyloid precursor protein metabolic process | 0.00154409 | 3.09054326 | 0.0116332 |
| GO:1902882 | regulation of response to oxidative stress | 0.00155595 | 2.637362637 | 0.01170268 |
| GO:0001894 | tissue homeostasis | 0.0015688 | 1.917602996 | 0.01177927 |
| GO:0046626 | regulation of insulin receptor signaling pathway | 0.00159902 | 2.898954704 | 0.01196837 |
| GO:0010043 | response to zinc ion | 0.00160114 | 3.585434174 | 0.01196837 |
| GO:0001501 | skeletal system development | 0.00160209 | 1.581809194 | 0.01196837 |
| GO:0071398 | cellular response to fatty acid | 0.00161025 | 3.297423888 | 0.01200914 |
| GO:0022612 | gland morphogenesis | 0.00163766 | 2.285714286 | 0.01219298 |
| GO:0045935 | positive regulation of nucleobase-containing compound metabolic process | 0.00170694 | 1.27554007 | 0.01268747 |
| GO:1905155 | positive regulation of membrane invagination | 0.00171837 | 6.453781513 | 0.01272976 |
| GO:0060100 | positive regulation of phagocytosis, engulfment | 0.00171837 | 6.453781513 | 0.01272976 |
| GO:0045601 | regulation of endothelial cell differentiation | 0.00173422 | 3.918367347 | 0.01280538 |
| GO:1903307 | positive regulation of regulated secretory pathway | 0.00173436 | 3.047619048 | 0.01280538 |
| GO:0050858 | negative regulation of antigen receptor-mediated signaling pathway | 0.00174106 | 4.432900433 | 0.01281492 |
| GO:0051222 | positive regulation of protein transport | 0.00174144 | 1.771265771 | 0.01281492 |
| GO:0001912 | positive regulation of leukocyte mediated cytotoxicity | 0.00176329 | 2.723404255 | 0.01295417 |
| GO:0032388 | positive regulation of intracellular transport | 0.00178398 | 1.928571429 | 0.01308451 |

| | | | | |
|---|---|---|---|---|
| GO:0060760 | positive regulation of response to cytokine stimulus | 0.00182771 | 3.244239631 | 0.01338309 |
| GO:1905037 | autophagosome organization | 0.0018703 | 2.587601078 | 0.01364982 |
| GO:1903036 | positive regulation of response to wounding | 0.0018703 | 2.587601078 | 0.01364982 |
| GO:0097396 | response to interleukin-17 | 0.0018938 | 5.12 | 0.01377595 |
| GO:1901739 | regulation of myoblast fusion | 0.0018938 | 5.12 | 0.01377595 |
| GO:1900407 | regulation of cellular response to oxidative stress | 0.00194296 | 2.694736842 | 0.01408997 |
| GO:0032481 | positive regulation of type I interferon production | 0.00194333 | 3.005870841 | 0.01408997 |
| GO:0010594 | regulation of endothelial cell migration | 0.0020281 | 2.133333333 | 0.01464834 |
| GO:2000249 | regulation of actin cytoskeleton reorganization | 0.00203027 | 3.827242525 | 0.01464834 |
| GO:0046839 | phospholipid dephosphorylation | 0.00203027 | 3.827242525 | 0.01464834 |
| GO:0008217 | regulation of blood pressure | 0.00204826 | 1.97044335 | 0.01473019 |
| GO:0045732 | positive regulation of protein catabolic process | 0.00204826 | 1.97044335 | 0.01473019 |
| GO:0007611 | learning or memory | 0.00207091 | 1.769585253 | 0.0148689 |
| GO:0001952 | regulation of cell-matrix adhesion | 0.00210867 | 2.372955289 | 0.01511557 |
| GO:0007520 | myoblast fusion | 0.00211557 | 3.450134771 | 0.01514049 |
| GO:0002437 | inflammatory response to antigenic stimulus | 0.00218653 | 2.796638655 | 0.01562313 |
| GO:0032386 | regulation of intracellular transport | 0.0021909 | 1.582741384 | 0.01562913 |
| GO:0002820 | negative regulation of adaptive immune response | 0.00233372 | 3.142857143 | 0.01654797 |
| GO:0060348 | bone development | 0.00233542 | 1.891625616 | 0.01654797 |
| GO:0033043 | regulation of organelle organization | 0.00233749 | 1.321115077 | 0.01654797 |
| GO:0036336 | dendritic cell migration | 0.00234959 | 4.923076923 | 0.01654797 |
| GO:0050765 | negative regulation of phagocytosis | 0.00234959 | 4.923076923 | 0.01654797 |
| GO:0050860 | negative regulation of T cell receptor signaling pathway | 0.00234959 | 4.923076923 | 0.01654797 |
| GO:1903306 | negative regulation of regulated secretory pathway | 0.00234959 | 4.923076923 | 0.01654797 |
| GO:0050857 | positive regulation of antigen receptor-mediated signaling pathway | 0.00234959 | 4.923076923 | 0.01654797 |
| GO:0030194 | positive regulation of blood coagulation | 0.00236466 | 3.74025974 | 0.01660136 |
| GO:1900048 | positive regulation of hemostasis | 0.00236466 | 3.74025974 | 0.01660136 |
| GO:0030336 | negative regulation of cell migration | 0.00237413 | 1.793851718 | 0.01661515 |
| GO:0051402 | neuron apoptotic process | 0.00237413 | 1.793851718 | 0.01661515 |
| GO:0009101 | glycoprotein biosynthetic process | 0.00240021 | 1.754152824 | 0.01677119 |
| GO:1904705 | regulation of vascular smooth muscle cell proliferation | 0.00242275 | 2.925714286 | 0.016902 |
| GO:1903319 | positive regulation of protein maturation | 0.00249189 | 4.179591837 | 0.01735706 |
| GO:2000146 | negative regulation of cell motility | 0.00251274 | 1.767803194 | 0.0174748 |
| GO:0045596 | negative regulation of cell differentiation | 0.00252894 | 1.418719212 | 0.01755986 |
| GO:0071363 | cellular response to growth factor stimulus | 0.00253643 | 1.476639108 | 0.01758433 |
| GO:0046850 | regulation of bone remodeling | 0.00262582 | 3.094505495 | 0.01817557 |
| GO:0010001 | glial cell differentiation | 0.00264393 | 1.848375451 | 0.01827239 |
| GO:0032729 | positive regulation of interferon-gamma production | 0.00266825 | 2.732348112 | 0.01841174 |
| GO:0010605 | negative regulation of macromolecule metabolic process | 0.00285366 | 1.205584242 | 0.01966049 |
| GO:0032845 | negative regulation of homeostatic process | 0.00286433 | 1.920768307 | 0.01970337 |
| GO:1901223 | negative regulation of NIK/NF-kappaB signaling | 0.00288319 | 4.740740741 | 0.0198024 |
| GO:1901550 | regulation of endothelial cell development | 0.00294781 | 5.77443609 | 0.02015249 |
| GO:0045779 | negative regulation of bone resorption | 0.00294781 | 5.77443609 | 0.02015249 |
| GO:1901741 | positive regulation of myoblast fusion | 0.00294781 | 5.77443609 | 0.02015249 |
| GO:0006733 | oxidoreduction coenzyme metabolic process | 0.00298099 | 2.064516129 | 0.02034791 |
| GO:0045625 | regulation of T-helper 1 cell differentiation | 0.0030796 | 7.619047619 | 0.02098864 |
| GO:0005996 | monosaccharide metabolic process | 0.00313569 | 1.781076067 | 0.02133811 |
| GO:0097300 | programmed necrotic cell death | 0.00316183 | 3.577639752 | 0.02148301 |
| GO:0044088 | regulation of vacuole organization | 0.00329755 | 3.002132196 | 0.02237079 |
| GO:1990874 | vascular smooth muscle cell proliferation | 0.00331701 | 2.813186813 | 0.02246844 |
| GO:0007259 | JAK-STAT cascade | 0.00335717 | 2.001421464 | 0.02270572 |
| GO:0044242 | cellular lipid catabolic process | 0.00336608 | 1.896858328 | 0.0227313 |
| GO:0051962 | positive regulation of nervous system development | 0.00342002 | 1.477633478 | 0.02306039 |
| GO:0033005 | positive regulation of mast cell activation | 0.00346966 | 3.953667954 | 0.0232888 |
| GO:0090022 | regulation of neutrophil chemotaxis | 0.00346966 | 3.953667954 | 0.0232888 |
| GO:1902837 | amino acid import into cell | 0.00346966 | 3.953667954 | 0.0232888 |
| GO:0034104 | negative regulation of tissue remodeling | 0.00350245 | 4.571428571 | 0.02343784 |
| GO:1900017 | positive regulation of cytokine production involved in inflammatory response | 0.00350245 | 4.571428571 | 0.02343784 |
| GO:0019318 | hexose metabolic process | 0.00351981 | 1.835369092 | 0.02351847 |
| GO:0051592 | response to calcium ion | 0.00353824 | 2.131463628 | 0.02360602 |
| GO:0045637 | regulation of myeloid cell differentiation | 0.00354896 | 1.889020071 | 0.02364194 |
| GO:0051149 | positive regulation of muscle cell differentiation | 0.00357812 | 2.252587992 | 0.02380041 |
| GO:0050769 | positive regulation of neurogenesis | 0.00363553 | 1.509015257 | 0.02413572 |
| GO:0031345 | negative regulation of cell projection organization | 0.00363943 | 1.916406737 | 0.02413572 |
| GO:0036473 | cell death in response to oxidative stress | 0.00366869 | 2.509803922 | 0.02429344 |
| GO:1904406 | negative regulation of nitric oxide metabolic process | 0.00375576 | 5.485714286 | 0.02472219 |
| GO:0010572 | positive regulation of platelet activation | 0.00375576 | 5.485714286 | 0.02472219 |
| GO:0045623 | negative regulation of T-helper cell differentiation | 0.00375576 | 5.485714286 | 0.02472219 |
| GO:0045019 | negative regulation of nitric oxide biosynthetic process | 0.00375576 | 5.485714286 | 0.02472219 |
| GO:0050714 | positive regulation of protein secretion | 0.00380888 | 2.066182405 | 0.02503461 |
| GO:0090303 | positive regulation of wound healing | 0.00389025 | 2.612244898 | 0.02553157 |
| GO:0031669 | cellular response to nutrient levels | 0.00394044 | 1.8735363 | 0.02582273 |
| GO:0060443 | mammary gland morphogenesis | 0.00398594 | 3.15270936 | 0.02608232 |
| GO:1904951 | positive regulation of establishment of protein localization | 0.00400163 | 1.68030888 | 0.02614633 |
| GO:0098586 | cellular response to virus | 0.00404776 | 2.742857143 | 0.02636998 |
| GO:1903201 | regulation of oxidative stress-induced cell death | 0.00404776 | 2.742857143 | 0.02636998 |

| | | | | |
|---|---|---|---|---|
| GO:0071559 | response to transforming growth factor beta | 0.00408676 | 1.81512605 | 0.02658496 |
| GO:0090023 | positive regulation of neutrophil chemotaxis | 0.00421531 | 4.413793103 | 0.02738099 |
| GO:0043301 | negative regulation of leukocyte degranulation | 0.00425756 | 7.032967033 | 0.02761489 |
| GO:0006886 | intracellular protein transport | 0.00427103 | 1.370983447 | 0.02766177 |
| GO:0002823 | negative regulation of adaptive immune response based on somatic recombination of immune receptors built from immunoglobulin superfamily domains | 0.00448056 | 3.099273608 | 0.02885014 |
| GO:0042987 | amyloid precursor protein catabolic process | 0.00448056 | 3.099273608 | 0.02885014 |
| GO:0043536 | positive regulation of blood vessel endothelial cell migration | 0.00448056 | 3.099273608 | 0.02885014 |
| GO:0071622 | regulation of granulocyte chemotaxis | 0.00448056 | 3.099273608 | 0.02885014 |
| GO:0048468 | cell development | 0.00451486 | 1.206843583 | 0.0290288 |
| GO:0035313 | wound healing, spreading of epidermal cells | 0.00471073 | 5.224489796 | 0.03005383 |
| GO:0046851 | negative regulation of bone remodeling | 0.00471073 | 5.224489796 | 0.03005383 |
| GO:1902170 | cellular response to reactive nitrogen species | 0.00471073 | 5.224489796 | 0.03005383 |
| GO:0002281 | macrophage activation involved in immune response | 0.00471073 | 5.224489796 | 0.03005383 |
| GO:0014910 | regulation of smooth muscle cell migration | 0.00471474 | 2.438095238 | 0.03005383 |
| GO:0045920 | negative regulation of exocytosis | 0.00471499 | 3.750915751 | 0.03005383 |
| GO:0010975 | regulation of neuron projection development | 0.00478447 | 1.481085892 | 0.03045289 |
| GO:0043315 | positive regulation of neutrophil degranulation | 0.0048211 | 10.44897959 | 0.03059809 |
| GO:1903980 | positive regulation of microglial cell activation | 0.0048211 | 10.44897959 | 0.03059809 |
| GO:2001238 | positive regulation of extrinsic apoptotic signaling pathway | 0.00502168 | 3.047619048 | 0.0317862 |
| GO:0001919 | regulation of receptor recycling | 0.00502983 | 4.266666667 | 0.0317862 |
| GO:0032691 | negative regulation of interleukin-1 beta production | 0.00502983 | 4.266666667 | 0.0317862 |
| GO:0046888 | negative regulation of hormone secretion | 0.00511235 | 2.41509434 | 0.03226171 |
| GO:0071407 | cellular response to organic cyclic compound | 0.0051259 | 1.457556936 | 0.03230119 |
| GO:0034284 | response to monosaccharide | 0.00514708 | 1.807713199 | 0.03235655 |
| GO:0014074 | response to purine-containing compound | 0.0051566 | 2.009419152 | 0.03235655 |
| GO:0009267 | cellular response to starvation | 0.0051566 | 2.009419152 | 0.03235655 |
| GO:0045861 | negative regulation of proteolysis | 0.0052609 | 1.649109511 | 0.03296432 |
| GO:0010770 | positive regulation of cell morphogenesis involved in differentiation | 0.00538939 | 1.959183673 | 0.03372172 |
| GO:0019722 | calcium-mediated signaling | 0.00557378 | 1.915646259 | 0.03477663 |
| GO:0042307 | positive regulation of protein import into nucleus | 0.00558153 | 2.793650794 | 0.03477663 |
| GO:0015914 | phospholipid transport | 0.00558153 | 2.793650794 | 0.03477663 |
| GO:0034332 | adherens junction organization | 0.00567028 | 2.216450216 | 0.03527998 |
| GO:0033004 | negative regulation of mast cell activation | 0.0057055 | 6.530612245 | 0.03544931 |
| GO:0044257 | cellular protein catabolic process | 0.00579686 | 1.401539778 | 0.03596648 |
| GO:0099515 | actin filament-based transport | 0.00582618 | 4.987012987 | 0.03604746 |
| GO:0006896 | Golgi to vacuole transport | 0.00582618 | 4.987012987 | 0.03604746 |
| GO:0032647 | regulation of interferon-alpha production | 0.00595406 | 4.129032258 | 0.03658321 |
| GO:1905523 | positive regulation of macrophage migration | 0.00595406 | 4.129032258 | 0.03658321 |
| GO:0034123 | positive regulation of toll-like receptor signaling pathway | 0.00595406 | 4.129032258 | 0.03658321 |
| GO:0090330 | regulation of platelet aggregation | 0.00595406 | 4.129032258 | 0.03658321 |
| GO:0071624 | positive regulation of granulocyte chemotaxis | 0.00595406 | 4.129032258 | 0.03658321 |
| GO:0030279 | negative regulation of ossification | 0.00598808 | 2.37037037 | 0.03674128 |
| GO:0001889 | liver development | 0.00602492 | 1.939393939 | 0.03691615 |
| GO:0048259 | regulation of receptor-mediated endocytosis | 0.00607777 | 2.199785177 | 0.03718856 |
| GO:0001960 | negative regulation of cytokine-mediated signaling pathway | 0.00615902 | 2.755381605 | 0.03760806 |
| GO:0051650 | establishment of vesicle localization | 0.00616331 | 1.662337662 | 0.03760806 |
| GO:0050792 | regulation of viral process | 0.00617638 | 1.716134998 | 0.03763599 |
| GO:0032411 | positive regulation of transporter activity | 0.00619559 | 2.129158513 | 0.03765946 |
| GO:0010638 | positive regulation of organelle organization | 0.00619899 | 1.431711146 | 0.03765946 |
| GO:0045444 | fat cell differentiation | 0.00620573 | 1.806888763 | 0.03765946 |
| GO:0051931 | regulation of sensory perception | 0.00625475 | 2.949308756 | 0.03780157 |
| GO:0071715 | icosanoid transport | 0.00625475 | 2.949308756 | 0.03780157 |
| GO:1901571 | fatty acid derivative transport | 0.00625475 | 2.949308756 | 0.03780157 |
| GO:0032743 | positive regulation of interleukin-2 production | 0.0062701 | 3.567944251 | 0.03784272 |
| GO:0090276 | regulation of peptide hormone secretion | 0.00675894 | 1.792717087 | 0.04073753 |
| GO:0001558 | regulation of cell growth | 0.00680469 | 1.549636804 | 0.04095755 |
| GO:0097352 | autophagosome maturation | 0.00683798 | 3.164835165 | 0.04104639 |
| GO:0070527 | platelet aggregation | 0.00683798 | 3.164835165 | 0.04104639 |
| GO:0042176 | regulation of protein catabolic process | 0.00687204 | 1.593912141 | 0.04119505 |
| GO:0019932 | second-messenger-mediated signaling | 0.00691319 | 1.682734443 | 0.04138573 |
| GO:0044060 | regulation of endocrine process | 0.0069525 | 2.902494331 | 0.04151796 |
| GO:0043331 | response to dsRNA | 0.0069634 | 2.167195767 | 0.04151796 |
| GO:1903729 | regulation of plasma membrane organization | 0.0069634 | 2.167195767 | 0.04151796 |
| GO:0032958 | inositol phosphate biosynthetic process | 0.00699608 | 4 | 0.04165675 |
| GO:0034109 | homotypic cell-cell adhesion | 0.00703254 | 2.551495017 | 0.04181765 |
| GO:0002761 | regulation of myeloid leukocyte differentiation | 0.00704734 | 2.1003861 | 0.04184951 |
| GO:0007044 | cell-substrate junction assembly | 0.00707904 | 2.425655977 | 0.0419253 |
| GO:0000045 | autophagosome assembly | 0.00707904 | 2.425655977 | 0.0419253 |
| GO:0061008 | hepaticobiliary system development | 0.00709223 | 1.910447761 | 0.04194736 |
| GO:0050650 | chondroitin sulfate proteoglycan biosynthetic process | 0.00711529 | 4.770186335 | 0.04197169 |
| GO:0051770 | positive regulation of nitric-oxide synthase biosynthetic process | 0.00711529 | 4.770186335 | 0.04197169 |
| GO:0034110 | regulation of homotypic cell-cell adhesion | 0.00717728 | 3.482993197 | 0.0422249 |
| GO:0032689 | negative regulation of interferon-gamma production | 0.00717728 | 3.482993197 | 0.0422249 |
| GO:0042100 | B cell proliferation | 0.00727559 | 2.229965157 | 0.04274649 |
| GO:0051153 | regulation of striated muscle cell differentiation | 0.0074434 | 2.151260504 | 0.04364348 |
| GO:0032308 | positive regulation of prostaglandin secretion | 0.00744796 | 6.095238095 | 0.04364348 |

| | | | | |
|---|---|---|---|---|
| GO:0043525 | positive regulation of neuron apoptotic process | 0.00766421 | 2.522167488 | 0.04479217 |
| GO:0019229 | regulation of vasoconstriction | 0.00766421 | 2.522167488 | 0.04479217 |
| GO:0061756 | leukocyte adhesion to vascular endothelial cell | 0.00767752 | 3.105121294 | 0.0448108 |
| GO:0009895 | negative regulation of catabolic process | 0.00779751 | 1.726273726 | 0.04545128 |
| GO:0048015 | phosphatidylinositol-mediated signaling | 0.00794773 | 2.019281332 | 0.04626601 |
| GO:0014909 | smooth muscle cell migration | 0.00809641 | 2.285714286 | 0.04700797 |
| GO:0045921 | positive regulation of exocytosis | 0.00809641 | 2.285714286 | 0.04700797 |
| GO:0002828 | regulation of type 2 immune response | 0.00816386 | 3.878787879 | 0.04717308 |
| GO:0001881 | receptor recycling | 0.00816386 | 3.878787879 | 0.04717308 |
| GO:0048873 | homeostasis of number of cells within a tissue | 0.00817809 | 3.401993355 | 0.04717308 |
| GO:0098751 | bone cell development | 0.00817809 | 3.401993355 | 0.04717308 |
| GO:1903427 | negative regulation of reactive oxygen species biosynthetic process | 0.00817809 | 3.401993355 | 0.04717308 |
| GO:0051806 | entry into cell of other organism involved in symbiotic interaction | 0.00829174 | 2.377142857 | 0.04770444 |
| GO:0044409 | entry into host | 0.00829174 | 2.377142857 | 0.04770444 |
| GO:0050864 | regulation of B cell activation | 0.00851482 | 1.657315494 | 0.04892431 |
| GO:0007584 | response to nutrient | 0.00855898 | 1.91473448 | 0.04909283 |
| GO:1903522 | regulation of blood circulation | 0.00856665 | 1.735140772 | 0.04909283 |
| GO:0043550 | regulation of lipid kinase activity | 0.00859222 | 3.047619048 | 0.04909283 |
| GO:1903362 | regulation of cellular protein catabolic process | 0.00860927 | 1.783972125 | 0.04909283 |
| GO:0030208 | dermatan sulfate biosynthetic process | 0.00862172 | 18.28571429 | 0.04909283 |
| GO:0032764 | negative regulation of mast cell cytokine production | 0.00862172 | 18.28571429 | 0.04909283 |
| GO:0002605 | negative regulation of dendritic cell antigen processing and presentation | 0.00862172 | 18.28571429 | 0.04909283 |
| GO:0031346 | positive regulation of cell projection organization | 0.00872405 | 1.517640254 | 0.04961172 |
| GO:0051224 | negative regulation of protein transport | 0.00877225 | 1.951845907 | 0.04982187 |
| GO:0006605 | protein targeting | 0.0088537 | 1.505882353 | 0.05022008 |
| GO:0060761 | negative regulation of response to cytokine stimulus | 0.00894637 | 2.612244898 | 0.05054153 |
| GO:0008625 | extrinsic apoptotic signaling pathway via death domain receptors | 0.00894637 | 2.612244898 | 0.05054153 |
| GO:2000378 | negative regulation of reactive oxygen species metabolic process | 0.00894637 | 2.612244898 | 0.05054153 |
| GO:0051828 | entry into other organism involved in symbiotic interaction | 0.00895601 | 2.353606789 | 0.05054153 |
| GO:0032414 | positive regulation of ion transmembrane transporter activity | 0.00904365 | 2.104830421 | 0.05097121 |
| GO:0043491 | protein kinase B signaling | 0.00920966 | 1.86407767 | 0.05177512 |
| GO:0030308 | negative regulation of cell growth | 0.00920966 | 1.86407767 | 0.05177512 |
| GO:2000785 | regulation of autophagosome assembly | 0.00927792 | 3.324675325 | 0.05209278 |
| GO:0007188 | adenylate cyclase-modulating G-protein coupled receptor signaling pathway | 0.0093253 | 1.828571429 | 0.05229249 |
| GO:0002011 | morphogenesis of an epithelial sheet | 0.00940597 | 2.770562771 | 0.05264724 |
| GO:0048017 | inositol lipid-mediated signaling | 0.00946488 | 1.982788296 | 0.05264724 |
| GO:0001779 | natural killer cell differentiation | 0.00946528 | 3.764705882 | 0.05264724 |
| GO:1990776 | response to angiotensin | 0.00946528 | 3.764705882 | 0.05264724 |
| GO:0032607 | interferon-alpha production | 0.00946528 | 3.764705882 | 0.05264724 |
| GO:1904706 | negative regulation of vascular smooth muscle cell proliferation | 0.00946528 | 3.764705882 | 0.05264724 |
| GO:0035587 | purinergic receptor signaling pathway | 0.0095074 | 5.714285714 | 0.05264724 |
| GO:0048102 | autophagic cell death | 0.0095074 | 5.714285714 | 0.05264724 |
| GO:0032957 | inositol trisphosphate metabolic process | 0.0095074 | 5.714285714 | 0.05264724 |
| GO:0002866 | positive regulation of acute inflammatory response to antigenic stimulus | 0.0095074 | 5.714285714 | 0.05264724 |
| GO:0097576 | vacuole fusion | 0.00958602 | 2.992207792 | 0.0530163 |
| GO:1904591 | positive regulation of protein import | 0.00977422 | 2.578754579 | 0.05398977 |
| GO:0010769 | regulation of cell morphogenesis involved in differentiation | 0.00984249 | 1.607535322 | 0.05429916 |
| GO:0043523 | regulation of neuron apoptotic process | 0.01002483 | 1.690802348 | 0.0552363 |
| GO:0032409 | regulation of transporter activity | 0.01014065 | 1.651980418 | 0.0557968 |
| GO:0031330 | negative regulation of cellular catabolic process | 0.01018774 | 1.846153846 | 0.0557968 |
| GO:0006986 | response to unfolded protein | 0.01021501 | 2.142857143 | 0.0557968 |
| GO:0006639 | acylglycerol metabolic process | 0.01025383 | 2.074974671 | 0.0557968 |
| GO:0035855 | megakaryocyte development | 0.0102651 | 4.388571429 | 0.0557968 |
| GO:0033008 | positive regulation of mast cell activation involved in immune response | 0.0102651 | 4.388571429 | 0.0557968 |
| GO:0033622 | integrin activation | 0.0102651 | 4.388571429 | 0.0557968 |
| GO:0002864 | regulation of acute inflammatory response to antigenic stimulus | 0.0102651 | 4.388571429 | 0.0557968 |
| GO:0043306 | positive regulation of mast cell degranulation | 0.0102651 | 4.388571429 | 0.0557968 |
| GO:0043011 | myeloid dendritic cell differentiation | 0.0102651 | 4.388571429 | 0.0557968 |
| GO:0010884 | positive regulation of lipid storage | 0.0102651 | 4.388571429 | 0.0557968 |
| GO:0007219 | Notch signaling pathway | 0.01033777 | 1.91949487 | 0.05612294 |
| GO:2000273 | positive regulation of receptor activity | 0.01035383 | 2.729211087 | 0.05614131 |
| GO:0070741 | response to interleukin-6 | 0.01048215 | 3.250793651 | 0.05656019 |
| GO:1902930 | regulation of alcohol biosynthetic process | 0.01048215 | 3.250793651 | 0.05656019 |
| GO:0048246 | macrophage chemotaxis | 0.01048215 | 3.250793651 | 0.05656019 |
| GO:0006984 | ER-nucleus signaling pathway | 0.01048215 | 3.250793651 | 0.05656019 |
| GO:0010827 | regulation of glucose transport | 0.01064274 | 2.411302983 | 0.05704869 |
| GO:0038089 | positive regulation of cell migration by vascular endothelial growth factor signaling pathway | 0.0106559 | 8.126984127 | 0.05704869 |
| GO:0014004 | microglia differentiation | 0.0106559 | 8.126984127 | 0.05704869 |
| GO:0048260 | positive regulation of receptor-mediated endocytosis | 0.0106592 | 2.546112116 | 0.05704869 |
| GO:0060147 | regulation of posttranscriptional gene silencing | 0.01066282 | 2.93877551 | 0.05704869 |
| GO:0042149 | cellular response to glucose starvation | 0.01066282 | 2.93877551 | 0.05704869 |
| GO:0001961 | positive regulation of cytokine-mediated signaling pathway | 0.01066282 | 2.93877551 | 0.05704869 |
| GO:0050796 | regulation of insulin secretion | 0.01070711 | 1.837320574 | 0.05714759 |
| GO:0002793 | positive regulation of peptide secretion | 0.01070711 | 1.837320574 | 0.05714759 |
| GO:0048639 | positive regulation of developmental growth | 0.01078195 | 1.803971813 | 0.05747778 |
| GO:0003158 | endothelium development | 0.01090493 | 2.060362173 | 0.05780222 |

| | | | | |
|---|---|---|---|---|
| GO:1904950 | negative regulation of establishment of protein localization | 0.01090644 | 1.908948195 | 0.05780222 |
| GO:0002693 | positive regulation of cellular extravasation | 0.01090805 | 3.657142857 | 0.05780222 |
| GO:0060142 | regulation of syncytium formation by plasma membrane fusion | 0.01090805 | 3.657142857 | 0.05780222 |
| GO:0031294 | lymphocyte costimulation | 0.01090805 | 3.657142857 | 0.05780222 |
| GO:0022008 | neurogenesis | 0.01102852 | 1.221534203 | 0.0583708 |
| GO:1901888 | regulation of cell junction assembly | 0.01119947 | 2.285714286 | 0.05920483 |
| GO:0000904 | cell morphogenesis involved in differentiation | 0.01127731 | 1.361702128 | 0.05954525 |
| GO:0042594 | response to starvation | 0.01130617 | 1.795918367 | 0.05955571 |
| GO:0008361 | regulation of cell size | 0.01130617 | 1.795918367 | 0.05955571 |
| GO:0048525 | negative regulation of viral process | 0.01143911 | 1.992673993 | 0.06018438 |
| GO:0045824 | negative regulation of innate immune response | 0.01150993 | 2.385093168 | 0.06048515 |
| GO:0006638 | neutral lipid metabolic process | 0.01158805 | 2.045954046 | 0.06082352 |
| GO:0072659 | protein localization to plasma membrane | 0.01161089 | 1.691916624 | 0.06086234 |
| GO:0051897 | positive regulation of protein kinase B signaling | 0.01162292 | 2.10989011 | 0.06086234 |
| GO:1904645 | response to beta-amyloid | 0.01179605 | 3.180124224 | 0.06162097 |
| GO:0002691 | regulation of cellular extravasation | 0.01179605 | 3.180124224 | 0.06162097 |
| GO:0051955 | regulation of amino acid transport | 0.01182654 | 2.887218045 | 0.06162097 |
| GO:0060966 | regulation of gene silencing by RNA | 0.01182654 | 2.887218045 | 0.06162097 |
| GO:0006140 | regulation of nucleotide metabolic process | 0.01183735 | 1.936134454 | 0.06162097 |
| GO:0001845 | phagolysosome assembly | 0.01190403 | 5.378151261 | 0.06167817 |
| GO:0045064 | T-helper 2 cell differentiation | 0.01190403 | 5.378151261 | 0.06167817 |
| GO:0001921 | positive regulation of receptor recycling | 0.01190403 | 5.378151261 | 0.06167817 |
| GO:0030206 | chondroitin sulfate biosynthetic process | 0.01190403 | 5.378151261 | 0.06167817 |
| GO:0015696 | ammonium transport | 0.01203621 | 2.263945578 | 0.06229019 |
| GO:0010829 | negative regulation of glucose transport | 0.01214975 | 4.21978022 | 0.06273121 |
| GO:0035743 | CD4-positive, alpha-beta T cell cytokine production | 0.01214975 | 4.21978022 | 0.06273121 |
| GO:0048041 | focal adhesion assembly | 0.01246258 | 2.65010352 | 0.06419679 |
| GO:1903391 | regulation of adherens junction organization | 0.01246258 | 2.65010352 | 0.06419679 |
| GO:0046466 | membrane lipid catabolic process | 0.01249967 | 3.555555556 | 0.06431308 |
| GO:1901184 | regulation of ERBB signaling pathway | 0.01261008 | 2.48324515 | 0.06480584 |
| GO:0017038 | protein import | 0.01262539 | 1.679959616 | 0.06480934 |
| GO:0044265 | cellular macromolecule catabolic process | 0.01265083 | 1.308550186 | 0.06486481 |
| GO:0002792 | negative regulation of peptide secretion | 0.0129201 | 2.242587601 | 0.06609245 |
| GO:0001938 | positive regulation of endothelial cell proliferation | 0.0129201 | 2.242587601 | 0.06609245 |
| GO:0015749 | monosaccharide transport | 0.01314892 | 2.151260504 | 0.06718534 |
| GO:1903827 | regulation of cellular protein localization | 0.0131677 | 1.4271777 | 0.06720382 |
| GO:0002832 | negative regulation of response to biotic stimulus | 0.01322484 | 3.112462006 | 0.06741776 |
| GO:0010977 | negative regulation of neuron projection development | 0.0134341 | 1.867895545 | 0.06840583 |
| GO:0072594 | establishment of protein localization to organelle | 0.01359406 | 1.500771367 | 0.06914084 |
| GO:0006029 | proteoglycan metabolic process | 0.01368073 | 2.452961672 | 0.06942227 |
| GO:0002704 | negative regulation of leukocyte mediated immunity | 0.01368073 | 2.452961672 | 0.06942227 |
| GO:0031623 | receptor internalization | 0.01403394 | 2.133333333 | 0.07105185 |
| GO:0010522 | regulation of calcium ion transport into cytosol | 0.01403394 | 2.133333333 | 0.07105185 |
| GO:0098927 | vesicle-mediated transport between endosomal compartments | 0.01424742 | 3.459459459 | 0.07192837 |
| GO:0060143 | positive regulation of syncytium formation by plasma membrane fusion | 0.01425578 | 4.063492063 | 0.07192837 |
| GO:0032753 | positive regulation of interleukin-4 production | 0.01425578 | 4.063492063 | 0.07192837 |
| GO:0051489 | regulation of filopodium assembly | 0.01443013 | 2.789346247 | 0.07254152 |
| GO:0031532 | actin cytoskeleton reorganization | 0.0144367 | 2.309774436 | 0.07254152 |
| GO:0007399 | nervous system development | 0.01454982 | 1.175952291 | 0.07254152 |
| GO:0070391 | response to lipoteichoic acid | 0.01461132 | 7.314285714 | 0.07254152 |
| GO:0033029 | regulation of neutrophil apoptotic process | 0.01461132 | 7.314285714 | 0.07254152 |
| GO:0002317 | plasma cell differentiation | 0.01461132 | 7.314285714 | 0.07254152 |
| GO:1901724 | positive regulation of cell proliferation involved in kidney development | 0.01461132 | 7.314285714 | 0.07254152 |
| GO:0002315 | marginal zone B cell differentiation | 0.01461132 | 7.314285714 | 0.07254152 |
| GO:0070666 | regulation of mast cell proliferation | 0.01461132 | 7.314285714 | 0.07254152 |
| GO:0071223 | cellular response to lipoteichoic acid | 0.01461132 | 7.314285714 | 0.07254152 |
| GO:1903223 | positive regulation of oxidative stress-induced neuron death | 0.01461132 | 7.314285714 | 0.07254152 |
| GO:1903265 | positive regulation of tumor necrosis factor-mediated signaling pathway | 0.01461132 | 7.314285714 | 0.07254152 |
| GO:0010919 | regulation of inositol phosphate biosynthetic process | 0.01465568 | 5.079365079 | 0.07254152 |
| GO:0070102 | interleukin-6-mediated signaling pathway | 0.01465568 | 5.079365079 | 0.07254152 |
| GO:0045342 | MHC class II biosynthetic process | 0.01465568 | 5.079365079 | 0.07254152 |
| GO:0002730 | regulation of dendritic cell cytokine production | 0.01465568 | 5.079365079 | 0.07254152 |
| GO:2000319 | regulation of T-helper 17 cell differentiation | 0.01465568 | 5.079365079 | 0.07254152 |
| GO:0019058 | viral life cycle | 0.01483009 | 1.585466557 | 0.07330632 |
| GO:0051896 | regulation of protein kinase B signaling | 0.01485984 | 1.848024316 | 0.07330632 |
| GO:0060402 | calcium ion transport into cytosol | 0.01485984 | 1.848024316 | 0.07330632 |
| GO:0099565 | chemical synaptic transmission, postsynaptic | 0.01552716 | 2.285714286 | 0.07642815 |
| GO:0050709 | negative regulation of protein secretion | 0.01552716 | 2.285714286 | 0.07642815 |
| GO:1990778 | protein localization to cell periphery | 0.0155861 | 1.56590371 | 0.07663312 |
| GO:0045666 | positive regulation of neuron differentiation | 0.01605926 | 1.473435655 | 0.07887195 |
| GO:0045954 | positive regulation of natural killer cell mediated cytotoxicity | 0.01615826 | 3.368421053 | 0.07909512 |
| GO:0006884 | cell volume homeostasis | 0.01615826 | 3.368421053 | 0.07909512 |
| GO:0050996 | positive regulation of lipid catabolic process | 0.01615826 | 3.368421053 | 0.07909512 |
| GO:0032728 | positive regulation of interferon-beta production | 0.01644733 | 2.985422741 | 0.08033263 |
| GO:0002686 | negative regulation of leukocyte migration | 0.01644733 | 2.985422741 | 0.08033263 |
| GO:0051354 | negative regulation of oxidoreductase activity | 0.01659346 | 3.918367347 | 0.08063977 |

| | | | | |
|---|---|---|---|---|
| GO:1904466 | positive regulation of matrix metallopeptidase secretion | 0.01661944 | 13.71428571 | 0.08063977 |
| GO:0035624 | receptor transactivation | 0.01661944 | 13.71428571 | 0.08063977 |
| GO:0070164 | negative regulation of adiponectin secretion | 0.01661944 | 13.71428571 | 0.08063977 |
| GO:1904151 | positive regulation of microglial cell mediated cytotoxicity | 0.01661944 | 13.71428571 | 0.08063977 |
| GO:2000520 | regulation of immunological synapse formation | 0.01661944 | 13.71428571 | 0.08063977 |
| GO:1903524 | positive regulation of blood circulation | 0.01667829 | 2.262150221 | 0.08074842 |
| GO:0043535 | regulation of blood vessel endothelial cell migration | 0.01667829 | 2.262150221 | 0.08074842 |
| GO:0060079 | excitatory postsynaptic potential | 0.01730192 | 2.366386555 | 0.0836763 |
| GO:1905114 | cell surface receptor signaling pathway involved in cell-cell signaling | 0.01753746 | 1.432214416 | 0.08470438 |
| GO:0072657 | protein localization to membrane | 0.01755274 | 1.406593407 | 0.08470438 |
| GO:0033500 | carbohydrate homeostasis | 0.01766884 | 1.595015576 | 0.08517188 |
| GO:0032700 | negative regulation of interleukin-17 production | 0.01777778 | 4.812030075 | 0.08541817 |
| GO:1900225 | regulation of NLRP3 inflammasome complex assembly | 0.01777778 | 4.812030075 | 0.08541817 |
| GO:0050855 | regulation of B cell receptor signaling pathway | 0.01777778 | 4.812030075 | 0.08541817 |
| GO:0034620 | cellular response to unfolded protein | 0.01789198 | 2.239067055 | 0.08587377 |
| GO:0051961 | negative regulation of nervous system development | 0.01792082 | 1.535808024 | 0.08591908 |
| GO:0007269 | neurotransmitter secretion | 0.01797221 | 1.849117175 | 0.08607232 |
| GO:0051482 | positive regulation of cytosolic calcium ion concentration involved in phospholipase C-activating G-protein coupled signaling pathway | 0.01823885 | 3.282051282 | 0.08703032 |
| GO:0002724 | regulation of T cell cytokine production | 0.01823885 | 3.282051282 | 0.08703032 |
| GO:1903727 | positive regulation of phospholipid metabolic process | 0.01825083 | 2.925714286 | 0.08703032 |
| GO:0045646 | regulation of erythrocyte differentiation | 0.01825083 | 2.925714286 | 0.08703032 |
| GO:0050768 | negative regulation of neurogenesis | 0.01838193 | 1.558441558 | 0.08756124 |
| GO:0099643 | signal release from synapse | 0.01888526 | 1.838786911 | 0.0898622 |
| GO:0031644 | regulation of neurological system process | 0.01916083 | 2.048 | 0.09074042 |
| GO:0046718 | viral entry into host cell | 0.01917016 | 2.216450216 | 0.09074042 |
| GO:0071402 | cellular response to lipoprotein particle stimulus | 0.01917224 | 3.783251232 | 0.09074042 |
| GO:0044090 | positive regulation of vacuole organization | 0.01917224 | 3.783251232 | 0.09074042 |
| GO:0071677 | positive regulation of mononuclear cell migration | 0.01917224 | 3.783251232 | 0.09074042 |
| GO:0002246 | wound healing involved in inflammatory response | 0.0192861 | 6.649350649 | 0.0909784 |
| GO:0007220 | Notch receptor processing | 0.0192861 | 6.649350649 | 0.0909784 |
| GO:0090160 | Golgi to lysosome transport | 0.0192861 | 6.649350649 | 0.0909784 |
| GO:1900182 | positive regulation of protein localization to nucleus | 0.01930467 | 2.12244898 | 0.0909784 |
| GO:0007187 | G-protein coupled receptor signaling pathway, coupled to cyclic nucleotide second messenger | 0.01991748 | 1.675583381 | 0.09376668 |
| GO:0071359 | cellular response to dsRNA | 0.02008029 | 2.311986864 | 0.09443188 |
| GO:0032846 | positive regulation of homeostatic process | 0.02010141 | 1.611622276 | 0.09443188 |
| GO:0003254 | regulation of membrane depolarization | 0.02018874 | 2.868347339 | 0.09474164 |
| GO:0030522 | intracellular receptor signaling pathway | 0.02044979 | 1.721973094 | 0.09586515 |
| GO:0002717 | positive regulation of natural killer cell mediated immunity | 0.02049551 | 3.2 | 0.09587655 |
| GO:0034105 | positive regulation of tissue remodeling | 0.02049551 | 3.2 | 0.09587655 |
| GO:1903076 | regulation of protein localization to plasma membrane | 0.020564 | 2.103666245 | 0.09609549 |
| GO:0034767 | positive regulation of ion transmembrane transport | 0.02067002 | 1.749829118 | 0.09648912 |
| GO:0050994 | regulation of lipid catabolic process | 0.02070575 | 2.438095238 | 0.09655419 |
| GO:0010812 | negative regulation of cell-substrate adhesion | 0.02084658 | 2.612244898 | 0.09710868 |
| GO:0097320 | plasma membrane tubulation | 0.02128328 | 4.571428571 | 0.09883117 |
| GO:0045986 | negative regulation of smooth muscle contraction | 0.02128328 | 4.571428571 | 0.09883117 |
| GO:0030050 | vesicle transport along actin filament | 0.02128328 | 4.571428571 | 0.09883117 |
| GO:0014812 | muscle cell migration | 0.02156468 | 2.015748031 | 0.09984101 |
| GO:0050688 | regulation of defense response to virus | 0.02158556 | 2.285714286 | 0.09984101 |
| GO:0002637 | regulation of immunoglobulin production | 0.02158556 | 2.285714286 | 0.09984101 |
| GO:0051147 | regulation of muscle cell differentiation | 0.0215909 | 1.662337662 | 0.09984101 |
| GO:0007179 | transforming growth factor beta receptor signaling pathway | 0.02177894 | 1.772594752 | 0.10060555 |
| GO:1904894 | positive regulation of STAT cascade | 0.02188334 | 2.085213033 | 0.1009825 |
| GO:0014015 | positive regulation of gliogenesis | 0.02192741 | 2.172560113 | 0.10108055 |
| GO:0010226 | response to lithium ion | 0.02200073 | 3.657142857 | 0.10131312 |
| GO:0031333 | negative regulation of protein complex assembly | 0.02266025 | 1.887557604 | 0.10424187 |
| GO:0043114 | regulation of vascular permeability | 0.02293415 | 3.12195122 | 0.10528318 |
| GO:0009311 | oligosaccharide metabolic process | 0.02293415 | 3.12195122 | 0.10528318 |
| GO:0060349 | bone morphogenesis | 0.02326424 | 2.067080745 | 0.10668799 |
| GO:0010721 | negative regulation of cell development | 0.02359114 | 1.48994709 | 0.10807523 |
| GO:0034250 | positive regulation of cellular amide metabolic process | 0.02384223 | 1.754689755 | 0.10900009 |
| GO:0006643 | membrane lipid metabolic process | 0.02384223 | 1.754689755 | 0.10900009 |
| GO:0042058 | regulation of epidermal growth factor receptor signaling pathway | 0.02416842 | 2.374768089 | 0.11026374 |
| GO:0061180 | mammary gland epithelium development | 0.02416842 | 2.374768089 | 0.11026374 |
| GO:0097400 | interleukin-17-mediated signaling pathway | 0.02468827 | 6.095238095 | 0.11230041 |
| GO:0090109 | regulation of cell-substrate junction assembly | 0.02471623 | 2.531868132 | 0.11230041 |
| GO:0051893 | regulation of focal adhesion assembly | 0.02471623 | 2.531868132 | 0.11230041 |
| GO:0042269 | regulation of natural killer cell mediated cytotoxicity | 0.02471623 | 2.531868132 | 0.11230041 |
| GO:0015748 | organophosphate ester transport | 0.02496474 | 2.13037448 | 0.1133133 |
| GO:0035767 | endothelial cell chemotaxis | 0.02508662 | 3.539170507 | 0.11363368 |
| GO:0045932 | negative regulation of muscle contraction | 0.02508662 | 3.539170507 | 0.11363368 |
| GO:0002524 | hypersensitivity | 0.0251827 | 4.353741497 | 0.11372006 |
| GO:0050665 | hydrogen peroxide biosynthetic process | 0.0251827 | 4.353741497 | 0.11372006 |
| GO:0010042 | response to manganese ion | 0.0251827 | 4.353741497 | 0.11372006 |
| GO:0051259 | protein oligomerization | 0.02555407 | 1.613445378 | 0.11519033 |
| GO:0002861 | regulation of inflammatory response to antigenic stimulus | 0.02556029 | 3.047619048 | 0.11519033 |
| GO:1904627 | response to phorbol 13-acetate 12-myristate | 0.02670144 | 10.97142857 | 0.11972406 |

| | | | | |
|---|---|---|---|---|
| GO:0038157 | granulocyte-macrophage colony-stimulating factor signaling pathway | 0.02670144 | 10.97142857 | 0.11972406 |
| GO:0035771 | interleukin-4-mediated signaling pathway | 0.02670144 | 10.97142857 | 0.11972406 |
| GO:1904628 | cellular response to phorbol 13-acetate 12-myristate | 0.02670144 | 10.97142857 | 0.11972406 |
| GO:0033031 | positive regulation of neutrophil apoptotic process | 0.02670144 | 10.97142857 | 0.11972406 |
| GO:0010883 | regulation of lipid storage | 0.02682799 | 2.493506494 | 0.12015821 |
| GO:0046596 | regulation of viral entry into host cell | 0.02685251 | 2.708994709 | 0.12015821 |
| GO:0032355 | response to estradiol | 0.02771811 | 1.840071878 | 0.12390641 |
| GO:0006109 | regulation of carbohydrate metabolic process | 0.02793198 | 1.693121693 | 0.12461094 |
| GO:0034764 | positive regulation of transmembrane transport | 0.02793198 | 1.693121693 | 0.12461094 |
| GO:0030149 | sphingolipid catabolic process | 0.02843672 | 3.428571429 | 0.12648061 |
| GO:0034694 | response to prostaglandin | 0.02843672 | 3.428571429 | 0.12648061 |
| GO:0060148 | positive regulation of posttranscriptional gene silencing | 0.02843672 | 3.428571429 | 0.12648061 |
| GO:0046890 | regulation of lipid biosynthetic process | 0.02910897 | 1.685319289 | 0.12934074 |
| GO:0060149 | negative regulation of posttranscriptional gene silencing | 0.02948415 | 4.155844156 | 0.13048432 |
| GO:0002863 | positive regulation of inflammatory response to antigenic stimulus | 0.02948415 | 4.155844156 | 0.13048432 |
| GO:0060967 | negative regulation of gene silencing by RNA | 0.02948415 | 4.155844156 | 0.13048432 |
| GO:1903209 | positive regulation of oxidative stress-induced cell death | 0.02948415 | 4.155844156 | 0.13048432 |
| GO:0006836 | neurotransmitter transport | 0.03026935 | 1.628687102 | 0.13382555 |
| GO:0045628 | regulation of T-helper 2 cell differentiation | 0.03081748 | 5.626373626 | 0.13476945 |
| GO:0001768 | establishment of T cell polarity | 0.03081748 | 5.626373626 | 0.13476945 |
| GO:0055119 | relaxation of cardiac muscle | 0.03081748 | 5.626373626 | 0.13476945 |
| GO:0032725 | positive regulation of granulocyte macrophage colony-stimulating factor production | 0.03081748 | 5.626373626 | 0.13476945 |
| GO:0001767 | establishment of lymphocyte polarity | 0.03081748 | 5.626373626 | 0.13476945 |
| GO:0036005 | response to macrophage colony-stimulating factor | 0.03081748 | 5.626373626 | 0.13476945 |
| GO:0032650 | regulation of interleukin-1 alpha production | 0.03081748 | 5.626373626 | 0.13476945 |
| GO:0023035 | CD40 signaling pathway | 0.03081748 | 5.626373626 | 0.13476945 |
| GO:0042590 | antigen processing and presentation of exogenous peptide antigen via MHC class I | 0.03081748 | 5.626373626 | 0.13476945 |
| GO:0060368 | regulation of Fc receptor mediated stimulatory signaling pathway | 0.03081748 | 5.626373626 | 0.13476945 |
| GO:0043320 | natural killer cell degranulation | 0.03081748 | 5.626373626 | 0.13476945 |
| GO:0030282 | bone mineralization | 0.03114469 | 1.980952381 | 0.13606604 |
| GO:0048147 | negative regulation of fibroblast proliferation | 0.03139499 | 2.909090909 | 0.13700059 |
| GO:0002715 | regulation of natural killer cell mediated immunity | 0.03142045 | 2.420168067 | 0.13700059 |
| GO:0043436 | oxoacid metabolic process | 0.03145771 | 1.25819135 | 0.1370282 |
| GO:0022898 | regulation of transmembrane transporter activity | 0.03178446 | 1.543599258 | 0.13831549 |
| GO:0032814 | regulation of natural killer cell activation | 0.03204307 | 2.612244898 | 0.13916746 |
| GO:0014003 | oligodendrocyte development | 0.03204307 | 2.612244898 | 0.13916746 |
| GO:0061045 | negative regulation of wound healing | 0.03229264 | 2.257495591 | 0.140114 |
| GO:0001936 | regulation of endothelial cell proliferation | 0.03327413 | 1.756255044 | 0.1442313 |
| GO:1903900 | regulation of viral life cycle | 0.03419139 | 1.654815772 | 0.14778215 |
| GO:0071731 | response to nitric oxide | 0.03419339 | 3.97515528 | 0.14778215 |
| GO:1901889 | negative regulation of cell junction assembly | 0.03419339 | 3.97515528 | 0.14778215 |
| GO:0046824 | positive regulation of nucleocytoplasmic transport | 0.03448802 | 2.117293233 | 0.14876525 |
| GO:1903035 | negative regulation of response to wounding | 0.03448802 | 2.117293233 | 0.14876525 |
| GO:0031326 | regulation of cellular biosynthetic process | 0.034589 | 1.10362063 | 0.14905572 |
| GO:0034113 | heterotypic cell-cell adhesion | 0.03487339 | 2.56641604 | 0.14998942 |
| GO:0071548 | response to dexamethasone | 0.03487339 | 2.56641604 | 0.14998942 |
| GO:0050678 | regulation of epithelial cell proliferation | 0.03508826 | 1.425389755 | 0.1507672 |
| GO:1905475 | regulation of protein localization to membrane | 0.03573287 | 1.705403405 | 0.15338821 |
| GO:1903205 | regulation of hydrogen peroxide-induced cell death | 0.03595198 | 3.226890756 | 0.15358465 |
| GO:0032673 | regulation of interleukin-4 production | 0.03595198 | 3.226890756 | 0.15358465 |
| GO:0070528 | protein kinase C signaling | 0.03595198 | 3.226890756 | 0.15358465 |
| GO:0045022 | early endosome to late endosome transport | 0.03595198 | 3.226890756 | 0.15358465 |
| GO:0070269 | pyroptosis | 0.03595198 | 3.226890756 | 0.15358465 |
| GO:2001242 | regulation of intrinsic apoptotic signaling pathway | 0.03634618 | 1.736632083 | 0.15496976 |
| GO:1902904 | negative regulation of supramolecular fiber organization | 0.03634618 | 1.736632083 | 0.15496976 |
| GO:0032412 | regulation of ion transmembrane transporter activity | 0.0364282 | 1.539201539 | 0.15517011 |
| GO:0043462 | regulation of ATPase activity | 0.03652315 | 2.351020408 | 0.15542514 |
| GO:1903901 | negative regulation of viral life cycle | 0.03670499 | 1.932636469 | 0.15604904 |
| GO:0001890 | placenta development | 0.03725787 | 1.696612666 | 0.15824773 |
| GO:0050679 | positive regulation of epithelial cell proliferation | 0.03731375 | 1.613445378 | 0.15831243 |
| GO:0002467 | germinal center formation | 0.03766621 | 5.224489796 | 0.15831243 |
| GO:0042362 | fat-soluble vitamin biosynthetic process | 0.03766621 | 5.224489796 | 0.15831243 |
| GO:0072683 | T cell extravasation | 0.03766621 | 5.224489796 | 0.15831243 |
| GO:0002829 | negative regulation of type 2 immune response | 0.03766621 | 5.224489796 | 0.15831243 |
| GO:1902902 | negative regulation of autophagosome assembly | 0.03766621 | 5.224489796 | 0.15831243 |
| GO:0032610 | interleukin-1 alpha production | 0.03766621 | 5.224489796 | 0.15831243 |
| GO:0070344 | regulation of fat cell proliferation | 0.03766621 | 5.224489796 | 0.15831243 |
| GO:0010936 | negative regulation of macrophage cytokine production | 0.03766621 | 5.224489796 | 0.15831243 |
| GO:0051712 | positive regulation of killing of cells of other organism | 0.03766621 | 5.224489796 | 0.15831243 |
| GO:0090594 | inflammatory response to wounding | 0.03766621 | 5.224489796 | 0.15831243 |
| GO:0048538 | thymus development | 0.03786449 | 2.522167488 | 0.15899498 |
| GO:0030879 | mammary gland development | 0.03795717 | 1.726984127 | 0.15910685 |
| GO:1901031 | regulation of response to reactive oxygen species | 0.0380348 | 2.782608696 | 0.15910685 |
| GO:0048536 | spleen development | 0.0380348 | 2.782608696 | 0.15910685 |
| GO:0035886 | vascular smooth muscle cell differentiation | 0.0380348 | 2.782608696 | 0.15910685 |
| GO:0071608 | macrophage inflammatory protein-1 alpha production | 0.03861653 | 9.142857143 | 0.15913602 |

| | | | | |
|---|---|---|---|---|
| GO:0014005 | microglia development | 0.03861653 | 9.142857143 | 0.15913602 |
| GO:0033025 | regulation of mast cell apoptotic process | 0.03861653 | 9.142857143 | 0.15913602 |
| GO:1903969 | regulation of response to macrophage colony-stimulating factor | 0.03861653 | 9.142857143 | 0.15913602 |
| GO:2000321 | positive regulation of T-helper 17 cell differentiation | 0.03861653 | 9.142857143 | 0.15913602 |
| GO:2000427 | positive regulation of apoptotic cell clearance | 0.03861653 | 9.142857143 | 0.15913602 |
| GO:1902262 | apoptotic process involved in blood vessel morphogenesis | 0.03861653 | 9.142857143 | 0.15913602 |
| GO:0002677 | negative regulation of chronic inflammatory response | 0.03861653 | 9.142857143 | 0.15913602 |
| GO:0043545 | molybdopterin cofactor metabolic process | 0.03861653 | 9.142857143 | 0.15913602 |
| GO:0070668 | positive regulation of mast cell proliferation | 0.03861653 | 9.142857143 | 0.15913602 |
| GO:0071640 | regulation of macrophage inflammatory protein 1 alpha production | 0.03861653 | 9.142857143 | 0.15913602 |
| GO:0006777 | Mo-molybdopterin cofactor biosynthetic process | 0.03861653 | 9.142857143 | 0.15913602 |
| GO:1990773 | matrix metallopeptidase secretion | 0.03861653 | 9.142857143 | 0.15913602 |
| GO:0051189 | prosthetic group metabolic process | 0.03861653 | 9.142857143 | 0.15913602 |
| GO:0019720 | Mo-molybdopterin cofactor metabolic process | 0.03861653 | 9.142857143 | 0.15913602 |
| GO:1903972 | regulation of cellular response to macrophage colony-stimulating factor stimulus | 0.03861653 | 9.142857143 | 0.15913602 |
| GO:0006665 | sphingolipid metabolic process | 0.03886254 | 1.804511278 | 0.15985242 |
| GO:0045598 | regulation of fat cell differentiation | 0.03886254 | 1.804511278 | 0.15985242 |
| GO:0008211 | glucocorticoid metabolic process | 0.0393139 | 3.80952381 | 0.16140924 |
| GO:0046629 | gamma-delta T cell activation | 0.0393139 | 3.80952381 | 0.16140924 |
| GO:1902115 | regulation of organelle assembly | 0.03946624 | 1.654421769 | 0.16188467 |
| GO:0051494 | negative regulation of cytoskeleton organization | 0.03961917 | 1.717442778 | 0.16236161 |
| GO:0016051 | carbohydrate biosynthetic process | 0.03990232 | 1.625396825 | 0.16337086 |
| GO:0010758 | regulation of macrophage chemotaxis | 0.04012597 | 3.134693878 | 0.16368145 |
| GO:0070670 | response to interleukin-4 | 0.04012597 | 3.134693878 | 0.16368145 |
| GO:0050654 | chondroitin sulfate proteoglycan metabolic process | 0.04012597 | 3.134693878 | 0.16368145 |
| GO:0090218 | positive regulation of lipid kinase activity | 0.04012597 | 3.134693878 | 0.16368145 |
| GO:0051283 | negative regulation of sequestering of calcium ion | 0.04018828 | 1.976833977 | 0.1637848 |
| GO:1900180 | regulation of protein localization to nucleus | 0.04044905 | 1.679300292 | 0.16469604 |
| GO:1904375 | regulation of protein localization to cell periphery | 0.0407806 | 1.901714286 | 0.16589352 |
| GO:0031175 | neuron projection development | 0.040866 | 1.239989667 | 0.16608842 |
| GO:0032964 | collagen biosynthetic process | 0.04101912 | 2.479418886 | 0.16625315 |
| GO:0070918 | production of small RNA involved in gene silencing by RNA | 0.04101912 | 2.479418886 | 0.16625315 |
| GO:0045010 | actin nucleation | 0.04101912 | 2.479418886 | 0.16625315 |
| GO:0061138 | morphogenesis of a branching epithelium | 0.04143267 | 1.618204804 | 0.16777579 |
| GO:0042692 | muscle cell differentiation | 0.04152425 | 1.367156208 | 0.16799307 |
| GO:0030574 | collagen catabolic process | 0.04166552 | 2.723404255 | 0.16825731 |
| GO:0046365 | monosaccharide catabolic process | 0.04166552 | 2.723404255 | 0.16825731 |
| GO:1901215 | negative regulation of neuron death | 0.04187944 | 1.551924091 | 0.16896715 |
| GO:0010821 | regulation of mitochondrion organization | 0.04208025 | 1.741496599 | 0.16931089 |
| GO:0090316 | positive regulation of intracellular protein transport | 0.04208025 | 1.741496599 | 0.16931089 |
| GO:0006096 | glycolytic process | 0.04213733 | 2.151260504 | 0.16931089 |
| GO:0032272 | negative regulation of protein polymerization | 0.04213733 | 2.151260504 | 0.16931089 |
| GO:0060135 | maternal process involved in female pregnancy | 0.04215574 | 2.285714286 | 0.16931089 |
| GO:0007041 | lysosomal transport | 0.04293264 | 1.886621315 | 0.172275 |
| GO:0001678 | cellular glucose homeostasis | 0.04383296 | 1.662337662 | 0.17572851 |
| GO:0001570 | vasculogenesis | 0.0438879 | 2.031746032 | 0.17578968 |
| GO:0051353 | positive regulation of oxidoreductase activity | 0.0443397 | 2.438095238 | 0.17705104 |
| GO:0048679 | regulation of axon regeneration | 0.04458183 | 3.047619048 | 0.17705104 |
| GO:0036474 | cell death in response to hydrogen peroxide | 0.04458183 | 3.047619048 | 0.17705104 |
| GO:1903206 | negative regulation of hydrogen peroxide-induced cell death | 0.044847 | 3.657142857 | 0.17705104 |
| GO:0048569 | post-embryonic animal organ development | 0.044847 | 3.657142857 | 0.17705104 |
| GO:0055098 | response to low-density lipoprotein particle | 0.044847 | 3.657142857 | 0.17705104 |
| GO:1990000 | amyloid fibril formation | 0.044847 | 3.657142857 | 0.17705104 |
| GO:0002544 | chronic inflammatory response | 0.044847 | 3.657142857 | 0.17705104 |
| GO:1900409 | positive regulation of cellular response to oxidative stress | 0.044847 | 3.657142857 | 0.17705104 |
| GO:0071404 | cellular response to low-density lipoprotein particle stimulus | 0.044847 | 3.657142857 | 0.17705104 |
| GO:1901623 | regulation of lymphocyte chemotaxis | 0.044847 | 3.657142857 | 0.17705104 |
| GO:0034114 | regulation of heterotypic cell-cell adhesion | 0.044847 | 3.657142857 | 0.17705104 |
| GO:0034695 | response to prostaglandin E | 0.044847 | 3.657142857 | 0.17705104 |
| GO:1901032 | negative regulation of response to reactive oxygen species | 0.044847 | 3.657142857 | 0.17705104 |
| GO:0051279 | regulation of release of sequestered calcium ion into cytosol | 0.04488224 | 2.126245847 | 0.17705104 |
| GO:0051966 | regulation of synaptic transmission, glutamatergic | 0.04488224 | 2.126245847 | 0.17705104 |
| GO:0030808 | regulation of nucleotide biosynthetic process | 0.04488224 | 2.126245847 | 0.17705104 |
| GO:0000079 | regulation of cyclin-dependent protein serine/threonine kinase activity | 0.04517609 | 2.254403131 | 0.17791119 |
| GO:0014049 | positive regulation of glutamate secretion | 0.04522077 | 4.876190476 | 0.17791119 |
| GO:0070943 | neutrophil mediated killing of symbiont cell | 0.04522077 | 4.876190476 | 0.17791119 |
| GO:0010762 | regulation of fibroblast migration | 0.04550722 | 2.666666667 | 0.17840443 |
| GO:0032008 | positive regulation of TOR signaling | 0.04550722 | 2.666666667 | 0.17840443 |
| GO:0043277 | apoptotic cell clearance | 0.04550722 | 2.666666667 | 0.17840443 |
| GO:0032768 | regulation of monooxygenase activity | 0.04550722 | 2.666666667 | 0.17840443 |
| GO:0010976 | positive regulation of neuron projection development | 0.04675476 | 1.43605086 | 0.18313316 |
| GO:1903829 | positive regulation of cellular protein localization | 0.0468633 | 1.458689459 | 0.18339612 |
| GO:0045834 | positive regulation of lipid metabolic process | 0.04741476 | 1.645714286 | 0.18539047 |
| GO:0046620 | regulation of organ growth | 0.04747075 | 1.857142857 | 0.18544572 |
| GO:0050922 | negative regulation of chemotaxis | 0.04782837 | 2.398126464 | 0.18667813 |
| GO:0048662 | negative regulation of smooth muscle cell proliferation | 0.04833497 | 2.223938224 | 0.18848935 |

| | | | | |
|---|---|---|---|---|
| GO:0007200 | phospholipase C-activating G-protein coupled receptor signaling pathway | 0.04918355 | 1.991513437 | 0.19162984 |
| GO:0034122 | negative regulation of toll-like receptor signaling pathway | 0.04932158 | 2.965250965 | 0.19199876 |
| GO:0097106 | postsynaptic density organization | 0.05079196 | 3.516483516 | 0.19634238 |
| GO:0032727 | positive regulation of interferon-alpha production | 0.05079196 | 3.516483516 | 0.19634238 |
| GO:1903798 | regulation of production of miRNAs involved in gene silencing by miRNA | 0.05079196 | 3.516483516 | 0.19634238 |
| GO:0002726 | positive regulation of T cell cytokine production | 0.05079196 | 3.516483516 | 0.19634238 |
| GO:1903077 | negative regulation of protein localization to plasma membrane | 0.05079196 | 3.516483516 | 0.19634238 |
| GO:1903975 | regulation of glial cell migration | 0.05079196 | 3.516483516 | 0.19634238 |
| GO:0014048 | regulation of glutamate secretion | 0.05079196 | 3.516483516 | 0.19634238 |
| GO:0042908 | xenobiotic transport | 0.05079196 | 3.516483516 | 0.19634238 |
| GO:0009887 | animal organ morphogenesis | 0.05146591 | 1.215969216 | 0.19868228 |
| GO:0046847 | filopodium assembly | 0.05148697 | 2.359447005 | 0.19868228 |
| GO:0060558 | regulation of calcidiol 1-monooxygenase activity | 0.0521347 | 7.836734694 | 0.20011366 |
| GO:0072679 | thymocyte migration | 0.0521347 | 7.836734694 | 0.20011366 |
| GO:1904139 | regulation of microglial cell migration | 0.0521347 | 7.836734694 | 0.20011366 |
| GO:0002408 | myeloid dendritic cell chemotaxis | 0.0521347 | 7.836734694 | 0.20011366 |
| GO:0002578 | negative regulation of antigen processing and presentation | 0.0521347 | 7.836734694 | 0.20011366 |
| GO:0032747 | positive regulation of interleukin-23 production | 0.0521347 | 7.836734694 | 0.20011366 |
| GO:0006913 | nucleocytoplasmic transport | 0.05221928 | 1.421169504 | 0.20011366 |
| GO:0051169 | nuclear transport | 0.05221928 | 1.421169504 | 0.20011366 |
| GO:0046165 | alcohol biosynthetic process | 0.05232767 | 1.828571429 | 0.20035573 |
| GO:0019377 | glycolipid catabolic process | 0.05346231 | 4.571428571 | 0.20399488 |
| GO:0032736 | positive regulation of interleukin-13 production | 0.05346231 | 4.571428571 | 0.20399488 |
| GO:1901722 | regulation of cell proliferation involved in kidney development | 0.05346231 | 4.571428571 | 0.20399488 |
| GO:0032645 | regulation of granulocyte macrophage colony-stimulating factor production | 0.05346231 | 4.571428571 | 0.20399488 |
| GO:0002064 | epithelial cell development | 0.05365949 | 1.49271137 | 0.20457104 |
| GO:1901343 | negative regulation of vasculature development | 0.05418771 | 1.765517241 | 0.2064072 |
| GO:0001953 | negative regulation of cell-matrix adhesion | 0.05434632 | 2.887218045 | 0.20647874 |
| GO:0014912 | negative regulation of smooth muscle cell migration | 0.05434632 | 2.887218045 | 0.20647874 |
| GO:0033198 | response to ATP | 0.05434632 | 2.887218045 | 0.20647874 |
| GO:0035239 | tube morphogenesis | 0.05467298 | 1.378523915 | 0.2075418 |
| GO:0006672 | ceramide metabolic process | 0.05489041 | 1.952843273 | 0.20818881 |
| GO:0046622 | positive regulation of organ growth | 0.05531701 | 2.321995465 | 0.20944818 |
| GO:0090183 | regulation of kidney development | 0.05531701 | 2.321995465 | 0.20944818 |
| GO:0060562 | epithelial tube morphogenesis | 0.05579284 | 1.40311174 | 0.2110694 |
| GO:0007156 | homophilic cell adhesion via plasma membrane adhesion molecules | 0.05642847 | 1.671836735 | 0.2132919 |
| GO:0007205 | protein kinase C-activating G-protein coupled receptor signaling pathway | 0.05714607 | 3.386243386 | 0.21508676 |
| GO:1903513 | endoplasmic reticulum to cytosol transport | 0.05714607 | 3.386243386 | 0.21508676 |
| GO:0002523 | leukocyte migration involved in inflammatory response | 0.05714607 | 3.386243386 | 0.21508676 |
| GO:0030970 | retrograde protein transport, ER to cytosol | 0.05714607 | 3.386243386 | 0.21508676 |
| GO:1904996 | positive regulation of leukocyte adhesion to vascular endothelial cell | 0.05714607 | 3.386243386 | 0.21508676 |
| GO:0010565 | regulation of cellular ketone metabolic process | 0.0579004 | 1.703637977 | 0.21774088 |
| GO:0048193 | Golgi vesicle transport | 0.05828828 | 1.496695475 | 0.21893794 |
| GO:0007157 | heterophilic cell-cell adhesion via plasma membrane cell adhesion molecules | 0.05831755 | 2.509803922 | 0.21893794 |
| GO:0030968 | endoplasmic reticulum unfolded protein response | 0.05865786 | 2.13729128 | 0.21984289 |
| GO:1904029 | regulation of cyclin-dependent protein kinase activity | 0.05865786 | 2.13729128 | 0.21984289 |
| GO:0006753 | nucleoside phosphate metabolic process | 0.05889622 | 1.298059965 | 0.22054968 |
| GO:1904892 | regulation of STAT cascade | 0.05919867 | 1.627524308 | 0.22138703 |
| GO:0046686 | response to cadmium ion | 0.05931973 | 2.285714286 | 0.22138703 |
| GO:0048255 | mRNA stabilization | 0.05931973 | 2.285714286 | 0.22138703 |
| GO:0060393 | regulation of pathway-restricted SMAD protein phosphorylation | 0.05931973 | 2.285714286 | 0.22138703 |
| GO:0090162 | establishment of epithelial cell polarity | 0.05965625 | 2.813186813 | 0.22245553 |
| GO:0030500 | regulation of bone mineralization | 0.06038662 | 2.009419152 | 0.22498969 |
| GO:0051603 | proteolysis involved in cellular protein catabolic process | 0.06063112 | 1.264313152 | 0.22571082 |
| GO:0097479 | synaptic vesicle localization | 0.06116204 | 1.543098252 | 0.22749607 |
| GO:0045346 | regulation of MHC class II biosynthetic process | 0.06236766 | 4.302521008 | 0.231268 |
| GO:0061621 | canonical glycolysis | 0.06236766 | 4.302521008 | 0.231268 |
| GO:0014889 | muscle atrophy | 0.06236766 | 4.302521008 | 0.231268 |
| GO:0014068 | positive regulation of phosphatidylinositol 3-kinase signaling | 0.06238493 | 2.10989011 | 0.231268 |
| GO:0048705 | skeletal system morphogenesis | 0.06244894 | 1.501066098 | 0.2313117 |
| GO:0048016 | inositol phosphate-mediated signaling | 0.06301999 | 2.461538462 | 0.23284284 |
| GO:1902743 | regulation of lamellipodium organization | 0.06301999 | 2.461538462 | 0.23284284 |
| GO:0043618 | regulation of transcription from RNA polymerase II promoter in response to stress | 0.06301999 | 2.461538462 | 0.23284284 |
| GO:0043470 | regulation of carbohydrate catabolic process | 0.06349601 | 2.250549451 | 0.23396567 |
| GO:0030856 | regulation of epithelial cell differentiation | 0.06349784 | 1.643659711 | 0.23396567 |
| GO:0033157 | regulation of intracellular protein transport | 0.06359115 | 1.480885312 | 0.23396567 |
| GO:0051216 | cartilage development | 0.06361246 | 1.557975657 | 0.23396567 |
| GO:0045981 | positive regulation of nucleotide metabolic process | 0.06385167 | 1.98757764 | 0.23396567 |
| GO:0045778 | positive regulation of ossification | 0.06388894 | 1.828571429 | 0.23396567 |
| GO:0035967 | cellular response to topologically incorrect protein | 0.06388894 | 1.828571429 | 0.23396567 |
| GO:0070920 | regulation of production of small RNA involved in gene silencing by RNA | 0.06390484 | 3.265306122 | 0.23396567 |
| GO:1902884 | positive regulation of response to oxidative stress | 0.06390484 | 3.265306122 | 0.23396567 |
| GO:0060252 | positive regulation of glial cell proliferation | 0.06390484 | 3.265306122 | 0.23396567 |
| GO:1904376 | negative regulation of protein localization to cell periphery | 0.06390484 | 3.265306122 | 0.23396567 |
| GO:0014066 | regulation of phosphatidylinositol 3-kinase signaling | 0.06424 | 1.897574124 | 0.2349985 |
| GO:0006775 | fat-soluble vitamin metabolic process | 0.06525064 | 2.742857143 | 0.23849861 |

| | | | | |
|---|---|---|---|---|
| GO:0043271 | negative regulation of ion transport | 0.0661777 | 1.602356406 | 0.24168774 |
| GO:0051952 | regulation of amine transport | 0.06698522 | 1.813459268 | 0.24206345 |
| GO:0045833 | negative regulation of lipid metabolic process | 0.06698522 | 1.813459268 | 0.24206345 |
| GO:0032741 | positive regulation of interleukin-18 production | 0.06704557 | 6.857142857 | 0.24206345 |
| GO:0050861 | positive regulation of B cell receptor signaling pathway | 0.06704557 | 6.857142857 | 0.24206345 |
| GO:1904124 | microglial cell migration | 0.06704557 | 6.857142857 | 0.24206345 |
| GO:1904464 | regulation of matrix metallopeptidase secretion | 0.06704557 | 6.857142857 | 0.24206345 |
| GO:0048549 | positive regulation of pinocytosis | 0.06704557 | 6.857142857 | 0.24206345 |
| GO:0072126 | positive regulation of glomerular mesangial cell proliferation | 0.06704557 | 6.857142857 | 0.24206345 |
| GO:0060696 | regulation of phospholipid catabolic process | 0.06704557 | 6.857142857 | 0.24206345 |
| GO:1903976 | negative regulation of glial cell migration | 0.06704557 | 6.857142857 | 0.24206345 |
| GO:1903979 | negative regulation of microglial cell activation | 0.06704557 | 6.857142857 | 0.24206345 |
| GO:0045627 | positive regulation of T-helper 1 cell differentiation | 0.06704557 | 6.857142857 | 0.24206345 |
| GO:2000643 | positive regulation of early endosome to late endosome transport | 0.06704557 | 6.857142857 | 0.24206345 |
| GO:0071224 | cellular response to peptidoglycan | 0.06704557 | 6.857142857 | 0.24206345 |
| GO:2001259 | positive regulation of cation channel activity | 0.06744017 | 1.966205837 | 0.24309191 |
| GO:1905897 | regulation of response to endoplasmic reticulum stress | 0.06744017 | 1.966205837 | 0.24309191 |
| GO:0019217 | regulation of fatty acid metabolic process | 0.06757033 | 1.879839786 | 0.24336307 |
| GO:0060964 | regulation of gene silencing by miRNA | 0.0679393 | 2.41509434 | 0.24429472 |
| GO:0060251 | regulation of glial cell proliferation | 0.0679393 | 2.41509434 | 0.24429472 |
| GO:0046928 | regulation of neurotransmitter secretion | 0.06887972 | 1.74789916 | 0.24747538 |
| GO:0043255 | regulation of carbohydrate biosynthetic process | 0.07017683 | 1.798594848 | 0.25193139 |
| GO:0045780 | positive regulation of bone resorption | 0.07106206 | 3.15270936 | 0.25370106 |
| GO:0046852 | positive regulation of bone remodeling | 0.07106206 | 3.15270936 | 0.25370106 |
| GO:0006270 | DNA replication initiation | 0.07106206 | 3.15270936 | 0.25370106 |
| GO:0030204 | chondroitin sulfate metabolic process | 0.07106206 | 3.15270936 | 0.25370106 |
| GO:1901185 | negative regulation of ERBB signaling pathway | 0.07106206 | 3.15270936 | 0.25370106 |
| GO:0007176 | regulation of epidermal growth factor-activated receptor activity | 0.07106206 | 3.15270936 | 0.25370106 |
| GO:0043524 | negative regulation of neuron apoptotic process | 0.0711265 | 1.586005831 | 0.25370106 |
| GO:0070570 | regulation of neuron projection regeneration | 0.07112793 | 2.675958188 | 0.25370106 |
| GO:0009117 | nucleotide metabolic process | 0.07124816 | 1.283708491 | 0.25392546 |
| GO:0061051 | positive regulation of cell growth involved in cardiac muscle cell development | 0.07191006 | 4.063492063 | 0.25493597 |
| GO:0001821 | histamine secretion | 0.07191006 | 4.063492063 | 0.25493597 |
| GO:0071732 | cellular response to nitric oxide | 0.07191006 | 4.063492063 | 0.25493597 |
| GO:0071801 | regulation of podosome assembly | 0.07191006 | 4.063492063 | 0.25493597 |
| GO:0050862 | positive regulation of T cell receptor signaling pathway | 0.07191006 | 4.063492063 | 0.25493597 |
| GO:0061430 | bone trabecula morphogenesis | 0.07191006 | 4.063492063 | 0.25493597 |
| GO:0050829 | defense response to Gram-negative bacterium | 0.07193453 | 1.735140772 | 0.25493597 |
| GO:0014911 | positive regulation of smooth muscle cell migration | 0.07237134 | 2.18336887 | 0.25627899 |
| GO:0051588 | regulation of neurotransmitter transport | 0.07303699 | 1.684210526 | 0.25794653 |
| GO:0072666 | establishment of protein localization to vacuole | 0.07307515 | 2.37037037 | 0.25794653 |
| GO:1903573 | negative regulation of response to endoplasmic reticulum stress | 0.07307515 | 2.37037037 | 0.25794653 |
| GO:0045907 | positive regulation of vasoconstriction | 0.07307515 | 2.37037037 | 0.25794653 |
| GO:0070167 | regulation of biomineral tissue development | 0.0745554 | 1.845347313 | 0.26296213 |
| GO:0097480 | establishment of synaptic vesicle localization | 0.07760078 | 1.517155334 | 0.27326826 |
| GO:0048489 | synaptic vesicle transport | 0.07760078 | 1.517155334 | 0.27326826 |
| GO:0051209 | release of sequestered calcium ion into cytosol | 0.07821097 | 1.828571429 | 0.27485565 |
| GO:0006939 | smooth muscle contraction | 0.07821097 | 1.828571429 | 0.27485565 |
| GO:0046467 | membrane lipid biosynthetic process | 0.07830254 | 1.710174717 | 0.27485565 |
| GO:0010712 | regulation of collagen metabolic process | 0.07842673 | 2.327272727 | 0.27485565 |
| GO:2000727 | positive regulation of cardiac muscle cell differentiation | 0.07860996 | 3.047619048 | 0.27485565 |
| GO:0045332 | phospholipid translocation | 0.07860996 | 3.047619048 | 0.27485565 |
| GO:0072574 | hepatocyte proliferation | 0.07860996 | 3.047619048 | 0.27485565 |
| GO:1903649 | regulation of cytoplasmic transport | 0.07860996 | 3.047619048 | 0.27485565 |
| GO:0090200 | positive regulation of release of cytochrome c from mitochondria | 0.07860996 | 3.047619048 | 0.27485565 |
| GO:0051145 | smooth muscle cell differentiation | 0.07874352 | 2.006968641 | 0.27488874 |
| GO:1903670 | regulation of sprouting angiogenesis | 0.07874352 | 2.006968641 | 0.27488874 |
| GO:0031640 | killing of cells of other organism | 0.08032805 | 1.755428571 | 0.28019941 |
| GO:0060491 | regulation of cell projection assembly | 0.08102196 | 1.53089701 | 0.28239756 |
| GO:0045071 | negative regulation of viral genome replication | 0.08194396 | 2.120082816 | 0.28467125 |
| GO:0032456 | endocytic recycling | 0.08194396 | 2.120082816 | 0.28467125 |
| GO:0008347 | glial cell migration | 0.08194396 | 2.120082816 | 0.28467125 |
| GO:0030837 | negative regulation of actin filament polymerization | 0.08194396 | 2.120082816 | 0.28467125 |
| GO:2000370 | positive regulation of clathrin-dependent endocytosis | 0.08205986 | 3.84962406 | 0.28467125 |
| GO:0033623 | regulation of integrin activation | 0.08205986 | 3.84962406 | 0.28467125 |
| GO:0031214 | biomineral tissue development | 0.08224445 | 1.581467181 | 0.28508836 |
| GO:0006664 | glycolipid metabolic process | 0.08303809 | 1.885125184 | 0.28534862 |
| GO:0006685 | sphingomyelin catabolic process | 0.0831569 | 6.095238095 | 0.28534862 |
| GO:0060753 | regulation of mast cell chemotaxis | 0.0831569 | 6.095238095 | 0.28534862 |
| GO:0031622 | positive regulation of fever generation | 0.0831569 | 6.095238095 | 0.28534862 |
| GO:0051126 | negative regulation of actin nucleation | 0.0831569 | 6.095238095 | 0.28534862 |
| GO:0031946 | regulation of glucocorticoid biosynthetic process | 0.0831569 | 6.095238095 | 0.28534862 |
| GO:0060556 | regulation of vitamin D biosynthetic process | 0.0831569 | 6.095238095 | 0.28534862 |
| GO:1990456 | mitochondrion-ER tethering | 0.0831569 | 6.095238095 | 0.28534862 |
| GO:0097527 | necroptotic signaling pathway | 0.0831569 | 6.095238095 | 0.28534862 |
| GO:0010727 | negative regulation of hydrogen peroxide metabolic process | 0.0831569 | 6.095238095 | 0.28534862 |

| | | | | |
|---|---|---|---|---|
| GO:0070424 | regulation of nucleotide-binding oligomerization domain containing signaling pathway | 0.0831569 | 6.095238095 | 0.28534862 |
| GO:0044857 | plasma membrane raft organization | 0.0831569 | 6.095238095 | 0.28534862 |
| GO:1900125 | regulation of hyaluronan biosynthetic process | 0.0831569 | 6.095238095 | 0.28534862 |
| GO:0071526 | semaphorin-plexin signaling pathway | 0.08372081 | 2.551495017 | 0.28706128 |
| GO:0048145 | regulation of fibroblast proliferation | 0.083905 | 1.741496599 | 0.28724818 |
| GO:0015837 | amine transport | 0.083905 | 1.741496599 | 0.28724818 |
| GO:0032291 | axon ensheathment in central nervous system | 0.08653934 | 2.949308756 | 0.29467277 |
| GO:1903393 | positive regulation of adherens junction organization | 0.08653934 | 2.949308756 | 0.29467277 |
| GO:2000637 | positive regulation of gene silencing by miRNA | 0.08653934 | 2.949308756 | 0.29467277 |
| GO:0043552 | positive regulation of phosphatidylinositol 3-kinase activity | 0.08653934 | 2.949308756 | 0.29467277 |
| GO:0090025 | regulation of monocyte chemotaxis | 0.08653934 | 2.949308756 | 0.29467277 |
| GO:0051957 | positive regulation of amino acid transport | 0.08653934 | 2.949308756 | 0.29467277 |
| GO:0032801 | receptor catabolic process | 0.08653934 | 2.949308756 | 0.29467277 |
| GO:0019752 | carboxylic acid metabolic process | 0.08728933 | 1.203247401 | 0.29699826 |
| GO:0010469 | regulation of receptor activity | 0.08819241 | 1.564553094 | 0.29984067 |
| GO:0051928 | positive regulation of calcium ion transport | 0.08882607 | 1.630573248 | 0.30176341 |
| GO:0048666 | neuron development | 0.08959903 | 1.169840061 | 0.30404269 |
| GO:0030810 | positive regulation of nucleotide biosynthetic process | 0.08977152 | 2.245614035 | 0.30404269 |
| GO:0010559 | regulation of glycoprotein biosynthetic process | 0.08977152 | 2.245614035 | 0.30404269 |
| GO:1902883 | negative regulation of response to oxidative stress | 0.08977152 | 2.245614035 | 0.30404269 |
| GO:0002021 | response to dietary excess | 0.09042924 | 2.493506494 | 0.30580271 |
| GO:2000008 | regulation of protein localization to cell surface | 0.09042924 | 2.493506494 | 0.30580271 |
| GO:0002040 | sprouting angiogenesis | 0.09134972 | 1.714285714 | 0.30867984 |
| GO:0045600 | positive regulation of fat cell differentiation | 0.09220994 | 2.060362173 | 0.31092146 |
| GO:0031102 | neuron projection regeneration | 0.09220994 | 2.060362173 | 0.31092146 |
| GO:0055021 | regulation of cardiac muscle tissue growth | 0.09253969 | 1.936134454 | 0.31092146 |
| GO:0010468 | regulation of gene expression | 0.09258656 | 1.07088419 | 0.31092146 |
| GO:0071379 | cellular response to prostaglandin stimulus | 0.09278514 | 3.657142857 | 0.31092146 |
| GO:1903392 | negative regulation of adherens junction organization | 0.09278514 | 3.657142857 | 0.31092146 |
| GO:0060965 | negative regulation of gene silencing by miRNA | 0.09278514 | 3.657142857 | 0.31092146 |
| GO:0034383 | low-density lipoprotein particle clearance | 0.09278514 | 3.657142857 | 0.31092146 |
| GO:0090036 | regulation of protein kinase C signaling | 0.09278514 | 3.657142857 | 0.31092146 |
| GO:0006670 | sphingosine metabolic process | 0.09278514 | 3.657142857 | 0.31092146 |
| GO:0090030 | regulation of steroid hormone biosynthetic process | 0.09278514 | 3.657142857 | 0.31092146 |
| GO:0050848 | regulation of calcium-mediated signaling | 0.09392454 | 1.764411028 | 0.31450167 |
| GO:0071322 | cellular response to carbohydrate stimulus | 0.09440665 | 1.547996977 | 0.31587723 |
| GO:0034204 | lipid translocation | 0.09483971 | 2.857142857 | 0.31613238 |
| GO:0072576 | liver morphogenesis | 0.09483971 | 2.857142857 | 0.31613238 |
| GO:1903846 | positive regulation of cellular response to transforming growth factor beta stimulus | 0.09483971 | 2.857142857 | 0.31613238 |
| GO:0030511 | positive regulation of transforming growth factor beta receptor signaling pathway | 0.09483971 | 2.857142857 | 0.31613238 |
| GO:0001945 | lymph vessel development | 0.09483971 | 2.857142857 | 0.31613238 |
| GO:1990089 | response to nerve growth factor | 0.09576079 | 2.206896552 | 0.3182448 |
| GO:0001659 | temperature homeostasis | 0.09576079 | 2.206896552 | 0.3182448 |
| GO:0097035 | regulation of membrane lipid distribution | 0.09576079 | 2.206896552 | 0.3182448 |
| GO:0060337 | type I interferon signaling pathway | 0.09576079 | 2.206896552 | 0.3182448 |
| GO:0030148 | sphingolipid biosynthetic process | 0.09604627 | 1.828571429 | 0.31895425 |
| GO:0071804 | cellular potassium ion transport | 0.09640386 | 1.516376307 | 0.3196625 |
| GO:0071805 | potassium ion transmembrane transport | 0.09640386 | 1.516376307 | 0.3196625 |
| GO:0010765 | positive regulation of sodium ion transport | 0.09740635 | 2.438095238 | 0.3225743 |
| GO:0050810 | regulation of steroid biosynthetic process | 0.09742763 | 1.913621262 | 0.3225743 |
| GO:0048754 | branching morphogenesis of an epithelial tube | 0.09761388 | 1.539849624 | 0.32294957 |
| GO:0014897 | striated muscle hypertrophy | 0.09812555 | 1.749068323 | 0.32440014 |

**Appendix Table S4.** Enriched KEGG pathways in up-regulated genes in Ptf1a RGC.

| Term | description | PValue | Fold Enrichment | FDR |
|---|---|---|---|---|
| mmu04512 | ECM-receptor interaction | 7.28E-10 | 5.474025974 | 2.02E-07 |
| mmu04510 | Focal adhesion | 2.45E-09 | 3.537821099 | 2.77E-07 |
| mmu00190 | Oxidative phosphorylation | 2.99E-09 | 4.247921391 | 2.77E-07 |
| mmu04974 | Protein digestion and absorption | 5.61E-09 | 4.67271353 | 3.90E-07 |
| mmu05415 | Diabetic cardiomyopathy | 3.07E-08 | 3.261437276 | 1.71E-06 |
| mmu05208 | Chemical carcinogenesis - reactive oxygen species | 9.57E-08 | 3.099834528 | 4.43E-06 |
| mmu05016 | Huntington disease | 2.91E-07 | 2.65846736 | 1.15E-05 |
| mmu04714 | Thermogenesis | 2.47E-06 | 2.780457638 | 8.59E-05 |
| mmu04260 | Cardiac muscle contraction | 2.04E-05 | 3.954961295 | 6.29E-04 |
| mmu05014 | Amyotrophic lateral sclerosis | 2.57E-05 | 2.175764615 | 7.15E-04 |
| mmu05012 | Parkinson disease | 2.96E-05 | 2.432900433 | 7.48E-04 |
| mmu04151 | PI3K-Akt signaling pathway | 3.58E-05 | 2.172474561 | 8.29E-04 |
| mmu05020 | Prion disease | 1.01E-04 | 2.31099604 | 2.17E-03 |
| mmu05165 | Human papillomavirus infection | 2.26E-04 | 2.027737062 | 4.49E-03 |
| mmu05022 | Pathways of neurodegeneration - multiple diseases | 3.21E-04 | 1.850686772 | 5.96E-03 |
| mmu05205 | Proteoglycans in cancer | 5.89E-04 | 2.349825784 | 1.02E-02 |
| mmu04360 | Axon guidance | 8.76E-04 | 2.407937761 | 1.43E-02 |
| mmu05200 | Pathways in cancer | 1.28E-03 | 1.689780885 | 1.97E-02 |
| mmu00260 | Glycine, serine and threonine metabolism | 1.50E-03 | 4.587755102 | 2.19E-02 |
| mmu04723 | Retrograde endocannabinoid signaling | 1.91E-03 | 2.479867623 | 2.66E-02 |
| mmu04964 | Proximal tubule bicarbonate reclamation | 2.16E-03 | 6.256029685 | 2.86E-02 |
| mmu05010 | Alzheimer disease | 2.48E-03 | 1.796770928 | 3.14E-02 |
| mmu04932 | Non-alcoholic fatty liver disease | 3.20E-03 | 2.352694924 | 3.86E-02 |
| mmu04971 | Gastric acid secretion | 5.01E-03 | 3.058503401 | 5.81E-02 |
| mmu05146 | Amoebiasis | 6.66E-03 | 2.572572954 | 7.41E-02 |
| mmu05222 | Small cell lung cancer | 6.96E-03 | 2.713188501 | 7.44E-02 |
| mmu04024 | cAMP signaling pathway | 7.35E-03 | 1.981076067 | 7.57E-02 |
| mmu04550 | Signaling pathways regulating pluripotency of stem cells | 7.83E-03 | 2.293877551 | 7.77E-02 |
| mmu05224 | Breast cancer | 1.16E-02 | 2.184645287 | 1.11E-01 |
| mmu04080 | Neuroactive ligand-receptor interaction | 1.77E-02 | 1.604525748 | 1.64E-01 |
| mmu04390 | Hippo signaling pathway | 1.92E-02 | 2.045495905 | 0.171755276 |
| mmu04350 | TGF-beta signaling pathway | 2.20E-02 | 2.414607948 | 0.191454248 |
| mmu04933 | AGE-RAGE signaling pathway in diabetic complications | 3.13E-02 | 2.271165892 | 0.263518918 |
| mmu04022 | cGMP-PKG signaling pathway | 3.83E-02 | 1.856317093 | 0.303181164 |
| mmu03320 | PPAR signaling pathway | 3.93E-02 | 2.319651456 | 0.303181164 |
| mmu04727 | GABAergic synapse | 3.93E-02 | 2.319651456 | 0.303181164 |
| mmu04330 | Notch signaling pathway | 4.52E-02 | 2.676190476 | 0.33963235 |
| mmu03020 | RNA polymerase | 4.84E-02 | 3.584183673 | 0.354402754 |
| mmu04926 | Relaxin signaling pathway | 5.42E-02 | 1.956019617 | 0.386118866 |
| mmu04919 | Thyroid hormone signaling pathway | 7.73E-02 | 1.911564626 | 0.537370644 |
| mmu04960 | Aldosterone-regulated sodium reabsorption | 8.14E-02 | 3.018259936 | 0.552015416 |
| mmu05033 | Nicotine addiction | 9.43E-02 | 2.867346939 | 0.616339537 |
| mmu04810 | Regulation of actin cytoskeleton | 9.87E-02 | 1.564007421 | 0.616339537 |
| mmu04514 | Cell adhesion molecules | 0.098803721 | 1.638483965 | 0.616339537 |

**Appendix Table S5.** Enriched GO terms in up-regulated genes in Ptf1a RGC.

| Term | description | PValue | Fold Enrichment | FDR |
|---|---|---|---|---|
| GO:0035556 | intracellular signal transduction | 2.53E-43 | 2.142857143 | 1.12E-39 |
| GO:0031347 | regulation of defense response | 3.09E-34 | 3.171066908 | 6.85E-31 |
| GO:0046649 | lymphocyte activation | 7.79E-29 | 2.647308588 | 1.15E-25 |
| GO:1902531 | regulation of intracellular signal transduction | 2.52E-28 | 2.136779752 | 2.79E-25 |
| GO:0009966 | regulation of signal transduction | 1.70E-27 | 1.801984768 | 1.51E-24 |
| GO:0002366 | leukocyte activation involved in immune response | 3.24E-26 | 4.123781263 | 2.39E-23 |
| GO:0016477 | cell migration | 1.01E-25 | 2.19521985 | 6.39E-23 |
| GO:0002274 | myeloid leukocyte activation | 3.63E-25 | 4.520064205 | 2.01E-22 |
| GO:0002694 | regulation of leukocyte activation | 1.21E-24 | 2.732858879 | 5.96E-22 |
| GO:0050900 | leukocyte migration | 1.37E-24 | 3.731778426 | 6.05E-22 |
| GO:0009967 | positive regulation of signal transduction | 6.30E-24 | 2.087950895 | 2.54E-21 |
| GO:0007159 | leukocyte cell-cell adhesion | 1.64E-23 | 3.396482567 | 6.07E-21 |
| GO:0007166 | cell surface receptor signaling pathway | 2.46E-23 | 1.772496097 | 8.39E-21 |
| GO:1902533 | positive regulation of intracellular signal transduction | 6.78E-23 | 2.355828221 | 2.14E-20 |
| GO:0031349 | positive regulation of defense response | 1.15E-22 | 3.361050328 | 3.40E-20 |
| GO:0007249 | I-kappaB kinase/NF-kappaB signaling | 1.57E-22 | 4.591218306 | 4.36E-20 |
| GO:0050727 | regulation of inflammatory response | 4.80E-22 | 3.447664507 | 1.25E-19 |
| GO:0001819 | positive regulation of cytokine production | 1.17E-21 | 3.064456722 | 2.87E-19 |
| GO:0045088 | regulation of innate immune response | 2.98E-21 | 3.523809524 | 6.62E-19 |
| GO:0019221 | cytokine-mediated signaling pathway | 3.04E-21 | 3.317136378 | 6.62E-19 |
| GO:0002521 | leukocyte differentiation | 3.14E-21 | 2.764119601 | 6.62E-19 |
| GO:1903555 | regulation of tumor necrosis factor superfamily cytokine production | 1.34E-20 | 4.758017493 | 2.71E-18 |
| GO:0050778 | positive regulation of immune response | 3.59E-20 | 2.435433562 | 6.91E-18 |
| GO:0048534 | hematopoietic or lymphoid organ development | 5.40E-20 | 2.259301587 | 9.96E-18 |
| GO:0032103 | positive regulation of response to external stimulus | 5.70E-20 | 3.462095238 | 1.01E-17 |
| GO:0016310 | phosphorylation | 1.01E-19 | 1.781175346 | 1.72E-17 |
| GO:0002520 | immune system development | 1.42E-19 | 2.202912505 | 2.33E-17 |
| GO:0006897 | endocytosis | 1.77E-19 | 2.432597842 | 2.80E-17 |
| GO:0019220 | regulation of phosphate metabolic process | 1.95E-19 | 1.903744947 | 2.98E-17 |
| GO:0001818 | negative regulation of cytokine production | 2.21E-19 | 3.766492644 | 3.26E-17 |
| GO:0032680 | regulation of tumor necrosis factor production | 2.47E-19 | 4.642487047 | 3.53E-17 |
| GO:0051174 | regulation of phosphorus metabolic process | 2.62E-19 | 1.898676403 | 3.62E-17 |
| GO:0030097 | hemopoiesis | 3.49E-19 | 2.262325581 | 4.68E-17 |
| GO:0032640 | tumor necrosis factor production | 5.87E-19 | 4.64399093 | 7.65E-17 |
| GO:0010647 | positive regulation of cell communication | 1.13E-18 | 1.832128961 | 1.43E-16 |
| GO:0032675 | regulation of interleukin-6 production | 1.50E-18 | 4.840336134 | 1.84E-16 |
| GO:0031325 | positive regulation of cellular metabolic process | 4.74E-18 | 1.562323391 | 5.68E-16 |
| GO:0030334 | regulation of cell migration | 1.05E-17 | 2.249433107 | 1.22E-15 |
| GO:0010942 | positive regulation of cell death | 1.08E-17 | 2.484734263 | 1.23E-15 |
| GO:1903037 | regulation of leukocyte cell-cell adhesion | 1.26E-17 | 3.112462006 | 1.39E-15 |
| GO:0050866 | negative regulation of cell activation | 1.30E-17 | 4.090225564 | 1.41E-15 |
| GO:0002695 | negative regulation of leukocyte activation | 1.92E-17 | 4.281533101 | 2.02E-15 |
| GO:2000145 | regulation of cell motility | 2.51E-17 | 2.189496191 | 2.59E-15 |
| GO:0002285 | lymphocyte activation involved in immune response | 4.49E-17 | 4.04551201 | 4.52E-15 |
| GO:0006915 | apoptotic process | 5.56E-17 | 1.795027976 | 5.48E-15 |
| GO:0097529 | myeloid leukocyte migration | 6.22E-17 | 3.881049563 | 5.99E-15 |
| GO:0002685 | regulation of leukocyte migration | 1.01E-16 | 3.901972504 | 9.54E-15 |
| GO:0050764 | regulation of phagocytosis | 1.11E-16 | 5.285714286 | 1.02E-14 |
| GO:0043122 | regulation of I-kappaB kinase/NF-kappaB signaling | 2.04E-16 | 4.044766294 | 1.85E-14 |
| GO:0071345 | cellular response to cytokine stimulus | 2.13E-16 | 2.167450611 | 1.88E-14 |
| GO:0000165 | MAPK cascade | 2.58E-16 | 2.355146676 | 2.20E-14 |
| GO:0023014 | signal transduction by protein phosphorylation | 2.58E-16 | 2.355146676 | 2.20E-14 |
| GO:0043068 | positive regulation of programmed cell death | 3.10E-16 | 2.505622657 | 2.59E-14 |
| GO:0050863 | regulation of T cell activation | 3.36E-16 | 3.134693878 | 2.76E-14 |
| GO:0002699 | positive regulation of immune effector process | 4.44E-16 | 3.334780721 | 3.57E-14 |
| GO:0002286 | T cell activation involved in immune response | 5.32E-16 | 5.087003222 | 4.21E-14 |
| GO:0043065 | positive regulation of apoptotic process | 5.84E-16 | 2.49716803 | 4.54E-14 |
| GO:0060627 | regulation of vesicle-mediated transport | 6.87E-16 | 2.550828482 | 5.25E-14 |
| GO:0010562 | positive regulation of phosphorus metabolic process | 2.26E-15 | 2.012303486 | 1.67E-13 |
| GO:0045937 | positive regulation of phosphate metabolic process | 2.26E-15 | 2.012303486 | 1.67E-13 |
| GO:0030335 | positive regulation of cell migration | 3.83E-15 | 2.56427379 | 2.78E-13 |
| GO:0070663 | regulation of leukocyte proliferation | 5.73E-15 | 3.46122449 | 4.09E-13 |
| GO:0051249 | regulation of lymphocyte activation | 6.32E-15 | 2.401937046 | 4.44E-13 |
| GO:0045089 | positive regulation of innate immune response | 6.77E-15 | 3.352380952 | 4.69E-13 |
| GO:0050867 | positive regulation of cell activation | 7.11E-15 | 2.519101644 | 4.85E-13 |
| GO:0043408 | regulation of MAPK cascade | 7.97E-15 | 2.314943369 | 5.35E-13 |
| GO:0046631 | alpha-beta T cell activation | 1.02E-14 | 4.053019146 | 6.71E-13 |
| GO:2000377 | regulation of reactive oxygen species metabolic process | 1.26E-14 | 3.736645963 | 8.24E-13 |
| GO:0022409 | positive regulation of cell-cell adhesion | 1.31E-14 | 3.216931217 | 8.30E-13 |
| GO:0030217 | T cell differentiation | 1.31E-14 | 3.216931217 | 8.30E-13 |
| GO:0051246 | regulation of protein metabolic process | 2.33E-14 | 1.555473098 | 1.45E-12 |
| GO:0051250 | negative regulation of lymphocyte activation | 2.76E-14 | 4.203612479 | 1.70E-12 |
| GO:2000147 | positive regulation of cell motility | 4.63E-14 | 2.453674121 | 2.81E-12 |
| GO:1903039 | positive regulation of leukocyte cell-cell adhesion | 5.33E-14 | 3.378881988 | 3.19E-12 |
| GO:0030036 | actin cytoskeleton organization | 5.54E-14 | 2.330035615 | 3.27E-12 |
| GO:0030100 | regulation of endocytosis | 6.43E-14 | 3.141104294 | 3.75E-12 |
| GO:0043067 | regulation of programmed cell death | 7.19E-14 | 1.776326531 | 4.14E-12 |
| GO:0031399 | regulation of protein modification process | 7.74E-14 | 1.712009106 | 4.40E-12 |
| GO:0051272 | positive regulation of cellular component movement | 9.01E-14 | 2.409745293 | 5.05E-12 |
| GO:0046651 | lymphocyte proliferation | 1.04E-13 | 2.996825397 | 5.76E-12 |
| GO:0043410 | positive regulation of MAPK cascade | 1.56E-13 | 2.518783542 | 8.44E-12 |
| GO:1901652 | response to peptide | 1.56E-13 | 2.518783542 | 8.44E-12 |
| GO:1903557 | positive regulation of tumor necrosis factor superfamily cytokine production | 2.14E-13 | 4.876190476 | 1.14E-11 |
| GO:0050729 | positive regulation of inflammatory response | 2.26E-13 | 4.136054422 | 1.19E-11 |
| GO:0032496 | response to lipopolysaccharide | 2.99E-13 | 2.650660264 | 1.54E-11 |
| GO:0002696 | positive regulation of leukocyte activation | 3.00E-13 | 2.452583071 | 1.54E-11 |
| GO:0032268 | regulation of cellular protein metabolic process | 3.07E-13 | 1.551783859 | 1.56E-11 |
| GO:0042981 | regulation of apoptotic process | 3.59E-13 | 1.75864923 | 1.81E-11 |
| GO:0032270 | positive regulation of cellular protein metabolic process | 4.30E-13 | 1.754582989 | 2.14E-11 |
| GO:0002718 | regulation of cytokine production involved in immune response | 4.68E-13 | 4.606324973 | 2.30E-11 |
| GO:0060326 | cell chemotaxis | 5.78E-13 | 3.057054401 | 2.82E-11 |
| GO:0010648 | negative regulation of cell communication | 6.03E-13 | 1.832336504 | 2.90E-11 |
| GO:0030595 | leukocyte chemotaxis | 6.28E-13 | 3.486682809 | 2.99E-11 |
| GO:0032760 | positive regulation of tumor necrosis factor production | 8.20E-13 | 4.803874092 | 3.86E-11 |
| GO:0050670 | regulation of lymphocyte proliferation | 9.94E-13 | 3.331118494 | 4.63E-11 |
| GO:0051247 | positive regulation of protein metabolic process | 1.10E-12 | 1.718053375 | 5.07E-11 |
| GO:0043549 | regulation of kinase activity | 1.22E-12 | 2.186841149 | 5.58E-11 |
| GO:0032944 | regulation of mononuclear cell proliferation | 2.01E-12 | 3.267789245 | 9.09E-11 |
| GO:0002292 | T cell differentiation involved in immune response | 2.45E-12 | 5.786618445 | 1.09E-10 |
| GO:0061082 | myeloid leukocyte cytokine production | 2.46E-12 | 6.372294372 | 1.09E-10 |
| GO:0050766 | positive regulation of phagocytosis | 2.56E-12 | 5.30875576 | 1.12E-10 |
| GO:0031098 | stress-activated protein kinase signaling cascade | 2.77E-12 | 3.099273608 | 1.20E-10 |

| | | | | |
|---|---|---|---|---|
| GO:0002221 | pattern recognition receptor signaling pathway | 2.97E-12 | 3.817896389 | 1.28E-10 |
| GO:0002703 | regulation of leukocyte mediated immunity | 3.11E-12 | 3.047619048 | 1.30E-10 |
| GO:0072358 | cardiovascular system development | 3.12E-12 | 2.13037448 | 1.30E-10 |
| GO:0001944 | vasculature development | 3.12E-12 | 2.13037448 | 1.30E-10 |
| GO:0002275 | myeloid cell activation involved in immune response | 3.28E-12 | 4.571428571 | 1.36E-10 |
| GO:0002758 | innate immune response-activating signal transduction | 4.24E-12 | 3.695040711 | 1.74E-10 |
| GO:0042119 | neutrophil activation | 4.34E-12 | 7.896103896 | 1.76E-10 |
| GO:0001568 | blood vessel development | 4.44E-12 | 2.159545204 | 1.79E-10 |
| GO:0050870 | positive regulation of T cell activation | 4.91E-12 | 3.291428571 | 1.95E-10 |
| GO:0002446 | neutrophil mediated immunity | 4.94E-12 | 8.43956044 | 1.95E-10 |
| GO:0051092 | positive regulation of NF-kappaB transcription factor activity | 5.12E-12 | 4.025157233 | 2.01E-10 |
| GO:0032755 | positive regulation of interleukin-6 production | 5.82E-12 | 4.777348777 | 2.26E-10 |
| GO:2000379 | positive regulation of reactive oxygen species metabolic process | 6.32E-12 | 4.46344207 | 2.42E-10 |
| GO:0002253 | activation of immune response | 6.34E-12 | 2.268424465 | 2.42E-10 |
| GO:0002757 | immune response-activating signal transduction | 7.58E-12 | 2.382293763 | 2.87E-10 |
| GO:0002764 | immune response-regulating signaling pathway | 8.51E-12 | 2.360462257 | 3.19E-10 |
| GO:0002687 | positive regulation of leukocyte migration | 9.39E-12 | 3.84962406 | 3.49E-10 |
| GO:0010604 | positive regulation of macromolecule metabolic process | 1.28E-11 | 1.443072265 | 4.71E-10 |
| GO:0006644 | phospholipid metabolic process | 1.74E-11 | 2.657283603 | 6.38E-10 |
| GO:0051403 | stress-activated MAPK cascade | 1.91E-11 | 3.069387755 | 6.95E-10 |
| GO:0009617 | response to bacterium | 1.98E-11 | 1.868389567 | 7.12E-10 |
| GO:0043123 | positive regulation of I-kappaB kinase/NF-kappaB signaling | 2.31E-11 | 4.022857143 | 8.24E-10 |
| GO:0006464 | cellular protein modification process | 3.58E-11 | 1.405879581 | 1.27E-09 |
| GO:0036211 | protein modification process | 3.71E-11 | 1.405166479 | 1.30E-09 |
| GO:0032940 | secretion by cell | 4.76E-11 | 1.962553442 | 1.66E-09 |
| GO:0001525 | angiogenesis | 5.38E-11 | 2.317907445 | 1.86E-09 |
| GO:0031401 | positive regulation of protein modification process | 5.50E-11 | 1.781684982 | 1.88E-09 |
| GO:0042093 | T-helper cell differentiation | 5.52E-11 | 5.830227743 | 1.88E-09 |
| GO:0002218 | activation of innate immune response | 5.81E-11 | 3.27386151 | 1.97E-09 |
| GO:1903530 | regulation of secretion by cell | 6.49E-11 | 2.142259414 | 2.18E-09 |
| GO:0006909 | phagocytosis | 6.55E-11 | 2.523594053 | 2.18E-09 |
| GO:0002294 | CD4-positive, alpha-beta T cell differentiation involved in immune response | 7.49E-11 | 5.746938776 | 2.48E-09 |
| GO:0070304 | positive regulation of stress-activated protein kinase signaling cascade | 8.15E-11 | 3.577639752 | 2.67E-09 |
| GO:0051046 | regulation of secretion | 8.66E-11 | 2.054320988 | 2.82E-09 |
| GO:0002532 | production of molecular mediator involved in inflammatory response | 8.81E-11 | 4.968944099 | 2.85E-09 |
| GO:0032655 | regulation of interleukin-12 production | 9.27E-11 | 6 | 2.97E-09 |
| GO:0009968 | negative regulation of signal transduction | 9.82E-11 | 1.790796808 | 3.13E-09 |
| GO:0002293 | alpha-beta T cell differentiation involved in immune response | 1.01E-10 | 5.665995976 | 3.19E-09 |
| GO:0002287 | alpha-beta T cell activation involved in immune response | 1.35E-10 | 5.587301587 | 4.25E-09 |
| GO:0002819 | regulation of adaptive immune response | 1.52E-10 | 3.12195122 | 4.73E-09 |
| GO:0007162 | negative regulation of cell adhesion | 1.64E-10 | 2.808777429 | 5.09E-09 |
| GO:0051130 | positive regulation of cellular component organization | 1.66E-10 | 1.69829153 | 5.12E-09 |
| GO:0031348 | negative regulation of defense response | 1.71E-10 | 3.059477487 | 5.23E-09 |
| GO:0042098 | T cell proliferation | 2.51E-10 | 3.180124224 | 7.62E-09 |
| GO:0001776 | leukocyte homeostasis | 2.66E-10 | 4.110741971 | 8.03E-09 |
| GO:1901653 | cellular response to peptide | 2.91E-10 | 2.605352431 | 8.72E-09 |
| GO:0002700 | regulation of production of molecular mediator of immune response | 3.12E-10 | 3.277628032 | 9.27E-09 |
| GO:0045597 | positive regulation of cell differentiation | 3.29E-10 | 1.793007641 | 9.71E-09 |
| GO:1902532 | negative regulation of intracellular signal transduction | 3.57E-10 | 2.253521127 | 1.04E-08 |
| GO:0022408 | negative regulation of cell-cell adhesion | 3.58E-10 | 3.262240107 | 1.04E-08 |
| GO:0032102 | negative regulation of response to external stimulus | 3.61E-10 | 2.648627139 | 1.04E-08 |
| GO:1902105 | regulation of leukocyte differentiation | 4.86E-10 | 2.62667719 | 1.40E-08 |
| GO:0051347 | positive regulation of transferase activity | 6.85E-10 | 2.190634657 | 1.96E-08 |
| GO:0043434 | response to peptide hormone | 7.26E-10 | 2.356994357 | 2.06E-08 |
| GO:0046903 | secretion | 7.39E-10 | 1.80638772 | 2.09E-08 |
| GO:0010934 | macrophage cytokine production | 7.67E-10 | 6.453781513 | 2.15E-08 |
| GO:0055082 | cellular chemical homeostasis | 7.73E-10 | 1.919604205 | 2.15E-08 |
| GO:0045619 | regulation of lymphocyte differentiation | 8.07E-10 | 3.114160948 | 2.23E-08 |
| GO:0051345 | positive regulation of hydrolase activity | 8.32E-10 | 2.166975881 | 2.29E-08 |
| GO:0050728 | negative regulation of inflammatory response | 8.47E-10 | 3.704948646 | 2.32E-08 |
| GO:1901342 | regulation of vasculature development | 8.54E-10 | 2.557105164 | 2.32E-08 |
| GO:1904018 | positive regulation of vasculature development | 9.16E-10 | 3.158441558 | 2.47E-08 |
| GO:1990266 | neutrophil migration | 9.65E-10 | 3.899159664 | 2.59E-08 |
| GO:0031329 | regulation of cellular catabolic process | 1.22E-09 | 2.072555852 | 3.25E-08 |
| GO:1900015 | regulation of cytokine production involved in inflammatory response | 1.25E-09 | 5.888619855 | 3.32E-08 |
| GO:0097190 | apoptotic signaling pathway | 1.43E-09 | 2.127583109 | 3.77E-08 |
| GO:0045807 | positive regulation of endocytosis | 1.72E-09 | 3.211149826 | 4.52E-08 |
| GO:0045055 | regulated exocytosis | 1.95E-09 | 2.752030578 | 5.07E-08 |
| GO:0050777 | negative regulation of immune response | 2.26E-09 | 3.117840685 | 5.85E-08 |
| GO:0001666 | response to hypoxia | 2.40E-09 | 2.597101449 | 6.19E-08 |
| GO:1903038 | negative regulation of leukocyte cell-cell adhesion | 2.67E-09 | 3.632923368 | 6.84E-08 |
| GO:0043300 | regulation of leukocyte degranulation | 3.06E-09 | 5.603686636 | 7.78E-08 |
| GO:0002822 | regulation of adaptive immune response based on somatic recombination of immune receptors built from immunoglobulin superfamily domains | 3.24E-09 | 3.021118012 | 8.21E-08 |
| GO:0050801 | ion homeostasis | 3.32E-09 | 1.888854003 | 8.37E-08 |
| GO:0030162 | regulation of proteolysis | 3.40E-09 | 1.974293059 | 8.52E-08 |
| GO:0031331 | positive regulation of cellular catabolic process | 3.50E-09 | 2.432090077 | 8.72E-08 |
| GO:1903706 | regulation of hemopoiesis | 3.87E-09 | 2.27638484 | 9.58E-08 |
| GO:0051051 | negative regulation of transport | 4.39E-09 | 2.168394437 | 1.08E-07 |
| GO:0051050 | positive regulation of transport | 4.46E-09 | 1.779591837 | 1.09E-07 |
| GO:0038061 | NIK/NF-kappaB signaling | 4.48E-09 | 4.155844156 | 1.09E-07 |
| GO:0006873 | cellular ion homeostasis | 4.70E-09 | 2.001172562 | 1.14E-07 |
| GO:0050868 | negative regulation of T cell activation | 5.23E-09 | 3.737226277 | 1.26E-07 |
| GO:0070302 | regulation of stress-activated protein kinase signaling cascade | 5.33E-09 | 2.868347339 | 1.28E-07 |
| GO:0045621 | positive regulation of lymphocyte differentiation | 6.07E-09 | 3.607385811 | 1.44E-07 |
| GO:0030099 | myeloid cell differentiation | 6.20E-09 | 2.305590062 | 1.47E-07 |
| GO:1903426 | regulation of reactive oxygen species biosynthetic process | 8.38E-09 | 3.896955504 | 1.96E-07 |
| GO:0032652 | regulation of interleukin-1 production | 8.38E-09 | 3.896955504 | 1.96E-07 |
| GO:0002283 | neutrophil activation involved in immune response | 9.06E-09 | 9.540372671 | 2.11E-07 |
| GO:0045766 | positive regulation of angiogenesis | 1.06E-08 | 3.124192391 | 2.45E-07 |
| GO:0010506 | regulation of autophagy | 1.21E-08 | 2.629318394 | 2.80E-07 |
| GO:0032872 | regulation of stress-activated MAPK cascade | 1.25E-08 | 2.829931973 | 2.87E-07 |
| GO:0045862 | positive regulation of proteolysis | 1.27E-08 | 2.468319559 | 2.90E-07 |
| GO:0071375 | cellular response to peptide hormone stimulus | 1.32E-08 | 2.587601078 | 3.00E-07 |
| GO:0032695 | negative regulation of interleukin-12 production | 1.34E-08 | 10.58646617 | 3.02E-07 |
| GO:0007015 | actin filament organization | 1.85E-08 | 2.256029685 | 4.16E-07 |
| GO:0071219 | cellular response to molecule of bacterial origin | 2.25E-08 | 2.5108742 | 5.04E-07 |
| GO:0043269 | regulation of ion transport | 2.62E-08 | 1.897142857 | 5.83E-07 |
| GO:0044267 | cellular protein metabolic process | 2.68E-08 | 1.27647748 | 5.93E-07 |
| GO:0048513 | animal organ development | 3.07E-08 | 1.327837932 | 6.76E-07 |
| GO:0010563 | negative regulation of phosphorus metabolic process | 3.31E-08 | 2.073637703 | 7.22E-07 |
| GO:0045936 | negative regulation of phosphate metabolic process | 3.31E-08 | 2.073637703 | 7.22E-07 |
| GO:0009896 | positive regulation of catabolic process | 3.55E-08 | 2.195800059 | 7.70E-07 |
| GO:0008154 | actin polymerization or depolymerization | 3.68E-08 | 2.862111801 | 7.96E-07 |
| GO:0002720 | positive regulation of cytokine production involved in immune response | 4.15E-08 | 4.363636364 | 8.93E-07 |
| GO:0043254 | regulation of protein complex assembly | 4.19E-08 | 2.119092123 | 8.97E-07 |
| GO:0045765 | regulation of angiogenesis | 4.21E-08 | 2.459482038 | 8.97E-07 |

| | | | | |
|---|---|---|---|---|
| GO:0051251 | positive regulation of lymphocyte activation | 4.33E-08 | 2.132750615 | 9.18E-07 |
| GO:0007169 | transmembrane receptor protein tyrosine kinase signaling pathway | 4.55E-08 | 2.05597469 | 9.61E-07 |
| GO:0007596 | blood coagulation | 4.95E-08 | 3.131807419 | 1.04E-06 |
| GO:0051248 | negative regulation of protein metabolic process | 5.54E-08 | 1.663641457 | 1.16E-06 |
| GO:0002886 | regulation of myeloid leukocyte mediated immunity | 6.44E-08 | 4.694980695 | 1.34E-06 |
| GO:0032956 | regulation of actin cytoskeleton organization | 6.96E-08 | 2.339425587 | 1.44E-06 |
| GO:0071417 | cellular response to organonitrogen compound | 6.98E-08 | 1.928872815 | 1.44E-06 |
| GO:0002702 | positive regulation of production of molecular mediator of immune response | 7.59E-08 | 3.404926108 | 1.56E-06 |
| GO:0071222 | cellular response to lipopolysaccharide | 7.82E-08 | 2.468010517 | 1.60E-06 |
| GO:0070665 | positive regulation of leukocyte proliferation | 8.22E-08 | 3.213852814 | 1.67E-06 |
| GO:0032611 | interleukin-1 beta production | 8.28E-08 | 4.022857143 | 1.67E-06 |
| GO:0010935 | regulation of macrophage cytokine production | 8.98E-08 | 5.962732919 | 1.81E-06 |
| GO:0043087 | regulation of GTPase activity | 1.02E-07 | 2.361067504 | 2.03E-06 |
| GO:0033003 | regulation of mast cell activation | 1.11E-07 | 5.096018735 | 2.22E-06 |
| GO:0042554 | superoxide anion generation | 1.21E-07 | 5.835866261 | 2.41E-06 |
| GO:1902563 | regulation of neutrophil activation | 1.30E-07 | 10.15873016 | 2.58E-06 |
| GO:0006887 | exocytosis | 1.38E-07 | 2.221628838 | 2.72E-06 |
| GO:1902107 | positive regulation of leukocyte differentiation | 1.42E-07 | 2.813186813 | 2.78E-06 |
| GO:0008284 | positive regulation of cell proliferation | 1.57E-07 | 1.692208959 | 3.07E-06 |
| GO:0016485 | protein processing | 1.66E-07 | 2.499047619 | 3.22E-06 |
| GO:1901222 | regulation of NIK/NF-kappaB signaling | 1.69E-07 | 3.721871049 | 3.27E-06 |
| GO:0098542 | defense response to other organism | 1.92E-07 | 1.781474964 | 3.70E-06 |
| GO:0001959 | regulation of cytokine-mediated signaling pathway | 1.94E-07 | 3.348088531 | 3.72E-06 |
| GO:0070664 | negative regulation of leukocyte proliferation | 1.99E-07 | 3.831292517 | 3.81E-06 |
| GO:0042542 | response to hydrogen peroxide | 2.04E-07 | 3.160493827 | 3.88E-06 |
| GO:0032269 | negative regulation of cellular protein metabolic process | 2.25E-07 | 1.651319683 | 4.26E-06 |
| GO:0002456 | T cell mediated immunity | 2.30E-07 | 3.226890756 | 4.34E-06 |
| GO:1903556 | negative regulation of tumor necrosis factor superfamily cytokine production | 2.37E-07 | 4.571428571 | 4.45E-06 |
| GO:0030183 | B cell differentiation | 2.59E-07 | 3.047619048 | 4.84E-06 |
| GO:0050672 | negative regulation of lymphocyte proliferation | 2.74E-07 | 3.918367347 | 5.11E-06 |
| GO:0002705 | positive regulation of leukocyte mediated immunity | 2.83E-07 | 2.840499307 | 5.24E-06 |
| GO:0072359 | circulatory system development | 3.25E-07 | 1.620616246 | 6.00E-06 |
| GO:0032945 | negative regulation of mononuclear cell proliferation | 3.27E-07 | 3.878787879 | 6.00E-06 |
| GO:0001667 | ameboidal-type cell migration | 3.47E-07 | 2.118945432 | 6.35E-06 |
| GO:0032651 | regulation of interleukin-1 beta production | 3.87E-07 | 3.84 | 7.06E-06 |
| GO:0043030 | regulation of macrophage activation | 4.31E-07 | 4.958837772 | 7.83E-06 |
| GO:0045428 | regulation of nitric oxide biosynthetic process | 4.44E-07 | 4.388571429 | 8.03E-06 |
| GO:0046486 | glycerolipid metabolic process | 4.52E-07 | 2.243938829 | 8.12E-06 |
| GO:2000401 | regulation of lymphocyte migration | 4.53E-07 | 4.639658849 | 8.12E-06 |
| GO:0070555 | response to interleukin-1 | 4.70E-07 | 3.401993355 | 8.39E-06 |
| GO:0031663 | lipopolysaccharide-mediated signaling pathway | 4.87E-07 | 5.274725275 | 8.66E-06 |
| GO:0006809 | nitric oxide biosynthetic process | 5.02E-07 | 4.136054422 | 8.89E-06 |
| GO:0051129 | negative regulation of cellular component organization | 5.21E-07 | 1.754272815 | 9.20E-06 |
| GO:0002688 | regulation of leukocyte chemotaxis | 5.42E-07 | 3.375824176 | 9.52E-06 |
| GO:2001233 | regulation of apoptotic signaling pathway | 5.85E-07 | 2.138925295 | 1.02E-05 |
| GO:0031400 | negative regulation of protein modification process | 5.99E-07 | 1.905354366 | 1.04E-05 |
| GO:0051604 | protein maturation | 6.23E-07 | 2.292226292 | 1.08E-05 |
| GO:0043124 | negative regulation of I-kappaB kinase/NF-kappaB signaling | 6.29E-07 | 5.175202156 | 1.09E-05 |
| GO:1903825 | organic acid transmembrane transport | 6.38E-07 | 3.72815534 | 1.10E-05 |
| GO:0002526 | acute inflammatory response | 6.49E-07 | 3.148533586 | 1.11E-05 |
| GO:0008064 | regulation of actin polymerization or depolymerization | 6.50E-07 | 2.792399719 | 1.11E-05 |
| GO:0080171 | lytic vacuole organization | 6.62E-07 | 4.27458256 | 1.13E-05 |
| GO:0007252 | I-kappaB phosphorylation | 6.78E-07 | 8.707482993 | 1.15E-05 |
| GO:1903305 | regulation of regulated secretory pathway | 7.24E-07 | 2.778711485 | 1.22E-05 |
| GO:0043069 | negative regulation of programmed cell death | 7.82E-07 | 1.654740141 | 1.32E-05 |
| GO:0032715 | negative regulation of interleukin-6 production | 8.06E-07 | 5.079365079 | 1.35E-05 |
| GO:0072538 | T-helper 17 type immune response | 8.88E-07 | 5.446808511 | 1.48E-05 |
| GO:0030832 | regulation of actin filament length | 8.93E-07 | 2.751733703 | 1.49E-05 |
| GO:0030833 | regulation of actin filament polymerization | 9.35E-07 | 2.866409266 | 1.55E-05 |
| GO:0016236 | macroautophagy | 9.58E-07 | 2.549800797 | 1.58E-05 |
| GO:0010631 | epithelial cell migration | 1.07E-06 | 2.299401198 | 1.77E-05 |
| GO:0071621 | granulocyte chemotaxis | 1.08E-06 | 3.250793651 | 1.77E-05 |
| GO:0006508 | proteolysis | 1.25E-06 | 1.472876712 | 2.04E-05 |
| GO:0032653 | regulation of interleukin-10 production | 1.33E-06 | 4.571428571 | 2.17E-05 |
| GO:0015031 | protein transport | 1.39E-06 | 1.480666559 | 2.26E-05 |
| GO:0032479 | regulation of type I interferon production | 1.40E-06 | 3.555555556 | 2.26E-05 |
| GO:1903708 | positive regulation of hemopoiesis | 1.45E-06 | 2.465489567 | 2.33E-05 |
| GO:0002698 | negative regulation of immune effector process | 1.48E-06 | 2.93877551 | 2.37E-05 |
| GO:0042886 | amide transport | 1.52E-06 | 1.464036232 | 2.43E-05 |
| GO:0034341 | response to interferon-gamma | 1.54E-06 | 3.009041591 | 2.46E-05 |
| GO:0034614 | cellular response to reactive oxygen species | 1.56E-06 | 2.860335196 | 2.47E-05 |
| GO:0032649 | regulation of interferon-gamma production | 1.58E-06 | 3.285714286 | 2.50E-05 |
| GO:0060548 | negative regulation of cell death | 1.70E-06 | 1.590708479 | 2.68E-05 |
| GO:0061081 | positive regulation of myeloid leukocyte cytokine production involved in immune response | 2.11E-06 | 5.528239203 | 3.30E-05 |
| GO:0032733 | positive regulation of interleukin-10 production | 2.11E-06 | 5.528239203 | 3.30E-05 |
| GO:0031334 | positive regulation of protein complex assembly | 2.16E-06 | 2.292879534 | 3.37E-05 |
| GO:0032271 | regulation of protein polymerization | 2.25E-06 | 2.496844521 | 3.49E-05 |
| GO:0080135 | regulation of cellular response to stress | 2.25E-06 | 1.732100034 | 3.49E-05 |
| GO:0045453 | bone resorption | 2.28E-06 | 4.144761905 | 3.51E-05 |
| GO:0009620 | response to fungus | 2.28E-06 | 4.144761905 | 3.51E-05 |
| GO:0015833 | peptide transport | 2.34E-06 | 1.459657893 | 3.59E-05 |
| GO:0042088 | T-helper 1 type immune response | 2.45E-06 | 5.019607843 | 3.74E-05 |
| GO:0050854 | regulation of antigen receptor-mediated signaling pathway | 2.48E-06 | 4.36673774 | 3.77E-05 |
| GO:0072539 | T-helper 17 cell differentiation | 2.50E-06 | 6.704761905 | 3.78E-05 |
| GO:0043032 | positive regulation of macrophage activation | 2.50E-06 | 6.704761905 | 3.78E-05 |
| GO:0032868 | response to insulin | 2.51E-06 | 2.33958634 | 3.78E-05 |
| GO:0051493 | regulation of cytoskeleton organization | 2.57E-06 | 1.880123194 | 3.86E-05 |
| GO:0000902 | cell morphogenesis | 2.63E-06 | 1.605418287 | 3.93E-05 |
| GO:0007229 | integrin-mediated signaling pathway | 2.64E-06 | 3.297423888 | 3.93E-05 |
| GO:1905954 | positive regulation of lipid localization | 2.64E-06 | 3.297423888 | 3.93E-05 |
| GO:0006461 | protein complex assembly | 2.67E-06 | 1.587407584 | 3.95E-05 |
| GO:0001774 | microglial cell activation | 2.75E-06 | 5.402597403 | 4.07E-05 |
| GO:0017157 | regulation of exocytosis | 2.86E-06 | 2.393766234 | 4.21E-05 |
| GO:0006820 | anion transport | 3.02E-06 | 1.848792598 | 4.44E-05 |
| GO:0006812 | cation transport | 3.56E-06 | 1.572481572 | 5.20E-05 |
| GO:1903034 | regulation of response to wounding | 3.57E-06 | 2.576623377 | 5.20E-05 |
| GO:0009891 | positive regulation of biosynthetic process | 3.97E-06 | 1.405941904 | 5.77E-05 |
| GO:0015711 | organic anion transport | 4.07E-06 | 1.969802555 | 5.90E-05 |
| GO:0032720 | negative regulation of tumor necrosis factor production | 4.43E-06 | 4.179591837 | 6.40E-05 |
| GO:0006650 | glycerophospholipid metabolic process | 4.69E-06 | 2.342653787 | 6.74E-05 |
| GO:0002313 | mature B cell differentiation involved in immune response | 4.86E-06 | 6.285714286 | 6.97E-05 |
| GO:0034220 | ion transmembrane transport | 5.03E-06 | 1.629420085 | 7.19E-05 |
| GO:0043066 | negative regulation of apoptotic process | 5.81E-06 | 1.601917188 | 8.27E-05 |
| GO:1902622 | regulation of neutrophil migration | 6.09E-06 | 4.654545455 | 8.65E-05 |
| GO:0030258 | lipid modification | 6.38E-06 | 2.344322344 | 9.03E-05 |
| GO:0097191 | extrinsic apoptotic signaling pathway | 6.46E-06 | 2.458583433 | 9.11E-05 |

| | | | | |
|---|---|---|---|---|
| GO:0071396 | cellular response to lipid | 6.49E-06 | 1.726852707 | 9.13E-05 |
| GO:0032608 | interferon-beta production | 7.13E-06 | 4.285714286 | 9.99E-05 |
| GO:0002833 | positive regulation of response to biotic stimulus | 7.66E-06 | 4.007827789 | 1.07E-04 |
| GO:0050920 | regulation of chemotaxis | 7.67E-06 | 2.438095238 | 1.07E-04 |
| GO:0050671 | positive regulation of lymphocyte proliferation | 7.69E-06 | 2.906338694 | 1.07E-04 |
| GO:0044070 | regulation of anion transport | 8.12E-06 | 2.982776089 | 1.12E-04 |
| GO:0072676 | lymphocyte migration | 8.50E-06 | 3.294723295 | 1.17E-04 |
| GO:0009306 | protein secretion | 8.86E-06 | 2.031746032 | 1.22E-04 |
| GO:1901224 | positive regulation of NIK/NF-kappaB signaling | 9.14E-06 | 3.745266781 | 1.25E-04 |
| GO:0002534 | cytokine production involved in inflammatory response | 9.29E-06 | 4.49122807 | 1.27E-04 |
| GO:2000404 | regulation of T cell migration | 9.35E-06 | 4.851311953 | 1.27E-04 |
| GO:0009615 | response to virus | 1.02E-05 | 2.080120937 | 1.39E-04 |
| GO:0032731 | positive regulation of interleukin-1 beta production | 1.04E-05 | 4.155844156 | 1.41E-04 |
| GO:0032946 | positive regulation of mononuclear cell proliferation | 1.07E-05 | 2.849721707 | 1.45E-04 |
| GO:0002262 | myeloid cell homeostasis | 1.08E-05 | 2.585858586 | 1.45E-04 |
| GO:0090322 | regulation of superoxide metabolic process | 1.10E-05 | 5.224489796 | 1.48E-04 |
| GO:1902903 | regulation of supramolecular fiber organization | 1.12E-05 | 2.031746032 | 1.49E-04 |
| GO:0032480 | negative regulation of type I interferon production | 1.13E-05 | 6.530612245 | 1.50E-04 |
| GO:0007009 | plasma membrane organization | 1.13E-05 | 2.012303486 | 1.50E-04 |
| GO:0007167 | enzyme linked receptor protein signaling pathway | 1.14E-05 | 1.616209074 | 1.52E-04 |
| GO:0010508 | positive regulation of autophagy | 1.35E-05 | 2.737382378 | 1.78E-04 |
| GO:2001236 | regulation of extrinsic apoptotic signaling pathway | 1.35E-05 | 2.737382378 | 1.78E-04 |
| GO:0030168 | platelet activation | 1.37E-05 | 3.464661654 | 1.80E-04 |
| GO:0033619 | membrane protein proteolysis | 1.39E-05 | 4.338983051 | 1.82E-04 |
| GO:1902624 | positive regulation of neutrophil migration | 1.40E-05 | 5.102990033 | 1.84E-04 |
| GO:0030838 | positive regulation of actin filament polymerization | 1.43E-05 | 3.308843537 | 1.86E-04 |
| GO:0032648 | regulation of interferon-beta production | 1.50E-05 | 4.033613445 | 1.94E-04 |
| GO:2000191 | regulation of fatty acid transport | 1.57E-05 | 5.587301587 | 2.02E-04 |
| GO:0002719 | negative regulation of cytokine production involved in immune response | 1.57E-05 | 5.587301587 | 2.02E-04 |
| GO:0002706 | regulation of lymphocyte mediated immunity | 1.59E-05 | 2.391802291 | 2.05E-04 |
| GO:1903793 | positive regulation of anion transport | 1.78E-05 | 3.750915751 | 2.29E-04 |
| GO:0002888 | positive regulation of myeloid leukocyte mediated immunity | 1.80E-05 | 4.571428571 | 2.30E-04 |
| GO:0043313 | regulation of neutrophil degranulation | 1.82E-05 | 10.66666667 | 2.33E-04 |
| GO:0002821 | positive regulation of adaptive immune response | 1.83E-05 | 2.68907563 | 2.33E-04 |
| GO:0032732 | positive regulation of interleukin-1 production | 2.09E-05 | 3.703435805 | 2.65E-04 |
| GO:0032735 | positive regulation of interleukin-12 production | 2.24E-05 | 4.876190476 | 2.83E-04 |
| GO:0032660 | regulation of interleukin-17 production | 2.24E-05 | 4.876190476 | 2.83E-04 |
| GO:0061041 | regulation of wound healing | 2.41E-05 | 2.583850932 | 3.03E-04 |
| GO:0008285 | negative regulation of cell proliferation | 2.48E-05 | 1.666546112 | 3.11E-04 |
| GO:0032692 | negative regulation of interleukin-1 production | 2.64E-05 | 5.293233083 | 3.29E-04 |
| GO:0050869 | negative regulation of B cell activation | 2.64E-05 | 5.293233083 | 3.29E-04 |
| GO:0050708 | regulation of protein secretion | 2.78E-05 | 2.105627706 | 3.46E-04 |
| GO:0045123 | cellular extravasation | 2.85E-05 | 3.611992945 | 3.54E-04 |
| GO:0007264 | small GTPase mediated signal transduction | 2.89E-05 | 1.938117182 | 3.58E-04 |
| GO:0045806 | negative regulation of endocytosis | 2.96E-05 | 3.80952381 | 3.65E-04 |
| GO:0010634 | positive regulation of epithelial cell migration | 3.02E-05 | 2.675958188 | 3.72E-04 |
| GO:0048660 | regulation of smooth muscle cell proliferation | 3.05E-05 | 2.493506494 | 3.73E-04 |
| GO:0032890 | regulation of organic acid transport | 3.05E-05 | 3.416012559 | 3.73E-04 |
| GO:0031328 | positive regulation of cellular biosynthetic process | 3.06E-05 | 1.368613386 | 3.73E-04 |
| GO:1902905 | positive regulation of supramolecular fiber organization | 3.23E-05 | 2.388674389 | 3.93E-04 |
| GO:0002690 | positive regulation of leukocyte chemotaxis | 3.57E-05 | 3.226890756 | 4.33E-04 |
| GO:0051047 | positive regulation of secretion | 3.62E-05 | 1.920417482 | 4.38E-04 |
| GO:2000193 | positive regulation of fatty acid transport | 3.63E-05 | 6.582857143 | 4.38E-04 |
| GO:0043112 | receptor metabolic process | 3.94E-05 | 2.568218299 | 4.75E-04 |
| GO:0051223 | regulation of protein transport | 3.97E-05 | 1.738736699 | 4.76E-04 |
| GO:0052547 | regulation of peptidase activity | 3.97E-05 | 1.897193314 | 4.76E-04 |
| GO:0070486 | leukocyte aggregation | 4.02E-05 | 7.69924812 | 4.80E-04 |
| GO:0032273 | positive regulation of protein polymerization | 4.08E-05 | 2.774384236 | 4.86E-04 |
| GO:0008643 | carbohydrate transport | 4.10E-05 | 2.695970696 | 4.87E-04 |
| GO:0043302 | positive regulation of leukocyte degranulation | 4.28E-05 | 5.028571429 | 5.06E-04 |
| GO:0045622 | regulation of T-helper cell differentiation | 4.28E-05 | 5.028571429 | 5.06E-04 |
| GO:1900077 | negative regulation of cellular response to insulin stimulus | 4.29E-05 | 4.571428571 | 5.06E-04 |
| GO:0022604 | regulation of cell morphogenesis | 4.41E-05 | 1.795324675 | 5.18E-04 |
| GO:0060284 | regulation of cell development | 4.45E-05 | 1.509544073 | 5.21E-04 |
| GO:0051495 | positive regulation of cytoskeleton organization | 4.47E-05 | 2.267428571 | 5.23E-04 |
| GO:0032870 | cellular response to hormone stimulus | 4.53E-05 | 1.750117869 | 5.29E-04 |
| GO:0009395 | phospholipid catabolic process | 4.79E-05 | 4.170426065 | 5.56E-04 |
| GO:0010628 | positive regulation of gene expression | 4.79E-05 | 1.347195482 | 5.56E-04 |
| GO:0043409 | negative regulation of MAPK cascade | 4.95E-05 | 2.476190476 | 5.73E-04 |
| GO:0032722 | positive regulation of chemokine production | 5.13E-05 | 3.442016807 | 5.92E-04 |
| GO:0030593 | neutrophil chemotaxis | 5.23E-05 | 3.134693878 | 6.01E-04 |
| GO:1903531 | negative regulation of secretion by cell | 5.40E-05 | 2.463360474 | 6.20E-04 |
| GO:0046942 | carboxylic acid transport | 5.68E-05 | 1.992993383 | 6.47E-04 |
| GO:1902106 | negative regulation of leukocyte differentiation | 5.68E-05 | 2.995073892 | 6.47E-04 |
| GO:0010720 | positive regulation of cell development | 5.68E-05 | 1.674044266 | 6.47E-04 |
| GO:0043373 | CD4-positive, alpha-beta T cell lineage commitment | 5.89E-05 | 7.314285714 | 6.69E-04 |
| GO:0071346 | cellular response to interferon-gamma | 5.97E-05 | 2.879640045 | 6.76E-04 |
| GO:0010543 | regulation of platelet activation | 6.42E-05 | 4.388571429 | 7.25E-04 |
| GO:0033006 | regulation of mast cell activation involved in immune response | 6.73E-05 | 4.789115646 | 7.59E-04 |
| GO:0015849 | organic acid transport | 6.77E-05 | 1.976833977 | 7.60E-04 |
| GO:1903532 | positive regulation of secretion by cell | 6.77E-05 | 1.976833977 | 7.60E-04 |
| GO:0046434 | organophosphate catabolic process | 7.31E-05 | 2.596119929 | 8.16E-04 |
| GO:0008360 | regulation of cell shape | 7.31E-05 | 2.596119929 | 8.16E-04 |
| GO:0002790 | peptide secretion | 7.60E-05 | 1.832388905 | 8.46E-04 |
| GO:0090407 | organophosphate biosynthetic process | 7.76E-05 | 1.768508863 | 8.61E-04 |
| GO:0048661 | positive regulation of smooth muscle cell proliferation | 7.99E-05 | 2.919567827 | 8.85E-04 |
| GO:0002824 | positive regulation of adaptive immune response based on somatic recombination of immune receptors built from immunoglobulin superfamily domains | 8.03E-05 | 2.580192813 | 8.87E-04 |
| GO:0050832 | defense response to fungus | 8.14E-05 | 3.961904762 | 8.97E-04 |
| GO:1900026 | positive regulation of substrate adhesion-dependent cell spreading | 8.34E-05 | 4.677740864 | 9.15E-04 |
| GO:0046627 | negative regulation of insulin receptor signaling pathway | 8.34E-05 | 4.677740864 | 9.15E-04 |
| GO:0070498 | interleukin-1-mediated signaling pathway | 8.42E-05 | 6.965986395 | 9.21E-04 |
| GO:0032688 | negative regulation of interferon-beta production | 8.56E-05 | 8.533333333 | 9.32E-04 |
| GO:0032494 | response to peptidoglycan | 8.56E-05 | 8.533333333 | 9.32E-04 |
| GO:0002228 | natural killer cell mediated immunity | 8.88E-05 | 3.287319422 | 9.64E-04 |
| GO:0070371 | ERK1 and ERK2 cascade | 9.09E-05 | 2.013605442 | 9.85E-04 |
| GO:0010811 | positive regulation of cell-substrate adhesion | 9.19E-05 | 2.704225352 | 9.93E-04 |
| GO:0070301 | cellular response to hydrogen peroxide | 9.52E-05 | 2.992207792 | 0.001024199 |
| GO:2001235 | positive regulation of apoptotic signaling pathway | 9.53E-05 | 2.431610942 | 0.001024199 |
| GO:0032757 | positive regulation of interleukin-8 production | 9.64E-05 | 3.896955504 | 0.001032305 |
| GO:1902565 | positive regulation of neutrophil activation | 9.65E-05 | 10.97142857 | 0.001032305 |
| GO:0032088 | negative regulation of NF-kappaB transcription factor activity | 9.94E-05 | 3.428571429 | 0.001061375 |
| GO:0034762 | regulation of transmembrane transport | 1.01E-04 | 1.760846561 | 0.001075059 |
| GO:0051048 | negative regulation of secretion | 1.05E-04 | 2.275555556 | 0.001114023 |
| GO:1903428 | positive regulation of reactive oxygen species biosynthetic process | 1.08E-04 | 3.605633803 | 0.001148228 |
| GO:0009914 | hormone transport | 1.09E-04 | 1.896296296 | 0.001155162 |
| GO:1903317 | regulation of protein maturation | 1.12E-04 | 3.077793494 | 0.001184492 |

| | | | | |
|---|---|---|---|---|
| GO:0042267 | natural killer cell mediated cytotoxicity | 1.14E-04 | 3.386243386 | 0.001202203 |
| GO:0046879 | hormone secretion | 1.25E-04 | 1.902828136 | 0.001307745 |
| GO:2000021 | regulation of ion homeostasis | 1.28E-04 | 2.20952381 | 0.00134531 |
| GO:0001771 | immunological synapse formation | 1.31E-04 | 8 | 0.001362421 |
| GO:1903978 | regulation of microglial cell activation | 1.31E-04 | 8 | 0.001362421 |
| GO:0008654 | phospholipid biosynthetic process | 1.33E-04 | 2.380952381 | 0.001378788 |
| GO:0032642 | regulation of chemokine production | 1.34E-04 | 2.912768647 | 0.001387884 |
| GO:1904407 | positive regulation of nitric oxide metabolic process | 1.35E-04 | 4.063492063 | 0.001393507 |
| GO:0090087 | regulation of peptide transport | 1.41E-04 | 1.652486772 | 0.001451007 |
| GO:0070542 | response to fatty acid | 1.42E-04 | 3.018030513 | 0.001466593 |
| GO:0051173 | positive regulation of nitrogen compound metabolic process | 1.46E-04 | 1.321957791 | 0.001496235 |
| GO:0010595 | positive regulation of endothelial cell migration | 1.49E-04 | 2.887218045 | 0.001531654 |
| GO:0032892 | positive regulation of organic acid transport | 1.60E-04 | 3.98961039 | 0.00163778 |
| GO:0002791 | regulation of peptide secretion | 1.61E-04 | 1.917050691 | 0.001643248 |
| GO:0070201 | regulation of establishment of protein localization | 1.62E-04 | 1.65232358 | 0.00164862 |
| GO:0002675 | positive regulation of acute inflammatory response | 1.62E-04 | 4.812030075 | 0.00164862 |
| GO:0034765 | regulation of ion transmembrane transport | 1.76E-04 | 1.744820065 | 0.001782689 |
| GO:0006865 | amino acid transport | 1.78E-04 | 2.445182724 | 0.001801426 |
| GO:0010959 | regulation of metal ion transport | 1.79E-04 | 1.793481572 | 0.001809162 |
| GO:0050820 | positive regulation of coagulation | 1.85E-04 | 4.279635258 | 0.001859894 |
| GO:0019751 | polyol metabolic process | 1.85E-04 | 2.837438424 | 0.001861992 |
| GO:0002295 | T-helper cell lineage commitment | 1.93E-04 | 7.529411765 | 0.001931235 |
| GO:0019216 | regulation of lipid metabolic process | 1.95E-04 | 1.846153846 | 0.001953215 |
| GO:0051056 | regulation of small GTPase mediated signal transduction | 1.96E-04 | 2.155632985 | 0.001958803 |
| GO:0048535 | lymph node development | 1.98E-04 | 5.30875576 | 0.001966681 |
| GO:0034446 | substrate adhesion-dependent cell spreading | 2.01E-04 | 2.932614555 | 0.00199008 |
| GO:0043304 | regulation of mast cell degranulation | 2.01E-04 | 4.688644689 | 0.00199008 |
| GO:0002768 | immune response-regulating cell surface receptor signaling pathway | 2.06E-04 | 1.841726619 | 0.002033037 |
| GO:0031324 | negative regulation of cellular metabolic process | 2.08E-04 | 1.267201267 | 0.002052038 |
| GO:1903707 | negative regulation of hemopoiesis | 2.11E-04 | 2.417077176 | 0.002073367 |
| GO:0030163 | protein catabolic process | 2.14E-04 | 1.495618306 | 0.002097621 |
| GO:0045063 | T-helper 1 cell differentiation | 2.17E-04 | 6.095238095 | 0.002125822 |
| GO:0002279 | mast cell activation involved in immune response | 2.21E-04 | 3.368421053 | 0.002159605 |
| GO:0010557 | positive regulation of macromolecule biosynthetic process | 2.22E-04 | 1.335265141 | 0.002159605 |
| GO:0002429 | immune response-activating cell surface receptor signaling pathway | 2.22E-04 | 1.851146384 | 0.002159605 |
| GO:0097193 | intrinsic apoptotic signaling pathway | 2.24E-04 | 1.992673993 | 0.002178363 |
| GO:0042742 | defense response to bacterium | 2.28E-04 | 1.685031692 | 0.002214875 |
| GO:0042832 | defense response to protozoan | 2.47E-04 | 4.571428571 | 0.0023859 |
| GO:0045893 | positive regulation of transcription, DNA-templated | 2.58E-04 | 1.364120782 | 0.002488541 |
| GO:0002701 | negative regulation of production of molecular mediator of immune response | 2.65E-04 | 4.104956268 | 0.002551199 |
| GO:0050856 | regulation of T cell receptor signaling pathway | 2.65E-04 | 4.104956268 | 0.002551199 |
| GO:0070163 | regulation of adiponectin secretion | 2.71E-04 | 13.06122449 | 0.002597365 |
| GO:0070162 | adiponectin secretion | 2.71E-04 | 13.06122449 | 0.002597365 |
| GO:0070423 | nucleotide-binding oligomerization domain containing signaling pathway | 2.76E-04 | 7.111111111 | 0.002632416 |
| GO:0002448 | mast cell mediated immunity | 2.83E-04 | 3.495798319 | 0.002698528 |
| GO:0010507 | negative regulation of autophagy | 2.89E-04 | 3.282051282 | 0.002744079 |
| GO:0010632 | regulation of epithelial cell migration | 2.93E-04 | 2.10430839 | 0.002781588 |
| GO:0032370 | positive regulation of lipid transport | 3.01E-04 | 2.955266955 | 0.00284215 |
| GO:0030193 | regulation of blood coagulation | 3.01E-04 | 2.955266955 | 0.00284215 |
| GO:0032368 | regulation of lipid transport | 3.06E-04 | 2.477419355 | 0.002883899 |
| GO:1900024 | regulation of substrate adhesion-dependent cell spreading | 3.07E-04 | 3.719128329 | 0.002883899 |
| GO:1902680 | positive regulation of RNA biosynthetic process | 3.11E-04 | 1.356890459 | 0.002915412 |
| GO:1905521 | regulation of macrophage migration | 3.16E-04 | 4.022857143 | 0.002957805 |
| GO:0031344 | regulation of cell projection organization | 3.18E-04 | 1.540077953 | 0.00297389 |
| GO:0051607 | defense response to virus | 3.20E-04 | 2.003913894 | 0.002986258 |
| GO:0034612 | response to tumor necrosis factor | 3.26E-04 | 2.124481328 | 0.00302854 |
| GO:0032663 | regulation of interleukin-2 production | 3.26E-04 | 3.445134576 | 0.00302854 |
| GO:0032677 | regulation of interleukin-8 production | 3.29E-04 | 3.240506329 | 0.003044572 |
| GO:0015908 | fatty acid transport | 3.44E-04 | 2.800514801 | 0.00317828 |
| GO:0010803 | regulation of tumor necrosis factor-mediated signaling pathway | 3.65E-04 | 4.353741497 | 0.003357878 |
| GO:2000403 | positive regulation of lymphocyte migration | 3.65E-04 | 4.353741497 | 0.003357878 |
| GO:0051254 | positive regulation of RNA metabolic process | 3.67E-04 | 1.343482093 | 0.003369657 |
| GO:0002753 | cytoplasmic pattern recognition receptor signaling pathway | 3.74E-04 | 3.943977591 | 0.003413916 |
| GO:0006023 | aminoglycan biosynthetic process | 3.74E-04 | 3.395918367 | 0.003413916 |
| GO:0032303 | regulation of icosanoid secretion | 3.75E-04 | 5.626373626 | 0.003413916 |
| GO:0032930 | positive regulation of superoxide anion generation | 3.75E-04 | 5.626373626 | 0.003413916 |
| GO:0032963 | collagen metabolic process | 3.78E-04 | 2.675958188 | 0.003433951 |
| GO:0035872 | nucleotide-binding domain, leucine rich repeat containing receptor signaling pathway | 3.84E-04 | 6.736842105 | 0.003489872 |
| GO:0050921 | positive regulation of chemotaxis | 3.95E-04 | 2.430379747 | 0.003577909 |
| GO:0045620 | negative regulation of lymphocyte differentiation | 4.15E-04 | 3.597189696 | 0.003747957 |
| GO:0000768 | syncytium formation by plasma membrane fusion | 4.28E-04 | 3.348088531 | 0.003864455 |
| GO:0010940 | positive regulation of necrotic cell death | 4.29E-04 | 8.43956044 | 0.003864455 |
| GO:0050851 | antigen receptor-mediated signaling pathway | 4.38E-04 | 1.858712716 | 0.003935275 |
| GO:0045429 | positive regulation of nitric oxide biosynthetic process | 4.41E-04 | 3.868131868 | 0.003950305 |
| GO:0046883 | regulation of hormone secretion | 4.72E-04 | 1.912967033 | 0.004225352 |
| GO:1902582 | single-organism intracellular transport | 4.74E-04 | 1.516688919 | 0.004234413 |
| GO:0002673 | regulation of acute inflammatory response | 4.77E-04 | 3.12195122 | 0.004254709 |
| GO:0006691 | leukotriene metabolic process | 4.82E-04 | 5.417989418 | 0.004282282 |
| GO:2001234 | negative regulation of apoptotic signaling pathway | 4.84E-04 | 2.072874494 | 0.004282282 |
| GO:0043270 | positive regulation of ion transport | 4.84E-04 | 1.848555816 | 0.004282282 |
| GO:0051271 | negative regulation of cellular component movement | 4.84E-04 | 1.848555816 | 0.004282282 |
| GO:2000406 | positive regulation of T cell migration | 4.85E-04 | 4.702040816 | 0.004282282 |
| GO:1901214 | regulation of neuron death | 4.89E-04 | 1.797121797 | 0.004305448 |
| GO:0008630 | intrinsic apoptotic signaling pathway in response to DNA damage | 5.15E-04 | 2.70310559 | 0.004523948 |
| GO:0032495 | response to muramyl dipeptide | 5.24E-04 | 6.4 | 0.00458723 |
| GO:0030220 | platelet formation | 5.24E-04 | 6.4 | 0.00458723 |
| GO:0051208 | sequestering of calcium ion | 5.51E-04 | 2.59167604 | 0.004810042 |
| GO:0046834 | lipid phosphorylation | 5.53E-04 | 3.482993197 | 0.004810042 |
| GO:0030865 | cortical cytoskeleton organization | 5.53E-04 | 3.482993197 | 0.004810042 |
| GO:0006949 | syncytium formation | 5.57E-04 | 3.256360078 | 0.004824655 |
| GO:0034121 | regulation of toll-like receptor signaling pathway | 5.57E-04 | 3.256360078 | 0.004824655 |
| GO:0071674 | mononuclear cell migration | 5.65E-04 | 2.917933131 | 0.004888481 |
| GO:0002708 | positive regulation of lymphocyte mediated immunity | 5.93E-04 | 2.355828221 | 0.005117775 |
| GO:0032740 | positive regulation of interleukin-17 production | 6.13E-04 | 5.224489796 | 0.005279722 |
| GO:0006869 | lipid transport | 6.17E-04 | 1.759892689 | 0.00530763 |
| GO:0043316 | cytotoxic T cell degranulation | 6.24E-04 | 18.28571429 | 0.005348755 |
| GO:0002369 | T cell cytokine production | 6.25E-04 | 4.063492063 | 0.005348755 |
| GO:0001562 | response to protozoan | 6.25E-04 | 4.063492063 | 0.005348755 |
| GO:0098792 | xenophagy | 6.38E-04 | 7.836734694 | 0.005443164 |
| GO:0050767 | regulation of neurogenesis | 6.50E-04 | 1.454953691 | 0.005539455 |
| GO:0021782 | glial cell development | 6.60E-04 | 2.551495017 | 0.005616096 |
| GO:1903725 | regulation of phospholipid metabolic process | 6.80E-04 | 3.011764706 | 0.005768153 |
| GO:0032305 | positive regulation of icosanoid secretion | 7.00E-04 | 6.095238095 | 0.005916853 |
| GO:0060099 | regulation of phagocytosis, engulfment | 7.00E-04 | 6.095238095 | 0.005916853 |
| GO:0045124 | regulation of bone resorption | 7.02E-04 | 3.657142857 | 0.00592472 |
| GO:0033209 | tumor necrosis factor-mediated signaling pathway | 7.15E-04 | 3.16952381 | 0.006011134 |

| | | | | |
|---|---|---|---|---|
| GO:0044744 | protein targeting to nucleus | 7.18E-04 | 2.126245847 | 0.006011134 |
| GO:0006606 | protein import into nucleus | 7.18E-04 | 2.126245847 | 0.006011134 |
| GO:1902593 | single-organism nuclear import | 7.18E-04 | 2.126245847 | 0.006011134 |
| GO:1903364 | positive regulation of cellular protein catabolic process | 7.43E-04 | 2.44668008 | 0.006207365 |
| GO:0051767 | nitric-oxide synthase biosynthetic process | 7.70E-04 | 5.044334975 | 0.006410223 |
| GO:0051769 | regulation of nitric-oxide synthase biosynthetic process | 7.70E-04 | 5.044334975 | 0.006410223 |
| GO:0061515 | myeloid cell development | 7.79E-04 | 2.827687776 | 0.006475067 |
| GO:0035296 | regulation of tube diameter | 7.94E-04 | 2.247406225 | 0.006572298 |
| GO:0050880 | regulation of blood vessel size | 7.94E-04 | 2.247406225 | 0.006572298 |
| GO:0050830 | defense response to Gram-positive bacterium | 8.29E-04 | 2.19047619 | 0.006849435 |
| GO:0043303 | mast cell degranulation | 8.30E-04 | 3.324675325 | 0.006849435 |
| GO:0009100 | glycoprotein metabolic process | 8.43E-04 | 1.760846561 | 0.006944871 |
| GO:0045017 | glycerolipid biosynthetic process | 8.53E-04 | 2.234920635 | 0.007014331 |
| GO:0016482 | cytosolic transport | 8.71E-04 | 2.285714286 | 0.007100193 |
| GO:0090505 | epiboly involved in wound healing | 8.72E-04 | 4.330827068 | 0.007100193 |
| GO:0030866 | cortical actin cytoskeleton organization | 8.72E-04 | 4.330827068 | 0.007100193 |
| GO:0044319 | wound healing, spreading of cells | 8.72E-04 | 4.330827068 | 0.007100193 |
| GO:0071248 | cellular response to metal ion | 8.74E-04 | 2.096985583 | 0.007100193 |
| GO:0097696 | STAT cascade | 8.74E-04 | 2.096985583 | 0.007100193 |
| GO:0044724 | single-organism carbohydrate catabolic process | 8.76E-04 | 2.412698413 | 0.007111427 |
| GO:0032763 | regulation of mast cell cytokine production | 8.95E-04 | 10.15873016 | 0.007205059 |
| GO:1904415 | regulation of xenophagy | 8.95E-04 | 10.15873016 | 0.007205059 |
| GO:1904417 | positive regulation of xenophagy | 8.95E-04 | 10.15873016 | 0.007205059 |
| GO:0002291 | T cell activation via T cell receptor contact with antigen bound to MHC molecule on antigen presenting cell | 8.95E-04 | 10.15873016 | 0.007205059 |
| GO:1905153 | regulation of membrane invagination | 9.18E-04 | 5.818181818 | 0.007355569 |
| GO:0045624 | positive regulation of T-helper cell differentiation | 9.18E-04 | 5.818181818 | 0.007355569 |
| GO:0036344 | platelet morphogenesis | 9.18E-04 | 5.818181818 | 0.007355569 |
| GO:0090257 | regulation of muscle system process | 9.34E-04 | 1.956773853 | 0.007469157 |
| GO:0019362 | pyridine nucleotide metabolic process | 9.38E-04 | 2.272189349 | 0.00748869 |
| GO:0002009 | morphogenesis of an epithelium | 9.67E-04 | 1.591367582 | 0.007701038 |
| GO:0043623 | cellular protein complex assembly | 9.71E-04 | 1.552813219 | 0.007718951 |
| GO:0007265 | Ras protein signal transduction | 0.00104103 | 1.807909605 | 0.008264805 |
| GO:0010939 | regulation of necrotic cell death | 0.001044987 | 4.21978022 | 0.008281383 |
| GO:0016311 | dephosphorylation | 0.001051469 | 1.788819876 | 0.008317874 |
| GO:1901216 | positive regulation of neuron death | 0.001108592 | 2.438095238 | 0.008754119 |
| GO:0030203 | glycosaminoglycan metabolic process | 0.001133773 | 2.612244898 | 0.008937035 |
| GO:0051155 | positive regulation of striated muscle cell differentiation | 0.001148093 | 3.009041591 | 0.009033837 |
| GO:2001237 | negative regulation of extrinsic apoptotic signaling pathway | 0.001167701 | 2.715700141 | 0.009171835 |
| GO:0002507 | tolerance induction | 0.001178839 | 4.718894009 | 0.009242929 |
| GO:1900016 | negative regulation of cytokine production involved in inflammatory response | 0.001185378 | 5.565217391 | 0.009261422 |
| GO:0061760 | antifungal innate immune response | 0.001185378 | 5.565217391 | 0.009261422 |
| GO:0051346 | negative regulation of hydrolase activity | 0.001212848 | 1.741496599 | 0.009448747 |
| GO:0036294 | cellular response to decreased oxygen levels | 0.00121362 | 2.174517375 | 0.009448747 |
| GO:0051348 | negative regulation of transferase activity | 0.001236332 | 1.921325052 | 0.009588487 |
| GO:0071675 | regulation of mononuclear cell migration | 0.001238062 | 3.409200969 | 0.009588487 |
| GO:0043647 | inositol phosphate metabolic process | 0.001238062 | 3.409200969 | 0.009588487 |
| GO:0006937 | regulation of muscle contraction | 0.001346516 | 2.206896552 | 0.010410237 |
| GO:0048732 | gland development | 0.001382573 | 1.6 | 0.010670383 |
| GO:0051282 | regulation of sequestering of calcium ion | 0.001401536 | 2.467120181 | 0.010797922 |
| GO:0098739 | import across plasma membrane | 0.001408368 | 2.316190476 | 0.010831717 |
| GO:0032667 | regulation of interleukin-23 production | 0.001426574 | 9.142857143 | 0.01091489 |
| GO:0032762 | mast cell cytokine production | 0.001426574 | 9.142857143 | 0.01091489 |
| GO:0002604 | regulation of dendritic cell antigen processing and presentation | 0.001426574 | 9.142857143 | 0.01091489 |
| GO:0051960 | regulation of nervous system development | 0.001432742 | 1.393044784 | 0.010921807 |
| GO:0001885 | endothelial cell development | 0.001434874 | 2.934744268 | 0.010921807 |
| GO:0034333 | adherens junction assembly | 0.001434874 | 2.934744268 | 0.010921807 |
| GO:0002360 | T cell lineage commitment | 0.001438662 | 4.571428571 | 0.010931858 |
| GO:0002312 | B cell activation involved in immune response | 0.001444953 | 2.782608696 | 0.010960856 |
| GO:1904149 | regulation of microglial cell mediated cytotoxicity | 0.001496436 | 14.62857143 | 0.011331987 |
| GO:0031579 | membrane raft organization | 0.001508204 | 5.333333333 | 0.011401611 |
| GO:0046474 | glycerophospholipid biosynthetic process | 0.001532967 | 2.367934224 | 0.011569067 |
| GO:0042982 | amyloid precursor protein metabolic process | 0.00154409 | 3.09054326 | 0.011633196 |
| GO:1902882 | regulation of response to oxidative stress | 0.001555955 | 2.637362637 | 0.011702683 |
| GO:0001894 | tissue homeostasis | 0.001568797 | 1.917602996 | 0.011779273 |
| GO:0046626 | regulation of insulin receptor signaling pathway | 0.001599024 | 2.898954704 | 0.011968369 |
| GO:0010043 | response to zinc ion | 0.001601144 | 3.585434174 | 0.011968369 |
| GO:0001501 | skeletal system development | 0.001602086 | 1.581809194 | 0.011968369 |
| GO:0071398 | cellular response to fatty acid | 0.001610254 | 3.297423888 | 0.012009137 |
| GO:0022612 | gland morphogenesis | 0.001637657 | 2.285714286 | 0.012192976 |
| GO:0045935 | positive regulation of nucleobase-containing compound metabolic process | 0.001706938 | 1.27554007 | 0.012687473 |
| GO:1905155 | positive regulation of membrane invagination | 0.001718374 | 6.453781513 | 0.012729758 |
| GO:0060100 | positive regulation of phagocytosis, engulfment | 0.001718374 | 6.453781513 | 0.012729758 |
| GO:0045601 | regulation of endothelial cell differentiation | 0.001734219 | 3.918367347 | 0.012805382 |
| GO:1903307 | positive regulation of regulated secretory pathway | 0.001734363 | 3.047619048 | 0.012805382 |
| GO:0050858 | negative regulation of antigen receptor-mediated signaling pathway | 0.001741061 | 4.432900433 | 0.012814923 |
| GO:0051222 | positive regulation of protein transport | 0.001741441 | 1.771265771 | 0.012814923 |
| GO:0001912 | positive regulation of leukocyte mediated cytotoxicity | 0.001763288 | 2.723404255 | 0.012954174 |
| GO:0032388 | positive regulation of intracellular transport | 0.001783983 | 1.928571429 | 0.013084509 |
| GO:0060760 | positive regulation of response to cytokine stimulus | 0.001827713 | 3.244239631 | 0.013383088 |
| GO:1905037 | autophagosome organization | 0.001870302 | 2.587601078 | 0.013649819 |
| GO:1903036 | positive regulation of response to wounding | 0.001870302 | 2.587601078 | 0.013649819 |
| GO:0097396 | response to interleukin-17 | 0.001893805 | 5.12 | 0.013775951 |
| GO:1901739 | regulation of myoblast fusion | 0.001893805 | 5.12 | 0.013775951 |
| GO:1900407 | regulation of cellular response to oxidative stress | 0.001942961 | 2.694736842 | 0.014089972 |
| GO:0032481 | positive regulation of type I interferon production | 0.001943335 | 3.005870841 | 0.014089972 |
| GO:0010594 | regulation of endothelial cell migration | 0.002028097 | 2.133333333 | 0.014648338 |
| GO:2000249 | regulation of actin cytoskeleton reorganization | 0.002030266 | 3.827242525 | 0.014648338 |
| GO:0046839 | phospholipid dephosphorylation | 0.002030266 | 3.827242525 | 0.014648338 |
| GO:0008217 | regulation of blood pressure | 0.002048261 | 1.97044335 | 0.014730187 |
| GO:0045732 | positive regulation of protein catabolic process | 0.002048261 | 1.97044335 | 0.014730187 |
| GO:0007611 | learning or memory | 0.002070906 | 1.769585253 | 0.014868903 |
| GO:0001952 | regulation of cell-matrix adhesion | 0.002108674 | 2.372955289 | 0.015115573 |
| GO:0007520 | myoblast fusion | 0.002115567 | 3.450134771 | 0.015140486 |
| GO:0002437 | inflammatory response to antigenic stimulus | 0.002186533 | 2.796638655 | 0.01562313 |
| GO:0032386 | regulation of intracellular transport | 0.002190901 | 1.582741384 | 0.01562913 |
| GO:0002820 | negative regulation of adaptive immune response | 0.00233372 | 3.142857143 | 0.016547967 |
| GO:0060348 | bone development | 0.002335422 | 1.891625616 | 0.016547967 |
| GO:0033043 | regulation of organelle organization | 0.002337493 | 1.321115077 | 0.016547967 |
| GO:0036336 | dendritic cell migration | 0.002349587 | 4.923076923 | 0.016547967 |
| GO:0050765 | negative regulation of phagocytosis | 0.002349587 | 4.923076923 | 0.016547967 |
| GO:0050860 | negative regulation of T cell receptor signaling pathway | 0.002349587 | 4.923076923 | 0.016547967 |
| GO:1903306 | negative regulation of regulated secretory pathway | 0.002349587 | 4.923076923 | 0.016547967 |
| GO:0050857 | positive regulation of antigen receptor-mediated signaling pathway | 0.002349587 | 4.923076923 | 0.016547967 |
| GO:0030194 | positive regulation of blood coagulation | 0.002364663 | 3.74025974 | 0.016601355 |
| GO:1900048 | positive regulation of hemostasis | 0.002364663 | 3.74025974 | 0.016601355 |
| GO:0030336 | negative regulation of cell migration | 0.002374128 | 1.793851718 | 0.016615147 |

| | | | | |
|---|---|---|---|---|
| GO:0051402 | neuron apoptotic process | 0.002374128 | 1.793851718 | 0.016615147 |
| GO:0009101 | glycoprotein biosynthetic process | 0.002400211 | 1.754152824 | 0.016771194 |
| GO:1904705 | regulation of vascular smooth muscle cell proliferation | 0.002422748 | 2.925714286 | 0.016902005 |
| GO:1903319 | positive regulation of protein maturation | 0.002491894 | 4.179591837 | 0.017357057 |
| GO:2000146 | negative regulation of cell motility | 0.002512742 | 1.767803194 | 0.017474801 |
| GO:0045596 | negative regulation of cell differentiation | 0.002528936 | 1.418719212 | 0.017559857 |
| GO:0071363 | cellular response to growth factor stimulus | 0.002536431 | 1.476639108 | 0.017584335 |
| GO:0046850 | regulation of bone remodeling | 0.002625817 | 3.094505495 | 0.018175574 |
| GO:0010001 | glial cell differentiation | 0.002643927 | 1.848375451 | 0.018272385 |
| GO:0032729 | positive regulation of interferon-gamma production | 0.002668247 | 2.732348112 | 0.018411738 |
| GO:0010605 | negative regulation of macromolecule metabolic process | 0.002853656 | 1.205584242 | 0.019660492 |
| GO:0032845 | negative regulation of homeostatic process | 0.002864327 | 1.920768307 | 0.019703367 |
| GO:1901223 | negative regulation of NIK/NF-kappaB signaling | 0.002883193 | 4.740740741 | 0.019802398 |
| GO:1901550 | regulation of endothelial cell development | 0.002947813 | 5.77443609 | 0.020152485 |
| GO:0045779 | negative regulation of bone resorption | 0.002947813 | 5.77443609 | 0.020152485 |
| GO:1901741 | positive regulation of myoblast fusion | 0.002947813 | 5.77443609 | 0.020152485 |
| GO:0006733 | oxidoreduction coenzyme metabolic process | 0.002980991 | 2.064516129 | 0.020347906 |
| GO:0045625 | regulation of T-helper 1 cell differentiation | 0.003079597 | 7.619047619 | 0.020988639 |
| GO:0005996 | monosaccharide metabolic process | 0.003135691 | 1.781076067 | 0.02133811 |
| GO:0097300 | programmed necrotic cell death | 0.003161834 | 3.577639752 | 0.021483013 |
| GO:0044088 | regulation of vacuole organization | 0.003297545 | 3.002132196 | 0.02237079 |
| GO:1990874 | vascular smooth muscle cell proliferation | 0.003317012 | 2.813186813 | 0.022468443 |
| GO:0007259 | JAK-STAT cascade | 0.003357167 | 2.001421464 | 0.022705722 |
| GO:0044242 | cellular lipid catabolic process | 0.00336608 | 1.896858328 | 0.0227313 |
| GO:0051962 | positive regulation of nervous system development | 0.003420017 | 1.477633478 | 0.023060391 |
| GO:0033005 | positive regulation of mast cell activation | 0.003469664 | 3.953667954 | 0.023288803 |
| GO:0090022 | regulation of neutrophil chemotaxis | 0.003469664 | 3.953667954 | 0.023288803 |
| GO:1902837 | amino acid import into cell | 0.003469664 | 3.953667954 | 0.023288803 |
| GO:0034104 | negative regulation of tissue remodeling | 0.003502449 | 4.571428571 | 0.023437837 |
| GO:1900017 | positive regulation of cytokine production involved in inflammatory response | 0.003502449 | 4.571428571 | 0.023437837 |
| GO:0019318 | hexose metabolic process | 0.003519807 | 1.835369092 | 0.023518471 |
| GO:0051592 | response to calcium ion | 0.003538239 | 2.131463628 | 0.02360602 |
| GO:0045637 | regulation of myeloid cell differentiation | 0.003548959 | 1.889020071 | 0.023641941 |
| GO:0051149 | positive regulation of muscle cell differentiation | 0.003578121 | 2.252587992 | 0.023800414 |
| GO:0050769 | positive regulation of neurogenesis | 0.00363553 | 1.509015257 | 0.024135724 |
| GO:0031345 | negative regulation of cell projection organization | 0.003639427 | 1.916406737 | 0.024135724 |
| GO:0036473 | cell death in response to oxidative stress | 0.003668694 | 2.509803922 | 0.024293444 |
| GO:1904406 | negative regulation of nitric oxide metabolic process | 0.003755764 | 5.485714286 | 0.024722191 |
| GO:0010572 | positive regulation of platelet activation | 0.003755764 | 5.485714286 | 0.024722191 |
| GO:0045623 | negative regulation of T-helper cell differentiation | 0.003755764 | 5.485714286 | 0.024722191 |
| GO:0045019 | negative regulation of nitric oxide biosynthetic process | 0.003755764 | 5.485714286 | 0.024722191 |
| GO:0050714 | positive regulation of protein secretion | 0.003808878 | 2.066182405 | 0.025034613 |
| GO:0090303 | positive regulation of wound healing | 0.00389025 | 2.612244898 | 0.025531569 |
| GO:0031669 | cellular response to nutrient levels | 0.003940443 | 1.8735363 | 0.025822726 |
| GO:0060443 | mammary gland morphogenesis | 0.003985944 | 3.15270936 | 0.026082323 |
| GO:1904951 | positive regulation of establishment of protein localization | 0.004001628 | 1.68030888 | 0.026146331 |
| GO:0098586 | cellular response to virus | 0.004047762 | 2.742857143 | 0.026369981 |
| GO:1903201 | regulation of oxidative stress-induced cell death | 0.004047762 | 2.742857143 | 0.026369981 |
| GO:0071559 | response to transforming growth factor beta | 0.004086762 | 1.81512605 | 0.026584957 |
| GO:0090023 | positive regulation of neutrophil chemotaxis | 0.004215312 | 4.413793103 | 0.027380987 |
| GO:0043301 | negative regulation of leukocyte degranulation | 0.004257556 | 7.032967033 | 0.027614893 |
| GO:0006886 | intracellular protein transport | 0.004271028 | 1.370983447 | 0.027661773 |
| GO:0002823 | negative regulation of adaptive immune response based on somatic recombination of immune receptors built from immunoglobulin superfamily domains | 0.004480563 | 3.099273608 | 0.028850136 |
| GO:0042987 | amyloid precursor protein catabolic process | 0.004480563 | 3.099273608 | 0.028850136 |
| GO:0043536 | positive regulation of blood vessel endothelial cell migration | 0.004480563 | 3.099273608 | 0.028850136 |
| GO:0071622 | regulation of granulocyte chemotaxis | 0.004480563 | 3.099273608 | 0.028850136 |
| GO:0048468 | cell development | 0.004514863 | 1.206843583 | 0.029028797 |
| GO:0035313 | wound healing, spreading of epidermal cells | 0.00471073 | 5.224489796 | 0.030053834 |
| GO:0046851 | negative regulation of bone remodeling | 0.00471073 | 5.224489796 | 0.030053834 |
| GO:1902170 | cellular response to reactive nitrogen species | 0.00471073 | 5.224489796 | 0.030053834 |
| GO:0002281 | macrophage activation involved in immune response | 0.00471073 | 5.224489796 | 0.030053834 |
| GO:0014910 | regulation of smooth muscle cell migration | 0.004714742 | 2.438095238 | 0.030053834 |
| GO:0045920 | negative regulation of exocytosis | 0.004714992 | 3.750915751 | 0.030053834 |
| GO:0010975 | regulation of neuron projection development | 0.004784473 | 1.481085892 | 0.030452893 |
| GO:0043315 | positive regulation of neutrophil degranulation | 0.004821098 | 10.44897959 | 0.030598088 |
| GO:1903980 | positive regulation of microglial cell activation | 0.004821098 | 10.44897959 | 0.030598088 |
| GO:2001238 | positive regulation of extrinsic apoptotic signaling pathway | 0.005021679 | 3.047619048 | 0.031786201 |
| GO:0001919 | regulation of receptor recycling | 0.005029826 | 4.266666667 | 0.031786201 |
| GO:0032691 | negative regulation of interleukin-1 beta production | 0.005029826 | 4.266666667 | 0.031786201 |
| GO:0046888 | negative regulation of hormone secretion | 0.005112352 | 2.41509434 | 0.032261708 |
| GO:0071407 | cellular response to organic cyclic compound | 0.0051259 | 1.457556936 | 0.03230119 |
| GO:0034284 | response to monosaccharide | 0.005147085 | 1.807713199 | 0.032356548 |
| GO:0014074 | response to purine-containing compound | 0.005156597 | 2.009419152 | 0.032356548 |
| GO:0009267 | cellular response to starvation | 0.005156597 | 2.009419152 | 0.032356548 |
| GO:0045861 | negative regulation of proteolysis | 0.005260897 | 1.649109511 | 0.032964321 |
| GO:0010770 | positive regulation of cell morphogenesis involved in differentiation | 0.005389385 | 1.959183673 | 0.033721717 |
| GO:0019722 | calcium-mediated signaling | 0.005573782 | 1.915646259 | 0.034776633 |
| GO:0042307 | positive regulation of protein import into nucleus | 0.005581532 | 2.793650794 | 0.034776633 |
| GO:0015914 | phospholipid transport | 0.005581532 | 2.793650794 | 0.034776633 |
| GO:0034332 | adherens junction organization | 0.005670282 | 2.216450216 | 0.035279983 |
| GO:0033004 | negative regulation of mast cell activation | 0.005705499 | 6.530612245 | 0.035449313 |
| GO:0044257 | cellular protein catabolic process | 0.005796855 | 1.401539778 | 0.035966482 |
| GO:0099515 | actin filament-based transport | 0.00582618 | 4.987012987 | 0.036047456 |
| GO:0006896 | Golgi to vacuole transport | 0.00582618 | 4.987012987 | 0.036047456 |
| GO:0032647 | regulation of interferon-alpha production | 0.005954062 | 4.129032258 | 0.036583213 |
| GO:1905523 | positive regulation of macrophage migration | 0.005954062 | 4.129032258 | 0.036583213 |
| GO:0034123 | positive regulation of toll-like receptor signaling pathway | 0.005954062 | 4.129032258 | 0.036583213 |
| GO:0090330 | regulation of platelet aggregation | 0.005954062 | 4.129032258 | 0.036583213 |
| GO:0071624 | positive regulation of granulocyte chemotaxis | 0.005954062 | 4.129032258 | 0.036583213 |
| GO:0030279 | negative regulation of ossification | 0.005988082 | 2.37037037 | 0.036741278 |
| GO:0001889 | liver development | 0.006024916 | 1.939393939 | 0.036916154 |
| GO:0048259 | regulation of receptor-mediated endocytosis | 0.00607777 | 2.199785177 | 0.037188564 |
| GO:0001960 | negative regulation of cytokine-mediated signaling pathway | 0.006159023 | 2.755381605 | 0.037608057 |
| GO:0051650 | establishment of vesicle localization | 0.006163307 | 1.662337662 | 0.037608057 |
| GO:0050792 | regulation of viral process | 0.00617638 | 1.716134998 | 0.03763599 |
| GO:0032411 | positive regulation of transporter activity | 0.006195594 | 2.129158513 | 0.037659457 |
| GO:0010638 | positive regulation of organelle organization | 0.006198993 | 1.431711146 | 0.037659457 |
| GO:0045444 | fat cell differentiation | 0.006205734 | 1.806888763 | 0.037659457 |
| GO:0051931 | regulation of sensory perception | 0.006254753 | 2.949308756 | 0.037801575 |
| GO:0071715 | icosanoid transport | 0.006254753 | 2.949308756 | 0.037801575 |
| GO:1901571 | fatty acid derivative transport | 0.006254753 | 2.949308756 | 0.037801575 |
| GO:0032743 | positive regulation of interleukin-2 production | 0.006270103 | 3.567944251 | 0.037842719 |
| GO:0090276 | regulation of peptide hormone secretion | 0.006758936 | 1.792717087 | 0.040737532 |
| GO:0001558 | regulation of cell growth | 0.006804686 | 1.549636804 | 0.040957554 |
| GO:0097352 | autophagosome maturation | 0.006837977 | 3.164835165 | 0.041046391 |
| GO:0070527 | platelet aggregation | 0.006837977 | 3.164835165 | 0.041046391 |

| | | | | |
|---|---|---|---|---|
| GO:0042176 | regulation of protein catabolic process | 0.006872041 | 1.593912141 | 0.041195051 |
| GO:0019932 | second-messenger-mediated signaling | 0.006913192 | 1.682734443 | 0.041385733 |
| GO:0044060 | regulation of endocrine process | 0.006952497 | 2.902494331 | 0.041517965 |
| GO:0043331 | response to dsRNA | 0.006963397 | 2.167195767 | 0.041517965 |
| GO:1903729 | regulation of plasma membrane organization | 0.006963397 | 2.167195767 | 0.041517965 |
| GO:0032958 | inositol phosphate biosynthetic process | 0.006996078 | 4 | 0.041656754 |
| GO:0034109 | homotypic cell-cell adhesion | 0.00703254 | 2.551495017 | 0.041817652 |
| GO:0002761 | regulation of myeloid leukocyte differentiation | 0.007047344 | 2.1003861 | 0.041849511 |
| GO:0007044 | cell-substrate junction assembly | 0.007079035 | 2.425655977 | 0.041925301 |
| GO:0000045 | autophagosome assembly | 0.007079035 | 2.425655977 | 0.041925301 |
| GO:0061008 | hepaticobiliary system development | 0.007092229 | 1.910447761 | 0.04194736 |
| GO:0050650 | chondroitin sulfate proteoglycan biosynthetic process | 0.007115292 | 4.770186335 | 0.041971695 |
| GO:0051770 | positive regulation of nitric-oxide synthase biosynthetic process | 0.007115292 | 4.770186335 | 0.041971695 |
| GO:0034110 | regulation of homotypic cell-cell adhesion | 0.00717728 | 3.482993197 | 0.042224898 |
| GO:0032689 | negative regulation of interferon-gamma production | 0.00717728 | 3.482993197 | 0.042224898 |
| GO:0042100 | B cell proliferation | 0.007275588 | 2.229965157 | 0.042746494 |
| GO:0051153 | regulation of striated muscle cell differentiation | 0.0074434 | 2.151260504 | 0.043643482 |
| GO:0032308 | positive regulation of prostaglandin secretion | 0.007447962 | 6.095238095 | 0.043643482 |
| GO:0043525 | positive regulation of neuron apoptotic process | 0.007664214 | 2.522167488 | 0.044792172 |
| GO:0019229 | regulation of vasoconstriction | 0.007664214 | 2.522167488 | 0.044792172 |
| GO:0061756 | leukocyte adhesion to vascular endothelial cell | 0.007677516 | 3.105121294 | 0.044810799 |
| GO:0009895 | negative regulation of catabolic process | 0.007797511 | 1.726273726 | 0.045451284 |
| GO:0048015 | phosphatidylinositol-mediated signaling | 0.007947728 | 2.019281332 | 0.046266015 |
| GO:0014909 | smooth muscle cell migration | 0.008096407 | 2.285714286 | 0.047007973 |
| GO:0045921 | positive regulation of exocytosis | 0.008096407 | 2.285714286 | 0.047007973 |
| GO:0002828 | regulation of type 2 immune response | 0.008163859 | 3.878787879 | 0.047173084 |
| GO:0001881 | receptor recycling | 0.008163859 | 3.878787879 | 0.047173084 |
| GO:0048873 | homeostasis of number of cells within a tissue | 0.008178088 | 3.401993355 | 0.047173084 |
| GO:0098751 | bone cell development | 0.008178088 | 3.401993355 | 0.047173084 |
| GO:1903427 | negative regulation of reactive oxygen species biosynthetic process | 0.008178088 | 3.401993355 | 0.047173084 |
| GO:0051806 | entry into cell of other organism involved in symbiotic interaction | 0.008291743 | 2.377142857 | 0.047704444 |
| GO:0044409 | entry into host | 0.008291743 | 2.377142857 | 0.047704444 |
| GO:0050864 | regulation of B cell activation | 0.008514817 | 1.657315494 | 0.048924306 |
| GO:0007584 | response to nutrient | 0.008558981 | 1.91473448 | 0.049092829 |
| GO:1903522 | regulation of blood circulation | 0.008566646 | 1.735140772 | 0.049092829 |
| GO:0043550 | regulation of lipid kinase activity | 0.008592221 | 3.047619048 | 0.049092829 |
| GO:1903362 | regulation of cellular protein catabolic process | 0.008609266 | 1.783972125 | 0.049092829 |
| GO:0030208 | dermatan sulfate biosynthetic process | 0.00862172 | 18.28571429 | 0.049092829 |
| GO:0032764 | negative regulation of mast cell cytokine production | 0.00862172 | 18.28571429 | 0.049092829 |
| GO:0002605 | negative regulation of dendritic cell antigen processing and presentation | 0.00862172 | 18.28571429 | 0.049092829 |
| GO:0031346 | positive regulation of cell projection organization | 0.008724048 | 1.517640254 | 0.049611725 |
| GO:0051224 | negative regulation of protein transport | 0.008772248 | 1.951845907 | 0.049821869 |
| GO:0006605 | protein targeting | 0.008853698 | 1.505882353 | 0.050220081 |
| GO:0060761 | negative regulation of response to cytokine stimulus | 0.008946374 | 2.612244898 | 0.050541533 |
| GO:0008625 | extrinsic apoptotic signaling pathway via death domain receptors | 0.008946374 | 2.612244898 | 0.050541533 |
| GO:2000378 | negative regulation of reactive oxygen species metabolic process | 0.008946374 | 2.612244898 | 0.050541533 |
| GO:0051828 | entry into other organism involved in symbiotic interaction | 0.008956005 | 2.353606789 | 0.050541533 |
| GO:0032414 | positive regulation of ion transmembrane transporter activity | 0.009043651 | 2.104830421 | 0.050971214 |
| GO:0043491 | protein kinase B signaling | 0.009209661 | 1.86407767 | 0.051775124 |
| GO:0030308 | negative regulation of cell growth | 0.009209661 | 1.86407767 | 0.051775124 |
| GO:2000785 | regulation of autophagosome assembly | 0.009277925 | 3.324675325 | 0.052092784 |
| GO:0007188 | adenylate cyclase-modulating G-protein coupled receptor signaling pathway | 0.009325298 | 1.828571429 | 0.052292494 |
| GO:0002011 | morphogenesis of an epithelial sheet | 0.009405967 | 2.770562771 | 0.052647239 |
| GO:0048017 | inositol lipid-mediated signaling | 0.009464884 | 1.982788296 | 0.052647239 |
| GO:0001779 | natural killer cell differentiation | 0.009465279 | 3.764705882 | 0.052647239 |
| GO:1990776 | response to angiotensin | 0.009465279 | 3.764705882 | 0.052647239 |
| GO:0032607 | interferon-alpha production | 0.009465279 | 3.764705882 | 0.052647239 |
| GO:1904706 | negative regulation of vascular smooth muscle cell proliferation | 0.009465279 | 3.764705882 | 0.052647239 |
| GO:0035587 | purinergic receptor signaling pathway | 0.009507402 | 5.714285714 | 0.052647239 |
| GO:0048102 | autophagic cell death | 0.009507402 | 5.714285714 | 0.052647239 |
| GO:0032957 | inositol trisphosphate metabolic process | 0.009507402 | 5.714285714 | 0.052647239 |
| GO:0002866 | positive regulation of acute inflammatory response to antigenic stimulus | 0.009507402 | 5.714285714 | 0.052647239 |
| GO:0097576 | vacuole fusion | 0.009586017 | 2.992207792 | 0.053016299 |
| GO:1904591 | positive regulation of protein import | 0.009774221 | 2.578754579 | 0.053989774 |
| GO:0010769 | regulation of cell morphogenesis involved in differentiation | 0.009842489 | 1.607535322 | 0.05429916 |
| GO:0043523 | regulation of neuron apoptotic process | 0.010024828 | 1.690802348 | 0.055236305 |
| GO:0032409 | regulation of transporter activity | 0.010140649 | 1.651980418 | 0.055796803 |
| GO:0031330 | negative regulation of cellular catabolic process | 0.010187741 | 1.846153846 | 0.055796803 |
| GO:0006986 | response to unfolded protein | 0.010215007 | 2.142857143 | 0.055796803 |
| GO:0006639 | acylglycerol metabolic process | 0.010253833 | 2.074974671 | 0.055796803 |
| GO:0035855 | megakaryocyte development | 0.0102651 | 4.388571429 | 0.055796803 |
| GO:0033008 | positive regulation of mast cell activation involved in immune response | 0.0102651 | 4.388571429 | 0.055796803 |
| GO:0033622 | integrin activation | 0.0102651 | 4.388571429 | 0.055796803 |
| GO:0002864 | regulation of acute inflammatory response to antigenic stimulus | 0.0102651 | 4.388571429 | 0.055796803 |
| GO:0043306 | positive regulation of mast cell degranulation | 0.0102651 | 4.388571429 | 0.055796803 |
| GO:0043011 | myeloid dendritic cell differentiation | 0.0102651 | 4.388571429 | 0.055796803 |
| GO:0010884 | positive regulation of lipid storage | 0.0102651 | 4.388571429 | 0.055796803 |
| GO:0007219 | Notch signaling pathway | 0.01033777 | 1.91949487 | 0.056122941 |
| GO:2000273 | positive regulation of receptor activity | 0.010353826 | 2.729211087 | 0.056141308 |
| GO:0070741 | response to interleukin-6 | 0.010482148 | 3.250793651 | 0.05656019 |
| GO:1902930 | regulation of alcohol biosynthetic process | 0.010482148 | 3.250793651 | 0.05656019 |
| GO:0048246 | macrophage chemotaxis | 0.010482148 | 3.250793651 | 0.05656019 |
| GO:0006984 | ER-nucleus signaling pathway | 0.010482148 | 3.250793651 | 0.05656019 |
| GO:0010827 | regulation of glucose transport | 0.010642735 | 2.411302983 | 0.057048686 |
| GO:0038089 | positive regulation of cell migration by vascular endothelial growth factor signaling pathway | 0.010655897 | 8.126984127 | 0.057048686 |
| GO:0014004 | microglia differentiation | 0.010655897 | 8.126984127 | 0.057048686 |
| GO:0048260 | positive regulation of receptor-mediated endocytosis | 0.0106592 | 2.546112116 | 0.057048686 |
| GO:0060147 | regulation of posttranscriptional gene silencing | 0.010662824 | 2.93877551 | 0.057048686 |
| GO:0042149 | cellular response to glucose starvation | 0.010662824 | 2.93877551 | 0.057048686 |
| GO:0001961 | positive regulation of cytokine-mediated signaling pathway | 0.010662824 | 2.93877551 | 0.057048686 |
| GO:0050796 | regulation of insulin secretion | 0.01070711 | 1.837320574 | 0.057147589 |
| GO:0002793 | positive regulation of peptide secretion | 0.01070711 | 1.837320574 | 0.057147589 |
| GO:0048639 | positive regulation of developmental growth | 0.010781949 | 1.803971813 | 0.05747778 |
| GO:0003158 | endothelium development | 0.010904928 | 2.060362173 | 0.057802219 |
| GO:1904950 | negative regulation of establishment of protein localization | 0.010906435 | 1.908948195 | 0.057802219 |
| GO:0002693 | positive regulation of cellular extravasation | 0.010908049 | 3.657142857 | 0.057802219 |
| GO:0060142 | regulation of syncytium formation by plasma membrane fusion | 0.010908049 | 3.657142857 | 0.057802219 |
| GO:0031294 | lymphocyte costimulation | 0.010908049 | 3.657142857 | 0.057802219 |
| GO:0022008 | neurogenesis | 0.011028524 | 1.221534203 | 0.058370803 |
| GO:1901888 | regulation of cell junction assembly | 0.011199469 | 2.285714286 | 0.059204829 |
| GO:0000904 | cell morphogenesis involved in differentiation | 0.011277305 | 1.361702128 | 0.059545246 |
| GO:0042594 | response to starvation | 0.011306174 | 1.795918367 | 0.059555706 |
| GO:0008361 | regulation of cell size | 0.011306174 | 1.795918367 | 0.059555706 |
| GO:0048525 | negative regulation of viral process | 0.011439108 | 1.992673993 | 0.06018438 |
| GO:0045824 | negative regulation of innate immune response | 0.011509928 | 2.385093168 | 0.060485148 |
| GO:0006638 | neutral lipid metabolic process | 0.011588048 | 2.045954046 | 0.060823521 |

| | | | | |
|---|---|---|---|---|
| GO:0072659 | protein localization to plasma membrane | 0.011610894 | 1.691916624 | 0.06086234 |
| GO:0051897 | positive regulation of protein kinase B signaling | 0.011622921 | 2.10989011 | 0.06086234 |
| GO:1904645 | response to beta-amyloid | 0.011796051 | 3.180124224 | 0.061620975 |
| GO:0002691 | regulation of cellular extravasation | 0.011796051 | 3.180124224 | 0.061620975 |
| GO:0051955 | regulation of amino acid transport | 0.011826542 | 2.887218045 | 0.061620975 |
| GO:0060966 | regulation of gene silencing by RNA | 0.011826542 | 2.887218045 | 0.061620975 |
| GO:0006140 | regulation of nucleotide metabolic process | 0.011837348 | 1.936134454 | 0.061620975 |
| GO:0001845 | phagolysosome assembly | 0.011904026 | 5.378151261 | 0.061678171 |
| GO:0045064 | T-helper 2 cell differentiation | 0.011904026 | 5.378151261 | 0.061678171 |
| GO:0001921 | positive regulation of receptor recycling | 0.011904026 | 5.378151261 | 0.061678171 |
| GO:0030206 | chondroitin sulfate biosynthetic process | 0.011904026 | 5.378151261 | 0.061678171 |
| GO:0015696 | ammonium transport | 0.012036208 | 2.263945578 | 0.062290187 |
| GO:0010829 | negative regulation of glucose transport | 0.012149746 | 4.21978022 | 0.062731208 |
| GO:0035743 | CD4-positive, alpha-beta T cell cytokine production | 0.012149746 | 4.21978022 | 0.062731208 |
| GO:0048041 | focal adhesion assembly | 0.012462581 | 2.65010352 | 0.064196785 |
| GO:1903391 | regulation of adherens junction organization | 0.012462581 | 2.65010352 | 0.064196785 |
| GO:0046466 | membrane lipid catabolic process | 0.012499675 | 3.555555556 | 0.064313076 |
| GO:1901184 | regulation of ERBB signaling pathway | 0.012610076 | 2.48324515 | 0.064805842 |
| GO:0017038 | protein import | 0.012625386 | 1.679959616 | 0.064809341 |
| GO:0044265 | cellular macromolecule catabolic process | 0.012650834 | 1.308550186 | 0.06486481 |
| GO:0002792 | negative regulation of peptide secretion | 0.012920104 | 2.242587601 | 0.06609245 |
| GO:0001938 | positive regulation of endothelial cell proliferation | 0.012920104 | 2.242587601 | 0.06609245 |
| GO:0015749 | monosaccharide transport | 0.013148915 | 2.151260504 | 0.067185344 |
| GO:1903827 | regulation of cellular protein localization | 0.013167702 | 1.4271777 | 0.067203823 |
| GO:0002832 | negative regulation of response to biotic stimulus | 0.013224839 | 3.112462006 | 0.067417764 |
| GO:0010977 | negative regulation of neuron projection development | 0.013434102 | 1.867895545 | 0.068405828 |
| GO:0072594 | establishment of protein localization to organelle | 0.013594057 | 1.500771367 | 0.069140842 |
| GO:0006029 | proteoglycan metabolic process | 0.013680732 | 2.452961672 | 0.069422269 |
| GO:0002704 | negative regulation of leukocyte mediated immunity | 0.013680732 | 2.452961672 | 0.069422269 |
| GO:0031623 | receptor internalization | 0.014033944 | 2.133333333 | 0.071051852 |
| GO:0010522 | regulation of calcium ion transport into cytosol | 0.014033944 | 2.133333333 | 0.071051852 |
| GO:0098927 | vesicle-mediated transport between endosomal compartments | 0.014247419 | 3.459459459 | 0.071928371 |
| GO:0060143 | positive regulation of syncytium formation by plasma membrane fusion | 0.014255781 | 4.063492063 | 0.071928371 |
| GO:0032753 | positive regulation of interleukin-4 production | 0.014255781 | 4.063492063 | 0.071928371 |
| GO:0051489 | regulation of filopodium assembly | 0.014430132 | 2.789346247 | 0.072541519 |
| GO:0031532 | actin cytoskeleton reorganization | 0.014436698 | 2.309774436 | 0.072541519 |
| GO:0007399 | nervous system development | 0.014549821 | 1.175952291 | 0.072541519 |
| GO:0070391 | response to lipoteichoic acid | 0.014611322 | 7.314285714 | 0.072541519 |
| GO:0033029 | regulation of neutrophil apoptotic process | 0.014611322 | 7.314285714 | 0.072541519 |
| GO:0002317 | plasma cell differentiation | 0.014611322 | 7.314285714 | 0.072541519 |
| GO:1901724 | positive regulation of cell proliferation involved in kidney development | 0.014611322 | 7.314285714 | 0.072541519 |
| GO:0002315 | marginal zone B cell differentiation | 0.014611322 | 7.314285714 | 0.072541519 |
| GO:0070666 | regulation of mast cell proliferation | 0.014611322 | 7.314285714 | 0.072541519 |
| GO:0071223 | cellular response to lipoteichoic acid | 0.014611322 | 7.314285714 | 0.072541519 |
| GO:1903223 | positive regulation of oxidative stress-induced neuron death | 0.014611322 | 7.314285714 | 0.072541519 |
| GO:1903265 | positive regulation of tumor necrosis factor-mediated signaling pathway | 0.014611322 | 7.314285714 | 0.072541519 |
| GO:0010919 | regulation of inositol phosphate biosynthetic process | 0.014655679 | 5.079365079 | 0.072541519 |
| GO:0070102 | interleukin-6-mediated signaling pathway | 0.014655679 | 5.079365079 | 0.072541519 |
| GO:0045342 | MHC class II biosynthetic process | 0.014655679 | 5.079365079 | 0.072541519 |
| GO:0002730 | regulation of dendritic cell cytokine production | 0.014655679 | 5.079365079 | 0.072541519 |
| GO:2000319 | regulation of T-helper 17 cell differentiation | 0.014655679 | 5.079365079 | 0.072541519 |
| GO:0019058 | viral life cycle | 0.014830094 | 1.585466557 | 0.073306318 |
| GO:0051896 | regulation of protein kinase B signaling | 0.014859836 | 1.848024316 | 0.073306318 |
| GO:0060402 | calcium ion transport into cytosol | 0.014859836 | 1.848024316 | 0.073306318 |
| GO:0099565 | chemical synaptic transmission, postsynaptic | 0.015527163 | 2.285714286 | 0.076428145 |
| GO:0050709 | negative regulation of protein secretion | 0.015527163 | 2.285714286 | 0.076428145 |
| GO:1990778 | protein localization to cell periphery | 0.015586105 | 1.56590371 | 0.076633123 |
| GO:0045666 | positive regulation of neuron differentiation | 0.016059255 | 1.473435655 | 0.078871953 |
| GO:0045954 | positive regulation of natural killer cell mediated cytotoxicity | 0.016158258 | 3.368421053 | 0.079095121 |
| GO:0006884 | cell volume homeostasis | 0.016158258 | 3.368421053 | 0.079095121 |
| GO:0050996 | positive regulation of lipid catabolic process | 0.016158258 | 3.368421053 | 0.079095121 |
| GO:0032728 | positive regulation of interferon-beta production | 0.016447335 | 2.985422741 | 0.080332628 |
| GO:0002686 | negative regulation of leukocyte migration | 0.016447335 | 2.985422741 | 0.080332628 |
| GO:0051354 | negative regulation of oxidoreductase activity | 0.016593462 | 3.918367347 | 0.080639766 |
| GO:1904466 | positive regulation of matrix metallopeptidase secretion | 0.016619437 | 13.71428571 | 0.080639766 |
| GO:0035624 | receptor transactivation | 0.016619437 | 13.71428571 | 0.080639766 |
| GO:0070164 | negative regulation of adiponectin secretion | 0.016619437 | 13.71428571 | 0.080639766 |
| GO:1904151 | positive regulation of microglial cell mediated cytotoxicity | 0.016619437 | 13.71428571 | 0.080639766 |
| GO:2000520 | regulation of immunological synapse formation | 0.016619437 | 13.71428571 | 0.080639766 |
| GO:1903524 | positive regulation of blood circulation | 0.016678286 | 2.262150221 | 0.080748424 |
| GO:0043535 | regulation of blood vessel endothelial cell migration | 0.016678286 | 2.262150221 | 0.080748424 |
| GO:0060079 | excitatory postsynaptic potential | 0.017301917 | 2.366386555 | 0.083676301 |
| GO:1905114 | cell surface receptor signaling pathway involved in cell-cell signaling | 0.017537462 | 1.432214416 | 0.08470438 |
| GO:0072657 | protein localization to membrane | 0.017552736 | 1.406593407 | 0.08470438 |
| GO:0033500 | carbohydrate homeostasis | 0.017668839 | 1.595015576 | 0.085171879 |
| GO:0032700 | negative regulation of interleukin-17 production | 0.017777777 | 4.812030075 | 0.08541817 |
| GO:1900225 | regulation of NLRP3 inflammasome complex assembly | 0.017777777 | 4.812030075 | 0.08541817 |
| GO:0050855 | regulation of B cell receptor signaling pathway | 0.017777777 | 4.812030075 | 0.08541817 |
| GO:0034620 | cellular response to unfolded protein | 0.017891985 | 2.239067055 | 0.085873772 |
| GO:0051961 | negative regulation of nervous system development | 0.01792082 | 1.535808024 | 0.085919084 |
| GO:0007269 | neurotransmitter secretion | 0.017972211 | 1.849117175 | 0.086072318 |
| GO:0051482 | positive regulation of cytosolic calcium ion concentration involved in phospholipase C-activating G-protein coupled signaling pathway | 0.018238852 | 3.282051282 | 0.08703032 |
| GO:0002724 | regulation of T cell cytokine production | 0.018238852 | 3.282051282 | 0.08703032 |
| GO:1903727 | positive regulation of phospholipid metabolic process | 0.018250828 | 2.925714286 | 0.08703032 |
| GO:0045646 | regulation of erythrocyte differentiation | 0.018250828 | 2.925714286 | 0.08703032 |
| GO:0050768 | negative regulation of neurogenesis | 0.01838193 | 1.558441558 | 0.087561236 |
| GO:0099643 | signal release from synapse | 0.018885262 | 1.838786911 | 0.089862202 |
| GO:0031644 | regulation of neurological system process | 0.019160826 | 2.048 | 0.090740425 |
| GO:0046718 | viral entry into host cell | 0.019170157 | 2.216450216 | 0.090740425 |
| GO:0071402 | cellular response to lipoprotein particle stimulus | 0.019172243 | 3.783251232 | 0.090740425 |
| GO:0044090 | positive regulation of vacuole organization | 0.019172243 | 3.783251232 | 0.090740425 |
| GO:0071677 | positive regulation of mononuclear cell migration | 0.019172243 | 3.783251232 | 0.090740425 |
| GO:0002246 | wound healing involved in inflammatory response | 0.019286101 | 6.649350649 | 0.090978401 |
| GO:0007220 | Notch receptor processing | 0.019286101 | 6.649350649 | 0.090978401 |
| GO:0090160 | Golgi to lysosome transport | 0.019286101 | 6.649350649 | 0.090978401 |
| GO:1900182 | positive regulation of protein localization to nucleus | 0.019304672 | 2.12244898 | 0.090978401 |
| GO:0007187 | G-protein coupled receptor signaling pathway, coupled to cyclic nucleotide second messenger | 0.019917483 | 1.675583381 | 0.093766683 |
| GO:0071359 | cellular response to dsRNA | 0.020080285 | 2.311986864 | 0.094431882 |
| GO:0032846 | positive regulation of homeostatic process | 0.020101414 | 1.611622276 | 0.094431882 |
| GO:0003254 | regulation of membrane depolarization | 0.020188737 | 2.868347339 | 0.094741638 |
| GO:0030522 | intracellular receptor signaling pathway | 0.020449789 | 1.721973094 | 0.095865147 |
| GO:0002717 | positive regulation of natural killer cell mediated immunity | 0.020495507 | 3.2 | 0.095876552 |
| GO:0034105 | positive regulation of tissue remodeling | 0.020495507 | 3.2 | 0.095876552 |
| GO:1903076 | regulation of protein localization to plasma membrane | 0.020564001 | 2.103666245 | 0.096095488 |
| GO:0034767 | positive regulation of ion transmembrane transport | 0.020670018 | 1.749829118 | 0.096489123 |
| GO:0050994 | regulation of lipid catabolic process | 0.020705751 | 2.438095238 | 0.096554187 |

| | | | | |
|---|---|---|---|---|
| GO:0010812 | negative regulation of cell-substrate adhesion | 0.020846581 | 2.612244898 | 0.097108678 |
| GO:0097320 | plasma membrane tubulation | 0.021283281 | 4.571428571 | 0.098831166 |
| GO:0045986 | negative regulation of smooth muscle contraction | 0.021283281 | 4.571428571 | 0.098831166 |
| GO:0030050 | vesicle transport along actin filament | 0.021283281 | 4.571428571 | 0.098831166 |
| GO:0014812 | muscle cell migration | 0.021564677 | 2.015748031 | 0.09984101 |
| GO:0050688 | regulation of defense response to virus | 0.021585557 | 2.285714286 | 0.09984101 |
| GO:0002637 | regulation of immunoglobulin production | 0.021585557 | 2.285714286 | 0.09984101 |
| GO:0051147 | regulation of muscle cell differentiation | 0.0215909 | 1.662337662 | 0.09984101 |
| GO:0007179 | transforming growth factor beta receptor signaling pathway | 0.021778943 | 1.772594752 | 0.100605545 |
| GO:1904894 | positive regulation of STAT cascade | 0.021883341 | 2.085213033 | 0.100982502 |
| GO:0014015 | positive regulation of gliogenesis | 0.021927406 | 2.172560113 | 0.101080552 |
| GO:0010226 | response to lithium ion | 0.022000727 | 3.657142857 | 0.101313118 |
| GO:0031333 | negative regulation of protein complex assembly | 0.022660252 | 1.887557604 | 0.104241866 |
| GO:0043114 | regulation of vascular permeability | 0.022934147 | 3.12195122 | 0.105283185 |
| GO:0009311 | oligosaccharide metabolic process | 0.022934147 | 3.12195122 | 0.105283185 |
| GO:0060349 | bone morphogenesis | 0.023264243 | 2.067080745 | 0.106687988 |
| GO:0010721 | negative regulation of cell development | 0.023591139 | 1.48994709 | 0.108075229 |
| GO:0034250 | positive regulation of cellular amide metabolic process | 0.023842231 | 1.754689755 | 0.109000088 |
| GO:0006643 | membrane lipid metabolic process | 0.023842231 | 1.754689755 | 0.109000088 |
| GO:0042058 | regulation of epidermal growth factor receptor signaling pathway | 0.024168417 | 2.374768089 | 0.110263735 |
| GO:0061180 | mammary gland epithelium development | 0.024168417 | 2.374768089 | 0.110263735 |
| GO:0097400 | interleukin-17-mediated signaling pathway | 0.02468827 | 6.095238095 | 0.112300409 |
| GO:0090109 | regulation of cell-substrate junction assembly | 0.02471623 | 2.531868132 | 0.112300409 |
| GO:0051893 | regulation of focal adhesion assembly | 0.02471623 | 2.531868132 | 0.112300409 |
| GO:0042269 | regulation of natural killer cell mediated cytotoxicity | 0.02471623 | 2.531868132 | 0.112300409 |
| GO:0015748 | organophosphate ester transport | 0.024964736 | 2.13037448 | 0.113313299 |
| GO:0035767 | endothelial cell chemotaxis | 0.025086624 | 3.539170507 | 0.113633684 |
| GO:0045932 | negative regulation of muscle contraction | 0.025086624 | 3.539170507 | 0.113633684 |
| GO:0002524 | hypersensitivity | 0.025182704 | 4.353741497 | 0.11372006 |
| GO:0050665 | hydrogen peroxide biosynthetic process | 0.025182704 | 4.353741497 | 0.11372006 |
| GO:0010042 | response to manganese ion | 0.025182704 | 4.353741497 | 0.11372006 |
| GO:0051259 | protein oligomerization | 0.025554073 | 1.613445378 | 0.115190329 |
| GO:0002861 | regulation of inflammatory response to antigenic stimulus | 0.025560292 | 3.047619048 | 0.115190329 |
| GO:1904627 | response to phorbol 13-acetate 12-myristate | 0.026701438 | 10.97142857 | 0.119724058 |
| GO:0038157 | granulocyte-macrophage colony-stimulating factor signaling pathway | 0.026701438 | 10.97142857 | 0.119724058 |
| GO:0035771 | interleukin-4-mediated signaling pathway | 0.026701438 | 10.97142857 | 0.119724058 |
| GO:1904628 | cellular response to phorbol 13-acetate 12-myristate | 0.026701438 | 10.97142857 | 0.119724058 |
| GO:0033031 | positive regulation of neutrophil apoptotic process | 0.026701438 | 10.97142857 | 0.119724058 |
| GO:0010883 | regulation of lipid storage | 0.02682799 | 2.493506494 | 0.120158205 |
| GO:0046596 | regulation of viral entry into host cell | 0.026852511 | 2.708994709 | 0.120158205 |
| GO:0032355 | response to estradiol | 0.027718115 | 1.840071878 | 0.123906406 |
| GO:0006109 | regulation of carbohydrate metabolic process | 0.027931978 | 1.693121693 | 0.124610938 |
| GO:0034764 | positive regulation of transmembrane transport | 0.027931978 | 1.693121693 | 0.124610938 |
| GO:0030149 | sphingolipid catabolic process | 0.028436725 | 3.428571429 | 0.126480614 |
| GO:0034694 | response to prostaglandin | 0.028436725 | 3.428571429 | 0.126480614 |
| GO:0060148 | positive regulation of posttranscriptional gene silencing | 0.028436725 | 3.428571429 | 0.126480614 |
| GO:0046890 | regulation of lipid biosynthetic process | 0.029108965 | 1.685319289 | 0.129340738 |
| GO:0060149 | negative regulation of posttranscriptional gene silencing | 0.029484154 | 4.155844156 | 0.130484317 |
| GO:0002863 | positive regulation of inflammatory response to antigenic stimulus | 0.029484154 | 4.155844156 | 0.130484317 |
| GO:0060967 | negative regulation of gene silencing by RNA | 0.029484154 | 4.155844156 | 0.130484317 |
| GO:1903209 | positive regulation of oxidative stress-induced cell death | 0.029484154 | 4.155844156 | 0.130484317 |
| GO:0006836 | neurotransmitter transport | 0.030269346 | 1.628687102 | 0.13382555 |
| GO:0045628 | regulation of T-helper 2 cell differentiation | 0.030817484 | 5.626373626 | 0.13476945 |
| GO:0001768 | establishment of T cell polarity | 0.030817484 | 5.626373626 | 0.13476945 |
| GO:0055119 | relaxation of cardiac muscle | 0.030817484 | 5.626373626 | 0.13476945 |
| GO:0032725 | positive regulation of granulocyte macrophage colony-stimulating factor production | 0.030817484 | 5.626373626 | 0.13476945 |
| GO:0001767 | establishment of lymphocyte polarity | 0.030817484 | 5.626373626 | 0.13476945 |
| GO:0036005 | response to macrophage colony-stimulating factor | 0.030817484 | 5.626373626 | 0.13476945 |
| GO:0032650 | regulation of interleukin-1 alpha production | 0.030817484 | 5.626373626 | 0.13476945 |
| GO:0023035 | CD40 signaling pathway | 0.030817484 | 5.626373626 | 0.13476945 |
| GO:0042590 | antigen processing and presentation of exogenous peptide antigen via MHC class I | 0.030817484 | 5.626373626 | 0.13476945 |
| GO:0060368 | regulation of Fc receptor mediated stimulatory signaling pathway | 0.030817484 | 5.626373626 | 0.13476945 |
| GO:0043320 | natural killer cell degranulation | 0.030817484 | 5.626373626 | 0.13476945 |
| GO:0030282 | bone mineralization | 0.031144687 | 1.980952381 | 0.136066041 |
| GO:0048147 | negative regulation of fibroblast proliferation | 0.031394992 | 2.909090909 | 0.137000594 |
| GO:0002715 | regulation of natural killer cell mediated immunity | 0.031420452 | 2.420168067 | 0.137000594 |
| GO:0043436 | oxoacid metabolic process | 0.031457715 | 1.25819135 | 0.137028198 |
| GO:0022898 | regulation of transmembrane transporter activity | 0.031784463 | 1.543599258 | 0.13831549 |
| GO:0032814 | regulation of natural killer cell activation | 0.032043072 | 2.612244898 | 0.139167461 |
| GO:0014003 | oligodendrocyte development | 0.032043072 | 2.612244898 | 0.139167461 |
| GO:0061045 | negative regulation of wound healing | 0.03229264 | 2.257495591 | 0.140113999 |
| GO:0001936 | regulation of endothelial cell proliferation | 0.033274128 | 1.756255044 | 0.144231299 |
| GO:1903900 | regulation of viral life cycle | 0.034191393 | 1.654815772 | 0.147782155 |
| GO:0071731 | response to nitric oxide | 0.034193388 | 3.97515528 | 0.147782155 |
| GO:1901889 | negative regulation of cell junction assembly | 0.034193388 | 3.97515528 | 0.147782155 |
| GO:0046824 | positive regulation of nucleocytoplasmic transport | 0.034488017 | 2.117293233 | 0.148765253 |
| GO:1903035 | negative regulation of response to wounding | 0.034488017 | 2.117293233 | 0.148765253 |
| GO:0031326 | regulation of cellular biosynthetic process | 0.034589003 | 1.10362063 | 0.149055721 |
| GO:0034113 | heterotypic cell-cell adhesion | 0.034873386 | 2.56641604 | 0.149989419 |
| GO:0071548 | response to dexamethasone | 0.034873386 | 2.56641604 | 0.149989419 |
| GO:0050678 | regulation of epithelial cell proliferation | 0.035088257 | 1.425389755 | 0.150767196 |
| GO:1905475 | regulation of protein localization to membrane | 0.035732875 | 1.705403405 | 0.153388212 |
| GO:1903205 | regulation of hydrogen peroxide-induced cell death | 0.035951983 | 3.226890756 | 0.153584652 |
| GO:0032673 | regulation of interleukin-4 production | 0.035951983 | 3.226890756 | 0.153584652 |
| GO:0070528 | protein kinase C signaling | 0.035951983 | 3.226890756 | 0.153584652 |
| GO:0045022 | early endosome to late endosome transport | 0.035951983 | 3.226890756 | 0.153584652 |
| GO:0070269 | pyroptosis | 0.035951983 | 3.226890756 | 0.153584652 |
| GO:2001242 | regulation of intrinsic apoptotic signaling pathway | 0.036346181 | 1.736632083 | 0.154969761 |
| GO:1902904 | negative regulation of supramolecular fiber organization | 0.036346181 | 1.736632083 | 0.154969761 |
| GO:0032412 | regulation of ion transmembrane transporter activity | 0.036428199 | 1.539201539 | 0.155170115 |
| GO:0043462 | regulation of ATPase activity | 0.036523154 | 2.351020408 | 0.15542514 |
| GO:1903901 | negative regulation of viral life cycle | 0.036704988 | 1.932636469 | 0.156049037 |
| GO:0001890 | placenta development | 0.037257874 | 1.696612666 | 0.158247729 |
| GO:0050679 | positive regulation of epithelial cell proliferation | 0.03731375 | 1.613445378 | 0.158312426 |
| GO:0002467 | germinal center formation | 0.037666207 | 5.224489796 | 0.158312426 |
| GO:0042362 | fat-soluble vitamin biosynthetic process | 0.037666207 | 5.224489796 | 0.158312426 |
| GO:0072683 | T cell extravasation | 0.037666207 | 5.224489796 | 0.158312426 |
| GO:0002829 | negative regulation of type 2 immune response | 0.037666207 | 5.224489796 | 0.158312426 |
| GO:1902902 | negative regulation of autophagosome assembly | 0.037666207 | 5.224489796 | 0.158312426 |
| GO:0032610 | interleukin-1 alpha production | 0.037666207 | 5.224489796 | 0.158312426 |
| GO:0070344 | regulation of fat cell proliferation | 0.037666207 | 5.224489796 | 0.158312426 |
| GO:0010936 | negative regulation of macrophage cytokine production | 0.037666207 | 5.224489796 | 0.158312426 |
| GO:0051712 | positive regulation of killing of cells of other organism | 0.037666207 | 5.224489796 | 0.158312426 |
| GO:0090594 | inflammatory response to wounding | 0.037666207 | 5.224489796 | 0.158312426 |
| GO:0048538 | thymus development | 0.037864493 | 2.522167488 | 0.15899498 |
| GO:0030879 | mammary gland development | 0.037957171 | 1.726984127 | 0.159106848 |

| | | | | |
|---|---|---|---|---|
| GO:1901031 | regulation of response to reactive oxygen species | 0.038034797 | 2.782608696 | 0.159106848 |
| GO:0048536 | spleen development | 0.038034797 | 2.782608696 | 0.159106848 |
| GO:0035886 | vascular smooth muscle cell differentiation | 0.038034797 | 2.782608696 | 0.159106848 |
| GO:0071608 | macrophage inflammatory protein-1 alpha production | 0.038616529 | 9.142857143 | 0.159136023 |
| GO:0014005 | microglia development | 0.038616529 | 9.142857143 | 0.159136023 |
| GO:0033025 | regulation of mast cell apoptotic process | 0.038616529 | 9.142857143 | 0.159136023 |
| GO:1903969 | regulation of response to macrophage colony-stimulating factor | 0.038616529 | 9.142857143 | 0.159136023 |
| GO:2000321 | positive regulation of T-helper 17 cell differentiation | 0.038616529 | 9.142857143 | 0.159136023 |
| GO:2000427 | positive regulation of apoptotic cell clearance | 0.038616529 | 9.142857143 | 0.159136023 |
| GO:1902262 | apoptotic process involved in blood vessel morphogenesis | 0.038616529 | 9.142857143 | 0.159136023 |
| GO:0002677 | negative regulation of chronic inflammatory response | 0.038616529 | 9.142857143 | 0.159136023 |
| GO:0043545 | molybdopterin cofactor metabolic process | 0.038616529 | 9.142857143 | 0.159136023 |
| GO:0070668 | positive regulation of mast cell proliferation | 0.038616529 | 9.142857143 | 0.159136023 |
| GO:0071640 | regulation of macrophage inflammatory protein 1 alpha production | 0.038616529 | 9.142857143 | 0.159136023 |
| GO:0006777 | Mo-molybdopterin cofactor biosynthetic process | 0.038616529 | 9.142857143 | 0.159136023 |
| GO:1990773 | matrix metallopeptidase secretion | 0.038616529 | 9.142857143 | 0.159136023 |
| GO:0051189 | prosthetic group metabolic process | 0.038616529 | 9.142857143 | 0.159136023 |
| GO:0019720 | Mo-molybdopterin cofactor metabolic process | 0.038616529 | 9.142857143 | 0.159136023 |
| GO:1903972 | regulation of cellular response to macrophage colony-stimulating factor stimulus | 0.038616529 | 9.142857143 | 0.159136023 |
| GO:0006665 | sphingolipid metabolic process | 0.038862541 | 1.804511278 | 0.159852419 |
| GO:0045598 | regulation of fat cell differentiation | 0.038862541 | 1.804511278 | 0.159852419 |
| GO:0008211 | glucocorticoid metabolic process | 0.039313897 | 3.80952381 | 0.161409236 |
| GO:0046629 | gamma-delta T cell activation | 0.039313897 | 3.80952381 | 0.161409236 |
| GO:1902115 | regulation of organelle assembly | 0.039466241 | 1.654421769 | 0.161884674 |
| GO:0051494 | negative regulation of cytoskeleton organization | 0.039619165 | 1.717442778 | 0.162361611 |
| GO:0016051 | carbohydrate biosynthetic process | 0.039902319 | 1.625396825 | 0.163370864 |
| GO:0010758 | regulation of macrophage chemotaxis | 0.040125972 | 3.134693878 | 0.163681451 |
| GO:0070670 | response to interleukin-4 | 0.040125972 | 3.134693878 | 0.163681451 |
| GO:0050654 | chondroitin sulfate proteoglycan metabolic process | 0.040125972 | 3.134693878 | 0.163681451 |
| GO:0090218 | positive regulation of lipid kinase activity | 0.040125972 | 3.134693878 | 0.163681451 |
| GO:0051283 | negative regulation of sequestering of calcium ion | 0.040188279 | 1.976833977 | 0.1637848 |
| GO:1900180 | regulation of protein localization to nucleus | 0.040449049 | 1.679300292 | 0.164696036 |
| GO:1904375 | regulation of protein localization to cell periphery | 0.040780596 | 1.901714286 | 0.165893518 |
| GO:0031175 | neuron projection development | 0.040866 | 1.239989667 | 0.166088422 |
| GO:0032964 | collagen biosynthetic process | 0.041019119 | 2.479418886 | 0.166253152 |
| GO:0070918 | production of small RNA involved in gene silencing by RNA | 0.041019119 | 2.479418886 | 0.166253152 |
| GO:0045010 | actin nucleation | 0.041019119 | 2.479418886 | 0.166253152 |
| GO:0061138 | morphogenesis of a branching epithelium | 0.041432666 | 1.618204804 | 0.167775786 |
| GO:0042692 | muscle cell differentiation | 0.041524247 | 1.367156208 | 0.167993071 |
| GO:0030574 | collagen catabolic process | 0.041665523 | 2.723404255 | 0.168257308 |
| GO:0046365 | monosaccharide catabolic process | 0.041665523 | 2.723404255 | 0.168257308 |
| GO:1901215 | negative regulation of neuron death | 0.041879442 | 1.551924091 | 0.168967149 |
| GO:0010821 | regulation of mitochondrion organization | 0.042080249 | 1.741496599 | 0.169310894 |
| GO:0090316 | positive regulation of intracellular protein transport | 0.042080249 | 1.741496599 | 0.169310894 |
| GO:0006096 | glycolytic process | 0.042137335 | 2.151260504 | 0.169310894 |
| GO:0032272 | negative regulation of protein polymerization | 0.042137335 | 2.151260504 | 0.169310894 |
| GO:0060135 | maternal process involved in female pregnancy | 0.042155737 | 2.285714286 | 0.169310894 |
| GO:0007041 | lysosomal transport | 0.042932641 | 1.886621315 | 0.172275 |
| GO:0001678 | cellular glucose homeostasis | 0.043832958 | 1.662337662 | 0.17572851 |
| GO:0001570 | vasculogenesis | 0.043887899 | 2.031746032 | 0.175789685 |
| GO:0051353 | positive regulation of oxidoreductase activity | 0.044339698 | 2.438095238 | 0.177051042 |
| GO:0048679 | regulation of axon regeneration | 0.044581826 | 3.047619048 | 0.177051042 |
| GO:0036474 | cell death in response to hydrogen peroxide | 0.044581826 | 3.047619048 | 0.177051042 |
| GO:1903206 | negative regulation of hydrogen peroxide-induced cell death | 0.044847002 | 3.657142857 | 0.177051042 |
| GO:0048569 | post-embryonic animal organ development | 0.044847002 | 3.657142857 | 0.177051042 |
| GO:0055098 | response to low-density lipoprotein particle | 0.044847002 | 3.657142857 | 0.177051042 |
| GO:1990000 | amyloid fibril formation | 0.044847002 | 3.657142857 | 0.177051042 |
| GO:0002544 | chronic inflammatory response | 0.044847002 | 3.657142857 | 0.177051042 |
| GO:1900409 | positive regulation of cellular response to oxidative stress | 0.044847002 | 3.657142857 | 0.177051042 |
| GO:0071404 | cellular response to low-density lipoprotein particle stimulus | 0.044847002 | 3.657142857 | 0.177051042 |
| GO:1901623 | regulation of lymphocyte chemotaxis | 0.044847002 | 3.657142857 | 0.177051042 |
| GO:0034114 | regulation of heterotypic cell-cell adhesion | 0.044847002 | 3.657142857 | 0.177051042 |
| GO:0034695 | response to prostaglandin E | 0.044847002 | 3.657142857 | 0.177051042 |
| GO:1901032 | negative regulation of response to reactive oxygen species | 0.044847002 | 3.657142857 | 0.177051042 |
| GO:0051279 | regulation of release of sequestered calcium ion into cytosol | 0.044882239 | 2.126245847 | 0.177051042 |
| GO:0051966 | regulation of synaptic transmission, glutamatergic | 0.044882239 | 2.126245847 | 0.177051042 |
| GO:0030808 | regulation of nucleotide biosynthetic process | 0.044882239 | 2.126245847 | 0.177051042 |
| GO:0000079 | regulation of cyclin-dependent protein serine/threonine kinase activity | 0.045176091 | 2.254403131 | 0.177911192 |
| GO:0014049 | positive regulation of glutamate secretion | 0.045220768 | 4.876190476 | 0.177911192 |
| GO:0070943 | neutrophil mediated killing of symbiont cell | 0.045220768 | 4.876190476 | 0.177911192 |
| GO:0010762 | regulation of fibroblast migration | 0.045507225 | 2.666666667 | 0.178404429 |
| GO:0032008 | positive regulation of TOR signaling | 0.045507225 | 2.666666667 | 0.178404429 |
| GO:0043277 | apoptotic cell clearance | 0.045507225 | 2.666666667 | 0.178404429 |
| GO:0032768 | regulation of monooxygenase activity | 0.045507225 | 2.666666667 | 0.178404429 |
| GO:0010976 | positive regulation of neuron projection development | 0.046754763 | 1.43605086 | 0.183133158 |
| GO:1903829 | positive regulation of cellular protein localization | 0.046863297 | 1.458689459 | 0.183396119 |
| GO:0045834 | positive regulation of lipid metabolic process | 0.047414764 | 1.645714286 | 0.185390472 |
| GO:0046620 | regulation of organ growth | 0.047470755 | 1.857142857 | 0.185445716 |
| GO:0050922 | negative regulation of chemotaxis | 0.047828369 | 2.398126464 | 0.186678129 |
| GO:0048662 | negative regulation of smooth muscle cell proliferation | 0.048334966 | 2.223938224 | 0.188489349 |
| GO:0007200 | phospholipase C-activating G-protein coupled receptor signaling pathway | 0.049183551 | 1.991513437 | 0.191629844 |
| GO:0034122 | negative regulation of toll-like receptor signaling pathway | 0.049321578 | 2.965250965 | 0.19199876 |
| GO:0097106 | postsynaptic density organization | 0.050791957 | 3.516483516 | 0.196342382 |
| GO:0032727 | positive regulation of interferon-alpha production | 0.050791957 | 3.516483516 | 0.196342382 |
| GO:1903798 | regulation of production of miRNAs involved in gene silencing by miRNA | 0.050791957 | 3.516483516 | 0.196342382 |
| GO:0002726 | positive regulation of T cell cytokine production | 0.050791957 | 3.516483516 | 0.196342382 |
| GO:1903077 | negative regulation of protein localization to plasma membrane | 0.050791957 | 3.516483516 | 0.196342382 |
| GO:1903975 | regulation of glial cell migration | 0.050791957 | 3.516483516 | 0.196342382 |
| GO:0014048 | regulation of glutamate secretion | 0.050791957 | 3.516483516 | 0.196342382 |
| GO:0042908 | xenobiotic transport | 0.050791957 | 3.516483516 | 0.196342382 |
| GO:0009887 | animal organ morphogenesis | 0.05146591 | 1.215969216 | 0.198682282 |
| GO:0046847 | filopodium assembly | 0.051486966 | 2.359447005 | 0.198682282 |
| GO:0060558 | regulation of calcidiol 1-monooxygenase activity | 0.052134702 | 7.836734694 | 0.200113661 |
| GO:0072679 | thymocyte migration | 0.052134702 | 7.836734694 | 0.200113661 |
| GO:1904139 | regulation of microglial cell migration | 0.052134702 | 7.836734694 | 0.200113661 |
| GO:0002408 | myeloid dendritic cell chemotaxis | 0.052134702 | 7.836734694 | 0.200113661 |
| GO:0002578 | negative regulation of antigen processing and presentation | 0.052134702 | 7.836734694 | 0.200113661 |
| GO:0032747 | positive regulation of interleukin-23 production | 0.052134702 | 7.836734694 | 0.200113661 |
| GO:0006913 | nucleocytoplasmic transport | 0.052219276 | 1.421169504 | 0.200113661 |
| GO:0051169 | nuclear transport | 0.052219276 | 1.421169504 | 0.200113661 |
| GO:0046165 | alcohol biosynthetic process | 0.05232767 | 1.828571429 | 0.200355729 |
| GO:0019377 | glycolipid catabolic process | 0.053462314 | 4.571428571 | 0.203994876 |
| GO:0032736 | positive regulation of interleukin-13 production | 0.053462314 | 4.571428571 | 0.203994876 |
| GO:1901722 | regulation of cell proliferation involved in kidney development | 0.053462314 | 4.571428571 | 0.203994876 |
| GO:0032645 | regulation of granulocyte macrophage colony-stimulating factor production | 0.053462314 | 4.571428571 | 0.203994876 |
| GO:0002064 | epithelial cell development | 0.053659491 | 1.49271137 | 0.204571039 |

| | | | | |
|---|---|---|---|---|
| GO:1901343 | negative regulation of vasculature development | 0.054187713 | 1.765517241 | 0.206407196 |
| GO:0001953 | negative regulation of cell-matrix adhesion | 0.054346323 | 2.887218045 | 0.20647874 |
| GO:0014912 | negative regulation of smooth muscle cell migration | 0.054346323 | 2.887218045 | 0.20647874 |
| GO:0033198 | response to ATP | 0.054346323 | 2.887218045 | 0.20647874 |
| GO:0035239 | tube morphogenesis | 0.054672976 | 1.378523915 | 0.207541804 |
| GO:0006672 | ceramide metabolic process | 0.054890412 | 1.952843273 | 0.208188807 |
| GO:0046622 | positive regulation of organ growth | 0.055317014 | 2.321995465 | 0.209448183 |
| GO:0090183 | regulation of kidney development | 0.055317014 | 2.321995465 | 0.209448183 |
| GO:0060562 | epithelial tube morphogenesis | 0.055792838 | 1.40311174 | 0.211069404 |
| GO:0007156 | homophilic cell adhesion via plasma membrane adhesion molecules | 0.056428466 | 1.671836735 | 0.213291898 |
| GO:0007205 | protein kinase C-activating G-protein coupled receptor signaling pathway | 0.057146075 | 3.386243386 | 0.215086755 |
| GO:1903513 | endoplasmic reticulum to cytosol transport | 0.057146075 | 3.386243386 | 0.215086755 |
| GO:0002523 | leukocyte migration involved in inflammatory response | 0.057146075 | 3.386243386 | 0.215086755 |
| GO:0030970 | retrograde protein transport, ER to cytosol | 0.057146075 | 3.386243386 | 0.215086755 |
| GO:1904996 | positive regulation of leukocyte adhesion to vascular endothelial cell | 0.057146075 | 3.386243386 | 0.215086755 |
| GO:0010565 | regulation of cellular ketone metabolic process | 0.057900398 | 1.703637977 | 0.217740884 |
| GO:0048193 | Golgi vesicle transport | 0.058288284 | 1.496695475 | 0.218937936 |
| GO:0007157 | heterophilic cell-cell adhesion via plasma membrane cell adhesion molecules | 0.058317554 | 2.509803922 | 0.218937936 |
| GO:0030968 | endoplasmic reticulum unfolded protein response | 0.058657855 | 2.13729128 | 0.219842893 |
| GO:1904029 | regulation of cyclin-dependent protein kinase activity | 0.058657855 | 2.13729128 | 0.219842893 |
| GO:0006753 | nucleoside phosphate metabolic process | 0.058896223 | 1.298059965 | 0.220549676 |
| GO:1904892 | regulation of STAT cascade | 0.059198675 | 1.627524308 | 0.221387027 |
| GO:0046686 | response to cadmium ion | 0.059319729 | 2.285714286 | 0.221387027 |
| GO:0048255 | mRNA stabilization | 0.059319729 | 2.285714286 | 0.221387027 |
| GO:0060393 | regulation of pathway-restricted SMAD protein phosphorylation | 0.059319729 | 2.285714286 | 0.221387027 |
| GO:0090162 | establishment of epithelial cell polarity | 0.059656245 | 2.813186813 | 0.222455527 |
| GO:0030500 | regulation of bone mineralization | 0.060386622 | 2.009419152 | 0.224989686 |
| GO:0051603 | proteolysis involved in cellular protein catabolic process | 0.060631123 | 1.264313152 | 0.225710819 |
| GO:0097479 | synaptic vesicle localization | 0.061162036 | 1.543098252 | 0.227496069 |
| GO:0045346 | regulation of MHC class II biosynthetic process | 0.062367661 | 4.302521008 | 0.231267997 |
| GO:0061621 | canonical glycolysis | 0.062367661 | 4.302521008 | 0.231267997 |
| GO:0014889 | muscle atrophy | 0.062367661 | 4.302521008 | 0.231267997 |
| GO:0014068 | positive regulation of phosphatidylinositol 3-kinase signaling | 0.062384934 | 2.10989011 | 0.231267997 |
| GO:0048705 | skeletal system morphogenesis | 0.062448937 | 1.501066098 | 0.231311699 |
| GO:0048016 | inositol phosphate-mediated signaling | 0.063019993 | 2.461538462 | 0.232842843 |
| GO:1902743 | regulation of lamellipodium organization | 0.063019993 | 2.461538462 | 0.232842843 |
| GO:0043618 | regulation of transcription from RNA polymerase II promoter in response to stress | 0.063019993 | 2.461538462 | 0.232842843 |
| GO:0043470 | regulation of carbohydrate catabolic process | 0.063496014 | 2.250549451 | 0.233965666 |
| GO:0030856 | regulation of epithelial cell differentiation | 0.063497842 | 1.643659711 | 0.233965666 |
| GO:0033157 | regulation of intracellular protein transport | 0.063591148 | 1.480885312 | 0.233965666 |
| GO:0051216 | cartilage development | 0.063612463 | 1.557975657 | 0.233965666 |
| GO:0045981 | positive regulation of nucleotide metabolic process | 0.063851672 | 1.98757764 | 0.233965666 |
| GO:0045778 | positive regulation of ossification | 0.063888936 | 1.828571429 | 0.233965666 |
| GO:0035967 | cellular response to topologically incorrect protein | 0.063888936 | 1.828571429 | 0.233965666 |
| GO:0070920 | regulation of production of small RNA involved in gene silencing by RNA | 0.063904843 | 3.265306122 | 0.233965666 |
| GO:1902884 | positive regulation of response to oxidative stress | 0.063904843 | 3.265306122 | 0.233965666 |
| GO:0060252 | positive regulation of glial cell proliferation | 0.063904843 | 3.265306122 | 0.233965666 |
| GO:1904376 | negative regulation of protein localization to cell periphery | 0.063904843 | 3.265306122 | 0.233965666 |
| GO:0014066 | regulation of phosphatidylinositol 3-kinase signaling | 0.064239996 | 1.897574124 | 0.234998499 |
| GO:0006775 | fat-soluble vitamin metabolic process | 0.065250636 | 2.742857143 | 0.238498613 |
| GO:0043271 | negative regulation of ion transport | 0.066177705 | 1.602356406 | 0.241687743 |
| GO:0051952 | regulation of amine transport | 0.066985224 | 1.813459268 | 0.242063453 |
| GO:0045833 | negative regulation of lipid metabolic process | 0.066985224 | 1.813459268 | 0.242063453 |
| GO:0032741 | positive regulation of interleukin-18 production | 0.067045566 | 6.857142857 | 0.242063453 |
| GO:0050861 | positive regulation of B cell receptor signaling pathway | 0.067045566 | 6.857142857 | 0.242063453 |
| GO:1904124 | microglial cell migration | 0.067045566 | 6.857142857 | 0.242063453 |
| GO:1904464 | regulation of matrix metallopeptidase secretion | 0.067045566 | 6.857142857 | 0.242063453 |
| GO:0048549 | positive regulation of pinocytosis | 0.067045566 | 6.857142857 | 0.242063453 |
| GO:0072126 | positive regulation of glomerular mesangial cell proliferation | 0.067045566 | 6.857142857 | 0.242063453 |
| GO:0060696 | regulation of phospholipid catabolic process | 0.067045566 | 6.857142857 | 0.242063453 |
| GO:1903976 | negative regulation of glial cell migration | 0.067045566 | 6.857142857 | 0.242063453 |
| GO:1903979 | negative regulation of microglial cell activation | 0.067045566 | 6.857142857 | 0.242063453 |
| GO:0045627 | positive regulation of T-helper 1 cell differentiation | 0.067045566 | 6.857142857 | 0.242063453 |
| GO:2000643 | positive regulation of early endosome to late endosome transport | 0.067045566 | 6.857142857 | 0.242063453 |
| GO:0071224 | cellular response to peptidoglycan | 0.067045566 | 6.857142857 | 0.242063453 |
| GO:2001259 | positive regulation of cation channel activity | 0.067440171 | 1.966205837 | 0.24309191 |
| GO:1905897 | regulation of response to endoplasmic reticulum stress | 0.067440171 | 1.966205837 | 0.24309191 |
| GO:0019217 | regulation of fatty acid metabolic process | 0.067570333 | 1.879839786 | 0.24336307 |
| GO:0060964 | regulation of gene silencing by miRNA | 0.067939298 | 2.41509434 | 0.244294716 |
| GO:0060251 | regulation of glial cell proliferation | 0.067939298 | 2.41509434 | 0.244294716 |
| GO:0046928 | regulation of neurotransmitter secretion | 0.068879715 | 1.74789916 | 0.247475376 |
| GO:0043255 | regulation of carbohydrate biosynthetic process | 0.070176826 | 1.798594848 | 0.251931394 |
| GO:0045780 | positive regulation of bone resorption | 0.071062059 | 3.15270936 | 0.253701058 |
| GO:0046852 | positive regulation of bone remodeling | 0.071062059 | 3.15270936 | 0.253701058 |
| GO:0006270 | DNA replication initiation | 0.071062059 | 3.15270936 | 0.253701058 |
| GO:0030204 | chondroitin sulfate metabolic process | 0.071062059 | 3.15270936 | 0.253701058 |
| GO:1901185 | negative regulation of ERBB signaling pathway | 0.071062059 | 3.15270936 | 0.253701058 |
| GO:0007176 | regulation of epidermal growth factor-activated receptor activity | 0.071062059 | 3.15270936 | 0.253701058 |
| GO:0043524 | negative regulation of neuron apoptotic process | 0.071126501 | 1.586005831 | 0.253701058 |
| GO:0070570 | regulation of neuron projection regeneration | 0.071127926 | 2.675958188 | 0.253701058 |
| GO:0009117 | nucleotide metabolic process | 0.071248158 | 1.283708491 | 0.253925455 |
| GO:0061051 | positive regulation of cell growth involved in cardiac muscle cell development | 0.071910058 | 4.063492063 | 0.254935972 |
| GO:0001821 | histamine secretion | 0.071910058 | 4.063492063 | 0.254935972 |
| GO:0071732 | cellular response to nitric oxide | 0.071910058 | 4.063492063 | 0.254935972 |
| GO:0071801 | regulation of podosome assembly | 0.071910058 | 4.063492063 | 0.254935972 |
| GO:0050862 | positive regulation of T cell receptor signaling pathway | 0.071910058 | 4.063492063 | 0.254935972 |
| GO:0061430 | bone trabecula morphogenesis | 0.071910058 | 4.063492063 | 0.254935972 |
| GO:0050829 | defense response to Gram-negative bacterium | 0.071934529 | 1.735140772 | 0.254935972 |
| GO:0014911 | positive regulation of smooth muscle cell migration | 0.072371337 | 2.18336887 | 0.256278994 |
| GO:0051588 | regulation of neurotransmitter transport | 0.073036986 | 1.684210526 | 0.257946533 |
| GO:0072666 | establishment of protein localization to vacuole | 0.073075147 | 2.37037037 | 0.257946533 |
| GO:1903573 | negative regulation of response to endoplasmic reticulum stress | 0.073075147 | 2.37037037 | 0.257946533 |
| GO:0045907 | positive regulation of vasoconstriction | 0.073075147 | 2.37037037 | 0.257946533 |
| GO:0070167 | regulation of biomineral tissue development | 0.074555403 | 1.845347313 | 0.262962131 |
| GO:0097480 | establishment of synaptic vesicle localization | 0.077600784 | 1.517155334 | 0.27326826 |
| GO:0048489 | synaptic vesicle transport | 0.077600784 | 1.517155334 | 0.27326826 |
| GO:0051209 | release of sequestered calcium ion into cytosol | 0.078210971 | 1.828571429 | 0.27485565 |
| GO:0006939 | smooth muscle contraction | 0.078210971 | 1.828571429 | 0.27485565 |
| GO:0046467 | membrane lipid biosynthetic process | 0.078302543 | 1.710174717 | 0.27485565 |
| GO:0010712 | regulation of collagen metabolic process | 0.078426731 | 2.327272727 | 0.27485565 |
| GO:2000727 | positive regulation of cardiac muscle cell differentiation | 0.078609957 | 3.047619048 | 0.27485565 |
| GO:0045332 | phospholipid translocation | 0.078609957 | 3.047619048 | 0.27485565 |
| GO:0072574 | hepatocyte proliferation | 0.078609957 | 3.047619048 | 0.27485565 |
| GO:1903649 | regulation of cytoplasmic transport | 0.078609957 | 3.047619048 | 0.27485565 |
| GO:0090200 | positive regulation of release of cytochrome c from mitochondria | 0.078609957 | 3.047619048 | 0.27485565 |
| GO:0051145 | smooth muscle cell differentiation | 0.078743524 | 2.006968641 | 0.274888739 |

| | | | | |
|---|---|---|---|---|
| GO:1903670 | regulation of sprouting angiogenesis | 0.078743524 | 2.006968641 | 0.274888739 |
| GO:0031640 | killing of cells of other organism | 0.080328048 | 1.755428571 | 0.280199411 |
| GO:0060491 | regulation of cell projection assembly | 0.081021963 | 1.53089701 | 0.282397557 |
| GO:0045071 | negative regulation of viral genome replication | 0.081943961 | 2.120082816 | 0.284671247 |
| GO:0032456 | endocytic recycling | 0.081943961 | 2.120082816 | 0.284671247 |
| GO:0008347 | glial cell migration | 0.081943961 | 2.120082816 | 0.284671247 |
| GO:0030837 | negative regulation of actin filament polymerization | 0.081943961 | 2.120082816 | 0.284671247 |
| GO:2000370 | positive regulation of clathrin-dependent endocytosis | 0.08205986 | 3.84962406 | 0.284671247 |
| GO:0033623 | regulation of integrin activation | 0.08205986 | 3.84962406 | 0.284671247 |
| GO:0031214 | biomineral tissue development | 0.082244454 | 1.581467181 | 0.285088365 |
| GO:0006664 | glycolipid metabolic process | 0.083038091 | 1.885125184 | 0.285348618 |
| GO:0006685 | sphingomyelin catabolic process | 0.0831569 | 6.095238095 | 0.285348618 |
| GO:0060753 | regulation of mast cell chemotaxis | 0.0831569 | 6.095238095 | 0.285348618 |
| GO:0031622 | positive regulation of fever generation | 0.0831569 | 6.095238095 | 0.285348618 |
| GO:0051126 | negative regulation of actin nucleation | 0.0831569 | 6.095238095 | 0.285348618 |
| GO:0031946 | regulation of glucocorticoid biosynthetic process | 0.0831569 | 6.095238095 | 0.285348618 |
| GO:0060556 | regulation of vitamin D biosynthetic process | 0.0831569 | 6.095238095 | 0.285348618 |
| GO:1990456 | mitochondrion-ER tethering | 0.0831569 | 6.095238095 | 0.285348618 |
| GO:0097527 | necroptotic signaling pathway | 0.0831569 | 6.095238095 | 0.285348618 |
| GO:0010727 | negative regulation of hydrogen peroxide metabolic process | 0.0831569 | 6.095238095 | 0.285348618 |
| GO:0070424 | regulation of nucleotide-binding oligomerization domain containing signaling pathway | 0.0831569 | 6.095238095 | 0.285348618 |
| GO:0044857 | plasma membrane raft organization | 0.0831569 | 6.095238095 | 0.285348618 |
| GO:1900125 | regulation of hyaluronan biosynthetic process | 0.0831569 | 6.095238095 | 0.285348618 |
| GO:0071526 | semaphorin-plexin signaling pathway | 0.083720805 | 2.551495017 | 0.287061275 |
| GO:0048145 | regulation of fibroblast proliferation | 0.083905 | 1.741496599 | 0.287248184 |
| GO:0015837 | amine transport | 0.083905 | 1.741496599 | 0.287248184 |
| GO:0032291 | axon ensheathment in central nervous system | 0.086539339 | 2.949308756 | 0.29467277 |
| GO:1903393 | positive regulation of adherens junction organization | 0.086539339 | 2.949308756 | 0.29467277 |
| GO:2000637 | positive regulation of gene silencing by miRNA | 0.086539339 | 2.949308756 | 0.29467277 |
| GO:0043552 | positive regulation of phosphatidylinositol 3-kinase activity | 0.086539339 | 2.949308756 | 0.29467277 |
| GO:0090025 | regulation of monocyte chemotaxis | 0.086539339 | 2.949308756 | 0.29467277 |
| GO:0051957 | positive regulation of amino acid transport | 0.086539339 | 2.949308756 | 0.29467277 |
| GO:0032801 | receptor catabolic process | 0.086539339 | 2.949308756 | 0.29467277 |
| GO:0019752 | carboxylic acid metabolic process | 0.087289331 | 1.203247401 | 0.296998263 |
| GO:0010469 | regulation of receptor activity | 0.088192413 | 1.564553094 | 0.299840668 |
| GO:0051928 | positive regulation of calcium ion transport | 0.088826069 | 1.630573248 | 0.301763408 |
| GO:0048666 | neuron development | 0.089599027 | 1.169840061 | 0.304042689 |
| GO:0030810 | positive regulation of nucleotide biosynthetic process | 0.089771521 | 2.245614035 | 0.304042689 |
| GO:0010559 | regulation of glycoprotein biosynthetic process | 0.089771521 | 2.245614035 | 0.304042689 |
| GO:1902883 | negative regulation of response to oxidative stress | 0.089771521 | 2.245614035 | 0.304042689 |
| GO:0002021 | response to dietary excess | 0.090429243 | 2.493506494 | 0.305802707 |
| GO:2000008 | regulation of protein localization to cell surface | 0.090429243 | 2.493506494 | 0.305802707 |
| GO:0002040 | sprouting angiogenesis | 0.091349721 | 1.714285714 | 0.308679837 |
| GO:0045600 | positive regulation of fat cell differentiation | 0.092209942 | 2.060362173 | 0.310921458 |
| GO:0031102 | neuron projection regeneration | 0.092209942 | 2.060362173 | 0.310921458 |
| GO:0055021 | regulation of cardiac muscle tissue growth | 0.09253969 | 1.936134454 | 0.310921458 |
| GO:0010468 | regulation of gene expression | 0.092586557 | 1.07088419 | 0.310921458 |
| GO:0071379 | cellular response to prostaglandin stimulus | 0.092785139 | 3.657142857 | 0.310921458 |
| GO:1903392 | negative regulation of adherens junction organization | 0.092785139 | 3.657142857 | 0.310921458 |
| GO:0060965 | negative regulation of gene silencing by miRNA | 0.092785139 | 3.657142857 | 0.310921458 |
| GO:0034383 | low-density lipoprotein particle clearance | 0.092785139 | 3.657142857 | 0.310921458 |
| GO:0090036 | regulation of protein kinase C signaling | 0.092785139 | 3.657142857 | 0.310921458 |
| GO:0006670 | sphingosine metabolic process | 0.092785139 | 3.657142857 | 0.310921458 |
| GO:0090030 | regulation of steroid hormone biosynthetic process | 0.092785139 | 3.657142857 | 0.310921458 |
| GO:0050848 | regulation of calcium-mediated signaling | 0.093924538 | 1.764411028 | 0.314501665 |
| GO:0071322 | cellular response to carbohydrate stimulus | 0.094406649 | 1.547996977 | 0.315877233 |
| GO:0034204 | lipid translocation | 0.094839713 | 2.857142857 | 0.316132377 |
| GO:0072576 | liver morphogenesis | 0.094839713 | 2.857142857 | 0.316132377 |
| GO:1903846 | positive regulation of cellular response to transforming growth factor beta stimulus | 0.094839713 | 2.857142857 | 0.316132377 |
| GO:0030511 | positive regulation of transforming growth factor beta receptor signaling pathway | 0.094839713 | 2.857142857 | 0.316132377 |
| GO:0001945 | lymph vessel development | 0.094839713 | 2.857142857 | 0.316132377 |
| GO:1990089 | response to nerve growth factor | 0.095760793 | 2.206896552 | 0.318244797 |
| GO:0001659 | temperature homeostasis | 0.095760793 | 2.206896552 | 0.318244797 |
| GO:0097035 | regulation of membrane lipid distribution | 0.095760793 | 2.206896552 | 0.318244797 |
| GO:0060337 | type I interferon signaling pathway | 0.095760793 | 2.206896552 | 0.318244797 |
| GO:0030148 | sphingolipid biosynthetic process | 0.096046268 | 1.828571429 | 0.318954249 |
| GO:0071804 | cellular potassium ion transport | 0.096403859 | 1.516376307 | 0.319662497 |
| GO:0071805 | potassium ion transmembrane transport | 0.096403859 | 1.516376307 | 0.319662497 |
| GO:0010765 | positive regulation of sodium ion transport | 0.097406354 | 2.438095238 | 0.322574304 |
| GO:0050810 | regulation of steroid biosynthetic process | 0.097427634 | 1.913621262 | 0.322574304 |
| GO:0048754 | branching morphogenesis of an epithelial tube | 0.097613878 | 1.539849624 | 0.322949572 |
| GO:0014897 | striated muscle hypertrophy | 0.098125551 | 1.749068323 | 0.324400141 |

**Appendix Table S6.** Enriched KEGG pathway in down-regulated genes in Ptf1a RGC.

| Term | description | PValue | Fold Enrichment | FDR |
|---|---|---|---|---|
| mmu04380 | Osteoclast differentiation | 4.68E-15 | 4.501686341 | 1.13E-12 |
| mmu05135 | Yersinia infection | 1.91E-11 | 3.847474264 | 2.30E-09 |
| mmu05417 | Lipid and atherosclerosis | 3.45E-11 | 3.088876397 | 2.77E-09 |
| mmu04670 | Leukocyte transendothelial migration | 1.65E-09 | 3.726641324 | 9.52E-08 |
| mmu05140 | Leishmaniasis | 1.97E-09 | 4.765695013 | 9.52E-08 |
| mmu04064 | NF-kappa B signaling pathway | 2.39E-09 | 3.899205011 | 9.60E-08 |
| mmu04662 | B cell receptor signaling pathway | 6.47E-09 | 4.305706493 | 2.23E-07 |
| mmu04142 | Lysosome | 9.75E-09 | 3.369683343 | 2.78E-07 |
| mmu04625 | C-type lectin receptor signaling pathway | 1.04E-08 | 3.655504698 | 2.78E-07 |
| mmu04668 | TNF signaling pathway | 1.27E-08 | 3.623155099 | 3.05E-07 |
| mmu05132 | Salmonella infection | 5.65E-08 | 2.517273327 | 1.15E-06 |
| mmu05152 | Tuberculosis | 5.85E-08 | 2.864230841 | 1.15E-06 |
| mmu05161 | Hepatitis B | 6.19E-08 | 2.976898168 | 1.15E-06 |
| mmu04621 | NOD-like receptor signaling pathway | 1.26E-07 | 2.634048247 | 2.17E-06 |
| mmu05145 | Toxoplasmosis | 5.58E-07 | 3.308416373 | 8.96E-06 |
| mmu05142 | Chagas disease | 6.68E-07 | 3.386041029 | 1.01E-05 |
| mmu05235 | PD-L1 expression and PD-1 checkpoint pathway in cancer | 7.57E-07 | 3.618580408 | 1.07E-05 |
| mmu04722 | Neurotrophin signaling pathway | 8.77E-07 | 3.13297005 | 1.12E-05 |
| mmu04062 | Chemokine signaling pathway | 8.80E-07 | 2.60623946 | 1.12E-05 |
| mmu05169 | Epstein-Barr virus infection | 9.80E-07 | 2.428797734 | 1.18E-05 |
| mmu04666 | Fc gamma R-mediated phagocytosis | 1.60E-06 | 3.461250825 | 1.84E-05 |
| mmu04072 | Phospholipase D signaling pathway | 3.94E-06 | 2.74776192 | 4.31E-05 |
| mmu04611 | Platelet activation | 4.93E-06 | 2.934885492 | 5.17E-05 |
| mmu04015 | Rap1 signaling pathway | 9.59E-06 | 2.338308301 | 9.63E-05 |
| mmu04659 | Th17 cell differentiation | 1.35E-05 | 3.032715008 | 1.30E-04 |
| mmu04010 | MAPK signaling pathway | 1.91E-05 | 2.063071434 | 1.77E-04 |
| mmu04620 | Toll-like receptor signaling pathway | 2.29E-05 | 3.032715008 | 2.05E-04 |
| mmu05162 | Measles | 2.60E-05 | 2.596502576 | 2.23E-04 |
| mmu05163 | Human cytomegalovirus infection | 2.70E-05 | 2.13237774 | 2.23E-04 |
| mmu05205 | Proteoglycans in cancer | 2.78E-05 | 2.293028421 | 2.23E-04 |
| mmu05134 | Legionellosis | 3.09E-05 | 3.728747961 | 0.000240075 |
| mmu04066 | HIF-1 signaling pathway | 4.70E-05 | 2.793290139 | 0.000354059 |
| mmu05167 | Kaposi sarcoma-associated herpesvirus infection | 6.15E-05 | 2.166225006 | 0.000449163 |
| mmu04664 | Fc epsilon RI signaling pathway | 7.82E-05 | 3.446267055 | 0.000554237 |
| mmu04060 | Cytokine-cytokine receptor interaction | 8.55E-05 | 1.973341958 | 0.000588752 |
| mmu05166 | Human T-cell leukemia virus 1 infection | 9.25E-05 | 2.062246206 | 0.000619117 |
| mmu05133 | Pertussis | 1.23E-04 | 3.150872736 | 0.000798319 |
| mmu04650 | Natural killer cell mediated cytotoxicity | 1.66E-04 | 2.637143486 | 0.001051839 |
| mmu05146 | Amoebiasis | 1.94E-04 | 2.692597437 | 0.001197495 |
| mmu04210 | Apoptosis | 2.05E-04 | 2.452931257 | 0.001236005 |
| mmu04630 | JAK-STAT signaling pathway | 2.52E-04 | 2.256484381 | 0.001466085 |
| mmu05418 | Fluid shear stress and atherosclerosis | 2.56E-04 | 2.356501527 | 0.001466085 |
| mmu04657 | IL-17 signaling pathway | 3.34E-04 | 2.771836298 | 0.001869233 |
| mmu04936 | Alcoholic liver disease | 0.000341271 | 2.365947879 | 0.001869233 |
| mmu05164 | Influenza A | 0.00039392 | 2.19126807 | 0.002109661 |
| mmu04071 | Sphingolipid signaling pathway | 0.000449006 | 2.44573791 | 0.0023524 |
| mmu04070 | Phosphatidylinositol signaling system | 0.000482685 | 2.685216414 | 0.002475044 |
| mmu04658 | Th1 and Th2 cell differentiation | 0.000562265 | 2.757013644 | 0.002823039 |
| mmu05200 | Pathways in cancer | 0.000786566 | 1.563830944 | 0.003868621 |
| mmu04933 | AGE-RAGE signaling pathway in diabetic complications | 0.000858086 | 2.552284908 | 0.004135976 |
| mmu05144 | Malaria | 0.001050289 | 3.192331588 | 0.004963133 |
| mmu04370 | VEGF signaling pathway | 0.001220795 | 3.137291388 | 0.005588478 |
| mmu04810 | Regulation of actin cytoskeleton | 0.001229001 | 1.929909551 | 0.005588478 |
| mmu04622 | RIG-I-like receptor signaling pathway | 0.001862433 | 2.816092508 | 0.008311968 |
| mmu05321 | Inflammatory bowel disease | 0.002145653 | 2.934885492 | 0.009327536 |
| mmu05170 | Human immunodeficiency virus 1 infection | 0.002167394 | 1.832265318 | 0.009327536 |
| mmu04613 | Neutrophil extracellular trap formation | 0.002295108 | 1.904603628 | 0.009703879 |
| mmu00562 | Inositol phosphate metabolism | 0.002385118 | 2.737867716 | 0.009910576 |
| mmu04660 | T cell receptor signaling pathway | 0.002900846 | 2.355506803 | 0.011849219 |
| mmu05171 | Coronavirus disease - COVID-19 | 0.003290846 | 1.78033877 | 0.013218233 |
| mmu05143 | African trypanosomiasis | 0.003353186 | 3.499286548 | 0.013247832 |
| mmu04935 | Growth hormone synthesis, secretion and action | 0.003731677 | 2.222248067 | 0.014505391 |
| mmu04931 | Insulin resistance | 0.005463506 | 2.205610915 | 0.020900078 |
| mmu04520 | Adherens junction | 0.006326072 | 2.562857754 | 0.023821614 |
| mmu00532 | Glycosaminoglycan biosynthesis - chondroitin sulfate / dermatan sulfate | 0.008233727 | 4.549072513 | 0.030482071 |
| mmu04145 | Phagosome | 0.008347787 | 1.83295962 | 0.030482071 |
| mmu04061 | Viral protein interaction with cytokine and cytokine receptor | 0.009084065 | 2.234632111 | 0.032675516 |
| mmu05100 | Bacterial invasion of epithelial cells | 0.010521879 | 2.394248691 | 0.037290776 |
| mmu04928 | Parathyroid hormone synthesis, secretion and action | 0.011045097 | 2.106052089 | 0.038577802 |
| mmu05323 | Rheumatoid arthritis | 0.011236501 | 2.265821558 | 0.038685667 |
| mmu04140 | Autophagy - animal | 0.011805462 | 1.922143315 | 0.040029799 |

| | | | | |
|---|---|---|---|---|
| mmu04068 | FoxO signaling pathway | 0.01195911 | 1.967792181 | 0.040029799 |
| mmu04216 | Ferroptosis | 0.014394709 | 3.032715008 | 0.04752226 |
| mmu05221 | Acute myeloid leukemia | 0.015685002 | 2.382847507 | 0.051082237 |
| mmu04920 | Adipocytokine signaling pathway | 0.017219253 | 2.349286274 | 0.0553312 |
| mmu04640 | Hematopoietic cell lineage | 0.019969861 | 2.097090165 | 0.063325482 |
| mmu04623 | Cytosolic DNA-sensing pathway | 0.021330475 | 2.406916673 | 0.066761616 |
| mmu04926 | Relaxin signaling pathway | 0.022099297 | 1.880753494 | 0.068281162 |
| mmu04970 | Salivary secretion | 0.023060848 | 2.140740006 | 0.070350182 |
| mmu00511 | Other glycan degradation | 0.027144858 | 4.212104178 | 0.081773885 |
| mmu05215 | Prostate cancer | 0.028766914 | 1.991176521 | 0.085590447 |
| mmu04912 | GnRH signaling pathway | 0.033545998 | 2.021810006 | 0.097892585 |
| mmu01521 | EGFR tyrosine kinase inhibitor resistance | 0.033714044 | 2.111383867 | 0.097892585 |
| mmu04750 | Inflammatory mediator regulation of TRP channels | 0.039332313 | 1.79097343 | 0.112846278 |
| mmu04979 | Cholesterol metabolism | 0.039972764 | 2.475685721 | 0.113334544 |
| mmu04217 | Necroptosis | 0.041121117 | 1.636976851 | 0.115234758 |
| mmu04510 | Focal adhesion | 0.041714995 | 1.584254109 | 0.115555332 |
| mmu04020 | Calcium signaling pathway | 0.044105043 | 1.516357504 | 0.120787674 |
| mmu04917 | Prolactin signaling pathway | 0.052937371 | 2.049131762 | 0.143347264 |
| mmu05231 | Choline metabolism in cancer | 0.056672669 | 1.856764291 | 0.151756813 |
| mmu05212 | Pancreatic cancer | 0.060878544 | 1.995207242 | 0.161227791 |
| mmu05415 | Diabetic cardiomyopathy | 0.063103679 | 1.509170976 | 0.164872681 |
| mmu05202 | Transcriptional misregulation in cancer | 0.063623068 | 1.489279692 | 0.164872681 |
| mmu05160 | Hepatitis C | 0.076466639 | 1.562307732 | 0.196047447 |
| mmu05222 | Small cell lung cancer | 0.084381053 | 1.793541134 | 0.214061407 |
| mmu04144 | Endocytosis | 0.086993736 | 1.393710941 | 0.218390525 |
| mmu04024 | cAMP signaling pathway | 0.087991366 | 1.447432163 | 0.218617723 |
| mmu04932 | Non-alcoholic fatty liver disease | 0.089114563 | 1.555238466 | 0.219149078 |
| mmu04514 | Cell adhesion molecules | 0.090756937 | 1.499694235 | 0.220933554 |
| mmu00740 | Riboflavin metabolism | 0.093009026 | 5.686340641 | 0.22263456 |
| mmu04930 | Type II diabetes mellitus | 0.09330328 | 2.211354694 | 0.22263456 |
| mmu04919 | Thyroid hormone signaling pathway | 0.096065967 | 1.64272063 | 0.226979393 |
| mmu04218 | Cellular senescence | 0.097875946 | 1.483393211 | 0.229010708 |
| mmu05223 | Non-small cell lung cancer | 0.099595699 | 1.89544688 | 0.230793879 |
